# Dormant viral pathways underlie space-induced neural senescence

**DOI:** 10.1101/2025.11.02.686043

**Authors:** Aline M.A. Martins, Hideyuki Nakashima, Angela Macia, Taichi Umeyama, Satoshi Miyashita, Livia Luz de Souza Nascimento, Luisa Bulcao Vieira Coelho, Pinar Mesci, Erik Viirre, Nicole Suarez, Aurian Saleh, Alex Savchenko, Beatriz Freitas, Aline L. Leite, Diego M. Assis, Masataka Ishizu, Tomoo Owa, Mikio Hoshino, John Yates, Kinichi Nakashima, Alysson R. Muotri

**Author notes:** These authors contributed equally to this work (Aline M.A. Martins, Hideyuki Nakashima, Angela Macia).

## Abstract

Long-duration spaceflight is associated with neurological symptoms in astronauts, yet the underlying molecular mechanisms remain unclear. Using human brain organoids cultured aboard the International Space Station, we analyzed three independent spaceflights to demonstrate that exposure to the space environment triggers Space-Induced Neural Senescence (SINS), characterized by chromatin remodeling, mitochondrial dysfunction, and activation of viral-like transcriptional programs in the absence of infection. Multi-omics analyses identified upregulation of endogenous LINE-1 (L1) retroelements, whose activity was markedly enhanced in organoids lacking MECP2, a known L1 repressor implicated in Rett syndrome. The resulting accumulation of cytoplasmic L1 DNA elicited an IL-6-mediated inflammatory and neurotoxic response, which was reversed by reverse transcriptase inhibitors (RTi) such as lamivudine or stavudine. Parallel preclinical experiments in *Mecp2*-deficient mice confirmed that RTi treatment restored neuronal morphology, synaptogenesis, function, cognition, and survival. These findings reveal that the space environment reactivates dormant genomic retroelements, providing an unexpected mechanistic insight into astronaut neurobiology and identifying a potential therapeutic strategy for both space-induced and terrestrial neurological conditions. Our pioneering study demonstrates the value of space-enabling research in accelerating drug discovery and disease treatment on Earth.

## INTRODUCTION

In recent years, human space exploration efforts have increased substantially, including the Artemis mission to land on the Moon in preparation for missions to Mars, as well as numerous private companies preparing to build the next generation of space stations in Low-Earth Orbit (LEO) to replace the International Space Station (ISS) ^1^. As humans push farther into space, they face hazards like cosmic radiation and prolonged microgravity, which can cause a spectrum of physiological issues, including DNA damage, musculoskeletal degeneration, and neurological problems ^2^. Space exposure is known to affect the astronaut’s brain, as evidenced by changes such as increased ventricular volume ^3,4^, altered global white matter volume ^5,6^, and persistent cognitive decline post-flight ^2–4,7–9^. However, the exact mechanisms remain unknown, largely due to limitations of current noninvasive brain imaging technologies. Therefore, successful long-duration exploration requires a comprehensive understanding of spaceflight’s impact to develop effective countermeasures ^10^. The SpaceX lnspiration4 mission highlighted the value of biomedical research in private space missions by collecting extensive health data, including multi-omic analyses. This contributes to the expanding biomedical database ^11^ and supports precision astronaut medicine. This “Second Space Age,” characterized by commercial participation ^12^, provides diverse participant data from non-professional astronauts. Recent studies utilize multi-omics and systems biology to characterize the effects of spaceflight. Authors aggregated transcriptomic, proteomic, metabolomic, and epigenetic profiles, revealing alterations converging on mitochondrial and metabolic processes, innate immunity, chronic inflammation, cell cycle regulation, circadian rhythm, and olfactory functions ^1–13^. Confirmed by NASA’s Twin Study, mitochondrial dysregulation emerged as a consistent phenotype across multiple biological layers ^14–16^.

Although microgravity is the most recognized abiotic event in the space environment, radiation is among the foremost health risks facing astronauts during space missions, particularly long-duration exploration missions ^14^. The chronic low-dose radiation exposure increases their risk of developing cancer, induces DNA damage, and triggers oxidative stress ^14,17^. Radiation also contributes to neurological disruptions, such as neurovascular damage and neurodegeneration, in simulated models ^18,19^, potentially accelerating aging ^18^.

To understand how the space environment affects the brain, Masarapu and colleagues used single-nucleus multi-omics and spatial transcriptomics on mouse brains flown on the ISS for around 40 days ^20^. This study revealed neuroinflammation, oxidative stress, and impaired synaptic signaling, consistent with findings in neurodegenerative disorders and radiation studies. However, similar studies cannot be conducted in humans; thus, the translatability of these observations is unclear.

Due to the inability to study astronaut brains at the cellular level, we used human brain organoids as a model system. These organoids mimic the functional and cellular features of the human brain ^21^, providing a unique experimental model for investigating the consequences of space exposure ^22^. We developed an automated device to sustain organoid culture on the ISS and return live samples, with an Earth replica serving as ground control. Next, both proteomics and transcriptomics analyses were performed on the brain organoid samples returned from the ISS. One of the most representative molecular pathways upregulated in the flight samples relative to the ground control was associated with viral infection. In the absence of any additional viral infections, we hypothesized that this activation resulted from an increase in endogenous Long Interspersed Nuclear Elements-1 (LINE-1, or L1) retroelements. L1 elements are a class of retrotransposons comprising approximately 20% of the mouse and human genomes ^23,24^. Several neurological and psychiatric diseases exhibited variation in L1 expression, suggesting L1s could be detrimental when de-regulated ^25–31^, especially in age-related neurodegenerative diseases ^32,33^. The upregulation of this pathway is linked to a series of histone modifications, relaxing the chromatin, and alterations in MECP2, a well-known L1 repressor ^34–38^.

To translate our observations in space to a clinically relevant model on Earth, we studied L1 upregulation in Rett syndrome (RTT, OMIM#312750), caused by MECP2 mutations ^39^. We show that increased L1 expression in MECP2-mutant human astrocytes triggers an Interleukin 6 (IL6)-mediated inflammatory neurotoxic response. Blocking L1 reverse transcription using reverse transcriptase inhibitors (RTi) ameliorates neurotoxicity, improving cognition and survival. Our results contribute to understanding the molecular mechanisms underlying the neurological consequences of spaceflight and offer potential neuroprotective countermeasures for astronauts and therapeutic opportunities for neurological conditions on Earth.

## RESULTS

### An autonomous platform for human brain organoids at ISS

To investigate how the space environment affects human brain organoids, we used a multi-modal approach, combining proteomics (mass spectrometry-based) and transcriptomics in two different levels of information (bulk and single-nuclei analysis). We used human brain organoids generated using a semi-guided protocol, a robust approach that yields consistent results across different iPSC lines ^1,2^. Derivation of brain organoids using this approach yields clear neural rosettes with a spatially organized arrangement of proliferative cells, intermediate progenitor cells, glial cells, and lower- and upper-layer cortical neurons. Based on cell population annotation, 1-month organoids consisted of approximately 70% progenitor and radial glial cells (expressing markers such as SOX2) that would mature into functional neurons (NeuN2-positive) and astrocytes (GFAP-positive). Over the course of organoid maturation, neurons exhibit a characteristic pyramidal morphology, and mature synaptic ultrastructure develops, enabling the formation of functional populations of inhibitory interneurons, which in turn facilitate the establishment of complex neural circuitry. One-month-old human brain organoids from 3 iPSC lines derived from healthy individuals with different genetic backgrounds, plus a cell line with mutated *MECP2* (MECP2-KO), were loaded into a 9U Cubelab™ hardware (Space Tango) on three independent Commercial Resupply Services (CRS) using SpaceX (SpX) rockets missions (CRS-SpX18, CRS-SpX30; CRS-SpX31), each lasting around 30 days at the ISS (**Fig. 1**), returning with two months old. Space Tango’s flight and ground hardware supported 12 wells, each equipped with impellers and automated operations (fluid conveyance, fluid routing, thermal control, environmental monitoring, imaging, and sampling), containing approximately 30 organoids (**Fig. 1** and **Fig. S1A).** The organoids were monitored in real time in each well in both environments, the ground and the ISS units **(Fig. S1A).** Cubelab environmental telemetry (CO2 levels, culture medium temperature, and radiation) was recorded during operations **(Fig. S1B and E,** and **Table S1**). As they return from the ISS at 2 months old, brain organoids are self-organized in a structure containing five major cell types - progenitors, intermediate progenitors, glial cells, glutamatergic neurons, and GABAergic neurons, as previously described **(Fig. S1C).** We also visually tracked the organoids’ viability and overall health in both environments, measuring their diameter during the experimental weeks **(Fig. S1D).** ISS and ground samples were run synchronously, and scheduled operations were initiated simultaneously. Upon return to Earth, the samples were processed for proteomic and transcriptomic analyses.

**Fig. 1.**
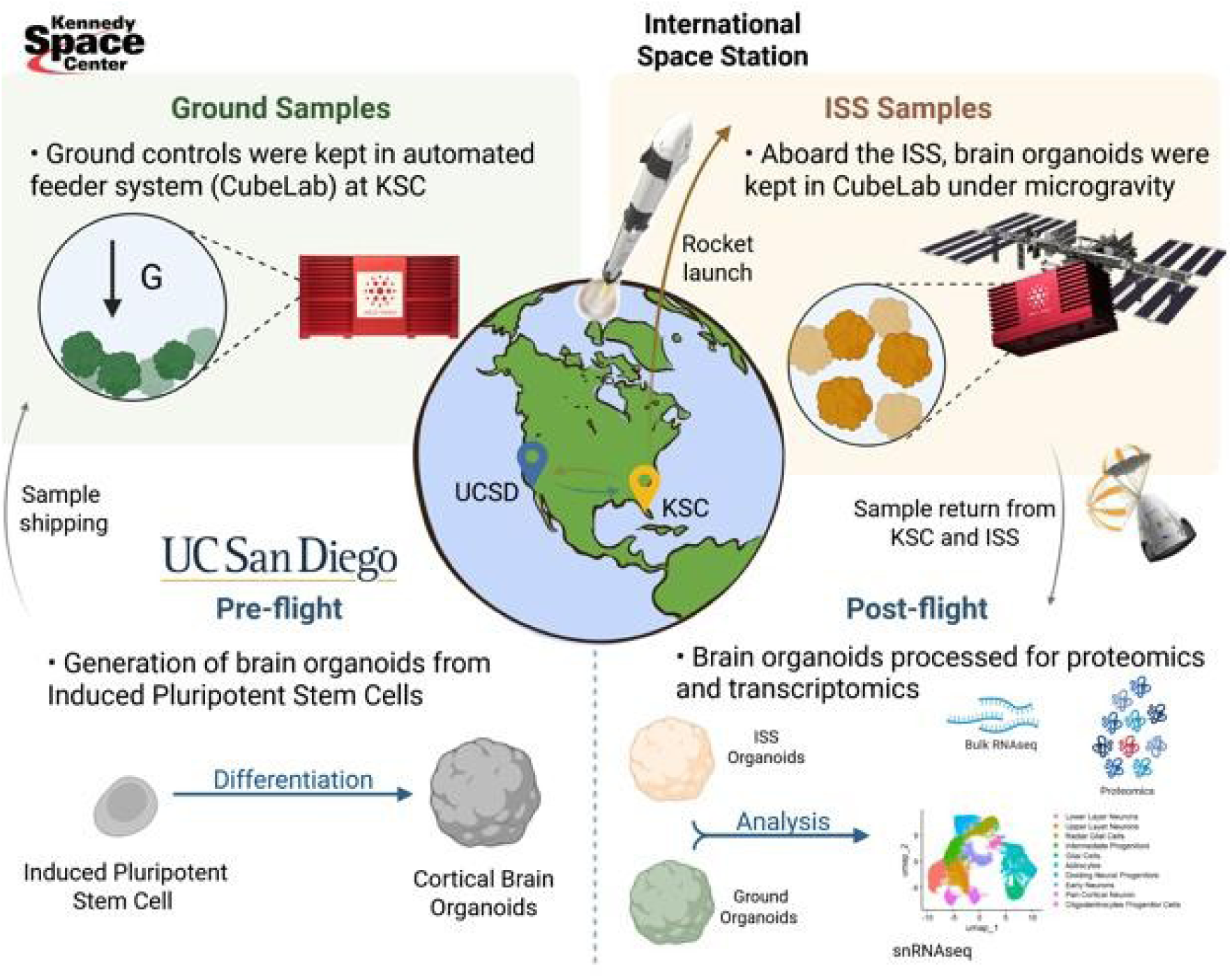
Schematic brain organoids experiments sent to the International Space Station (ISS). This schematic depicts the process and experimental setup for studying brain organoids under space environment conditions aboard the International Space Station (ISS). Induced pluripotent stem cells were differentiated into brain organoids. These brain organoids were launched on SpaceX CRS-SpX18, CRS-SpX30; CRS-SpX31 missions (on average 30 days) and maintained aboard the ISS in an automated system (Cubelab), exposed to the space environment. A synchronous ground control group of brain organoids was maintained in parallel under identical conditions in another Cubelab to enable comparative studies. Following their return from ISS, comprehensive molecular analyses, including proteomic and transcriptomics assessments, were conducted on both ISS and Earth-bound brain organoids to evaluate the impact of space environment conditions on molecular and cellular processes and development.

### Space-induced neural senescence - SINS

The effects of the space environment on brain organoids revealed significant alterations in both proteomic and transcriptomic profiles. A summary of all experiments performed in this study, including flight missions and details of their statistical analyses, is provided in **Table S2.** In the proteomics analysis, we identified an average of 6000 proteins per run and 4059 differentially expressed proteins between brain organoids on Earth (ground control) and those exposed to the space environment at the ISS (p≤0.01; fold change 1.2) **(Table S3).** Proteomic analysis of samples at the ISS-exposed to space demonstrated a distinct set of upregulated proteins (**Fig. 2A**). Proteins such as tenascin (TNC); neurofilament light polypeptide (NEFL); mitochondrial import inner membrane translocase (TIMM8A), and subunit of cytochrome c oxidase (HIGD1A), show significant fold changes and p-values, implying alterations on the metabolism of the brain organoids. Reactome pathway analysis identified key affected pathways, including RNA metabolism, intracellular protein transport, and viral infection (**Fig. 2B**).

**Fig. 2.**
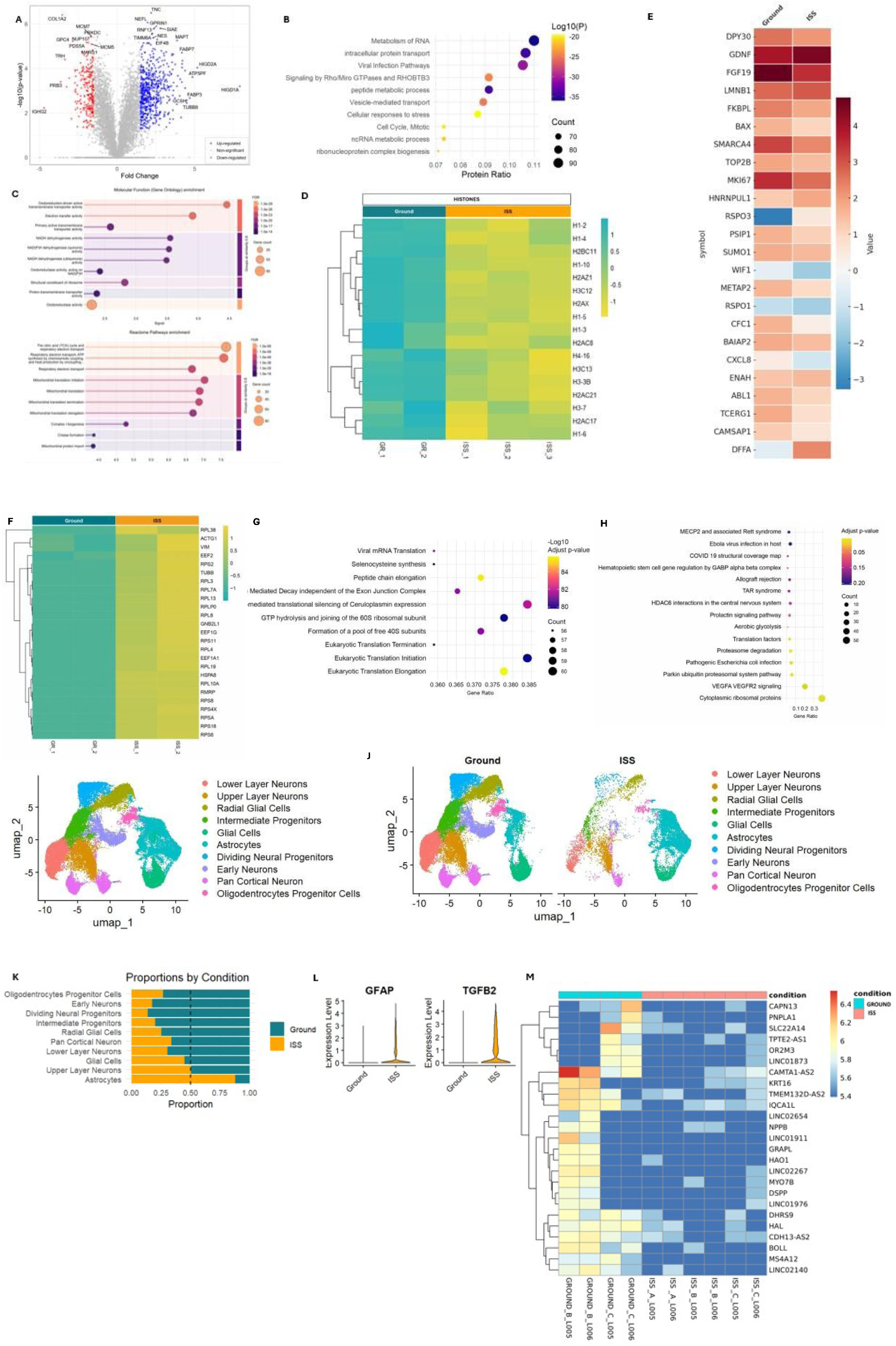
Proteomics and transcriptomics analysis data from healthy brain organoids at the International Space Station (ISS). **(A)** Volcano plot of proteomics data showing proteins upregulated (in blue) and downregulated (in red) in brain organoids at ISS, when compared to ground control. **(B)** Reactome pathway analysis of the proteins upregulated in the proteomics experiment. **(C)** Gene Ontology (GO) analysis for the mitochondria framework. 629 downregulated proteins are significantly enriched in functions and pathways crucial for cellular energy production (respiration) and the synthesis of mitochondrial proteins. **(D)** Heatmap of a framework of general downregulation in histone showing global chromatin remodeling. **(E)** Secretome panel on a subsequent flight (Spx31). The media of a batch of healthy brain organoids with 2 months at arrival were evaluated using Olink Reveal® panel technology (Thermo Fisher Scientific). Sample controls were used to determine precision within and between plates, triplicate negative controls samples set the limit of detection (LOD) and triplicate samples were used to standardize protein levels within a plate. Secretome sample run 1:1. Counts of known sequences were translated into Normalized Protein eXpression (NPX) values as their log-2 scaled quantification. **(F)** Heatmap of the top 25 Differentially Expressed Genes (DEGs) in ISS *vs.* ground conditions. The colors represent fold changes, with red indicating genes that are downregulated and blue indicating genes that are upregulated. **(G** Reactome pathway analysis of the upregulated genes from the transcriptomics experiment (Panel D). **(H)** WikiPathway enrichment analysis of the transcriptomic data (Panel D). **(I)** The UMAP plot shows the distribution of various neural cell types identified across both ground and ISS conditions. Each cluster is color-coded to represent a specific cell population: lower layer neurons (red), upper layer neurons (orange), radial glial cells (yellow-green), intermediate progenitors (green), glial cells (teal), astrocytes (cyan), dividing neural progenitors (light blue), early neurons (purple), pan cortical neuron (pink), and oligodendrocytes progenitor cells (magenta). The plot highlights the spatial organization of cell clusters, emphasizing the diversity of cell types within the dataset. **(J)** UMAP analysis comparing brain organoids samples from ground and ISS. **(K)** Stacked bar plots representing the relative proportions of different cell types in the ground and ISS. Each bar represents a specific cell type, with the dashed line indicating equal proportion mark. (L) *GFAP* and *TGFB2* expression in cells across ground and ISS conditions. Violin plots show the distribution of gene expression levels for GFAP and TGFB2 in cells from the ground and ISS environments. Each violin represents the kernel density estimation of log expression values, indicating higher levels of expression in the ISS condition compared to the ground condition. **(M)** Heatmap of healthy brain organoids, using bulk transcriptomics analysis, in a subsequent flight (SpX30). For proteomics analysis: p≤0.01; fc≥1.2. Test T Student for comparisons. For transcriptomics bulk analysis: p≤0.05; fc≥1.2. For pathway analysis: *Homo sapiens* as the reference organism, minimum overlap of 3 genes, enrichment factor > 1.5, and p < 0.01.

To evaluate the oxidation/phosphorylation (OxPHOS) state of the experiment, we performed an enrichment analysis and observed the downregulation of the mitochondrial-related proteins in the ISS (**Fig. 2C**). We found 629 downregulated proteins are significantly enriched in functions and pathways crucial for cellular energy production (respiration) and the synthesis of mitochondrial proteins, demonstrating that space environment causes a deep alteration in the energetic machinery in the brain organoids. The most significantly affected molecular functions are related to the electron transport chain and oxidoreductase activity. The most enriched biological pathway is associated with the citric acid (TCA) cycle and respiratory electron transport, indicating that multiple aspects of mitochondrial protein synthesis, including translation initiation, elongation, and termination, are significantly downregulated, suggesting reduced capacity for building and repairing mitochondrial components (**Fig. 2C**).

We also observed alterations in epigenetic marks. Histones play a crucial role in regulating gene expression by influencing chromatin structure and accessibility. Reduced histone levels can alter chromatin dynamics, making DNA more accessible to transcription factors and enzymes involved in DNA repair and replication ^3^. Such alterations might reflect cellular responses to stress conditions and cellular senescence^4^. Histones are globally downregulated in the ISS when compared to ground control brain organoids (**Fig. 2D**). Specifically, the histone acetyltransferases (HAT1 - histone acetyltransferase type B; KAT8 - histone acetyltransferase KAT8; KAT? - histone acetyltransferase KAT?), usually linked to open chromatin, were upregulated at the ISS **(Fig. S2A,** upper panel). At the same time, we observed downregulation of deacetylases at the ISS, indicating a withdrawal of acetyl groups **(Fig. S2A,** bottom panel). Together, these observations suggest that cells in brain organoids exposed to the ISS undergo chromatin remodeling, thereby facilitating genomic accessibility and overall transcription.

To further complement our proteomics analysis, we evaluated the secretome panel on a subsequent flight (Spx31), aiming to understand how the space environment altered the pattern of secreted effectors (**Fig. 2E**). The conditioned media of a batch of control brain organoids were analyzed using Olink Reveal® panel technology, a Proximity Extension Assay technology (PEA). The readouts were compared with an aliquot of unconditioned medium (blank) used to grow the organoids in the ISS, and this negative control was compared with each of the tested conditioned media. The data analysis revealed that the secretome of brain organoids exposed to a space environment exhibited a coordinated response of protein effectors in the ISS relative to ground control. The secretome panel highlight: *i)* modulation of neural development activity: the upregulation of secreted Glial Cell Line-Derived Neurotrophic Factor (GDNF) and sustained expression of Fibroblast Growth Factor 19 (FGF19) suggests an intrinsic adaptive mechanism where supportive glial cells respond to the unique demands of the space environment by enhancing trophic support and developmental cues, against space-induced cellular insults ^5^_;_ *ii)* alteration of proliferative state, impacting epigenetic regulation: downregulation in the ISS brain organoids of DPY30 (Dpy30 histone methyltransferase complex subunit), SMARCA4 (SWI/SNF related, matrix associated, actin dependent regulator of chromatin, subfamily A, member 4), TOP2B (DNA Topoisomerase II Beta) and MKl67 (marker of proliferation Ki-67) emphasizes changes in chromatin remodeling and gene expression regulation in the space environment, and a coordination of a robust epigenetic response^6^_;_ *iii)* alterations in cytoskeletal dynamics and trafficking: together, the downregulation compared to ground samples of BAIAP2 (brain-specific angiogenesis inhibitor 1-associated protein 2), CAMSAP1 (calmodulin regulated spectrin associated protein 1) and ABL1 (ABL proto-oncogene 1, non-receptor tyrosine kinase) indicates a significant disruption or alteration in actin cytoskeleton organization and dynamics, that could lead to changes in cell shape, migration, and the formation of cellular protrusions essential for neuronal connectivity^7,8^.

The bulk transcriptomic data also showed significant differential gene expression between the ground control and organoids at the ISS. We identified 150 differentially expressed genes (DEGs) with a 1.5-fold change and p-values <0.05. **(Table S4).** Most DEGs are involved in viral RNA translation, followed by selenocysteine synthesis and peptide chain elongation (**Fig. 2F**). Consistent with the proteomic pathway analyses, the transcriptional Reactome pathway analyses also uncovered activation of viral response pathways (**Fig. 2G**). Moreover, WikiPathway analyses revealed significant alterations in MECP2-related gene expression under space environment conditions (**Fig. 2H**). To determine whether the viral pathway could also be detected in the blood of astronauts, we compared our DEGs with those from the NASA Twins Project using Ingenuity Pathway Analysis (IPA). As expected, pathways related to the hematopoietic system (e.g., T-cell activation) are present only in the blood astronaut samples **(Fig. S2B).** The top hits across biological and disease pathways include several viral-related pathways, such as viral infection, retrovirus replication, and HIV infection, among others **(Fig. S2B).**

To determine which cell-type-specific responses were affected by the space environment at the ISS, we performed a single-nucleus transcriptomics experiment (snRNA-seq). We identify 10 cell populations in the brain organoids: lower-layer neurons, upper-layer neurons, radial glial cells, intermediate progenitors, glial cells, astrocytes, dividing neural progenitors, early neurons, pan-cortical neurons, and oligodendrocyte progenitor cells (**Fig. 2I** and **Fig. S2C).** Changes in cell populations were observed when we compared ground control and brain organoids exposed to the space environment (**Fig. 2J**). Mainly, we observed a higher number of active astrocytes (glial fibrillary acidic protein (GFAP)-expressing cells and transforming growth factor beta 2 (TGFβ2)-expressing cells) in organoids exposed to the space environment (**Figs. 2K-L**). We generate a heatmap of differentially expressed key marker genes across cell types. We observed distinct expression patterns across cell clusters, with GFAP expression with a higher intensity in astrocytes; forkhead-box G1 (FOXG1), 8-cell lymphoma/leukemia 11b (BCL11b) and (glutamic acid decarboxylase) GAD1, with high intensity in upper- and lower-layer neurons and intermediate progenitors **(Fig. S2C).**

We evaluated human brain organoids from another independent spaceflight (SpX30) to validate our previous findings using bulk transcriptomics. The subsequent spaceflight also revealed similar signaling pathways altered in the ISS, complementing the original observations from the SpX18 flight (**Fig. 2M**). The analysis of the gene expression differences between brain organoids in the ground experiments *vs.* brain organoids exposed to the ISS revealed: *i)* downregulation of non-coding RNAs: the downregulation of LINC RNAs (Long Interspersed Non-Coding RNAs), such as LINC01873, LINC02654, LINC01911. Interestingly, the primary function of these LINC RNAs is to suppress the activity of retrotransposons, such as L1 elements; *ii)* cellular stress: the decreased expression of CAPN13 (calpain 13), PNPLA1 (patatin-like phospholipase 1), and SLC22A14 (transmembrane solute carrier family 22 member 14) indicates impairment of neurogenesis and reduction in cell proliferation and long-term survival of neurons; suppressed oxidative phosphorylation, specially through mitochondria deficiency in ATP production, leading to impaired in fatty acid beta-oxidation (FAO) and reduction in acyl-carnitines and TCA cycle; *iii)* altered neurodevelopment and impaired plasticity: the coordinated downregulation of key developmental genes - NPPB (natriuretic peptide B), MYO7B (myosin VIIB), and CDH13-AS2 (long non-coding RNA (lncRNA) that plays a role in regulating the expression of the CDH13 gene, also known as T-cadherin) indicates that the space environment may hinder the brain’s developmental genesis (**Fig. 2M**).

Collectively, all the changes in transcriptomics and proteomics described above represent a novel phenomenon not observed on Earth. Thus, we coined the term “Space-Induced Neural Senescence” (SINS) to distinguish it from the linear aging process observed on the ground. These findings provide insights into the neurological effects of spaceflight, advancing our understanding of the molecular mechanisms underlying the negative impact on the human brain caused by the space environment.

### Space environment activates retroelement transcription in mice

To independently validate our main observations in a different model system, we analyzed publicly available RNA-seq data from the retinas of mice exposed to spaceflight^10^. Although there was no overall significant change in global gene expression between the ISS samples and ground controls (**Fig. 3A-C**), gene set enrichment analysis (GSEA) revealed enrichment of genes associated with retrotransposon silencing, transposition, IL6-type cytokine receptor interaction, inflammasome activation, and phagocytosis in the ISS samples. In addition, mitochondrial dysfunction in ISS was suggested by downregulation of the aerobic electron transport chain and mitochondrial biogenesis, and by upregulation of cytochrome c release (**Fig. 3D, bottom panel),** corroborating our previous mitochondrial findings in the human model. Based on these observations, we inferred that retrotransposons are upregulated in the mouse retina during spaceflight. Supporting this, we found an increased expression of class I retrotransposable elements: LINE, SINE, and LTR **(Fig. S3A).** Notably, multiple L1 subfamilies were significantly upregulated in the ISS samples **(Fig. S3B).** Retrotransposable elements are also upregulated following exposure to simulated 5-ion galactic cosmic radiation (GCR), which models the effect of cosmic rays **(Fig. S3C-D).** Furthermore, both evolutionary young and ancient L1 subfamilies have been previously shown to be upregulated in the mouse lung under combined GCR and microgravity exposure^11^. Together, these findings suggest that environmental factors specific to space, including microgravity and cosmic radiation, can trigger the activation of endogenous L1 retroelements in both mouse and human brains.

**Fig. 3.**
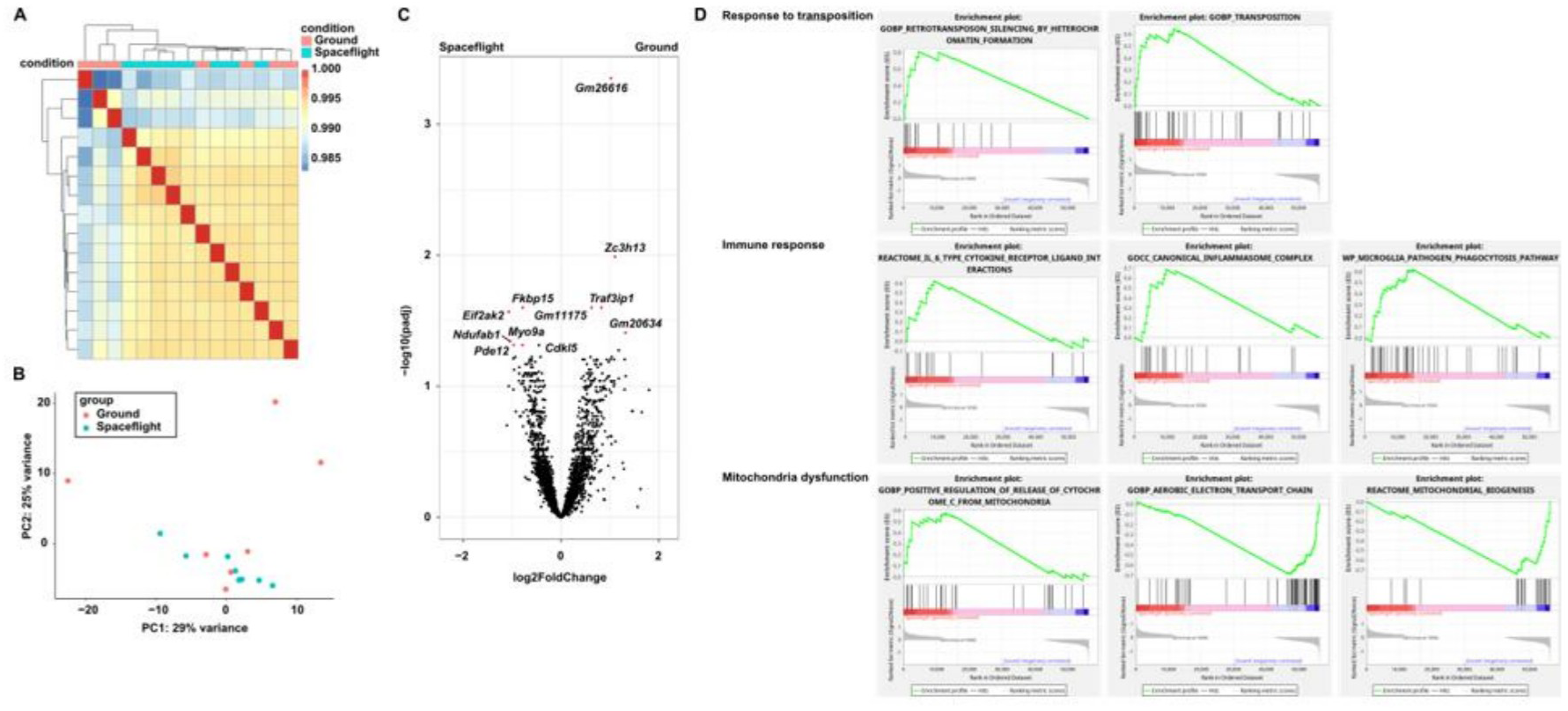
Reanalysis of publicly available RNAseq data from ISS and ground control mouse retinas. **(A)** Heatmap of ISS and ground control retinas. **(B)** Principal coordinates analysis (PCoA) of ISS and ground control retinas. **(C)** Volcano plot of differentially expressed genes. **(D)** GSEA analysis of spaceflight and ground control retinas.

### MECP2-KO brain organoids exacerbate the viral pathway at the ISS

In the absence of external viral infections and based on our previous results, we hypothesized that activation of endogenous L1 retroelements led to the upregulation of the viral pathway observed upon space exposure in human brain organoids. L1s can reverse-transcribe their RNA sequences and accumulate in the cytoplasm of astrocytes, thereby triggering an inflammatory response ^12,13^. To test this hypothesis, we used brain organoids lacking functional MeCP2 (MECP2-KO^9^), a known **L**1 repressor^14–16^. Moreover, our transcriptomic results indicate alterations in MECP2-related gene expression under spaceflight conditions (**Fig. 2H**). MECP2-KO and control brain organoids were exposed to the space environment and subjected to bulk proteomics analysis (**Fig. 4A** and **Table S3).** The Reactome analysis showed that the most upregulated proteins are related to cellular stress responses, including viral pathways (**Fig. 4B** and **Fig. S4).** The viral pathway was highly enriched (-log_10_(p) = −9.975, ratio = 0.044328553), comprising 34 proteins **(Table S5).** The viral pathway was further increased in the absence of the MECP2 gene (**Fig. 4B** and **Fig. S4A),** confirming MeCP2’s role in silencing this pathway. We evaluate the bulk transcriptome of the same MeCP2-KO model brain organoids in a subsequent flight (SpX30). The heatmap shows that the genes listed are overall more highly expressed in the MeCP2-KO ISS samples and show significantly lower expression in the healthy brain organoids samples (ISS control) (**Fig. 4C**). A functional analysis of the most highly upregulated genes reveals a pro-inflammatory and immune-active state in the MeCP2-KO ISS samples, corroborating the first spaceflight (SpX18). Genes related to *i)* inflammation: SELE (E-selectin), OSMR (oncostatin M Receptor); *ii)* immune activation: CCR? (C-C motif chemokine receptor 7); *iii)* tissue stress and remodeling: TGFBI; and *iv)* neurological stress response: NPY (neuropeptide Y) are highlighted in the ISS condition. To reinforce our findings, we also compared, from the same flight, the ground MeCP2-KO brain organoids *vs.* ISS MeCP2-KO brain organoids using a bulk transcriptomics approach **(Fig. S4B).** In this evaluation, we highlight the following findings: *i)* upregulation of excitatory and inhibitory circuits in the brain: upregulation of NPAS4 (neuronal PAS domain-containing protein 4) in the ISS samples strongly suggests that the combined stress of the MeCP2-deficient background and spaceflight triggers a state of significant neurological distress or aberrant neuronal activity. This highlights the brain’s vulnerability in a system where a key epigenetic safeguard (MeCP2) is already absent; *ii)* activation of inflammatory and tissue remodeling: LRRC15 (leucine rich repeat containing 15) is implicated in tissue remodeling and its upregulation is a response to stress or injury and has been described with an essential role in response to viral infections, including as a key determinant of host’s susceptibility/resistance to SARS-CoV-2^17^, IL36B (interleukin 36 beta): a pro-inflammatory cytokine belonging to the IL-1 family, indicating an active inflammatory response, when compared to the ground samples.

**Fig. 4.**
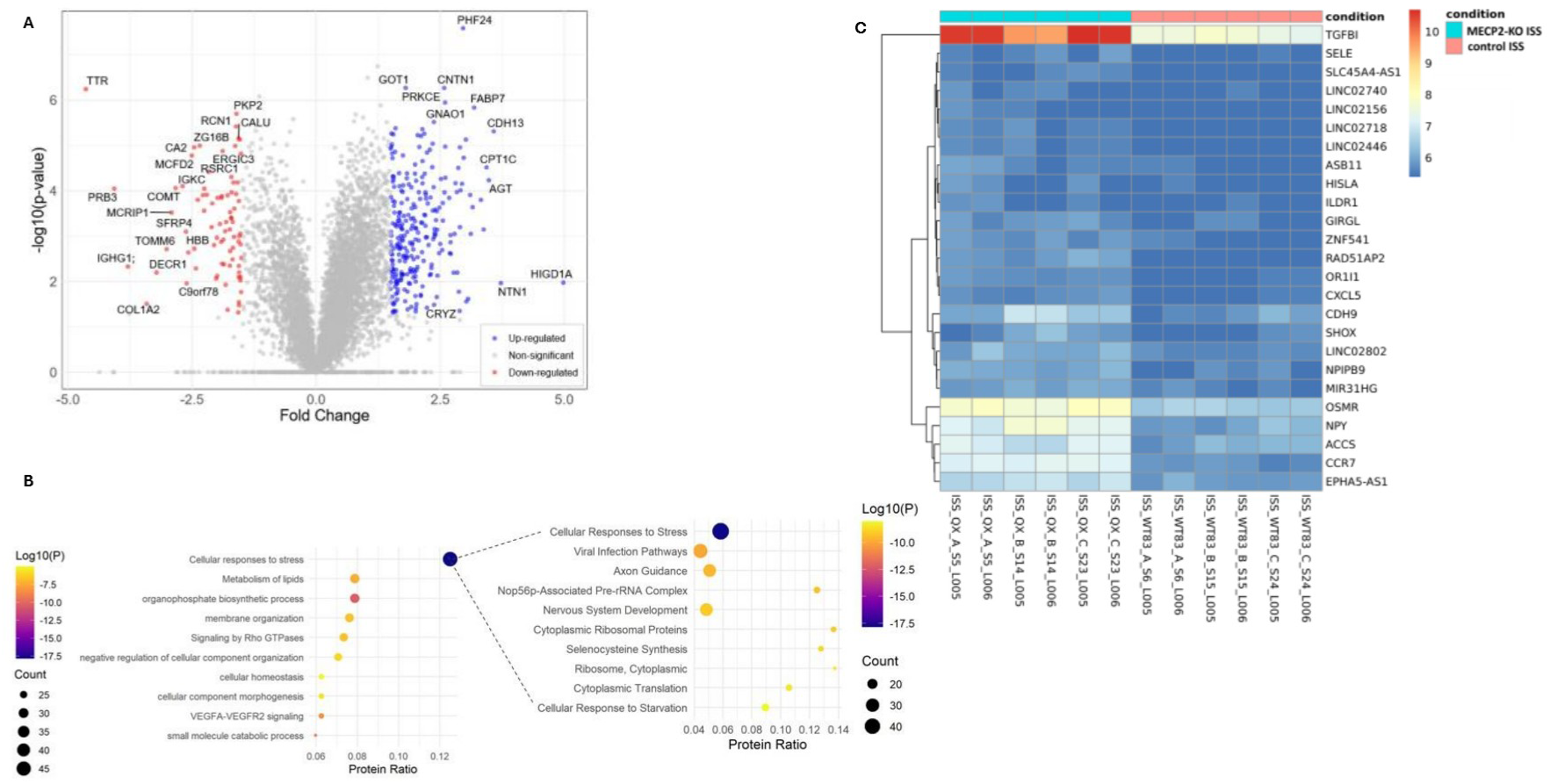
Proteomics and transcriptomics analysis overview of MECP2-KO brain organoids at the International Space Station (ISS). **(A)** Volcano plot displaying proteins upregulated in MECP2-KO cortical brain organoids (in blue) and downregulated (in red), when compare to control brain organoids, both at ISS. **(B)** Reactome pathway analysis from proteins upregulated in MECP2-KO brain organoids at ISS. This panel includes an overall pathway visualization followed by a focusing on the “cellular response to stress” pathway, highlighting the viral pathway_0_ **(C)** Heatmap from a subsequent spaceflight (SpX30) demonstrating a robust and consistent difference between the MECP2-KO brain organoids vs. healthy brain (control) organoids, both submitted to the space environment, at ISS. The hierarchical clustering indicates a distinct gene expression signature associated with the MeCP2-KO condition at ISS. Each experiment was done in biological duplicates. For proteomics bulk analysis: p≤0.01; fc≥1.2. Test T Student for comparisons. For pathway analysis: *Homo sapiens* as the reference organism, minimum overlap of 3 genes, enrichment factor> 1.5, and p < 0.01.

### Translating to Earth: accumulated L1 transcripts in astrocytes promote neurotoxicity

To translate the observations in space to Earth, we decided to conduct a series of groundwork studies in a clinically relevant human condition. A lack of functional MECP2 in the nervous system can cause Rett syndrome (RTT) and other neurodevelopmental conditions^19^. Accumulating L1-derived single-stranded DNA (ssDNA) in human astrocytes can trigger a neurotoxic inflammatory response in another neurodevelopmental disorder^12^. Using the MECP2-KO and control lines, we checked for L1 expression and found that endogenous L1 transcripts and L1-derived ssDNA are enriched in iPSC-derived astrocytes lacking MECP2 **(Fig. 5A, B** and **Fig. S5A).** Cytokines are important signaling molecules produced by glial cells and are responsible for cell maturation and inflammation during neurodevelopment^18^. We found several cytokines that were upregulated in astrocytes lacking MECP2 **(Fig. S5B),** including an increase in secreted IL6 **(Fig. 5C** and **Figs. S5C-E).** High levels of secreted astrocytic IL6 have recently been implicated in RTT, affecting synaptogenesis in a mouse model ^21^. We validated these observations in our human system, using astrocyte-conditioned media (ACM) and by adding recombinant human IL6 (rhll-6) to the culture media of iPSC-derived control neurons at the same concentration found in MECP2-KO astrocytes (2 ng/ml) for two weeks **(Fig. 5D** and **Figs. S5F, G).** Moreover, we also found a two-fold increase in the expression of the IL6 family cytokine receptor’s signal transducer, gp130 receptor (IL6ST) in postmortem cortical RTT samples compared with age-matched control samples **(Fig. S5E).** More importantly, postmortem cortical tissues showed a fivefold increase in MECP2-KO IL6 protein levels compared with controls, consistent with the in vitro observations **(Fig. S5C).** We observed a significant decrease in levels of synapsin1 puncta in control iPSC-derived neurons exposed to MECP2-KO ACM **(Fig. 5D).** These alterations were associated with functional synaptic defects in neurons exposed to conditioned media from MECP2-KO astrocytes. We also observed a significant reduction in synapsin1 puncta and total dendritic length in neurons when rhll-6 was added to the medium **(Figs. S5F-G).** Moreover, control neurons treated with rhll-6 that matured in multi-electrode array (MEA) plates decreased their spontaneous activity **(Fig. S5H).** Finally, we co-cultured independently generated human neurons and astrocytes. MECP2-KO astrocytes negatively affected the dendritic morphology of control neurons, reducing total dendritic length, the number of segments per neuron, and the spine density **(Figs. S5I-L).**

### L1 inhibition rescues MECP2-KO astrocytes defects

To confirm that the deleterious effect of MECP2-KO astrocytes is due to the accumulation of L1 elements, we treated our cultures with FDA-approved antiretroviral reverse transcriptase (RTi) drugs Lamivudine (3TC) and Stavudine (d4T), known to inhibit L1 reverse transcription^40^. We additionally used another antiretroviral drug, Nevirapine (NVP), as a negative control since no effect on L1 reverse transcription has been previously shown^40,41^. When MECP2-KO astrocytes were chronically treated with RTi, cytoplasmic L1 copies, and IL6 expression decreased to control levels (**Figs. 5B-C**). Similarly, we observed reductions in other pro-inflammatory cytokines and related factors, including CXCL10 (interferon-gamma-induced protein 10), CCL2 (monocyte chemoattractant protein 1), and TMEM173 (stimulator of interferon genes) **(Figs. S5M-0).** Previous studies have reported elevated levels of the excitatory neurotransmitter glutamate in the cerebrospinal fluid of patients with MECP2 mutations ^22,24^. MeCP2 has been shown to modulate glutamate clearance by regulating various astroglial genes^23^. Indeed, we found that several genes related to glutamate metabolism were misregulated in MECP2-KO astrocytes **(Fig. S5P).** MECP2-KO astrocytes treated with RTi restored glutamate, reactive oxygen species (ROS), and glutathione levels to control cells **(Figs. 5E-G).**

**Fig. 5.**
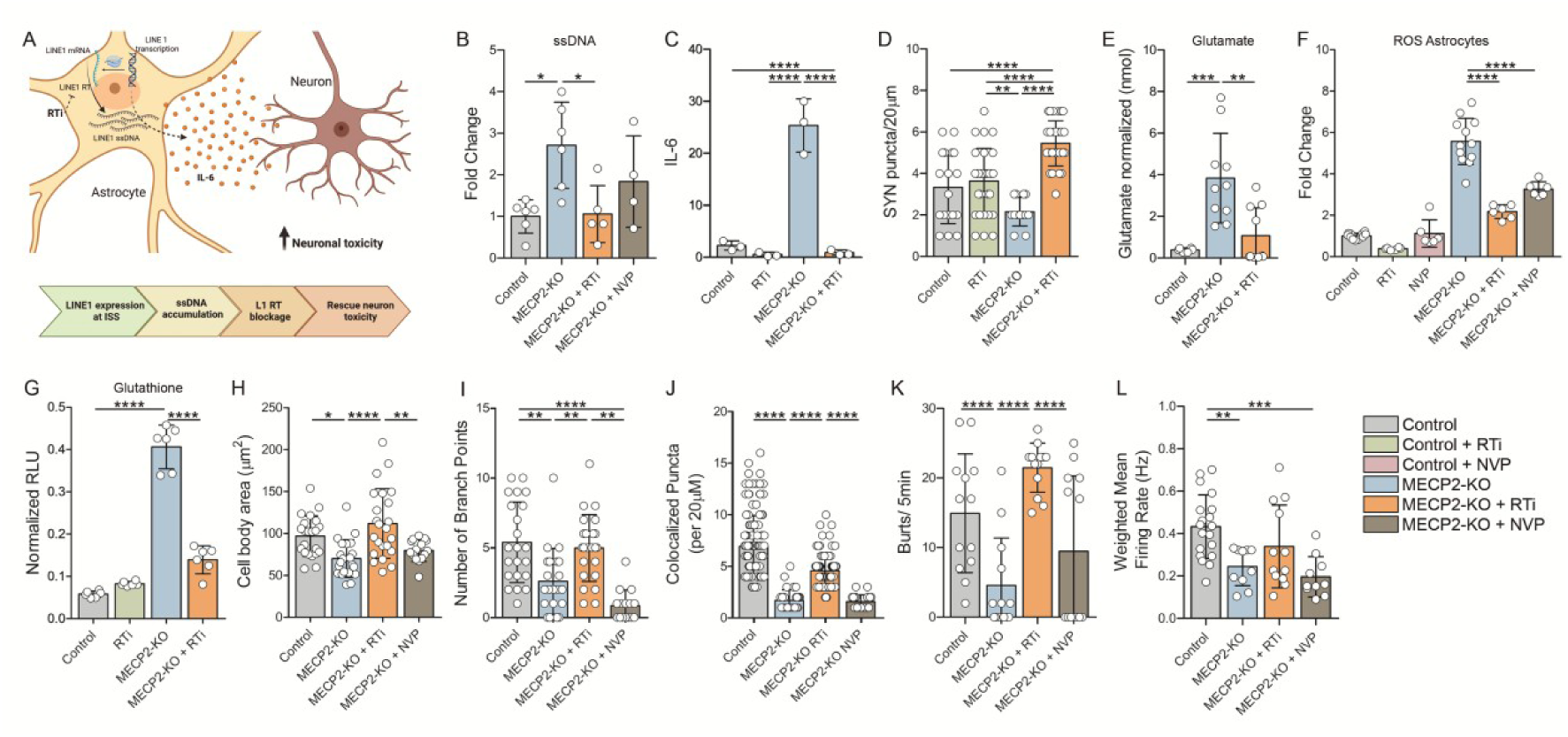
L1 expression in MECP2-KO astrocytes causes neurotoxicity that can be rescued with reverse transcriptase inhibitors (RTi), but not nevirapine (NVP). **(A)** Schematic figure on how ssDNA accumulates in the astrocytes, leading to increased neurotoxicity in neurons. **(B)** Quantification of extrachromosomal **L**1-derived single-stranded DNA (ssDNA) on MECP2-KO astrocytes. **(C)** Quantification of released IL-6 from astrocytes by enzyme-linked immunosorbent assay (ELISA). **(D)** Quantification of synaptic puncta per 20 µm on control and MECP2-KO neurons treated with astrocyte conditioned media (ACM). **(E)** Quantification of glutamate buffering in MECP2-KO astrocytes. Data are represented as mean ±SEM. Levels are normalized to control equaling one. **(F)** Quantification of reactive oxygen species (ROS) levels in MECP2-KO astrocytes. Data are represented as mean ±SEM. Levels are normalized to control without treatment equaling one. **(G)** Quantification of intracellular glutathione in MECP2-KO astrocytes. Data are represented as mean ±SEM. Levels are normalized to control without treatment equaling one. **(H)** Quantification of neuronal cell body area. **(I)** Quantification of neuronal dendritic complexity by number of branch points. **(J)** Quantification of co-localized synaptic puncta (defined by SYN1+/PSD95+ on MAP2+ dendrite) per 20 µm on neurons from each condition. **(K)** Quantification of the total number of spikes of control neurons matured on MEA plates. **(L)** Quantification of weighted mean firing rate from organoids in each condition. Reverse-transcriptase inhibitor (RTi) or nevirapine (NVP) treatment is indicated. Each color represents one condition: control (black), MECP2-KO (blue), MECP2-KO+RTi (orange), and MECP2-KO+NVP (green). Each experiment was done in biological triplicates. The number of measurements (n) for these experiments is indicated directly by the symbols (o) on each bar graph. Data are represented as mean ±SEM. *p<0.5* (*), p<0.01 (**), p<0.001 (***) and p<0.0001 (****); ordinary one-way ANOVA.

### L1 inhibition rescues MECP2-KO neuronal defects in mixed cultures

We also tested if RTi could restore neuronal function in mixed brain organoid cultures containing MECP2-KO neurons and astrocytes. Treating MECP2-KO-derived cultures with RTi, rescued neuronal cell body area and dendritic complexity **(Figs. 5H-I** and **Figs. S5Q-S).** To determine if the control of L1 activity by RTi can ameliorate synapse formation in MECP2-KO neurons, co-localization of SYN1 (synapsin I) and PSD95 (postsynaptic density protein 95) on a MAP2 (microtubule-associated protein 2) positive neurite was quantified. MECP2-KO neurons in the RTi-treated culture displayed a significant increase in co-localized puncta compared to those without RTi-treatment, which is not seen in the NVP-treated culture **(Fig. 5J).** At the functional level, neuronal network activity measured by the number of bursts and firing rate in MECP2-KO-derived organoids was also rescued in cultures treated with RTi (**Figs. 5K-L**).

### L1 reverse transcriptase inhibition rescued survival and behavior in a Mecp2-KO mouse model

Hemizygous Mecp2-KO males have a median lifespan of approximately 10 weeks and exhibit pronounced behavioral and physiological abnormalities ^25,29,30,42^. We treated these animals with 3TC or d4T in the drinking water (physiological dose of 2 mg/ml^26^ starting at 4 weeks old. Both 3TC and d4T ameliorated the overall aspect of the Mecp2-KO animals and extended their lifespan (**Fig. 6A-B**). Next, we tested whether any behavior would be rescued following the animals’ extended lifespan. The open-field test revealed that 3TC treatment improves the mobility of Mecp2-KO mice (**Figs. 6C-D** and **Fig. S6A).** Previous studies have demonstrated that this Mecp2-KO mouse model exhibits decreased anxiety behavior ^27^. We found no effect of 3TC treatment on the anxiety-related behavior of Mecp2-KO mice (**Fig. 6E** and **Fig. S6B, Supp. Videos 1-3).** There are conflicting results regarding the sociability of these rodent models ^31,33^. In our hands, Mecp2-KO mice have impaired social memory, and this phenotype can be rescued with 3TC treatment (**Fig. 6F**). Following behavioral analysis, the animals’ brains were fixed and subjected to Golgi staining for morphological analysis. We focus on the prefrontal cortex, as this region is critical for social behavior. We found that the 3TC treatment rescued neuronal soma size and spine density of Mecp2-KO neurons in the cingulate cortex (**Fig. 6G-H** and **Fig. S6C-D).**

**Fig. 6.**
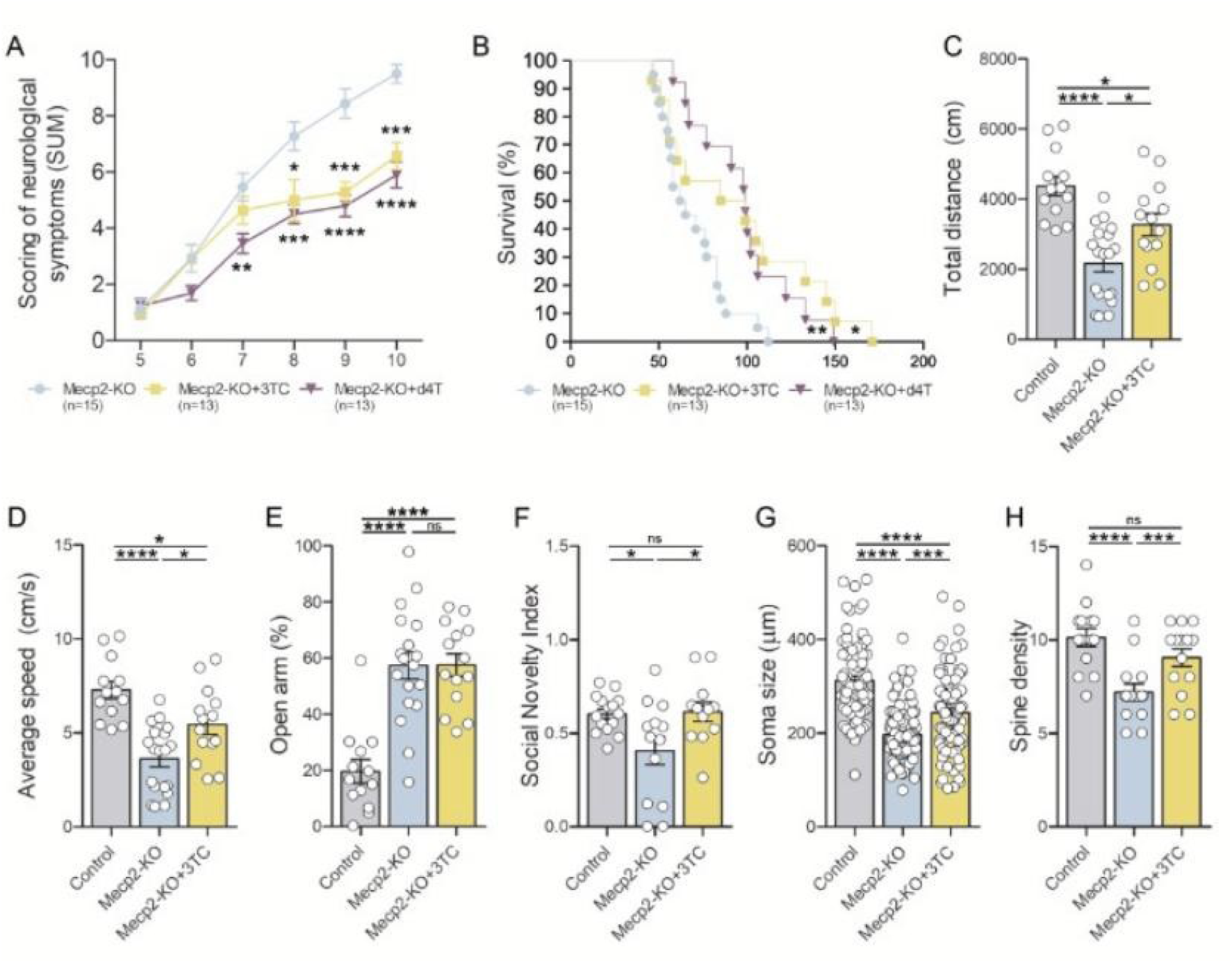
Treatment of Mecp2-KO with reverse transcriptase inhibitors (RTis) in a mouse model. **(A)** Time plot of scores from a neurological symptoms assessment of Mecp2-KO mice with and without treatment of RTis lamivudine (3TC) and stavudine (d4T) at 2 mg/ml. The time plot of neurological symptoms scores depicts 5 weeks after RTi treatment was introduced in drinking water (weeks 5-10). **(B)** Time plot of percent survival depicts mice’s lifespan in a number of days. **(C)** Quantification of total distance traveled and average speed of WT, Mecp2-KO, and 3TC-treated Mecp2-KO mice in the open-field test. **(D)** Quantification of average speed (cm/seconds) by WT, Mecp2-KO, and 3TC-treated Mecp2-KO mice in an open arm chamber during the elevated plus maze test. **(E)** Quantification of time spent (in percentage of total time) by WT, Mecp2-KO, and 3TC-treated Mecp2-KO mice in an open arm chamber during the elevated plus maze test. **(F)** Quantification of social novelty index of WT, Mecp2-KO, and 3TC-treated Mecp2-KO mice in a three-chamber test. Social novelty index is measured as time spent in the chamber with an intruder divided by the total time spent in both chambers. **(G)** Quantification of soma size and spine density. **(H)** Quantification of neurons from the cingulate cortex in brains fixed from WT, Mecp2-KO, and 3TC-treated Mecp2-KO mice. Spine density is measured by number of spines per 10 µm of dendrite. The number of measurements *(n)* for these experiments is indicated directly by the symbols (o) on each bar graph. Data are represented as mean ±SEM. *p<0.5* (*), p<0.01 (**), p<0.001 (***) and p<0.0001 (****); ordinary one-way ANOVA.

### L1 reverse transcriptase inhibition recovered abnormal gene expression in both astrocytes and neurons in the brains of Mecp2-KO mice

Given that behavioral impairments and aberrant neuronal morphologies in Mecp2-KO mice were restored by 3TC treatment, we next examined whether these restorations are also detected at the molecular level. To this end, we conducted spatial transcriptomics analysis (Xenium, 5k) on forebrain sections from 8-9-week-old WT, Mecp2-KO, and Mecp2-KO mice treated with 3TC. We applied unsupervised clustering on the combined dataset of 270,741 cells that passed quality control to define cell clusters. We identified 10 unique clusters and annotated each cluster (excitatory neurons, inhibitory neurons, astrocytes, oligodendrocyte progenitor cells (OPC), oligodendrocytes, microglia, endothelial cells, pericytes, fibroblasts, smooth muscle cells) with the expression profiles of canonical cell type marker genes and spatial information (**Fig. 7A-C**). Since we have demonstrated that astrocytes play critical roles in neuronal morphology and function (**Figs. 5 and S5),** we focused on astrocytes and reanalyzed the astrocyte cluster using a different clustering method. We found that the cell number and spatial distribution of each astrocyte subcluster in the forebrain were not substantially altered in Mecp2-KO compared to WT (**Figure 7D-E**). However, a comparison of gene expression across the entire astrocyte cluster between WT and Mecp2-KO identified 173 genes as significantly up-regulated and 173 as significantly down-regulated under Mecp2-KO conditions **(Table S6).** Gene ontology (GO) analysis of the upregulated genes revealed enrichment of terms related to oxidative stress and interleukin signal (**Fig. 7F**). Intriguingly, significant overlap was observed between the genes upregulated in these Mecp2-KO astrocytes and the 30 genes included in the gene list (5006 genes) in the Xenium panel of 150 genes **(Table S4)** that were upregulated under ISS conditions (**Fig. 7G**). Among the 4 overlapping genes, ALDOA (aldolase A), an enzyme involved in cellular metabolism and previously implicated in inflammation ^32^ and DNA damage responses ^43^, was significantly downregulated in Mecp2-KO mouse brains following 3TC treatment. Furthermore, we found that 3TC treatment restored the impaired expression of inflammation-related genes implicated in these GO terms (**Fig. 7H**). In addition, consistent with these transcriptomic findings, immunostaining results showed that complement component 3 (C3), a key immune marker associated with reactive astrocytes, was upregulated in Mecp2-KO astrocytes and reduced by 3TC treatment (**Fig. 7I-J**).

**Fig. 7.**
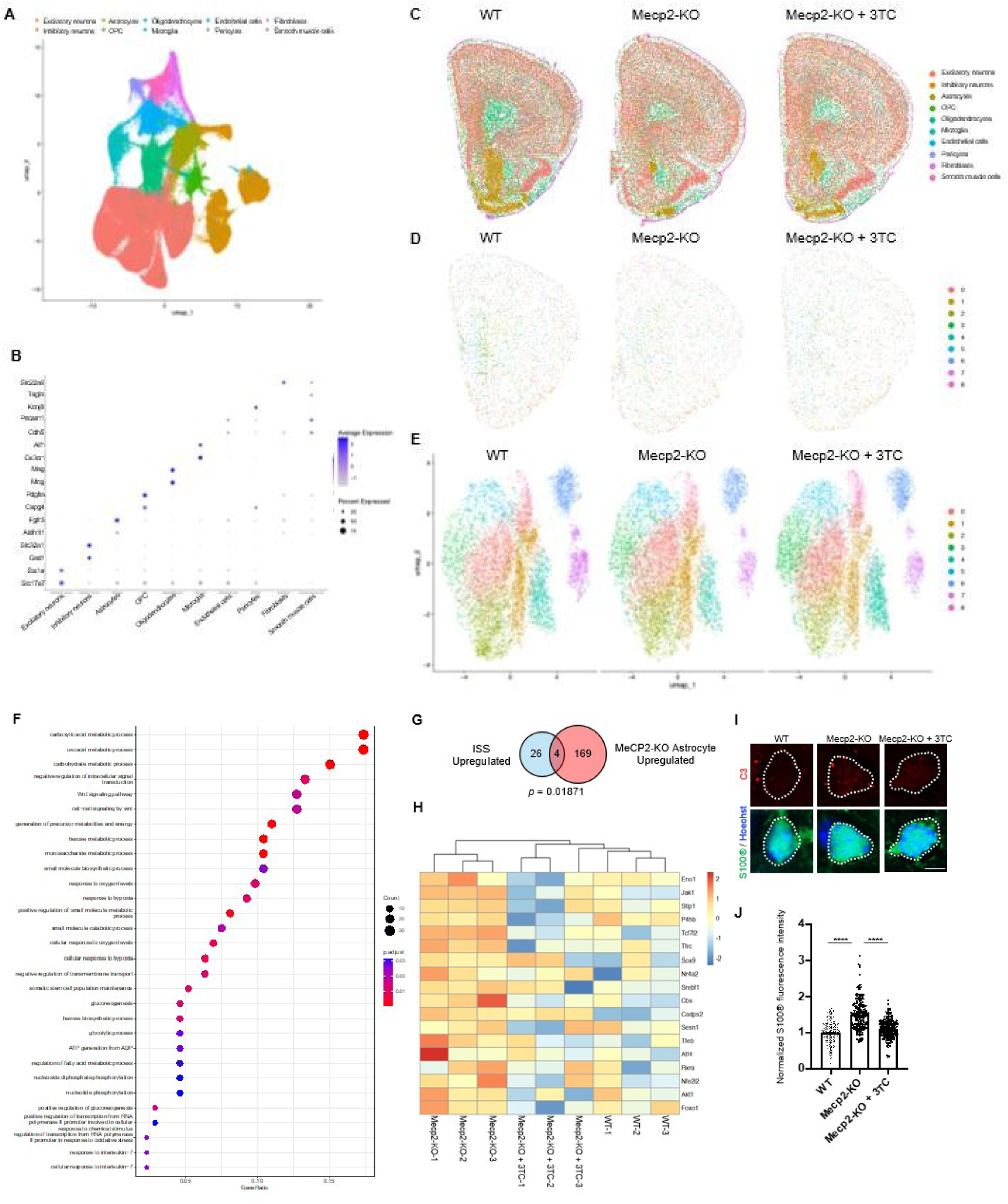
Spatial transcriptomics analysis of WT, Mecp2-KO and 3TC-treated Mecp2-KO astrocytes. **(A)** UMAP plot showing 270741 cells annotated into 10 clusters from the brains, including PFC region, of WT (n=3), Mecp2-KO (n=3) and 3TC-treated Mecp2-KO (n=3) mice. **(B)** Scaled dot plot of canonical cell type marker genes used to define each cluster. **(C)** Spatial representative image plots of cell types in (A) across coronal hemisection of the WT, Mecp2-KO and 3TC-treated Mecp2-KO mice forebrains. **(D)** Spatial representative image plots of astrocyte subtypes in coronal hemisections of WT, Mecp2-KO and 3TC-treated Mecp2-KO mouse forebrains. **(E)** UMAP analysis comparing astrocyte subtypes across WT, Mecp2-KO and 3TC-treated Mecp2-KO mouse forebrains. **(F)** GO analysis of upregulated genes in Mecp2-KO astrocytes compared to WT astrocytes. (G) Overlap between ISS upregulated genes and Mecp2-KO astrocyte upregulated genes. **(H)** Heatmap of differentially expressed response to interleukin-7 (GO:0098760) and cellular response to external stimulus (GO:0071496) genes that were upregulated in Mecp2-ko astrocytes compared to WT astrocytes and downregulated in 3TC- treated Mecp2-KO astrocytes. *(I)* lmmunofluorescence staining for C3 (red) and S100β (green) in the PFC regions of 8-week-old WT, Mecp2-KO and 3TC-treated Mecp2-KO mouse brains. Nuclei are counterstained with Hoechst (blue). Scale bar, 5 µm. **(J)** Quantification of C3 signal intensity within the soma of S100β+ astrocytes. The number of individual measurements (n) for each experimental group is indicated by the symbols (o) on the bar graph. Data are presented as mean ± SEM. ****p<0.001 by ordinary one-way ANOVA.

Based on the recovery of gene expression in astrocytes, we next compared gene expression in clusters of excitatory neurons in WT, Mecp2-KO, and Mecp2-KO mice treated with 3TC and performed GO analysis. We found that downregulated genes in Mecp2-KO relative to WT were enriched for GO terms associated with synaptic signaling **(Fig. S7A).** In addition, genes upregulated with 3TC treatment in Mecp2-KO mice were enriched in similar GO categories **(Fig. S7B).** Moreover, when we examined genes in a specific GO term (synaptic signaling), expression of those genes was reduced in Mecp2-KO but recovered with 3TC treatment **(Fig. S7C),** suggesting that 3TC treatment recovered impaired gene expression not only in astrocytes but also in neurons.

## DISCUSSION

The modern cell has evolved to coexist with autonomous L1 sequences by activating defense mechanisms within the cell’s intrinsic immune system ^44^. Moreover, DNA hypermethylation responses were significantly pronounced in evolutionarily younger retrotransposons, suggesting that recent genomic insertions might be particularly sensitive indicators of space exposure^11^. By employing a multi-omics approach that integrates proteomics, bulk transcriptomics, and single-nucleus transcriptomics, we provided a comprehensive view of the molecular alterations that affect human brain cells across space. This led us to reveal a complex interplay between viral pathways, astrocyte reactivity, and inflammatory responses, a novel phenomenon that we call Space-Induced Neural Senescence (SINS).

Multiple studies on neurological and psychiatric disorders have revealed increased L1 expression in the brain ^39^. We previously reported that the accumulation of L1-derived ssDNA in human iPSC-derived astrocytes triggered pathological inflammation in a severe progeroid neurological condition, Aicardi-Goutieres Syndrome (AGS) ^12^. There is substantial evidence that transposable elements, particularly L1 retrotransposons, are aberrantly reactivated in the aging brain and across various neurodegenerative diseases, including Amyotrophic Lateral Sclerosis (ALS), Alzheimer’s Disease (AD), Parkinson’s Disease (PD), and Huntington’s Disease^45–49^. Our findings demonstrated that the histone proteins and deacetylation processes are altered by the space environment, likely contributing to an “epigenetic rewiring” and the activation of viral pathways in space. Histone downregulation is frequently observed under conditions of cellular stress or in disease states, indicating alterations in chromatin dynamics that affect the accessibility and stability of genomic DNA ^50^. Reduced histone expression impairs nucleosome assembly, leading to chromatin relaxation and aberrant activation of otherwise silenced genomic regions, such as transposable elements^51^. This chromatin disruption often results in compromised epigenetic regulation, enabling the transcriptional activation of repetitive sequences, including L1s. Histone depletion thus facilitates L1 mobilization, reflecting a critical link between chromatin remodeling and potential genome instability^52^. The activation of L1s following histone downregulation has profound biological implications, particularly in the context of aging and neurodegeneration. Aberrant L1 activation is closely associated with age-related neurodegenerative diseases such as Alzheimer’s disease, where increased retrotransposition events correlate with neuronal loss and cognitive decline^53^. It is interesting to note that our re-analyses of the blood transcriptomic data from the NASA Twins Project ^54^ have also revealed an increase in viral molecular pathways, suggesting that the reactivation of endogenous retroviruses is likely not specific to brain tissue. However, due to the brain’s limited regenerative capacity, the impact of this phenomenon is expected to be more pronounced. Our findings are in agreement with the space medicine literature^7,55–57^ that dysregulation in OxPHOS proteins corroborates the idea that mitochondrial impairment is a shared pathological mechanism underlying diverse neurodegenerative processes, including Parkinson’s disease (PD), Huntington’s, and Alzheimer’s diseases^58,59^. Also, corroborated with the genomic findings from the lnspiration4 mission, which also evaluated the OxPHOS genomic framework, revealing a sustained downregulation of PD-associated OxPHOS genes in astronauts’ peripheral blood mononuclear cells (PBMCs)^60,61^.

We and others have previously revealed that brain tissue isolated from RTT patients is more susceptible to L1 expression^14,16^. Here, we found that the accumulation of L1 transcripts in the absence of MeCP2 induces IL6 release by astrocytes. Several groups have previously reported interleukin abnormalities and cytokine release in RTT ^18,21,62,63^. However, in the absence of a stimulating agent, this observation was not linked to a potential disease contribution. Activation of transposable elements exacerbates aging and neurodegeneration ^45,64,65^, promoting genomic instability, neuroinflammation, and aberrant immune signaling, as observed in our experiments in LEO. Moreover, recent studies have revealed a significant upregulation of transposable elements, including L1s, in excitatory neurons and astrocytes in AD brains^66,67^, demonstrating that an increase in TE expression in AD is related to immune activation, synaptic dysfunction, and cellular senescence, similar to the findings associated to the SINS phenomenon, described in this study.

Thus, we hypothesized that the cytokine abnormalities in RTT are not a direct consequence of MECP2 mutations, but rather a pathological condition exacerbated by the accumulation of cytoplasmic L1 copies in astrocytes, which triggers the IL6-mediated neurotoxic response. Our findings linked L1 expression in RTT astrocytes to disease phenotypes, thereby enabling us to test whether L1 inhibition by RTi could reduce inflammation and mitigate neurotoxicity. The analyses from independent spaceflights corroborate that, in LEO, brain organoids activate sterile-mediated cellular responses, resulting in the activation of an immune response and epigenetic derangement that can activate viral pathways, inducing SINS, and immune dysfunction, offering a new framework for understanding how “dormant” genomic elements become pathogenic under certain conditions. Notably, increased levels of IL6 are also associated with neuropsychiatric conditions such as depression, Alzheimer’s Disease, and aging ^68,69^. Thus, we compared *in vitro* and *in vivo* non-treated and RTi-treated models in MECP2-KO brain organoids and Mecp2-KO mice, respectively. Our findings are summarized in **Fig. 8**. *In vitro,* the loss of MeCP2 results in increased transcription of L1 elements, leading to elevated inflammation, characterized by increased levels of IL6, ROS, glutamate, and inflammatory markers such as CXCL10, CCL2, and TMEM173. The molecular findings in the secretome panel and mitochondrial framework analysis indicate that these signatures are consistent with known space-environment stressors, including activation of the viral pathway, leading to oxidative stress and impairment of mitochondrial homeostasis. Importantly, they also align with recent literature describing spaceflight-induced activation of retrotransposons in mice ^10^ and the cellular senescence pathway in hematopoietic stem cells ^70–72^. These findings collectively suggest a potentially space-environment-sensitive state, characterized by shifts in differentiation toward an inflammatory and viral-responsive pattern of transcription and translation.

**Fig. 8.**
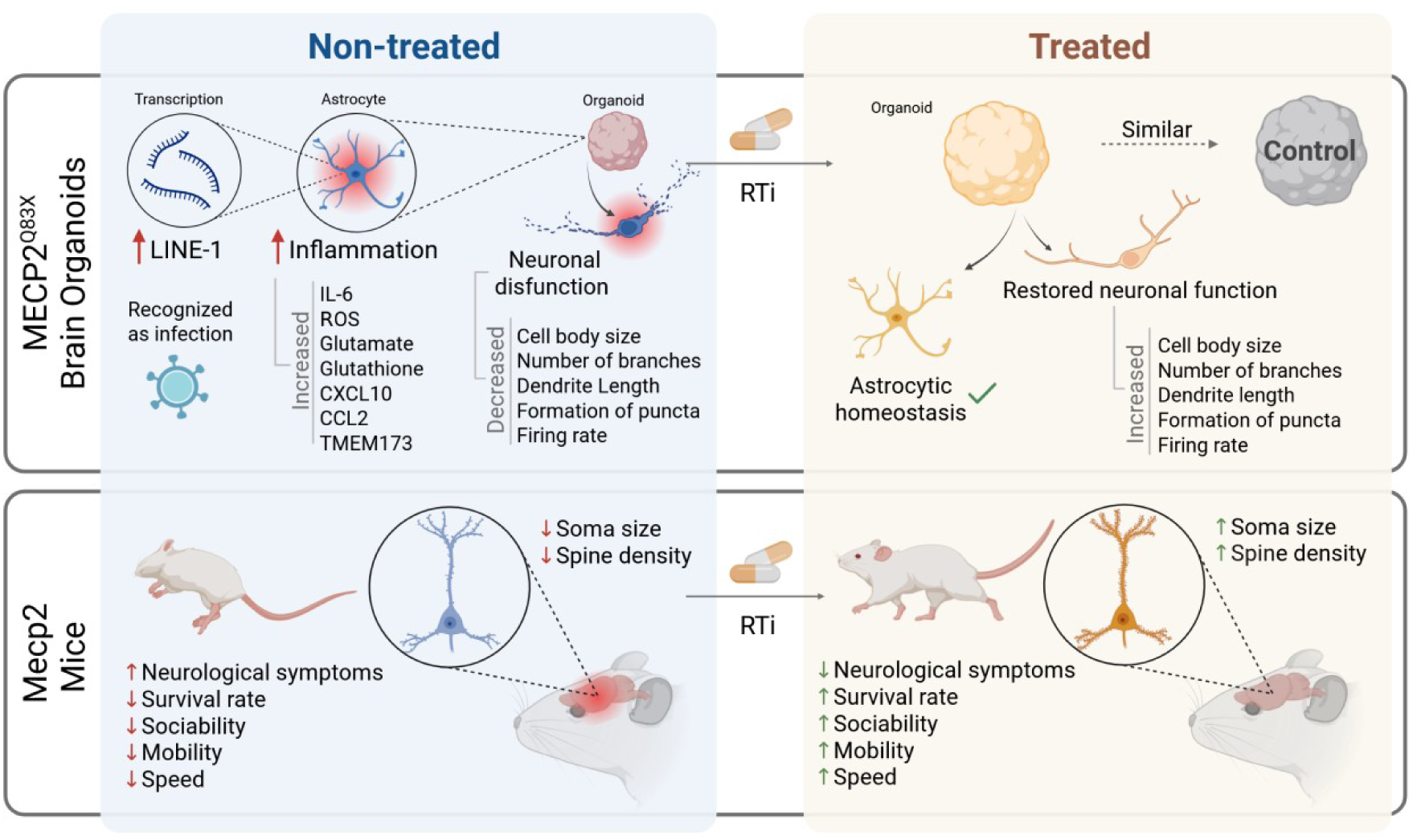
Summary comparing non-treated and RTi-treated models in MECP2-KO brain organoids and Mecp2 mice. Top Panel: Brain Organoids (MeCP2-KO). The left side depicts non-treated MeCP2-KO brain organoids, where the loss of MeCP2 results in increased transcription of LINE-1 elements, which are recognized as infection triggers. This leads to elevated inflammation, characterized by increased levels of IL-6, ROS, glutamate, and inflammatory markers such as CXCL10, CCL2, and TMEM173. In parallel, neuronal dysfunction is observed, including reductions in cell body size, dendritic branching, dendrite length, the formation of synaptic puncta, and firing rates. The right side represents organoids treated with RTi (reverse transcriptase inhibitor). Treatment restores neuronal function and promotes astrocytic homeostasis. Neuronal characteristics, such as cell body size, number of dendritic branches, dendrite length, synaptic puncta formation, and firing rates, are restored to levels comparable to the control condition, indicating a significant recovery in cellular and synaptic function. **Bottom Panel: Mecp2-KO Mice.** The left side shows non-treated Mecp2 mice, where a decrease in soma size and spine density is observed, accompanied by severe neurological symptoms, including reduced survival rate, sociability, mobility, and speed. The right side illustrates the effects of RTi treatment in Mecp2 mice. Treatment leads to increased soma size and spine density, accompanied by improvements in neurological symptoms. Treated mice show enhanced survival rate, sociability, mobility, and speed compared to non-treated controls.

MECP2-KO brain organoids treated with RTi restore neuronal function, promoting astrocytic homeostasis and indicating a significant recovery in cellular and synaptic function. *In vivo,* untreated Mecp2-KO mice exhibited severe neurological deficits, including reduced survival, sociability, mobility, and speed. When Mecp2-KO mice were subjected to RTi treatment, we observed improved neurological symptoms and increased survival, as well as increased sociability, mobility, and speed, compared with untreated controls. Furthermore, we confirmed these therapeutic effects also at the molecular level. We demonstrated that RTi treatment suppresses cytokine responses and reduces inflammatory markers typically associated with senescent cells and aged mice. These effects underscore a causal link between retroelement activation and space-related inflammation, as observed in our experiments on the ISS during independent spaceflights.

Radiation and microgravity stressors do not act in isolation but rather appear to have additive or even synergistic effects on brain function. Since we cannot isolate the various factors acting at the ISS, we refer to the combined effects of cosmic radiation, microgravity, and other unmeasured factors as the “space environment”. Due to its nature, space radiation - especially GCR (galactic cosmic radiation) - plays a crucial role in biological systems. Rodents subjected to a complex 33-beam GCR simulation exhibited deficits in attentional processing, along with disrupted dopaminergic neurotransmission in the prefrontal cortex, despite maintaining reward-related behaviors ^73^, illustrating that cosmic radiation inflicts structural and functional damage on neural circuits. In this study, the radiation measured in all experiments at ISS, specifically the GCR dose recorded, averaged 0.119 mGy/day (absorbed dose) and 0.126 mGy/day (dose equivalent). In contrast, the South Atlantic Anomaly (SAA) doses were about 0.084 mGy/day (absorbed dose) and 0.086 mGy/day (dose equivalent). Given these values, the recorded doses from our spaceflights fall within the lower end of the expected radiation exposure range for ISS missions. For shorter missions, such as typical CRS resupply flights, the biological impact is likely negligible, as the total absorbed dose remains low and below critical thresholds that can trigger acute health effects. Moreover, we have previously investigated the role of radiation in brain organoids submitted to proton radiation (0.5 and 2 Gy)^74^. We found dose- and time-dependent increases in DNA damage, oxidative stress, and activation of DNA damage checkpoint genes, particularly at 48 hours post-irradiation. Other significant findings included declines in mitochondrial function and neural identity, corroborating the results of this study; however, we acknowledge the limitations of attributing the phenotypic findings solely to this stressor.

The collective findings on the secretome panel suggest that the space environment may be triggering a “recalibration” of the brain organoids’ developmental program, causing these models to strengthen their cellular defenses to address the unique challenges of space. This may serve as a mechanism to achieve a more resilient, faster-differentiated state, leading to a SINS state. The combined findings of the three spaceflights validate the results described in this study, demonstrating that the space environment leads to biological alterations in the brain organoids model, resulting in a phenotype that displays profound cellular stress, impaired genome regulation, and inflammation, potentially mediated by cGAS/STING pathway, a mechanism previously described by our group^12^, and which is activated by presence of viral DNA^75^ and retroelements^75,76^, especially in neurodegenerative diseases^77,78^. It is well established in the literature that transposable element activation is not merely a bystander effect in aging and senescence but a potentially causal mechanism in disease progression, contributing to cell-to-cell variability and neuronal vulnerability ^79,80^.

The successful preclinical data highlight RTi’s immediate utility in therapeutic regimens for **RTT** and other neuroinflammatory disorders, revealing a potential novel neuroprotective treatment for astronauts on long spaceflight missions as a translational venue. The proven safety and tolerability profile of 3TC makes it an attractive candidate for drug repurposing strategies aimed at preventive treatment in astronauts, particularly given the altered immune function and potential viral reactivation that can occur during extended space missions ^81,82^. Prophylactic and therapeutic administration of 3TC drugs could theoretically mitigate latent retroviral or endogenous retroviral elements whose activation may be potentiated by the stress-induced immunosuppression associated with spaceflight^83^, and its known efficacy against viral replication during spaceflight has already been documented in the literature^84^. The broad anti-inflammatory and immunomodulatory effects of RTis, particularly their capacity to mitigate inflammation driven by endogenous retroviruses, offer a novel and powerful mechanism to counter the pervasive immune dysregulation observed in microgravity.

Our study’s findings align with recent astronaut ^60,85^ and mouse ^20^ data on the space environment, underscoring the critical need for a comprehensive approach to protecting astronaut health during long-duration space missions. This could include implementing antiviral and anti-inflammatory strategies to ensure the well-being of future generations of explorers.

## LIMITATIONS OF THE STUDY

Limitations of the study are attributable to inherent difficulties in the space environment and in LEO. The experiments are extremely precious and constrained by the payload capacity of each spaceflight, making it difficult and costly to send multiple biological replicates on a single flight.

## RESOURCE AVAILABILITY

### Lead Contact

Requests for further information and resources should be directed to and will be fulfilled by the lead contact, Dr. Alysson R. Muotri.

### Materials Availability

This study did not generate new unique reagents or materials

### Data and code availability

- The mass spectrometry proteomics data have been deposited to the ProteomeXchange Consortium via the PRIDE (https://www.ebi.ac.uk/pride/) partner repository with the dataset identifier PXD055001 and also in the NASA Genelab repository, https://osdr.nasa.gov/bio/submission/rsa/RDSA-939 and are publicly available as of the date of publication.
- The transcriptomics data is deposit in the NASA Genelab repository under the study number OSD-765 and also in the Zenda (10.5281/zenodo.16388786 and 10.5281/zenodo.16389553) and are publicly available as of the date of publication.
- This paper does not report original code.
- Any additional information required to reanalyze the data reported in this paper is available from the lead contact upon request.

## ACKNOWLEDGMENTS

We would like to thank the invaluable efforts of the ISSCOR team: Natalia Chermont, Isabelle Luz, Paulo Carvalho, Juliana Fisher, and Blake Tsu. Roberto Herai, Luiz Pedro Petroski, and Adriano Ferrasa for all the insights on RNAseq data. We acknowledge the invaluable efforts of Space Tango, our implementation partner, in providing telematric data from all missions. We acknowledge the Space Radiation Analysis Group (SRAG) for providing the radiation data used in this work. We acknowledge Daniel McClatchy and Claire Delahunty from Yates’s lab for their experimental advice, Richard Waite, Narayanaganesh “Ganesh” Balasubramanian, Leslie Sandoval, Michael Krawitzky, and Matt Willetts from Bruker for their support in this work. This work was supported by grants from the NSF/CASIS grant (CBET 2126309) and NASA (80JSC018M0005) to A.R.M.; AMED (JP20gm1310008, JP24wm0625306), JSPS KAKENHI (JP23H00391) and an Intramural Research Grant (3-9) for Neurological and Psychiatric Disorders of NCNP Grant to K.N.; JSPS KAKENHI (JP22K15201, JP23H04170, JP25H01866) and AMED (23gm1310008) to H.N. This research was also supported by NPO the Rett Syndrome Support Organization to H.N.(2019-2-10) and an NIH grant (#S10 OD026929) from Dr. K. Jepsen at the UCSD Institute of Genomic Medicine. A.R.M., H.N., and K.N. are inventors on a pending U.S. patent application for the use of reverse transcriptase inhibitors for Rett syndrome (U.S. Application No. US18/569,721).

## AUTHOR CONTRIBUTIONS

Conceptualization, A.M., H.N., A.M., P.M., E.V., K.N., A.R.M.; Methodology, A.M., K.N., J.Y., A.M., P.M., A.R.M.; Investigation, A.M., H.N., A.M., T.U., S.M., P.M., L.L., L.C., E.V., N.S., A.S., A.S., B.F., A.L., D.A., M.I., J.Y.,T.O., M.H., K.N., A.R.M. Visualization, A.M., P.M., L.L., L.C., A.S., K.N., A.R.M. Funding acquisition, K.M., H.N., A.R.M.; Project administration, K.N., A.R.M.; Supervision, K.N., A.R.M. Writing - original draft, A.R.M.; Writing - review & editing, A.M., H.N., A.M., T.U., P.M., L.L., L.C., E.V., N.S., A.S., A.S., B.F., A.L., D.A., M.I., J.Y., K.N., A.R.M.

## SUPPLEMENTAL INFORMATION

Figures S1-S7, Tables S1-6, Supplementary Videos 1-3.

## SUPPLEMENTAL FIGURE LEGENDS

**Fig. S1.**
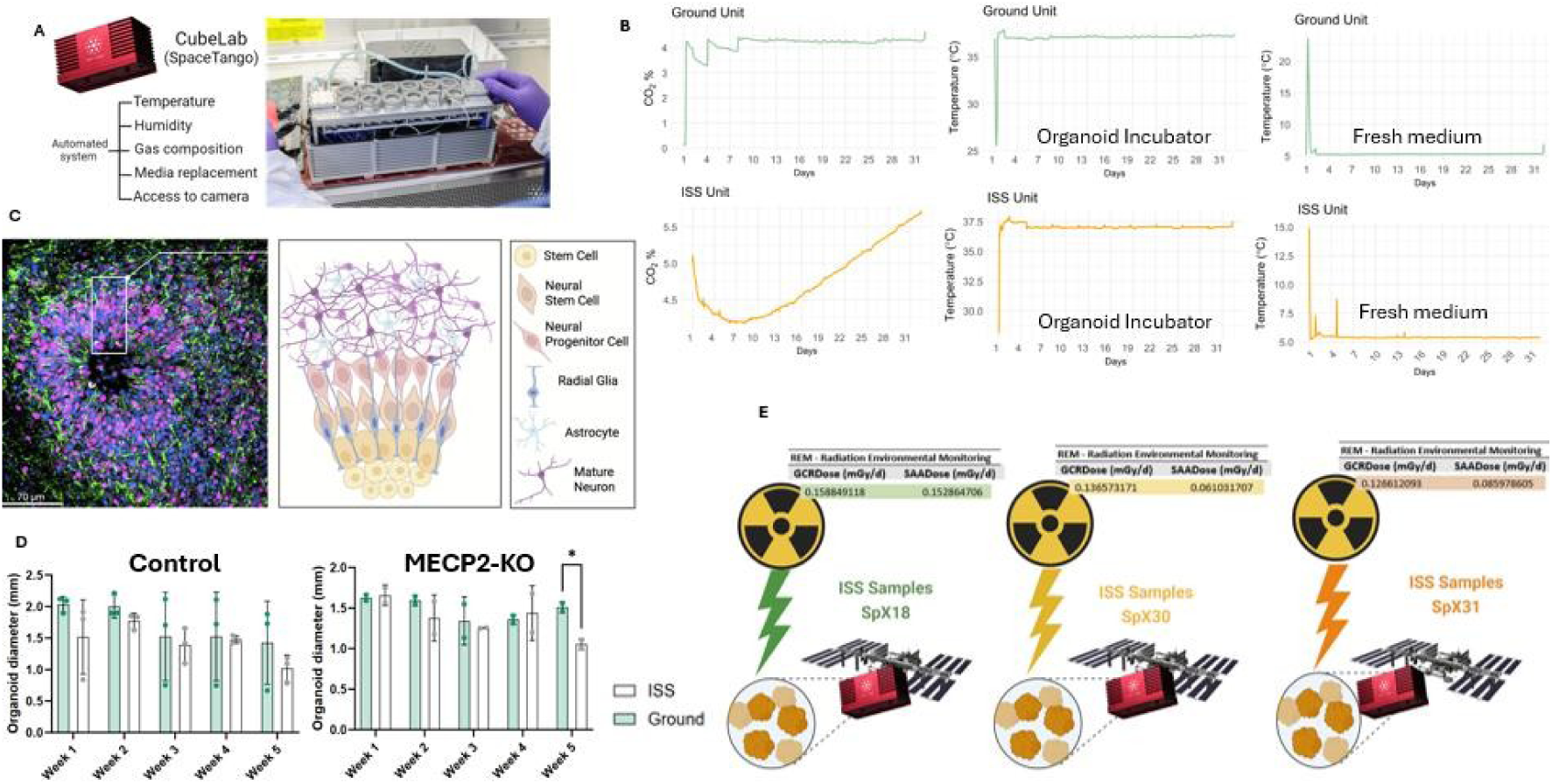
CubeLab and environmental conditions and functionality. **(A)** Illustration representation of the Cubelab capabilities monitored during the space environment experiment on the International Space Station (ISS). The outer view and internal setup of the Cubelab, which houses the wells containing the organoids, are depicted, as well the organoids inside the well in the ISS. The Cubelab is designed to control temperature, humidity, and gas composition, automatically exchanging the cell medium. Additionally, the Cubelab is equipped with cameras that allow real-time observation of the organoids from Earth during spaceflight. **(B)** Environmental parameters monitored during the space environment experiment on the International Space Station (ISS) compared to ground control. The X-axis represents the days of the mission, while the Y-axis shows values for CO_2_ concentration, organoid incubator temperature and fresh culture medium temperature, in both environments. The CO_2_ concentration was maintained at around 5%; the organoid incubator temperature was kept around 37°C; and the fresh culture medium temperature at 4°C until it was used to feed the organoids every two days. Duplicate measurements were taken for each temperature to ensure accurate readings. **(C)** Schematic view of the brain organoid used in this work. On the left, a confocal image of a human cortical organoid stained for stem cell marker SOX2 and neural progenitor marker nestin.

**Fig. S2.**
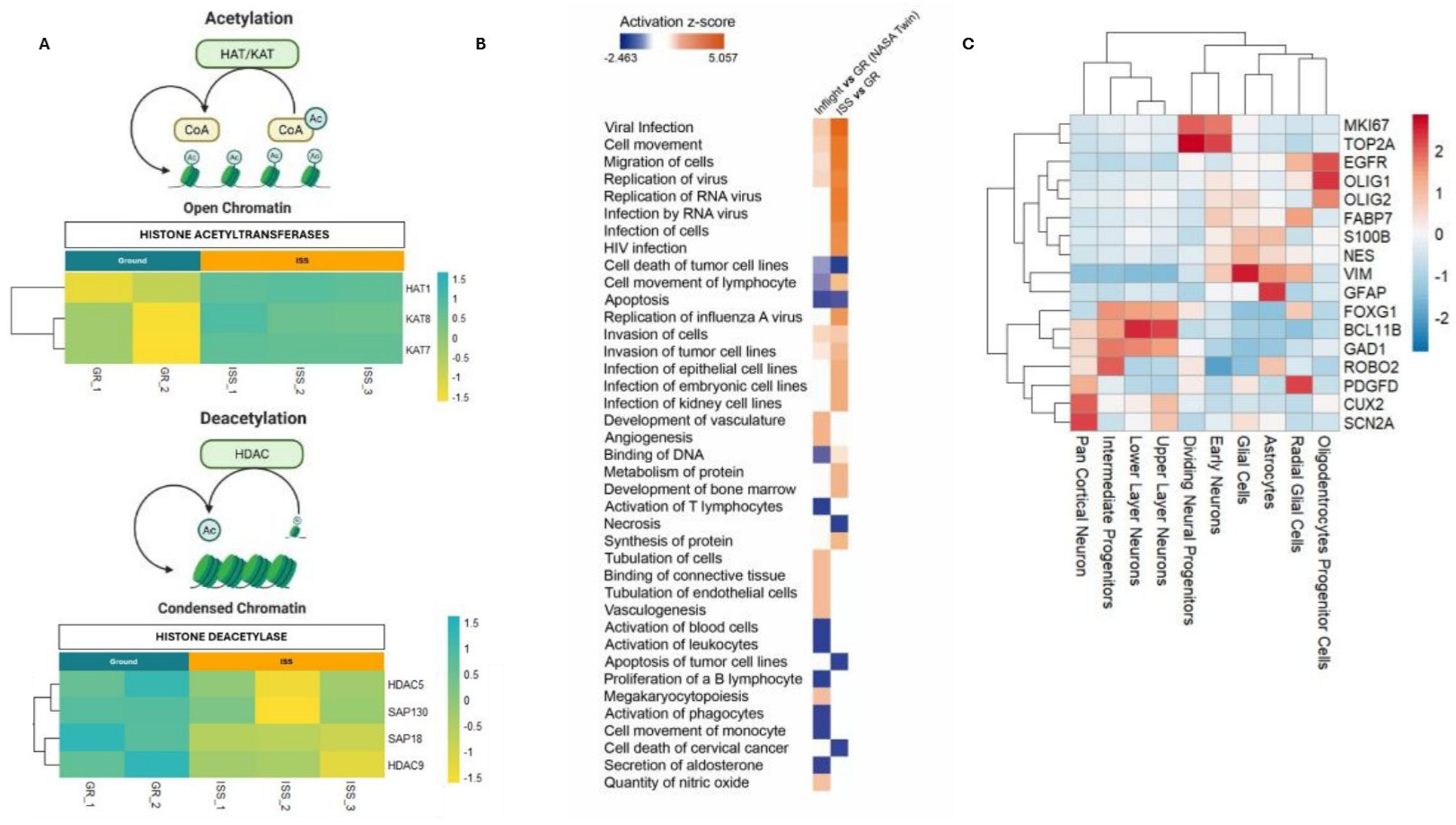
Proteomics and transcriptomics approach for viral pathway evaluation. **(A)** Heatmap of a framework of histone proteins showing upregulation of acetyltransferases and downregulation of deacetylases at ISS. **(B)** Differentially expressed genes (DEG) comparison between this study and NASA Twin study using Ingenuity Pathway Analysis (IPA), showing canonical pathways (on the left) and diseases and biological pathways (on the right). The colors correspond to activation z-score (negative score in blue, positive in orange). For pathway analysis: *Homo sapiens* as the reference organism, minimum overlap of 3 genes, enrichment factor> 1.5, and p < 0.01. **(C)** Heatmap of differentially expressed genes across cell types. The heatmap shows the normalized expression levels of key marker genes across different neural cell populations. Warmer colors (red) indicate higher expression levels, while cooler colors (blue) represent lower expression. Key genes exhibit distinct expression patterns across cell clusters.

**Fig. S3.**
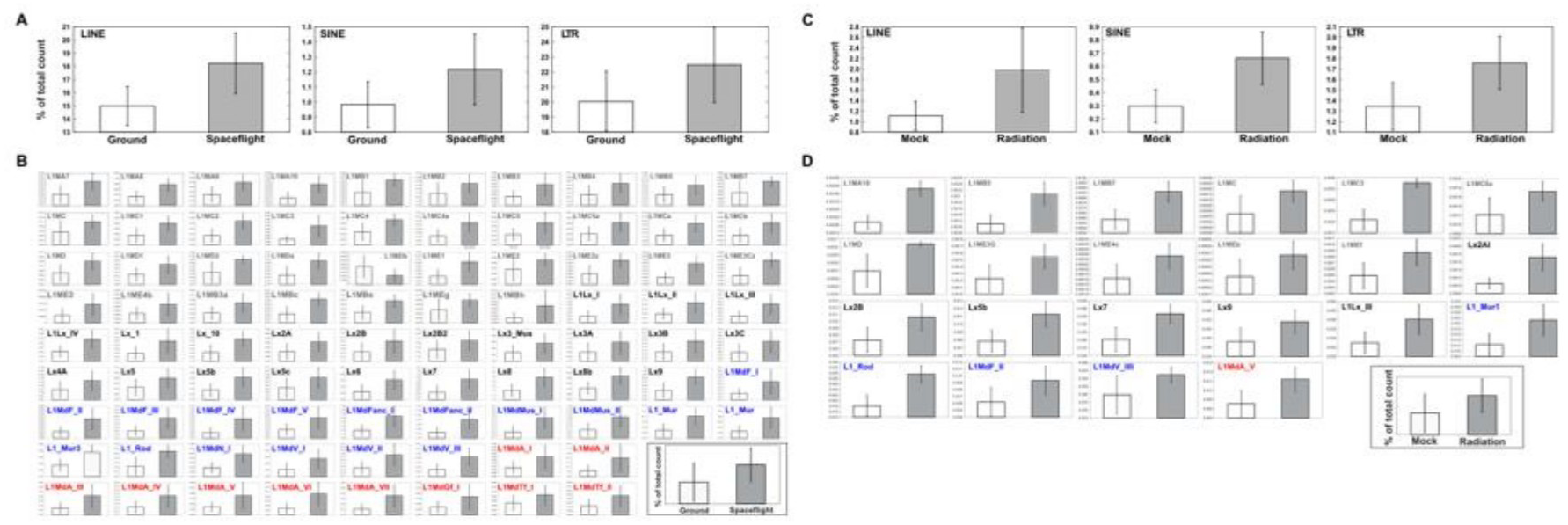
Retroelement expression analysis using publicly available RNA-seq data. **(A)** Expression levels of transposable elements in the retina of mice exposed to space flight (GSE159771). **(B)** Subfamily-level expression of L1 elements in the spaceflight retina. **(C)** Expression of transposable elements in mouse lung following simulated five-ion galactic cosmic radiation (GSE199680) **(D)** L1 subfamily expression profiles in mouse lung under five-ion radiation condition. **(B, D)** The color of L1 subfamily members represents evolutionary age: ancient members in gray, old members in black, middle members in blue, and young members in red.

**Fig. S4.**
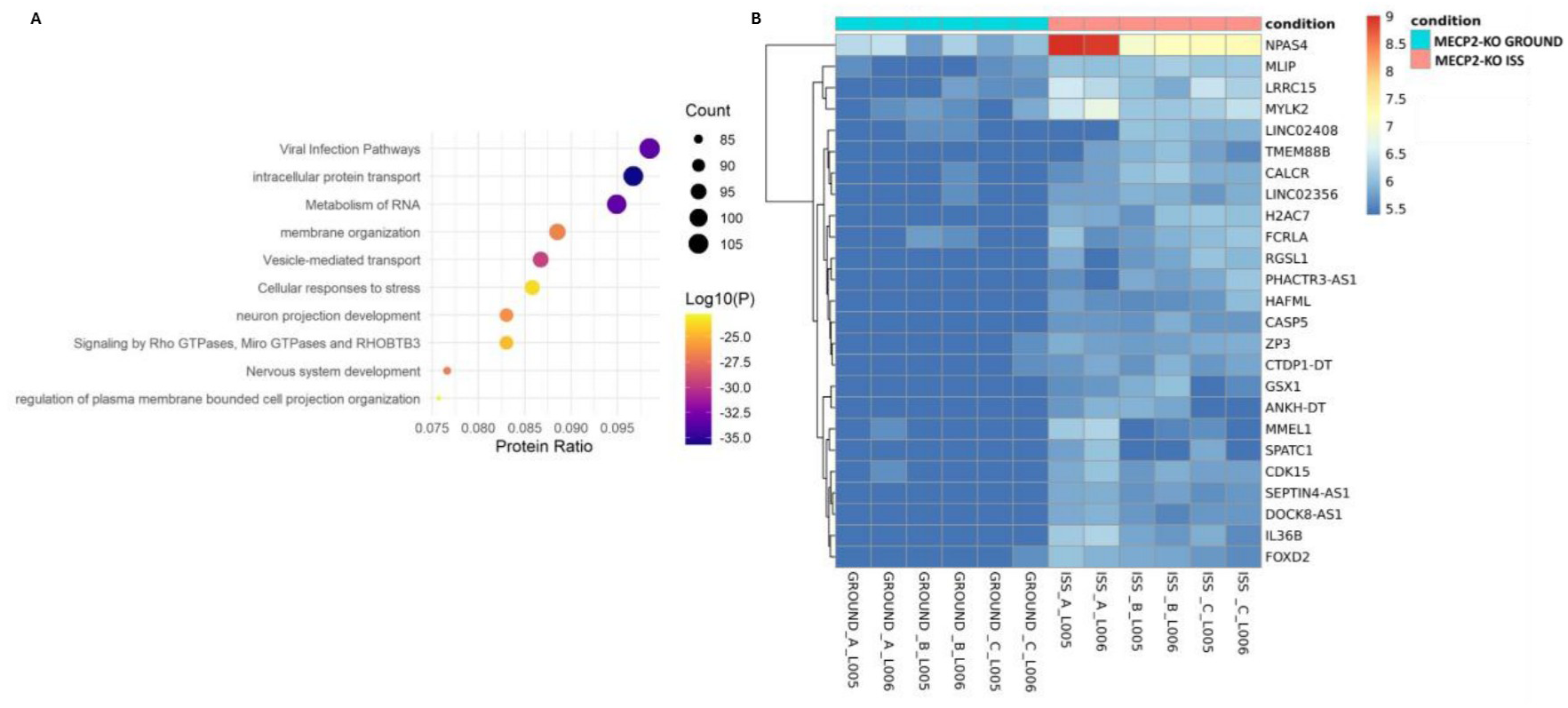
Proteomics and transcriptomics analysis of MECP2-KO brain organoids experiments. **(A)** Reactome from the proteomic analysis comparing control cells to MECP2-KO cells in ground conditions, emphasizing that MeCP2-KO cells demonstrate upregulation of the viral pathway. This analysis highlights the intrinsic alterations in MECP2-KO cells that predispose them to enhanced viral pathway activity and nervous system development even without the influence of the space environment. **(B)** Heatmap of bulk transcriptomics comparing ground MECP2-KO brain organoids vs. ground MECP2-KO brain organoids. For proteomics bulk analysis: p≤0.01; fc≥1.2. Test T Student for comparisons. For pathway analysis: *Homo sapiens* as the reference organism, minimum overlap of 3 genes, enrichment factor> 1.5, and p < 0.01.

**Fig. S5.**
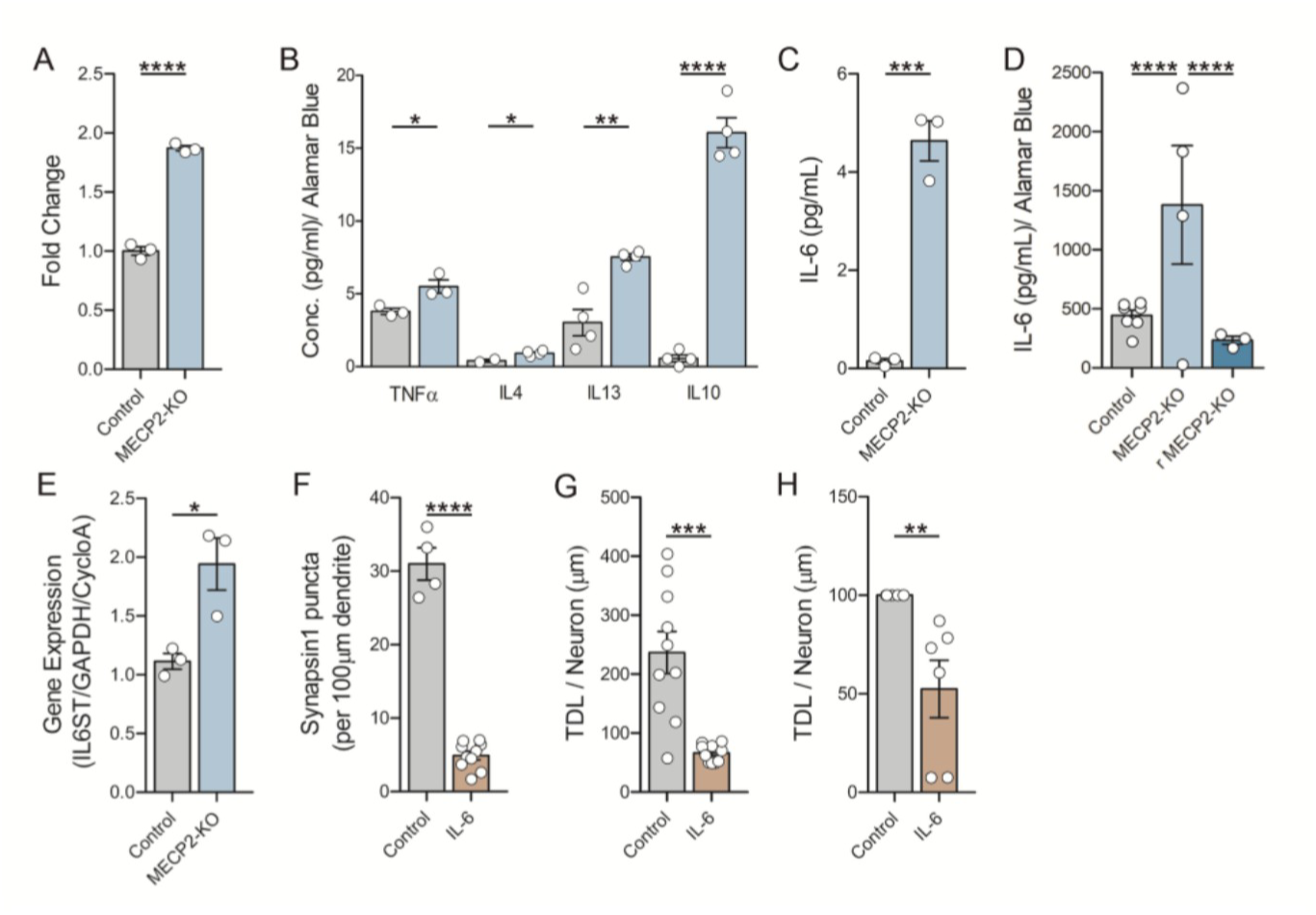
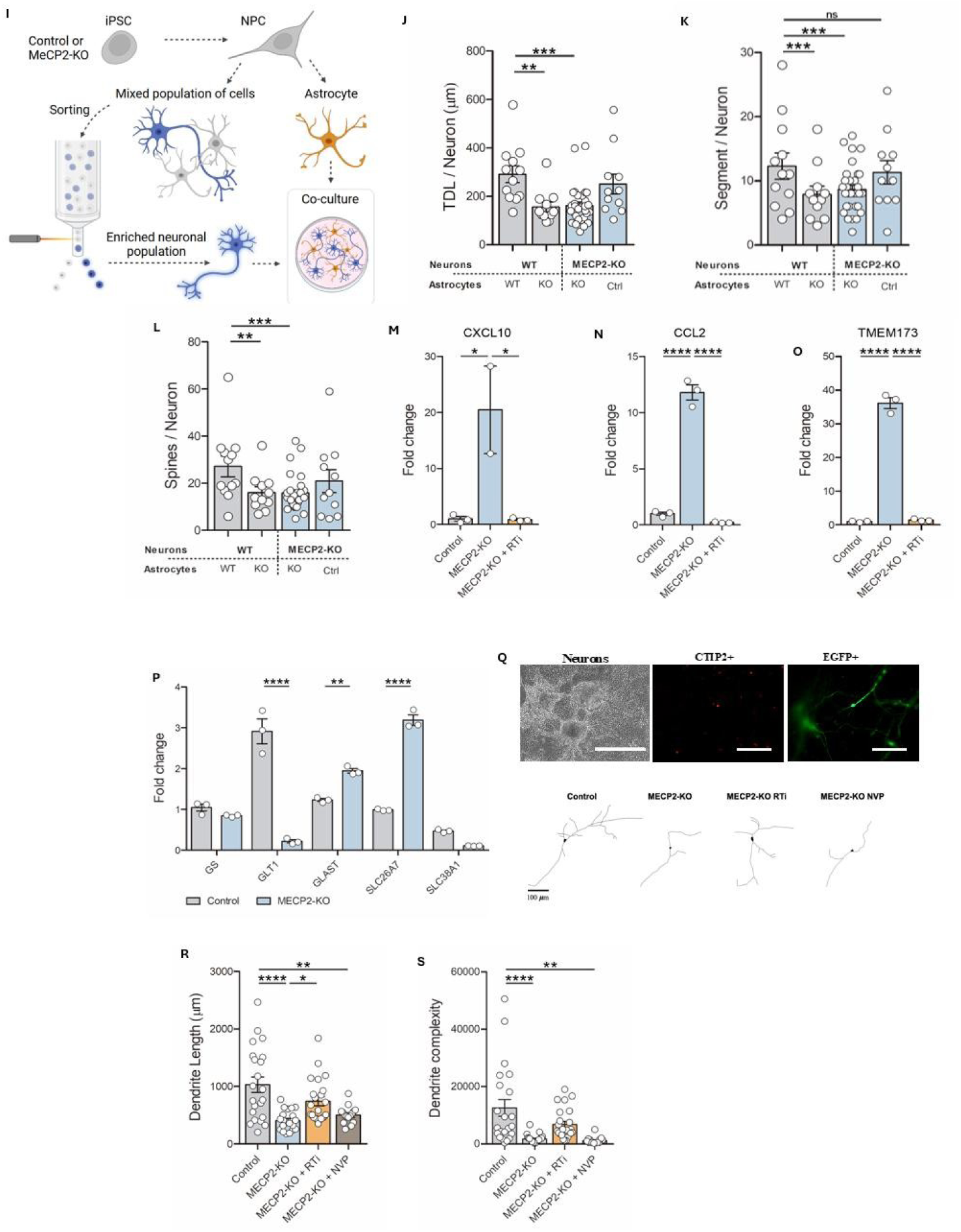
MECP2-KO astrocytes phenotypes and interplay with neurons. **(A)** Increase **L1** endogenous retroelement expression in MECP2-KO astrocytes measured by qPCR. **(B)** Quantification of TNFa, IL-4, IL-13, and IL-10 released in the media from control and MECP2-KO astrocytes by enzyme-linked immunosorbent assay (ELISA). **(C)** IL-6 protein level in post-mortem **RTT** brain sample compared to an age-matched control sample by ELISA. **(D)** Increased IL-6 released from RTT-derived astrocytes was rescued when MECP2 was re-expressed in the cells. The dotted red line indicates the background buffer signal for the ELISA assay. **(E)** Expression levels of the IL-6 family cytokine receptor’s signal transducer, gp130 receptor (IL6ST in post-mortem RTT brain samples compared to an age-matched control sample by qPCR. **(F)** Reduction of synapsin1 puncta in control neurons upon IL-6 treatment. **(G)** Reduction of Total Dendritic Length (TDL) in control neurons upon IL-6 treatment. **(H)** Reduction of neuronal activity measured by the number of control neurons upon IL-6 treatment, measured through MEA over a 260-second time course. (I) Schematic diagram of the experimental design. iPSC-derived NPCs were used to produced enriched populations of neurons and astrocytes for co-cultured experiments. **(J)** Co-culture experiments of neurons and astrocytes from the two genetic backgrounds (control and MECP2-KO) showing the negative impact of MECP2-KO astrocytes in neurons regarding total dendritic length **(TDL),** number of segments **(K),** and number of spines **(L). (M)** Quantification of CXCL10, CCL2 **(N),** and TMEM173 **(0)** expression in MECP2-KO astrocytes treated with RTis by enzyme-linked immunosorbent assay (ELISA). **(P)** Quantification of GS, GLT1, GLAST, and SLC26A7 gene expression in control and MECP2-KO astrocytes compare to control. **(Q) Top:** Representative image of neurons. SYN::EGFP positive cells co-stained with a cortical marker CTIP2 and EGFP to be further traced. Scale bar, 1000 µm for neurons; Scale bar, 100 µm for CTIP+ and EGFP+ neurons. **Bottom:** Representative images of tracings from control, MECP2-KO, RTi-treated MECP2-KO, and NVP-treated MECP2-KO neurons. **(R)** Quantification of dendritic length and complexity **(S)** in mixed cultures of neurons and astrocytes. Reverse-transcriptase inhibitor (RTi) or nevirapine (NVP)-treatment is indicated. Each color represents one condition: control (black), MECP2-KO (blue), MECP2-KO+RTi (orange), and MECP2-KO+NVP (green). Each experiment was done in biological triplicates. The number of measurements (n) for these experiments is indicated directly by the symbols (o) on each bar graph. Data are represented as mean ±SEM; *p<0.5* (*), p<0.01 (**), p<0.001 (***) and p<0.0001 (****).

**Fig. S6.**
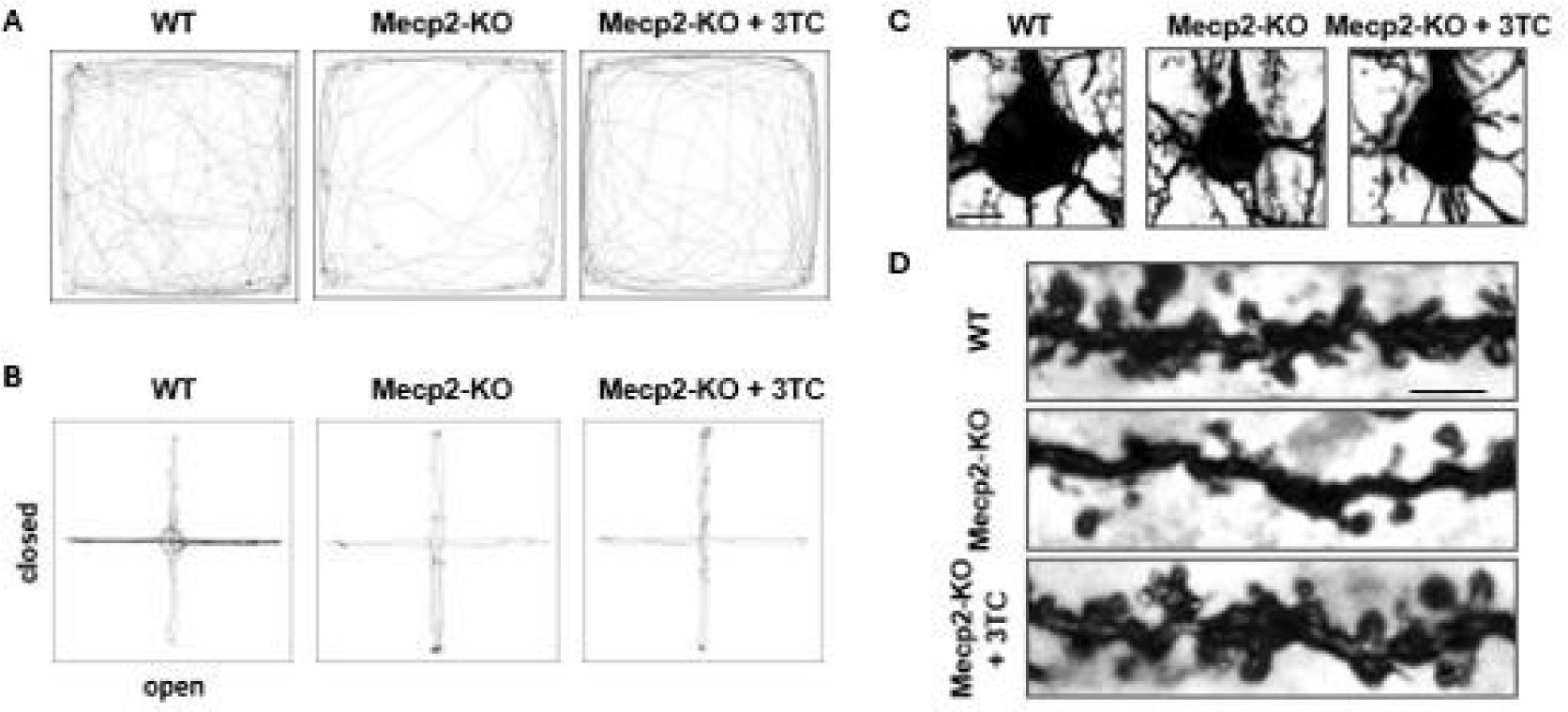
RTT-like phenotypic analysis of reverse transcriptase inhibitor-treated Mecp2-KO mice. **(A)** Digital tracking of WT, Mecp2-ko and 3TC-treated Mecp2-KO mice exposed to the open-field test. Representative traces of mice activity obtained from video tracking. **(B)** Digital tracking of WT, Mecp2-ko and 3TC-treated Mecp2-KO mice exposed to the elevated plus maze test. Representative traces of mice activity obtained from video tracking. **(C)** Representative images of somata of Golgi-stained layer 2/3 pyramidal neurons in PFC of WT, Mecp2-ko and 3TC-treated Mecp2-KO mice. Scale bar, 10 µm. **(D)** Representative images of dendrites of Golgi-stained layer 2/3 pyramidal neurons in PFC of WT, Mecp2-ko and 3TC-treated Mecp2-KO mice. Scale bar, 5 µm.

**Fig. S7.**
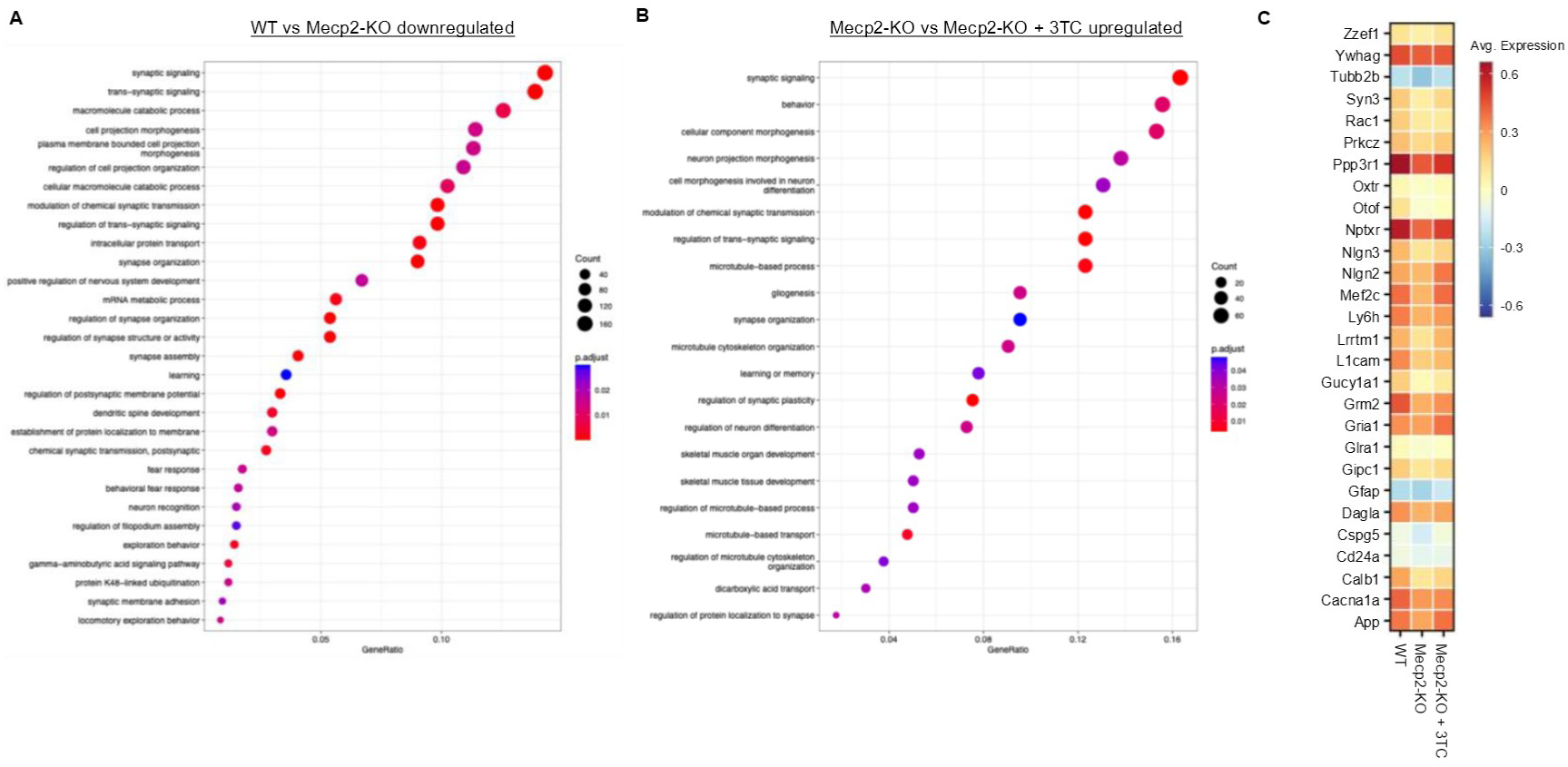
Spatial transcriptomics analysis of WT, Mecp2-KO and 3TC-treated Mecp2-KO excitatory neurons. (A) GO analysis of downregulated genes in Mecp2-KO excitatory neurons compared to WT excitatory neurons. **(B)** GO analysis of upregulated genes in 3TC-treated Mecp2-KO excitatory neurons compared to Mecp2-KO excitatory neurons. **(C)** Gene expression profiles of synaptic signaling (GO:0099536) genes that were significantly downregulated in Mecp2-KO excitatory neurons compared to WT excitatory neurons and whose expression was significantly improved by 3TC treatment.

Supplementary Video 1. Representative video of WT mice performing the open field test.

Supplementary Video 2. Representative video of Mecp2-KO mice performing the open field test.

Supplementary Video 3. Representative video of Mecp2-KO+3TC mice performing the open field test.

## MATERIAL AND METHODS

### IPSC tissue culture

All iPSCs were previously characterized^86^. In addition, we also used MECP2 isogenic cell lines using CRISPR/Cas9 genome editing (RTT/C9)^86^. Two isolated clones for each cell line were selected and expanded for further differentiation and experimentation. Mycoplasma and CNV analyses were performed regularly in all cultured cells. For consistency, “Control” or “MECP2-KO” were used in the manuscript. All iPSCs were cultured and manually passaged onto Matrigel-coated (Corning) and fed daily with mTESR1 (StemCell Technologies). iPSCs were differentiated into NPCs as described^87^. NPCs were cultured on the plate coated with 10 µg/mL of poly-L-ornithine (Sigma-Aldrich) and 2.5 µg/mL laminin (Life Technologies). Media was changed every other day with NG medium (DMEM/F12 50/50 1X (Corning Cellgro), 1% HEPES (VWR International), 0.5% Penicillin Streptomycin (Pen-Strep) (Life Technologies), 1X GlutaMAX (Life Technologies), 0.5% N2 NeuroPlex (Gemini Bio-Products) and 1% Gem21 NeuroPlex (Gemini Bio-Products) was supplemented with 20 ng/mL basic human fibroblast growth factor (bFGF) (R&D Systems, 414-TE).

### Neural differentiation

For neural differentiation, bFGF was withdrawn from the NPCs containing NG media, and Rock inhibitor (Y27632) was added at a concentration of 10ng/mL for 48 hours. Cells matured for 5 to 8 weeks, replacing the media every 3 to 4 days. IPSC-derived cortical organoids were differentiated following^1^. Briefly, iPSCs were cultured, dissociated using a 1:1 Dulbecco’s phosphate-buffered saline (DPBS, Fisher Scientific) and StemPro Accutase solution, transferred to 6-well plates, and kept in suspension. Neural induction media consisted of DMEM/F12, 1% Glutamax, 1% N2 Neuroplex, 1% non-essential amino acids (Gibco), 1% Pen-Strep, 1µM of Dorsomorphin (R&D Systems), and 10µM SB431542 (SB, Stemgent). Neural proliferation media consisted of Neurobasal media (Life Technologies), 2% Gem21 Neuroplex, 1% non-essential amino acids, 1% Glutamax, 20 ng/mL epidermal growth factor (EGF, Peprotech), and 20 ng/mL bFGF. For neuronal maturation, the cells were kept in the same media, but the growth factors were withdrawn. Organoid results are combined from four separate batches of differentiation.

Astrocytes were differentiated as described^88^. Briefly, monolayer NPCs are dissociated into single-cell suspension using StemPro Accutase. To form astrospheres, cells were seeded at a density of 4-5 million cells per well in a 6-well plate (Genesee, 25-105) shaking at 95 r.p.m. at 37°C in NSCR EL medium containing DMEM/F12 (Gibco) with 1% N2 supplement (Gibco) and 0.1% SM1 supplement, 1% nonessential amino acids (Gibco), 1% penicillin/streptomycin (Gibco), 1% GlutaMAX solution (Gibco), supplemented with 20 ng/ml epidermal growth factor 1 (EGF-1, Peprotech) and 20 ng/mL leukemia inhibitory factor (LIF, Peprotech). Following 2-4 weeks in the NSCR EL media, the astrospheres were expanded in NSCR media now supplemented with 20 ng/mL EGF and 20 ng/mL FGF (NSCR EF) for 2-4 weeks. Following this expansion phase, astrospheres were plated on 20 µg/mL poly-L-ornithine and 5 µg/mL laminin and remained in NSCR EF media, allowing astrocyte progenitor cells (APCs) to migrate outward until 70% confluency. For astrocyte assays, progenitors were expanded and seeded at 50,000 cells/well in a 96-well plate. These progenitor cells can be expanded and frozen in cryovials for later use. For differentiation into mature astrocytes, media was changed to contain half of the NSCR basal media and half Astromed CNTF media. Astromed media contained Neurobasal media (Gibco) with 0.2% SM1, 1% NEAA, 1% penicillin/streptomycin, 1% Glutamax and 10ng/mL of ciliary neurotrophic factor (CNTF, Peprotech). After 48 hours, cells were transitioned into full Astromed CNTF media for 14 days.

### Cubelab operations (flight and ground)

Space Tango’s flight and ground hardware support 12 wells containing impellers and automated operations (fluid conveyance, fluid routing, thermal control, environmental monitoring, imaging, and sampling) within each 9U Cubelab_TM._ All wetted materials selected within the Cubelab were certified biocompatible and sterilizable. Wetted materials that come in contact with fluids or cells were sterilized, assembled, maintained sterile, tested, and confirmed to be free of microbial contamination prior to loading experimental fluids and biology. Science and the Flight system Cubelab were prepared at the Kennedy Space Center (KSC) Space Station Processing Facility (SSPF) in Brevard County, FL. Scheduled operations began shortly after science loading, continuing through launch, ascent, and on-orbit, and ended upon fixation and transfer to cold stowage. Space Tango’s Powered Ascent Utility Locker (PAUL) allows Cubelabs to be fully powered while installed in the SpaceX Cargo Dragon vehicle. The PAUL provides the optimal culture environment and capabilities for the entire time the science remains within the Cubelab, including while on the launch pad. Flight and Ground systems were run synchronously. The Ground system scheduled operations were initiated 9 days after the Flight system.

The Flight system Cubelab culture environment was maintained at 37°C and 5-7% CO_2_, and culture media was held at 2-8°C, providing optimal culture conditions. Media exchanges were performed every 3 - 4 days using microfluidic routing and pumping. Organoids were exposed to the Cubelab environmental gases through a 10:1 (base:curing agent) polydimethylsiloxane (Sylgard 184, Dow) interface within each well cap. Impeller-driven agitation was required for the Flight system on Earth, through ascent and installation, and during media exchanges. During all other times, no agitation was performed. Imaging of wells was performed twice per week for 30 days, beginning after installation and ending prior to cold stowage transfer. SpaceX mission CRS-25 was 30 days. On the 30^th^ day of on-orbit operations, 4% paraformaldehyde fixation of 6 wells and RNALater fixation of the remaining 6 wells was performed. The Flight system was stored at approximately 4°C for 6 days until science removal at the SSPF. Additionally, Space Tango’s cryovial flight hardware contains 6 stainless steel boxes, each supporting up to 98 1 ml cryovials (Nunc, 374088) for up to 588 1 ml cryovials per 9U Cubelab ™. The Flight system Cubelab culture environment was maintained at 37°C and 5% CO2. providing optimal environmental culture conditions. SpaceX CRS-25 mission was 30 days. Ground control samples were loaded into 1x stainless steel cryovial box and maintained within a humidified incubator set to 37°C and 5% CO2, located at the SSPF, until the flight samples returned to Earth. Flight and ground samples were run synchronously, and scheduled operations were initiated at the exact same time.

The Ground system experiment was performed at the Space Tango Headquarters in Lexington, KY. The Cubelab culture environment was maintained at 37°C, and 5-7% CO2, and culture media was held at 2-8°C, providing optimal culture conditions. Media exchanges were performed every 3-4 days using microfluidic routing and pumping. Impeller-driven agitation was required for the Ground system throughout the entire experiment on Earth. Imaging of wells was performed during media exchanges only due to impeller movement. On the 30^th^ day of matched on-orbit operations, 4% paraformaldehyde fixation of 6 wells and RNALater fixation of 6 wells was performed. The Ground system was stored at approximately 4°C for 17 days until scientific removal. Cubelab environmental telemetry (culture environment temperature, temperature of culture media, Cubelab CO2, Cubelab humidity) was recorded during powered operations. Telemetry data was available in near real-time through Space Tango’s customer dashboard.

### Imaging operations and post-processing (flight and ground)

During the ascent phase, agitation was enabled continuously on all wells. 13-second videos were recorded once per hour for each well during this time. Once installed into the ISS, the constant agitation period was discontinued. This was changed to record videos during the agitation event during the media exchanges, which occurred every 3.5 days. To get more insight into how well the organoids were growing, a health check was added after the first 2 media exchanges. The health check involved capturing a photo of each well once per hour for the remainder of the mission. Post-processing included adding a scale bar onto each image that represents the size of 1mm in the image. Images were uploaded to Space Tango’s data portal. The ground study payload had constant agitation enabled from the experiment start until fixation. During this time, 13-second videos were recorded once per hour for each well. Additionally, health check images were added before and after each media exchange at an interval of 3.5 days. Post-processing included adding a scale bar onto each image that represents the size of 1mm in the image. Images were uploaded Space Tango’s data portal.

### Mass spectrometry bulk analysis

A total of 500 ng of each sample was randomized loaded onto individual Evotips trap columns following manufacture’s protocol. The peptides were separated using an Evosep One LC system (Evosep, Denmark) in 44 min gradient (30 samples per day predefined Evosep proprietary method) on a PepSep 8 (8 cm × 150 µm × 1.5 µm C18 column, Bruker Daltonics) at 50°C. The mobile phases were 0.1% FA in LC-MS grade water (buffer A) and 0.1% FA in 99.9% ACN (buffer B). The LC system was coupled with timsTOF Pro 2, a hybrid trap ion mobility spectrometer and quadrupole time-of-flight instrument (Bruker Daltonics) operated in positive mode previously calibrated for mass (m/z) and ion mobility dimension (1/K0), using three ions of Agilent ESI-Low Tuning Mix (m/z 622.0289; 922.0097 and 1221.9906). The nano-electrospray source (CaptiveSpray source, Bruker Daltonics) was operated at 1600 V of Capillary voltage, 3.0 Umin of dry gas, and 180 °C of dry temperature. Data was acquired in dia-PASEF acquisition mode with a cycle time estimate of 1.8 s46. To cover the mass range of 400 to 1200 m/z, 32 isolation windows of 25 m/z width were used. Ramp Time and accumulation time were set to 100 ms, and collision energy ramped linearly as a function of the mobility from 59 eV at 1/K0 = 1.6 Vs. cm-2 to 20 eV at 1/K0 = 0.6 Vs. cm-2.

### Data processing for mass spectrometry-based bulk proteomics

All diaPASEF data was imported in Spectronaut (v.19, Biognosys) and processed using directDIA workflow which does not require spectral library from DOA runs. In brief, Spectronaut Pulsar search engine firstly generates ODA-like pseudospectra from the fasta database and then performs searches. The human protein reference database (UP000005640 reviewed canonical proteins) was used with Pulsar and the following settings: Trypsin/Pas specific enzyme; peptide length from 7 to 52; max missed cleavages 2; toggle N-terminal M turned on; Carbamidomethyl on C as fixed modification; Oxidation on Mand Acetyl at protein N-terminus as variable modifications; FDR at PSM, peptide and protein group level set to 0.01; minimum fragment relative intensity 1%; 3-6 fragments kept for each precursor; the extraction of the ion chromatograms and ion mobility window was set as “dynamic” with correction factor 1.

### Functional analysis for mass spectrometry-based bulk proteomics

The interpretation of systems-level studies was done on Metascape, an online platform that provides a comprehensive resource for annotating and analyzing gene/protein lists^53^. In this study, we used enrichment analysis to compare thousands of protein sets defined by their involvement in specific biological processes, protein localization, enzymatic function, pathway membership, and protein network analysis and interactions. All data was post-processed to generate Kappa similarities, and all pairs of enriched terms were computed into clusters of comparable terms^54^.

The proteins quantified in at least 70% of the samples within each group were subjected to statistical and graphical analyses using the R programming environment. Abundance values were log10-transformed prior to analysis. Boxplots were generated using the ggplot2, dplyr, and tidyr packages^89^. Heatmaps were constructed using the heatmap REF package, based on ANOVA p-values to highlight differential expression patterns. Volcano plots were created using the ggplot2 and ggrepel REF packages. Gene Ontology (GO) enrichment analyses were performed using Cytoscape^90^ and STRING^91^.

Protein identifiers and associated statistics (log_2_ fold change and adjusted p-values) were submitted to Metascape (www.metascape.org) for pathway enrichment analysis. Parameters were set to default values: *Homo sapiens* as the reference organism, minimum overlap of 3 genes, enrichment factor > 1.5, and p < 0.01. The resulting output was used for downstream processing, focusing on the table containing pathway description, enrichment ratio, −log_10_(p-value), and the number of proteins contributing to each term. Data visualization was performed in R (version 4.3.2) using the readxl, ggplot2, and viridis packages. The enriched pathways were ordered according to their enrichment ratio, and a dot plot was generated to display the top biological processes. In the plot, the x-axis represents the enrichment ratio (GeneRatio), the y-axis lists pathway descriptions, dot size corresponds to the number of input proteins (Count), and dot color encodes statistical significance based on −log_10_(p-value).

### Secretome panel evaluation by Proximity Extension Assay (PEA) Olink® Reveal (Thermo Fisher Scientific)

The Olink Reveal technology uses Proximity Extension Assay (PEA)^92^, where a matched pair of antibodies labelled with unique complementary oligonucleotides (proximity probes) bind to the respective target protein in a sample. As a result, the probes come into close proximity and hybridize to each one, enabling DNA amplification of the protein signal, which is quantified on a next generation sequencing read-out which has been described in detail previously. Antibodies targeting 1034 unique proteins are distributed across 384-plex panel. Assays are run in a 96-well format. Each plate consisted of 32 secretome samples (in triplicates) from brain organoids that belong to the ground experiment and the ISS experiment, using 20 µL of media sample. In brief, pairs of oligonucleotide-labeled antibody probes designed for each protein bind to their target, bringing the complementary oligonucleotides in close proximity and allowing for their hybridization. The addition of a DNA polymerase leads to the extension of the hybridized oligonucleotides, generating a unique protein identification “barcode”. Next, library preparation adds sample identification indexes and the required nucleotides for lllumina sequencing. Prior to sequencing using the lllumina® NovaSeq™ 6000/NextSeq™ 550/NextSeq™ 2000, libraries go through a bead-based purification step and the quality is assessed using the Agilent 2100 Bioanalyzer (Agilent Technologies, Palo Alto, CA). The raw output data is quality controlled, normalized and converted into NPX®, values as their log-2 scaled quantification unit of relative abundance. Data normalization is performed using an internal extension control and an external plate control to adjust for intra- and inter-run variation. All assay validation data (detection limits, intra- and inter-assay precision data, predefined values, etc.) are available on manufacturer’s website (www.olink.com). Data was processed by Olink’s MyData Cloud Software.

### RNA sequencing and transcriptomic analysis

For transcriptomic analyses, we combined brain organoids from the three healthy donors for ground processing for RNA sequencing. Ribodepletion RNA sequencing was performed at the UC San Diego IGM core facility. Data was analyzed by ROSALIND® (https://rosalind.bio/), with a HyperScale architecture developed by ROSALIND, Inc. (San Diego, CA). Briefly, reads were trimmed using cutadapt. Quality scores were assessed using FastQC. Reads were aligned to the Homo sapiens genome build hg19 using STAR. Individual sample reads were quantified using HTseq and normalized via Relative Log Expression (RLE) using DESeq2 R library. Read Distribution percentages, violin plots, identity heatmaps, and sample MOS plots were generated as part of the QC step using RSeQC. DEseq2 was also used to calculate fold changes and p-values and perform optional covariate correction. The clustering of genes for the final heatmap of differentially expressed genes was done using the PAM (Partitioning Around Medoids) method using the fpc R library. Hypergeometric distribution was used to analyze the enrichment of pathways, gene ontology, domain structure, and other ontologies. The topGO R library was used to determine local similarities and dependencies between GO terms to perform Elim pruning correction. Several databases were referenced for enrichment analysis, including lnterpro, NCBI, MSigDB, REACTOME, WikiPathways. Enrichment was calculated relative to a set of background genes relevant to the experiment. Functional enrichment analysis of pathways, gene ontology, domain structure, and other ontologies was performed using HOMER.

In our study, we employed the DESeq2 package in R for differential expression analysis, utilizing a significance threshold of p < 0.05. Raw count data, obtained from experimental samples representing ground and ISS conditions, were processed and normalized. Subsequently, differential expression analysis was performed, allowing for the identification of genes significantly altered between the two conditions, with a fold change threshold of 1.5. Additionally, to gain insights into the biological pathways underlying these expression changes, Reactome pathway analysis was conducted on the differentially expressed genes, with a significance threshold of p < 0.05. Furthermore, we performed additional pathway analysis using Wiki Pathways library, employing a false discovery rate (FDR) adjusted p-value cutoff of 0.1, to further elucidate the enriched biological processes associated with the observed gene expression alterations.

### Xenium experiment

Mice were anesthetized and perfused trans-cardiac with PBS followed by 4% buffered paraformaldehyde (PFA). Brains were removed and fixed with 4% PFA overnight at 4°C. The fixed tissue was cryoprotected with 30% sucrose and frozen in optimal cutting temperature medium (OCT). Coronal sections were obtained by sectioning the tissue at 10 µm using Leica CM3050S cryostat. Sections were washed with PBS and placed onto Xenium slides.

The Xenium 5K Mouse Pan-Tissue and Pathways Panel (1Ox Genomics) was used for transcriptomic profiling. Probe hybridization, RNase treatment, ligation, amplification, and cell staining were performed according to the manufacturer’s protocol. Data acquisition and on-board analysis were conducted using the Xenium Analyzer (v3.3).

### Xenium data processing

Regions of interest (ROls) in the WT, Mecp2-KO, and 3TC-treated Mecp2-KO brains were manually selected using Xenium Explorer 3 (v3.1.1). Only cells within ROls with more than 250 detected genes and more than 350 transcripts were included in downstream analysis. Data normalization, principal component analysis (PCA), UMAP-based dimensionality reduction, and cell clustering were performed using the Seurat package (v5.0.1). Cell type annotations were manually assigned based on the expression of established marker genes. Differential gene expression analysis was performed using the FindAIIMarkers function in Seurat. Over-representation analysis of genes included in the predefined gene list of the Xenium panel in the transcriptome data of brain organoid samples returned from the ISS was performed using Fisher’s Exact Test. Gene Ontology (GO) analysis of Biological Processes was conducted using the clusterProfiler package (v4.4.4).

### Reverse Transcriptase inhibitors

The reverse-transcriptase inhibitors for cultured cells were prepared as follows: Lamivudine (3TC, Sigma-Aldrich) was prepared in dimethyl sulfoxide (DMSO, Sigma-Aldrich), and 10 µM of 3TC was used in the media. Stavudine (d4T, Sigma-Aldrich) was re-suspended in water, and 1µM of d4T was used in the media in combination with 3TC. Nevirapine (Sigma Aldrich, SML0097) was re-suspended in DMSO and 400 nM nevirapine was used in media. Control cells contained 10 µM of DMSO as a vehicular control.

### lmmunofluorescence

Cells cultured in monolayer were washed with DPBS, fixed in 4% paraformaldehyde (PFA, Core Bio) for 20 minutes at room temperature, and permeabilized with a solution of DPBS and 0.1% Triton X-100 (Promega) for 10 minutes at room temperature. Cells were incubated at room temperature with blocking solution for 30 minutes (DPBS, 10% donkey or goat serum (Fisher Scientific), and 0.005% Triton X-100). Primary and secondary antibodies were diluted in a solution containing DPBS, 1% donkey or goat serum, and 0.005% Triton X-100. Cells were incubated with the primary antibody overnight at 4°C. The next day, cells were washed with DPBS three times for 5 minutes and incubated with secondary antibodies for 30 minutes at room temperature. Cells were washed twice with DPBS, and nuclei were stained using DAPI (VWR International) for 10 minutes at room temperature. Finally, cells were washed once with DPBS for 5 minutes, and the coverslips were mounted using Prolong Gold antifade mounting (Life Technologies) onto glass slides (Fisher Scientific).

For cortical brain organoids, cells were washed with DPBS and placed in a 1.5 ml microcentrifuge tube with 4% PFA overnight at 4°C. Cortical organoids were washed with DPBS and incubated in 30% sucrose (Sigma-Aldrich) for 48 hours at 4°C. The tissue was removed from the sucrose solution, placed with optimal cutting temperature compound (OCT, VWR) within an embedding mold (Fisher Scientific), and incubated at −80°C overnight. Using a cryostat (Leica), the molds containing the organoids were sliced at 10.0 µm and placed onto a glass slide which was left to dry at room temperature for at least 4 hours. The slides were lined with a hydrophobic barrier pen (Vector Laboratories). Organoid slides were permeabilized, stained, and mounted as explained above.

### lmmunohistochemistry

Mice were anesthetized and perfused trans-cardiac with PBS followed by 4% buffered paraformaldehyde (PFA). Brains were removed and fixed with 4% PFA overnight at 4°C. The fixed tissue was cryoprotected with 30% sucrose and frozen in optimal cutting temperature medium (OCT). Coronal sections were obtained by sectioning the tissue at 40 µm using Leica CM3050S cryostat. Sections were washed with PBS to remove the OCT, permeabilized and blocked with blocking buffer at RT for 1 h, and then incubated with primary antibody diluted in blocking buffer overnight at 4°C. After three washes with PBS, the sections were incubated with corresponding secondary antibodies at RT for 2 h. The following primary and secondary antibodies were used in this experiment: rabbit anti-hC3D (1:500; DAKO, Cat.# A0063), mouse anti-S100p (1:500; Sigma, Cat.#S2532); CF488 donkey anti-mouse lgG (H+L), highly cross-adsorbed (1:500; Biotium), CF555 donkey anti-rabbit lgG (H+L), highly cross-adsorbed (1:500; Biotium). Hoechst 33258 (1:500) was used for nuclear staining. After three washes with PBS, the sections were mounted on glass slides. Fluorescence images of astrocyte were acquired using Zeiss LSM 800 confocal microscope.

### Magnetic cell sorting

Briefly, iPSC-derived cells were dissociated using the neurosphere dissociation kit (MACS), incubated with the CD44-PE and CD184-PE (both antibodies were from BO Biosciences) for 15 minutes in the dark at 4°C and washed twice with dPBS to remove unbound antibodies before sorting. Next, the cells were incubated with PE antibodies and conjugated with magnetic beads (MACS). Following the antibody incubation, cells are loaded into a magnetic column, and the cells that are CD44 and CD184 positive are retained in the column. The remaining enriched neuronal population is obtained and counted for viability and seeding.

### Astrocyte calcium wave propagation

Calcium waves were induced by focal mechanical stimulation of single astrocytes in the center of the field of view. A 175 W Xenon lamp and a wavelength switcher (Lambda DG-4, Sutter Instrument) provided 485 nm fluorescence excitation. The emission from Fluo-4 loaded cells is detected at a wavelength of 520 nm using a F-Fluar 40X/1.3 NA oil immersion objective and an attached 12-bit CCD camera. Changes in Fluo-4 fluorescence intensities emitted at 520 nm were acquired at 0.67 Hz. The calcium imaging data were visualized and analyzed offline using MetaMorph software (Universal Imaging Corporation). The fluorescent intensity of Fluo-4 was determined in individual cells in every astrocyte soma in the field of view. The distance that calcium waves propagated within a certain time was determined by measuring the distance from the stimulating point to the farthest responding cell in the field of view.

### Morphological analysis of iPSC-derived neurons

Cultured NPCs were transduced with a lentivirus backbone containing Synapsin 1 (SYN) promoter upstream of an EGFP reporter^93^. The transduction multiplicity of infection (M.O.1.) was equal to 2. Cells transduced with a SYN::GFP lentivirus were differentiated into neurons for 5-6 weeks, fixed, and stained with GFP and CTIP2 antibodies. Differentiated neurons were traced using Neurolucida v.2017 (MBF Bioscience, Williston, VT) connected to a Zeiss Axio lmager 2 microscope at a 40X oil objective. Only neurons that were both GFP- and CTIP2-positive were included in this analysis. The morphology of the neurons was quantified using Neurolucida Explorer v.11 (MBF Bioscience, Williston, VT), and the results reflect the summed length of all neurites and dendrites per traced neuron. Sholl analysis was performed using Neurolucida Explorer’s Sholl analysis. This analysis specified a center point within the soma and created a grid of concentric rings around it with radii increasing in 10 µm increments. Neuronal complexity was determined by recording the number of intersections within each ring.

### Synaptic puncta quantification

5-6-week-old neurons were fixed and stained for the following markers: SYN1, postsynaptic density protein 95 (PSD-95), and MAP2 (refer to **lmmunocytochemistry** for protocol). Using a fluorescence microscope (Z1 Axio Observer Apotome Zeiss), the slides were imaged by compiling Z stack images taken from incremental focus distances. Co-localized SYN1 (presynaptic) and PSD-95 (postsynaptic) were quantified using the compiled Z stack image. Only co-localized puncta that were in proximity of a MAP2-positive neurite were included in this analysis.

### Measurement of cortical organoid diameter

Cortical organoids were imaged on an Evos FL Imagine System (Thermo Fisher Scientific) using a 4X magnification. The images were uploaded to Image J, and the diameter of each organoid was determined by the program’s length measurement.

### RNA extraction and qPCR

RNA was extracted using the RNAse Plus Mini Kit (Qiagen). Briefly, adherent cells were washed with DPBS and further lysed using RLT buffer with B-mercaptoethanol (Thermo Fisher Scientific) and dissociated using a 1ml syringe (Fisher Scientific, 1482330) and a needle (VWR, BD- 305196). RNA from lysates was extracted following the manufacturer’s instructions. cDNA was generated using the Qiagen Quantitect Reverse Transcription Kit. Protocol was followed on 1 µg of the RNA samples. qPCR was performed using selected primers and SYBR Green master mix (BioRad). Diluted cDNA was used in a 96-well or 384-well plates (BioRad) and ran on a BioRad CFX machine. PCR conditions were as follows: initial cycle of 95°C for 32 min, followed by 40 cycles of 15s at 95°C, 30s at 58°C, 30s at 72°C, with a final step of 5s at 57°C and 5s at 95°C for the melting curve. 3 plates were run to obtain triplicates per condition. The L1 primers used were: ORF1: ATGGGGAAAAAACAGAACAGAAAAACTGGAAACTCTAAAACGCAGAGCGCCTCTCCTCCTCCAA AGGAACGCAGTTCCTC; ORF2:

GCTCATGGGTAGGAAGAATCAATATCGTGAAAATGGCCATACTGCCCAAGGTAATTTACAGATTCAATGCCATCCCCATC; ORF2-3’UTR:

TGGAAACCATCATTCTCAGTAAACTATCGCAAGAACAAAAAACCAAACACCGCATATTCTCACTCA TAGGTGGGAA TTGA

### PCR array

A PCR array was performed using RT2 Profiler PCR Array 384 format (Qiagen), following the manufacturer’s instructions. Briefly, 400 ng of total RNA was used for reverse transcription using the RT2 First Strand Kit (Qiagen). Genomic DNA was eliminated prior to reverse transcription. Real-time PCR was performed using RT2 SYBR Green Mastermix and cDNA mix, divided into 384 wells. Plates were sealed with Optical Adhesive Film and ran into a CFX384 (BioRad). Run conditions: 1 cycle of 10 min at 95°C, followed by 40 cycles of 15s at 95°C and 1 minute at 60°C, where data collection was performed. For data analysis, GeneGlobe Data Analysis Center was used. Reference genes used: peptidyl-prolyl cis-trans isomerase H (PPIH), glyceraldehyde-3-phosphate dehydrogenase (GAPDH), Non-POU Domain Containing Octamer Binding (NONO), Lactate dehydrogenase A (LDHA), Actin Beta (ACTS). Significant changes in gene expression were set to a min of 2fold differences.

### Multi-electrode array (MEA)

12-well MEA plates (Axion Biosystems) were coated with 100 µg/mL of poly-L-ornithine (Sigma-Aldrich) and 5 µg/mL laminin (Life Technologies). Cultures were placed in the center of the MEA well to fully cover the electrodes. Media was changed twice a week, and recordings were measured one day after the media was changed. Recordings were performed once a week using a Maestro MEA System and AxlS Software (Axion Biosystems). A band-pass filter with 10 Hz and 2.5 kHz cutoff frequencies was used, and a spike detector threshold was set at 5.5 times the standard deviation. For each recording, the plate was left untouched in the Maestro for 3 minutes prior to recording, then 3 minutes of data were recorded. Analysis was performed using Axion Biosystems Neural Metrics Tool. The criteria to detect an active electrode was set to 5 spikes per minute, and the criteria to detect a burst electrode was set to 5 bursts per minute.

### TUNEL assay

Cells were cultured with the different treatments on 96-well plates, and once confluent, the cells were fixed with 4% PFA and permeabilized with 0.1% Triton X-100 for 15 minutes. Cells were blocked with 3% bovine serum album (Gemini) for 1 hour. Cells were incubated with primary antibodies of Nestin and TUNEL (Click-iT TUNEL assay kit, Life Technologies) overnight at 4’C. The following day, cells were incubated with secondary antibodies for 30 minutes at room temperature and with DAPI for 5 minutes at room temperature. Cells were mounted using Prolong Gold Antifade mounting, and images were taken and manually quantified.

### IL6 Elisa

Astrocyte progenitor cellss were dissociated using 1:1 Accutase:PBS for three minutes and seeded at 50.000 cells/well in a 96-well plate. Following 14 days of differentiation with CNTF, astrocyte media was conditioned for 48 hours and tested for the presence of IL6 without exogenous stimulation (R&D Systems). All samples were assayed in triplicate following the manufacturer’s instructions for cell culture supernatant. To confirm the functionality of the iPSCs-derived astrocytes, selected wells were stimulated with 1Ong/ml interleukin-1-β (IL-β) during 48 hours prior to IL-6 analysis. Optical density was measured at 450 nm with wavelength correction set to 540 nm using the Tecan Infinite Pro 200 plate reader. Standards were assayed in duplicates, and samples were assayed in triplicates. The corresponding plate was fixed and stained for normalization. Images were taken using Z1 Axio Observer Apotome Zeiss, and 8-10 pictures were taken per condition. GFAP-positive cells were manually counted, and **DAPI** was quantified using Fiji-lmageJ imaging. Recombinant human IL6 was added as a positive control.

### ROS Assay

Astrocyte progenitors were plated at a density of 50,000 cells/well in a white-walled, clear bottomed 96 well plate (Corning) and differentiated for 14 days with CNTF. ROS-Glo H2O2 (Promega) Homogenous (Lytic) protocol was followed according to the manufacturer’s instructions. Astrocytes and RTI treatments were conditioned in the Astromed media for 48 hours. The H2O2 substrate solution was added to the plate and treated for 4 hours before ROS Glo detection solution was added. The relative luminescence was recorded using a Tecan Infinite Pro 200 plate reader. Six replicates were run per condition.

### Glutathione Assay

GSH-GLO Glutathione Assay (Promega V6911) protocol for adherent mammalian cells was followed according to the manufacturer’s instructions to measure glutathione levels present in astrocytes. Cells were seeded at 50,000 cells/well on a white-walled 96 well plate. The medium was removed, 1X GSH-Glo reagent was added to the cells and incubated for 30 minutes at room temperature. Following the addition of Luciferin Detection reagent, relative luminescence was recorded using Tecan Infinite Pro 200 luminometer. Values were normalized to GFAP+ expressing cells.

### Glutamate Assay

The glutamate assay (ab83389) was used to detect extracellular glutamate levels. Astrocytes differentiated with CNTF for 14 days were plated in 96-well plates to confluency. The media was conditioned for 48 hours.

### Ca^2+^ Flux Assay

Rhod-4 No Wash Calcium Assay Kit was performed following the manufacturer’s instructions. iPSCs-derived neurons were grown in a 96-well plate and cultured in NG media. Media was changed 48 hours prior to the experiment.1 00ul of Rhod-4 Dye-Loading Solution was added per well and incubated at room temperature for 1 hour. Calcium flux assay was run by monitoring the fluorescence intensity at Ex/Em = 540/590 nm. Images were acquired for 20 seconds (approximately 600 frames) per field. For analysis, calcium fluorescence traces were retrieved using customized software, Netcal (Orlandi et al., 2017) based on Matlab software®. First, a manual selection of Regions of Interest (ROls) was carried out to track the activity of all the cells in the culture, both neurons and astrocytes. After analyzing the recording, further post-processing allows the division of the traces into 2 groups with different firing behavior: neurons that display a sharp and fast increase of fluorescence followed by a rapid decline in the wave function. These astrocytes have a slow progression in the fluorescence intensity and a similar slow decrease in intensity. The classification of the traces was done by training the Netcal program with the same parameters for each experiment. The analysis of each group highlights specific characteristics that all the traces in the group share together: Average fluorescence trace, Average number of bursts, Burst intensity, and Duration of bursts.

### Animals

Mecp2-KO mice^25^ (C57BL/6 background) were obtained from Mutant Mouse Resource and Research Centers (MMRRC). They were maintained on a 12-hour light/dark cycle with free access to food and water. All aspects of animal care and treatment were carried out according to the guidelines of the Experimental Animal Care Committee of Kyushu University.

### 3TC and d4T *in vivo* treatments

Mecp2-KO mice were administered 2 mg/ml 3TC (QA-4615, Combi-Blocks) or d4T (D3580, TCI) in drinking water starting at 4 weeks. All mice were observed daily and weighed once per week.

### Scoring of symptoms

Neurological symptoms of mice were scored as previously described^94^. In brief, mice were scored blind to genotype and treatment status, focusing on mobility, gait, hindlimb clasping, tremor, breathing, and general condition. Each of the 6 symptoms was scored from Oto 2; 0 corresponds to the symptom being absent or the same as in the WT, 1 to the symptom being present, and 2 to the symptom being severe.

### Open-field test

The open-field test was performed with 8-week-old WT, Mecp2-KO, and 3TC-treated Mecp2-KO male mice. Each mouse was placed in the corner of the open-field apparatus (50 cm × 50 cm × 30 cm), which was illuminated at 100 lux. Total distance traveled (cm) and average speed (cm/s) were calculated using the automated video tracking software TimeOFCR2 (O’Hara & Co., Tokyo, Japan). Data were collected for 10 min.

### Elevated plus maze test

The elevated plus maze consisted of two opposite open arms (25 cm × 5 cm × 0.25 cm) and two enclosed arms (25 cm × 5 cm × 15 cm), illuminated at 30 lux and placed 50 cm above the floor. Each mouse was placed in the central square of the maze (5 × 5 cm), facing one of the closed arms. Mouse behavior was recorded for 10 min, and the time spent on open arm was analyzed using the automated video tracking software TimeEP1 (O’Hara & Co., Tokyo, Japan). The percentages of time spent in open arm over total time were calculated.

### Three-chambers social interaction

The three-chambers apparatus is a rectangular non-transparent gray Plexiglas box (60 × 40 × 22 cm), partitioned into three equal-sized (20 × 40 × 22 cm) chambers by transparent Plexiglas plates with small openings (5 × 3 cm) allowing access into each chamber. One quarter-cylinder cage made of wire was placed in the corner of each of the side chambers. The box was illuminated at 20 lux. The subject mouse was placed into the central chamber and allowed to freely explore the three chambers for 5 minutes. In the first testing section, an age-matched stranger mouse (S1) of the same strain was placed in one of the cages, and the subject mouse was allowed to explore the three chambers without restrictions for 10 min. In the second section of preference for social novelty, a second stranger mouse (S2) was placed in the opposite cage, and the subject mouse was allowed to explore again for 10 minutes. All tests were video-recorded, and the time spent in the quadrant around the cage was analyzed using the automated video tracking software TimeCSI (O’Hara & Co., Tokyo, Japan). The social novelty index was calculated by the following formula: [Time spent interacting with stranger 2 in the second section)/ [Time spent interacting with stranger 1+stranger 2 in the second section).

### Golgi staining

Golgi staining was carried out using the FD Rapid GolgiStain Kit (FD NeuroTechnologies, Cat.# PK401) according to the manufacturer’s instructions. Briefly, mice at 8-9 weeks of age were euthanized, and brains were quickly removed and rinsed in double distilled water to remove blood from the surface. Brains were immersed in impregnation solution prepared by mixing equal volumes of Solutions A and B, and stored at RT for 2 weeks in the dark. They were then transferred into Solution C and stored at RT for 1 week in the dark, subsequently embedded in TFM tissue freezing medium (General Data Healthcare, Cat.# 15-183-1) and frozen at −80°C. Brains were sectioned serially at a thickness of 100 µm using a Cryostat (Leica, Cat.# CM1950). The sections were mounted on gelatin-coated glass slides and dried naturally overnight at RT. The dry-mounted sections were rinsed with double distilled water twice for 4 min. They were placed in the staining solution consisting of one part Solution D, one part Solution E, and 2 parts double distilled water for 10 min in the dark. They were further rinsed in double distilled water two times for 4 min and dehydrated in sequential rinses of 50, 75, and 95% ethanol for 4min each rinse. After further dehydration by 100% ethanol, the sections were cleaned in xylene three times for 4 min and finally coverslipped with Permount. Brightfield images of Golgi-stained neurons were acquired using a KEYENCE BZ-X700 fluorescence microscope.

### Measurement of dendritic soma size *in vivo*

Three mice of each group (WT, Mecp2-KO, and 3TC-treated Mecp2-KO) were examined. For images of Golgi-stained neurons, Z-series brightfield images of layer 2/3 pyramidal neurons in PFC were acquired at 20X objective using a KEYENCE BZ-X700 fluorescence microscope. To profile entire cell morphology in the Z direction, Z-series images were acquired every 1 µm. Sequential images of each field were reconstructed using the maximum intensity projection method. Soma size was quantified by tracing the outline of the cell body using lmageJ.

### Measurement of dendritic spine density *in vivo*

Three mice of each group (WT, Mecp2-KO, and 3TC-treated Mecp2-KO were examined. Z-series brightfield images of layer 2/3 pyramidal neurons in PFC were acquired every 0.2 µm at 100X -oil objective using KEYENCE BZ-X700 fluorescence microscope. Sequential images of each field were reconstructed using the maximum intensity projection method. The number of spines of secondary apical dendrites was manually counted, and then spine density per 10 µm was calculated.

### Quantification and Statistical analysis

Error bars for figures are standard error of the mean (S.E.M.) using GraphPad Prism v6 (GraphPad Software Inc). Technical replicates were used to determine the standard error.

### Publicly available RNA-seq data analysis

Publicly available RNA-seq data were obtained from the Gene Expression Omnibus (GSE159771 for spaceflight mouse retina; GSE199680 for mouse lung exposed to simulated GCR). The GCR exposure included five ion species: 1 GeV 1**H** (34.8%), 600 MeV/U 28Si (1.1%), 250 MeV/U 4He (18.0%), 350 MeV/U 160 (5.8%), 600 MeV/U 56Fe (1.0%), and 250 MeV 1H (39.2%). Raw FASTQ files were retrieved using SRA Toolkit. Adapter trimming and quality filtering were performed using Trimmomatic and Cutadapt. Reads were aligned to the mouse T2T genome (mhaESC) using STAR-2.7.11b, and transposable elements expression was quantified using TEtranscripts, with annotations from RepeatMasker. Differential expression analysis was conducted using DESeq2, and gene set enrichment analysis was performed with GSEA (gene set database: msigdb.v2023.2.Mm.symbols.gmt).

