## Supplementary material for "Dormant viral pathways underlie space-induced neural senescence": Table_S1

**Table 1. Radiation measurement of all spaceflights in his study**

| | RAD - Radiation Absorbed Dose | | REM - Radiation Environmental Monitoring | |
|---|---|---|---|---|
| **Spaceflight** | **GCRDose (mGy/d)** | **SAADose (mGy/d)** | **GCRDose (mGy/d)** | **SAADose (mGy/d)** |
| **SpX18 (mean)** | **NA** | **NA** | 0.158849118 | 0.152864706 |
| SD | **NA** | **NA** | 0.002032176 | 0.011423318 |
| Cumulative dosage | **NA** | **NA** | 5.40087 | 5.1974 |
| **SpX30 (mean)** | 0.128014634 | 0.095702195 | 0.136573171 | 0.061031707 |
| SD | 0.002211937 | 0.012065307 | 0.001963572 | 0.007339247 |
| Cumulative dosage | 5.2486 | 3.92379 | 5.5995 | 2.5023 |
| **SpX31 (mean)** | 0.118621395 | 0.084363721 | 0.126612093 | 0.085978605 |
| SD | 0.003259008 | 0.010492391 | 0.006561856 | 0.029145775 |
| Cumulative dosage | 5.103979008 | 3.638132391 | 5.44432 | 3.69708 |
