## Supplementary material for "Dormant viral pathways underlie space-induced neural senescence": Table_S2

**Table 2. Summary of analysis and statistical tests used in all experiments**

| Figure panel | Analysis | Statistical test | Groups being compared |
|---|---|---|---|
| **Fig. S1D** | Organoids diameter comparison | *T-Student* | control ISS<br>control ground |
| **Fig. 2A** | *Proteomics* | *T-Student / Volcano plot* | control ISS<br>control ground |
| **Fig. 2B** | *Proteomics* | T-Student / Reactome | control ISS<br>control ground |
| **Fig. 2C** | *Proteomics* | T-Student / Heatmap | control ISS<br>control ground |
| **Fig. 2D** | Proteomics | GO enrichment | control ISS<br>control ground |
| **Fig. 2E** | Proteomics | GO enrichment | control ISS<br>control ground |
| **Fig. 2F** | Transcriptomics | *T-Student / Heatmap* | control ISS<br>control ground |
| **Fig. 2G** | Transcriptomics | T-Student / Reactome | control ISS<br>control ground |
| **Fig. 2H** | Transcriptomics | T-Student / Reactome | control ISS<br>control ground |
| **Fig. 2I** | Transcriptomics | single cell | pool of control organoids in<br>ISS + ground |
| **Fig. 2J** | Transcriptomics | single cell | control ISS<br>control ground |

| | | | |
|---|---|---|---|
| **Fig. 2K** | Transcriptomics | *T-Student / single cell* | control ISS<br>control ground |
| **Fig. 2L** | Transcriptomics | *T-Student / single cell* | control ISS<br>control ground |
| **Fig. 2M** | Transcriptomics | T-Student / Heatmap | control ISS<br>control ground |
| **Fig. S2A** | Proteomics | *T-Student / Heatmap* | control ISS<br>control ground |
| **Fig. S2B** | Transcriptomics | *T-Student / Heatmap* | control ISS<br>control ground |
| **Fig. S2C** | Transcriptomics | *T-Student / Heatmap* | NASA TWINS DEGs<br>our study DEGs |
| **Fig. 3A** | Transcriptomics | *bulk/Heatmap* | ISS<br>control ground |
| **Fig. 3B** | Transcriptomics | *bulk/PcoA* | ISS<br>control ground |
| **Fig. 3C** | Transcriptomics | *bulk/Volcano* | ISS<br>control ground |
| **Fig. 3D** | Transcriptomics | *bulk/GSEA* | ISS<br>control ground |
| **Fig. 4A** | Proteomics | *T-Student / Volcano plot* | MeCP2-KO ISS<br>control ISS |
| **Fig. 4B** | Proteomics | T-Student / Reactome | MeCP2-KO ISS<br>control ISS |
| **Fig. 4C** | Transcriptomics | *T-Student / Heatmap* | MeCP2-KO ISS<br>control ISS |
| **Fig. S4A** | Proteomics | T-Student / Reactome | MeCP2-KO ISS<br>control ISS |
| **Fig. S4B** | Transcriptomics | *T-Student / Heatmap* | MeCP2-KO ISS<br>control ISS |

| | | | |
|---|---|---|---|
| **Fig. 5A** | ssDNA | one-way ANOVA | control |
| | | | MECP2-KO |
| | | | MECP2-KO + RTi |
| | | | MECP2-KO + NVP |
| **Fig. 5B** | IL-6 | one-way ANOVA | control |
| | | | control + RTi |
| | | | MECP2-KO |
| | | | MECP2-KO + RTi |
| **Fig. 5C** | SYN puncta/20um | one-way ANOVA | control (ACM / RTi -) |
| | | | control (ACM / RTi +) |
| | | | MECP2-KO (ACM / RTi -) |
| | | | MECP2-KO (ACM / RTi +) |
| **Fig. 5D** | Glutamate | one-way ANOVA | control |
| | | | MECP2-KO |
| | | | MECP2-KO + Rti |
| **Fig. 5E** | ROS Astrocytes | one-way ANOVA | control RTi - / NVP - |
| | | | control RTi + / NVP - |
| | | | control RTi - / NVP + |
| | | | MECP2-KO RTi - / NVP - |
| | | | MECP2-KO RTi + / NVP - |
| | | | MECP2-KO RTi - / NVP + |
| **Fig. 5F** | Glutathione | one-way ANOVA | control |
| | | | control + RTi |
| | | | MECP2-KO |
| | | | MECP2-KO + RTi |
| **Fig. 5G** | Cell body area (µm2) | one-way ANOVA | control |
| | | | MECP2-KO |
| | | | MECP2-KO + RTi |
| | | | MECP2-KO + NVP |
| **Fig. 5H** | Number of branch points | one-way ANOVA | control |
| | | | MECP2-KO |
| | | | MECP2-KO + RTi |
| | | | MECP2-KO + NVP |
| **Fig. 5I** | Colocalized puncta (per20µm) | one-way ANOVA | control |
| | | | MECP2-KO |
| | | | MECP2-KO + RTi |
| | | | MECP2-KO + NVP |

| | | | |
|---|---|---|---|
| **Fig. 5J** | Burst/5min | one-way ANOVA | control |
| | | | MECP2-KO |
| | | | MECP2-KO + RTi |
| | | | MECP2-KO + NVP |
| **Fig. 5K** | Firing rate (Hz) | one-way ANOVA | control |
| | | | MECP2-KO |
| | | | MECP2-KO + RTi |
| | | | MECP2-KO + NVP |
| **Fig. S5A** | Endogenous L1 | *T-Student* | control |
| | | | MeCP2-KO |
| **Fig. S5B** | Concentration | T-Student | control TNFα |
| | | | MECP2-KO TNFα |
| | | | control IL-4 |
| | | | MeCP2-KO IL-4 |
| | | | control IL-13 |
| | | | MeCP2-KO IL-13 |
| | | | control IL-10 |
| | | | MeCP2-KO IL-10 |
| **Fig. S5C** | IL-6 (pg/µL) | *T-Student* | control |
| | | | MeCP2-KO |
| **Fig. S5D** | IL-6 (pg/µL) | one-way ANOVA | control |
| | | | RTT |
| | | | rRRT |
| **Fig. S5E** | Gene expression | *T-Student* | control |
| | | | MeCP2-KO |
| **Fig. S5F** | Synapsin1 puncta (per 100µm dendrite) | *T-Student* | control |
| | | | control IL-6 |
| **Fig. S5G** | TDL/neuron (µm) | *T-Student* | control |
| | | | control IL-6 |
| **Fig. S5H** | % number of spikes | *T-Student* | before IL-6 |
| | | | after IL-6 |
| | | | Neurons control Ctrl |

| | | | |
|---|---|---|---|
| **Fig. S5J** | TDL/neuron (µm) | one-way ANOVA | Neurons control KO |
| | | | Neurons MECP2-KO KO |
| | | | Neurons MECP2-KO Ctrl |
| | | | Astrocytes control Ctrl |
| | | | Astrocytes control KO |
| | | | Astrocytes MECP2-KO KO |
| | | | Astrocytes MECP2-KO Ctrl |
| **Fig. S5K** | Segment/neuron (µm) | one-way ANOVA | Neurons control Ctrl |
| | | | Neurons control KO |
| | | | Neurons MECP2-KO KO |
| | | | Neurons MECP2-KO Ctrl |
| | | | Astrocytes control Ctrl |
| | | | Astrocytes control KO |
| | | | Astrocytes MECP2-KO KO |
| | | | Astrocytes MECP2-KO Ctrl |
| **Fig. S5L** | Spines/neuron (µm) | one-way ANOVA | Neurons control Ctrl |
| | | | Neurons control RTT |
| | | | Neurons RTT RTT |
| | | | Neurons RTT Ctrl |
| | | | Astrocytes control Ctrl |
| | | | Astrocytes control RTT |
| | | | Astrocytes RTT RTT |
| | | | Astrocytes RTT Ctrl |
| **Fig. S5M** | CXCL10 | one-way ANOVA | control |
| | | | MeCP2-KO |
| | | | MeCP2-KO + RTi |
| **Fig. S5N** | CXCL2 | one-way ANOVA | control |
| | | | MeCP2-KO |
| | | | MeCP2-KO + RTi |
| **Fig. S5O** | TMEN173 | one-way ANOVA | control |
| | | | MeCP2-KO |
| | | | MeCP2-KO + RTi |
| **Fig. S5P** | Relative gene | one-way ANOVA | control GS |
| | | | MeCP2-KO GS |
| | | | control GLT1 |
| | | | MeCP2-KO GLT1 |
| | | | control GLAST |

| | | | |
|---|---|---|---|
| Fig. S5P | expression | one-way ANOVA | MeCP2-KO GLAST |
| | | | control SLC26A7 |
| | | | MeCP2-KO SLC26A7 |
| | | | control SLC38A1 |
| | | | MeCP2-KO SLC38A1 |
| **Fig. S5R** | Dendrite lengh (µm) | one-way ANOVA | control |
| | | | MECP2-KO |
| | | | MECP2-KO + RTi |
| | | | MECP2-KO + NVP |
| **Fig. S5S** | Dendrite complexity | one-way ANOVA | control |
| | | | MECP2-KO |
| | | | MECP2-KO + RTi |
| | | | MECP2-KO + NVP |
| **Fig. 6A** | Scoring of neurological symptons (SUM) | one-way ANOVA | Mecp2-KO |
| | | | Mecp2-KO + 3TC |
| | | | Mecp2-KO + d4T |
| **Fig. 6B** | lifespan | log rank test | Mecp2-KO |
| | | | Mecp2-KO + 3TC |
| | | | Mecp2-KO + d4T |
| **Fig. 6C** | Total distance (cm) | one-way ANOVA | WT |
| | | | Mecp2-KO |
| | | | Mecp2-KO + 3TC |
| **Fig. 6D** | Average speed (cm/s) | one-way ANOVA | WT |
| | | | Mecp2-KO |
| | | | Mecp2-KO + 3TC |
| **Fig. 6E** | Opem arm (%) | one-way ANOVA | WT |
| | | | Mecp2-KO |
| | | | Mecp2-KO + 3TC |
| | | | WT |

| | | | |
|---|---|---|---|
| **Fig. 6F** | Social novelty index | one-way ANOVA | Mecp2-KO |
| | | | Mecp2-KO + 3TC |
| **Fig. 6G** | Soma size (µm) | one-way ANOVA | WT |
| | | | Mecp2-KO |
| | | | Mecp2-KO + 3TC |
| **Fig. 6H** | Spine density | one-way ANOVA | WT |
| | | | Mecp2-KO |
| | | | Mecp2-KO + 3TC |
| **Fig. 7F** | Spatial transcriptomics | wilcox | WT |
| | | | Mecp2-KO |
| **Fig. 7G** | Overlap genes | Fisher's Exact | ISS upregulated |
| | | | Mecp2-KO astrocyte upregulated |
| **Fig. 7J** | Intensity | one-way ANOVA | WT |
| | | | Mecp2-KO |
| | | | Mecp2-KO + 3TC |
| **Fig. S7A** | Spatial transcriptomics | wilcox | WT |
| | | | Mecp2-KO |
| **Fig. S7B** | Spatial transcriptomics | wilcox | Mecp2-KO |
| | | | Mecp2-KO + 3TC |

| Numbers of samples and replicates | | | |
|---|---|---|---|
| Number of samples (organoid, cells, mice) per experiment | Technical replicates | p value or FDR | Mission number |
| >10 | 3 | | |
| >10 | 3 | p≤ 0.05 | CRS-SpX25 |
| >10 | 3 | | |
| >10 | 3 | p≤ 0.01 | CRS-SpX25 |
| >10 | 3 | p≤ 0.01 | |
| >10 | 3 | | CRS-SpX25 |
| >10 | 3 | | |
| >10 | 3 | p≤ 0.01 | CRS-SpX31 |
| >10 | 3 | | |
| | 3 | FDR = 1.00E-68 | CRS-SpX25 |
| >10 | 3 | | |
| | 3 | p≤ 0.01 | CRS-SpX25 |
| >10 | 2 | | |
| >10 | 2 | p≤ 0.05 | CRS-SpX25 |
| >10 | 2 | | |
| >10 | 2 | p≤ 0.05 | CRS-SpX25 |
| >10 | 2 | | |
| >10 | 2 | p≤ 0.05 | CRS-SpX25 |
| >10 | 2 | not applicable | |
| >10 | 2 | not applicable | CRS-SpX25 |
| >10 | 2 | not applicable | |
| >10 | 2 | not applicable | CRS-SpX25 |

| | | | |
|---|---|---|---|
| >10 | 2 | not applicable | CRS-SpX25 |
| >10 | 2 | not applicable | |

| | | | |
|---|---|---|---|
| >10 | 2 | not applicable | CRS-SpX25 |
| >10 | 2 | not applicable | |

| | | | |
|---|---|---|---|
| 3 | 3 | $p \leq 0.05$ | CRS-SpX30 |
| 3 | 3 | | |

| | | | |
|---|---|---|---|
| >10 | 3 | $p \leq 0.01$ | CRS-SpX25 |
| >10 | 3 | | |

| | | | |
|---|---|---|---|
| >10 | 2 | $p \leq 0.05$ | CRS-SpX25 |
| >10 | 2 | | |

| | | | |
|---|---|---|---|
| 2 astronauts | 3 | $p \leq 0.05$ | Expeditions 43, 44, 45, and 46 |
| 3 organoids | 3 | | CRS-SpX25 |

| | | | |
|---|---|---|---|
| 8 | 1 | not applicable | not applicable |
| 7 | 1 | | |

| | | | |
|---|---|---|---|
| 8 | 1 | not applicable | not applicable |
| 7 | 1 | | |

| | | | |
|---|---|---|---|
| 8 | 1 | $p < 0.05$, fold change>1.5 | not applicable |
| 7 | 1 | | |

| | | | |
|---|---|---|---|
| 8 | 1 | Enrichment Score > 0.5 | not applicable |
| 7 | 1 | | |

| | | | |
|---|---|---|---|
| >10 | 3 | $p \leq 0.01$ | CRS-SpX25 |
| >10 | 3 | | |

| | | | |
|---|---|---|---|
| >10 | 3 | $p \leq 0.05$ | CRS-SpX25 |
| >10 | 3 | | |

| | | | |
|---|---|---|---|
| >10 | 3 | $p \leq 0.05$ | CRS-SpX30 |
| >10 | 3 | | |

| | | | |
|---|---|---|---|
| >10 | 2 | $p \leq 0.01$ | CRS-SpX25 |
| >10 | 2 | | |

| | | | |
|---|---|---|---|
| 3 | 3 | $p \leq 0.05$ | CRS-SpX30 |
| 3 | 3 | | |

| | | | |
|---|---|---|---|
| 6 | 3 | p<0.05 (*) | not applicable |
| 6 | 3 | | |
| 5 | 3 | | |
| 4 | 3 | | |

| | | | |
|---|---|---|---|
| 3 | 3 | p<0.0001 (****) | not applicable |
| 3 | 3 | | |
| 3 | 3 | | |
| 3 | 3 | | |

| | | | |
|---|---|---|---|
| >10 | 3 | p<0.01 (**) and p<0.0001 (****) | not applicable |
| >10 | 3 | | |
| >10 | 3 | | |
| >10 | 3 | | |

| | | | |
|---|---|---|---|
| 10 | 3 | p<0.01 (**) and p<0.001 (***) | not applicable |
| 10 | 3 | | |
| 10 | 3 | | |

| | | | |
|---|---|---|---|
| >10 | 3 | p<0.0001 (****) | not applicable |
| >10 | 3 | | |
| >10 | 3 | | |
| >10 | 3 | | |
| >10 | 3 | | |
| >10 | 3 | | |

| | | | |
|---|---|---|---|
| 6 | 3 | p<0.0001 (****) | not applicable |
| 6 | 3 | | |
| 6 | 3 | | |
| 6 | 3 | | |

| | | | |
|---|---|---|---|
| >10 | 3 | p<0.05 (*), p<0.01 (**), and p<0.0001 (****) | not applicable |
| >10 | 3 | | |
| >10 | 3 | | |
| >10 | 3 | | |

| | | | |
|---|---|---|---|
| >10 | 3 | p<0.01 (**), and p<0.0001 (****) | not applicable |
| >10 | 3 | | |
| >10 | 3 | | |
| >10 | 3 | | |

| | | | |
|---|---|---|---|
| >10 | 3 | p<0.0001 (****) | not applicable |
| >10 | 3 | | |
| >10 | 3 | | |
| >10 | 3 | | |

| | | | |
|---|---|---|---|
| >10 | 3 | p<0.01 (**) and p<0.0001 (****) | not applicable |
| >10 | 3 | | |
| >10 | 3 | | |
| >10 | 3 | | |
| >10 | 3 | p<0.01 (**), and p<0.001 (***) | not applicable |
| >10 | 3 | | |
| >10 | 3 | | |
| >10 | 3 | | |
| 3 | 3 | p<0.0001 (****) | not applicable |
| 3 | 3 | | |
| 3 | 3 | p<0.05 (*), p<0.01 (**), and p<0.0001 (****) | not applicable |
| 3 | 3 | | |
| 3 | 3 | | |
| 3 | 3 | | |
| 3 | 3 | | |
| 3 | 3 | | |
| 3 | 3 | | |
| 3 | 3 | | |
| 3 | 3 | p<0.001 (***) | not applicable |
| 3 | 3 | | |
| 5 | 3 | p<0.0001 (****) | not applicable |
| 10 | 3 | | |
| 8 | 3 | | |
| 3 | 3 | p<0.05 (*) | not applicable |
| 3 | 3 | | |
| 4 | 3 | p<0.0001 (****) | not applicable |
| 10 | 3 | | |
| 10 | 3 | p<0.001 (***) | not applicable |
| 10 | 3 | | |
| 6 | 3 | p<0.01 (**) | not applicable |
| 6 | 3 | | |
| >10 | 3 | | |

| | | | |
|---|---|---|---|
| >10 | 3 | | |
| >10 | 3 | | |
| >10 | 3 | | |
| >10 | 3 | | |
| >10 | 3 | | |
| >10 | 3 | | |
| >10 | 3 | p<0.01 (**), and p<0.001 (***) | not applicable |
| >10 | 3 | | |
| >10 | 3 | | |
| >10 | 3 | | |
| >10 | 3 | | |
| >10 | 3 | | |
| >10 | 3 | | |
| >10 | 3 | | |
| >10 | 3 | p<0.001 (***) | not applicable |
| >10 | 3 | | |
| >10 | 3 | | |
| >10 | 3 | | |
| >10 | 3 | | |
| >10 | 3 | | |
| >10 | 3 | | |
| >10 | 3 | | |
| >10 | 3 | p<0.01 (**), and p<0.001 (***) | not applicable |
| 3 | 3 | | |
| 3 | 3 | | |
| 3 | 3 | p<0.05 (*) | not applicable |
| 3 | 3 | | |
| 3 | 3 | | |
| 3 | 3 | p<0.05 (*) | not applicable |
| 3 | 3 | | |
| 3 | 3 | | |
| 3 | 3 | p<0.0001 (****) | Do not apply |
| 3 | 3 | | |
| 3 | 3 | | |
| 3 | 3 | | |
| 3 | 3 | | |
| 3 | 3 | | |

| | | | |
|---|---|---|---|
| 3 | 3 | p<0.01 (**), and p<0.0001 (****) | not applicable |
| 3 | 3 | | |
| 3 | 3 | | |
| 3 | 3 | | |
| 3 | 3 | | |
| >10 | 3 | p<0.05 (*), p<0.01 (**), and p<0.0001 (****) | not applicable |
| >10 | 3 | | |
| >10 | 3 | | |
| >10 | 3 | | |
| >10 | 3 | p<0.001 (***) | not applicable |
| >10 | 3 | | |
| >10 | 3 | | |
| >10 | 3 | | |
| 15 | 1 | p<0.05 (*), p<0.01 (**), p<0.001 (***) and p<0.0001 (****) | not applicable |
| 13 | 1 | | |
| 13 | 1 | | |
| 20 | 1 | p<0.05 (*), and p<0.01 (**) | not applicable |
| 14 | 1 | | |
| 13 | 1 | | |
| 13 | 1 | p<0.05 (*), and p<0.0001 (****) | not applicable |
| 19 | 1 | | |
| 14 | 1 | | |
| 13 | 1 | p<0.05 (*), and p<0.0001 (****) | not applicable |
| 19 | 1 | | |
| 14 | 1 | | |
| 13 | 1 | p<0.0001 (****) | not applicable |
| 18 | 1 | | |
| 14 | 1 | | |
| 13 | 1 | | |

| | | | |
|---|---|---|---|
| 13 | 1 | p<0.05 (*) | not applicable |
| 12 | 1 | | |
| >10 | 3 | p<0.01 (***) and p<0.001 (****) | not applicable |
| >10 | 3 | | |
| >10 | 3 | | |
| >4 | 3 | p<0.05 (*) and p<0.001 (***) | not applicable |
| >4 | 3 | | |
| >4 | 3 | | |
| >10 | 1 | p≤ 0.05 | not applicable |
| >10 | 1 | | |
| >10 | 1 | p≤ 0.05 | not applicable |
| >10 | 1 | | |
| >10 | 3 | p<0.001 (****) | not applicable |
| >10 | 3 | | |
| >10 | 3 | | |
| >10 | 1 | p≤ 0.05 | not applicable |
| >10 | 1 | | |
| >10 | 1 | p≤ 0.05 | not applicable |
| >10 | 1 | | |
