## Supplementary material for "Dormant viral pathways underlie space-induced neural senescence": Table_S3

| Significant | #NAME? | Difference | PG.Protein | PG.Protein | PG.Genes | PG.Protein | Symbol co | Symbol typ | Symbol siz |
|---|---|---|---|---|---|---|---|---|---|
| | 0.299066 | 0.057296 | A0A0B4J2I | A0A0B4J2I | GATD3B;G | Putative gl | Color2 [Da | 1 | 4 |
| | 0 | NaN | A0A0B4J2I | A0A0B4J2I | PIGBOS1 | Protein PI( | Color2 [Da | 1 | 4 |
| | 0 | NaN | A0A0U1RF | A0A0U1RF | NBDY | Negative r | Color2 [Da | 1 | 4 |
| | 0.388948 | 0.269781 | A0A0U1RF | A0A0U1RF | MMP24OS | Protein MI | Color2 [Da | 1 | 4 |
| + | 2.100954 | 0.99824 | A0A1B0GU | A0A1B0GU | BRD3OS | Putative u | Color2 [Da | 1 | 4 |
| | 0.166302 | 0.216235 | A0A2Z4LIS | A0A2Z4LIS | FOXO3B | Forkhead I | Color2 [Da | 1 | 4 |
| + | 2.270267 | -1.62145 | A0A3B3IR\ | A0A3B3IR\ | MCTS2 | Malignant | Color2 [Da | 1 | 4 |
| | 0 | NaN | A0AV96 | A0AV96 | RBM47 | RNA-bindi | Color2 [Da | 1 | 4 |
| | 0 | -0.37376 | A0AVF1 | A0AVF1 | TTC26 | Intraflagell | Color2 [Da | 1 | 4 |
| + | 3.634882 | -1.18349 | A0AVT1 | A0AVT1 | UBA6 | Ubiquitin-l | Color2 [Da | 1 | 4 |
| + | 1.496817 | -0.44768 | A0FGR8 | A0FGR8 | ESYT2 | Extended s | Color2 [Da | 1 | 4 |
| + | 0.890233 | -0.71906 | A0JNW5 | A0JNW5 | UHRF1BP1 | UHRF1-bin | Color2 [Da | 1 | 4 |
| + | 3.650331 | 1.571396 | A0MZ66 | A0MZ66 | SHTN1 | Shootin-1 | Color2 [Da | 1 | 4 |
| | 0 | 1.587512 | A1KXE4 | A1KXE4 | FAM168B | Myelin-ass | Color2 [Da | 1 | 4 |
| | 0.333991 | 0.227264 | A1L020 | A1L020 | MEX3A | RNA-bindi | Color2 [Da | 1 | 4 |
| + | 1.040957 | -0.45296 | A1L0T0 | A1L0T0 | ILVBL | 2-hydroxy | Color2 [Da | 1 | 4 |
| | 0 | NaN | A1L157 | A1L157 | TSPAN11 | Tetraspani | Color2 [Da | 1 | 4 |
| + | 1.067695 | 0.631742 | A1L188 | A1L188 | NDUFAF8 | NADH deh | Color2 [Da | 1 | 4 |
| + | 1.735607 | -0.57067 | A1X283 | A1X283 | SH3PXD2B | SH3 and P. | Color2 [Da | 1 | 4 |
| | 0 | 0.204012 | A1Z1Q3 | A1Z1Q3 | MACROD2 | ADP-ribose | Color2 [Da | 1 | 4 |
| | 0.619447 | 0.955244 | A2A3N6 | A2A3N6 | PIPSL | Putative PI | Color2 [Da | 1 | 4 |
| + | 0.963074 | 1.019785 | A2AJT9 | A2AJT9 | BCLAF3 | BCLAF1 an | Color2 [Da | 1 | 4 |
| + | 2.532215 | -0.64615 | A2RRP1 | A2RRP1 | NBAS | NBAS subu | Color2 [Da | 1 | 4 |
| | 0.611004 | -0.14782 | A2RTX5 | A2RTX5 | TARS3 | Threonine- | Color2 [Da | 1 | 4 |
| + | 2.076433 | -0.79157 | A3KMH1 | A3KMH1 | VWA8 | von Willeb | Color2 [Da | 1 | 4 |
| + | 1.022556 | -0.35298 | A3KN83 | A3KN83 | SBNO1 | Protein str | Color2 [Da | 1 | 4 |
| | 0.523823 | 0.169783 | A4D161 | A4D161 | FAM221A | Protein FA | Color2 [Da | 1 | 4 |
| + | 0.844873 | 0.703641 | A4D1E9 | A4D1E9 | GTPBP10 | GTP-bindir | Color2 [Da | 1 | 4 |
| + | 0.975879 | -0.41523 | A4D1P6 | A4D1P6 | WDR91 | WD repeat | Color2 [Da | 1 | 4 |
| | 0 | -0.84939 | A4D2B0 | A4D2B0 | MBLAC1 | Metallo-be | Color2 [Da | 1 | 4 |
| + | 1.073743 | 0.52118 | A5D8V6 | A5D8V6 | VPS37C | Vacuolar p | Color2 [Da | 1 | 4 |
| + | 2.073976 | 0.816865 | A5PKW4 | A5PKW4 | PSD | PH and SE( | Color2 [Da | 1 | 4 |
| | 0 | NaN | A5PLL7 | A5PLL7 | PEDS1 | Plasmanyl | Color2 [Da | 1 | 4 |
| + | 2.686526 | -0.87026 | A5YKK6 | A5YKK6 | CNOT1 | CCR4-NOT | Color2 [Da | 1 | 4 |
| + | 3.141672 | 3.033856 | A6NCE7;Q | A6NCE7;Q | MAP1LC3E | Microtubu | Color2 [Da | 1 | 4 |
| + | 1.890141 | 3.062805 | A6NCW0;/ | A6NCW0;/ | USP17L3;U | Ubiquitin c | Color2 [Da | 1 | 4 |
| + | 1.382652 | 0.90685 | A6NDB9 | A6NDB9 | PALM3 | Paralemm | Color2 [Da | 1 | 4 |
| | 0 | 0.539616 | A6NDG6 | A6NDG6 | PGP | Glycerol-3- | Color2 [Da | 1 | 4 |
| + | 1.250776 | -0.75455 | A6NDU8 | A6NDU8 | RIMOC1 | RAB7A-inte | Color2 [Da | 1 | 4 |
| | 0.481952 | 0.117369 | A6NE02 | A6NE02 | BTBD17 | BTB/POZ d | Color2 [Da | 1 | 4 |
| | 0.15386 | 0.173303 | A6NFI3 | A6NFI3 | ZNF316 | Zinc finger | Color2 [Da | 1 | 4 |
| + | 2.040435 | 2.782016 | A6NFN3 | A6NFN3 | RBFOX3 | RNA bindir | Color2 [Da | 1 | 4 |
| + | 1.648921 | 1.994894 | A6NGB9 | A6NGB9 | WIPF3 | WAS/WAS | Color2 [Da | 1 | 4 |
| + | 1.182004 | 0.543834 | A6NGN9 | A6NGN9 | IGLON5 | IgLON fam | Color2 [Da | 1 | 4 |
| + | 2.393978 | -0.55939 | A6NHL2 | A6NHL2 | TUBAL3 | Tubulin alp | Color2 [Da | 1 | 4 |
| + | 3.155811 | 1.481319 | A6NHQ2 | A6NHQ2 | FBLL1 | rRNA/tRN/ | Color2 [Da | 1 | 4 |

-Bruker Confidential-

| | | | | | | | | | |
|---|---|---|---|---|---|---|---|---|---|
| + | 2.355923 | -0.90211 | A6NHR9 | A6NHR9 | SMCHD1 | Structural | Color2 [Da | 1 | 4 |
| | 0.72128 | 0.153662 | A6NHX0 | A6NHX0 | CASTOR2 | Cytosolic a | Color2 [Da | 1 | 4 |
| | 0.598495 | 1.689575 | A6NI47;Q6 | A6NI47;Q6 | POTEM;PC | Putative P( | Color2 [Da | 1 | 4 |
| + | 2.413641 | 1.269435 | A6NIH7 | A6NIH7 | UNC119B | Protein un | Color2 [Da | 1 | 4 |
| + | 0.997626 | 0.885016 | A6NJ78 | A6NJ78 | METTL15 | 12S rRNA I | Color2 [Da | 1 | 4 |
| | 0.447219 | 0.17351 | A6NKD9 | A6NKD9 | CCDC85C | Coiled-coil | Color2 [Da | 1 | 4 |
| | 0 | NaN | A6NKG5 | A6NKG5 | RTL1 | Retrotrans | Color2 [Da | 1 | 4 |
| | 0 | NaN | A6NKN8 | A6NKN8 | PCP4L1 | Purkinje ce | Color2 [Da | 1 | 4 |
| + | 2.119525 | 0.874879 | A6ZKI3 | A6ZKI3 | RTL8C | Retrotrans | Color2 [Da | 1 | 4 |
| + | 1.53074 | 0.566989 | A7E2V4 | A7E2V4 | ZSWIM8 | Zinc finger | Color2 [Da | 1 | 4 |
| | 0.16225 | 0.136316 | A7KAX9 | A7KAX9 | ARHGAP32 | Rho GTPas | Color2 [Da | 1 | 4 |
| | 0.064857 | 0.043378 | A7MBM2 | A7MBM2 | DISP2 | Protein dis | Color2 [Da | 1 | 4 |
| + | 1.177578 | -0.66012 | A7MD48 | A7MD48 | SRRM4 | Serine/arg | Color2 [Da | 1 | 4 |
| | 0 | NaN | A8K2U0 | A8K2U0 | A2ML1 | Alpha-2-m | Color2 [Da | 1 | 4 |
| | 0 | NaN | A8MTJ3;P1 | A8MTJ3;P1 | GNAT3;GN | Guanine n | Color2 [Da | 1 | 4 |
| + | 1.743172 | 0.405529 | A8MVW0 | A8MVW0 | FAM171A2 | Protein FA | Color2 [Da | 1 | 4 |
| + | 1.290898 | 0.610665 | A8MW92 | A8MW92 | PHF20L1 | PHD finger | Color2 [Da | 1 | 4 |
| + | 1.294831 | 1.001955 | A8MWD9; | A8MWD9; | SNRPGP15 | Putative sr | Color2 [Da | 1 | 4 |
| | 0 | NaN | A8MYZ6 | A8MYZ6 | FOXO6 | Forkhead I | Color2 [Da | 1 | 4 |
| + | 0.901279 | -0.79343 | A9UHW6 | A9UHW6 | MIF4GD | MIF4G dor | Color2 [Da | 1 | 4 |
| | 0 | NaN | A9Z1Z3 | A9Z1Z3 | FER1L4 | Fer-1-like | Color2 [Da | 1 | 4 |
| | 0 | 1.592289 | B7ZAP0 | B7ZAP0 | RABGAP1L | Rab GTPas | Color2 [Da | 1 | 4 |
| | 0 | NaN | B7ZAQ6;P( | B7ZAQ6;P( | GPR89A;G | Golgi pH re | Color2 [Da | 1 | 4 |
| + | 1.802543 | 1.65305 | C4AMC7;C | C4AMC7;C | WASH3P;W | Putative W | Color2 [Da | 1 | 4 |
| | 0 | -0.90363 | C9J798;O4 | C9J798;O4 | RASA4B;R/ | Ras GTPas | Color2 [Da | 1 | 4 |
| + | 2.293161 | 1.717101 | C9J7I0 | C9J7I0 | UMAD1 | UBAP1-MV | Color2 [Da | 1 | 4 |
| | 0.125653 | -0.11258 | C9JLW8 | C9JLW8 | MCRIP1 | Mapk-regu | Color2 [Da | 1 | 4 |
| | 0.519192 | 1.401182 | C9JRZ8 | C9JRZ8 | AKR1B15 | Aldo-keto | Color2 [Da | 1 | 4 |
| + | 3.519358 | 2.068393 | E9PAV3;Q1 | E9PAV3;Q1 | NACA | Nascent pc | Color2 [Da | 1 | 4 |
| + | 2.489889 | 1.503535 | E9PRG8 | E9PRG8 | C11orf98 | Uncharact | Color2 [Da | 1 | 4 |
| | 0.193972 | 0.549091 | F8WCM5;I | F8WCM5;I | INS-IGF2;II | Insulin, iso | Color2 [Da | 1 | 4 |
| | 0 | NaN | H3BS89 | H3BS89 | TMEM178 | Transmem | Color2 [Da | 1 | 4 |
| + | 1.554144 | 1.758169 | L0R6Q1 | L0R6Q1 | SLC35A4 | SLC35A4 u | Color2 [Da | 1 | 4 |
| + | 2.698104 | 1.378172 | L0R819 | L0R819 | ASDURF | ASNSD1 u | Color2 [Da | 1 | 4 |
| + | 3.813378 | 2.388368 | O00115 | O00115 | DNASE2 | Deoxyribo | Color2 [Da | 1 | 4 |
| + | 3.957175 | -1.16563 | O00116 | O00116 | AGPS | Alkyldihyd | Color2 [Da | 1 | 4 |
| + | 1.453826 | -0.43437 | O00139 | O00139 | KIF2A | Kinesin-lik | Color2 [Da | 1 | 4 |
| | 0 | NaN | O00142 | O00142 | TK2 | Thymidine | Color2 [Da | 1 | 4 |
| + | 3.238186 | -1.35206 | O00148 | O00148 | DDX39A | ATP-deper | Color2 [Da | 1 | 4 |
| + | 1.095033 | 0.534473 | O00151 | O00151 | PDLIM1 | PDZ and LI | Color2 [Da | 1 | 4 |
| + | 3.364867 | 0.625831 | O00154 | O00154 | ACOT7 | Cytosolic a | Color2 [Da | 1 | 4 |
| | 0.217993 | -0.12689 | O00159 | O00159 | MYO1C | Unconvent | Color2 [Da | 1 | 4 |
| + | 2.386396 | 1.358979 | O00161 | O00161 | SNAP23 | Synaptoso | Color2 [Da | 1 | 4 |
| + | 2.175191 | 0.810217 | O00165 | O00165 | HAX1 | HCLS1-ass | Color2 [Da | 1 | 4 |
| | 0.776265 | -0.22686 | O00170 | O00170 | AIP | AH recept | Color2 [Da | 1 | 4 |
| + | 0.935803 | -0.50855 | O00178 | O00178 | GTPBP1 | GTP-bindir | Color2 [Da | 1 | 4 |
| | 0.477523 | -0.26757 | O00186 | O00186 | STXBP3 | Syntaxin-b | Color2 [Da | 1 | 4 |

| | | | | | | | | | |
|---|---|---|---|---|---|---|---|---|---|
| | 1.399913 | -0.18268 | O00192 | O00192 | ARVCF | Armadillo | Color2 [Da | 1 | 4 |
| + | 2.088846 | 2.309632 | O00193 | O00193 | SMAP | Small acidi | Color2 [Da | 1 | 4 |
| + | 2.01296 | 1.327701 | O00194 | O00194 | RAB27B | Ras-relate | Color2 [Da | 1 | 4 |
| | 0.40013 | -0.06274 | O00203 | O00203 | AP3B1 | AP-3 comp | Color2 [Da | 1 | 4 |
| | 1.535034 | 0.162284 | O00213 | O00213 | APBB1 | Amyloid b | Color2 [Da | 1 | 4 |
| + | 3.555703 | 0.994631 | O00217 | O00217 | NDUFS8 | NADH deh | Color2 [Da | 1 | 4 |
| + | 2.3031 | -0.38631 | O00231 | O00231 | PSMD11 | 26S protea | Color2 [Da | 1 | 4 |
| + | 3.140881 | -0.70295 | O00232 | O00232 | PSMD12 | 26S protea | Color2 [Da | 1 | 4 |
| + | 1.595165 | 1.237686 | O00233 | O00233 | PSMD9 | 26S protea | Color2 [Da | 1 | 4 |
| + | 3.097813 | 3.037525 | O00244 | O00244 | ATOX1 | Copper tra | Color2 [Da | 1 | 4 |
| + | 2.432351 | -0.95279 | O00255 | O00255 | MEN1 | Menin | Color2 [Da | 1 | 4 |
| | 0.786981 | 0.270444 | O00257 | O00257 | CBX4 | E3 SUMO-| | Color2 [Da | 1 | 4 |
| | 0 | NaN | O00258 | O00258 | GET1 | Guided en | Color2 [Da | 1 | 4 |
| + | 2.54209 | 0.870886 | O00264 | O00264 | PGRMC1 | Membrane | Color2 [Da | 1 | 4 |
| + | 1.399905 | 0.226623 | O00267 | O00267 | SUPT5H | Transcripti | Color2 [Da | 1 | 4 |
| + | 1.877419 | 0.62765 | O00273 | O00273 | DFFA | DNA fragm | Color2 [Da | 1 | 4 |
| + | 1.728001 | 0.694285 | O00287 | O00287 | RFXAP | Regulatory | Color2 [Da | 1 | 4 |
| + | 2.111231 | -0.41687 | O00291 | O00291 | HIP1 | Huntingtin | Color2 [Da | 1 | 4 |
| + | 3.685659 | -0.90313 | O00299 | O00299 | CLIC1 | Chloride ir | Color2 [Da | 1 | 4 |
| | 0.15615 | -0.07524 | O00303 | O00303 | EIF3F | Eukaryotic | Color2 [Da | 1 | 4 |
| + | 1.553255 | 0.873943 | O00305 | O00305 | CACNB4 | Voltage-de | Color2 [Da | 1 | 4 |
| + | 1.706855 | 1.275965 | O00308 | O00308 | WWP2 | NEDD4-lik | Color2 [Da | 1 | 4 |
| + | 2.098866 | 0.88368 | O00330 | O00330 | PDHX | Pyruvate d | Color2 [Da | 1 | 4 |
| | 0 | NaN | O00339 | O00339 | MATN2 | Matrilin-2 | Color2 [Da | 1 | 4 |
| | 0.046299 | -0.02867 | O00399 | O00399 | DCTN6 | Dynactin s | Color2 [Da | 1 | 4 |
| + | 1.714014 | 0.357294 | O00401 | O00401 | WASL | Neural Wis | Color2 [Da | 1 | 4 |
| | 0 | -0.5066 | O00408 | O00408 | PDE2A | cGMP-dep | Color2 [Da | 1 | 4 |
| + | 3.615151 | -1.84187 | O00410 | O00410 | IPO5 | Importin-5 | Color2 [Da | 1 | 4 |
| | 0 | NaN | O00411 | O00411 | POLRMT | DNA-direc | Color2 [Da | 1 | 4 |
| + | 2.275045 | 1.266575 | O00422 | O00422 | SAP18 | Histone de | Color2 [Da | 1 | 4 |
| + | 1.176629 | -0.9133 | O00423 | O00423 | EML1 | Echinoder | Color2 [Da | 1 | 4 |
| + | 2.871209 | -0.97307 | O00425 | O00425 | IGF2BP3 | Insulin-like | Color2 [Da | 1 | 4 |
| + | 3.286916 | 0.711596 | O00429 | O00429 | DNM1L | Dynamin-1 | Color2 [Da | 1 | 4 |
| | 0.790779 | -0.32745 | O00442 | O00442 | RTCA | RNA 3'-ter | Color2 [Da | 1 | 4 |
| + | 1.490183 | -0.44488 | O00443 | O00443 | PIK3C2A | Phosphati | Color2 [Da | 1 | 4 |
| | 0.17385 | 0.412163 | O00445 | O00445 | SYT5 | Synaptota | Color2 [Da | 1 | 4 |
| | 0 | -0.2376 | O00451 | O00451 | GFRA2 | GDNF fam | Color2 [Da | 1 | 4 |
| + | 2.615846 | -1.54343 | O00458 | O00458 | IFRD1 | Interferon | Color2 [Da | 1 | 4 |
| + | 2.421201 | 0.999592 | O00459 | O00459 | PIK3R2 | Phosphati | Color2 [Da | 1 | 4 |
| + | 1.956199 | -0.59022 | O00461 | O00461 | GOLIM4 | Golgi integ | Color2 [Da | 1 | 4 |
| | 0.003437 | -0.00131 | O00462 | O00462 | MANBA | Beta-manr | Color2 [Da | 1 | 4 |
| + | 1.458916 | -0.33487 | O00468 | O00468 | AGRN | Agrin | Color2 [Da | 1 | 4 |
| | 0 | NaN | O00469 | O00469 | PLOD2 | Procollage | Color2 [Da | 1 | 4 |
| + | 2.472811 | 0.624805 | O00470 | O00470 | MEIS1 | Homeobo | Color2 [Da | 1 | 4 |
| | 0.375924 | -0.34046 | O00471 | O00471 | EXOC5 | Exocyst co | Color2 [Da | 1 | 4 |
| | 0 | 0.848275 | O00479 | O00479 | HMGN4 | High mobi | Color2 [Da | 1 | 4 |
| + | 1.002354 | 0.473911 | O00483 | O00483 | NDUFA4 | Cytochron | Color2 [Da | 1 | 4 |

| | | | | | | | | | |
|---|---|---|---|---|---|---|---|---|---|
| | 0.634083 | 0.211908 | O00487 | O00487 | PSMD14 | 26S protea | Color2 [Da | 1 | 4 |
| | 0.192878 | 0.104804 | O00488 | O00488 | ZNF593 | Zinc finger | Color2 [Da | 1 | 4 |
| + | 4.394442 | 1.993045 | O00499 | O00499 | BIN1 | Myc box-d | Color2 [Da | 1 | 4 |
| + | 1.473911 | -0.64033 | O00505 | O00505 | KPNA3 | Importin s | Color2 [Da | 1 | 4 |
| + | 2.634871 | -1.49703 | O00506 | O00506 | STK25 | Serine/thr | Color2 [Da | 1 | 4 |
| | 0.171152 | -0.13598 | O00512 | O00512 | BCL9 | B-cell CLL/ | Color2 [Da | 1 | 4 |
| | 0.61 | 0.474928 | O00519 | O00519 | FAAH | Fatty-acid | Color2 [Da | 1 | 4 |
| | 0 | NaN | O00533 | O00533 | CHL1 | Neural cell | Color2 [Da | 1 | 4 |
| | 0.269165 | -0.09195 | O00534 | O00534 | VWA5A | von Willeb | Color2 [Da | 1 | 4 |
| + | 0.759003 | -1.06053 | O00541 | O00541 | PES1 | Pescadillo | Color2 [Da | 1 | 4 |
| + | 1.427562 | 0.646894 | O00560 | O00560 | SDCBP | Syntenin-1 | Color2 [Da | 1 | 4 |
| | 0.016316 | 0.007284 | O00562 | O00562 | PITPNM1 | Membrane | Color2 [Da | 1 | 4 |
| + | 1.353049 | -0.47131 | O00566 | O00566 | MPHOSPH | U3 small n | Color2 [Da | 1 | 4 |
| + | 1.537097 | -0.49865 | O00567 | O00567 | NOP56 | Nucleolar | Color2 [Da | 1 | 4 |
| + | 3.6097 | -1.21612 | O00571 | O00571 | DDX3X | ATP-deper | Color2 [Da | 1 | 4 |
| + | 2.47609 | 1.579966 | O00584 | O00584 | RNASET2 | Ribonuclea | Color2 [Da | 1 | 4 |
| + | 2.122863 | -1.19503 | O00592 | O00592 | PODXL | Podocalyxi | Color2 [Da | 1 | 4 |
| | 0 | -1.27895 | O00622 | O00622 | CCN1 | CCN family | Color2 [Da | 1 | 4 |
| | 0 | -2.05058 | O00623 | O00623 | PEX12 | Peroxisom | Color2 [Da | 1 | 4 |
| + | 2.243225 | 0.912383 | O00625 | O00625 | PIR | Pirin | Color2 [Da | 1 | 4 |
| + | 1.500368 | -1.33165 | O00629 | O00629 | KPNA4 | Importin s | Color2 [Da | 1 | 4 |
| | 0 | NaN | O00634 | O00634 | NTN3 | Netrin-3 | Color2 [Da | 1 | 4 |
| | 0.280006 | -0.15467 | O00712 | O00712 | NFIB | Nuclear fa | Color2 [Da | 1 | 4 |
| | 0 | NaN | O00716 | O00716 | E2F3 | Transcripti | Color2 [Da | 1 | 4 |
| | 0.387417 | -0.54951 | O00743 | O00743 | PPP6C | Serine/thr | Color2 [Da | 1 | 4 |
| + | 2.269865 | 0.422077 | O00746 | O00746 | NME4 | Nucleoside | Color2 [Da | 1 | 4 |
| | 0.473531 | 1.131312 | O00748 | O00748 | CES2 | Cocaine es | Color2 [Da | 1 | 4 |
| | 0.145447 | -0.13731 | O00750 | O00750 | PIK3C2B | Phosphatic | Color2 [Da | 1 | 4 |
| + | 3.160104 | 1.089663 | O00754 | O00754 | MAN2B1 | Lysosomal | Color2 [Da | 1 | 4 |
| + | 0.949439 | 0.378871 | O00755 | O00755 | WNT7A | Protein Wi | Color2 [Da | 1 | 4 |
| | 0.359718 | 0.24552 | O00762 | O00762 | UBE2C | Ubiquitin-c | Color2 [Da | 1 | 4 |
| + | 2.594374 | 1.059338 | O00764 | O00764 | PDXK | Pyridoxal k | Color2 [Da | 1 | 4 |
| | 0 | NaN | O00767 | O00767 | SCD | Stearoyl-C | Color2 [Da | 1 | 4 |
| | 0 | NaN | O14493;O | O14493;O | CLDN4;CLD | Claudin-4; | Color2 [Da | 1 | 4 |
| | 0.158519 | 0.040318 | O14497 | O14497 | ARID1A | AT-rich int | Color2 [Da | 1 | 4 |
| | 0 | 1.249571 | O14508 | O14508 | SOCS2 | Suppresso | Color2 [Da | 1 | 4 |
| + | 1.435783 | 0.705489 | O14523 | O14523 | C2CD2L | Phospholip | Color2 [Da | 1 | 4 |
| | 0 | NaN | O14525 | O14525 | ASTN1 | Astrotactir | Color2 [Da | 1 | 4 |
| | 0 | 0.693691 | O14526 | O14526 | FCHO1 | F-BAR don | Color2 [Da | 1 | 4 |
| + | 2.015097 | 1.297873 | O14530 | O14530 | TXNDC9 | Thioredoxi | Color2 [Da | 1 | 4 |
| | 0.678162 | -0.19996 | O14531 | O14531 | DPYSL4 | Dihydropy | Color2 [Da | 1 | 4 |
| + | 1.841898 | 3.024381 | O14548 | O14548 | COX7A2L | Cytochron | Color2 [Da | 1 | 4 |
| | 0 | 1.055398 | O14558 | O14558 | HSPB6 | Heat shocl | Color2 [Da | 1 | 4 |
| | 0.235763 | -0.09449 | O14559 | O14559 | ARHGAP33 | Rho GTPas | Color2 [Da | 1 | 4 |
| + | 1.676147 | 0.674828 | O14561 | O14561 | NDUFAB1 | Acyl carrie | Color2 [Da | 1 | 4 |
| + | 2.093267 | 1.084924 | O14562 | O14562 | UBFD1 | Ubiquitin c | Color2 [Da | 1 | 4 |
| + | 2.611008 | 1.297664 | O14576 | O14576 | DYNC1I1 | Cytoplasm | Color2 [Da | 1 | 4 |

| | | | | | | | | | |
|---|---|---|---|---|---|---|---|---|---|
| | 0 | -0.46844 | O14578 | O14578 | CIT | Citron Rho | Color2 [Da | 1 | 4 |
| | 1.138337 | 0.223733 | O14579 | O14579 | COPE | Coatomer | Color2 [Da | 1 | 4 |
| + | 1.326099 | 0.921119 | O14594 | O14594 | NCAN | Neurocan | Color2 [Da | 1 | 4 |
| | 0 | NaN | O14602 | O14602 | EIF1AY | Eukaryotic | Color2 [Da | 1 | 4 |
| + | 2.167709 | -0.39495 | O14617 | O14617 | AP3D1 | AP-3 comp | Color2 [Da | 1 | 4 |
| + | 2.943843 | 0.617589 | O14618 | O14618 | CCS | Copper ch | Color2 [Da | 1 | 4 |
| + | 3.272577 | 1.239056 | O14639 | O14639 | ABLIM1 | Actin-bind | Color2 [Da | 1 | 4 |
| + | 1.364515 | 1.164411 | O14640;P5 | O14640;P5 | DVL1;DVL1 | Segment p | Color2 [Da | 1 | 4 |
| + | 1.108842 | 0.388704 | O14641 | O14641 | DVL2 | Segment p | Color2 [Da | 1 | 4 |
| + | 4.02165 | -1.93408 | O14646 | O14646 | CHD1 | Chromodo | Color2 [Da | 1 | 4 |
| | 0.518467 | -0.41565 | O14647 | O14647 | CHD2 | Chromodo | Color2 [Da | 1 | 4 |
| | 0.219863 | 0.435235 | O14653 | O14653 | GOSR2 | Golgi SNAI | Color2 [Da | 1 | 4 |
| + | 1.01778 | -0.63702 | O14656 | O14656 | TOR1A | Torsin-1A | Color2 [Da | 1 | 4 |
| | 0 | 0.926862 | O14657 | O14657 | TOR1B | Torsin-1B | Color2 [Da | 1 | 4 |
| + | 1.324662 | 0.555175 | O14662 | O14662 | STX16 | Syntaxin-1 | Color2 [Da | 1 | 4 |
| | 0.001405 | -0.00121 | O14672 | O14672 | ADAM10 | Disintegrir | Color2 [Da | 1 | 4 |
| | 0.696983 | -0.06856 | O14683 | O14683 | TP53I11 | Tumor pro | Color2 [Da | 1 | 4 |
| + | 1.268271 | 0.378384 | O14686 | O14686 | KMT2D | Histone-ly | Color2 [Da | 1 | 4 |
| | 0 | NaN | O14727 | O14727 | APAF1 | Apoptotic | Color2 [Da | 1 | 4 |
| + | 2.115775 | -0.93209 | O14730 | O14730 | RIOK3 | Serine/thr | Color2 [Da | 1 | 4 |
| + | 3.028689 | -1.34265 | O14734 | O14734 | ACOT8 | Acyl-coenz | Color2 [Da | 1 | 4 |
| | 0.59658 | -0.47695 | O14735 | O14735 | CDIPT | CDP-diacyl | Color2 [Da | 1 | 4 |
| + | 4.500754 | 2.183016 | O14737 | O14737 | PDCD5 | Programm | Color2 [Da | 1 | 4 |
| + | 3.611675 | -1.42022 | O14744 | O14744 | PRMT5 | Protein arg | Color2 [Da | 1 | 4 |
| + | 2.563275 | 0.955358 | O14745 | O14745 | SLC9A3R1 | Na(+)/H(+) | Color2 [Da | 1 | 4 |
| + | 1.515716 | 0.749979 | O14763 | O14763 | TNFRSF10I | Tumor nec | Color2 [Da | 1 | 4 |
| + | 0.781285 | 1.039481 | O14770 | O14770 | MEIS2 | Homeobox | Color2 [Da | 1 | 4 |
| + | 2.68852 | 0.682231 | O14773 | O14773 | TPP1 | Tripeptidyl | Color2 [Da | 1 | 4 |
| + | 1.963394 | 0.312151 | O14775 | O14775 | GNB5 | Guanine n | Color2 [Da | 1 | 4 |
| + | 1.975276 | -0.21261 | O14776 | O14776 | TCERG1 | Transcripti | Color2 [Da | 1 | 4 |
| | 0.003499 | 0.002124 | O14777 | O14777 | NDC80 | Kinetochoi | Color2 [Da | 1 | 4 |
| | 0 | NaN | O14782 | O14782 | KIF3C | Kinesin-like | Color2 [Da | 1 | 4 |
| | 0 | -1.20658 | O14786 | O14786 | NRP1 | Neuropilin | Color2 [Da | 1 | 4 |
| + | 2.083619 | -1.31195 | O14787 | O14787 | TNPO2 | Transporti | Color2 [Da | 1 | 4 |
| + | 1.614739 | -0.79356 | O14795 | O14795 | UNC13B | Protein un | Color2 [Da | 1 | 4 |
| | 0 | 0.370815 | O14802 | O14802 | POLR3A | DNA-direc | Color2 [Da | 1 | 4 |
| + | 1.577229 | 0.614492 | O14807 | O14807 | MRAS | Ras-relate | Color2 [Da | 1 | 4 |
| | 0 | NaN | O14810 | O14810 | CPLX1 | Complexin | Color2 [Da | 1 | 4 |
| + | 3.14874 | 2.034689 | O14818 | O14818 | PSMA7 | Proteasom | Color2 [Da | 1 | 4 |
| + | 0.848189 | 1.199565 | O14828 | O14828 | SCAMP3 | Secretory | Color2 [Da | 1 | 4 |
| + | 2.887337 | -0.81776 | O14874 | O14874 | BCKDK | [3-methyl- | Color2 [Da | 1 | 4 |
| | 0.59931 | -0.35083 | O14880 | O14880 | MGST3 | Microsoma | Color2 [Da | 1 | 4 |
| | 0.357923 | 0.146891 | O14907 | O14907 | TAX1BP3 | Tax1-bindi | Color2 [Da | 1 | 4 |
| + | 2.413151 | 1.151274 | O14908 | O14908 | GIPC1 | PDZ doma | Color2 [Da | 1 | 4 |
| + | 2.691316 | 1.441243 | O14910 | O14910 | LIN7A | Protein lin | Color2 [Da | 1 | 4 |
| | 0.582714 | 0.601243 | O14917 | O14917 | PCDH17 | Protocadh | Color2 [Da | 1 | 4 |
| | 0.716838 | 0.168039 | O14924 | O14924 | RGS12 | Regulator | Color2 [Da | 1 | 4 |

| | | | | | | | | | | |
|---|---|---|---|---|---|---|---|---|---|---|
| + | 1.745849 | 2.734691 | O14925;Q | O14925;Q | TIMM23;T | Mitochonc | Color2 | [Da | 1 | 4 |
| + | 2.422902 | -2.59905 | O14929 | O14929 | HAT1 | Histone ac | Color2 | [Da | 1 | 4 |
| + | 2.066796 | 0.69046 | O14933 | O14933 | UBE2L6 | Ubiquitin/ | Color2 | [Da | 1 | 4 |
| + | 1.789749 | -0.44347 | O14936 | O14936 | CASK | Peripheral | Color2 | [Da | 1 | 4 |
| + | 2.869344 | 1.647064 | O14949 | O14949 | UQCRQ | Cytochron | Color2 | [Da | 1 | 4 |
| + | 3.015531 | 1.23726 | O14950;P1 | O14950;P1 | MYL12B;M | Myosin re | Color2 | [Da | 1 | 4 |
| + | 4.214801 | 4.191343 | O14957 | O14957 | UQCR11 | Cytochron | Color2 | [Da | 1 | 4 |
| + | 3.086711 | 1.050524 | O14964 | O14964 | HGS | Hepatocyt | Color2 | [Da | 1 | 4 |
| | 0.44255 | 0.435072 | O14967 | O14967 | CLGN | Calmegin | Color2 | [Da | 1 | 4 |
| + | 4.761274 | 1.435794 | O14972 | O14972 | VPS26C | Vacuolar p | Color2 | [Da | 1 | 4 |
| + | 1.32615 | 0.520464 | O14974 | O14974 | PPP1R12A | Protein ph | Color2 | [Da | 1 | 4 |
| | 0.558468 | 0.158145 | O14976 | O14976 | GAK | Cyclin-G-a | Color2 | [Da | 1 | 4 |
| + | 3.94309 | 1.707035 | O14979 | O14979 | HNRNPDL | Heterogen | Color2 | [Da | 1 | 4 |
| + | 3.435406 | -1.77408 | O14980 | O14980 | XPO1 | Exportin-1 | Color2 | [Da | 1 | 4 |
| + | 1.661509 | 0.347348 | O14981 | O14981 | BTAF1 | TATA-bind | Color2 | [Da | 1 | 4 |
| + | 3.074272 | 1.594961 | O14994 | O14994 | SYN3 | Synapsin-3 | Color2 | [Da | 1 | 4 |
| | 0.335847 | -0.2212 | O15013 | O15013 | ARHGEF10 | Rho guanir | Color2 | [Da | 1 | 4 |
| | 0.233066 | 0.331858 | O15014 | O15014 | ZNF609 | Zinc finger | Color2 | [Da | 1 | 4 |
| | 1.959885 | -0.16698 | O15020 | O15020 | SPTBN2 | Spectrin b | Color2 | [Da | 1 | 4 |
| | 0.092186 | 0.021948 | O15027 | O15027 | SEC16A | Protein tra | Color2 | [Da | 1 | 4 |
| + | 2.237723 | -0.45289 | O15031 | O15031 | PLXNB2 | Plexin-B2 | Color2 | [Da | 1 | 4 |
| + | 1.677093 | 2.524646 | O15034 | O15034 | RIMBP2 | RIMS-bind | Color2 | [Da | 1 | 4 |
| | 0.00635 | -0.00646 | O15040 | O15040 | TECPR2 | Tectonin b | Color2 | [Da | 1 | 4 |
| + | 2.169281 | 1.084861 | O15041 | O15041 | SEMA3E | Semaphor | Color2 | [Da | 1 | 4 |
| + | 1.869682 | -0.72497 | O15042 | O15042 | U2SURP | U2 snRNP- | Color2 | [Da | 1 | 4 |
| + | 1.471477 | 0.625435 | O15047 | O15047 | SETD1A | Histone-ly | Color2 | [Da | 1 | 4 |
| + | 2.789164 | 1.285994 | O15061 | O15061 | SYNM | Synemin | Color2 | [Da | 1 | 4 |
| + | 2.629491 | 0.834008 | O15066 | O15066 | KIF3B | Kinesin-lik | Color2 | [Da | 1 | 4 |
| + | 3.643745 | -1.14483 | O15067 | O15067 | PFAS | Phosphoril | Color2 | [Da | 1 | 4 |
| | 0 | NaN | O15068 | O15068 | MCF2L | Guanine n | Color2 | [Da | 1 | 4 |
| | 0.822082 | -0.20758 | O15069 | O15069 | NACAD | NAC-alpha | Color2 | [Da | 1 | 4 |
| | 0 | NaN | O15072 | O15072 | ADAMTS3 | A disinteg | Color2 | [Da | 1 | 4 |
| + | 2.453122 | 0.35597 | O15075 | O15075 | DCLK1 | Serine/thr | Color2 | [Da | 1 | 4 |
| + | 1.086791 | 0.906027 | O15083 | O15083 | ERC2 | ERC protei | Color2 | [Da | 1 | 4 |
| | 0.283687 | -0.25454 | O15084 | O15084 | ANKRD28 | Serine/thr | Color2 | [Da | 1 | 4 |
| + | 1.19506 | 0.30433 | O15085 | O15085 | ARHGEF11 | Rho guanir | Color2 | [Da | 1 | 4 |
| | 0 | NaN | O15091 | O15091 | PRORP | Mitochonc | Color2 | [Da | 1 | 4 |
| | 0 | NaN | O15111 | O15111 | CHUK | Inhibitor o | Color2 | [Da | 1 | 4 |
| + | 3.370585 | 1.891297 | O15116 | O15116 | LSM1 | U6 snRNA- | Color2 | [Da | 1 | 4 |
| + | 1.544099 | 0.638659 | O15118 | O15118 | NPC1 | NPC intrac | Color2 | [Da | 1 | 4 |
| | 0 | -1.31315 | O15121 | O15121 | DEGS1 | Sphingolip | Color2 | [Da | 1 | 4 |
| | 0.182284 | 0.102688 | O15126 | O15126 | SCAMP1 | Secretory | Color2 | [Da | 1 | 4 |
| + | 2.580963 | -1.02131 | O15131 | O15131 | KPNA5 | Importin s | Color2 | [Da | 1 | 4 |
| | 0 | -2.55016 | O15143 | O15143 | ARPC1B | Actin-relat | Color2 | [Da | 1 | 4 |
| | 0.99046 | -0.18938 | O15144 | O15144 | ARPC2 | Actin-relat | Color2 | [Da | 1 | 4 |
| | 0.193574 | -0.08404 | O15145 | O15145 | ARPC3 | Actin-relat | Color2 | [Da | 1 | 4 |
| | 0.412674 | -0.60757 | O15155 | O15155 | BET1 | BET1 hom | Color2 | [Da | 1 | 4 |

| | | | | | | | | | |
|---|---|---|---|---|---|---|---|---|---|
| | 0.002905 | 0.004357 | O15160 | O15160 | POLR1C | DNA-direc | Color2 [Da | 1 | 4 |
| + | 2.182463 | -1.66809 | O15164 | O15164 | TRIM24 | Transcripti | Color2 [Da | 1 | 4 |
| + | 1.827423 | 0.630456 | O15173 | O15173 | PGRMC2 | Membrane | Color2 [Da | 1 | 4 |
| + | 2.840587 | -1.08878 | O15182 | O15182 | CETN3 | Centrin-3 | Color2 [Da | 1 | 4 |
| | 0.713487 | 0.417073 | O15198 | O15198 | SMAD9 | Mothers a | Color2 [Da | 1 | 4 |
| + | 0.929997 | -0.54701 | O15211 | O15211 | RGL2 | Ral guanin | Color2 [Da | 1 | 4 |
| + | 1.533633 | 1.017519 | O15212 | O15212 | PFDN6 | Prefoldin s | Color2 [Da | 1 | 4 |
| + | 2.229944 | -2.03861 | O15213 | O15213 | WDR46 | WD repeat | Color2 [Da | 1 | 4 |
| + | 3.006025 | 1.408765 | O15217 | O15217 | GSTA4 | Glutathion | Color2 [Da | 1 | 4 |
| | 0.030408 | 0.030514 | O15226 | O15226 | NKRF | NF-kappa- | Color2 [Da | 1 | 4 |
| + | 2.326005 | -1.28581 | O15228 | O15228 | GNPAT | Dihydroxy | Color2 [Da | 1 | 4 |
| + | 1.004529 | -1.10227 | O15230 | O15230 | LAMA5 | Laminin su | Color2 [Da | 1 | 4 |
| | 0.859257 | 0.357842 | O15231 | O15231 | ZNF185 | Zinc finger | Color2 [Da | 1 | 4 |
| + | 2.291563 | 1.607912 | O15234 | O15234 | CASC3 | Protein CA | Color2 [Da | 1 | 4 |
| | 0 | 0.450851 | O15239 | O15239 | NDUFA1 | NADH deh | Color2 [Da | 1 | 4 |
| + | 2.426247 | 1.123956 | O15240 | O15240 | VGF | Neurosecr | Color2 [Da | 1 | 4 |
| + | 2.212277 | -0.99148 | O15254 | O15254 | ACOX3 | Peroxisom | Color2 [Da | 1 | 4 |
| + | 1.096421 | -0.46065 | O15258 | O15258 | RER1 | Protein RE | Color2 [Da | 1 | 4 |
| + | 1.625011 | -0.64678 | O15260 | O15260 | SURF4 | Surfeit loc | Color2 [Da | 1 | 4 |
| + | 3.334193 | -1.97699 | O15269 | O15269 | SPTLC1 | Serine palr | Color2 [Da | 1 | 4 |
| + | 1.241229 | -1.31873 | O15270 | O15270 | SPTLC2 | Serine palr | Color2 [Da | 1 | 4 |
| + | 3.222695 | -1.3752 | O15294 | O15294 | OGT | UDP-N-ace | Color2 [Da | 1 | 4 |
| | 0 | NaN | O15297 | O15297 | PPM1D | Protein ph | Color2 [Da | 1 | 4 |
| + | 1.821413 | -1.22546 | O15305 | O15305 | PMM2 | Phosphom | Color2 [Da | 1 | 4 |
| | 0 | -0.98763 | O15344 | O15344 | MID1 | E3 ubiquiti | Color2 [Da | 1 | 4 |
| | 0.000637 | -0.00058 | O15347 | O15347 | HMGB3 | High mobil | Color2 [Da | 1 | 4 |
| + | 2.423638 | -0.68733 | O15355 | O15355 | PPM1G | Protein ph | Color2 [Da | 1 | 4 |
| + | 1.212695 | 0.262528 | O15357 | O15357 | INPPL1 | Phosphati | Color2 [Da | 1 | 4 |
| + | 1.158725 | -0.58698 | O15371 | O15371 | EIF3D | Eukaryotic | Color2 [Da | 1 | 4 |
| + | 3.537498 | -0.6289 | O15372 | O15372 | EIF3H | Eukaryotic | Color2 [Da | 1 | 4 |
| | 0 | NaN | O15379 | O15379 | HDAC3 | Histone de | Color2 [Da | 1 | 4 |
| | 0 | NaN | O15381 | O15381 | NVL | Nuclear va | Color2 [Da | 1 | 4 |
| + | 1.952285 | 0.470134 | O15382 | O15382 | BCAT2 | Branched- | Color2 [Da | 1 | 4 |
| | 0.950395 | -0.30584 | O15394 | O15394 | NCAM2 | Neural cell | Color2 [Da | 1 | 4 |
| + | 1.882364 | -0.92512 | O15397 | O15397 | IPO8 | Importin-8 | Color2 [Da | 1 | 4 |
| + | 3.302796 | 1.911105 | O15400 | O15400 | STX7 | Syntaxin-7 | Color2 [Da | 1 | 4 |
| | 0.141551 | -0.15772 | O15405 | O15405 | TOX3 | TOX high r | Color2 [Da | 1 | 4 |
| | 0 | -1.7354 | O15427 | O15427 | SLC16A3 | Monocarb | Color2 [Da | 1 | 4 |
| | 0 | -1.15875 | O15439 | O15439 | ABCC4 | ATP-bindir | Color2 [Da | 1 | 4 |
| | 0.241943 | -0.36134 | O15446 | O15446 | POLR1G | DNA-direc | Color2 [Da | 1 | 4 |
| | 0 | -1.22369 | O15460 | O15460 | P4HA2 | Prolyl 4-hy | Color2 [Da | 1 | 4 |
| | 0.526049 | -0.08726 | O15484 | O15484 | CAPN5 | Calpain-5 | Color2 [Da | 1 | 4 |
| + | 0.808303 | 0.459676 | O15498 | O15498 | YKT6 | Synaptobr | Color2 [Da | 1 | 4 |
| | 0.678748 | 0.537394 | O15504 | O15504 | NUP42 | Nucleopor | Color2 [Da | 1 | 4 |
| + | 3.594694 | 1.061978 | O15511 | O15511 | ARPC5 | Actin-relat | Color2 [Da | 1 | 4 |
| + | 3.413362 | 2.271056 | O15514 | O15514 | POLR2D | DNA-direc | Color2 [Da | 1 | 4 |
| + | 1.479966 | -0.99986 | O15530 | O15530 | PDPK1 | 3-phospho | Color2 [Da | 1 | 4 |

| | | | | | | | | | |
|---|---|---|---|---|---|---|---|---|---|
| + | 4.715588 | 4.180407 | O15540 | O15540 | FABP7 | Fatty acid- | Color2 [Da | 1 | 4 |
| | 0 | NaN | O15550 | O15550 | KDM6A | Lysine-spe | Color2 [Da | 1 | 4 |
| + | 4.000207 | -1.17448 | O43143 | O43143 | DHX15 | ATP-deper | Color2 [Da | 1 | 4 |
| | 0.079599 | 0.069527 | O43148 | O43148 | RNMT | mRNA cap | Color2 [Da | 1 | 4 |
| | 0 | 0.722633 | O43149 | O43149 | ZZEF1 | Zinc finger | Color2 [Da | 1 | 4 |
| | 0.631931 | 0.90094 | O43150 | O43150 | ASAP2 | Arf-GAP w | Color2 [Da | 1 | 4 |
| | 0 | -1.04166 | O43156 | O43156 | TTI1 | TELO2-inte | Color2 [Da | 1 | 4 |
| + | 0.966913 | -0.62427 | O43157 | O43157 | PLXNB1 | Plexin-B1 | Color2 [Da | 1 | 4 |
| + | 1.11565 | -0.92965 | O43159 | O43159 | RRP8 | Ribosomal | Color2 [Da | 1 | 4 |
| + | 1.614327 | -0.61696 | O43164 | O43164 | PJA2 | E3 ubiquiti | Color2 [Da | 1 | 4 |
| + | 1.019442 | 1.242103 | O43169 | O43169 | CYB5B | Cytochrom | Color2 [Da | 1 | 4 |
| + | 1.726679 | -1.00904 | O43172 | O43172 | PRPF4 | U4/U6 sm | Color2 [Da | 1 | 4 |
| + | 1.769409 | -0.57561 | O43175 | O43175 | PHGDH | D-3-phosp | Color2 [Da | 1 | 4 |
| | 0.439208 | 0.513722 | O43181 | O43181 | NDUFS4 | NADH deh | Color2 [Da | 1 | 4 |
| | 0 | NaN | O43189 | O43189 | PHF1 | PHD finger | Color2 [Da | 1 | 4 |
| | 0.58671 | 2.110582 | O43236 | O43236 | SEPTIN4 | Septin-4 | Color2 [Da | 1 | 4 |
| + | 1.756305 | 0.652769 | O43237 | O43237 | DYNC1LI2 | Cytoplasm | Color2 [Da | 1 | 4 |
| + | 1.693215 | -0.52535 | O43242 | O43242 | PSMD3 | 26S protea | Color2 [Da | 1 | 4 |
| + | 2.054787 | 1.070704 | O43251 | O43251 | RBFOX2 | RNA bindir | Color2 [Da | 1 | 4 |
| + | 0.944456 | -0.74945 | O43252 | O43252 | PAPSS1 | Bifunction | Color2 [Da | 1 | 4 |
| + | 2.947086 | -1.05509 | O43264 | O43264 | ZW10 | Centromer | Color2 [Da | 1 | 4 |
| | 0 | NaN | O43272 | O43272 | PRODH | Proline del | Color2 [Da | 1 | 4 |
| | 0 | -1.52627 | O43281 | O43281 | EFS | Embryonal | Color2 [Da | 1 | 4 |
| | 0.149104 | 0.030508 | O43290 | O43290 | SART1 | U4/U6.U5 | Color2 [Da | 1 | 4 |
| + | 2.688688 | -1.41711 | O43292 | O43292 | GPAA1 | Glycosylph | Color2 [Da | 1 | 4 |
| + | 1.759059 | -0.49072 | O43295 | O43295 | SRGAP3 | SLIT-ROBO | Color2 [Da | 1 | 4 |
| | 0 | -0.97911 | O43299 | O43299 | AP5Z1 | AP-5 comp | Color2 [Da | 1 | 4 |
| + | 3.135158 | 0.833812 | O43301 | O43301 | HSPA12A | Heat shocl | Color2 [Da | 1 | 4 |
| + | 2.602457 | 1.782284 | O43303 | O43303 | CCP110 | Centriolar | Color2 [Da | 1 | 4 |
| | 0 | 2.03349 | O43306 | O43306 | ADCY6 | Adenylate | Color2 [Da | 1 | 4 |
| | 0 | -1.61654 | O43307;Q | O43307;Q | ARHGEF9;S | Rho guanir | Color2 [Da | 1 | 4 |
| | 0.434423 | 0.254154 | O43310 | O43310 | CTIF | CBP80/20- | Color2 [Da | 1 | 4 |
| | 0 | 2.128996 | O43312 | O43312 | MTSS1 | Protein M | Color2 [Da | 1 | 4 |
| + | 0.996737 | -0.52906 | O43314 | O43314 | PPIP5K2 | Inositol he | Color2 [Da | 1 | 4 |
| + | 1.712622 | 0.888094 | O43318 | O43318 | MAP3K7 | Mitogen-a | Color2 [Da | 1 | 4 |
| + | 2.206294 | 0.841049 | O43324 | O43324 | EEF1E1 | Eukaryotic | Color2 [Da | 1 | 4 |
| | 0.922988 | 0.204858 | O43325 | O43325 | LYRM1 | LYR motif- | Color2 [Da | 1 | 4 |
| | 0.087445 | 0.028389 | O43347 | O43347 | MSI1 | RNA-bindi | Color2 [Da | 1 | 4 |
| + | 2.298513 | 0.249468 | O43390 | O43390 | HNRNPR | Heterogen | Color2 [Da | 1 | 4 |
| | 0.234553 | 0.137562 | O43395 | O43395 | PRPF3 | U4/U6 sm | Color2 [Da | 1 | 4 |
| | 0.112558 | 0.033607 | O43396 | O43396 | TXNL1 | Thioredoxi | Color2 [Da | 1 | 4 |
| + | 3.281174 | 2.825058 | O43399 | O43399 | TPD52L2 | Tumor pro | Color2 [Da | 1 | 4 |
| + | 1.442164 | -0.50892 | O43402 | O43402 | EMC8 | ER membr | Color2 [Da | 1 | 4 |
| | 0 | 0.048004 | O43424 | O43424 | GRID2 | Glutamate | Color2 [Da | 1 | 4 |
| | 0.374487 | -0.1122 | O43426 | O43426 | SYNJ1 | Synaptojar | Color2 [Da | 1 | 4 |
| + | 3.083719 | -0.99236 | O43427 | O43427 | FIBP | Acidic fibr | Color2 [Da | 1 | 4 |
| + | 2.609978 | -0.65833 | O43432 | O43432 | EIF4G3 | Eukaryotic | Color2 [Da | 1 | 4 |

| | | | | | | | | | |
|---|---|---|---|---|---|---|---|---|---|
| + | 1.131272 | 0.44416 | O43447 | O43447 | PPIH | Peptidyl-p | Color2 [Da | 1 | 4 |
| + | 3.667765 | 2.234061 | O43464 | O43464 | HTRA2 | Serine pro | Color2 [Da | 1 | 4 |
| | 0.475938 | -0.31793 | O43488 | O43488 | AKR7A2 | Aflatoxin E | Color2 [Da | 1 | 4 |
| + | 2.153884 | 0.576122 | O43491 | O43491 | EPB41L2 | Band 4.1-li | Color2 [Da | 1 | 4 |
| | 0.570114 | 0.486789 | O43493 | O43493 | TGOLN2 | Trans-Golg | Color2 [Da | 1 | 4 |
| + | 1.312904 | 1.483366 | O43504 | O43504 | LAMTOR5 | Ragulator | Color2 [Da | 1 | 4 |
| + | 1.429708 | -0.95523 | O43505 | O43505 | B4GAT1 | Beta-1,4-g | Color2 [Da | 1 | 4 |
| | 0.691446 | -0.59958 | O43513 | O43513 | MED7 | Mediator | Color2 [Da | 1 | 4 |
| | 0 | 1.634023 | O43516 | O43516 | WIPF1 | WAS/WAS | Color2 [Da | 1 | 4 |
| + | 2.775058 | 1.647291 | O43524 | O43524 | FOXO3 | Forkhead | Color2 [Da | 1 | 4 |
| + | 1.25267 | 0.366489 | O43526 | O43526 | KCNQ2 | Potassium | Color2 [Da | 1 | 4 |
| | 0 | NaN | O43548 | O43548 | TGM5 | Protein-glu | Color2 [Da | 1 | 4 |
| + | 5.576315 | 2.273644 | O43567 | O43567 | RNF13 | E3 ubiquiti | Color2 [Da | 1 | 4 |
| | 0 | 0.666032 | O43570 | O43570 | CA12 | Carbonic a | Color2 [Da | 1 | 4 |
| + | 3.251387 | 2.060341 | O43581 | O43581 | SYT7 | Synaptota | Color2 [Da | 1 | 4 |
| + | 0.891701 | 0.694606 | O43583 | O43583 | DENR | Density-re | Color2 [Da | 1 | 4 |
| + | 2.530345 | -1.36062 | O43592 | O43592 | XPOT | Exportin-T | Color2 [Da | 1 | 4 |
| + | 2.424246 | 1.742085 | O43598 | O43598 | DNPH1 | 2'-deoxynu | Color2 [Da | 1 | 4 |
| + | 2.745368 | 0.959705 | O43602 | O43602 | DCX | Neuronal r | Color2 [Da | 1 | 4 |
| | 0.297325 | 0.112943 | O43615 | O43615 | TIMM44 | Mitochonc | Color2 [Da | 1 | 4 |
| | 0.24032 | 0.370684 | O43617 | O43617 | TRAPPC3 | Trafficking | Color2 [Da | 1 | 4 |
| + | 4.716886 | 1.788395 | O43633 | O43633 | CHMP2A | Charged m | Color2 [Da | 1 | 4 |
| + | 0.992474 | 0.62925 | O43639 | O43639 | NCK2 | Cytoplasm | Color2 [Da | 1 | 4 |
| + | 2.446289 | -2.34693 | O43657 | O43657 | TSPAN6 | Tetraspani | Color2 [Da | 1 | 4 |
| + | 3.09995 | -1.06344 | O43660 | O43660 | PLRG1 | Pleiotropic | Color2 [Da | 1 | 4 |
| + | 2.152133 | 1.919491 | O43670 | O43670 | ZNF207 | BUB3-inte | Color2 [Da | 1 | 4 |
| + | 2.326158 | 1.407237 | O43674 | O43674 | NDUFB5 | NADH deh | Color2 [Da | 1 | 4 |
| + | 1.398686 | 2.112631 | O43676 | O43676 | NDUFB3 | NADH deh | Color2 [Da | 1 | 4 |
| | 0 | NaN | O43677 | O43677 | NDUFC1 | NADH deh | Color2 [Da | 1 | 4 |
| + | 2.616113 | 2.358158 | O43678 | O43678 | NDUFA2 | NADH deh | Color2 [Da | 1 | 4 |
| | 0.315559 | 0.132349 | O43681 | O43681 | GET3 | ATPase GE | Color2 [Da | 1 | 4 |
| + | 2.147336 | -0.88115 | O43684 | O43684 | BUB3 | Mitotic ch | Color2 [Da | 1 | 4 |
| + | 2.667604 | 2.173052 | O43687;Q | O43687;Q | AKAP7 | A-kinase a | Color2 [Da | 1 | 4 |
| + | 3.508147 | 0.70159 | O43692 | O43692 | PI15 | Peptidase | Color2 [Da | 1 | 4 |
| + | 3.497285 | -1.10739 | O43707 | O43707 | ACTN4 | Alpha-acti | Color2 [Da | 1 | 4 |
| + | 1.525611 | 0.441963 | O43708 | O43708 | GSTZ1 | Maleylace | Color2 [Da | 1 | 4 |
| | 0 | -0.89844 | O43709 | O43709 | BUD23 | Probable 1 | Color2 [Da | 1 | 4 |
| + | 2.343503 | 2.324287 | O43716 | O43716 | GATC | Glutamyl-t | Color2 [Da | 1 | 4 |
| + | 3.46193 | -0.38682 | O43719 | O43719 | HTATSF1 | HIV Tat-sp | Color2 [Da | 1 | 4 |
| | 0.577668 | 0.529433 | O43741 | O43741 | PRKAB2 | 5'-AMP-ac | Color2 [Da | 1 | 4 |
| | 0.773362 | -0.14178 | O43747 | O43747 | AP1G1 | AP-1 comp | Color2 [Da | 1 | 4 |
| + | 1.974166 | 0.921964 | O43752 | O43752 | STX6 | Syntaxin-6 | Color2 [Da | 1 | 4 |
| + | 1.062849 | -0.51736 | O43759 | O43759 | SYNGR1 | Synaptogy | Color2 [Da | 1 | 4 |
| | 0.016496 | 0.032233 | O43761 | O43761 | SYNGR3 | Synaptogy | Color2 [Da | 1 | 4 |
| + | 1.587703 | 0.583392 | O43765 | O43765 | SGTA | Small gluta | Color2 [Da | 1 | 4 |
| + | 3.121662 | 4.390554 | O43768 | O43768 | ENSA | Alpha-end | Color2 [Da | 1 | 4 |
| + | 1.997283 | -0.6607 | O43772 | O43772 | SLC25A20 | Mitochonc | Color2 [Da | 1 | 4 |

| | | | | | | | | | |
|---|---|---|---|---|---|---|---|---|---|
| + | 2.490491 | -1.12868 | O43776 | O43776 | NARS1 | Asparagine | Color2 | [Da | 1 | 4 |
| | 0 | 3.594513 | O43790;Q | O43790;Q | KRT86;KRT | Keratin, ty | Color2 | [Da | 1 | 4 |
| + | 2.877138 | -0.69577 | O43795 | O43795 | MYO1B | Unconvent | Color2 | [Da | 1 | 4 |
| + | 1.631118 | 1.486743 | O43805 | O43805 | SSNA1 | Sjoegren s | Color2 | [Da | 1 | 4 |
| | 0.094278 | -0.09505 | O43808 | O43808 | SLC25A17 | Peroxisom | Color2 | [Da | 1 | 4 |
| + | 1.4227 | 0.747107 | O43809 | O43809 | NUDT21 | Cleavage a | Color2 | [Da | 1 | 4 |
| + | 3.311098 | -1.93701 | O43813 | O43813 | LANCL1 | Glutathion | Color2 | [Da | 1 | 4 |
| | 1.045008 | 0.17914 | O43815 | O43815 | STRN | Striatin | Color2 | [Da | 1 | 4 |
| + | 1.21894 | -0.82038 | O43818 | O43818 | RRP9 | U3 small n | Color2 | [Da | 1 | 4 |
| + | 1.458201 | 1.541989 | O43819 | O43819 | SCO2 | Protein SC | Color2 | [Da | 1 | 4 |
| | 0 | -1.31918 | O43822 | O43822 | CFAP410 | Cilia- and f | Color2 | [Da | 1 | 4 |
| | 0.748692 | -0.38177 | O43823 | O43823 | AKAP8 | A-kinase a | Color2 | [Da | 1 | 4 |
| | 0.862833 | 0.18165 | O43837 | O43837 | IDH3B | Isocitrate ( | Color2 | [Da | 1 | 4 |
| | 0 | -1.36385 | O43847 | O43847 | NRDC | Nardilysin | Color2 | [Da | 1 | 4 |
| + | 2.163174 | 0.506231 | O43852 | O43852 | CALU | Calumenin | Color2 | [Da | 1 | 4 |
| + | 2.992972 | 1.488416 | O43854 | O43854 | EDIL3 | EGF-like re | Color2 | [Da | 1 | 4 |
| + | 1.993716 | -0.7039 | O43865 | O43865 | AHCYL1 | S-adenosy | Color2 | [Da | 1 | 4 |
| | 0.134317 | -0.16403 | O43896 | O43896 | KIF1C | Kinesin-lik | Color2 | [Da | 1 | 4 |
| | 0 | NaN | O43909 | O43909 | EXTL3 | Exostosin- | Color2 | [Da | 1 | 4 |
| | 0 | 0.971154 | O43913 | O43913 | ORC5 | Origin recc | Color2 | [Da | 1 | 4 |
| + | 0.83052 | 0.69749 | O43920 | O43920 | NDUFS5 | NADH deh | Color2 | [Da | 1 | 4 |
| + | 2.234712 | 1.193501 | O43924 | O43924 | PDE6D | Retinal roc | Color2 | [Da | 1 | 4 |
| | 0 | -2.11451 | O43929 | O43929 | ORC4 | Origin recc | Color2 | [Da | 1 | 4 |
| + | 1.872752 | 0.437575 | O60216 | O60216 | RAD21 | Double-str | Color2 | [Da | 1 | 4 |
| | 0 | NaN | O60218 | O60218 | AKR1B10 | Aldo-keto | Color2 | [Da | 1 | 4 |
| + | 5.544202 | 2.246156 | O60220 | O60220 | TIMM8A | Mitochonc | Color2 | [Da | 1 | 4 |
| + | 1.118621 | -0.97107 | O60229 | O60229 | KALRN | Kalirin | Color2 | [Da | 1 | 4 |
| + | 2.624845 | -1.49598 | O60231 | O60231 | DHX16 | Pre-mRNA | Color2 | [Da | 1 | 4 |
| + | 2.932544 | 2.67999 | O60232 | O60232 | ZNRD2 | Protein ZN | Color2 | [Da | 1 | 4 |
| + | 1.887226 | 2.633347 | O60238 | O60238 | BNIP3L | BCL2/ader | Color2 | [Da | 1 | 4 |
| + | 2.301296 | -0.82019 | O60242 | O60242 | ADGRB3 | Adhesion ( | Color2 | [Da | 1 | 4 |
| + | 0.698429 | 1.078592 | O60243 | O60243 | HS6ST1 | Heparan-s | Color2 | [Da | 1 | 4 |
| + | 2.686998 | -1.72514 | O60244 | O60244 | MED14 | Mediator ( | Color2 | [Da | 1 | 4 |
| | 0.092082 | 0.021127 | O60245 | O60245 | PCDH7 | Protocadh | Color2 | [Da | 1 | 4 |
| | 1.029722 | 0.126672 | O60256 | O60256 | PRPSAP2 | Phosphoril | Color2 | [Da | 1 | 4 |
| + | 0.700652 | 1.010404 | O60262 | O60262 | GNG7 | Guanine n | Color2 | [Da | 1 | 4 |
| + | 2.312408 | -0.71982 | O60264 | O60264 | SMARCA5 | SWI/SNF-r | Color2 | [Da | 1 | 4 |
| + | 1.23229 | 0.389917 | O60268 | O60268 | KIAA0513 | Uncharact | Color2 | [Da | 1 | 4 |
| + | 1.366454 | 0.234542 | O60271 | O60271 | SPAG9 | C-Jun-amir | Color2 | [Da | 1 | 4 |
| + | 3.592326 | -0.67402 | O60282 | O60282 | KIF5C | Kinesin he | Color2 | [Da | 1 | 4 |
| | 0 | 1.905364 | O60287 | O60287 | URB1 | Nucleolar | Color2 | [Da | 1 | 4 |
| | 0.813136 | -0.28637 | O60291 | O60291 | MGRN1 | E3 ubiquiti | Color2 | [Da | 1 | 4 |
| + | 1.666819 | 0.310819 | O60292 | O60292 | SIPA1L3 | Signal-indu | Color2 | [Da | 1 | 4 |
| + | 1.140961 | -0.25662 | O60306 | O60306 | AQR | RNA helica | Color2 | [Da | 1 | 4 |
| + | 1.294534 | 0.300591 | O60313 | O60313 | OPA1 | Dynamin-l | Color2 | [Da | 1 | 4 |
| + | 3.753269 | 0.822709 | O60315 | O60315 | ZEB2 | Zinc finger | Color2 | [Da | 1 | 4 |
| | 0 | -0.34683 | O60318 | O60318 | MCM3AP | Germinal-( | Color2 | [Da | 1 | 4 |

| | | | | | | | | | |
|---|---|---|---|---|---|---|---|---|---|
| | 0.704861 | 0.397171 | O60331 | O60331 | PIP5K1C | Phosphatic | Color2 [Da | 1 | 4 |
| + | 1.135729 | 0.265905 | O60333 | O60333 | KIF1B | Kinesin-lik | Color2 [Da | 1 | 4 |
| + | 2.810537 | -1.18395 | O60341 | O60341 | KDM1A | Lysine-spe | Color2 [Da | 1 | 4 |
| + | 0.89131 | -0.9637 | O60343 | O60343 | TBC1D4 | TBC1 dom | Color2 [Da | 1 | 4 |
| + | 1.43633 | 1.78982 | O60346 | O60346 | PHLPP1 | PH domair | Color2 [Da | 1 | 4 |
| + | 1.283613 | 0.69409 | O60361 | O60361 | NME2P1 | Putative n | Color2 [Da | 1 | 4 |
| | 0 | -2.52834 | O60427 | O60427 | FADS1 | Acyl-CoA ( | Color2 [Da | 1 | 4 |
| | 0 | -0.67145 | O60437 | O60437 | PPL | Periplakin | Color2 [Da | 1 | 4 |
| + | 1.773275 | 0.576852 | O60443 | O60443 | GSDME | Gasdermir | Color2 [Da | 1 | 4 |
| + | 1.090721 | -0.43336 | O60462 | O60462 | NRP2 | Neuropilin | Color2 [Da | 1 | 4 |
| + | 3.208167 | -1.71288 | O60476 | O60476 | MAN1A2 | Mannosyl- | Color2 [Da | 1 | 4 |
| | 0.415051 | -0.17467 | O60486 | O60486 | PLXNC1 | Plexin-C1 | Color2 [Da | 1 | 4 |
| + | 2.310392 | -0.89651 | O60488 | O60488 | ACSL4 | Long-chair | Color2 [Da | 1 | 4 |
| + | 2.361345 | 0.799578 | O60493 | O60493 | SNX3 | Sorting ne: | Color2 [Da | 1 | 4 |
| | 0.303068 | -0.09815 | O60502 | O60502 | OGA | Protein O- | Color2 [Da | 1 | 4 |
| + | 1.455033 | 0.461964 | O60506 | O60506 | SYNCRIP | Heterogen | Color2 [Da | 1 | 4 |
| | 0.327186 | 0.530071 | O60507 | O60507 | TPST1 | Protein-tyı | Color2 [Da | 1 | 4 |
| + | 1.656026 | -1.16489 | O60508 | O60508 | CDC40 | Pre-mRNA | Color2 [Da | 1 | 4 |
| | 0 | -0.27286 | O60518 | O60518 | RANBP6 | Ran-bindir | Color2 [Da | 1 | 4 |
| | 0.019431 | -0.00579 | O60524 | O60524 | NEMF | Ribosome | Color2 [Da | 1 | 4 |
| + | 2.874613 | -1.1649 | O60547 | O60547 | GMDS | GDP-manr | Color2 [Da | 1 | 4 |
| + | 3.188336 | -0.78976 | O60551 | O60551 | NMT2 | Glycylpept | Color2 [Da | 1 | 4 |
| + | 3.036399 | -1.40761 | O60568 | O60568 | PLOD3 | Multifunct | Color2 [Da | 1 | 4 |
| + | 1.757459 | 0.833304 | O60573 | O60573 | EIF4E2 | Eukaryotic | Color2 [Da | 1 | 4 |
| + | 1.865219 | -0.35678 | O60610 | O60610 | DIAPH1 | Protein dia | Color2 [Da | 1 | 4 |
| + | 2.181991 | 1.404327 | O60613 | O60613 | SELENOF | Selenopro | Color2 [Da | 1 | 4 |
| + | 3.46399 | 0.900816 | O60641 | O60641 | SNAP91 | Clathrin cc | Color2 [Da | 1 | 4 |
| | 0.611088 | -0.14314 | O60645 | O60645 | EXOC3 | Exocyst co | Color2 [Da | 1 | 4 |
| + | 3.077357 | 1.845155 | O60664 | O60664 | PLIN3 | Perilipin-3 | Color2 [Da | 1 | 4 |
| | 0 | NaN | O60678 | O60678 | PRMT3 | Protein arg | Color2 [Da | 1 | 4 |
| | 0 | NaN | O60683 | O60683 | PEX10 | Peroxisom | Color2 [Da | 1 | 4 |
| + | 2.095157 | -0.34486 | O60684 | O60684 | KPNA6 | Importin s | Color2 [Da | 1 | 4 |
| + | 2.840126 | -1.45446 | O60701 | O60701 | UGDH | UDP-gluco | Color2 [Da | 1 | 4 |
| + | 1.860646 | 0.389896 | O60716 | O60716 | CTNND1 | Catenin de | Color2 [Da | 1 | 4 |
| + | 3.727738 | 2.138916 | O60739 | O60739 | EIF1B | Eukaryotic | Color2 [Da | 1 | 4 |
| + | 2.840838 | 0.531713 | O60749 | O60749 | SNX2 | Sorting ne: | Color2 [Da | 1 | 4 |
| | 0.957589 | 0.103959 | O60762 | O60762 | DPM1 | Dolichol-pl | Color2 [Da | 1 | 4 |
| + | 3.022177 | -1.09187 | O60763 | O60763 | USO1 | General ve | Color2 [Da | 1 | 4 |
| + | 0.851363 | 0.52397 | O60783 | O60783 | MRPS14 | 28S riboso | Color2 [Da | 1 | 4 |
| + | 2.660867 | 0.478048 | O60784 | O60784 | TOM1 | Target of M | Color2 [Da | 1 | 4 |
| | 0 | 0.232007 | O60825 | O60825 | PFKFB2 | 6-phospho | Color2 [Da | 1 | 4 |
| + | 1.604669 | -0.48353 | O60826 | O60826 | CCDC22 | Coiled-coil | Color2 [Da | 1 | 4 |
| + | 2.649878 | 2.542903 | O60828 | O60828 | PQBP1 | Polyglutan | Color2 [Da | 1 | 4 |
| + | 2.115856 | 1.850901 | O60830 | O60830 | TIMM17B | Mitochonc | Color2 [Da | 1 | 4 |
| | 0.146054 | 0.251196 | O60831 | O60831 | PRAF2 | PRA1 fami | Color2 [Da | 1 | 4 |
| + | 2.543221 | -1.60108 | O60832 | O60832 | DKC1 | H/ACA ribc | Color2 [Da | 1 | 4 |
| + | 2.58941 | -0.79173 | O60841 | O60841 | EIF5B | Eukaryotic | Color2 [Da | 1 | 4 |

| | | | | | | | | | |
|---|---|---|---|---|---|---|---|---|---|
| | 0 | 0.263865 | O60858 | O60858 | TRIM13 | E3 ubiquiti | Color2 [Da | 1 | 4 |
| | 0.64913 | 0.312989 | O60861 | O60861 | GAS7 | Growth ar | Color2 [Da | 1 | 4 |
| + | 2.332708 | 1.131888 | O60869 | O60869 | EDF1 | Endothelia | Color2 [Da | 1 | 4 |
| | 0.780772 | 0.257367 | O60870 | O60870 | KIN | DNA/RNA- | Color2 [Da | 1 | 4 |
| | 0 | -0.03276 | O60879 | O60879 | DIAPH2 | Protein dia | Color2 [Da | 1 | 4 |
| + | 3.839609 | 0.853011 | O60884 | O60884 | DNAJA2 | DnaJ homo | Color2 [Da | 1 | 4 |
| + | 2.004347 | 0.771915 | O60885 | O60885 | BRD4 | Bromodon | Color2 [Da | 1 | 4 |
| + | 3.195214 | 2.664865 | O60888 | O60888 | CUTA | Protein Cu | Color2 [Da | 1 | 4 |
| | 0 | NaN | O60890 | O60890 | OPHN1 | Oligophrer | Color2 [Da | 1 | 4 |
| | 0 | 0.030876 | O60911 | O60911 | CTSV | Cathepsin | Color2 [Da | 1 | 4 |
| | 0 | NaN | O60921 | O60921 | HUS1 | Checkpoin | Color2 [Da | 1 | 4 |
| + | 3.269736 | 1.663864 | O60925 | O60925 | PFDN1 | Prefoldin s | Color2 [Da | 1 | 4 |
| + | 2.616601 | 2.149832 | O60927 | O60927 | PPP1R11 | E3 ubiquiti | Color2 [Da | 1 | 4 |
| + | 1.714395 | -0.76706 | O60934 | O60934 | NBN | Nibrin | Color2 [Da | 1 | 4 |
| + | 2.807504 | 2.704112 | O60936 | O60936 | NOL3 | Nucleolar | | Color2 [Da | 1 | 4 |
| | 0 | 1.839419 | O60941 | O60941 | DTNB | Dystrobrev | Color2 [Da | 1 | 4 |
| | 0 | -0.52981 | O60942 | O60942 | RNGTT | mRNA-cap | Color2 [Da | 1 | 4 |
| + | 1.973215 | 0.237741 | O75027 | O75027 | ABCB7 | Iron-sulfur | Color2 [Da | 1 | 4 |
| | 0.542662 | -0.13947 | O75037 | O75037 | KIF21B | Kinesin-lik | Color2 [Da | 1 | 4 |
| | 0.719129 | 0.172243 | O75044 | O75044 | SRGAP2 | SLIT-ROBO | Color2 [Da | 1 | 4 |
| | 0.388028 | -0.07235 | O75051 | O75051 | PLXNA2 | Plexin-A2 | Color2 [Da | 1 | 4 |
| + | 3.182913 | -1.26977 | O75054 | O75054 | IGSF3 | Immunogl | Color2 [Da | 1 | 4 |
| + | 1.8119 | 1.277184 | O75056 | O75056 | SDC3 | Syndecan- | Color2 [Da | 1 | 4 |
| + | 3.319319 | 1.931539 | O75061 | O75061 | DNAJC6 | Putative ty | Color2 [Da | 1 | 4 |
| | 0 | -1.14946 | O75063 | O75063 | FAM20B | Glycosami | Color2 [Da | 1 | 4 |
| | 0 | 0.249442 | O75064 | O75064 | DENND4B | DENN dom | Color2 [Da | 1 | 4 |
| + | 2.369525 | -0.78741 | O75083 | O75083 | WDR1 | WD repeat | Color2 [Da | 1 | 4 |
| + | 3.070522 | 1.147596 | O75093 | O75093 | SLIT1 | Slit homol | Color2 [Da | 1 | 4 |
| | 0 | NaN | O75094 | O75094 | SLIT3 | Slit homol | Color2 [Da | 1 | 4 |
| | 0 | NaN | O75096 | O75096 | LRP4 | Low-densi | Color2 [Da | 1 | 4 |
| + | 2.656455 | -1.2049 | O75110 | O75110 | ATP9A | Probable p | Color2 [Da | 1 | 4 |
| | 0.339888 | 0.134863 | O75113 | O75113 | N4BP1 | NEDD4-bir | Color2 [Da | 1 | 4 |
| + | 1.161732 | -0.29601 | O75116 | O75116 | ROCK2 | Rho-associ | Color2 [Da | 1 | 4 |
| + | 1.145829 | 0.286328 | O75122 | O75122 | CLASP2 | CLIP-assoc | Color2 [Da | 1 | 4 |
| + | 1.266401 | -0.37276 | O75127 | O75127 | PTCD1 | Pentatrico | Color2 [Da | 1 | 4 |
| + | 1.355726 | -0.51783 | O75131 | O75131 | CPNE3 | Copine-3 | Color2 [Da | 1 | 4 |
| + | 1.810963 | 0.692452 | O75143 | O75143 | ATG13 | Autophagy | Color2 [Da | 1 | 4 |
| + | 3.28582 | 0.515505 | O75145 | O75145 | PPFIA3 | Liprin-alph | Color2 [Da | 1 | 4 |
| | 0.48391 | 0.251828 | O75146 | O75146 | HIP1R | Huntingtin | Color2 [Da | 1 | 4 |
| | 0.464788 | 0.068369 | O75150 | O75150 | RNF40 | E3 ubiquiti | Color2 [Da | 1 | 4 |
| | 0.64046 | -0.23659 | O75151 | O75151 | PHF2 | Lysine-spe | Color2 [Da | 1 | 4 |
| + | 1.245964 | -0.41715 | O75152 | O75152 | ZC3H11A | Zinc finger | Color2 [Da | 1 | 4 |
| | 0.507905 | 0.516527 | O75153 | O75153 | CLUH | Clustered | Color2 [Da | 1 | 4 |
| + | 1.506092 | -0.26915 | O75155 | O75155 | CAND2 | Cullin-asso | Color2 [Da | 1 | 4 |
| + | 2.374229 | 0.567082 | O75157 | O75157 | TSC22D2 | TSC22 don | Color2 [Da | 1 | 4 |
| + | 2.611297 | -1.04155 | O75165 | O75165 | DNAJC13 | DnaJ homo | Color2 [Da | 1 | 4 |
| | 0 | 0.008808 | O75167 | O75167 | PHACTR2 | Phosphata | Color2 [Da | 1 | 4 |

| | | | | | | | | | |
|---|---|---|---|---|---|---|---|---|---|
| + | 1.224817 | -0.49224 | O75170 | O75170 | PPP6R2 | Serine/thr | Color2 [Da | 1 | 4 |
| + | 1.28282 | 0.823305 | O75175 | O75175 | CNOT3 | CCR4-NOT | Color2 [Da | 1 | 4 |
| + | 1.271968 | 1.24339 | O75177 | O75177 | SS18L1 | Calcium-re | Color2 [Da | 1 | 4 |
| | 0.018542 | -0.0091 | O75179 | O75179 | ANKRD17 | Ankyrin re | Color2 [Da | 1 | 4 |
| + | 1.713979 | -0.66964 | O75182 | O75182 | SIN3B | Paired am | Color2 [Da | 1 | 4 |
| + | 3.022078 | 1.769102 | O75190 | O75190 | DNAJB6 | DnaJ hom | Color2 [Da | 1 | 4 |
| + | 4.072655 | 1.68367 | O75208 | O75208 | COQ9 | Ubiquinon | Color2 [Da | 1 | 4 |
| + | 2.863854 | 1.764056 | O75223 | O75223 | GGCT | Gamma-gl | Color2 [Da | 1 | 4 |
| + | 2.575157 | -0.89571 | O75251 | O75251 | NDUFS7 | NADH deh | Color2 [Da | 1 | 4 |
| + | 4.009021 | -1.40667 | O75306 | O75306 | NDUFS2 | NADH deh | Color2 [Da | 1 | 4 |
| + | 1.247956 | 0.603011 | O75312 | O75312 | ZPR1 | Zinc finger | Color2 [Da | 1 | 4 |
| + | 1.694562 | 0.544638 | O75323 | O75323 | NIPSNAP2 | Protein Ni | Color2 [Da | 1 | 4 |
| | 0 | -0.07076 | O75324 | O75324 | SNN | Stannin | Color2 [Da | 1 | 4 |
| + | 3.03725 | 0.847789 | O75334 | O75334 | PPFIA2 | Liprin-alph | Color2 [Da | 1 | 4 |
| + | 1.396715 | -0.61986 | O75340 | O75340 | PDCD6 | Programm | Color2 [Da | 1 | 4 |
| + | 3.025407 | 3.15403 | O75347 | O75347 | TBCA | Tubulin-sp | Color2 [Da | 1 | 4 |
| + | 3.849361 | 1.196947 | O75348 | O75348 | ATP6V1G1 | V-type pro | Color2 [Da | 1 | 4 |
| | 0.204867 | -0.13219 | O75351 | O75351 | VPS4B | Vacuolar p | Color2 [Da | 1 | 4 |
| + | 4.07699 | -2.17122 | O75352 | O75352 | MPDU1 | Mannose-I | Color2 [Da | 1 | 4 |
| | 0 | NaN | O75354 | O75354 | ENTPD6 | Ectonucle | Color2 [Da | 1 | 4 |
| | 0 | NaN | O75356 | O75356 | ENTPD5 | Nucleoside | Color2 [Da | 1 | 4 |
| + | 0.955548 | -0.3464 | O75367 | O75367 | MACROH2 | Core histo | Color2 [Da | 1 | 4 |
| + | 4.347779 | 4.215085 | O75368 | O75368 | SH3BGRL | SH3 domai | Color2 [Da | 1 | 4 |
| + | 3.8473 | -1.21172 | O75369 | O75369 | FLNB | Filamin-B | Color2 [Da | 1 | 4 |
| | 0.254203 | -0.12456 | O75376 | O75376 | NCOR1 | Nuclear re | Color2 [Da | 1 | 4 |
| + | 2.067274 | 1.051215 | O75380 | O75380 | NDUFS6 | NADH deh | Color2 [Da | 1 | 4 |
| + | 4.35151 | 2.91243 | O75381 | O75381 | PEX14 | Peroxisom | Color2 [Da | 1 | 4 |
| | 0.970485 | 0.292221 | O75382 | O75382 | TRIM3 | Tripartite | Color2 [Da | 1 | 4 |
| | 0.166401 | 0.026156 | O75390 | O75390 | CS | Citrate syn | Color2 [Da | 1 | 4 |
| + | 2.273949 | 0.818628 | O75391 | O75391 | SPAG7 | Sperm-ass | Color2 [Da | 1 | 4 |
| + | 3.477026 | -1.3896 | O75396 | O75396 | SEC22B | Vesicle-tra | Color2 [Da | 1 | 4 |
| + | 1.876739 | -0.22745 | O75400 | O75400 | PRPF40A | Pre-mRNA | Color2 [Da | 1 | 4 |
| | 0.451453 | 0.337374 | O75420 | O75420 | GIGYF1 | GRB10-int | Color2 [Da | 1 | 4 |
| + | 1.816371 | -0.76666 | O75425 | O75425 | MOSPD3 | Motile spe | Color2 [Da | 1 | 4 |
| | 0.594427 | -0.18234 | O75431 | O75431 | MTX2 | Metaxin-2 | Color2 [Da | 1 | 4 |
| + | 1.395654 | -0.39657 | O75436 | O75436 | VPS26A | Vacuolar p | Color2 [Da | 1 | 4 |
| + | 2.561531 | 2.002609 | O75438 | O75438 | NDUFB1 | NADH deh | Color2 [Da | 1 | 4 |
| + | 2.410656 | -1.03333 | O75439 | O75439 | PMPCB | Mitochon | Color2 [Da | 1 | 4 |
| | 0 | -0.6334 | O75443 | O75443 | TECTA | Alpha-tect | Color2 [Da | 1 | 4 |
| | 0.783016 | 0.428838 | O75448 | O75448 | MED24 | Mediator | Color2 [Da | 1 | 4 |
| | 0 | NaN | O75449 | O75449 | KATNA1 | Katanin p6 | Color2 [Da | 1 | 4 |
| + | 3.233352 | 1.001216 | O75475 | O75475 | PSIP1 | PC4 and SI | Color2 [Da | 1 | 4 |
| + | 1.715187 | -0.66162 | O75477 | O75477 | ERLIN1 | Erlin-1 | Color2 [Da | 1 | 4 |
| | 0 | 0.339034 | O75486 | O75486 | SUPT3H | Transcripti | Color2 [Da | 1 | 4 |
| + | 5.22248 | -2.85659 | O75487 | O75487 | GPC4 | Glypican-4 | Color2 [Da | 1 | 4 |
| + | 3.028511 | 1.624136 | O75489 | O75489 | NDUFS3 | NADH deh | Color2 [Da | 1 | 4 |
| | 0.463992 | -0.18218 | O75493 | O75493 | CA11 | Carbonic a | Color2 [Da | 1 | 4 |

| | | | | | | | | | |
|---|---|---|---|---|---|---|---|---|---|
| + | 1.420612 | 0.663459 | O75494 | O75494 | SRSF10 | Serine/arg | Color2 [Da | 1 | 4 |
| + | 1.425727 | 1.131605 | O75496 | O75496 | GMNN | Geminin | Color2 [Da | 1 | 4 |
| + | 1.072229 | 0.76198 | O75503 | O75503 | CLN5 | Ceroid-lipo | Color2 [Da | 1 | 4 |
| + | 1.549364 | 0.722001 | O75506 | O75506 | HSBP1 | Heat shock | Color2 [Da | 1 | 4 |
| + | 1.872576 | -0.65621 | O75521 | O75521 | ECI2 | Enoyl-CoA | Color2 [Da | 1 | 4 |
| | 0.04868 | -0.02516 | O75525 | O75525 | KHDRBS3 | KH domain | Color2 [Da | 1 | 4 |
| + | 1.01736 | -1.79364 | O75530 | O75530 | EED | Polycomb | Color2 [Da | 1 | 4 |
| + | 2.719089 | 0.905018 | O75531 | O75531 | BANF1 | Barrier-to- | Color2 [Da | 1 | 4 |
| + | 3.635204 | -1.063 | O75533 | O75533 | SF3B1 | Splicing fac | Color2 [Da | 1 | 4 |
| + | 2.1927 | -0.38891 | O75534 | O75534 | CSDE1 | Cold shock | Color2 [Da | 1 | 4 |
| | 0.31488 | 0.211286 | O75553 | O75553 | DAB1 | Disabled h | Color2 [Da | 1 | 4 |
| + | 1.774457 | -0.58488 | O75554 | O75554 | WBP4 | WW doma | Color2 [Da | 1 | 4 |
| | 0.024888 | 0.010393 | O75569 | O75569 | PRKRA | Interferon | Color2 [Da | 1 | 4 |
| + | 1.847288 | -1.57624 | O75582 | O75582 | RPS6KA5 | Ribosomal | Color2 [Da | 1 | 4 |
| + | 1.489771 | -1.09817 | O75592 | O75592 | MYCBP2 | E3 ubiquiti | Color2 [Da | 1 | 4 |
| + | 1.435581 | -0.62183 | O75600 | O75600 | GCAT | 2-amino-3 | Color2 [Da | 1 | 4 |
| + | 2.449669 | -0.44419 | O75607 | O75607 | NPM3 | Nucleoplas | Color2 [Da | 1 | 4 |
| | 0.665262 | 0.510552 | O75608 | O75608 | LYPLA1 | Acyl-prote | Color2 [Da | 1 | 4 |
| + | 3.015084 | 3.632198 | O75629 | O75629 | CREG1 | Protein CR | Color2 [Da | 1 | 4 |
| | 0 | -1.28652 | O75635 | O75635 | SERPINB7 | Serpin B7 | Color2 [Da | 1 | 4 |
| + | 4.161576 | -1.03792 | O75643 | O75643 | SNRNP200 | U5 small n | Color2 [Da | 1 | 4 |
| + | 4.121465 | -0.76782 | O75663 | O75663 | TIPRL | TIP41-like | Color2 [Da | 1 | 4 |
| + | 1.797566 | 0.888373 | O75674 | O75674 | TOM1L1 | TOM1-like | Color2 [Da | 1 | 4 |
| + | 1.251243 | 0.615252 | O75683 | O75683 | SURF6 | Surfeit locu | Color2 [Da | 1 | 4 |
| | 0.744229 | -0.27539 | O75688 | O75688 | PPM1B | Protein ph | Color2 [Da | 1 | 4 |
| + | 1.439608 | 0.538033 | O75689 | O75689 | ADAP1 | Arf-GAP w | Color2 [Da | 1 | 4 |
| | 0 | 3.337077 | O75691 | O75691 | UTP20 | Small subu | Color2 [Da | 1 | 4 |
| + | 2.629134 | -1.49609 | O75694 | O75694 | NUP155 | Nuclear po | Color2 [Da | 1 | 4 |
| + | 1.20507 | -0.59179 | O75695 | O75695 | RP2 | Protein XR | Color2 [Da | 1 | 4 |
| | 0 | NaN | O75717 | O75717 | WDHD1 | WD repeat | Color2 [Da | 1 | 4 |
| + | 1.278984 | -0.67673 | O75718 | O75718 | CRTAP | Cartilage-a | Color2 [Da | 1 | 4 |
| + | 1.582849 | 0.229788 | O75746 | O75746 | SLC25A12 | Calcium-bi | Color2 [Da | 1 | 4 |
| + | 4.341778 | 2.195434 | O75781 | O75781 | PALM | Paralemm | Color2 [Da | 1 | 4 |
| + | 3.685747 | 0.76148 | O75787 | O75787 | ATP6AP2 | Renin rece | Color2 [Da | 1 | 4 |
| | 0 | NaN | O75794 | O75794 | CDC123 | Cell divisio | Color2 [Da | 1 | 4 |
| + | 1.21505 | 1.046868 | O75817 | O75817 | POP7 | Ribonuclea | Color2 [Da | 1 | 4 |
| | 0 | NaN | O75818 | O75818 | RPP40 | Ribonuclea | Color2 [Da | 1 | 4 |
| | 0 | -1.51778 | O75820 | O75820 | ZNF189 | Zinc finger | Color2 [Da | 1 | 4 |
| + | 2.848414 | 0.97145 | O75821 | O75821 | EIF3G | Eukaryotic | Color2 [Da | 1 | 4 |
| + | 4.855935 | 1.527284 | O75822 | O75822 | EIF3J | Eukaryotic | Color2 [Da | 1 | 4 |
| + | 1.732894 | 1.046436 | O75832 | O75832 | PSMD10 | 26S protea | Color2 [Da | 1 | 4 |
| | 0.579236 | -1.79747 | O75844 | O75844 | ZMPSTE24 | CAAX pren | Color2 [Da | 1 | 4 |
| | 0 | NaN | O75865 | O75865 | TRAPPC6A | Trafficking | Color2 [Da | 1 | 4 |
| | 0.58546 | 0.209815 | O75874 | O75874 | IDH1 | Isocitrate d | Color2 [Da | 1 | 4 |
| + | 1.437566 | 1.152647 | O75880 | O75880 | SCO1 | Protein SC | Color2 [Da | 1 | 4 |
| | 0.76397 | -0.30939 | O75882 | O75882 | ATRN | Attractin | Color2 [Da | 1 | 4 |
| + | 0.730143 | 0.577016 | O75884 | O75884 | RBBP9 | Serine hyd | Color2 [Da | 1 | 4 |

| | | | | | | | | | |
|---|---|---|---|---|---|---|---|---|---|
| + | 2.031612 | 1.458254 | O75886 | O75886 | STAM2 | Signal tran | Color2 [Da | 1 | 4 |
| | 0 | -0.39247 | O75891 | O75891 | ALDH1L1 | Cytosolic 1 | Color2 [Da | 1 | 4 |
| + | 2.983273 | 1.492505 | O75896 | O75896 | TUSC2 | Tumor sup | Color2 [Da | 1 | 4 |
| | 0 | -0.12816 | O75899 | O75899 | GABBR2 | Gamma-ar | Color2 [Da | 1 | 4 |
| | 0 | -0.92033 | O75907 | O75907 | DGAT1 | Diacylglyce | Color2 [Da | 1 | 4 |
| | 0 | -0.51771 | O75909 | O75909 | CCNK | Cyclin-K | Color2 [Da | 1 | 4 |
| + | 2.28169 | -0.49095 | O75912 | O75912 | DGKI | Diacylglyce | Color2 [Da | 1 | 4 |
| + | 2.309587 | 1.820494 | O75914 | O75914 | PAK3 | Serine/thr | Color2 [Da | 1 | 4 |
| + | 3.096761 | -0.63832 | O75915 | O75915 | ARL6IP5 | PRA1 fami | Color2 [Da | 1 | 4 |
| | 0 | NaN | O75928 | O75928 | PIAS2 | E3 SUMO-| | Color2 [Da | 1 | 4 |
| + | 2.360063 | 0.728354 | O75934 | O75934 | BCAS2 | Pre-mRNA | Color2 [Da | 1 | 4 |
| + | 2.793749 | 0.954333 | O75935 | O75935 | DCTN3 | Dynactin s | Color2 [Da | 1 | 4 |
| | 0.71045 | -0.10017 | O75937 | O75937 | DNAJC8 | DnaJ homo | Color2 [Da | 1 | 4 |
| + | 1.525349 | 1.459507 | O75940 | O75940 | SMNDC1 | Survival of | Color2 [Da | 1 | 4 |
| + | 3.706104 | 2.044723 | O75947 | O75947 | ATP5PD | ATP syntha | Color2 [Da | 1 | 4 |
| | 0.613893 | 0.304955 | O75953 | O75953 | DNAJB5 | DnaJ homo | Color2 [Da | 1 | 4 |
| | 1.221012 | -0.23044 | O75955 | O75955 | FLOT1 | Flotillin-1 | Color2 [Da | 1 | 4 |
| + | 2.989708 | -0.84627 | O75962 | O75962 | TRIO | Triple func | Color2 [Da | 1 | 4 |
| + | 3.452026 | 2.66101 | O75964 | O75964 | ATP5MG | ATP syntha | Color2 [Da | 1 | 4 |
| | 1.233872 | 0.194893 | O75970 | O75970 | MPDZ | Multiple P | Color2 [Da | 1 | 4 |
| + | 1.893306 | -0.27318 | O75976 | O75976 | CPD | Carboxype | Color2 [Da | 1 | 4 |
| + | 1.632118 | -0.28888 | O76003 | O76003 | GLRX3 | Glutaredo: | Color2 [Da | 1 | 4 |
| + | 1.797233 | -0.97758 | O76021 | O76021 | RSL1D1 | Ribosomal | Color2 [Da | 1 | 4 |
| + | 0.8844 | 0.509291 | O76024 | O76024 | WFS1 | Wolframin | Color2 [Da | 1 | 4 |
| | 0.227398 | -0.09522 | O76031 | O76031 | CLPX | ATP-deper | Color2 [Da | 1 | 4 |
| + | 2.697332 | 1.961716 | O76038 | O76038 | SCGN | Secretago | Color2 [Da | 1 | 4 |
| + | 1.914495 | 1.423322 | O76041 | O76041 | NEBL | Nebulette | Color2 [Da | 1 | 4 |
| + | 1.10366 | -0.56053 | O76054 | O76054 | SEC14L2 | SEC14-like | Color2 [Da | 1 | 4 |
| + | 2.568745 | -1.38674 | O76062 | O76062 | TM7SF2 | Delta(14)-: | Color2 [Da | 1 | 4 |
| + | 3.178661 | 3.389265 | O76070 | O76070 | SNCG | Gamma-sy | Color2 [Da | 1 | 4 |
| | 0 | -1.87409 | O76071 | O76071 | CIAO1 | Probable c | Color2 [Da | 1 | 4 |
| + | 2.191211 | -0.93126 | O76094 | O76094 | SRP72 | Signal reco | Color2 [Da | 1 | 4 |
| + | 2.334378 | 1.200469 | O94759 | O94759 | TRPM2 | Transient r | Color2 [Da | 1 | 4 |
| + | 3.576749 | 1.609598 | O94760 | O94760 | DDAH1 | N(G),N(G)- | Color2 [Da | 1 | 4 |
| | 0.022904 | -0.00549 | O94763 | O94763 | URI1 | Unconvent | Color2 [Da | 1 | 4 |
| | 0.977238 | -0.25151 | O94766 | O94766 | B3GAT3 | Galactosyl | Color2 [Da | 1 | 4 |
| | 0.48042 | 0.547827 | O94772 | O94772 | LY6H | Lymphocy | Color2 [Da | 1 | 4 |
| + | 2.931495 | -0.68862 | O94776 | O94776 | MTA2 | Metastasis | Color2 [Da | 1 | 4 |
| | 0 | 0.590396 | O94779 | O94779 | CNTN5 | Contactin- | Color2 [Da | 1 | 4 |
| | 0.148589 | -0.19597 | O94805 | O94805 | ACTL6B | Actin-like | Color2 [Da | 1 | 4 |
| | 0 | 1.818868 | O94808 | O94808 | GFPT2 | Glutamine | Color2 [Da | 1 | 4 |
| + | 3.415948 | 3.071573 | O94811 | O94811 | TPPP | Tubulin po | Color2 [Da | 1 | 4 |
| | 0.492184 | 0.126769 | O94813 | O94813 | SLIT2 | Slit homolo | Color2 [Da | 1 | 4 |
| | 0.387184 | 0.151875 | O94817 | O94817 | ATG12 | Ubiquitin-l | Color2 [Da | 1 | 4 |
| | 0.084181 | -0.15201 | O94818 | O94818 | NOL4 | Nucleolar | Color2 [Da | 1 | 4 |
| + | 0.852088 | 0.48037 | O94819 | O94819 | KBTBD11 | Kelch repe | Color2 [Da | 1 | 4 |
| + | 2.732057 | -2.09922 | O94822 | O94822 | LTN1 | E3 ubiquiti | Color2 [Da | 1 | 4 |

| | | | | | | | | | |
|---|---|---|---|---|---|---|---|---|---|
| + | 1.767898 | -0.52335 | O94826 | O94826 | TOMM70 | Mitochonc | Color2 [Da | 1 | 4 |
| | 0 | -1.26737 | O94829 | O94829 | IPO13 | Importin-1 | Color2 [Da | 1 | 4 |
| | 0.092818 | 0.040853 | O94830 | O94830 | DDHD2 | Phospholip | Color2 [Da | 1 | 4 |
| | 0.331795 | -0.06598 | O94842 | O94842 | TOX4 | TOX high n | Color2 [Da | 1 | 4 |
| | 0 | -2.3912 | O94855 | O94855 | SEC24D | Protein tra | Color2 [Da | 1 | 4 |
| | 0.308706 | -0.24056 | O94856 | O94856 | NFASC | Neurofasci | Color2 [Da | 1 | 4 |
| | 0 | -0.86635 | O94864 | O94864 | SUPT7L | STAGA cor | Color2 [Da | 1 | 4 |
| + | 2.414929 | -1.16344 | O94874 | O94874 | UFL1 | E3 UFM1-p | Color2 [Da | 1 | 4 |
| | 0.15323 | 0.061515 | O94875 | O94875 | SORBS2 | Sorbin and | Color2 [Da | 1 | 4 |
| | 1.049007 | -0.15564 | O94876 | O94876 | TMCC1 | Transmem | Color2 [Da | 1 | 4 |
| + | 1.642695 | -1.03108 | O94880 | O94880 | PHF14 | PHD finger | Color2 [Da | 1 | 4 |
| | 0.063543 | 0.025384 | O94888 | O94888 | UBXN7 | UBX doma | Color2 [Da | 1 | 4 |
| | 0 | NaN | O94900 | O94900 | TOX | Thymocyte | Color2 [Da | 1 | 4 |
| + | 0.996904 | -1.23948 | O94901 | O94901 | SUN1 | SUN doma | Color2 [Da | 1 | 4 |
| | 0.221295 | -0.07836 | O94903 | O94903 | PLPBP | Pyridoxal p | Color2 [Da | 1 | 4 |
| + | 1.918829 | -0.67507 | O94905 | O94905 | ERLIN2 | Erlin-2 | Color2 [Da | 1 | 4 |
| + | 2.781469 | -0.99878 | O94906 | O94906 | PRPF6 | Pre-mRNA | Color2 [Da | 1 | 4 |
| | 0 | NaN | O94907 | O94907 | DKK1 | Dickkopf-r | Color2 [Da | 1 | 4 |
| | 0.452077 | -0.20293 | O94910 | O94910 | ADGRL1 | Adhesion ( | Color2 [Da | 1 | 4 |
| | 0.33374 | -0.0819 | O94913 | O94913 | PCF11 | Pre-mRNA | Color2 [Da | 1 | 4 |
| + | 2.289078 | -0.81654 | O94915 | O94915 | FRYL | Protein fur | Color2 [Da | 1 | 4 |
| + | 1.806629 | -1.28085 | O94919 | O94919 | ENDOD1 | Endonucle | Color2 [Da | 1 | 4 |
| | 0.000492 | 0.001816 | O94921 | O94921 | CDK14 | Cyclin-dep | Color2 [Da | 1 | 4 |
| | 0 | 0.573138 | O94923 | O94923 | GLCE | D-glucuror | Color2 [Da | 1 | 4 |
| + | 1.822473 | 0.727178 | O94925 | O94925 | GLS | Glutamina | Color2 [Da | 1 | 4 |
| | 0 | NaN | O94927 | O94927 | HAUS5 | HAUS augr | Color2 [Da | 1 | 4 |
| + | 1.863518 | 0.964861 | O94929 | O94929 | ABLIM3 | Actin-bind | Color2 [Da | 1 | 4 |
| | 0 | -0.77161 | O94952 | O94952 | FBXO21 | F-box only | Color2 [Da | 1 | 4 |
| | 0.611704 | -1.04379 | O94953 | O94953 | KDM4B | Lysine-spe | Color2 [Da | 1 | 4 |
| | 0.2806 | 0.19025 | O94964 | O94964 | SOGA1 | Protein SO | Color2 [Da | 1 | 4 |
| + | 2.213891 | -0.6709 | O94966 | O94966 | USP19 | Ubiquitin c | Color2 [Da | 1 | 4 |
| + | 1.774593 | -0.29018 | O94967 | O94967 | WDR47 | WD repeat | Color2 [Da | 1 | 4 |
| + | 3.839822 | -1.02226 | O94973 | O94973 | AP2A2 | AP-2 comp | Color2 [Da | 1 | 4 |
| + | 2.396494 | -0.92421 | O94979 | O94979 | SEC31A | Protein tra | Color2 [Da | 1 | 4 |
| + | 0.920747 | 0.510494 | O94985 | O94985 | CLSTN1 | Calsynteni | Color2 [Da | 1 | 4 |
| | 0.704536 | -0.43247 | O94992 | O94992 | HEXIM1 | Protein HE | Color2 [Da | 1 | 4 |
| + | 2.517577 | 1.500401 | O95057 | O95057 | DIRAS1 | GTP-bindir | Color2 [Da | 1 | 4 |
| | 0 | NaN | O95070 | O95070 | YIF1A | Protein YIF | Color2 [Da | 1 | 4 |
| | 0.222499 | 0.205377 | O95071 | O95071 | UBR5 | E3 ubiquiti | Color2 [Da | 1 | 4 |
| | 0.108912 | 0.030555 | O95073 | O95073 | FSBP | Fibrinogen | Color2 [Da | 1 | 4 |
| | 0.593093 | 0.292216 | O95084 | O95084 | PRSS23 | Serine pro | Color2 [Da | 1 | 4 |
| + | 1.512043 | 1.029496 | O95104 | O95104 | SCAF4 | SR-related | Color2 [Da | 1 | 4 |
| + | 4.747435 | 3.167068 | O95139 | O95139 | NDUFB6 | NADH deh | Color2 [Da | 1 | 4 |
| | 0.423678 | 0.215014 | O95140 | O95140 | MFN2 | Mitofusin- | Color2 [Da | 1 | 4 |
| | 0.326056 | 0.232304 | O95155 | O95155 | UBE4B | Ubiquitin c | Color2 [Da | 1 | 4 |
| + | 1.07503 | -0.92816 | O95163 | O95163 | ELP1 | Elongator c | Color2 [Da | 1 | 4 |
| | 0 | 0.116783 | O95164 | O95164 | UBL3 | Ubiquitin-l | Color2 [Da | 1 | 4 |

| | | | | | | | | | | |
|---|---|---|---|---|---|---|---|---|---|---|
| + | 1.716951 | 1.8053 | O95166 | O95166 | GABARAP | Gamma-ar | Color2 | [Da | 1 | 4 |
| + | 2.243359 | 2.574581 | O95167 | O95167 | NDUFA3 | NADH deh | Color2 | [Da | 1 | 4 |
| + | 1.819511 | 1.51997 | O95168 | O95168 | NDUFB4 | NADH deh | Color2 | [Da | 1 | 4 |
| | 0.256947 | 0.272521 | O95169 | O95169 | NDUFB8 | NADH deh | Color2 | [Da | 1 | 4 |
| | 0 | NaN | O95196 | O95196 | CSPG5 | Chondroiti | Color2 | [Da | 1 | 4 |
| + | 1.254919 | -0.40881 | O95197 | O95197 | RTN3 | Reticulon-: | Color2 | [Da | 1 | 4 |
| + | 1.219797 | 0.336765 | O95202 | O95202 | LETM1 | Mitochonc | Color2 | [Da | 1 | 4 |
| + | 3.714194 | 1.917323 | O95208 | O95208 | EPN2 | Epsin-2 | Color2 | [Da | 1 | 4 |
| + | 3.31536 | 0.592793 | O95218 | O95218 | ZRANB2 | Zinc finger | Color2 | [Da | 1 | 4 |
| | 0.940965 | -0.22566 | O95219 | O95219 | SNX4 | Sorting ne: | Color2 | [Da | 1 | 4 |
| + | 1.835781 | -0.95109 | O95232 | O95232 | LUC7L3 | Luc7-like p | Color2 | [Da | 1 | 4 |
| | 0 | 1.315851 | O95235 | O95235 | KIF20A | Kinesin-lik | Color2 | [Da | 1 | 4 |
| | 0.776902 | -0.24256 | O95239 | O95239 | KIF4A | Chromoso | Color2 | [Da | 1 | 4 |
| + | 1.70433 | -0.24717 | O95248 | O95248 | SBF1 | Myotubula | Color2 | [Da | 1 | 4 |
| | 0.433476 | 0.710973 | O95249 | O95249 | GOSR1 | Golgi SNAI | Color2 | [Da | 1 | 4 |
| + | 1.271477 | -0.80819 | O95251 | O95251 | KAT7 | Histone ac | Color2 | [Da | 1 | 4 |
| + | 3.061772 | -0.79748 | O95260 | O95260 | ATE1 | Arginyl-tRI | Color2 | [Da | 1 | 4 |
| + | 2.464474 | -1.53552 | O95273 | O95273 | CCNDBP1 | Cyclin-D1-I | Color2 | [Da | 1 | 4 |
| + | 1.610838 | -0.90628 | O95278 | O95278 | EPM2A | Laforin | Color2 | [Da | 1 | 4 |
| + | 2.299096 | 0.554176 | O95292 | O95292 | VAPB | Vesicle-ass | Color2 | [Da | 1 | 4 |
| + | 1.929382 | 1.937431 | O95295 | O95295 | SNAPIN | SNARE-ass | Color2 | [Da | 1 | 4 |
| | 0.060676 | -0.12458 | O95297 | O95297 | MPZL1 | Myelin prc | Color2 | [Da | 1 | 4 |
| + | 0.973816 | 0.590846 | O95298 | O95298 | NDUFC2 | NADH deh | Color2 | [Da | 1 | 4 |
| + | 1.6112 | -1.10869 | O95299 | O95299 | NDUFA10 | NADH deh | Color2 | [Da | 1 | 4 |
| | 0.269589 | 0.139133 | O95302 | O95302 | FKBP9 | Peptidyl-pi | Color2 | [Da | 1 | 4 |
| + | 1.260209 | 0.423176 | O95319 | O95319 | CELF2 | CUGBP Ela | Color2 | [Da | 1 | 4 |
| + | 2.98259 | 0.877244 | O95336 | O95336 | PGLS | 6-phospho | Color2 | [Da | 1 | 4 |
| + | 3.20644 | -0.5895 | O95347 | O95347 | SMC2 | Structural | Color2 | [Da | 1 | 4 |
| + | 1.963897 | -1.33302 | O95352 | O95352 | ATG7 | Ubiquitin-l | Color2 | [Da | 1 | 4 |
| | 0.355907 | -0.06144 | O95359 | O95359 | TACC2 | Transform | Color2 | [Da | 1 | 4 |
| + | 1.766822 | -0.85637 | O95363 | O95363 | FARS2 | Phenylalar | Color2 | [Da | 1 | 4 |
| | 0.835337 | -0.24698 | O95372 | O95372 | LYPLA2 | Acyl-prote | Color2 | [Da | 1 | 4 |
| + | 2.730495 | -1.04513 | O95373 | O95373 | IPO7 | Importin-7 | Color2 | [Da | 1 | 4 |
| + | 3.427696 | -1.3211 | O95376 | O95376 | ARIH2 | E3 ubiquiti | Color2 | [Da | 1 | 4 |
| + | 1.564451 | 1.461023 | O95390 | O95390 | GDF11 | Growth/di | Color2 | [Da | 1 | 4 |
| + | 1.568305 | -0.41852 | O95391 | O95391 | SLU7 | Pre-mRNA | Color2 | [Da | 1 | 4 |
| + | 2.306952 | -0.70264 | O95394 | O95394 | PGM3 | Phosphoac | Color2 | [Da | 1 | 4 |
| | 0 | 0.104293 | O95396 | O95396 | MOCS3 | Adenylyltr | Color2 | [Da | 1 | 4 |
| + | 2.819516 | 1.28487 | O95400 | O95400 | CD2BP2 | CD2 antige | Color2 | [Da | 1 | 4 |
| + | 1.964788 | 1.036096 | O95425 | O95425 | SVIL | Supervillin | Color2 | [Da | 1 | 4 |
| | 0.277154 | -0.40791 | O95429 | O95429 | BAG4 | BAG family | Color2 | [Da | 1 | 4 |
| | 0.307604 | 0.175975 | O95433 | O95433 | AHSA1 | Activator c | Color2 | [Da | 1 | 4 |
| | 0 | 0.077006 | O95453 | O95453 | PARN | Poly(A)-sp | Color2 | [Da | 1 | 4 |
| + | 1.949551 | -0.98629 | O95456 | O95456 | PSMG1 | Proteasom | Color2 | [Da | 1 | 4 |
| | 0.354978 | 0.164644 | O95466 | O95466 | FMNL1 | Formin-lik | Color2 | [Da | 1 | 4 |
| + | 2.435042 | -1.51904 | O95470 | O95470 | SGPL1 | Sphingosir | Color2 | [Da | 1 | 4 |
| + | 2.015766 | -0.95235 | O95478 | O95478 | NSA2 | Ribosome | Color2 | [Da | 1 | 4 |

| | | | | | | | | | |
|---|---|---|---|---|---|---|---|---|---|
| + | 2.442039 | -0.59347 | O95479 | O95479 | H6PD | GDH/6PGL | Color2 [Da | 1 | 4 |
| + | 2.957589 | -1.14393 | O95486 | O95486 | SEC24A | Protein tra | Color2 [Da | 1 | 4 |
| + | 1.415495 | 0.706328 | O95487 | O95487 | SEC24B | Protein tra | Color2 [Da | 1 | 4 |
| + | 3.883722 | -1.9736 | O95490 | O95490 | ADGRL2 | Adhesion ( | Color2 [Da | 1 | 4 |
| + | 2.068272 | 0.956192 | O95502 | O95502 | NPTXR | Neuronal p | Color2 [Da | 1 | 4 |
| | 0 | 0.413128 | O95551 | O95551 | TDP2 | Tyrosyl-DN | Color2 [Da | 1 | 4 |
| + | 3.326241 | 2.286662 | O95563 | O95563 | MPC2 | Mitochonc | Color2 [Da | 1 | 4 |
| + | 1.533147 | 1.260422 | O95571 | O95571 | ETHE1 | Persulfide | Color2 [Da | 1 | 4 |
| + | 2.996721 | -0.79519 | O95573 | O95573 | ACSL3 | Fatty acid | Color2 [Da | 1 | 4 |
| | 0 | NaN | O95602 | O95602 | POLR1A | DNA-direc | Color2 [Da | 1 | 4 |
| | 0 | -0.57659 | O95613 | O95613 | PCNT | Pericentrir | Color2 [Da | 1 | 4 |
| + | 1.186121 | 0.654693 | O95619 | O95619 | YEATS4 | YEATS don | Color2 [Da | 1 | 4 |
| | 0 | 0.183016 | O95622 | O95622 | ADCY5 | Adenylate | Color2 [Da | 1 | 4 |
| + | 1.478991 | 0.580271 | O95630 | O95630 | STAMBP | STAM-binc | Color2 [Da | 1 | 4 |
| + | 0.615796 | 1.542351 | O95631 | O95631 | NTN1 | Netrin-1 | Color2 [Da | 1 | 4 |
| + | 2.471798 | 1.546395 | O95670 | O95670 | ATP6V1G2 | V-type pro | Color2 [Da | 1 | 4 |
| + | 1.391769 | -0.508 | O95671 | O95671 | ASMTL | Probable b | Color2 [Da | 1 | 4 |
| + | 3.974084 | -1.75673 | O95674 | O95674 | CDS2 | Phosphatic | Color2 [Da | 1 | 4 |
| | 0.765647 | 0.477991 | O95678 | O95678 | KRT75 | Keratin, ty | Color2 [Da | 1 | 4 |
| + | 1.317016 | 0.279368 | O95684 | O95684 | CEP43 | Centrosom | Color2 [Da | 1 | 4 |
| | 0.229822 | 0.326334 | O95696 | O95696 | BRD1 | Bromodon | Color2 [Da | 1 | 4 |
| | 0.489743 | -0.56258 | O95714 | O95714 | HERC2 | E3 ubiquiti | Color2 [Da | 1 | 4 |
| | 0 | 0.50489 | O95715 | O95715 | CXCL14 | C-X-C moti | Color2 [Da | 1 | 4 |
| | 0.405383 | -0.14899 | O95716 | O95716 | RAB3D | Ras-relate | Color2 [Da | 1 | 4 |
| + | 1.600022 | 0.477887 | O95721 | O95721 | SNAP29 | Synaptoso | Color2 [Da | 1 | 4 |
| | 0 | -2.28346 | O95741 | O95741 | CPNE6 | Copine-6 | Color2 [Da | 1 | 4 |
| | 0.859067 | -0.25343 | O95747 | O95747 | OXSR1 | Serine/thr | Color2 [Da | 1 | 4 |
| | 0.987468 | 0.288689 | O95749 | O95749 | GGPS1 | Geranylge | Color2 [Da | 1 | 4 |
| + | 1.684223 | -0.42696 | O95757 | O95757 | HSPA4L | Heat shocl | Color2 [Da | 1 | 4 |
| | 0.840115 | 0.109554 | O95758 | O95758 | PTBP3 | Polypyrimi | Color2 [Da | 1 | 4 |
| + | 1.90773 | -1.96395 | O95772 | O95772 | STARD3NL | STARD3 N- | Color2 [Da | 1 | 4 |
| + | 1.432087 | 1.10471 | O95777 | O95777 | LSM8 | U6 snRNA- | Color2 [Da | 1 | 4 |
| + | 1.480365 | -0.25659 | O95782 | O95782 | AP2A1 | AP-2 comp | Color2 [Da | 1 | 4 |
| | 0.40999 | 0.427916 | O95785 | O95785 | WIZ | Protein Wi | Color2 [Da | 1 | 4 |
| + | 2.879181 | 1.495533 | O95793 | O95793 | STAU1 | Double-str | Color2 [Da | 1 | 4 |
| | 0 | -1.11565 | O95801 | O95801 | TTC4 | Tetratricop | Color2 [Da | 1 | 4 |
| + | 1.083144 | 0.375527 | O95816 | O95816 | BAG2 | BAG family | Color2 [Da | 1 | 4 |
| + | 3.958559 | 3.583772 | O95817 | O95817 | BAG3 | BAG family | Color2 [Da | 1 | 4 |
| + | 1.974276 | -0.61493 | O95819 | O95819 | MAP4K4 | Mitogen-a | Color2 [Da | 1 | 4 |
| + | 1.593212 | -0.88863 | O95822 | O95822 | MLYCD | Malonyl-C | Color2 [Da | 1 | 4 |
| + | 2.080271 | 0.962353 | O95825 | O95825 | CRYZL1 | Quinone o | Color2 [Da | 1 | 4 |
| + | 2.842163 | 1.001732 | O95831 | O95831 | AIFM1 | Apoptosis- | Color2 [Da | 1 | 4 |
| | 0 | 1.78552 | O95834 | O95834 | EML2 | Echinoderi | Color2 [Da | 1 | 4 |
| + | 2.006521 | 0.667915 | O95848 | O95848 | NUDT14 | Uridine dip | Color2 [Da | 1 | 4 |
| | 0.558682 | 0.228399 | O95861 | O95861 | BPNT1 | 3'(2'),5'-bis | Color2 [Da | 1 | 4 |
| + | 1.154296 | -0.81821 | O95864 | O95864 | FADS2 | Acyl-CoA 6 | Color2 [Da | 1 | 4 |
| + | 1.428286 | 0.591745 | O95865 | O95865 | DDAH2 | N(G),N(G)- | Color2 [Da | 1 | 4 |

| | | | | | | | | | |
|---|---|---|---|---|---|---|---|---|---|
| + | 1.555535 | -1.06671 | O95870 | O95870 | ABHD16A | Phosphati | Color2 [Da | 1 | 4 |
| + | 2.433895 | 1.591896 | O95881 | O95881 | TXNDC12 | Thioredoxi | Color2 [Da | 1 | 4 |
| | 0.161575 | 0.06296 | O95897 | O95897 | OLFM2 | Noelin-2 | Color2 [Da | 1 | 4 |
| | 0.073836 | 0.160948 | O95926 | O95926 | SYF2 | Pre-mRNA | Color2 [Da | 1 | 4 |
| + | 1.801239 | -0.89246 | O95948 | O95948 | ONECUT2 | One cut dc | Color2 [Da | 1 | 4 |
| | 0.487833 | -0.04324 | O95983 | O95983 | MBD3 | Methyl-Cp | Color2 [Da | 1 | 4 |
| | 0 | NaN | O95985 | O95985 | TOP3B | DNA topoi | Color2 [Da | 1 | 4 |
| + | 2.141658 | 0.811271 | O95989 | O95989 | NUDT3 | Diphosphc | Color2 [Da | 1 | 4 |
| + | 0.707586 | 1.520825 | O95996 | O95996 | APC2 | Adenomat | Color2 [Da | 1 | 4 |
| | 0.635012 | 1.002382 | O95999 | O95999 | BCL10 | B-cell lymp | Color2 [Da | 1 | 4 |
| + | 1.740228 | 0.627672 | O96000 | O96000 | NDUFB10 | NADH deh | Color2 [Da | 1 | 4 |
| | 0.331954 | -0.22692 | O96005 | O96005 | CLPTM1 | Cleft lip an | Color2 [Da | 1 | 4 |
| + | 2.211183 | 0.854528 | O96006 | O96006 | ZBED1 | E3 SUMO-| | Color2 [Da | 1 | 4 |
| + | 3.230245 | 1.518532 | O96007 | O96007 | MOCS2 | Molybdop | Color2 [Da | 1 | 4 |
| | 0.241533 | 0.111579 | O96008 | O96008 | TOMM40 | Mitochonc | Color2 [Da | 1 | 4 |
| + | 1.932813 | -0.71957 | O96011 | O96011 | PEX11B | Peroxisom | Color2 [Da | 1 | 4 |
| | 0 | 0.118791 | O96013 | O96013 | PAK4 | Serine/thr | Color2 [Da | 1 | 4 |
| + | 3.389967 | -0.89227 | O96019 | O96019 | ACTL6A | Actin-like p | Color2 [Da | 1 | 4 |
| + | 2.14927 | -0.62819 | O96028 | O96028 | NSD2 | Histone-ly | Color2 [Da | 1 | 4 |
| | 0.466483 | 0.644869 | P00156 | P00156 | MT-CYB | Cytochron | Color2 [Da | 1 | 4 |
| + | 3.581231 | 2.154087 | P00167 | P00167 | CYB5A | Cytochron | Color2 [Da | 1 | 4 |
| + | 2.4787 | -1.25257 | P00338 | P00338 | LDHA | L-lactate d | Color2 [Da | 1 | 4 |
| | 0.64608 | -0.52952 | P00352 | P00352 | ALDH1A1 | Aldehyde ( | Color2 [Da | 1 | 4 |
| | 0.350747 | -0.09718 | P00367 | P00367 | GLUD1 | Glutamate | Color2 [Da | 1 | 4 |
| + | 2.291556 | 0.789291 | P00374 | P00374 | DHFR | Dihydrofol | Color2 [Da | 1 | 4 |
| + | 2.121589 | -1.23441 | P00387 | P00387 | CYB5R3 | NADH-cyto | Color2 [Da | 1 | 4 |
| + | 4.535034 | 2.335866 | P00390 | P00390 | GSR | Glutathion | Color2 [Da | 1 | 4 |
| + | 1.869237 | -1.49205 | P00395 | P00395 | MT-CO1 | Cytochron | Color2 [Da | 1 | 4 |
| + | 3.0276 | -1.17184 | P00403 | P00403 | MT-CO2 | Cytochron | Color2 [Da | 1 | 4 |
| | 0 | NaN | P00414 | P00414 | MT-CO3 | Cytochron | Color2 [Da | 1 | 4 |
| + | 3.179324 | 3.316597 | P00441 | P00441 | SOD1 | Superoxide | Color2 [Da | 1 | 4 |
| + | 1.184856 | 0.741765 | P00450 | P00450 | CP | Ceruloplas | Color2 [Da | 1 | 4 |
| | 0 | NaN | P00480 | P00480 | OTC | Ornithine t | Color2 [Da | 1 | 4 |
| + | 2.130798 | 1.564116 | P00491 | P00491 | PNP | Purine nuc | Color2 [Da | 1 | 4 |
| + | 1.137463 | 1.362581 | P00492 | P00492 | HPRT1 | Hypoxanth | Color2 [Da | 1 | 4 |
| + | 3.091078 | 1.669937 | P00505 | P00505 | GOT2 | Aspartate | Color2 [Da | 1 | 4 |
| + | 1.020983 | 1.063732 | P00519 | P00519 | ABL1 | Tyrosine-p | Color2 [Da | 1 | 4 |
| + | 3.124286 | -2.30231 | P00533 | P00533 | EGFR | Epidermal | Color2 [Da | 1 | 4 |
| | 0.91321 | -0.16136 | P00558 | P00558 | PGK1 | Phosphogl | Color2 [Da | 1 | 4 |
| + | 2.023236 | 1.262935 | P00568 | P00568 | AK1 | Adenylate | Color2 [Da | 1 | 4 |
| | 0 | -0.23588 | P00734 | P00734 | F2 | Prothromb | Color2 [Da | 1 | 4 |
| | 0 | NaN | P00738 | P00738 | HP | Haptoglob | Color2 [Da | 1 | 4 |
| | 0 | NaN | P00750 | P00750 | PLAT | Tissue-typ | Color2 [Da | 1 | 4 |
| + | 2.12748 | 1.89253 | P00813 | P00813 | ADA | Adenosine | Color2 [Da | 1 | 4 |
| | 0 | -0.03422 | P00846 | P00846 | MT-ATP6 | ATP syntha | Color2 [Da | 1 | 4 |
| | 0 | NaN | P00915 | P00915 | CA1 | Carbonic a | Color2 [Da | 1 | 4 |
| + | 2.76828 | 0.7875 | P00918 | P00918 | CA2 | Carbonic a | Color2 [Da | 1 | 4 |

| | | | | | | | | | |
|---|---|---|---|---|---|---|---|---|---|
| + | 3.722065 | -1.8049 | P00966 | P00966 | ASS1 | Argininosu | Color2 [Da | 1 | 4 |
| + | 3.732955 | 2.302872 | P01008 | P01008 | SERPINC1 | Antithrom | Color2 [Da | 1 | 4 |
| + | 2.536186 | 2.590112 | P01019 | P01019 | AGT | Angiotensi | Color2 [Da | 1 | 4 |
| + | 1.994375 | -0.66431 | P01023 | P01023 | A2M | Alpha-2-m | Color2 [Da | 1 | 4 |
| | 0.508397 | 1.207373 | P01024 | P01024 | C3 | Compleme | Color2 [Da | 1 | 4 |
| | 0.223587 | -0.18142 | P01034 | P01034 | CST3 | Cystatin-C | Color2 [Da | 1 | 4 |
| + | 1.666881 | 0.728158 | P01040 | P01040 | CSTA | Cystatin-A | Color2 [Da | 1 | 4 |
| + | 1.860753 | 0.940247 | P01111 | P01111 | NRAS | GTPase NF | Color2 [Da | 1 | 4 |
| | 0.182703 | -0.06685 | P01112 | P01112 | HRAS | GTPase HF | Color2 [Da | 1 | 4 |
| + | 1.232002 | 0.696567 | P01116 | P01116 | KRAS | GTPase KR | Color2 [Da | 1 | 4 |
| | 0 | NaN | P01130 | P01130 | LDLR | Low-densi | Color2 [Da | 1 | 4 |
| | 0 | 2.205301 | P01210 | P01210 | PENK | Proenkeph | Color2 [Da | 1 | 4 |
| + | 1.598607 | 0.867863 | P01303 | P01303 | NPY | Pro-neuro | Color2 [Da | 1 | 4 |
| | 0.985655 | -0.30404 | P01591 | P01591 | JCHAIN | Immunogl | Color2 [Da | 1 | 4 |
| | 0.630758 | -0.93058 | P01833 | P01833 | PIGR | Polymeric | Color2 [Da | 1 | 4 |
| + | 2.294928 | -1.31461 | P01834 | P01834 | IGKC | Immunogl | Color2 [Da | 1 | 4 |
| + | 2.223723 | -2.55525 | P01857;P0 | P01857;P0 | IGHG1; | Immunogl | Color2 [Da | 1 | 4 |
| + | 2.229738 | -4.70105 | P01859 | P01859 | IGHG2 | Immunogl | Color2 [Da | 1 | 4 |
| | 0 | NaN | P01871;P0 | P01871;P0 | IGHM; | Immunogl | Color2 [Da | 1 | 4 |
| + | 3.337093 | -1.18523 | P01876 | P01876 | IGHA1 | Immunogl | Color2 [Da | 1 | 4 |
| + | 2.636784 | -1.64442 | P01877 | P01877 | IGHA2 | Immunogl | Color2 [Da | 1 | 4 |
| + | 2.224119 | -1.12818 | P01889 | P01889 | HLA-B | HLA class I | Color2 [Da | 1 | 4 |
| | 0 | 0.068346 | P02042 | P02042 | HBD | Hemoglob | Color2 [Da | 1 | 4 |
| + | 3.298771 | 2.022357 | P02100;P6 | P02100;P6 | HBE1;HBG | Hemoglob | Color2 [Da | 1 | 4 |
| + | 1.391481 | -1.56068 | P02452 | P02452 | COL1A1 | Collagen a | Color2 [Da | 1 | 4 |
| + | 1.8939 | -0.9098 | P02458 | P02458 | COL2A1 | Collagen a | Color2 [Da | 1 | 4 |
| | 0 | NaN | P02461 | P02461 | COL3A1 | Collagen a | Color2 [Da | 1 | 4 |
| + | 3.138817 | 1.992338 | P02511 | P02511 | CRYAB | Alpha-crys | Color2 [Da | 1 | 4 |
| + | 3.261916 | 0.982002 | P02533 | P02533 | KRT14 | Keratin, ty | Color2 [Da | 1 | 4 |
| + | 3.176623 | 0.258351 | P02538 | P02538 | KRT6A | Keratin, ty | Color2 [Da | 1 | 4 |
| | 0.433345 | -0.07762 | P02545 | P02545 | LMNA | Prelamin-A | Color2 [Da | 1 | 4 |
| + | 1.579545 | -0.62631 | P02647 | P02647 | APOA1 | Apolipopro | Color2 [Da | 1 | 4 |
| + | 4.616709 | -1.90025 | P02649 | P02649 | APOE | Apolipopro | Color2 [Da | 1 | 4 |
| | 0 | NaN | P02655 | P02655 | APOC2 | Apolipopro | Color2 [Da | 1 | 4 |
| | 0.506948 | 1.797911 | P02656 | P02656 | APOC3 | Apolipopro | Color2 [Da | 1 | 4 |
| | 0 | -2.80309 | P02679 | P02679 | FGG | Fibrinogen | Color2 [Da | 1 | 4 |
| | 0 | NaN | P02689 | P02689 | PMP2 | Myelin P2 | Color2 [Da | 1 | 4 |
| | 0 | NaN | P02748 | P02748 | C9 | Compleme | Color2 [Da | 1 | 4 |
| + | 2.305154 | -0.78198 | P02751 | P02751 | FN1 | Fibronecti | Color2 [Da | 1 | 4 |
| + | 3.57008 | -3.08514 | P02766 | P02766 | TTR | Transthyre | Color2 [Da | 1 | 4 |
| + | 2.488429 | -0.92069 | P02768 | P02768 | ALB | Albumin | Color2 [Da | 1 | 4 |
| | 0 | NaN | P02774 | P02774 | GC | Vitamin D- | Color2 [Da | 1 | 4 |
| + | 3.349883 | -1.785 | P02786 | P02786 | TFRC | Transferrir | Color2 [Da | 1 | 4 |
| + | 2.624104 | 0.651307 | P02787 | P02787 | TF | Serotransf | Color2 [Da | 1 | 4 |
| + | 1.691053 | -1.18329 | P02788 | P02788 | LTF | Lactotrans | Color2 [Da | 1 | 4 |
| | 0 | -0.69403 | P02790 | P02790 | HPX | Hemopexi | Color2 [Da | 1 | 4 |
| + | 2.795029 | 3.588432 | P02792 | P02792 | FTL | Ferritin ligl | Color2 [Da | 1 | 4 |

-Bruker Confidential-

| | | | | | | | | | |
|---|---|---|---|---|---|---|---|---|---|
| + | 3.414765 | 3.221547 | P02794 | P02794 | FTH1 | Ferritin he | Color2 [Da | 1 | 4 |
| | 0 | NaN | P03891 | P03891 | MT-ND2 | NADH-ubi | Color2 [Da | 1 | 4 |
| + | 2.667632 | 1.857055 | P03928 | P03928 | MT-ATP8 | ATP syntha | Color2 [Da | 1 | 4 |
| | 0.516923 | -0.22403 | P04004 | P04004 | VTN | Vitronectir | Color2 [Da | 1 | 4 |
| + | 1.478555 | 0.292503 | P04040 | P04040 | CAT | Catalase | Color2 [Da | 1 | 4 |
| + | 0.714646 | -0.64329 | P04049 | P04049 | RAF1 | RAF proto- | Color2 [Da | 1 | 4 |
| | 0.859373 | 0.094152 | P04062 | P04062 | GBA | Lysosomal | Color2 [Da | 1 | 4 |
| + | 1.170086 | 0.808605 | P04066 | P04066 | FUCA1 | Tissue alpł | Color2 [Da | 1 | 4 |
| + | 3.372561 | -0.7316 | P04075 | P04075 | ALDOA | Fructose-b | Color2 [Da | 1 | 4 |
| + | 1.615196 | 2.535986 | P04080 | P04080 | CSTB | Cystatin-B | Color2 [Da | 1 | 4 |
| + | 2.551637 | 1.213535 | P04083 | P04083 | ANXA1 | Annexin A: | Color2 [Da | 1 | 4 |
| + | 1.674037 | 1.363087 | P04156 | P04156 | PRNP | Major prio | Color2 [Da | 1 | 4 |
| + | 2.376332 | 2.169827 | P04179 | P04179 | SOD2 | Superoxide | Color2 [Da | 1 | 4 |
| + | 1.405144 | 0.282811 | P04181 | P04181 | OAT | Ornithine a | Color2 [Da | 1 | 4 |
| + | 0.69894 | 1.550653 | P04216 | P04216 | THY1 | Thy-1 men | Color2 [Da | 1 | 4 |
| + | 2.025125 | 1.006777 | P04259 | P04259 | KRT6B | Keratin, ty | Color2 [Da | 1 | 4 |
| + | 3.373026 | 1.340842 | P04264 | P04264 | KRT1 | Keratin, ty | Color2 [Da | 1 | 4 |
| + | 2.675308 | 2.322995 | P04271 | P04271 | S100B | Protein S1 | Color2 [Da | 1 | 4 |
| + | 1.502695 | 0.269139 | P04350 | P04350 | TUBB4A | Tubulin be | Color2 [Da | 1 | 4 |
| + | 2.892347 | 0.997677 | P04406 | P04406 | GAPDH | Glyceralde | Color2 [Da | 1 | 4 |
| + | 2.473484 | 1.250719 | P04424 | P04424 | ASL | Argininosu | Color2 [Da | 1 | 4 |
| | 0.403756 | -0.10178 | P04439 | P04439 | HLA-A | HLA class I | Color2 [Da | 1 | 4 |
| | 0 | NaN | P04626 | P04626 | ERBB2 | Receptor t | Color2 [Da | 1 | 4 |
| | 0 | NaN | P04628 | P04628 | WNT1 | Proto-onco | Color2 [Da | 1 | 4 |
| + | 5.367964 | 0.777387 | P04632 | P04632 | CAPNS1 | Calpain sm | Color2 [Da | 1 | 4 |
| + | 3.045935 | 2.07187 | P04792 | P04792 | HSPB1 | Heat shocl | Color2 [Da | 1 | 4 |
| + | 1.234565 | -1.14572 | P04818 | P04818 | TYMS | Thymidyla | Color2 [Da | 1 | 4 |
| + | 3.521536 | -1.49687 | P04843 | P04843 | RPN1 | Dolichyl-di | Color2 [Da | 1 | 4 |
| + | 3.53957 | -1.52579 | P04844 | P04844 | RPN2 | Dolichyl-di | Color2 [Da | 1 | 4 |
| + | 2.698977 | -1.24016 | P04899 | P04899 | GNAI2 | Guanine n | Color2 [Da | 1 | 4 |
| + | 1.546791 | 0.956028 | P04908;Q7 | P04908;Q7 | H2AC8;H2. | Histone H2 | Color2 [Da | 1 | 4 |
| + | 1.963443 | -0.98235 | P04920 | P04920 | SLC4A2 | Anion exch | Color2 [Da | 1 | 4 |
| | 0 | NaN | P05019 | P05019 | IGF1 | Insulin-like | Color2 [Da | 1 | 4 |
| + | 2.933178 | -1.07191 | P05023 | P05023 | ATP1A1 | Sodium/pc | Color2 [Da | 1 | 4 |
| | 0.454413 | -0.24401 | P05026 | P05026 | ATP1B1 | Sodium/pc | Color2 [Da | 1 | 4 |
| + | 2.776829 | 1.132537 | P05060 | P05060 | CHGB | Secretogra | Color2 [Da | 1 | 4 |
| | 0.434701 | -0.48176 | P05062 | P05062 | ALDOB | Fructose-b | Color2 [Da | 1 | 4 |
| + | 1.864598 | 0.587792 | P05067 | P05067 | APP | Amyloid-b | Color2 [Da | 1 | 4 |
| + | 2.530207 | 1.147222 | P05089 | P05089 | ARG1 | Arginase-1 | Color2 [Da | 1 | 4 |
| | 0.976647 | -0.17993 | P05091 | P05091 | ALDH2 | Aldehyde c | Color2 [Da | 1 | 4 |
| | 0.064609 | 0.087852 | P05109 | P05109 | S100A8 | Protein S1 | Color2 [Da | 1 | 4 |
| | 0.004386 | -0.02014 | P05114 | P05114 | HMGN1 | Non-histor | Color2 [Da | 1 | 4 |
| | 0 | -0.62839 | P05120 | P05120 | SERPINB2 | Plasminog | Color2 [Da | 1 | 4 |
| | 0 | NaN | P05129 | P05129 | PRKCG | Protein kir | Color2 [Da | 1 | 4 |
| | 0.450608 | 0.153104 | P05141 | P05141 | SLC25A5 | ADP/ATP t | Color2 [Da | 1 | 4 |
| | 0 | NaN | P05156 | P05156 | CFI | Compleme | Color2 [Da | 1 | 4 |
| | 0.006044 | 0.00605 | P05161 | P05161 | ISG15 | Ubiquitin-l | Color2 [Da | 1 | 4 |

| | | | | | | | | | |
|---|---|---|---|---|---|---|---|---|---|
| | 0 | -3.19693 | P05164 | P05164 | MPO | Myeloperc | Color2 [Da | 1 | 4 |
| + | 2.914736 | -1.08599 | P05165 | P05165 | PCCA | Propionyl- | Color2 [Da | 1 | 4 |
| + | 2.945318 | -0.77104 | P05166 | P05166 | PCCB | Propionyl- | Color2 [Da | 1 | 4 |
| | 0 | NaN | P05186 | P05186 | ALPL | Alkaline pł | Color2 [Da | 1 | 4 |
| + | 1.557592 | -0.37166 | P05198 | P05198 | EIF2S1 | Eukaryotic | Color2 [Da | 1 | 4 |
| | 0 | NaN | P05230 | P05230 | FGF1 | Fibroblast | Color2 [Da | 1 | 4 |
| | 0 | NaN | P05362 | P05362 | ICAM1 | Intercellul | Color2 [Da | 1 | 4 |
| + | 2.742133 | 2.149473 | P05386 | P05386 | RPLP1 | 60S acidic | Color2 [Da | 1 | 4 |
| + | 2.696167 | 2.054402 | P05387 | P05387 | RPLP2 | 60S acidic | Color2 [Da | 1 | 4 |
| | 0.929685 | -0.29541 | P05388 | P05388 | RPLP0 | 60S acidic | Color2 [Da | 1 | 4 |
| | 0.183021 | -0.2027 | P05408 | P05408 | SCG5 | Neuroend | Color2 [Da | 1 | 4 |
| + | 2.671975 | 4.683861 | P05413 | P05413 | FABP3 | Fatty acid- | Color2 [Da | 1 | 4 |
| + | 0.906416 | -0.51431 | P05423 | P05423 | POLR3D | DNA-direc | Color2 [Da | 1 | 4 |
| + | 2.193649 | 0.687353 | P05455 | P05455 | SSB | Lupus La p | Color2 [Da | 1 | 4 |
| + | 2.043181 | -2.33032 | P05496;P4 | P05496;P4 | ATP5MC1; | ATP syntha | Color2 [Da | 1 | 4 |
| | 0 | NaN | P05546 | P05546 | SERPIND1 | Heparin cc | Color2 [Da | 1 | 4 |
| | 0 | NaN | P05549;Q6 | P05549;Q6 | TFAP2A;TF | Transcripti | Color2 [Da | 1 | 4 |
| + | 2.041267 | -0.88218 | P05556 | P05556 | ITGB1 | Integrin be | Color2 [Da | 1 | 4 |
| | 0.293595 | 0.258233 | P05771 | P05771 | PRKCB | Protein kir | Color2 [Da | 1 | 4 |
| + | 1.162567 | 0.436756 | P05783 | P05783 | KRT18 | Keratin, ty | Color2 [Da | 1 | 4 |
| + | 2.403399 | 0.82174 | P05787 | P05787 | KRT8 | Keratin, ty | Color2 [Da | 1 | 4 |
| + | 2.599206 | -1.49593 | P05937 | P05937 | CALB1 | Calbindin | Color2 [Da | 1 | 4 |
| | 0.436595 | 0.781406 | P05997 | P05997 | COL5A2 | Collagen a | Color2 [Da | 1 | 4 |
| + | 1.969034 | 0.806693 | P06132 | P06132 | UROD | Uroporphy | Color2 [Da | 1 | 4 |
| + | 0.931941 | -0.45955 | P06241 | P06241 | FYN | Tyrosine-p | Color2 [Da | 1 | 4 |
| | 0 | NaN | P06280 | P06280 | GLA | Alpha-gala | Color2 [Da | 1 | 4 |
| + | 1.272868 | 0.239411 | P06396 | P06396 | GSN | Gelsolin | Color2 [Da | 1 | 4 |
| | 0 | NaN | P06400 | P06400 | RB1 | Retinoblas | Color2 [Da | 1 | 4 |
| + | 4.176495 | 2.063467 | P06454 | P06454 | PTMA | Prothymos | Color2 [Da | 1 | 4 |
| + | 2.400337 | -0.83713 | P06493 | P06493 | CDK1 | Cyclin-dep | Color2 [Da | 1 | 4 |
| + | 5.068946 | 2.175912 | P06576 | P06576 | ATP5F1B | ATP syntha | Color2 [Da | 1 | 4 |
| | 0.078273 | -0.04156 | P06702 | P06702 | S100A9 | Protein S1 | Color2 [Da | 1 | 4 |
| + | 1.946514 | 0.9484 | P06703 | P06703 | S100A6 | Protein S1 | Color2 [Da | 1 | 4 |
| | 0.274206 | 0.13478 | P06730 | P06730 | EIF4E | Eukaryotic | Color2 [Da | 1 | 4 |
| | 0.167322 | -0.03587 | P06732 | P06732 | CKM | Creatine ki | Color2 [Da | 1 | 4 |
| + | 1.519032 | -0.42146 | P06733 | P06733 | ENO1 | Alpha-eno | Color2 [Da | 1 | 4 |
| + | 1.993428 | -0.78851 | P06737 | P06737 | PYGL | Glycogen p | Color2 [Da | 1 | 4 |
| + | 2.148985 | -0.62364 | P06744 | P06744 | GPI | Glucose-6- | Color2 [Da | 1 | 4 |
| | 0.170573 | -0.04555 | P06746 | P06746 | POLB | DNA polyn | Color2 [Da | 1 | 4 |
| | 0.926798 | -0.16348 | P06748 | P06748 | NPM1 | Nucleopho | Color2 [Da | 1 | 4 |
| + | 2.70551 | 1.201421 | P06753 | P06753 | TPM3 | Tropomyo | Color2 [Da | 1 | 4 |
| + | 1.05124 | -0.4858 | P06756 | P06756 | ITGAV | Integrin al | Color2 [Da | 1 | 4 |
| + | 3.096755 | 2.857593 | P06850 | P06850 | CRH | Corticolibe | Color2 [Da | 1 | 4 |
| + | 4.663784 | 2.345784 | P06858 | P06858 | LPL | Lipoprotei | Color2 [Da | 1 | 4 |
| + | 2.697842 | 0.740373 | P06865 | P06865 | HEXA | Beta-hexo | Color2 [Da | 1 | 4 |
| + | 3.304884 | 0.860547 | P06899 | P06899 | H2BC11 | Histone H2 | Color2 [Da | 1 | 4 |
| + | 3.201568 | 0.679642 | P07093 | P07093 | SERPINE2 | Glia-derive | Color2 [Da | 1 | 4 |

| | | | | | | | | | |
|---|---|---|---|---|---|---|---|---|---|
| + | 3.04396 | -1.59572 | P07099 | P07099 | EPHX1 | Epoxide hy | Color2 [Da | 1 | 4 |
| + | 2.778926 | 2.649203 | P07108 | P07108 | DBI | Acyl-CoA-b | Color2 [Da | 1 | 4 |
| + | 2.507366 | 0.413446 | P07195 | P07195 | LDHB | L-lactate d | Color2 [Da | 1 | 4 |
| + | 6.071365 | 1.975877 | P07196 | P07196 | NEFL | Neurofilan | Color2 [Da | 1 | 4 |
| + | 3.851574 | 1.279015 | P07197 | P07197 | NEFM | Neurofilan | Color2 [Da | 1 | 4 |
| | 0.094048 | -0.09751 | P07199 | P07199 | CENPB | Major cent | Color2 [Da | 1 | 4 |
| + | 1.71699 | 0.699109 | P07203 | P07203 | GPX1 | Glutathion | Color2 [Da | 1 | 4 |
| + | 1.625964 | -0.601 | P07205 | P07205 | PGK2 | Phosphogl | Color2 [Da | 1 | 4 |
| + | 2.109177 | -0.20248 | P07237 | P07237 | P4HB | Protein dis | Color2 [Da | 1 | 4 |
| | 0.105251 | 0.057002 | P07305 | P07305 | H1-0 | Histone H1 | Color2 [Da | 1 | 4 |
| + | 2.749212 | 2.071888 | P07311 | P07311 | ACYP1 | Acylphosp | Color2 [Da | 1 | 4 |
| + | 3.887375 | 1.471416 | P07339 | P07339 | CTSD | Cathepsin | Color2 [Da | 1 | 4 |
| | 0.268626 | -0.05303 | P07355 | P07355 | ANXA2 | Annexin A | Color2 [Da | 1 | 4 |
| + | 1.835223 | -0.17796 | P07384 | P07384 | CAPN1 | Calpain-1 c | Color2 [Da | 1 | 4 |
| + | 1.284434 | -0.36914 | P07437 | P07437 | TUBB | Tubulin be | Color2 [Da | 1 | 4 |
| | 0.40899 | 1.743872 | P07477;P0 | P07477;P0 | PRSS1;PRS | Serine pro | Color2 [Da | 1 | 4 |
| | 0 | 2.389619 | P07585 | P07585 | DCN | Decorin | Color2 [Da | 1 | 4 |
| + | 3.580407 | 1.253001 | P07602 | P07602 | PSAP | Prosaposir | Color2 [Da | 1 | 4 |
| + | 2.98666 | 1.436706 | P07686 | P07686 | HEXB | Beta-hexo | Color2 [Da | 1 | 4 |
| | 1.111842 | 0.168748 | P07737 | P07737 | PFN1 | Profilin-1 | Color2 [Da | 1 | 4 |
| | 0.676101 | 0.592835 | P07738 | P07738 | BPGM | Bisphosph | Color2 [Da | 1 | 4 |
| + | 2.713477 | 0.970477 | P07741 | P07741 | APRT | Adenine pl | Color2 [Da | 1 | 4 |
| + | 3.591644 | -1.43874 | P07814 | P07814 | EPRS1 | Bifunction | Color2 [Da | 1 | 4 |
| + | 0.986584 | 0.474392 | P07858 | P07858 | CTSB | Cathepsin | Color2 [Da | 1 | 4 |
| + | 3.264746 | -1.04255 | P07900 | P07900 | HSP90AA1 | Heat shock | Color2 [Da | 1 | 4 |
| + | 1.591771 | -0.61235 | P07902 | P07902 | GALT | Galactose- | Color2 [Da | 1 | 4 |
| + | 2.794981 | 1.260823 | P07910 | P07910 | HNRNPC | Heterogen | Color2 [Da | 1 | 4 |
| + | 1.072259 | 0.531071 | P07919 | P07919 | UQCRH | Cytochrom | Color2 [Da | 1 | 4 |
| | 0 | NaN | P07942 | P07942 | LAMB1 | Laminin su | Color2 [Da | 1 | 4 |
| + | 2.292305 | -0.89224 | P07947 | P07947 | YES1 | Tyrosine-p | Color2 [Da | 1 | 4 |
| + | 1.430282 | 1.694175 | P07951 | P07951 | TPM2 | Tropomyo | Color2 [Da | 1 | 4 |
| + | 3.770708 | 1.496848 | P07954 | P07954 | FH | Fumarate | Color2 [Da | 1 | 4 |
| + | 2.242965 | -1.74194 | P07996 | P07996 | THBS1 | Thrombos | Color2 [Da | 1 | 4 |
| + | 2.611914 | -1.77232 | P08069 | P08069 | IGF1R | Insulin-like | Color2 [Da | 1 | 4 |
| + | 6.405336 | -3.54001 | P08123 | P08123 | COL1A2 | Collagen a | Color2 [Da | 1 | 4 |
| + | 3.577119 | -0.58497 | P08133 | P08133 | ANXA6 | Annexin A | Color2 [Da | 1 | 4 |
| | 0.502012 | -0.28033 | P08134 | P08134 | RHOC | Rho-relate | Color2 [Da | 1 | 4 |
| | 1.1235 | -0.17488 | P08195 | P08195 | SLC3A2 | 4F2 cell-su | Color2 [Da | 1 | 4 |
| + | 3.175702 | 2.311478 | P08236 | P08236 | GUSB | Beta-glucu | Color2 [Da | 1 | 4 |
| + | 2.029896 | -0.6407 | P08237 | P08237 | PFKM | ATP-deper | Color2 [Da | 1 | 4 |
| + | 2.957482 | -0.90918 | P08238 | P08238 | HSP90AB1 | Heat shock | Color2 [Da | 1 | 4 |
| + | 3.526857 | -1.18594 | P08240 | P08240 | SRPRA | Signal reco | Color2 [Da | 1 | 4 |
| + | 1.603644 | -0.76014 | P08243 | P08243 | ASNS | Asparagine | Color2 [Da | 1 | 4 |
| | 0 | NaN | P08246 | P08246 | ELANE | Neutrophil | Color2 [Da | 1 | 4 |
| | 0.365783 | 0.620051 | P08247 | P08247 | SYP | Synaptoph | Color2 [Da | 1 | 4 |
| | 0.19853 | 0.391236 | P08254 | P08254 | MMP3 | Stromelysi | Color2 [Da | 1 | 4 |
| | 0 | NaN | P08311 | P08311 | CTSG | Cathepsin | Color2 [Da | 1 | 4 |

| | | | | | | | | | |
|---|---|---|---|---|---|---|---|---|---|
| + | 1.643781 | 1.156712 | P08397 | P08397 | HMBS | Porphobili | Color2 [Da | 1 | 4 |
| + | 2.387875 | -0.95584 | P08559 | P08559 | PDHA1 | Pyruvate d | Color2 [Da | 1 | 4 |
| + | 2.32522 | -1.93257 | P08572 | P08572 | COL4A2 | Collagen a | Color2 [Da | 1 | 4 |
| + | 1.560802 | -0.89223 | P08574 | P08574 | CYC1 | Cytochron | Color2 [Da | 1 | 4 |
| + | 3.009091 | 2.013176 | P08579 | P08579 | SNRPB2 | U2 small n | Color2 [Da | 1 | 4 |
| + | 2.288548 | 0.561844 | P08621 | P08621 | SNRNP70 | U1 small n | Color2 [Da | 1 | 4 |
| | 0 | -1.50609 | P08648 | P08648 | ITGA5 | Integrin al | Color2 [Da | 1 | 4 |
| + | 5.151777 | 2.003112 | P08670 | P08670 | VIM | Vimentin | Color2 [Da | 1 | 4 |
| + | 2.744966 | 1.614199 | P08708 | P08708 | RPS17 | 40S riboso | Color2 [Da | 1 | 4 |
| + | 1.542155 | -0.51975 | P08727 | P08727 | KRT19 | Keratin, ty | Color2 [Da | 1 | 4 |
| + | 3.833576 | -2.00977 | P08754 | P08754 | GNAI3 | Guanine n | Color2 [Da | 1 | 4 |
| + | 1.831081 | 0.624431 | P08758 | P08758 | ANXA5 | Annexin A! | Color2 [Da | 1 | 4 |
| + | 1.667488 | -0.41033 | P08779 | P08779 | KRT16 | Keratin, ty | Color2 [Da | 1 | 4 |
| | 0 | NaN | P08842 | P08842 | STS | Steryl-sulf | Color2 [Da | 1 | 4 |
| + | 3.051838 | 0.970382 | P08865 | P08865 | RPSA | 40S riboso | Color2 [Da | 1 | 4 |
| | 0 | NaN | P08913 | P08913 | ADRA2A | Alpha-2A a | Color2 [Da | 1 | 4 |
| + | 1.790143 | 1.600104 | P08949 | P08949 | NMB | Neuromed | Color2 [Da | 1 | 4 |
| + | 3.588057 | -1.43466 | P08962 | P08962 | CD63 | CD63 antig | Color2 [Da | 1 | 4 |
| + | 3.464296 | 2.254949 | P09012 | P09012 | SNRPA | U1 small n | Color2 [Da | 1 | 4 |
| | 0.678906 | -0.41969 | P09038 | P09038 | FGF2 | Fibroblast | Color2 [Da | 1 | 4 |
| | 0.249702 | 0.252898 | P09086 | P09086 | POU2F2 | POU doma | Color2 [Da | 1 | 4 |
| + | 3.865869 | 1.135501 | P09104 | P09104 | ENO2 | Gamma-er | Color2 [Da | 1 | 4 |
| | 0.040721 | -0.01425 | P09110 | P09110 | ACAA1 | 3-ketoacyl | Color2 [Da | 1 | 4 |
| | 0.014828 | -0.03101 | P09132 | P09132 | SRP19 | Signal recc | Color2 [Da | 1 | 4 |
| + | 1.088179 | 0.543142 | P09211 | P09211 | GSTP1 | Glutathion | Color2 [Da | 1 | 4 |
| + | 1.314252 | 1.509909 | P09234 | P09234 | SNRPC | U1 small n | Color2 [Da | 1 | 4 |
| + | 1.341303 | 0.426881 | P09382 | P09382 | LGALS1 | Galectin-1 | Color2 [Da | 1 | 4 |
| + | 3.660893 | 0.830438 | P09417 | P09417 | QDPR | Dihydropt | Color2 [Da | 1 | 4 |
| | 0.692061 | -0.21514 | P09429 | P09429 | HMGB1 | High mobil | Color2 [Da | 1 | 4 |
| + | 1.832919 | 0.488477 | P09455 | P09455 | RBP1 | Retinol-bir | Color2 [Da | 1 | 4 |
| | 0.056423 | -0.01363 | P09471 | P09471 | GNAO1 | Guanine n | Color2 [Da | 1 | 4 |
| | 0.004584 | 0.005547 | P09486 | P09486 | SPARC | SPARC | Color2 [Da | 1 | 4 |
| | 0 | NaN | P09488 | P09488 | GSTM1 | Glutathion | Color2 [Da | 1 | 4 |
| + | 1.21994 | 1.114839 | P09493 | P09493 | TPM1 | Tropomyo | Color2 [Da | 1 | 4 |
| + | 2.462357 | 1.462066 | P09496 | P09496 | CLTA | Clathrin lig | Color2 [Da | 1 | 4 |
| + | 3.23166 | 3.033575 | P09497 | P09497 | CLTB | Clathrin lig | Color2 [Da | 1 | 4 |
| + | 1.750857 | 0.205738 | P09525 | P09525 | ANXA4 | Annexin A | Color2 [Da | 1 | 4 |
| | 0.42932 | -0.092 | P09543 | P09543 | CNP | 2',3'-cyclic | Color2 [Da | 1 | 4 |
| | 0.795086 | 0.451984 | P09601 | P09601 | HMOX1 | Heme oxyg | Color2 [Da | 1 | 4 |
| | 0 | NaN | P09619 | P09619 | PDGFRB | Platelet-de | Color2 [Da | 1 | 4 |
| + | 3.592598 | 1.284878 | P09622 | P09622 | DLD | Dihydrolip | Color2 [Da | 1 | 4 |
| + | 1.358606 | 0.805104 | P09651 | P09651 | HNRNPA1 | Heterogen | Color2 [Da | 1 | 4 |
| + | 3.456162 | 2.87548 | P09661 | P09661 | SNRPA1 | U2 small n | Color2 [Da | 1 | 4 |
| + | 2.483862 | 1.550586 | P09669 | P09669 | COX6C | Cytochron | Color2 [Da | 1 | 4 |
| + | 2.44987 | -0.51278 | P09874 | P09874 | PARP1 | Poly [ADP- | Color2 [Da | 1 | 4 |
| + | 3.10312 | 0.675546 | P09936 | P09936 | UCHL1 | Ubiquitin c | Color2 [Da | 1 | 4 |
| + | 2.078144 | -0.70656 | P09960 | P09960 | LTA4H | Leukotrien | Color2 [Da | 1 | 4 |

| | | | | | | | | | |
|---|---|---|---|---|---|---|---|---|---|
| + | 3.345059 | 1.561437 | P09972 | P09972 | ALDOC | Fructose-b | Color2 [Da | 1 | 4 |
| | 0 | NaN | P0C0L4 | P0C0L4 | C4A | Compleme | Color2 [Da | 1 | 4 |
| + | 2.639623 | 1.592293 | P0C0S5;Q7 | P0C0S5;Q7 | H2AZ1;H2A | Histone H2 | Color2 [Da | 1 | 4 |
| + | 1.741568 | 1.065497 | P0C0S8;Q9 | P0C0S8;Q9 | H2AC17;H2 | Histone H2 | Color2 [Da | 1 | 4 |
| + | 1.372371 | 0.491894 | P0C870 | P0C870 | JMJD7 | Bifunction | Color2 [Da | 1 | 4 |
| + | 0.939067 | -0.35707 | P0CAP2 | P0CAP2 | POLR2M | DNA-direc | Color2 [Da | 1 | 4 |
| + | 3.716523 | 3.265781 | P0CG34;P0 | P0CG34;P0 | TMSB15A;T | Thymosin | Color2 [Da | 1 | 4 |
| | 0.218366 | 0.282368 | P0CG38 | P0CG38 | POTEI | POTE anky | Color2 [Da | 1 | 4 |
| + | 4.049668 | 2.695968 | P0CG47;P0 | P0CG47;P0 | UBB;UBC;R | Polyubiqui | Color2 [Da | 1 | 4 |
| | 0 | NaN | P0CW18 | P0CW18 | PRSS56 | Serine pro | Color2 [Da | 1 | 4 |
| + | 1.767224 | 1.088679 | P0DI81;P0 | P0DI81;P0 | TRAPPC2;T | Trafficking | Color2 [Da | 1 | 4 |
| + | 2.940061 | 1.629961 | P0DJ93 | P0DJ93 | SMIM13 | Small integ | Color2 [Da | 1 | 4 |
| | 0.162272 | -0.02352 | P0DMV8;P | P0DMV8;P | HSPA1A;HS | Heat shock | Color2 [Da | 1 | 4 |
| + | 1.774344 | -0.73787 | P0DOY2;P0 | P0DOY2;P0 | IGLC2;IGLC | Immunoglo | Color2 [Da | 1 | 4 |
| + | 2.523768 | 1.089447 | P0DP23;P0 | P0DP23;P0 | CALM1;CA | Calmodulin | Color2 [Da | 1 | 4 |
| | 0 | NaN | P0DP57 | P0DP57 | SLURP2 | Secreted L | Color2 [Da | 1 | 4 |
| + | 2.604749 | 1.977569 | P0DPH7;P0 | P0DPH7;P0 | TUBA3C;TU | Tubulin alp | Color2 [Da | 1 | 4 |
| | 0 | 1.934186 | P0DTE7;P0 | P0DTE7;P0 | AMY1B;AM | Alpha-amy | Color2 [Da | 1 | 4 |
| | 0.466015 | 0.586923 | P10109 | P10109 | FDX1 | Adrenodox | Color2 [Da | 1 | 4 |
| + | 3.263946 | 0.711864 | P10114 | P10114 | RAP2A | Ras-relate | Color2 [Da | 1 | 4 |
| + | 2.275265 | -0.34993 | P10155 | P10155 | RO60 | RNA-bindi | Color2 [Da | 1 | 4 |
| | 0 | -4.98159 | P10163 | P10163 | PRB4 | Basic saliva | Color2 [Da | 1 | 4 |
| + | 2.314246 | 0.867973 | P10253 | P10253 | GAA | Lysosomal | Color2 [Da | 1 | 4 |
| + | 1.49306 | -0.70431 | P10301 | P10301 | RRAS | Ras-relate | Color2 [Da | 1 | 4 |
| | 0.162244 | 0.187784 | P10321 | P10321 | HLA-C | HLA class I | Color2 [Da | 1 | 4 |
| | 0 | NaN | P10398 | P10398 | ARAF | Serine/thr | Color2 [Da | 1 | 4 |
| + | 1.871876 | 0.782505 | P10412 | P10412 | H1-4 | Histone H1 | Color2 [Da | 1 | 4 |
| | 0 | NaN | P10415 | P10415 | BCL2 | Apoptosis | Color2 [Da | 1 | 4 |
| | 0.381582 | 0.990035 | P10451 | P10451 | SPP1 | Osteopont | Color2 [Da | 1 | 4 |
| | 0.54273 | 0.257733 | P10515 | P10515 | DLAT | Dihydrolip | Color2 [Da | 1 | 4 |
| | 0.93761 | 0.091952 | P10586 | P10586 | PTPRF | Receptor-t | Color2 [Da | 1 | 4 |
| + | 3.805078 | -2.41485 | P10589 | P10589 | NR2F1 | COUP tran | Color2 [Da | 1 | 4 |
| + | 2.78585 | 1.488793 | P10599 | P10599 | TXN | Thioredoxi | Color2 [Da | 1 | 4 |
| + | 2.300598 | 2.456383 | P10606 | P10606 | COX5B | Cytochrom | Color2 [Da | 1 | 4 |
| | 0.539382 | 0.767581 | P10619 | P10619 | CTSA | Lysosomal | Color2 [Da | 1 | 4 |
| + | 1.314495 | 0.998751 | P10620 | P10620 | MGST1 | Microsoma | Color2 [Da | 1 | 4 |
| + | 5.262328 | 3.866859 | P10636 | P10636 | MAPT | Microtubu | Color2 [Da | 1 | 4 |
| | 0.172945 | 0.050415 | P10644 | P10644 | PRKAR1A | cAMP-dep | Color2 [Da | 1 | 4 |
| + | 3.460229 | 0.944455 | P10645 | P10645 | CHGA | Chromogra | Color2 [Da | 1 | 4 |
| + | 3.100237 | -1.43376 | P10768 | P10768 | ESD | S-formylgl | Color2 [Da | 1 | 4 |
| + | 3.024228 | 1.106863 | P10809 | P10809 | HSPD1 | 60 kDa hea | Color2 [Da | 1 | 4 |
| + | 3.219726 | 1.528055 | P10909 | P10909 | CLU | Clusterin | Color2 [Da | 1 | 4 |
| | 1.586574 | 0.178628 | P11021 | P11021 | HSPA5 | Endoplasm | Color2 [Da | 1 | 4 |
| + | 3.020953 | -1.9645 | P11047 | P11047 | LAMC1 | Laminin su | Color2 [Da | 1 | 4 |
| | 0.287282 | 0.184167 | P11117 | P11117 | ACP2 | Lysosomal | Color2 [Da | 1 | 4 |
| + | 3.60454 | 1.551427 | P11137 | P11137 | MAP2 | Microtubu | Color2 [Da | 1 | 4 |
| | 1.374886 | 0.209675 | P11142 | P11142 | HSPA8 | Heat shock | Color2 [Da | 1 | 4 |

| | | | | | | | | | |
|---|---|---|---|---|---|---|---|---|---|
| + | 3.377759 | -2.05775 | P11166 | P11166 | SLC2A1 | Solute carr | Color2 [Da | 1 | 4 |
| + | 2.853602 | -1.68352 | P11169 | P11169 | SLC2A3 | Solute carr | Color2 [Da | 1 | 4 |
| | 0.824419 | -0.34052 | P11171 | P11171 | EPB41 | Protein 4.1 | Color2 [Da | 1 | 4 |
| + | 2.197644 | -1.61845 | P11172 | P11172 | UMPS | Uridine 5'- | Color2 [Da | 1 | 4 |
| + | 1.634258 | 0.407988 | P11177 | P11177 | PDHB | Pyruvate d | Color2 [Da | 1 | 4 |
| + | 1.695844 | -0.18921 | P11182 | P11182 | DBT | Lipoamide | Color2 [Da | 1 | 4 |
| + | 2.139324 | -0.78631 | P11216 | P11216 | PYGB | Glycogen p | Color2 [Da | 1 | 4 |
| + | 1.956099 | 0.936878 | P11217 | P11217 | PYGM | Glycogen p | Color2 [Da | 1 | 4 |
| + | 2.247981 | 1.002387 | P11233 | P11233 | RALA | Ras-relate | Color2 [Da | 1 | 4 |
| + | 2.668806 | 0.826073 | P11234 | P11234 | RALB | Ras-relate | Color2 [Da | 1 | 4 |
| + | 2.364431 | 0.662128 | P11274 | P11274 | BCR | Breakpoint | Color2 [Da | 1 | 4 |
| + | 2.130066 | 0.831411 | P11279 | P11279 | LAMP1 | Lysosome- | Color2 [Da | 1 | 4 |
| | 0.112225 | 0.023176 | P11310 | P11310 | ACADM | Medium-c | Color2 [Da | 1 | 4 |
| | 0 | NaN | P11362 | P11362 | FGFR1 | Fibroblast | Color2 [Da | 1 | 4 |
| + | 2.79508 | -1.27997 | P11387 | P11387 | TOP1 | DNA topoi | Color2 [Da | 1 | 4 |
| + | 3.468971 | -1.37502 | P11388 | P11388 | TOP2A | DNA topoi | Color2 [Da | 1 | 4 |
| + | 3.446336 | -0.79667 | P11413 | P11413 | G6PD | Glucose-6- | Color2 [Da | 1 | 4 |
| + | 2.662205 | 1.362638 | P11441 | P11441 | UBL4A | Ubiquitin-l | Color2 [Da | 1 | 4 |
| | 0.31251 | 0.095333 | P11498 | P11498 | PC | Pyruvate c | Color2 [Da | 1 | 4 |
| + | 2.77909 | 0.730077 | P11532 | P11532 | DMD | Dystrophin | Color2 [Da | 1 | 4 |
| + | 3.25398 | -0.78179 | P11586 | P11586 | MTHFD1 | C-1-tetrah | Color2 [Da | 1 | 4 |
| + | 3.268326 | -2.0917 | P11717 | P11717 | IGF2R | Cation-ind | Color2 [Da | 1 | 4 |
| + | 3.021237 | -1.53316 | P11766 | P11766 | ADH5 | Alcohol de | Color2 [Da | 1 | 4 |
| | 0.397757 | 0.299872 | P11908 | P11908 | PRPS2 | Ribose-pho | Color2 [Da | 1 | 4 |
| + | 1.765777 | 0.501017 | P11940 | P11940 | PABPC1 | Polyadeny | Color2 [Da | 1 | 4 |
| + | 1.72915 | -0.61948 | P12004 | P12004 | PCNA | Proliferatir | Color2 [Da | 1 | 4 |
| | 0.435111 | 0.273417 | P12036 | P12036 | NEFH | Neurofilan | Color2 [Da | 1 | 4 |
| + | 1.115249 | 0.922114 | P12074 | P12074 | COX6A1 | Cytochrom | Color2 [Da | 1 | 4 |
| + | 0.945439 | -0.38543 | P12081 | P12081 | HARS1 | Histidine-- | Color2 [Da | 1 | 4 |
| + | 1.991342 | -1.39502 | P12107 | P12107 | COL11A1 | Collagen a | Color2 [Da | 1 | 4 |
| + | 3.678539 | -1.65271 | P12109 | P12109 | COL6A1 | Collagen a | Color2 [Da | 1 | 4 |
| | 0 | NaN | P12110 | P12110 | COL6A2 | Collagen a | Color2 [Da | 1 | 4 |
| + | 0.927761 | -0.4324 | P12111 | P12111 | COL6A3 | Collagen a | Color2 [Da | 1 | 4 |
| + | 0.967722 | 0.565739 | P12235 | P12235 | SLC25A4 | ADP/ATP t | Color2 [Da | 1 | 4 |
| | 0.128757 | -0.11153 | P12236 | P12236 | SLC25A6 | ADP/ATP t | Color2 [Da | 1 | 4 |
| + | 2.718505 | 1.590193 | P12259 | P12259 | F5 | Coagulatio | Color2 [Da | 1 | 4 |
| + | 2.442921 | -1.16754 | P12268 | P12268 | IMPDH2 | Inosine-5'- | Color2 [Da | 1 | 4 |
| + | 2.328607 | -0.34444 | P12270 | P12270 | TPR | Nucleopro | Color2 [Da | 1 | 4 |
| + | 1.614199 | 0.944817 | P12273 | P12273 | PIP | Prolactin-i | Color2 [Da | 1 | 4 |
| + | 4.000443 | 0.855046 | P12277 | P12277 | CKB | Creatine ki | Color2 [Da | 1 | 4 |
| + | 0.691771 | -1.59504 | P12429 | P12429 | ANXA3 | Annexin A3 | Color2 [Da | 1 | 4 |
| + | 2.79557 | 1.720256 | P12532 | P12532 | CKMT1B | Creatine ki | Color2 [Da | 1 | 4 |
| | 0 | NaN | P12645 | P12645 | BMP3 | Bone morp | Color2 [Da | 1 | 4 |
| | 0.058362 | -0.01743 | P12694 | P12694 | BCKDHA | 2-oxoisova | Color2 [Da | 1 | 4 |
| + | 3.79649 | -1.62161 | P12814 | P12814 | ACTN1 | Alpha-actir | Color2 [Da | 1 | 4 |
| | 0 | 0.844569 | P12830 | P12830 | CDH1 | Cadherin-1 | Color2 [Da | 1 | 4 |
| | 0.624079 | 1.179079 | P12882 | P12882 | MYH1 | Myosin-1 | Color2 [Da | 1 | 4 |

| | | | | | | | | | |
|---|---|---|---|---|---|---|---|---|---|
| | 0 | NaN | P12883 | P12883 | MYH7 | Myosin-7 | Color2 [Da | 1 | 4 |
| + | 2.30838 | 0.413203 | P12931 | P12931 | SRC | Proto-onc | Color2 [Da | 1 | 4 |
| + | 2.677203 | 1.790003 | P12955 | P12955 | PEPD | Xaa-Pro di | Color2 [Da | 1 | 4 |
| + | 2.599908 | -0.69293 | P12956 | P12956 | XRCC6 | X-ray repa | Color2 [Da | 1 | 4 |
| + | 3.353471 | -1.4009 | P13010 | P13010 | XRCC5 | X-ray repa | Color2 [Da | 1 | 4 |
| | 0.170378 | -0.38019 | P13051 | P13051 | UNG | Uracil-DN/ | Color2 [Da | 1 | 4 |
| + | 3.297579 | 1.942339 | P13073 | P13073 | COX4I1 | Cytochron | Color2 [Da | 1 | 4 |
| | 0 | NaN | P13164;Q( | P13164;Q( | IFITM1;IFI | Interferon | Color2 [Da | 1 | 4 |
| + | 1.804846 | 0.981382 | P13473 | P13473 | LAMP2 | Lysosome- | Color2 [Da | 1 | 4 |
| + | 1.529066 | -0.53636 | P13489 | P13489 | RNH1 | Ribonuclea | Color2 [Da | 1 | 4 |
| | 0.018599 | 0.02891 | P13497 | P13497 | BMP1 | Bone morp | Color2 [Da | 1 | 4 |
| + | 1.448599 | -0.21306 | P13521 | P13521 | SCG2 | Secretogra | Color2 [Da | 1 | 4 |
| + | 2.845717 | -0.78245 | P13591 | P13591 | NCAM1 | Neural cell | Color2 [Da | 1 | 4 |
| | 0.928393 | -0.29948 | P13611 | P13611 | VCAN | Versican c | Color2 [Da | 1 | 4 |
| + | 3.006167 | -0.66623 | P13637 | P13637 | ATP1A3 | Sodium/pc | Color2 [Da | 1 | 4 |
| + | 3.731151 | -1.14594 | P13639 | P13639 | EEF2 | Elongation | Color2 [Da | 1 | 4 |
| + | 4.249386 | 1.921652 | P13645 | P13645 | KRT10 | Keratin, ty | Color2 [Da | 1 | 4 |
| + | 1.998432 | 1.111518 | P13646 | P13646 | KRT13 | Keratin, ty | Color2 [Da | 1 | 4 |
| + | 2.783657 | 1.192495 | P13647 | P13647 | KRT5 | Keratin, ty | Color2 [Da | 1 | 4 |
| | 0.66405 | 0.00975 | P13667 | P13667 | PDIA4 | Protein dis | Color2 [Da | 1 | 4 |
| + | 3.85587 | -2.27531 | P13674 | P13674 | P4HA1 | Prolyl 4-hy | Color2 [Da | 1 | 4 |
| | 0.669869 | 0.22791 | P13693 | P13693 | TPT1 | Translatior | Color2 [Da | 1 | 4 |
| + | 2.666834 | 1.598702 | P13716 | P13716 | ALAD | Delta-amir | Color2 [Da | 1 | 4 |
| | 0 | NaN | P13796 | P13796 | LCP1 | Plastin-2 | Color2 [Da | 1 | 4 |
| | 0.187216 | -0.17894 | P13797 | P13797 | PLS3 | Plastin-3 | Color2 [Da | 1 | 4 |
| | 1.132034 | 0.200467 | P13798 | P13798 | APEH | Acylamino | Color2 [Da | 1 | 4 |
| + | 1.807528 | 0.431031 | P13804 | P13804 | ETFA | Electron tr | Color2 [Da | 1 | 4 |
| + | 2.648988 | -1.18168 | P13807 | P13807 | GYS1 | Glycogen [ | Color2 [Da | 1 | 4 |
| | 1.098542 | 0.202663 | P13861 | P13861 | PRKAR2A | cAMP-dep | Color2 [Da | 1 | 4 |
| + | 0.774845 | 2.296749 | P13929 | P13929 | ENO3 | Beta-enola | Color2 [Da | 1 | 4 |
| | 0.985343 | 0.281495 | P13984 | P13984 | GTF2F2 | General tr | Color2 [Da | 1 | 4 |
| + | 1.440084 | 0.604343 | P13987 | P13987 | CD59 | CD59 glyc | Color2 [Da | 1 | 4 |
| | 0.030252 | -0.01564 | P13995 | P13995 | MTHFD2 | Bifunction | Color2 [Da | 1 | 4 |
| | 0 | NaN | P14136 | P14136 | GFAP | Glial fibrill | Color2 [Da | 1 | 4 |
| + | 3.602174 | 2.031764 | P14174 | P14174 | MIF | Macropha | Color2 [Da | 1 | 4 |
| + | 2.392789 | 3.852094 | P14209 | P14209 | CD99 | CD99 antig | Color2 [Da | 1 | 4 |
| + | 1.473129 | 0.3136 | P14314 | P14314 | PRKCSH | Glucosidas | Color2 [Da | 1 | 4 |
| | 0 | NaN | P14317 | P14317 | HCLS1 | Hematopo | Color2 [Da | 1 | 4 |
| | 0.260282 | -0.24315 | P14324 | P14324 | FDPS | Farnesyl p | Color2 [Da | 1 | 4 |
| + | 2.261165 | 1.586689 | P14406 | P14406 | COX7A2 | Cytochron | Color2 [Da | 1 | 4 |
| + | 0.984403 | -0.57785 | P14415 | P14415 | ATP1B2 | Sodium/pc | Color2 [Da | 1 | 4 |
| + | 1.420703 | -0.39023 | P14550 | P14550 | AKR1A1 | Aldo-keto | Color2 [Da | 1 | 4 |
| + | 3.503004 | -1.19245 | P14618 | P14618 | PKM | Pyruvate k | Color2 [Da | 1 | 4 |
| + | 2.595968 | 2.248594 | P14621 | P14621 | ACYP2 | Acylphosp | Color2 [Da | 1 | 4 |
| + | 2.637856 | -0.62597 | P14625 | P14625 | HSP90B1 | Endoplasm | Color2 [Da | 1 | 4 |
| | 0 | NaN | P14635 | P14635 | CCNB1 | G2/mitotic | Color2 [Da | 1 | 4 |
| + | 1.847671 | 1.073148 | P14649 | P14649 | MYL6B | Myosin ligl | Color2 [Da | 1 | 4 |

| | | | | | | | | | |
|---|---|---|---|---|---|---|---|---|---|
| + | 1.41597 | -0.43324 | P14678 | P14678 | SNRPB | Small nucl | Color2 [Da | 1 | 4 |
| + | 0.963004 | -0.54857 | P14735 | P14735 | IDE | Insulin-deg | Color2 [Da | 1 | 4 |
| + | 1.311576 | 0.730787 | P14854 | P14854 | COX6B1 | Cytochrom | Color2 [Da | 1 | 4 |
| + | 1.569026 | -0.50365 | P14859 | P14859 | POU2F1 | POU doma | Color2 [Da | 1 | 4 |
| + | 3.715977 | -0.89899 | P14866 | P14866 | HNRNPL | Heterogen | Color2 [Da | 1 | 4 |
| + | 1.985705 | -0.65844 | P14868 | P14868 | DARS1 | Aspartate- | Color2 [Da | 1 | 4 |
| | 0.352641 | 0.10629 | P14923 | P14923 | JUP | Junction p | Color2 [Da | 1 | 4 |
| + | 2.88801 | 1.577733 | P14927 | P14927 | UQCRB | Cytochrom | Color2 [Da | 1 | 4 |
| | 0.40242 | -0.16731 | P15056 | P15056 | BRAF | Serine/thr | Color2 [Da | 1 | 4 |
| | 0.611859 | -1.12452 | P15104 | P15104 | GLUL | Glutamine | Color2 [Da | 1 | 4 |
| | 0.650535 | -0.19332 | P15121 | P15121 | AKR1B1 | Aldo-keto | Color2 [Da | 1 | 4 |
| | 0 | NaN | P15151 | P15151 | PVR | Poliovirus | Color2 [Da | 1 | 4 |
| + | 2.563338 | -0.87933 | P15170 | P15170 | GSPT1 | Eukaryotic | Color2 [Da | 1 | 4 |
| + | 3.779173 | 0.6833 | P15289 | P15289 | ARSA | Arylsulfata | Color2 [Da | 1 | 4 |
| + | 2.852329 | 0.782597 | P15311 | P15311 | EZR | Ezrin | Color2 [Da | 1 | 4 |
| | 0.209902 | -0.95195 | P15313 | P15313 | ATP6V1B1 | V-type pro | Color2 [Da | 1 | 4 |
| + | 1.562064 | 0.8235 | P15336 | P15336 | ATF2 | Cyclic AMI | Color2 [Da | 1 | 4 |
| + | 1.803023 | -0.53216 | P15374 | P15374 | UCHL3 | Ubiquitin c | Color2 [Da | 1 | 4 |
| + | 2.03944 | 0.528377 | P15531 | P15531 | NME1 | Nucleoside | Color2 [Da | 1 | 4 |
| + | 0.864731 | 0.545162 | P15559 | P15559 | NQO1 | NAD(P)H d | Color2 [Da | 1 | 4 |
| + | 3.08999 | 1.25262 | P15586 | P15586 | GNS | N-acetylglu | Color2 [Da | 1 | 4 |
| + | 3.61399 | 1.932491 | P15848 | P15848 | ARSB | Arylsulfata | Color2 [Da | 1 | 4 |
| + | 1.197219 | 0.501784 | P15880 | P15880 | RPS2 | 40S riboso | Color2 [Da | 1 | 4 |
| + | 2.085995 | -0.51394 | P15882 | P15882 | CHN1 | N-chimaer | Color2 [Da | 1 | 4 |
| + | 1.430042 | 0.544513 | P15924 | P15924 | DSP | Desmoplal | Color2 [Da | 1 | 4 |
| + | 2.796383 | 0.48298 | P15927 | P15927 | RPA2 | Replicatior | Color2 [Da | 1 | 4 |
| + | 2.402539 | 0.522867 | P16035 | P16035 | TIMP2 | Metallopro | Color2 [Da | 1 | 4 |
| + | 2.281421 | 1.414263 | P16070 | P16070 | CD44 | CD44 antig | Color2 [Da | 1 | 4 |
| + | 1.841535 | 0.884458 | P16083 | P16083 | NQO2 | Ribosyldih | Color2 [Da | 1 | 4 |
| + | 3.003981 | 2.57089 | P16104 | P16104 | H2AX | Histone H2 | Color2 [Da | 1 | 4 |
| + | 1.394424 | 0.421153 | P16152 | P16152 | CBR1 | Carbonyl r | Color2 [Da | 1 | 4 |
| + | 1.716371 | 1.738275 | P16219 | P16219 | ACADS | Short-chai | Color2 [Da | 1 | 4 |
| + | 1.478082 | 1.13545 | P16220 | P16220 | CREB1 | Cyclic AMI | Color2 [Da | 1 | 4 |
| | 0.212194 | -0.06237 | P16278 | P16278 | GLB1 | Beta-galac | Color2 [Da | 1 | 4 |
| + | 1.309232 | 0.237749 | P16298 | P16298 | PPP3CB | Serine/thr | Color2 [Da | 1 | 4 |
| | 1.176767 | 0.229932 | P16333 | P16333 | NCK1 | Cytoplasm | Color2 [Da | 1 | 4 |
| + | 2.679345 | 1.46242 | P16401 | P16401 | H1-5 | Histone H1 | Color2 [Da | 1 | 4 |
| + | 1.734604 | 1.174869 | P16402 | P16402 | H1-3 | Histone H1 | Color2 [Da | 1 | 4 |
| + | 1.959627 | 0.90347 | P16403 | P16403 | H1-2 | Histone H1 | Color2 [Da | 1 | 4 |
| + | 3.021952 | -1.15056 | P16435 | P16435 | POR | NADPH--cy | Color2 [Da | 1 | 4 |
| | 0.195136 | -0.16116 | P16455 | P16455 | MGMT | Methylate | Color2 [Da | 1 | 4 |
| | 0.055553 | -0.06676 | P16591 | P16591 | FER | Tyrosine-p | Color2 [Da | 1 | 4 |
| + | 2.467132 | -1.21036 | P16615 | P16615 | ATP2A2 | Sarcoplasn | Color2 [Da | 1 | 4 |
| + | 1.596268 | 0.456616 | P16870 | P16870 | CPE | Carboxype | Color2 [Da | 1 | 4 |
| + | 2.22264 | 1.319171 | P16930 | P16930 | FAH | Fumarylac | Color2 [Da | 1 | 4 |
| + | 3.128657 | 2.598456 | P16949 | P16949 | STMN1 | Stathmin | Color2 [Da | 1 | 4 |
| + | 3.232097 | 3.237997 | P16989 | P16989 | YBX3 | Y-box-binc | Color2 [Da | 1 | 4 |

| | | | | | | | | | |
|---|---|---|---|---|---|---|---|---|---|
| | 0.783979 | -0.2006 | P17028 | P17028 | ZNF24 | Zinc finger | Color2 [Da | 1 | 4 |
| + | 1.540633 | -0.56987 | P17029 | P17029 | ZKSCAN1 | Zinc finger | Color2 [Da | 1 | 4 |
| | 0.861687 | -0.24994 | P17050 | P17050 | NAGA | Alpha-N-ac | Color2 [Da | 1 | 4 |
| | 0 | NaN | P17066 | P17066 | HSPA6 | Heat shock | Color2 [Da | 1 | 4 |
| + | 1.596423 | 0.922251 | P17096 | P17096 | HMGA1 | High mobil | Color2 [Da | 1 | 4 |
| + | 0.829002 | -0.97667 | P17098 | P17098 | ZNF8 | Zinc finger | Color2 [Da | 1 | 4 |
| + | 1.505294 | -0.5346 | P17152 | P17152 | TMEM11 | Transmem | Color2 [Da | 1 | 4 |
| + | 3.506227 | 2.434305 | P17174 | P17174 | GOT1 | Aspartate | Color2 [Da | 1 | 4 |
| | 0 | NaN | P17213 | P17213 | BPI | Bactericida | Color2 [Da | 1 | 4 |
| + | 1.098318 | -0.43397 | P17252 | P17252 | PRKCA | Protein kir | Color2 [Da | 1 | 4 |
| | 0 | -1.70011 | P17301 | P17301 | ITGA2 | Integrin al | Color2 [Da | 1 | 4 |
| + | 0.98816 | -0.49069 | P17302 | P17302 | GJA1 | Gap juncti | Color2 [Da | 1 | 4 |
| + | 1.243827 | 0.929946 | P17405 | P17405 | SMPD1 | Sphingomy | Color2 [Da | 1 | 4 |
| + | 2.695091 | -0.48854 | P17480 | P17480 | UBTF | Nucleolar t | Color2 [Da | 1 | 4 |
| | 0.137036 | -0.20877 | P17535 | P17535 | JUND | Transcripti | Color2 [Da | 1 | 4 |
| | 0 | NaN | P17544 | P17544 | ATF7 | Cyclic AMF | Color2 [Da | 1 | 4 |
| | 0.157514 | 0.077098 | P17568 | P17568 | NDUFB7 | NADH deh | Color2 [Da | 1 | 4 |
| + | 3.427291 | 2.563097 | P17600 | P17600 | SYN1 | Synapsin-1 | Color2 [Da | 1 | 4 |
| + | 2.096645 | -0.57382 | P17612 | P17612 | PRKACA | cAMP-dep | Color2 [Da | 1 | 4 |
| + | 3.283167 | -1.01345 | P17655 | P17655 | CAPN2 | Calpain-2 c | Color2 [Da | 1 | 4 |
| | 0 | -0.37039 | P17661 | P17661 | DES | Desmin | Color2 [Da | 1 | 4 |
| + | 4.004381 | 2.253804 | P17677 | P17677 | GAP43 | Neuromoc | Color2 [Da | 1 | 4 |
| + | 5.430731 | -1.2695 | P17812 | P17812 | CTPS1 | CTP syntha | Color2 [Da | 1 | 4 |
| + | 3.107659 | -1.24801 | P17844 | P17844 | DDX5 | Probable A | Color2 [Da | 1 | 4 |
| + | 2.785261 | -0.95499 | P17858 | P17858 | PFKL | ATP-deper | Color2 [Da | 1 | 4 |
| + | 2.539507 | 0.813562 | P17900 | P17900 | GM2A | Gangliosid | Color2 [Da | 1 | 4 |
| + | 1.667217 | -1.97151 | P17931 | P17931 | LGALS3 | Galectin-3 | Color2 [Da | 1 | 4 |
| | 0.103621 | -0.12911 | P17936 | P17936 | IGFBP3 | Insulin-like | Color2 [Da | 1 | 4 |
| + | 3.399335 | 0.58877 | P17980 | P17980 | PSMC3 | 26S protea | Color2 [Da | 1 | 4 |
| + | 1.492055 | -0.46912 | P17987 | P17987 | TCP1 | T-complex | Color2 [Da | 1 | 4 |
| + | 2.358027 | -1.35226 | P18031 | P18031 | PTPN1 | Tyrosine-p | Color2 [Da | 1 | 4 |
| | 0.596542 | 0.401177 | P18065 | P18065 | IGFBP2 | Insulin-like | Color2 [Da | 1 | 4 |
| | 0 | NaN | P18074 | P18074 | ERCC2 | General tr | Color2 [Da | 1 | 4 |
| + | 1.730701 | -1.06543 | P18075 | P18075 | BMP7 | Bone morp | Color2 [Da | 1 | 4 |
| + | 1.965262 | 1.50224 | P18077 | P18077 | RPL35A | 60S riboso | Color2 [Da | 1 | 4 |
| + | 2.736688 | -0.48755 | P18085 | P18085 | ARF4 | ADP-ribosy | Color2 [Da | 1 | 4 |
| + | 1.662923 | -0.61901 | P18124 | P18124 | RPL7 | 60S riboso | Color2 [Da | 1 | 4 |
| + | 2.078494 | -0.29467 | P18206 | P18206 | VCL | Vinculin | Color2 [Da | 1 | 4 |
| + | 3.251853 | -0.77408 | P18433 | P18433 | PTPRA | Receptor-t | Color2 [Da | 1 | 4 |
| + | 1.601566 | -2.15885 | P18440 | P18440 | NAT1 | Arylamine | Color2 [Da | 1 | 4 |
| | 0 | 0.34401 | P18507 | P18507 | GABRG2 | Gamma-ar | Color2 [Da | 1 | 4 |
| | 0.212694 | 0.046135 | P18583 | P18583 | SON | Protein SO | Color2 [Da | 1 | 4 |
| + | 1.752106 | 0.464012 | P18615 | P18615 | NELFE | Negative e | Color2 [Da | 1 | 4 |
| | 0.253286 | -0.08373 | P18621 | P18621 | RPL17 | 60S riboso | Color2 [Da | 1 | 4 |
| + | 1.389132 | 0.507563 | P18669 | P18669 | PGAM1 | Phosphogl | Color2 [Da | 1 | 4 |
| | 1.02381 | -0.2201 | P18754 | P18754 | RCC1 | Regulator | Color2 [Da | 1 | 4 |
| + | 1.554441 | 1.781912 | P18827 | P18827 | SDC1 | Syndecan- | Color2 [Da | 1 | 4 |

| | | | | | | | | | |
|---|---|---|---|---|---|---|---|---|---|
| | 0.428899 | 0.401712 | P18858 | P18858 | LIG1 | DNA ligase | Color2 [Da | 1 | 4 |
| + | 3.616416 | 4.864026 | P18859 | P18859 | ATP5PF | ATP syntha | Color2 [Da | 1 | 4 |
| + | 1.01432 | 0.449515 | P18887 | P18887 | XRCC1 | DNA repai | Color2 [Da | 1 | 4 |
| + | 1.740118 | 2.216426 | P19012 | P19012 | KRT15 | Keratin, ty | Color2 [Da | 1 | 4 |
| | 0 | 0.500836 | P19013 | P19013 | KRT4 | Keratin, ty | Color2 [Da | 1 | 4 |
| | 0.37759 | 0.101955 | P19021 | P19021 | PAM | Peptidyl-gl | Color2 [Da | 1 | 4 |
| + | 3.249468 | 0.82534 | P19022 | P19022 | CDH2 | Cadherin-2 | Color2 [Da | 1 | 4 |
| | 1.027151 | -0.15025 | P19086 | P19086 | GNAZ | Guanine n | Color2 [Da | 1 | 4 |
| + | 1.934182 | -1.1218 | P19174 | P19174 | PLCG1 | 1-phospha | Color2 [Da | 1 | 4 |
| | 0 | NaN | P19320 | P19320 | VCAM1 | Vascular c | Color2 [Da | 1 | 4 |
| + | 1.106549 | 0.474257 | P19338 | P19338 | NCL | Nucleolin | Color2 [Da | 1 | 4 |
| + | 5.521219 | -0.63608 | P19367 | P19367 | HK1 | Hexokinas | Color2 [Da | 1 | 4 |
| | 0.817291 | 0.383843 | P19387 | P19387 | POLR2C | DNA-direc | Color2 [Da | 1 | 4 |
| + | 1.77359 | 1.01151 | P19388 | P19388 | POLR2E | DNA-direc | Color2 [Da | 1 | 4 |
| + | 2.978872 | 1.95123 | P19404 | P19404 | NDUFV2 | NADH deh | Color2 [Da | 1 | 4 |
| | 0 | -2.35397 | P19447 | P19447 | ERCC3 | General tr | Color2 [Da | 1 | 4 |
| + | 2.511879 | -0.41938 | P19525 | P19525 | EIF2AK2 | Interferon | Color2 [Da | 1 | 4 |
| | 0.471868 | -0.45374 | P19532 | P19532 | TFE3 | Transcripti | Color2 [Da | 1 | 4 |
| + | 0.878446 | -0.3864 | P19623 | P19623 | SRM | Spermidin | Color2 [Da | 1 | 4 |
| + | 0.938106 | 0.808041 | P19634 | P19634 | SLC9A1 | Sodium/hy | Color2 [Da | 1 | 4 |
| + | 2.337825 | -0.96467 | P19784 | P19784 | CSNK2A2 | Casein kin | Color2 [Da | 1 | 4 |
| + | 1.181639 | 1.570681 | P19823 | P19823 | ITIH2 | Inter-alpha | Color2 [Da | 1 | 4 |
| + | 0.924747 | -1.14822 | P19838 | P19838 | NFKB1 | Nuclear fa | Color2 [Da | 1 | 4 |
| + | 3.407845 | -1.03623 | P20020 | P20020 | ATP2B1 | Plasma me | Color2 [Da | 1 | 4 |
| + | 2.894107 | 0.648578 | P20042 | P20042 | EIF2S2 | Eukaryotic | Color2 [Da | 1 | 4 |
| | 0 | NaN | P20061 | P20061 | TCN1 | Transcoba | Color2 [Da | 1 | 4 |
| + | 2.880918 | 0.61477 | P20073 | P20073 | ANXA7 | Annexin A | Color2 [Da | 1 | 4 |
| + | 0.989836 | -0.42002 | P20226 | P20226 | TBP | TATA-box- | Color2 [Da | 1 | 4 |
| + | 1.737735 | 0.714054 | P20264 | P20264 | POU3F3 | POU doma | Color2 [Da | 1 | 4 |
| | 0.703768 | 0.195741 | P20265 | P20265 | POU3F2 | POU doma | Color2 [Da | 1 | 4 |
| + | 1.862649 | 1.134967 | P20290 | P20290 | BTF3 | Transcripti | Color2 [Da | 1 | 4 |
| | 0.801042 | -0.2537 | P20336 | P20336 | RAB3A | Ras-relate | Color2 [Da | 1 | 4 |
| + | 2.001923 | -1.01121 | P20337 | P20337 | RAB3B | Ras-relate | Color2 [Da | 1 | 4 |
| | 0.331439 | 0.133318 | P20338 | P20338 | RAB4A | Ras-relate | Color2 [Da | 1 | 4 |
| + | 2.424944 | 0.849123 | P20339 | P20339 | RAB5A | Ras-relate | Color2 [Da | 1 | 4 |
| + | 1.774279 | 0.914264 | P20340 | P20340 | RAB6A | Ras-relate | Color2 [Da | 1 | 4 |
| | 0 | -0.71463 | P20366 | P20366 | TAC1 | Protachyki | Color2 [Da | 1 | 4 |
| + | 4.40206 | -3.3622 | P20396 | P20396 | TRH | Pro-thyrot | Color2 [Da | 1 | 4 |
| + | 2.132057 | 1.201096 | P20618 | P20618 | PSMB1 | Proteasom | Color2 [Da | 1 | 4 |
| + | 2.186109 | -0.58929 | P20645 | P20645 | M6PR | Cation-dep | Color2 [Da | 1 | 4 |
| + | 1.482047 | 1.62022 | P20674 | P20674 | COX5A | Cytochrom | Color2 [Da | 1 | 4 |
| + | 1.853798 | -0.26099 | P20700 | P20700 | LMNB1 | Lamin-B1 | Color2 [Da | 1 | 4 |
| + | 3.042102 | 3.359442 | P20810 | P20810 | CAST | Calpastatir | Color2 [Da | 1 | 4 |
| + | 1.077707 | -0.6417 | P20839 | P20839 | IMPDH1 | Inosine-5'- | Color2 [Da | 1 | 4 |
| | 0.769958 | 0.218694 | P20930 | P20930 | FLG | Filaggrin | Color2 [Da | 1 | 4 |
| + | 1.731569 | 3.611685 | P20933 | P20933 | AGA | N(4)-(beta | Color2 [Da | 1 | 4 |
| | 0.230025 | -0.18081 | P20936 | P20936 | RASA1 | Ras GTPas | Color2 [Da | 1 | 4 |

| | | | | | | | | | |
|---|---|---|---|---|---|---|---|---|---|
| + | 2.9434 | 2.535832 | P20962 | P20962 | PTMS | Parathymo | Color2 | [Da | 1 | 4 |
| | 1.12814 | -0.24619 | P21127;Q9 | P21127;Q9 | CDK11B;CI | Cyclin-dep | Color2 | [Da | 1 | 4 |
| + | 0.961872 | 0.692978 | P21246 | P21246 | PTN | Pleiotroph | Color2 | [Da | 1 | 4 |
| + | 2.893037 | -1.2353 | P21266 | P21266 | GSTM3 | Glutathion | Color2 | [Da | 1 | 4 |
| | 1.492395 | 0.07238 | P21281 | P21281 | ATP6V1B2 | V-type pro | Color2 | [Da | 1 | 4 |
| | 0.822794 | 0.254568 | P21283 | P21283 | ATP6V1C1 | V-type pro | Color2 | [Da | 1 | 4 |
| | 0.720496 | 0.32954 | P21291 | P21291 | CSRP1 | Cysteine a | Color2 | [Da | 1 | 4 |
| + | 4.002788 | -0.87947 | P21333 | P21333 | FLNA | Filamin-A | Color2 | [Da | 1 | 4 |
| + | 1.867101 | -0.68726 | P21359 | P21359 | NF1 | Neurofibro | Color2 | [Da | 1 | 4 |
| | 0.687515 | 0.3846 | P21397 | P21397 | MAOA | Amine oxi | Color2 | [Da | 1 | 4 |
| + | 2.224725 | -0.51987 | P21399 | P21399 | ACO1 | Cytoplasm | Color2 | [Da | 1 | 4 |
| + | 2.866342 | 1.548947 | P21579 | P21579 | SYT1 | Synaptota | Color2 | [Da | 1 | 4 |
| + | 2.435158 | 1.632521 | P21583 | P21583 | KITLG | Kit ligand | Color2 | [Da | 1 | 4 |
| + | 1.891294 | -0.54754 | P21741 | P21741 | MDK | Midkine | Color2 | [Da | 1 | 4 |
| + | 0.861781 | 0.448013 | P21796 | P21796 | VDAC1 | Voltage-de | Color2 | [Da | 1 | 4 |
| | 0 | NaN | P21802;P2 | P21802;P2 | FGFR2;FGF | Fibroblast | Color2 | [Da | 1 | 4 |
| + | 2.13249 | -1.77123 | P21810 | P21810 | BGN | Biglycan | Color2 | [Da | 1 | 4 |
| + | 2.312439 | -1.05563 | P21912 | P21912 | SDHB | Succinate | Color2 | [Da | 1 | 4 |
| | 0 | -0.34349 | P21926 | P21926 | CD9 | CD9 antige | Color2 | [Da | 1 | 4 |
| | 0 | -0.88117 | P21953 | P21953 | BCKDHB | 2-oxoisova | Color2 | [Da | 1 | 4 |
| + | 2.098473 | -0.39964 | P21964 | P21964 | COMT | Catechol O | Color2 | [Da | 1 | 4 |
| + | 2.218876 | -1.25944 | P22033 | P22033 | MMUT | Methylma | Color2 | [Da | 1 | 4 |
| | 1.248787 | -0.22669 | P22059 | P22059 | OSBP | Oxysterol- | Color2 | [Da | 1 | 4 |
| + | 3.164506 | 1.908431 | P22061 | P22061 | PCMT1 | Protein-L-i | Color2 | [Da | 1 | 4 |
| + | 1.07795 | 0.424008 | P22087 | P22087 | FBL | rRNA 2'-O- | Color2 | [Da | 1 | 4 |
| + | 2.928899 | -0.75538 | P22102 | P22102 | GART | Trifunctior | Color2 | [Da | 1 | 4 |
| + | 2.725056 | -1.18893 | P22234 | P22234 | PAICS | Bifunction | Color2 | [Da | 1 | 4 |
| + | 3.749035 | 1.42777 | P22304 | P22304 | IDS | Iduronate | Color2 | [Da | 1 | 4 |
| + | 3.653806 | 0.926802 | P22307 | P22307 | SCP2 | Sterol carr | Color2 | [Da | 1 | 4 |
| + | 3.868759 | -1.07682 | P22314 | P22314 | UBA1 | Ubiquitin-l | Color2 | [Da | 1 | 4 |
| + | 3.680301 | 1.081206 | P22392 | P22392 | NME2 | Nucleoside | Color2 | [Da | 1 | 4 |
| + | 1.94832 | 1.513305 | P22415 | P22415 | USF1 | Upstream | Color2 | [Da | 1 | 4 |
| + | 1.617118 | 1.134904 | P22492 | P22492 | H1-6 | Histone H1 | Color2 | [Da | 1 | 4 |
| + | 1.793996 | 0.46019 | P22570 | P22570 | FDXR | NADPH:ad | Color2 | [Da | 1 | 4 |
| + | 3.172363 | 0.489877 | P22626 | P22626 | HNRNPA2I | Heterogen | Color2 | [Da | 1 | 4 |
| | 0.335418 | 0.171976 | P22670 | P22670 | RFX1 | MHC class | Color2 | [Da | 1 | 4 |
| + | 2.134981 | 0.873678 | P22676 | P22676 | CALB2 | Calretinin | Color2 | [Da | 1 | 4 |
| | 0.08901 | 0.035805 | P22681 | P22681 | CBL | E3 ubiquiti | Color2 | [Da | 1 | 4 |
| + | 3.442828 | -1.01916 | P22694 | P22694 | PRKACB | cAMP-dep | Color2 | [Da | 1 | 4 |
| + | 1.394437 | -0.24376 | P22695 | P22695 | UQCRC2 | Cytochron | Color2 | [Da | 1 | 4 |
| + | 1.955483 | 1.199701 | P22735 | P22735 | TGM1 | Protein-glu | Color2 | [Da | 1 | 4 |
| | 0.448709 | 0.302757 | P22830 | P22830 | FECH | Ferrochela | Color2 | [Da | 1 | 4 |
| | 0.02612 | 0.031566 | P23025 | P23025 | XPA | DNA repair | Color2 | [Da | 1 | 4 |
| | 0 | -1.19278 | P23142 | P23142 | FBLN1 | Fibulin-1 | Color2 | [Da | 1 | 4 |
| + | 1.860177 | 0.813423 | P23193 | P23193 | TCEA1 | Transcripti | Color2 | [Da | 1 | 4 |
| + | 2.422789 | -1.34119 | P23229 | P23229 | ITGA6 | Integrin al | Color2 | [Da | 1 | 4 |
| + | 2.756789 | 0.345641 | P23246 | P23246 | SFPQ | Splicing fa | Color2 | [Da | 1 | 4 |

| | | | | | | | | | |
|---|---|---|---|---|---|---|---|---|---|
| + | 0.909348 | -0.55383 | P23258 | P23258 | TUBG1 | Tubulin ga | Color2 [Da | 1 | 4 |
| + | 1.205373 | 0.634866 | P23284 | P23284 | PPIB | Peptidyl-pı | Color2 [Da | 1 | 4 |
| | 0 | NaN | P23297 | P23297 | S100A1 | Protein S1( | Color2 [Da | 1 | 4 |
| | 0.764592 | 0.342931 | P23327 | P23327 | HRC | Sarcoplasn | Color2 [Da | 1 | 4 |
| | 0 | NaN | P23352 | P23352 | ANOS1 | Anosmin-1 | Color2 [Da | 1 | 4 |
| + | 3.203305 | -1.21643 | P23368 | P23368 | ME2 | NAD-depe | Color2 [Da | 1 | 4 |
| + | 3.678923 | -1.05444 | P23378 | P23378 | GLDC | Glycine de | Color2 [Da | 1 | 4 |
| + | 2.849519 | -0.80729 | P23381 | P23381 | WARS1 | Tryptopha | Color2 [Da | 1 | 4 |
| | 0.273167 | 0.07829 | P23396 | P23396 | RPS3 | 40S riboso | Color2 [Da | 1 | 4 |
| + | 2.575437 | 4.499806 | P23434 | P23434 | GCSH | Glycine cle | Color2 [Da | 1 | 4 |
| + | 1.209421 | -0.56764 | P23435 | P23435 | CBLN1 | Cerebellin· | Color2 [Da | 1 | 4 |
| | 0 | 0.867373 | P23443 | P23443 | RPS6KB1 | Ribosomal | Color2 [Da | 1 | 4 |
| + | 1.94451 | 0.627799 | P23468 | P23468 | PTPRD | Receptor-t | Color2 [Da | 1 | 4 |
| | 0 | -1.1267 | P23470 | P23470 | PTPRG | Receptor-t | Color2 [Da | 1 | 4 |
| + | 3.059695 | 1.020865 | P23471 | P23471 | PTPRZ1 | Receptor-t | Color2 [Da | 1 | 4 |
| | 0.065415 | 0.14204 | P23508 | P23508 | MCC | Colorectal | Color2 [Da | 1 | 4 |
| + | 1.801688 | 1.296694 | P23511 | P23511 | NFYA | Nuclear tra | Color2 [Da | 1 | 4 |
| + | 2.909326 | -0.91913 | P23526 | P23526 | AHCY | Adenosylh | Color2 [Da | 1 | 4 |
| + | 2.034698 | 0.64333 | P23528 | P23528 | CFL1 | Cofilin-1 | Color2 [Da | 1 | 4 |
| + | 5.311577 | 2.587238 | P23588 | P23588 | EIF4B | Eukaryotic | Color2 [Da | 1 | 4 |
| + | 1.697998 | -1.08615 | P23610 | P23610 | F8A3 | 40-kDa hu | Color2 [Da | 1 | 4 |
| + | 1.280647 | -0.47075 | P23634 | P23634 | ATP2B4 | Plasma me | Color2 [Da | 1 | 4 |
| | 0 | -1.25282 | P23743 | P23743 | DGKA | Diacylglyce | Color2 [Da | 1 | 4 |
| + | 0.86762 | 1.091957 | P23763 | P23763 | VAMP1 | Vesicle-ass | Color2 [Da | 1 | 4 |
| + | 2.009906 | -1.59235 | P23786 | P23786 | CPT2 | Carnitine ( | Color2 [Da | 1 | 4 |
| + | 2.39159 | -1.20336 | P23919 | P23919 | DTYMK | Thymidyla | Color2 [Da | 1 | 4 |
| + | 2.778109 | -0.82113 | P23921 | P23921 | RRM1 | Ribonucle( | Color2 [Da | 1 | 4 |
| | 0 | -2.69833 | P24158 | P24158 | PRTN3 | Myeloblas | Color2 [Da | 1 | 4 |
| | 0 | NaN | P24390 | P24390 | KDELR1 | ER lumen ( | Color2 [Da | 1 | 4 |
| | 0.923812 | 0.187805 | P24534 | P24534 | EEF1B2 | Elongation | Color2 [Da | 1 | 4 |
| + | 2.899618 | 0.891067 | P24539 | P24539 | ATP5PB | ATP syntha | Color2 [Da | 1 | 4 |
| + | 2.532637 | 2.659395 | P24588 | P24588 | AKAP5 | A-kinase a | Color2 [Da | 1 | 4 |
| + | 1.98198 | 1.872845 | P24593 | P24593 | IGFBP5 | Insulin-like | Color2 [Da | 1 | 4 |
| | 0.039308 | -0.01403 | P24666 | P24666 | ACP1 | Low molec | Color2 [Da | 1 | 4 |
| + | 1.927726 | 0.243289 | P24752 | P24752 | ACAT1 | Acetyl-CoA | Color2 [Da | 1 | 4 |
| + | 6.502883 | 2.198115 | P24821 | P24821 | TNC | Tenascin | Color2 [Da | 1 | 4 |
| | 0.076705 | 0.176607 | P24844 | P24844 | MYL9 | Myosin re( | Color2 [Da | 1 | 4 |
| + | 1.76581 | -0.23528 | P24928 | P24928 | POLR2A | DNA-direc | Color2 [Da | 1 | 4 |
| + | 2.632831 | -1.29471 | P24941 | P24941 | CDK2 | Cyclin-dep | Color2 [Da | 1 | 4 |
| + | 0.888563 | 0.396418 | P25054 | P25054 | APC | Adenomat | Color2 [Da | 1 | 4 |
| | 0.614821 | -0.23622 | P25098 | P25098 | GRK2 | Beta-adrer | Color2 [Da | 1 | 4 |
| + | 2.694401 | -0.75931 | P25205 | P25205 | MCM3 | DNA replic | Color2 [Da | 1 | 4 |
| | 0.509074 | 0.915931 | P25208 | P25208 | NFYB | Nuclear tra | Color2 [Da | 1 | 4 |
| + | 2.351221 | 1.029623 | P25311 | P25311 | AZGP1 | Zinc-alpha | Color2 [Da | 1 | 4 |
| + | 1.16743 | -0.83091 | P25325 | P25325 | MPST | 3-mercapt | Color2 [Da | 1 | 4 |
| | 0 | -1.45993 | P25391 | P25391 | LAMA1 | Laminin su | Color2 [Da | 1 | 4 |
| | 0.08167 | -0.13091 | P25398 | P25398 | RPS12 | 40S riboso | Color2 [Da | 1 | 4 |

| | | | | | | | | | |
|---|---|---|---|---|---|---|---|---|---|
| | 0.753738 | 0.335758 | P25440 | P25440 | BRD2 | Bromodon | Color2 [Da | 1 | 4 |
| | 0 | NaN | P25445 | P25445 | FAS | Tumor nec | Color2 [Da | 1 | 4 |
| + | 2.215953 | 0.890706 | P25490 | P25490 | YY1 | Transcripti | Color2 [Da | 1 | 4 |
| | 2.028379 | 0.149312 | P25685 | P25685 | DNAJB1 | DnaJ homo | Color2 [Da | 1 | 4 |
| + | 2.766037 | 1.593949 | P25686 | P25686 | DNAJB2 | DnaJ homo | Color2 [Da | 1 | 4 |
| + | 2.442993 | 1.048978 | P25705 | P25705 | ATP5F1A | ATP syntha | Color2 [Da | 1 | 4 |
| + | 3.798388 | 1.698247 | P25786 | P25786 | PSMA1 | Proteasom | Color2 [Da | 1 | 4 |
| + | 2.766635 | 2.10116 | P25787 | P25787 | PSMA2 | Proteasom | Color2 [Da | 1 | 4 |
| + | 2.059786 | 1.509289 | P25788 | P25788 | PSMA3 | Proteasom | Color2 [Da | 1 | 4 |
| + | 2.381197 | 1.017969 | P25789 | P25789 | PSMA4 | Proteasom | Color2 [Da | 1 | 4 |
| | 0 | NaN | P25815 | P25815 | S100P | Protein S1 | Color2 [Da | 1 | 4 |
| | 0 | -0.88941 | P26006 | P26006 | ITGA3 | Integrin al | Color2 [Da | 1 | 4 |
| + | 1.815789 | -1.37741 | P26012 | P26012 | ITGB8 | Integrin be | Color2 [Da | 1 | 4 |
| | 0 | NaN | P26022 | P26022 | PTX3 | Pentraxin- | Color2 [Da | 1 | 4 |
| + | 3.158181 | 0.599747 | P26038 | P26038 | MSN | Moesin | Color2 [Da | 1 | 4 |
| + | 3.690999 | -0.86005 | P26196 | P26196 | DDX6 | Probable A | Color2 [Da | 1 | 4 |
| + | 1.737502 | -0.5454 | P26232 | P26232 | CTNNA2 | Catenin al | Color2 [Da | 1 | 4 |
| + | 1.972935 | -0.8539 | P26358 | P26358 | DNMT1 | DNA (cytos | Color2 [Da | 1 | 4 |
| + | 1.75391 | 0.703313 | P26368 | P26368 | U2AF2 | Splicing fa | Color2 [Da | 1 | 4 |
| | 0.16208 | 0.107666 | P26373 | P26373 | RPL13 | 60S riboso | Color2 [Da | 1 | 4 |
| | 0 | -0.55308 | P26374 | P26374 | CHML | Rab protei | Color2 [Da | 1 | 4 |
| + | 3.598532 | 1.096615 | P26378 | P26378 | ELAVL4 | ELAV-like p | Color2 [Da | 1 | 4 |
| | 0.896567 | 0.333563 | P26440 | P26440 | IVD | Isovaleryl- | Color2 [Da | 1 | 4 |
| + | 1.76827 | -0.616 | P26583 | P26583 | HMGB2 | High mobil | Color2 [Da | 1 | 4 |
| | 0.76643 | 0.221097 | P26599 | P26599 | PTBP1 | Polypyrimi | Color2 [Da | 1 | 4 |
| + | 3.257322 | -1.46469 | P26639 | P26639 | TARS1 | Threonine- | Color2 [Da | 1 | 4 |
| + | 4.499559 | -1.21099 | P26640 | P26640 | VARS1 | Valine--tRI | Color2 [Da | 1 | 4 |
| + | 2.745247 | -1.06105 | P26641 | P26641 | EEF1G | Elongation | Color2 [Da | 1 | 4 |
| + | 2.285064 | 0.942157 | P26885 | P26885 | FKBP2 | Peptidyl-p | Color2 [Da | 1 | 4 |
| + | 1.71088 | -0.28839 | P26992 | P26992 | CNTFR | Ciliary neu | Color2 [Da | 1 | 4 |
| | 0 | NaN | P27037;Q1 | P27037;Q1 | ACVR2A;A | Activin rec | Color2 [Da | 1 | 4 |
| | 0.594086 | -1.15579 | P27105 | P27105 | STOM | Stomatin | Color2 [Da | 1 | 4 |
| + | 1.449785 | 1.065343 | P27144 | P27144 | AK4 | Adenylate | Color2 [Da | 1 | 4 |
| | 0.092295 | 0.070977 | P27338 | P27338 | MAOB | Amine oxi | Color2 [Da | 1 | 4 |
| | 0.273814 | -0.13922 | P27348 | P27348 | YWHAQ | 14-3-3 pro | Color2 [Da | 1 | 4 |
| + | 1.843177 | -0.69563 | P27361 | P27361 | MAPK3 | Mitogen-a | Color2 [Da | 1 | 4 |
| + | 1.620087 | 0.3102 | P27448 | P27448 | MARK3 | MAP/micr | Color2 [Da | 1 | 4 |
| + | 1.510699 | -1.76302 | P27449 | P27449 | ATP6V0C | V-type pro | Color2 [Da | 1 | 4 |
| + | 2.274249 | 0.4826 | P27482 | P27482 | CALML3 | Calmoduli | Color2 [Da | 1 | 4 |
| | 0.593034 | -0.40388 | P27544 | P27544 | CERS1 | Ceramide | Color2 [Da | 1 | 4 |
| + | 3.672018 | -2.19167 | P27635 | P27635 | RPL10 | 60S riboso | Color2 [Da | 1 | 4 |
| | 0 | NaN | P27658 | P27658 | COL8A1 | Collagen a | Color2 [Da | 1 | 4 |
| + | 2.730859 | -0.54357 | P27694 | P27694 | RPA1 | Replicatio | Color2 [Da | 1 | 4 |
| | 0.139036 | 0.101374 | P27695 | P27695 | APEX1 | DNA-(apur | Color2 [Da | 1 | 4 |
| | 0 | NaN | P27701 | P27701 | CD82 | CD82 antig | Color2 [Da | 1 | 4 |
| + | 1.366101 | -1.93881 | P27707 | P27707 | DCK | Deoxycytid | Color2 [Da | 1 | 4 |
| + | 2.557258 | -1.58455 | P27708 | P27708 | CAD | CAD prote | Color2 [Da | 1 | 4 |

| | | | | | | | | | |
|---|---|---|---|---|---|---|---|---|---|
| | 0.593926 | 0.128711 | P27797 | P27797 | CALR | Calreticulin | Color2 [Da | 1 | 4 |
| + | 4.689113 | 3.012419 | P27816 | P27816 | MAP4 | Microtubu | Color2 [Da | 1 | 4 |
| + | 4.119435 | -1.24394 | P27824 | P27824 | CANX | Calnexin | Color2 [Da | 1 | 4 |
| + | 2.169419 | 0.61168 | P27986 | P27986 | PIK3R1 | Phosphatid | Color2 [Da | 1 | 4 |
| + | 2.367938 | -0.89444 | P27987 | P27987 | ITPKB | Inositol-tri | Color2 [Da | 1 | 4 |
| + | 3.302469 | 2.021698 | P28066 | P28066 | PSMA5 | Proteasom | Color2 [Da | 1 | 4 |
| + | 3.662755 | 3.035748 | P28070 | P28070 | PSMB4 | Proteasom | Color2 [Da | 1 | 4 |
| + | 2.684804 | 2.192803 | P28072 | P28072 | PSMB6 | Proteasom | Color2 [Da | 1 | 4 |
| + | 2.472486 | 1.577563 | P28074 | P28074 | PSMB5 | Proteasom | Color2 [Da | 1 | 4 |
| | 1.049742 | -0.24255 | P28161 | P28161 | GSTM2 | Glutathion | Color2 [Da | 1 | 4 |
| + | 3.652506 | -1.02245 | P28288 | P28288 | ABCD3 | ATP-bindir | Color2 [Da | 1 | 4 |
| + | 1.114375 | 1.077906 | P28289 | P28289 | TMOD1 | Tropomod | Color2 [Da | 1 | 4 |
| | 0 | -0.87254 | P28290 | P28290 | ITPRID2 | Protein ITF | Color2 [Da | 1 | 4 |
| + | 2.257727 | -0.87937 | P28331 | P28331 | NDUFS1 | NADH-ubiq | Color2 [Da | 1 | 4 |
| | 0 | NaN | P28340 | P28340 | POLD1 | DNA polyn | Color2 [Da | 1 | 4 |
| + | 2.638971 | -1.23456 | P28370 | P28370 | SMARCA1 | Probable g | Color2 [Da | 1 | 4 |
| + | 1.364505 | 1.195887 | P28472 | P28472 | GABRB3 | Gamma-ar | Color2 [Da | 1 | 4 |
| + | 1.486029 | -0.42937 | P28482 | P28482 | MAPK1 | Mitogen-a | Color2 [Da | 1 | 4 |
| + | 1.237682 | 0.561982 | P28676 | P28676 | GCA | Grancalcin | Color2 [Da | 1 | 4 |
| | 0 | -0.71097 | P28702 | P28702 | RXRB | Retinoic ac | Color2 [Da | 1 | 4 |
| | 0.024561 | -0.02039 | P28715 | P28715 | ERCC5 | DNA excisi | Color2 [Da | 1 | 4 |
| | 0.279596 | 0.546097 | P28799 | P28799 | GRN | Progranuli | Color2 [Da | 1 | 4 |
| + | 2.863055 | -0.50867 | P28838 | P28838 | LAP3 | Cytosol am | Color2 [Da | 1 | 4 |
| | 0.606449 | -0.20477 | P29083 | P29083 | GTF2E1 | General tra | Color2 [Da | 1 | 4 |
| | 0.406172 | -0.18518 | P29084 | P29084 | GTF2E2 | Transcripti | Color2 [Da | 1 | 4 |
| + | 1.322925 | 0.412534 | P29120 | P29120 | PCSK1 | Neuroendo | Color2 [Da | 1 | 4 |
| + | 2.053077 | -0.97787 | P29144 | P29144 | TPP2 | Tripeptidyl | Color2 [Da | 1 | 4 |
| | 0.372014 | -0.09063 | P29218 | P29218 | IMPA1 | Inositol mo | Color2 [Da | 1 | 4 |
| + | 0.828898 | -0.52566 | P29279 | P29279 | CCN2 | CCN family | Color2 [Da | 1 | 4 |
| + | 1.850149 | -2.46452 | P29317 | P29317 | EPHA2 | Ephrin type | Color2 [Da | 1 | 4 |
| + | 0.986833 | -0.5538 | P29323 | P29323 | EPHB2 | Ephrin type | Color2 [Da | 1 | 4 |
| | 0 | -0.34596 | P29353 | P29353 | SHC1 | SHC-transf | Color2 [Da | 1 | 4 |
| + | 0.811804 | 0.706246 | P29372 | P29372 | MPG | DNA-3-me | Color2 [Da | 1 | 4 |
| + | 1.71629 | -0.51477 | P29373 | P29373 | CRABP2 | Cellular re | Color2 [Da | 1 | 4 |
| | 0.256409 | 0.680241 | P29375 | P29375 | KDM5A | Lysine-spe | Color2 [Da | 1 | 4 |
| | 0.148082 | 0.047798 | P29401 | P29401 | TKT | Transketol | Color2 [Da | 1 | 4 |
| + | 0.981885 | 1.087774 | P29475 | P29475 | NOS1 | Nitric oxid | Color2 [Da | 1 | 4 |
| + | 1.216938 | 0.365 | P29508 | P29508 | SERPINB3 | Serpin B3 | Color2 [Da | 1 | 4 |
| | 0.588891 | -0.68171 | P29558;Q6 | P29558;Q6 | RBMS1;RB | RNA-bindi | Color2 [Da | 1 | 4 |
| + | 0.75978 | -0.85972 | P29590 | P29590 | PML | Protein PM | Color2 [Da | 1 | 4 |
| + | 2.302836 | 0.65325 | P29692 | P29692 | EEF1D | Elongation | Color2 [Da | 1 | 4 |
| + | 1.687902 | 0.610804 | P29762 | P29762 | CRABP1 | Cellular re | Color2 [Da | 1 | 4 |
| + | 3.256998 | 1.078182 | P29966 | P29966 | MARCKS | Myristoyla | Color2 [Da | 1 | 4 |
| | 0.393456 | -0.05892 | P29992 | P29992 | GNA11 | Guanine nu | Color2 [Da | 1 | 4 |
| + | 2.010802 | 0.905477 | P30038 | P30038 | ALDH4A1 | Delta-1-py | Color2 [Da | 1 | 4 |
| + | 1.623384 | 1.177602 | P30039 | P30039 | PBLD | Phenazine | Color2 [Da | 1 | 4 |
| + | 2.809208 | 1.138516 | P30040 | P30040 | ERP29 | Endoplasm | Color2 [Da | 1 | 4 |

| | | | | | | | | | | |
|---|---|---|---|---|---|---|---|---|---|---|
| + | 3.973789 | 1.310592 | P30041 | P30041 | PRDX6 | Peroxiredc | Color2 [Da | | 1 | 4 |
| + | 1.118422 | 0.773946 | P30043 | P30043 | BLVRB | Flavin redu | Color2 [Da | | 1 | 4 |
| + | 2.3692 | 1.477749 | P30044 | P30044 | PRDX5 | Peroxiredc | Color2 [Da | | 1 | 4 |
| + | 3.523702 | 2.962065 | P30046 | P30046 | DDT | D-dopachr | Color2 [Da | | 1 | 4 |
| + | 1.798323 | 4.126657 | P30047 | P30047 | GCHFR | GTP cycloh | Color2 [Da | | 1 | 4 |
| + | 2.698145 | 1.897897 | P30048 | P30048 | PRDX3 | Thioredoxi | Color2 [Da | | 1 | 4 |
| + | 3.240445 | 1.229661 | P30049 | P30049 | ATP5F1D | ATP syntha | Color2 [Da | | 1 | 4 |
| | 0.427253 | 0.268091 | P30050 | P30050 | RPL12 | 60S riboso | Color2 [Da | | 1 | 4 |
| + | 1.7421 | 0.616025 | P30084 | P30084 | ECHS1 | Enoyl-CoA | Color2 [Da | | 1 | 4 |
| + | 2.284424 | 0.579915 | P30085 | P30085 | CMPK1 | UMP-CMP | Color2 [Da | | 1 | 4 |
| + | 2.889441 | 2.635274 | P30086 | P30086 | PEBP1 | Phosphatic | Color2 [Da | | 1 | 4 |
| | 0.351623 | -0.06378 | P30101 | P30101 | PDIA3 | Protein dis | Color2 [Da | | 1 | 4 |
| + | 3.096606 | -1.25393 | P30153 | P30153 | PPP2R1A | Serine/thr | Color2 [Da | | 1 | 4 |
| | 0 | -2.91724 | P30154 | P30154 | PPP2R1B | Serine/thr | Color2 [Da | | 1 | 4 |
| | 0.131679 | 0.418527 | P30260 | P30260 | CDC27 | Cell divisio | Color2 [Da | | 1 | 4 |
| + | 3.823224 | 2.131617 | P30405 | P30405 | PPIF | Peptidyl-p | Color2 [Da | | 1 | 4 |
| | 0.379918 | 0.340964 | P30414 | P30414 | NKTR | NK-tumor | Color2 [Da | | 1 | 4 |
| + | 0.915207 | -0.46246 | P30419 | P30419 | NMT1 | Glycylpept | Color2 [Da | | 1 | 4 |
| + | 1.802664 | 0.618254 | P30519 | P30519 | HMOX2 | Heme oxyg | Color2 [Da | | 1 | 4 |
| | 0.242558 | 0.053916 | P30520 | P30520 | ADSS2 | Adenylosu | Color2 [Da | | 1 | 4 |
| | 0 | NaN | P30531 | P30531 | SLC6A1 | Sodium- ar | Color2 [Da | | 1 | 4 |
| + | 2.292149 | 0.661684 | P30533 | P30533 | LRPAP1 | Alpha-2-m | Color2 [Da | | 1 | 4 |
| + | 1.238684 | -0.98127 | P30566 | P30566 | ADSL | Adenylosu | Color2 [Da | | 1 | 4 |
| + | 2.556947 | 0.790586 | P30622 | P30622 | CLIP1 | CAP-Gly do | Color2 [Da | | 1 | 4 |
| | 0.674728 | 0.346041 | P30626 | P30626 | SRI | Sorcin | Color2 [Da | | 1 | 4 |
| | 0 | NaN | P30711 | P30711 | GSTT1 | Glutathion | Color2 [Da | | 1 | 4 |
| + | 1.135075 | 0.745699 | P30740 | P30740 | SERPINB1 | Leukocyte | Color2 [Da | | 1 | 4 |
| | 0.33763 | -0.09933 | P30837 | P30837 | ALDH1B1 | Aldehyde ( | Color2 [Da | | 1 | 4 |
| + | 2.475479 | -0.90231 | P30876 | P30876 | POLR2B | DNA-direc | Color2 [Da | | 1 | 4 |
| | 0 | NaN | P30990 | P30990 | NTS | Neurotens | Color2 [Da | | 1 | 4 |
| | 0 | 1.672164 | P31025 | P31025 | LCN1 | Lipocalin-1 | Color2 [Da | | 1 | 4 |
| + | 1.322073 | -0.30476 | P31040 | P31040 | SDHA | Succinate ( | Color2 [Da | | 1 | 4 |
| | 0.429282 | 0.074152 | P31146 | P31146 | CORO1A | Coronin-1A | Color2 [Da | | 1 | 4 |
| + | 2.930844 | -0.59157 | P31150 | P31150 | GDI1 | Rab GDP d | Color2 [Da | | 1 | 4 |
| + | 1.948429 | 1.220383 | P31151 | P31151 | S100A7 | Protein S1 | Color2 [Da | | 1 | 4 |
| | 0 | NaN | P31152;Q1 | P31152;Q1 | MAPK4;MA | Mitogen-a | Color2 [Da | | 1 | 4 |
| | 0.59414 | -0.41766 | P31153 | P31153 | MAT2A | S-adenosy | Color2 [Da | | 1 | 4 |
| + | 1.855189 | 0.501856 | P31323 | P31323 | PRKAR2B | cAMP-dep | Color2 [Da | | 1 | 4 |
| + | 1.07373 | -0.65512 | P31327 | P31327 | CPS1 | Carbamoyl | Color2 [Da | | 1 | 4 |
| + | 1.388556 | -0.56508 | P31350 | P31350 | RRM2 | Ribonucle | Color2 [Da | | 1 | 4 |
| | 0.115447 | 0.06397 | P31483 | P31483 | TIA1 | Cytotoxic g | Color2 [Da | | 1 | 4 |
| + | 2.213534 | -0.92734 | P31689 | P31689 | DNAJA1 | DnaJ homo | Color2 [Da | | 1 | 4 |
| + | 2.226583 | -1.37278 | P31749 | P31749 | AKT1 | RAC-alpha | Color2 [Da | | 1 | 4 |
| | 0 | -0.81934 | P31751 | P31751 | AKT2 | RAC-beta s | Color2 [Da | | 1 | 4 |
| + | 2.988242 | -1.22993 | P31930 | P31930 | UQCRC1 | Cytochrom | Color2 [Da | | 1 | 4 |
| | 0.54752 | -0.2871 | P31937 | P31937 | HIBADH | 3-hydroxyi | Color2 [Da | | 1 | 4 |
| + | 3.517387 | -1.11806 | P31939 | P31939 | ATIC | Bifunction | Color2 [Da | | 1 | 4 |

| | | | | | | | | | | |
|---|---|---|---|---|---|---|---|---|---|---|
| + | 3.296511 | 1.118568 | P31942 | P31942 | HNRNPH3 | Heterogen | Color2 [Da | | 1 | 4 |
| + | 2.096515 | 0.413778 | P31943 | P31943 | HNRNPH1 | Heterogen | Color2 [Da | | 1 | 4 |
| + | 2.496109 | 1.13652 | P31944 | P31944 | CASP14 | Caspase-1 | Color2 [Da | | 1 | 4 |
| | 0.714011 | 0.57425 | P31946 | P31946 | YWHAB | 14-3-3 pro | Color2 [Da | | 1 | 4 |
| | 0.742713 | 0.390073 | P31947 | P31947 | SFN | 14-3-3 pro | Color2 [Da | | 1 | 4 |
| | 0.522577 | -0.03534 | P31948 | P31948 | STIP1 | Stress-indu | Color2 [Da | | 1 | 4 |
| + | 2.618818 | 1.11777 | P31949 | P31949 | S100A11 | Protein S1 | Color2 [Da | | 1 | 4 |
| + | 1.201878 | -0.41619 | P32004 | P32004 | L1CAM | Neural cell | Color2 [Da | | 1 | 4 |
| + | 3.821593 | 1.027636 | P32119 | P32119 | PRDX2 | Peroxiredo | Color2 [Da | | 1 | 4 |
| | 0.274395 | 0.052572 | P32121 | P32121 | ARRB2 | Beta-arres | Color2 [Da | | 1 | 4 |
| + | 2.504237 | -1.05451 | P32189 | P32189 | GK | Glycerol ki | Color2 [Da | | 1 | 4 |
| | 0 | -0.68094 | P32242 | P32242 | OTX1 | Homeobox | Color2 [Da | | 1 | 4 |
| | 0 | NaN | P32243 | P32243 | OTX2 | Homeobox | Color2 [Da | | 1 | 4 |
| + | 2.778388 | -1.38452 | P32321 | P32321 | DCTD | Deoxycytic | Color2 [Da | | 1 | 4 |
| + | 1.306711 | -0.55087 | P32322 | P32322 | PYCR1 | Pyrroline-5 | Color2 [Da | | 1 | 4 |
| + | 3.274972 | 1.715917 | P32418 | P32418 | SLC8A1 | Sodium/ca | Color2 [Da | | 1 | 4 |
| + | 0.980623 | 0.403982 | P32780 | P32780 | GTF2H1 | General tr | Color2 [Da | | 1 | 4 |
| | 0.142326 | 0.146313 | P32856 | P32856 | STX2 | Syntaxin-2 | Color2 [Da | | 1 | 4 |
| + | 2.64251 | 1.478904 | P32969 | P32969 | RPL9P9 | 60S riboso | Color2 [Da | | 1 | 4 |
| | 0.527657 | -0.18165 | P33121 | P33121 | ACSL1 | Long-chain | Color2 [Da | | 1 | 4 |
| + | 3.212609 | -1.21839 | P33176 | P33176 | KIF5B | Kinesin-1 h | Color2 [Da | | 1 | 4 |
| + | 2.723759 | 1.367181 | P33240 | P33240 | CSTF2 | Cleavage s | Color2 [Da | | 1 | 4 |
| | 0 | NaN | P33241 | P33241 | LSP1 | Lymphocyt | Color2 [Da | | 1 | 4 |
| + | 2.598772 | 2.136689 | P33316 | P33316 | DUT | Deoxyurid | Color2 [Da | | 1 | 4 |
| | 0 | NaN | P33402 | P33402 | GUCY1A2 | Guanylate | Color2 [Da | | 1 | 4 |
| + | 1.842929 | -0.81221 | P33527 | P33527 | ABCC1 | Multidrug | Color2 [Da | | 1 | 4 |
| + | 3.235629 | -0.98911 | P33552 | P33552 | CKS2 | Cyclin-dep | Color2 [Da | | 1 | 4 |
| | 0 | NaN | P33897 | P33897 | ABCD1 | ATP-bindir | Color2 [Da | | 1 | 4 |
| | 0 | NaN | P33947 | P33947 | KDELR2 | ER lumen p | Color2 [Da | | 1 | 4 |
| | 0 | 0.188497 | P33981 | P33981 | TTK | Dual speci | Color2 [Da | | 1 | 4 |
| + | 2.971408 | -0.87955 | P33991 | P33991 | MCM4 | DNA replic | Color2 [Da | | 1 | 4 |
| + | 5.100448 | -1.64796 | P33992 | P33992 | MCM5 | DNA replic | Color2 [Da | | 1 | 4 |
| + | 5.454736 | -1.67777 | P33993 | P33993 | MCM7 | DNA replic | Color2 [Da | | 1 | 4 |
| + | 4.336868 | 2.155327 | P34059 | P34059 | GALNS | N-acetylga | Color2 [Da | | 1 | 4 |
| + | 2.271225 | -0.91995 | P34741 | P34741 | SDC2 | Syndecan- | Color2 [Da | | 1 | 4 |
| + | 2.37931 | -1.28464 | P34897 | P34897 | SHMT2 | Serine hyd | Color2 [Da | | 1 | 4 |
| + | 1.859767 | 0.585266 | P34903 | P34903 | GABRA3 | Gamma-ar | Color2 [Da | | 1 | 4 |
| + | 1.945585 | -1.74786 | P34913 | P34913 | EPHX2 | Bifunction | Color2 [Da | | 1 | 4 |
| | 0 | NaN | P34931 | P34931 | HSPA1L | Heat shock | Color2 [Da | | 1 | 4 |
| + | 3.525804 | -0.48535 | P34932 | P34932 | HSPA4 | Heat shock | Color2 [Da | | 1 | 4 |
| | 0.188515 | -0.13553 | P34947 | P34947 | GRK5 | G protein- | Color2 [Da | | 1 | 4 |
| + | 1.841555 | -0.67541 | P34949 | P34949 | MPI | Mannose-( | Color2 [Da | | 1 | 4 |
| | 0 | 0.441864 | P35030 | P35030 | PRSS3 | Trypsin-3 | Color2 [Da | | 1 | 4 |
| | 0.004374 | -0.00267 | P35052 | P35052 | GPC1 | Glypican-1 | Color2 [Da | | 1 | 4 |
| + | 2.973215 | 1.00371 | P35080 | P35080 | PFN2 | Profilin-2 | Color2 [Da | | 1 | 4 |
| + | 1.922285 | -0.53571 | P35221 | P35221 | CTNNA1 | Catenin alp | Color2 [Da | | 1 | 4 |
| + | 2.660846 | -1.13098 | P35222 | P35222 | CTNNB1 | Catenin be | Color2 [Da | | 1 | 4 |

| | | | | | | | | | |
|---|---|---|---|---|---|---|---|---|---|
| | 0 | NaN | P35226 | P35226 | BMI1 | Polycomb | Color2 [Da | 1 | 4 |
| + | 4.314563 | 0.971371 | P35232 | P35232 | PHB1 | Prohibitin | Color2 [Da | 1 | 4 |
| + | 2.042866 | 3.0843 | P35237 | P35237 | SERPINB6 | Serpin B6 | Color2 [Da | 1 | 4 |
| + | 1.661643 | -0.68439 | P35240 | P35240 | NF2 | Merlin | Color2 [Da | 1 | 4 |
| + | 3.248432 | 1.099736 | P35241 | P35241 | RDX | Radixin | Color2 [Da | 1 | 4 |
| + | 3.077006 | 1.225327 | P35244 | P35244 | RPA3 | Replicatior | Color2 [Da | 1 | 4 |
| + | 2.26466 | -0.95205 | P35249 | P35249 | RFC4 | Replicatior | Color2 [Da | 1 | 4 |
| | 0.459223 | -0.45018 | P35250 | P35250 | RFC2 | Replicatior | Color2 [Da | 1 | 4 |
| | 0.902985 | -0.27809 | P35251 | P35251 | RFC1 | Replicatior | Color2 [Da | 1 | 4 |
| + | 2.540191 | 1.289805 | P35268 | P35268 | RPL22 | 60S riboso | Color2 [Da | 1 | 4 |
| + | 1.434722 | 0.562626 | P35269 | P35269 | GTF2F1 | General tra | Color2 [Da | 1 | 4 |
| + | 0.864203 | 1.357193 | P35270 | P35270 | SPR | Sepiapterii | Color2 [Da | 1 | 4 |
| + | 1.421886 | -0.62967 | P35442 | P35442 | THBS2 | Thrombos | Color2 [Da | 1 | 4 |
| | 0 | NaN | P35443 | P35443 | THBS4 | Thrombos | Color2 [Da | 1 | 4 |
| + | 1.359067 | 0.291906 | P35475 | P35475 | IDUA | Alpha-L-idu | Color2 [Da | 1 | 4 |
| + | 2.725521 | -1.61254 | P35520 | P35520 | CBS | Cystathion | Color2 [Da | 1 | 4 |
| + | 4.39293 | 1.682596 | P35527 | P35527 | KRT9 | Keratin, ty | Color2 [Da | 1 | 4 |
| + | 1.205158 | -1.36895 | P35555 | P35555 | FBN1 | Fibrillin-1 | Color2 [Da | 1 | 4 |
| + | 1.365828 | -0.69339 | P35556 | P35556 | FBN2 | Fibrillin-2 | Color2 [Da | 1 | 4 |
| + | 0.857749 | -0.42215 | P35573 | P35573 | AGL | Glycogen c | Color2 [Da | 1 | 4 |
| + | 4.090615 | -1.63228 | P35579 | P35579 | MYH9 | Myosin-9 | Color2 [Da | 1 | 4 |
| + | 2.441999 | -0.41861 | P35580 | P35580 | MYH10 | Myosin-10 | Color2 [Da | 1 | 4 |
| + | 4.209369 | -1.40801 | P35606 | P35606 | COPB2 | Coatomer | Color2 [Da | 1 | 4 |
| | 0 | -0.71339 | P35610 | P35610 | SOAT1 | Sterol O-ac | Color2 [Da | 1 | 4 |
| | 0.043971 | -0.00768 | P35611 | P35611 | ADD1 | Alpha-add | Color2 [Da | 1 | 4 |
| + | 2.413438 | 1.118116 | P35612 | P35612 | ADD2 | Beta-addu | Color2 [Da | 1 | 4 |
| + | 2.095849 | -0.9945 | P35613 | P35613 | BSG | Basigin | Color2 [Da | 1 | 4 |
| | 0.146879 | -0.10503 | P35625 | P35625 | TIMP3 | Metallopro | Color2 [Da | 1 | 4 |
| + | 1.807907 | 0.995344 | P35637 | P35637 | FUS | RNA-bindi | Color2 [Da | 1 | 4 |
| | 0.019661 | -0.00573 | P35658 | P35658 | NUP214 | Nuclear pc | Color2 [Da | 1 | 4 |
| + | 1.811843 | -0.64849 | P35659 | P35659 | DEK | Protein DE | Color2 [Da | 1 | 4 |
| | 0 | -0.3283 | P35711 | P35711 | SOX5 | Transcripti | Color2 [Da | 1 | 4 |
| | 0.475096 | 0.46837 | P35716 | P35716 | SOX11 | Transcripti | Color2 [Da | 1 | 4 |
| | 0 | 2.392768 | P35754 | P35754 | GLRX | Glutaredo: | Color2 [Da | 1 | 4 |
| | 0.681299 | 0.14246 | P35813 | P35813 | PPM1A | Protein ph | Color2 [Da | 1 | 4 |
| + | 3.858252 | 1.765755 | P35908 | P35908 | KRT2 | Keratin, ty | Color2 [Da | 1 | 4 |
| | 0.670425 | 0.47584 | P35914 | P35914 | HMGCL | Hydroxym | Color2 [Da | 1 | 4 |
| | 0.003209 | 0.001011 | P35998 | P35998 | PSMC2 | 26S protea | Color2 [Da | 1 | 4 |
| | 0 | -1.04169 | P36383 | P36383 | GJC1 | Gap juncti | Color2 [Da | 1 | 4 |
| + | 2.751381 | 1.536447 | P36404 | P36404 | ARL2 | ADP-ribosy | Color2 [Da | 1 | 4 |
| + | 3.521527 | 1.135019 | P36405 | P36405 | ARL3 | ADP-ribosy | Color2 [Da | 1 | 4 |
| + | 1.509295 | -0.47398 | P36507 | P36507 | MAP2K2 | Dual speci | Color2 [Da | 1 | 4 |
| + | 1.834589 | 1.595592 | P36542 | P36542 | ATP5F1C | ATP syntha | Color2 [Da | 1 | 4 |
| + | 5.146197 | 1.70081 | P36543 | P36543 | ATP6V1E1 | V-type pro | Color2 [Da | 1 | 4 |
| + | 0.998099 | -0.5583 | P36551 | P36551 | CPOX | Oxygen-de | Color2 [Da | 1 | 4 |
| + | 1.181615 | -0.53608 | P36578 | P36578 | RPL4 | 60S riboso | Color2 [Da | 1 | 4 |
| | 0.153904 | 0.100592 | P36639 | P36639 | NUDT1 | Oxidized p | Color2 [Da | 1 | 4 |

| | | | | | | | | | |
|---|---|---|---|---|---|---|---|---|---|
| + | 1.654162 | -0.4275 | P36776 | P36776 | LONP1 | Lon protea | Color2 [Da | 1 | 4 |
| + | 2.527991 | 0.689484 | P36871 | P36871 | PGM1 | Phosphogl | Color2 [Da | 1 | 4 |
| | 0.12421 | -0.07973 | P36873 | P36873 | PPP1CC | Serine/thr | Color2 [Da | 1 | 4 |
| | 0.514778 | 0.344495 | P36896 | P36896 | ACVR1B | Activin rec | Color2 [Da | 1 | 4 |
| | 0 | -1.29222 | P36897 | P36897 | TGFBR1 | TGF-beta r | Color2 [Da | 1 | 4 |
| | 0.224888 | -0.08347 | P36915 | P36915 | GNL1 | Guanine n | Color2 [Da | 1 | 4 |
| + | 0.823229 | -0.44519 | P36954 | P36954 | POLR2I | DNA-direc | Color2 [Da | 1 | 4 |
| + | 3.531681 | 1.344032 | P36957 | P36957 | DLST | Dihydrolip | Color2 [Da | 1 | 4 |
| + | 2.333022 | -0.81957 | P36969 | P36969 | GPX4 | Phospholip | Color2 [Da | 1 | 4 |
| | 0.97482 | 0.303367 | P37108 | P37108 | SRP14 | Signal reco | Color2 [Da | 1 | 4 |
| | 0.906182 | -0.15615 | P37198 | P37198 | NUP62 | Nuclear pc | Color2 [Da | 1 | 4 |
| + | 1.734498 | 0.828811 | P37235 | P37235 | HPCAL1 | Hippocalci | Color2 [Da | 1 | 4 |
| + | 2.383059 | -1.23396 | P37268 | P37268 | FDFT1 | Squalene s | Color2 [Da | 1 | 4 |
| + | 3.129259 | 1.769316 | P37802 | P37802 | TAGLN2 | Transgelin | Color2 [Da | 1 | 4 |
| + | 2.150982 | 1.049748 | P37837 | P37837 | TALDO1 | Transaldol | Color2 [Da | 1 | 4 |
| + | 2.701582 | 2.978119 | P37840 | P37840 | SNCA | Alpha-synu | Color2 [Da | 1 | 4 |
| | 0.03781 | -0.01955 | P38117 | P38117 | ETFB | Electron tr | Color2 [Da | 1 | 4 |
| + | 2.464135 | 1.244531 | P38159 | P38159 | RBMX | RNA-bindi | Color2 [Da | 1 | 4 |
| | 0 | -0.40785 | P38405 | P38405 | GNAL | Guanine n | Color2 [Da | 1 | 4 |
| | 0.055399 | 0.032507 | P38432 | P38432 | COIL | Coilin | Color2 [Da | 1 | 4 |
| | 0 | NaN | P38435 | P38435 | GGCX | Vitamin K- | Color2 [Da | 1 | 4 |
| | 0.0875 | 0.142958 | P38570 | P38570 | ITGAE | Integrin al | Color2 [Da | 1 | 4 |
| | 0.820167 | -0.30544 | P38571 | P38571 | LIPA | Lysosomal | Color2 [Da | 1 | 4 |
| | 0.38854 | -0.13281 | P38606 | P38606 | ATP6V1A | V-type pro | Color2 [Da | 1 | 4 |
| | 0.156686 | 0.013925 | P38646 | P38646 | HSPA9 | Stress-70 p | Color2 [Da | 1 | 4 |
| + | 2.250191 | -0.77883 | P38919 | P38919 | EIF4A3 | Eukaryotic | Color2 [Da | 1 | 4 |
| | 0 | 1.307991 | P38936 | P38936 | CDKN1A | Cyclin-dep | Color2 [Da | 1 | 4 |
| + | 1.777185 | 1.046166 | P39019 | P39019 | RPS19 | 40S riboso | Color2 [Da | 1 | 4 |
| + | 3.140907 | -1.10064 | P39023 | P39023 | RPL3 | 60S riboso | Color2 [Da | 1 | 4 |
| | 0.437687 | -0.60352 | P39060 | P39060 | COL18A1 | Collagen a | Color2 [Da | 1 | 4 |
| + | 2.420951 | -1.15539 | P39656 | P39656 | DDOST | Dolichyl-di | Color2 [Da | 1 | 4 |
| + | 1.937016 | 1.138441 | P39687 | P39687 | ANP32A | Acidic leuc | Color2 [Da | 1 | 4 |
| + | 2.731011 | -1.28444 | P39748 | P39748 | FEN1 | Flap endor | Color2 [Da | 1 | 4 |
| | 0.008318 | -0.00212 | P39880 | P39880 | CUX1 | Homeobox | Color2 [Da | 1 | 4 |
| | 0 | 0.936034 | P40121 | P40121 | CAPG | Macropha | Color2 [Da | 1 | 4 |
| | 0.015775 | -0.0045 | P40123 | P40123 | CAP2 | Adenylyl c | Color2 [Da | 1 | 4 |
| + | 1.182572 | -0.60854 | P40222 | P40222 | TXLNA | Alpha-taxil | Color2 [Da | 1 | 4 |
| | 0.166892 | 0.05607 | P40227 | P40227 | CCT6A | T-complex | Color2 [Da | 1 | 4 |
| + | 2.309777 | -0.26636 | P40424 | P40424 | PBX1 | Pre-B-cell l | Color2 [Da | 1 | 4 |
| + | 1.574576 | -0.70087 | P40425 | P40425 | PBX2 | Pre-B-cell l | Color2 [Da | 1 | 4 |
| + | 2.12224 | 0.825295 | P40426 | P40426 | PBX3 | Pre-B-cell l | Color2 [Da | 1 | 4 |
| | 0.374438 | -0.12901 | P40429 | P40429 | RPL13A | 60S riboso | Color2 [Da | 1 | 4 |
| | 0.155528 | -0.17911 | P40616 | P40616 | ARL1 | ADP-ribosy | Color2 [Da | 1 | 4 |
| | 0.164565 | 0.138996 | P40692 | P40692 | MLH1 | DNA mism | Color2 [Da | 1 | 4 |
| + | 2.322376 | -1.09049 | P40763 | P40763 | STAT3 | Signal tran | Color2 [Da | 1 | 4 |
| | 0.465231 | 0.140519 | P40818 | P40818 | USP8 | Ubiquitin c | Color2 [Da | 1 | 4 |
| | 0.076962 | -0.02691 | P40855 | P40855 | PEX19 | Peroxisom | Color2 [Da | 1 | 4 |

| | | | | | | | | | |
|---|---|---|---|---|---|---|---|---|---|
| + | 2.20953 | 1.279258 | P40925 | P40925 | MDH1 | Malate del | Color2 [Da | 1 | 4 |
| + | 3.529181 | 1.616145 | P40926 | P40926 | MDH2 | Malate del | Color2 [Da | 1 | 4 |
| + | 1.932586 | -0.62863 | P40937 | P40937 | RFC5 | Replicatior | Color2 [Da | 1 | 4 |
| | 0 | -3.74625 | P40938 | P40938 | RFC3 | Replicatior | Color2 [Da | 1 | 4 |
| + | 3.913192 | -1.29342 | P40939 | P40939 | HADHA | Trifunctior | Color2 [Da | 1 | 4 |
| + | 3.790093 | -1.01728 | P41091 | P41091 | EIF2S3 | Eukaryotic | Color2 [Da | 1 | 4 |
| + | 4.558664 | 1.423466 | P41208 | P41208 | CETN2 | Centrin-2 | Color2 [Da | 1 | 4 |
| | 0 | 1.806177 | P41214 | P41214 | EIF2D | Eukaryotic | Color2 [Da | 1 | 4 |
| | 0.503964 | 0.552763 | P41217 | P41217 | CD200 | OX-2 mem | Color2 [Da | 1 | 4 |
| | 0.886252 | 0.092443 | P41219 | P41219 | PRPH | Peripherin | Color2 [Da | 1 | 4 |
| | 0 | NaN | P41222 | P41222 | PTGDS | Prostaglan | Color2 [Da | 1 | 4 |
| + | 3.096962 | -1.33071 | P41223 | P41223 | BUD31 | Protein BU | Color2 [Da | 1 | 4 |
| + | 2.118663 | 2.116935 | P41225 | P41225 | SOX3 | Transcripti | Color2 [Da | 1 | 4 |
| + | 1.457727 | 0.45593 | P41227 | P41227 | NAA10 | N-alpha-ac | Color2 [Da | 1 | 4 |
| | 0 | -2.03061 | P41229 | P41229 | KDM5C | Lysine-spe | Color2 [Da | 1 | 4 |
| + | 2.676697 | 2.010157 | P41236 | P41236 | PPP1R2 | Protein ph | Color2 [Da | 1 | 4 |
| | 0.783133 | -0.29023 | P41240 | P41240 | CSK | Tyrosine-p | Color2 [Da | 1 | 4 |
| + | 3.022489 | -0.89697 | P41250 | P41250 | GARS1 | Glycine--tF | Color2 [Da | 1 | 4 |
| + | 2.879261 | -1.37033 | P41252 | P41252 | IARS1 | Isoleucine- | Color2 [Da | 1 | 4 |
| | 0 | NaN | P41271 | P41271 | NBL1 | Neuroblas | Color2 [Da | 1 | 4 |
| + | 1.934246 | 1.065059 | P41567 | P41567 | EIF1 | Eukaryotic | Color2 [Da | 1 | 4 |
| | 0 | -0.16848 | P41594 | P41594 | GRM5 | Metabotrc | Color2 [Da | 1 | 4 |
| + | 4.759044 | -1.87104 | P41732 | P41732 | TSPAN7 | Tetraspani | Color2 [Da | 1 | 4 |
| + | 1.708503 | -0.7752 | P41743 | P41743 | PRKCI | Protein kir | Color2 [Da | 1 | 4 |
| + | 2.58166 | 0.706785 | P42025 | P42025 | ACTR1B | Beta-centr | Color2 [Da | 1 | 4 |
| + | 3.435999 | 1.213229 | P42126 | P42126 | ECI1 | Enoyl-CoA | Color2 [Da | 1 | 4 |
| | 1.482154 | 0.175004 | P42166 | P42166 | TMPO | Lamina-as: | Color2 [Da | 1 | 4 |
| | 0.69438 | -0.18009 | P42167 | P42167 | TMPO | Lamina-as: | Color2 [Da | 1 | 4 |
| + | 2.667051 | -1.21577 | P42224 | P42224 | STAT1 | Signal tran | Color2 [Da | 1 | 4 |
| | 0 | NaN | P42226 | P42226 | STAT6 | Signal tran | Color2 [Da | 1 | 4 |
| + | 1.454151 | -0.65857 | P42261 | P42261 | GRIA1 | Glutamate | Color2 [Da | 1 | 4 |
| + | 3.389436 | -1.20256 | P42262 | P42262 | GRIA2 | Glutamate | Color2 [Da | 1 | 4 |
| + | 1.506622 | -0.53527 | P42285 | P42285 | MTREX | Exosome F | Color2 [Da | 1 | 4 |
| + | 1.176451 | -0.51429 | P42330 | P42330 | AKR1C3 | Aldo-keto | Color2 [Da | 1 | 4 |
| | 0 | -0.42962 | P42336 | P42336 | PIK3CA | Phosphati | Color2 [Da | 1 | 4 |
| | 0 | NaN | P42338 | P42338 | PIK3CB | Phosphati | Color2 [Da | 1 | 4 |
| + | 1.697539 | -0.56223 | P42345 | P42345 | MTOR | Serine/thr | Color2 [Da | 1 | 4 |
| | 0.354493 | -0.11527 | P42356 | P42356 | PI4KA | Phosphati | Color2 [Da | 1 | 4 |
| | 0.517298 | 0.696946 | P42357 | P42357 | HAL | Histidine a | Color2 [Da | 1 | 4 |
| + | 1.613253 | -0.51487 | P42566 | P42566 | EPS15 | Epidermal | Color2 [Da | 1 | 4 |
| + | 1.404108 | -1.0019 | P42574 | P42574 | CASP3 | Caspase-3 | Color2 [Da | 1 | 4 |
| | 0 | NaN | P42658 | P42658 | DPP6 | Dipeptidyl | Color2 [Da | 1 | 4 |
| + | 3.443336 | -1.3478 | P42677 | P42677 | RPS27 | 40S riboso | Color2 [Da | 1 | 4 |
| | 0.100521 | -0.24491 | P42684 | P42684 | ABL2 | Tyrosine-p | Color2 [Da | 1 | 4 |
| | 0.817257 | -0.31064 | P42694 | P42694 | HELZ | Probable h | Color2 [Da | 1 | 4 |
| | 0 | 1.976188 | P42695 | P42695 | NCAPD3 | Condensin | Color2 [Da | 1 | 4 |
| | 0.572627 | 1.550868 | P42696 | P42696 | RBM34 | RNA-bindi | Color2 [Da | 1 | 4 |

| | | | | | | | | | |
|---|---|---|---|---|---|---|---|---|---|
| + | 2.959861 | -1.03415 | P42704 | P42704 | LRPPRC | Leucine-ric | Color2 [Da | 1 | 4 |
| + | 1.284388 | -0.35948 | P42765 | P42765 | ACAA2 | 3-ketoacyl | Color2 [Da | 1 | 4 |
| | 0.294514 | -0.10102 | P42766 | P42766 | RPL35 | 60S riboso | Color2 [Da | 1 | 4 |
| | 0.450045 | -0.12044 | P42785 | P42785 | PRCP | Lysosomal | Color2 [Da | 1 | 4 |
| | 0.282468 | 0.229659 | P42857 | P42857 | NSG1 | Neuronal v | Color2 [Da | 1 | 4 |
| + | 2.195144 | -0.53392 | P42858 | P42858 | HTT | Huntingtin | Color2 [Da | 1 | 4 |
| + | 1.062692 | -0.71208 | P42892 | P42892 | ECE1 | Endothelir | Color2 [Da | 1 | 4 |
| + | 0.902329 | -1.83832 | P42898 | P42898 | MTHFR | Methylene | Color2 [Da | 1 | 4 |
| | 0.010868 | -0.00329 | P43003 | P43003 | SLC1A3 | Excitatory | Color2 [Da | 1 | 4 |
| | 0.338256 | 0.95765 | P43004 | P43004 | SLC1A2 | Excitatory | Color2 [Da | 1 | 4 |
| + | 1.614704 | -0.59403 | P43007 | P43007 | SLC1A4 | Neutral an | Color2 [Da | 1 | 4 |
| + | 1.868486 | -0.42231 | P43034 | P43034 | PAFAH1B1 | Platelet-ac | Color2 [Da | 1 | 4 |
| + | 1.513369 | -0.66729 | P43121 | P43121 | MCAM | Cell surfac | Color2 [Da | 1 | 4 |
| + | 2.097016 | -0.60919 | P43146 | P43146 | DCC | Netrin rece | Color2 [Da | 1 | 4 |
| + | 1.180632 | -0.28443 | P43155 | P43155 | CRAT | Carnitine O | Color2 [Da | 1 | 4 |
| + | 1.448609 | 0.314016 | P43243 | P43243 | MATR3 | Matrin-3 | Color2 [Da | 1 | 4 |
| + | 2.824239 | -0.90893 | P43246 | P43246 | MSH2 | DNA mism | Color2 [Da | 1 | 4 |
| + | 5.407642 | -1.49132 | P43304 | P43304 | GPD2 | Glycerol-3- | Color2 [Da | 1 | 4 |
| + | 1.063344 | 0.399132 | P43307 | P43307 | SSR1 | Translocor | Color2 [Da | 1 | 4 |
| + | 1.76373 | -0.61916 | P43378 | P43378 | PTPN9 | Tyrosine-p | Color2 [Da | 1 | 4 |
| + | 1.537665 | 0.790271 | P43487 | P43487 | RANBP1 | Ran-specif | Color2 [Da | 1 | 4 |
| + | 2.552903 | -1.13078 | P43490 | P43490 | NAMPT | Nicotinam | Color2 [Da | 1 | 4 |
| | 0 | NaN | P43652 | P43652 | AFM | Afamin | Color2 [Da | 1 | 4 |
| + | 3.369602 | 0.831807 | P43686 | P43686 | PSMC4 | 26S protea | Color2 [Da | 1 | 4 |
| | 0.349654 | -0.35587 | P43897 | P43897 | TSFM | Elongation | Color2 [Da | 1 | 4 |
| | 0 | NaN | P45378 | P45378 | TNNT3 | Troponin T | Color2 [Da | 1 | 4 |
| | 0 | -0.39159 | P45877 | P45877 | PPIC | Peptidyl-pr | Color2 [Da | 1 | 4 |
| + | 2.335689 | -0.62959 | P45880 | P45880 | VDAC2 | Voltage-de | Color2 [Da | 1 | 4 |
| + | 1.160943 | 0.396311 | P45954 | P45954 | ACADSB | Short/bran | Color2 [Da | 1 | 4 |
| + | 2.786564 | 0.936405 | P45973 | P45973 | CBX5 | Chromobo | Color2 [Da | 1 | 4 |
| | 0.385361 | -0.07175 | P45974 | P45974 | USP5 | Ubiquitin c | Color2 [Da | 1 | 4 |
| + | 1.343478 | -0.50724 | P45983 | P45983 | MAPK8 | Mitogen-a | Color2 [Da | 1 | 4 |
| | 0 | -0.66479 | P45984 | P45984 | MAPK9 | Mitogen-a | Color2 [Da | 1 | 4 |
| + | 1.657182 | -0.94662 | P45985 | P45985 | MAP2K4 | Dual speci | Color2 [Da | 1 | 4 |
| + | 1.593142 | -0.60648 | P46013 | P46013 | MKI67 | Proliferatio | Color2 [Da | 1 | 4 |
| | 0 | -0.46494 | P46019 | P46019 | PHKA2 | Phosphory | Color2 [Da | 1 | 4 |
| | 0.722711 | -0.29142 | P46060 | P46060 | RANGAP1 | Ran GTPas | Color2 [Da | 1 | 4 |
| + | 2.509468 | -1.04164 | P46063 | P46063 | RECQL | ATP-deper | Color2 [Da | 1 | 4 |
| | 0.415008 | 0.345795 | P46087 | P46087 | NOP2 | Probable 2 | Color2 [Da | 1 | 4 |
| | 0.351382 | -0.10494 | P46100 | P46100 | ATRX | Transcripti | Color2 [Da | 1 | 4 |
| + | 2.956551 | 1.390544 | P46108 | P46108 | CRK | Adapter m | Color2 [Da | 1 | 4 |
| + | 2.397475 | 1.130609 | P46109 | P46109 | CRKL | Crk-like pr | Color2 [Da | 1 | 4 |
| + | 1.573742 | 1.21307 | P46199 | P46199 | MTIF2 | Translation | Color2 [Da | 1 | 4 |
| + | 2.590393 | 0.368675 | P46379 | P46379 | BAG6 | Large proli | Color2 [Da | 1 | 4 |
| + | 1.512 | 0.353754 | P46459 | P46459 | NSF | Vesicle-fus | Color2 [Da | 1 | 4 |
| + | 2.69677 | 2.487513 | P46527 | P46527 | CDKN1B | Cyclin-dep | Color2 [Da | 1 | 4 |
| + | 1.081272 | -1.07822 | P46734 | P46734 | MAP2K3 | Dual speci | Color2 [Da | 1 | 4 |

| | | | | | | | | | |
|---|---|---|---|---|---|---|---|---|---|
| + | 2.294793 | -0.51015 | P46736 | P46736 | BRCC3 | Lys-63-spe | Color2 [Da | 1 | 4 |
| | 0.211346 | -0.13 | P46776 | P46776 | RPL27A | 60S riboso | Color2 [Da | 1 | 4 |
| + | 2.0211 | -0.77493 | P46777 | P46777 | RPL5 | 60S riboso | Color2 [Da | 1 | 4 |
| + | 1.280991 | 0.466478 | P46778 | P46778 | RPL21 | 60S riboso | Color2 [Da | 1 | 4 |
| + | 1.766829 | 0.647218 | P46779 | P46779 | RPL28 | 60S riboso | Color2 [Da | 1 | 4 |
| | 0.5041 | -0.2208 | P46781 | P46781 | RPS9 | 40S riboso | Color2 [Da | 1 | 4 |
| + | 1.433778 | 0.337618 | P46782 | P46782 | RPS5 | 40S riboso | Color2 [Da | 1 | 4 |
| + | 3.019759 | 1.361932 | P46783 | P46783 | RPS10 | 40S riboso | Color2 [Da | 1 | 4 |
| + | 4.646232 | 1.42789 | P46821 | P46821 | MAP1B | Microtubu | Color2 [Da | 1 | 4 |
| + | 3.457707 | 2.417142 | P46926 | P46926 | GNPDA1 | Glucosami | Color2 [Da | 1 | 4 |
| | 0 | 0.469468 | P46934 | P46934 | NEDD4 | E3 ubiquiti | Color2 [Da | 1 | 4 |
| + | 3.378372 | 2.049548 | P46937 | P46937 | YAP1 | Transcripti | Color2 [Da | 1 | 4 |
| + | 1.662484 | -0.26719 | P46939 | P46939 | UTRN | Utrophin | Color2 [Da | 1 | 4 |
| + | 3.145125 | -1.36305 | P46940 | P46940 | IQGAP1 | Ras GTPase | Color2 [Da | 1 | 4 |
| | 0.264223 | -0.07129 | P46976 | P46976 | GYG1 | Glycogenir | Color2 [Da | 1 | 4 |
| + | 2.721697 | -1.14791 | P46977 | P46977 | STT3A | Dolichyl-di | Color2 [Da | 1 | 4 |
| + | 1.716777 | 1.319184 | P47736 | P47736 | RAP1GAP | Rap1 GTPa | Color2 [Da | 1 | 4 |
| + | 2.416617 | 1.720551 | P47755 | P47755 | CAPZA2 | F-actin-cap | Color2 [Da | 1 | 4 |
| + | 3.418045 | -0.85516 | P47756 | P47756 | CAPZB | F-actin-cap | Color2 [Da | 1 | 4 |
| + | 2.5723 | 1.849757 | P47813 | P47813 | EIF1AX | Eukaryotic | Color2 [Da | 1 | 4 |
| | 0 | NaN | P47870 | P47870 | GABRB2 | Gamma-ar | Color2 [Da | 1 | 4 |
| + | 2.634089 | -0.79833 | P47897 | P47897 | QARS1 | Glutamine | Color2 [Da | 1 | 4 |
| + | 2.582722 | 1.498427 | P47914 | P47914 | RPL29 | 60S riboso | Color2 [Da | 1 | 4 |
| | 0.405726 | 0.275279 | P47929 | P47929 | LGALS7B | Galectin-7 | Color2 [Da | 1 | 4 |
| + | 0.989709 | 0.941645 | P47972 | P47972 | NPTX2 | Neuronal p | Color2 [Da | 1 | 4 |
| + | 4.728987 | 1.798011 | P47985 | P47985 | UQCRFS1 | Cytochrom | Color2 [Da | 1 | 4 |
| + | 3.032986 | 1.524049 | P48047 | P48047 | ATP5PO | ATP syntha | Color2 [Da | 1 | 4 |
| | 0.353465 | 0.36808 | P48058 | P48058 | GRIA4 | Glutamate | Color2 [Da | 1 | 4 |
| + | 2.181986 | -0.89973 | P48147 | P48147 | PREP | Prolyl endo | Color2 [Da | 1 | 4 |
| + | 0.790223 | 0.52448 | P48200 | P48200 | IREB2 | Iron-respo | Color2 [Da | 1 | 4 |
| | 0.012703 | 0.004042 | P48380 | P48380 | RFX3 | Transcripti | Color2 [Da | 1 | 4 |
| | 0.289442 | 0.265873 | P48382 | P48382 | RFX5 | DNA-bindi | Color2 [Da | 1 | 4 |
| + | 1.572711 | -0.92619 | P48426 | P48426 | PIP4K2A | Phosphatic | Color2 [Da | 1 | 4 |
| + | 1.889433 | 1.964254 | P48431 | P48431 | SOX2 | Transcripti | Color2 [Da | 1 | 4 |
| + | 2.495579 | 1.968476 | P48436 | P48436 | SOX9 | Transcripti | Color2 [Da | 1 | 4 |
| + | 1.889719 | -0.39763 | P48444 | P48444 | ARCN1 | Coatomer | Color2 [Da | 1 | 4 |
| + | 3.011136 | -1.46587 | P48449 | P48449 | LSS | Lanosterol | Color2 [Da | 1 | 4 |
| | 0.062267 | -0.09378 | P48454 | P48454 | PPP3CC | Serine/thr | Color2 [Da | 1 | 4 |
| + | 4.737015 | -1.31396 | P48506 | P48506 | GCLC | Glutamate | Color2 [Da | 1 | 4 |
| + | 0.995416 | 0.871149 | P48507 | P48507 | GCLM | Glutamate | Color2 [Da | 1 | 4 |
| | 0 | NaN | P48509 | P48509 | CD151 | CD151 ant | Color2 [Da | 1 | 4 |
| + | 2.526347 | 2.677207 | P48539 | P48539 | PCP4 | Calmoduli | Color2 [Da | 1 | 4 |
| | 0 | NaN | P48553 | P48553 | TRAPPC10 | Trafficking | Color2 [Da | 1 | 4 |
| | 0.339114 | -0.13637 | P48556 | P48556 | PSMD8 | 26S protea | Color2 [Da | 1 | 4 |
| | 0 | NaN | P48594 | P48594 | SERPINB4 | Serpin B4 | Color2 [Da | 1 | 4 |
| + | 1.344756 | 1.167991 | P48634 | P48634 | PRRC2A | Protein PR | Color2 [Da | 1 | 4 |
| + | 4.382223 | 2.536245 | P48637 | P48637 | GSS | Glutathion | Color2 [Da | 1 | 4 |

| | | | | | | | | | |
|---|---|---|---|---|---|---|---|---|---|
| + | 3.52144 | -0.6533 | P48643 | P48643 | CCT5 | T-complex | Color2 [Da | 1 | 4 |
| | 0 | NaN | P48645 | P48645 | NMU | Neuromed | Color2 [Da | 1 | 4 |
| | 0 | -0.35485 | P48651 | P48651 | PTDSS1 | Phosphatic | Color2 [Da | 1 | 4 |
| + | 5.384047 | 2.329609 | P48681 | P48681 | NES | Nestin | Color2 [Da | 1 | 4 |
| + | 1.521857 | 0.830312 | P48723 | P48723 | HSPA13 | Heat shock | Color2 [Da | 1 | 4 |
| | 0 | NaN | P48728 | P48728 | AMT | Aminomet | Color2 [Da | 1 | 4 |
| + | 0.944942 | -0.3562 | P48729 | P48729 | CSNK1A1 | Casein kina | Color2 [Da | 1 | 4 |
| + | 3.82928 | -1.1061 | P48730 | P48730 | CSNK1D | Casein kina | Color2 [Da | 1 | 4 |
| + | 2.181099 | -0.87624 | P48735 | P48735 | IDH2 | Isocitrate ( | Color2 [Da | 1 | 4 |
| + | 3.249463 | -0.47169 | P48739 | P48739 | PITPNB | Phosphatic | Color2 [Da | 1 | 4 |
| | 0 | NaN | P48742;Q9 | P48742;Q9 | LHX1;LHX5 | LIM/home | Color2 [Da | 1 | 4 |
| + | 2.852453 | 1.25212 | P49006 | P49006 | MARCKSL1 | MARCKS-re | Color2 [Da | 1 | 4 |
| + | 0.893486 | 0.688991 | P49023 | P49023 | PXN | Paxillin | Color2 [Da | 1 | 4 |
| + | 1.396014 | 0.703985 | P49069 | P49069 | CAMLG | Guided en | Color2 [Da | 1 | 4 |
| + | 2.397997 | -0.74517 | P49116 | P49116 | NR2C2 | Nuclear re | Color2 [Da | 1 | 4 |
| + | 1.76747 | -0.52572 | P49189 | P49189 | ALDH9A1 | 4-trimethy | Color2 [Da | 1 | 4 |
| | 0.206774 | 0.092039 | P49207 | P49207 | RPL34 | 60S riboso | Color2 [Da | 1 | 4 |
| + | 1.105359 | 0.298663 | P49247 | P49247 | RPIA | Ribose-5-p | Color2 [Da | 1 | 4 |
| + | 3.853976 | -1.81789 | P49257 | P49257 | LMAN1 | Protein ER | Color2 [Da | 1 | 4 |
| | 0.163821 | -0.05597 | P49321 | P49321 | NASP | Nuclear au | Color2 [Da | 1 | 4 |
| + | 3.330992 | -1.14954 | P49327 | P49327 | FASN | Fatty acid : | Color2 [Da | 1 | 4 |
| + | 1.803694 | -0.38329 | P49354 | P49354 | FNTA | Protein far | Color2 [Da | 1 | 4 |
| | 0.13662 | -0.11752 | P49356 | P49356 | FNTB | Protein far | Color2 [Da | 1 | 4 |
| | 1.039939 | -0.09083 | P49366 | P49366 | DHPS | Deoxyhypu | Color2 [Da | 1 | 4 |
| + | 2.352247 | -0.75537 | P49368 | P49368 | CCT3 | T-complex | Color2 [Da | 1 | 4 |
| + | 2.382065 | 0.800348 | P49406 | P49406 | MRPL19 | 39S riboso | Color2 [Da | 1 | 4 |
| + | 1.696017 | 0.307336 | P49407 | P49407 | ARRB1 | Beta-arres | Color2 [Da | 1 | 4 |
| | 0.45411 | 0.098611 | P49411 | P49411 | TUFM | Elongation | Color2 [Da | 1 | 4 |
| + | 3.690498 | 2.356073 | P49418 | P49418 | AMPH | Amphiphy: | Color2 [Da | 1 | 4 |
| + | 2.859116 | -0.66998 | P49419 | P49419 | ALDH7A1 | Alpha-ami | Color2 [Da | 1 | 4 |
| | 0.694603 | 0.266217 | P49427 | P49427 | CDC34 | Ubiquitin-( | Color2 [Da | 1 | 4 |
| + | 5.120085 | 0.454041 | P49441 | P49441 | INPP1 | Inositol po | Color2 [Da | 1 | 4 |
| | 0.583796 | -0.6262 | P49450 | P49450 | CENPA | Histone H3 | Color2 [Da | 1 | 4 |
| | 0 | NaN | P49454 | P49454 | CENPF | Centromer | Color2 [Da | 1 | 4 |
| + | 1.370254 | 0.80228 | P49458 | P49458 | SRP9 | Signal reco | Color2 [Da | 1 | 4 |
| | 0.347984 | -0.51201 | P49459 | P49459 | UBE2A | Ubiquitin-( | Color2 [Da | 1 | 4 |
| | 0.06326 | -0.04875 | P49585 | P49585 | PCYT1A | Choline-ph | Color2 [Da | 1 | 4 |
| + | 1.591004 | -0.23991 | P49588 | P49588 | AARS1 | Alanine--tI | Color2 [Da | 1 | 4 |
| + | 2.374138 | -0.75565 | P49589 | P49589 | CARS1 | Cysteine--t | Color2 [Da | 1 | 4 |
| + | 1.013432 | -0.39806 | P49590 | P49590 | HARS2 | Histidine-- | Color2 [Da | 1 | 4 |
| + | 2.661935 | -0.98893 | P49591 | P49591 | SARS1 | Serine--tRI | Color2 [Da | 1 | 4 |
| + | 1.191962 | 0.394014 | P49593 | P49593 | PPM1F | Protein ph | Color2 [Da | 1 | 4 |
| + | 0.850621 | -0.52173 | P49641 | P49641 | MAN2A2 | Alpha-mar | Color2 [Da | 1 | 4 |
| | 0 | NaN | P49643 | P49643 | PRIM2 | DNA prima | Color2 [Da | 1 | 4 |
| | 0.026335 | 0.022031 | P49674 | P49674 | CSNK1E | Casein kina | Color2 [Da | 1 | 4 |
| | 0 | NaN | P49703 | P49703 | ARL4D | ADP-ribosy | Color2 [Da | 1 | 4 |
| + | 1.868696 | -1.15269 | P49711 | P49711 | CTCF | Transcripti | Color2 [Da | 1 | 4 |

| | | | | | | | | | |
|---|---|---|---|---|---|---|---|---|---|
| + | 2.649836 | 1.780398 | P49720 | P49720 | PSMB3 | Proteasom | Color2 [Da | 1 | 4 |
| + | 3.883664 | 1.749445 | P49721 | P49721 | PSMB2 | Proteasom | Color2 [Da | 1 | 4 |
| + | 4.222037 | -1.08833 | P49736 | P49736 | MCM2 | DNA replic | Color2 [Da | 1 | 4 |
| | 0 | NaN | P49747 | P49747 | COMP | Cartilage o | Color2 [Da | 1 | 4 |
| + | 1.425379 | -0.43396 | P49748 | P49748 | ACADVL | Very long- | Color2 [Da | 1 | 4 |
| | 0.014563 | 0.00366 | P49750 | P49750 | YLPM1 | YLP motif- | Color2 [Da | 1 | 4 |
| | 0 | -3.98234 | P49754 | P49754 | VPS41 | Vacuolar p | Color2 [Da | 1 | 4 |
| + | 1.746758 | -0.63896 | P49755 | P49755 | TMED10 | Transmem | Color2 [Da | 1 | 4 |
| + | 2.507405 | -0.39126 | P49756 | P49756 | RBM25 | RNA-bindi | Color2 [Da | 1 | 4 |
| + | 3.196296 | 1.670411 | P49758 | P49758 | RGS6 | Regulator | Color2 [Da | 1 | 4 |
| + | 3.015626 | -0.89809 | P49770 | P49770 | EIF2B2 | Translation | Color2 [Da | 1 | 4 |
| + | 3.493249 | 1.150756 | P49773 | P49773 | HINT1 | Adenosine | Color2 [Da | 1 | 4 |
| | 0.664732 | 0.530497 | P49789 | P49789 | FHIT | Bis(5'-ader | Color2 [Da | 1 | 4 |
| + | 1.089269 | -1.09587 | P49790 | P49790 | NUP153 | Nuclear pc | Color2 [Da | 1 | 4 |
| + | 3.226856 | -0.8553 | P49792 | P49792 | RANBP2 | E3 SUMO- | Color2 [Da | 1 | 4 |
| + | 1.794404 | 2.404338 | P49796 | P49796 | RGS3 | Regulator | Color2 [Da | 1 | 4 |
| + | 3.112453 | 2.576608 | P49802 | P49802 | RGS7 | Regulator | Color2 [Da | 1 | 4 |
| | 0.755345 | -0.4776 | P49815 | P49815 | TSC2 | Tuberin | Color2 [Da | 1 | 4 |
| + | 3.658574 | -1.7241 | P49821 | P49821 | NDUFV1 | NADH deh | Color2 [Da | 1 | 4 |
| | 0.098271 | -0.04642 | P49840 | P49840 | GSK3A | Glycogen s | Color2 [Da | 1 | 4 |
| + | 1.96153 | -0.76369 | P49841 | P49841 | GSK3B | Glycogen s | Color2 [Da | 1 | 4 |
| | 0.565797 | 0.219778 | P49842 | P49842 | STK19 | Serine/thr | Color2 [Da | 1 | 4 |
| + | 1.336458 | -1.24165 | P49848 | P49848 | TAF6 | Transcripti | Color2 [Da | 1 | 4 |
| | 0 | NaN | P49862 | P49862 | KLK7 | Kallikrein-7 | Color2 [Da | 1 | 4 |
| + | 3.545069 | -1.98928 | P49902 | P49902 | NT5C2 | Cytosolic p | Color2 [Da | 1 | 4 |
| + | 2.511452 | -0.88453 | P49903 | P49903 | SEPHS1 | Selenide, v | Color2 [Da | 1 | 4 |
| | 0 | NaN | P49913 | P49913 | CAMP | Cathelicidi | Color2 [Da | 1 | 4 |
| + | 2.677924 | -1.56201 | P49915 | P49915 | GMPS | GMP synth | Color2 [Da | 1 | 4 |
| + | 2.021399 | -0.44909 | P49916 | P49916 | LIG3 | DNA ligase | Color2 [Da | 1 | 4 |
| | 0.368741 | -0.2095 | P49959 | P49959 | MRE11 | Double-str | Color2 [Da | 1 | 4 |
| | 0.106805 | -0.06542 | P50148 | P50148 | GNAQ | Guanine n | Color2 [Da | 1 | 4 |
| + | 3.541371 | 1.649636 | P50150 | P50150 | GNG4 | Guanine n | Color2 [Da | 1 | 4 |
| | 0 | NaN | P50151 | P50151 | GNG10 | Guanine n | Color2 [Da | 1 | 4 |
| + | 1.350973 | 0.514752 | P50213 | P50213 | IDH3A | Isocitrate | Color2 [Da | 1 | 4 |
| + | 3.918862 | -2.22476 | P50225;P5 | P50225;P5 | SULT1A1;S | Sulfotrans | Color2 [Da | 1 | 4 |
| | 0 | -0.62317 | P50281 | P50281 | MMP14 | Matrix me | Color2 [Da | 1 | 4 |
| + | 1.595905 | -1.11116 | P50336 | P50336 | PPOX | Protoporp | Color2 [Da | 1 | 4 |
| + | 2.067499 | -0.6493 | P50395 | P50395 | GDI2 | Rab GDP d | Color2 [Da | 1 | 4 |
| | 0.168505 | -0.0226 | P50402 | P50402 | EMD | Emerin | Color2 [Da | 1 | 4 |
| + | 3.833447 | -1.73856 | P50416 | P50416 | CPT1A | Carnitine O | Color2 [Da | 1 | 4 |
| | 0.520847 | 0.131469 | P50440 | P50440 | GATM | Glycine an | Color2 [Da | 1 | 4 |
| + | 0.880296 | 1.910096 | P50452 | P50452 | SERPINB8 | Serpin B8 | Color2 [Da | 1 | 4 |
| + | 1.333351 | 0.495448 | P50453 | P50453 | SERPINB9 | Serpin B9 | Color2 [Da | 1 | 4 |
| | 0.065719 | 0.020287 | P50454 | P50454 | SERPINH1 | Serpin H1 | Color2 [Da | 1 | 4 |
| + | 3.196141 | 0.887151 | P50502 | P50502 | ST13 | Hsc70-inte | Color2 [Da | 1 | 4 |
| + | 3.016546 | -0.66648 | P50542 | P50542 | PEX5 | Peroxisom | Color2 [Da | 1 | 4 |
| | 0.082381 | 0.03909 | P50552 | P50552 | VASP | Vasodilato | Color2 [Da | 1 | 4 |

| | | | | | | | | | |
|---|---|---|---|---|---|---|---|---|---|
| + | 1.400037 | 0.338971 | P50570 | P50570 | DNM2 | Dynamin-2 | Color2 [Da | 1 | 4 |
| | 0.169552 | -0.08646 | P50579 | P50579 | METAP2 | Methionin | Color2 [Da | 1 | 4 |
| | 0.868458 | 0.250767 | P50583 | P50583 | NUDT2 | Bis(5'-nucl | Color2 [Da | 1 | 4 |
| + | 1.151078 | 1.790695 | P50607 | P50607 | TUB | Tubby pro | Color2 [Da | 1 | 4 |
| + | 1.509527 | -1.27512 | P50613 | P50613 | CDK7 | Cyclin-dep | Color2 [Da | 1 | 4 |
| + | 0.956064 | -0.49899 | P50749 | P50749 | RASSF2 | Ras associ | Color2 [Da | 1 | 4 |
| | 0.351969 | -0.19386 | P50750 | P50750 | CDK9 | Cyclin-dep | Color2 [Da | 1 | 4 |
| + | 1.027865 | -0.70338 | P50851 | P50851 | LRBA | Lipopolysa | Color2 [Da | 1 | 4 |
| | 0.057286 | -0.09292 | P50895 | P50895 | BCAM | Basal cell a | Color2 [Da | 1 | 4 |
| | 0.545164 | 0.500123 | P50897 | P50897 | PPT1 | Palmitoyl-| | Color2 [Da | 1 | 4 |
| + | 2.187972 | 2.03565 | P50914 | P50914 | RPL14 | 60S riboso | Color2 [Da | 1 | 4 |
| + | 2.657412 | -1.01817 | P50990 | P50990 | CCT8 | T-complex | Color2 [Da | 1 | 4 |
| + | 1.232048 | -0.24519 | P50991 | P50991 | CCT4 | T-complex | Color2 [Da | 1 | 4 |
| | 0 | NaN | P50993 | P50993 | ATP1A2 | Sodium/pc | Color2 [Da | 1 | 4 |
| + | 3.311827 | 1.731656 | P50995 | P50995 | ANXA11 | Annexin A | Color2 [Da | 1 | 4 |
| + | 2.47732 | -0.93094 | P51003 | P51003 | PAPOLA | Poly(A) po | Color2 [Da | 1 | 4 |
| | 0.144068 | 0.035411 | P51114 | P51114 | FXR1 | Fragile X m | Color2 [Da | 1 | 4 |
| | 0.62732 | -0.18345 | P51116 | P51116 | FXR2 | Fragile X m | Color2 [Da | 1 | 4 |
| + | 2.796884 | 1.146235 | P51148 | P51148 | RAB5C | Ras-relate | Color2 [Da | 1 | 4 |
| + | 1.931189 | 1.053167 | P51149 | P51149 | RAB7A | Ras-relate | Color2 [Da | 1 | 4 |
| + | 2.737362 | 0.728292 | P51151 | P51151 | RAB9A | Ras-relate | Color2 [Da | 1 | 4 |
| + | 1.59017 | -1.35165 | P51153 | P51153 | RAB13 | Ras-relate | Color2 [Da | 1 | 4 |
| | 0.302234 | 0.535569 | P51157 | P51157 | RAB28 | Ras-relate | Color2 [Da | 1 | 4 |
| + | 2.06933 | 2.775754 | P51161 | P51161 | FABP6 | Gastrotrop | Color2 [Da | 1 | 4 |
| + | 2.924795 | -1.13944 | P51178 | P51178 | PLCD1 | 1-phospha | Color2 [Da | 1 | 4 |
| + | 1.344513 | -0.61969 | P51398 | P51398 | DAP3 | 28S riboso | Color2 [Da | 1 | 4 |
| + | 1.512183 | 0.652888 | P51452 | P51452 | DUSP3 | Dual speci | Color2 [Da | 1 | 4 |
| + | 1.952903 | 1.115647 | P51511 | P51511 | MMP15 | Matrix me | Color2 [Da | 1 | 4 |
| + | 3.567235 | 1.894407 | P51513 | P51513 | NOVA1 | RNA-bindi | Color2 [Da | 1 | 4 |
| | 0.045313 | 0.028398 | P51531 | P51531 | SMARCA2 | Probable g | Color2 [Da | 1 | 4 |
| + | 3.112499 | -0.84927 | P51532 | P51532 | SMARCA4 | Transcripti | Color2 [Da | 1 | 4 |
| | 0.117331 | 0.049282 | P51553 | P51553 | IDH3G | Isocitrate | Color2 [Da | 1 | 4 |
| | 0.679184 | -0.12469 | P51570 | P51570 | GALK1 | Galactokin | Color2 [Da | 1 | 4 |
| + | 2.686952 | -0.8422 | P51571 | P51571 | SSR4 | Translocor | Color2 [Da | 1 | 4 |
| + | 2.231082 | -0.80424 | P51572 | P51572 | BCAP31 | B-cell rece | Color2 [Da | 1 | 4 |
| + | 2.358371 | -1.10975 | P51580 | P51580 | TPMT | Thiopurine | Color2 [Da | 1 | 4 |
| + | 2.893839 | 2.089594 | P51608 | P51608 | MECP2 | Methyl-Cp | Color2 [Da | 1 | 4 |
| | 1.028312 | -0.08822 | P51610 | P51610 | HCFC1 | Host cell fa | Color2 [Da | 1 | 4 |
| + | 3.152697 | -1.54281 | P51648 | P51648 | ALDH3A2 | Aldehyde | Color2 [Da | 1 | 4 |
| + | 2.384406 | 0.887229 | P51649 | P51649 | ALDH5A1 | Succinate- | Color2 [Da | 1 | 4 |
| + | 1.79987 | -0.26133 | P51659 | P51659 | HSD17B4 | Peroxisom | Color2 [Da | 1 | 4 |
| + | 2.073875 | 1.136755 | P51665 | P51665 | PSMD7 | 26S protea | Color2 [Da | 1 | 4 |
| | 0.940854 | -0.13189 | P51668;Q9 | P51668;Q9 | UBE2D1;U | Ubiquitin- | Color2 [Da | 1 | 4 |
| + | 2.002634 | -1.12814 | P51674 | P51674 | GPM6A | Neuronal r | Color2 [Da | 1 | 4 |
| | 0.249794 | -0.12871 | P51687 | P51687 | SUOX | Sulfite oxi | Color2 [Da | 1 | 4 |
| + | 4.645028 | 3.625046 | P51688 | P51688 | SGSH | N-sulphogl | Color2 [Da | 1 | 4 |
| | 0.396452 | 0.184513 | P51692 | P51692 | STAT5B | Signal tran | Color2 [Da | 1 | 4 |

| | | | | | | | | | |
|---|---|---|---|---|---|---|---|---|---|
| | 0.497802 | 0.330179 | P51693 | P51693 | APLP1 | Amyloid b | Color2 [Da | 1 | 4 |
| + | 2.191424 | -0.99245 | P51784 | P51784 | USP11 | Ubiquitin c | Color2 [Da | 1 | 4 |
| | 0 | NaN | P51790 | P51790 | CLCN3 | H(+)/Cl(-) e | Color2 [Da | 1 | 4 |
| | 0 | NaN | P51793;P5 | P51793;P5 | CLCN4;CLC | H(+)/Cl(-) e | Color2 [Da | 1 | 4 |
| | 0.60789 | 0.44632 | P51797 | P51797 | CLCN6 | H(+)/Cl(-) e | Color2 [Da | 1 | 4 |
| | 0.144117 | -0.29206 | P51798 | P51798 | CLCN7 | H(+)/Cl(-) e | Color2 [Da | 1 | 4 |
| + | 1.4575 | -0.58748 | P51805 | P51805 | PLXNA3 | Plexin-A3 | Color2 [Da | 1 | 4 |
| + | 1.891678 | -0.67529 | P51809 | P51809 | VAMP7 | Vesicle-ass | Color2 [Da | 1 | 4 |
| + | 1.823161 | -0.43382 | P51812 | P51812 | RPS6KA3 | Ribosomal | Color2 [Da | 1 | 4 |
| | 0 | NaN | P51817 | P51817 | PRKX | cAMP-dep | Color2 [Da | 1 | 4 |
| + | 1.485829 | -0.75895 | P51826 | P51826 | AFF3 | AF4/FMR2 | Color2 [Da | 1 | 4 |
| + | 2.839902 | 2.117735 | P51858 | P51858 | HDGF | Hepatoma | Color2 [Da | 1 | 4 |
| | 0 | NaN | P51946 | P51946 | CCNH | Cyclin-H | Color2 [Da | 1 | 4 |
| | 0.199363 | 0.393398 | P51948 | P51948 | MNAT1 | CDK-activa | Color2 [Da | 1 | 4 |
| + | 1.915681 | 0.965691 | P51956 | P51956 | NEK3 | Serine/thr | Color2 [Da | 1 | 4 |
| + | 1.844692 | 0.588868 | P51970 | P51970 | NDUFA8 | NADH deh | Color2 [Da | 1 | 4 |
| + | 1.77033 | 0.322324 | P51991 | P51991 | HNRNPA3 | Heterogen | Color2 [Da | 1 | 4 |
| + | 2.683184 | -1.14982 | P52209 | P52209 | PGD | 6-phospho | Color2 [Da | 1 | 4 |
| | 0.179567 | 0.054539 | P52272 | P52272 | HNRNPM | Heterogen | Color2 [Da | 1 | 4 |
| + | 2.988223 | -1.88522 | P52292 | P52292 | KPNA2 | Importin s | Color2 [Da | 1 | 4 |
| + | 0.879022 | -0.78669 | P52294 | P52294 | KPNA1 | Importin s | Color2 [Da | 1 | 4 |
| | 0.236754 | 0.591708 | P52298 | P52298 | NCBP2 | Nuclear ca | Color2 [Da | 1 | 4 |
| + | 2.857889 | -0.96419 | P52306 | P52306 | RAP1GDS1 | Rap1 GTPa | Color2 [Da | 1 | 4 |
| | 0.509914 | -0.26085 | P52429 | P52429 | DGKE | Diacylglyce | Color2 [Da | 1 | 4 |
| | 0.090346 | -0.05289 | P52434 | P52434 | POLR2H | DNA-direc | Color2 [Da | 1 | 4 |
| | 0.147787 | 0.242959 | P52435;Q9 | P52435;Q9 | POLR2J;PC | DNA-direc | Color2 [Da | 1 | 4 |
| + | 2.871498 | -1.5935 | P52564 | P52564 | MAP2K6 | Dual speci | Color2 [Da | 1 | 4 |
| + | 1.941046 | 0.561528 | P52565 | P52565 | ARHGDIA | Rho GDP-d | Color2 [Da | 1 | 4 |
| | 0 | NaN | P52566 | P52566 | ARHGDIB | Rho GDP-d | Color2 [Da | 1 | 4 |
| + | 2.505955 | 0.731999 | P52594 | P52594 | AGFG1 | Arf-GAP do | Color2 [Da | 1 | 4 |
| | 0.397303 | -0.14388 | P52597 | P52597 | HNRNPF | Heterogen | Color2 [Da | 1 | 4 |
| | 0.129058 | 0.251109 | P52630 | P52630 | STAT2 | Signal tran | Color2 [Da | 1 | 4 |
| + | 2.758876 | 3.222572 | P52655 | P52655 | GTF2A1 | Transcripti | Color2 [Da | 1 | 4 |
| + | 3.834638 | -1.60271 | P52701 | P52701 | MSH6 | DNA mism | Color2 [Da | 1 | 4 |
| + | 0.983952 | -0.72837 | P52732 | P52732 | KIF11 | Kinesin-lik | Color2 [Da | 1 | 4 |
| + | 4.177235 | -1.85398 | P52735 | P52735 | VAV2 | Guanine n | Color2 [Da | 1 | 4 |
| | 0.171697 | 0.430686 | P52739 | P52739 | ZNF131 | Zinc finger | Color2 [Da | 1 | 4 |
| | 0.238919 | 0.37014 | P52747 | P52747 | ZNF143 | Zinc finger | Color2 [Da | 1 | 4 |
| | 0.400856 | 0.129496 | P52756 | P52756 | RBM5 | RNA-bindi | Color2 [Da | 1 | 4 |
| + | 3.765921 | 3.486598 | P52758 | P52758 | RIDA | 2-iminobu | Color2 [Da | 1 | 4 |
| + | 1.973363 | -0.68745 | P52788 | P52788 | SMS | Spermine s | Color2 [Da | 1 | 4 |
| | 0 | NaN | P52789 | P52789 | HK2 | Hexokinas | Color2 [Da | 1 | 4 |
| | 0 | 0.160808 | P52797 | P52797 | EFNA3 | Ephrin-A3 | Color2 [Da | 1 | 4 |
| + | 1.642508 | -0.53229 | P52799 | P52799 | EFNB2 | Ephrin-B2 | Color2 [Da | 1 | 4 |
| + | 3.682932 | 3.372627 | P52815 | P52815 | MRPL12 | 39S riboso | Color2 [Da | 1 | 4 |
| | 0 | NaN | P52824 | P52824 | DGKQ | Diacylglyce | Color2 [Da | 1 | 4 |
| | 0.201201 | 0.274365 | P52849 | P52849 | NDST2 | Bifunction | Color2 [Da | 1 | 4 |

| | | | | | | | | | |
|---|---|---|---|---|---|---|---|---|---|
| + | 2.593939 | -1.12629 | P52888 | P52888 | THOP1 | Thimet oli | Color2 [Da | 1 | 4 |
| + | 1.26178 | -0.5249 | P52895 | P52895 | AKR1C2 | Aldo-keto | Color2 [Da | 1 | 4 |
| + | 2.065572 | 1.296022 | P52907 | P52907 | CAPZA1 | F-actin-cap | Color2 [Da | 1 | 4 |
| | 0.056867 | -0.01561 | P52926 | P52926 | HMGA2 | High mobi | Color2 [Da | 1 | 4 |
| + | 1.17104 | 0.761139 | P52943 | P52943 | CRIP2 | Cysteine-ri | Color2 [Da | 1 | 4 |
| | 1.602843 | 0.124485 | P52948 | P52948 | NUP98 | Nuclear pc | Color2 [Da | 1 | 4 |
| + | 1.207275 | -0.52906 | P53004 | P53004 | BLVRA | Biliverdin r | Color2 [Da | 1 | 4 |
| + | 1.576723 | -0.89201 | P53007 | P53007 | SLC25A1 | Tricarboxy | Color2 [Da | 1 | 4 |
| + | 2.543122 | -0.79128 | P53041 | P53041 | PPP5C | Serine/thr | Color2 [Da | 1 | 4 |
| | 0.217477 | -0.32197 | P53355 | P53355 | DAPK1 | Death-ass | Color2 [Da | 1 | 4 |
| + | 4.702139 | 0.798798 | P53365 | P53365 | ARFIP2 | Arfaptin-2 | Color2 [Da | 1 | 4 |
| + | 3.169318 | 1.695865 | P53367 | P53367 | ARFIP1 | Arfaptin-1 | Color2 [Da | 1 | 4 |
| + | 0.981126 | -0.89335 | P53384 | P53384 | NUBP1 | Cytosolic F | Color2 [Da | 1 | 4 |
| + | 5.270757 | -1.33243 | P53396 | P53396 | ACLY | ATP-citrate | Color2 [Da | 1 | 4 |
| | 0.561949 | -0.25034 | P53582 | P53582 | METAP1 | Methionin | Color2 [Da | 1 | 4 |
| + | 0.990936 | 0.335612 | P53597 | P53597 | SUCLG1 | Succinate- | Color2 [Da | 1 | 4 |
| | 0.602841 | -0.34447 | P53602 | P53602 | MVD | Diphosphc | Color2 [Da | 1 | 4 |
| + | 2.029071 | -1.61045 | P53611 | P53611 | RABGGTB | Geranylge | Color2 [Da | 1 | 4 |
| + | 5.456932 | -1.26603 | P53618 | P53618 | COPB1 | Coatomer | Color2 [Da | 1 | 4 |
| + | 4.018138 | -1.77283 | P53621 | P53621 | COPA | Coatomer | Color2 [Da | 1 | 4 |
| | 1.181131 | -0.13441 | P53634 | P53634 | CTSC | Dipeptidyl | Color2 [Da | 1 | 4 |
| | 0 | -0.63052 | P53667 | P53667 | LIMK1 | LIM domai | Color2 [Da | 1 | 4 |
| + | 0.771126 | 0.525642 | P53677 | P53677 | AP3M2 | AP-3 comp | Color2 [Da | 1 | 4 |
| + | 3.874721 | -1.56934 | P53680 | P53680 | AP2S1 | AP-2 comp | Color2 [Da | 1 | 4 |
| + | 1.610656 | -1.46676 | P53701 | P53701 | HCCS | Holocytocl | Color2 [Da | 1 | 4 |
| | 0.678564 | 0.326597 | P53779 | P53779 | MAPK10 | Mitogen-a | Color2 [Da | 1 | 4 |
| + | 0.991588 | 0.530056 | P53805 | P53805 | RCAN1 | Calcipressi | Color2 [Da | 1 | 4 |
| + | 2.670083 | -1.46783 | P53985 | P53985 | SLC16A1 | Monocarb | Color2 [Da | 1 | 4 |
| + | 2.356403 | 0.585084 | P53990 | P53990 | IST1 | IST1 homo | Color2 [Da | 1 | 4 |
| + | 3.893146 | -1.92758 | P53992 | P53992 | SEC24C | Protein tra | Color2 [Da | 1 | 4 |
| + | 2.998409 | 0.817707 | P53999 | P53999 | SUB1 | Activated I | Color2 [Da | 1 | 4 |
| + | 1.313173 | 1.217698 | P54105 | P54105 | CLNS1A | Methylosc | Color2 [Da | 1 | 4 |
| + | 3.995095 | -1.3164 | P54136 | P54136 | RARS1 | Arginine--t | Color2 [Da | 1 | 4 |
| | 0 | NaN | P54198 | P54198 | HIRA | Protein HII | Color2 [Da | 1 | 4 |
| + | 3.362453 | -0.6872 | P54252 | P54252 | ATXN3 | Ataxin-3 | Color2 [Da | 1 | 4 |
| | 0.379632 | -0.31983 | P54277 | P54277 | PMS1 | PMS1 prot | Color2 [Da | 1 | 4 |
| | 0 | 0.193743 | P54278 | P54278 | PMS2 | Mismatch | Color2 [Da | 1 | 4 |
| + | 2.927014 | 1.768268 | P54284 | P54284 | CACNB3 | Voltage-de | Color2 [Da | 1 | 4 |
| + | 2.788046 | -1.44562 | P54289 | P54289 | CACNA2D1 | Voltage-de | Color2 [Da | 1 | 4 |
| | 0.367208 | 0.120165 | P54577 | P54577 | YARS1 | Tyrosine--t | Color2 [Da | 1 | 4 |
| + | 1.937373 | -0.80623 | P54578 | P54578 | USP14 | Ubiquitin c | Color2 [Da | 1 | 4 |
| + | 3.739057 | -1.19735 | P54619 | P54619 | PRKAG1 | 5'-AMP-ac | Color2 [Da | 1 | 4 |
| | 0.443046 | -0.75178 | P54646 | P54646 | PRKAA2 | 5'-AMP-ac | Color2 [Da | 1 | 4 |
| + | 2.241354 | -0.67697 | P54687 | P54687 | BCAT1 | Branched- | Color2 [Da | 1 | 4 |
| + | 3.579312 | -1.73207 | P54709 | P54709 | ATP1B3 | Sodium/pc | Color2 [Da | 1 | 4 |
| + | 3.105238 | 2.916419 | P54725 | P54725 | RAD23A | UV excisio | Color2 [Da | 1 | 4 |
| + | 4.860565 | 2.437254 | P54727 | P54727 | RAD23B | UV excisio | Color2 [Da | 1 | 4 |

| | | | | | | | | | |
|---|---|---|---|---|---|---|---|---|---|
| | 0.382192 | -0.42972 | P54750 | P54750 | PDE1A | Dual speci | Color2 [Da | 1 | 4 |
| | 0 | 0.098372 | P54753 | P54753 | EPHB3 | Ephrin typ | Color2 [Da | 1 | 4 |
| | 0.595211 | 0.557398 | P54756 | P54756 | EPHA5 | Ephrin typ | Color2 [Da | 1 | 4 |
| | 0 | -0.65342 | P54762 | P54762 | EPHB1 | Ephrin typ | Color2 [Da | 1 | 4 |
| | 0.156545 | 0.06849 | P54764 | P54764 | EPHA4 | Ephrin typ | Color2 [Da | 1 | 4 |
| | 0.501129 | 0.132233 | P54802 | P54802 | NAGLU | Alpha-N-ac | Color2 [Da | 1 | 4 |
| | 0.44093 | 0.136628 | P54803 | P54803 | GALC | Galactocer | Color2 [Da | 1 | 4 |
| + | 2.661259 | 1.480283 | P54819 | P54819 | AK2 | Adenylate | Color2 [Da | 1 | 4 |
| | 0 | 0.246491 | P54829 | P54829 | PTPN5 | Tyrosine-p | Color2 [Da | 1 | 4 |
| | 0 | NaN | P54868 | P54868 | HMGCS2 | Hydroxym | Color2 [Da | 1 | 4 |
| + | 2.723889 | -1.2074 | P54886 | P54886 | ALDH18A1 | Delta-1-py | Color2 [Da | 1 | 4 |
| | 0.440624 | -0.10253 | P54920 | P54920 | NAPA | Alpha-solu | Color2 [Da | 1 | 4 |
| | 0.274637 | 0.450533 | P55001 | P55001 | MFAP2 | Microfibril | Color2 [Da | 1 | 4 |
| + | 2.332813 | -0.57804 | P55010 | P55010 | EIF5 | Eukaryotic | Color2 [Da | 1 | 4 |
| + | 1.927719 | 0.682661 | P55011 | P55011 | SLC12A2 | Solute carr | Color2 [Da | 1 | 4 |
| + | 2.294383 | 1.33743 | P55036 | P55036 | PSMD4 | 26S protea | Color2 [Da | 1 | 4 |
| + | 1.484657 | -0.93154 | P55039 | P55039 | DRG2 | Developm | Color2 [Da | 1 | 4 |
| | 0.618362 | -0.18774 | P55058 | P55058 | PLTP | Phospholip | Color2 [Da | 1 | 4 |
| + | 4.455021 | -1.5589 | P55060 | P55060 | CSE1L | Exportin-2 | Color2 [Da | 1 | 4 |
| | 0.079388 | -0.09564 | P55061 | P55061 | TMBIM6 | Bax inhibit | Color2 [Da | 1 | 4 |
| + | 3.289064 | -0.92173 | P55072 | P55072 | VCP | Transitiona | Color2 [Da | 1 | 4 |
| + | 3.158805 | 1.104516 | P55081 | P55081 | MFAP1 | Microfibril | Color2 [Da | 1 | 4 |
| + | 3.218747 | 1.282404 | P55084 | P55084 | HADHB | Trifunctior | Color2 [Da | 1 | 4 |
| | 0 | 1.921772 | P55087 | P55087 | AQP4 | Aquaporin | Color2 [Da | 1 | 4 |
| | 0 | NaN | P55107 | P55107 | GDF10 | Growth/di | Color2 [Da | 1 | 4 |
| + | 2.552019 | 0.687676 | P55145 | P55145 | MANF | Mesencep | Color2 [Da | 1 | 4 |
| | 0.348569 | -0.18647 | P55157 | P55157 | MTTP | Microsoma | Color2 [Da | 1 | 4 |
| + | 2.507953 | 0.653678 | P55196 | P55196 | AFDN | Afadin | Color2 [Da | 1 | 4 |
| | 0 | NaN | P55201 | P55201 | BRPF1 | Peregrin | Color2 [Da | 1 | 4 |
| | 0.477691 | 0.13998 | P55209 | P55209 | NAP1L1 | Nucleoson | Color2 [Da | 1 | 4 |
| + | 4.434209 | -1.36039 | P55210 | P55210 | CASP7 | Caspase-7 | Color2 [Da | 1 | 4 |
| + | 4.056947 | -1.69235 | P55212 | P55212 | CASP6 | Caspase-6 | Color2 [Da | 1 | 4 |
| | 0.101644 | -0.04553 | P55263 | P55263 | ADK | Adenosine | Color2 [Da | 1 | 4 |
| + | 1.613292 | -0.79011 | P55265 | P55265 | ADAR | Double-str | Color2 [Da | 1 | 4 |
| + | 2.734551 | -1.53175 | P55268 | P55268 | LAMB2 | Laminin su | Color2 [Da | 1 | 4 |
| + | 1.820143 | 1.624744 | P55273 | P55273 | CDKN2D | Cyclin-dep | Color2 [Da | 1 | 4 |
| + | 1.374588 | 0.571135 | P55283 | P55283 | CDH4 | Cadherin-4 | Color2 [Da | 1 | 4 |
| | 0 | NaN | P55285 | P55285 | CDH6 | Cadherin-6 | Color2 [Da | 1 | 4 |
| + | 1.942851 | 1.707345 | P55286 | P55286 | CDH8 | Cadherin-8 | Color2 [Da | 1 | 4 |
| + | 4.64849 | 3.546703 | P55290 | P55290 | CDH13 | Cadherin-1 | Color2 [Da | 1 | 4 |
| | 0 | 3.677623 | P55316 | P55316 | FOXG1 | Forkhead I | Color2 [Da | 1 | 4 |
| + | 2.251981 | 1.851299 | P55327 | P55327 | TPD52 | Tumor pro | Color2 [Da | 1 | 4 |
| + | 2.877305 | -1.02048 | P55345 | P55345 | PRMT2 | Protein arg | Color2 [Da | 1 | 4 |
| + | 0.662544 | -1.44337 | P55347 | P55347 | PKNOX1 | Homeobox | Color2 [Da | 1 | 4 |
| + | 1.113655 | -0.3759 | P55735 | P55735 | SEC13 | Protein SE | Color2 [Da | 1 | 4 |
| + | 2.847145 | -0.62014 | P55769 | P55769 | SNU13 | NHP2-like | Color2 [Da | 1 | 4 |
| + | 2.92787 | -0.44432 | P55786 | P55786 | NPEPPS | Puromycin | Color2 [Da | 1 | 4 |

| | | | | | | | | | |
|---|---|---|---|---|---|---|---|---|---|
| | 0 | 0.669211 | P55789 | P55789 | GFER | FAD-linkec | Color2 [Da | 1 | 4 |
| + | 1.320692 | 0.538856 | P55795 | P55795 | HNRNPH2 | Heterogen | Color2 [Da | 1 | 4 |
| + | 4.022822 | 0.740658 | P55809 | P55809 | OXCT1 | Succinyl-C | Color2 [Da | 1 | 4 |
| + | 2.761392 | -0.92114 | P55884 | P55884 | EIF3B | Eukaryotic | Color2 [Da | 1 | 4 |
| + | 1.985767 | 1.605856 | P55957 | P55957 | BID | BH3-intera | Color2 [Da | 1 | 4 |
| + | 2.738145 | 0.61836 | P56134 | P56134 | ATP5MF | ATP syntha | Color2 [Da | 1 | 4 |
| | 0 | NaN | P56177 | P56177 | DLX1 | Homeobox | Color2 [Da | 1 | 4 |
| | 0 | NaN | P56178 | P56178 | DLX5 | Homeobox | Color2 [Da | 1 | 4 |
| | 0 | NaN | P56179 | P56179 | DLX6 | Homeobox | Color2 [Da | 1 | 4 |
| | 0.248739 | 0.327656 | P56181 | P56181 | NDUFV3 | NADH deh | Color2 [Da | 1 | 4 |
| + | 1.261133 | 0.581947 | P56182 | P56182 | RRP1 | Ribosomal | Color2 [Da | 1 | 4 |
| + | 4.835341 | -1.57887 | P56192 | P56192 | MARS1 | Methionin | Color2 [Da | 1 | 4 |
| + | 3.521844 | 4.070459 | P56211 | P56211 | ARPP19 | cAMP-regu | Color2 [Da | 1 | 4 |
| + | 1.515549 | 1.315668 | P56270 | P56270 | MAZ | Myc-assoc | Color2 [Da | 1 | 4 |
| + | 2.271256 | -1.49881 | P56373 | P56373 | P2RX3 | P2X purinc | Color2 [Da | 1 | 4 |
| + | 1.624453 | -1.32793 | P56377 | P56377 | AP1S2 | AP-1 comp | Color2 [Da | 1 | 4 |
| + | 2.838782 | 2.528066 | P56378 | P56378 | ATP5MJ | ATP syntha | Color2 [Da | 1 | 4 |
| | 0 | 1.441239 | P56381;Q5 | P56381;Q5 | ATP5F1E;A | ATP syntha | Color2 [Da | 1 | 4 |
| + | 3.391936 | 4.173568 | P56385 | P56385 | ATP5ME | ATP syntha | Color2 [Da | 1 | 4 |
| | 0 | -0.10349 | P56524 | P56524 | HDAC4 | Histone de | Color2 [Da | 1 | 4 |
| + | 1.856977 | 0.242286 | P56537 | P56537 | EIF6 | Eukaryotic | Color2 [Da | 1 | 4 |
| | 0.15161 | -0.33011 | P56545 | P56545 | CTBP2 | C-terminal | Color2 [Da | 1 | 4 |
| + | 1.766701 | 1.014225 | P56556 | P56556 | NDUFA6 | NADH deh | Color2 [Da | 1 | 4 |
| | 0 | 0.602192 | P56559 | P56559 | ARL4C | ADP-ribosy | Color2 [Da | 1 | 4 |
| | 0 | NaN | P56589 | P56589 | PEX3 | Peroxisom | Color2 [Da | 1 | 4 |
| | 0 | -0.42998 | P56703 | P56703 | WNT3 | Proto-onco | Color2 [Da | 1 | 4 |
| | 0 | 0.428721 | P56817 | P56817 | BACE1 | Beta-secre | Color2 [Da | 1 | 4 |
| | 0 | NaN | P56937 | P56937 | HSD17B7 | 3-keto-ste | Color2 [Da | 1 | 4 |
| + | 2.758351 | 1.058836 | P56945 | P56945 | BCAR1 | Breast can | Color2 [Da | 1 | 4 |
| + | 2.021536 | 1.102379 | P56962 | P56962 | STX17 | Syntaxin-1 | Color2 [Da | 1 | 4 |
| + | 1.220852 | -0.65331 | P57071 | P57071 | PRDM15 | PR domain | Color2 [Da | 1 | 4 |
| + | 1.261405 | -0.39774 | P57076 | P57076 | CFAP298 | Cilia- and f | Color2 [Da | 1 | 4 |
| | 0 | 1.77764 | P57081 | P57081 | WDR4 | tRNA (guar | Color2 [Da | 1 | 4 |
| + | 2.244193 | -0.88632 | P57088 | P57088 | TMEM33 | Transmem | Color2 [Da | 1 | 4 |
| + | 2.635468 | 1.901519 | P57105 | P57105 | SYNJ2BP | Synaptojar | Color2 [Da | 1 | 4 |
| + | 1.349764 | -1.4686 | P57678 | P57678 | GEMIN4 | Gem-assoc | Color2 [Da | 1 | 4 |
| + | 2.038848 | 2.12111 | P57721 | P57721 | PCBP3 | Poly(rC)-bi | Color2 [Da | 1 | 4 |
| | 0 | -1.63541 | P57723 | P57723 | PCBP4 | Poly(rC)-bi | Color2 [Da | 1 | 4 |
| + | 3.975478 | -1.86979 | P57737 | P57737 | CORO7 | Coronin-7 | Color2 [Da | 1 | 4 |
| + | 5.159626 | -1.85016 | P57740 | P57740 | NUP107 | Nuclear pc | Color2 [Da | 1 | 4 |
| | 0.148075 | -0.05401 | P57772 | P57772 | EEFSEC | Selenocyst | Color2 [Da | 1 | 4 |
| | 0.021604 | -0.00707 | P58004 | P58004 | SESN2 | Sestrin-2 | Color2 [Da | 1 | 4 |
| | 0.046689 | -0.00735 | P58107 | P58107 | EPPK1 | Epiplakin | Color2 [Da | 1 | 4 |
| | 0 | NaN | P58166 | P58166 | INHBE | Inhibin bet | Color2 [Da | 1 | 4 |
| + | 1.951011 | 1.065664 | P58401 | P58401 | NRXN2 | Neurexin-2 | Color2 [Da | 1 | 4 |
| | 0 | NaN | P58417 | P58417 | NXPH1 | Neurexoph | Color2 [Da | 1 | 4 |
| + | 2.612401 | 2.533351 | P58511 | P58511 | SMIM11 | Small integ | Color2 [Da | 1 | 4 |

| | | | | | | | | | |
|---|---|---|---|---|---|---|---|---|---|
| + | 3.524153 | 1.747393 | P58546 | P58546 | MTPN | Myotrophi | Color2 [Da | 1 | 4 |
| + | 1.5676 | 1.186566 | P58549 | P58549 | FXYD7 | FXYD dom | Color2 [Da | 1 | 4 |
| | 1.490476 | 0.191828 | P58876;Q5 | P58876;Q5 | H2BC5;H2I | Histone H2 | Color2 [Da | 1 | 4 |
| | 0.45496 | 0.831092 | P59190 | P59190 | RAB15 | Ras-relate | Color2 [Da | 1 | 4 |
| | 0 | NaN | P59665;P5 | P59665;P5 | DEFA1B;DI | Neutrophi | Color2 [Da | 1 | 4 |
| + | 2.934005 | 1.782436 | P59768 | P59768 | GNG2 | Guanine n | Color2 [Da | 1 | 4 |
| | 0 | 1.314965 | P59780 | P59780 | AP3S2 | AP-3 comp | Color2 [Da | 1 | 4 |
| | 0.318039 | -0.18473 | P59923 | P59923 | ZNF445 | Zinc finger | Color2 [Da | 1 | 4 |
| | 0.628645 | 0.436059 | P59998 | P59998 | ARPC4 | Actin-relat | Color2 [Da | 1 | 4 |
| + | 2.747404 | 1.402572 | P60006 | P60006 | ANAPC15 | Anaphase- | Color2 [Da | 1 | 4 |
| | 0.366755 | 0.407696 | P60033 | P60033 | CD81 | CD81 antig | Color2 [Da | 1 | 4 |
| + | 3.84995 | 3.396094 | P60059 | P60059 | SEC61G | Protein tra | Color2 [Da | 1 | 4 |
| + | 3.809725 | 1.87504 | P60174 | P60174 | TPI1 | Triosephos | Color2 [Da | 1 | 4 |
| + | 2.486486 | -1.04402 | P60228 | P60228 | EIF3E | Eukaryotic | Color2 [Da | 1 | 4 |
| + | 1.897072 | 1.208553 | P60468 | P60468 | SEC61B | Protein tra | Color2 [Da | 1 | 4 |
| | 0.049913 | -0.16365 | P60484 | P60484 | PTEN | Phosphatic | Color2 [Da | 1 | 4 |
| + | 0.914936 | -0.58 | P60510 | P60510 | PPP4C | Serine/thr | Color2 [Da | 1 | 4 |
| + | 2.658016 | 1.695019 | P60520 | P60520 | GABARAPL | Gamma-ar | Color2 [Da | 1 | 4 |
| + | 2.337679 | 2.49324 | P60602 | P60602 | ROMO1 | Reactive o | Color2 [Da | 1 | 4 |
| | 0.3964 | -0.07036 | P60660 | P60660 | MYL6 | Myosin ligl | Color2 [Da | 1 | 4 |
| + | 1.503847 | -0.44595 | P60709 | P60709 | ACTB | Actin, cyto | Color2 [Da | 1 | 4 |
| | 0.098076 | -0.08125 | P60763 | P60763 | RAC3 | Ras-relate | Color2 [Da | 1 | 4 |
| + | 3.087477 | -0.95378 | P60842 | P60842 | EIF4A1 | Eukaryotic | Color2 [Da | 1 | 4 |
| + | 1.78677 | 0.280472 | P60866 | P60866 | RPS20 | 40S riboso | Color2 [Da | 1 | 4 |
| + | 2.328565 | 1.406487 | P60880 | P60880 | SNAP25 | Synaptoso | Color2 [Da | 1 | 4 |
| + | 0.995625 | -0.56696 | P60891 | P60891 | PRPS1 | Ribose-pho | Color2 [Da | 1 | 4 |
| + | 4.132873 | 1.480369 | P60900 | P60900 | PSMA6 | Proteasom | Color2 [Da | 1 | 4 |
| + | 3.220676 | 2.643798 | P60903 | P60903 | S100A10 | Protein S1 | Color2 [Da | 1 | 4 |
| | 0.365601 | -0.32493 | P60953 | P60953 | CDC42 | Cell divisio | Color2 [Da | 1 | 4 |
| | 0.20652 | 0.099511 | P60981 | P60981 | DSTN | Destrin | Color2 [Da | 1 | 4 |
| + | 1.633238 | 0.872559 | P60983 | P60983 | GMFB | Glia matur | Color2 [Da | 1 | 4 |
| | 0 | NaN | P60985 | P60985 | KRTDAP | Keratinocy | Color2 [Da | 1 | 4 |
| | 0.736967 | -0.41565 | P61006 | P61006 | RAB8A | Ras-relate | Color2 [Da | 1 | 4 |
| + | 1.424863 | 0.962727 | P61009 | P61009 | SPCS3 | Signal pep | Color2 [Da | 1 | 4 |
| + | 3.287923 | -0.74533 | P61011 | P61011 | SRP54 | Signal reco | Color2 [Da | 1 | 4 |
| | 0.602584 | 0.422785 | P61018 | P61018 | RAB4B | Ras-relate | Color2 [Da | 1 | 4 |
| + | 1.226662 | 0.242642 | P61019 | P61019 | RAB2A | Ras-relate | Color2 [Da | 1 | 4 |
| + | 3.409555 | 1.247139 | P61020 | P61020 | RAB5B | Ras-relate | Color2 [Da | 1 | 4 |
| + | 3.606444 | -1.37725 | P61024 | P61024 | CKS1B | Cyclin-dep | Color2 [Da | 1 | 4 |
| + | 2.763646 | -0.7715 | P61026 | P61026 | RAB10 | Ras-relate | Color2 [Da | 1 | 4 |
| | 0.700255 | 0.461305 | P61077;P6 | P61077;P6 | UBE2D3;U | Ubiquitin-c | Color2 [Da | 1 | 4 |
| | 0.887844 | 0.149048 | P61081 | P61081 | UBE2M | NEDD8-co | Color2 [Da | 1 | 4 |
| + | 3.594828 | 0.850626 | P61086 | P61086 | UBE2K | Ubiquitin-c | Color2 [Da | 1 | 4 |
| + | 3.814429 | 2.030768 | P61088 | P61088 | UBE2N | Ubiquitin-c | Color2 [Da | 1 | 4 |
| + | 2.457608 | 0.885046 | P61106 | P61106 | RAB14 | Ras-relate | Color2 [Da | 1 | 4 |
| + | 1.546337 | -0.52854 | P61129 | P61129 | ZC3H6 | Zinc finger | Color2 [Da | 1 | 4 |
| + | 2.809051 | -1.10229 | P61158 | P61158 | ACTR3 | Actin-relat | Color2 [Da | 1 | 4 |

| | | | | | | | | | |
|---|---|---|---|---|---|---|---|---|---|
| + | 2.010514 | -0.67931 | P61160 | P61160 | ACTR2 | Actin-relat | Color2 [Da | 1 | 4 |
| + | 2.592829 | 0.591096 | P61163 | P61163 | ACTR1A | Alpha-cent | Color2 [Da | 1 | 4 |
| + | 2.7752 | -1.30339 | P61201 | P61201 | COPS2 | COP9 signa | Color2 [Da | 1 | 4 |
| + | 1.977394 | -1.03351 | P61221 | P61221 | ABCE1 | ATP-bindir | Color2 [Da | 1 | 4 |
| + | 2.4559 | 1.225703 | P61224 | P61224 | RAP1B | Ras-relate | Color2 [Da | 1 | 4 |
| | 0.502377 | 0.215979 | P61225 | P61225 | RAP2B | Ras-relate | Color2 [Da | 1 | 4 |
| | 0 | NaN | P61244 | P61244 | MAX | Protein ma | Color2 [Da | 1 | 4 |
| + | 1.971459 | -0.28586 | P61247 | P61247 | RPS3A | 40S riboso | Color2 [Da | 1 | 4 |
| + | 1.637315 | 1.014356 | P61254 | P61254 | RPL26 | 60S riboso | Color2 [Da | 1 | 4 |
| + | 2.028688 | 0.594327 | P61266 | P61266 | STX1B | Syntaxin-1 | Color2 [Da | 1 | 4 |
| + | 2.888652 | 0.768093 | P61289 | P61289 | PSME3 | Proteasom | Color2 [Da | 1 | 4 |
| + | 2.354508 | -1.37986 | P61313 | P61313 | RPL15 | 60S riboso | Color2 [Da | 1 | 4 |
| | 0 | NaN | P61326 | P61326 | MAGOH | Protein ma | Color2 [Da | 1 | 4 |
| + | 2.376254 | 0.829789 | P61353 | P61353 | RPL27 | 60S riboso | Color2 [Da | 1 | 4 |
| + | 3.658509 | -0.79833 | P61421 | P61421 | ATP6V0D1 | V-type pro | Color2 [Da | 1 | 4 |
| + | 2.566011 | 2.79073 | P61457 | P61457 | PCBD1 | Pterin-4-al | Color2 [Da | 1 | 4 |
| + | 0.931679 | 0.748579 | P61513 | P61513 | RPL37A | 60S riboso | Color2 [Da | 1 | 4 |
| + | 0.812928 | 0.643917 | P61586 | P61586 | RHOA | Transform | Color2 [Da | 1 | 4 |
| + | 2.049181 | -1.11073 | P61587 | P61587 | RND3 | Rho-relate | Color2 [Da | 1 | 4 |
| | 0.145546 | 0.15624 | P61599 | P61599 | NAA20 | N-alpha-ac | Color2 [Da | 1 | 4 |
| + | 1.432606 | 0.606561 | P61601 | P61601 | NCALD | Neurocalci | Color2 [Da | 1 | 4 |
| + | 3.530671 | 2.056973 | P61604 | P61604 | HSPE1 | 10 kDa hea | Color2 [Da | 1 | 4 |
| | 0 | -0.56794 | P61619 | P61619 | SEC61A1 | Protein tra | Color2 [Da | 1 | 4 |
| + | 2.638703 | -1.86006 | P61626 | P61626 | LYZ | Lysozyme | Color2 [Da | 1 | 4 |
| + | 2.203018 | 0.536651 | P61758 | P61758 | VBP1 | Prefoldin s | Color2 [Da | 1 | 4 |
| + | 2.797431 | 0.342157 | P61764 | P61764 | STXBP1 | Syntaxin-b | Color2 [Da | 1 | 4 |
| + | 1.122384 | -0.31448 | P61769 | P61769 | B2M | Beta-2-mic | Color2 [Da | 1 | 4 |
| | 0.371151 | -0.42241 | P61803 | P61803 | DAD1 | Dolichyl-di | Color2 [Da | 1 | 4 |
| + | 4.009538 | 1.90183 | P61916 | P61916 | NPC2 | NPC intrac | Color2 [Da | 1 | 4 |
| + | 4.490742 | -1.4689 | P61923 | P61923 | COPZ1 | Coatomer | Color2 [Da | 1 | 4 |
| + | 3.836186 | 1.422964 | P61925 | P61925 | PKIA | cAMP-dep | Color2 [Da | 1 | 4 |
| + | 2.713288 | 1.323167 | P61956 | P61956 | SUMO2 | Small ubiq | Color2 [Da | 1 | 4 |
| + | 3.119271 | 2.152689 | P61960 | P61960 | UFM1 | Ubiquitin-f | Color2 [Da | 1 | 4 |
| + | 0.872796 | -0.83106 | P61962 | P61962 | DCAF7 | DDB1- and | Color2 [Da | 1 | 4 |
| + | 1.303739 | -0.40114 | P61964 | P61964 | WDR5 | WD repeat | Color2 [Da | 1 | 4 |
| + | 2.55899 | -1.08712 | P61966 | P61966 | AP1S1 | AP-1 comp | Color2 [Da | 1 | 4 |
| + | 1.111953 | 0.314107 | P61970 | P61970 | NUTF2 | Nuclear tra | Color2 [Da | 1 | 4 |
| + | 2.555769 | 0.578965 | P61978 | P61978 | HNRNPK | Heterogen | Color2 [Da | 1 | 4 |
| + | 1.777549 | 0.79462 | P61981 | P61981 | YWHAG | 14-3-3 pro | Color2 [Da | 1 | 4 |
| | 0 | NaN | P62068 | P62068 | USP46 | Ubiquitin c | Color2 [Da | 1 | 4 |
| | 0.124185 | 0.10277 | P62070 | P62070 | RRAS2 | Ras-relate | Color2 [Da | 1 | 4 |
| + | 2.087412 | 0.965348 | P62072 | P62072 | TIMM10 | Mitochonc | Color2 [Da | 1 | 4 |
| + | 3.135509 | 1.6461 | P62081 | P62081 | RPS7 | 40S riboso | Color2 [Da | 1 | 4 |
| + | 1.436771 | -0.69266 | P62136 | P62136 | PPP1CA | Serine/thr | Color2 [Da | 1 | 4 |
| | 0.000915 | -0.00073 | P62140 | P62140 | PPP1CB | Serine/thr | Color2 [Da | 1 | 4 |
| + | 3.094647 | 1.479743 | P62166 | P62166 | NCS1 | Neuronal c | Color2 [Da | 1 | 4 |
| + | 2.35142 | 0.324331 | P62191 | P62191 | PSMC1 | 26S protea | Color2 [Da | 1 | 4 |

| | | | | | | | | | |
|---|---|---|---|---|---|---|---|---|---|
| + | 1.614057 | 0.390572 | P62195 | P62195 | PSMC5 | 26S protea | Color2 [Da | 1 | 4 |
| | 0.422452 | -0.50775 | P62241 | P62241 | RPS8 | 40S riboso | Color2 [Da | 1 | 4 |
| | 0.450677 | -0.24628 | P62244 | P62244 | RPS15A | 40S riboso | Color2 [Da | 1 | 4 |
| | 0.320356 | 0.111022 | P62249 | P62249 | RPS16 | 40S riboso | Color2 [Da | 1 | 4 |
| | 0.671574 | 0.310949 | P62253 | P62253 | UBE2G1 | Ubiquitin-c | Color2 [Da | 1 | 4 |
| + | 2.398706 | 0.69659 | P62256 | P62256 | UBE2H | Ubiquitin-c | Color2 [Da | 1 | 4 |
| + | 3.67199 | 0.782156 | P62258 | P62258 | YWHAE | 14-3-3 pro | Color2 [Da | 1 | 4 |
| | 0.217026 | -0.14084 | P62263 | P62263 | RPS14 | 40S riboso | Color2 [Da | 1 | 4 |
| + | 1.717663 | 1.060354 | P62266 | P62266 | RPS23 | 40S riboso | Color2 [Da | 1 | 4 |
| + | 3.170697 | 0.990394 | P62269 | P62269 | RPS18 | 40S riboso | Color2 [Da | 1 | 4 |
| + | 1.825609 | 1.085522 | P62273 | P62273 | RPS29 | 40S riboso | Color2 [Da | 1 | 4 |
| + | 3.026018 | 1.673046 | P62277 | P62277 | RPS13 | 40S riboso | Color2 [Da | 1 | 4 |
| | 0.034644 | 0.008876 | P62280 | P62280 | RPS11 | 40S riboso | Color2 [Da | 1 | 4 |
| + | 1.880693 | 0.565798 | P62304 | P62304 | SNRPE | Small nucl | Color2 [Da | 1 | 4 |
| + | 1.773676 | 0.430002 | P62306 | P62306 | SNRPF | Small nucl | Color2 [Da | 1 | 4 |
| + | 1.071642 | 1.29657 | P62310 | P62310 | LSM3 | U6 snRNA- | Color2 [Da | 1 | 4 |
| + | 0.938158 | 0.677011 | P62312 | P62312 | LSM6 | U6 snRNA- | Color2 [Da | 1 | 4 |
| + | 2.239665 | 1.525273 | P62314 | P62314 | SNRPD1 | Small nucl | Color2 [Da | 1 | 4 |
| | 0.389183 | 0.108296 | P62316 | P62316 | SNRPD2 | Small nucl | Color2 [Da | 1 | 4 |
| + | 2.906631 | 1.003296 | P62318 | P62318 | SNRPD3 | Small nucl | Color2 [Da | 1 | 4 |
| + | 3.226438 | 2.497869 | P62328 | P62328 | TMSB4X | Thymosin | Color2 [Da | 1 | 4 |
| + | 1.077181 | 0.373195 | P62330 | P62330 | ARF6 | ADP-ribosy | Color2 [Da | 1 | 4 |
| + | 2.253072 | -0.40306 | P62333 | P62333 | PSMC6 | 26S protea | Color2 [Da | 1 | 4 |
| + | 1.439419 | -1.0611 | P62341 | P62341 | SELENOT | Thioredoxi | Color2 [Da | 1 | 4 |
| + | 1.08642 | -0.69849 | P62380 | P62380 | TBPL1 | TATA box-l | Color2 [Da | 1 | 4 |
| + | 1.388277 | 0.339429 | P62424 | P62424 | RPL7A | 60S riboso | Color2 [Da | 1 | 4 |
| | 0.624869 | 0.966852 | P62487 | P62487 | POLR2G | DNA-direc | Color2 [Da | 1 | 4 |
| + | 2.28514 | -0.34405 | P62495 | P62495 | ETF1 | Eukaryotic | Color2 [Da | 1 | 4 |
| + | 3.304982 | 2.519295 | P62633 | P62633 | CNBP | CCHC-type | Color2 [Da | 1 | 4 |
| + | 2.850044 | 0.988039 | P62699 | P62699 | YPEL5 | Protein yip | Color2 [Da | 1 | 4 |
| + | 2.591054 | -1.35766 | P62701 | P62701 | RPS4X | 40S riboso | Color2 [Da | 1 | 4 |
| + | 2.060172 | -0.68067 | P62714;P6 | P62714;P6 | PPP2CB;PF | Serine/thr | Color2 [Da | 1 | 4 |
| + | 3.145098 | -0.56262 | P62736 | P62736 | ACTA2 | Actin, aort | Color2 [Da | 1 | 4 |
| + | 4.117338 | 1.355109 | P62745 | P62745 | RHOB | Rho-relate | Color2 [Da | 1 | 4 |
| | 0.454475 | 0.156247 | P62750 | P62750 | RPL23A | 60S riboso | Color2 [Da | 1 | 4 |
| + | 2.072315 | -0.25572 | P62753 | P62753 | RPS6 | 40S riboso | Color2 [Da | 1 | 4 |
| + | 2.815825 | 2.600964 | P62760 | P62760 | VSNL1 | Visinin-like | Color2 [Da | 1 | 4 |
| + | 1.554637 | 0.458205 | P62805 | P62805 | H4-16 | Histone H4 | Color2 [Da | 1 | 4 |
| + | 1.455329 | -0.61765 | P62820 | P62820 | RAB1A | Ras-relate | Color2 [Da | 1 | 4 |
| + | 3.787573 | -0.55703 | P62826 | P62826 | RAN | GTP-bindir | Color2 [Da | 1 | 4 |
| | 0.62343 | -0.23721 | P62829 | P62829 | RPL23 | 60S riboso | Color2 [Da | 1 | 4 |
| | 0 | 3.032207 | P62834 | P62834 | RAP1A | Ras-relate | Color2 [Da | 1 | 4 |
| + | 3.95866 | 1.410731 | P62841 | P62841 | RPS15 | 40S riboso | Color2 [Da | 1 | 4 |
| | 0.156243 | -0.05719 | P62847 | P62847 | RPS24 | 40S riboso | Color2 [Da | 1 | 4 |
| + | 1.464772 | 0.992786 | P62851 | P62851 | RPS25 | 40S riboso | Color2 [Da | 1 | 4 |
| + | 1.859396 | -0.65753 | P62854 | P62854 | RPS26 | 40S riboso | Color2 [Da | 1 | 4 |
| + | 4.651458 | 1.598619 | P62857 | P62857 | RPS28 | 40S riboso | Color2 [Da | 1 | 4 |

| | | | | | | | | | |
|---|---|---|---|---|---|---|---|---|---|
| + | 1.007181 | 1.564863 | P62861 | P62861 | FAU | FAU ubiqu | Color2 [Da | 1 | 4 |
| | 0.637297 | -0.22932 | P62873 | P62873 | GNB1 | Guanine n | Color2 [Da | 1 | 4 |
| | 0 | NaN | P62875 | P62875 | POLR2L | DNA-direc | Color2 [Da | 1 | 4 |
| + | 1.799131 | 1.175946 | P62877 | P62877 | RBX1 | E3 ubiquiti | Color2 [Da | 1 | 4 |
| | 0.404087 | 0.204353 | P62879 | P62879 | GNB2 | Guanine n | Color2 [Da | 1 | 4 |
| + | 1.930786 | 0.991134 | P62888 | P62888 | RPL30 | 60S riboso | Color2 [Da | 1 | 4 |
| | 0.169476 | 1.1129 | P62891;Q5 | P62891;Q5 | RPL39;RPL | 60S riboso | Color2 [Da | 1 | 4 |
| + | 3.923796 | 1.390033 | P62899 | P62899 | RPL31 | 60S riboso | Color2 [Da | 1 | 4 |
| + | 1.304135 | 0.260675 | P62906 | P62906 | RPL10A | 60S riboso | Color2 [Da | 1 | 4 |
| | 0.140562 | -0.02129 | P62910 | P62910 | RPL32 | 60S riboso | Color2 [Da | 1 | 4 |
| + | 2.453509 | -1.19023 | P62913 | P62913 | RPL11 | 60S riboso | Color2 [Da | 1 | 4 |
| | 1.40384 | -0.05076 | P62917 | P62917 | RPL8 | 60S riboso | Color2 [Da | 1 | 4 |
| + | 3.098199 | 0.860579 | P62937 | P62937 | PPIA | Peptidyl-p | Color2 [Da | 1 | 4 |
| + | 3.555609 | 1.834227 | P62942 | P62942 | FKBP1A | Peptidyl-p | Color2 [Da | 1 | 4 |
| + | 3.244691 | 2.223535 | P62993 | P62993 | GRB2 | Growth fa | Color2 [Da | 1 | 4 |
| + | 1.252122 | 0.656664 | P62995 | P62995 | TRA2B | Transform | Color2 [Da | 1 | 4 |
| | 0.388702 | -0.26969 | P63000 | P63000 | RAC1 | Ras-relate | Color2 [Da | 1 | 4 |
| + | 3.60238 | -0.79904 | P63010 | P63010 | AP2B1 | AP-2 comp | Color2 [Da | 1 | 4 |
| + | 1.44396 | 1.287317 | P63027 | P63027 | VAMP2 | Vesicle-ass | Color2 [Da | 1 | 4 |
| + | 2.549224 | -1.10252 | P63092;Q5 | P63092;Q5 | GNAS | Guanine n | Color2 [Da | 1 | 4 |
| + | 1.320217 | -0.58837 | P63096 | P63096 | GNAI1 | Guanine n | Color2 [Da | 1 | 4 |
| + | 3.14503 | 2.370089 | P63098 | P63098 | PPP3R1 | Calcineurir | Color2 [Da | 1 | 4 |
| | 0.021953 | 0.005986 | P63104 | P63104 | YWHAZ | 14-3-3 pro | Color2 [Da | 1 | 4 |
| | 0.377039 | 0.720011 | P63146 | P63146 | UBE2B | Ubiquitin-c | Color2 [Da | 1 | 4 |
| + | 2.554631 | -0.46818 | P63151 | P63151 | PPP2R2A | Serine/thr | Color2 [Da | 1 | 4 |
| | 0.01024 | -0.00487 | P63165 | P63165 | SUMO1 | Small ubiq | Color2 [Da | 1 | 4 |
| + | 4.519515 | 1.587856 | P63172 | P63172 | DYNLT1 | Dynein ligh | Color2 [Da | 1 | 4 |
| + | 1.201707 | 1.272359 | P63173 | P63173 | RPL38 | 60S riboso | Color2 [Da | 1 | 4 |
| + | 1.985635 | 1.444832 | P63208 | P63208 | SKP1 | S-phase kir | Color2 [Da | 1 | 4 |
| + | 1.576376 | -0.79319 | P63215 | P63215 | GNG3 | Guanine n | Color2 [Da | 1 | 4 |
| | 0.583689 | 1.030827 | P63218 | P63218 | GNG5 | Guanine n | Color2 [Da | 1 | 4 |
| + | 3.647726 | 1.323599 | P63220 | P63220 | RPS21 | 40S riboso | Color2 [Da | 1 | 4 |
| | 0.152014 | 0.077014 | P63241 | P63241 | EIF5A | Eukaryotic | Color2 [Da | 1 | 4 |
| + | 2.242307 | -0.86418 | P63244 | P63244 | RACK1 | Receptor c | Color2 [Da | 1 | 4 |
| + | 4.652874 | 0.92615 | P63261 | P63261 | ACTG1 | Actin, cyto | Color2 [Da | 1 | 4 |
| + | 1.354625 | -0.73581 | P63279 | P63279 | UBE2I | SUMO-cor | Color2 [Da | 1 | 4 |
| + | 0.936039 | 0.431407 | P63302 | P63302 | SELENOW | Selenopro | Color2 [Da | 1 | 4 |
| + | 1.94119 | 3.283644 | P63313 | P63313 | TMSB10 | Thymosin | Color2 [Da | 1 | 4 |
| + | 3.344739 | 1.020867 | P67809 | P67809 | YBX1 | Y-box-bind | Color2 [Da | 1 | 4 |
| + | 1.982228 | 0.692896 | P67812 | P67812 | SEC11A | Signal pep | Color2 [Da | 1 | 4 |
| + | 1.032 | 0.59651 | P67870 | P67870 | CSNK2B | Casein kin | Color2 [Da | 1 | 4 |
| + | 4.33629 | 1.419515 | P67936 | P67936 | TPM4 | Tropomyo | Color2 [Da | 1 | 4 |
| + | 2.532197 | 0.837123 | P68036 | P68036 | UBE2L3 | Ubiquitin-c | Color2 [Da | 1 | 4 |
| + | 4.344545 | -1.23249 | P68104 | P68104 | EEF1A1 | Elongation | Color2 [Da | 1 | 4 |
| | 0 | NaN | P68106 | P68106 | FKBP1B | Peptidyl-p | Color2 [Da | 1 | 4 |
| + | 1.322799 | 0.962522 | P68133 | P68133 | ACTA1 | Actin, alph | Color2 [Da | 1 | 4 |
| | 0.915372 | -0.10061 | P68363 | P68363 | TUBA1B | Tubulin alp | Color2 [Da | 1 | 4 |

| | | | | | | | | | |
|---|---|---|---|---|---|---|---|---|---|
| + | 2.386354 | 2.321394 | P68366 | P68366 | TUBA4A | Tubulin alp | Color2 [Da | 1 | 4 |
| + | 1.753368 | -0.28407 | P68371 | P68371 | TUBB4B | Tubulin be | Color2 [Da | 1 | 4 |
| | 0.332599 | -0.16471 | P68400 | P68400 | CSNK2A1 | Casein kina | Color2 [Da | 1 | 4 |
| + | 4.206499 | 0.956448 | P68402 | P68402 | PAFAH1B2 | Platelet-ac | Color2 [Da | 1 | 4 |
| + | 3.34331 | 0.829715 | P68431 | P68431 | H3C12 | Histone H3 | Color2 [Da | 1 | 4 |
| + | 1.492148 | 0.796024 | P68543 | P68543 | UBXN2A | UBX doma | Color2 [Da | 1 | 4 |
| + | 2.52309 | -2.55156 | P68871 | P68871 | HBB | Hemoglob | Color2 [Da | 1 | 4 |
| + | 1.824044 | -1.70602 | P69905 | P69905 | HBA2 | Hemoglob | Color2 [Da | 1 | 4 |
| | 0.616054 | -0.2651 | P78310 | P78310 | CXADR | Coxsackiev | Color2 [Da | 1 | 4 |
| + | 3.038436 | 1.352577 | P78318 | P78318 | IGBP1 | Immunogl | Color2 [Da | 1 | 4 |
| | 0.528653 | 0.297063 | P78324 | P78324 | SIRPA | Tyrosine-p | Color2 [Da | 1 | 4 |
| + | 2.19193 | 2.156326 | P78330 | P78330 | PSPH | Phosphose | Color2 [Da | 1 | 4 |
| + | 3.479865 | 0.621129 | P78332 | P78332 | RBM6 | RNA-bindi | Color2 [Da | 1 | 4 |
| + | 2.269614 | -0.74518 | P78344 | P78344 | EIF4G2 | Eukaryotic | Color2 [Da | 1 | 4 |
| | 0.12314 | -0.28876 | P78345 | P78345 | RPP38 | Ribonuclea | Color2 [Da | 1 | 4 |
| + | 1.30082 | -0.47691 | P78346 | P78346 | RPP30 | Ribonuclea | Color2 [Da | 1 | 4 |
| | 0.276083 | -0.03593 | P78347 | P78347 | GTF2I | General tra | Color2 [Da | 1 | 4 |
| | 0.387882 | -0.08217 | P78352 | P78352 | DLG4 | Disks large | Color2 [Da | 1 | 4 |
| | 0.39537 | -0.10493 | P78356 | P78356 | PIP4K2B | Phosphatic | Color2 [Da | 1 | 4 |
| + | 1.050048 | -0.61086 | P78357 | P78357 | CNTNAP1 | Contactin- | Color2 [Da | 1 | 4 |
| | 0.932097 | -0.23392 | P78362 | P78362 | SRPK2 | SRSF prote | Color2 [Da | 1 | 4 |
| | 1.082069 | 0.213447 | P78364 | P78364 | PHC1 | Polyhome | Color2 [Da | 1 | 4 |
| | 0 | NaN | P78368 | P78368 | CSNK1G2 | Casein kina | Color2 [Da | 1 | 4 |
| | 0.022142 | -0.00448 | P78371 | P78371 | CCT2 | T-complex | Color2 [Da | 1 | 4 |
| + | 2.170196 | -0.9527 | P78406 | P78406 | RAE1 | mRNA exp | Color2 [Da | 1 | 4 |
| + | 1.55867 | 0.250416 | P78417 | P78417 | GSTO1 | Glutathion | Color2 [Da | 1 | 4 |
| + | 5.418731 | -1.67692 | P78527 | P78527 | PRKDC | DNA-depe | Color2 [Da | 1 | 4 |
| | 0.340908 | 0.555648 | P78536 | P78536 | ADAM17 | Disintegrin | Color2 [Da | 1 | 4 |
| + | 0.952606 | 1.270379 | P78537 | P78537 | BLOC1S1 | Biogenesis | Color2 [Da | 1 | 4 |
| | 0 | NaN | P78539 | P78539 | SRPX | Sushi repe | Color2 [Da | 1 | 4 |
| + | 1.555064 | 1.41474 | P78540 | P78540 | ARG2 | Arginase-2 | Color2 [Da | 1 | 4 |
| + | 4.081297 | 1.538563 | P78559 | P78559 | MAP1A | Microtubu | Color2 [Da | 1 | 4 |
| + | 2.372008 | 1.383132 | P78560 | P78560 | CRADD | Death dom | Color2 [Da | 1 | 4 |
| + | 3.646992 | 0.9539 | P80303 | P80303 | NUCB2 | Nucleobin | Color2 [Da | 1 | 4 |
| | 0 | NaN | P80370 | P80370 | DLK1 | Protein de | Color2 [Da | 1 | 4 |
| + | 1.265232 | 0.234735 | P80404 | P80404 | ABAT | 4-aminobu | Color2 [Da | 1 | 4 |
| + | 2.688516 | 1.021969 | P80723 | P80723 | BASP1 | Brain acid | Color2 [Da | 1 | 4 |
| + | 1.684155 | 1.281874 | P81605 | P81605 | DCD | Dermcidin | Color2 [Da | 1 | 4 |
| + | 2.086877 | -0.71608 | P82094 | P82094 | TMF1 | TATA elem | Color2 [Da | 1 | 4 |
| + | 1.375502 | 0.985261 | P82650 | P82650 | MRPS22 | 28S riboso | Color2 [Da | 1 | 4 |
| | 0.331437 | 0.283718 | P82663 | P82663 | MRPS25 | 28S riboso | Color2 [Da | 1 | 4 |
| + | 1.89391 | 1.663086 | P82664 | P82664 | MRPS10 | 28S riboso | Color2 [Da | 1 | 4 |
| | 0.594572 | -0.16979 | P82673 | P82673 | MRPS35 | 28S riboso | Color2 [Da | 1 | 4 |
| | 0.682955 | -0.22469 | P82675 | P82675 | MRPS5 | 28S riboso | Color2 [Da | 1 | 4 |
| + | 1.678689 | 1.016174 | P82909 | P82909 | MRPS36 | Alpha-keto | Color2 [Da | 1 | 4 |
| + | 0.979369 | 0.620519 | P82912 | P82912 | MRPS11 | 28S riboso | Color2 [Da | 1 | 4 |
| + | 1.631924 | 0.406904 | P82914 | P82914 | MRPS15 | 28S riboso | Color2 [Da | 1 | 4 |

| | | | | | | | | | |
|---|---|---|---|---|---|---|---|---|---|
| + | 3.762962 | -0.49636 | P82921 | P82921 | MRPS21 | 28S riboso | Color2 [Da | 1 | 4 |
| + | 2.730349 | 1.416501 | P82930 | P82930 | MRPS34 | 28S riboso | Color2 [Da | 1 | 4 |
| + | 1.208881 | 0.744136 | P82932 | P82932 | MRPS6 | 28S riboso | Color2 [Da | 1 | 4 |
| | 0.656678 | -0.32531 | P82933 | P82933 | MRPS9 | 28S riboso | Color2 [Da | 1 | 4 |
| + | 1.731963 | 3.405848 | P82970 | P82970 | HMGN5 | High mobi | Color2 [Da | 1 | 4 |
| + | 3.146591 | 2.109628 | P82979 | P82979 | SARNP | SAP domai | Color2 [Da | 1 | 4 |
| + | 0.805365 | 0.691115 | P83111 | P83111 | LACTB | Serine bet | Color2 [Da | 1 | 4 |
| + | 1.409119 | -0.65384 | P83436 | P83436 | COG7 | Conserved | Color2 [Da | 1 | 4 |
| + | 0.913452 | 1.177798 | P83731 | P83731 | RPL24 | 60S riboso | Color2 [Da | 1 | 4 |
| + | 2.224524 | 1.864274 | P83876 | P83876 | TXNL4A | Thioredoxi | Color2 [Da | 1 | 4 |
| | 0.475384 | 0.412058 | P83881;Q9 | P83881;Q9 | RPL36A;RP | 60S riboso | Color2 [Da | 1 | 4 |
| + | 2.340007 | 0.836188 | P83916 | P83916 | CBX1 | Chromobo | Color2 [Da | 1 | 4 |
| + | 2.683738 | -0.9417 | P84022 | P84022 | SMAD3 | Mothers a | Color2 [Da | 1 | 4 |
| + | 1.152645 | 2.495543 | P84074 | P84074 | HPCA | Neuron-sp | Color2 [Da | 1 | 4 |
| + | 2.249884 | 0.178352 | P84077 | P84077 | ARF1 | ADP-ribosy | Color2 [Da | 1 | 4 |
| | 0.593341 | -0.33198 | P84085 | P84085 | ARF5 | ADP-ribosy | Color2 [Da | 1 | 4 |
| + | 0.832943 | 0.459823 | P84090 | P84090 | ERH | Enhancer c | Color2 [Da | 1 | 4 |
| | 0.455449 | -0.66117 | P84095 | P84095 | RHOG | Rho-relate | Color2 [Da | 1 | 4 |
| | 0.852792 | 0.212431 | P84098 | P84098 | RPL19 | 60S riboso | Color2 [Da | 1 | 4 |
| + | 1.792443 | 1.915815 | P84103 | P84103 | SRSF3 | Serine/arg | Color2 [Da | 1 | 4 |
| + | 1.903958 | 1.325944 | P84157 | P84157 | MXRA7 | Matrix-ren | Color2 [Da | 1 | 4 |
| + | 2.460811 | 3.233704 | P84243 | P84243 | H3-3B | Histone H3 | Color2 [Da | 1 | 4 |
| + | 1.13228 | 0.277189 | P85037 | P85037 | FOXK1 | Forkhead b | Color2 [Da | 1 | 4 |
| + | 1.921044 | 2.467528 | P86397 | P86397 | HTD2 | Hydroxyac | Color2 [Da | 1 | 4 |
| + | 2.035862 | -1.04123 | P86790;P8 | P86790;P8 | CCZ1B;CCZ | Vacuolar fu | Color2 [Da | 1 | 4 |
| | 0.220228 | -0.10743 | P98077 | P98077 | SHC2 | SHC-transf | Color2 [Da | 1 | 4 |
| | 0 | NaN | P98095 | P98095 | FBLN2 | Fibulin-2 | Color2 [Da | 1 | 4 |
| | 0 | NaN | P98155 | P98155 | VLDLR | Very low-c | Color2 [Da | 1 | 4 |
| | 0 | NaN | P98160 | P98160 | HSPG2 | Basement | Color2 [Da | 1 | 4 |
| + | 1.913886 | -0.62891 | P98164 | P98164 | LRP2 | Low-densi | Color2 [Da | 1 | 4 |
| + | 2.318954 | -0.98548 | P98170 | P98170 | XIAP | E3 ubiquiti | Color2 [Da | 1 | 4 |
| | 0.136103 | -0.07085 | P98172 | P98172 | EFNB1 | Ephrin-B1 | Color2 [Da | 1 | 4 |
| | 0.392055 | 0.595858 | P98174 | P98174 | FGD1 | FYVE, RhoG | Color2 [Da | 1 | 4 |
| + | 2.48821 | 0.56596 | P98175 | P98175 | RBM10 | RNA-bindi | Color2 [Da | 1 | 4 |
| | 0.522624 | 0.352586 | P98179 | P98179 | RBM3 | RNA-bindi | Color2 [Da | 1 | 4 |
| + | 2.352022 | -1.38675 | P98194 | P98194 | ATP2C1 | Calcium-tr | Color2 [Da | 1 | 4 |
| + | 3.870042 | 2.789243 | P99999 | P99999 | CYCS | Cytochrom | Color2 [Da | 1 | 4 |
| + | 1.971888 | 0.950543 | Q00013 | Q00013 | MPP1 | 55 kDa ery | Color2 [Da | 1 | 4 |
| + | 3.609313 | 0.987713 | Q00059 | Q00059 | TFAM | Transcripti | Color2 [Da | 1 | 4 |
| + | 2.279103 | 0.615489 | Q00169 | Q00169 | PITPNA | Phosphatic | Color2 [Da | 1 | 4 |
| + | 3.270275 | -1.14454 | Q00325 | Q00325 | SLC25A3 | Phosphate | Color2 [Da | 1 | 4 |
| + | 2.471631 | -0.75374 | Q00341 | Q00341 | HDLBP | Vigilin | Color2 [Da | 1 | 4 |
| | 0.314162 | -0.02224 | Q00403 | Q00403 | GTF2B | Transcripti | Color2 [Da | 1 | 4 |
| | 0 | -1.92741 | Q00534 | Q00534 | CDK6 | Cyclin-dep | Color2 [Da | 1 | 4 |
| + | 3.293587 | -1.33521 | Q00535 | Q00535 | CDK5 | Cyclin-dep | Color2 [Da | 1 | 4 |
| + | 2.939804 | -1.07538 | Q00536 | Q00536 | CDK16 | Cyclin-dep | Color2 [Da | 1 | 4 |
| | 0 | NaN | Q00537 | Q00537 | CDK17 | Cyclin-dep | Color2 [Da | 1 | 4 |

-Bruker Confidential-

| | | | | | | | | | |
|---|---|---|---|---|---|---|---|---|---|
| + | 2.207811 | 1.651844 | Q00577 | Q00577 | PURA | Transcripti | Color2 [Da | 1 | 4 |
| + | 1.968457 | 3.286388 | Q00587 | Q00587 | CDC42EP1 | Cdc42 effe | Color2 [Da | 1 | 4 |
| + | 5.038809 | -1.30766 | Q00610 | Q00610 | CLTC | Clathrin he | Color2 [Da | 1 | 4 |
| | 0.947815 | -0.27917 | Q00613 | Q00613 | HSF1 | Heat shock | Color2 [Da | 1 | 4 |
| | 0 | NaN | Q00653 | Q00653 | NFKB2 | Nuclear fa | Color2 [Da | 1 | 4 |
| + | 2.925099 | 1.397929 | Q00688 | Q00688 | FKBP3 | Peptidyl-p | Color2 [Da | 1 | 4 |
| + | 1.249582 | 1.153143 | Q00765 | Q00765 | REEP5 | Receptor e | Color2 [Da | 1 | 4 |
| + | 1.33649 | -0.89395 | Q00796 | Q00796 | SORD | Sorbitol de | Color2 [Da | 1 | 4 |
| | 0.589995 | -0.24889 | Q00839 | Q00839 | HNRNPU | Heterogen | Color2 [Da | 1 | 4 |
| | 0.922243 | -0.24575 | Q01081 | Q01081 | U2AF1 | Splicing fa | Color2 [Da | 1 | 4 |
| | 1.691345 | -0.08331 | Q01082 | Q01082 | SPTBN1 | Spectrin b | Color2 [Da | 1 | 4 |
| | 0.216413 | 0.115704 | Q01085 | Q01085 | TIAL1 | Nucleolysi | Color2 [Da | 1 | 4 |
| | 0.797532 | 0.339163 | Q01105 | Q01105 | SET | Protein SE | Color2 [Da | 1 | 4 |
| + | 3.465282 | 1.665795 | Q01130 | Q01130 | SRSF2 | Serine/arg | Color2 [Da | 1 | 4 |
| | 0.624312 | 0.486558 | Q01167 | Q01167 | FOXK2 | Forkhead I | Color2 [Da | 1 | 4 |
| | 0 | NaN | Q01415 | Q01415 | GALK2 | N-acetylga | Color2 [Da | 1 | 4 |
| + | 1.019274 | -0.38155 | Q01433 | Q01433 | AMPD2 | AMP dear | Color2 [Da | 1 | 4 |
| | 0.166558 | 0.094931 | Q01459 | Q01459 | CTBS | Di-N-acety | Color2 [Da | 1 | 4 |
| + | 2.614044 | 1.75198 | Q01469 | Q01469 | FABP5 | Fatty acid- | Color2 [Da | 1 | 4 |
| + | 3.059509 | 1.01817 | Q01484 | Q01484 | ANK2 | Ankyrin-2 | Color2 [Da | 1 | 4 |
| + | 1.669905 | 0.339503 | Q01518 | Q01518 | CAP1 | Adenylyl c | Color2 [Da | 1 | 4 |
| | 0.332189 | 0.374671 | Q01538 | Q01538 | MYT1 | Myelin tra | Color2 [Da | 1 | 4 |
| | 0 | NaN | Q01546 | Q01546 | KRT76 | Keratin, ty | Color2 [Da | 1 | 4 |
| + | 2.246447 | -0.65941 | Q01581 | Q01581 | HMGCS1 | Hydroxym | Color2 [Da | 1 | 4 |
| + | 1.298572 | -0.50568 | Q01650 | Q01650 | SLC7A5 | Large neut | Color2 [Da | 1 | 4 |
| + | 1.328871 | 1.920924 | Q01658 | Q01658 | DR1 | Protein Dr | Color2 [Da | 1 | 4 |
| + | 1.131472 | 0.896409 | Q01664 | Q01664 | TFAP4 | Transcripti | Color2 [Da | 1 | 4 |
| | 0.453952 | 0.223634 | Q01780 | Q01780 | EXOSC10 | Exosome c | Color2 [Da | 1 | 4 |
| + | 4.234806 | -1.6957 | Q01813 | Q01813 | PFKP | ATP-deper | Color2 [Da | 1 | 4 |
| | 0.307436 | 0.253385 | Q01814 | Q01814 | ATP2B2 | Plasma me | Color2 [Da | 1 | 4 |
| + | 1.123785 | -0.45422 | Q01831 | Q01831 | XPC | DNA repai | Color2 [Da | 1 | 4 |
| + | 3.819754 | 1.611588 | Q01844 | Q01844 | EWSR1 | RNA-bindi | Color2 [Da | 1 | 4 |
| | 0.722325 | -0.51913 | Q01850 | Q01850 | CDR2 | Cerebellar | Color2 [Da | 1 | 4 |
| | 0.205495 | -0.24534 | Q01968 | Q01968 | OCRL | Inositol po | Color2 [Da | 1 | 4 |
| | 0.307259 | -0.17822 | Q01970 | Q01970 | PLCB3 | 1-phospha | Color2 [Da | 1 | 4 |
| | 0 | NaN | Q01995 | Q01995 | TAGLN | Transgelin | Color2 [Da | 1 | 4 |
| | 0 | -1.4711 | Q02040 | Q02040 | AKAP17A | A-kinase a | Color2 [Da | 1 | 4 |
| | 0.582027 | -0.42408 | Q02086 | Q02086 | SP2 | Transcripti | Color2 [Da | 1 | 4 |
| | 0 | 1.007465 | Q02108 | Q02108 | GUCY1A1 | Guanylate | Color2 [Da | 1 | 4 |
| + | 4.105778 | 2.071008 | Q02127 | Q02127 | DHODH | Dihydroor | Color2 [Da | 1 | 4 |
| | 1.124359 | -0.24682 | Q02153 | Q02153 | GUCY1B1 | Guanylate | Color2 [Da | 1 | 4 |
| + | 2.003331 | 1.206665 | Q02156 | Q02156 | PRKCE | Protein kir | Color2 [Da | 1 | 4 |
| + | 3.741809 | -1.69656 | Q02218 | Q02218 | OGDH | 2-oxogluta | Color2 [Da | 1 | 4 |
| + | 2.365272 | -1.00865 | Q02241 | Q02241 | KIF23 | Kinesin-lik | Color2 [Da | 1 | 4 |
| + | 1.473728 | -0.55262 | Q02246 | Q02246 | CNTN2 | Contactin- | Color2 [Da | 1 | 4 |
| + | 2.552599 | 0.316818 | Q02252 | Q02252 | ALDH6A1 | Methylma | Color2 [Da | 1 | 4 |
| | 0.070993 | 0.047751 | Q02410 | Q02410 | APBA1 | Amyloid-b | Color2 [Da | 1 | 4 |

| | | | | | | | | | |
|---|---|---|---|---|---|---|---|---|---|
| + | 2.343609 | 1.157267 | Q02413 | Q02413 | DSG1 | Desmoglei | Color2 [Da | 1 | 4 |
| | 0.182489 | -0.51976 | Q02539 | Q02539 | H1-1 | Histone H1 | Color2 [Da | 1 | 4 |
| + | 2.198806 | -1.89355 | Q02543 | Q02543 | RPL18A | 60S riboso | Color2 [Da | 1 | 4 |
| | 0.060291 | 0.111003 | Q02641 | Q02641 | CACNB1 | Voltage-de | Color2 [Da | 1 | 4 |
| + | 1.084031 | -0.45081 | Q02750 | Q02750 | MAP2K1 | Dual speci | Color2 [Da | 1 | 4 |
| | 0.657214 | -0.11482 | Q02790 | Q02790 | FKBP4 | Peptidyl-pı | Color2 [Da | 1 | 4 |
| + | 4.385059 | -1.6952 | Q02809 | Q02809 | PLOD1 | Procollage | Color2 [Da | 1 | 4 |
| + | 3.320243 | 1.110654 | Q02818 | Q02818 | NUCB1 | Nucleobin | Color2 [Da | 1 | 4 |
| + | 3.211832 | 0.930471 | Q02878 | Q02878 | RPL6 | 60S riboso | Color2 [Da | 1 | 4 |
| + | 3.104322 | -0.90517 | Q02880 | Q02880 | TOP2B | DNA topoi | Color2 [Da | 1 | 4 |
| + | 4.480255 | 2.335861 | Q02952 | Q02952 | AKAP12 | A-kinase a | Color2 [Da | 1 | 4 |
| + | 1.714382 | 0.821541 | Q02978 | Q02978 | SLC25A11 | Mitochonc | Color2 [Da | 1 | 4 |
| + | 1.645047 | -0.37867 | Q03001 | Q03001 | DST | Dystonin | Color2 [Da | 1 | 4 |
| | 0 | -0.55951 | Q03135 | Q03135 | CAV1 | Caveolin-1 | Color2 [Da | 1 | 4 |
| + | 1.035532 | 0.615125 | Q03154 | Q03154 | ACY1 | Aminoacyl | Color2 [Da | 1 | 4 |
| + | 1.897877 | -0.93093 | Q03164 | Q03164 | KMT2A | Histone-ly | Color2 [Da | 1 | 4 |
| + | 0.727285 | -0.66222 | Q03181 | Q03181 | PPARD | Peroxisom | Color2 [Da | 1 | 4 |
| | 0 | 0.947894 | Q03188 | Q03188 | CENPC | Centromei | Color2 [Da | 1 | 4 |
| + | 2.19237 | 0.580536 | Q03252 | Q03252 | LMNB2 | Lamin-B2 | Color2 [Da | 1 | 4 |
| + | 1.462857 | 2.695726 | Q03393 | Q03393 | PTS | 6-pyruvoyl | Color2 [Da | 1 | 4 |
| + | 0.759632 | -0.62356 | Q03426 | Q03426 | MVK | Mevalonat | Color2 [Da | 1 | 4 |
| | 0.466323 | -0.19221 | Q03701 | Q03701 | CEBPZ | CCAAT/enl | Color2 [Da | 1 | 4 |
| + | 3.404081 | -3.63259 | Q04118 | Q04118 | PRB3 | Basic saliva | Color2 [Da | 1 | 4 |
| + | 0.828175 | 0.435532 | Q04206 | Q04206 | RELA | Transcripti | Color2 [Da | 1 | 4 |
| + | 1.704618 | 0.844603 | Q04323 | Q04323 | UBXN1 | UBX doma | Color2 [Da | 1 | 4 |
| + | 1.942186 | -1.16604 | Q04446 | Q04446 | GBE1 | 1,4-alpha-g | Color2 [Da | 1 | 4 |
| + | 2.852206 | -0.73667 | Q04637 | Q04637 | EIF4G1 | Eukaryotic | Color2 [Da | 1 | 4 |
| + | 1.564964 | 0.88127 | Q04695 | Q04695 | KRT17 | Keratin, ty | Color2 [Da | 1 | 4 |
| + | 0.84144 | 0.93893 | Q04721 | Q04721 | NOTCH2 | Neurogeni | Color2 [Da | 1 | 4 |
| + | 2.265527 | -0.88738 | Q04724 | Q04724 | TLE1 | Transducir | Color2 [Da | 1 | 4 |
| + | 2.944651 | 1.005044 | Q04726 | Q04726 | TLE3 | Transducir | Color2 [Da | 1 | 4 |
| + | 3.330948 | 0.58562 | Q04760 | Q04760 | GLO1 | Lactoylglut | Color2 [Da | 1 | 4 |
| + | 3.868187 | 1.162597 | Q04837 | Q04837 | SSBP1 | Single-stra | Color2 [Da | 1 | 4 |
| + | 2.209095 | 0.649051 | Q04917 | Q04917 | YWHAH | 14-3-3 pro | Color2 [Da | 1 | 4 |
| | 0 | -0.59279 | Q05048 | Q05048 | CSTF1 | Cleavage s | Color2 [Da | 1 | 4 |
| + | 1.784466 | -0.98666 | Q05086 | Q05086 | UBE3A | Ubiquitin-p | Color2 [Da | 1 | 4 |
| + | 2.821137 | 0.953362 | Q05193 | Q05193 | DNM1 | Dynamin-1 | Color2 [Da | 1 | 4 |
| + | 1.992152 | 1.414607 | Q05209 | Q05209 | PTPN12 | Tyrosine-p | Color2 [Da | 1 | 4 |
| | 0 | NaN | Q05329 | Q05329 | GAD2 | Glutamate | Color2 [Da | 1 | 4 |
| + | 3.917811 | 1.645061 | Q05397 | Q05397 | PTK2 | Focal adhe | Color2 [Da | 1 | 4 |
| | 0 | -1.23284 | Q05516 | Q05516 | ZBTB16 | Zinc finger | Color2 [Da | 1 | 4 |
| | 0.657799 | 0.273269 | Q05519 | Q05519 | SRSF11 | Serine/arg | Color2 [Da | 1 | 4 |
| + | 2.417866 | -0.46047 | Q05639 | Q05639 | EEF1A2 | Elongation | Color2 [Da | 1 | 4 |
| + | 3.218024 | 1.838939 | Q05682 | Q05682 | CALD1 | Caldesmor | Color2 [Da | 1 | 4 |
| + | 0.717955 | 0.739072 | Q05823 | Q05823 | RNASEL | 2-5A-depe | Color2 [Da | 1 | 4 |
| | 0 | -0.17127 | Q05D32 | Q05D32 | CTDSPL2 | CTD small | Color2 [Da | 1 | 4 |
| + | 1.249929 | -0.46387 | Q06124 | Q06124 | PTPN11 | Tyrosine-p | Color2 [Da | 1 | 4 |

| | | | | | | | | | |
|---|---|---|---|---|---|---|---|---|---|
| + | 1.505488 | -0.98406 | Q06136 | Q06136 | KDSR | 3-ketodihy | Color2 [Da | 1 | 4 |
| | 0 | NaN | Q06190 | Q06190 | PPP2R3A | Serine/thr | Color2 [Da | 1 | 4 |
| + | 0.860034 | -0.40815 | Q06203 | Q06203 | PPAT | Amidopho | Color2 [Da | 1 | 4 |
| + | 2.623976 | -0.53767 | Q06210 | Q06210 | GFPT1 | Glutamine | Color2 [Da | 1 | 4 |
| + | 0.919861 | -0.71546 | Q06265 | Q06265 | EXOSC9 | Exosome c | Color2 [Da | 1 | 4 |
| | 0 | NaN | Q06278 | Q06278 | AOX1 | Aldehyde c | Color2 [Da | 1 | 4 |
| + | 1.651955 | 0.804762 | Q06323 | Q06323 | PSME1 | Proteasom | Color2 [Da | 1 | 4 |
| + | 2.394361 | -0.88732 | Q06330 | Q06330 | RBPJ | Recombini | Color2 [Da | 1 | 4 |
| | 0.091971 | 0.039545 | Q06455 | Q06455 | RUNX1T1 | Protein CB | Color2 [Da | 1 | 4 |
| + | 2.889862 | 0.868299 | Q06481 | Q06481 | APLP2 | Amyloid be | Color2 [Da | 1 | 4 |
| + | 3.137796 | -0.60986 | Q06546 | Q06546 | GABPA | GA-binding | Color2 [Da | 1 | 4 |
| | 0.495672 | 0.770908 | Q06547 | Q06547 | GABPB1 | GA-binding | Color2 [Da | 1 | 4 |
| | 0.109932 | -0.26258 | Q06587 | Q06587 | RING1 | E3 ubiquiti | Color2 [Da | 1 | 4 |
| + | 1.852624 | -0.61079 | Q06787 | Q06787 | FMR1 | Synaptic fu | Color2 [Da | 1 | 4 |
| + | 2.214534 | 0.680658 | Q06830 | Q06830 | PRDX1 | Peroxiredo | Color2 [Da | 1 | 4 |
| | 0.427277 | -0.27348 | Q07020 | Q07020 | RPL18 | 60S riboso | Color2 [Da | 1 | 4 |
| + | 3.233836 | 1.365181 | Q07021 | Q07021 | C1QBP | Compleme | Color2 [Da | 1 | 4 |
| + | 3.013919 | -0.62769 | Q07065 | Q07065 | CKAP4 | Cytoskelet | Color2 [Da | 1 | 4 |
| | 0.842642 | 0.140932 | Q07157 | Q07157 | TJP1 | Tight junct | Color2 [Da | 1 | 4 |
| | 0.632274 | -0.55811 | Q07343 | Q07343 | PDE4B | cAMP-spec | Color2 [Da | 1 | 4 |
| | 0.528445 | 0.376331 | Q07352 | Q07352 | ZFP36L1 | mRNA dec | Color2 [Da | 1 | 4 |
| + | 1.144391 | 0.495393 | Q07666 | Q07666 | KHDRBS1 | KH domair | Color2 [Da | 1 | 4 |
| + | 2.605871 | -1.58856 | Q07812 | Q07812 | BAX | Apoptosis | Color2 [Da | 1 | 4 |
| + | 2.859732 | -0.67188 | Q07866 | Q07866 | KLC1 | Kinesin ligl | Color2 [Da | 1 | 4 |
| | 0 | -1.08205 | Q07889 | Q07889 | SOS1 | Son of sev | Color2 [Da | 1 | 4 |
| + | 2.357274 | -0.73911 | Q07954 | Q07954 | LRP1 | Prolow-de | Color2 [Da | 1 | 4 |
| + | 3.21946 | 0.903008 | Q07955 | Q07955 | SRSF1 | Serine/arg | Color2 [Da | 1 | 4 |
| + | 4.672838 | 0.903534 | Q07960 | Q07960 | ARHGAP1 | Rho GTPas | Color2 [Da | 1 | 4 |
| + | 0.983721 | 1.287782 | Q08117 | Q08117 | TLE5 | TLE family | Color2 [Da | 1 | 4 |
| | 0.834709 | 0.207531 | Q08170 | Q08170 | SRSF4 | Serine/arg | Color2 [Da | 1 | 4 |
| | 0.379554 | 0.217312 | Q08174 | Q08174 | PCDH1 | Protocadh | Color2 [Da | 1 | 4 |
| + | 2.399939 | 1.339814 | Q08188 | Q08188 | TGM3 | Protein-glu | Color2 [Da | 1 | 4 |
| + | 2.064861 | 0.500507 | Q08209 | Q08209 | PPP3CA | Protein ph | Color2 [Da | 1 | 4 |
| + | 3.83721 | -1.1734 | Q08211 | Q08211 | DHX9 | ATP-deper | Color2 [Da | 1 | 4 |
| + | 1.090509 | 3.848514 | Q08257 | Q08257 | CRYZ | Quinone o | Color2 [Da | 1 | 4 |
| + | 2.051422 | -1.97326 | Q08345 | Q08345 | DDR1 | Epithelial c | Color2 [Da | 1 | 4 |
| + | 1.626085 | 0.906885 | Q08357 | Q08357 | SLC20A2 | Sodium-de | Color2 [Da | 1 | 4 |
| + | 2.037885 | -0.24922 | Q08378 | Q08378 | GOLGA3 | Golgin sub | Color2 [Da | 1 | 4 |
| + | 4.640059 | -0.57132 | Q08379 | Q08379 | GOLGA2 | Golgin sub | Color2 [Da | 1 | 4 |
| | 1.113618 | 0.229721 | Q08380 | Q08380 | LGALS3BP | Galectin-3 | Color2 [Da | 1 | 4 |
| + | 3.495013 | -0.97988 | Q08431 | Q08431 | MFGE8 | Lactadheri | Color2 [Da | 1 | 4 |
| | 0 | NaN | Q08462 | Q08462 | ADCY2 | Adenylate | Color2 [Da | 1 | 4 |
| + | 2.61446 | 1.626857 | Q08495 | Q08495 | DMTN | Dematin | Color2 [Da | 1 | 4 |
| + | 3.096774 | 1.180025 | Q08554 | Q08554 | DSC1 | Desmocoll | Color2 [Da | 1 | 4 |
| + | 0.910512 | 1.947914 | Q08623 | Q08623 | PUDP | Pseudouri | Color2 [Da | 1 | 4 |
| | 0 | 1.353225 | Q08629 | Q08629 | SPOCK1 | Testican-1 | Color2 [Da | 1 | 4 |
| + | 2.137219 | -2.13827 | Q08722 | Q08722 | CD47 | Leukocyte | Color2 [Da | 1 | 4 |

| | | | | | | | | | |
|---|---|---|---|---|---|---|---|---|---|
| + | 1.701782 | -0.38629 | Q08752 | Q08752 | PPID | Peptidyl-pı | Color2 [Da | 1 | 4 |
| + | 3.102509 | -0.75176 | Q08945 | Q08945 | SSRP1 | FACT comp | Color2 [Da | 1 | 4 |
| | 0.560687 | -0.58924 | Q08999 | Q08999 | RBL2 | Retinoblas | Color2 [Da | 1 | 4 |
| + | 1.933253 | 0.410995 | Q08AD1 | Q08AD1 | CAMSAP2 | Calmodulir | Color2 [Da | 1 | 4 |
| + | 1.090825 | 0.396038 | Q08AE8 | Q08AE8 | SPIRE1 | Protein spi | Color2 [Da | 1 | 4 |
| + | 0.90873 | 1.068461 | Q08AI8 | Q08AI8 | MAB21L4 | Protein ma | Color2 [Da | 1 | 4 |
| + | 3.729223 | -1.11297 | Q08AM6 | Q08AM6 | VAC14 | Protein VA | Color2 [Da | 1 | 4 |
| | 0.19203 | -0.06729 | Q08J23 | Q08J23 | NSUN2 | RNA cytosi | Color2 [Da | 1 | 4 |
| | 0 | NaN | Q09019 | Q09019 | DMWD | Dystrophia | Color2 [Da | 1 | 4 |
| + | 1.409271 | -0.70805 | Q09028 | Q09028 | RBBP4 | Histone-bi | Color2 [Da | 1 | 4 |
| + | 3.893987 | -2.20455 | Q09161 | Q09161 | NCBP1 | Nuclear ca | Color2 [Da | 1 | 4 |
| | 0.267044 | 0.154375 | Q09472 | Q09472 | EP300 | Histone ac | Color2 [Da | 1 | 4 |
| + | 3.294493 | 1.125709 | Q09666 | Q09666 | AHNAK | Neuroblas | Color2 [Da | 1 | 4 |
| + | 1.587283 | -0.37081 | Q0VDF9 | Q0VDF9 | HSPA14 | Heat shocl | Color2 [Da | 1 | 4 |
| + | 2.573841 | -0.9595 | Q0VDG4 | Q0VDG4 | SCRN3 | Secernin-3 | Color2 [Da | 1 | 4 |
| + | 0.729917 | 0.856255 | Q0VGL1 | Q0VGL1 | LAMTOR4 | Ragulator | Color2 [Da | 1 | 4 |
| + | 1.325559 | -0.69336 | Q10471 | Q10471 | GALNT2 | Polypeptic | Color2 [Da | 1 | 4 |
| | 0.37554 | -0.10248 | Q10567 | Q10567 | AP1B1 | AP-1 comp | Color2 [Da | 1 | 4 |
| + | 3.792962 | -1.42028 | Q10570 | Q10570 | CPSF1 | Cleavage a | Color2 [Da | 1 | 4 |
| + | 2.305832 | -0.59733 | Q10713 | Q10713 | PMPCA | Mitochonc | Color2 [Da | 1 | 4 |
| + | 1.237134 | -0.23659 | Q12756 | Q12756 | KIF1A | Kinesin-lik | Color2 [Da | 1 | 4 |
| + | 2.346133 | -0.48313 | Q12765 | Q12765 | SCRN1 | Secernin-1 | Color2 [Da | 1 | 4 |
| + | 3.896463 | -1.36674 | Q12768 | Q12768 | WASHC5 | WASH com | Color2 [Da | 1 | 4 |
| + | 1.496141 | -1.10106 | Q12769 | Q12769 | NUP160 | Nuclear pc | Color2 [Da | 1 | 4 |
| + | 1.151695 | 0.373132 | Q12770 | Q12770 | SCAP | Sterol regu | Color2 [Da | 1 | 4 |
| | 0 | -1.82352 | Q12788 | Q12788 | TBL3 | Transducir | Color2 [Da | 1 | 4 |
| | 0.054168 | -0.02378 | Q12789 | Q12789 | GTF3C1 | General tra | Color2 [Da | 1 | 4 |
| + | 2.069423 | 0.182614 | Q12792 | Q12792 | TWF1 | Twinfilin-1 | Color2 [Da | 1 | 4 |
| + | 2.409272 | -0.60917 | Q12797 | Q12797 | ASPH | Aspartyl/a | Color2 [Da | 1 | 4 |
| + | 0.866977 | -0.41614 | Q12800 | Q12800 | TFCP2 | Alpha-glob | Color2 [Da | 1 | 4 |
| + | 3.502555 | -2.07915 | Q12802 | Q12802 | AKAP13 | A-kinase a | Color2 [Da | 1 | 4 |
| + | 2.155323 | 0.476474 | Q12824 | Q12824 | SMARCB1 | SWI/SNF-r | Color2 [Da | 1 | 4 |
| + | 4.303191 | -1.25862 | Q12830 | Q12830 | BPTF | Nucleosom | Color2 [Da | 1 | 4 |
| | 0.289672 | -0.10978 | Q12840 | Q12840 | KIF5A | Kinesin he | Color2 [Da | 1 | 4 |
| + | 3.270211 | 0.94849 | Q12841 | Q12841 | FSTL1 | Follistatin- | Color2 [Da | 1 | 4 |
| + | 1.350884 | -0.8582 | Q12846 | Q12846 | STX4 | Syntaxin-4 | Color2 [Da | 1 | 4 |
| + | 1.832685 | 0.385847 | Q12849 | Q12849 | GRSF1 | G-rich seq | Color2 [Da | 1 | 4 |
| | 0 | -0.09047 | Q12857 | Q12857 | NFIA | Nuclear fa | Color2 [Da | 1 | 4 |
| + | 2.996313 | 0.675645 | Q12860 | Q12860 | CNTN1 | Contactin- | Color2 [Da | 1 | 4 |
| + | 1.101185 | -1.14532 | Q12872 | Q12872 | SFSWAP | Splicing fa | Color2 [Da | 1 | 4 |
| | 1.091033 | -0.25211 | Q12873 | Q12873 | CHD3 | Chromodo | Color2 [Da | 1 | 4 |
| | 0.497689 | -0.46552 | Q12874 | Q12874 | SF3A3 | Splicing fa | Color2 [Da | 1 | 4 |
| + | 1.53863 | 0.250001 | Q12888 | Q12888 | TP53BP1 | TP53-bindi | Color2 [Da | 1 | 4 |
| | 0.179906 | 0.139578 | Q12904 | Q12904 | AIMP1 | Aminoacyl | Color2 [Da | 1 | 4 |
| + | 1.977585 | -0.6032 | Q12905 | Q12905 | ILF2 | Interleukir | Color2 [Da | 1 | 4 |
| + | 1.373356 | -0.2734 | Q12906 | Q12906 | ILF3 | Interleukir | Color2 [Da | 1 | 4 |
| + | 2.835176 | -1.67609 | Q12907 | Q12907 | LMAN2 | Vesicular i | Color2 [Da | 1 | 4 |

| | | | | | | | | | |
|---|---|---|---|---|---|---|---|---|---|
| | 0.780217 | -0.33879 | Q12913 | Q12913 | PTPRJ | Receptor-t | Color2 [Da | 1 | 4 |
| + | 2.446353 | 1.82387 | Q12926 | Q12926 | ELAVL2 | ELAV-like p | Color2 [Da | 1 | 4 |
| | 0.387367 | 0.078559 | Q12929 | Q12929 | EPS8 | Epidermal | Color2 [Da | 1 | 4 |
| + | 2.480511 | -0.4201 | Q12931 | Q12931 | TRAP1 | Heat shock | Color2 [Da | 1 | 4 |
| + | 3.611032 | -1.47984 | Q12933 | Q12933 | TRAF2 | TNF recept | Color2 [Da | 1 | 4 |
| | 1.280692 | -0.17963 | Q12955 | Q12955 | ANK3 | Ankyrin-3 | Color2 [Da | 1 | 4 |
| | 0.343822 | 0.206183 | Q12959 | Q12959 | DLG1 | Disks large | Color2 [Da | 1 | 4 |
| + | 1.92968 | 2.715086 | Q12962 | Q12962 | TAF10 | Transcripti | Color2 [Da | 1 | 4 |
| | 0 | -0.45734 | Q12965 | Q12965 | MYO1E | Unconvent | Color2 [Da | 1 | 4 |
| + | 2.026286 | 0.767284 | Q12972 | Q12972 | PPP1R8 | Nuclear inl | Color2 [Da | 1 | 4 |
| | 0.168666 | -0.06027 | Q12979 | Q12979 | ABR | Active brea | Color2 [Da | 1 | 4 |
| + | 2.552356 | 1.722061 | Q12980 | Q12980 | NPRL3 | GATOR cor | Color2 [Da | 1 | 4 |
| + | 0.814958 | 1.590986 | Q12983 | Q12983 | BNIP3 | BCL2/ader | Color2 [Da | 1 | 4 |
| + | 1.109694 | -0.2777 | Q12996 | Q12996 | CSTF3 | Cleavage s | Color2 [Da | 1 | 4 |
| + | 3.606299 | 1.314223 | Q13011 | Q13011 | ECH1 | Delta(3,5)- | Color2 [Da | 1 | 4 |
| + | 2.489774 | -1.09691 | Q13015 | Q13015 | MLLT11 | Protein AF | Color2 [Da | 1 | 4 |
| + | 2.29556 | -0.73558 | Q13017 | Q13017 | ARHGAP5 | Rho GTPas | Color2 [Da | 1 | 4 |
| + | 1.782177 | 0.849106 | Q13033 | Q13033 | STRN3 | Striatin-3 | Color2 [Da | 1 | 4 |
| | 0 | -1.39175 | Q13042 | Q13042 | CDC16 | Cell divisio | Color2 [Da | 1 | 4 |
| + | 1.0529 | -0.28151 | Q13043 | Q13043 | STK4 | Serine/thr | Color2 [Da | 1 | 4 |
| + | 3.861594 | -1.87871 | Q13045 | Q13045 | FLII | Protein flig | Color2 [Da | 1 | 4 |
| | 0 | -0.56645 | Q13049 | Q13049 | TRIM32 | E3 ubiquiti | Color2 [Da | 1 | 4 |
| | 0.361884 | -0.15869 | Q13057 | Q13057 | COASY | Bifunction | Color2 [Da | 1 | 4 |
| + | 1.761073 | -0.6544 | Q13084 | Q13084 | MRPL28 | 39S riboso | Color2 [Da | 1 | 4 |
| | 1.12335 | -0.23544 | Q13085 | Q13085 | ACACA | Acetyl-CoA | Color2 [Da | 1 | 4 |
| + | 2.379609 | -1.03 | Q13098 | Q13098 | GPS1 | COP9 signa | Color2 [Da | 1 | 4 |
| | 0 | -0.67573 | Q13107 | Q13107 | USP4 | Ubiquitin c | Color2 [Da | 1 | 4 |
| | 0 | NaN | Q13111 | Q13111 | CHAF1A | Chromatin | Color2 [Da | 1 | 4 |
| | 0 | -0.00337 | Q13112 | Q13112 | CHAF1B | Chromatin | Color2 [Da | 1 | 4 |
| + | 1.651249 | -0.49908 | Q13114 | Q13114 | TRAF3 | TNF recept | Color2 [Da | 1 | 4 |
| | 0.444066 | 0.198913 | Q13123 | Q13123 | IK | Protein Re | Color2 [Da | 1 | 4 |
| + | 1.792877 | -0.60367 | Q13126 | Q13126 | MTAP | S-methyl-5 | Color2 [Da | 1 | 4 |
| + | 1.968442 | -0.82404 | Q13131 | Q13131 | PRKAA1 | 5'-AMP-ac | Color2 [Da | 1 | 4 |
| | 0.971425 | -0.25999 | Q13136 | Q13136 | PPFIA1 | Liprin-alph | Color2 [Da | 1 | 4 |
| | 0.660119 | -0.65876 | Q13144 | Q13144 | EIF2B5 | Translation | Color2 [Da | 1 | 4 |
| + | 1.958906 | -0.42685 | Q13148 | Q13148 | TARDBP | TAR DNA-b | Color2 [Da | 1 | 4 |
| + | 1.19013 | 0.351419 | Q13151 | Q13151 | HNRNPA0 | Heterogen | Color2 [Da | 1 | 4 |
| + | 2.022126 | 0.570804 | Q13153 | Q13153 | PAK1 | Serine/thr | Color2 [Da | 1 | 4 |
| + | 2.484381 | -2.11039 | Q13155 | Q13155 | AIMP2 | Aminoacyl | Color2 [Da | 1 | 4 |
| | 0.540401 | 0.617407 | Q13158 | Q13158 | FADD | FAS-associ | Color2 [Da | 1 | 4 |
| | 0.144689 | -0.05929 | Q13162 | Q13162 | PRDX4 | Peroxiredc | Color2 [Da | 1 | 4 |
| | 0 | 1.385289 | Q13164 | Q13164 | MAPK7 | Mitogen-a | Color2 [Da | 1 | 4 |
| + | 1.89309 | 0.471368 | Q13177 | Q13177 | PAK2 | Serine/thr | Color2 [Da | 1 | 4 |
| + | 2.119166 | 0.890719 | Q13185 | Q13185 | CBX3 | Chromobo | Color2 [Da | 1 | 4 |
| | 0.523598 | 0.688343 | Q13188 | Q13188 | STK3 | Serine/thr | Color2 [Da | 1 | 4 |
| + | 1.163239 | 0.402636 | Q13190 | Q13190 | STX5 | Syntaxin-5 | Color2 [Da | 1 | 4 |
| | 0.037358 | -0.03846 | Q13191 | Q13191 | CBLB | E3 ubiquiti | Color2 [Da | 1 | 4 |

| | | | | | | | | | |
|---|---|---|---|---|---|---|---|---|---|
| + | 2.839756 | -1.08814 | Q13200 | Q13200 | PSMD2 | 26S protea | Color2 [Da | 1 | 4 |
| + | 3.180157 | -3.25706 | Q13201 | Q13201 | MMRN1 | Multimerin | Color2 [Da | 1 | 4 |
| + | 2.25682 | -0.97238 | Q13206 | Q13206 | DDX10 | Probable A | Color2 [Da | 1 | 4 |
| + | 1.581535 | -0.52068 | Q13217 | Q13217 | DNAJC3 | DnaJ homo | Color2 [Da | 1 | 4 |
| + | 2.758043 | 1.324669 | Q13228 | Q13228 | SELENBP1 | Methaneth | Color2 [Da | 1 | 4 |
| + | 1.000719 | -0.47161 | Q13232 | Q13232 | NME3 | Nucleoside | Color2 [Da | 1 | 4 |
| + | 1.901015 | 0.871536 | Q13242 | Q13242 | SRSF9 | Serine/arg | Color2 [Da | 1 | 4 |
| + | 2.115356 | 1.167567 | Q13243 | Q13243 | SRSF5 | Serine/arg | Color2 [Da | 1 | 4 |
| + | 1.281607 | 0.902335 | Q13247 | Q13247 | SRSF6 | Serine/arg | Color2 [Da | 1 | 4 |
| | 0.639345 | 0.32122 | Q13257 | Q13257 | MAD2L1 | Mitotic spi | Color2 [Da | 1 | 4 |
| | 0.581213 | 0.07824 | Q13263 | Q13263 | TRIM28 | Transcripti | Color2 [Da | 1 | 4 |
| + | 1.239165 | 0.339911 | Q13283 | Q13283 | G3BP1 | Ras GTPase | Color2 [Da | 1 | 4 |
| | 0 | -0.97485 | Q13303 | Q13303 | KCNAB2 | Voltage-ga | Color2 [Da | 1 | 4 |
| + | 3.04081 | -1.11415 | Q13308 | Q13308 | PTK7 | Inactive ty | Color2 [Da | 1 | 4 |
| | 0.591537 | 0.185026 | Q13310 | Q13310 | PABPC4 | Polyadeny | Color2 [Da | 1 | 4 |
| | 0 | 0.545785 | Q13315 | Q13315 | ATM | Serine-pro | Color2 [Da | 1 | 4 |
| | 0 | -1.59535 | Q13322 | Q13322 | GRB10 | Growth fac | Color2 [Da | 1 | 4 |
| | 0.602344 | -0.44658 | Q13325 | Q13325 | IFIT5 | Interferon- | Color2 [Da | 1 | 4 |
| + | 2.406932 | -0.98304 | Q13330 | Q13330 | MTA1 | Metastasis | Color2 [Da | 1 | 4 |
| + | 2.330809 | 1.083297 | Q13332 | Q13332 | PTPRS | Receptor-t | Color2 [Da | 1 | 4 |
| + | 2.305155 | -0.52314 | Q13347 | Q13347 | EIF3I | Eukaryotic | Color2 [Da | 1 | 4 |
| + | 1.338176 | -0.9426 | Q13356 | Q13356 | PPIL2 | RING-type | Color2 [Da | 1 | 4 |
| + | 2.278404 | -0.97558 | Q13362 | Q13362 | PPP2R5C | Serine/thr | Color2 [Da | 1 | 4 |
| + | 2.234645 | -1.02234 | Q13363 | Q13363 | CTBP1 | C-terminal | Color2 [Da | 1 | 4 |
| | 1.230812 | 0.112864 | Q13367 | Q13367 | AP3B2 | AP-3 comp | Color2 [Da | 1 | 4 |
| + | 2.065793 | 0.815702 | Q13368 | Q13368 | MPP3 | MAGUK p5 | Color2 [Da | 1 | 4 |
| + | 2.502468 | -0.95427 | Q13394 | Q13394 | MAB21L1 | Putative n | Color2 [Da | 1 | 4 |
| + | 2.592348 | 1.848973 | Q13404 | Q13404 | UBE2V1 | Ubiquitin-c | Color2 [Da | 1 | 4 |
| + | 2.27792 | 0.813082 | Q13405 | Q13405 | MRPL49 | 39S riboso | Color2 [Da | 1 | 4 |
| + | 2.898151 | 0.421921 | Q13409 | Q13409 | DYNC1I2 | Cytoplasm | Color2 [Da | 1 | 4 |
| | 0 | -1.0843 | Q13416 | Q13416 | ORC2 | Origin reco | Color2 [Da | 1 | 4 |
| + | 2.526174 | -0.9397 | Q13418 | Q13418 | ILK | Integrin-lir | Color2 [Da | 1 | 4 |
| + | 2.655153 | -0.41807 | Q13423 | Q13423 | NNT | NAD(P) tra | Color2 [Da | 1 | 4 |
| + | 1.175231 | 1.402772 | Q13424 | Q13424 | SNTA1 | Alpha-1-sy | Color2 [Da | 1 | 4 |
| + | 2.304415 | 1.438254 | Q13425 | Q13425 | SNTB2 | Beta-2-syn | Color2 [Da | 1 | 4 |
| | 0.081887 | -0.1062 | Q13427 | Q13427 | PPIG | Peptidyl-p | Color2 [Da | 1 | 4 |
| + | 2.350503 | 0.959044 | Q13428 | Q13428 | TCOF1 | Treacle pro | Color2 [Da | 1 | 4 |
| + | 2.080472 | 1.497989 | Q13432 | Q13432 | UNC119 | Protein un | Color2 [Da | 1 | 4 |
| + | 1.99666 | 0.662591 | Q13433 | Q13433 | SLC39A6 | Zinc transp | Color2 [Da | 1 | 4 |
| + | 2.572038 | 0.866228 | Q13435 | Q13435 | SF3B2 | Splicing fac | Color2 [Da | 1 | 4 |
| + | 1.301628 | -0.36601 | Q13438 | Q13438 | OS9 | Protein OS | Color2 [Da | 1 | 4 |
| | 0.616223 | -0.08844 | Q13439 | Q13439 | GOLGA4 | Golgin sub | Color2 [Da | 1 | 4 |
| + | 3.788067 | 2.988655 | Q13442 | Q13442 | PDAP1 | 28 kDa hea | Color2 [Da | 1 | 4 |
| | 0.177459 | -0.14661 | Q13445 | Q13445 | TMED1 | Transmem | Color2 [Da | 1 | 4 |
| + | 2.123652 | 0.667769 | Q13449 | Q13449 | LSAMP | Limbic syst | Color2 [Da | 1 | 4 |
| + | 2.382726 | -1.94214 | Q13451 | Q13451 | FKBP5 | Peptidyl-pi | Color2 [Da | 1 | 4 |
| | 0 | 0.903247 | Q13454 | Q13454 | TUSC3 | Tumor sup | Color2 [Da | 1 | 4 |

| | | | | | | | | | |
|---|---|---|---|---|---|---|---|---|---|
| + | 1.220938 | -0.46529 | Q13459 | Q13459 | MYO9B | Unconvent | Color2 [Da | 1 | 4 |
| | 0.151977 | 0.087463 | Q13464 | Q13464 | ROCK1 | Rho-associ | Color2 [Da | 1 | 4 |
| + | 2.774499 | -1.47888 | Q13491 | Q13491 | GPM6B | Neuronal r | Color2 [Da | 1 | 4 |
| | 1.129496 | -0.22421 | Q13492 | Q13492 | PICALM | Phosphatic | Color2 [Da | 1 | 4 |
| + | 1.642451 | -2.06984 | Q13496 | Q13496 | MTM1 | Myotubula | Color2 [Da | 1 | 4 |
| | 0.018394 | -0.01157 | Q13501 | Q13501 | SQSTM1 | Sequestos | Color2 [Da | 1 | 4 |
| | 0.389848 | -0.15166 | Q13505 | Q13505 | MTX1 | Metaxin-1 | Color2 [Da | 1 | 4 |
| | 0.059485 | 0.013718 | Q13509 | Q13509 | TUBB3 | Tubulin be | Color2 [Da | 1 | 4 |
| + | 5.035233 | 1.640111 | Q13510 | Q13510 | ASAH1 | Acid ceram | Color2 [Da | 1 | 4 |
| + | 2.411284 | 2.4841 | Q13522 | Q13522 | PPP1R1A | Protein ph | Color2 [Da | 1 | 4 |
| | 0.722395 | -0.23875 | Q13523 | Q13523 | PRPF4B | Serine/thr | Color2 [Da | 1 | 4 |
| + | 1.277452 | 0.269067 | Q13526 | Q13526 | PIN1 | Peptidyl-p | Color2 [Da | 1 | 4 |
| + | 1.078265 | 2.005212 | Q13541 | Q13541 | EIF4EBP1 | Eukaryotic | Color2 [Da | 1 | 4 |
| | 0 | NaN | Q13542 | Q13542 | EIF4EBP2 | Eukaryotic | Color2 [Da | 1 | 4 |
| + | 1.801387 | -0.51991 | Q13546 | Q13546 | RIPK1 | Receptor-i | Color2 [Da | 1 | 4 |
| + | 3.202357 | -1.63044 | Q13547 | Q13547 | HDAC1 | Histone de | Color2 [Da | 1 | 4 |
| + | 1.279043 | 0.301874 | Q13554 | Q13554 | CAMK2B | Calcium/ca | Color2 [Da | 1 | 4 |
| + | 0.832148 | 0.765416 | Q13555 | Q13555 | CAMK2G | Calcium/ca | Color2 [Da | 1 | 4 |
| + | 1.137201 | -0.31675 | Q13557 | Q13557 | CAMK2D | Calcium/ca | Color2 [Da | 1 | 4 |
| + | 4.60957 | 2.123311 | Q13561 | Q13561 | DCTN2 | Dynactin s | Color2 [Da | 1 | 4 |
| + | 1.625601 | -0.23408 | Q13564 | Q13564 | NAE1 | NEDD8-act | Color2 [Da | 1 | 4 |
| | 0.228812 | -0.1111 | Q13572 | Q13572 | ITPK1 | Inositol-te | Color2 [Da | 1 | 4 |
| | 1.064332 | -0.15033 | Q13573 | Q13573 | SNW1 | SNW doma | Color2 [Da | 1 | 4 |
| + | 2.165418 | -1.09292 | Q13574 | Q13574 | DGKZ | Diacylglyce | Color2 [Da | 1 | 4 |
| + | 2.697022 | 0.528413 | Q13576 | Q13576 | IQGAP2 | Ras GTPase | Color2 [Da | 1 | 4 |
| | 0.574479 | -0.4181 | Q13586 | Q13586 | STIM1 | Stromal in | Color2 [Da | 1 | 4 |
| + | 1.133716 | 0.315285 | Q13595 | Q13595 | TRA2A | Transform | Color2 [Da | 1 | 4 |
| | 0.615695 | -0.10452 | Q13596 | Q13596 | SNX1 | Sorting ne | Color2 [Da | 1 | 4 |
| + | 2.963997 | 1.733832 | Q13601 | Q13601 | KRR1 | KRR1 smal | Color2 [Da | 1 | 4 |
| + | 1.191854 | -0.51726 | Q13610 | Q13610 | PWP1 | Periodic tr | Color2 [Da | 1 | 4 |
| + | 2.226129 | -1.15264 | Q13613 | Q13613 | MTMR1 | Myotubula | Color2 [Da | 1 | 4 |
| + | 1.736678 | -0.741 | Q13614 | Q13614 | MTMR2 | Myotubula | Color2 [Da | 1 | 4 |
| + | 1.374592 | -0.80058 | Q13615 | Q13615 | MTMR3 | Myotubula | Color2 [Da | 1 | 4 |
| + | 2.553927 | -1.55204 | Q13616 | Q13616 | CUL1 | Cullin-1 | Color2 [Da | 1 | 4 |
| + | 1.668816 | -0.70142 | Q13617 | Q13617 | CUL2 | Cullin-2 | Color2 [Da | 1 | 4 |
| + | 3.387142 | -1.13883 | Q13618 | Q13618 | CUL3 | Cullin-3 | Color2 [Da | 1 | 4 |
| + | 2.670864 | -1.26755 | Q13619 | Q13619 | CUL4A | Cullin-4A | Color2 [Da | 1 | 4 |
| + | 2.404824 | -1.00288 | Q13620 | Q13620 | CUL4B | Cullin-4B | Color2 [Da | 1 | 4 |
| + | 1.922769 | -0.21826 | Q13625 | Q13625 | TP53BP2 | Apoptosis- | Color2 [Da | 1 | 4 |
| + | 2.136378 | -1.30146 | Q13627 | Q13627 | DYRK1A | Dual speci | Color2 [Da | 1 | 4 |
| | 0.246433 | -0.27296 | Q13630 | Q13630 | GFUS | GDP-L-fuc | Color2 [Da | 1 | 4 |
| | 0.48876 | 0.179847 | Q13636 | Q13636 | RAB31 | Ras-relate | Color2 [Da | 1 | 4 |
| + | 1.057078 | -0.84136 | Q13637 | Q13637 | RAB32 | Ras-relate | Color2 [Da | 1 | 4 |
| | 0 | -0.2853 | Q13641 | Q13641 | TPBG | Trophobla | Color2 [Da | 1 | 4 |
| + | 2.616999 | 0.519262 | Q13642 | Q13642 | FHL1 | Four and a | Color2 [Da | 1 | 4 |
| | 0 | -2.01634 | Q13671 | Q13671 | RIN1 | Ras and Ra | Color2 [Da | 1 | 4 |
| | 0 | -0.7963 | Q13683 | Q13683 | ITGA7 | Integrin al | Color2 [Da | 1 | 4 |

| | | | | | | | | | |
|---|---|---|---|---|---|---|---|---|---|
| + | 0.826418 | -0.79638 | Q13685 | Q13685 | AAMP | Angio-assc | Color2 [Da | 1 | 4 |
| + | 2.255504 | -0.74749 | Q13724 | Q13724 | MOGS | Mannosyl- | Color2 [Da | 1 | 4 |
| + | 1.256255 | 0.506529 | Q13740 | Q13740 | ALCAM | CD166 ant | Color2 [Da | 1 | 4 |
| + | 2.131561 | -1.44965 | Q13769 | Q13769 | THOC5 | THO comp | Color2 [Da | 1 | 4 |
| + | 2.110817 | 0.659957 | Q13796 | Q13796 | SHROOM2 | Protein Sh | Color2 [Da | 1 | 4 |
| + | 3.69241 | 0.397223 | Q13813 | Q13813 | SPTAN1 | Spectrin al | Color2 [Da | 1 | 4 |
| + | 2.026824 | -0.5473 | Q13823 | Q13823 | GNL2 | Nucleolar ( | Color2 [Da | 1 | 4 |
| | 0.451596 | 0.245228 | Q13825 | Q13825 | AUH | Methylglut | Color2 [Da | 1 | 4 |
| | 0.069229 | -0.05106 | Q13835 | Q13835 | PKP1 | Plakophilir | Color2 [Da | 1 | 4 |
| + | 3.764969 | -1.23958 | Q13838 | Q13838 | DDX39B | Spliceoson | Color2 [Da | 1 | 4 |
| + | 1.616501 | 0.444591 | Q13867 | Q13867 | BLMH | Bleomycin | Color2 [Da | 1 | 4 |
| + | 1.405908 | 0.4469 | Q13868 | Q13868 | EXOSC2 | Exosome c | Color2 [Da | 1 | 4 |
| | 0 | -0.33421 | Q13873 | Q13873 | BMPR2 | Bone morp | Color2 [Da | 1 | 4 |
| + | 1.504857 | 1.07196 | Q13884 | Q13884 | SNTB1 | Beta-1-syn | Color2 [Da | 1 | 4 |
| + | 1.256842 | 1.156813 | Q13885 | Q13885 | TUBB2A | Tubulin be | Color2 [Da | 1 | 4 |
| | 0.335467 | 0.180475 | Q13895 | Q13895 | BYSL | Bystin | Color2 [Da | 1 | 4 |
| | 0.235349 | -0.10261 | Q13907 | Q13907 | IDI1 | Isopenteny | Color2 [Da | 1 | 4 |
| + | 2.600257 | -2.3504 | Q13938 | Q13938 | CAPS | Calcyphosi | Color2 [Da | 1 | 4 |
| | 0 | NaN | Q13948 | Q13948 | CUX1 | Protein CA | Color2 [Da | 1 | 4 |
| + | 1.490507 | -0.62877 | Q13951 | Q13951 | CBFB | Core-bindi | Color2 [Da | 1 | 4 |
| + | 2.584095 | 1.607577 | Q13952 | Q13952 | NFYC | Nuclear tra | Color2 [Da | 1 | 4 |
| | 0.461523 | -0.70833 | Q14004 | Q14004 | CDK13 | Cyclin-dep | Color2 [Da | 1 | 4 |
| + | 2.793678 | -0.73559 | Q14008 | Q14008 | CKAP5 | Cytoskelet | Color2 [Da | 1 | 4 |
| + | 2.794983 | 1.802727 | Q14011 | Q14011 | CIRBP | Cold-induc | Color2 [Da | 1 | 4 |
| + | 1.29135 | 0.610309 | Q14012 | Q14012 | CAMK1 | Calcium/ca | Color2 [Da | 1 | 4 |
| + | 2.160856 | 0.667662 | Q14019 | Q14019 | COTL1 | Coactosin- | Color2 [Da | 1 | 4 |
| + | 3.308088 | 2.512689 | Q14061 | Q14061 | COX17 | Cytochron | Color2 [Da | 1 | 4 |
| + | 0.676873 | 1.434216 | Q14088 | Q14088 | RAB33A | Ras-relate | Color2 [Da | 1 | 4 |
| + | 3.243219 | 1.691926 | Q14103 | Q14103 | HNRNPD | Heterogen | Color2 [Da | 1 | 4 |
| + | 3.297515 | -1.45374 | Q14108 | Q14108 | SCARB2 | Lysosome | Color2 [Da | 1 | 4 |
| | 0 | NaN | Q14112 | Q14112 | NID2 | Nidogen-2 | Color2 [Da | 1 | 4 |
| | 0 | NaN | Q14117 | Q14117 | DPYS | Dihydropy | Color2 [Da | 1 | 4 |
| + | 3.453706 | 1.098738 | Q14118 | Q14118 | DAG1 | Dystroglyc | Color2 [Da | 1 | 4 |
| + | 1.940153 | -0.73016 | Q14123 | Q14123 | PDE1C | Dual speci | Color2 [Da | 1 | 4 |
| | 0 | -1.07979 | Q14126 | Q14126 | DSG2 | Desmoglei | Color2 [Da | 1 | 4 |
| | 0.683839 | -0.73814 | Q14137 | Q14137 | BOP1 | Ribosome | Color2 [Da | 1 | 4 |
| + | 2.297159 | -1.83418 | Q14139 | Q14139 | UBE4A | Ubiquitin c | Color2 [Da | 1 | 4 |
| + | 1.334466 | 0.477914 | Q14141 | Q14141 | SEPTIN6 | Septin-6 | Color2 [Da | 1 | 4 |
| | 0 | NaN | Q14149 | Q14149 | MORC3 | MORC fam | Color2 [Da | 1 | 4 |
| + | 1.459498 | 0.36222 | Q14151 | Q14151 | SAFB2 | Scaffold at | Color2 [Da | 1 | 4 |
| + | 3.222318 | -0.7278 | Q14152 | Q14152 | EIF3A | Eukaryotic | Color2 [Da | 1 | 4 |
| + | 1.242545 | 0.382277 | Q14155 | Q14155 | ARHGEF7 | Rho guanir | Color2 [Da | 1 | 4 |
| + | 3.28063 | 0.860022 | Q14157 | Q14157 | UBAP2L | Ubiquitin-a | Color2 [Da | 1 | 4 |
| | 0.035488 | -0.00939 | Q14160 | Q14160 | SCRIB | Protein scr | Color2 [Da | 1 | 4 |
| | 0.517056 | 0.14987 | Q14161 | Q14161 | GIT2 | ARF GTPas | Color2 [Da | 1 | 4 |
| + | 2.598634 | -1.25873 | Q14165 | Q14165 | MLEC | Malectin | Color2 [Da | 1 | 4 |
| + | 2.195329 | -1.21579 | Q14166 | Q14166 | TTLL12 | Tubulin--ty | Color2 [Da | 1 | 4 |

| | | | | | | | | | |
|---|---|---|---|---|---|---|---|---|---|
| | 0.392162 | 0.176701 | Q14168 | Q14168 | MPP2 | MAGUK p5 | Color2 [Da | 1 | 4 |
| | 0 | NaN | Q14181 | Q14181 | POLA2 | DNA polyn | Color2 [Da | 1 | 4 |
| + | 1.598618 | -0.5506 | Q14185 | Q14185 | DOCK1 | Dedicator | Color2 [Da | 1 | 4 |
| | 0 | -0.74921 | Q14186 | Q14186 | TFDP1 | Transcripti | Color2 [Da | 1 | 4 |
| | 0 | NaN | Q14188 | Q14188 | TFDP2 | Transcripti | Color2 [Da | 1 | 4 |
| | 0 | NaN | Q14191 | Q14191 | WRN | Werner sy | Color2 [Da | 1 | 4 |
| + | 2.028905 | 0.434665 | Q14194 | Q14194 | CRMP1 | Dihydropy | Color2 [Da | 1 | 4 |
| | 0.907654 | 0.305804 | Q14195 | Q14195 | DPYSL3 | Dihydropy | Color2 [Da | 1 | 4 |
| + | 1.753281 | 0.713764 | Q14197 | Q14197 | MRPL58 | Peptidyl-tF | Color2 [Da | 1 | 4 |
| | 0.228482 | 0.378247 | Q14202 | Q14202 | ZMYM3 | Zinc finger | Color2 [Da | 1 | 4 |
| | 1.281516 | 0.20833 | Q14203 | Q14203 | DCTN1 | Dynactin s | Color2 [Da | 1 | 4 |
| + | 3.910634 | -1.26027 | Q14204 | Q14204 | DYNC1H1 | Cytoplasm | Color2 [Da | 1 | 4 |
| + | 2.770332 | -1.47098 | Q14232 | Q14232 | EIF2B1 | Translatior | Color2 [Da | 1 | 4 |
| + | 1.395422 | -0.67019 | Q14240 | Q14240 | EIF4A2 | Eukaryotic | Color2 [Da | 1 | 4 |
| + | 1.974611 | -0.71669 | Q14241 | Q14241 | ELOA | Elongin-A | Color2 [Da | 1 | 4 |
| + | 2.022978 | 2.533578 | Q14244 | Q14244 | MAP7 | Ensconsin | Color2 [Da | 1 | 4 |
| + | 2.576913 | 1.393323 | Q14247 | Q14247 | CTTN | Src substra | Color2 [Da | 1 | 4 |
| | 0.976608 | -0.30678 | Q14254 | Q14254 | FLOT2 | Flotillin-2 | Color2 [Da | 1 | 4 |
| + | 1.979339 | 0.975541 | Q14257 | Q14257 | RCN2 | Reticuloca | Color2 [Da | 1 | 4 |
| + | 1.012425 | -0.90901 | Q14258 | Q14258 | TRIM25 | E3 ubiquiti | Color2 [Da | 1 | 4 |
| + | 2.431218 | -0.63433 | Q14315 | Q14315 | FLNC | Filamin-C | Color2 [Da | 1 | 4 |
| + | 1.538275 | -0.79522 | Q14318 | Q14318 | FKBP8 | Peptidyl-pı | Color2 [Da | 1 | 4 |
| | 0.308297 | 0.120191 | Q14320 | Q14320 | FAM50A | Protein FA | Color2 [Da | 1 | 4 |
| | 0.288055 | -0.39111 | Q14331 | Q14331 | FRG1 | Protein FR | Color2 [Da | 1 | 4 |
| + | 0.92349 | -0.52488 | Q14344 | Q14344 | GNA13 | Guanine nı | Color2 [Da | 1 | 4 |
| + | 2.111931 | 0.822323 | Q14353 | Q14353 | GAMT | Guanidino | Color2 [Da | 1 | 4 |
| + | 2.992378 | -0.21021 | Q14376 | Q14376 | GALE | UDP-gluco | Color2 [Da | 1 | 4 |
| | 0 | -0.63764 | Q14432 | Q14432 | PDE3A | cGMP-inhi | Color2 [Da | 1 | 4 |
| | 0.178755 | -0.08828 | Q14444 | Q14444 | CAPRIN1 | Caprin-1 | Color2 [Da | 1 | 4 |
| | 0.922324 | 0.209946 | Q14457 | Q14457 | BECN1 | Beclin-1 | Color2 [Da | 1 | 4 |
| | 0.694103 | -0.11493 | Q14498 | Q14498 | RBM39 | RNA-bindiı | Color2 [Da | 1 | 4 |
| | 0.085671 | -0.04773 | Q14515 | Q14515 | SPARCL1 | SPARC-like | Color2 [Da | 1 | 4 |
| + | 2.126007 | -0.61177 | Q14517 | Q14517 | FAT1 | Protocadh | Color2 [Da | 1 | 4 |
| | 1.044128 | -0.24364 | Q14527 | Q14527 | HLTF | Helicase-lil | Color2 [Da | 1 | 4 |
| | 0 | -1.31669 | Q14532 | Q14532 | KRT32 | Keratin, ty | Color2 [Da | 1 | 4 |
| | 0.283018 | 0.249416 | Q14534 | Q14534 | SQLE | Squalene r | Color2 [Da | 1 | 4 |
| + | 2.502109 | -1.47341 | Q14554 | Q14554 | PDIA5 | Protein dis | Color2 [Da | 1 | 4 |
| + | 1.204636 | -0.33876 | Q14558 | Q14558 | PRPSAP1 | Phosphoril | Color2 [Da | 1 | 4 |
| | 0.762826 | -0.29214 | Q14562 | Q14562 | DHX8 | ATP-deper | Color2 [Da | 1 | 4 |
| | 0.454925 | 0.143686 | Q14563 | Q14563 | SEMA3A | Semaphor | Color2 [Da | 1 | 4 |
| + | 2.555118 | -1.5076 | Q14566 | Q14566 | MCM6 | DNA replic | Color2 [Da | 1 | 4 |
| | 0.663706 | -0.84612 | Q14568 | Q14568 | HSP90AA2 | Heat shocl | Color2 [Da | 1 | 4 |
| | 0 | 1.649159 | Q14574 | Q14574 | DSC3 | Desmocoll | Color2 [Da | 1 | 4 |
| + | 2.602573 | 1.26428 | Q14576 | Q14576 | ELAVL3 | ELAV-like p | Color2 [Da | 1 | 4 |
| | 0 | NaN | Q14590 | Q14590 | ZNF235 | Zinc finger | Color2 [Da | 1 | 4 |
| + | 1.656396 | 1.028517 | Q14624 | Q14624 | ITIH4 | Inter-alpha | Color2 [Da | 1 | 4 |
| | 0 | -0.99776 | Q14642 | Q14642 | INPP5A | Inositol po | Color2 [Da | 1 | 4 |

| | | | | | | | | | |
|---|---|---|---|---|---|---|---|---|---|
| | 0 | NaN | Q14651 | Q14651 | PLS1 | Plastin-1 | Color2 [Da | 1 | 4 |
| | 0 | -0.5929 | Q14653 | Q14653 | IRF3 | Interferon | Color2 [Da | 1 | 4 |
| | 0.05575 | -0.05942 | Q14657 | Q14657 | LAGE3 | EKC/KEOP: | Color2 [Da | 1 | 4 |
| | 0 | NaN | Q14667 | Q14667 | KIAA0100 | Protein KI/ | Color2 [Da | 1 | 4 |
| + | 2.159463 | -0.69383 | Q14669 | Q14669 | TRIP12 | E3 ubiquiti | Color2 [Da | 1 | 4 |
| + | 1.892163 | -0.57141 | Q14671 | Q14671 | PUM1 | Pumilio ho | Color2 [Da | 1 | 4 |
| | 0.344262 | -0.111 | Q14676 | Q14676 | MDC1 | Mediator ( | Color2 [Da | 1 | 4 |
| + | 1.724532 | 0.844428 | Q14677 | Q14677 | CLINT1 | Clathrin in | Color2 [Da | 1 | 4 |
| | 0 | NaN | Q14678 | Q14678 | KANK1 | KN motif a | Color2 [Da | 1 | 4 |
| | 0.818114 | 0.334632 | Q14681 | Q14681 | KCTD2 | BTB/POZ d | Color2 [Da | 1 | 4 |
| + | 2.155022 | -0.49278 | Q14683 | Q14683 | SMC1A | Structural | Color2 [Da | 1 | 4 |
| + | 1.286605 | -0.54965 | Q14684 | Q14684 | RRP1B | Ribosomal | Color2 [Da | 1 | 4 |
| | 0.726566 | -0.56074 | Q14687 | Q14687 | GSE1 | Genetic su | Color2 [Da | 1 | 4 |
| | 0.570024 | -0.05544 | Q14689 | Q14689 | DIP2A | Disco-inte | Color2 [Da | 1 | 4 |
| | 0.73727 | -0.33242 | Q14690 | Q14690 | PDCD11 | Protein RR | Color2 [Da | 1 | 4 |
| | 0 | NaN | Q14691 | Q14691 | GINS1 | DNA replic | Color2 [Da | 1 | 4 |
| + | 2.131664 | -1.10114 | Q14692 | Q14692 | BMS1 | Ribosome | Color2 [Da | 1 | 4 |
| + | 2.33458 | -0.65273 | Q14694 | Q14694 | USP10 | Ubiquitin ( | Color2 [Da | 1 | 4 |
| + | 2.285714 | 1.039142 | Q14696 | Q14696 | MESD | LRP chape | Color2 [Da | 1 | 4 |
| + | 2.711859 | -1.13587 | Q14697 | Q14697 | GANAB | Neutral alp | Color2 [Da | 1 | 4 |
| + | 1.707327 | 0.593176 | Q14699 | Q14699 | RFTN1 | Raftlin | Color2 [Da | 1 | 4 |
| + | 3.989358 | -2.1553 | Q14728 | Q14728 | MFSD10 | Major facil | Color2 [Da | 1 | 4 |
| + | 2.036874 | -0.57301 | Q14738 | Q14738 | PPP2R5D | Serine/thr | Color2 [Da | 1 | 4 |
| | 0.570301 | 0.297254 | Q14739 | Q14739 | LBR | Delta(14)-: | Color2 [Da | 1 | 4 |
| + | 3.254448 | -0.99153 | Q14746 | Q14746 | COG2 | Conserved | Color2 [Da | 1 | 4 |
| + | 0.786625 | -0.50345 | Q14766 | Q14766 | LTBP1 | Latent-trai | Color2 [Da | 1 | 4 |
| + | 2.296527 | -0.49853 | Q14789 | Q14789 | GOLGB1 | Golgin sub | Color2 [Da | 1 | 4 |
| + | 2.780327 | -1.58222 | Q14807 | Q14807 | KIF22 | Kinesin-lik | Color2 [Da | 1 | 4 |
| | 0 | NaN | Q14831 | Q14831 | GRM7 | Metabotro | Color2 [Da | 1 | 4 |
| + | 1.223953 | -0.34394 | Q14839 | Q14839 | CHD4 | Chromodo | Color2 [Da | 1 | 4 |
| + | 3.556838 | 2.000818 | Q14847 | Q14847 | LASP1 | LIM and SI | Color2 [Da | 1 | 4 |
| + | 0.850803 | 1.060148 | Q14894 | Q14894 | CRYM | Ketimine r | Color2 [Da | 1 | 4 |
| + | 2.753587 | -1.26648 | Q14914 | Q14914 | PTGR1 | Prostaglan | Color2 [Da | 1 | 4 |
| + | 2.512554 | 1.364475 | Q14919 | Q14919 | DRAP1 | Dr1-associ | Color2 [Da | 1 | 4 |
| | 0.0488 | 0.296227 | Q14956 | Q14956 | GPNMB | Transmem | Color2 [Da | 1 | 4 |
| + | 1.617133 | -0.35177 | Q14966 | Q14966 | ZNF638 | Zinc finger | Color2 [Da | 1 | 4 |
| + | 3.027411 | -1.50058 | Q14974 | Q14974 | KPNB1 | Importin s | Color2 [Da | 1 | 4 |
| + | 2.8325 | 2.153514 | Q14978 | Q14978 | NOLC1 | Nucleolar a | Color2 [Da | 1 | 4 |
| + | 2.536412 | -0.85014 | Q14980 | Q14980 | NUMA1 | Nuclear m | Color2 [Da | 1 | 4 |
| + | 1.784846 | 2.175943 | Q14982 | Q14982 | OPCML | Opioid-bin | Color2 [Da | 1 | 4 |
| | 0.600684 | -0.38705 | Q14997 | Q14997 | PSME4 | Proteasom | Color2 [Da | 1 | 4 |
| | 0 | -0.86234 | Q14BN4 | Q14BN4 | SLMAP | Sarcolemn | Color2 [Da | 1 | 4 |
| + | 1.831127 | -1.15241 | Q14C86 | Q14C86 | GAPVD1 | GTPase-ac | Color2 [Da | 1 | 4 |
| + | 2.555554 | 0.631722 | Q14CS0 | Q14CS0 | UBXN2B | UBX doma | Color2 [Da | 1 | 4 |
| + | 1.410611 | -0.68242 | Q14CX7 | Q14CX7 | NAA25 | N-alpha-ac | Color2 [Da | 1 | 4 |
| | 0 | NaN | Q14CZ8 | Q14CZ8 | HEPACAM | Hepatocyt | Color2 [Da | 1 | 4 |
| + | 2.135147 | -1.38569 | Q15003 | Q15003 | NCAPH | Condensin | Color2 [Da | 1 | 4 |

| | | | | | | | | | | |
|---|---|---|---|---|---|---|---|---|---|---|
| + | 2.339405 | -1.66477 | Q15005 | Q15005 | SPCS2 | Signal pep | Color2 | [Da | 1 | 4 |
| + | 3.455635 | -0.79658 | Q15006 | Q15006 | EMC2 | ER membr | Color2 | [Da | 1 | 4 |
| | 0.569743 | 0.194343 | Q15007 | Q15007 | WTAP | Pre-mRNA | Color2 | [Da | 1 | 4 |
| + | 3.497251 | -2.12695 | Q15008 | Q15008 | PSMD6 | 26S protea | Color2 | [Da | 1 | 4 |
| | 0 | -1.32952 | Q15013 | Q15013 | MAD2L1BI | MAD2L1-b | Color2 | [Da | 1 | 4 |
| + | 2.748776 | 0.842165 | Q15014 | Q15014 | MORF4L2 | Mortality f | Color2 | [Da | 1 | 4 |
| + | 1.020293 | 0.604569 | Q15018 | Q15018 | ABRAXAS2 | BRISC com | Color2 | [Da | 1 | 4 |
| + | 4.295995 | 1.431895 | Q15019 | Q15019 | SEPTIN2 | Septin-2 | Color2 | [Da | 1 | 4 |
| + | 2.091112 | -0.82061 | Q15020 | Q15020 | SART3 | Squamous | Color2 | [Da | 1 | 4 |
| + | 3.522983 | -1.19712 | Q15021 | Q15021 | NCAPD2 | Condensin | Color2 | [Da | 1 | 4 |
| + | 0.777334 | -0.92501 | Q15024 | Q15024 | EXOSC7 | Exosome c | Color2 | [Da | 1 | 4 |
| + | 3.323142 | -0.89143 | Q15029 | Q15029 | EFTUD2 | 116 kDa U | Color2 | [Da | 1 | 4 |
| + | 2.14027 | -0.88072 | Q15031 | Q15031 | LARS2 | Leucine--tI | Color2 | [Da | 1 | 4 |
| + | 2.023434 | -0.57898 | Q15036 | Q15036 | SNX17 | Sorting ne | Color2 | [Da | 1 | 4 |
| | 0 | -2.67508 | Q15041 | Q15041 | ARL6IP1 | ADP-ribosy | Color2 | [Da | 1 | 4 |
| + | 2.218109 | -0.83216 | Q15042 | Q15042 | RAB3GAP1 | Rab3 GTPa | Color2 | [Da | 1 | 4 |
| | 0 | NaN | Q15043 | Q15043 | SLC39A14 | Metal cati | Color2 | [Da | 1 | 4 |
| + | 2.528971 | -0.83342 | Q15046 | Q15046 | KARS1 | Lysine--tRI | Color2 | [Da | 1 | 4 |
| | 0 | 0.272314 | Q15047 | Q15047 | SETDB1 | Histone-ly | Color2 | [Da | 1 | 4 |
| + | 2.329425 | -1.48173 | Q15048 | Q15048 | LRRC14 | Leucine-ric | Color2 | [Da | 1 | 4 |
| + | 2.368092 | 0.713434 | Q15050 | Q15050 | RRS1 | Ribosome | Color2 | [Da | 1 | 4 |
| | 0 | 0.990793 | Q15051 | Q15051 | IQCB1 | IQ calmod | Color2 | [Da | 1 | 4 |
| | 0.068962 | 0.047907 | Q15054 | Q15054 | POLD3 | DNA polyn | Color2 | [Da | 1 | 4 |
| + | 3.207976 | 1.705172 | Q15056 | Q15056 | EIF4H | Eukaryotic | Color2 | [Da | 1 | 4 |
| | 0.371882 | 0.105398 | Q15059 | Q15059 | BRD3 | Bromodon | Color2 | [Da | 1 | 4 |
| | 0.587273 | 0.382853 | Q15061 | Q15061 | WDR43 | WD repeat | Color2 | [Da | 1 | 4 |
| | 0 | NaN | Q15063 | Q15063 | POSTN | Periostin | Color2 | [Da | 1 | 4 |
| + | 1.367131 | -0.44892 | Q15067 | Q15067 | ACOX1 | Peroxisom | Color2 | [Da | 1 | 4 |
| + | 2.161888 | 0.317471 | Q15075 | Q15075 | EEA1 | Early endo | Color2 | [Da | 1 | 4 |
| + | 2.664759 | 0.911069 | Q15084 | Q15084 | PDIA6 | Protein dis | Color2 | [Da | 1 | 4 |
| + | 2.562082 | 1.457299 | Q15102 | Q15102 | PAFAH1B3 | Platelet-ac | Color2 | [Da | 1 | 4 |
| + | 3.377728 | 1.028713 | Q15111 | Q15111 | PLCL1 | Inactive pl | Color2 | [Da | 1 | 4 |
| | 0.573356 | 0.1457 | Q15118 | Q15118 | PDK1 | [Pyruvate | Color2 | [Da | 1 | 4 |
| + | 0.967252 | -0.53645 | Q15120 | Q15120 | PDK3 | [Pyruvate | Color2 | [Da | 1 | 4 |
| + | 3.916234 | 1.66909 | Q15121 | Q15121 | PEA15 | Astrocytic | Color2 | [Da | 1 | 4 |
| + | 1.2951 | -1.88831 | Q15125 | Q15125 | EBP | 3-beta-hyc | Color2 | [Da | 1 | 4 |
| | 0.991553 | 0.295567 | Q15126 | Q15126 | PMVK | Phosphom | Color2 | [Da | 1 | 4 |
| | 0 | 0.860736 | Q15139 | Q15139 | PRKD1 | Serine/thr | Color2 | [Da | 1 | 4 |
| | 0.133251 | 0.049654 | Q15147 | Q15147 | PLCB4 | 1-phospha | Color2 | [Da | 1 | 4 |
| + | 2.180206 | -0.26004 | Q15149 | Q15149 | PLEC | Plectin | Color2 | [Da | 1 | 4 |
| | 0.585898 | -0.09104 | Q15154 | Q15154 | PCM1 | Pericentric | Color2 | [Da | 1 | 4 |
| + | 2.419948 | -1.04418 | Q15155 | Q15155 | NOMO1 | Nodal moc | Color2 | [Da | 1 | 4 |
| + | 1.423575 | 0.981401 | Q15165 | Q15165 | PON2 | Serum par | Color2 | [Da | 1 | 4 |
| | 0.374338 | -0.31832 | Q15170 | Q15170 | TCEAL1 | Transcripti | Color2 | [Da | 1 | 4 |
| | 0 | NaN | Q15172 | Q15172 | PPP2R5A | Serine/thr | Color2 | [Da | 1 | 4 |
| | 0.355076 | -0.23693 | Q15173 | Q15173 | PPP2R5B | Serine/thr | Color2 | [Da | 1 | 4 |
| + | 2.886919 | 0.81849 | Q15181 | Q15181 | PPA1 | Inorganic p | Color2 | [Da | 1 | 4 |

| | | | | | | | | | |
|---|---|---|---|---|---|---|---|---|---|
| | 0.207315 | 0.046925 | Q15185 | Q15185 | PTGES3 | Prostaglan | Color2 [Da | 1 | 4 |
| + | 1.468643 | -0.50297 | Q15208 | Q15208 | STK38 | Serine/thr | Color2 [Da | 1 | 4 |
| + | 2.730663 | 1.659842 | Q15223 | Q15223 | NECTIN1 | Nectin-1 | Color2 [Da | 1 | 4 |
| + | 2.566504 | 0.483337 | Q15233 | Q15233 | NONO | Non-POU ( | Color2 [Da | 1 | 4 |
| | 1.089019 | -0.22667 | Q15257 | Q15257 | PTPA | Serine/thr | Color2 [Da | 1 | 4 |
| + | 3.367968 | -1.93952 | Q15269 | Q15269 | PWP2 | Periodic tr | Color2 [Da | 1 | 4 |
| + | 1.541409 | 2.46476 | Q15274 | Q15274 | QPRT | Nicotinate | Color2 [Da | 1 | 4 |
| | 0.612843 | 0.185222 | Q15276 | Q15276 | RABEP1 | Rab GTPas | Color2 [Da | 1 | 4 |
| | 0 | NaN | Q15283 | Q15283 | RASA2 | Ras GTPas | Color2 [Da | 1 | 4 |
| + | 1.335204 | -0.48826 | Q15286 | Q15286 | RAB35 | Ras-relate | Color2 [Da | 1 | 4 |
| + | 2.60863 | 1.936299 | Q15287 | Q15287 | RNPS1 | RNA-bindi | Color2 [Da | 1 | 4 |
| + | 2.775526 | -1.81727 | Q15291 | Q15291 | RBBP5 | Retinoblas | Color2 [Da | 1 | 4 |
| + | 1.649182 | 0.250792 | Q15293 | Q15293 | RCN1 | Reticuloca | Color2 [Da | 1 | 4 |
| | 0.282261 | 0.120429 | Q15303 | Q15303 | ERBB4 | Receptor t | Color2 [Da | 1 | 4 |
| | 0.736562 | 0.468091 | Q15311 | Q15311 | RALBP1 | RalA-bindi | Color2 [Da | 1 | 4 |
| + | 2.734055 | -1.54346 | Q15334 | Q15334 | LLGL1 | Lethal(2) g | Color2 [Da | 1 | 4 |
| | 0 | NaN | Q15349 | Q15349 | RPS6KA2 | Ribosomal | Color2 [Da | 1 | 4 |
| + | 1.481631 | -0.51091 | Q15363 | Q15363 | TMED2 | Transmem | Color2 [Da | 1 | 4 |
| | 0.208077 | -0.06086 | Q15365 | Q15365 | PCBP1 | Poly(rC)-bi | Color2 [Da | 1 | 4 |
| + | 1.713539 | -0.50915 | Q15366 | Q15366 | PCBP2 | Poly(rC)-bi | Color2 [Da | 1 | 4 |
| + | 1.808439 | 0.701599 | Q15369 | Q15369 | ELOC | Elongin-C | Color2 [Da | 1 | 4 |
| + | 2.467657 | 0.868689 | Q15370 | Q15370 | ELOB | Elongin-B | Color2 [Da | 1 | 4 |
| | 0 | NaN | Q15375 | Q15375 | EPHA7 | Ephrin typ | Color2 [Da | 1 | 4 |
| + | 2.893205 | 1.977668 | Q15382 | Q15382 | RHEB | GTP-bindir | Color2 [Da | 1 | 4 |
| | 0 | NaN | Q15386 | Q15386 | UBE3C | Ubiquitin-p | Color2 [Da | 1 | 4 |
| | 0.137871 | -0.40439 | Q15388 | Q15388 | TOMM20 | Mitochonc | Color2 [Da | 1 | 4 |
| | 0.017647 | -0.04946 | Q15392 | Q15392 | DHCR24 | Delta(24)-s | Color2 [Da | 1 | 4 |
| + | 3.833079 | -2.11551 | Q15393 | Q15393 | SF3B3 | Splicing fa | Color2 [Da | 1 | 4 |
| + | 1.462055 | -0.98638 | Q15397 | Q15397 | PUM3 | Pumilio ho | Color2 [Da | 1 | 4 |
| + | 2.462417 | 1.46769 | Q15404 | Q15404 | RSU1 | Ras suppre | Color2 [Da | 1 | 4 |
| | 0.161528 | 0.054247 | Q15417 | Q15417 | CNN3 | Calponin-3 | Color2 [Da | 1 | 4 |
| + | 2.516072 | -1.15931 | Q15418 | Q15418 | RPS6KA1 | Ribosomal | Color2 [Da | 1 | 4 |
| + | 1.513233 | 0.833918 | Q15424 | Q15424 | SAFB | Scaffold at | Color2 [Da | 1 | 4 |
| + | 2.562625 | 1.129454 | Q15427 | Q15427 | SF3B4 | Splicing fa | Color2 [Da | 1 | 4 |
| + | 3.011876 | 1.122388 | Q15428 | Q15428 | SF3A2 | Splicing fa | Color2 [Da | 1 | 4 |
| + | 3.52316 | 1.679392 | Q15435 | Q15435 | PPP1R7 | Protein ph | Color2 [Da | 1 | 4 |
| + | 3.706204 | -1.44036 | Q15436 | Q15436 | SEC23A | Protein tra | Color2 [Da | 1 | 4 |
| + | 4.070908 | -1.64769 | Q15437 | Q15437 | SEC23B | Protein tra | Color2 [Da | 1 | 4 |
| + | 2.776748 | 0.741432 | Q15459 | Q15459 | SF3A1 | Splicing fa | Color2 [Da | 1 | 4 |
| | 0 | NaN | Q15465 | Q15465 | SHH | Sonic hedg | Color2 [Da | 1 | 4 |
| | 0.126786 | 0.138722 | Q15477 | Q15477 | SKIV2L | Helicase SI | Color2 [Da | 1 | 4 |
| | 0.109959 | -0.02789 | Q15506 | Q15506 | SPA17 | Sperm surf | Color2 [Da | 1 | 4 |
| + | 2.306323 | 2.209961 | Q15517 | Q15517 | CDSN | Corneodes | Color2 [Da | 1 | 4 |
| + | 3.624181 | 2.32574 | Q15526 | Q15526 | SURF1 | Surfeit locu | Color2 [Da | 1 | 4 |
| | 0.610665 | 0.168841 | Q15528 | Q15528 | MED22 | Mediator ( | Color2 [Da | 1 | 4 |
| + | 1.597661 | 0.800641 | Q15532 | Q15532 | SS18 | Protein SS | Color2 [Da | 1 | 4 |
| | 0 | NaN | Q15542 | Q15542 | TAF5 | Transcripti | Color2 [Da | 1 | 4 |

| | | | | | | | | | |
|---|---|---|---|---|---|---|---|---|---|
| + | 1.161892 | 0.368389 | Q15545 | Q15545 | TAF7 | Transcripti | Color2 [Da | 1 | 4 |
| | 0.744935 | 0.118018 | Q15554 | Q15554 | TERF2 | Telomeric | Color2 [Da | 1 | 4 |
| + | 4.127083 | 1.006716 | Q15555 | Q15555 | MAPRE2 | Microtubu | Color2 [Da | 1 | 4 |
| | 0 | NaN | Q15582 | Q15582 | TGFBI | Transform | Color2 [Da | 1 | 4 |
| + | 0.792116 | -0.79348 | Q15596 | Q15596 | NCOA2 | Nuclear re | Color2 [Da | 1 | 4 |
| + | 2.239507 | 1.40438 | Q15599 | Q15599 | SLC9A3R2 | Na(+)/H(+) | Color2 [Da | 1 | 4 |
| | 0 | NaN | Q15629 | Q15629 | TRAM1 | Translocat | Color2 [Da | 1 | 4 |
| + | 2.820113 | -1.89319 | Q15631 | Q15631 | TSN | Translin | Color2 [Da | 1 | 4 |
| | 0.38853 | 0.206015 | Q15633 | Q15633 | TARBP2 | RISC-loadi | Color2 [Da | 1 | 4 |
| + | 3.559226 | 1.199783 | Q15637 | Q15637 | SF1 | Splicing fa | Color2 [Da | 1 | 4 |
| + | 1.607363 | -0.75931 | Q15642 | Q15642 | TRIP10 | Cdc42-inte | Color2 [Da | 1 | 4 |
| | 0.34534 | -0.13305 | Q15643 | Q15643 | TRIP11 | Thyroid re | Color2 [Da | 1 | 4 |
| | 0 | NaN | Q15645 | Q15645 | TRIP13 | Pachytene | Color2 [Da | 1 | 4 |
| + | 2.311043 | -1.04329 | Q15648 | Q15648 | MED1 | Mediator ( | Color2 [Da | 1 | 4 |
| + | 1.38711 | -0.25751 | Q15650 | Q15650 | TRIP4 | Activating | Color2 [Da | 1 | 4 |
| | 0.434253 | 0.75151 | Q15651 | Q15651 | HMGN3 | High mobi | Color2 [Da | 1 | 4 |
| | 0.492335 | -0.22481 | Q15653 | Q15653 | NFKBIB | NF-kappa- | Color2 [Da | 1 | 4 |
| + | 1.409441 | 1.168866 | Q15654 | Q15654 | TRIP6 | Thyroid re | Color2 [Da | 1 | 4 |
| + | 2.34131 | 1.488115 | Q15691 | Q15691 | MAPRE1 | Microtubu | Color2 [Da | 1 | 4 |
| + | 1.815721 | 0.723462 | Q15700 | Q15700 | DLG2 | Disks large | Color2 [Da | 1 | 4 |
| + | 3.120958 | 3.243686 | Q15714 | Q15714 | TSC22D1 | TSC22 don | Color2 [Da | 1 | 4 |
| + | 2.196269 | 0.824256 | Q15717 | Q15717 | ELAVL1 | ELAV-like p | Color2 [Da | 1 | 4 |
| | 0.20779 | 0.213093 | Q15738 | Q15738 | NSDHL | Sterol-4-al | Color2 [Da | 1 | 4 |
| | 0.228291 | -0.20605 | Q15742 | Q15742 | NAB2 | NGFI-A-bir | Color2 [Da | 1 | 4 |
| | 0 | NaN | Q15746 | Q15746 | MYLK | Myosin ligl | Color2 [Da | 1 | 4 |
| | 0.573943 | -0.10182 | Q15750 | Q15750 | TAB1 | TGF-beta-a | Color2 [Da | 1 | 4 |
| + | 0.790411 | 1.704021 | Q15751 | Q15751 | HERC1 | Probable E | Color2 [Da | 1 | 4 |
| + | 2.200123 | -1.85465 | Q15758 | Q15758 | SLC1A5 | Neutral an | Color2 [Da | 1 | 4 |
| + | 1.474012 | 0.681967 | Q15768 | Q15768 | EFNB3 | Ephrin-B3 | Color2 [Da | 1 | 4 |
| | 0.039346 | 0.056228 | Q15771 | Q15771 | RAB30 | Ras-relate | Color2 [Da | 1 | 4 |
| + | 2.157681 | 0.70976 | Q15773 | Q15773 | MLF2 | Myeloid le | Color2 [Da | 1 | 4 |
| | 0.554945 | -0.35549 | Q15776 | Q15776 | ZKSCAN8 | Zinc finger | Color2 [Da | 1 | 4 |
| | 0 | NaN | Q15777 | Q15777 | MPPED2 | Metalloph | Color2 [Da | 1 | 4 |
| | 0.751121 | 0.376885 | Q15785 | Q15785 | TOMM34 | Mitochonc | Color2 [Da | 1 | 4 |
| | 0.44809 | 0.277464 | Q15796 | Q15796 | SMAD2 | Mothers a | Color2 [Da | 1 | 4 |
| + | 2.610368 | -1.65823 | Q15800 | Q15800 | MSMO1 | Methylste | Color2 [Da | 1 | 4 |
| + | 2.319935 | 0.717222 | Q15811 | Q15811 | ITSN1 | Intersectin | Color2 [Da | 1 | 4 |
| + | 0.787175 | -0.49197 | Q15813 | Q15813 | TBCE | Tubulin-sp | Color2 [Da | 1 | 4 |
| + | 3.165154 | 0.949092 | Q15814 | Q15814 | TBCC | Tubulin-sp | Color2 [Da | 1 | 4 |
| + | 3.113391 | 1.602295 | Q15819 | Q15819 | UBE2V2 | Ubiquitin-( | Color2 [Da | 1 | 4 |
| | 0 | NaN | Q15828 | Q15828 | CST6 | Cystatin-M | Color2 [Da | 1 | 4 |
| | 0 | -1.14682 | Q15833 | Q15833 | STXBP2 | Syntaxin-b | Color2 [Da | 1 | 4 |
| | 0 | 4.644248 | Q15836 | Q15836 | VAMP3 | Vesicle-ass | Color2 [Da | 1 | 4 |
| + | 2.714591 | 2.533958 | Q15843 | Q15843 | NEDD8 | NEDD8 | Color2 [Da | 1 | 4 |
| + | 1.833248 | 0.778978 | Q15853 | Q15853 | USF2 | Upstream | Color2 [Da | 1 | 4 |
| + | 2.944543 | 1.363561 | Q15904 | Q15904 | ATP6AP1 | V-type pro | Color2 [Da | 1 | 4 |
| + | 4.055428 | 1.684659 | Q15907 | Q15907 | RAB11B | Ras-relate | Color2 [Da | 1 | 4 |

| | | | | | | | | | |
|---|---|---|---|---|---|---|---|---|---|
| | 0 | NaN | Q15910 | Q15910 | EZH2 | Histone-ly: | Color2 [Da | 1 | 4 |
| + | 1.677915 | -0.96671 | Q15911 | Q15911 | ZFHX3 | Zinc finger | Color2 [Da | 1 | 4 |
| + | 3.329155 | 2.671086 | Q15942 | Q15942 | ZYX | Zyxin | Color2 [Da | 1 | 4 |
| | 0.021996 | -0.0156 | Q16134 | Q16134 | ETFDH | Electron tr | Color2 [Da | 1 | 4 |
| + | 2.770184 | 4.03751 | Q16143 | Q16143 | SNCB | Beta-synu( | Color2 [Da | 1 | 4 |
| + | 2.222833 | -0.57011 | Q16181 | Q16181 | SEPTIN7 | Septin-7 | Color2 [Da | 1 | 4 |
| + | 1.718129 | 0.877817 | Q16186 | Q16186 | ADRM1 | Proteasom | Color2 [Da | 1 | 4 |
| + | 2.803511 | 0.540243 | Q16204 | Q16204 | CCDC6 | Coiled-coil | Color2 [Da | 1 | 4 |
| + | 1.577522 | -1.33001 | Q16222 | Q16222 | UAP1 | UDP-N-ace | Color2 [Da | 1 | 4 |
| + | 1.839353 | -1.20748 | Q16270 | Q16270 | IGFBP7 | Insulin-like | Color2 [Da | 1 | 4 |
| + | 3.787343 | 1.373444 | Q16352 | Q16352 | INA | Alpha-inte | Color2 [Da | 1 | 4 |
| + | 3.21699 | -1.57805 | Q16378 | Q16378 | PRR4 | Proline-ric | Color2 [Da | 1 | 4 |
| | 0 | 0.573461 | Q16401 | Q16401 | PSMD5 | 26S protea | Color2 [Da | 1 | 4 |
| + | 2.846204 | -1.27652 | Q16512 | Q16512 | PKN1 | Serine/thr | Color2 [Da | 1 | 4 |
| + | 3.300161 | -0.59877 | Q16513 | Q16513 | PKN2 | Serine/thr | Color2 [Da | 1 | 4 |
| + | 1.092031 | -0.43158 | Q16514 | Q16514 | TAF12 | Transcripti | Color2 [Da | 1 | 4 |
| | 0 | NaN | Q16518 | Q16518 | RPE65 | Retinoid is | Color2 [Da | 1 | 4 |
| + | 3.04565 | -0.75082 | Q16527 | Q16527 | CSRP2 | Cysteine a | Color2 [Da | 1 | 4 |
| + | 3.707135 | -1.24996 | Q16531 | Q16531 | DDB1 | DNA dama | Color2 [Da | 1 | 4 |
| + | 3.60036 | -1.33332 | Q16537 | Q16537 | PPP2R5E | Serine/thr | Color2 [Da | 1 | 4 |
| | 0 | NaN | Q16539 | Q16539 | MAPK14 | Mitogen-a | Color2 [Da | 1 | 4 |
| + | 3.000248 | 2.297294 | Q16540 | Q16540 | MRPL23 | 39S riboso | Color2 [Da | 1 | 4 |
| | 1.589285 | -0.16551 | Q16543 | Q16543 | CDC37 | Hsp90 co-( | Color2 [Da | 1 | 4 |
| + | 2.30266 | 0.457322 | Q16555 | Q16555 | DPYSL2 | Dihydropy | Color2 [Da | 1 | 4 |
| + | 0.79655 | 1.00081 | Q16566 | Q16566 | CAMK4 | Calcium/ca | Color2 [Da | 1 | 4 |
| | 0.308361 | 0.153817 | Q16576 | Q16576 | RBBP7 | Histone-bi | Color2 [Da | 1 | 4 |
| | 0.059371 | 0.0393 | Q16594 | Q16594 | TAF9 | Transcripti | Color2 [Da | 1 | 4 |
| + | 3.375277 | 2.250213 | Q16595 | Q16595 | FXN | Frataxin, n | Color2 [Da | 1 | 4 |
| | 0 | 1.10845 | Q16610 | Q16610 | ECM1 | Extracellul | Color2 [Da | 1 | 4 |
| + | 1.813567 | -0.50336 | Q16620 | Q16620 | NTRK2 | BDNF/NT-: | Color2 [Da | 1 | 4 |
| + | 1.167907 | 0.411847 | Q16623 | Q16623 | STX1A | Syntaxin-1 | Color2 [Da | 1 | 4 |
| | 0.755563 | 0.306381 | Q16625 | Q16625 | OCLN | Occludin | Color2 [Da | 1 | 4 |
| + | 3.296319 | 3.020858 | Q16626 | Q16626 | MEA1 | Male-enha | Color2 [Da | 1 | 4 |
| + | 1.861718 | 0.737473 | Q16629 | Q16629 | SRSF7 | Serine/arg | Color2 [Da | 1 | 4 |
| + | 2.875471 | 0.723207 | Q16630 | Q16630 | CPSF6 | Cleavage a | Color2 [Da | 1 | 4 |
| + | 1.808272 | 0.824107 | Q16637 | Q16637 | SMN2 | Survival m | Color2 [Da | 1 | 4 |
| + | 3.779635 | 0.803468 | Q16643 | Q16643 | DBN1 | Drebrin | Color2 [Da | 1 | 4 |
| | 0.600849 | 1.533658 | Q16656 | Q16656 | NRF1 | Nuclear re | Color2 [Da | 1 | 4 |
| + | 1.746089 | -0.54647 | Q16658 | Q16658 | FSCN1 | Fascin | Color2 [Da | 1 | 4 |
| + | 1.841556 | 1.577561 | Q16666 | Q16666 | IFI16 | Gamma-in | Color2 [Da | 1 | 4 |
| + | 1.430633 | 0.248904 | Q16698 | Q16698 | DECR1 | 2,4-dienoy | Color2 [Da | 1 | 4 |
| | 0.407951 | 0.444358 | Q16718 | Q16718 | NDUFA5 | NADH deh | Color2 [Da | 1 | 4 |
| + | 1.415374 | -0.75021 | Q16720 | Q16720 | ATP2B3 | Plasma me | Color2 [Da | 1 | 4 |
| + | 3.104199 | 0.955763 | Q16740 | Q16740 | CLPP | ATP-deper | Color2 [Da | 1 | 4 |
| + | 1.303901 | -0.83502 | Q16762 | Q16762 | TST | Thiosulfate | Color2 [Da | 1 | 4 |
| + | 1.991456 | 0.623291 | Q16763 | Q16763 | UBE2S | Ubiquitin-( | Color2 [Da | 1 | 4 |
| + | 1.602058 | 0.52647 | Q16773 | Q16773 | KYAT1 | Kynurenin | Color2 [Da | 1 | 4 |

| | | | | | | | | | |
|---|---|---|---|---|---|---|---|---|---|
| + | 3.154027 | 1.319291 | Q16774 | Q16774 | GUK1 | Guanylate | Color2 [Da | 1 | 4 |
| + | 3.993641 | 1.381434 | Q16775 | Q16775 | HAGH | Hydroxyac | Color2 [Da | 1 | 4 |
| | 0.254746 | 0.52711 | Q16777 | Q16777 | H2AC20 | Histone H2 | Color2 [Da | 1 | 4 |
| + | 2.375301 | 0.721019 | Q16795 | Q16795 | NDUFA9 | NADH deh | Color2 [Da | 1 | 4 |
| + | 2.854052 | 1.670421 | Q16799 | Q16799 | RTN1 | Reticulon-1 | Color2 [Da | 1 | 4 |
| + | 1.101472 | -1.49038 | Q16822 | Q16822 | PCK2 | Phosphoe | Color2 [Da | 1 | 4 |
| + | 0.915809 | -0.6887 | Q16827 | Q16827 | PTPRO | Receptor-t | Color2 [Da | 1 | 4 |
| + | 3.115537 | 1.673319 | Q16836 | Q16836 | HADH | Hydroxyac | Color2 [Da | 1 | 4 |
| | 0 | -0.43132 | Q16842 | Q16842 | ST3GAL2 | CMP-N-ac | Color2 [Da | 1 | 4 |
| + | 1.358399 | 1.045446 | Q16849 | Q16849 | PTPRN | Receptor-t | Color2 [Da | 1 | 4 |
| + | 1.55507 | -1.23322 | Q16850 | Q16850 | CYP51A1 | Lanosterol | Color2 [Da | 1 | 4 |
| + | 1.850235 | 0.39839 | Q16851 | Q16851 | UGP2 | UTP--gluc | Color2 [Da | 1 | 4 |
| | 0.331729 | -0.15601 | Q16854 | Q16854 | DGUOK | Deoxyguar | Color2 [Da | 1 | 4 |
| + | 2.879446 | 1.731179 | Q16864 | Q16864 | ATP6V1F | V-type pro | Color2 [Da | 1 | 4 |
| + | 0.760502 | 0.808254 | Q16878 | Q16878 | CDO1 | Cysteine d | Color2 [Da | 1 | 4 |
| | 0.004249 | 0.001239 | Q16881 | Q16881 | TXNRD1 | Thioredoxi | Color2 [Da | 1 | 4 |
| | 0.217498 | 0.17761 | Q16890 | Q16890 | TPD52L1 | Tumor pro | Color2 [Da | 1 | 4 |
| + | 4.710502 | 1.044689 | Q16891 | Q16891 | IMMT | MICOS cor | Color2 [Da | 1 | 4 |
| + | 2.293517 | 1.099286 | Q17R31 | Q17R31 | TATDN3 | Putative d | Color2 [Da | 1 | 4 |
| | 0 | 1.700104 | Q17R89 | Q17R89 | ARHGAP44 | Rho GTPas | Color2 [Da | 1 | 4 |
| | 0.159947 | -0.12888 | Q17RY0 | Q17RY0 | CPEB4 | Cytoplasm | Color2 [Da | 1 | 4 |
| + | 1.275226 | -0.53858 | Q1ED39 | Q1ED39 | KNOP1 | Lysine-rich | Color2 [Da | 1 | 4 |
| + | 3.481833 | -0.68659 | Q1KMD3 | Q1KMD3 | HNRNPUL2 | Heterogen | Color2 [Da | 1 | 4 |
| | 0.122411 | 0.112438 | Q1L5Z9 | Q1L5Z9 | LONRF2 | LON pepti | Color2 [Da | 1 | 4 |
| | 0.586798 | 0.493751 | Q24JP5 | Q24JP5 | TMEM132 | Transmem | Color2 [Da | 1 | 4 |
| | 0 | -2.52609 | Q27J81 | Q27J81 | INF2 | Inverted fo | Color2 [Da | 1 | 4 |
| + | 5.169251 | -1.97133 | Q29RF7 | Q29RF7 | PDS5A | Sister chro | Color2 [Da | 1 | 4 |
| + | 1.449913 | -1.45269 | Q2KHR3 | Q2KHR3 | QSER1 | Glutamine | Color2 [Da | 1 | 4 |
| + | 1.36589 | -2.175 | Q2M1K9 | Q2M1K9 | ZNF423 | Zinc finger | Color2 [Da | 1 | 4 |
| + | 1.071124 | 0.330157 | Q2M2I8 | Q2M2I8 | AAK1 | AP2-associ | Color2 [Da | 1 | 4 |
| + | 2.401216 | 1.722497 | Q2M2W7 | Q2M2W7 | C17orf58 | UPF0450 p | Color2 [Da | 1 | 4 |
| + | 2.463991 | -1.60054 | Q2M389 | Q2M389 | WASHC4 | WASH con | Color2 [Da | 1 | 4 |
| + | 2.342814 | 1.438347 | Q2MJR0 | Q2MJR0 | SPRED3 | Sprouty-re | Color2 [Da | 1 | 4 |
| | 0 | NaN | Q2MV58 | Q2MV58 | TCTN1 | Tectonic-1 | Color2 [Da | 1 | 4 |
| | 0 | NaN | Q2NKX8 | Q2NKX8 | ERCC6L | DNA excisi | Color2 [Da | 1 | 4 |
| | 0.526803 | -0.22625 | Q2NL82 | Q2NL82 | TSR1 | Pre-rRNA-p | Color2 [Da | 1 | 4 |
| + | 2.367138 | -1.17012 | Q2PZI1 | Q2PZI1 | DPY19L1 | Probable C | Color2 [Da | 1 | 4 |
| | 0 | NaN | Q2Q1W2 | Q2Q1W2 | TRIM71 | E3 ubiquiti | Color2 [Da | 1 | 4 |
| | 0 | NaN | Q2T9J0 | Q2T9J0 | TYSND1 | Peroxisom | Color2 [Da | 1 | 4 |
| + | 0.892585 | -0.48337 | Q2TAL8 | Q2TAL8 | QRICH1 | Transcripti | Color2 [Da | 1 | 4 |
| + | 1.639687 | -0.56938 | Q2TAY7 | Q2TAY7 | SMU1 | WD40 rep | Color2 [Da | 1 | 4 |
| | 0 | NaN | Q2TAZ0 | Q2TAZ0 | ATG2A | Autophagy | Color2 [Da | 1 | 4 |
| | 0 | NaN | Q2TBE0 | Q2TBE0 | CWF19L2 | CWF19-lik | Color2 [Da | 1 | 4 |
| | 0 | NaN | Q2VPB7 | Q2VPB7 | AP5B1 | AP-5 comp | Color2 [Da | 1 | 4 |
| | 0 | NaN | Q2VWP7 | Q2VWP7 | PRTG | Protogenir | Color2 [Da | 1 | 4 |
| + | 2.449846 | -0.64834 | Q2Y0W8 | Q2Y0W8 | SLC4A8 | Electroneu | Color2 [Da | 1 | 4 |
| + | 2.042469 | 1.111988 | Q32MZ4 | Q32MZ4 | LRRFIP1 | Leucine-ric | Color2 [Da | 1 | 4 |

| | | | | | | | | | |
|---|---|---|---|---|---|---|---|---|---|
| + | 0.990561 | 0.5537 | Q32NB8 | Q32NB8 | PGS1 | CDP-diacyl | Color2 [Da | 1 | 4 |
| + | 2.42084 | -1.10894 | Q32P28 | Q32P28 | P3H1 | Prolyl 3-hy | Color2 [Da | 1 | 4 |
| + | 0.891932 | 0.753912 | Q32P41 | Q32P41 | TRMT5 | tRNA (guai | Color2 [Da | 1 | 4 |
| + | 1.59106 | -1.03264 | Q3B726 | Q3B726 | POLR1F | DNA-direc | Color2 [Da | 1 | 4 |
| | 0 | NaN | Q3B7J2 | Q3B7J2 | GFOD2 | Glucose-fr | Color2 [Da | 1 | 4 |
| | 0 | -0.97963 | Q3B7T1 | Q3B7T1 | EDRF1 | Erythroid ( | Color2 [Da | 1 | 4 |
| + | 3.296803 | 1.58478 | Q3KQU3 | Q3KQU3 | MAP7D1 | MAP7 don | Color2 [Da | 1 | 4 |
| | 0 | NaN | Q3L8U1 | Q3L8U1 | CHD9 | Chromodo | Color2 [Da | 1 | 4 |
| + | 2.055989 | -0.69472 | Q3LXA3 | Q3LXA3 | TKFC | Triokinase, | Color2 [Da | 1 | 4 |
| + | 0.745353 | 0.942187 | Q3MHD2 | Q3MHD2 | LSM12 | Protein LSI | Color2 [Da | 1 | 4 |
| + | 1.738586 | -0.82019 | Q3SXM5 | Q3SXM5 | HSDL1 | Inactive hy | Color2 [Da | 1 | 4 |
| | 0 | NaN | Q3SY84 | Q3SY84 | KRT71 | Keratin, ty | Color2 [Da | 1 | 4 |
| | 0.004677 | 0.001425 | Q3V6T2 | Q3V6T2 | CCDC88A | Girdin | Color2 [Da | 1 | 4 |
| + | 3.022326 | 0.981053 | Q3YEC7 | Q3YEC7 | RABL6 | Rab-like pr | Color2 [Da | 1 | 4 |
| | 0.210242 | -0.24239 | Q3ZAQ7 | Q3ZAQ7 | VMA21 | Vacuolar A | Color2 [Da | 1 | 4 |
| + | 2.453884 | 4.489401 | Q3ZCM7 | Q3ZCM7 | TUBB8 | Tubulin be | Color2 [Da | 1 | 4 |
| | 0.061949 | -0.03352 | Q3ZCQ8 | Q3ZCQ8 | TIMM50 | Mitochonc | Color2 [Da | 1 | 4 |
| + | 1.8165 | 1.135743 | Q3ZCW2 | Q3ZCW2 | LGALSL | Galectin-re | Color2 [Da | 1 | 4 |
| + | 1.044827 | -0.48522 | Q495W5 | Q495W5 | FUT11 | Alpha-(1,3 | Color2 [Da | 1 | 4 |
| + | 1.213607 | 0.985512 | Q496J9 | Q496J9 | SV2C | Synaptic v | Color2 [Da | 1 | 4 |
| + | 2.379362 | -0.72945 | Q49A26 | Q49A26 | GLYR1 | Putative o: | Color2 [Da | 1 | 4 |
| | 0 | NaN | Q49AR2 | Q49AR2 | C5orf22 | UPF0489 p | Color2 [Da | 1 | 4 |
| | 0 | -1.59049 | Q4AE62 | Q4AE62 | GTDC1 | Glycosyltra | Color2 [Da | 1 | 4 |
| + | 2.580499 | 0.757404 | Q4G0F5 | Q4G0F5 | VPS26B | Vacuolar p | Color2 [Da | 1 | 4 |
| + | 1.012592 | -0.68463 | Q4G0I0 | Q4G0I0 | CCSMST1 | Protein CC | Color2 [Da | 1 | 4 |
| | 0.091066 | -0.05362 | Q4G0J3 | Q4G0J3 | LARP7 | La-related | Color2 [Da | 1 | 4 |
| + | 1.771297 | 0.909868 | Q4G0N4 | Q4G0N4 | NADK2 | NAD kinas | Color2 [Da | 1 | 4 |
| | 0.380221 | -0.37842 | Q4G0X4 | Q4G0X4 | KCTD21 | BTB/POZ d | Color2 [Da | 1 | 4 |
| | 0 | -1.485 | Q4G148 | Q4G148 | GXYLT1 | Glucoside | Color2 [Da | 1 | 4 |
| | 0.01142 | -0.01196 | Q4G176 | Q4G176 | ACSF3 | Malonate- | Color2 [Da | 1 | 4 |
| + | 1.133666 | -0.7278 | Q4J6C6 | Q4J6C6 | PREPL | Prolyl endo | Color2 [Da | 1 | 4 |
| + | 3.076149 | 1.668521 | Q4KMP7 | Q4KMP7 | TBC1D10B | TBC1 dom | Color2 [Da | 1 | 4 |
| | 0.083075 | 0.217719 | Q4LE39 | Q4LE39 | ARID4B | AT-rich int | Color2 [Da | 1 | 4 |
| + | 0.982092 | 0.785953 | Q4U2R6 | Q4U2R6 | MRPL51 | 39S riboso | Color2 [Da | 1 | 4 |
| | 1.009394 | 0.128775 | Q4V328 | Q4V328 | GRIPAP1 | GRIP1-asso | Color2 [Da | 1 | 4 |
| + | 1.664896 | 1.658773 | Q4VC05 | Q4VC05 | BCL7A | B-cell CLL/ | Color2 [Da | 1 | 4 |
| + | 1.50082 | 0.910576 | Q4VC31 | Q4VC31 | MIX23 | Protein MI | Color2 [Da | 1 | 4 |
| + | 0.840034 | -0.72478 | Q4VCS5 | Q4VCS5 | AMOT | Angiomoti | Color2 [Da | 1 | 4 |
| | 0 | -0.92168 | Q4ZIN3 | Q4ZIN3 | TMEM259 | Membralir | Color2 [Da | 1 | 4 |
| | 0 | NaN | Q504Q3 | Q504Q3 | PAN2 | PAN2-PAN | Color2 [Da | 1 | 4 |
| | 0.842453 | -0.21556 | Q52LJ0 | Q52LJ0 | FAM98B | Protein FA | Color2 [Da | 1 | 4 |
| | 0 | NaN | Q52MB2 | Q52MB2 | CCDC184 | Coiled-coil | Color2 [Da | 1 | 4 |
| + | 2.36098 | -2.07407 | Q53EL6 | Q53EL6 | PDCD4 | Programm | Color2 [Da | 1 | 4 |
| | 0 | NaN | Q53EP0 | Q53EP0 | FNDC3B | Fibronectii | Color2 [Da | 1 | 4 |
| + | 2.362333 | 1.416749 | Q53ET0 | Q53ET0 | CRTC2 | CREB-regu | Color2 [Da | 1 | 4 |
| + | 0.833491 | 0.540194 | Q53F19 | Q53F19 | NCBP3 | Nuclear ca | Color2 [Da | 1 | 4 |
| | 0.116591 | -0.13054 | Q53FP2 | Q53FP2 | TMEM35A | Novel acet | Color2 [Da | 1 | 4 |

| | | | | | | | | | |
|---|---|---|---|---|---|---|---|---|---|
| | 0.111974 | -0.06015 | Q53FT3 | Q53FT3 | HIKESHI | Protein Hil | Color2 [Da | 1 | 4 |
| + | 4.188042 | 3.746186 | Q53GG5 | Q53GG5 | PDLIM3 | PDZ and LI | Color2 [Da | 1 | 4 |
| + | 1.693689 | 0.36603 | Q53GL0 | Q53GL0 | PLEKHO1 | Pleckstrin | Color2 [Da | 1 | 4 |
| | 0.208433 | -0.03266 | Q53GQ0 | Q53GQ0 | HSD17B12 | Very-long- | Color2 [Da | 1 | 4 |
| + | 1.852921 | -0.58865 | Q53GS7 | Q53GS7 | GLE1 | mRNA exp | Color2 [Da | 1 | 4 |
| + | 1.855889 | -1.18533 | Q53GS9 | Q53GS9 | USP39 | U4/U6.U5 | Color2 [Da | 1 | 4 |
| | 0 | -1.85247 | Q53GT1 | Q53GT1 | KLHL22 | Kelch-like | Color2 [Da | 1 | 4 |
| | 0.835202 | 0.378559 | Q53H12 | Q53H12 | AGK | Acylglycer | Color2 [Da | 1 | 4 |
| | 0.060968 | 0.012854 | Q53H47 | Q53H47 | SETMAR | Histone-ly | Color2 [Da | 1 | 4 |
| + | 2.186392 | 0.653514 | Q53H82 | Q53H82 | LACTB2 | Endoribon | Color2 [Da | 1 | 4 |
| | 0 | -0.56553 | Q53H96 | Q53H96 | PYCR3 | Pyrroline-5 | Color2 [Da | 1 | 4 |
| + | 1.379881 | 0.550973 | Q53HC0 | Q53HC0 | CCDC92 | Coiled-coil | Color2 [Da | 1 | 4 |
| + | 2.553004 | 0.829193 | Q53HC9 | Q53HC9 | EIPR1 | EARP and ( | Color2 [Da | 1 | 4 |
| | 0.658087 | -0.37099 | Q53HV7 | Q53HV7 | SMUG1 | Single-stra | Color2 [Da | 1 | 4 |
| | 0.45174 | 0.212233 | Q53LP3 | Q53LP3 | SOWAHC | Ankyrin re | Color2 [Da | 1 | 4 |
| | 0 | 0.574466 | Q53QV2 | Q53QV2 | LBH | Protein LB | Color2 [Da | 1 | 4 |
| | 0 | -0.45817 | Q53RT3 | Q53RT3 | ASPRV1 | Retroviral- | Color2 [Da | 1 | 4 |
| + | 1.85626 | 0.655281 | Q53S08 | Q53S08 | RAB6D | Ras-relate | Color2 [Da | 1 | 4 |
| + | 0.944002 | 0.972329 | Q53S33 | Q53S33 | BOLA3 | BolA-like p | Color2 [Da | 1 | 4 |
| + | 1.133264 | -0.80669 | Q53SF7 | Q53SF7 | COBLL1 | Cordon-ble | Color2 [Da | 1 | 4 |
| + | 1.770385 | 0.948825 | Q53T59 | Q53T59 | HS1BP3 | HCLS1-bin | Color2 [Da | 1 | 4 |
| | 0 | -1.19024 | Q562E7 | Q562E7 | WDR81 | WD repeat | Color2 [Da | 1 | 4 |
| + | 2.88863 | -0.28334 | Q562R1 | Q562R1 | ACTBL2 | Beta-actin | Color2 [Da | 1 | 4 |
| | 0.897461 | -0.3255 | Q567U6 | Q567U6 | CCDC93 | Coiled-coil | Color2 [Da | 1 | 4 |
| | 0 | -0.43599 | Q56VL3 | Q56VL3 | OCIAD2 | OCIA doma | Color2 [Da | 1 | 4 |
| | 0 | 0.00172 | Q587I9 | Q587I9 | SFT2D3 | Vesicle tra | Color2 [Da | 1 | 4 |
| + | 1.481033 | -0.59057 | Q58EX2 | Q58EX2 | SDK2 | Protein sid | Color2 [Da | 1 | 4 |
| | 0.010581 | 0.022911 | Q58FF6 | Q58FF6 | HSP90AB4 | Putative h | Color2 [Da | 1 | 4 |
| | 0.219963 | -0.10431 | Q58FF8 | Q58FF8 | HSP90AB2 | Putative h | Color2 [Da | 1 | 4 |
| | 0.034357 | -0.09772 | Q58FG0 | Q58FG0 | HSP90AA5 | Putative h | Color2 [Da | 1 | 4 |
| + | 1.729426 | -1.11229 | Q58FG1 | Q58FG1 | HSP90AA4 | Putative h | Color2 [Da | 1 | 4 |
| | 0.200613 | 0.169054 | Q59EK9 | Q59EK9 | RUNDC3A | RUN doma | Color2 [Da | 1 | 4 |
| | 0 | -0.27117 | Q5BJD5 | Q5BJD5 | TMEM41B | Transmem | Color2 [Da | 1 | 4 |
| + | 2.856589 | -1.68554 | Q5BJF2 | Q5BJF2 | TMEM97 | Sigma intr | Color2 [Da | 1 | 4 |
| | 0 | NaN | Q5BJF6 | Q5BJF6 | ODF2 | Outer den | Color2 [Da | 1 | 4 |
| | 0 | 0.386772 | Q5BJH2 | Q5BJH2 | TMEM128 | Transmem | Color2 [Da | 1 | 4 |
| | 0 | 0.501774 | Q5BJH7 | Q5BJH7 | YIF1B | Protein YIF | Color2 [Da | 1 | 4 |
| | 0 | -0.84438 | Q5BKT4;Q | Q5BKT4;Q | ALG10;ALG | Dol-P-Glc:( | Color2 [Da | 1 | 4 |
| + | 1.172128 | 0.454114 | Q5BKY9 | Q5BKY9 | FAM133B | Protein FA | Color2 [Da | 1 | 4 |
| + | 1.616499 | -0.45942 | Q5BKZ1 | Q5BKZ1 | ZNF326 | DBIRD con | Color2 [Da | 1 | 4 |
| + | 1.575575 | 0.887231 | Q5D862 | Q5D862 | FLG2 | Filaggrin-2 | Color2 [Da | 1 | 4 |
| | 0.165698 | 0.07678 | Q5EB52 | Q5EB52 | MEST | Mesoderm | Color2 [Da | 1 | 4 |
| + | 2.127078 | 0.281175 | Q5EBL4 | Q5EBL4 | RILPL1 | RILP-like p | Color2 [Da | 1 | 4 |
| + | 1.353441 | 0.420491 | Q5EBL8 | Q5EBL8 | PDZD11 | PDZ doma | Color2 [Da | 1 | 4 |
| | 0 | NaN | Q5GH72 | Q5GH72 | XKR7 | XK-related | Color2 [Da | 1 | 4 |
| | 0 | -0.38056 | Q5GH76 | Q5GH76 | XKR4 | XK-related | Color2 [Da | 1 | 4 |
| | 0.273567 | -0.32608 | Q5GLZ8 | Q5GLZ8 | HERC4 | Probable E | Color2 [Da | 1 | 4 |

| | | | | | | | | | |
|---|---|---|---|---|---|---|---|---|---|
| | 0 | 3.099983 | Q5H9L2 | Q5H9L2 | TCEAL5 | Transcripti | Color2 [Da | 1 | 4 |
| + | 1.497095 | -0.69583 | Q5H9R7 | Q5H9R7 | PPP6R3 | Serine/thr | Color2 [Da | 1 | 4 |
| + | 2.312039 | 0.631822 | Q5HYI8 | Q5HYI8 | RABL3 | Rab-like pr | Color2 [Da | 1 | 4 |
| | 0.193242 | 0.126548 | Q5HYJ3 | Q5HYJ3 | FAM76B | Protein FA | Color2 [Da | 1 | 4 |
| | 0.661578 | -0.19863 | Q5HYK3 | Q5HYK3 | COQ5 | 2-methoxy | Color2 [Da | 1 | 4 |
| + | 2.828657 | 1.161692 | Q5HYW2 | Q5HYW2 | NHSL2 | NHS-like p | Color2 [Da | 1 | 4 |
| + | 3.045082 | -1.16739 | Q5IJ48 | Q5IJ48 | CRB2 | Protein cru | Color2 [Da | 1 | 4 |
| + | 1.827627 | -1.20397 | Q5J8M3 | Q5J8M3 | EMC4 | ER membr | Color2 [Da | 1 | 4 |
| | 0 | -0.87136 | Q5JPH6 | Q5JPH6 | EARS2 | Probable g | Color2 [Da | 1 | 4 |
| | 0.615448 | -0.52544 | Q5JPI3 | Q5JPI3 | C3orf38 | Uncharact | Color2 [Da | 1 | 4 |
| | 0 | -0.19191 | Q5JPI9 | Q5JPI9 | EEF1AKMT | EEF1A lysin | Color2 [Da | 1 | 4 |
| + | 3.072519 | -0.44519 | Q5JRA6 | Q5JRA6 | MIA3 | Transport | Color2 [Da | 1 | 4 |
| | 0.555048 | -0.45898 | Q5JRX3 | Q5JRX3 | PITRM1 | Presequen | Color2 [Da | 1 | 4 |
| + | 2.489053 | 2.850708 | Q5JS37 | Q5JS37 | NHLRC3 | NHL repea | Color2 [Da | 1 | 4 |
| + | 1.00256 | 0.434584 | Q5JS54 | Q5JS54 | PSMG4 | Proteasom | Color2 [Da | 1 | 4 |
| + | 3.076709 | 0.793818 | Q5JSH3 | Q5JSH3 | WDR44 | WD repeat | Color2 [Da | 1 | 4 |
| | 0.169622 | -0.24673 | Q5JSZ5 | Q5JSZ5 | PRRC2B | Protein PR | Color2 [Da | 1 | 4 |
| + | 1.537805 | -1.14677 | Q5JTH9 | Q5JTH9 | RRP12 | RRP12-like | Color2 [Da | 1 | 4 |
| + | 3.439559 | 2.753273 | Q5JTJ3 | Q5JTJ3 | COA6 | Cytochrom | Color2 [Da | 1 | 4 |
| + | 1.563568 | 0.222516 | Q5JTV8 | Q5JTV8 | TOR1AIP1 | Torsin-1A- | Color2 [Da | 1 | 4 |
| | 0.707227 | -0.31895 | Q5JTZ9 | Q5JTZ9 | AARS2 | Alanine--tR | Color2 [Da | 1 | 4 |
| | 0.035737 | -0.02109 | Q5JU69 | Q5JU69 | TOR2A | Torsin-2A | Color2 [Da | 1 | 4 |
| | 0 | NaN | Q5JU85 | Q5JU85 | IQSEC2 | IQ motif ar | Color2 [Da | 1 | 4 |
| + | 1.882396 | 0.693292 | Q5JUX0 | Q5JUX0 | SPIN3 | Spindlin-3 | Color2 [Da | 1 | 4 |
| | 0 | -0.81463 | Q5JVF3 | Q5JVF3 | PCID2 | PCI domain | Color2 [Da | 1 | 4 |
| | 0.544246 | 1.238454 | Q5JVS0 | Q5JVS0 | HABP4 | Intracellula | Color2 [Da | 1 | 4 |
| + | 1.793419 | -1.48547 | Q5K4L6 | Q5K4L6 | SLC27A3 | Solute carr | Color2 [Da | 1 | 4 |
| | 0 | NaN | Q5KU26 | Q5KU26 | COLEC12 | Collectin-1 | Color2 [Da | 1 | 4 |
| + | 2.906673 | 2.06901 | Q5M775 | Q5M775 | SPECC1 | Cytospin-B | Color2 [Da | 1 | 4 |
| | 0.307788 | 0.85226 | Q5MIZ7 | Q5MIZ7 | PPP4R3B | Serine/thr | Color2 [Da | 1 | 4 |
| | 0.00262 | -0.00302 | Q5MNZ6 | Q5MNZ6 | WDR45B | WD repeat | Color2 [Da | 1 | 4 |
| + | 1.154175 | 0.295102 | Q5PRF9 | Q5PRF9 | SAMD4B | Protein Sm | Color2 [Da | 1 | 4 |
| | 0.695502 | -0.22323 | Q5PSV4 | Q5PSV4 | BRMS1L | Breast can | Color2 [Da | 1 | 4 |
| | 0.040753 | -0.0354 | Q5QJ74 | Q5QJ74 | TBCEL | Tubulin-sp | Color2 [Da | 1 | 4 |
| + | 0.958316 | 0.352947 | Q5QJE6 | Q5QJE6 | DNTTIP2 | Deoxynucl | Color2 [Da | 1 | 4 |
| | 0.353966 | -0.18425 | Q5R372 | Q5R372 | RABGAP1L | Rab GTPas | Color2 [Da | 1 | 4 |
| | 0 | 0.393313 | Q5R3F8 | Q5R3F8 | ELFN2 | Protein ph | Color2 [Da | 1 | 4 |
| + | 1.990361 | -0.83317 | Q5R3I4 | Q5R3I4 | TTC38 | Tetratricop | Color2 [Da | 1 | 4 |
| + | 1.5102 | 0.844804 | Q5RI15 | Q5RI15 | COX20 | Cytochrom | Color2 [Da | 1 | 4 |
| + | 2.134975 | -0.87625 | Q5RKV6 | Q5RKV6 | EXOSC6 | Exosome c | Color2 [Da | 1 | 4 |
| + | 2.546921 | 1.034026 | Q5SGD2 | Q5SGD2 | PPM1L | Protein ph | Color2 [Da | 1 | 4 |
| + | 1.409215 | 0.818812 | Q5SNT2 | Q5SNT2 | TMEM201 | Transmem | Color2 [Da | 1 | 4 |
| | 0 | NaN | Q5SQH8 | Q5SQH8 | C6orf136 | Uncharact | Color2 [Da | 1 | 4 |
| + | 2.67483 | 0.748984 | Q5SQI0 | Q5SQI0 | ATAT1 | Alpha-tubu | Color2 [Da | 1 | 4 |
| + | 2.962133 | -1.36475 | Q5SRE5 | Q5SRE5 | NUP188 | Nucleopor | Color2 [Da | 1 | 4 |
| | 1.036175 | 0.037624 | Q5SSJ5 | Q5SSJ5 | HP1BP3 | Heterochro | Color2 [Da | 1 | 4 |
| | 0 | NaN | Q5ST30 | Q5ST30 | VARS2 | Valine--tRI | Color2 [Da | 1 | 4 |

| | | | | | | | | | |
|---|---|---|---|---|---|---|---|---|---|
| | 0 | NaN | Q5SVZ6 | Q5SVZ6 | ZMYM1 | Zinc finger | Color2 [Da | 1 | 4 |
| + | 3.217281 | 0.661707 | Q5SW79 | Q5SW79 | CEP170 | Centrosom | Color2 [Da | 1 | 4 |
| + | 1.372509 | 1.846245 | Q5SWA1 | Q5SWA1 | PPP1R15B | Protein ph | Color2 [Da | 1 | 4 |
| | 0.440937 | -0.13468 | Q5SWX8 | Q5SWX8 | ODR4 | Protein od | Color2 [Da | 1 | 4 |
| + | 3.077056 | 3.095928 | Q5SXM8 | Q5SXM8 | DNLZ | DNL-type z | Color2 [Da | 1 | 4 |
| | 0.230772 | -0.25675 | Q5SY16 | Q5SY16 | NOL9 | Polynuclec | Color2 [Da | 1 | 4 |
| | 0.628155 | 0.78138 | Q5SYC1 | Q5SYC1 | CLVS2 | Clavesin-2 | Color2 [Da | 1 | 4 |
| | 0 | NaN | Q5SYE7 | Q5SYE7 | NHSL1 | NHS-like p | Color2 [Da | 1 | 4 |
| + | 2.328946 | -0.78428 | Q5SZK8 | Q5SZK8 | FREM2 | FRAS1-rela | Color2 [Da | 1 | 4 |
| | 1.141498 | -0.12541 | Q5SZQ8 | Q5SZQ8 | CELF3 | CUGBP Ela | Color2 [Da | 1 | 4 |
| | 0 | 1.592019 | Q5T013 | Q5T013 | HYI | Putative hy | Color2 [Da | 1 | 4 |
| + | 1.601519 | 0.597862 | Q5T0D9 | Q5T0D9 | TPRG1L | Tumor pro | Color2 [Da | 1 | 4 |
| + | 1.530982 | -0.77475 | Q5T0F9 | Q5T0F9 | CC2D1B | Coiled-coil | Color2 [Da | 1 | 4 |
| + | 2.832881 | 0.945853 | Q5T0N5 | Q5T0N5 | FNBP1L | Formin-bir | Color2 [Da | 1 | 4 |
| | 0.308407 | -0.15092 | Q5T160 | Q5T160 | RARS2 | Probable a | Color2 [Da | 1 | 4 |
| + | 1.881923 | 2.524283 | Q5T1J5;Q9 | Q5T1J5;Q9 | CHCHD2P9 | Putative cc | Color2 [Da | 1 | 4 |
| + | 1.759785 | -0.3309 | Q5T1M5 | Q5T1M5 | FKBP15 | FK506-binc | Color2 [Da | 1 | 4 |
| + | 1.472147 | -1.11581 | Q5T1Q4 | Q5T1Q4 | SLC35F1 | Solute carr | Color2 [Da | 1 | 4 |
| | 0 | NaN | Q5T1V6 | Q5T1V6 | DDX59 | Probable A | Color2 [Da | 1 | 4 |
| + | 0.970893 | 0.6703 | Q5T200 | Q5T200 | ZC3H13 | Zinc finger | Color2 [Da | 1 | 4 |
| | 0 | 0.111298 | Q5T2E6 | Q5T2E6 | ARMH3 | Armadillo- | Color2 [Da | 1 | 4 |
| | 0 | -0.95715 | Q5T2T1 | Q5T2T1 | MPP7 | MAGUK p5 | Color2 [Da | 1 | 4 |
| | 0 | -0.51305 | Q5T3F8 | Q5T3F8 | TMEM63B | CSC1-like p | Color2 [Da | 1 | 4 |
| | 0.76273 | 0.335243 | Q5T3J3 | Q5T3J3 | LRIF1 | Ligand-dep | Color2 [Da | 1 | 4 |
| + | 2.283153 | 1.887601 | Q5T440 | Q5T440 | IBA57 | Putative tr | Color2 [Da | 1 | 4 |
| | 0 | -1.37345 | Q5T447 | Q5T447 | HECTD3 | E3 ubiquiti | Color2 [Da | 1 | 4 |
| | 0 | 0.566797 | Q5T4D3 | Q5T4D3 | TMTC4 | Protein O- | Color2 [Da | 1 | 4 |
| + | 3.012829 | -0.85924 | Q5T4S7 | Q5T4S7 | UBR4 | E3 ubiquiti | Color2 [Da | 1 | 4 |
| + | 2.056905 | -0.80845 | Q5T5P2 | Q5T5P2 | KIAA1217 | Sickle tail p | Color2 [Da | 1 | 4 |
| | 1.004377 | -0.14865 | Q5T5U3 | Q5T5U3 | ARHGAP21 | Rho GTPas | Color2 [Da | 1 | 4 |
| | 0 | NaN | Q5T5X7 | Q5T5X7 | BEND3 | BEN doma | Color2 [Da | 1 | 4 |
| | 0.726621 | -0.25822 | Q5T5Y3 | Q5T5Y3 | CAMSAP1 | Calmodulin | Color2 [Da | 1 | 4 |
| | 0.061363 | -0.03758 | Q5T653 | Q5T653 | MRPL2 | 39S riboso | Color2 [Da | 1 | 4 |
| + | 1.125565 | -0.92938 | Q5T6F2 | Q5T6F2 | UBAP2 | Ubiquitin-a | Color2 [Da | 1 | 4 |
| + | 1.526399 | 0.977799 | Q5T749 | Q5T749 | KPRP | Keratinocy | Color2 [Da | 1 | 4 |
| + | 1.864672 | -0.68049 | Q5T7N2 | Q5T7N2 | L1TD1 | LINE-1 typ | Color2 [Da | 1 | 4 |
| | 0 | 0.687346 | Q5T7P8 | Q5T7P8 | SYT6 | Synaptotag | Color2 [Da | 1 | 4 |
| | 0.075829 | -0.02224 | Q5T7V8 | Q5T7V8 | GORAB | RAB6-inter | Color2 [Da | 1 | 4 |
| | 0.541144 | -0.70711 | Q5T7W0 | Q5T7W0 | ZNF618 | Zinc finger | Color2 [Da | 1 | 4 |
| + | 2.03029 | 0.622581 | Q5T8D3 | Q5T8D3 | ACBD5 | Acyl-CoA-b | Color2 [Da | 1 | 4 |
| | 0.078068 | -0.06068 | Q5T8P6 | Q5T8P6 | RBM26 | RNA-bindir | Color2 [Da | 1 | 4 |
| + | 2.071026 | 1.535314 | Q5T9A4 | Q5T9A4 | ATAD3B | ATPase far | Color2 [Da | 1 | 4 |
| + | 2.071166 | -1.55498 | Q5T9L3 | Q5T9L3 | WLS | Protein wr | Color2 [Da | 1 | 4 |
| + | 1.8394 | -1.31798 | Q5TA45 | Q5TA45 | INTS11 | Integrator | Color2 [Da | 1 | 4 |
| + | 1.186638 | -0.41478 | Q5TAP6 | Q5TAP6 | UTP14C | U3 small n | Color2 [Da | 1 | 4 |
| + | 2.014075 | -1.35148 | Q5TAQ9 | Q5TAQ9 | DCAF8 | DDB1- and | Color2 [Da | 1 | 4 |
| | 0 | -1.42483 | Q5TBA9 | Q5TBA9 | FRY | Protein fur | Color2 [Da | 1 | 4 |

| | | | | | | | | | |
|---|---|---|---|---|---|---|---|---|---|
| | 0.11107 | -0.08832 | Q5TC12 | Q5TC12 | ATPAF1 | ATP syntha | Color2 [Da | 1 | 4 |
| | 0.37476 | -0.18798 | Q5TC82 | Q5TC82 | RC3H1 | Roquin-1 | Color2 [Da | 1 | 4 |
| + | 1.293678 | 1.02003 | Q5TCQ9 | Q5TCQ9 | MAGI3 | Membrane | Color2 [Da | 1 | 4 |
| + | 1.103 | 0.844473 | Q5TCY1 | Q5TCY1 | TTBK1 | Tau-tubuli | Color2 [Da | 1 | 4 |
| | 0.179826 | -0.11366 | Q5TDH0 | Q5TDH0 | DDI2 | Protein DD | Color2 [Da | 1 | 4 |
| + | 1.315641 | 0.745552 | Q5TEC6 | Q5TEC6 | H3-7 | Histone H3 | Color2 [Da | 1 | 4 |
| + | 1.225156 | 0.625569 | Q5TF21 | Q5TF21 | SOGA3 | Protein SO | Color2 [Da | 1 | 4 |
| | 0.439115 | -0.75348 | Q5TFE4 | Q5TFE4 | NT5DC1 | 5'-nucleoti | Color2 [Da | 1 | 4 |
| | 0.420781 | 0.174661 | Q5TGY3 | Q5TGY3 | AHDC1 | AT-hook D | Color2 [Da | 1 | 4 |
| + | 2.676111 | 1.608918 | Q5TGZ0 | Q5TGZ0 | MICOS10 | MICOS cor | Color2 [Da | 1 | 4 |
| + | 1.090422 | -0.48772 | Q5TH69 | Q5TH69 | ARFGEF3 | Brefeldin A | Color2 [Da | 1 | 4 |
| + | 0.773025 | 0.782727 | Q5THJ4 | Q5THJ4 | VPS13D | Intermemb | Color2 [Da | 1 | 4 |
| + | 2.104159 | -1.40822 | Q5TKA1 | Q5TKA1 | LIN9 | Protein lin | Color2 [Da | 1 | 4 |
| + | 1.419482 | -0.22746 | Q5TZA2 | Q5TZA2 | CROCC | Rootletin | Color2 [Da | 1 | 4 |
| | 0.628089 | 0.540176 | Q5TZF3 | Q5TZF3 | ANKRD45 | Ankyrin re | Color2 [Da | 1 | 4 |
| + | 2.863065 | 1.227778 | Q5U5X0 | Q5U5X0 | LYRM7 | Complex II | Color2 [Da | 1 | 4 |
| + | 1.045708 | 1.285189 | Q5UCC4 | Q5UCC4 | EMC10 | ER membr | Color2 [Da | 1 | 4 |
| + | 0.733729 | 0.583567 | Q5VIR6 | Q5VIR6 | VPS53 | Vacuolar p | Color2 [Da | 1 | 4 |
| + | 2.14229 | -0.66186 | Q5VSL9 | Q5VSL9 | STRIP1 | Striatin-int | Color2 [Da | 1 | 4 |
| | 0 | -1.7096 | Q5VST6 | Q5VST6 | ABHD17B | Alpha/beta | Color2 [Da | 1 | 4 |
| | 0.427206 | 0.14275 | Q5VT06 | Q5VT06 | CEP350 | Centrosom | Color2 [Da | 1 | 4 |
| + | 1.025739 | -0.52882 | Q5VT25 | Q5VT25 | CDC42BPA | Serine/thr | Color2 [Da | 1 | 4 |
| + | 1.738907 | 1.19622 | Q5VT52 | Q5VT52 | RPRD2 | Regulation | Color2 [Da | 1 | 4 |
| + | 2.175748 | -1.55625 | Q5VTL8 | Q5VTL8 | PRPF38B | Pre-mRNA | Color2 [Da | 1 | 4 |
| | 0.597679 | -0.08139 | Q5VTR2 | Q5VTR2 | RNF20 | E3 ubiquiti | Color2 [Da | 1 | 4 |
| + | 1.032382 | 0.45048 | Q5VU43 | Q5VU43 | PDE4DIP | Myomegal | Color2 [Da | 1 | 4 |
| | 0 | -2.08544 | Q5VU97 | Q5VU97 | CACHD1 | VWFA and | Color2 [Da | 1 | 4 |
| | 0.496224 | -0.27271 | Q5VUB5 | Q5VUB5 | FAM171A1 | Protein FA | Color2 [Da | 1 | 4 |
| | 0 | NaN | Q5VUD6 | Q5VUD6 | DIPK1B | Divergent | Color2 [Da | 1 | 4 |
| | 0.233534 | -0.25256 | Q5VUJ6 | Q5VUJ6 | LRCH2 | Leucine-ric | Color2 [Da | 1 | 4 |
| | 0 | NaN | Q5VUM1 | Q5VUM1 | SDHAF4 | Succinate | Color2 [Da | 1 | 4 |
| + | 4.138467 | -1.5864 | Q5VV42 | Q5VV42 | CDKAL1 | Threonylca | Color2 [Da | 1 | 4 |
| | 0 | 0.597461 | Q5VVQ6 | Q5VVQ6 | YOD1 | Ubiquitin t | Color2 [Da | 1 | 4 |
| + | 1.62787 | -1.07117 | Q5VW32 | Q5VW32 | BROX | BRO1 dom | Color2 [Da | 1 | 4 |
| | 0 | 0.785221 | Q5VW36 | Q5VW36 | FOCAD | Focadhesin | Color2 [Da | 1 | 4 |
| + | 1.216617 | 0.979394 | Q5VW38 | Q5VW38 | GPR107 | Protein GP | Color2 [Da | 1 | 4 |
| + | 1.18955 | 0.486455 | Q5VWJ9 | Q5VWJ9 | SNX30 | Sorting ne | Color2 [Da | 1 | 4 |
| + | 1.336742 | 1.539806 | Q5VWP3 | Q5VWP3 | MLIP | Muscular L | Color2 [Da | 1 | 4 |
| + | 1.916801 | -1.08132 | Q5VWQ0 | Q5VWQ0 | RSBN1 | Lysine-spe | Color2 [Da | 1 | 4 |
| | 0.130941 | 0.087594 | Q5VWQ8 | Q5VWQ8 | DAB2IP | Disabled h | Color2 [Da | 1 | 4 |
| + | 2.955933 | 1.963976 | Q5VWX1 | Q5VWX1 | KHDRBS2 | KH domain | Color2 [Da | 1 | 4 |
| + | 1.384951 | -0.50948 | Q5VWZ2 | Q5VWZ2 | LYPLAL1 | Lysophosp | Color2 [Da | 1 | 4 |
| + | 3.485321 | -0.95078 | Q5VYK3 | Q5VYK3 | ECPAS | Proteasom | Color2 [Da | 1 | 4 |
| | 0 | NaN | Q5VYS8 | Q5VYS8 | TUT7 | Terminal u | Color2 [Da | 1 | 4 |
| | 0 | NaN | Q5VZ46 | Q5VZ46 | KIAA1614 | Uncharact | Color2 [Da | 1 | 4 |
| | 0 | NaN | Q5VZ89 | Q5VZ89 | DENND4C | DENN dom | Color2 [Da | 1 | 4 |
| | 0.661271 | -0.40665 | Q5VZE5 | Q5VZE5 | NAA35 | N-alpha-ac | Color2 [Da | 1 | 4 |

| | | | | | | | | | |
|---|---|---|---|---|---|---|---|---|---|
| | 0.149462 | 0.277517 | Q5VZK9 | Q5VZK9 | CARMIL1 | F-actin-un | Color2 [Da | 1 | 4 |
| + | 2.377701 | -1.13238 | Q5VZL5 | Q5VZL5 | ZMYM4 | Zinc finger | Color2 [Da | 1 | 4 |
| | 0.012725 | 0.008718 | Q5VZM2 | Q5VZM2 | RRAGB | Ras-relate | Color2 [Da | 1 | 4 |
| | 0 | -0.55857 | Q5W0V3 | Q5W0V3 | FHIP2A | FHF compl | Color2 [Da | 1 | 4 |
| + | 3.361497 | 2.825501 | Q5XKP0 | Q5XKP0 | MICOS13 | MICOS cor | Color2 [Da | 1 | 4 |
| | 0 | NaN | Q5XKR4 | Q5XKR4 | OTP | Homeobo› | Color2 [Da | 1 | 4 |
| | 0 | 0.209626 | Q5XPI4 | Q5XPI4 | RNF123 | E3 ubiquiti | Color2 [Da | 1 | 4 |
| + | 1.084633 | -0.83782 | Q5XUX1 | Q5XUX1 | FBXW9 | F-box/WD | Color2 [Da | 1 | 4 |
| + | 1.066039 | -0.94064 | Q5ZPR3 | Q5ZPR3 | CD276 | CD276 ant | Color2 [Da | 1 | 4 |
| | 0 | NaN | Q63HR2;Q | Q63HR2;Q | TNS2;TNS3 | Tensin-2;T | Color2 [Da | 1 | 4 |
| + | 2.813048 | 1.046217 | Q641Q2 | Q641Q2 | WASHC2A | WASH con | Color2 [Da | 1 | 4 |
| | 0.241327 | -0.37302 | Q641Q3 | Q641Q3 | METRNL | Meteorin-l | Color2 [Da | 1 | 4 |
| | 0 | NaN | Q643R3 | Q643R3 | LPCAT4 | Lysophosp | Color2 [Da | 1 | 4 |
| + | 2.149719 | -1.29583 | Q658Y4 | Q658Y4 | FAM91A1 | Protein FA | Color2 [Da | 1 | 4 |
| | 0 | -0.07484 | Q659C4 | Q659C4 | LARP1B | La-related | Color2 [Da | 1 | 4 |
| + | 2.535402 | -1.39022 | Q66K14 | Q66K14 | TBC1D9B | TBC1 dom | Color2 [Da | 1 | 4 |
| + | 2.836312 | -0.45418 | Q66K74 | Q66K74 | MAP1S | Microtubu | Color2 [Da | 1 | 4 |
| | 0.591548 | -0.18664 | Q66LE6 | Q66LE6 | PPP2R2D | Serine/thr | Color2 [Da | 1 | 4 |
| + | 1.18532 | 0.780149 | Q66PJ3 | Q66PJ3 | ARL6IP4 | ADP-ribosy | Color2 [Da | 1 | 4 |
| + | 2.722284 | 1.457922 | Q674X7 | Q674X7 | KAZN | Kazrin | Color2 [Da | 1 | 4 |
| | 0.671712 | -0.26222 | Q676U5 | Q676U5 | ATG16L1 | Autophagy | Color2 [Da | 1 | 4 |
| + | 1.514766 | 0.857562 | Q684P5 | Q684P5 | RAP1GAP2 | Rap1 GTPa | Color2 [Da | 1 | 4 |
| | 0 | -0.85256 | Q68BL8 | Q68BL8 | OLFML2B | Olfactome | Color2 [Da | 1 | 4 |
| + | 1.521011 | 2.330638 | Q68CP9 | Q68CP9 | ARID2 | AT-rich int | Color2 [Da | 1 | 4 |
| + | 1.229033 | 0.587989 | Q68CQ7 | Q68CQ7 | GLT8D1 | Glycosyltra | Color2 [Da | 1 | 4 |
| + | 3.048128 | 2.105835 | Q68D91 | Q68D91 | MBLAC2 | Acyl-coenz | Color2 [Da | 1 | 4 |
| | 0 | NaN | Q68DK2 | Q68DK2 | ZFYVE26 | Zinc finger | Color2 [Da | 1 | 4 |
| + | 0.916501 | 0.826367 | Q68DU8 | Q68DU8 | KCTD16 | BTB/POZ d | Color2 [Da | 1 | 4 |
| + | 2.848487 | -2.54003 | Q68E01 | Q68E01 | INTS3 | Integrator | Color2 [Da | 1 | 4 |
| + | 2.109494 | -0.79621 | Q68EM7 | Q68EM7 | ARHGAP17 | Rho GTPas | Color2 [Da | 1 | 4 |
| | 0 | NaN | Q69YH5 | Q69YH5 | CDCA2 | Cell divisio | Color2 [Da | 1 | 4 |
| | 0 | 0.443442 | Q69YL0 | Q69YL0 | NCBP2AS2 | Protein NC | Color2 [Da | 1 | 4 |
| + | 1.113967 | -0.62089 | Q69YN4 | Q69YN4 | VIRMA | Protein vir | Color2 [Da | 1 | 4 |
| + | 1.761685 | 0.470528 | Q69YQ0 | Q69YQ0 | SPECC1L | Cytospin-A | Color2 [Da | 1 | 4 |
| | 0 | NaN | Q6AI08 | Q6AI08 | HEATR6 | HEAT repe | Color2 [Da | 1 | 4 |
| + | 3.104729 | 0.939391 | Q6BDS2 | Q6BDS2 | UHRF1BP1 | UHRF1-bin | Color2 [Da | 1 | 4 |
| + | 1.50101 | 0.954953 | Q6DD87 | Q6DD87 | ZNF787 | Zinc finger | Color2 [Da | 1 | 4 |
| | 0 | NaN | Q6DD88 | Q6DD88 | ATL3 | Atlastin-3 | Color2 [Da | 1 | 4 |
| | 0.047164 | 0.021027 | Q6DKI1 | Q6DKI1 | RPL7L1 | 60S riboso | Color2 [Da | 1 | 4 |
| + | 2.686108 | -1.31984 | Q6DKJ4 | Q6DKJ4 | NXN | Nucleored | Color2 [Da | 1 | 4 |
| | 0.440462 | 0.160962 | Q6DKK2 | Q6DKK2 | TTC19 | Tetratrico | Color2 [Da | 1 | 4 |
| + | 1.538367 | 0.578848 | Q6DN90 | Q6DN90 | IQSEC1 | IQ motif a | Color2 [Da | 1 | 4 |
| + | 1.942225 | 2.000171 | Q6F5E8 | Q6F5E8 | CARMIL2 | Capping pr | Color2 [Da | 1 | 4 |
| | 0 | -2.4696 | Q6FHJ7 | Q6FHJ7 | SFRP4 | Secreted fı | Color2 [Da | 1 | 4 |
| | 0.78378 | 0.237729 | Q6FI81 | Q6FI81 | CIAPIN1 | Anamorsir | Color2 [Da | 1 | 4 |
| | 0 | 0.762617 | Q6FIF0 | Q6FIF0 | ZFAND6 | AN1-type z | Color2 [Da | 1 | 4 |
| + | 1.335622 | -0.91699 | Q6GMV2 | Q6GMV2 | SMYD5 | Histone-lys | Color2 [Da | 1 | 4 |

| | | | | | | | | | |
|---|---|---|---|---|---|---|---|---|---|
| | 0.411694 | 0.402939 | Q6GMV3 | Q6GMV3 | PTRHD1 | Putative p | Color2 [Da | 1 | 4 |
| + | 2.085541 | -1.02042 | Q6GQQ9 | Q6GQQ9 | OTUD7B | OTU doma | Color2 [Da | 1 | 4 |
| + | 1.999607 | -0.45849 | Q6GYQ0 | Q6GYQ0 | RALGAPA1 | Ral GTPase | Color2 [Da | 1 | 4 |
| + | 1.515427 | 1.713936 | Q6H8Q1 | Q6H8Q1 | ABLIM2 | Actin-bind | Color2 [Da | 1 | 4 |
| + | 1.647489 | 1.40313 | Q6I9Y2 | Q6I9Y2 | THOC7 | THO comp | Color2 [Da | 1 | 4 |
| | 0 | NaN | Q6IA69 | Q6IA69 | NADSYN1 | Glutamine | Color2 [Da | 1 | 4 |
| | 0 | NaN | Q6IA86 | Q6IA86 | ELP2 | Elongator | Color2 [Da | 1 | 4 |
| + | 2.991883 | 1.633544 | Q6IAA8 | Q6IAA8 | LAMTOR1 | Ragulator | Color2 [Da | 1 | 4 |
| + | 0.943837 | 0.392236 | Q6IBS0 | Q6IBS0 | TWF2 | Twinfilin-2 | Color2 [Da | 1 | 4 |
| | 0 | NaN | Q6IBW4 | Q6IBW4 | NCAPH2 | Condensin | Color2 [Da | 1 | 4 |
| | 0.045789 | -0.01775 | Q6IC98 | Q6IC98 | GRAMD4 | GRAM dor | Color2 [Da | 1 | 4 |
| + | 2.867846 | 1.260454 | Q6ICG6 | Q6ICG6 | KIAA0930 | Uncharact | Color2 [Da | 1 | 4 |
| | 0 | NaN | Q6IE81 | Q6IE81 | JADE1 | Protein Jac | Color2 [Da | 1 | 4 |
| + | 1.227701 | 1.072426 | Q6IEE7 | Q6IEE7 | TMEM132 | Transmem | Color2 [Da | 1 | 4 |
| + | 1.469284 | -0.97376 | Q6IN85 | Q6IN85 | PPP4R3A | Serine/thr | Color2 [Da | 1 | 4 |
| | 0 | NaN | Q6IPR1 | Q6IPR1 | ETFRF1 | Electron tr | Color2 [Da | 1 | 4 |
| + | 2.355689 | 0.947703 | Q6IQ22 | Q6IQ22 | RAB12 | Ras-relate | Color2 [Da | 1 | 4 |
| + | 1.323992 | -1.31437 | Q6IQ23 | Q6IQ23 | PLEKHA7 | Pleckstrin | Color2 [Da | 1 | 4 |
| | 0.359316 | -0.1349 | Q6IQ49 | Q6IQ49 | SDE2 | Replicatior | Color2 [Da | 1 | 4 |
| | 0.577332 | -0.65384 | Q6IS24 | Q6IS24 | GALNT17 | Polypeptid | Color2 [Da | 1 | 4 |
| | 0 | NaN | Q6JQN1 | Q6JQN1 | ACAD10 | Acyl-CoA d | Color2 [Da | 1 | 4 |
| + | 2.610238 | 1.021096 | Q6KB66 | Q6KB66 | KRT80 | Keratin, ty | Color2 [Da | 1 | 4 |
| + | 1.139862 | 0.372788 | Q6KC79 | Q6KC79 | NIPBL | Nipped-B-l | Color2 [Da | 1 | 4 |
| + | 1.124876 | 0.442395 | Q6KCM7 | Q6KCM7 | SLC25A25 | Calcium-bi | Color2 [Da | 1 | 4 |
| | 0.542208 | -0.38514 | Q6L8Q7 | Q6L8Q7 | PDE12 | 2',5'-phosp | Color2 [Da | 1 | 4 |
| | 0.133771 | -0.08342 | Q6MZP7 | Q6MZP7 | LIN54 | Protein lin | Color2 [Da | 1 | 4 |
| | 0 | -0.06419 | Q6N022 | Q6N022 | TENM4 | Teneurin-4 | Color2 [Da | 1 | 4 |
| | 0 | NaN | Q6N069 | Q6N069 | NAA16 | N-alpha-ac | Color2 [Da | 1 | 4 |
| + | 0.908735 | 0.586077 | Q6NT76 | Q6NT76 | HMBOX1 | Homeobox | Color2 [Da | 1 | 4 |
| + | 3.061127 | -2.67196 | Q6NTF9 | Q6NTF9 | RHBDD2 | Rhomboid | Color2 [Da | 1 | 4 |
| + | 2.219423 | -0.21162 | Q6NUK1 | Q6NUK1 | SLC25A24 | Calcium-bi | Color2 [Da | 1 | 4 |
| + | 2.191881 | -1.86681 | Q6NUM9 | Q6NUM9 | RETSAT | All-trans-re | Color2 [Da | 1 | 4 |
| | 0 | -2.56131 | Q6NUQ4 | Q6NUQ4 | TMEM214 | Transmem | Color2 [Da | 1 | 4 |
| | 0 | NaN | Q6NUS6 | Q6NUS6 | TCTN3 | Tectonic-3 | Color2 [Da | 1 | 4 |
| | 1.332069 | -0.14213 | Q6NVY1 | Q6NVY1 | HIBCH | 3-hydroxyi | Color2 [Da | 1 | 4 |
| + | 2.656296 | 2.357572 | Q6NW29 | Q6NW29 | RWDD4 | RWD doma | Color2 [Da | 1 | 4 |
| | 0.31127 | 0.415337 | Q6NXE6 | Q6NXE6 | ARMC6 | Armadillo | Color2 [Da | 1 | 4 |
| | 0 | NaN | Q6NXR4 | Q6NXR4 | TTI2 | TELO2-inte | Color2 [Da | 1 | 4 |
| | 0 | -1.03265 | Q6NXT6 | Q6NXT6 | TAPT1 | Transmem | Color2 [Da | 1 | 4 |
| + | 2.195981 | -0.62968 | Q6NYC1 | Q6NYC1 | JMJD6 | Bifunction | Color2 [Da | 1 | 4 |
| + | 1.897318 | 0.652275 | Q6NYC8 | Q6NYC8 | PPP1R18 | Phostensir | Color2 [Da | 1 | 4 |
| + | 1.171178 | 0.966435 | Q6NZ67;Q | Q6NZ67;Q | MZT2B;MZ | Mitotic-sp | Color2 [Da | 1 | 4 |
| | 0.374226 | -0.26427 | Q6NZI2 | Q6NZI2 | CAVIN1 | Caveolae-a | Color2 [Da | 1 | 4 |
| + | 0.805999 | -0.67238 | Q6NZY4 | Q6NZY4 | ZCCHC8 | Zinc finger | Color2 [Da | 1 | 4 |
| | 0 | -1.03592 | Q6P158 | Q6P158 | DHX57 | Putative A | Color2 [Da | 1 | 4 |
| + | 1.004792 | 0.724531 | Q6P161 | Q6P161 | MRPL54 | 39S riboso | Color2 [Da | 1 | 4 |
| | 0 | NaN | Q6P1A2 | Q6P1A2 | LPCAT3 | Lysophosp | Color2 [Da | 1 | 4 |

| | | | | | | | | | |
|---|---|---|---|---|---|---|---|---|---|
| + | 2.723781 | 0.547968 | Q6P1J9 | Q6P1J9 | CDC73 | Parafibron | Color2 [Da | 1 | 4 |
| + | 3.035882 | 0.702438 | Q6P1K2 | Q6P1K2 | PMF1 | Polyamine | Color2 [Da | 1 | 4 |
| + | 1.313895 | 1.60107 | Q6P1L5 | Q6P1L5 | FAM117B | Protein FA | Color2 [Da | 1 | 4 |
| + | 1.461183 | 1.05037 | Q6P1L8 | Q6P1L8 | MRPL14 | 39S riboso | Color2 [Da | 1 | 4 |
| + | 1.832454 | -0.55301 | Q6P1M0 | Q6P1M0 | SLC27A4 | Long-chair | Color2 [Da | 1 | 4 |
| + | 1.304194 | 0.492621 | Q6P1N0 | Q6P1N0 | CC2D1A | Coiled-coil | Color2 [Da | 1 | 4 |
| + | 1.683116 | 0.66027 | Q6P1N9 | Q6P1N9 | TATDN1 | Deoxyribo | Color2 [Da | 1 | 4 |
| | 0 | NaN | Q6P1Q9 | Q6P1Q9 | METTL2B | tRNA N(3)- | Color2 [Da | 1 | 4 |
| + | 1.030906 | -0.68879 | Q6P1R3 | Q6P1R3 | MSANTD2 | Myb/SANT | Color2 [Da | 1 | 4 |
| + | 0.812561 | 0.522274 | Q6P2C8 | Q6P2C8 | MED27 | Mediator c | Color2 [Da | 1 | 4 |
| + | 2.961655 | -0.63906 | Q6P2E9 | Q6P2E9 | EDC4 | Enhancer c | Color2 [Da | 1 | 4 |
| + | 0.804682 | 0.453567 | Q6P2I3 | Q6P2I3 | FAHD2B | Fumarylac | Color2 [Da | 1 | 4 |
| + | 2.364376 | -1.11323 | Q6P2Q9 | Q6P2Q9 | PRPF8 | Pre-mRNA | Color2 [Da | 1 | 4 |
| | 0.834685 | 0.317952 | Q6P3S6 | Q6P3S6 | FBXO42 | F-box only | Color2 [Da | 1 | 4 |
| + | 2.68946 | -1.13179 | Q6P3W7 | Q6P3W7 | SCYL2 | SCY1-like p | Color2 [Da | 1 | 4 |
| | 0 | -0.22075 | Q6P3X3 | Q6P3X3 | TTC27 | Tetratricop | Color2 [Da | 1 | 4 |
| + | 0.917748 | -1.02696 | Q6P4A7 | Q6P4A7 | SFXN4 | Sideroflexi | Color2 [Da | 1 | 4 |
| | 0.050003 | -0.0367 | Q6P4E1 | Q6P4E1 | GOLM2 | Protein GC | Color2 [Da | 1 | 4 |
| | 0 | NaN | Q6P4F2 | Q6P4F2 | FDX2 | Ferredoxir | Color2 [Da | 1 | 4 |
| + | 1.432574 | 0.683935 | Q6P4Q7 | Q6P4Q7 | CNNM4 | Metal tran | Color2 [Da | 1 | 4 |
| | 0.630316 | -0.18768 | Q6P4R8 | Q6P4R8 | NFRKB | Nuclear fa | Color2 [Da | 1 | 4 |
| + | 2.08711 | 2.304174 | Q6P587 | Q6P587 | FAHD1 | Acylpyruva | Color2 [Da | 1 | 4 |
| | 0 | NaN | Q6P6B7 | Q6P6B7 | ANKRD16 | Ankyrin re | Color2 [Da | 1 | 4 |
| | 0.271458 | -0.17431 | Q6P6C2 | Q6P6C2 | ALKBH5 | RNA deme | Color2 [Da | 1 | 4 |
| + | 2.060958 | 0.560335 | Q6P995 | Q6P995 | FAM171B | Protein FA | Color2 [Da | 1 | 4 |
| | 0.819798 | -0.28846 | Q6P996 | Q6P996 | PDXDC1 | Pyridoxal-c | Color2 [Da | 1 | 4 |
| | 0.217118 | 0.662094 | Q6P9B6 | Q6P9B6 | MEAK7 | MTOR-ass | Color2 [Da | 1 | 4 |
| | 0 | NaN | Q6P9B9 | Q6P9B9 | INTS5 | Integrator | Color2 [Da | 1 | 4 |
| + | 1.032585 | -0.58755 | Q6PCB0 | Q6PCB0 | VWA1 | von Willeb | Color2 [Da | 1 | 4 |
| + | 1.772643 | -0.33605 | Q6PCB7 | Q6PCB7 | SLC27A1 | Long-chair | Color2 [Da | 1 | 4 |
| + | 1.841509 | -0.60146 | Q6PCE3 | Q6PCE3 | PGM2L1 | Glucose 1, | Color2 [Da | 1 | 4 |
| | 0 | 1.941181 | Q6PCT2 | Q6PCT2 | FBXL19 | F-box/LRR- | Color2 [Da | 1 | 4 |
| + | 1.29525 | -1.34804 | Q6PD62 | Q6PD62 | CTR9 | RNA polyn | Color2 [Da | 1 | 4 |
| + | 0.911018 | 0.605517 | Q6PD74 | Q6PD74 | AAGAB | Alpha- and | Color2 [Da | 1 | 4 |
| | 0.534699 | 0.956018 | Q6PFW1 | Q6PFW1 | PPIP5K1 | Inositol he | Color2 [Da | 1 | 4 |
| + | 2.639892 | -0.76454 | Q6PGN9 | Q6PGN9 | PSRC1 | Proline/se | Color2 [Da | 1 | 4 |
| + | 2.642858 | -0.97518 | Q6PGP7 | Q6PGP7 | TTC37 | Tetratricop | Color2 [Da | 1 | 4 |
| | 0 | -0.02418 | Q6PHR2 | Q6PHR2 | ULK3 | Serine/thr | Color2 [Da | 1 | 4 |
| + | 2.662293 | -1.45409 | Q6PI48 | Q6PI48 | DARS2 | Aspartate- | Color2 [Da | 1 | 4 |
| | 0.030064 | -0.03057 | Q6PI78 | Q6PI78 | TMEM65 | Transmem | Color2 [Da | 1 | 4 |
| | 0.540038 | 0.384904 | Q6PI98 | Q6PI98 | INO80C | INO80 con | Color2 [Da | 1 | 4 |
| + | 2.06494 | 1.32542 | Q6PID6 | Q6PID6 | TTC33 | Tetratricop | Color2 [Da | 1 | 4 |
| | 0.028359 | -0.02728 | Q6PII3 | Q6PII3 | CCDC174 | Coiled-coil | Color2 [Da | 1 | 4 |
| | 0 | 0.211791 | Q6PIJ6 | Q6PIJ6 | FBXO38 | F-box only | Color2 [Da | 1 | 4 |
| | 0.08424 | -0.07497 | Q6PIU2 | Q6PIU2 | NCEH1 | Neutral ch | Color2 [Da | 1 | 4 |
| | 0.589994 | -0.18989 | Q6PJG6 | Q6PJG6 | BRAT1 | BRCA1-ass | Color2 [Da | 1 | 4 |
| + | 1.321689 | -0.4407 | Q6PJT7 | Q6PJT7 | ZC3H14 | Zinc finger | Color2 [Da | 1 | 4 |

| | | | | | | | | | |
|---|---|---|---|---|---|---|---|---|---|
| | 0 | NaN | Q6PJW8 | Q6PJW8 | CNST | Consortin | Color2 [Da | 1 | 4 |
| + | 2.384886 | 0.594113 | Q6PKG0 | Q6PKG0 | LARP1 | La-related | Color2 [Da | 1 | 4 |
| | 0.617707 | -0.41471 | Q6PL18 | Q6PL18 | ATAD2 | ATPase far | Color2 [Da | 1 | 4 |
| + | 1.119076 | 0.672317 | Q6PL24 | Q6PL24 | TMED8 | Protein TM | Color2 [Da | 1 | 4 |
| + | 1.051514 | -0.42978 | Q6PML9 | Q6PML9 | SLC30A9 | Zinc transp | Color2 [Da | 1 | 4 |
| + | 2.617508 | 3.401728 | Q6PUV4 | Q6PUV4 | CPLX2 | Complexin | Color2 [Da | 1 | 4 |
| + | 1.209 | 1.025744 | Q6QNY0 | Q6QNY0 | BLOC1S3 | Biogenesis | Color2 [Da | 1 | 4 |
| + | 2.006348 | 1.484445 | Q6QNY1 | Q6QNY1 | BLOC1S2 | Biogenesis | Color2 [Da | 1 | 4 |
| | 0 | NaN | Q6R327 | Q6R327 | RICTOR | Rapamycir | Color2 [Da | 1 | 4 |
| + | 0.663553 | -0.97053 | Q6RFH5 | Q6RFH5 | WDR74 | WD repeat | Color2 [Da | 1 | 4 |
| | 0.273676 | -1.09923 | Q6SA08 | Q6SA08 | TSSK4 | Testis-spec | Color2 [Da | 1 | 4 |
| + | 1.019582 | 0.409834 | Q6SPF0 | Q6SPF0 | SAMD1 | Sterile alpl | Color2 [Da | 1 | 4 |
| + | 1.594018 | -0.43711 | Q6STE5 | Q6STE5 | SMARCD3 | SWI/SNF-r | Color2 [Da | 1 | 4 |
| + | 1.789864 | -0.61755 | Q6SZW1 | Q6SZW1 | SARM1 | NAD(+) hy | Color2 [Da | 1 | 4 |
| + | 3.169455 | 1.250622 | Q6T4P5 | Q6T4P5 | PLPPR3 | Phospholip | Color2 [Da | 1 | 4 |
| | 0.591708 | 0.363873 | Q6U841 | Q6U841 | SLC4A10 | Sodium-dr | Color2 [Da | 1 | 4 |
| | 1.146235 | -0.19645 | Q6UB35 | Q6UB35 | MTHFD1L | Monofunc | Color2 [Da | 1 | 4 |
| + | 1.802289 | 1.172675 | Q6ULP2 | Q6ULP2 | AFTPH | Aftiphilin | Color2 [Da | 1 | 4 |
| + | 2.574137 | -0.60273 | Q6UN15 | Q6UN15 | FIP1L1 | Pre-mRNA | Color2 [Da | 1 | 4 |
| + | 1.272904 | -0.40766 | Q6UUV7 | Q6UUV7 | CRTC3 | CREB-regu | Color2 [Da | 1 | 4 |
| + | 2.007322 | 0.998601 | Q6UUV9 | Q6UUV9 | CRTC1 | CREB-regu | Color2 [Da | 1 | 4 |
| + | 1.276517 | 1.229634 | Q6UVY6 | Q6UVY6 | MOXD1 | DBH-like m | Color2 [Da | 1 | 4 |
| | 0 | -1.24595 | Q6UW63 | Q6UW63 | POGLUT2 | Protein O-; | Color2 [Da | 1 | 4 |
| + | 4.030209 | -2.25378 | Q6UW68 | Q6UW68 | TMEM205 | Transmem | Color2 [Da | 1 | 4 |
| + | 2.027105 | 2.7557 | Q6UW78 | Q6UW78 | UQCC3 | Ubiquinol- | Color2 [Da | 1 | 4 |
| + | 1.632738 | 0.406594 | Q6UWE0 | Q6UWE0 | LRSAM1 | E3 ubiquiti | Color2 [Da | 1 | 4 |
| | 0 | -1.05635 | Q6UWP7 | Q6UWP7 | LCLAT1 | Lysocardio | Color2 [Da | 1 | 4 |
| | 0.359256 | 0.168895 | Q6UWP8 | Q6UWP8 | SBSN | Suprabasir | Color2 [Da | 1 | 4 |
| | 0 | NaN | Q6UWU4 | Q6UWU4 | C6orf89 | Bombesin | Color2 [Da | 1 | 4 |
| | 0 | NaN | Q6UX01;Q | Q6UX01;Q | LMBR1L;LI | Protein LM | Color2 [Da | 1 | 4 |
| + | 1.281478 | 0.393068 | Q6UX04 | Q6UX04 | CWC27 | Spliceoson | Color2 [Da | 1 | 4 |
| + | 2.091803 | -0.63763 | Q6UX71 | Q6UX71 | PLXDC2 | Plexin dom | Color2 [Da | 1 | 4 |
| | 0 | NaN | Q6UXB8 | Q6UXB8 | PI16 | Peptidase | Color2 [Da | 1 | 4 |
| + | 3.860756 | -1.11582 | Q6UXD5 | Q6UXD5 | SEZ6L2 | Seizure 6-l | Color2 [Da | 1 | 4 |
| + | 1.266295 | 0.720967 | Q6UXH1 | Q6UXH1 | CRELD2 | Protein dis | Color2 [Da | 1 | 4 |
| + | 1.841355 | 0.864763 | Q6UXK2 | Q6UXK2 | ISLR2 | Immunogl | Color2 [Da | 1 | 4 |
| | 0 | 1.493906 | Q6UXK5 | Q6UXK5 | LRRN1 | Leucine-ric | Color2 [Da | 1 | 4 |
| + | 3.299423 | -0.53896 | Q6UXN9 | Q6UXN9 | WDR82 | WD repeat | Color2 [Da | 1 | 4 |
| + | 2.448232 | 0.929067 | Q6UXV4 | Q6UXV4 | APOOL | MICOS cor | Color2 [Da | 1 | 4 |
| + | 1.83206 | 0.897702 | Q6V0I7 | Q6V0I7 | FAT4 | Protocadh | Color2 [Da | 1 | 4 |
| + | 3.219325 | 1.102623 | Q6VY07 | Q6VY07 | PACS1 | Phosphofu | Color2 [Da | 1 | 4 |
| + | 1.977038 | -0.50989 | Q6WCQ1 | Q6WCQ1 | MPRIP | Myosin ph | Color2 [Da | 1 | 4 |
| | 0.697994 | -0.34903 | Q6WKZ4 | Q6WKZ4 | RAB11FIP1 | Rab11 fam | Color2 [Da | 1 | 4 |
| | 0 | NaN | Q6X4U4 | Q6X4U4 | SOSTDC1 | Sclerostin | Color2 [Da | 1 | 4 |
| + | 1.068195 | -1.08601 | Q6XQN6 | Q6XQN6 | NAPRT | Nicotinate | Color2 [Da | 1 | 4 |
| + | 4.252309 | -2.3568 | Q6Y1H2 | Q6Y1H2 | HACD2 | Very-long- | Color2 [Da | 1 | 4 |
| | 0.568782 | -0.29386 | Q6Y288 | Q6Y288 | B3GLCT | Beta-1,3-g | Color2 [Da | 1 | 4 |

| | | | | | | | | | |
|---|---|---|---|---|---|---|---|---|---|
| + | 1.082887 | 0.359014 | Q6Y7W6 | Q6Y7W6 | GIGYF2 | GRB10-int | Color2 [Da | 1 | 4 |
| | 0 | NaN | Q6YHK3 | Q6YHK3 | CD109 | CD109 ant | Color2 [Da | 1 | 4 |
| | 0.113119 | 0.04189 | Q6YN16 | Q6YN16 | HSDL2 | Hydroxyst | Color2 [Da | 1 | 4 |
| | 0.219686 | -0.1414 | Q6YP21 | Q6YP21 | KYAT3 | Kynurenin | Color2 [Da | 1 | 4 |
| | 0 | -1.53812 | Q6ZMG9 | Q6ZMG9 | CERS6 | Ceramide | Color2 [Da | 1 | 4 |
| + | 1.567781 | -0.30972 | Q6ZMI0 | Q6ZMI0 | PPP1R21 | Protein ph | Color2 [Da | 1 | 4 |
| | 0 | NaN | Q6ZMK1 | Q6ZMK1 | CYHR1 | Cysteine a | Color2 [Da | 1 | 4 |
| + | 2.090444 | 2.164056 | Q6ZMP0 | Q6ZMP0 | THSD4 | Thrombos | Color2 [Da | 1 | 4 |
| | 0.297557 | 0.201732 | Q6ZN17 | Q6ZN17 | LIN28B | Protein lin | Color2 [Da | 1 | 4 |
| + | 2.719897 | 3.333498 | Q6ZN30 | Q6ZN30 | BNC2 | Zinc finger | Color2 [Da | 1 | 4 |
| + | 1.069484 | 0.929835 | Q6ZN55 | Q6ZN55 | ZNF574 | Zinc finger | Color2 [Da | 1 | 4 |
| + | 2.425253 | -1.83963 | Q6ZPD9 | Q6ZPD9 | DPY19L3 | Probable C | Color2 [Da | 1 | 4 |
| | 0 | -1.06683 | Q6ZRP7 | Q6ZRP7 | QSOX2 | Sulfhydryl | Color2 [Da | 1 | 4 |
| | 0 | 1.159394 | Q6ZRS2 | Q6ZRS2 | SRCAP | Helicase SI | Color2 [Da | 1 | 4 |
| | 0.064517 | 0.030928 | Q6ZS17 | Q6ZS17 | RIPOR1 | Rho family | Color2 [Da | 1 | 4 |
| + | 1.332647 | 1.236891 | Q6ZSJ8 | Q6ZSJ8 | C1orf122 | Uncharact | Color2 [Da | 1 | 4 |
| + | 0.949176 | -0.33309 | Q6ZSR9 | Q6ZSR9 | | Uncharact | Color2 [Da | 1 | 4 |
| + | 0.736643 | 0.718082 | Q6ZSY5 | Q6ZSY5 | PPP1R3F | Protein ph | Color2 [Da | 1 | 4 |
| | 0.804689 | 0.32087 | Q6ZSZ5 | Q6ZSZ5 | ARHGEF18 | Rho guanir | Color2 [Da | 1 | 4 |
| | 0.214137 | -0.16288 | Q6ZT21 | Q6ZT21 | TMPPE | Transmem | Color2 [Da | 1 | 4 |
| | 0.023588 | 0.017329 | Q6ZTA4 | Q6ZTA4 | TRIM67 | Tripartite | Color2 [Da | 1 | 4 |
| + | 1.160165 | -0.5508 | Q6ZTN6 | Q6ZTN6 | ANKRD13D | Ankyrin re | Color2 [Da | 1 | 4 |
| | 0 | 0.210238 | Q6ZTW0 | Q6ZTW0 | TPGS1 | Tubulin po | Color2 [Da | 1 | 4 |
| + | 4.278519 | 1.336464 | Q6ZU35 | Q6ZU35 | CRACD | Capping pr | Color2 [Da | 1 | 4 |
| + | 1.818304 | 0.821218 | Q6ZUT1 | Q6ZUT1 | NKAPD1 | Uncharact | Color2 [Da | 1 | 4 |
| + | 0.707277 | 0.82043 | Q6ZUT9 | Q6ZUT9 | DENND5B | DENN dom | Color2 [Da | 1 | 4 |
| + | 1.034926 | 0.439044 | Q6ZVC0 | Q6ZVC0 | NYAP1 | Neuronal t | Color2 [Da | 1 | 4 |
| + | 2.404954 | 1.631004 | Q6ZVF9 | Q6ZVF9 | GPRIN3 | G protein- | Color2 [Da | 1 | 4 |
| + | 1.819577 | 1.551279 | Q6ZVL6 | Q6ZVL6 | KIAA1549L | UPF0606 p | Color2 [Da | 1 | 4 |
| + | 2.857039 | 0.966581 | Q6ZVM7 | Q6ZVM7 | TOM1L2 | TOM1-like | Color2 [Da | 1 | 4 |
| | 0 | 2.252661 | Q6ZVX7 | Q6ZVX7 | NCCRP1 | F-box only | Color2 [Da | 1 | 4 |
| | 0 | NaN | Q6ZW49 | Q6ZW49 | PAXIP1 | PAX-intera | Color2 [Da | 1 | 4 |
| + | 2.508638 | -2.68395 | Q6ZWT7 | Q6ZWT7 | MBOAT2 | Lysophosp | Color2 [Da | 1 | 4 |
| + | 3.07939 | -1.25732 | Q6ZXV5 | Q6ZXV5 | TMTC3 | Protein O- | Color2 [Da | 1 | 4 |
| + | 2.904363 | -0.64312 | Q709C8 | Q709C8 | VPS13C | Intermeml | Color2 [Da | 1 | 4 |
| | 0 | NaN | Q709F0 | Q709F0 | ACAD11 | Acyl-CoA d | Color2 [Da | 1 | 4 |
| | 0.921732 | -0.20568 | Q70E73 | Q70E73 | RAPH1 | Ras-associ | Color2 [Da | 1 | 4 |
| | 0.370675 | 0.290314 | Q70IA6 | Q70IA6 | MOB2 | MOB kinas | Color2 [Da | 1 | 4 |
| | 0.02096 | 0.021257 | Q70UQ0 | Q70UQ0 | IKBIP | Inhibitor o | Color2 [Da | 1 | 4 |
| | 0 | NaN | Q70Z35 | Q70Z35 | PREX2 | Phosphatic | Color2 [Da | 1 | 4 |
| + | 1.018232 | 0.597718 | Q712K3 | Q712K3 | UBE2R2 | Ubiquitin-c | Color2 [Da | 1 | 4 |
| + | 1.629763 | 1.904451 | Q71DI3 | Q71DI3 | H3C13 | Histone H3 | Color2 [Da | 1 | 4 |
| | 0.111275 | -0.11883 | Q71H61 | Q71H61 | ILDR2 | Immunogl | Color2 [Da | 1 | 4 |
| | 0.455389 | 0.245612 | Q71RC2 | Q71RC2 | LARP4 | La-related | Color2 [Da | 1 | 4 |
| | 0 | NaN | Q71SY5 | Q71SY5 | MED25 | Mediator c | Color2 [Da | 1 | 4 |
| + | 0.959168 | -0.65519 | Q71UM5 | Q71UM5 | RPS27L | 40S riboso | Color2 [Da | 1 | 4 |
| | 0.02011 | 0.052649 | Q75N03 | Q75N03 | CBLL1 | E3 ubiquiti | Color2 [Da | 1 | 4 |

| | | | | | | | | | |
|---|---|---|---|---|---|---|---|---|---|
| | 0 | NaN | Q75N90 | Q75N90 | FBN3 | Fibrillin-3 | Color2 [Da | 1 | 4 |
| + | 4.137175 | -0.84146 | Q75T13 | Q75T13 | PGAP1 | GPI inosito | Color2 [Da | 1 | 4 |
| + | 3.20917 | 2.645768 | Q765P7 | Q765P7 | MTSS2 | Protein M | Color2 [Da | 1 | 4 |
| + | 3.500397 | 1.147511 | Q76I76 | Q76I76 | SSH2 | Protein ph | Color2 [Da | 1 | 4 |
| + | 1.734549 | 0.673117 | Q76L83 | Q76L83 | ASXL2 | Putative P | Color2 [Da | 1 | 4 |
| | 0 | NaN | Q76M96 | Q76M96 | CCDC80 | Coiled-coil | Color2 [Da | 1 | 4 |
| | 0.604973 | -0.34453 | Q76N89 | Q76N89 | HECW1 | E3 ubiquiti | Color2 [Da | 1 | 4 |
| + | 2.405818 | -1.35104 | Q7KZ85 | Q7KZ85 | SUPT6H | Transcripti | Color2 [Da | 1 | 4 |
| + | 2.857869 | -0.86558 | Q7KZF4 | Q7KZF4 | SND1 | Staphylocc | Color2 [Da | 1 | 4 |
| | 0.86631 | -0.3039 | Q7KZI7 | Q7KZI7 | MARK2 | Serine/thr | Color2 [Da | 1 | 4 |
| + | 2.0093 | -1.28772 | Q7KZN9 | Q7KZN9 | COX15 | Cytochron | Color2 [Da | 1 | 4 |
| + | 1.87992 | -0.61099 | Q7L014 | Q7L014 | DDX46 | Probable A | Color2 [Da | 1 | 4 |
| + | 3.948429 | 0.274926 | Q7L099 | Q7L099 | RUFY3 | Protein RU | Color2 [Da | 1 | 4 |
| | 0.598328 | 0.292542 | Q7L0J3 | Q7L0J3 | SV2A | Synaptic v | Color2 [Da | 1 | 4 |
| + | 2.036986 | -0.31913 | Q7L0Y3 | Q7L0Y3 | TRMT10C | tRNA meth | Color2 [Da | 1 | 4 |
| + | 2.996119 | -1.93426 | Q7L190 | Q7L190 | DPPA4 | Developm | Color2 [Da | 1 | 4 |
| + | 3.051478 | -1.5658 | Q7L1Q6 | Q7L1Q6 | BZW1 | eIF5-mimi | Color2 [Da | 1 | 4 |
| | 0 | 1.693606 | Q7L1V2 | Q7L1V2 | MON1B | Vacuolar f | Color2 [Da | 1 | 4 |
| + | 0.799298 | -1.26775 | Q7L1W4 | Q7L1W4 | LRRC8D | Volume-re | Color2 [Da | 1 | 4 |
| + | 3.729607 | 3.087749 | Q7L266 | Q7L266 | ASRGL1 | Isoasparty | Color2 [Da | 1 | 4 |
| + | 1.186988 | -0.87971 | Q7L2E3 | Q7L2E3 | DHX30 | ATP-deper | Color2 [Da | 1 | 4 |
| + | 3.278157 | -1.04157 | Q7L2H7 | Q7L2H7 | EIF3M | Eukaryotic | Color2 [Da | 1 | 4 |
| + | 0.943357 | -0.9719 | Q7L2J0 | Q7L2J0 | MEPCE | 7SK snRNA | Color2 [Da | 1 | 4 |
| | 0.756444 | -0.38402 | Q7L311 | Q7L311 | ARMCX2 | Armadillo | Color2 [Da | 1 | 4 |
| | 0 | NaN | Q7L3B6 | Q7L3B6 | CDC37L1 | Hsp90 co-c | Color2 [Da | 1 | 4 |
| | 0 | -0.45221 | Q7L3T8 | Q7L3T8 | PARS2 | Probable p | Color2 [Da | 1 | 4 |
| | 0.574056 | 0.220316 | Q7L4I2 | Q7L4I2 | RSRC2 | Arginine/s | Color2 [Da | 1 | 4 |
| | 0.430014 | -0.4189 | Q7L4P6 | Q7L4P6 | BEND5 | BEN doma | Color2 [Da | 1 | 4 |
| + | 1.63326 | -1.0024 | Q7L523 | Q7L523 | RRAGA | Ras-relate | Color2 [Da | 1 | 4 |
| + | 4.367266 | -1.54212 | Q7L576 | Q7L576 | CYFIP1 | Cytoplasm | Color2 [Da | 1 | 4 |
| | 0.080452 | -0.08072 | Q7L592 | Q7L592 | NDUFAF7 | Protein arg | Color2 [Da | 1 | 4 |
| + | 3.652751 | -0.99421 | Q7L5D6 | Q7L5D6 | GET4 | Golgi to EF | Color2 [Da | 1 | 4 |
| + | 2.064617 | -1.18238 | Q7L5N1 | Q7L5N1 | COPS6 | COP9 signa | Color2 [Da | 1 | 4 |
| | 0 | NaN | Q7L5Y1 | Q7L5Y1 | ENOSF1 | Mitochonc | Color2 [Da | 1 | 4 |
| | 0 | NaN | Q7L5Y9 | Q7L5Y9 | MAEA | E3 ubiquiti | Color2 [Da | 1 | 4 |
| + | 1.66896 | -0.87108 | Q7L775 | Q7L775 | EPM2AIP1 | EPM2A-int | Color2 [Da | 1 | 4 |
| + | 1.135679 | -0.2725 | Q7L7X3 | Q7L7X3 | TAOK1 | Serine/thr | Color2 [Da | 1 | 4 |
| + | 2.614614 | 2.632765 | Q7L804 | Q7L804 | RAB11FIP2 | Rab11 fam | Color2 [Da | 1 | 4 |
| + | 0.995257 | -0.55891 | Q7L8A9 | Q7L8A9 | VASH1 | Tubulinyl-T | Color2 [Da | 1 | 4 |
| | 0.250373 | 0.196854 | Q7L8J4 | Q7L8J4 | SH3BP5L | SH3 domai | Color2 [Da | 1 | 4 |
| | 0 | NaN | Q7L8L6 | Q7L8L6 | FASTKD5 | FAST kinas | Color2 [Da | 1 | 4 |
| | 0 | NaN | Q7L985;Q9 | Q7L985;Q9 | LINGO2;LII | Leucine-ric | Color2 [Da | 1 | 4 |
| | 0.446662 | 0.287023 | Q7L9L4;Q9 | Q7L9L4;Q9 | MOB1B;M | MOB kinas | Color2 [Da | 1 | 4 |
| + | 2.631361 | -0.48384 | Q7LBC6 | Q7LBC6 | KDM3B | Lysine-spe | Color2 [Da | 1 | 4 |
| + | 1.888913 | 0.732058 | Q7LBR1 | Q7LBR1 | CHMP1B | Charged m | Color2 [Da | 1 | 4 |
| | 0 | NaN | Q7LFL8 | Q7LFL8 | CXXC5 | CXXC-type | Color2 [Da | 1 | 4 |
| + | 1.887129 | -0.98089 | Q7LG56 | Q7LG56 | RRM2B | Ribonucle | Color2 [Da | 1 | 4 |

| | | | | | | | | | |
|---|---|---|---|---|---|---|---|---|---|
| + | 1.133066 | -1.23983 | Q7LGA3 | Q7LGA3 | HS2ST1 | Heparan su | Color2 [Da | 1 | 4 |
| | 0.154964 | 0.167293 | Q7LGC8 | Q7LGC8 | CHST3 | Carbohydr | Color2 [Da | 1 | 4 |
| | 0 | NaN | Q7RTP0 | Q7RTP0 | NIPA1 | Magnesiur | Color2 [Da | 1 | 4 |
| + | 2.295635 | 0.784856 | Q7RTP6 | Q7RTP6 | MICAL3 | [F-actin]-m | Color2 [Da | 1 | 4 |
| + | 2.629231 | -0.95363 | Q7Z2K6 | Q7Z2K6 | ERMP1 | Endoplasm | Color2 [Da | 1 | 4 |
| + | 5.790671 | 2.365942 | Q7Z2K8 | Q7Z2K8 | GPRIN1 | G protein-i | Color2 [Da | 1 | 4 |
| | 0 | -1.41965 | Q7Z2T5 | Q7Z2T5 | TRMT1L | TRMT1-like | Color2 [Da | 1 | 4 |
| + | 1.777545 | 1.044384 | Q7Z2W9 | Q7Z2W9 | MRPL21 | 39S riboso | Color2 [Da | 1 | 4 |
| + | 2.494217 | -0.81841 | Q7Z2Z2 | Q7Z2Z2 | EFL1 | Elongation | Color2 [Da | 1 | 4 |
| + | 1.085418 | 0.441597 | Q7Z392 | Q7Z392 | TRAPPC11 | Trafficking | Color2 [Da | 1 | 4 |
| + | 1.796927 | 0.963903 | Q7Z3B1 | Q7Z3B1 | NEGR1 | Neuronal g | Color2 [Da | 1 | 4 |
| + | 2.788443 | 0.962632 | Q7Z3B4 | Q7Z3B4 | NUP54 | Nucleopor | Color2 [Da | 1 | 4 |
| + | 1.129426 | 0.503152 | Q7Z3C6 | Q7Z3C6 | ATG9A | Autophagy | Color2 [Da | 1 | 4 |
| + | 1.107005 | 0.408886 | Q7Z3D6 | Q7Z3D6 | DGLUCY | D-glutama | Color2 [Da | 1 | 4 |
| | 0 | -1.8791 | Q7Z3E5 | Q7Z3E5 | ARMC9 | LisH doma | Color2 [Da | 1 | 4 |
| | 0.487276 | 0.30606 | Q7Z3G6 | Q7Z3G6 | PRICKLE2 | Prickle-like | Color2 [Da | 1 | 4 |
| + | 2.215749 | -1.18977 | Q7Z3J2 | Q7Z3J2 | VPS35L | VPS35 end | Color2 [Da | 1 | 4 |
| | 0.924835 | -0.20304 | Q7Z3K3 | Q7Z3K3 | POGZ | Pogo trans | Color2 [Da | 1 | 4 |
| | 0 | 0.706748 | Q7Z3T8 | Q7Z3T8 | ZFYVE16 | Zinc finger | Color2 [Da | 1 | 4 |
| + | 0.910046 | -0.59158 | Q7Z3U7 | Q7Z3U7 | MON2 | Protein MO | Color2 [Da | 1 | 4 |
| | 0 | -1.44715 | Q7Z3V4 | Q7Z3V4 | UBE3B | Ubiquitin-p | Color2 [Da | 1 | 4 |
| + | 2.019164 | 3.34682 | Q7Z3Y8 | Q7Z3Y8 | KRT27 | Keratin, ty | Color2 [Da | 1 | 4 |
| + | 1.543892 | 0.44656 | Q7Z401 | Q7Z401 | DENND4A | C-myc pro | Color2 [Da | 1 | 4 |
| + | 3.149677 | -1.0632 | Q7Z406 | Q7Z406 | MYH14 | Myosin-14 | Color2 [Da | 1 | 4 |
| + | 0.922986 | 0.597436 | Q7Z417 | Q7Z417 | NUFIP2 | Nuclear fra | Color2 [Da | 1 | 4 |
| + | 1.747561 | 2.787826 | Q7Z422 | Q7Z422 | SZRD1 | SUZ domai | Color2 [Da | 1 | 4 |
| + | 2.557243 | 1.311519 | Q7Z434 | Q7Z434 | MAVS | Mitochonc | Color2 [Da | 1 | 4 |
| | 0 | 0.722418 | Q7Z449 | Q7Z449 | CYP2U1 | Cytochrom | Color2 [Da | 1 | 4 |
| + | 3.367359 | -0.79751 | Q7Z460 | Q7Z460 | CLASP1 | CLIP-assoc | Color2 [Da | 1 | 4 |
| | 0.020515 | -0.0191 | Q7Z478 | Q7Z478 | DHX29 | ATP-deper | Color2 [Da | 1 | 4 |
| + | 0.932017 | 0.53489 | Q7Z4G1 | Q7Z4G1 | COMMD6 | COMM do | Color2 [Da | 1 | 4 |
| + | 1.81509 | -0.97172 | Q7Z4G4 | Q7Z4G4 | TRMT11 | tRNA (guar | Color2 [Da | 1 | 4 |
| + | 1.747798 | 1.209447 | Q7Z4H3 | Q7Z4H3 | HDDC2 | 5'-deoxynu | Color2 [Da | 1 | 4 |
| + | 1.25424 | -1.11497 | Q7Z4H7 | Q7Z4H7 | HAUS6 | HAUS augr | Color2 [Da | 1 | 4 |
| | 0 | NaN | Q7Z4H8 | Q7Z4H8 | POGLUT3 | Protein O-g | Color2 [Da | 1 | 4 |
| | 0.115487 | -0.05131 | Q7Z4K8 | Q7Z4K8 | TRIM46 | Tripartite r | Color2 [Da | 1 | 4 |
| | 0 | -1.03755 | Q7Z4L5 | Q7Z4L5 | TTC21B | Tetratricop | Color2 [Da | 1 | 4 |
| | 0 | NaN | Q7Z4P5 | Q7Z4P5 | GDF7 | Growth/di | Color2 [Da | 1 | 4 |
| + | 1.821989 | -1.44602 | Q7Z4Q2 | Q7Z4Q2 | HEATR3 | HEAT repe | Color2 [Da | 1 | 4 |
| + | 1.350033 | -1.21184 | Q7Z4R8 | Q7Z4R8 | C6orf120 | UPF0669 p | Color2 [Da | 1 | 4 |
| | 1.554525 | 0.194909 | Q7Z4S6 | Q7Z4S6 | KIF21A | Kinesin-like | Color2 [Da | 1 | 4 |
| | 0.462106 | -0.09748 | Q7Z4V5 | Q7Z4V5 | HDGFL2 | Hepatoma | Color2 [Da | 1 | 4 |
| | 0.186319 | -0.09573 | Q7Z4W1 | Q7Z4W1 | DCXR | L-xylulose | Color2 [Da | 1 | 4 |
| | 0 | NaN | Q7Z569 | Q7Z569 | BRAP | BRCA1-ass | Color2 [Da | 1 | 4 |
| | 1.104535 | 0.109038 | Q7Z589 | Q7Z589 | EMSY | BRCA2-inte | Color2 [Da | 1 | 4 |
| | 0.948065 | -0.19452 | Q7Z5G4 | Q7Z5G4 | GOLGA7 | Golgin sub | Color2 [Da | 1 | 4 |
| + | 1.338115 | 1.065144 | Q7Z5J4 | Q7Z5J4 | RAI1 | Retinoic ac | Color2 [Da | 1 | 4 |

| | | | | | | | | | |
|---|---|---|---|---|---|---|---|---|---|
| | 0.126373 | -0.0423 | Q7Z5K2 | Q7Z5K2 | WAPL | Wings apa | Color2 [Da | 1 | 4 |
| + | 1.195765 | 1.829343 | Q7Z5L7 | Q7Z5L7 | PODN | Podocan | Color2 [Da | 1 | 4 |
| + | 2.155366 | 0.926661 | Q7Z5L9 | Q7Z5L9 | IRF2BP2 | Interferon | Color2 [Da | 1 | 4 |
| + | 1.030088 | -0.60956 | Q7Z5N4 | Q7Z5N4 | SDK1 | Protein sid | Color2 [Da | 1 | 4 |
| | 0.666682 | 0.375851 | Q7Z5Q1 | Q7Z5Q1 | CPEB2 | Cytoplasm | Color2 [Da | 1 | 4 |
| | 0 | NaN | Q7Z6B0 | Q7Z6B0 | CCDC91 | Coiled-coil | Color2 [Da | 1 | 4 |
| + | 2.477481 | -1.00618 | Q7Z6B7 | Q7Z6B7 | SRGAP1 | SLIT-ROBO | Color2 [Da | 1 | 4 |
| | 0.832839 | 0.102483 | Q7Z6E9 | Q7Z6E9 | RBBP6 | E3 ubiquiti | Color2 [Da | 1 | 4 |
| | 0.139811 | -0.07849 | Q7Z6G3 | Q7Z6G3 | NECAB2 | N-termina | Color2 [Da | 1 | 4 |
| + | 0.786112 | -0.88379 | Q7Z6G8 | Q7Z6G8 | ANKS1B | Ankyrin re | Color2 [Da | 1 | 4 |
| | 0.242039 | -0.11448 | Q7Z6J0 | Q7Z6J0 | SH3RF1 | E3 ubiquiti | Color2 [Da | 1 | 4 |
| + | 1.514882 | 1.035363 | Q7Z6K5 | Q7Z6K5 | ARPIN | Arpin | Color2 [Da | 1 | 4 |
| + | 2.655829 | 2.375634 | Q7Z6L0 | Q7Z6L0 | PRRT2 | Proline-ric | Color2 [Da | 1 | 4 |
| + | 0.869165 | -0.71459 | Q7Z6L1 | Q7Z6L1 | TECPR1 | Tectonin b | Color2 [Da | 1 | 4 |
| | 0 | 0.321996 | Q7Z6M1 | Q7Z6M1 | RABEPK | Rab9 effec | Color2 [Da | 1 | 4 |
| + | 2.202377 | -0.41185 | Q7Z6Z7 | Q7Z6Z7 | HUWE1 | E3 ubiquiti | Color2 [Da | 1 | 4 |
| | 0.258591 | -0.12522 | Q7Z739 | Q7Z739 | YTHDF3 | YTH doma | Color2 [Da | 1 | 4 |
| + | 1.537034 | 1.258794 | Q7Z794 | Q7Z794 | KRT77 | Keratin, ty | Color2 [Da | 1 | 4 |
| | 0 | -0.41828 | Q7Z7A4 | Q7Z7A4 | PXK | PX domain | Color2 [Da | 1 | 4 |
| | 0.135492 | 0.013337 | Q7Z7E8 | Q7Z7E8 | UBE2Q1 | Ubiquitin-c | Color2 [Da | 1 | 4 |
| + | 1.935877 | 0.867348 | Q7Z7F0 | Q7Z7F0 | KHDC4 | KH homolo | Color2 [Da | 1 | 4 |
| | 0.489653 | 0.300531 | Q7Z7F7 | Q7Z7F7 | MRPL55 | 39S riboso | Color2 [Da | 1 | 4 |
| + | 1.853462 | 0.278979 | Q7Z7H5 | Q7Z7H5 | TMED4 | Transmem | Color2 [Da | 1 | 4 |
| | 0 | 1.189033 | Q7Z7H8 | Q7Z7H8 | MRPL10 | 39S riboso | Color2 [Da | 1 | 4 |
| + | 1.96784 | -0.78583 | Q7Z7K6 | Q7Z7K6 | CENPV | Centromer | Color2 [Da | 1 | 4 |
| | 0 | NaN | Q7Z7L7 | Q7Z7L7 | ZER1 | Protein zer | Color2 [Da | 1 | 4 |
| | 0 | 0.007635 | Q7Z7M0 | Q7Z7M0 | MEGF8 | Multiple ep | Color2 [Da | 1 | 4 |
| + | 1.801861 | 1.627623 | Q86SE5 | Q86SE5 | RALYL | RNA-bindi | Color2 [Da | 1 | 4 |
| + | 2.189961 | -1.85922 | Q86SK9 | Q86SK9 | SCD5 | Stearoyl-C | Color2 [Da | 1 | 4 |
| | 0.467875 | 1.095086 | Q86SQ7 | Q86SQ7 | SDCCAG8 | Serologica | Color2 [Da | 1 | 4 |
| + | 0.683668 | 2.26796 | Q86SX6 | Q86SX6 | GLRX5 | Glutaredox | Color2 [Da | 1 | 4 |
| | 0.537229 | -0.77452 | Q86T03 | Q86T03 | PIP4P1 | Type 1 pho | Color2 [Da | 1 | 4 |
| | 0.133766 | 0.220471 | Q86TB9 | Q86TB9 | PATL1 | Protein PA | Color2 [Da | 1 | 4 |
| + | 1.210205 | 0.376373 | Q86TG7 | Q86TG7 | PEG10 | Retrotrans | Color2 [Da | 1 | 4 |
| | 0.409451 | -0.15064 | Q86TI0 | Q86TI0 | TBC1D1 | TBC1 doma | Color2 [Da | 1 | 4 |
| | 0 | NaN | Q86TI2 | Q86TI2 | DPP9 | Dipeptidyl | Color2 [Da | 1 | 4 |
| + | 1.134991 | 0.401033 | Q86TN4 | Q86TN4 | TRPT1 | tRNA 2'-ph | Color2 [Da | 1 | 4 |
| + | 4.242863 | -0.90988 | Q86TP1 | Q86TP1 | PRUNE1 | Exopolyph | Color2 [Da | 1 | 4 |
| + | 2.472123 | 1.504047 | Q86TS9 | Q86TS9 | MRPL52 | 39S riboso | Color2 [Da | 1 | 4 |
| + | 1.747988 | -0.65745 | Q86TU7 | Q86TU7 | SETD3 | Actin-histi | Color2 [Da | 1 | 4 |
| + | 2.778837 | -1.23171 | Q86TV6 | Q86TV6 | TTC7B | Tetratricop | Color2 [Da | 1 | 4 |
| | 0.353471 | -0.37792 | Q86TW2 | Q86TW2 | ADCK1 | AarF doma | Color2 [Da | 1 | 4 |
| + | 1.067299 | -0.51451 | Q86U06 | Q86U06 | RBM23 | Probable F | Color2 [Da | 1 | 4 |
| + | 2.089324 | 0.659451 | Q86U28 | Q86U28 | ISCA2 | Iron-sulfur | Color2 [Da | 1 | 4 |
| + | 1.410752 | -1.12657 | Q86U38 | Q86U38 | NOP9 | Nucleolar p | Color2 [Da | 1 | 4 |
| + | 4.038489 | 1.365954 | Q86U42 | Q86U42 | PABPN1 | Polyadeny | Color2 [Da | 1 | 4 |
| + | 1.503974 | -0.69965 | Q86U44 | Q86U44 | METTL3 | N6-adenos | Color2 [Da | 1 | 4 |

| | | | | | | | | | |
|---|---|---|---|---|---|---|---|---|---|
| | 0.730956 | -0.55024 | Q86U70 | Q86U70 | LDB1 | LIM domai | Color2 [Da | 1 | 4 |
| + | 1.442918 | -0.40894 | Q86U86 | Q86U86 | PBRM1 | Protein po | Color2 [Da | 1 | 4 |
| + | 1.874991 | -0.35518 | Q86U90 | Q86U90 | YRDC | Threonylca | Color2 [Da | 1 | 4 |
| + | 1.245771 | -0.39083 | Q86UA1 | Q86UA1 | PRPF39 | Pre-mRNA | Color2 [Da | 1 | 4 |
| | 0.753889 | 0.400701 | Q86UE4 | Q86UE4 | MTDH | Protein LYI | Color2 [Da | 1 | 4 |
| | 0.204448 | -0.48987 | Q86UE8 | Q86UE8 | TLK2 | Serine/thr | Color2 [Da | 1 | 4 |
| | 0 | NaN | Q86UL3 | Q86UL3 | GPAT4 | Glycerol-3- | Color2 [Da | 1 | 4 |
| + | 1.646924 | 1.204398 | Q86UL8 | Q86UL8 | MAGI2 | Membrane | Color2 [Da | 1 | 4 |
| + | 3.715258 | -0.59685 | Q86UP2 | Q86UP2 | KTN1 | Kinectin | Color2 [Da | 1 | 4 |
| + | 3.494237 | 1.769187 | Q86UR5 | Q86UR5 | RIMS1 | Regulating | Color2 [Da | 1 | 4 |
| | 0 | -0.36731 | Q86US8 | Q86US8 | SMG6 | Telomeras | Color2 [Da | 1 | 4 |
| + | 0.961457 | 0.671342 | Q86UU0 | Q86UU0 | BCL9L | B-cell CLL/ | Color2 [Da | 1 | 4 |
| + | 2.604797 | 0.675304 | Q86UU1 | Q86UU1 | PHLDB1 | Pleckstrin | Color2 [Da | 1 | 4 |
| + | 2.576023 | -1.44001 | Q86UV5 | Q86UV5 | USP48 | Ubiquitin c | Color2 [Da | 1 | 4 |
| + | 0.94976 | 1.250724 | Q86UX2 | Q86UX2 | ITIH5 | Inter-alpha | Color2 [Da | 1 | 4 |
| | 0.091658 | -0.12969 | Q86UX6 | Q86UX6 | STK32C | Serine/thr | Color2 [Da | 1 | 4 |
| + | 3.790007 | -1.48344 | Q86UY6 | Q86UY6 | NAA40 | N-alpha-ac | Color2 [Da | 1 | 4 |
| + | 1.749818 | 0.584493 | Q86UY8 | Q86UY8 | NT5DC3 | 5'-nucleoti | Color2 [Da | 1 | 4 |
| | 0 | NaN | Q86V15 | Q86V15 | CASZ1 | Zinc finger | Color2 [Da | 1 | 4 |
| | 0.351507 | 0.161975 | Q86V48 | Q86V48 | LUZP1 | Leucine zip | Color2 [Da | 1 | 4 |
| + | 4.52461 | -1.59036 | Q86V59 | Q86V59 | PNMA8A | Paraneopla | Color2 [Da | 1 | 4 |
| + | 3.353447 | 1.074346 | Q86V81 | Q86V81 | ALYREF | THO comp | Color2 [Da | 1 | 4 |
| | 0 | NaN | Q86V85 | Q86V85 | GPR180 | Integral m | Color2 [Da | 1 | 4 |
| + | 1.37253 | 3.184138 | Q86V88 | Q86V88 | MDP1 | Magnesiur | Color2 [Da | 1 | 4 |
| + | 1.517647 | -0.63168 | Q86V97 | Q86V97 | KBTBD6 | Kelch repe | Color2 [Da | 1 | 4 |
| + | 2.315892 | 2.109854 | Q86VD7 | Q86VD7 | SLC25A42 | Mitochonc | Color2 [Da | 1 | 4 |
| | 0.078373 | -0.02153 | Q86VM9 | Q86VM9 | ZC3H18 | Zinc finger | Color2 [Da | 1 | 4 |
| | 0.437255 | -0.20247 | Q86VN1 | Q86VN1 | VPS36 | Vacuolar p | Color2 [Da | 1 | 4 |
| | 0 | -1.28696 | Q86VP1 | Q86VP1 | TAX1BP1 | Tax1-bindi | Color2 [Da | 1 | 4 |
| | 0.134674 | -0.11399 | Q86VP3 | Q86VP3 | PACS2 | Phosphofu | Color2 [Da | 1 | 4 |
| + | 2.912473 | -1.38124 | Q86VP6 | Q86VP6 | CAND1 | Cullin-asso | Color2 [Da | 1 | 4 |
| | 0 | 0.058246 | Q86VQ0 | Q86VQ0 | LCA5 | Lebercilin | Color2 [Da | 1 | 4 |
| + | 2.388743 | 0.914985 | Q86VQ1 | Q86VQ1 | GLCCI1 | Glucocorti | Color2 [Da | 1 | 4 |
| + | 2.113636 | -1.20501 | Q86VR2 | Q86VR2 | RETREG3 | Reticuloph | Color2 [Da | 1 | 4 |
| + | 1.223676 | -0.65006 | Q86VR8 | Q86VR8 | FJX1 | Four-jointe | Color2 [Da | 1 | 4 |
| + | 1.329805 | -0.24598 | Q86VS8 | Q86VS8 | HOOK3 | Protein Hc | Color2 [Da | 1 | 4 |
| + | 0.876014 | -1.08041 | Q86VW0 | Q86VW0 | SESTD1 | SEC14 don | Color2 [Da | 1 | 4 |
| | 0 | -0.11143 | Q86VX2 | Q86VX2 | COMMD7 | COMM do | Color2 [Da | 1 | 4 |
| + | 2.127356 | -0.67106 | Q86W42 | Q86W42 | THOC6 | THO comp | Color2 [Da | 1 | 4 |
| + | 1.321342 | -0.78863 | Q86W50 | Q86W50 | METTL16 | RNA N6-ac | Color2 [Da | 1 | 4 |
| | 0.672362 | -0.69169 | Q86W56 | Q86W56 | PARG | Poly(ADP-r | Color2 [Da | 1 | 4 |
| + | 1.77568 | 1.166691 | Q86WA6 | Q86WA6 | BPHL | Valacyclov | Color2 [Da | 1 | 4 |
| | 0.822883 | 0.150774 | Q86WA8 | Q86WA8 | LONP2 | Lon protea | Color2 [Da | 1 | 4 |
| + | 2.057755 | -1.08238 | Q86WB0 | Q86WB0 | ZC3HC1 | Zinc finger | Color2 [Da | 1 | 4 |
| | 0.240308 | 0.292245 | Q86WC4 | Q86WC4 | OSTM1 | Osteopetr | Color2 [Da | 1 | 4 |
| | 0.172663 | -0.17617 | Q86WG3 | Q86WG3 | ATCAY | Caytaxin | Color2 [Da | 1 | 4 |
| | 0.511142 | -0.12975 | Q86WG5 | Q86WG5 | SBF2 | Myotubula | Color2 [Da | 1 | 4 |

-Bruker Confidential-

| | | | | | | | | | |
|---|---|---|---|---|---|---|---|---|---|
| | 0.797348 | 0.356711 | Q86WJ1 | Q86WJ1 | CHD1L | Chromodo | Color2 [Da | 1 | 4 |
| | 0 | -1.29656 | Q86WN1 | Q86WN1 | FCHSD1 | F-BAR and | Color2 [Da | 1 | 4 |
| | 0.035202 | -0.04217 | Q86WQ0 | Q86WQ0 | NR2C2AP | Nuclear re | Color2 [Da | 1 | 4 |
| | 0.347777 | -0.08635 | Q86WR0 | Q86WR0 | CCDC25 | Coiled-coil | Color2 [Da | 1 | 4 |
| | 0 | -1.01764 | Q86WT1;C | Q86WT1;C | TTC30A;TT | Tetratricoן | Color2 [Da | 1 | 4 |
| | 0.227972 | -0.07764 | Q86X02 | Q86X02 | CDR2L | Cerebellar | Color2 [Da | 1 | 4 |
| | 1.114278 | -0.12236 | Q86X10 | Q86X10 | RALGAPB | Ral GTPase | Color2 [Da | 1 | 4 |
| + | 1.865479 | -1.11665 | Q86X27 | Q86X27 | RALGPS2 | Ras-specifi | Color2 [Da | 1 | 4 |
| + | 0.918392 | -0.48415 | Q86X29 | Q86X29 | LSR | Lipolysis-st | Color2 [Da | 1 | 4 |
| + | 1.054583 | 0.770441 | Q86X52 | Q86X52 | CHSY1 | Chondroiti | Color2 [Da | 1 | 4 |
| + | 2.214251 | -0.91025 | Q86X55 | Q86X55 | CARM1 | Histone-ar | Color2 [Da | 1 | 4 |
| + | 0.915893 | -0.60544 | Q86X67 | Q86X67 | NUDT13 | NAD(P)H p | Color2 [Da | 1 | 4 |
| + | 2.373193 | -1.22919 | Q86X76 | Q86X76 | NIT1 | Deaminate | Color2 [Da | 1 | 4 |
| | 0.218749 | 0.111464 | Q86X83 | Q86X83 | COMMD2 | COMM do | Color2 [Da | 1 | 4 |
| | 0 | -1.56181 | Q86XA9 | Q86XA9 | HEATR5A | HEAT repe | Color2 [Da | 1 | 4 |
| | 0 | 0.704637 | Q86XD5 | Q86XD5 | FAM131B | Protein FA | Color2 [Da | 1 | 4 |
| | 0.477374 | 0.15801 | Q86XE3 | Q86XE3 | MICU3 | Calcium up | Color2 [Da | 1 | 4 |
| | 0 | NaN | Q86XI2 | Q86XI2 | NCAPG2 | Condensin | Color2 [Da | 1 | 4 |
| + | 1.105389 | -0.71787 | Q86XL3 | Q86XL3 | ANKLE2 | Ankyrin re | Color2 [Da | 1 | 4 |
| | 0 | NaN | Q86XN8 | Q86XN8 | MEX3D | RNA-bindi | Color2 [Da | 1 | 4 |
| + | 2.248717 | -0.66387 | Q86XP3 | Q86XP3 | DDX42 | ATP-deper | Color2 [Da | 1 | 4 |
| | 0.520032 | -0.2314 | Q86XX4 | Q86XX4 | FRAS1 | Extracellul | Color2 [Da | 1 | 4 |
| + | 1.988903 | -0.6138 | Q86XZ4 | Q86XZ4 | SPATS2 | Spermatog | Color2 [Da | 1 | 4 |
| + | 1.259466 | -1.13749 | Q86Y37 | Q86Y37 | CACUL1 | CDK2-asso | Color2 [Da | 1 | 4 |
| + | 1.815048 | 1.383471 | Q86Y39 | Q86Y39 | NDUFA11 | NADH deh | Color2 [Da | 1 | 4 |
| + | 2.23296 | -1.28393 | Q86Y56 | Q86Y56 | DNAAF5 | Dynein axo | Color2 [Da | 1 | 4 |
| + | 3.32789 | 2.533842 | Q86Y82 | Q86Y82 | STX12 | Syntaxin-1 | Color2 [Da | 1 | 4 |
| | 0 | -0.68029 | Q86YD1 | Q86YD1 | PTOV1 | Prostate tu | Color2 [Da | 1 | 4 |
| + | 2.671987 | 0.99957 | Q86YL5 | Q86YL5 | TDRP | Testis deve | Color2 [Da | 1 | 4 |
| + | 3.393478 | 1.137641 | Q86YM7 | Q86YM7 | HOMER1 | Homer pro | Color2 [Da | 1 | 4 |
| | 0 | -1.21268 | Q86YN1 | Q86YN1 | DOLPP1 | Dolichyldip | Color2 [Da | 1 | 4 |
| + | 2.633842 | -0.37878 | Q86YP4 | Q86YP4 | GATAD2A | Transcripti | Color2 [Da | 1 | 4 |
| | 0.146142 | -0.1492 | Q86YQ8 | Q86YQ8 | CPNE8 | Copine-8 | Color2 [Da | 1 | 4 |
| | 0.302019 | 0.112864 | Q86YR5 | Q86YR5 | GPSM1 | G-protein- | Color2 [Da | 1 | 4 |
| + | 2.668346 | -0.8605 | Q86YS7 | Q86YS7 | C2CD5 | C2 domain | Color2 [Da | 1 | 4 |
| + | 2.101594 | 0.530142 | Q86YZ3 | Q86YZ3 | HRNR | Hornerin | Color2 [Da | 1 | 4 |
| | 0.862473 | 0.306965 | Q86Z20 | Q86Z20 | CCDC125 | Coiled-coil | Color2 [Da | 1 | 4 |
| + | 1.109417 | -0.81636 | Q86Z23 | Q86Z23 | C1QL4 | Compleme | Color2 [Da | 1 | 4 |
| | 0.895788 | 0.279569 | Q8IU81 | Q8IU81 | IRF2BP1 | Interferon | Color2 [Da | 1 | 4 |
| + | 3.011077 | 0.997941 | Q8IU85 | Q8IU85 | CAMK1D | Calcium/ca | Color2 [Da | 1 | 4 |
| | 0.550536 | -0.5319 | Q8IUC4 | Q8IUC4 | RHPN2 | Rhophilin-: | Color2 [Da | 1 | 4 |
| + | 4.389274 | 1.086207 | Q8IUD2 | Q8IUD2 | ERC1 | ELKS/Rab6 | Color2 [Da | 1 | 4 |
| + | 2.146259 | 2.189896 | Q8IUE6 | Q8IUE6 | H2AC21 | Histone H2 | Color2 [Da | 1 | 4 |
| | 0.906848 | -0.28384 | Q8IUF1;Q9 | Q8IUF1;Q9 | CBWD2;CB | COBW dor | Color2 [Da | 1 | 4 |
| | 0.256208 | 0.086108 | Q8IUH3 | Q8IUH3 | RBM45 | RNA-bindi | Color2 [Da | 1 | 4 |
| | 0 | NaN | Q8IUH5 | Q8IUH5 | ZDHHC17 | Palmitoylt | Color2 [Da | 1 | 4 |
| + | 0.916369 | 1.244287 | Q8IUQ0 | Q8IUQ0 | CLVS1 | Clavesin-1 | Color2 [Da | 1 | 4 |

| | | | | | | | | | |
|---|---|---|---|---|---|---|---|---|---|
| + | 1.269812 | 3.040733 | Q8IUR6 | Q8IUR6 | CREBRF | CREB3 reg | Color2 [Da | 1 | 4 |
| | 0.202826 | -0.15426 | Q8IUR7 | Q8IUR7 | ARMC8 | Armadillo | Color2 [Da | 1 | 4 |
| | 0 | NaN | Q8IUW5 | Q8IUW5 | RELL1 | RELT-like p | Color2 [Da | 1 | 4 |
| + | 3.532921 | -2.26295 | Q8IUX7 | Q8IUX7 | AEBP1 | Adipocyte | Color2 [Da | 1 | 4 |
| + | 2.368938 | 0.80573 | Q8IV08 | Q8IV08 | PLD3 | 5'-3' exonu | Color2 [Da | 1 | 4 |
| + | 2.171969 | -1.13812 | Q8IV32 | Q8IV32 | CCDC71 | Coiled-coil | Color2 [Da | 1 | 4 |
| | 0.740479 | 0.131334 | Q8IV36 | Q8IV36 | HID1 | Protein HII | Color2 [Da | 1 | 4 |
| + | 1.044602 | -0.81921 | Q8IV38 | Q8IV38 | ANKMY2 | Ankyrin re | Color2 [Da | 1 | 4 |
| + | 2.483273 | 2.391672 | Q8IVD9 | Q8IVD9 | NUDCD3 | NudC dom | Color2 [Da | 1 | 4 |
| + | 2.19588 | 0.969337 | Q8IVF2 | Q8IVF2 | AHNAK2 | Protein AH | Color2 [Da | 1 | 4 |
| | 0 | NaN | Q8IVF5 | Q8IVF5 | TIAM2 | Rho guanir | Color2 [Da | 1 | 4 |
| | 0.001963 | 0.00072 | Q8IVF7 | Q8IVF7 | FMNL3 | Formin-like | Color2 [Da | 1 | 4 |
| | 0.156418 | -0.06774 | Q8IVH4 | Q8IVH4 | MMAA | Methylma | Color2 [Da | 1 | 4 |
| | 0 | NaN | Q8IVH8 | Q8IVH8 | MAP4K3 | Mitogen-a | Color2 [Da | 1 | 4 |
| | 0.001246 | 0.000713 | Q8IVL1 | Q8IVL1 | NAV2 | Neuron na | Color2 [Da | 1 | 4 |
| | 0 | -1.05368 | Q8IVL6 | Q8IVL6 | P3H3 | Prolyl 3-hy | Color2 [Da | 1 | 4 |
| + | 2.778355 | 0.846019 | Q8IVM0 | Q8IVM0 | CCDC50 | Coiled-coil | Color2 [Da | 1 | 4 |
| + | 1.676228 | -0.80123 | Q8IVS2 | Q8IVS2 | MCAT | Malonyl-C | Color2 [Da | 1 | 4 |
| + | 1.413434 | 0.602566 | Q8IVT5 | Q8IVT5 | KSR1 | Kinase sup | Color2 [Da | 1 | 4 |
| | 0 | NaN | Q8IVW6 | Q8IVW6 | ARID3B | AT-rich int | Color2 [Da | 1 | 4 |
| | 0.243777 | -0.1863 | Q8IW35 | Q8IW35 | CEP97 | Centrosom | Color2 [Da | 1 | 4 |
| + | 2.757218 | 0.78604 | Q8IW45 | Q8IW45 | NAXD | ATP-deper | Color2 [Da | 1 | 4 |
| + | 0.862444 | 1.040958 | Q8IW52 | Q8IW52 | SLITRK4 | SLIT and N | Color2 [Da | 1 | 4 |
| + | 0.874254 | 0.973458 | Q8IW75 | Q8IW75 | SERPINA12 | Serpin A12 | Color2 [Da | 1 | 4 |
| + | 1.649776 | -0.54473 | Q8IWA0 | Q8IWA0 | WDR75 | WD repeat | Color2 [Da | 1 | 4 |
| + | 1.1877 | -1.03359 | Q8IWA4 | Q8IWA4 | MFN1 | Mitofusin- | Color2 [Da | 1 | 4 |
| + | 1.829006 | -0.83672 | Q8IWA5 | Q8IWA5 | SLC44A2 | Choline tra | Color2 [Da | 1 | 4 |
| | 0 | NaN | Q8IWB1 | Q8IWB1 | ITPRIP | Inositol 1,4 | Color2 [Da | 1 | 4 |
| + | 0.833865 | -0.49614 | Q8IWB7 | Q8IWB7 | WDFY1 | WD repeat | Color2 [Da | 1 | 4 |
| | 0.951343 | -0.15269 | Q8IWE2 | Q8IWE2 | FAM114A1 | Protein NC | Color2 [Da | 1 | 4 |
| | 0 | 0.226848 | Q8IWE5 | Q8IWE5 | PLEKHM2 | Pleckstrin | Color2 [Da | 1 | 4 |
| | 0.202034 | 0.227768 | Q8IWJ2 | Q8IWJ2 | GCC2 | GRIP and c | Color2 [Da | 1 | 4 |
| | 0.23424 | -0.50682 | Q8IWL3 | Q8IWL3 | HSCB | Iron-sulfur | Color2 [Da | 1 | 4 |
| + | 1.419284 | 0.691847 | Q8IWQ3 | Q8IWQ3 | BRSK2 | Serine/thr | Color2 [Da | 1 | 4 |
| + | 1.09841 | 0.343086 | Q8IWR0 | Q8IWR0 | ZC3H7A | Zinc finger | Color2 [Da | 1 | 4 |
| + | 0.840245 | -0.98734 | Q8IWS0 | Q8IWS0 | PHF6 | PHD finger | Color2 [Da | 1 | 4 |
| + | 1.395296 | -0.88335 | Q8IWT0 | Q8IWT0 | ZBTB8OS | Protein arc | Color2 [Da | 1 | 4 |
| | 0 | 1.371442 | Q8IWT6 | Q8IWT6 | LRRC8A | Volume-re | Color2 [Da | 1 | 4 |
| + | 1.433732 | 0.820048 | Q8IWU5 | Q8IWU5 | SULF2 | Extracellul | Color2 [Da | 1 | 4 |
| | 0.571485 | 0.336353 | Q8IWU6 | Q8IWU6 | SULF1 | Extracellul | Color2 [Da | 1 | 4 |
| + | 1.470015 | -1.65716 | Q8IWV7 | Q8IWV7 | UBR1 | E3 ubiquiti | Color2 [Da | 1 | 4 |
| | 0 | NaN | Q8IWV8 | Q8IWV8 | UBR2 | E3 ubiquiti | Color2 [Da | 1 | 4 |
| + | 2.494792 | 0.845687 | Q8IWW6 | Q8IWW6 | ARHGAP12 | Rho GTPas | Color2 [Da | 1 | 4 |
| | 0 | NaN | Q8IWW8 | Q8IWW8 | ADHFE1 | Hydroxyac | Color2 [Da | 1 | 4 |
| | 0.78863 | 0.121209 | Q8IWX8 | Q8IWX8 | CHERP | Calcium hc | Color2 [Da | 1 | 4 |
| + | 0.975576 | 0.805987 | Q8IWY4;Q | Q8IWY4;Q | SCUBE1;SC | Signal pep | Color2 [Da | 1 | 4 |
| + | 2.034319 | -0.37099 | Q8IWY9 | Q8IWY9 | CDAN1 | Codanin-1 | Color2 [Da | 1 | 4 |

| | | | | | | | | | |
|---|---|---|---|---|---|---|---|---|---|
| | 0.257385 | 0.371003 | Q8IWZ3 | Q8IWZ3 | ANKHD1 | Ankyrin re | Color2 [Da | 1 | 4 |
| | 0.324784 | -0.08991 | Q8IWZ8 | Q8IWZ8 | SUGP1 | SURP and ( | Color2 [Da | 1 | 4 |
| | 0.473993 | -0.10694 | Q8IX01 | Q8IX01 | SUGP2 | SURP and ( | Color2 [Da | 1 | 4 |
| + | 1.210284 | 0.466972 | Q8IX03 | Q8IX03 | WWC1 | Protein KIE | Color2 [Da | 1 | 4 |
| | 0.246296 | 0.096782 | Q8IX12 | Q8IX12 | CCAR1 | Cell divisio | Color2 [Da | 1 | 4 |
| | 0.299711 | -0.25576 | Q8IX18 | Q8IX18 | DHX40 | Probable A | Color2 [Da | 1 | 4 |
| | 0.008487 | 0.007657 | Q8IX30 | Q8IX30 | SCUBE3 | Signal pep | Color2 [Da | 1 | 4 |
| + | 2.261262 | -0.80299 | Q8IXB1 | Q8IXB1 | DNAJC10 | DnaJ homo | Color2 [Da | 1 | 4 |
| + | 1.494364 | 1.518601 | Q8IXF0 | Q8IXF0 | NPAS3 | Neuronal P | Color2 [Da | 1 | 4 |
| + | 2.770282 | -1.14167 | Q8IXH7 | Q8IXH7 | NELFCD | Negative e | Color2 [Da | 1 | 4 |
| | 0.425181 | -0.32697 | Q8IXI1 | Q8IXI1 | RHOT2 | Mitochonc | Color2 [Da | 1 | 4 |
| | 0.002766 | -0.00393 | Q8IXI2 | Q8IXI2 | RHOT1 | Mitochonc | Color2 [Da | 1 | 4 |
| + | 1.02102 | 0.393971 | Q8IXJ6 | Q8IXJ6 | SIRT2 | NAD-depe | Color2 [Da | 1 | 4 |
| + | 1.403155 | -0.57993 | Q8IXK0 | Q8IXK0 | PHC2 | Polyhome | Color2 [Da | 1 | 4 |
| + | 0.852839 | 0.838395 | Q8IXM2 | Q8IXM2 | BAP18 | Chromatin | Color2 [Da | 1 | 4 |
| + | 4.963437 | 1.581746 | Q8IXM3 | Q8IXM3 | MRPL41 | 39S riboso | Color2 [Da | 1 | 4 |
| | 0 | -0.77896 | Q8IXM6 | Q8IXM6 | NRM | Nurim | Color2 [Da | 1 | 4 |
| | 0.377862 | -0.78267 | Q8IXQ3 | Q8IXQ3 | C9orf40 | Uncharact | Color2 [Da | 1 | 4 |
| + | 2.520212 | 1.904202 | Q8IXQ4 | Q8IXQ4 | GPALPP1 | GPALPP m | Color2 [Da | 1 | 4 |
| + | 3.101517 | 2.265014 | Q8IXS6 | Q8IXS6 | PALM2 | Paralemm | Color2 [Da | 1 | 4 |
| | 0 | NaN | Q8IXS8 | Q8IXS8 | FAM126B | Protein FA | Color2 [Da | 1 | 4 |
| + | 1.342128 | -0.43676 | Q8IXT5 | Q8IXT5 | RBM12B | RNA-bindi | Color2 [Da | 1 | 4 |
| | 0 | NaN | Q8IXW5 | Q8IXW5 | RPAP2 | Putative R | Color2 [Da | 1 | 4 |
| + | 0.875456 | -0.78309 | Q8IY17 | Q8IY17 | PNPLA6 | Patatin-lik | Color2 [Da | 1 | 4 |
| + | 2.062527 | -0.65475 | Q8IY22 | Q8IY22 | CMIP | C-Maf-ind | Color2 [Da | 1 | 4 |
| | 0 | -0.2453 | Q8IY26 | Q8IY26 | PLPP6 | Polyisopre | Color2 [Da | 1 | 4 |
| | 0.167178 | -0.17477 | Q8IY31 | Q8IY31 | IFT20 | Intraflagell | Color2 [Da | 1 | 4 |
| | 0.033605 | 0.122749 | Q8IY33 | Q8IY33 | MICALL2 | MICAL-like | Color2 [Da | 1 | 4 |
| | 0.38988 | -0.15249 | Q8IY37 | Q8IY37 | DHX37 | Probable A | Color2 [Da | 1 | 4 |
| | 0.216588 | 0.201976 | Q8IY63 | Q8IY63 | AMOTL1 | Angiomoti | Color2 [Da | 1 | 4 |
| + | 2.30654 | -1.3332 | Q8IY67 | Q8IY67 | RAVER1 | Ribonucle | Color2 [Da | 1 | 4 |
| | 0.444312 | -0.07424 | Q8IY81 | Q8IY81 | FTSJ3 | pre-rRNA 2 | Color2 [Da | 1 | 4 |
| | 0 | -0.96682 | Q8IY95 | Q8IY95 | TMEM192 | Transmem | Color2 [Da | 1 | 4 |
| + | 1.364826 | 0.256528 | Q8IYB3 | Q8IYB3 | SRRM1 | Serine/arg | Color2 [Da | 1 | 4 |
| | 0.09654 | -0.08257 | Q8IYB5 | Q8IYB5 | SMAP1 | Stromal m | Color2 [Da | 1 | 4 |
| + | 1.389234 | -0.57653 | Q8IYB7 | Q8IYB7 | DIS3L2 | DIS3-like e | Color2 [Da | 1 | 4 |
| + | 1.35567 | -0.93893 | Q8IYB8 | Q8IYB8 | SUPV3L1 | ATP-deper | Color2 [Da | 1 | 4 |
| + | 1.065172 | -0.40299 | Q8IYD1 | Q8IYD1 | GSPT2 | Eukaryotic | Color2 [Da | 1 | 4 |
| | 0.124895 | -0.04746 | Q8IYI6 | Q8IYI6 | EXOC8 | Exocyst co | Color2 [Da | 1 | 4 |
| | 0 | NaN | Q8IYK4 | Q8IYK4 | COLGALT2 | Procollage | Color2 [Da | 1 | 4 |
| + | 1.404876 | -0.57219 | Q8IYK8 | Q8IYK8 | REM2 | GTP-bindir | Color2 [Da | 1 | 4 |
| | 0 | NaN | Q8IYM9 | Q8IYM9 | TRIM22 | E3 ubiquiti | Color2 [Da | 1 | 4 |
| | 0 | -1.90279 | Q8IYQ7 | Q8IYQ7 | THNSL1 | Threonine | Color2 [Da | 1 | 4 |
| | 0 | -0.22445 | Q8IYR6 | Q8IYR6 | TMEFF1 | Tomoregu | Color2 [Da | 1 | 4 |
| + | 2.983669 | -2.7131 | Q8IYS2 | Q8IYS2 | KIAA2013 | Uncharact | Color2 [Da | 1 | 4 |
| | 0 | NaN | Q8IYU8 | Q8IYU8 | MICU2 | Calcium up | Color2 [Da | 1 | 4 |
| + | 4.597658 | -0.8108 | Q8IZ07 | Q8IZ07 | ANKRD13A | Ankyrin re | Color2 [Da | 1 | 4 |

| | | | | | | | | | |
|---|---|---|---|---|---|---|---|---|---|
| | 0 | NaN | Q8IZ13 | Q8IZ13 | ZBED8 | Protein ZB | Color2 [Da | 1 | 4 |
| + | 2.682928 | 1.641532 | Q8IZ21 | Q8IZ21 | PHACTR4 | Phosphata | Color2 [Da | 1 | 4 |
| + | 1.194956 | 0.586111 | Q8IZ40 | Q8IZ40 | RCOR2 | REST corep | Color2 [Da | 1 | 4 |
| | 0.573107 | 0.143881 | Q8IZ52 | Q8IZ52 | CHPF | Chondroiti | Color2 [Da | 1 | 4 |
| | 0 | -0.7415 | Q8IZ69 | Q8IZ69 | TRMT2A | tRNA (urac | Color2 [Da | 1 | 4 |
| + | 1.420345 | -0.87113 | Q8IZ81 | Q8IZ81 | ELMOD2 | ELMO dom | Color2 [Da | 1 | 4 |
| + | 1.570169 | -0.65778 | Q8IZ83 | Q8IZ83 | ALDH16A1 | Aldehyde ( | Color2 [Da | 1 | 4 |
| + | 1.529801 | -0.37032 | Q8IZA0 | Q8IZA0 | KIAA0319L | Dyslexia-as | Color2 [Da | 1 | 4 |
| | 0 | NaN | Q8IZD0 | Q8IZD0 | SAMD14 | Sterile alpl | Color2 [Da | 1 | 4 |
| + | 2.817107 | -0.76662 | Q8IZD4 | Q8IZD4 | DCP1B | mRNA-dec | Color2 [Da | 1 | 4 |
| | 0.427301 | -0.24047 | Q8IZD9 | Q8IZD9 | DOCK3 | Dedicator | Color2 [Da | 1 | 4 |
| + | 1.486374 | -0.27002 | Q8IZL8 | Q8IZL8 | PELP1 | Proline-, gl | Color2 [Da | 1 | 4 |
| | 0 | NaN | Q8IZM8 | Q8IZM8 | ZNF654 | Zinc finger | Color2 [Da | 1 | 4 |
| + | 5.13977 | 1.924281 | Q8IZP0 | Q8IZP0 | ABI1 | Abl interac | Color2 [Da | 1 | 4 |
| | 0.640299 | 0.532079 | Q8IZP9 | Q8IZP9 | ADGRG2 | Adhesion ( | Color2 [Da | 1 | 4 |
| | 0 | -1.19973 | Q8IZQ1 | Q8IZQ1 | WDFY3 | WD repeat | Color2 [Da | 1 | 4 |
| | 0.204771 | 0.080877 | Q8IZQ5 | Q8IZQ5 | SELENOH | Selenoprot | Color2 [Da | 1 | 4 |
| | 0 | NaN | Q8IZS8 | Q8IZS8 | CACNA2D3 | Voltage-de | Color2 [Da | 1 | 4 |
| | 0 | 1.696162 | Q8IZT6 | Q8IZT6 | ASPM | Abnormal | Color2 [Da | 1 | 4 |
| + | 4.74662 | -2.24078 | Q8N0U8 | Q8N0U8 | VKORC1L1 | Vitamin K ( | Color2 [Da | 1 | 4 |
| | 0.512912 | 0.257413 | Q8N0V4 | Q8N0V4 | LGI2 | Leucine-ric | Color2 [Da | 1 | 4 |
| | 0 | NaN | Q8N0W3 | Q8N0W3 | FCSK | L-fucose ki | Color2 [Da | 1 | 4 |
| + | 1.020965 | -0.40198 | Q8N0W4;( | Q8N0W4;( | NLGN4X;N | Neuroligin | Color2 [Da | 1 | 4 |
| + | 1.974933 | -0.64468 | Q8N0X4 | Q8N0X4 | CLYBL | Citramalyl- | Color2 [Da | 1 | 4 |
| | 0.051942 | -0.0262 | Q8N0X7 | Q8N0X7 | SPART | Spartin | Color2 [Da | 1 | 4 |
| | 0 | 0.606615 | Q8N0Z3 | Q8N0Z3 | SPICE1 | Spindle an | Color2 [Da | 1 | 4 |
| + | 1.847816 | -0.74187 | Q8N0Z6 | Q8N0Z6 | TTC5 | Tetratricop | Color2 [Da | 1 | 4 |
| + | 3.526017 | 2.790845 | Q8N111 | Q8N111 | CEND1 | Cell cycle e | Color2 [Da | 1 | 4 |
| | 0.296955 | 0.112739 | Q8N122 | Q8N122 | RPTOR | Regulatory | Color2 [Da | 1 | 4 |
| | 0.090974 | -0.01681 | Q8N126 | Q8N126 | CADM3 | Cell adhesi | Color2 [Da | 1 | 4 |
| + | 3.828397 | 1.507668 | Q8N128 | Q8N128 | FAM177A1 | Protein FA | Color2 [Da | 1 | 4 |
| | 0.087721 | 0.064288 | Q8N129 | Q8N129 | CNPY4 | Protein car | Color2 [Da | 1 | 4 |
| + | 1.763623 | -1.45587 | Q8N138 | Q8N138 | ORMDL3 | ORM1-like | Color2 [Da | 1 | 4 |
| | 0.521747 | -0.24987 | Q8N142 | Q8N142 | ADSS1 | Adenylosu | Color2 [Da | 1 | 4 |
| | 0 | NaN | Q8N145 | Q8N145 | LGI3 | Leucine-ric | Color2 [Da | 1 | 4 |
| + | 2.845007 | -1.43403 | Q8N158 | Q8N158 | GPC2 | Glypican-2 | Color2 [Da | 1 | 4 |
| | 0.240616 | -0.09873 | Q8N163 | Q8N163 | CCAR2 | Cell cycle a | Color2 [Da | 1 | 4 |
| + | 3.814699 | 2.74189 | Q8N183 | Q8N183 | NDUFAF2 | NADH deh | Color2 [Da | 1 | 4 |
| | 0.414696 | -0.48474 | Q8N1B4 | Q8N1B4 | VPS52 | Vacuolar p | Color2 [Da | 1 | 4 |
| | 0.438472 | -0.20365 | Q8N1F7 | Q8N1F7 | NUP93 | Nuclear pc | Color2 [Da | 1 | 4 |
| + | 1.085118 | -0.77957 | Q8N1F8 | Q8N1F8 | STK11IP | Serine/thr | Color2 [Da | 1 | 4 |
| + | 2.055463 | -0.60476 | Q8N1G0 | Q8N1G0 | ZNF687 | Zinc finger | Color2 [Da | 1 | 4 |
| + | 1.627218 | -1.38221 | Q8N1G2 | Q8N1G2 | CMTR1 | Cap-specif | Color2 [Da | 1 | 4 |
| | 0.163908 | 0.058816 | Q8N1G4 | Q8N1G4 | LRRC47 | Leucine-ric | Color2 [Da | 1 | 4 |
| + | 0.910892 | 0.588536 | Q8N1I0 | Q8N1I0 | DOCK4 | Dedicator | Color2 [Da | 1 | 4 |
| + | 1.045867 | 1.500396 | Q8N1N4 | Q8N1N4 | KRT78 | Keratin, ty | Color2 [Da | 1 | 4 |
| | 0.05204 | 0.025265 | Q8N2F6 | Q8N2F6 | ARMC10 | Armadillo | Color2 [Da | 1 | 4 |

| | | | | | | | | | |
|---|---|---|---|---|---|---|---|---|---|
| | 0.572285 | 1.670908 | Q8N2G4 | Q8N2G4 | LYPD1 | Ly6/PLAUF | Color2 | [Da | 1 | 4 |
| | 0.124645 | -0.02219 | Q8N2K0 | Q8N2K0 | ABHD12 | Lysophosp | Color2 | [Da | 1 | 4 |
| | 0.369305 | 0.926717 | Q8N2U0 | Q8N2U0 | TMEM256 | Transmem | Color2 | [Da | 1 | 4 |
| | 0.512127 | -0.83805 | Q8N302 | Q8N302 | AGGF1 | Angiogenic | Color2 | [Da | 1 | 4 |
| + | 1.416048 | -0.48784 | Q8N335 | Q8N335 | GPD1L | Glycerol-3- | Color2 | [Da | 1 | 4 |
| + | 1.707987 | 0.880517 | Q8N350 | Q8N350 | CBARP | Voltage-de | Color2 | [Da | 1 | 4 |
| | 0.223176 | -0.14849 | Q8N357 | Q8N357 | SLC35F6 | Solute carr | Color2 | [Da | 1 | 4 |
| | 0 | NaN | Q8N370 | Q8N370 | SLC43A2 | Large neut | Color2 | [Da | 1 | 4 |
| | 0 | NaN | Q8N392 | Q8N392 | ARHGAP18 | Rho GTPas | Color2 | [Da | 1 | 4 |
| + | 0.78723 | -0.64641 | Q8N3C0 | Q8N3C0 | ASCC3 | Activating | Color2 | [Da | 1 | 4 |
| | 0 | NaN | Q8N3E9 | Q8N3E9 | PLCD3 | 1-phospha | Color2 | [Da | 1 | 4 |
| + | 3.677342 | 3.91375 | Q8N3F0 | Q8N3F0 | MTURN | Maturin | Color2 | [Da | 1 | 4 |
| + | 0.77483 | 0.62758 | Q8N3F8 | Q8N3F8 | MICALL1 | MICAL-like | Color2 | [Da | 1 | 4 |
| | 0.295893 | -0.24348 | Q8N3J6 | Q8N3J6 | CADM2 | Cell adhesi | Color2 | [Da | 1 | 4 |
| | 0 | -0.86396 | Q8N3P4 | Q8N3P4 | VPS8 | Vacuolar p | Color2 | [Da | 1 | 4 |
| + | 1.511041 | 0.263096 | Q8N3R9 | Q8N3R9 | PALS1 | Protein PA | Color2 | [Da | 1 | 4 |
| | 0 | -0.49807 | Q8N3U4 | Q8N3U4 | STAG2 | Cohesin su | Color2 | [Da | 1 | 4 |
| + | 1.062253 | 0.405678 | Q8N3X1 | Q8N3X1 | FNBP4 | Formin-bir | Color2 | [Da | 1 | 4 |
| | 0 | -0.83431 | Q8N414 | Q8N414 | PGBD5 | PiggyBac tr | Color2 | [Da | 1 | 4 |
| + | 1.813711 | -0.48998 | Q8N428 | Q8N428 | GALNT16 | Polypeptid | Color2 | [Da | 1 | 4 |
| | 0 | NaN | Q8N461 | Q8N461 | FBXL16 | F-box/LRR- | Color2 | [Da | 1 | 4 |
| | 0.242103 | -0.09117 | Q8N465 | Q8N465 | D2HGDH | D-2-hydro: | Color2 | [Da | 1 | 4 |
| | 0.114251 | 0.118983 | Q8N474 | Q8N474 | SFRP1 | Secreted fr | Color2 | [Da | 1 | 4 |
| + | 2.356145 | 1.178418 | Q8N488 | Q8N488 | RYBP | RING1 and | Color2 | [Da | 1 | 4 |
| + | 1.489099 | 1.252388 | Q8N490 | Q8N490 | PNKD | Probable h | Color2 | [Da | 1 | 4 |
| + | 1.807628 | 1.855663 | Q8N4C8 | Q8N4C8 | MINK1 | Misshapen | Color2 | [Da | 1 | 4 |
| + | 3.941746 | 2.032108 | Q8N4H5 | Q8N4H5 | TOMM5 | Mitochonc | Color2 | [Da | 1 | 4 |
| + | 1.98905 | -1.58953 | Q8N4L2 | Q8N4L2 | PIP4P2 | Type 2 pho | Color2 | [Da | 1 | 4 |
| + | 2.36181 | 2.295595 | Q8N4P3 | Q8N4P3 | HDDC3 | Guanosine | Color2 | [Da | 1 | 4 |
| + | 1.207587 | 0.748459 | Q8N4Q0 | Q8N4Q0 | PTGR3 | Prostaglan | Color2 | [Da | 1 | 4 |
| + | 2.648462 | 2.816748 | Q8N4Q1 | Q8N4Q1 | CHCHD4 | Mitochonc | Color2 | [Da | 1 | 4 |
| + | 1.739646 | 1.07861 | Q8N4T8 | Q8N4T8 | CBR4 | 3-oxoacyl- | Color2 | [Da | 1 | 4 |
| + | 1.186652 | 0.802948 | Q8N4V1 | Q8N4V1 | MMGT1 | ER membr | Color2 | [Da | 1 | 4 |
| | 0 | NaN | Q8N4V2 | Q8N4V2 | SVOP | Synaptic ve | Color2 | [Da | 1 | 4 |
| | 0.32651 | 0.221977 | Q8N556 | Q8N556 | AFAP1 | Actin filam | Color2 | [Da | 1 | 4 |
| + | 1.45228 | -0.43886 | Q8N568 | Q8N568 | DCLK2 | Serine/thr | Color2 | [Da | 1 | 4 |
| + | 2.130597 | 1.122628 | Q8N573 | Q8N573 | OXR1 | Oxidation | Color2 | [Da | 1 | 4 |
| | 0 | -0.74988 | Q8N594 | Q8N594 | MPND | MPN doma | Color2 | [Da | 1 | 4 |
| | 0.126665 | -0.12458 | Q8N5A5 | Q8N5A5 | ZGPAT | Zinc finger | Color2 | [Da | 1 | 4 |
| + | 0.85961 | 0.525578 | Q8N5C8 | Q8N5C8 | TAB3 | TGF-beta-a | Color2 | [Da | 1 | 4 |
| | 0.240713 | -0.31996 | Q8N5D0 | Q8N5D0 | WDTC1 | WD and te | Color2 | [Da | 1 | 4 |
| + | 0.817406 | 0.872568 | Q8N5F7 | Q8N5F7 | NKAP | NF-kappa- | Color2 | [Da | 1 | 4 |
| + | 2.673198 | 3.731733 | Q8N5G0 | Q8N5G0 | SMIM20 | Small integ | Color2 | [Da | 1 | 4 |
| | 0 | -0.78641 | Q8N5I4 | Q8N5I4 | DHRSX | Dehydroge | Color2 | [Da | 1 | 4 |
| + | 1.104561 | 0.547734 | Q8N5I9 | Q8N5I9 | NOPCHAP1 | NOP prote | Color2 | [Da | 1 | 4 |
| + | 0.735815 | 0.830464 | Q8N5K1 | Q8N5K1 | CISD2 | CDGSH iro | Color2 | [Da | 1 | 4 |
| | 0.869332 | 0.157392 | Q8N5M1 | Q8N5M1 | ATPAF2 | ATP syntha | Color2 | [Da | 1 | 4 |

| | | | | | | | | | |
|---|---|---|---|---|---|---|---|---|---|
| + | 1.830326 | 0.81002 | Q8N5M4 | Q8N5M4 | TTC9C | Tetratricoן | Color2 [Da | 1 | 4 |
| | 0.144965 | -0.48036 | Q8N5M9 | Q8N5M9 | JAGN1 | Protein jag | Color2 [Da | 1 | 4 |
| + | 3.222628 | 1.316967 | Q8N5N7 | Q8N5N7 | MRPL50 | 39S riboso | Color2 [Da | 1 | 4 |
| | 0.089639 | 0.128284 | Q8N5Y2 | Q8N5Y2 | MSL3 | Male-spec | Color2 [Da | 1 | 4 |
| | 0 | NaN | Q8N5Y8 | Q8N5Y8 | PARP16 | Protein mc | Color2 [Da | 1 | 4 |
| | 0.527359 | 0.545672 | Q8N5Z0 | Q8N5Z0 | AADAT | Kynurenin | Color2 [Da | 1 | 4 |
| | 0.218531 | 0.255419 | Q8N608 | Q8N608 | DPP10 | Inactive di | Color2 [Da | 1 | 4 |
| + | 2.043097 | 0.667984 | Q8N668 | Q8N668 | COMMD1 | COMM do | Color2 [Da | 1 | 4 |
| | 0.121556 | 0.095063 | Q8N680 | Q8N680 | ZBTB2 | Zinc finger | Color2 [Da | 1 | 4 |
| + | 2.092024 | 1.061234 | Q8N684 | Q8N684 | CPSF7 | Cleavage a | Color2 [Da | 1 | 4 |
| | 0 | NaN | Q8N6C5 | Q8N6C5 | IGSF1 | Immunogl | Color2 [Da | 1 | 4 |
| + | 1.796057 | -0.1855 | Q8N6H7 | Q8N6H7 | ARFGAP2 | ADP-ribos | Color2 [Da | 1 | 4 |
| | 0 | -1.6109 | Q8N6L1 | Q8N6L1 | KRTCAP2 | Keratinocy | Color2 [Da | 1 | 4 |
| + | 1.079758 | -0.41492 | Q8N6M0 | Q8N6M0 | OTUD6B | Deubiquiti | Color2 [Da | 1 | 4 |
| + | 0.693444 | -0.72008 | Q8N6N2 | Q8N6N2 | TTC9B | Tetratricoן | Color2 [Da | 1 | 4 |
| + | 2.85376 | 2.072511 | Q8N6N3 | Q8N6N3 | C1orf52 | UPF0690 p | Color2 [Da | 1 | 4 |
| + | 1.408563 | 0.772034 | Q8N6N7 | Q8N6N7 | ACBD7 | Acyl-CoA-b | Color2 [Da | 1 | 4 |
| + | 1.027317 | -0.4489 | Q8N6R0 | Q8N6R0 | METTL13 | eEF1A lysi | Color2 [Da | 1 | 4 |
| + | 1.375545 | 0.959133 | Q8N6T3 | Q8N6T3 | ARFGAP1 | ADP-ribos | Color2 [Da | 1 | 4 |
| | 0 | NaN | Q8N6Y2 | Q8N6Y2 | LRRC17 | Leucine-ric | Color2 [Da | 1 | 4 |
| + | 3.453858 | -1.19551 | Q8N766 | Q8N766 | EMC1 | ER membr | Color2 [Da | 1 | 4 |
| + | 1.256362 | 0.382568 | Q8N7H5 | Q8N7H5 | PAF1 | RNA polyn | Color2 [Da | 1 | 4 |
| + | 1.843611 | 1.183081 | Q8N7J2 | Q8N7J2 | AMER2 | APC memb | Color2 [Da | 1 | 4 |
| | 0.095988 | -0.12569 | Q8N806 | Q8N806 | UBR7 | Putative E3 | Color2 [Da | 1 | 4 |
| | 0 | NaN | Q8N8D1 | Q8N8D1 | PDCD7 | Programm | Color2 [Da | 1 | 4 |
| | 0 | -0.99127 | Q8N8E3 | Q8N8E3 | CEP112 | Centrosom | Color2 [Da | 1 | 4 |
| + | 2.510778 | 0.853571 | Q8N8L6 | Q8N8L6 | ARL10 | ADP-ribos | Color2 [Da | 1 | 4 |
| | 0.009978 | -0.00645 | Q8N8N7 | Q8N8N7 | PTGR2 | Prostaglan | Color2 [Da | 1 | 4 |
| | 0 | 0.753548 | Q8N8R5 | Q8N8R5 | C2orf69 | Mitochonc | Color2 [Da | 1 | 4 |
| + | 2.477461 | 1.136799 | Q8N8S7 | Q8N8S7 | ENAH | Protein en | Color2 [Da | 1 | 4 |
| | 0 | -1.60661 | Q8N8Y2 | Q8N8Y2 | ATP6V0D2 | V-type pro | Color2 [Da | 1 | 4 |
| + | 1.25964 | 0.655531 | Q8N954 | Q8N954 | GPATCH11 | G patch dc | Color2 [Da | 1 | 4 |
| | 0 | NaN | Q8N960 | Q8N960 | CEP120 | Centrosom | Color2 [Da | 1 | 4 |
| | 0 | NaN | Q8N961 | Q8N961 | ABTB2 | Ankyrin re | Color2 [Da | 1 | 4 |
| + | 0.931904 | 1.587811 | Q8N983 | Q8N983 | MRPL43 | 39S riboso | Color2 [Da | 1 | 4 |
| + | 2.36728 | -1.28012 | Q8N9F7 | Q8N9F7 | GDPD1 | Lysophosp | Color2 [Da | 1 | 4 |
| | 0 | NaN | Q8N9I0 | Q8N9I0 | SYT2 | Synaptota | Color2 [Da | 1 | 4 |
| | 0.419317 | 0.304056 | Q8N9I9 | Q8N9I9 | DTX3 | Probable E | Color2 [Da | 1 | 4 |
| | 0.085075 | -0.0645 | Q8N9M1 | Q8N9M1 | C19orf47 | Uncharact | Color2 [Da | 1 | 4 |
| + | 1.047705 | 0.446789 | Q8N9N2 | Q8N9N2 | ASCC1 | Activating | Color2 [Da | 1 | 4 |
| + | 1.512512 | 0.779987 | Q8N9N7 | Q8N9N7 | LRRC57 | Leucine-ric | Color2 [Da | 1 | 4 |
| + | 2.318876 | 0.917973 | Q8N9N8 | Q8N9N8 | EIF1AD | Probable F | Color2 [Da | 1 | 4 |
| | 0 | -0.14073 | Q8N9Q2 | Q8N9Q2 | SREK1IP1 | Protein SR | Color2 [Da | 1 | 4 |
| + | 1.598847 | -0.70855 | Q8N9R8 | Q8N9R8 | SCAI | Protein SC | Color2 [Da | 1 | 4 |
| + | 2.027652 | -0.96831 | Q8N9T8 | Q8N9T8 | KRI1 | Protein KR | Color2 [Da | 1 | 4 |
| + | 2.074119 | -0.42422 | Q8NAF0 | Q8NAF0 | ZNF579 | Zinc finger | Color2 [Da | 1 | 4 |
| + | 1.056317 | -0.72897 | Q8NAT1 | Q8NAT1 | POMGNT2 | Protein O- | Color2 [Da | 1 | 4 |

| | | | | | | | | | |
|---|---|---|---|---|---|---|---|---|---|
| | 0.526955 | -0.29281 | Q8NAV1 | Q8NAV1 | PRPF38A | Pre-mRNA | Color2 [Da | 1 | 4 |
| | 0.400039 | 0.118961 | Q8NB37 | Q8NB37 | GATD1 | Glutamine | Color2 [Da | 1 | 4 |
| | 0 | NaN | Q8NB49 | Q8NB49 | ATP11C | Phospholip | Color2 [Da | 1 | 4 |
| | 0.559048 | -0.60074 | Q8NB78 | Q8NB78 | KDM1B | Lysine-spe | Color2 [Da | 1 | 4 |
| + | 0.883235 | -0.42958 | Q8NB90 | Q8NB90 | SPATA5 | ATPase far | Color2 [Da | 1 | 4 |
| | 0.74604 | 0.214047 | Q8NBF2 | Q8NBF2 | NHLRC2 | NHL repea | Color2 [Da | 1 | 4 |
| + | 1.921028 | 0.5082 | Q8NBF6 | Q8NBF6 | AVL9 | Late secret | Color2 [Da | 1 | 4 |
| + | 2.442323 | -1.32719 | Q8NBI6 | Q8NBI6 | XXYLT1 | Xyloside xy | Color2 [Da | 1 | 4 |
| + | 2.088422 | 0.460209 | Q8NBJ4 | Q8NBJ4 | GOLM1 | Golgi mem | Color2 [Da | 1 | 4 |
| + | 2.840155 | -1.42764 | Q8NBJ5 | Q8NBJ5 | COLGALT1 | Procollage | Color2 [Da | 1 | 4 |
| + | 2.695272 | 1.220062 | Q8NBJ7 | Q8NBJ7 | SUMF2 | Inactive C- | Color2 [Da | 1 | 4 |
| | 0 | 0.038085 | Q8NBK3 | Q8NBK3 | SUMF1 | Formylglyc | Color2 [Da | 1 | 4 |
| + | 2.643045 | -0.82025 | Q8NBL1 | Q8NBL1 | POGLUT1 | Protein O-; | Color2 [Da | 1 | 4 |
| + | 3.702276 | -2.32874 | Q8NBM4 | Q8NBM4 | UBAC2 | Ubiquitin-a | Color2 [Da | 1 | 4 |
| + | 1.017807 | -0.5247 | Q8NBM8 | Q8NBM8 | PCYOX1L | Prenylcyst | Color2 [Da | 1 | 4 |
| | 0.073938 | -0.21104 | Q8NBN3 | Q8NBN3 | TMEM87A | Transmem | Color2 [Da | 1 | 4 |
| + | 1.689296 | -0.32682 | Q8NBN7 | Q8NBN7 | RDH13 | Retinol del | Color2 [Da | 1 | 4 |
| | 0 | -0.68332 | Q8NBQ5 | Q8NBQ5 | HSD17B11 | Estradiol 1 | Color2 [Da | 1 | 4 |
| | 0 | NaN | Q8NBR6 | Q8NBR6 | MINDY2 | Ubiquitin c | Color2 [Da | 1 | 4 |
| + | 3.702551 | -0.94304 | Q8NBS9 | Q8NBS9 | TXNDC5 | Thioredoxi | Color2 [Da | 1 | 4 |
| + | 1.02384 | 0.32866 | Q8NBT2 | Q8NBT2 | SPC24 | Kinetochor | Color2 [Da | 1 | 4 |
| | 0.286981 | -0.17154 | Q8NBU5 | Q8NBU5 | ATAD1 | Outer mitc | Color2 [Da | 1 | 4 |
| | 0.710228 | -0.18918 | Q8NBX0 | Q8NBX0 | SCCPDH | Saccharop | Color2 [Da | 1 | 4 |
| + | 1.237366 | 0.844605 | Q8NBZ7 | Q8NBZ7 | UXS1 | UDP-glucu | Color2 [Da | 1 | 4 |
| + | 0.751984 | 1.039938 | Q8NC44 | Q8NC44 | RETREG2 | Reticuloph | Color2 [Da | 1 | 4 |
| + | 4.100599 | 2.80606 | Q8NC51 | Q8NC51 | SERBP1 | Plasminog | Color2 [Da | 1 | 4 |
| | 0 | -1.47857 | Q8NC54 | Q8NC54 | KCT2 | Keratinocy | Color2 [Da | 1 | 4 |
| | 0.475253 | -0.57203 | Q8NC56 | Q8NC56 | LEMD2 | LEM doma | Color2 [Da | 1 | 4 |
| + | 3.133105 | 2.422762 | Q8NC96 | Q8NC96 | NECAP1 | Adaptin ea | Color2 [Da | 1 | 4 |
| + | 1.435232 | -0.63214 | Q8NCA5 | Q8NCA5 | FAM98A | Protein FA | Color2 [Da | 1 | 4 |
| + | 1.329256 | 1.975597 | Q8NCB2 | Q8NCB2 | CAMKV | CaM kinas | Color2 [Da | 1 | 4 |
| | 0.071415 | 0.063715 | Q8NCC3 | Q8NCC3 | PLA2G15 | Phospholip | Color2 [Da | 1 | 4 |
| | 0.924824 | -0.14934 | Q8NCE2 | Q8NCE2 | MTMR14 | Myotubula | Color2 [Da | 1 | 4 |
| + | 0.894585 | 0.377658 | Q8NCF5 | Q8NCF5 | NFATC2IP | NFATC2-in | Color2 [Da | 1 | 4 |
| | 0 | 0.637276 | Q8NCG7 | Q8NCG7 | DAGLB | Diacylglyce | Color2 [Da | 1 | 4 |
| | 0 | -0.31097 | Q8NCH0 | Q8NCH0 | CHST14 | Carbohydr | Color2 [Da | 1 | 4 |
| + | 0.797984 | 0.841553 | Q8NCM8 | Q8NCM8 | DYNC2H1 | Cytoplasm | Color2 [Da | 1 | 4 |
| + | 2.774108 | -1.10704 | Q8NCN5 | Q8NCN5 | PDPR | Pyruvate d | Color2 [Da | 1 | 4 |
| + | 2.514317 | 0.728955 | Q8NCW5 | Q8NCW5 | NAXE | NAD(P)H-h | Color2 [Da | 1 | 4 |
| | 0 | NaN | Q8NCW6 | Q8NCW6 | GALNT11 | Polypeptid | Color2 [Da | 1 | 4 |
| | 0.472635 | 0.262616 | Q8NCY6 | Q8NCY6 | MSANTD4 | Myb/SANT | Color2 [Da | 1 | 4 |
| | 0.014908 | -0.01725 | Q8ND04 | Q8ND04 | SMG8 | Nonsense- | Color2 [Da | 1 | 4 |
| | 0.440194 | 0.350207 | Q8ND23 | Q8ND23 | CARMIL3 | Capping pr | Color2 [Da | 1 | 4 |
| | 0.059182 | 0.028613 | Q8ND24 | Q8ND24 | RNF214 | RING finge | Color2 [Da | 1 | 4 |
| + | 1.129253 | 0.365009 | Q8ND56 | Q8ND56 | LSM14A | Protein LSI | Color2 [Da | 1 | 4 |
| + | 2.12266 | -0.76712 | Q8ND76 | Q8ND76 | CCNY | Cyclin-Y | Color2 [Da | 1 | 4 |
| + | 2.932324 | -2.65076 | Q8ND82 | Q8ND82 | ZNF280C | Zinc finger | Color2 [Da | 1 | 4 |

| | | | | | | | | | |
|---|---|---|---|---|---|---|---|---|---|
| | 0.795105 | -0.39536 | Q8ND83 | Q8ND83 | SLAIN1 | SLAIN mot | Color2 [Da | 1 | 4 |
| | 0.474836 | -0.19623 | Q8ND90 | Q8ND90 | PNMA1 | Paraneopl | Color2 [Da | 1 | 4 |
| | 0 | NaN | Q8NDA8 | Q8NDA8 | MROH1 | Maestro h | Color2 [Da | 1 | 4 |
| + | 4.792611 | 2.857024 | Q8NDC0 | Q8NDC0 | MAPK1IP1 | MAPK-inte | Color2 [Da | 1 | 4 |
| + | 1.394348 | 1.15854 | Q8NDC4 | Q8NDC4 | MORN4 | MORN rep | Color2 [Da | 1 | 4 |
| + | 2.94958 | 0.909401 | Q8NDI1 | Q8NDI1 | EHBP1 | EH domair | Color2 [Da | 1 | 4 |
| | 0.327574 | 0.117938 | Q8NDT2 | Q8NDT2 | RBM15B | Putative R | Color2 [Da | 1 | 4 |
| | 0 | -1.02623 | Q8NDV7 | Q8NDV7 | TNRC6A | Trinucleoti | Color2 [Da | 1 | 4 |
| | 0.379735 | 0.284327 | Q8NDX5 | Q8NDX5 | PHC3 | Polyhome | Color2 [Da | 1 | 4 |
| + | 1.482764 | -0.93022 | Q8NDZ4 | Q8NDZ4 | DIPK2A | Divergent | Color2 [Da | 1 | 4 |
| + | 1.818489 | 1.151088 | Q8NE01 | Q8NE01 | CNNM3 | Metal tran | Color2 [Da | 1 | 4 |
| + | 2.334255 | -0.67246 | Q8NE71 | Q8NE71 | ABCF1 | ATP-bindir | Color2 [Da | 1 | 4 |
| | 0.486174 | -0.219 | Q8NE86 | Q8NE86 | MCU | Calcium ur | Color2 [Da | 1 | 4 |
| + | 2.776949 | -1.62906 | Q8NEB9 | Q8NEB9 | PIK3C3 | Phosphati | Color2 [Da | 1 | 4 |
| | 0 | NaN | Q8NEC7 | Q8NEC7 | GSTCD | Glutathion | Color2 [Da | 1 | 4 |
| + | 1.786243 | 1.121363 | Q8NEF9 | Q8NEF9 | SRFBP1 | Serum res | Color2 [Da | 1 | 4 |
| + | 1.301869 | 0.555805 | Q8NEJ9 | Q8NEJ9 | NGDN | Neuroguid | Color2 [Da | 1 | 4 |
| + | 1.248831 | 0.327803 | Q8NEL9 | Q8NEL9 | DDHD1 | Phospholi | Color2 [Da | 1 | 4 |
| | 0.544059 | 0.466622 | Q8NEN9 | Q8NEN9 | PDZD8 | PDZ doma | Color2 [Da | 1 | 4 |
| | 0.628104 | -0.2004 | Q8NES3 | Q8NES3 | LFNG | Beta-1,3-N | Color2 [Da | 1 | 4 |
| | 0.902654 | 0.334855 | Q8NEU8 | Q8NEU8 | APPL2 | DCC-intera | Color2 [Da | 1 | 4 |
| | 1.49222 | 0.169091 | Q8NEY1 | Q8NEY1 | NAV1 | Neuron na | Color2 [Da | 1 | 4 |
| + | 2.27083 | 0.583881 | Q8NEZ2 | Q8NEZ2 | VPS37A | Vacuolar p | Color2 [Da | 1 | 4 |
| | 0 | NaN | Q8NEZ4 | Q8NEZ4 | KMT2C | Histone-ly | Color2 [Da | 1 | 4 |
| + | 1.044229 | -0.76366 | Q8NEZ5 | Q8NEZ5 | FBXO22 | F-box only | Color2 [Da | 1 | 4 |
| + | 4.472005 | -1.58041 | Q8NF37 | Q8NF37 | LPCAT1 | Lysophosp | Color2 [Da | 1 | 4 |
| | 0.707969 | -0.38777 | Q8NF91 | Q8NF91 | SYNE1 | Nesprin-1 | Color2 [Da | 1 | 4 |
| | 0.402476 | 0.666374 | Q8NFA0 | Q8NFA0 | USP32 | Ubiquitin c | Color2 [Da | 1 | 4 |
| | 0.052577 | -0.02126 | Q8NFD5 | Q8NFD5 | ARID1B | AT-rich int | Color2 [Da | 1 | 4 |
| + | 0.829881 | -0.62439 | Q8NFF5 | Q8NFF5 | FLAD1 | FAD synth | Color2 [Da | 1 | 4 |
| | 0 | -0.8731 | Q8NFG4 | Q8NFG4 | FLCN | Folliculin | Color2 [Da | 1 | 4 |
| | 0.667956 | -0.32921 | Q8NFH3 | Q8NFH3 | NUP43 | Nucleopor | Color2 [Da | 1 | 4 |
| + | 1.816985 | -0.67107 | Q8NFH4 | Q8NFH4 | NUP37 | Nucleopor | Color2 [Da | 1 | 4 |
| + | 3.037373 | 1.724433 | Q8NFH5 | Q8NFH5 | NUP35 | Nucleopor | Color2 [Da | 1 | 4 |
| + | 2.76335 | 1.231836 | Q8NFH8 | Q8NFH8 | REPS2 | RalBP1-ass | Color2 [Da | 1 | 4 |
| | 0.685173 | -0.63757 | Q8NFI3 | Q8NFI3 | ENGASE | Cytosolic e | Color2 [Da | 1 | 4 |
| | 0 | NaN | Q8NFJ9 | Q8NFJ9 | BBS1 | Bardet-Bie | Color2 [Da | 1 | 4 |
| | 0.024677 | -0.00817 | Q8NFP7 | Q8NFP7 | NUDT10 | Diphosphc | Color2 [Da | 1 | 4 |
| + | 1.823854 | -0.61873 | Q8NFP9 | Q8NFP9 | NBEA | Neurobea | Color2 [Da | 1 | 4 |
| | 0.405497 | -0.07149 | Q8NFQ8 | Q8NFQ8 | TOR1AIP2 | Torsin-1A- | Color2 [Da | 1 | 4 |
| + | 1.812005 | 0.935236 | Q8NFV4 | Q8NFV4 | ABHD11 | Protein AB | Color2 [Da | 1 | 4 |
| + | 1.959333 | -0.70748 | Q8NFW8 | Q8NFW8 | CMAS | N-acylneu | Color2 [Da | 1 | 4 |
| + | 2.07256 | 0.282919 | Q8NFZ4 | Q8NFZ4 | NLGN2 | Neuroligin | Color2 [Da | 1 | 4 |
| + | 0.701223 | 0.796295 | Q8NFZ8 | Q8NFZ8 | CADM4 | Cell adhesi | Color2 [Da | 1 | 4 |
| | 0 | NaN | Q8NG11 | Q8NG11 | TSPAN14 | Tetraspani | Color2 [Da | 1 | 4 |
| | 0.362534 | -0.13063 | Q8NG27 | Q8NG27 | PJA1 | E3 ubiquiti | Color2 [Da | 1 | 4 |
| | 0 | NaN | Q8NG68 | Q8NG68 | TTL | Tubulin--ty | Color2 [Da | 1 | 4 |

| | | | | | | | | | |
|---|---|---|---|---|---|---|---|---|---|
| + | 2.758099 | 0.98442 | Q8NHG7 | Q8NHG7 | SVIP | Small VCP/ | Color2 [Da | 1 | 4 |
| | 0.325166 | 0.653489 | Q8NHG8 | Q8NHG8 | ZNRF2 | E3 ubiquiti | Color2 [Da | 1 | 4 |
| + | 2.668898 | -0.76165 | Q8NHH9 | Q8NHH9 | ATL2 | Atlastin-2 | Color2 [Da | 1 | 4 |
| + | 0.871712 | -1.09208 | Q8NHP6 | Q8NHP6 | MOSPD2 | Motile spe | Color2 [Da | 1 | 4 |
| + | 3.486927 | 2.188698 | Q8NHP8 | Q8NHP8 | PLBD2 | Putative pl | Color2 [Da | 1 | 4 |
| | 0.254766 | 0.243658 | Q8NHV4 | Q8NHV4 | NEDD1 | Protein NE | Color2 [Da | 1 | 4 |
| + | 1.349301 | 0.648149 | Q8NHZ8 | Q8NHZ8 | CDC26 | Anaphase- | Color2 [Da | 1 | 4 |
| + | 1.77259 | 1.909687 | Q8NI08 | Q8NI08 | NCOA7 | Nuclear re | Color2 [Da | 1 | 4 |
| + | 2.504072 | -1.4486 | Q8NI22 | Q8NI22 | MCFD2 | Multiple co | Color2 [Da | 1 | 4 |
| + | 1.899718 | -0.96667 | Q8NI27 | Q8NI27 | THOC2 | THO comp | Color2 [Da | 1 | 4 |
| + | 1.973176 | -1.29823 | Q8NI36 | Q8NI36 | WDR36 | WD repeat | Color2 [Da | 1 | 4 |
| + | 0.875749 | -1.58784 | Q8TAC9 | Q8TAC9 | SCAMP5 | Secretory ( | Color2 [Da | 1 | 4 |
| | 0 | NaN | Q8TAD7 | Q8TAD7 | OCC1 | Overexpre | Color2 [Da | 1 | 4 |
| + | 1.499488 | 0.420599 | Q8TAD8 | Q8TAD8 | SNIP1 | Smad nucl | Color2 [Da | 1 | 4 |
| + | 3.305501 | 2.354418 | Q8TAE6 | Q8TAE6 | PPP1R14C | Protein ph | Color2 [Da | 1 | 4 |
| + | 2.296282 | 2.152611 | Q8TAE8 | Q8TAE8 | GADD45GI | Growth ar | Color2 [Da | 1 | 4 |
| + | 3.420945 | -1.0045 | Q8TAF3 | Q8TAF3 | WDR48 | WD repeat | Color2 [Da | 1 | 4 |
| | 0.486128 | -0.33055 | Q8TAM2 | Q8TAM2 | TTC8 | Tetratricop | Color2 [Da | 1 | 4 |
| + | 4.083198 | 0.43919 | Q8TAQ2 | Q8TAQ2 | SMARCC2 | SWI/SNF c | Color2 [Da | 1 | 4 |
| | 0.212589 | -0.21303 | Q8TAT2 | Q8TAT2 | FGFBP3 | Fibroblast | Color2 [Da | 1 | 4 |
| + | 2.666388 | -0.90129 | Q8TAT6 | Q8TAT6 | NPLOC4 | Nuclear pr | Color2 [Da | 1 | 4 |
| | 0 | -0.24461 | Q8TAV0 | Q8TAV0 | FAM76A | Protein FA | Color2 [Da | 1 | 4 |
| | 0 | NaN | Q8TB03 | Q8TB03 | CXorf38 | Uncharact | Color2 [Da | 1 | 4 |
| | 0 | -1.47251 | Q8TB22 | Q8TB22 | SPATA20 | Spermatog | Color2 [Da | 1 | 4 |
| + | 1.882264 | -0.6882 | Q8TB36 | Q8TB36 | GDAP1 | Gangliosid | Color2 [Da | 1 | 4 |
| + | 2.780482 | 1.103813 | Q8TB37 | Q8TB37 | NUBPL | Iron-sulfur | Color2 [Da | 1 | 4 |
| | 0.062419 | -0.13281 | Q8TB52 | Q8TB52 | FBXO30 | F-box only | Color2 [Da | 1 | 4 |
| | 0.637769 | -0.34525 | Q8TB72 | Q8TB72 | PUM2 | Pumilio ho | Color2 [Da | 1 | 4 |
| | 1.041319 | 0.160097 | Q8TB96 | Q8TB96 | ITFG1 | T-cell imm | Color2 [Da | 1 | 4 |
| + | 1.074548 | -0.47736 | Q8TBA6 | Q8TBA6 | GOLGA5 | Golgin sub | Color2 [Da | 1 | 4 |
| | 0.206443 | -0.1275 | Q8TBB6 | Q8TBB6 | SLC7A14 | Probable c | Color2 [Da | 1 | 4 |
| + | 4.198674 | -1.10915 | Q8TBC4 | Q8TBC4 | UBA3 | NEDD8-act | Color2 [Da | 1 | 4 |
| + | 3.509963 | 1.541991 | Q8TBC5 | Q8TBC5 | ZSCAN18 | Zinc finger | Color2 [Da | 1 | 4 |
| | 0.558534 | 0.170233 | Q8TBF2 | Q8TBF2 | PRXL2B | Prostamid | Color2 [Da | 1 | 4 |
| | 0.441193 | 0.857169 | Q8TBF4 | Q8TBF4 | ZCRB1 | Zinc finger | Color2 [Da | 1 | 4 |
| + | 0.982502 | 1.0523 | Q8TBP5 | Q8TBP5 | FAM174A | Membrane | Color2 [Da | 1 | 4 |
| | 0 | NaN | Q8TBP6 | Q8TBP6 | SLC25A40 | Probable r | Color2 [Da | 1 | 4 |
| + | 3.805114 | -1.44425 | Q8TBQ9 | Q8TBQ9 | TMEM167 | Protein kis | Color2 [Da | 1 | 4 |
| + | 2.142128 | -0.4931 | Q8TBX8 | Q8TBX8 | PIP4K2C | Phosphatic | Color2 [Da | 1 | 4 |
| + | 2.132524 | -1.13308 | Q8TC07 | Q8TC07 | TBC1D15 | TBC1 dom | Color2 [Da | 1 | 4 |
| + | 3.209815 | -0.65756 | Q8TC12 | Q8TC12 | RDH11 | Retinol del | Color2 [Da | 1 | 4 |
| + | 1.875477 | 1.040172 | Q8TC26 | Q8TC26 | TMEM163 | Transmem | Color2 [Da | 1 | 4 |
| + | 3.184631 | 2.677212 | Q8TCA0 | Q8TCA0 | LRRC20 | Leucine-ric | Color2 [Da | 1 | 4 |
| | 0.194692 | 0.256374 | Q8TCC3 | Q8TCC3 | MRPL30 | 39S riboso | Color2 [Da | 1 | 4 |
| + | 1.764702 | 0.730725 | Q8TCD5 | Q8TCD5 | NT5C | 5'(3')-deox | Color2 [Da | 1 | 4 |
| | 0.618312 | -0.26478 | Q8TCE6 | Q8TCE6 | DENND10 | DENN dom | Color2 [Da | 1 | 4 |
| | 0 | -1.9309 | Q8TCG1 | Q8TCG1 | CIP2A | Protein CIF | Color2 [Da | 1 | 4 |

| | | | | | | | | | |
|---|---|---|---|---|---|---|---|---|---|
| | 0 | NaN | Q8TCG5 | Q8TCG5 | CPT1C | Carnitine ( | Color2 [Da | 1 | 4 |
| + | 2.563351 | -1.43205 | Q8TCJ2 | Q8TCJ2 | STT3B | Dolichyl-di | Color2 [Da | 1 | 4 |
| + | 1.607872 | -0.56535 | Q8TCS8 | Q8TCS8 | PNPT1 | Polyribonu | Color2 [Da | 1 | 4 |
| | 0.335597 | 0.873813 | Q8TCT7 | Q8TCT7 | SPPL2B | Signal pep | Color2 [Da | 1 | 4 |
| + | 2.201902 | -1.75209 | Q8TCT9 | Q8TCT9 | HM13 | Minor hist | Color2 [Da | 1 | 4 |
| + | 2.307171 | -1.2943 | Q8TCU4 | Q8TCU4 | ALMS1 | Alstrom sy | Color2 [Da | 1 | 4 |
| + | 3.469904 | -1.51161 | Q8TCU6 | Q8TCU6 | PREX1 | Phosphatic | Color2 [Da | 1 | 4 |
| | 0 | -0.85715 | Q8TCX1 | Q8TCX1 | DYNC2LI1 | Cytoplasm | Color2 [Da | 1 | 4 |
| + | 2.520541 | 2.049915 | Q8TCZ2 | Q8TCZ2 | CD99L2 | CD99 antig | Color2 [Da | 1 | 4 |
| | 0.725411 | 0.476238 | Q8TD16 | Q8TD16 | BICD2 | Protein bic | Color2 [Da | 1 | 4 |
| + | 2.407331 | -1.49119 | Q8TD19 | Q8TD19 | NEK9 | Serine/thr | Color2 [Da | 1 | 4 |
| + | 1.063575 | -1.33076 | Q8TD22 | Q8TD22 | SFXN5 | Sideroflexi | Color2 [Da | 1 | 4 |
| | 0.032128 | -0.02178 | Q8TD30 | Q8TD30 | GPT2 | Alanine an | Color2 [Da | 1 | 4 |
| | 0 | NaN | Q8TD43 | Q8TD43 | TRPM4 | Transient r | Color2 [Da | 1 | 4 |
| + | 1.917663 | 1.372504 | Q8TD55 | Q8TD55 | PLEKHO2 | Pleckstrin | Color2 [Da | 1 | 4 |
| + | 3.177749 | 1.381134 | Q8TDC3 | Q8TDC3 | BRSK1 | Serine/thr | Color2 [Da | 1 | 4 |
| | 0.14015 | -0.05667 | Q8TDD1 | Q8TDD1 | DDX54 | ATP-deper | Color2 [Da | 1 | 4 |
| | 0.318387 | -0.18342 | Q8TDH9 | Q8TDH9 | BLOC1S5 | Biogenesis | Color2 [Da | 1 | 4 |
| | 0.120978 | -0.33271 | Q8TDI0 | Q8TDI0 | CHD5 | Chromodo | Color2 [Da | 1 | 4 |
| + | 2.822867 | 0.722114 | Q8TDJ6 | Q8TDJ6 | DMXL2 | DmX-like p | Color2 [Da | 1 | 4 |
| | 0 | -2.02569 | Q8TDL5 | Q8TDL5 | BPIFB1 | BPI fold-cc | Color2 [Da | 1 | 4 |
| | 0.437925 | 0.123428 | Q8TDM6 | Q8TDM6 | DLG5 | Disks large | Color2 [Da | 1 | 4 |
| + | 1.13129 | -0.71527 | Q8TDN6 | Q8TDN6 | BRIX1 | Ribosome | Color2 [Da | 1 | 4 |
| + | 3.281727 | 2.224623 | Q8TDQ7 | Q8TDQ7 | GNPDA2 | Glucosami | Color2 [Da | 1 | 4 |
| + | 2.339666 | 1.055734 | Q8TDR0 | Q8TDR0 | TRAF3IP1 | TRAF3-inte | Color2 [Da | 1 | 4 |
| | 0.808347 | 0.212126 | Q8TDW7 | Q8TDW7 | FAT3 | Protocadh | Color2 [Da | 1 | 4 |
| + | 2.840194 | -1.29307 | Q8TDX7 | Q8TDX7 | NEK7 | Serine/thr | Color2 [Da | 1 | 4 |
| | 0.072337 | -0.06875 | Q8TDY2 | Q8TDY2 | RB1CC1 | RB1-induci | Color2 [Da | 1 | 4 |
| | 0 | -1.98119 | Q8TDY4 | Q8TDY4 | ASAP3 | Arf-GAP w | Color2 [Da | 1 | 4 |
| + | 1.15403 | -0.39355 | Q8TDZ2 | Q8TDZ2 | MICAL1 | [F-actin]-m | Color2 [Da | 1 | 4 |
| | 0 | -1.56101 | Q8TE04 | Q8TE04 | PANK1 | Pantothen | Color2 [Da | 1 | 4 |
| | 0 | NaN | Q8TE58 | Q8TE58 | ADAMTS15 | A disintegr | Color2 [Da | 1 | 4 |
| | 0 | NaN | Q8TE99 | Q8TE99 | PXYLP1 | 2-phospho | Color2 [Da | 1 | 4 |
| | 0 | -0.11623 | Q8TEA7 | Q8TEA7 | TBCK | TBC domai | Color2 [Da | 1 | 4 |
| + | 3.960416 | 1.694578 | Q8TEA8 | Q8TEA8 | DTD1 | D-aminoac | Color2 [Da | 1 | 4 |
| | 0.235145 | 0.246806 | Q8TEB1 | Q8TEB1 | DCAF11 | DDB1- and | Color2 [Da | 1 | 4 |
| + | 1.738529 | 0.626329 | Q8TED0 | Q8TED0 | UTP15 | U3 small n | Color2 [Da | 1 | 4 |
| + | 2.271043 | -0.75768 | Q8TED1 | Q8TED1 | GPX8 | Probable g | Color2 [Da | 1 | 4 |
| + | 3.003561 | -1.14103 | Q8TEM1 | Q8TEM1 | NUP210 | Nuclear pc | Color2 [Da | 1 | 4 |
| | 0 | NaN | Q8TEQ0 | Q8TEQ0 | SNX29 | Sorting ne | Color2 [Da | 1 | 4 |
| + | 1.530943 | -1.11913 | Q8TEQ6 | Q8TEQ6 | GEMIN5 | Gem-assoc | Color2 [Da | 1 | 4 |
| + | 1.040863 | -0.4835 | Q8TEQ8 | Q8TEQ8 | PIGO | GPI ethanc | Color2 [Da | 1 | 4 |
| + | 0.883175 | 0.387786 | Q8TEW0 | Q8TEW0 | PARD3 | Partitionin | Color2 [Da | 1 | 4 |
| | 0.001981 | -0.0051 | Q8TEW8 | Q8TEW8 | PARD3B | Partitionin | Color2 [Da | 1 | 4 |
| + | 2.110022 | -1.98224 | Q8TEX9 | Q8TEX9 | IPO4 | Importin-4 | Color2 [Da | 1 | 4 |
| + | 2.184435 | 0.779532 | Q8TF01 | Q8TF01 | PNISR | Arginine/s | Color2 [Da | 1 | 4 |
| + | 0.807983 | 0.599044 | Q8TF05 | Q8TF05 | PPP4R1 | Serine/thr | Color2 [Da | 1 | 4 |

| | | | | | | | | | |
|---|---|---|---|---|---|---|---|---|---|
| | 0.073223 | 0.111308 | Q8TF42 | Q8TF42 | UBASH3B | Ubiquitin-a | Color2 [Da | 1 | 4 |
| + | 2.583262 | 0.935024 | Q8TF44 | Q8TF44 | C2CD4C | C2 calcium | Color2 [Da | 1 | 4 |
| + | 3.282692 | 1.231991 | Q8TF61 | Q8TF61 | FBXO41 | F-box only | Color2 [Da | 1 | 4 |
| | 0 | 0.773647 | Q8TF68 | Q8TF68 | ZNF384 | Zinc finger | Color2 [Da | 1 | 4 |
| | 0.239 | 1.471622 | Q8TF72 | Q8TF72 | SHROOM3 | Protein Sh | Color2 [Da | 1 | 4 |
| | 0.573211 | 1.05548 | Q8TF74 | Q8TF74 | WIPF2 | WAS/WAS | Color2 [Da | 1 | 4 |
| | 0.099779 | 0.437708 | Q8WTR2 | Q8WTR2 | DUSP19 | Dual speci | Color2 [Da | 1 | 4 |
| | 0.094225 | 0.045139 | Q8WTS6 | Q8WTS6 | SETD7 | Histone-ly: | Color2 [Da | 1 | 4 |
| + | 2.024868 | -1.20077 | Q8WTT2 | Q8WTT2 | NOC3L | Nucleolar ( | Color2 [Da | 1 | 4 |
| + | 2.619344 | -1.08514 | Q8WTW3 | Q8WTW3 | COG1 | Conserved | Color2 [Da | 1 | 4 |
| + | 2.780263 | 0.615285 | Q8WU20 | Q8WU20 | FRS2 | Fibroblast | Color2 [Da | 1 | 4 |
| + | 1.389681 | -0.61646 | Q8WU76 | Q8WU76 | SCFD2 | Sec1 famil | Color2 [Da | 1 | 4 |
| | 0.878139 | 0.232605 | Q8WU79 | Q8WU79 | SMAP2 | Stromal m | Color2 [Da | 1 | 4 |
| | 0.075463 | 0.018263 | Q8WU90 | Q8WU90 | ZC3H15 | Zinc finger | Color2 [Da | 1 | 4 |
| + | 1.042752 | -0.51515 | Q8WUA2 | Q8WUA2 | PPIL4 | Peptidyl-p | Color2 [Da | 1 | 4 |
| + | 1.250404 | 0.535512 | Q8WUA4 | Q8WUA4 | GTF3C2 | General tr | Color2 [Da | 1 | 4 |
| + | 1.613621 | -1.26297 | Q8WUA8 | Q8WUA8 | TSKU | Tsukushi | Color2 [Da | 1 | 4 |
| + | 2.412965 | 1.201361 | Q8WUB8 | Q8WUB8 | PHF10 | PHD finger | Color2 [Da | 1 | 4 |
| + | 1.662534 | -0.88953 | Q8WUD1 | Q8WUD1 | RAB2B | Ras-relate | Color2 [Da | 1 | 4 |
| | 0 | 1.276163 | Q8WUD4 | Q8WUD4 | CCDC12 | Coiled-coil | Color2 [Da | 1 | 4 |
| + | 4.098835 | 0.625224 | Q8WUF8 | Q8WUF8 | FAM172A | Cotranscri | Color2 [Da | 1 | 4 |
| | 0 | NaN | Q8WUG5 | Q8WUG5 | SLC22A17 | Solute carr | Color2 [Da | 1 | 4 |
| | 0 | 0.517347 | Q8WUH2 | Q8WUH2 | TGFBRAP1 | Transform | Color2 [Da | 1 | 4 |
| + | 2.830899 | 1.244505 | Q8WUH6 | Q8WUH6 | TMEM263 | Transmem | Color2 [Da | 1 | 4 |
| + | 2.913169 | 1.200147 | Q8WUK0 | Q8WUK0 | PTPMT1 | Phosphati | Color2 [Da | 1 | 4 |
| + | 2.210945 | -1.00986 | Q8WUM0 | Q8WUM0 | NUP133 | Nuclear pc | Color2 [Da | 1 | 4 |
| + | 2.89599 | -1.01073 | Q8WUM4 | Q8WUM4 | PDCD6IP | Programm | Color2 [Da | 1 | 4 |
| + | 1.52705 | 0.456889 | Q8WUN7 | Q8WUN7 | UBTD2 | Ubiquitin c | Color2 [Da | 1 | 4 |
| | 0.904977 | 0.146459 | Q8WUQ7 | Q8WUQ7 | CACTIN | Cactin | Color2 [Da | 1 | 4 |
| + | 1.728764 | -0.70365 | Q8WUU5 | Q8WUU5 | GATAD1 | GATA zinc | Color2 [Da | 1 | 4 |
| + | 3.281864 | 1.394557 | Q8WUW1 | Q8WUW1 | BRK1 | Protein BR | Color2 [Da | 1 | 4 |
| + | 2.541561 | 0.785477 | Q8WUX9 | Q8WUX9 | CHMP7 | Charged m | Color2 [Da | 1 | 4 |
| | 0 | NaN | Q8WUY3 | Q8WUY3 | PRUNE2 | Protein pr | Color2 [Da | 1 | 4 |
| | 0.10757 | 0.320496 | Q8WUY8 | Q8WUY8 | NAT14 | Probable N | Color2 [Da | 1 | 4 |
| + | 2.651497 | 2.233215 | Q8WUZ0 | Q8WUZ0 | BCL7C | B-cell CLL/ | Color2 [Da | 1 | 4 |
| | 0 | NaN | Q8WV22 | Q8WV22 | NSMCE1 | Non-struct | Color2 [Da | 1 | 4 |
| | 0 | NaN | Q8WV74 | Q8WV74 | NUDT8 | Mitochonc | Color2 [Da | 1 | 4 |
| + | 2.475001 | 0.828573 | Q8WVC0 | Q8WVC0 | LEO1 | RNA polym | Color2 [Da | 1 | 4 |
| + | 2.190355 | -0.73927 | Q8WVC6 | Q8WVC6 | DCAKD | Dephosph | Color2 [Da | 1 | 4 |
| | 0.276521 | -0.08588 | Q8WVF1 | Q8WVF1 | OSCP1 | Protein OS | Color2 [Da | 1 | 4 |
| | 0.330277 | 0.236265 | Q8WVI0 | Q8WVI0 | SMIM4 | Small integ | Color2 [Da | 1 | 4 |
| + | 2.449527 | 0.78886 | Q8WVJ2 | Q8WVJ2 | NUDCD2 | NudC dom | Color2 [Da | 1 | 4 |
| | 0.697465 | 0.637465 | Q8WVK2 | Q8WVK2 | SNRNP27 | U4/U6.U5 | Color2 [Da | 1 | 4 |
| | 0.059349 | -0.04499 | Q8WVK7 | Q8WVK7 | SKA2 | Spindle an | Color2 [Da | 1 | 4 |
| + | 1.444139 | -1.54308 | Q8WVM0 | Q8WVM0 | TFB1M | Dimethyla | Color2 [Da | 1 | 4 |
| | 0.269879 | -0.02701 | Q8WVM7 | Q8WVM7 | STAG1 | Cohesin su | Color2 [Da | 1 | 4 |
| + | 3.259304 | -0.7004 | Q8WVM8 | Q8WVM8 | SCFD1 | Sec1 famil | Color2 [Da | 1 | 4 |

| | | | | | | | | | |
|---|---|---|---|---|---|---|---|---|---|
| + | 0.908274 | -0.52273 | Q8WVQ1 | Q8WVQ1 | CANT1 | Soluble cal | Color2 | [Da | 1 | 4 |
| | 0 | -0.60505 | Q8WVT3 | Q8WVT3 | TRAPPC12 | Trafficking | Color2 | [Da | 1 | 4 |
| | 0 | -0.22554 | Q8WVV4 | Q8WVV4 | POF1B | Protein PC | Color2 | [Da | 1 | 4 |
| + | 2.934625 | -1.24031 | Q8WVV9 | Q8WVV9 | HNRNPLL | Heterogen | Color2 | [Da | 1 | 4 |
| | 0.543554 | 0.164917 | Q8WVY7 | Q8WVY7 | UBLCP1 | Ubiquitin-l | Color2 | [Da | 1 | 4 |
| + | 3.17374 | 1.516872 | Q8WW12 | Q8WW12 | PCNP | PEST prote | Color2 | [Da | 1 | 4 |
| + | 2.266243 | -1.0703 | Q8WW22 | Q8WW22 | DNAJA4 | DnaJ homo | Color2 | [Da | 1 | 4 |
| | 0 | -1.36796 | Q8WW35 | Q8WW35 | DYNLT2B | Dynein ligh | Color2 | [Da | 1 | 4 |
| + | 2.845057 | 1.048998 | Q8WW59 | Q8WW59 | SPRYD4 | SPRY doma | Color2 | [Da | 1 | 4 |
| | 0.320084 | -0.07854 | Q8WWC4 | Q8WWC4 | MAIP1 | m-AAA pro | Color2 | [Da | 1 | 4 |
| + | 2.754022 | 0.594196 | Q8WWH5 | Q8WWH5 | TRUB1 | Pseudouri | Color2 | [Da | 1 | 4 |
| | 0 | NaN | Q8WWI5 | Q8WWI5 | SLC44A1 | Choline tra | Color2 | [Da | 1 | 4 |
| + | 3.29484 | -1.59684 | Q8WWK9 | Q8WWK9 | CKAP2 | Cytoskelet | Color2 | [Da | 1 | 4 |
| + | 1.849678 | 0.51458 | Q8WWM7 | Q8WWM7 | ATXN2L | Ataxin-2-li | Color2 | [Da | 1 | 4 |
| | 0 | 2.692512 | Q8WWM9 | Q8WWM9 | CYGB | Cytoglobin | Color2 | [Da | 1 | 4 |
| + | 3.120233 | -0.36321 | Q8WWQ0 | Q8WWQ0 | PHIP | PH-interac | Color2 | [Da | 1 | 4 |
| | 0 | NaN | Q8WWW8 | Q8WWW8 | GAB3 | GRB2-asso | Color2 | [Da | 1 | 4 |
| | 0.485321 | 0.091856 | Q8WWX9 | Q8WWX9 | SELENOM | Selenopro | Color2 | [Da | 1 | 4 |
| + | 1.596941 | -0.5223 | Q8WWY3 | Q8WWY3 | PRPF31 | U4/U6 sma | Color2 | [Da | 1 | 4 |
| | 0 | -1.0248 | Q8WX77 | Q8WX77 | IGFBPL1 | Insulin-like | Color2 | [Da | 1 | 4 |
| + | 1.578958 | -1.11269 | Q8WX92 | Q8WX92 | NELFB | Negative e | Color2 | [Da | 1 | 4 |
| | 0.677244 | -0.35703 | Q8WX93 | Q8WX93 | PALLD | Palladin | Color2 | [Da | 1 | 4 |
| + | 2.398338 | -0.23297 | Q8WX94 | Q8WX94 | NLRP7 | NACHT, LR | Color2 | [Da | 1 | 4 |
| | 0.091414 | 0.043375 | Q8WXA3 | Q8WXA3 | RUFY2 | RUN and F | Color2 | [Da | 1 | 4 |
| + | 1.13105 | 0.722689 | Q8WXA9 | Q8WXA9 | SREK1 | Splicing re | Color2 | [Da | 1 | 4 |
| | 0 | NaN | Q8WXB1 | Q8WXB1 | METTL21A | Protein N- | Color2 | [Da | 1 | 4 |
| + | 1.231747 | -1.29043 | Q8WXC6 | Q8WXC6 | COPS9 | COP9 signa | Color2 | [Da | 1 | 4 |
| + | 3.56841 | 0.902296 | Q8WXD2 | Q8WXD2 | SCG3 | Secretogra | Color2 | [Da | 1 | 4 |
| | 0 | 0.317591 | Q8WXD5 | Q8WXD5 | GEMIN6 | Gem-assoc | Color2 | [Da | 1 | 4 |
| + | 2.413051 | 1.182691 | Q8WXD9 | Q8WXD9 | CASKIN1 | Caskin-1 | Color2 | [Da | 1 | 4 |
| | 0.672492 | -0.63485 | Q8WXE9 | Q8WXE9 | STON2 | Stonin-2 | Color2 | [Da | 1 | 4 |
| | 0.642311 | 0.318997 | Q8WXF0 | Q8WXF0 | SRSF12 | Serine/arg | Color2 | [Da | 1 | 4 |
| + | 1.906626 | 0.673943 | Q8WXF1 | Q8WXF1 | PSPC1 | Paraspeckl | Color2 | [Da | 1 | 4 |
| + | 1.360176 | -0.47356 | Q8WXF7 | Q8WXF7 | ATL1 | Atlastin-1 | Color2 | [Da | 1 | 4 |
| + | 1.075282 | 0.335226 | Q8WXG6 | Q8WXG6 | MADD | MAP kinas | Color2 | [Da | 1 | 4 |
| + | 2.795236 | -0.44071 | Q8WXH0 | Q8WXH0 | SYNE2 | Nesprin-2 | Color2 | [Da | 1 | 4 |
| | 0.455051 | -0.28508 | Q8WXI2 | Q8WXI2 | CNKSR2 | Connector | Color2 | [Da | 1 | 4 |
| | 0.076393 | 0.040904 | Q8WXI9 | Q8WXI9 | GATAD2B | Transcripti | Color2 | [Da | 1 | 4 |
| | 0.700465 | -0.23086 | Q8WXX5 | Q8WXX5 | DNAJC9 | DnaJ homo | Color2 | [Da | 1 | 4 |
| | 0 | NaN | Q8WY21 | Q8WY21 | SORCS1 | VPS10 don | Color2 | [Da | 1 | 4 |
| + | 3.655059 | -2.66181 | Q8WY22 | Q8WY22 | BRI3BP | BRI3-bindi | Color2 | [Da | 1 | 4 |
| | 0.800008 | -0.34798 | Q8WY54 | Q8WY54 | PPM1E | Protein ph | Color2 | [Da | 1 | 4 |
| + | 1.30329 | 0.466865 | Q8WYA0 | Q8WYA0 | IFT81 | Intraflagell | Color2 | [Da | 1 | 4 |
| + | 0.981333 | -0.46934 | Q8WYA6 | Q8WYA6 | CTNNBL1 | Beta-cater | Color2 | [Da | 1 | 4 |
| | 0.294762 | 0.283283 | Q8WYB5 | Q8WYB5 | KAT6B | Histone ac | Color2 | [Da | 1 | 4 |
| | 0.738048 | -0.27378 | Q8WYH8 | Q8WYH8 | ING5 | Inhibitor o | Color2 | [Da | 1 | 4 |
| | 0.913709 | 0.158902 | Q8WYP5 | Q8WYP5 | AHCTF1 | Protein EL' | Color2 | [Da | 1 | 4 |

| | | | | | | | | | |
|---|---|---|---|---|---|---|---|---|---|
| | 0 | -0.16225 | Q8WYQ5 | Q8WYQ5 | DGCR8 | Microproc | Color2 [Da | 1 | 4 |
| + | 1.28411 | 1.026891 | Q8WYR4 | Q8WYR4 | RSPH1 | Radial spo | Color2 [Da | 1 | 4 |
| | 1.138109 | -0.11718 | Q8WZ42 | Q8WZ42 | TTN | Titin | Color2 [Da | 1 | 4 |
| + | 1.911921 | 0.945846 | Q8WZ74 | Q8WZ74 | CTTNBP2 | Cortactin-I | Color2 [Da | 1 | 4 |
| | 0 | -1.56443 | Q8WZ82 | Q8WZ82 | OVCA2 | Esterase O | Color2 [Da | 1 | 4 |
| + | 2.883345 | 1.978909 | Q8WZA0 | Q8WZA0 | LZIC | Protein LZI | Color2 [Da | 1 | 4 |
| + | 1.440934 | 0.435466 | Q8WZA9 | Q8WZA9 | IRGQ | Immunity- | Color2 [Da | 1 | 4 |
| + | 2.353141 | 1.189786 | Q92186 | Q92186 | ST8SIA2 | Alpha-2,8- | Color2 [Da | 1 | 4 |
| | 0.279993 | 0.142344 | Q92187 | Q92187 | ST8SIA4 | CMP-N-ace | Color2 [Da | 1 | 4 |
| + | 3.799202 | -0.70256 | Q92499 | Q92499 | DDX1 | ATP-deper | Color2 [Da | 1 | 4 |
| | 0 | -2.44741 | Q92504 | Q92504 | SLC39A7 | Zinc transp | Color2 [Da | 1 | 4 |
| + | 1.815436 | 0.732553 | Q92506 | Q92506 | HSD17B8 | (3R)-3-hyd | Color2 [Da | 1 | 4 |
| + | 2.412525 | -0.39269 | Q92520 | Q92520 | FAM3C | Protein FA | Color2 [Da | 1 | 4 |
| + | 3.039863 | 1.920978 | Q92522 | Q92522 | H1-10 | Histone H1 | Color2 [Da | 1 | 4 |
| + | 3.002465 | 1.372218 | Q92530 | Q92530 | PSMF1 | Proteasom | Color2 [Da | 1 | 4 |
| | 0 | NaN | Q92536 | Q92536 | SLC7A6 | Y+L amino | Color2 [Da | 1 | 4 |
| + | 1.957771 | -0.82288 | Q92538 | Q92538 | GBF1 | Golgi-spec | Color2 [Da | 1 | 4 |
| + | 1.316282 | 0.821021 | Q92539 | Q92539 | LPIN2 | Phosphatid | Color2 [Da | 1 | 4 |
| | 0.531519 | -0.09535 | Q92540 | Q92540 | SMG7 | Nonsense- | Color2 [Da | 1 | 4 |
| + | 2.038693 | 0.593796 | Q92541 | Q92541 | RTF1 | RNA polym | Color2 [Da | 1 | 4 |
| + | 3.989516 | 0.787751 | Q92542 | Q92542 | NCSTN | Nicastrin | Color2 [Da | 1 | 4 |
| + | 1.116467 | -1.04739 | Q92544 | Q92544 | TM9SF4 | Transmem | Color2 [Da | 1 | 4 |
| | 0 | NaN | Q92545 | Q92545 | TMEM131 | Transmem | Color2 [Da | 1 | 4 |
| | 0 | NaN | Q92547 | Q92547 | TOPBP1 | DNA topoi | Color2 [Da | 1 | 4 |
| | 0 | 0.136904 | Q92551 | Q92551 | IP6K1 | Inositol he | Color2 [Da | 1 | 4 |
| + | 2.876002 | -0.56733 | Q92552 | Q92552 | MRPS27 | 28S riboso | Color2 [Da | 1 | 4 |
| | 0.118993 | -0.04708 | Q92556 | Q92556 | ELMO1 | Engulfmen | Color2 [Da | 1 | 4 |
| + | 1.972962 | 0.684777 | Q92558 | Q92558 | WASF1 | Wiskott-Al | Color2 [Da | 1 | 4 |
| | 0.007864 | -0.01346 | Q92560 | Q92560 | BAP1 | Ubiquitin c | Color2 [Da | 1 | 4 |
| + | 2.254807 | -0.5614 | Q92562 | Q92562 | FIG4 | Polyphosp | Color2 [Da | 1 | 4 |
| + | 1.793169 | 0.756599 | Q92563 | Q92563 | SPOCK2 | Testican-2 | Color2 [Da | 1 | 4 |
| + | 1.90363 | 2.48103 | Q92567 | Q92567 | FAM168A | Protein FA | Color2 [Da | 1 | 4 |
| + | 0.730859 | -0.96248 | Q92572 | Q92572 | AP3S1 | AP-3 comp | Color2 [Da | 1 | 4 |
| | 2.588858 | 0.109722 | Q92574 | Q92574 | TSC1 | Hamartin | Color2 [Da | 1 | 4 |
| + | 3.273282 | 1.905289 | Q92575 | Q92575 | UBXN4 | UBX doma | Color2 [Da | 1 | 4 |
| + | 2.993214 | -1.29101 | Q92576 | Q92576 | PHF3 | PHD finger | Color2 [Da | 1 | 4 |
| | 0.904565 | -0.24616 | Q92581 | Q92581 | SLC9A6 | Sodium/hy | Color2 [Da | 1 | 4 |
| + | 1.002163 | -0.35085 | Q92597 | Q92597 | NDRG1 | Protein ND | Color2 [Da | 1 | 4 |
| + | 3.319255 | -0.54937 | Q92598 | Q92598 | HSPH1 | Heat shock | Color2 [Da | 1 | 4 |
| | 0.57421 | -0.21165 | Q92599 | Q92599 | SEPTIN8 | Septin-8 | Color2 [Da | 1 | 4 |
| | 0.670942 | -0.44908 | Q92600 | Q92600 | CNOT9 | CCR4-NOT | Color2 [Da | 1 | 4 |
| | 0 | NaN | Q92604 | Q92604 | LPGAT1 | Acyl-CoA:l | Color2 [Da | 1 | 4 |
| | 0.732701 | 0.331023 | Q92609 | Q92609 | TBC1D5 | TBC1 doma | Color2 [Da | 1 | 4 |
| + | 2.452439 | -0.59018 | Q92614 | Q92614 | MYO18A | Unconvent | Color2 [Da | 1 | 4 |
| + | 2.611581 | 0.714417 | Q92615 | Q92615 | LARP4B | La-related | Color2 [Da | 1 | 4 |
| + | 3.241934 | -1.17066 | Q92616 | Q92616 | GCN1 | eIF-2-alpha | Color2 [Da | 1 | 4 |
| + | 2.417944 | -0.96978 | Q92620 | Q92620 | DHX38 | Pre-mRNA | Color2 [Da | 1 | 4 |

| | | | | | | | | | | |
|---|---|---|---|---|---|---|---|---|---|---|
| + | 3.040702 | -0.8245 | Q92621 | Q92621 | NUP205 | Nuclear pc | Color2 | [Da | 1 | 4 |
| + | 0.810703 | 0.914847 | Q92623 | Q92623 | TTC9 | Tetratricop | Color2 | [Da | 1 | 4 |
| | 1.309922 | -0.17425 | Q92625 | Q92625 | ANKS1A | Ankyrin re | Color2 | [Da | 1 | 4 |
| + | 3.268267 | -2.1478 | Q92626 | Q92626 | PXDN | Peroxidasi | Color2 | [Da | 1 | 4 |
| + | 2.043314 | -1.10935 | Q92643 | Q92643 | PIGK | GPI-ancho | Color2 | [Da | 1 | 4 |
| + | 3.165945 | 1.321778 | Q92665 | Q92665 | MRPS31 | 28S riboso | Color2 | [Da | 1 | 4 |
| + | 3.2419 | 1.103918 | Q92667 | Q92667 | AKAP1 | A-kinase a | Color2 | [Da | 1 | 4 |
| | 0 | NaN | Q92685 | Q92685 | ALG3 | Dol-P-Man | Color2 | [Da | 1 | 4 |
| + | 1.195772 | -0.54965 | Q92688 | Q92688 | ANP32B | Acidic leuc | Color2 | [Da | 1 | 4 |
| + | 1.334893 | -0.25806 | Q92692 | Q92692 | NECTIN2 | Nectin-2 | Color2 | [Da | 1 | 4 |
| + | 0.985869 | -0.50347 | Q92696 | Q92696 | RABGGTA | Geranylger | Color2 | [Da | 1 | 4 |
| | 0.358445 | -0.2507 | Q92733 | Q92733 | PRCC | Proline-ric | Color2 | [Da | 1 | 4 |
| + | 2.802025 | 1.876852 | Q92734 | Q92734 | TFG | Protein TF | Color2 | [Da | 1 | 4 |
| | 0 | -0.01989 | Q92738 | Q92738 | USP6NL | USP6 N-ter | Color2 | [Da | 1 | 4 |
| + | 3.578666 | 0.883904 | Q92743 | Q92743 | HTRA1 | Serine pro | Color2 | [Da | 1 | 4 |
| + | 1.673001 | -0.93023 | Q92747 | Q92747 | ARPC1A | Actin-relat | Color2 | [Da | 1 | 4 |
| | 0.02006 | 0.023733 | Q92752 | Q92752 | TNR | Tenascin-R | Color2 | [Da | 1 | 4 |
| + | 3.522445 | -1.43943 | Q92769 | Q92769 | HDAC2 | Histone de | Color2 | [Da | 1 | 4 |
| + | 1.814496 | 1.494506 | Q92777 | Q92777 | SYN2 | Synapsin-2 | Color2 | [Da | 1 | 4 |
| | 0.201929 | 0.31266 | Q92782 | Q92782 | DPF1 | Zinc finger | Color2 | [Da | 1 | 4 |
| + | 3.865111 | 1.713044 | Q92783 | Q92783 | STAM | Signal tran | Color2 | [Da | 1 | 4 |
| + | 0.792526 | 0.511882 | Q92784 | Q92784 | DPF3 | Zinc finger | Color2 | [Da | 1 | 4 |
| + | 1.626944 | -0.3647 | Q92785 | Q92785 | DPF2 | Zinc finger | Color2 | [Da | 1 | 4 |
| + | 1.180911 | 0.285009 | Q92796 | Q92796 | DLG3 | Disks large | Color2 | [Da | 1 | 4 |
| + | 2.17124 | -0.99177 | Q92797 | Q92797 | SYMPK | Symplekin | Color2 | [Da | 1 | 4 |
| + | 2.813505 | 1.691284 | Q92804 | Q92804 | TAF15 | TATA-bind | Color2 | [Da | 1 | 4 |
| | 0.4187 | -0.1591 | Q92805 | Q92805 | GOLGA1 | Golgin sub | Color2 | [Da | 1 | 4 |
| + | 4.018693 | 2.231729 | Q92820 | Q92820 | GGH | Gamma-gl | Color2 | [Da | 1 | 4 |
| | 0.864184 | 0.357901 | Q92823 | Q92823 | NRCAM | Neuronal c | Color2 | [Da | 1 | 4 |
| + | 2.024027 | -0.98518 | Q92824 | Q92824 | PCSK5 | Proproteir | Color2 | [Da | 1 | 4 |
| | 0 | NaN | Q92828 | Q92828 | CORO2A | Coronin-2A | Color2 | [Da | 1 | 4 |
| + | 2.673073 | -0.85827 | Q92841 | Q92841 | DDX17 | Probable A | Color2 | [Da | 1 | 4 |
| + | 1.795567 | -0.61691 | Q92845 | Q92845 | KIFAP3 | Kinesin-ass | Color2 | [Da | 1 | 4 |
| | 0.192622 | -0.0296 | Q92859 | Q92859 | NEO1 | Neogenin | Color2 | [Da | 1 | 4 |
| + | 2.374091 | -1.11775 | Q92878 | Q92878 | RAD50 | DNA repai | Color2 | [Da | 1 | 4 |
| | 0.679826 | -0.19201 | Q92879 | Q92879 | CELF1 | CUGBP Ela | Color2 | [Da | 1 | 4 |
| + | 3.367515 | 1.647396 | Q92882 | Q92882 | OSTF1 | Osteoclast | Color2 | [Da | 1 | 4 |
| | 0.719908 | -0.27329 | Q92888 | Q92888 | ARHGEF1 | Rho guanir | Color2 | [Da | 1 | 4 |
| | 0 | -1.85097 | Q92889 | Q92889 | ERCC4 | DNA repai | Color2 | [Da | 1 | 4 |
| + | 1.911838 | 1.053961 | Q92890 | Q92890 | UFD1 | Ubiquitin r | Color2 | [Da | 1 | 4 |
| + | 1.709574 | -1.23157 | Q92896 | Q92896 | GLG1 | Golgi appa | Color2 | [Da | 1 | 4 |
| + | 3.99884 | -1.199 | Q92900 | Q92900 | UPF1 | Regulator | Color2 | [Da | 1 | 4 |
| | 0.929749 | -0.22898 | Q92905 | Q92905 | COPS5 | COP9 signa | Color2 | [Da | 1 | 4 |
| + | 1.675641 | 1.433118 | Q92913 | Q92913 | FGF13 | Fibroblast | Color2 | [Da | 1 | 4 |
| + | 1.380878 | 1.142562 | Q92917 | Q92917 | GPKOW | G-patch dc | Color2 | [Da | 1 | 4 |
| | 0.090536 | -0.0566 | Q92922 | Q92922 | SMARCC1 | SWI/SNF c | Color2 | [Da | 1 | 4 |
| + | 1.411528 | -0.90231 | Q92925 | Q92925 | SMARCD2 | SWI/SNF-r | Color2 | [Da | 1 | 4 |

| | | | | | | | | | |
|---|---|---|---|---|---|---|---|---|---|
| | 0.082321 | 0.046613 | Q92930 | Q92930 | RAB8B | Ras-relate | Color2 [Da | 1 | 4 |
| + | 2.419037 | 0.85 | Q92932 | Q92932 | PTPRN2 | Receptor-t | Color2 [Da | 1 | 4 |
| + | 4.140982 | 2.407287 | Q92934 | Q92934 | BAD | Bcl2-assoc | Color2 [Da | 1 | 4 |
| + | 2.039647 | 0.948243 | Q92945 | Q92945 | KHSRP | Far upstre | Color2 [Da | 1 | 4 |
| + | 1.57116 | -0.37535 | Q92947 | Q92947 | GCDH | Glutaryl-C | Color2 [Da | 1 | 4 |
| + | 1.486594 | -0.61203 | Q92963 | Q92963 | RIT1 | GTP-bindir | Color2 [Da | 1 | 4 |
| + | 3.621221 | -1.77233 | Q92973 | Q92973 | TNPO1 | Transporti | Color2 [Da | 1 | 4 |
| + | 2.416068 | -0.63322 | Q92974 | Q92974 | ARHGEF2 | Rho guanir | Color2 [Da | 1 | 4 |
| | 0 | 0.442778 | Q92979 | Q92979 | EMG1 | Ribosomal | Color2 [Da | 1 | 4 |
| + | 1.606065 | -0.72688 | Q92990 | Q92990 | GLMN | Glomulin | Color2 [Da | 1 | 4 |
| | 0 | NaN | Q92993 | Q92993 | KAT5 | Histone ac | Color2 [Da | 1 | 4 |
| + | 0.810764 | 0.58467 | Q92997 | Q92997 | DVL3 | Segment p | Color2 [Da | 1 | 4 |
| + | 4.636131 | -1.46906 | Q93008 | Q93008 | USP9X | Probable u | Color2 [Da | 1 | 4 |
| + | 2.750843 | -1.30799 | Q93009 | Q93009 | USP7 | Ubiquitin c | Color2 [Da | 1 | 4 |
| + | 0.74445 | 0.54545 | Q93015 | Q93015 | NAA80 | N-alpha-ac | Color2 [Da | 1 | 4 |
| + | 3.79254 | -0.8907 | Q93034 | Q93034 | CUL5 | Cullin-5 | Color2 [Da | 1 | 4 |
| + | 1.323683 | 1.38732 | Q93045 | Q93045 | STMN2 | Stathmin-2 | Color2 [Da | 1 | 4 |
| + | 1.897791 | -0.42744 | Q93050 | Q93050 | ATP6V0A1 | V-type pro | Color2 [Da | 1 | 4 |
| | 0.360223 | -0.4551 | Q93052 | Q93052 | LPP | Lipoma-pr | Color2 [Da | 1 | 4 |
| | 0.232328 | -0.21991 | Q93062 | Q93062 | RBPMS | RNA-bindi | Color2 [Da | 1 | 4 |
| | 0 | -0.19705 | Q93074 | Q93074 | MED12 | Mediator c | Color2 [Da | 1 | 4 |
| + | 1.539692 | 0.421638 | Q93088 | Q93088 | BHMT | Betaine--h | Color2 [Da | 1 | 4 |
| | 0.855629 | -0.349 | Q93096 | Q93096 | PTP4A1 | Protein tyr | Color2 [Da | 1 | 4 |
| | 0 | NaN | Q93097 | Q93097 | WNT2B | Protein W | Color2 [Da | 1 | 4 |
| | 0.203704 | -0.38545 | Q93100 | Q93100 | PHKB | Phosphory | Color2 [Da | 1 | 4 |
| | 0 | NaN | Q969E2 | Q969E2 | SCAMP4 | Secretory | Color2 [Da | 1 | 4 |
| + | 4.281665 | 3.54737 | Q969E4 | Q969E4 | TCEAL3 | Transcripti | Color2 [Da | 1 | 4 |
| + | 1.927755 | -0.31952 | Q969E8 | Q969E8 | TSR2 | Pre-rRNA-p | Color2 [Da | 1 | 4 |
| + | 1.783967 | 0.822738 | Q969G3 | Q969G3 | SMARCE1 | SWI/SNF-r | Color2 [Da | 1 | 4 |
| + | 2.5994 | 1.70624 | Q969G6 | Q969G6 | RFK | Riboflavin | Color2 [Da | 1 | 4 |
| + | 4.082644 | 1.583466 | Q969H8 | Q969H8 | MYDGF | Myeloid-d | Color2 [Da | 1 | 4 |
| + | 3.015641 | 1.940056 | Q969J3 | Q969J3 | BORCS5 | BLOC-1-rel | Color2 [Da | 1 | 4 |
| | 0 | NaN | Q969L2 | Q969L2 | MAL2 | Protein M/ | Color2 [Da | 1 | 4 |
| + | 2.532838 | 1.529769 | Q969L4 | Q969L4 | LSM10 | U7 snRNA- | Color2 [Da | 1 | 4 |
| | 0.757702 | -0.45309 | Q969M1 | Q969M1 | TOMM40L | Mitochonc | Color2 [Da | 1 | 4 |
| + | 1.007136 | -1.29833 | Q969M3 | Q969M3 | YIPF5 | Protein YIF | Color2 [Da | 1 | 4 |
| + | 2.447895 | -1.35848 | Q969N2 | Q969N2 | PIGT | GPI transa | Color2 [Da | 1 | 4 |
| | 0.456991 | -0.26284 | Q969P0 | Q969P0 | IGSF8 | Immunogl | Color2 [Da | 1 | 4 |
| + | 0.936731 | -0.35691 | Q969Q5 | Q969Q5 | RAB24 | Ras-relate | Color2 [Da | 1 | 4 |
| | 0 | NaN | Q969R2 | Q969R2 | OSBP2 | Oxysterol- | Color2 [Da | 1 | 4 |
| + | 1.32308 | 1.519907 | Q969R8 | Q969R8 | ITFG2 | KICSTOR c | Color2 [Da | 1 | 4 |
| | 0.243793 | -0.12186 | Q969S2 | Q969S2 | NEIL2 | Endonucle | Color2 [Da | 1 | 4 |
| + | 1.127011 | 0.487259 | Q969S3 | Q969S3 | ZNF622 | Zinc finger | Color2 [Da | 1 | 4 |
| | 0.579891 | -0.17193 | Q969T4 | Q969T4 | UBE2E3 | Ubiquitin-c | Color2 [Da | 1 | 4 |
| + | 3.173866 | 1.070728 | Q969T9 | Q969T9 | WBP2 | WW doma | Color2 [Da | 1 | 4 |
| + | 1.066677 | -0.29288 | Q969V3 | Q969V3 | NCLN | Nicalin | Color2 [Da | 1 | 4 |
| | 0 | NaN | Q969V5 | Q969V5 | MUL1 | Mitochonc | Color2 [Da | 1 | 4 |

| | | | | | | | | | |
|---|---|---|---|---|---|---|---|---|---|
| | 0.422538 | -0.13913 | Q969V6 | Q969V6 | MRTFA | Myocardin | Color2 [Da | 1 | 4 |
| + | 1.287682 | -1.0189 | Q969X5 | Q969X5 | ERGIC1 | Endoplasm | Color2 [Da | 1 | 4 |
| + | 1.543432 | -1.12916 | Q969X6 | Q969X6 | UTP4 | U3 small n | Color2 [Da | 1 | 4 |
| + | 1.387235 | -0.44065 | Q969Y2 | Q969Y2 | GTPBP3 | tRNA mod | Color2 [Da | 1 | 4 |
| + | 0.954295 | -0.6705 | Q969Z0 | Q969Z0 | TBRG4 | FAST kinas | Color2 [Da | 1 | 4 |
| + | 1.780848 | -0.96635 | Q96A23 | Q96A23 | CPNE4 | Copine-4 | Color2 [Da | 1 | 4 |
| + | 2.898207 | 1.469488 | Q96A26 | Q96A26 | FAM162A | Protein FA | Color2 [Da | 1 | 4 |
| + | 1.038759 | -0.44392 | Q96A33 | Q96A33 | CCDC47 | PAT compl | Color2 [Da | 1 | 4 |
| + | 1.287991 | 0.603103 | Q96A35 | Q96A35 | MRPL24 | 39S riboso | Color2 [Da | 1 | 4 |
| + | 1.55104 | 0.874698 | Q96A49 | Q96A49 | SYAP1 | Synapse-as | Color2 [Da | 1 | 4 |
| + | 3.041128 | -1.42953 | Q96A65 | Q96A65 | EXOC4 | Exocyst co | Color2 [Da | 1 | 4 |
| | 1.104997 | -0.19599 | Q96A72 | Q96A72 | MAGOHB | Protein ma | Color2 [Da | 1 | 4 |
| + | 3.857575 | 2.524587 | Q96A73 | Q96A73 | KIAA1191 | Putative m | Color2 [Da | 1 | 4 |
| + | 2.151644 | 1.261616 | Q96A83 | Q96A83 | COL26A1 | Collagen a | Color2 [Da | 1 | 4 |
| + | 3.420398 | 1.484065 | Q96A84 | Q96A84 | EMID1 | EMI doma | Color2 [Da | 1 | 4 |
| + | 2.936627 | 1.389356 | Q96AB3 | Q96AB3 | ISOC2 | Isochorism | Color2 [Da | 1 | 4 |
| + | 4.455134 | -1.18383 | Q96AC1 | Q96AC1 | FERMT2 | Fermitin fa | Color2 [Da | 1 | 4 |
| + | 4.07539 | 1.540204 | Q96AE4 | Q96AE4 | FUBP1 | Far upstre | Color2 [Da | 1 | 4 |
| + | 1.917192 | 0.597629 | Q96AG4 | Q96AG4 | LRRC59 | Leucine-ric | Color2 [Da | 1 | 4 |
| + | 1.388218 | 0.931619 | Q96AJ1 | Q96AJ1 | CLUAP1 | Clusterin-a | Color2 [Da | 1 | 4 |
| + | 1.491999 | 0.916857 | Q96AJ9 | Q96AJ9 | VTI1A | Vesicle tra | Color2 [Da | 1 | 4 |
| + | 2.739029 | -0.4377 | Q96AQ6 | Q96AQ6 | PBXIP1 | Pre-B-cell l | Color2 [Da | 1 | 4 |
| + | 1.549723 | 1.338643 | Q96AT9 | Q96AT9 | RPE | Ribulose-p | Color2 [Da | 1 | 4 |
| + | 3.542286 | -1.33115 | Q96AX1 | Q96AX1 | VPS33A | Vacuolar p | Color2 [Da | 1 | 4 |
| + | 3.63346 | -1.88242 | Q96AY3 | Q96AY3 | FKBP10 | Peptidyl-pı | Color2 [Da | 1 | 4 |
| + | 2.344414 | -1.25643 | Q96AY4 | Q96AY4 | TTC28 | Tetratricop | Color2 [Da | 1 | 4 |
| + | 3.171942 | 1.09636 | Q96B23 | Q96B23 | C18orf25 | Uncharact | Color2 [Da | 1 | 4 |
| + | 1.599342 | -1.16938 | Q96B26 | Q96B26 | EXOSC8 | Exosome c | Color2 [Da | 1 | 4 |
| + | 2.554061 | 1.698416 | Q96B36 | Q96B36 | AKT1S1 | Proline-ric | Color2 [Da | 1 | 4 |
| + | 1.26225 | 0.975297 | Q96B45 | Q96B45 | BORCS7 | BLOC-1-rel | Color2 [Da | 1 | 4 |
| + | 4.496641 | 2.746724 | Q96B49 | Q96B49 | TOMM6 | Mitochonc | Color2 [Da | 1 | 4 |
| + | 4.093905 | 3.319692 | Q96B54 | Q96B54 | ZNF428 | Zinc finger | Color2 [Da | 1 | 4 |
| + | 1.59296 | 1.542962 | Q96B86 | Q96B86 | RGMA | Repulsive g | Color2 [Da | 1 | 4 |
| + | 2.377284 | 1.290677 | Q96B97 | Q96B97 | SH3KBP1 | SH3 domai | Color2 [Da | 1 | 4 |
| | 0.266625 | 0.087375 | Q96BD5 | Q96BD5 | PHF21A | PHD finger | Color2 [Da | 1 | 4 |
| | 0 | NaN | Q96BD8 | Q96BD8 | SKA1 | Spindle an | Color2 [Da | 1 | 4 |
| | 0.64401 | -0.51059 | Q96BH1 | Q96BH1 | RNF25 | E3 ubiquiti | Color2 [Da | 1 | 4 |
| | 0.018106 | 0.0078 | Q96BJ3 | Q96BJ3 | AIDA | Axin intera | Color2 [Da | 1 | 4 |
| + | 0.67034 | -1.08618 | Q96BK5 | Q96BK5 | PINX1 | PIN2/TERF | Color2 [Da | 1 | 4 |
| + | 2.703803 | -0.67047 | Q96BM9 | Q96BM9 | ARL8A | ADP-ribosy | Color2 [Da | 1 | 4 |
| | 0 | NaN | Q96BP2 | Q96BP2 | CHCHD1 | Coiled-coil | Color2 [Da | 1 | 4 |
| + | 2.265196 | -0.85472 | Q96BP3 | Q96BP3 | PPWD1 | Peptidylpr | Color2 [Da | 1 | 4 |
| | 0.047498 | 0.012407 | Q96BW1 | Q96BW1 | UPRT | Uracil pho | Color2 [Da | 1 | 4 |
| + | 2.390819 | -0.76474 | Q96BW9 | Q96BW9 | TAMM41 | Phosphatic | Color2 [Da | 1 | 4 |
| + | 2.542371 | 0.663777 | Q96BY7 | Q96BY7 | ATG2B | Autophagy | Color2 [Da | 1 | 4 |
| + | 1.868505 | -1.8392 | Q96BY9 | Q96BY9 | SARAF | Store-oper | Color2 [Da | 1 | 4 |
| + | 1.171847 | 1.225267 | Q96C01 | Q96C01 | FAM136A | Protein FA | Color2 [Da | 1 | 4 |

| | | | | | | | | | |
|---|---|---|---|---|---|---|---|---|---|
| + | 4.156679 | 2.996111 | Q96C19 | Q96C19 | EFHD2 | EF-hand do | Color2 [Da | 1 | 4 |
| + | 2.070674 | 1.663288 | Q96C23 | Q96C23 | GALM | Galactose | Color2 [Da | 1 | 4 |
| + | 1.682116 | 0.710454 | Q96C34 | Q96C34 | RUNDC1 | RUN doma | Color2 [Da | 1 | 4 |
| | 0.094011 | 0.146035 | Q96C36 | Q96C36 | PYCR2 | Pyrroline-5 | Color2 [Da | 1 | 4 |
| + | 1.936583 | 1.688081 | Q96C57 | Q96C57 | CUSTOS | Protein CU | Color2 [Da | 1 | 4 |
| + | 2.1604 | -0.46695 | Q96C86 | Q96C86 | DCPS | m7GpppX | Color2 [Da | 1 | 4 |
| + | 2.584382 | 1.789406 | Q96C90 | Q96C90 | PPP1R14B | Protein ph | Color2 [Da | 1 | 4 |
| + | 1.058538 | -0.42617 | Q96CM4 | Q96CM4 | NXNL1 | Nucleored | Color2 [Da | 1 | 4 |
| + | 1.014111 | -1.05663 | Q96CM8 | Q96CM8 | ACSF2 | Medium-c | Color2 [Da | 1 | 4 |
| + | 1.541831 | 0.446809 | Q96CN4 | Q96CN4 | EVI5L | EVI5-like p | Color2 [Da | 1 | 4 |
| + | 1.521129 | 0.443982 | Q96CN7 | Q96CN7 | ISOC1 | Isochorism | Color2 [Da | 1 | 4 |
| + | 2.538381 | 0.772118 | Q96CN9 | Q96CN9 | GCC1 | GRIP and c | Color2 [Da | 1 | 4 |
| + | 3.767061 | 1.339955 | Q96CP2 | Q96CP2 | FLYWCH2 | FLYWCH fa | Color2 [Da | 1 | 4 |
| | 0 | -0.57708 | Q96CP6 | Q96CP6 | GRAMD1A | Protein As | Color2 [Da | 1 | 4 |
| | 0.030617 | 0.044072 | Q96CQ1;Q | Q96CQ1;Q | SLC25A36; | Solute carr | Color2 [Da | 1 | 4 |
| | 0.126065 | 0.041301 | Q96CS2 | Q96CS2 | HAUS1 | HAUS augr | Color2 [Da | 1 | 4 |
| + | 4.468329 | -1.1269 | Q96CS3 | Q96CS3 | FAF2 | FAS-associ | Color2 [Da | 1 | 4 |
| + | 1.623565 | 1.324166 | Q96CT7 | Q96CT7 | CCDC124 | Coiled-coil | Color2 [Da | 1 | 4 |
| + | 2.000508 | -0.99575 | Q96CU9 | Q96CU9 | FOXRED1 | FAD-deper | Color2 [Da | 1 | 4 |
| + | 1.142311 | -0.55138 | Q96CV9 | Q96CV9 | OPTN | Optineurir | Color2 [Da | 1 | 4 |
| + | 2.736275 | -1.34758 | Q96CW1 | Q96CW1 | AP2M1 | AP-2 comp | Color2 [Da | 1 | 4 |
| | 0.784439 | -0.30897 | Q96CW5 | Q96CW5 | TUBGCP3 | Gamma-tu | Color2 [Da | 1 | 4 |
| | 0.048923 | 0.083025 | Q96CW6 | Q96CW6 | SLC7A6OS | Probable F | Color2 [Da | 1 | 4 |
| + | 2.1295 | -0.40365 | Q96CX2 | Q96CX2 | KCTD12 | BTB/POZ d | Color2 [Da | 1 | 4 |
| + | 1.014165 | 0.621417 | Q96D05 | Q96D05 | FAM241B | Protein FA | Color2 [Da | 1 | 4 |
| + | 4.22392 | 3.217669 | Q96D09 | Q96D09 | GPRASP2 | G-protein | Color2 [Da | 1 | 4 |
| + | 1.071214 | -0.44975 | Q96D15 | Q96D15 | RCN3 | Reticuloca | Color2 [Da | 1 | 4 |
| | 0 | -1.98159 | Q96D46 | Q96D46 | NMD3 | 60S riboso | Color2 [Da | 1 | 4 |
| + | 2.124268 | -0.65565 | Q96D53 | Q96D53 | COQ8B | Atypical ki | Color2 [Da | 1 | 4 |
| + | 1.72416 | 0.734926 | Q96D71 | Q96D71 | REPS1 | RalBP1-ass | Color2 [Da | 1 | 4 |
| + | 3.929075 | -2.69582 | Q96DA0 | Q96DA0 | ZG16B | Zymogen g | Color2 [Da | 1 | 4 |
| + | 2.82292 | -1.41724 | Q96DA2 | Q96DA2 | RAB39B | Ras-relate | Color2 [Da | 1 | 4 |
| + | 4.897655 | 2.334621 | Q96DA6 | Q96DA6 | DNAJC19 | Mitochonc | Color2 [Da | 1 | 4 |
| + | 3.211939 | 2.219758 | Q96DB5 | Q96DB5 | RMDN1 | Regulator | Color2 [Da | 1 | 4 |
| | 0 | -1.82102 | Q96DC8 | Q96DC8 | ECHDC3 | Enoyl-CoA | Color2 [Da | 1 | 4 |
| + | 1.935501 | 0.884522 | Q96DE0 | Q96DE0 | NUDT16 | U8 snoRNA | Color2 [Da | 1 | 4 |
| + | 1.948967 | 0.42061 | Q96DF8 | Q96DF8 | ESS2 | Splicing fac | Color2 [Da | 1 | 4 |
| + | 1.407009 | -0.63525 | Q96DG6 | Q96DG6 | CMBL | Carboxyme | Color2 [Da | 1 | 4 |
| + | 3.446984 | 0.912177 | Q96DH6 | Q96DH6 | MSI2 | RNA-bindi | Color2 [Da | 1 | 4 |
| + | 2.243237 | -0.8906 | Q96DI7 | Q96DI7 | SNRNP40 | U5 small n | Color2 [Da | 1 | 4 |
| + | 3.33426 | -0.60821 | Q96DM3 | Q96DM3 | RMC1 | Regulator | Color2 [Da | 1 | 4 |
| + | 1.0114 | 1.134705 | Q96DV4 | Q96DV4 | MRPL38 | 39S riboso | Color2 [Da | 1 | 4 |
| + | 1.498637 | -0.81186 | Q96DZ1 | Q96DZ1 | ERLEC1 | Endoplasm | Color2 [Da | 1 | 4 |
| | 0.115492 | 0.067587 | Q96E09 | Q96E09 | PABIR1 | PPP2R1A-F | Color2 [Da | 1 | 4 |
| + | 1.136712 | 1.038215 | Q96E11 | Q96E11 | MRRF | Ribosome- | Color2 [Da | 1 | 4 |
| | 0.216587 | 0.032094 | Q96E17 | Q96E17 | RAB3C | Ras-relate | Color2 [Da | 1 | 4 |
| + | 1.066253 | 1.064919 | Q96E39 | Q96E39 | RBMXL1 | RNA bindir | Color2 [Da | 1 | 4 |

| | | | | | | | | | |
|---|---|---|---|---|---|---|---|---|---|
| + | 0.852115 | 0.86495 | Q96EA4 | Q96EA4 | SPDL1 | Protein Sp | Color2 [Da | 1 | 4 |
| | 0.327207 | 0.129611 | Q96EB1 | Q96EB1 | ELP4 | Elongator ( | Color2 [Da | 1 | 4 |
| | 0 | -0.16302 | Q96EB6 | Q96EB6 | SIRT1 | NAD-depe | Color2 [Da | 1 | 4 |
| + | 2.841565 | -1.22773 | Q96EE3 | Q96EE3 | SEH1L | Nucleopor | Color2 [Da | 1 | 4 |
| + | 1.656347 | 0.984473 | Q96EE4 | Q96EE4 | CCDC126 | Coiled-coil | Color2 [Da | 1 | 4 |
| + | 1.671278 | 2.102419 | Q96EG1 | Q96EG1 | ARSG | Arylsulfata | Color2 [Da | 1 | 4 |
| + | 2.760646 | 1.969284 | Q96EI5 | Q96EI5 | TCEAL4 | Transcripti | Color2 [Da | 1 | 4 |
| + | 1.225669 | 1.523367 | Q96EK4 | Q96EK4 | THAP11 | THAP dom | Color2 [Da | 1 | 4 |
| | 0.696358 | -0.51026 | Q96EK5 | Q96EK5 | KIFBP | KIF-binding | Color2 [Da | 1 | 4 |
| + | 1.04331 | -0.50273 | Q96EK6 | Q96EK6 | GNPNAT1 | Glucosami | Color2 [Da | 1 | 4 |
| + | 1.340176 | -0.74748 | Q96EK7 | Q96EK7 | FAM120B | Constitutiv | Color2 [Da | 1 | 4 |
| | 0.319148 | 0.411939 | Q96EK9 | Q96EK9 | KTI12 | Protein KT | Color2 [Da | 1 | 4 |
| | 0.101852 | -0.0809 | Q96EL2 | Q96EL2 | MRPS24 | 28S riboso | Color2 [Da | 1 | 4 |
| + | 1.022759 | 1.600704 | Q96EL3 | Q96EL3 | MRPL53 | 39S riboso | Color2 [Da | 1 | 4 |
| + | 2.583167 | 1.85814 | Q96EM0 | Q96EM0 | L3HYPDH | Trans-3-hy | Color2 [Da | 1 | 4 |
| + | 0.964757 | 0.422101 | Q96EP0 | Q96EP0 | RNF31 | E3 ubiquiti | Color2 [Da | 1 | 4 |
| + | 0.943116 | 0.415139 | Q96EP5 | Q96EP5 | DAZAP1 | DAZ-assoc | Color2 [Da | 1 | 4 |
| + | 1.920652 | 1.047671 | Q96EQ0 | Q96EQ0 | SGTB | Small gluta | Color2 [Da | 1 | 4 |
| | 0.404492 | -0.38093 | Q96ER3 | Q96ER3 | SAAL1 | Protein SA | Color2 [Da | 1 | 4 |
| + | 2.826522 | 0.545727 | Q96ER9 | Q96ER9 | CCDC51 | Mitochonc | Color2 [Da | 1 | 4 |
| + | 0.768923 | 0.530307 | Q96ES7 | Q96ES7 | SGF29 | SAGA-asso | Color2 [Da | 1 | 4 |
| + | 1.084277 | 0.286983 | Q96EV2 | Q96EV2 | RBM33 | RNA-bindi | Color2 [Da | 1 | 4 |
| | 0.406117 | 0.20571 | Q96EX1 | Q96EX1 | SMIM12 | Small integ | Color2 [Da | 1 | 4 |
| | 0 | NaN | Q96EX2 | Q96EX2 | RNFT2 | RING finge | Color2 [Da | 1 | 4 |
| + | 3.812827 | 1.477742 | Q96EY1 | Q96EY1 | DNAJA3 | DnaJ homo | Color2 [Da | 1 | 4 |
| + | 1.890545 | -1.06306 | Q96EY4 | Q96EY4 | TMA16 | Translatior | Color2 [Da | 1 | 4 |
| + | 1.175123 | -0.74755 | Q96EY7 | Q96EY7 | PTCD3 | Pentatrico | Color2 [Da | 1 | 4 |
| + | 3.43913 | 0.819699 | Q96EY8 | Q96EY8 | MMAB | Corrinoid a | Color2 [Da | 1 | 4 |
| | 0 | -1.73954 | Q96EZ8 | Q96EZ8 | MCRS1 | Microsphe | Color2 [Da | 1 | 4 |
| + | 3.622081 | -1.23935 | Q96F07 | Q96F07 | CYFIP2 | Cytoplasm | Color2 [Da | 1 | 4 |
| | 0.223588 | -0.26086 | Q96F10 | Q96F10 | SAT2 | Thialysine | Color2 [Da | 1 | 4 |
| | 0.312023 | 0.193465 | Q96F63 | Q96F63 | CCDC97 | Coiled-coil | Color2 [Da | 1 | 4 |
| + | 3.394447 | 1.841448 | Q96F85 | Q96F85 | CNRIP1 | CB1 canna | Color2 [Da | 1 | 4 |
| + | 2.197351 | -0.74272 | Q96F86 | Q96F86 | EDC3 | Enhancer c | Color2 [Da | 1 | 4 |
| | 0.184191 | -0.04767 | Q96FC7 | Q96FC7 | PHYHIPL | Phytanoyl- | Color2 [Da | 1 | 4 |
| + | 1.055309 | -1.76444 | Q96FF9 | Q96FF9 | CDCA5 | Sororin | Color2 [Da | 1 | 4 |
| + | 1.932117 | 1.46137 | Q96FH0 | Q96FH0 | BORCS8 | BLOC-1-rel | Color2 [Da | 1 | 4 |
| | 0.607423 | 0.528636 | Q96FJ2 | Q96FJ2 | DYNLL2 | Dynein ligh | Color2 [Da | 1 | 4 |
| | 0.594432 | 0.395141 | Q96FN4 | Q96FN4 | CPNE2 | Copine-2 | Color2 [Da | 1 | 4 |
| | 0 | 2.176045 | Q96FN9 | Q96FN9 | DTD2 | D-aminoac | Color2 [Da | 1 | 4 |
| + | 1.668479 | 1.807313 | Q96FQ6 | Q96FQ6 | S100A16 | Protein S1( | Color2 [Da | 1 | 4 |
| + | 1.352194 | 1.00609 | Q96FT9 | Q96FT9 | IFT43 | Intraflagell | Color2 [Da | 1 | 4 |
| + | 2.683446 | -1.17647 | Q96FV2 | Q96FV2 | SCRN2 | Secernin-2 | Color2 [Da | 1 | 4 |
| | 0.403902 | -0.38593 | Q96FV9 | Q96FV9 | THOC1 | THO comp | Color2 [Da | 1 | 4 |
| + | 3.018092 | -0.92517 | Q96FW1 | Q96FW1 | OTUB1 | Ubiquitin t | Color2 [Da | 1 | 4 |
| | 0.517585 | -0.46727 | Q96FX7 | Q96FX7 | TRMT61A | tRNA (ader | Color2 [Da | 1 | 4 |
| | 0 | NaN | Q96FZ2 | Q96FZ2 | HMCES | Abasic site | Color2 [Da | 1 | 4 |

| | | | | | | | | | |
|---|---|---|---|---|---|---|---|---|---|
| + | 0.932436 | 0.897725 | Q96FZ7 | Q96FZ7 | CHMP6 | Charged m | Color2 [Da | 1 | 4 |
| + | 2.548556 | 1.177681 | Q96G01 | Q96G01 | BICD1 | Protein bic | Color2 [Da | 1 | 4 |
| + | 1.317418 | -0.61527 | Q96G03 | Q96G03 | PGM2 | Phosphop‹ | Color2 [Da | 1 | 4 |
| + | 0.685494 | 0.867628 | Q96G21 | Q96G21 | IMP4 | U3 small n | Color2 [Da | 1 | 4 |
| | 0 | -2.66576 | Q96G23 | Q96G23 | CERS2 | Ceramide : | Color2 [Da | 1 | 4 |
| | 0.540548 | 0.245456 | Q96G25 | Q96G25 | MED8 | Mediator ( | Color2 [Da | 1 | 4 |
| + | 2.537495 | 0.835902 | Q96G28 | Q96G28 | CFAP36 | Cilia- and f | Color2 [Da | 1 | 4 |
| | 0 | -0.14625 | Q96G46 | Q96G46 | DUS3L | tRNA-dihy( | Color2 [Da | 1 | 4 |
| | 0.096646 | 0.282007 | Q96G61 | Q96G61 | NUDT11 | Diphospho | Color2 [Da | 1 | 4 |
| | 0 | -0.52105 | Q96G74 | Q96G74 | OTUD5 | OTU doma | Color2 [Da | 1 | 4 |
| + | 1.980975 | -1.96821 | Q96G97 | Q96G97 | BSCL2 | Seipin | Color2 [Da | 1 | 4 |
| + | 2.354509 | -1.55774 | Q96GA3 | Q96GA3 | LTV1 | Protein LT' | Color2 [Da | 1 | 4 |
| + | 2.595045 | -0.54884 | Q96GA7 | Q96GA7 | SDSL | Serine deh | Color2 [Da | 1 | 4 |
| + | 1.970951 | 1.211448 | Q96GC5 | Q96GC5 | MRPL48 | 39S riboso | Color2 [Da | 1 | 4 |
| | 0 | NaN | Q96GC9 | Q96GC9 | VMP1 | Vacuole m | Color2 [Da | 1 | 4 |
| + | 1.685544 | 0.526121 | Q96GD0 | Q96GD0 | PDXP | Chronophi | Color2 [Da | 1 | 4 |
| | 0 | -0.11811 | Q96GD4 | Q96GD4 | AURKB | Aurora kin | Color2 [Da | 1 | 4 |
| + | 2.460883 | 0.790379 | Q96GE9 | Q96GE9 | DMAC1 | Distal men | Color2 [Da | 1 | 4 |
| | 0 | NaN | Q96GF1 | Q96GF1 | RNF185 | E3 ubiquiti | Color2 [Da | 1 | 4 |
| | 0.314879 | 0.16918 | Q96GG9 | Q96GG9 | DCUN1D1 | DCN1-like | Color2 [Da | 1 | 4 |
| + | 0.752899 | 0.585289 | Q96GK7 | Q96GK7 | FAHD2A | Fumarylac | Color2 [Da | 1 | 4 |
| + | 2.888613 | -0.62938 | Q96GM5 | Q96GM5 | SMARCD1 | SWI/SNF-r | Color2 [Da | 1 | 4 |
| | 0 | NaN | Q96GP6 | Q96GP6 | SCARF2 | Scavenger | Color2 [Da | 1 | 4 |
| + | 1.05162 | -0.87939 | Q96GQ5 | Q96GQ5 | RUSF1 | RUS family | Color2 [Da | 1 | 4 |
| | 0.423661 | -0.59899 | Q96GQ7 | Q96GQ7 | DDX27 | Probable A | Color2 [Da | 1 | 4 |
| + | 1.835726 | 1.695663 | Q96GS4 | Q96GS4 | BORCS6 | BLOC-1-rel | Color2 [Da | 1 | 4 |
| + | 0.966236 | -0.80745 | Q96GV9 | Q96GV9 | MACIR | Macropha | Color2 [Da | 1 | 4 |
| + | 0.883814 | 0.714453 | Q96GW7 | Q96GW7 | BCAN | Brevican c | Color2 [Da | 1 | 4 |
| | 0.435694 | -0.43124 | Q96GW9 | Q96GW9 | MARS2 | Methionin | Color2 [Da | 1 | 4 |
| | 0.058971 | 0.044058 | Q96GX2 | Q96GX2 | ATXN7L3B | Ataxin-7-li | Color2 [Da | 1 | 4 |
| | 0.704844 | 0.503915 | Q96GX9 | Q96GX9 | APIP | Methylthic | Color2 [Da | 1 | 4 |
| + | 2.302051 | 1.0962 | Q96GY0 | Q96GY0 | ZC2HC1A | Zinc finger | Color2 [Da | 1 | 4 |
| + | 1.309291 | -0.85386 | Q96GZ6 | Q96GZ6 | SLC41A3 | Solute carr | Color2 [Da | 1 | 4 |
| + | 1.859115 | 0.499251 | Q96H20 | Q96H20 | SNF8 | Vacuolar-s | Color2 [Da | 1 | 4 |
| + | 1.399767 | 0.507959 | Q96HC4 | Q96HC4 | PDLIM5 | PDZ and LI | Color2 [Da | 1 | 4 |
| | 0.039703 | 0.06776 | Q96HD1 | Q96HD1 | CRELD1 | Protein dis | Color2 [Da | 1 | 4 |
| + | 2.537601 | -1.19345 | Q96HE7 | Q96HE7 | ERO1A | ERO1-like | Color2 [Da | 1 | 4 |
| + | 2.088767 | -1.20343 | Q96HF1 | Q96HF1 | SFRP2 | Secreted fi | Color2 [Da | 1 | 4 |
| | 0 | NaN | Q96HH4 | Q96HH4 | TMEM169 | Transmem | Color2 [Da | 1 | 4 |
| + | 1.946448 | 0.964253 | Q96HJ3 | Q96HJ3 | CCDC34 | Coiled-coil | Color2 [Da | 1 | 4 |
| + | 0.888485 | 0.650018 | Q96HJ9 | Q96HJ9 | FMC1 | Protein FM | Color2 [Da | 1 | 4 |
| + | 3.206441 | 3.858189 | Q96HN2 | Q96HN2 | AHCYL2 | Adenosylh | Color2 [Da | 1 | 4 |
| | 0.427687 | -0.21035 | Q96HP0 | Q96HP0 | DOCK6 | Dedicator | Color2 [Da | 1 | 4 |
| | 0 | -3.45327 | Q96HQ2 | Q96HQ2 | CDKN2AIP | CDKN2AIP | Color2 [Da | 1 | 4 |
| | 1.159085 | 0.204441 | Q96HR3 | Q96HR3 | MED30 | Mediator ( | Color2 [Da | 1 | 4 |
| | 0 | -0.48367 | Q96HR9 | Q96HR9 | REEP6 | Receptor ( | Color2 [Da | 1 | 4 |
| | 0.936932 | -0.19728 | Q96HS1 | Q96HS1 | PGAM5 | Serine/thr | Color2 [Da | 1 | 4 |

| | | | | | | | | | |
|---|---|---|---|---|---|---|---|---|---|
| + | 2.236408 | 0.968951 | Q96HU8 | Q96HU8 | DIRAS2 | GTP-bindir | Color2 [Da | 1 | 4 |
| + | 1.187365 | -0.87427 | Q96HW7 | Q96HW7 | INTS4 | Integrator | Color2 [Da | 1 | 4 |
| + | 2.32302 | 1.283814 | Q96HY6 | Q96HY6 | DDRGK1 | DDRGK do | Color2 [Da | 1 | 4 |
| + | 0.874293 | -0.48749 | Q96HY7 | Q96HY7 | DHTKD1 | 2-oxoadipa | Color2 [Da | 1 | 4 |
| + | 3.596595 | -0.91776 | Q96I15 | Q96I15 | SCLY | Selenocyst | Color2 [Da | 1 | 4 |
| | 0.254867 | -0.086 | Q96I24 | Q96I24 | FUBP3 | Far upstre | Color2 [Da | 1 | 4 |
| + | 3.935134 | 0.733371 | Q96I25 | Q96I25 | RBM17 | Splicing fa | Color2 [Da | 1 | 4 |
| | 0 | NaN | Q96I51 | Q96I51 | RCC1L | RCC1-like ( | Color2 [Da | 1 | 4 |
| + | 3.868486 | -1.4704 | Q96I59 | Q96I59 | NARS2 | Probable a | Color2 [Da | 1 | 4 |
| | 0.27939 | 0.244591 | Q96I99 | Q96I99 | SUCLG2 | Succinate- | Color2 [Da | 1 | 4 |
| | 0 | NaN | Q96ID5 | Q96ID5 | IGSF21 | Immunogl | Color2 [Da | 1 | 4 |
| | 0 | NaN | Q96IG2 | Q96IG2 | FBXL20 | F-box/LRR | Color2 [Da | 1 | 4 |
| + | 1.422743 | -0.60682 | Q96IJ6 | Q96IJ6 | GMPPA | Mannose-1 | Color2 [Da | 1 | 4 |
| + | 2.480083 | 1.57692 | Q96IK1 | Q96IK1 | BOD1 | Biorientati | Color2 [Da | 1 | 4 |
| + | 4.135867 | 1.01387 | Q96IU4 | Q96IU4 | ABHD14B | Putative pr | Color2 [Da | 1 | 4 |
| + | 1.56254 | 0.473091 | Q96IV0 | Q96IV0 | NGLY1 | Peptide-N( | Color2 [Da | 1 | 4 |
| + | 2.324857 | 0.815173 | Q96IW2 | Q96IW2 | SHD | SH2 domai | Color2 [Da | 1 | 4 |
| + | 0.905 | 0.457886 | Q96IW7 | Q96IW7 | SEC22A | Vesicle-tra | Color2 [Da | 1 | 4 |
| + | 1.80728 | 2.151259 | Q96IX5 | Q96IX5 | ATP5MK | ATP syntha | Color2 [Da | 1 | 4 |
| + | 3.33277 | 2.252408 | Q96IZ0 | Q96IZ0 | PAWR | PRKC apop | Color2 [Da | 1 | 4 |
| + | 1.439447 | -0.65279 | Q96IZ7 | Q96IZ7 | RSRC1 | Serine/Arg | Color2 [Da | 1 | 4 |
| + | 1.718233 | -0.94658 | Q96J01 | Q96J01 | THOC3 | THO comp | Color2 [Da | 1 | 4 |
| + | 1.287428 | -0.61236 | Q96J02 | Q96J02 | ITCH | E3 ubiquiti | Color2 [Da | 1 | 4 |
| + | 1.474179 | -0.94451 | Q96J42 | Q96J42 | TXNDC15 | Thioredoxi | Color2 [Da | 1 | 4 |
| + | 2.445734 | -0.22849 | Q96J84 | Q96J84 | KIRREL1 | Kin of IRRE | Color2 [Da | 1 | 4 |
| | 0.610655 | 0.581279 | Q96JA1 | Q96JA1 | LRIG1 | Leucine-ric | Color2 [Da | 1 | 4 |
| | 0.167534 | 0.032578 | Q96JB1 | Q96JB1 | DNAH8 | Dynein axo | Color2 [Da | 1 | 4 |
| + | 1.535986 | -1.10101 | Q96JB2 | Q96JB2 | COG3 | Conserved | Color2 [Da | 1 | 4 |
| + | 1.520219 | 0.282406 | Q96JB5 | Q96JB5 | CDK5RAP3 | CDK5 regu | Color2 [Da | 1 | 4 |
| | 0 | NaN | Q96JC1 | Q96JC1 | VPS39 | Vam6/Vps | Color2 [Da | 1 | 4 |
| + | 0.890232 | -1.07584 | Q96JC9 | Q96JC9 | EAF1 | ELL-associa | Color2 [Da | 1 | 4 |
| + | 3.651579 | 2.728742 | Q96JE9 | Q96JE9 | MAP6 | Microtubu | Color2 [Da | 1 | 4 |
| | 0.493633 | -0.3137 | Q96JG6 | Q96JG6 | VPS50 | Syndetin | Color2 [Da | 1 | 4 |
| | 1.206645 | 0.179792 | Q96JG8 | Q96JG8 | MAGED4B | Melanoma | Color2 [Da | 1 | 4 |
| + | 1.368745 | -0.39307 | Q96JH7 | Q96JH7 | VCPIP1 | Deubiquiti | Color2 [Da | 1 | 4 |
| + | 1.115255 | 0.788591 | Q96JH8 | Q96JH8 | RADIL | Ras-associ | Color2 [Da | 1 | 4 |
| | 0 | 0.381037 | Q96JI7 | Q96JI7 | SPG11 | Spatacsin | Color2 [Da | 1 | 4 |
| + | 3.165047 | -2.0741 | Q96JJ3 | Q96JJ3 | ELMO2 | Engulfmen | Color2 [Da | 1 | 4 |
| + | 2.414823 | -1.4143 | Q96JJ7 | Q96JJ7 | TMX3 | Protein dis | Color2 [Da | 1 | 4 |
| + | 1.735632 | 0.900911 | Q96JK2 | Q96JK2 | DCAF5 | DDB1- and | Color2 [Da | 1 | 4 |
| + | 1.661511 | -0.57337 | Q96JM2 | Q96JM2 | ZNF462 | Zinc finger | Color2 [Da | 1 | 4 |
| | 0.450597 | 0.262412 | Q96JM3 | Q96JM3 | CHAMP1 | Chromoso | Color2 [Da | 1 | 4 |
| | 0.214596 | -0.15665 | Q96JN0 | Q96JN0 | LCOR | Ligand-dep | Color2 [Da | 1 | 4 |
| | 0 | NaN | Q96JN2 | Q96JN2 | CCDC136 | Coiled-coil | Color2 [Da | 1 | 4 |
| + | 2.110242 | 1.392746 | Q96JP5 | Q96JP5 | ZFP91 | E3 ubiquiti | Color2 [Da | 1 | 4 |
| + | 3.468781 | -0.70956 | Q96JQ0 | Q96JQ0 | DCHS1 | Protocadh | Color2 [Da | 1 | 4 |
| | 0 | NaN | Q96JQ2 | Q96JQ2 | CLMN | Calmin | Color2 [Da | 1 | 4 |

| | | | | | | | | | |
|---|---|---|---|---|---|---|---|---|---|
| | 0 | NaN | Q96JX3 | Q96JX3 | SERAC1 | Protein SE | Color2 [Da | 1 | 4 |
| | 0 | NaN | Q96JY6 | Q96JY6 | PDLIM2 | PDZ and LI | Color2 [Da | 1 | 4 |
| + | 2.092524 | 0.988638 | Q96K17 | Q96K17 | BTF3L4 | Transcripti | Color2 [Da | 1 | 4 |
| | 0.105504 | 0.065287 | Q96K19 | Q96K19 | RNF170 | E3 ubiquiti | Color2 [Da | 1 | 4 |
| + | 1.397888 | 0.317008 | Q96K21 | Q96K21 | ZFYVE19 | Abscission | Color2 [Da | 1 | 4 |
| + | 1.984271 | -0.59345 | Q96K76 | Q96K76 | USP47 | Ubiquitin c | Color2 [Da | 1 | 4 |
| + | 2.165148 | -1.7061 | Q96K83 | Q96K83 | ZNF521 | Zinc finger | Color2 [Da | 1 | 4 |
| | 0 | -0.6062 | Q96KA5 | Q96KA5 | CLPTM1L | Cleft lip an | Color2 [Da | 1 | 4 |
| | 0.28359 | -0.18387 | Q96KB5 | Q96KB5 | PBK | Lymphokir | Color2 [Da | 1 | 4 |
| | 0 | NaN | Q96KG7 | Q96KG7 | MEGF10 | Multiple e | Color2 [Da | 1 | 4 |
| + | 3.079922 | -1.02196 | Q96KG9 | Q96KG9 | SCYL1 | N-termina | Color2 [Da | 1 | 4 |
| + | 1.627001 | 0.371148 | Q96KM6 | Q96KM6 | ZNF512B | Zinc finger | Color2 [Da | 1 | 4 |
| + | 2.656169 | -1.05789 | Q96KN1 | Q96KN1 | LRATD2 | Protein LR | Color2 [Da | 1 | 4 |
| + | 0.857223 | 0.648487 | Q96KN3 | Q96KN3 | PKNOX2 | Homeobox | Color2 [Da | 1 | 4 |
| + | 2.904597 | -0.72785 | Q96KP1 | Q96KP1 | EXOC2 | Exocyst co | Color2 [Da | 1 | 4 |
| | 1.519488 | -0.17619 | Q96KP4 | Q96KP4 | CNDP2 | Cytosolic r | Color2 [Da | 1 | 4 |
| | 0 | NaN | Q96KQ7 | Q96KQ7 | EHMT2 | Histone-ly | Color2 [Da | 1 | 4 |
| + | 3.233562 | -1.07221 | Q96KR1 | Q96KR1 | ZFR | Zinc finger | Color2 [Da | 1 | 4 |
| + | 3.757155 | 2.284868 | Q96KR6 | Q96KR6 | FAM210B | Protein FA | Color2 [Da | 1 | 4 |
| | 0 | NaN | Q96L14 | Q96L14 | CEP170P1 | Cep170-lik | Color2 [Da | 1 | 4 |
| | 0 | -1.29325 | Q96L58 | Q96L58 | B3GALT6 | Beta-1,3-g | Color2 [Da | 1 | 4 |
| | 0.492908 | -0.22689 | Q96L91 | Q96L91 | EP400 | E1A-bindir | Color2 [Da | 1 | 4 |
| + | 1.528035 | -0.48298 | Q96L92 | Q96L92 | SNX27 | Sorting ne | Color2 [Da | 1 | 4 |
| | 0.038438 | -0.01762 | Q96LB3 | Q96LB3 | IFT74 | Intraflagell | Color2 [Da | 1 | 4 |
| + | 0.707489 | 1.711636 | Q96LD8 | Q96LD8 | SENP8 | Sentrin-sp | Color2 [Da | 1 | 4 |
| + | 1.320019 | -1.46332 | Q96LI5;Q9 | Q96LI5;Q9 | CNOT6L;CI | CCR4-NOT | Color2 [Da | 1 | 4 |
| | 0 | 0.220643 | Q96LT4 | Q96LT4 | SAMD8 | Sphingomy | Color2 [Da | 1 | 4 |
| + | 2.067146 | -1.67885 | Q96LT7 | Q96LT7 | C9orf72 | Guanine n | Color2 [Da | 1 | 4 |
| | 0.545442 | -0.62229 | Q96LZ7 | Q96LZ7 | RMDN2 | Regulator | Color2 [Da | 1 | 4 |
| | 0.89748 | 0.309419 | Q96M27 | Q96M27 | PRRC1 | Protein PR | Color2 [Da | 1 | 4 |
| + | 1.39429 | -0.33533 | Q96M96 | Q96M96 | FGD4 | FYVE, Rho | Color2 [Da | 1 | 4 |
| + | 1.719179 | 1.331433 | Q96MC5 | Q96MC5 | BMERB1 | bMERB do | Color2 [Da | 1 | 4 |
| | 0 | -0.40789 | Q96ME1 | Q96ME1 | FBXL18 | F-box/LRR | Color2 [Da | 1 | 4 |
| + | 1.141108 | 0.615106 | Q96ME7 | Q96ME7 | ZNF512 | Zinc finger | Color2 [Da | 1 | 4 |
| + | 2.208533 | 1.178113 | Q96MG7 | Q96MG7 | NSMCE3 | Non-struct | Color2 [Da | 1 | 4 |
| + | 2.005676 | -1.62798 | Q96MH6 | Q96MH6 | TMEM68 | Transmem | Color2 [Da | 1 | 4 |
| | 0 | NaN | Q96MM6 | Q96MM6 | HSPA12B | Heat shock | Color2 [Da | 1 | 4 |
| | 0 | 1.519769 | Q96MM7 | Q96MM7 | HS6ST2 | Heparan-s | Color2 [Da | 1 | 4 |
| | 0.328525 | 0.321669 | Q96MP8 | Q96MP8 | KCTD7 | BTB/POZ d | Color2 [Da | 1 | 4 |
| | 0 | NaN | Q96MT8 | Q96MT8 | CEP63 | Centrosom | Color2 [Da | 1 | 4 |
| | 0.60428 | -0.55616 | Q96MU7 | Q96MU7 | YTHDC1 | YTH doma | Color2 [Da | 1 | 4 |
| | 0.041223 | 0.038191 | Q96MW1 | Q96MW1 | CCDC43 | Coiled-coil | Color2 [Da | 1 | 4 |
| | 0 | -2.60078 | Q96MW5 | Q96MW5 | COG8 | Conserved | Color2 [Da | 1 | 4 |
| | 0 | NaN | Q96MX0 | Q96MX0 | CMTM3 | CKLF-like M | Color2 [Da | 1 | 4 |
| + | 1.378001 | -0.70005 | Q96MX6 | Q96MX6 | DNAAF10 | Dynein axo | Color2 [Da | 1 | 4 |
| + | 0.992957 | 0.566213 | Q96MY1 | Q96MY1 | NOL4L | Nucleolar | Color2 [Da | 1 | 4 |
| + | 1.042049 | -0.69419 | Q96MZ0 | Q96MZ0 | GDAP1L1 | Gangliosid | Color2 [Da | 1 | 4 |

| | | | | | | | | | |
|---|---|---|---|---|---|---|---|---|---|
| | 0 | 1.219705 | Q96N03 | Q96N03 | VSTM2L | V-set and t | Color2 [Da | 1 | 4 |
| + | 1.186516 | 1.274232 | Q96N06 | Q96N06 | SPATA33 | Spermatog | Color2 [Da | 1 | 4 |
| | 0 | NaN | Q96N11 | Q96N11 | INTS15 | Integrator | Color2 [Da | 1 | 4 |
| + | 2.049891 | 1.145504 | Q96N16 | Q96N16 | JAKMIP1 | Janus kina: | Color2 [Da | 1 | 4 |
| + | 2.404332 | 0.428628 | Q96N64 | Q96N64 | PWWP2A | PWWP doi | Color2 [Da | 1 | 4 |
| + | 2.263381 | -1.56051 | Q96N66 | Q96N66 | MBOAT7 | Lysophosp | Color2 [Da | 1 | 4 |
| + | 2.159727 | -0.80484 | Q96N67 | Q96N67 | DOCK7 | Dedicator | Color2 [Da | 1 | 4 |
| | 0.139014 | -0.31093 | Q96NB3 | Q96NB3 | ZNF830 | Zinc finger | Color2 [Da | 1 | 4 |
| + | 1.829935 | 1.59105 | Q96ND0 | Q96ND0 | FAM210A | Protein FA | Color2 [Da | 1 | 4 |
| | 0 | NaN | Q96NL0 | Q96NL0 | RUNDC3B | RUN doma | Color2 [Da | 1 | 4 |
| | 0 | NaN | Q96NM4 | Q96NM4 | TOX2 | TOX high n | Color2 [Da | 1 | 4 |
| | 0 | NaN | Q96NT0 | Q96NT0 | CCDC115 | Coiled-coil | Color2 [Da | 1 | 4 |
| + | 2.549647 | 0.843546 | Q96NT1 | Q96NT1 | NAP1L5 | Nucleosom | Color2 [Da | 1 | 4 |
| | 0 | 1.140378 | Q96NY7 | Q96NY7 | CLIC6 | Chloride ir | Color2 [Da | 1 | 4 |
| + | 2.290739 | -0.56296 | Q96P11 | Q96P11 | NSUN5 | 28S rRNA ( | Color2 [Da | 1 | 4 |
| | 0.328272 | 0.109615 | Q96P16 | Q96P16 | RPRD1A | Regulation | Color2 [Da | 1 | 4 |
| | 0.53213 | -0.17751 | Q96P47 | Q96P47 | AGAP3 | Arf-GAP w | Color2 [Da | 1 | 4 |
| + | 1.255231 | -0.53368 | Q96P48 | Q96P48 | ARAP1 | Arf-GAP w | Color2 [Da | 1 | 4 |
| + | 1.223562 | 1.113607 | Q96P63 | Q96P63 | SERPINB12 | Serpin B12 | Color2 [Da | 1 | 4 |
| + | 5.049818 | -1.41201 | Q96P70 | Q96P70 | IPO9 | Importin-9 | Color2 [Da | 1 | 4 |
| + | 1.715052 | -0.33533 | Q96PC5 | Q96PC5 | MIA2 | Melanoma | Color2 [Da | 1 | 4 |
| + | 1.929886 | -0.72936 | Q96PE3 | Q96PE3 | INPP4A | Inositol po | Color2 [Da | 1 | 4 |
| + | 0.748594 | -1.10223 | Q96PE7 | Q96PE7 | MCEE | Methylma | Color2 [Da | 1 | 4 |
| + | 1.459265 | 0.381122 | Q96PK6 | Q96PK6 | RBM14 | RNA-bindi | Color2 [Da | 1 | 4 |
| | 0.38195 | -0.14874 | Q96PM9 | Q96PM9 | ZNF385A | Zinc finger | Color2 [Da | 1 | 4 |
| + | 1.032225 | 1.193746 | Q96PN7 | Q96PN7 | TRERF1 | Transcripti | Color2 [Da | 1 | 4 |
| | 0 | NaN | Q96PQ0 | Q96PQ0 | SORCS2 | VPS10 don | Color2 [Da | 1 | 4 |
| + | 1.957127 | 0.668751 | Q96PU5 | Q96PU5 | NEDD4L | E3 ubiquiti | Color2 [Da | 1 | 4 |
| + | 1.977796 | 0.874062 | Q96PU8 | Q96PU8 | QKI | Protein qu | Color2 [Da | 1 | 4 |
| + | 1.432554 | 0.986065 | Q96PV0 | Q96PV0 | SYNGAP1 | Ras/Rap G | Color2 [Da | 1 | 4 |
| | 0.076569 | -0.04252 | Q96PY5 | Q96PY5 | FMNL2 | Formin-like | Color2 [Da | 1 | 4 |
| + | 0.99926 | -0.73446 | Q96PZ0 | Q96PZ0 | PUS7 | Pseudouri | Color2 [Da | 1 | 4 |
| + | 1.647778 | 0.475323 | Q96Q04 | Q96Q04 | LMTK3 | Serine/thr | Color2 [Da | 1 | 4 |
| | 0.125193 | -0.14051 | Q96Q05 | Q96Q05 | TRAPPC9 | Trafficking | Color2 [Da | 1 | 4 |
| + | 2.990003 | 1.574754 | Q96Q11 | Q96Q11 | TRNT1 | CCA tRNA | Color2 [Da | 1 | 4 |
| + | 3.5012 | -1.57123 | Q96Q15 | Q96Q15 | SMG1 | Serine/thr | Color2 [Da | 1 | 4 |
| | 0 | -0.56814 | Q96Q42 | Q96Q42 | ALS2 | Alsin | Color2 [Da | 1 | 4 |
| + | 0.704416 | 0.8951 | Q96QA5 | Q96QA5 | GSDMA | Gasdermir | Color2 [Da | 1 | 4 |
| | 1.017868 | 0.235471 | Q96QC0 | Q96QC0 | PPP1R10 | Serine/thr | Color2 [Da | 1 | 4 |
| + | 1.297812 | 0.597202 | Q96QD9 | Q96QD9 | FYTTD1 | UAP56-int | Color2 [Da | 1 | 4 |
| | 0 | NaN | Q96QE2 | Q96QE2 | SLC2A13 | Proton my | Color2 [Da | 1 | 4 |
| + | 2.818331 | -1.88566 | Q96QG7 | Q96QG7 | MTMR9 | Myotubula | Color2 [Da | 1 | 4 |
| + | 2.81575 | -1.16022 | Q96QK1 | Q96QK1 | VPS35 | Vacuolar p | Color2 [Da | 1 | 4 |
| + | 1.484772 | 0.34709 | Q96QR8 | Q96QR8 | PURB | Transcripti | Color2 [Da | 1 | 4 |
| | 0 | -1.81449 | Q96QU8 | Q96QU8 | XPO6 | Exportin-6 | Color2 [Da | 1 | 4 |
| + | 1.845156 | 0.726733 | Q96QZ7 | Q96QZ7 | MAGI1 | Membrane | Color2 [Da | 1 | 4 |
| | 0 | -0.93493 | Q96RD7 | Q96RD7 | PANX1 | Pannexin-1 | Color2 [Da | 1 | 4 |

| | | | | | | | | | |
|---|---|---|---|---|---|---|---|---|---|
| + | 1.122165 | -0.84641 | Q96RE7 | Q96RE7 | NACC1 | Nucleus ac | Color2 [Da | 1 | 4 |
| + | 1.071471 | 1.096048 | Q96RF0 | Q96RF0 | SNX18 | Sorting ne: | Color2 [Da | 1 | 4 |
| + | 0.997494 | -0.72348 | Q96RL1 | Q96RL1 | UIMC1 | BRCA1-A c | Color2 [Da | 1 | 4 |
| + | 1.423506 | 0.991432 | Q96RL7 | Q96RL7 | VPS13A | Intermeml | Color2 [Da | 1 | 4 |
| | 0.004341 | -0.00124 | Q96RN5 | Q96RN5 | MED15 | Mediator ( | Color2 [Da | 1 | 4 |
| + | 2.833317 | -0.98319 | Q96RP9 | Q96RP9 | GFM1 | Elongation | Color2 [Da | 1 | 4 |
| + | 1.930053 | -0.93132 | Q96RQ1 | Q96RQ1 | ERGIC2 | Endoplasm | Color2 [Da | 1 | 4 |
| | 0 | 0.211831 | Q96RQ3 | Q96RQ3 | MCCC1 | Methylcro | Color2 [Da | 1 | 4 |
| + | 3.451585 | -0.93237 | Q96RR4 | Q96RR4 | CAMKK2 | Calcium/ca | Color2 [Da | 1 | 4 |
| + | 3.249503 | -1.54838 | Q96RS6 | Q96RS6 | NUDCD1 | NudC dom | Color2 [Da | 1 | 4 |
| | 0.530892 | 0.254923 | Q96RT1 | Q96RT1 | ERBIN | Erbin | Color2 [Da | 1 | 4 |
| | 0 | 0.314501 | Q96RT8 | Q96RT8 | TUBGCP5 | Gamma-tu | Color2 [Da | 1 | 4 |
| | 0 | NaN | Q96RU2 | Q96RU2 | USP28 | Ubiquitin ( | Color2 [Da | 1 | 4 |
| | 0 | NaN | Q96RW7 | Q96RW7 | HMCN1 | Hemicenti | Color2 [Da | 1 | 4 |
| + | 2.812495 | -1.47655 | Q96S06 | Q96S06 | LMF1 | Lipase mat | Color2 [Da | 1 | 4 |
| + | 1.064403 | -0.48549 | Q96S19 | Q96S19 | METTL26 | Methyltra | Color2 [Da | 1 | 4 |
| | 0.495768 | 0.252373 | Q96S44 | Q96S44 | TP53RK | EKC/KEOP: | Color2 [Da | 1 | 4 |
| | 0.576979 | -0.17962 | Q96S52 | Q96S52 | PIGS | GPI transa | Color2 [Da | 1 | 4 |
| + | 1.594435 | -0.52541 | Q96S55 | Q96S55 | WRNIP1 | ATPase WI | Color2 [Da | 1 | 4 |
| + | 0.82226 | -0.46735 | Q96S59 | Q96S59 | RANBP9 | Ran-bindir | Color2 [Da | 1 | 4 |
| | 0.541289 | -0.28492 | Q96S66 | Q96S66 | CLCC1 | Chloride cl | Color2 [Da | 1 | 4 |
| + | 1.614302 | 0.951064 | Q96S82 | Q96S82 | UBL7 | Ubiquitin-l | Color2 [Da | 1 | 4 |
| + | 1.688247 | 1.457616 | Q96S90 | Q96S90 | LYSMD1 | LysM and | | Color2 [Da | 1 | 4 |
| | 0 | NaN | Q96S97 | Q96S97 | MYADM | Myeloid-a: | Color2 [Da | 1 | 4 |
| + | 3.050294 | 0.847622 | Q96SB3 | Q96SB3 | PPP1R9B | Neurabin-: | Color2 [Da | 1 | 4 |
| + | 1.64378 | -0.6149 | Q96SB4 | Q96SB4 | SRPK1 | SRSF prote | Color2 [Da | 1 | 4 |
| + | 2.766075 | -2.3438 | Q96SB8 | Q96SB8 | SMC6 | Structural | Color2 [Da | 1 | 4 |
| + | 1.293736 | -1.22184 | Q96SI1 | Q96SI1 | KCTD15 | BTB/POZ d | Color2 [Da | 1 | 4 |
| | 0.192395 | 0.126672 | Q96SI9 | Q96SI9 | STRBP | Spermatid | Color2 [Da | 1 | 4 |
| | 0.289282 | -0.15327 | Q96SK2 | Q96SK2 | TMEM209 | Transmem | Color2 [Da | 1 | 4 |
| + | 1.509828 | -0.66015 | Q96SM3 | Q96SM3 | CPXM1 | Probable c | Color2 [Da | 1 | 4 |
| + | 2.907973 | 0.879359 | Q96ST2 | Q96ST2 | IWS1 | Protein IW | Color2 [Da | 1 | 4 |
| + | 2.034549 | -0.51476 | Q96ST3 | Q96ST3 | SIN3A | Paired am| | Color2 [Da | 1 | 4 |
| + | 2.114305 | -0.6856 | Q96SU4 | Q96SU4 | OSBPL9 | Oxysterol- | Color2 [Da | 1 | 4 |
| + | 1.999091 | -0.86056 | Q96SW2 | Q96SW2 | CRBN | Protein cer | Color2 [Da | 1 | 4 |
| + | 2.623401 | -1.57319 | Q96SY0 | Q96SY0 | INTS14 | Integrator | Color2 [Da | 1 | 4 |
| + | 0.806727 | 0.476476 | Q96SZ5 | Q96SZ5 | ADO | 2-aminoet | Color2 [Da | 1 | 4 |
| + | 1.943479 | -0.90882 | Q96T23 | Q96T23 | RSF1 | Remodelir | Color2 [Da | 1 | 4 |
| | 0 | NaN | Q96T25 | Q96T25 | ZIC5 | Zinc finger | Color2 [Da | 1 | 4 |
| + | 3.579531 | 0.847199 | Q96T37 | Q96T37 | RBM15 | RNA-bindi | Color2 [Da | 1 | 4 |
| + | 1.322236 | -0.69907 | Q96T51 | Q96T51 | RUFY1 | RUN and F | Color2 [Da | 1 | 4 |
| | 0.393758 | 0.263605 | Q96T58 | Q96T58 | SPEN | Msx2-inter | Color2 [Da | 1 | 4 |
| | 0 | NaN | Q96T60 | Q96T60 | PNKP | Bifunction | Color2 [Da | 1 | 4 |
| + | 0.86885 | -1.36733 | Q96T76 | Q96T76 | MMS19 | MMS19 nu | Color2 [Da | 1 | 4 |
| | 0.151259 | -0.07969 | Q96T88 | Q96T88 | UHRF1 | E3 ubiquiti | Color2 [Da | 1 | 4 |
| + | 1.672 | -0.86089 | Q96TA1 | Q96TA1 | NIBAN2 | Protein Nil | Color2 [Da | 1 | 4 |
| + | 1.653069 | 0.614546 | Q96TA2 | Q96TA2 | YME1L1 | ATP-deper | Color2 [Da | 1 | 4 |

| | | | | | | | | | |
|---|---|---|---|---|---|---|---|---|---|
| + | 1.44107 | 0.538327 | Q96TC7 | Q96TC7 | RMDN3 | Regulator | Color2 [Da | 1 | 4 |
| | 0.112419 | -0.06784 | Q99081 | Q99081 | TCF12 | Transcripti | Color2 [Da | 1 | 4 |
| | 0.081011 | 0.10799 | Q99250 | Q99250 | SCN2A | Sodium ch | Color2 [Da | 1 | 4 |
| + | 2.878564 | 1.423776 | Q99417 | Q99417 | MYCBP | c-Myc-bin | Color2 [Da | 1 | 4 |
| | 0.154759 | -0.15561 | Q99418 | Q99418 | CYTH2 | Cytohesin- | Color2 [Da | 1 | 4 |
| | 0.202842 | 0.135317 | Q99426 | Q99426 | TBCB | Tubulin-fo | Color2 [Da | 1 | 4 |
| + | 3.149206 | 1.84798 | Q99436 | Q99436 | PSMB7 | Proteasom | Color2 [Da | 1 | 4 |
| + | 1.008935 | -0.54711 | Q99439 | Q99439 | CNN2 | Calponin-2 | Color2 [Da | 1 | 4 |
| | 0.532519 | -0.67488 | Q99442 | Q99442 | SEC62 | Translocat | Color2 [Da | 1 | 4 |
| | 0.485438 | 0.081885 | Q99447 | Q99447 | PCYT2 | Ethanolam | Color2 [Da | 1 | 4 |
| + | 1.415539 | -0.37296 | Q99459 | Q99459 | CDC5L | Cell divisio | Color2 [Da | 1 | 4 |
| + | 4.474779 | -1.00302 | Q99460 | Q99460 | PSMD1 | 26S protea | Color2 [Da | 1 | 4 |
| + | 1.5797 | 0.570037 | Q99466 | Q99466 | NOTCH4 | Neurogeni | Color2 [Da | 1 | 4 |
| | 0.013362 | 0.003915 | Q99470 | Q99470 | SDF2 | Stromal ce | Color2 [Da | 1 | 4 |
| + | 3.07955 | 1.249268 | Q99471 | Q99471 | PFDN5 | Prefoldin s | Color2 [Da | 1 | 4 |
| | 0 | -1.32685 | Q99487 | Q99487 | PAFAH2 | Platelet-ac | Color2 [Da | 1 | 4 |
| + | 2.272288 | 0.848747 | Q99490 | Q99490 | AGAP2 | Arf-GAP w | Color2 [Da | 1 | 4 |
| + | 1.390595 | 0.735197 | Q99496 | Q99496 | RNF2 | E3 ubiquiti | Color2 [Da | 1 | 4 |
| + | 3.301684 | 2.646028 | Q99497 | Q99497 | PARK7 | Parkinson | Color2 [Da | 1 | 4 |
| | 0 | NaN | Q99504 | Q99504 | EYA3 | Eyes abser | Color2 [Da | 1 | 4 |
| | 0.573682 | 0.393016 | Q99519 | Q99519 | NEU1 | Sialidase-1 | Color2 [Da | 1 | 4 |
| + | 1.842228 | -0.78308 | Q99523 | Q99523 | SORT1 | Sortilin | Color2 [Da | 1 | 4 |
| + | 1.814049 | -0.60601 | Q99536 | Q99536 | VAT1 | Synaptic v | Color2 [Da | 1 | 4 |
| + | 3.042854 | 1.097479 | Q99538 | Q99538 | LGMN | Legumain | Color2 [Da | 1 | 4 |
| | 0.345871 | -0.09231 | Q99541 | Q99541 | PLIN2 | Perilipin-2 | Color2 [Da | 1 | 4 |
| + | 1.907351 | -1.08529 | Q99543 | Q99543 | DNAJC2 | DnaJ hom | Color2 [Da | 1 | 4 |
| + | 1.519395 | -0.61775 | Q99547 | Q99547 | MPHOSPH | M-phase p | Color2 [Da | 1 | 4 |
| + | 3.840179 | -1.30305 | Q99567 | Q99567 | NUP88 | Nuclear pc | Color2 [Da | 1 | 4 |
| + | 1.205705 | -1.4884 | Q99569 | Q99569 | PKP4 | Plakophilir | Color2 [Da | 1 | 4 |
| + | 1.203341 | -0.46462 | Q99570 | Q99570 | PIK3R4 | Phosphoin | Color2 [Da | 1 | 4 |
| + | 3.4672 | 3.112753 | Q99574 | Q99574 | SERPINI1 | Neuroserp | Color2 [Da | 1 | 4 |
| + | 2.544507 | -1.77924 | Q99575 | Q99575 | POP1 | Ribonucle | Color2 [Da | 1 | 4 |
| + | 2.993867 | 1.815091 | Q99576 | Q99576 | TSC22D3 | TSC22 don | Color2 [Da | 1 | 4 |
| + | 3.162529 | 3.328368 | Q99584 | Q99584 | S100A13 | Protein S1 | Color2 [Da | 1 | 4 |
| | 0 | -0.70218 | Q99590 | Q99590 | SCAF11 | Protein SC | Color2 [Da | 1 | 4 |
| + | 2.264185 | 1.360677 | Q99598 | Q99598 | TSNAX | Translin-as | Color2 [Da | 1 | 4 |
| | 0.111626 | -0.0234 | Q99608 | Q99608 | NDN | Necdin | Color2 [Da | 1 | 4 |
| | 0.274607 | -0.42678 | Q99611 | Q99611 | SEPHS2 | Selenide, v | Color2 [Da | 1 | 4 |
| + | 4.061406 | -1.48039 | Q99613 | Q99613 | EIF3C | Eukaryotic | Color2 [Da | 1 | 4 |
| + | 1.869839 | 1.301906 | Q99614 | Q99614 | TTC1 | Tetratrico | Color2 [Da | 1 | 4 |
| + | 0.961277 | -1.00862 | Q99615 | Q99615 | DNAJC7 | DnaJ hom | Color2 [Da | 1 | 4 |
| + | 3.191928 | 1.008986 | Q99622 | Q99622 | C12orf57 | Protein C1 | Color2 [Da | 1 | 4 |
| + | 5.127632 | 2.069357 | Q99623 | Q99623 | PHB2 | Prohibitin- | Color2 [Da | 1 | 4 |
| + | 2.21117 | 0.846372 | Q99627 | Q99627 | COPS8 | COP9 sign | Color2 [Da | 1 | 4 |
| + | 2.00939 | 0.267678 | Q99633 | Q99633 | PRPF18 | Pre-mRNA | Color2 [Da | 1 | 4 |
| | 0.045531 | 0.048975 | Q99638 | Q99638 | RAD9A | Cell cycle c | Color2 [Da | 1 | 4 |
| | 0.590808 | -0.40583 | Q99643 | Q99643 | SDHC | Succinate | Color2 [Da | 1 | 4 |

| | | | | | | | | | |
|---|---|---|---|---|---|---|---|---|---|
| + | 2.024937 | 1.578513 | Q99653 | Q99653 | CHP1 | Calcineurir | Color2 [Da | 1 | 4 |
| + | 0.746372 | -0.63421 | Q99661 | Q99661 | KIF2C | Kinesin-like | Color2 [Da | 1 | 4 |
| + | 2.300137 | 1.42374 | Q99674 | Q99674 | CGREF1 | Cell growtl | Color2 [Da | 1 | 4 |
| + | 3.440909 | 1.074572 | Q99689 | Q99689 | FEZ1 | Fasciculati | Color2 [Da | 1 | 4 |
| | 0.555122 | 0.539815 | Q99700 | Q99700 | ATXN2 | Ataxin-2 | Color2 [Da | 1 | 4 |
| | 0 | -0.31537 | Q99707 | Q99707 | MTR | Methionin | Color2 [Da | 1 | 4 |
| + | 1.791143 | 0.69859 | Q99714 | Q99714 | HSD17B10 | 3-hydroxy | Color2 [Da | 1 | 4 |
| | 0 | -1.92382 | Q99715 | Q99715 | COL12A1 | Collagen a | Color2 [Da | 1 | 4 |
| + | 1.70762 | 1.308615 | Q99717 | Q99717 | SMAD5 | Mothers a | Color2 [Da | 1 | 4 |
| | 0.156039 | 0.091407 | Q99719 | Q99719 | SEPTIN5 | Septin-5 | Color2 [Da | 1 | 4 |
| + | 2.373135 | -1.38128 | Q99720 | Q99720 | SIGMAR1 | Sigma non | Color2 [Da | 1 | 4 |
| + | 4.278107 | 1.943189 | Q99729 | Q99729 | HNRNPAB | Heterogen | Color2 [Da | 1 | 4 |
| + | 2.47322 | 1.254732 | Q99733 | Q99733 | NAP1L4 | Nucleosom | Color2 [Da | 1 | 4 |
| | 0 | -1.55864 | Q99735 | Q99735 | MGST2 | Microsom | Color2 [Da | 1 | 4 |
| | 0.047276 | -0.01579 | Q99747 | Q99747 | NAPG | Gamma-sc | Color2 [Da | 1 | 4 |
| | 0.350177 | -0.39872 | Q99755 | Q99755 | PIP5K1A | Phosphatid | Color2 [Da | 1 | 4 |
| + | 3.347614 | 2.78453 | Q99757 | Q99757 | TXN2 | Thioredoxi | Color2 [Da | 1 | 4 |
| | 0 | -0.56609 | Q99758 | Q99758 | ABCA3 | Phospholip | Color2 [Da | 1 | 4 |
| + | 2.198645 | 0.645145 | Q99766 | Q99766 | DMAC2L | ATP syntha | Color2 [Da | 1 | 4 |
| | 0.675442 | -0.41417 | Q99767 | Q99767 | APBA2 | Amyloid-b | Color2 [Da | 1 | 4 |
| | 0.553371 | 0.234298 | Q99784 | Q99784 | OLFM1 | Noelin | Color2 [Da | 1 | 4 |
| + | 2.769392 | -0.67473 | Q99798 | Q99798 | ACO2 | Aconitate | Color2 [Da | 1 | 4 |
| + | 2.08875 | -1.31845 | Q99805 | Q99805 | TM9SF2 | Transmem | Color2 [Da | 1 | 4 |
| + | 2.210371 | 1.199092 | Q99807 | Q99807 | COQ7 | 5-demetho | Color2 [Da | 1 | 4 |
| + | 2.712461 | 0.979177 | Q99816 | Q99816 | TSG101 | Tumor sus | Color2 [Da | 1 | 4 |
| + | 2.527629 | -1.06517 | Q99829 | Q99829 | CPNE1 | Copine-1 | Color2 [Da | 1 | 4 |
| + | 2.771425 | -1.07859 | Q99832 | Q99832 | CCT7 | T-complex | Color2 [Da | 1 | 4 |
| + | 2.30427 | 0.93377 | Q99848 | Q99848 | EBNA1BP2 | Probable r | Color2 [Da | 1 | 4 |
| | 0 | -1.09334 | Q99856 | Q99856 | ARID3A | AT-rich int | Color2 [Da | 1 | 4 |
| | 0.528831 | 0.113788 | Q99865;Q | Q99865;Q | SPIN2A;SP | Spindlin-2 | Color2 [Da | 1 | 4 |
| + | 2.792017 | -2.1113 | Q99873 | Q99873 | PRMT1 | Protein arg | Color2 [Da | 1 | 4 |
| + | 1.692186 | 0.804832 | Q99933 | Q99933 | BAG1 | BAG family | Color2 [Da | 1 | 4 |
| + | 1.836864 | -1.21388 | Q99942 | Q99942 | RNF5 | E3 ubiquiti | Color2 [Da | 1 | 4 |
| | 0.382763 | 0.210148 | Q99943 | Q99943 | AGPAT1 | 1-acyl-sn-g | Color2 [Da | 1 | 4 |
| | 0 | NaN | Q99946 | Q99946 | PRRT1 | Proline-ric | Color2 [Da | 1 | 4 |
| + | 4.042424 | -1.33573 | Q99959 | Q99959 | PKP2 | Plakophilir | Color2 [Da | 1 | 4 |
| + | 2.74557 | 1.281434 | Q99961 | Q99961 | SH3GL1 | Endophilin | Color2 [Da | 1 | 4 |
| + | 3.730582 | 1.872907 | Q99962 | Q99962 | SH3GL2 | Endophilin | Color2 [Da | 1 | 4 |
| + | 3.749971 | 2.450985 | Q99963 | Q99963 | SH3GL3 | Endophilin | Color2 [Da | 1 | 4 |
| | 0 | NaN | Q99969 | Q99969 | RARRES2 | Retinoic ac | Color2 [Da | 1 | 4 |
| + | 1.442866 | -1.02053 | Q99985 | Q99985 | SEMA3C | Semaphor | Color2 [Da | 1 | 4 |
| + | 1.088855 | -0.57796 | Q99986 | Q99986 | VRK1 | Serine/thr | Color2 [Da | 1 | 4 |
| + | 3.521373 | 2.393849 | Q99988 | Q99988 | GDF15 | Growth/di | Color2 [Da | 1 | 4 |
| | 0.921826 | -0.15475 | Q99996 | Q99996 | AKAP9 | A-kinase a | Color2 [Da | 1 | 4 |
| | 0.852434 | 0.155003 | Q9BPU6 | Q9BPU6 | DPYSL5 | Dihydropy | Color2 [Da | 1 | 4 |
| + | 3.562563 | 1.307448 | Q9BPW8 | Q9BPW8 | NIPSNAP1 | Protein Nip | Color2 [Da | 1 | 4 |
| | 0 | -2.71696 | Q9BPX3 | Q9BPX3 | NCAPG | Condensin | Color2 [Da | 1 | 4 |

| | | | | | | | | | |
|---|---|---|---|---|---|---|---|---|---|
| + | 2.657798 | 1.551602 | Q9BPX5 | Q9BPX5 | ARPC5L | Actin-relat | Color2 [Da | 1 | 4 |
| | 0.169986 | 0.16561 | Q9BPY3 | Q9BPY3 | FAM118B | Protein FA | Color2 [Da | 1 | 4 |
| + | 1.482632 | 0.288258 | Q9BPZ3 | Q9BPZ3 | PAIP2 | Polyadeny | Color2 [Da | 1 | 4 |
| + | 1.969989 | -0.37358 | Q9BPZ7 | Q9BPZ7 | MAPKAP1 | Target of r | Color2 [Da | 1 | 4 |
| | 0.005927 | -0.00178 | Q9BQ04 | Q9BQ04 | RBM4B | RNA-bindi | Color2 [Da | 1 | 4 |
| + | 1.788514 | 1.116912 | Q9BQ16 | Q9BQ16 | SPOCK3 | Testican-3 | Color2 [Da | 1 | 4 |
| | 0.739595 | -0.12335 | Q9BQ39 | Q9BQ39 | DDX50 | ATP-deper | Color2 [Da | 1 | 4 |
| | 0 | NaN | Q9BQ48 | Q9BQ48 | MRPL34 | 39S riboso | Color2 [Da | 1 | 4 |
| | 0 | NaN | Q9BQ52 | Q9BQ52 | ELAC2 | Zinc phosp | Color2 [Da | 1 | 4 |
| + | 3.002762 | 2.778014 | Q9BQ61 | Q9BQ61 | TRIR | Telomeras | Color2 [Da | 1 | 4 |
| + | 2.6028 | -0.70774 | Q9BQ67 | Q9BQ67 | GRWD1 | Glutamate | Color2 [Da | 1 | 4 |
| | 0.122463 | 0.185881 | Q9BQ69 | Q9BQ69 | MACROD1 | ADP-ribose | Color2 [Da | 1 | 4 |
| + | 0.850633 | -0.86858 | Q9BQ70 | Q9BQ70 | TCF25 | Transcripti | Color2 [Da | 1 | 4 |
| + | 0.87868 | 0.454624 | Q9BQ95 | Q9BQ95 | ECSIT | Evolutiona | Color2 [Da | 1 | 4 |
| + | 1.939584 | -0.56109 | Q9BQA1 | Q9BQA1 | WDR77 | Methylosc | Color2 [Da | 1 | 4 |
| | 0 | -0.48927 | Q9BQA9 | Q9BQA9 | CYBC1 | Cytochrom | Color2 [Da | 1 | 4 |
| + | 5.17711 | 1.404325 | Q9BQD3 | Q9BQD3 | KXD1 | KxDL moti | Color2 [Da | 1 | 4 |
| + | 1.489486 | -1.16763 | Q9BQE3 | Q9BQE3 | TUBA1C | Tubulin alp | Color2 [Da | 1 | 4 |
| | 0.405405 | 0.394018 | Q9BQE5 | Q9BQE5 | APOL2 | Apolipopr | Color2 [Da | 1 | 4 |
| + | 2.240555 | -0.84651 | Q9BQG0 | Q9BQG0 | MYBBP1A | Myb-bindi | Color2 [Da | 1 | 4 |
| | 0.759273 | -0.38772 | Q9BQG2 | Q9BQG2 | NUDT12 | NAD-capp | Color2 [Da | 1 | 4 |
| | 1.06698 | 0.252709 | Q9BQI0 | Q9BQI0 | AIF1L | Allograft ir | Color2 [Da | 1 | 4 |
| + | 2.193161 | 0.860317 | Q9BQI5 | Q9BQI5 | SGIP1 | SH3-conta | Color2 [Da | 1 | 4 |
| + | 1.027328 | 1.125937 | Q9BQQ3 | Q9BQQ3 | GORASP1 | Golgi reass | Color2 [Da | 1 | 4 |
| | 0 | -1.2445 | Q9BQS7 | Q9BQS7 | HEPH | Hephaesti | Color2 [Da | 1 | 4 |
| | 0 | NaN | Q9BQS8 | Q9BQS8 | FYCO1 | FYVE and c | Color2 [Da | 1 | 4 |
| | 0 | NaN | Q9BQT8 | Q9BQT8 | SLC25A21 | Mitochonc | Color2 [Da | 1 | 4 |
| | 0.239737 | 0.315546 | Q9BQT9 | Q9BQT9 | CLSTN3 | Calsynteni | Color2 [Da | 1 | 4 |
| + | 1.516372 | -0.6611 | Q9BR01 | Q9BR01 | SULT4A1 | Sulfotrans | Color2 [Da | 1 | 4 |
| | 0.086844 | -0.14653 | Q9BR61 | Q9BR61 | ACBD6 | Acyl-CoA-b | Color2 [Da | 1 | 4 |
| | 0.693379 | 0.158777 | Q9BR76 | Q9BR76 | CORO1B | Coronin-1B | Color2 [Da | 1 | 4 |
| + | 2.754583 | 0.744747 | Q9BRA0 | Q9BRA0 | NAA38 | N-alpha-ac | Color2 [Da | 1 | 4 |
| | 0.205721 | -0.19452 | Q9BRA2 | Q9BRA2 | TXNDC17 | Thioredoxi | Color2 [Da | 1 | 4 |
| | 0.547282 | 0.642679 | Q9BRD0 | Q9BRD0 | BUD13 | BUD13 hor | Color2 [Da | 1 | 4 |
| + | 3.517822 | 1.236964 | Q9BRF8 | Q9BRF8 | CPPED1 | Serine/thr | Color2 [Da | 1 | 4 |
| + | 2.349054 | 0.655162 | Q9BRG1 | Q9BRG1 | VPS25 | Vacuolar p | Color2 [Da | 1 | 4 |
| | 0.144386 | -0.28285 | Q9BRJ2 | Q9BRJ2 | MRPL45 | 39S riboso | Color2 [Da | 1 | 4 |
| + | 1.309592 | 0.257817 | Q9BRJ6 | Q9BRJ6 | C7orf50 | Uncharact | Color2 [Da | 1 | 4 |
| + | 1.050839 | -0.29077 | Q9BRJ7 | Q9BRJ7 | NUDT16L1 | Tudor-inte | Color2 [Da | 1 | 4 |
| + | 0.880311 | 1.049902 | Q9BRK0 | Q9BRK0 | REEP2 | Receptor e | Color2 [Da | 1 | 4 |
| + | 3.476591 | 0.522923 | Q9BRK5 | Q9BRK5 | SDF4 | 45 kDa cal | Color2 [Da | 1 | 4 |
| + | 2.47007 | 0.642105 | Q9BRL6 | Q9BRL6 | SRSF8 | Serine/arg | Color2 [Da | 1 | 4 |
| + | 2.510984 | -0.89492 | Q9BRP4 | Q9BRP4 | PAAF1 | Proteasom | Color2 [Da | 1 | 4 |
| + | 3.907064 | 4.387382 | Q9BRP8 | Q9BRP8 | PYM1 | Partner of | Color2 [Da | 1 | 4 |
| + | 3.361777 | 1.297882 | Q9BRQ6 | Q9BRQ6 | CHCHD6 | MICOS cor | Color2 [Da | 1 | 4 |
| + | 1.358577 | -0.8438 | Q9BRR6 | Q9BRR6 | ADPGK | ADP-deper | Color2 [Da | 1 | 4 |
| + | 0.822642 | 0.670269 | Q9BRS8 | Q9BRS8 | LARP6 | La-related | Color2 [Da | 1 | 4 |

| | | | | | | | | | |
|---|---|---|---|---|---|---|---|---|---|
| + | 1.574688 | 2.00331 | Q9BRT2 | Q9BRT2 | UQCC2 | Ubiquinol- | Color2 [Da | 1 | 4 |
| + | 2.534825 | 1.500448 | Q9BRT3 | Q9BRT3 | MIEN1 | Migration | Color2 [Da | 1 | 4 |
| | 0.234896 | 0.661603 | Q9BRT6 | Q9BRT6 | LLPH | Protein LLI | Color2 [Da | 1 | 4 |
| + | 2.837292 | 1.093461 | Q9BRT9 | Q9BRT9 | GINS4 | DNA replic | Color2 [Da | 1 | 4 |
| | 0.200919 | 0.108886 | Q9BRV8 | Q9BRV8 | SIKE1 | Suppresso | Color2 [Da | 1 | 4 |
| | 0.689126 | -0.55807 | Q9BRX2 | Q9BRX2 | PELO | Protein pe | Color2 [Da | 1 | 4 |
| | 0.544269 | -0.15279 | Q9BRX5 | Q9BRX5 | GINS3 | DNA replic | Color2 [Da | 1 | 4 |
| + | 3.299337 | 1.287622 | Q9BRX8 | Q9BRX8 | PRXL2A | Peroxiredc | Color2 [Da | 1 | 4 |
| | 0 | -0.37044 | Q9BRZ2 | Q9BRZ2 | TRIM56 | E3 ubiquiti | Color2 [Da | 1 | 4 |
| + | 0.923615 | -0.5878 | Q9BS18 | Q9BS18 | ANAPC13 | Anaphase- | Color2 [Da | 1 | 4 |
| + | 1.889014 | -0.58237 | Q9BS26 | Q9BS26 | ERP44 | Endoplasm | Color2 [Da | 1 | 4 |
| | 1.603398 | 0.168415 | Q9BS40 | Q9BS40 | LXN | Latexin | Color2 [Da | 1 | 4 |
| | 0.444989 | 0.676811 | Q9BSC4 | Q9BSC4 | NOL10 | Nucleolar | | Color2 [Da | 1 | 4 |
| + | 2.158505 | -0.41098 | Q9BSD7 | Q9BSD7 | NTPCR | Cancer-rel | Color2 [Da | 1 | 4 |
| + | 2.508478 | 0.627872 | Q9BSE4 | Q9BSE4 | HERPUD2 | Homocyst | Color2 [Da | 1 | 4 |
| | 0.047497 | 0.070416 | Q9BSF4 | Q9BSF4 | TIMM29 | Mitochonc | Color2 [Da | 1 | 4 |
| + | 1.528288 | 0.732269 | Q9BSH4 | Q9BSH4 | TACO1 | Translatior | Color2 [Da | 1 | 4 |
| | 0 | NaN | Q9BSH5 | Q9BSH5 | HDHD3 | Haloacid d | Color2 [Da | 1 | 4 |
| + | 1.764561 | -0.68655 | Q9BSJ2 | Q9BSJ2 | TUBGCP2 | Gamma-tu | Color2 [Da | 1 | 4 |
| | 0 | -0.94612 | Q9BSJ6 | Q9BSJ6 | PIMREG | Protein PII | Color2 [Da | 1 | 4 |
| + | 2.909595 | -1.40434 | Q9BSJ8 | Q9BSJ8 | ESYT1 | Extended s | Color2 [Da | 1 | 4 |
| + | 1.72399 | 0.787535 | Q9BSL1 | Q9BSL1 | UBAC1 | Ubiquitin-a | Color2 [Da | 1 | 4 |
| | 0 | -0.45028 | Q9BSQ5 | Q9BSQ5 | CCM2 | Cerebral ca | Color2 [Da | 1 | 4 |
| | 0 | NaN | Q9BSU1 | Q9BSU1 | PHAF1 | Phagosom | Color2 [Da | 1 | 4 |
| + | 2.486276 | 0.880294 | Q9BSV6 | Q9BSV6 | TSEN34 | tRNA-splic | Color2 [Da | 1 | 4 |
| + | 1.651934 | 1.578418 | Q9BSY4 | Q9BSY4 | CHCHD5 | Coiled-coil | Color2 [Da | 1 | 4 |
| + | 1.632735 | 0.849755 | Q9BSY9 | Q9BSY9 | DESI2 | Deubiquiti | Color2 [Da | 1 | 4 |
| + | 2.056877 | 0.454637 | Q9BT09 | Q9BT09 | CNPY3 | Protein ca | Color2 [Da | 1 | 4 |
| | 0 | -2.48751 | Q9BT22 | Q9BT22 | ALG1 | Chitobiosy | Color2 [Da | 1 | 4 |
| | 0.230814 | 1.290455 | Q9BT23 | Q9BT23 | LIMD2 | LIM domai | Color2 [Da | 1 | 4 |
| | 0 | 0.369792 | Q9BT40 | Q9BT40 | INPP5K | Inositol po | Color2 [Da | 1 | 4 |
| + | 1.420456 | 0.514043 | Q9BT43 | Q9BT43 | POLR3GL | DNA-direc | Color2 [Da | 1 | 4 |
| | 0.047238 | 0.091211 | Q9BT73 | Q9BT73 | PSMG3 | Proteasom | Color2 [Da | 1 | 4 |
| + | 1.597901 | -0.23739 | Q9BT78 | Q9BT78 | COPS4 | COP9 signa | Color2 [Da | 1 | 4 |
| + | 1.733882 | 1.126433 | Q9BT88 | Q9BT88 | SYT11 | Synaptota | Color2 [Da | 1 | 4 |
| | 0.452497 | -0.13779 | Q9BTA9 | Q9BTA9 | WAC | WW doma | Color2 [Da | 1 | 4 |
| | 0.465114 | 0.185305 | Q9BTC0 | Q9BTC0 | DIDO1 | Death-indu | Color2 [Da | 1 | 4 |
| | 0 | -0.08814 | Q9BTC8 | Q9BTC8 | MTA3 | Metastasis | Color2 [Da | 1 | 4 |
| + | 2.451487 | 1.637952 | Q9BTD8 | Q9BTD8 | RBM42 | RNA-bindi | Color2 [Da | 1 | 4 |
| | 0.483326 | 0.326 | Q9BTE1 | Q9BTE1 | DCTN5 | Dynactin s | Color2 [Da | 1 | 4 |
| | 0 | -0.93408 | Q9BTE3 | Q9BTE3 | MCMBP | Mini-chror | Color2 [Da | 1 | 4 |
| | 0.251209 | 0.130908 | Q9BTE6 | Q9BTE6 | AARSD1 | Alanyl-tRN | Color2 [Da | 1 | 4 |
| + | 0.892175 | 0.452539 | Q9BTE7 | Q9BTE7 | DCUN1D5 | DCN1-like | Color2 [Da | 1 | 4 |
| + | 3.21343 | 3.49264 | Q9BTL3 | Q9BTL3 | RAMAC | RNA guani | Color2 [Da | 1 | 4 |
| + | 0.747263 | 0.815972 | Q9BTM9 | Q9BTM9 | URM1 | Ubiquitin-r | Color2 [Da | 1 | 4 |
| + | 2.682643 | -0.63235 | Q9BTT0 | Q9BTT0 | ANP32E | Acidic leuc | Color2 [Da | 1 | 4 |
| + | 1.068684 | 0.539302 | Q9BTT4 | Q9BTT4 | MED10 | Mediator c | Color2 [Da | 1 | 4 |

| | | | | | | | | | |
|---|---|---|---|---|---|---|---|---|---|
| + | 1.475417 | -0.48388 | Q9BTU6 | Q9BTU6 | PI4K2A | Phosphatic | Color2 [Da | 1 | 4 |
| + | 2.587172 | -0.77233 | Q9BTV4 | Q9BTV4 | TMEM43 | Transmem | Color2 [Da | 1 | 4 |
| + | 2.293601 | -0.64408 | Q9BTV5 | Q9BTV5 | FSD1 | Fibronecti | Color2 [Da | 1 | 4 |
| + | 1.386522 | 0.809129 | Q9BTV7 | Q9BTV7 | CABLES2 | CDK5 and | Color2 [Da | 1 | 4 |
| + | 3.219077 | -1.5195 | Q9BTW9 | Q9BTW9 | TBCD | Tubulin-sp | Color2 [Da | 1 | 4 |
| + | 2.277345 | 0.923365 | Q9BTX1 | Q9BTX1 | NDC1 | Nucleopor | Color2 [Da | 1 | 4 |
| + | 2.655032 | -0.91807 | Q9BTY2 | Q9BTY2 | FUCA2 | Plasma alp | Color2 [Da | 1 | 4 |
| | 0.637938 | -0.49831 | Q9BTY7 | Q9BTY7 | HGH1 | Protein HG | Color2 [Da | 1 | 4 |
| + | 0.797575 | 0.94563 | Q9BU02 | Q9BU02 | THTPA | Thiamine-t | Color2 [Da | 1 | 4 |
| | 0 | 2.344456 | Q9BU23 | Q9BU23 | LMF2 | Lipase mat | Color2 [Da | 1 | 4 |
| + | 1.899164 | 0.95063 | Q9BU40 | Q9BU40 | CHRDL1 | Chordin-lik | Color2 [Da | 1 | 4 |
| + | 1.874217 | 1.039085 | Q9BU61 | Q9BU61 | NDUFAF3 | NADH deh | Color2 [Da | 1 | 4 |
| | 0.175462 | -0.30792 | Q9BU76 | Q9BU76 | MMTAG2 | Multiple m | Color2 [Da | 1 | 4 |
| + | 2.205489 | 1.168719 | Q9BU89 | Q9BU89 | DOHH | Deoxyhypu | Color2 [Da | 1 | 4 |
| | 0.480758 | 0.177081 | Q9BUA3 | Q9BUA3 | SPINDOC | Spindlin in | Color2 [Da | 1 | 4 |
| + | 2.286179 | 1.28827 | Q9BUB7 | Q9BUB7 | TMEM70 | Transmem | Color2 [Da | 1 | 4 |
| + | 1.213503 | -0.84298 | Q9BUE0 | Q9BUE0 | MED18 | Mediator c | Color2 [Da | 1 | 4 |
| | 0.354324 | -0.29247 | Q9BUE6 | Q9BUE6 | ISCA1 | Iron-sulfur | Color2 [Da | 1 | 4 |
| + | 1.487047 | -0.37563 | Q9BUF5 | Q9BUF5 | TUBB6 | Tubulin be | Color2 [Da | 1 | 4 |
| | 0.057707 | -0.04304 | Q9BUH6 | Q9BUH6 | PAXX | Protein PA | Color2 [Da | 1 | 4 |
| + | 2.187327 | -0.95753 | Q9BUI4 | Q9BUI4 | POLR3C | DNA-direc | Color2 [Da | 1 | 4 |
| + | 1.028169 | 0.358474 | Q9BUJ2 | Q9BUJ2 | HNRNPUL1 | Heterogen | Color2 [Da | 1 | 4 |
| | 0 | NaN | Q9BUK6 | Q9BUK6 | MSTO1 | Protein mi | Color2 [Da | 1 | 4 |
| + | 3.674264 | 2.057605 | Q9BUL8 | Q9BUL8 | PDCD10 | Programm | Color2 [Da | 1 | 4 |
| | 0.081619 | 0.315821 | Q9BUL9 | Q9BUL9 | RPP25 | Ribonuclea | Color2 [Da | 1 | 4 |
| + | 0.81793 | 2.233382 | Q9BUN5 | Q9BUN5 | CCDC28B | Coiled-coil | Color2 [Da | 1 | 4 |
| + | 0.997367 | -0.36284 | Q9BUN8 | Q9BUN8 | DERL1 | Derlin-1 | Color2 [Da | 1 | 4 |
| | 0.289754 | 0.597588 | Q9BUP0 | Q9BUP0 | EFHD1 | EF-hand do | Color2 [Da | 1 | 4 |
| + | 1.12431 | -0.26933 | Q9BUQ8 | Q9BUQ8 | DDX23 | Probable A | Color2 [Da | 1 | 4 |
| | 0.047584 | -0.0267 | Q9BUR4 | Q9BUR4 | WRAP53 | Telomeras | Color2 [Da | 1 | 4 |
| + | 3.621182 | 2.492325 | Q9BUR5 | Q9BUR5 | APOO | MICOS cor | Color2 [Da | 1 | 4 |
| + | 2.255538 | -0.47173 | Q9BUT1 | Q9BUT1 | BDH2 | 3-hydroxyl | Color2 [Da | 1 | 4 |
| + | 1.664145 | 0.525057 | Q9BV19 | Q9BV19 | C1orf50 | Uncharact | Color2 [Da | 1 | 4 |
| + | 3.229606 | -1.26549 | Q9BV20 | Q9BV20 | MRI1 | Methylthic | Color2 [Da | 1 | 4 |
| + | 1.480543 | -1.23507 | Q9BV23 | Q9BV23 | ABHD6 | Monoacylg | Color2 [Da | 1 | 4 |
| + | 0.885596 | 0.483473 | Q9BV29 | Q9BV29 | CCDC32 | Coiled-coil | Color2 [Da | 1 | 4 |
| + | 1.903795 | 0.739761 | Q9BV35 | Q9BV35 | SLC25A23 | Calcium-bi | Color2 [Da | 1 | 4 |
| + | 1.038078 | -0.37004 | Q9BV38 | Q9BV38 | WDR18 | WD repeat | Color2 [Da | 1 | 4 |
| | 0 | -0.15621 | Q9BV44 | Q9BV44 | THUMPD3 | tRNA (guar | Color2 [Da | 1 | 4 |
| + | 2.06934 | 0.266017 | Q9BV57 | Q9BV57 | ADI1 | Acireducto | Color2 [Da | 1 | 4 |
| + | 0.865389 | 0.635449 | Q9BV73 | Q9BV73 | CEP250 | Centrosom | Color2 [Da | 1 | 4 |
| | 0.346823 | -0.14013 | Q9BV79 | Q9BV79 | MECR | Enoyl-[acy | Color2 [Da | 1 | 4 |
| | 0.597044 | 0.563727 | Q9BV86 | Q9BV86 | NTMT1 | N-termina | Color2 [Da | 1 | 4 |
| + | 1.048346 | 0.698585 | Q9BVA0 | Q9BVA0 | KATNB1 | Katanin p8 | Color2 [Da | 1 | 4 |
| | 0.018386 | -0.06566 | Q9BVA1 | Q9BVA1 | TUBB2B | Tubulin be | Color2 [Da | 1 | 4 |
| + | 1.335077 | 0.256767 | Q9BVC4 | Q9BVC4 | MLST8 | Target of r | Color2 [Da | 1 | 4 |
| | 0.037955 | -0.04902 | Q9BVC5 | Q9BVC5 | C2orf49 | Ashwin | Color2 [Da | 1 | 4 |

| | | | | | | | | | |
|---|---|---|---|---|---|---|---|---|---|
| | 0.014036 | -0.02286 | Q9BVC6 | Q9BVC6 | TMEM109 | Transmem | Color2 [Da | 1 | 4 |
| + | 3.238581 | 1.900589 | Q9BVG4 | Q9BVG4 | PBDC1 | Protein PB | Color2 [Da | 1 | 4 |
| + | 1.345986 | -1.94777 | Q9BVG9 | Q9BVG9 | PTDSS2 | Phosphatic | Color2 [Da | 1 | 4 |
| + | 1.082078 | -0.55748 | Q9BVH7 | Q9BVH7 | ST6GALNA | Alpha-N-ac | Color2 [Da | 1 | 4 |
| | 0 | -1.14389 | Q9BVI0 | Q9BVI0 | PHF20 | PHD finger | Color2 [Da | 1 | 4 |
| | 0.153356 | 0.088634 | Q9BVI4 | Q9BVI4 | NOC4L | Nucleolar ( | Color2 [Da | 1 | 4 |
| + | 2.060135 | -0.44098 | Q9BVJ6 | Q9BVJ6 | UTP14A | U3 small n | Color2 [Da | 1 | 4 |
| + | 0.907637 | -0.35793 | Q9BVK6 | Q9BVK6 | TMED9 | Transmem | Color2 [Da | 1 | 4 |
| + | 1.434515 | 0.742626 | Q9BVL2 | Q9BVL2 | NUP58 | Nucleopor | Color2 [Da | 1 | 4 |
| | 0 | NaN | Q9BVL4 | Q9BVL4 | SELENOO | Protein ad | Color2 [Da | 1 | 4 |
| + | 0.981568 | 0.541223 | Q9BVM2 | Q9BVM2 | DPCD | Protein DP | Color2 [Da | 1 | 4 |
| + | 1.241418 | 1.442196 | Q9BVM4 | Q9BVM4 | GGACT | Gamma-gl | Color2 [Da | 1 | 4 |
| + | 3.056343 | -1.14993 | Q9BVP2 | Q9BVP2 | GNL3 | Guanine n | Color2 [Da | 1 | 4 |
| | 0 | -1.15208 | Q9BVS5 | Q9BVS5 | TRMT61B | tRNA (ade | Color2 [Da | 1 | 4 |
| | 0.670898 | 0.239129 | Q9BVT8 | Q9BVT8 | TMUB1 | Transmem | Color2 [Da | 1 | 4 |
| | 0 | NaN | Q9BVV7 | Q9BVV7 | TIMM21 | Mitochonc | Color2 [Da | 1 | 4 |
| | 0 | -1.37237 | Q9BW19 | Q9BW19 | KIFC1 | Kinesin-lik | Color2 [Da | 1 | 4 |
| + | 3.677186 | -1.63857 | Q9BW27 | Q9BW27 | NUP85 | Nuclear pc | Color2 [Da | 1 | 4 |
| + | 3.065774 | 0.902845 | Q9BW30 | Q9BW30 | TPPP3 | Tubulin po | Color2 [Da | 1 | 4 |
| + | 1.573789 | 0.393221 | Q9BW61 | Q9BW61 | DDA1 | DET1- and | Color2 [Da | 1 | 4 |
| + | 1.027265 | -0.34352 | Q9BW62 | Q9BW62 | KATNAL1 | Katanin p6 | Color2 [Da | 1 | 4 |
| | 0.155381 | -0.30417 | Q9BW71 | Q9BW71 | HIRIP3 | HIRA-inter | Color2 [Da | 1 | 4 |
| + | 4.038566 | 5.173201 | Q9BW72 | Q9BW72 | HIGD2A | HIG1 doma | Color2 [Da | 1 | 4 |
| | 0.406555 | 0.655064 | Q9BW83 | Q9BW83 | IFT27 | Intraflagell | Color2 [Da | 1 | 4 |
| + | 2.932283 | 1.227968 | Q9BW85 | Q9BW85 | YJU2 | Splicing fac | Color2 [Da | 1 | 4 |
| + | 2.134072 | 0.627647 | Q9BW91 | Q9BW91 | NUDT9 | ADP-ribose | Color2 [Da | 1 | 4 |
| | 0 | -0.77473 | Q9BW92 | Q9BW92 | TARS2 | Threonine- | Color2 [Da | 1 | 4 |
| + | 0.790523 | 0.488588 | Q9BWD1 | Q9BWD1 | ACAT2 | Acetyl-CoA | Color2 [Da | 1 | 4 |
| + | 2.434169 | -1.28705 | Q9BWF3 | Q9BWF3 | RBM4 | RNA-bindi | Color2 [Da | 1 | 4 |
| | 0 | 0.888905 | Q9BWG6 | Q9BWG6 | SCNM1 | Sodium ch | Color2 [Da | 1 | 4 |
| + | 3.955694 | 1.501645 | Q9BWH2 | Q9BWH2 | FUNDC2 | FUN14 dor | Color2 [Da | 1 | 4 |
| + | 3.660214 | -0.43328 | Q9BWH6 | Q9BWH6 | RPAP1 | RNA polym | Color2 [Da | 1 | 4 |
| | 0.326049 | -0.08693 | Q9BWJ5 | Q9BWJ5 | SF3B5 | Splicing fac | Color2 [Da | 1 | 4 |
| | 1.004875 | 0.167772 | Q9BWM7 | Q9BWM7 | SFXN3 | Sideroflexi | Color2 [Da | 1 | 4 |
| | 1.032363 | -0.15148 | Q9BWN1 | Q9BWN1 | PRR14 | Proline-ric | Color2 [Da | 1 | 4 |
| + | 1.807762 | -0.38649 | Q9BWS9 | Q9BWS9 | CHID1 | Chitinase c | Color2 [Da | 1 | 4 |
| | 0 | -1.05955 | Q9BWT3 | Q9BWT3 | PAPOLG | Poly(A) po | Color2 [Da | 1 | 4 |
| | 0.317078 | -0.07908 | Q9BWU0 | Q9BWU0 | SLC4A1AP | Kanadaptir | Color2 [Da | 1 | 4 |
| | 0 | NaN | Q9BWV1 | Q9BWV1 | BOC | Brother of | Color2 [Da | 1 | 4 |
| | 0.43575 | -0.30949 | Q9BWW4 | Q9BWW4 | SSBP3 | Single-stra | Color2 [Da | 1 | 4 |
| + | 2.799549 | 1.178262 | Q9BX40 | Q9BX40 | LSM14B | Protein LSI | Color2 [Da | 1 | 4 |
| + | 2.457598 | 1.214074 | Q9BX66 | Q9BX66 | SORBS1 | Sorbin and | Color2 [Da | 1 | 4 |
| + | 1.137014 | 0.43128 | Q9BX67 | Q9BX67 | JAM3 | Junctional | Color2 [Da | 1 | 4 |
| + | 2.598481 | 1.732663 | Q9BX68 | Q9BX68 | HINT2 | Adenosine | Color2 [Da | 1 | 4 |
| | 0 | NaN | Q9BX70 | Q9BX70 | BTBD2 | BTB/POZ d | Color2 [Da | 1 | 4 |
| | 0 | -0.25625 | Q9BX74 | Q9BX74 | TM2D1 | TM2 doma | Color2 [Da | 1 | 4 |
| + | 1.509235 | -0.98574 | Q9BXA9 | Q9BXA9 | SALL3 | Sal-like prc | Color2 [Da | 1 | 4 |

| | | | | | | | | | |
|---|---|---|---|---|---|---|---|---|---|
| + | 2.770002 | -1.4582 | Q9BXB4 | Q9BXB4 | OSBPL11 | Oxysterol- | Color2 [Da | 1 | 4 |
| + | 1.245596 | -0.97657 | Q9BXC9 | Q9BXC9 | BBS2 | Bardet-Bie | Color2 [Da | 1 | 4 |
| | 0 | -3.28277 | Q9BXF3 | Q9BXF3 | CECR2 | Cat eye syı | Color2 [Da | 1 | 4 |
| + | 0.906668 | 0.921114 | Q9BXF6 | Q9BXF6 | RAB11FIP5 | Rab11 fam | Color2 [Da | 1 | 4 |
| | 0 | 1.734581 | Q9BXH1 | Q9BXH1 | BBC3 | Bcl-2-bindi | Color2 [Da | 1 | 4 |
| + | 2.238889 | 0.904308 | Q9BXJ0 | Q9BXJ0 | C1QTNF5 | Compleme | Color2 [Da | 1 | 4 |
| + | 1.970133 | 1.791259 | Q9BXJ3 | Q9BXJ3 | C1QTNF4 | Compleme | Color2 [Da | 1 | 4 |
| | 0 | NaN | Q9BXJ8 | Q9BXJ8 | TMEM120 | Ion channe | Color2 [Da | 1 | 4 |
| + | 2.049059 | -1.58543 | Q9BXJ9 | Q9BXJ9 | NAA15 | N-alpha-ac | Color2 [Da | 1 | 4 |
| + | 4.158306 | 1.560671 | Q9BXK5 | Q9BXK5 | BCL2L13 | Bcl-2-like p | Color2 [Da | 1 | 4 |
| + | 1.321708 | -0.85862 | Q9BXM9 | Q9BXM9 | FSD1L | FSD1-like p | Color2 [Da | 1 | 4 |
| | 0.444591 | -0.35316 | Q9BXP2 | Q9BXP2 | SLC12A9 | Solute carı | Color2 [Da | 1 | 4 |
| | 1.398728 | 0.172053 | Q9BXP5 | Q9BXP5 | SRRT | Serrate RN | Color2 [Da | 1 | 4 |
| + | 1.645529 | -1.16277 | Q9BXR0 | Q9BXR0 | QTRT1 | Queuine tI | Color2 [Da | 1 | 4 |
| + | 2.734825 | -1.23295 | Q9BXS5 | Q9BXS5 | AP1M1 | AP-1 comp | Color2 [Da | 1 | 4 |
| + | 2.480962 | 2.555519 | Q9BXV9 | Q9BXV9 | GON7 | EKC/KEOP: | Color2 [Da | 1 | 4 |
| + | 2.900605 | -1.40856 | Q9BXW6 | Q9BXW6 | OSBPL1A | Oxysterol- | Color2 [Da | 1 | 4 |
| | 0.403971 | -0.07059 | Q9BXW7 | Q9BXW7 | HDHD5 | Haloacid d | Color2 [Da | 1 | 4 |
| + | 1.016897 | -0.63319 | Q9BXY0 | Q9BXY0 | MAK16 | Protein M/ | Color2 [Da | 1 | 4 |
| + | 0.813893 | 1.226712 | Q9BY11 | Q9BY11 | PACSIN1 | Protein kir | Color2 [Da | 1 | 4 |
| | 0.981625 | -0.30212 | Q9BY12 | Q9BY12 | SCAPER | S phase cy | Color2 [Da | 1 | 4 |
| + | 1.152073 | -0.44054 | Q9BY32 | Q9BY32 | ITPA | Inosine tri | Color2 [Da | 1 | 4 |
| + | 0.872992 | -0.37701 | Q9BY42 | Q9BY42 | RTF2 | Replicatior | Color2 [Da | 1 | 4 |
| + | 2.606876 | 1.330255 | Q9BY43 | Q9BY43 | CHMP4A | Charged m | Color2 [Da | 1 | 4 |
| + | 1.197407 | -0.53427 | Q9BY44 | Q9BY44 | EIF2A | Eukaryotic | Color2 [Da | 1 | 4 |
| | 0 | -0.07092 | Q9BY49 | Q9BY49 | PECR | Peroxisom | Color2 [Da | 1 | 4 |
| + | 4.591978 | 3.544442 | Q9BY50 | Q9BY50 | SEC11C | Signal pep | Color2 [Da | 1 | 4 |
| | 0.60701 | 0.300438 | Q9BY67 | Q9BY67 | CADM1 | Cell adhesi | Color2 [Da | 1 | 4 |
| | 0.35959 | 0.415487 | Q9BY76 | Q9BY76 | ANGPTL4 | Angiopoiet | Color2 [Da | 1 | 4 |
| + | 1.89881 | 0.389885 | Q9BY77 | Q9BY77 | POLDIP3 | Polymeras | Color2 [Da | 1 | 4 |
| | 0.608494 | -0.9509 | Q9BYC5 | Q9BYC5 | FUT8 | Alpha-(1,6 | Color2 [Da | 1 | 4 |
| | 0 | -1.32727 | Q9BYC9 | Q9BYC9 | MRPL20 | 39S riboso | Color2 [Da | 1 | 4 |
| + | 2.638577 | 0.893638 | Q9BYD1 | Q9BYD1 | MRPL13 | 39S riboso | Color2 [Da | 1 | 4 |
| + | 2.251696 | 0.724726 | Q9BYD2 | Q9BYD2 | MRPL9 | 39S riboso | Color2 [Da | 1 | 4 |
| + | 2.913252 | 1.163774 | Q9BYD3 | Q9BYD3 | MRPL4 | 39S riboso | Color2 [Da | 1 | 4 |
| + | 3.1211 | 2.100579 | Q9BYD6 | Q9BYD6 | MRPL1 | 39S riboso | Color2 [Da | 1 | 4 |
| + | 1.292042 | -0.62186 | Q9BYG3 | Q9BYG3 | NIFK | MKI67 FH/ | Color2 [Da | 1 | 4 |
| + | 1.984247 | 2.743322 | Q9BYG4 | Q9BYG4 | PARD6G | Partitionin | Color2 [Da | 1 | 4 |
| + | 1.023756 | 0.451246 | Q9BYG5 | Q9BYG5 | PARD6B | Partitionin | Color2 [Da | 1 | 4 |
| + | 0.877607 | -0.38809 | Q9BYJ9 | Q9BYJ9 | YTHDF1 | YTH doma | Color2 [Da | 1 | 4 |
| | 0 | NaN | Q9BYK8 | Q9BYK8 | HELZ2 | Helicase w | Color2 [Da | 1 | 4 |
| | 0.160537 | -0.1814 | Q9BYM8 | Q9BYM8 | RBCK1 | RanBP-typ | Color2 [Da | 1 | 4 |
| + | 2.245589 | 1.747832 | Q9BYN0 | Q9BYN0 | SRXN1 | Sulfiredoxi | Color2 [Da | 1 | 4 |
| + | 1.899265 | 0.76769 | Q9BYN8 | Q9BYN8 | MRPS26 | 28S riboso | Color2 [Da | 1 | 4 |
| | 0.646015 | -0.14187 | Q9BYP7 | Q9BYP7 | WNK3 | Serine/thr | Color2 [Da | 1 | 4 |
| + | 2.057209 | -0.90988 | Q9BYT8 | Q9BYT8 | NLN | Neurolysin | Color2 [Da | 1 | 4 |
| + | 2.711667 | 0.774442 | Q9BYV8 | Q9BYV8 | CEP41 | Centrosom | Color2 [Da | 1 | 4 |

| | | | | | | | | | |
|---|---|---|---|---|---|---|---|---|---|
| + | 0.78687 | -1.05777 | Q9BYW2 | Q9BYW2 | SETD2 | Histone-ly: | Color2 [Da | 1 | 4 |
| | 0 | -1.22079 | Q9BZ23;Q | Q9BZ23;Q | PANK2;PAI | Pantothen | Color2 [Da | 1 | 4 |
| | 0 | NaN | Q9BZ71 | Q9BZ71 | PITPNM3 | Membrane | Color2 [Da | 1 | 4 |
| | 0.469694 | 0.299091 | Q9BZ95 | Q9BZ95 | NSD3 | Histone-ly: | Color2 [Da | 1 | 4 |
| | 0 | -1.25966 | Q9BZA5;Q | Q9BZA5;Q | TXLNGY;T> | Putative g | Color2 [Da | 1 | 4 |
| + | 2.60324 | 0.955056 | Q9BZC1 | Q9BZC1 | CELF4 | CUGBP Ela | Color2 [Da | 1 | 4 |
| | 0 | -0.7463 | Q9BZD4 | Q9BZD4 | NUF2 | Kinetochoi | Color2 [Da | 1 | 4 |
| | 0.276104 | -0.29593 | Q9BZE1 | Q9BZE1 | MRPL37 | 39S riboso | Color2 [Da | 1 | 4 |
| + | 3.675262 | -1.28082 | Q9BZE4 | Q9BZE4 | GTPBP4 | GTP-bindir | Color2 [Da | 1 | 4 |
| | 0.325844 | 0.34906 | Q9BZE9 | Q9BZE9 | ASPSCR1 | Tether cor | Color2 [Da | 1 | 4 |
| + | 3.017739 | -1.20491 | Q9BZF1 | Q9BZF1 | OSBPL8 | Oxysterol- | Color2 [Da | 1 | 4 |
| | 0.763467 | 0.331961 | Q9BZF3 | Q9BZF3 | OSBPL6 | Oxysterol- | Color2 [Da | 1 | 4 |
| | 0 | -0.43968 | Q9BZF9 | Q9BZF9 | UACA | Uveal autc | Color2 [Da | 1 | 4 |
| | 0 | NaN | Q9BZG1 | Q9BZG1 | RAB34 | Ras-relate | Color2 [Da | 1 | 4 |
| + | 2.974406 | 0.83434 | Q9BZH6 | Q9BZH6 | WDR11 | WD repeat | Color2 [Da | 1 | 4 |
| + | 1.765117 | 0.302691 | Q9BZI7 | Q9BZI7 | UPF3B | Regulator | Color2 [Da | 1 | 4 |
| + | 3.342518 | -0.90776 | Q9BZJ0 | Q9BZJ0 | CRNKL1 | Crooked n | Color2 [Da | 1 | 4 |
| | 0.523111 | 0.398624 | Q9BZK7 | Q9BZK7 | TBL1XR1 | F-box-like/ | Color2 [Da | 1 | 4 |
| + | 1.551868 | 0.725839 | Q9BZL4 | Q9BZL4 | PPP1R12C | Protein ph | Color2 [Da | 1 | 4 |
| | 0 | -2.27217 | Q9BZL6 | Q9BZL6 | PRKD2 | Serine/thr | Color2 [Da | 1 | 4 |
| | 0 | -1.41624 | Q9BZQ6 | Q9BZQ6 | EDEM3 | ER degrada | Color2 [Da | 1 | 4 |
| | 0 | -1.66929 | Q9BZQ8 | Q9BZQ8 | NIBAN1 | Protein Nil | Color2 [Da | 1 | 4 |
| + | 2.485036 | 0.160837 | Q9BZV1 | Q9BZV1 | UBXN6 | UBX doma | Color2 [Da | 1 | 4 |
| | 0.200401 | 0.3859 | Q9BZX2 | Q9BZX2 | UCK2 | Uridine-cy | Color2 [Da | 1 | 4 |
| + | 3.830124 | -1.66059 | Q9BZZ5 | Q9BZZ5 | API5 | Apoptosis | Color2 [Da | 1 | 4 |
| + | 2.107023 | 2.512033 | Q9C005 | Q9C005 | DPY30 | Protein dp | Color2 [Da | 1 | 4 |
| | 0 | NaN | Q9C010 | Q9C010 | PKIB | cAMP-dep | Color2 [Da | 1 | 4 |
| | 0.617098 | 0.5181 | Q9C026 | Q9C026 | TRIM9 | E3 ubiquiti | Color2 [Da | 1 | 4 |
| | 0 | 0.083725 | Q9C035 | Q9C035 | TRIM5 | Tripartite r | Color2 [Da | 1 | 4 |
| | 0.367902 | -0.07338 | Q9C040 | Q9C040 | TRIM2 | Tripartite r | Color2 [Da | 1 | 4 |
| | 0 | 1.563903 | Q9C075 | Q9C075 | KRT23 | Keratin, ty | Color2 [Da | 1 | 4 |
| + | 1.141647 | 0.383521 | Q9C0B0 | Q9C0B0 | UNK | RING finge | Color2 [Da | 1 | 4 |
| | 0.108955 | -0.09186 | Q9C0B1 | Q9C0B1 | FTO | Alpha-ketc | Color2 [Da | 1 | 4 |
| + | 2.520654 | 1.415476 | Q9C0B5 | Q9C0B5 | ZDHHC5 | Palmitoylt | Color2 [Da | 1 | 4 |
| + | 1.973429 | 0.930411 | Q9C0C2 | Q9C0C2 | TNKS1BP1 | 182 kDa ta | Color2 [Da | 1 | 4 |
| | 0 | 0.718767 | Q9C0C6 | Q9C0C6 | CIPC | CLOCK-inte | Color2 [Da | 1 | 4 |
| | 0.513004 | 0.195796 | Q9C0C7 | Q9C0C7 | AMBRA1 | Activating | Color2 [Da | 1 | 4 |
| + | 1.125289 | 0.274966 | Q9C0C9 | Q9C0C9 | UBE2O | (E3-indepe | Color2 [Da | 1 | 4 |
| + | 1.949488 | 0.672561 | Q9C0D0 | Q9C0D0 | PHACTR1 | Phosphata | Color2 [Da | 1 | 4 |
| | 0 | 0.21224 | Q9C0D3 | Q9C0D3 | ZYG11B | Protein zyg | Color2 [Da | 1 | 4 |
| + | 1.651566 | -1.09855 | Q9C0D9 | Q9C0D9 | SELENOI | Ethanolam | Color2 [Da | 1 | 4 |
| + | 0.937302 | -1.00933 | Q9C0E2 | Q9C0E2 | XPO4 | Exportin-4 | Color2 [Da | 1 | 4 |
| | 0.40805 | 0.279654 | Q9C0E8 | Q9C0E8 | LNPK | Endoplasm | Color2 [Da | 1 | 4 |
| + | 2.126643 | 3.593498 | Q9C0F1 | Q9C0F1 | CEP44 | Centrosom | Color2 [Da | 1 | 4 |
| | 0.247347 | -0.20396 | Q9C0H2 | Q9C0H2 | TTYH3 | Protein tw | Color2 [Da | 1 | 4 |
| | 0.532812 | -0.15871 | Q9C0H5 | Q9C0H5 | ARHGAP39 | Rho GTPas | Color2 [Da | 1 | 4 |
| + | 4.049665 | 1.42744 | Q9C0H9 | Q9C0H9 | SRCIN1 | SRC kinase | Color2 [Da | 1 | 4 |

| | | | | | | | | | |
|---|---|---|---|---|---|---|---|---|---|
| | 0 | -1.01776 | Q9C0I1 | Q9C0I1 | MTMR12 | Myotubula | Color2 [Da | 1 | 4 |
| + | 0.732759 | 1.246493 | Q9C0J8 | Q9C0J8 | WDR33 | pre-mRNA | Color2 [Da | 1 | 4 |
| | 0 | NaN | Q9C0K0 | Q9C0K0 | BCL11B | B-cell lymp | Color2 [Da | 1 | 4 |
| + | 2.031016 | -1.5199 | Q9GZL7 | Q9GZL7 | WDR12 | Ribosome | Color2 [Da | 1 | 4 |
| + | 3.568898 | 1.388261 | Q9GZM8 | Q9GZM8 | NDEL1 | Nuclear di: | Color2 [Da | 1 | 4 |
| | 0.780714 | -0.39463 | Q9GZN7 | Q9GZN7 | ROGDI | Protein ro | Color2 [Da | 1 | 4 |
| + | 0.844751 | 0.787165 | Q9GZN8 | Q9GZN8 | C20orf27 | UPF0687 p | Color2 [Da | 1 | 4 |
| + | 3.911052 | 2.207387 | Q9GZP4 | Q9GZP4 | PITHD1 | PITH doma | Color2 [Da | 1 | 4 |
| | 0 | -0.994 | Q9GZP9 | Q9GZP9 | DERL2 | Derlin-2 | Color2 [Da | 1 | 4 |
| + | 1.704862 | 1.509117 | Q9GZR2 | Q9GZR2 | REXO4 | RNA exonu | Color2 [Da | 1 | 4 |
| + | 1.310219 | -0.50246 | Q9GZR7 | Q9GZR7 | DDX24 | ATP-deper | Color2 [Da | 1 | 4 |
| + | 1.377379 | 0.730594 | Q9GZS3 | Q9GZS3 | WDR61 | WD repeat | Color2 [Da | 1 | 4 |
| + | 2.125129 | 0.895211 | Q9GZT3 | Q9GZT3 | SLIRP | SRA stem-l | Color2 [Da | 1 | 4 |
| | 0.392 | -0.11697 | Q9GZT4 | Q9GZT4 | SRR | Serine race | Color2 [Da | 1 | 4 |
| + | 2.618853 | 1.018758 | Q9GZT6 | Q9GZT6 | CCDC90B | Coiled-coil | Color2 [Da | 1 | 4 |
| + | 1.079476 | 0.808003 | Q9GZT8 | Q9GZT8 | NIF3L1 | NIF3-like p | Color2 [Da | 1 | 4 |
| | 0.100868 | 0.040266 | Q9GZT9 | Q9GZT9 | EGLN1 | Egl nine hc | Color2 [Da | 1 | 4 |
| | 0 | NaN | Q9GZU8 | Q9GZU8 | PSME3IP1 | PSME3-int | Color2 [Da | 1 | 4 |
| + | 1.508399 | 1.892552 | Q9GZX3 | Q9GZX3 | CHST6 | Carbohydr | Color2 [Da | 1 | 4 |
| | 0 | -0.3905 | Q9GZX9 | Q9GZX9 | TWSG1 | Twisted ga | Color2 [Da | 1 | 4 |
| | 0.151434 | 0.136908 | Q9GZY4 | Q9GZY4 | COA1 | Cytochron | Color2 [Da | 1 | 4 |
| + | 2.115128 | -0.2462 | Q9GZY8 | Q9GZY8 | MFF | Mitochonc | Color2 [Da | 1 | 4 |
| | 0.463044 | -0.1749 | Q9GZZ1 | Q9GZZ1 | NAA50 | N-alpha-ac | Color2 [Da | 1 | 4 |
| | 0 | NaN | Q9GZZ8 | Q9GZZ8 | LACRT | Extracellul | Color2 [Da | 1 | 4 |
| | 0.822752 | -0.13823 | Q9GZZ9 | Q9GZZ9 | UBA5 | Ubiquitin-l | Color2 [Da | 1 | 4 |
| | 0.49759 | 0.651828 | Q9H000 | Q9H000 | MKRN2 | E3 ubiquiti | Color2 [Da | 1 | 4 |
| + | 1.215321 | 2.028363 | Q9H008 | Q9H008 | LHPP | Phospholy | Color2 [Da | 1 | 4 |
| + | 2.712905 | 0.881064 | Q9H019 | Q9H019 | MTFR1L | Mitochonc | Color2 [Da | 1 | 4 |
| + | 2.43149 | 0.925206 | Q9H061 | Q9H061 | TMEM126 | Transmem | Color2 [Da | 1 | 4 |
| + | 1.014476 | 0.527375 | Q9H074 | Q9H074 | PAIP1 | Polyadeny | Color2 [Da | 1 | 4 |
| | 1.61004 | 0.178138 | Q9H078 | Q9H078 | CLPB | Caseinolyt | Color2 [Da | 1 | 4 |
| + | 0.857927 | 1.082789 | Q9H082 | Q9H082 | RAB33B | Ras-relate | Color2 [Da | 1 | 4 |
| + | 3.746402 | 3.491888 | Q9H098 | Q9H098 | FAM107B | Protein FA | Color2 [Da | 1 | 4 |
| + | 1.1455 | -0.86206 | Q9H0A0 | Q9H0A0 | NAT10 | RNA cytidi | Color2 [Da | 1 | 4 |
| + | 1.676914 | 0.926837 | Q9H0A8 | Q9H0A8 | COMMD4 | COMM do | Color2 [Da | 1 | 4 |
| + | 1.124633 | -0.36799 | Q9H0B6 | Q9H0B6 | KLC2 | Kinesin ligl | Color2 [Da | 1 | 4 |
| + | 2.913492 | -1.02057 | Q9H0C8 | Q9H0C8 | ILKAP | Integrin-lir | Color2 [Da | 1 | 4 |
| + | 2.386854 | -0.51757 | Q9H0D6 | Q9H0D6 | XRN2 | 5'-3' exoril | Color2 [Da | 1 | 4 |
| + | 2.288249 | 1.932604 | Q9H0E2 | Q9H0E2 | TOLLIP | Toll-intera | Color2 [Da | 1 | 4 |
| + | 0.798522 | 1.048423 | Q9H0E3 | Q9H0E3 | SAP130 | Histone de | Color2 [Da | 1 | 4 |
| | 0.608907 | -0.35062 | Q9H0F7 | Q9H0F7 | ARL6 | ADP-ribosy | Color2 [Da | 1 | 4 |
| | 0 | NaN | Q9H0H3 | Q9H0H3 | KLHL25 | Kelch-like | Color2 [Da | 1 | 4 |
| | 0.227842 | 0.104441 | Q9H0H5 | Q9H0H5 | RACGAP1 | Rac GTPas | Color2 [Da | 1 | 4 |
| | 0.490706 | 0.294005 | Q9H0L4 | Q9H0L4 | CSTF2T | Cleavage s | Color2 [Da | 1 | 4 |
| + | 2.418898 | 1.375682 | Q9H0N5 | Q9H0N5 | PCBD2 | Pterin-4-al | Color2 [Da | 1 | 4 |
| + | 1.304264 | 0.65747 | Q9H0P0 | Q9H0P0 | NT5C3A | Cytosolic 5 | Color2 [Da | 1 | 4 |
| + | 3.370983 | -1.84942 | Q9H0Q0 | Q9H0Q0 | CYRIA | CYFIP-relat | Color2 [Da | 1 | 4 |

| | | | | | | | | | |
|---|---|---|---|---|---|---|---|---|---|
| + | 1.475633 | 0.856896 | Q9H0Q3 | Q9H0Q3 | FXYD6 | FXYD dom | Color2 [Da | 1 | 4 |
| | 0 | 0.574448 | Q9H0R3 | Q9H0R3 | TMEM222 | Transmem | Color2 [Da | 1 | 4 |
| + | 2.239014 | 1.146224 | Q9H0R4 | Q9H0R4 | HDHD2 | Haloacid d | Color2 [Da | 1 | 4 |
| | 0.388223 | 0.109967 | Q9H0S4 | Q9H0S4 | DDX47 | Probable A | Color2 [Da | 1 | 4 |
| + | 1.174468 | -0.69783 | Q9H0U3 | Q9H0U3 | MAGT1 | Magnesiur | Color2 [Da | 1 | 4 |
| | 0.380724 | 0.107805 | Q9H0U4 | Q9H0U4 | RAB1B | Ras-relate | Color2 [Da | 1 | 4 |
| + | 2.223798 | 2.479625 | Q9H0U6 | Q9H0U6 | MRPL18 | 39S riboso | Color2 [Da | 1 | 4 |
| + | 1.333537 | -0.81477 | Q9H0V9 | Q9H0V9 | LMAN2L | VIP36-like | Color2 [Da | 1 | 4 |
| | 0.105323 | 0.075252 | Q9H0W8 | Q9H0W8 | SMG9 | Nonsense- | Color2 [Da | 1 | 4 |
| + | 1.361738 | 0.970191 | Q9H0W9 | Q9H0W9 | C11orf54 | Ester hydr | Color2 [Da | 1 | 4 |
| | 0 | 0.411017 | Q9H0X4 | Q9H0X4 | FAM234A | Protein FA | Color2 [Da | 1 | 4 |
| + | 1.338291 | -1.00918 | Q9H0X9 | Q9H0X9 | OSBPL5 | Oxysterol- | Color2 [Da | 1 | 4 |
| + | 1.49644 | 0.201993 | Q9H115 | Q9H115 | NAPB | Beta-solub | Color2 [Da | 1 | 4 |
| + | 1.172912 | 1.409012 | Q9H147 | Q9H147 | DNTTIP1 | Deoxynucl | Color2 [Da | 1 | 4 |
| + | 3.672783 | 1.352612 | Q9H169 | Q9H169 | STMN4 | Stathmin-4 | Color2 [Da | 1 | 4 |
| + | 1.112698 | -0.74654 | Q9H173 | Q9H173 | SIL1 | Nucleotide | Color2 [Da | 1 | 4 |
| | 0.135366 | 0.07559 | Q9H1A3 | Q9H1A3 | METTL9 | Protein-L-I | Color2 [Da | 1 | 4 |
| | 0 | NaN | Q9H1A4 | Q9H1A4 | ANAPC1 | Anaphase- | Color2 [Da | 1 | 4 |
| + | 1.783126 | 0.380023 | Q9H1B7 | Q9H1B7 | IRF2BPL | Probable E | Color2 [Da | 1 | 4 |
| | 0 | 0.382098 | Q9H1E1 | Q9H1E1 | RNASE7 | Ribonuclea | Color2 [Da | 1 | 4 |
| + | 1.793566 | 1.293125 | Q9H1E3 | Q9H1E3 | NUCKS1 | Nuclear ub | Color2 [Da | 1 | 4 |
| + | 2.746979 | -1.24651 | Q9H1E5 | Q9H1E5 | TMX4 | Thioredoxi | Color2 [Da | 1 | 4 |
| | 0 | -1.43542 | Q9H1H9 | Q9H1H9 | KIF13A | Kinesin-lik | Color2 [Da | 1 | 4 |
| | 0 | -0.24608 | Q9H1I8 | Q9H1I8 | ASCC2 | Activating | Color2 [Da | 1 | 4 |
| | 0.950152 | 0.201106 | Q9H1K0 | Q9H1K0 | RBSN | Rabenosyr | Color2 [Da | 1 | 4 |
| | 0.627426 | 0.551692 | Q9H1K1 | Q9H1K1 | ISCU | Iron-sulfur | Color2 [Da | 1 | 4 |
| + | 1.30016 | -0.43529 | Q9H1P3 | Q9H1P3 | OSBPL2 | Oxysterol- | Color2 [Da | 1 | 4 |
| | 0 | -0.58111 | Q9H1Y0 | Q9H1Y0 | ATG5 | Autophagy | Color2 [Da | 1 | 4 |
| | 0.20313 | -0.18903 | Q9H1Z4 | Q9H1Z4 | WDR13 | WD repeat | Color2 [Da | 1 | 4 |
| + | 2.271944 | 1.265025 | Q9H204 | Q9H204 | MED28 | Mediator c | Color2 [Da | 1 | 4 |
| | 0.678248 | -0.173 | Q9H223 | Q9H223 | EHD4 | EH domair | Color2 [Da | 1 | 4 |
| | 0 | NaN | Q9H227 | Q9H227 | GBA3 | Cytosolic b | Color2 [Da | 1 | 4 |
| + | 0.77781 | 0.550992 | Q9H254 | Q9H254 | SPTBN4 | Spectrin be | Color2 [Da | 1 | 4 |
| + | 0.969763 | -0.63852 | Q9H267 | Q9H267 | VPS33B | Vacuolar p | Color2 [Da | 1 | 4 |
| + | 2.281807 | -0.76429 | Q9H269 | Q9H269 | VPS16 | Vacuolar p | Color2 [Da | 1 | 4 |
| + | 2.476167 | -0.97166 | Q9H270 | Q9H270 | VPS11 | Vacuolar p | Color2 [Da | 1 | 4 |
| + | 2.517968 | 2.589817 | Q9H299 | Q9H299 | SH3BGRL3 | SH3 domai | Color2 [Da | 1 | 4 |
| + | 2.778014 | -0.65096 | Q9H2C0 | Q9H2C0 | GAN | Gigaxonin | Color2 [Da | 1 | 4 |
| + | 1.33648 | -0.85833 | Q9H2D6 | Q9H2D6 | TRIOBP | TRIO and F | Color2 [Da | 1 | 4 |
| | 0.287099 | 0.243413 | Q9H2E6 | Q9H2E6 | SEMA6A | Semaphor | Color2 [Da | 1 | 4 |
| + | 1.379936 | 0.364488 | Q9H2G2 | Q9H2G2 | SLK | STE20-like | Color2 [Da | 1 | 4 |
| | 0 | 0.217964 | Q9H2G4 | Q9H2G4 | TSPYL2 | Testis-spec | Color2 [Da | 1 | 4 |
| + | 2.019915 | 0.690725 | Q9H2H8 | Q9H2H8 | PPIL3 | Peptidyl-pr | Color2 [Da | 1 | 4 |
| + | 2.035762 | 0.478399 | Q9H2J4 | Q9H2J4 | PDCL3 | Phosducin | Color2 [Da | 1 | 4 |
| + | 1.271022 | 0.832394 | Q9H2K0 | Q9H2K0 | MTIF3 | Translatior | Color2 [Da | 1 | 4 |
| | 0 | NaN | Q9H2K8 | Q9H2K8 | TAOK3 | Serine/thr | Color2 [Da | 1 | 4 |
| | 0 | -3.63948 | Q9H2M3 | Q9H2M3 | BHMT2 | S-methylm | Color2 [Da | 1 | 4 |

| | | | | | | | | | | |
|---|---|---|---|---|---|---|---|---|---|---|
| + | 2.740592 | -1.22031 | Q9H2M9 | Q9H2M9 | RAB3GAP2 | Rab3 GTPa | Color2 | [Da | 1 | 4 |
| | 0.180789 | -0.05601 | Q9H2P0 | Q9H2P0 | ADNP | Activity-de | Color2 | [Da | 1 | 4 |
| | 1.300602 | 0.222012 | Q9H2P9 | Q9H2P9 | DPH5 | Diphthine | Color2 | [Da | 1 | 4 |
| + | 2.41231 | -2.31236 | Q9H2U1 | Q9H2U1 | DHX36 | ATP-deper | Color2 | [Da | 1 | 4 |
| | 0.934578 | -0.24955 | Q9H2U2 | Q9H2U2 | PPA2 | Inorganic p | Color2 | [Da | 1 | 4 |
| + | 0.997493 | -2.31035 | Q9H2V7 | Q9H2V7 | SPNS1 | Protein spi | Color2 | [Da | 1 | 4 |
| + | 1.539463 | 0.725407 | Q9H2W6 | Q9H2W6 | MRPL46 | 39S riboso | Color2 | [Da | 1 | 4 |
| | 0 | NaN | Q9H2X8 | Q9H2X8 | IFI27L2 | Interferon | Color2 | [Da | 1 | 4 |
| + | 2.365941 | 2.290624 | Q9H2X9 | Q9H2X9 | SLC12A5 | Solute carı | Color2 | [Da | 1 | 4 |
| + | 1.558731 | 3.383942 | Q9H305 | Q9H305 | CDIP1 | Cell death- | Color2 | [Da | 1 | 4 |
| + | 2.377436 | 0.616857 | Q9H307 | Q9H307 | PNN | Pinin | Color2 | [Da | 1 | 4 |
| | 0 | NaN | Q9H313 | Q9H313 | TTYH1 | Protein tw | Color2 | [Da | 1 | 4 |
| + | 0.747802 | -0.63138 | Q9H330 | Q9H330 | TMEM245 | Transmem | Color2 | [Da | 1 | 4 |
| | 0.146983 | -0.09661 | Q9H336 | Q9H336 | CRISPLD1 | Cysteine-ri | Color2 | [Da | 1 | 4 |
| | 0 | NaN | Q9H3F6 | Q9H3F6 | KCTD10 | BTB/POZ d | Color2 | [Da | 1 | 4 |
| + | 1.505631 | 0.707657 | Q9H3G5 | Q9H3G5 | CPVL | Probable s | Color2 | [Da | 1 | 4 |
| + | 2.385706 | 1.449077 | Q9H3H3 | Q9H3H3 | C11orf68 | UPF0696 p | Color2 | [Da | 1 | 4 |
| | 0 | NaN | Q9H3H5 | Q9H3H5 | DPAGT1 | UDP-N-ace | Color2 | [Da | 1 | 4 |
| + | 2.915429 | 1.599814 | Q9H3H9 | Q9H3H9 | TCEAL2 | Transcripti | Color2 | [Da | 1 | 4 |
| | 0 | -0.02507 | Q9H3K2 | Q9H3K2 | GHITM | Growth hc | Color2 | [Da | 1 | 4 |
| | 0.175574 | 0.097558 | Q9H3K6 | Q9H3K6 | BOLA2B | BolA-like p | Color2 | [Da | 1 | 4 |
| + | 4.602947 | -2.12424 | Q9H3M7 | Q9H3M7 | TXNIP | Thioredoxi | Color2 | [Da | 1 | 4 |
| + | 4.541448 | -1.97383 | Q9H3N1 | Q9H3N1 | TMX1 | Thioredoxi | Color2 | [Da | 1 | 4 |
| + | 1.477411 | 0.479145 | Q9H3P2 | Q9H3P2 | NELFA | Negative e | Color2 | [Da | 1 | 4 |
| | 0.682642 | -0.27574 | Q9H3P7 | Q9H3P7 | ACBD3 | Golgi resid | Color2 | [Da | 1 | 4 |
| + | 3.912809 | 1.517469 | Q9H3Q1 | Q9H3Q1 | CDC42EP4 | Cdc42 effe | Color2 | [Da | 1 | 4 |
| | 0.252966 | -0.50153 | Q9H3R5 | Q9H3R5 | CENPH | Centromeı | Color2 | [Da | 1 | 4 |
| | 0 | NaN | Q9H3S5 | Q9H3S5 | PIGM | GPI mannc | Color2 | [Da | 1 | 4 |
| + | 1.059143 | -0.37741 | Q9H3S7 | Q9H3S7 | PTPN23 | Tyrosine-p | Color2 | [Da | 1 | 4 |
| + | 1.441323 | -0.66233 | Q9H3U1 | Q9H3U1 | UNC45A | Protein un | Color2 | [Da | 1 | 4 |
| | 0 | 0.967726 | Q9H3Y8 | Q9H3Y8 | PPDPF | Pancreatic | Color2 | [Da | 1 | 4 |
| | 0.195398 | -0.27337 | Q9H3Z4 | Q9H3Z4 | DNAJC5 | DnaJ homc | Color2 | [Da | 1 | 4 |
| | 0 | NaN | Q9H410 | Q9H410 | DSN1 | Kinetochoı | Color2 | [Da | 1 | 4 |
| + | 3.108624 | 1.578121 | Q9H425 | Q9H425 | C1orf198 | Uncharact | Color2 | [Da | 1 | 4 |
| + | 4.581082 | 1.273543 | Q9H444 | Q9H444 | CHMP4B | Charged m | Color2 | [Da | 1 | 4 |
| + | 1.197526 | 1.036239 | Q9H446 | Q9H446 | RWDD1 | RWD dom | Color2 | [Da | 1 | 4 |
| | 0.134588 | -0.05031 | Q9H467 | Q9H467 | CUEDC2 | CUE doma | Color2 | [Da | 1 | 4 |
| | 0.633478 | -0.27271 | Q9H477 | Q9H477 | RBKS | Ribokinase | Color2 | [Da | 1 | 4 |
| + | 2.528476 | -0.40476 | Q9H479 | Q9H479 | FN3K | Fructosam | Color2 | [Da | 1 | 4 |
| + | 1.78317 | -0.93153 | Q9H488 | Q9H488 | POFUT1 | GDP-fucos | Color2 | [Da | 1 | 4 |
| | 0 | -2.60994 | Q9H490 | Q9H490 | PIGU | Phosphatic | Color2 | [Da | 1 | 4 |
| + | 2.782715 | 1.293034 | Q9H492 | Q9H492 | MAP1LC3A | Microtubu | Color2 | [Da | 1 | 4 |
| + | 1.462286 | 0.714382 | Q9H496 | Q9H496 | TOR1AIP2 | Torsin-1A- | Color2 | [Da | 1 | 4 |
| | 0 | NaN | Q9H497 | Q9H497 | TOR3A | Torsin-3A | Color2 | [Da | 1 | 4 |
| | 0.450316 | -0.24214 | Q9H4A3 | Q9H4A3 | WNK1 | Serine/thr | Color2 | [Da | 1 | 4 |
| + | 2.403071 | -0.89426 | Q9H4A4 | Q9H4A4 | RNPEP | Aminopep | Color2 | [Da | 1 | 4 |
| + | 0.818519 | 0.63503 | Q9H4A6 | Q9H4A6 | GOLPH3 | Golgi phos | Color2 | [Da | 1 | 4 |

| | | | | | | | | | |
|---|---|---|---|---|---|---|---|---|---|
| | 0 | -0.72414 | Q9H4F8 | Q9H4F8 | SMOC1 | SPARC-rela | Color2 [Da | 1 | 4 |
| + | 2.082585 | 0.758663 | Q9H4G0 | Q9H4G0 | EPB41L1 | Band 4.1-li | Color2 [Da | 1 | 4 |
| + | 2.72596 | 2.161899 | Q9H4G4 | Q9H4G4 | GLIPR2 | Golgi-asso | Color2 [Da | 1 | 4 |
| | 0 | 3.382837 | Q9H4I9 | Q9H4I9 | SMDT1 | Essential M | Color2 [Da | 1 | 4 |
| + | 1.796516 | -0.81281 | Q9H4L4 | Q9H4L4 | SENP3 | Sentrin-sp | Color2 [Da | 1 | 4 |
| | 0 | -0.83729 | Q9H4L5 | Q9H4L5 | OSBPL3 | Oxysterol- | Color2 [Da | 1 | 4 |
| | 0.378808 | 0.118388 | Q9H4L7 | Q9H4L7 | SMARCAD | SWI/SNF-r | Color2 [Da | 1 | 4 |
| | 0 | NaN | Q9H4M3 | Q9H4M3 | FBXO44 | F-box only | Color2 [Da | 1 | 4 |
| | 0.902349 | 0.133263 | Q9H4M9 | Q9H4M9 | EHD1 | EH domair | Color2 [Da | 1 | 4 |
| | 0.427703 | -0.34095 | Q9H4W6 | Q9H4W6 | EBF3 | Transcripti | Color2 [Da | 1 | 4 |
| + | 1.48652 | -0.70954 | Q9H4Z3 | Q9H4Z3 | PCIF1 | mRNA (2'- | Color2 [Da | 1 | 4 |
| + | 2.146038 | 0.545689 | Q9H501 | Q9H501 | ESF1 | ESF1 homo | Color2 [Da | 1 | 4 |
| + | 1.240669 | -1.38957 | Q9H553 | Q9H553 | ALG2 | Alpha-1,3/ | Color2 [Da | 1 | 4 |
| | 0.112758 | 0.08298 | Q9H579 | Q9H579 | MROH8 | Protein MI | Color2 [Da | 1 | 4 |
| | 0.064803 | -0.03809 | Q9H582 | Q9H582 | ZNF644 | Zinc finger | Color2 [Da | 1 | 4 |
| + | 1.919713 | -0.55151 | Q9H583 | Q9H583 | HEATR1 | HEAT repe | Color2 [Da | 1 | 4 |
| | 0 | -0.5431 | Q9H598 | Q9H598 | SLC32A1 | Vesicular i | Color2 [Da | 1 | 4 |
| + | 2.398705 | -1.31389 | Q9H5K3 | Q9H5K3 | POMK | Protein O- | Color2 [Da | 1 | 4 |
| + | 1.135799 | -0.52475 | Q9H5N1 | Q9H5N1 | RABEP2 | Rab GTPas | Color2 [Da | 1 | 4 |
| | 0 | 0.818825 | Q9H5V9 | Q9H5V9 | STEEP1 | STING ER e | Color2 [Da | 1 | 4 |
| | 0.52357 | -0.25554 | Q9H5X1 | Q9H5X1 | CIAO2A | Cytosolic i | Color2 [Da | 1 | 4 |
| | 0 | NaN | Q9H6A9 | Q9H6A9 | PCNX3 | Pecanex-lil | Color2 [Da | 1 | 4 |
| + | 2.743772 | 2.364358 | Q9H6E4 | Q9H6E4 | CCDC134 | Coiled-coil | Color2 [Da | 1 | 4 |
| + | 2.508653 | 0.896596 | Q9H6F5 | Q9H6F5 | CCDC86 | Coiled-coil | Color2 [Da | 1 | 4 |
| + | 0.960732 | -1.21607 | Q9H6K4 | Q9H6K4 | OPA3 | Optic atro | Color2 [Da | 1 | 4 |
| + | 2.374753 | 1.549811 | Q9H6K5 | Q9H6K5 | PRR36 | Proline-ric | Color2 [Da | 1 | 4 |
| + | 2.529844 | -1.52813 | Q9H6L2 | Q9H6L2 | TMEM231 | Transmem | Color2 [Da | 1 | 4 |
| | 0.560203 | -0.16071 | Q9H6R3 | Q9H6R3 | ACSS3 | Acyl-CoA s | Color2 [Da | 1 | 4 |
| | 0.153239 | 0.036282 | Q9H6R4 | Q9H6R4 | NOL6 | Nucleolar | Color2 [Da | 1 | 4 |
| | 0.575433 | -0.94931 | Q9H6R7 | Q9H6R7 | WDCP | WD repeat | Color2 [Da | 1 | 4 |
| | 0.459878 | 0.359744 | Q9H6T3 | Q9H6T3 | RPAP3 | RNA polyn | Color2 [Da | 1 | 4 |
| + | 1.301946 | -0.31834 | Q9H6U6 | Q9H6U6 | BCAS3 | BCAS3 mic | Color2 [Da | 1 | 4 |
| | 0 | -2.25599 | Q9H6U8 | Q9H6U8 | ALG9 | Alpha-1,2- | Color2 [Da | 1 | 4 |
| + | 3.533377 | -0.57946 | Q9H6V9 | Q9H6V9 | LDAH | Lipid dropl | Color2 [Da | 1 | 4 |
| | 0.37223 | -0.18483 | Q9H6W3 | Q9H6W3 | RIOX1 | Ribosomal | Color2 [Da | 1 | 4 |
| + | 1.668612 | 0.818435 | Q9H6X2 | Q9H6X2 | ANTXR1 | Anthrax to | Color2 [Da | 1 | 4 |
| | 0.34051 | 0.231664 | Q9H6Y7 | Q9H6Y7 | RNF167 | E3 ubiquiti | Color2 [Da | 1 | 4 |
| + | 2.893918 | 1.099391 | Q9H6Z4 | Q9H6Z4 | RANBP3 | Ran-bindir | Color2 [Da | 1 | 4 |
| + | 0.857088 | 0.498899 | Q9H772 | Q9H772 | GREM2 | Gremlin-2 | Color2 [Da | 1 | 4 |
| | 1.276998 | 0.218881 | Q9H773 | Q9H773 | DCTPP1 | dCTP pyro | Color2 [Da | 1 | 4 |
| + | 3.141783 | -1.43734 | Q9H7B2 | Q9H7B2 | RPF2 | Ribosome | Color2 [Da | 1 | 4 |
| + | 2.190773 | -1.49476 | Q9H7B4 | Q9H7B4 | SMYD3 | Histone-ly | Color2 [Da | 1 | 4 |
| + | 2.130014 | 1.82853 | Q9H7C9 | Q9H7C9 | AAMDC | Mth938 do | Color2 [Da | 1 | 4 |
| | 0.46046 | 1.020839 | Q9H7D0 | Q9H7D0 | DOCK5 | Dedicator | Color2 [Da | 1 | 4 |
| + | 1.130407 | -0.72312 | Q9H7D7 | Q9H7D7 | WDR26 | WD repeat | Color2 [Da | 1 | 4 |
| + | 1.047388 | -0.33545 | Q9H7E9 | Q9H7E9 | C8orf33 | UPF0488 p | Color2 [Da | 1 | 4 |
| + | 2.270747 | -0.37106 | Q9H7J1 | Q9H7J1 | PPP1R3E | Protein ph | Color2 [Da | 1 | 4 |

| | | | | | | | | | |
|---|---|---|---|---|---|---|---|---|---|
| | 0.002975 | -0.00135 | Q9H7L9 | Q9H7L9 | SUDS3 | Sin3 histor | Color2 [Da | 1 | 4 |
| + | 2.515278 | 0.742141 | Q9H7N4 | Q9H7N4 | SCAF1 | Splicing fa( | Color2 [Da | 1 | 4 |
| + | 3.146729 | 1.329755 | Q9H7P6 | Q9H7P6 | MVB12B | Multivesic | Color2 [Da | 1 | 4 |
| + | 1.048821 | 0.475025 | Q9H7X7 | Q9H7X7 | IFT22 | Intraflagell | Color2 [Da | 1 | 4 |
| + | 1.280659 | -0.76159 | Q9H7Z6 | Q9H7Z6 | KAT8 | Histone ac | Color2 [Da | 1 | 4 |
| + | 4.025514 | 2.467447 | Q9H7Z7 | Q9H7Z7 | PTGES2 | Prostaglan | Color2 [Da | 1 | 4 |
| + | 2.329913 | 0.700277 | Q9H814 | Q9H814 | PHAX | Phosphory | Color2 [Da | 1 | 4 |
| + | 1.843561 | -0.38862 | Q9H832 | Q9H832 | UBE2Z | Ubiquitin-( | Color2 [Da | 1 | 4 |
| + | 2.491044 | 1.574695 | Q9H840 | Q9H840 | GEMIN7 | Gem-asso( | Color2 [Da | 1 | 4 |
| + | 2.775255 | -0.71379 | Q9H845 | Q9H845 | ACAD9 | Complex I | Color2 [Da | 1 | 4 |
| | 0.031665 | -0.05492 | Q9H853 | Q9H853 | TUBA4B | Putative tu | Color2 [Da | 1 | 4 |
| + | 1.080076 | -0.48685 | Q9H857 | Q9H857 | NT5DC2 | 5'-nucleoti | Color2 [Da | 1 | 4 |
| | 0 | -0.53831 | Q9H871 | Q9H871 | RMND5A | E3 ubiquiti | Color2 [Da | 1 | 4 |
| | 0.155935 | -0.2168 | Q9H875 | Q9H875 | PRKRIP1 | PRKR-inter | Color2 [Da | 1 | 4 |
| | 0.741075 | -0.46978 | Q9H8G2 | Q9H8G2 | CAAP1 | Caspase ac | Color2 [Da | 1 | 4 |
| | 0 | NaN | Q9H8H2 | Q9H8H2 | DDX31 | Probable A | Color2 [Da | 1 | 4 |
| | 0.647929 | -0.6657 | Q9H8H3 | Q9H8H3 | METTL7A | Putative m | Color2 [Da | 1 | 4 |
| | 0.571888 | 0.53369 | Q9H8M2 | Q9H8M2 | BRD9 | Bromodon | Color2 [Da | 1 | 4 |
| + | 1.037416 | 0.565607 | Q9H8M5 | Q9H8M5 | CNNM2 | Metal tran | Color2 [Da | 1 | 4 |
| + | 2.283992 | -1.31 | Q9H8M7 | Q9H8M7 | MINDY3 | Ubiquitin ( | Color2 [Da | 1 | 4 |
| + | 2.844257 | -1.04314 | Q9H8T0 | Q9H8T0 | AKTIP | AKT-intera | Color2 [Da | 1 | 4 |
| | 0.152471 | 0.112847 | Q9H8Y5 | Q9H8Y5 | ANKZF1 | Ankyrin re | Color2 [Da | 1 | 4 |
| + | 2.981595 | 1.887126 | Q9H8Y8 | Q9H8Y8 | GORASP2 | Golgi reas | Color2 [Da | 1 | 4 |
| | 0 | NaN | Q9H900 | Q9H900 | ZWILCH | Protein zw | Color2 [Da | 1 | 4 |
| + | 2.405849 | 1.357968 | Q9H902 | Q9H902 | REEP1 | Receptor e | Color2 [Da | 1 | 4 |
| + | 2.098412 | 1.216499 | Q9H903 | Q9H903 | MTHFD2L | Bifunction | Color2 [Da | 1 | 4 |
| + | 3.365265 | 3.645299 | Q9H910 | Q9H910 | JPT2 | Jupiter mi( | Color2 [Da | 1 | 4 |
| | 0 | 0.194572 | Q9H920 | Q9H920 | RNF121 | RING finge | Color2 [Da | 1 | 4 |
| | 0.474182 | 0.325746 | Q9H936 | Q9H936 | SLC25A22 | Mitochonc | Color2 [Da | 1 | 4 |
| | 0 | -1.50999 | Q9H974 | Q9H974 | QTRT2 | Queuine tI | Color2 [Da | 1 | 4 |
| | 0 | 0.250152 | Q9H977 | Q9H977 | WDR54 | WD repeat | Color2 [Da | 1 | 4 |
| + | 3.171822 | -1.55167 | Q9H993 | Q9H993 | ARMT1 | Damage-co | Color2 [Da | 1 | 4 |
| + | 1.939355 | -0.95103 | Q9H9A5 | Q9H9A5 | CNOT10 | CCR4-NOT | Color2 [Da | 1 | 4 |
| + | 2.466405 | -0.45888 | Q9H9A6 | Q9H9A6 | LRRC40 | Leucine-ric | Color2 [Da | 1 | 4 |
| + | 1.666588 | -1.1148 | Q9H9B1 | Q9H9B1 | EHMT1 | Histone-ly: | Color2 [Da | 1 | 4 |
| | 0.109756 | -0.03194 | Q9H9B4 | Q9H9B4 | SFXN1 | Sideroflexi | Color2 [Da | 1 | 4 |
| | 0.255286 | 0.30119 | Q9H9C1 | Q9H9C1 | VIPAS39 | Spermatog | Color2 [Da | 1 | 4 |
| + | 2.468711 | -1.55481 | Q9H9E3 | Q9H9E3 | COG4 | Conserved | Color2 [Da | 1 | 4 |
| | 0 | NaN | Q9H9G7 | Q9H9G7 | AGO3 | Protein arg | Color2 [Da | 1 | 4 |
| + | 4.784992 | 2.274137 | Q9H9H4 | Q9H9H4 | VPS37B | Vacuolar p | Color2 [Da | 1 | 4 |
| + | 1.146217 | -0.29557 | Q9H9J2 | Q9H9J2 | MRPL44 | 39S riboso | Color2 [Da | 1 | 4 |
| + | 1.454181 | -0.71078 | Q9H9P8 | Q9H9P8 | L2HGDH | L-2-hydrox | Color2 [Da | 1 | 4 |
| + | 1.062222 | -0.77718 | Q9H9Q2 | Q9H9Q2 | COPS7B | COP9 signa | Color2 [Da | 1 | 4 |
| + | 1.693393 | -1.19082 | Q9H9Q4 | Q9H9Q4 | NHEJ1 | Non-homo | Color2 [Da | 1 | 4 |
| | 0 | 1.876624 | Q9H9R9 | Q9H9R9 | DBNDD1 | Dysbindin | Color2 [Da | 1 | 4 |
| + | 2.211196 | -1.13513 | Q9H9T3 | Q9H9T3 | ELP3 | Elongator ( | Color2 [Da | 1 | 4 |
| | 0.307969 | 0.194771 | Q9H9Y2 | Q9H9Y2 | RPF1 | Ribosome | Color2 [Da | 1 | 4 |

| | | | | | | | | | |
|---|---|---|---|---|---|---|---|---|---|
| + | 2.539029 | -2.43242 | Q9H9Y4 | Q9H9Y4 | GPN2 | GPN-loop ( | Color2 [Da | 1 | 4 |
| | 0 | NaN | Q9H9Y6 | Q9H9Y6 | POLR1B | DNA-direc | Color2 [Da | 1 | 4 |
| + | 2.063862 | -1.34796 | Q9H9Z2 | Q9H9Z2 | LIN28A | Protein lin | Color2 [Da | 1 | 4 |
| | 0.885234 | 0.302537 | Q9HA47 | Q9HA47 | UCK1 | Uridine-cy | Color2 [Da | 1 | 4 |
| + | 0.990728 | 0.506788 | Q9HA64 | Q9HA64 | FN3KRP | Ketosamin | Color2 [Da | 1 | 4 |
| | 0.031643 | -0.01694 | Q9HA65 | Q9HA65 | TBC1D17 | TBC1 doma | Color2 [Da | 1 | 4 |
| + | 2.380597 | -1.06149 | Q9HA77 | Q9HA77 | CARS2 | Probable c | Color2 [Da | 1 | 4 |
| + | 3.371933 | 1.898388 | Q9HAB8 | Q9HAB8 | PPCS | Phosphopa | Color2 [Da | 1 | 4 |
| | 0.094126 | -0.07521 | Q9HAC7 | Q9HAC7 | SUGCT | Succinate- | Color2 [Da | 1 | 4 |
| | 0.476443 | -0.54103 | Q9HAD4 | Q9HAD4 | WDR41 | WD repeat | Color2 [Da | 1 | 4 |
| | 0.005769 | -0.01101 | Q9HAF1 | Q9HAF1 | MEAF6 | Chromatin | Color2 [Da | 1 | 4 |
| | 0.009508 | -0.02226 | Q9HAN9 | Q9HAN9 | NMNAT1 | Nicotinam | Color2 [Da | 1 | 4 |
| + | 2.559387 | 3.348496 | Q9HAP6 | Q9HAP6 | LIN7B | Protein lin | Color2 [Da | 1 | 4 |
| + | 2.654182 | -0.95567 | Q9HAR2 | Q9HAR2 | ADGRL3 | Adhesion ( | Color2 [Da | 1 | 4 |
| | 0.542949 | -0.25475 | Q9HAS0 | Q9HAS0 | C17orf75 | Protein Nji | Color2 [Da | 1 | 4 |
| + | 5.806819 | 2.784213 | Q9HAT2 | Q9HAT2 | SIAE | Sialate O-a | Color2 [Da | 1 | 4 |
| | 0.644275 | -0.17737 | Q9HAU0 | Q9HAU0 | PLEKHA5 | Pleckstrin | Color2 [Da | 1 | 4 |
| + | 1.995545 | -1.18506 | Q9HAU5 | Q9HAU5 | UPF2 | Regulator | Color2 [Da | 1 | 4 |
| | 0.084643 | 0.111106 | Q9HAV0 | Q9HAV0 | GNB4 | Guanine n | Color2 [Da | 1 | 4 |
| + | 4.106992 | -1.42088 | Q9HAV4 | Q9HAV4 | XPO5 | Exportin-5 | Color2 [Da | 1 | 4 |
| + | 3.823149 | 1.2805 | Q9HAV7 | Q9HAV7 | GRPEL1 | GrpE prote | Color2 [Da | 1 | 4 |
| | 0 | NaN | Q9HAW0 | Q9HAW0 | BRF2 | Transcripti | Color2 [Da | 1 | 4 |
| | 0 | NaN | Q9HAZ1 | Q9HAZ1 | CLK4 | Dual speci | Color2 [Da | 1 | 4 |
| + | 1.371712 | -0.31614 | Q9HB07 | Q9HB07 | MYG1 | MYG1 exo | Color2 [Da | 1 | 4 |
| + | 1.437591 | 0.665655 | Q9HB21 | Q9HB21 | PLEKHA1 | Pleckstrin | Color2 [Da | 1 | 4 |
| | 0.201576 | 0.236018 | Q9HB40 | Q9HB40 | SCPEP1 | Retinoid-ir | Color2 [Da | 1 | 4 |
| + | 1.611732 | 0.512342 | Q9HB71 | Q9HB71 | CACYBP | Calcyclin-b | Color2 [Da | 1 | 4 |
| + | 3.536465 | -0.8125 | Q9HB90 | Q9HB90 | RRAGC | Ras-relate | Color2 [Da | 1 | 4 |
| | 0.219166 | -0.12596 | Q9HBD1 | Q9HBD1 | RC3H2 | Roquin-2 | Color2 [Da | 1 | 4 |
| | 0 | 0.172042 | Q9HBF4 | Q9HBF4 | ZFYVE1 | Zinc finger | Color2 [Da | 1 | 4 |
| | 0.468894 | -0.70855 | Q9HBH5 | Q9HBH5 | RDH14 | Retinol de | Color2 [Da | 1 | 4 |
| | 0 | NaN | Q9HBK9 | Q9HBK9 | AS3MT | Arsenite m | Color2 [Da | 1 | 4 |
| + | 1.574641 | 0.814842 | Q9HBL7 | Q9HBL7 | PLGRKT | Plasminog | Color2 [Da | 1 | 4 |
| + | 4.89343 | -1.39234 | Q9HBM0 | Q9HBM0 | VEZT | Vezatin | Color2 [Da | 1 | 4 |
| | 0.720876 | 0.301859 | Q9HBM1 | Q9HBM1 | SPC25 | Kinetochor | Color2 [Da | 1 | 4 |
| | 1.074729 | -0.23691 | Q9HBR0 | Q9HBR0 | SLC38A10 | Putative so | Color2 [Da | 1 | 4 |
| | 0 | NaN | Q9HBU6 | Q9HBU6 | ETNK1 | Ethanolam | Color2 [Da | 1 | 4 |
| | 0 | -1.68915 | Q9HC07 | Q9HC07 | TMEM165 | Transmem | Color2 [Da | 1 | 4 |
| | 0.026366 | -0.01975 | Q9HC35 | Q9HC35 | EML4 | Echinoder | Color2 [Da | 1 | 4 |
| + | 2.658789 | 1.485961 | Q9HC38 | Q9HC38 | GLOD4 | Glyoxalase | Color2 [Da | 1 | 4 |
| + | 2.058026 | 1.085164 | Q9HC52 | Q9HC52 | CBX8 | Chromobo | Color2 [Da | 1 | 4 |
| + | 1.468275 | 0.415257 | Q9HC56 | Q9HC56 | PCDH9 | Protocadh | Color2 [Da | 1 | 4 |
| | 0 | -0.68431 | Q9HC78 | Q9HC78 | ZBTB20 | Zinc finger | Color2 [Da | 1 | 4 |
| | 0 | NaN | Q9HC98 | Q9HC98 | NEK6 | Serine/thr | Color2 [Da | 1 | 4 |
| + | 2.708464 | 2.789542 | Q9HCB6 | Q9HCB6 | SPON1 | Spondin-1 | Color2 [Da | 1 | 4 |
| + | 3.521136 | -1.05865 | Q9HCC0 | Q9HCC0 | MCCC2 | Methylcro | Color2 [Da | 1 | 4 |
| | 0.418934 | -0.09748 | Q9HCD5 | Q9HCD5 | NCOA5 | Nuclear re | Color2 [Da | 1 | 4 |

| | | | | | | | | | |
|---|---|---|---|---|---|---|---|---|---|
| | 0.663634 | 0.151575 | Q9HCD6 | Q9HCD6 | TANC2 | Protein TA | Color2 [Da | 1 | 4 |
| | 0 | NaN | Q9HCE0 | Q9HCE0 | EPG5 | Ectopic P g | Color2 [Da | 1 | 4 |
| + | 1.556667 | -0.32054 | Q9HCE3 | Q9HCE3 | ZNF532 | Zinc finger | Color2 [Da | 1 | 4 |
| + | 2.178285 | -1.65754 | Q9HCE5 | Q9HCE5 | METTL14 | N6-adenos | Color2 [Da | 1 | 4 |
| | 0.302366 | 0.265217 | Q9HCE7 | Q9HCE7 | SMURF1 | E3 ubiquiti | Color2 [Da | 1 | 4 |
| + | 1.975018 | -1.42809 | Q9HCG8 | Q9HCG8 | CWC22 | Pre-mRNA | Color2 [Da | 1 | 4 |
| + | 1.229435 | 1.310787 | Q9HCH3 | Q9HCH3 | CPNE5 | Copine-5 | Color2 [Da | 1 | 4 |
| | 0 | NaN | Q9HCJ0 | Q9HCJ0 | TNRC6C | Trinucleoti | Color2 [Da | 1 | 4 |
| | 0 | NaN | Q9HCJ1 | Q9HCJ1 | ANKH | Progressiv | Color2 [Da | 1 | 4 |
| + | 2.528696 | 0.313882 | Q9HCJ6 | Q9HCJ6 | VAT1L | Synaptic v | Color2 [Da | 1 | 4 |
| + | 2.636116 | -1.17184 | Q9HCK4 | Q9HCK4 | ROBO2 | Roundabo | Color2 [Da | 1 | 4 |
| + | 0.78377 | 0.764678 | Q9HCK5 | Q9HCK5 | AGO4 | Protein arg | Color2 [Da | 1 | 4 |
| + | 1.480778 | -0.5672 | Q9HCK8 | Q9HCK8 | CHD8 | Chromodo | Color2 [Da | 1 | 4 |
| + | 1.030726 | 0.969875 | Q9HCM2 | Q9HCM2 | PLXNA4 | Plexin-A4 | Color2 [Da | 1 | 4 |
| + | 2.221709 | 0.756318 | Q9HCM3 | Q9HCM3 | KIAA1549 | UPF0606 p | Color2 [Da | 1 | 4 |
| + | 2.219085 | -1.11397 | Q9HCM4 | Q9HCM4 | EPB41L5 | Band 4.1-li | Color2 [Da | 1 | 4 |
| | 0 | -1.21356 | Q9HCN4 | Q9HCN4 | GPN1 | GPN-loop ( | Color2 [Da | 1 | 4 |
| + | 2.80261 | -0.76657 | Q9HCN8 | Q9HCN8 | SDF2L1 | Stromal ce | Color2 [Da | 1 | 4 |
| + | 1.433044 | -1.83885 | Q9HCS4 | Q9HCS4 | TCF7L1 | Transcripti | Color2 [Da | 1 | 4 |
| | 0 | NaN | Q9HCS5 | Q9HCS5 | EPB41L4A | Band 4.1-li | Color2 [Da | 1 | 4 |
| + | 1.122953 | -0.93508 | Q9HCS7 | Q9HCS7 | XAB2 | Pre-mRNA | Color2 [Da | 1 | 4 |
| + | 1.711324 | 0.421704 | Q9HCU4 | Q9HCU4 | CELSR2 | Cadherin E | Color2 [Da | 1 | 4 |
| + | 3.509264 | -1.1295 | Q9HCU5 | Q9HCU5 | PREB | Prolactin r | Color2 [Da | 1 | 4 |
| + | 3.838984 | 1.586934 | Q9HD15 | Q9HD15 | SRA1 | Steroid rec | Color2 [Da | 1 | 4 |
| + | 2.958789 | -1.063 | Q9HD20 | Q9HD20 | ATP13A1 | Endoplasm | Color2 [Da | 1 | 4 |
| | 0 | NaN | Q9HD26 | Q9HD26 | GOPC | Golgi-asso | Color2 [Da | 1 | 4 |
| + | 2.42664 | 1.097151 | Q9HD33 | Q9HD33 | MRPL47 | 39S riboso | Color2 [Da | 1 | 4 |
| + | 1.298807 | 1.159897 | Q9HD34 | Q9HD34 | LYRM4 | LYR motif- | Color2 [Da | 1 | 4 |
| | 0 | NaN | Q9HD40 | Q9HD40 | SEPSECS | O-phospho | Color2 [Da | 1 | 4 |
| + | 3.353642 | 1.43539 | Q9HD42 | Q9HD42 | CHMP1A | Charged m | Color2 [Da | 1 | 4 |
| + | 2.552092 | -1.41243 | Q9HD45 | Q9HD45 | TM9SF3 | Transmem | Color2 [Da | 1 | 4 |
| | 0.332538 | 0.033393 | Q9HDC9 | Q9HDC9 | APMAP | Adipocyte | Color2 [Da | 1 | 4 |
| + | 2.159102 | -2.67899 | Q9NNW5 | Q9NNW5 | WDR6 | WD repeat | Color2 [Da | 1 | 4 |
| + | 0.836178 | 1.637485 | Q9NNW7 | Q9NNW7 | TXNRD2 | Thioredoxi | Color2 [Da | 1 | 4 |
| + | 3.358107 | -2.71618 | Q9NP55 | Q9NP55 | BPIFA1 | BPI fold-cc | Color2 [Da | 1 | 4 |
| + | 3.177035 | -1.16882 | Q9NP58 | Q9NP58 | ABCB6 | ATP-bindir | Color2 [Da | 1 | 4 |
| + | 1.81706 | -0.68068 | Q9NP61 | Q9NP61 | ARFGAP3 | ADP-ribosy | Color2 [Da | 1 | 4 |
| | 0.13776 | -0.09292 | Q9NP64 | Q9NP64 | ZCCHC17 | Zinc finger | Color2 [Da | 1 | 4 |
| + | 3.238446 | 1.217345 | Q9NP66 | Q9NP66 | HMG20A | High mobil | Color2 [Da | 1 | 4 |
| + | 1.124281 | 0.26992 | Q9NP72 | Q9NP72 | RAB18 | Ras-relate | Color2 [Da | 1 | 4 |
| + | 1.240153 | -0.58258 | Q9NP73 | Q9NP73 | ALG13 | Putative bi | Color2 [Da | 1 | 4 |
| + | 2.473134 | 2.176772 | Q9NP74 | Q9NP74 | PALMD | Palmdelph | Color2 [Da | 1 | 4 |
| + | 1.775893 | -1.06474 | Q9NP77 | Q9NP77 | SSU72 | RNA polym | Color2 [Da | 1 | 4 |
| | 0.618469 | -0.26246 | Q9NP79 | Q9NP79 | VTA1 | Vacuolar p | Color2 [Da | 1 | 4 |
| | 0.211135 | -0.15661 | Q9NP80 | Q9NP80 | PNPLA8 | Calcium-in | Color2 [Da | 1 | 4 |
| | 0.074123 | -0.02839 | Q9NP81 | Q9NP81 | SARS2 | Serine--tRI | Color2 [Da | 1 | 4 |
| | 0.453332 | 0.080348 | Q9NP90 | Q9NP90 | RAB9B | Ras-relate | Color2 [Da | 1 | 4 |

| | | | | | | | | | |
|---|---|---|---|---|---|---|---|---|---|
| | 0 | -1.51353 | Q9NP92 | Q9NP92 | MRPS30 | 39S riboso | Color2 [Da | 1 | 4 |
| + | 2.577422 | 1.444616 | Q9NP97 | Q9NP97 | DYNLRB1 | Dynein ligl | Color2 [Da | 1 | 4 |
| + | 1.686376 | 1.576574 | Q9NPA0 | Q9NPA0 | EMC7 | ER membr | Color2 [Da | 1 | 4 |
| + | 2.510996 | 1.582054 | Q9NPA8 | Q9NPA8 | ENY2 | Transcripti | Color2 [Da | 1 | 4 |
| | 0 | NaN | Q9NPB6 | Q9NPB6 | PARD6A | Partitionin | Color2 [Da | 1 | 4 |
| + | 1.323915 | -0.68655 | Q9NPB8 | Q9NPB8 | GPCPD1 | Glyceroph | Color2 [Da | 1 | 4 |
| | 0.199668 | -0.19761 | Q9NPD3 | Q9NPD3 | EXOSC4 | Exosome c | Color2 [Da | 1 | 4 |
| | 0 | -1.51519 | Q9NPD7 | Q9NPD7 | NRN1 | Neuritin | Color2 [Da | 1 | 4 |
| | 0 | -0.98499 | Q9NPD8 | Q9NPD8 | UBE2T | Ubiquitin-c | Color2 [Da | 1 | 4 |
| | 0.086448 | 0.022727 | Q9NPE2 | Q9NPE2 | NGRN | Neugrin | Color2 [Da | 1 | 4 |
| | 0 | NaN | Q9NPF0 | Q9NPF0 | CD320 | CD320 ant | Color2 [Da | 1 | 4 |
| + | 0.773566 | -0.79231 | Q9NPF4 | Q9NPF4 | OSGEP | tRNA N6-a | Color2 [Da | 1 | 4 |
| + | 1.536057 | 1.336169 | Q9NPF5 | Q9NPF5 | DMAP1 | DNA meth | Color2 [Da | 1 | 4 |
| | 0 | NaN | Q9NPG1 | Q9NPG1 | FZD3 | Frizzled-3 | Color2 [Da | 1 | 4 |
| + | 1.745932 | -2.94059 | Q9NPG3 | Q9NPG3 | UBN1 | Ubinuclein | Color2 [Da | 1 | 4 |
| | 0.099518 | 0.026138 | Q9NPH2 | Q9NPH2 | ISYNA1 | Inositol-3-| | Color2 [Da | 1 | 4 |
| | 0.177904 | 0.09091 | Q9NPI1 | Q9NPI1 | BRD7 | Bromodon | Color2 [Da | 1 | 4 |
| | 0.545536 | 0.170602 | Q9NPI6 | Q9NPI6 | DCP1A | mRNA-dec | Color2 [Da | 1 | 4 |
| + | 3.140883 | 1.486887 | Q9NPJ3 | Q9NPJ3 | ACOT13 | Acyl-coenz | Color2 [Da | 1 | 4 |
| + | 2.802376 | 0.867387 | Q9NPJ6 | Q9NPJ6 | MED4 | Mediator c | Color2 [Da | 1 | 4 |
| + | 2.924743 | 1.509396 | Q9NPL8 | Q9NPL8 | TIMMDC1 | Complex I | Color2 [Da | 1 | 4 |
| + | 3.295613 | -1.19224 | Q9NPQ8 | Q9NPQ8 | RIC8A | Synembryı | Color2 [Da | 1 | 4 |
| | 0 | NaN | Q9NQ11 | Q9NQ11 | ATP13A2 | Polyamine | Color2 [Da | 1 | 4 |
| + | 3.827436 | -0.77859 | Q9NQ29 | Q9NQ29 | LUC7L | Putative R | Color2 [Da | 1 | 4 |
| + | 1.671029 | 1.059381 | Q9NQ48 | Q9NQ48 | LZTFL1 | Leucine zip | Color2 [Da | 1 | 4 |
| + | 3.198782 | 1.328621 | Q9NQ50 | Q9NQ50 | MRPL40 | 39S riboso | Color2 [Da | 1 | 4 |
| | 0.041754 | 0.045874 | Q9NQ55 | Q9NQ55 | PPAN | Suppresso | Color2 [Da | 1 | 4 |
| + | 3.419546 | 0.995502 | Q9NQ66 | Q9NQ66 | PLCB1 | 1-phospha | Color2 [Da | 1 | 4 |
| | 0 | NaN | Q9NQ69 | Q9NQ69 | LHX9 | LIM/home | Color2 [Da | 1 | 4 |
| + | 3.191082 | -1.72573 | Q9NQ86 | Q9NQ86 | TRIM36 | E3 ubiquiti | Color2 [Da | 1 | 4 |
| + | 1.76746 | -0.64595 | Q9NQ88 | Q9NQ88 | TIGAR | Fructose-2 | Color2 [Da | 1 | 4 |
| | 0.736566 | 0.372159 | Q9NQ89 | Q9NQ89 | C12orf4 | Protein C1 | Color2 [Da | 1 | 4 |
| | 0 | NaN | Q9NQ92 | Q9NQ92 | COPRS | Coordinato | Color2 [Da | 1 | 4 |
| + | 1.987802 | 1.783686 | Q9NQA3 | Q9NQA3 | WASH6P | WAS prote | Color2 [Da | 1 | 4 |
| + | 3.066544 | 1.384311 | Q9NQC3 | Q9NQC3 | RTN4 | Reticulon-4 | Color2 [Da | 1 | 4 |
| | 0.034387 | -0.02558 | Q9NQC8 | Q9NQC8 | IFT46 | Intraflagell | Color2 [Da | 1 | 4 |
| | 0.022367 | 0.016 | Q9NQE9 | Q9NQE9 | HINT3 | Adenosine | Color2 [Da | 1 | 4 |
| | 0.255534 | 0.100212 | Q9NQG5 | Q9NQG5 | RPRD1B | Regulation | Color2 [Da | 1 | 4 |
| + | 1.765837 | 1.182065 | Q9NQG6 | Q9NQG6 | MIEF1 | Mitochonc | Color2 [Da | 1 | 4 |
| | 0.183233 | -0.12146 | Q9NQH7 | Q9NQH7 | XPNPEP3 | Xaa-Pro ar | Color2 [Da | 1 | 4 |
| + | 0.929465 | 0.555726 | Q9NQP4 | Q9NQP4 | PFDN4 | Prefoldin s | Color2 [Da | 1 | 4 |
| | 0.145261 | 0.120753 | Q9NQR4 | Q9NQR4 | NIT2 | Omega-am | Color2 [Da | 1 | 4 |
| + | 2.305878 | 0.934219 | Q9NQS1 | Q9NQS1 | AVEN | Cell death | Color2 [Da | 1 | 4 |
| + | 1.718531 | -1.0135 | Q9NQS7 | Q9NQS7 | INCENP | Inner centı | Color2 [Da | 1 | 4 |
| + | 1.218867 | -1.52036 | Q9NQT4 | Q9NQT4 | EXOSC5 | Exosome c | Color2 [Da | 1 | 4 |
| | 0.393783 | -0.2652 | Q9NQT5 | Q9NQT5 | EXOSC3 | Exosome c | Color2 [Da | 1 | 4 |
| | 0 | -0.63527 | Q9NQV6 | Q9NQV6 | PRDM10 | PR domain | Color2 [Da | 1 | 4 |

| | | | | | | | | | | |
|---|---|---|---|---|---|---|---|---|---|---|
| + | 1.647937 | -1.03613 | Q9NQW6 | Q9NQW6 | ANLN | Anillin | Color2 | [Da | 1 | 4 |
| + | 2.940682 | -0.7134 | Q9NQW7 | Q9NQW7 | XPNPEP1 | Xaa-Pro ar | Color2 | [Da | 1 | 4 |
| + | 3.130868 | 1.729907 | Q9NQX3 | Q9NQX3 | GPHN | Gephyrin | Color2 | [Da | 1 | 4 |
| + | 0.92596 | 1.041724 | Q9NQX5 | Q9NQX5 | NPDC1 | Neural pro | Color2 | [Da | 1 | 4 |
| | 0.220913 | 0.231671 | Q9NQX7 | Q9NQX7 | ITM2C | Integral m | Color2 | [Da | 1 | 4 |
| + | 2.519742 | 1.186435 | Q9NQY0 | Q9NQY0 | BIN3 | Bridging in | Color2 | [Da | 1 | 4 |
| | 0.23306 | -0.05034 | Q9NQZ2 | Q9NQZ2 | UTP3 | Something | Color2 | [Da | 1 | 4 |
| + | 0.714834 | -1.35806 | Q9NR09 | Q9NR09 | BIRC6 | Baculovira | Color2 | [Da | 1 | 4 |
| + | 3.220828 | -1.92862 | Q9NR12 | Q9NR12 | PDLIM7 | PDZ and LI | Color2 | [Da | 1 | 4 |
| | 0.410513 | -0.35406 | Q9NR19 | Q9NR19 | ACSS2 | Acetyl-coe | Color2 | [Da | 1 | 4 |
| + | 3.028413 | 1.879272 | Q9NR28 | Q9NR28 | DIABLO | Diablo IAP | Color2 | [Da | 1 | 4 |
| + | 3.056937 | -1.09998 | Q9NR30 | Q9NR30 | DDX21 | Nucleolar | Color2 | [Da | 1 | 4 |
| + | 1.774355 | 0.602929 | Q9NR31 | Q9NR31 | SAR1A | GTP-bindir | Color2 | [Da | 1 | 4 |
| | 0.313046 | 0.12191 | Q9NR33 | Q9NR33 | POLE4 | DNA polyn | Color2 | [Da | 1 | 4 |
| + | 2.427762 | -0.16735 | Q9NR45 | Q9NR45 | NANS | Sialic acid | Color2 | [Da | 1 | 4 |
| + | 1.832031 | 0.724415 | Q9NR46 | Q9NR46 | SH3GLB2 | Endophilin | Color2 | [Da | 1 | 4 |
| + | 2.14688 | 0.752054 | Q9NR48 | Q9NR48 | ASH1L | Histone-ly | Color2 | [Da | 1 | 4 |
| + | 1.099359 | -0.65268 | Q9NR50 | Q9NR50 | EIF2B3 | Translatior | Color2 | [Da | 1 | 4 |
| | 0 | NaN | Q9NR77 | Q9NR77 | PXMP2 | Peroxisom | Color2 | [Da | 1 | 4 |
| | 0 | -1.72628 | Q9NRA0 | Q9NRA0 | SPHK2 | Sphingosir | Color2 | [Da | 1 | 4 |
| | 0.106069 | -0.0558 | Q9NRA1 | Q9NRA1 | PDGFC | Platelet-de | Color2 | [Da | 1 | 4 |
| + | 1.982171 | 0.628435 | Q9NRA8 | Q9NRA8 | EIF4ENIF1 | Eukaryotic | Color2 | [Da | 1 | 4 |
| | 0 | NaN | Q9NRB3 | Q9NRB3 | CHST12 | Carbohydr | Color2 | [Da | 1 | 4 |
| | 0 | NaN | Q9NRC1 | Q9NRC1 | ST7 | Suppresso | Color2 | [Da | 1 | 4 |
| + | 2.433025 | -1.12233 | Q9NRD5 | Q9NRD5 | PICK1 | PRKCA-bin | Color2 | [Da | 1 | 4 |
| + | 1.223685 | 0.448814 | Q9NRF2 | Q9NRF2 | SH2B1 | SH2B adap | Color2 | [Da | 1 | 4 |
| + | 1.297762 | -0.5943 | Q9NRF8 | Q9NRF8 | CTPS2 | CTP syntha | Color2 | [Da | 1 | 4 |
| + | 2.299065 | 1.117647 | Q9NRF9 | Q9NRF9 | POLE3 | DNA polyn | Color2 | [Da | 1 | 4 |
| | 0.04339 | 0.117737 | Q9NRG0 | Q9NRG0 | CHRAC1 | Chromatin | Color2 | [Da | 1 | 4 |
| + | 1.251931 | 1.010044 | Q9NRG7 | Q9NRG7 | SDR39U1 | Epimerase | Color2 | [Da | 1 | 4 |
| + | 2.917687 | -0.82554 | Q9NRG9 | Q9NRG9 | AAAS | Aladin | Color2 | [Da | 1 | 4 |
| + | 1.227421 | -0.54303 | Q9NRJ7 | Q9NRJ7 | PCDHB16 | Protocadh | Color2 | [Da | 1 | 4 |
| + | 2.063753 | 0.693134 | Q9NRL3 | Q9NRL3 | STRN4 | Striatin-4 | Color2 | [Da | 1 | 4 |
| + | 1.596007 | -0.27754 | Q9NRN7 | Q9NRN7 | AASDHPPT | L-aminoad | Color2 | [Da | 1 | 4 |
| | 0.112148 | -0.06298 | Q9NRP2 | Q9NRP2 | CMC2 | COX assen | Color2 | [Da | 1 | 4 |
| + | 3.242428 | 1.22703 | Q9NRR5 | Q9NRR5 | UBQLN4 | Ubiquilin-4 | Color2 | [Da | 1 | 4 |
| + | 3.076378 | 1.031414 | Q9NRS6 | Q9NRS6 | SNX15 | Sorting ne | Color2 | [Da | 1 | 4 |
| + | 1.487073 | 0.680212 | Q9NRV9 | Q9NRV9 | HEBP1 | Heme-binc | Color2 | [Da | 1 | 4 |
| + | 2.908213 | 1.353595 | Q9NRW1 | Q9NRW1 | RAB6B | Ras-relate | Color2 | [Da | 1 | 4 |
| + | 2.593543 | -0.89354 | Q9NRW7 | Q9NRW7 | VPS45 | Vacuolar p | Color2 | [Da | 1 | 4 |
| + | 1.4402 | -0.96038 | Q9NRX1 | Q9NRX1 | PNO1 | RNA-bindi | Color2 | [Da | 1 | 4 |
| + | 1.20804 | 1.739537 | Q9NRX2 | Q9NRX2 | MRPL17 | 39S riboso | Color2 | [Da | 1 | 4 |
| + | 1.408942 | 0.937822 | Q9NRX4 | Q9NRX4 | PHPT1 | 14 kDa ph | Color2 | [Da | 1 | 4 |
| | 0.527975 | -0.32851 | Q9NRX5 | Q9NRX5 | SERINC1 | Serine inc | Color2 | [Da | 1 | 4 |
| | 0.328082 | 0.303821 | Q9NRY2 | Q9NRY2 | INIP | SOSS comp | Color2 | [Da | 1 | 4 |
| + | 0.921453 | -0.3876 | Q9NRY4 | Q9NRY4 | ARHGAP35 | Rho GTPas | Color2 | [Da | 1 | 4 |
| | 0.071126 | -0.01891 | Q9NRY5 | Q9NRY5 | FAM114A2 | Protein FA | Color2 | [Da | 1 | 4 |

| | | | | | | | | | |
|---|---|---|---|---|---|---|---|---|---|
| | 0 | NaN | Q9NRY6 | Q9NRY6 | PLSCR3 | Phospholi | Color2 [Da | 1 | 4 |
| + | 3.242471 | -2.56404 | Q9NRZ5 | Q9NRZ5 | AGPAT4 | 1-acyl-sn-g | Color2 [Da | 1 | 4 |
| + | 1.654244 | -1.67388 | Q9NRZ7 | Q9NRZ7 | AGPAT3 | 1-acyl-sn-g | Color2 [Da | 1 | 4 |
| | 0.119051 | -0.29243 | Q9NS56 | Q9NS56 | TOPORS | E3 ubiquiti | Color2 [Da | 1 | 4 |
| + | 3.773 | 2.043731 | Q9NS69 | Q9NS69 | TOMM22 | Mitochonc | Color2 [Da | 1 | 4 |
| + | 2.805765 | -0.79424 | Q9NS86 | Q9NS86 | LANCL2 | LanC-like p | Color2 [Da | 1 | 4 |
| | 0 | -0.13606 | Q9NS87 | Q9NS87 | KIF15 | Kinesin-like | Color2 [Da | 1 | 4 |
| | 0.169375 | 0.079265 | Q9NSA3 | Q9NSA3 | CTNNBIP1 | Beta-cater | Color2 [Da | 1 | 4 |
| | 0 | 2.061762 | Q9NSB8 | Q9NSB8 | HOMER2 | Homer prc | Color2 [Da | 1 | 4 |
| + | 1.162051 | 2.769757 | Q9NSC2 | Q9NSC2 | SALL1 | Sal-like prc | Color2 [Da | 1 | 4 |
| + | 1.918581 | 0.643148 | Q9NSC5 | Q9NSC5 | HOMER3 | Homer prc | Color2 [Da | 1 | 4 |
| + | 2.397584 | -0.63956 | Q9NSD9 | Q9NSD9 | FARSB | Phenylalar | Color2 [Da | 1 | 4 |
| + | 3.297582 | -1.02729 | Q9NSE4 | Q9NSE4 | IARS2 | Isoleucine- | Color2 [Da | 1 | 4 |
| | 0.085295 | 0.055845 | Q9NSI2 | Q9NSI2 | SLX9 | Ribosome | Color2 [Da | 1 | 4 |
| | 0.126586 | 0.050171 | Q9NSI6 | Q9NSI6 | BRWD1 | Bromodon | Color2 [Da | 1 | 4 |
| | 0.462364 | -0.15736 | Q9NSK0 | Q9NSK0 | KLC4 | Kinesin ligl | Color2 [Da | 1 | 4 |
| + | 2.243704 | 1.881556 | Q9NSK7 | Q9NSK7 | C19orf12 | Protein C1 | Color2 [Da | 1 | 4 |
| + | 0.990338 | -0.35718 | Q9NSY0 | Q9NSY0 | NRBP2 | Nuclear re | Color2 [Da | 1 | 4 |
| + | 1.596083 | -0.5849 | Q9NT22 | Q9NT22 | EMILIN3 | EMILIN-3 | Color2 [Da | 1 | 4 |
| + | 1.2147 | 0.613438 | Q9NT62 | Q9NT62 | ATG3 | Ubiquitin-l | Color2 [Da | 1 | 4 |
| + | 2.016133 | 0.721643 | Q9NT68 | Q9NT68 | TENM2 | Teneurin-2 | Color2 [Da | 1 | 4 |
| | 0.570176 | -0.86308 | Q9NT99 | Q9NT99 | LRRC4B | Leucine-ric | Color2 [Da | 1 | 4 |
| | 0.52917 | 0.217697 | Q9NTG7 | Q9NTG7 | SIRT3 | NAD-depe | Color2 [Da | 1 | 4 |
| | 0 | NaN | Q9NTI2 | Q9NTI2 | ATP8A2 | Phospholi | Color2 [Da | 1 | 4 |
| + | 1.976115 | -0.75353 | Q9NTI5 | Q9NTI5 | PDS5B | Sister chro | Color2 [Da | 1 | 4 |
| | 0.225769 | -0.37365 | Q9NTJ3 | Q9NTJ3 | SMC4 | Structural | Color2 [Da | 1 | 4 |
| + | 1.2427 | -0.61819 | Q9NTJ4 | Q9NTJ4 | MAN2C1 | Alpha-mar | Color2 [Da | 1 | 4 |
| + | 1.847733 | -1.06033 | Q9NTJ5 | Q9NTJ5 | SACM1L | Phosphatic | Color2 [Da | 1 | 4 |
| | 0.115769 | 0.032611 | Q9NTK5 | Q9NTK5 | OLA1 | Obg-like A | Color2 [Da | 1 | 4 |
| + | 3.522873 | 0.819826 | Q9NTX5 | Q9NTX5 | ECHDC1 | Ethylmalor | Color2 [Da | 1 | 4 |
| + | 1.324849 | -0.35212 | Q9NTX7 | Q9NTX7 | RNF146 | E3 ubiquiti | Color2 [Da | 1 | 4 |
| | 0.018531 | 0.006046 | Q9NTZ6 | Q9NTZ6 | RBM12 | RNA-bindi | Color2 [Da | 1 | 4 |
| + | 1.465624 | -0.89194 | Q9NU22 | Q9NU22 | MDN1 | Midasin | Color2 [Da | 1 | 4 |
| + | 2.467125 | 2.563771 | Q9NU23 | Q9NU23 | LYRM2 | LYR motif- | Color2 [Da | 1 | 4 |
| | 0 | -1.54884 | Q9NUB1 | Q9NUB1 | ACSS1 | Acetyl-coe | Color2 [Da | 1 | 4 |
| + | 1.091857 | -0.37801 | Q9NUD5 | Q9NUD5 | ZCCHC3 | Zinc finger | Color2 [Da | 1 | 4 |
| | 0 | NaN | Q9NUE0 | Q9NUE0 | ZDHHC18 | Palmitoylt | Color2 [Da | 1 | 4 |
| + | 1.376551 | 1.217772 | Q9NUG6 | Q9NUG6 | PDRG1 | p53 and D | Color2 [Da | 1 | 4 |
| + | 1.298693 | -1.46812 | Q9NUI1 | Q9NUI1 | DECR2 | Peroxisom | Color2 [Da | 1 | 4 |
| + | 2.442196 | 1.667948 | Q9NUJ1 | Q9NUJ1 | ABHD10 | Palmitoyl-| | Color2 [Da | 1 | 4 |
| + | 1.84892 | -0.62634 | Q9NUJ3 | Q9NUJ3 | TCP11L1 | T-complex | Color2 [Da | 1 | 4 |
| + | 2.45188 | 0.821504 | Q9NUL3 | Q9NUL3 | STAU2 | Double-str | Color2 [Da | 1 | 4 |
| + | 1.171911 | 1.789081 | Q9NUL5 | Q9NUL5 | SHFL | Shiftless ar | Color2 [Da | 1 | 4 |
| | 0 | NaN | Q9NUL7 | Q9NUL7 | DDX28 | Probable A | Color2 [Da | 1 | 4 |
| + | 1.313856 | 2.067575 | Q9NUM4 | Q9NUM4 | TMEM106 | Transmem | Color2 [Da | 1 | 4 |
| | 0 | NaN | Q9NUN5 | Q9NUN5 | LMBRD1 | Lysosomal | Color2 [Da | 1 | 4 |
| + | 2.391355 | 0.851567 | Q9NUP1 | Q9NUP1 | BLOC1S4 | Biogenesis | Color2 [Da | 1 | 4 |

| | | | | | | | | | |
|---|---|---|---|---|---|---|---|---|---|
| + | 2.479055 | 1.915493 | Q9NUP9 | Q9NUP9 | LIN7C | Protein lin | Color2 [Da | 1 | 4 |
| + | 1.660475 | -1.00314 | Q9NUQ2 | Q9NUQ2 | AGPAT5 | 1-acyl-sn-g | Color2 [Da | 1 | 4 |
| + | 1.200231 | -0.36842 | Q9NUQ6 | Q9NUQ6 | SPATS2L | SPATS2-lik | Color2 [Da | 1 | 4 |
| + | 1.790262 | -0.85078 | Q9NUQ7 | Q9NUQ7 | UFSP2 | Ufm1-spec | Color2 [Da | 1 | 4 |
| | 0.202495 | -0.06351 | Q9NUQ8 | Q9NUQ8 | ABCF3 | ATP-bindir | Color2 [Da | 1 | 4 |
| | 0.975229 | -0.09817 | Q9NUQ9 | Q9NUQ9 | CYRIB | CYFIP-relat | Color2 [Da | 1 | 4 |
| + | 0.873007 | -0.50006 | Q9NUT2 | Q9NUT2 | ABCB8 | Mitochonc | Color2 [Da | 1 | 4 |
| + | 1.828584 | -0.67827 | Q9NUU7 | Q9NUU7 | DDX19A | ATP-deper | Color2 [Da | 1 | 4 |
| + | 1.766113 | 0.766166 | Q9NUW8 | Q9NUW8 | TDP1 | Tyrosyl-DN | Color2 [Da | 1 | 4 |
| | 0 | -1.47082 | Q9NUX5 | Q9NUX5 | POT1 | Protection | Color2 [Da | 1 | 4 |
| + | 1.941671 | 0.47587 | Q9NV31 | Q9NV31 | IMP3 | U3 small n | Color2 [Da | 1 | 4 |
| | 0.034746 | -0.0784 | Q9NV35 | Q9NV35 | NUDT15 | Nucleotide | Color2 [Da | 1 | 4 |
| + | 2.478899 | 1.6666 | Q9NV56 | Q9NV56 | MRGBP | MRG/MOF | Color2 [Da | 1 | 4 |
| + | 1.883376 | -1.05859 | Q9NV70 | Q9NV70 | EXOC1 | Exocyst co | Color2 [Da | 1 | 4 |
| | 0.348716 | 0.220264 | Q9NV88 | Q9NV88 | INTS9 | Integrator | Color2 [Da | 1 | 4 |
| + | 0.862444 | -0.38759 | Q9NV96 | Q9NV96 | TMEM30A | Cell cycle c | Color2 [Da | 1 | 4 |
| + | 1.019418 | -1.07385 | Q9NVA1 | Q9NVA1 | UQCC1 | Ubiquinol- | Color2 [Da | 1 | 4 |
| | 1.059351 | 0.258507 | Q9NVA2 | Q9NVA2 | SEPTIN11 | Septin-11 | Color2 [Da | 1 | 4 |
| | 0 | NaN | Q9NVA4 | Q9NVA4 | TMEM184 | Transmem | Color2 [Da | 1 | 4 |
| | 0 | NaN | Q9NVC6 | Q9NVC6 | MED17 | Mediator c | Color2 [Da | 1 | 4 |
| + | 1.896367 | -1.31751 | Q9NVD7 | Q9NVD7 | PARVA | Alpha-parv | Color2 [Da | 1 | 4 |
| + | 2.788653 | -0.62421 | Q9NVE7 | Q9NVE7 | PANK4 | 4'-phosphc | Color2 [Da | 1 | 4 |
| + | 1.442925 | -0.27536 | Q9NVG8 | Q9NVG8 | TBC1D13 | TBC1 dom | Color2 [Da | 1 | 4 |
| + | 1.765505 | -0.61147 | Q9NVH0 | Q9NVH0 | EXD2 | Exonuclea | Color2 [Da | 1 | 4 |
| + | 1.18312 | -0.26329 | Q9NVH1 | Q9NVH1 | DNAJC11 | DnaJ homc | Color2 [Da | 1 | 4 |
| | 0.39426 | -0.12684 | Q9NVH2 | Q9NVH2 | INTS7 | Integrator | Color2 [Da | 1 | 4 |
| + | 1.557716 | -1.92308 | Q9NVH6 | Q9NVH6 | TMLHE | Trimethyll | Color2 [Da | 1 | 4 |
| + | 1.494303 | 0.869582 | Q9NVI7 | Q9NVI7 | ATAD3A | ATPase far | Color2 [Da | 1 | 4 |
| + | 1.686309 | -0.50539 | Q9NVJ2 | Q9NVJ2 | ARL8B | ADP-ribosy | Color2 [Da | 1 | 4 |
| | 0.292147 | 0.282713 | Q9NVK5 | Q9NVK5 | FGFR1OP2 | FGFR1 onc | Color2 [Da | 1 | 4 |
| | 0.128798 | 0.210604 | Q9NVM6 | Q9NVM6 | DNAJC17 | DnaJ homc | Color2 [Da | 1 | 4 |
| | 0.411597 | 0.332946 | Q9NVM9 | Q9NVM9 | INTS13 | Integrator | Color2 [Da | 1 | 4 |
| + | 2.229546 | -2.12798 | Q9NVN8 | Q9NVN8 | GNL3L | Guanine n | Color2 [Da | 1 | 4 |
| | 0.689133 | -0.17998 | Q9NVP1 | Q9NVP1 | DDX18 | ATP-deper | Color2 [Da | 1 | 4 |
| | 0.771451 | -0.2819 | Q9NVR2 | Q9NVR2 | INTS10 | Integrator | Color2 [Da | 1 | 4 |
| | 0.301481 | -0.2754 | Q9NVS2 | Q9NVS2 | MRPS18A | 39S riboso | Color2 [Da | 1 | 4 |
| + | 1.970825 | 1.804881 | Q9NVS9 | Q9NVS9 | PNPO | Pyridoxine | Color2 [Da | 1 | 4 |
| + | 2.371291 | 0.794893 | Q9NVT9 | Q9NVT9 | ARMC1 | Armadillo | Color2 [Da | 1 | 4 |
| | 0 | NaN | Q9NVU7 | Q9NVU7 | SDAD1 | Protein SD | Color2 [Da | 1 | 4 |
| | 0 | 0.302534 | Q9NVV0 | Q9NVV0 | TMEM38B | Trimeric in | Color2 [Da | 1 | 4 |
| | 0 | -1.97092 | Q9NVV4 | Q9NVV4 | MTPAP | Poly(A) RN | Color2 [Da | 1 | 4 |
| | 0.461029 | 0.483942 | Q9NVW2 | Q9NVW2 | RLIM | E3 ubiquiti | Color2 [Da | 1 | 4 |
| | 0 | NaN | Q9NVX2 | Q9NVX2 | NLE1 | Notchless | Color2 [Da | 1 | 4 |
| | 0.716267 | -0.43929 | Q9NVZ3 | Q9NVZ3 | NECAP2 | Adaptin ea | Color2 [Da | 1 | 4 |
| | 0.498628 | 0.570263 | Q9NW13 | Q9NW13 | RBM28 | RNA-bindi | Color2 [Da | 1 | 4 |
| + | 1.184117 | -1.47395 | Q9NW15 | Q9NW15 | ANO10 | Anoctamir | Color2 [Da | 1 | 4 |
| | 0.526638 | -0.30543 | Q9NW64 | Q9NW64 | RBM22 | Pre-mRNA | Color2 [Da | 1 | 4 |

| | | | | | | | | | | |
|---|---|---|---|---|---|---|---|---|---|---|
| + | 2.790227 | -0.50411 | Q9NW68 | Q9NW68 | BSDC1 | BSD doma | Color2 | [Da | 1 | 4 |
| + | 1.450019 | -1.1215 | Q9NW82 | Q9NW82 | WDR70 | WD repeat | Color2 | [Da | 1 | 4 |
| | 0.31449 | 0.559062 | Q9NWA0 | Q9NWA0 | MED9 | Mediator ( | Color2 | [Da | 1 | 4 |
| | 0.528023 | 1.378615 | Q9NWB1 | Q9NWB1 | RBFOX1 | RNA bindir | Color2 | [Da | 1 | 4 |
| + | 0.939642 | 0.646597 | Q9NWB6 | Q9NWB6 | ARGLU1 | Arginine ai | Color2 | [Da | 1 | 4 |
| + | 1.498225 | 0.562877 | Q9NWB7 | Q9NWB7 | IFT57 | Intraflagell | Color2 | [Da | 1 | 4 |
| + | 1.351204 | 1.465025 | Q9NWH9 | Q9NWH9 | SLTM | SAFB-like t | Color2 | [Da | 1 | 4 |
| | 0.353445 | 0.830497 | Q9NWM8 | Q9NWM8 | FKBP14 | Peptidyl-pı | Color2 | [Da | 1 | 4 |
| + | 1.90519 | 2.37991 | Q9NWQ8 | Q9NWQ8 | PAG1 | Phosphopı | Color2 | [Da | 1 | 4 |
| | 0.599883 | 0.237513 | Q9NWS0 | Q9NWS0 | PIH1D1 | PIH1 doma | Color2 | [Da | 1 | 4 |
| | 0 | NaN | Q9NWS6 | Q9NWS6 | FAM118A | Protein FA | Color2 | [Da | 1 | 4 |
| | 0.653834 | -0.17722 | Q9NWS8 | Q9NWS8 | RMND1 | Required f | Color2 | [Da | 1 | 4 |
| | 0.515665 | -0.34851 | Q9NWT1 | Q9NWT1 | PAK1IP1 | p21-activa | Color2 | [Da | 1 | 4 |
| + | 1.429608 | -0.69797 | Q9NWT6 | Q9NWT6 | HIF1AN | Hypoxia-in | Color2 | [Da | 1 | 4 |
| | 0.797759 | -0.2973 | Q9NWU1 | Q9NWU1 | OXSM | 3-oxoacyl- | Color2 | [Da | 1 | 4 |
| + | 1.609956 | 0.595389 | Q9NWU2 | Q9NWU2 | GID8 | Glucose-in | Color2 | [Da | 1 | 4 |
| | 0.134334 | 0.08336 | Q9NWU5 | Q9NWU5 | MRPL22 | 39S riboso | Color2 | [Da | 1 | 4 |
| | 0.205161 | 0.313346 | Q9NWV4 | Q9NWV4 | CZIB | CXXC moti | Color2 | [Da | 1 | 4 |
| + | 1.09499 | 0.718066 | Q9NWV8 | Q9NWV8 | BABAM1 | BRISC and | Color2 | [Da | 1 | 4 |
| | 0.275772 | -0.45381 | Q9NWX6 | Q9NWX6 | THG1L | Probable t | Color2 | [Da | 1 | 4 |
| + | 1.256604 | -0.79308 | Q9NWY4 | Q9NWY4 | HPF1 | Histone PA | Color2 | [Da | 1 | 4 |
| + | 1.10836 | -0.45006 | Q9NWZ5 | Q9NWZ5 | UCKL1 | Uridine-cy | Color2 | [Da | 1 | 4 |
| | 0 | -0.29865 | Q9NX00 | Q9NX00 | TMEM160 | Transmem | Color2 | [Da | 1 | 4 |
| + | 2.084476 | 1.53332 | Q9NX01 | Q9NX01 | TXNL4B | Thioredoxi | Color2 | [Da | 1 | 4 |
| | 0 | NaN | Q9NX02 | Q9NX02 | NLRP2 | NACHT, LR | Color2 | [Da | 1 | 4 |
| | 0 | -0.18467 | Q9NX05 | Q9NX05 | FAM120C | Constitutiv | Color2 | [Da | 1 | 4 |
| + | 1.288022 | 0.690216 | Q9NX08 | Q9NX08 | COMMD8 | COMM do | Color2 | [Da | 1 | 4 |
| | 0.218735 | -0.73183 | Q9NX14 | Q9NX14 | NDUFB11 | NADH deh | Color2 | [Da | 1 | 4 |
| + | 1.56198 | 2.318681 | Q9NX18 | Q9NX18 | SDHAF2 | Succinate ( | Color2 | [Da | 1 | 4 |
| + | 1.706751 | 1.413851 | Q9NX20 | Q9NX20 | MRPL16 | 39S riboso | Color2 | [Da | 1 | 4 |
| | 0.002385 | -0.00728 | Q9NX24 | Q9NX24 | NHP2 | H/ACA ribo | Color2 | [Da | 1 | 4 |
| + | 2.921195 | 1.480819 | Q9NX40 | Q9NX40 | OCIAD1 | OCIA dom | Color2 | [Da | 1 | 4 |
| | 0.414899 | -0.07332 | Q9NX46 | Q9NX46 | ADPRS | ADP-ribos | Color2 | [Da | 1 | 4 |
| + | 1.417208 | 1.278675 | Q9NX47 | Q9NX47 | MARCHF5 | E3 ubiquiti | Color2 | [Da | 1 | 4 |
| + | 3.855681 | 2.980246 | Q9NX55 | Q9NX55 | HYPK | Huntingtin | Color2 | [Da | 1 | 4 |
| + | 1.402726 | -0.36605 | Q9NX58 | Q9NX58 | LYAR | Cell growtl | Color2 | [Da | 1 | 4 |
| | 0.518114 | 0.700169 | Q9NX62 | Q9NX62 | BPNT2 | Golgi-resic | Color2 | [Da | 1 | 4 |
| + | 2.248957 | 0.657553 | Q9NX63 | Q9NX63 | CHCHD3 | MICOS cor | Color2 | [Da | 1 | 4 |
| | 0.258994 | 0.3865 | Q9NX70 | Q9NX70 | MED29 | Mediator ( | Color2 | [Da | 1 | 4 |
| + | 3.825953 | -0.6807 | Q9NX74 | Q9NX74 | DUS2 | tRNA-dihy | Color2 | [Da | 1 | 4 |
| | 0 | NaN | Q9NX76 | Q9NX76 | CMTM6 | CKLF-like N | Color2 | [Da | 1 | 4 |
| | 0.613294 | 0.824751 | Q9NX94 | Q9NX94 | WBP1L | WW doma | Color2 | [Da | 1 | 4 |
| + | 2.871232 | -0.66394 | Q9NX95 | Q9NX95 | SYBU | Syntabulin | Color2 | [Da | 1 | 4 |
| + | 2.151222 | -0.86969 | Q9NXA8 | Q9NXA8 | SIRT5 | NAD-depe | Color2 | [Da | 1 | 4 |
| + | 2.927851 | -0.9758 | Q9NXC5 | Q9NXC5 | MIOS | GATOR coı | Color2 | [Da | 1 | 4 |
| + | 2.225781 | -1.34395 | Q9NXE4 | Q9NXE4 | SMPD4 | Sphingomy | Color2 | [Da | 1 | 4 |
| + | 1.125135 | -0.31623 | Q9NXF1 | Q9NXF1 | TEX10 | Testis-expı | Color2 | [Da | 1 | 4 |

| | | | | | | | | | |
|---|---|---|---|---|---|---|---|---|---|
| + | 2.485303 | -0.85079 | Q9NXG2 | Q9NXG2 | THUMPD1 | THUMP dc | Color2 [Da | 1 | 4 |
| | 0.761723 | 0.309525 | Q9NXG6 | Q9NXG6 | P4HTM | Transmem | Color2 [Da | 1 | 4 |
| + | 3.642531 | -1.41284 | Q9NXH9 | Q9NXH9 | TRMT1 | tRNA (guai | Color2 [Da | 1 | 4 |
| | 0.21198 | -0.17609 | Q9NXN4 | Q9NXN4 | GDAP2 | Gangliosid | Color2 [Da | 1 | 4 |
| | 0.11132 | 0.184042 | Q9NXR1 | Q9NXR1 | NDE1 | Nuclear di: | Color2 [Da | 1 | 4 |
| + | 0.863625 | -1.09589 | Q9NXR7 | Q9NXR7 | BABAM2 | BRISC and | Color2 [Da | 1 | 4 |
| | 0.130772 | -0.17287 | Q9NXR8 | Q9NXR8 | ING3 | Inhibitor o | Color2 [Da | 1 | 4 |
| | 0.194177 | 0.198196 | Q9NXS2 | Q9NXS2 | QPCTL | Glutaminy | Color2 [Da | 1 | 4 |
| + | 1.504168 | 1.140318 | Q9NXU5 | Q9NXU5 | ARL15 | ADP-ribosy | Color2 [Da | 1 | 4 |
| | 0.730053 | -0.21011 | Q9NXV2 | Q9NXV2 | KCTD5 | BTB/POZ d | Color2 [Da | 1 | 4 |
| | 0.968145 | 0.313355 | Q9NXV6 | Q9NXV6 | CDKN2AIP | CDKN2A-ir | Color2 [Da | 1 | 4 |
| + | 1.86869 | 0.549263 | Q9NXX6 | Q9NXX6 | NSMCE4A | Non-struct | Color2 [Da | 1 | 4 |
| | 0.482498 | 0.385044 | Q9NY12 | Q9NY12 | GAR1 | H/ACA ribo | Color2 [Da | 1 | 4 |
| | 0.135756 | -0.0785 | Q9NY27 | Q9NY27 | PPP4R2 | Serine/thr | Color2 [Da | 1 | 4 |
| + | 1.212947 | 0.277595 | Q9NY33 | Q9NY33 | DPP3 | Dipeptidyl | Color2 [Da | 1 | 4 |
| | 0 | NaN | Q9NY35 | Q9NY35 | CLDND1 | Claudin do | Color2 [Da | 1 | 4 |
| | 0.103022 | -0.05 | Q9NY46 | Q9NY46 | SCN3A | Sodium ch | Color2 [Da | 1 | 4 |
| | 0.859445 | 0.222388 | Q9NY47 | Q9NY47 | CACNA2D2 | Voltage-de | Color2 [Da | 1 | 4 |
| + | 2.199984 | 1.521122 | Q9NY59 | Q9NY59 | SMPD3 | Sphingomy | Color2 [Da | 1 | 4 |
| + | 2.176317 | 0.56054 | Q9NY61 | Q9NY61 | AATF | Protein AA | Color2 [Da | 1 | 4 |
| | 0.070615 | 0.099616 | Q9NY72 | Q9NY72 | SCN3B | Sodium ch | Color2 [Da | 1 | 4 |
| | 0 | -0.27951 | Q9NY93 | Q9NY93 | DDX56 | Probable A | Color2 [Da | 1 | 4 |
| + | 1.931646 | 0.579571 | Q9NYB0 | Q9NYB0 | TERF2IP | Telomeric | Color2 [Da | 1 | 4 |
| + | 3.541328 | 1.922643 | Q9NYB9 | Q9NYB9 | ABI2 | Abl interac | Color2 [Da | 1 | 4 |
| + | 2.603878 | 0.725698 | Q9NYF8 | Q9NYF8 | BCLAF1 | Bcl-2-asso | Color2 [Da | 1 | 4 |
| | 0 | NaN | Q9NYG2 | Q9NYG2 | ZDHHC3 | Palmitoylt | Color2 [Da | 1 | 4 |
| | 0 | -0.17226 | Q9NYH9 | Q9NYH9 | UTP6 | U3 small n | Color2 [Da | 1 | 4 |
| + | 2.252782 | 1.094359 | Q9NYI0 | Q9NYI0 | PSD3 | PH and SE( | Color2 [Da | 1 | 4 |
| + | 1.946229 | 3.042789 | Q9NYJ1 | Q9NYJ1 | COA4 | Cytochron | Color2 [Da | 1 | 4 |
| | 0 | -2.15552 | Q9NYJ8 | Q9NYJ8 | TAB2 | TGF-beta-a | Color2 [Da | 1 | 4 |
| + | 2.19346 | -0.92469 | Q9NYK5 | Q9NYK5 | MRPL39 | 39S riboso | Color2 [Da | 1 | 4 |
| + | 1.610962 | 0.499806 | Q9NYL9 | Q9NYL9 | TMOD3 | Tropomod | Color2 [Da | 1 | 4 |
| + | 1.378532 | -0.88591 | Q9NYQ6 | Q9NYQ6 | CELSR1 | Cadherin E | Color2 [Da | 1 | 4 |
| | 0.650067 | -0.13797 | Q9NYQ7 | Q9NYQ7 | CELSR3 | Cadherin E | Color2 [Da | 1 | 4 |
| | 0.736286 | 0.122779 | Q9NYR9 | Q9NYR9 | NKIRAS2 | NF-kappa- | Color2 [Da | 1 | 4 |
| | 0.399841 | 0.404215 | Q9NYS0 | Q9NYS0 | NKIRAS1 | NF-kappa- | Color2 [Da | 1 | 4 |
| | 0 | NaN | Q9NYS7 | Q9NYS7 | WSB2 | WD repeat | Color2 [Da | 1 | 4 |
| + | 3.865716 | -1.10054 | Q9NYU2 | Q9NYU2 | UGGT1 | UDP-gluco | Color2 [Da | 1 | 4 |
| + | 1.962214 | -1.38456 | Q9NYV4 | Q9NYV4 | CDK12 | Cyclin-dep | Color2 [Da | 1 | 4 |
| | 0 | NaN | Q9NYY8 | Q9NYY8 | FASTKD2 | FAST kinas | Color2 [Da | 1 | 4 |
| + | 2.847314 | -1.79956 | Q9NZ01 | Q9NZ01 | TECR | Very-long- | Color2 [Da | 1 | 4 |
| + | 3.560247 | -2.11673 | Q9NZ08 | Q9NZ08 | ERAP1 | Endoplasm | Color2 [Da | 1 | 4 |
| + | 1.920291 | -0.72621 | Q9NZ32 | Q9NZ32 | ACTR10 | Actin-relat | Color2 [Da | 1 | 4 |
| | 0 | NaN | Q9NZ42 | Q9NZ42 | PSENEN | Gamma-se | Color2 [Da | 1 | 4 |
| | 0.048272 | 0.026312 | Q9NZ43 | Q9NZ43 | USE1 | Vesicle tra | Color2 [Da | 1 | 4 |
| + | 2.978189 | 1.209258 | Q9NZ45 | Q9NZ45 | CISD1 | CDGSH iro | Color2 [Da | 1 | 4 |
| | 0.519231 | 0.51683 | Q9NZ52 | Q9NZ52 | GGA3 | ADP-ribosy | Color2 [Da | 1 | 4 |

| | | | | | | | | | |
|---|---|---|---|---|---|---|---|---|---|
| | 0.6416 | 0.554821 | Q9NZ53 | Q9NZ53 | PODXL2 | Podocalyxi | Color2 [Da | 1 | 4 |
| + | 1.940098 | 1.271952 | Q9NZ56 | Q9NZ56 | FMN2 | Formin-2 | Color2 [Da | 1 | 4 |
| + | 1.387978 | -2.55921 | Q9NZ63 | Q9NZ63 | C9orf78 | Telomere l | Color2 [Da | 1 | 4 |
| + | 3.609891 | 2.988784 | Q9NZ72 | Q9NZ72 | STMN3 | Stathmin-3 | Color2 [Da | 1 | 4 |
| | 1.422106 | -0.15533 | Q9NZ94 | Q9NZ94 | NLGN3 | Neuroligin | Color2 [Da | 1 | 4 |
| | 0.14828 | 0.045123 | Q9NZB2 | Q9NZB2 | FAM120A | Constitutiv | Color2 [Da | 1 | 4 |
| | 0 | NaN | Q9NZC9 | Q9NZC9 | SMARCAL1 | SWI/SNF-r | Color2 [Da | 1 | 4 |
| + | 1.240027 | -0.47365 | Q9NZD2 | Q9NZD2 | GLTP | Glycolipid | Color2 [Da | 1 | 4 |
| | 0.562354 | -1.05711 | Q9NZD8 | Q9NZD8 | SPG21 | Maspardin | Color2 [Da | 1 | 4 |
| | 0 | NaN | Q9NZH8 | Q9NZH8 | IL36G | Interleukir | Color2 [Da | 1 | 4 |
| | 0 | 0.053317 | Q9NZI7 | Q9NZI7 | UBP1 | Upstream- | Color2 [Da | 1 | 4 |
| + | 1.85579 | -0.58552 | Q9NZI8 | Q9NZI8 | IGF2BP1 | Insulin-like | Color2 [Da | 1 | 4 |
| + | 2.565563 | -0.80348 | Q9NZJ4 | Q9NZJ4 | SACS | Sacsin | Color2 [Da | 1 | 4 |
| | 0 | -1.07645 | Q9NZJ5 | Q9NZJ5 | EIF2AK3 | Eukaryotic | Color2 [Da | 1 | 4 |
| + | 0.985766 | -0.55387 | Q9NZJ7 | Q9NZJ7 | MTCH1 | Mitochonc | Color2 [Da | 1 | 4 |
| | 0.318324 | 0.755137 | Q9NZJ9 | Q9NZJ9 | NUDT4 | Diphospho | Color2 [Da | 1 | 4 |
| + | 2.450937 | -0.83896 | Q9NZL4 | Q9NZL4 | HSPBP1 | Hsp70-bin | Color2 [Da | 1 | 4 |
| | 0.619396 | -0.22211 | Q9NZL9 | Q9NZL9 | MAT2B | Methionin | Color2 [Da | 1 | 4 |
| + | 2.103318 | -1.57412 | Q9NZM1 | Q9NZM1 | MYOF | Myoferlin | Color2 [Da | 1 | 4 |
| + | 1.499153 | 0.804985 | Q9NZM3 | Q9NZM3 | ITSN2 | Intersectin | Color2 [Da | 1 | 4 |
| + | 0.866646 | 0.823828 | Q9NZM4 | Q9NZM4 | BICRA | BRD4-inter | Color2 [Da | 1 | 4 |
| | 0.384437 | -0.63506 | Q9NZM5 | Q9NZM5 | NOP53 | Ribosome | Color2 [Da | 1 | 4 |
| + | 1.88792 | 0.607422 | Q9NZN3 | Q9NZN3 | EHD3 | EH domair | Color2 [Da | 1 | 4 |
| | 0.28839 | 0.095645 | Q9NZN5 | Q9NZN5 | ARHGEF12 | Rho guanir | Color2 [Da | 1 | 4 |
| | 0.229483 | -0.02903 | Q9NZN8 | Q9NZN8 | CNOT2 | CCR4-NOT | Color2 [Da | 1 | 4 |
| | 0.206577 | -0.07174 | Q9NZQ3 | Q9NZQ3 | NCKIPSD | NCK-intera | Color2 [Da | 1 | 4 |
| + | 4.428935 | 2.622098 | Q9NZR1 | Q9NZR1 | TMOD2 | Tropomod | Color2 [Da | 1 | 4 |
| + | 1.962904 | 0.984972 | Q9NZT1 | Q9NZT1 | CALML5 | Calmodulir | Color2 [Da | 1 | 4 |
| + | 2.499205 | 1.539116 | Q9NZT2 | Q9NZT2 | OGFR | Opioid gro | Color2 [Da | 1 | 4 |
| + | 2.211179 | -2.1798 | Q9NZU0 | Q9NZU0 | FLRT3 | Leucine-ric | Color2 [Da | 1 | 4 |
| + | 1.434949 | 0.713948 | Q9NZU5 | Q9NZU5 | LMCD1 | LIM and cy | Color2 [Da | 1 | 4 |
| | 0 | NaN | Q9NZV8 | Q9NZV8 | KCND2 | Potassium | Color2 [Da | 1 | 4 |
| | 0.841802 | 0.219745 | Q9NZW5 | Q9NZW5 | PALS2 | Protein PA | Color2 [Da | 1 | 4 |
| + | 3.494886 | 2.311907 | Q9NZZ3 | Q9NZZ3 | CHMP5 | Charged m | Color2 [Da | 1 | 4 |
| + | 2.369512 | 0.992724 | Q9P000 | Q9P000 | COMMD9 | COMM do | Color2 [Da | 1 | 4 |
| | 0 | -2.40133 | Q9P003 | Q9P003 | CNIH4 | Protein co | Color2 [Da | 1 | 4 |
| | 0.50176 | 0.5445 | Q9P013 | Q9P013 | CWC15 | Spliceoson | Color2 [Da | 1 | 4 |
| + | 1.445473 | 0.479884 | Q9P015 | Q9P015 | MRPL15 | 39S riboso | Color2 [Da | 1 | 4 |
| + | 1.237902 | 0.314709 | Q9P016 | Q9P016 | THYN1 | Thymocyte | Color2 [Da | 1 | 4 |
| + | 2.612889 | 0.684636 | Q9P032 | Q9P032 | NDUFAF4 | NADH deh | Color2 [Da | 1 | 4 |
| + | 3.670797 | -1.47711 | Q9P035 | Q9P035 | HACD3 | Very-long- | Color2 [Da | 1 | 4 |
| | 0.24618 | -0.18559 | Q9P086 | Q9P086 | MED11 | Mediator c | Color2 [Da | 1 | 4 |
| + | 1.148119 | 0.589413 | Q9P0I2 | Q9P0I2 | EMC3 | ER membr | Color2 [Da | 1 | 4 |
| + | 2.608238 | 1.777536 | Q9P0J0 | Q9P0J0 | NDUFA13 | NADH deh | Color2 [Da | 1 | 4 |
| + | 2.585637 | 0.40656 | Q9P0J1 | Q9P0J1 | PDP1 | [Pyruvate c | Color2 [Da | 1 | 4 |
| | 0.743656 | 0.337097 | Q9P0J7 | Q9P0J7 | KCMF1 | E3 ubiquiti | Color2 [Da | 1 | 4 |
| | 0.39471 | 0.418217 | Q9P0K1 | Q9P0K1 | ADAM22 | Disintegrir | Color2 [Da | 1 | 4 |

| | | | | | | | | | |
|---|---|---|---|---|---|---|---|---|---|
| + | 2.29935 | -0.7896 | Q9P0K7 | Q9P0K7 | RAI14 | Ankycorbi | Color2 [Da | 1 | 4 |
| | 0 | NaN | Q9P0K9 | Q9P0K9 | FRRS1L | DOMON d | Color2 [Da | 1 | 4 |
| + | 2.355342 | 0.793613 | Q9P0L0 | Q9P0L0 | VAPA | Vesicle-ass | Color2 [Da | 1 | 4 |
| | 0.354459 | 0.091113 | Q9P0L2 | Q9P0L2 | MARK1 | Serine/thr | Color2 [Da | 1 | 4 |
| | 0.856511 | 0.265504 | Q9P0M6 | Q9P0M6 | MACROH2 | Core histo | Color2 [Da | 1 | 4 |
| + | 1.446786 | 0.83856 | Q9P0M9 | Q9P0M9 | MRPL27 | 39S riboso | Color2 [Da | 1 | 4 |
| | 0 | -0.08384 | Q9P0N9 | Q9P0N9 | TBC1D7 | TBC1 dom | Color2 [Da | 1 | 4 |
| | 0.611438 | 0.517018 | Q9P0P0 | Q9P0P0 | RNF181 | E3 ubiquiti | Color2 [Da | 1 | 4 |
| + | 2.54976 | 1.881609 | Q9P0P8 | Q9P0P8 | MTRES1 | Mitochonc | Color2 [Da | 1 | 4 |
| + | 2.371561 | 2.022974 | Q9P0S2 | Q9P0S2 | COX16 | Cytochrom | Color2 [Da | 1 | 4 |
| + | 2.634049 | 1.829964 | Q9P0S3 | Q9P0S3 | ORMDL1 | ORM1-like | Color2 [Da | 1 | 4 |
| + | 3.594111 | 2.610872 | Q9P0S9 | Q9P0S9 | TMEM14C | Transmem | Color2 [Da | 1 | 4 |
| | 0.141841 | 0.406149 | Q9P0U1 | Q9P0U1 | TOMM7 | Mitochonc | Color2 [Da | 1 | 4 |
| + | 1.258626 | -0.93891 | Q9P0U4 | Q9P0U4 | CXXC1 | CXXC-type | Color2 [Da | 1 | 4 |
| + | 2.557762 | -1.19723 | Q9P0V3 | Q9P0V3 | SH3BP4 | SH3 domai | Color2 [Da | 1 | 4 |
| | 0.539318 | -0.40305 | Q9P0V9 | Q9P0V9 | SEPTIN10 | Septin-10 | Color2 [Da | 1 | 4 |
| | 0.643674 | -0.99212 | Q9P0Z9 | Q9P0Z9 | PIPOX | Peroxisom | Color2 [Da | 1 | 4 |
| + | 1.386162 | -1.62753 | Q9P107 | Q9P107 | GMIP | GEM-inter | Color2 [Da | 1 | 4 |
| + | 1.926487 | 2.196397 | Q9P121 | Q9P121 | NTM | Neurotrim | Color2 [Da | 1 | 4 |
| + | 1.790366 | 0.584814 | Q9P1F3 | Q9P1F3 | ABRACL | Costars fa | Color2 [Da | 1 | 4 |
| | 0.49353 | -0.72014 | Q9P1U1 | Q9P1U1 | ACTR3B | Actin-relat | Color2 [Da | 1 | 4 |
| + | 2.21863 | 0.696061 | Q9P1Y5 | Q9P1Y5 | CAMSAP3 | Calmoduli | Color2 [Da | 1 | 4 |
| | 0.140178 | -0.05909 | Q9P1Y6 | Q9P1Y6 | PHRF1 | PHD and R | Color2 [Da | 1 | 4 |
| | 0.257713 | 0.309122 | Q9P219 | Q9P219 | CCDC88C | Protein Da | Color2 [Da | 1 | 4 |
| + | 2.348638 | 1.511972 | Q9P227 | Q9P227 | ARHGAP23 | Rho GTPas | Color2 [Da | 1 | 4 |
| + | 1.181915 | 1.278803 | Q9P242 | Q9P242 | NYAP2 | Neuronal t | Color2 [Da | 1 | 4 |
| + | 2.112184 | 0.656404 | Q9P244 | Q9P244 | LRFN1 | Leucine-ric | Color2 [Da | 1 | 4 |
| + | 1.131773 | -0.57908 | Q9P246 | Q9P246 | STIM2 | Stromal in | Color2 [Da | 1 | 4 |
| + | 1.580488 | -0.8572 | Q9P253 | Q9P253 | VPS18 | Vacuolar p | Color2 [Da | 1 | 4 |
| + | 5.378694 | -1.26371 | Q9P258 | Q9P258 | RCC2 | Protein RC | Color2 [Da | 1 | 4 |
| + | 2.787312 | 0.639346 | Q9P260 | Q9P260 | RELCH | RAB11-bin | Color2 [Da | 1 | 4 |
| | 0.009981 | 0.005014 | Q9P265 | Q9P265 | DIP2B | Disco-inte | Color2 [Da | 1 | 4 |
| + | 2.040549 | 1.486511 | Q9P270 | Q9P270 | SLAIN2 | SLAIN mot | Color2 [Da | 1 | 4 |
| + | 1.254388 | 0.253382 | Q9P286 | Q9P286 | PAK5 | Serine/thr | Color2 [Da | 1 | 4 |
| + | 1.40767 | -0.36693 | Q9P287 | Q9P287 | BCCIP | BRCA2 anc | Color2 [Da | 1 | 4 |
| | 0 | NaN | Q9P291 | Q9P291 | ARMCX1 | Armadillo | Color2 [Da | 1 | 4 |
| + | 2.053289 | -0.6779 | Q9P2B2 | Q9P2B2 | PTGFRN | Prostaglan | Color2 [Da | 1 | 4 |
| + | 1.352576 | -0.2393 | Q9P2B4 | Q9P2B4 | CTTNBP2N | CTTNBP2 N | Color2 [Da | 1 | 4 |
| + | 1.478736 | -0.47933 | Q9P2D1 | Q9P2D1 | CHD7 | Chromodo | Color2 [Da | 1 | 4 |
| | 0.243115 | -0.20865 | Q9P2D3 | Q9P2D3 | HEATR5B | HEAT repe | Color2 [Da | 1 | 4 |
| + | 1.617628 | 0.869762 | Q9P2E7 | Q9P2E7 | PCDH10 | Protocadh | Color2 [Da | 1 | 4 |
| + | 1.742561 | -0.69713 | Q9P2E9 | Q9P2E9 | RRBP1 | Ribosome- | Color2 [Da | 1 | 4 |
| | 0.442347 | 0.116351 | Q9P2F8 | Q9P2F8 | SIPA1L2 | Signal-indu | Color2 [Da | 1 | 4 |
| + | 1.813552 | -1.03699 | Q9P2H5 | Q9P2H5 | USP35 | Ubiquitin c | Color2 [Da | 1 | 4 |
| | 0.750902 | -0.41047 | Q9P2I0 | Q9P2I0 | CPSF2 | Cleavage a | Color2 [Da | 1 | 4 |
| + | 2.759993 | -1.44644 | Q9P2J5 | Q9P2J5 | LARS1 | Leucine--tl | Color2 [Da | 1 | 4 |
| + | 2.155718 | 1.067023 | Q9P2K3 | Q9P2K3 | RCOR3 | REST corep | Color2 [Da | 1 | 4 |

| | | | | | | | | | |
|---|---|---|---|---|---|---|---|---|---|
| + | 2.284162 | 0.898069 | Q9P2K5 | Q9P2K5 | MYEF2 | Myelin exp | Color2 [Da | 1 | 4 |
| + | 3.106681 | -1.61906 | Q9P2K6 | Q9P2K6 | KLHL42 | Kelch-like | Color2 [Da | 1 | 4 |
| | 0.140442 | -0.23643 | Q9P2K8 | Q9P2K8 | EIF2AK4 | eIF-2-alpha | Color2 [Da | 1 | 4 |
| | 0 | 0.950483 | Q9P2L0 | Q9P2L0 | WDR35 | WD repeat | Color2 [Da | 1 | 4 |
| | 0.661526 | -0.09845 | Q9P2N5 | Q9P2N5 | RBM27 | RNA-bindi | Color2 [Da | 1 | 4 |
| | 0 | -0.53013 | Q9P2N6 | Q9P2N6 | KANSL3 | KAT8 regul | Color2 [Da | 1 | 4 |
| | 0.090849 | 0.342115 | Q9P2N7 | Q9P2N7 | KLHL13 | Kelch-like | Color2 [Da | 1 | 4 |
| | 0.366487 | -0.50745 | Q9P2P5 | Q9P2P5 | HECW2 | E3 ubiquiti | Color2 [Da | 1 | 4 |
| + | 1.307027 | 0.537218 | Q9P2Q2 | Q9P2Q2 | FRMD4A | FERM dom | Color2 [Da | 1 | 4 |
| + | 3.914133 | -1.18351 | Q9P2R3 | Q9P2R3 | ANKFY1 | Rabankyrin | Color2 [Da | 1 | 4 |
| + | 2.166769 | -0.42601 | Q9P2R7 | Q9P2R7 | SUCLA2 | Succinate- | Color2 [Da | 1 | 4 |
| | 0.549261 | 0.486236 | Q9P2S2 | Q9P2S2 | NRXN2 | Neurexin-2 | Color2 [Da | 1 | 4 |
| + | 1.099143 | 0.489898 | Q9P2T1 | Q9P2T1 | GMPR2 | GMP redu | Color2 [Da | 1 | 4 |
| | 0 | -0.19014 | Q9P2U8 | Q9P2U8 | SLC17A6 | Vesicular g | Color2 [Da | 1 | 4 |
| + | 0.804776 | -0.94957 | Q9P2W1 | Q9P2W1 | PSMC3IP | Homologo | Color2 [Da | 1 | 4 |
| | 0.083619 | 0.03101 | Q9P2W9 | Q9P2W9 | STX18 | Syntaxin-1 | Color2 [Da | 1 | 4 |
| + | 1.591496 | -1.86793 | Q9P2X0 | Q9P2X0 | DPM3 | Dolichol-p | Color2 [Da | 1 | 4 |
| | 0.705038 | -0.2371 | Q9P2X3 | Q9P2X3 | IMPACT | Protein IM | Color2 [Da | 1 | 4 |
| | 0.799271 | -0.31787 | Q9P2Y4 | Q9P2Y4 | ZNF219 | Zinc finger | Color2 [Da | 1 | 4 |
| + | 2.295645 | -0.68005 | Q9UBB4 | Q9UBB4 | ATXN10 | Ataxin-10 | Color2 [Da | 1 | 4 |
| + | 2.707014 | -0.72081 | Q9UBB6 | Q9UBB6 | NCDN | Neurochor | Color2 [Da | 1 | 4 |
| + | 1.223313 | -0.29595 | Q9UBB9 | Q9UBB9 | TFIP11 | Tuftelin-in | Color2 [Da | 1 | 4 |
| + | 3.114631 | 1.478161 | Q9UBC2 | Q9UBC2 | EPS15L1 | Epidermal | Color2 [Da | 1 | 4 |
| | 0 | NaN | Q9UBC9 | Q9UBC9 | SPRR3 | Small proli | Color2 [Da | 1 | 4 |
| + | 2.33991 | -0.96333 | Q9UBD5 | Q9UBD5 | ORC3 | Origin reco | Color2 [Da | 1 | 4 |
| + | 1.871086 | -0.28902 | Q9UBE0 | Q9UBE0 | SAE1 | SUMO-act | Color2 [Da | 1 | 4 |
| + | 2.072898 | -1.23654 | Q9UBF2 | Q9UBF2 | COPG2 | Coatomer | Color2 [Da | 1 | 4 |
| | 0 | NaN | Q9UBF8 | Q9UBF8 | PI4KB | Phosphatid | Color2 [Da | 1 | 4 |
| + | 1.608544 | -1.23862 | Q9UBG0 | Q9UBG0 | MRC2 | C-type ma | Color2 [Da | 1 | 4 |
| | 0.302078 | 0.147673 | Q9UBH6 | Q9UBH6 | XPR1 | Xenotropic | Color2 [Da | 1 | 4 |
| | 0.69254 | -0.1081 | Q9UBI1 | Q9UBI1 | COMMD3 | COMM do | Color2 [Da | 1 | 4 |
| + | 2.190621 | 1.798047 | Q9UBI6 | Q9UBI6 | GNG12 | Guanine n | Color2 [Da | 1 | 4 |
| + | 2.198301 | 0.427544 | Q9UBI9 | Q9UBI9 | HECA | Headcase | Color2 [Da | 1 | 4 |
| | 0.145964 | 0.028796 | Q9UBK7;Q | Q9UBK7;Q | RABL2A;RA | Rab-like pr | Color2 [Da | 1 | 4 |
| + | 1.953804 | 1.141461 | Q9UBK9 | Q9UBK9 | UXT | Protein UX | Color2 [Da | 1 | 4 |
| | 0.271807 | 0.172194 | Q9UBL3 | Q9UBL3 | ASH2L | Set1/Ash2 | Color2 [Da | 1 | 4 |
| + | 1.395639 | -1.4947 | Q9UBM7 | Q9UBM7 | DHCR7 | 7-dehydro | Color2 [Da | 1 | 4 |
| | 0 | -0.83884 | Q9UBM8 | Q9UBM8 | MGAT4C | Alpha-1,3- | Color2 [Da | 1 | 4 |
| + | 2.23645 | -1.7737 | Q9UBN7 | Q9UBN7 | HDAC6 | Histone de | Color2 [Da | 1 | 4 |
| + | 2.144399 | 0.788858 | Q9UBP0 | Q9UBP0 | SPAST | Spastin | Color2 [Da | 1 | 4 |
| + | 2.895 | 1.959935 | Q9UBP4 | Q9UBP4 | DKK3 | Dickkopf-r | Color2 [Da | 1 | 4 |
| | 0.127227 | -0.1413 | Q9UBP6 | Q9UBP6 | METTL1 | tRNA (guar | Color2 [Da | 1 | 4 |
| | 0 | -0.48332 | Q9UBP9 | Q9UBP9 | GULP1 | PTB domai | Color2 [Da | 1 | 4 |
| + | 1.578673 | 0.333454 | Q9UBQ0 | Q9UBQ0 | VPS29 | Vacuolar p | Color2 [Da | 1 | 4 |
| | 0.688345 | -0.48367 | Q9UBQ5 | Q9UBQ5 | EIF3K | Eukaryotic | Color2 [Da | 1 | 4 |
| + | 3.154679 | 0.537118 | Q9UBQ6 | Q9UBQ6 | EXTL2 | Exostosin- | Color2 [Da | 1 | 4 |
| + | 2.760219 | -1.02792 | Q9UBQ7 | Q9UBQ7 | GRHPR | Glyoxylate | Color2 [Da | 1 | 4 |

| | | | | | | | | | |
|---|---|---|---|---|---|---|---|---|---|
| | 0.446954 | 0.248208 | Q9UBS3 | Q9UBS3 | DNAJB9 | DnaJ homo | Color2 [Da | 1 | 4 |
| + | 2.901542 | 1.344433 | Q9UBS4 | Q9UBS4 | DNAJB11 | DnaJ homo | Color2 [Da | 1 | 4 |
| + | 1.714234 | -0.85857 | Q9UBT2 | Q9UBT2 | UBA2 | SUMO-act | Color2 [Da | 1 | 4 |
| | 0.627553 | -0.41002 | Q9UBU6 | Q9UBU6 | FAM8A1 | Protein FA | Color2 [Da | 1 | 4 |
| | 0.374565 | -0.06421 | Q9UBU8 | Q9UBU8 | MORF4L1 | Mortality f | Color2 [Da | 1 | 4 |
| + | 3.167927 | -0.80456 | Q9UBU9 | Q9UBU9 | NXF1 | Nuclear RN | Color2 [Da | 1 | 4 |
| + | 1.820116 | -0.97626 | Q9UBV2 | Q9UBV2 | SEL1L | Protein sel | Color2 [Da | 1 | 4 |
| + | 1.0837 | -0.29181 | Q9UBV8 | Q9UBV8 | PEF1 | Peflin | Color2 [Da | 1 | 4 |
| + | 0.956849 | -0.53722 | Q9UBW7 | Q9UBW7 | ZMYM2 | Zinc finger | Color2 [Da | 1 | 4 |
| + | 1.966143 | 0.578525 | Q9UBW8 | Q9UBW8 | COPS7A | COP9 signa | Color2 [Da | 1 | 4 |
| | 0.003151 | -0.00262 | Q9UBX3 | Q9UBX3 | SLC25A10 | Mitochond | Color2 [Da | 1 | 4 |
| | 0 | NaN | Q9UBY8 | Q9UBY8 | CLN8 | Protein CL | Color2 [Da | 1 | 4 |
| | 0.41816 | -0.30427 | Q9UDR5 | Q9UDR5 | AASS | Alpha-ami | Color2 [Da | 1 | 4 |
| + | 1.873622 | 0.444439 | Q9UDT6 | Q9UDT6 | CLIP2 | CAP-Gly do | Color2 [Da | 1 | 4 |
| + | 1.39355 | 1.410375 | Q9UDW1 | Q9UDW1 | UQCR10 | Cytochrom | Color2 [Da | 1 | 4 |
| | 0 | NaN | Q9UDW3 | Q9UDW3 | ZMAT5 | Zinc finger | Color2 [Da | 1 | 4 |
| + | 0.884677 | 0.910341 | Q9UDX5 | Q9UDX5 | MTFP1 | Mitochond | Color2 [Da | 1 | 4 |
| | 0.28007 | 0.252686 | Q9UDY2 | Q9UDY2 | TJP2 | Tight junct | Color2 [Da | 1 | 4 |
| | 1.004842 | -0.19262 | Q9UDY4 | Q9UDY4 | DNAJB4 | DnaJ homo | Color2 [Da | 1 | 4 |
| | 0 | NaN | Q9UDY8 | Q9UDY8 | MALT1 | Mucosa-as | Color2 [Da | 1 | 4 |
| + | 0.94343 | 0.748487 | Q9UEE9 | Q9UEE9 | CFDP1 | Craniofacia | Color2 [Da | 1 | 4 |
| | 0.447942 | -0.36106 | Q9UEG4 | Q9UEG4 | ZNF629 | Zinc finger | Color2 [Da | 1 | 4 |
| | 0.20692 | 0.696059 | Q9UER7 | Q9UER7 | DAXX | Death dom | Color2 [Da | 1 | 4 |
| + | 1.173866 | 0.442487 | Q9UEU0 | Q9UEU0 | VTI1B | Vesicle tra | Color2 [Da | 1 | 4 |
| | 0.080583 | -0.03524 | Q9UEW8 | Q9UEW8 | STK39 | STE20/SPS | Color2 [Da | 1 | 4 |
| + | 2.231341 | 0.571662 | Q9UEY8 | Q9UEY8 | ADD3 | Gamma-ad | Color2 [Da | 1 | 4 |
| + | 1.25474 | -0.30696 | Q9UFC0 | Q9UFC0 | LRWD1 | Leucine-ric | Color2 [Da | 1 | 4 |
| + | 3.333994 | 1.845443 | Q9UFG5 | Q9UFG5 | C19orf25 | UPF0449 p | Color2 [Da | 1 | 4 |
| + | 3.803953 | 1.887378 | Q9UFN0 | Q9UFN0 | NIPSNAP3A | Protein Nip | Color2 [Da | 1 | 4 |
| | 0 | NaN | Q9UFW8 | Q9UFW8 | CGGBP1 | CGG triplet | Color2 [Da | 1 | 4 |
| | 0 | -1.98826 | Q9UG01 | Q9UG01 | IFT172 | Intraflagell | Color2 [Da | 1 | 4 |
| | 0 | -1.04408 | Q9UG63 | Q9UG63 | ABCF2 | ATP-bindir | Color2 [Da | 1 | 4 |
| | 0 | -1.38247 | Q9UGC7 | Q9UGC7 | MTRF1L | Peptide ch | Color2 [Da | 1 | 4 |
| + | 1.36941 | 1.368572 | Q9UGI6 | Q9UGI6 | KCNN3 | Small conc | Color2 [Da | 1 | 4 |
| + | 1.116736 | -1.55061 | Q9UGJ1 | Q9UGJ1 | TUBGCP4 | Gamma-tu | Color2 [Da | 1 | 4 |
| | 0 | -0.44373 | Q9UGL1 | Q9UGL1 | KDM5B | Lysine-spe | Color2 [Da | 1 | 4 |
| + | 2.015997 | -0.89353 | Q9UGM3 | Q9UGM3 | DMBT1 | Deleted in | Color2 [Da | 1 | 4 |
| | 0.551628 | 0.202321 | Q9UGM6 | Q9UGM6 | WARS2 | Tryptopha | Color2 [Da | 1 | 4 |
| + | 2.364263 | -1.00438 | Q9UGN5 | Q9UGN5 | PARP2 | Poly [ADP- | Color2 [Da | 1 | 4 |
| + | 3.892192 | -2.70096 | Q9UGP4 | Q9UGP4 | LIMD1 | LIM domai | Color2 [Da | 1 | 4 |
| + | 2.311939 | -0.89384 | Q9UGP8 | Q9UGP8 | SEC63 | Translocat | Color2 [Da | 1 | 4 |
| | 0 | 0.63676 | Q9UGQ2 | Q9UGQ2 | CACFD1 | Calcium ch | Color2 [Da | 1 | 4 |
| | 0.546895 | 0.214823 | Q9UGR2 | Q9UGR2 | ZC3H7B | Zinc finger | Color2 [Da | 1 | 4 |
| | 0.265236 | -0.12891 | Q9UGU0 | Q9UGU0 | TCF20 | Transcripti | Color2 [Da | 1 | 4 |
| | 0.504479 | -0.72224 | Q9UGU5 | Q9UGU5 | HMGXB4 | HMG dom | Color2 [Da | 1 | 4 |
| + | 0.978312 | -0.74492 | Q9UGV2 | Q9UGV2 | NDRG3 | Protein ND | Color2 [Da | 1 | 4 |
| + | 1.184091 | 0.327329 | Q9UH03 | Q9UH03 | SEPTIN3 | Neuronal-s | Color2 [Da | 1 | 4 |

| | | | | | | | | | |
|---|---|---|---|---|---|---|---|---|---|
| + | 1.388215 | -0.77075 | Q9UH62 | Q9UH62 | ARMCX3 | Armadillo | Color2 [Da | 1 | 4 |
| + | 1.864415 | -1.06095 | Q9UH65 | Q9UH65 | SWAP70 | Switch-ass | Color2 [Da | 1 | 4 |
| + | 3.403343 | -1.60276 | Q9UH99 | Q9UH99 | SUN2 | SUN doma | Color2 [Da | 1 | 4 |
| + | 1.638704 | 0.744639 | Q9UHA2 | Q9UHA2 | SS18L2 | SS18-like p | Color2 [Da | 1 | 4 |
| + | 2.313302 | 2.089519 | Q9UHA4 | Q9UHA4 | LAMTOR3 | Ragulator | Color2 [Da | 1 | 4 |
| | 0 | NaN | Q9UHB4 | Q9UHB4 | NDOR1 | NADPH-de | Color2 [Da | 1 | 4 |
| + | 0.863131 | 0.661265 | Q9UHB6 | Q9UHB6 | LIMA1 | LIM domai | Color2 [Da | 1 | 4 |
| + | 0.892603 | -0.80718 | Q9UHB7 | Q9UHB7 | AFF4 | AF4/FMR2 | Color2 [Da | 1 | 4 |
| + | 3.945437 | -1.04161 | Q9UHB9 | Q9UHB9 | SRP68 | Signal recc | Color2 [Da | 1 | 4 |
| | 0.415997 | 0.232976 | Q9UHC6 | Q9UHC6 | CNTNAP2 | Contactin- | Color2 [Da | 1 | 4 |
| + | 1.084316 | -0.31549 | Q9UHD1 | Q9UHD1 | CHORDC1 | Cysteine a | Color2 [Da | 1 | 4 |
| | 0.483512 | -0.38208 | Q9UHD2 | Q9UHD2 | TBK1 | Serine/thr | Color2 [Da | 1 | 4 |
| | 0.210629 | -0.0731 | Q9UHD8 | Q9UHD8 | SEPTIN9 | Septin-9 | Color2 [Da | 1 | 4 |
| + | 2.926938 | 2.078807 | Q9UHD9 | Q9UHD9 | UBQLN2 | Ubiquilin-2 | Color2 [Da | 1 | 4 |
| | 0 | NaN | Q9UHF5 | Q9UHF5 | IL17B | Interleukir | Color2 [Da | 1 | 4 |
| | 0 | NaN | Q9UHG0 | Q9UHG0 | DCDC2 | Doublecor | Color2 [Da | 1 | 4 |
| + | 2.166548 | 1.159753 | Q9UHG2 | Q9UHG2 | PCSK1N | ProSAAS | Color2 [Da | 1 | 4 |
| + | 2.706987 | 1.016981 | Q9UHG3 | Q9UHG3 | PCYOX1 | Prenylcyst | Color2 [Da | 1 | 4 |
| + | 1.01305 | -1.38741 | Q9UHI6 | Q9UHI6 | DDX20 | Probable A | Color2 [Da | 1 | 4 |
| + | 1.303571 | -0.66491 | Q9UHJ6 | Q9UHJ6 | SHPK | Sedoheptu | Color2 [Da | 1 | 4 |
| + | 4.102558 | 1.461162 | Q9UHL4 | Q9UHL4 | DPP7 | Dipeptidyl | Color2 [Da | 1 | 4 |
| + | 1.719324 | -1.20772 | Q9UHN1 | Q9UHN1 | POLG2 | DNA polyn | Color2 [Da | 1 | 4 |
| | 0.145841 | -0.07294 | Q9UHQ9 | Q9UHQ9 | CYB5R1 | NADH-cyto | Color2 [Da | 1 | 4 |
| | 0 | NaN | Q9UHR4 | Q9UHR4 | BAIAP2L1 | Brain-spec | Color2 [Da | 1 | 4 |
| + | 1.98983 | 0.57351 | Q9UHR5 | Q9UHR5 | SAP30BP | SAP30-bin | Color2 [Da | 1 | 4 |
| | 0 | 0.273192 | Q9UHR6 | Q9UHR6 | ZNHIT2 | Zinc finger | Color2 [Da | 1 | 4 |
| + | 2.479288 | 1.049974 | Q9UHV9 | Q9UHV9 | PFDN2 | Prefoldin s | Color2 [Da | 1 | 4 |
| + | 1.337453 | 1.260063 | Q9UHW9 | Q9UHW9 | SLC12A6 | Solute carr | Color2 [Da | 1 | 4 |
| | 1.265988 | -0.16286 | Q9UHX1 | Q9UHX1 | PUF60 | Poly(U)-bir | Color2 [Da | 1 | 4 |
| + | 1.177003 | -0.57713 | Q9UHY1 | Q9UHY1 | NRBP1 | Nuclear re | Color2 [Da | 1 | 4 |
| + | 3.739215 | 1.892904 | Q9UHY7 | Q9UHY7 | ENOPH1 | Enolase-pł | Color2 [Da | 1 | 4 |
| + | 1.415479 | 0.512517 | Q9UHY8 | Q9UHY8 | FEZ2 | Fasciculati | Color2 [Da | 1 | 4 |
| + | 3.151066 | 1.392071 | Q9UI08 | Q9UI08 | EVL | Ena/VASP- | Color2 [Da | 1 | 4 |
| + | 3.816697 | 0.820094 | Q9UI09 | Q9UI09 | NDUFA12 | NADH deh | Color2 [Da | 1 | 4 |
| | 0 | -1.13435 | Q9UI10 | Q9UI10 | EIF2B4 | Translatior | Color2 [Da | 1 | 4 |
| | 0.496926 | -0.20137 | Q9UI12 | Q9UI12 | ATP6V1H | V-type pro | Color2 [Da | 1 | 4 |
| + | 1.049516 | 0.386672 | Q9UI15 | Q9UI15 | TAGLN3 | Transgelin | Color2 [Da | 1 | 4 |
| + | 3.471264 | -2.32211 | Q9UI26 | Q9UI26 | IPO11 | Importin-1 | Color2 [Da | 1 | 4 |
| | 0.41664 | 0.114496 | Q9UI30 | Q9UI30 | TRMT112 | Multifunct | Color2 [Da | 1 | 4 |
| + | 2.359399 | 0.259519 | Q9UI36 | Q9UI36 | DACH1 | Dachshunc | Color2 [Da | 1 | 4 |
| | 0.086724 | -0.28142 | Q9UI42 | Q9UI42 | CPA4 | Carboxype | Color2 [Da | 1 | 4 |
| + | 0.743588 | -0.62583 | Q9UI47 | Q9UI47 | CTNNA3 | Catenin alı | Color2 [Da | 1 | 4 |
| + | 4.079145 | -1.9915 | Q9UIA9 | Q9UIA9 | XPO7 | Exportin-7 | Color2 [Da | 1 | 4 |
| + | 1.270789 | 0.405205 | Q9UIC8 | Q9UIC8 | LCMT1 | Leucine ca | Color2 [Da | 1 | 4 |
| + | 2.446955 | -0.83862 | Q9UID3 | Q9UID3 | VPS51 | Vacuolar p | Color2 [Da | 1 | 4 |
| | 0 | -1.41424 | Q9UIF8 | Q9UIF8 | BAZ2B | Bromodon | Color2 [Da | 1 | 4 |
| | 0.347953 | -0.20183 | Q9UIF9 | Q9UIF9 | BAZ2A | Bromodon | Color2 [Da | 1 | 4 |

| | | | | | | | | | |
|---|---|---|---|---|---|---|---|---|---|
| + | 2.697175 | -1.30478 | Q9UIG0 | Q9UIG0 | BAZ1B | Tyrosine-p | Color2 [Da | 1 | 4 |
| | 0 | 2.931025 | Q9UII2 | Q9UII2 | ATP5IF1 | ATPase inh | Color2 [Da | 1 | 4 |
| + | 3.0588 | 0.991818 | Q9UIJ7 | Q9UIJ7 | AK3 | GTP:AMP | Color2 [Da | 1 | 4 |
| + | 2.737533 | 1.283527 | Q9UIL1 | Q9UIL1 | SCOC | Short coile | Color2 [Da | 1 | 4 |
| + | 1.447199 | 0.941505 | Q9UIQ6 | Q9UIQ6 | LNPEP | Leucyl-cyst | Color2 [Da | 1 | 4 |
| | 0 | -0.70941 | Q9UIS9 | Q9UIS9 | MBD1 | Methyl-Cp | Color2 [Da | 1 | 4 |
| + | 1.841995 | -1.47303 | Q9UIV1 | Q9UIV1 | CNOT7 | CCR4-NOT | Color2 [Da | 1 | 4 |
| | 0.903745 | -0.23446 | Q9UIW2 | Q9UIW2 | PLXNA1 | Plexin-A1 | Color2 [Da | 1 | 4 |
| + | 1.484076 | -0.53338 | Q9UJ14 | Q9UJ14 | GGT7 | Glutathion | Color2 [Da | 1 | 4 |
| + | 0.964987 | -0.32848 | Q9UJ41 | Q9UJ41 | RABGEF1 | Rab5 GDP/ | Color2 [Da | 1 | 4 |
| + | 1.296169 | 0.313992 | Q9UJ68 | Q9UJ68 | MSRA | Mitochonc | Color2 [Da | 1 | 4 |
| + | 1.495141 | -0.30145 | Q9UJ70 | Q9UJ70 | NAGK | N-acetyl-D | Color2 [Da | 1 | 4 |
| | 0 | NaN | Q9UJ83 | Q9UJ83 | HACL1 | 2-hydroxya | Color2 [Da | 1 | 4 |
| + | 1.315661 | 0.593669 | Q9UJA5 | Q9UJA5 | TRMT6 | tRNA (ade | Color2 [Da | 1 | 4 |
| | 0 | NaN | Q9UJA9 | Q9UJA9 | ENPP5 | Ectonuclec | Color2 [Da | 1 | 4 |
| + | 1.015335 | 0.883993 | Q9UJC3 | Q9UJC3 | HOOK1 | Protein Hc | Color2 [Da | 1 | 4 |
| + | 2.465093 | 2.062237 | Q9UJC5 | Q9UJC5 | SH3BGRL2 | SH3 domai | Color2 [Da | 1 | 4 |
| + | 1.200897 | 0.957171 | Q9UJD0 | Q9UJD0 | RIMS3 | Regulating | Color2 [Da | 1 | 4 |
| | 0.062021 | 0.028356 | Q9UJF2 | Q9UJF2 | RASAL2 | Ras GTPas | Color2 [Da | 1 | 4 |
| + | 1.195682 | -0.8768 | Q9UJH8 | Q9UJH8 | METRN | Meteorin | Color2 [Da | 1 | 4 |
| | 0 | NaN | Q9UJQ1 | Q9UJQ1 | LAMP5 | Lysosome- | Color2 [Da | 1 | 4 |
| + | 3.238848 | -1.12629 | Q9UJS0 | Q9UJS0 | SLC25A13 | Calcium-bi | Color2 [Da | 1 | 4 |
| + | 2.576683 | 0.959791 | Q9UJU6 | Q9UJU6 | DBNL | Drebrin-lik | Color2 [Da | 1 | 4 |
| + | 2.32089 | 0.889935 | Q9UJV8 | Q9UJV8 | PURG | Purine-rich | Color2 [Da | 1 | 4 |
| + | 1.957372 | -1.11494 | Q9UJV9 | Q9UJV9 | DDX41 | Probable A | Color2 [Da | 1 | 4 |
| | 0.269804 | 0.062334 | Q9UJW0 | Q9UJW0 | DCTN4 | Dynactin s | Color2 [Da | 1 | 4 |
| + | 1.916838 | -0.92624 | Q9UJX2 | Q9UJX2 | CDC23 | Cell divisio | Color2 [Da | 1 | 4 |
| | 0 | NaN | Q9UJX3 | Q9UJX3 | ANAPC7 | Anaphase- | Color2 [Da | 1 | 4 |
| + | 1.555171 | -0.40407 | Q9UJX4 | Q9UJX4 | ANAPC5 | Anaphase- | Color2 [Da | 1 | 4 |
| | 0 | NaN | Q9UJX5 | Q9UJX5 | ANAPC4 | Anaphase- | Color2 [Da | 1 | 4 |
| | 0 | NaN | Q9UJX6 | Q9UJX6 | ANAPC2 | Anaphase- | Color2 [Da | 1 | 4 |
| + | 1.625096 | 0.646409 | Q9UJY1 | Q9UJY1 | HSPB8 | Heat shocl | Color2 [Da | 1 | 4 |
| + | 0.874531 | -0.63894 | Q9UJY4 | Q9UJY4 | GGA2 | ADP-ribosy | Color2 [Da | 1 | 4 |
| + | 2.589432 | 0.599654 | Q9UJY5 | Q9UJY5 | GGA1 | ADP-ribosy | Color2 [Da | 1 | 4 |
| + | 3.011433 | 0.894071 | Q9UJZ1 | Q9UJZ1 | STOML2 | Stomatin-l | Color2 [Da | 1 | 4 |
| | 0.576086 | 0.177256 | Q9UK22 | Q9UK22 | FBXO2 | F-box only | Color2 [Da | 1 | 4 |
| | 0.722091 | 0.289897 | Q9UK41 | Q9UK41 | VPS28 | Vacuolar p | Color2 [Da | 1 | 4 |
| + | 1.320067 | 1.64793 | Q9UK45 | Q9UK45 | LSM7 | U6 snRNA- | Color2 [Da | 1 | 4 |
| + | 2.321545 | 0.742765 | Q9UK53 | Q9UK53 | ING1 | Inhibitor o | Color2 [Da | 1 | 4 |
| | 0 | 2.042021 | Q9UK55 | Q9UK55 | SERPINA1( | Protein Z-c | Color2 [Da | 1 | 4 |
| | 0 | -0.5881 | Q9UK58 | Q9UK58 | CCNL1 | Cyclin-L1 | Color2 [Da | 1 | 4 |
| + | 1.623362 | 0.97103 | Q9UK59 | Q9UK59 | DBR1 | Lariat debi | Color2 [Da | 1 | 4 |
| + | 4.045688 | 4.186778 | Q9UK76 | Q9UK76 | JPT1 | Jupiter mic | Color2 [Da | 1 | 4 |
| | 0 | NaN | Q9UK99 | Q9UK99 | FBXO3 | F-box only | Color2 [Da | 1 | 4 |
| + | 2.459213 | 0.817888 | Q9UKA9 | Q9UKA9 | PTBP2 | Polypyrimi | Color2 [Da | 1 | 4 |
| | 0 | 2.377212 | Q9UKB3 | Q9UKB3 | DNAJC12 | DnaJ homc | Color2 [Da | 1 | 4 |
| + | 0.907833 | -0.34799 | Q9UKD2 | Q9UKD2 | MRTO4 | mRNA turr | Color2 [Da | 1 | 4 |

| | | | | | | | | | |
|---|---|---|---|---|---|---|---|---|---|
| + | 2.187502 | 0.475963 | Q9UKE5 | Q9UKE5 | TNIK | TRAF2 and | Color2 [Da | 1 | 4 |
| + | 1.821071 | -1.3281 | Q9UKF6 | Q9UKF6 | CPSF3 | Cleavage a | Color2 [Da | 1 | 4 |
| + | 1.331039 | -1.20858 | Q9UKF7 | Q9UKF7 | PITPNC1 | Cytoplasm | Color2 [Da | 1 | 4 |
| | 0.024671 | 0.015338 | Q9UKG1 | Q9UKG1 | APPL1 | DCC-intera | Color2 [Da | 1 | 4 |
| | 0.068938 | -0.01996 | Q9UKG9 | Q9UKG9 | CROT | Peroxisom | Color2 [Da | 1 | 4 |
| | 0.055525 | -0.13296 | Q9UKI2 | Q9UKI2 | CDC42EP3 | Cdc42 effe | Color2 [Da | 1 | 4 |
| | 0.318923 | -0.06759 | Q9UKJ3 | Q9UKJ3 | GPATCH8 | G patch dc | Color2 [Da | 1 | 4 |
| + | 2.98293 | 1.244668 | Q9UKK9 | Q9UKK9 | NUDT5 | ADP-sugar | Color2 [Da | 1 | 4 |
| + | 1.990427 | 0.905438 | Q9UKL0 | Q9UKL0 | RCOR1 | REST corep | Color2 [Da | 1 | 4 |
| | 0 | -1.37332 | Q9UKM7 | Q9UKM7 | MAN1B1 | Endoplasm | Color2 [Da | 1 | 4 |
| + | 2.487551 | 0.629226 | Q9UKM9 | Q9UKM9 | RALY | RNA-bindi | Color2 [Da | 1 | 4 |
| + | 3.971944 | -1.03537 | Q9UKN8 | Q9UKN8 | GTF3C4 | General tr | Color2 [Da | 1 | 4 |
| | 0 | -0.64455 | Q9UKR5 | Q9UKR5 | ERG28 | Ergosterol | Color2 [Da | 1 | 4 |
| | 0.138223 | -0.08654 | Q9UKU0 | Q9UKU0 | ACSL6 | Long-chair | Color2 [Da | 1 | 4 |
| | 0.163568 | 0.100912 | Q9UKU9 | Q9UKU9 | ANGPTL2 | Angiopoiet | Color2 [Da | 1 | 4 |
| + | 1.415443 | 1.176849 | Q9UKV0 | Q9UKV0 | HDAC9 | Histone de | Color2 [Da | 1 | 4 |
| + | 2.170934 | 0.783729 | Q9UKV3 | Q9UKV3 | ACIN1 | Apoptotic | Color2 [Da | 1 | 4 |
| + | 3.939393 | -2.56177 | Q9UKV8 | Q9UKV8 | AGO2 | Protein arg | Color2 [Da | 1 | 4 |
| | 0 | NaN | Q9UKX2 | Q9UKX2 | MYH2 | Myosin-2 | Color2 [Da | 1 | 4 |
| + | 1.308831 | 1.099955 | Q9UKX7 | Q9UKX7 | NUP50 | Nuclear pc | Color2 [Da | 1 | 4 |
| + | 1.314804 | -0.76469 | Q9UKY0 | Q9UKY0 | PRND | Prion-like | Color2 [Da | 1 | 4 |
| + | 4.582731 | 3.036761 | Q9UKY7 | Q9UKY7 | CDV3 | Protein CD | Color2 [Da | 1 | 4 |
| + | 1.648787 | 1.139323 | Q9UKZ1 | Q9UKZ1 | CNOT11 | CCR4-NOT | Color2 [Da | 1 | 4 |
| | 0 | NaN | Q9UKZ4 | Q9UKZ4 | TENM1 | Teneurin-1 | Color2 [Da | 1 | 4 |
| + | 1.991442 | -1.14999 | Q9UL15 | Q9UL15 | BAG5 | BAG family | Color2 [Da | 1 | 4 |
| + | 1.004109 | -0.8361 | Q9UL18 | Q9UL18 | AGO1 | Protein arg | Color2 [Da | 1 | 4 |
| | 0.033246 | 0.023428 | Q9UL25 | Q9UL25 | RAB21 | Ras-relate | Color2 [Da | 1 | 4 |
| + | 1.659415 | -0.88497 | Q9UL26 | Q9UL26 | RAB22A | Ras-relate | Color2 [Da | 1 | 4 |
| + | 1.686056 | -0.49512 | Q9UL33 | Q9UL33 | TRAPPC2L | Trafficking | Color2 [Da | 1 | 4 |
| + | 2.12795 | -0.45209 | Q9UL40 | Q9UL40 | ZNF346 | Zinc finger | Color2 [Da | 1 | 4 |
| + | 1.105819 | -0.69559 | Q9UL42 | Q9UL42 | PNMA2 | Paraneopl | Color2 [Da | 1 | 4 |
| | 1.203247 | 0.213026 | Q9UL45 | Q9UL45 | BLOC1S6 | Biogenesis | Color2 [Da | 1 | 4 |
| | 1.090605 | -0.24495 | Q9UL46 | Q9UL46 | PSME2 | Proteasom | Color2 [Da | 1 | 4 |
| + | 2.734092 | 0.53731 | Q9UL54 | Q9UL54 | TAOK2 | Serine/thr | Color2 [Da | 1 | 4 |
| | 0 | -1.31492 | Q9UL63 | Q9UL63 | MKLN1 | Muskelin | Color2 [Da | 1 | 4 |
| + | 2.064991 | -0.80567 | Q9ULA0 | Q9ULA0 | DNPEP | Aspartyl ar | Color2 [Da | 1 | 4 |
| + | 1.04438 | 0.583429 | Q9ULB1 | Q9ULB1 | NRXN1 | Neurexin-1 | Color2 [Da | 1 | 4 |
| + | 0.73294 | 0.632591 | Q9ULC3 | Q9ULC3 | RAB23 | Ras-relate | Color2 [Da | 1 | 4 |
| + | 1.664401 | -0.49969 | Q9ULC4 | Q9ULC4 | MCTS1 | Malignant | Color2 [Da | 1 | 4 |
| | 0 | -1.22024 | Q9ULD0 | Q9ULD0 | OGDHL | 2-oxogluta | Color2 [Da | 1 | 4 |
| | 0 | NaN | Q9ULD6 | Q9ULD6 | INTU | Protein int | Color2 [Da | 1 | 4 |
| + | 0.756507 | -0.6019 | Q9ULD9 | Q9ULD9 | ZNF608 | Zinc finger | Color2 [Da | 1 | 4 |
| + | 0.687125 | 0.993243 | Q9ULE3 | Q9ULE3 | DENND2A | DENN dom | Color2 [Da | 1 | 4 |
| | 0.297813 | 0.78336 | Q9ULE4 | Q9ULE4 | FAM184B | Protein FA | Color2 [Da | 1 | 4 |
| | 0 | 0.874873 | Q9ULE6 | Q9ULE6 | PALD1 | Paladin | Color2 [Da | 1 | 4 |
| + | 1.824471 | 1.314546 | Q9ULF5 | Q9ULF5 | SLC39A10 | Zinc transp | Color2 [Da | 1 | 4 |
| | 0.042119 | 0.010814 | Q9ULH0 | Q9ULH0 | KIDINS220 | Kinase D-ir | Color2 [Da | 1 | 4 |

| | | | | | | | | | |
|---|---|---|---|---|---|---|---|---|---|
| | 0.895054 | -0.24416 | Q9ULH1 | Q9ULH1 | ASAP1 | Arf-GAP w | Color2 [Da | 1 | 4 |
| + | 1.959841 | -0.41326 | Q9ULH7 | Q9ULH7 | MRTFB | Myocardin | Color2 [Da | 1 | 4 |
| | 0 | 0.709235 | Q9ULI0 | Q9ULI0 | ATAD2B | ATPase far | Color2 [Da | 1 | 4 |
| | 0 | -1.76124 | Q9ULJ6 | Q9ULJ6 | ZMIZ1 | Zinc finger | Color2 [Da | 1 | 4 |
| + | 1.939074 | -1.22115 | Q9ULJ7 | Q9ULJ7 | ANKRD50 | Ankyrin re | Color2 [Da | 1 | 4 |
| + | 3.895916 | 0.869898 | Q9ULJ8 | Q9ULJ8 | PPP1R9A | Neurabin-: | Color2 [Da | 1 | 4 |
| + | 2.681721 | -1.74584 | Q9ULK4 | Q9ULK4 | MED23 | Mediator ( | Color2 [Da | 1 | 4 |
| + | 1.861004 | -0.56947 | Q9ULK5 | Q9ULK5 | VANGL2 | Vang-like p | Color2 [Da | 1 | 4 |
| | 0 | NaN | Q9ULL1 | Q9ULL1 | PLEKHG1 | Pleckstrin | Color2 [Da | 1 | 4 |
| + | 1.271758 | -0.79482 | Q9ULL5 | Q9ULL5 | PRR12 | Proline-ric | Color2 [Da | 1 | 4 |
| | 0.766961 | -0.26667 | Q9ULM3 | Q9ULM3 | YEATS2 | YEATS don | Color2 [Da | 1 | 4 |
| | 0.444701 | 0.499474 | Q9ULN7 | Q9ULN7 | PNMA8B | Paraneopl | Color2 [Da | 1 | 4 |
| + | 1.57928 | 0.580433 | Q9ULP0 | Q9ULP0 | NDRG4 | Protein ND | Color2 [Da | 1 | 4 |
| | 0.342698 | -0.14178 | Q9ULP9 | Q9ULP9 | TBC1D24 | TBC1 dom | Color2 [Da | 1 | 4 |
| + | 1.605088 | 0.576729 | Q9ULR0 | Q9ULR0 | ISY1 | Pre-mRNA | Color2 [Da | 1 | 4 |
| + | 0.964013 | -0.38842 | Q9ULR3 | Q9ULR3 | PPM1H | Protein ph | Color2 [Da | 1 | 4 |
| | 0 | NaN | Q9ULS5 | Q9ULS5 | TMCC3 | Transmem | Color2 [Da | 1 | 4 |
| + | 3.158701 | -1.08529 | Q9ULT8 | Q9ULT8 | HECTD1 | E3 ubiquiti | Color2 [Da | 1 | 4 |
| + | 1.612088 | 0.45344 | Q9ULU4 | Q9ULU4 | ZMYND8 | Protein kir | Color2 [Da | 1 | 4 |
| + | 1.717123 | -0.33734 | Q9ULU8 | Q9ULU8 | CADPS | Calcium-de | Color2 [Da | 1 | 4 |
| | 0 | NaN | Q9ULV0 | Q9ULV0 | MYO5B | Unconvent | Color2 [Da | 1 | 4 |
| + | 3.138387 | -1.46994 | Q9ULV3 | Q9ULV3 | CIZ1 | Cip1-intera | Color2 [Da | 1 | 4 |
| + | 2.848842 | -1.34803 | Q9ULV4 | Q9ULV4 | CORO1C | Coronin-1( | Color2 [Da | 1 | 4 |
| | 0 | 1.511267 | Q9ULX3 | Q9ULX3 | NOB1 | RNA-bindi | Color2 [Da | 1 | 4 |
| | 0.682517 | 0.230748 | Q9ULX6 | Q9ULX6 | AKAP8L | A-kinase a | Color2 [Da | 1 | 4 |
| + | 2.176578 | -2.36845 | Q9ULX7 | Q9ULX7 | CA14 | Carbonic a | Color2 [Da | 1 | 4 |
| + | 3.126771 | -1.88128 | Q9UM00 | Q9UM00 | TMCO1 | Calcium lo | Color2 [Da | 1 | 4 |
| + | 2.559178 | 1.325625 | Q9UM19 | Q9UM19 | HPCAL4 | Hippocalci | Color2 [Da | 1 | 4 |
| + | 3.26621 | 2.420644 | Q9UM22 | Q9UM22 | EPDR1 | Mammalia | Color2 [Da | 1 | 4 |
| | 0.742591 | -0.16232 | Q9UM47 | Q9UM47 | NOTCH3 | Neurogeni | Color2 [Da | 1 | 4 |
| + | 4.294793 | -1.60509 | Q9UM54 | Q9UM54 | MYO6 | Unconvent | Color2 [Da | 1 | 4 |
| + | 3.335308 | 1.518443 | Q9UMR2 | Q9UMR2 | DDX19B | ATP-deper | Color2 [Da | 1 | 4 |
| + | 1.298892 | 1.794487 | Q9UMR5 | Q9UMR5 | PPT2 | Lysosomal | Color2 [Da | 1 | 4 |
| + | 3.490772 | 2.19341 | Q9UMS0 | Q9UMS0 | NFU1 | NFU1 iron- | Color2 [Da | 1 | 4 |
| | 0.285802 | -0.14722 | Q9UMS4 | Q9UMS4 | PRPF19 | Pre-mRNA | Color2 [Da | 1 | 4 |
| + | 3.034336 | 1.846697 | Q9UMX0 | Q9UMX0 | UBQLN1 | Ubiquilin-1 | Color2 [Da | 1 | 4 |
| + | 2.62614 | 1.096295 | Q9UMX1 | Q9UMX1 | SUFU | Suppresso | Color2 [Da | 1 | 4 |
| + | 1.578016 | 1.190955 | Q9UMX5 | Q9UMX5 | NENF | Neudesin | Color2 [Da | 1 | 4 |
| | 0.825527 | 0.256772 | Q9UMY1 | Q9UMY1 | NOL7 | Nucleolar | | Color2 [Da | 1 | 4 |
| | 0.128066 | -0.06935 | Q9UMY4 | Q9UMY4 | SNX12 | Sorting ne: | Color2 [Da | 1 | 4 |
| + | 2.835008 | 0.880313 | Q9UMZ2 | Q9UMZ2 | SYNRG | Synergin g | Color2 [Da | 1 | 4 |
| + | 4.345821 | 0.57886 | Q9UN36 | Q9UN36 | NDRG2 | Protein ND | Color2 [Da | 1 | 4 |
| + | 1.123906 | 0.469166 | Q9UN37 | Q9UN37 | VPS4A | Vacuolar p | Color2 [Da | 1 | 4 |
| | 0 | NaN | Q9UN70 | Q9UN70 | PCDHGC3 | Protocadh | Color2 [Da | 1 | 4 |
| + | 2.405472 | 1.311139 | Q9UN73 | Q9UN73 | PCDHA6 | Protocadh | Color2 [Da | 1 | 4 |
| | 0.076483 | 0.134198 | Q9UN74;C | Q9UN74;C | PCDHA4;P | Protocadh | Color2 [Da | 1 | 4 |
| | 0 | -3.47892 | Q9UN81 | Q9UN81 | L1RE1 | LINE-1 retr | Color2 [Da | 1 | 4 |

| | | | | | | | | | |
|---|---|---|---|---|---|---|---|---|---|
| + | 3.215864 | 1.026226 | Q9UN86 | Q9UN86 | G3BP2 | Ras GTPas | Color2 [Da | 1 | 4 |
| | 0 | 0.484416 | Q9UNA4 | Q9UNA4 | POLI | DNA polyn | Color2 [Da | 1 | 4 |
| | 0.117737 | -0.04365 | Q9UNE7 | Q9UNE7 | STUB1 | E3 ubiquiti | Color2 [Da | 1 | 4 |
| + | 1.974359 | 0.43321 | Q9UNF0 | Q9UNF0 | PACSIN2 | Protein kir | Color2 [Da | 1 | 4 |
| | 0.191376 | 0.083717 | Q9UNF1 | Q9UNF1 | MAGED2 | Melanoma | Color2 [Da | 1 | 4 |
| + | 0.79169 | -0.61934 | Q9UNH6 | Q9UNH6 | SNX7 | Sorting ne: | Color2 [Da | 1 | 4 |
| | 0.43411 | 0.038881 | Q9UNH7 | Q9UNH7 | SNX6 | Sorting ne: | Color2 [Da | 1 | 4 |
| | 0.928487 | -0.25391 | Q9UNI6 | Q9UNI6 | DUSP12 | Dual speci | Color2 [Da | 1 | 4 |
| + | 1.679045 | -0.61605 | Q9UNK0 | Q9UNK0 | STX8 | Syntaxin-8 | Color2 [Da | 1 | 4 |
| | 0 | -1.25756 | Q9UNL2 | Q9UNL2 | SSR3 | Translocor | Color2 [Da | 1 | 4 |
| + | 2.705405 | -0.87726 | Q9UNM6 | Q9UNM6 | PSMD13 | 26S protea | Color2 [Da | 1 | 4 |
| + | 1.882403 | 0.460762 | Q9UNN5 | Q9UNN5 | FAF1 | FAS-associ | Color2 [Da | 1 | 4 |
| | 0.0727 | 0.02668 | Q9UNP9 | Q9UNP9 | PPIE | Peptidyl-p | Color2 [Da | 1 | 4 |
| | 0.470841 | 0.122206 | Q9UNQ2 | Q9UNQ2 | DIMT1 | Probable c | Color2 [Da | 1 | 4 |
| + | 3.133078 | -1.28196 | Q9UNS2 | Q9UNS2 | COPS3 | COP9 signa | Color2 [Da | 1 | 4 |
| + | 2.353616 | -0.76726 | Q9UNW1 | Q9UNW1 | MINPP1 | Multiple ir | Color2 [Da | 1 | 4 |
| + | 2.530826 | 0.38898 | Q9UNW9 | Q9UNW9 | NOVA2 | RNA-bindi | Color2 [Da | 1 | 4 |
| + | 2.176816 | 1.273718 | Q9UNX3 | Q9UNX3 | RPL26L1 | 60S riboso | Color2 [Da | 1 | 4 |
| | 0 | -0.26216 | Q9UNX4 | Q9UNX4 | WDR3 | WD repeat | Color2 [Da | 1 | 4 |
| + | 3.920262 | 2.063146 | Q9UNZ2 | Q9UNZ2 | NSFL1C | NSFL1 cofa | Color2 [Da | 1 | 4 |
| | 0 | -0.03664 | Q9UP65 | Q9UP65 | PLA2G4C | Cytosolic p | Color2 [Da | 1 | 4 |
| + | 1.339828 | -0.88957 | Q9UP83 | Q9UP83 | COG5 | Conserved | Color2 [Da | 1 | 4 |
| + | 2.509745 | 1.303313 | Q9UPA5 | Q9UPA5 | BSN | Protein ba | Color2 [Da | 1 | 4 |
| | 0 | 0.412623 | Q9UPM9 | Q9UPM9 | B9D1 | B9 domain | Color2 [Da | 1 | 4 |
| + | 2.996921 | -0.37929 | Q9UPN3 | Q9UPN3 | MACF1 | Microtubu | Color2 [Da | 1 | 4 |
| | 0.570508 | -0.26489 | Q9UPN4 | Q9UPN4 | CEP131 | Centrosom | Color2 [Da | 1 | 4 |
| | 0.849012 | -0.20585 | Q9UPN6 | Q9UPN6 | SCAF8 | SR-related | Color2 [Da | 1 | 4 |
| + | 0.932994 | 1.010975 | Q9UPN7 | Q9UPN7 | PPP6R1 | Serine/thr | Color2 [Da | 1 | 4 |
| | 0.105319 | -0.08269 | Q9UPN9 | Q9UPN9 | TRIM33 | E3 ubiquiti | Color2 [Da | 1 | 4 |
| | 0.218563 | -0.09071 | Q9UPQ0 | Q9UPQ0 | LIMCH1 | LIM and ca | Color2 [Da | 1 | 4 |
| | 0.866489 | 0.238619 | Q9UPQ3 | Q9UPQ3 | AGAP1 | Arf-GAP w | Color2 [Da | 1 | 4 |
| | 0 | -0.09327 | Q9UPQ7 | Q9UPQ7 | PDZRN3 | E3 ubiquiti | Color2 [Da | 1 | 4 |
| | 0 | NaN | Q9UPQ8 | Q9UPQ8 | DOLK | Dolichol ki | Color2 [Da | 1 | 4 |
| + | 1.334232 | -1.04504 | Q9UPQ9 | Q9UPQ9 | TNRC6B | Trinucleoti | Color2 [Da | 1 | 4 |
| + | 2.255961 | 0.825907 | Q9UPR0 | Q9UPR0 | PLCL2 | Inactive pl | Color2 [Da | 1 | 4 |
| | 0 | -0.77601 | Q9UPR3 | Q9UPR3 | SMG5 | Nonsense- | Color2 [Da | 1 | 4 |
| | 0.25897 | 0.190177 | Q9UPR5 | Q9UPR5 | SLC8A2 | Sodium/ca | Color2 [Da | 1 | 4 |
| + | 1.204162 | 1.116336 | Q9UPS6 | Q9UPS6 | SETD1B | Histone-ly: | Color2 [Da | 1 | 4 |
| | 0.87773 | -0.26986 | Q9UPT5 | Q9UPT5 | EXOC7 | Exocyst co | Color2 [Da | 1 | 4 |
| | 0.3193 | -0.067 | Q9UPT6 | Q9UPT6 | MAPK8IP3 | C-Jun-amir | Color2 [Da | 1 | 4 |
| | 0.241512 | 0.098365 | Q9UPT8 | Q9UPT8 | ZC3H4 | Zinc finger | Color2 [Da | 1 | 4 |
| + | 0.874986 | -0.9504 | Q9UPT9 | Q9UPT9 | USP22 | Ubiquitin c | Color2 [Da | 1 | 4 |
| + | 1.916034 | -1.52363 | Q9UPU5 | Q9UPU5 | USP24 | Ubiquitin c | Color2 [Da | 1 | 4 |
| + | 4.373607 | 3.064185 | Q9UPV7 | Q9UPV7 | PHF24 | PHD finger | Color2 [Da | 1 | 4 |
| + | 0.762753 | 1.495182 | Q9UPV9 | Q9UPV9 | TRAK1 | Trafficking | Color2 [Da | 1 | 4 |
| | 0.498698 | 0.24317 | Q9UPW5 | Q9UPW5 | AGTPBP1 | Cytosolic c | Color2 [Da | 1 | 4 |
| + | 1.160041 | 1.197177 | Q9UPW8 | Q9UPW8 | UNC13A | Protein un | Color2 [Da | 1 | 4 |

| | | | | | | | | | |
|---|---|---|---|---|---|---|---|---|---|
| + | 0.898386 | 0.611008 | Q9UPX8 | Q9UPX8 | SHANK2 | SH3 and m | Color2 [Da | 1 | 4 |
| | 0.622762 | 0.208925 | Q9UPY3 | Q9UPY3 | DICER1 | Endoribon | Color2 [Da | 1 | 4 |
| + | 3.214671 | 2.630482 | Q9UPY6 | Q9UPY6 | WASF3 | Wiskott-Al | Color2 [Da | 1 | 4 |
| + | 2.951642 | 1.238376 | Q9UPY8 | Q9UPY8 | MAPRE3 | Microtubu | Color2 [Da | 1 | 4 |
| | 0 | NaN | Q9UPZ3 | Q9UPZ3 | HPS5 | Hermansky | Color2 [Da | 1 | 4 |
| | 0.217305 | 0.196827 | Q9UPZ6 | Q9UPZ6 | THSD7A | Thrombos | Color2 [Da | 1 | 4 |
| | 0.915014 | 0.23166 | Q9UQ03 | Q9UQ03 | CORO2B | Coronin-2E | Color2 [Da | 1 | 4 |
| + | 1.163143 | 0.48093 | Q9UQ16 | Q9UQ16 | DNM3 | Dynamin-3 | Color2 [Da | 1 | 4 |
| + | 2.87752 | 0.377756 | Q9UQ35 | Q9UQ35 | SRRM2 | Serine/arg | Color2 [Da | 1 | 4 |
| | 0 | -0.8842 | Q9UQ49 | Q9UQ49 | NEU3 | Sialidase-3 | Color2 [Da | 1 | 4 |
| + | 2.736888 | -0.31195 | Q9UQ80 | Q9UQ80 | PA2G4 | Proliferatio | Color2 [Da | 1 | 4 |
| + | 1.321988 | 0.262616 | Q9UQ90 | Q9UQ90 | SPG7 | Paraplegin | Color2 [Da | 1 | 4 |
| | 0.602663 | -0.10031 | Q9UQB3 | Q9UQB3 | CTNND2 | Catenin de | Color2 [Da | 1 | 4 |
| | 0.341491 | -0.14154 | Q9UQB8 | Q9UQB8 | BAIAP2 | Brain-spec | Color2 [Da | 1 | 4 |
| | 0.019795 | 0.016882 | Q9UQC2 | Q9UQC2 | GAB2 | GRB2-asso | Color2 [Da | 1 | 4 |
| + | 3.456339 | -0.45458 | Q9UQE7 | Q9UQE7 | SMC3 | Structural | Color2 [Da | 1 | 4 |
| + | 1.120009 | 0.596842 | Q9UQF2 | Q9UQF2 | MAPK8IP1 | C-Jun-amir | Color2 [Da | 1 | 4 |
| + | 1.033999 | 0.80103 | Q9UQL6 | Q9UQL6 | HDAC5 | Histone de | Color2 [Da | 1 | 4 |
| + | 4.091405 | 0.931186 | Q9UQM7 | Q9UQM7 | CAMK2A | Calcium/ca | Color2 [Da | 1 | 4 |
| + | 1.506419 | 1.088289 | Q9UQN3 | Q9UQN3 | CHMP2B | Charged m | Color2 [Da | 1 | 4 |
| | 0.48734 | -0.63184 | Q9UQR0 | Q9UQR0 | SCML2 | Sex comb o | Color2 [Da | 1 | 4 |
| + | 1.220258 | -0.58673 | Q9UQR1 | Q9UQR1 | ZNF148 | Zinc finger | Color2 [Da | 1 | 4 |
| | 0 | 0.613816 | Q9Y217 | Q9Y217 | MTMR6 | Myotubula | Color2 [Da | 1 | 4 |
| | 0 | -0.00115 | Q9Y221 | Q9Y221 | NIP7 | 60S riboso | Color2 [Da | 1 | 4 |
| + | 2.278178 | -1.94821 | Q9Y223 | Q9Y223 | GNE | Bifunction | Color2 [Da | 1 | 4 |
| + | 3.853229 | 0.940162 | Q9Y224 | Q9Y224 | RTRAF | RNA transc | Color2 [Da | 1 | 4 |
| | 1.656477 | -0.1174 | Q9Y230 | Q9Y230 | RUVBL2 | RuvB-like 2 | Color2 [Da | 1 | 4 |
| | 0 | 0.620601 | Q9Y231 | Q9Y231 | FUT9 | 4-galactos | Color2 [Da | 1 | 4 |
| | 0 | -1.38775 | Q9Y232 | Q9Y232 | CDYL | Chromodo | Color2 [Da | 1 | 4 |
| | 0.557406 | -0.23245 | Q9Y233 | Q9Y233 | PDE10A | cAMP and | Color2 [Da | 1 | 4 |
| + | 1.346019 | 0.456274 | Q9Y237 | Q9Y237 | PIN4 | Peptidyl-p | Color2 [Da | 1 | 4 |
| + | 3.201839 | 7.911285 | Q9Y241 | Q9Y241 | HIGD1A | HIG1 doma | Color2 [Da | 1 | 4 |
| + | 1.965437 | -0.59514 | Q9Y243 | Q9Y243 | AKT3 | RAC-gamm | Color2 [Da | 1 | 4 |
| + | 0.745263 | 0.851412 | Q9Y244 | Q9Y244 | POMP | Proteasom | Color2 [Da | 1 | 4 |
| | 0.596394 | 0.676004 | Q9Y247 | Q9Y247 | FAM50B | Protein FA | Color2 [Da | 1 | 4 |
| | 0 | 0.437285 | Q9Y250 | Q9Y250 | LZTS1 | Leucine zip | Color2 [Da | 1 | 4 |
| | 0 | NaN | Q9Y259 | Q9Y259 | CHKB | Choline/et | Color2 [Da | 1 | 4 |
| + | 2.668584 | -0.65314 | Q9Y262 | Q9Y262 | EIF3L | Eukaryotic | Color2 [Da | 1 | 4 |
| + | 2.97659 | -0.78785 | Q9Y263 | Q9Y263 | PLAA | Phospholip | Color2 [Da | 1 | 4 |
| + | 3.417275 | -0.47545 | Q9Y265 | Q9Y265 | RUVBL1 | RuvB-like 1 | Color2 [Da | 1 | 4 |
| + | 3.945537 | 1.446871 | Q9Y266 | Q9Y266 | NUDC | Nuclear m | Color2 [Da | 1 | 4 |
| | 0.15577 | 0.007075 | Q9Y276 | Q9Y276 | BCS1L | Mitochonc | Color2 [Da | 1 | 4 |
| | 0.076618 | 0.022471 | Q9Y277 | Q9Y277 | VDAC3 | Voltage-de | Color2 [Da | 1 | 4 |
| + | 1.829696 | 1.474522 | Q9Y281 | Q9Y281 | CFL2 | Cofilin-2 | Color2 [Da | 1 | 4 |
| | 1.105822 | 0.185476 | Q9Y282 | Q9Y282 | ERGIC3 | Endoplasm | Color2 [Da | 1 | 4 |
| + | 3.153606 | 1.295499 | Q9Y285 | Q9Y285 | FARSA | Phenylalar | Color2 [Da | 1 | 4 |
| | 0.556895 | -0.32138 | Q9Y287 | Q9Y287 | ITM2B | Integral m | Color2 [Da | 1 | 4 |

| | | | | | | | | | |
|---|---|---|---|---|---|---|---|---|---|
| | 0 | NaN | Q9Y291 | Q9Y291 | MRPS33 | 28S riboso | Color2 [Da | 1 | 4 |
| + | 2.084912 | -1.3999 | Q9Y294 | Q9Y294 | ASF1A | Histone ch | Color2 [Da | 1 | 4 |
| + | 3.343024 | -0.75247 | Q9Y295 | Q9Y295 | DRG1 | Developm | Color2 [Da | 1 | 4 |
| + | 2.234299 | 0.997455 | Q9Y296 | Q9Y296 | TRAPPC4 | Trafficking | Color2 [Da | 1 | 4 |
| + | 3.51917 | -1.10024 | Q9Y2A7 | Q9Y2A7 | NCKAP1 | Nck-associ | Color2 [Da | 1 | 4 |
| | 0.688429 | -0.34845 | Q9Y2B0 | Q9Y2B0 | CNPY2 | Protein ca | Color2 [Da | 1 | 4 |
| + | 1.937789 | 0.963522 | Q9Y2B5 | Q9Y2B5 | VPS9D1 | VPS9 dom | Color2 [Da | 1 | 4 |
| + | 2.525884 | 1.559374 | Q9Y2C4 | Q9Y2C4 | EXOG | Nuclease E | Color2 [Da | 1 | 4 |
| | 0.654723 | -0.18538 | Q9Y2D4 | Q9Y2D4 | EXOC6B | Exocyst co | Color2 [Da | 1 | 4 |
| + | 2.125759 | -1.24714 | Q9Y2D5 | Q9Y2D5 | AKAP2 | A-kinase a | Color2 [Da | 1 | 4 |
| | 0.082229 | 0.067907 | Q9Y2E4 | Q9Y2E4 | DIP2C | Disco-inter | Color2 [Da | 1 | 4 |
| + | 1.23129 | 0.545193 | Q9Y2E5 | Q9Y2E5 | MAN2B2 | Epididymis | Color2 [Da | 1 | 4 |
| | 0.284532 | 0.203138 | Q9Y2G0 | Q9Y2G0 | EFR3B | Protein EF | Color2 [Da | 1 | 4 |
| | 0.188684 | -0.25236 | Q9Y2G2 | Q9Y2G2 | CARD8 | Caspase re | Color2 [Da | 1 | 4 |
| + | 2.281958 | -0.84527 | Q9Y2G5 | Q9Y2G5 | POFUT2 | GDP-fucos | Color2 [Da | 1 | 4 |
| | 0 | NaN | Q9Y2G8 | Q9Y2G8 | DNAJC16 | DnaJ homo | Color2 [Da | 1 | 4 |
| + | 2.505756 | 0.939979 | Q9Y2H0 | Q9Y2H0 | DLGAP4 | Disks large | Color2 [Da | 1 | 4 |
| | 0.477498 | -0.25532 | Q9Y2H1 | Q9Y2H1 | STK38L | Serine/thr | Color2 [Da | 1 | 4 |
| | 0.877617 | 0.246499 | Q9Y2H2 | Q9Y2H2 | INPP5F | Phosphatio | Color2 [Da | 1 | 4 |
| | 0.925149 | 0.193171 | Q9Y2H5 | Q9Y2H5 | PLEKHA6 | Pleckstrin | Color2 [Da | 1 | 4 |
| | 0.38738 | -0.18949 | Q9Y2H6 | Q9Y2H6 | FNDC3A | Fibronecti | Color2 [Da | 1 | 4 |
| + | 1.838114 | 0.881729 | Q9Y2H9 | Q9Y2H9 | MAST1 | Microtubu | Color2 [Da | 1 | 4 |
| + | 0.901283 | -0.47752 | Q9Y2I1 | Q9Y2I1 | NISCH | Nischarin | Color2 [Da | 1 | 4 |
| | 0.081192 | -0.04668 | Q9Y2I7 | Q9Y2I7 | PIKFYVE | 1-phospha | Color2 [Da | 1 | 4 |
| | 0.970167 | 0.264713 | Q9Y2I8 | Q9Y2I8 | WDR37 | WD repeat | Color2 [Da | 1 | 4 |
| | 0 | NaN | Q9Y2J0 | Q9Y2J0 | RPH3A | Rabphilin-3 | Color2 [Da | 1 | 4 |
| + | 1.910884 | 0.354058 | Q9Y2J2 | Q9Y2J2 | EPB41L3 | Band 4.1-li | Color2 [Da | 1 | 4 |
| | 0.177205 | 0.047959 | Q9Y2J4 | Q9Y2J4 | AMOTL2 | Angiomoti | Color2 [Da | 1 | 4 |
| + | 1.660629 | -1.96097 | Q9Y2K2 | Q9Y2K2 | SIK3 | Serine/thr | Color2 [Da | 1 | 4 |
| | 0.668771 | 0.238274 | Q9Y2K5 | Q9Y2K5 | R3HDM2 | R3H doma | Color2 [Da | 1 | 4 |
| + | 2.913978 | -0.94441 | Q9Y2L1 | Q9Y2L1 | DIS3 | Exosome c | Color2 [Da | 1 | 4 |
| | 0.457526 | 0.254462 | Q9Y2L5 | Q9Y2L5 | TRAPPC8 | Trafficking | Color2 [Da | 1 | 4 |
| | 0 | -0.7118 | Q9Y2L9 | Q9Y2L9 | LRCH1 | Leucine-ric | Color2 [Da | 1 | 4 |
| | 0 | -1.49084 | Q9Y2P8 | Q9Y2P8 | RCL1 | RNA 3'-ter | Color2 [Da | 1 | 4 |
| + | 1.448557 | -0.35157 | Q9Y2Q0 | Q9Y2Q0 | ATP8A1 | Phospholip | Color2 [Da | 1 | 4 |
| | 0.439081 | 0.231542 | Q9Y2Q3 | Q9Y2Q3 | GSTK1 | Glutathion | Color2 [Da | 1 | 4 |
| + | 3.228718 | 2.215597 | Q9Y2Q5 | Q9Y2Q5 | LAMTOR2 | Ragulator | Color2 [Da | 1 | 4 |
| + | 1.917206 | 0.710802 | Q9Y2Q9 | Q9Y2Q9 | MRPS28 | 28S riboso | Color2 [Da | 1 | 4 |
| + | 3.498475 | 3.163991 | Q9Y2R0 | Q9Y2R0 | COA3 | Cytochrom | Color2 [Da | 1 | 4 |
| + | 1.039348 | -0.49991 | Q9Y2R4 | Q9Y2R4 | DDX52 | Probable A | Color2 [Da | 1 | 4 |
| | 0.643017 | 0.277275 | Q9Y2R9 | Q9Y2R9 | MRPS7 | 28S riboso | Color2 [Da | 1 | 4 |
| + | 2.23329 | -0.56811 | Q9Y2S2 | Q9Y2S2 | CRYL1 | Lambda-cr | Color2 [Da | 1 | 4 |
| + | 2.38694 | 0.87045 | Q9Y2S7 | Q9Y2S7 | POLDIP2 | Polymeras | Color2 [Da | 1 | 4 |
| | 0.762146 | -0.36769 | Q9Y2T2 | Q9Y2T2 | AP3M1 | AP-3 comp | Color2 [Da | 1 | 4 |
| + | 2.985777 | 1.539931 | Q9Y2T7 | Q9Y2T7 | YBX2 | Y-box-bind | Color2 [Da | 1 | 4 |
| | 0 | NaN | Q9Y2U8 | Q9Y2U8 | LEMD3 | Inner nucle | Color2 [Da | 1 | 4 |
| + | 1.931073 | 0.664387 | Q9Y2V2 | Q9Y2V2 | CARHSP1 | Calcium-re | Color2 [Da | 1 | 4 |

| | | | | | | | | | |
|---|---|---|---|---|---|---|---|---|---|
| + | 1.07405 | -0.98131 | Q9Y2V7 | Q9Y2V7 | COG6 | Conserved | Color2 [Da | 1 | 4 |
| + | 2.173685 | 0.562493 | Q9Y2W1 | Q9Y2W1 | THRAP3 | Thyroid hc | Color2 [Da | 1 | 4 |
| + | 2.274744 | 1.516826 | Q9Y2W2 | Q9Y2W2 | WBP11 | WW doma | Color2 [Da | 1 | 4 |
| | 0.474043 | 0.367903 | Q9Y2W6 | Q9Y2W6 | TDRKH | Tudor and | Color2 [Da | 1 | 4 |
| | 0 | NaN | Q9Y2X0 | Q9Y2X0 | MED16 | Mediator ( | Color2 [Da | 1 | 4 |
| + | 3.243661 | -1.00166 | Q9Y2X3 | Q9Y2X3 | NOP58 | Nucleolar | | Color2 [Da | 1 | 4 |
| + | 1.817152 | 0.827982 | Q9Y2X7 | Q9Y2X7 | GIT1 | ARF GTPas | Color2 [Da | 1 | 4 |
| + | 1.605694 | -0.31022 | Q9Y2X9 | Q9Y2X9 | ZNF281 | Zinc finger | Color2 [Da | 1 | 4 |
| | 0.401627 | -0.21384 | Q9Y2Y0 | Q9Y2Y0 | ARL2BP | ADP-ribos\ | Color2 [Da | 1 | 4 |
| | 0 | -1.21829 | Q9Y2Y9 | Q9Y2Y9 | KLF13 | Krueppel-l | Color2 [Da | 1 | 4 |
| + | 2.598432 | 0.509443 | Q9Y2Z0 | Q9Y2Z0 | SUGT1 | Protein SG | Color2 [Da | 1 | 4 |
| + | 0.855222 | -0.53244 | Q9Y2Z4 | Q9Y2Z4 | YARS2 | Tyrosine--t | Color2 [Da | 1 | 4 |
| | 0 | 2.404527 | Q9Y2Z9 | Q9Y2Z9 | COQ6 | Ubiquinon | Color2 [Da | 1 | 4 |
| | 0 | -0.34483 | Q9Y303 | Q9Y303 | AMDHD2 | N-acetylglu | Color2 [Da | 1 | 4 |
| + | 1.582167 | -1.16649 | Q9Y305 | Q9Y305 | ACOT9 | Acyl-coenz | Color2 [Da | 1 | 4 |
| | 0.529857 | -0.24707 | Q9Y312 | Q9Y312 | AAR2 | Protein AA | Color2 [Da | 1 | 4 |
| + | 1.190667 | -0.65518 | Q9Y314 | Q9Y314 | NOSIP | Nitric oxid | Color2 [Da | 1 | 4 |
| + | 3.101174 | -0.98688 | Q9Y315 | Q9Y315 | DERA | Deoxyribo: | Color2 [Da | 1 | 4 |
| | 0.646045 | -0.642 | Q9Y316 | Q9Y316 | MEMO1 | Protein MI | Color2 [Da | 1 | 4 |
| + | 0.986639 | -0.58957 | Q9Y320 | Q9Y320 | TMX2 | Thioredoxi | Color2 [Da | 1 | 4 |
| | 0 | 1.538905 | Q9Y328 | Q9Y328 | NSG2 | Neuronal \ | Color2 [Da | 1 | 4 |
| + | 2.220628 | 1.267422 | Q9Y333 | Q9Y333 | LSM2 | U6 snRNA- | Color2 [Da | 1 | 4 |
| + | 2.757021 | 1.296695 | Q9Y343 | Q9Y343 | SNX24 | Sorting ne: | Color2 [Da | 1 | 4 |
| + | 1.164714 | 1.035263 | Q9Y366 | Q9Y366 | IFT52 | Intraflagell | Color2 [Da | 1 | 4 |
| + | 2.635843 | 0.71855 | Q9Y371 | Q9Y371 | SH3GLB1 | Endophilin | Color2 [Da | 1 | 4 |
| + | 2.072402 | 0.76913 | Q9Y375 | Q9Y375 | NDUFAF1 | Complex I | Color2 [Da | 1 | 4 |
| + | 3.173638 | 0.922153 | Q9Y376 | Q9Y376 | CAB39 | Calcium-bi | Color2 [Da | 1 | 4 |
| + | 1.817557 | -0.78863 | Q9Y383 | Q9Y383 | LUC7L2 | Putative R | Color2 [Da | 1 | 4 |
| + | 2.804799 | -1.86857 | Q9Y385 | Q9Y385 | UBE2J1 | Ubiquitin-( | Color2 [Da | 1 | 4 |
| | 0.351073 | 0.244916 | Q9Y388 | Q9Y388 | RBMX2 | RNA-bindi | Color2 [Da | 1 | 4 |
| + | 1.757452 | -0.65142 | Q9Y394 | Q9Y394 | DHRS7 | Dehydroge | Color2 [Da | 1 | 4 |
| + | 0.929791 | -0.48284 | Q9Y399 | Q9Y399 | MRPS2 | 28S riboso | Color2 [Da | 1 | 4 |
| + | 1.513077 | -0.93679 | Q9Y3A2 | Q9Y3A2 | UTP11 | Probable U | Color2 [Da | 1 | 4 |
| + | 1.341784 | 1.240228 | Q9Y3A3 | Q9Y3A3 | MOB4 | MOB-like p | Color2 [Da | 1 | 4 |
| | 0.2954 | 0.089498 | Q9Y3A5 | Q9Y3A5 | SBDS | Ribosome | Color2 [Da | 1 | 4 |
| + | 3.212174 | -1.24984 | Q9Y3A6 | Q9Y3A6 | TMED5 | Transmem | Color2 [Da | 1 | 4 |
| | 0.475108 | 0.325963 | Q9Y3B2 | Q9Y3B2 | EXOSC1 | Exosome c | Color2 [Da | 1 | 4 |
| + | 2.046039 | -0.84342 | Q9Y3B3 | Q9Y3B3 | TMED7 | Transmem | Color2 [Da | 1 | 4 |
| + | 2.221043 | 1.004585 | Q9Y3B4 | Q9Y3B4 | SF3B6 | Splicing fa | Color2 [Da | 1 | 4 |
| + | 1.815861 | 0.762219 | Q9Y3B7 | Q9Y3B7 | MRPL11 | 39S riboso | Color2 [Da | 1 | 4 |
| + | 3.101101 | 1.962646 | Q9Y3B8 | Q9Y3B8 | REXO2 | Oligoribon | Color2 [Da | 1 | 4 |
| + | 1.148312 | 0.597162 | Q9Y3B9 | Q9Y3B9 | RRP15 | RRP15-like | Color2 [Da | 1 | 4 |
| + | 2.279231 | 1.956929 | Q9Y3C0 | Q9Y3C0 | WASHC3 | WASH con | Color2 [Da | 1 | 4 |
| | 0.715014 | 0.379747 | Q9Y3C1 | Q9Y3C1 | NOP16 | Nucleolar | | Color2 [Da | 1 | 4 |
| | 0.147997 | 0.162778 | Q9Y3C4 | Q9Y3C4 | TPRKB | EKC/KEOP! | Color2 [Da | 1 | 4 |
| | 0.50263 | -0.25269 | Q9Y3C5 | Q9Y3C5 | RNF11 | RING finge | Color2 [Da | 1 | 4 |
| | 0.24688 | 0.082443 | Q9Y3C6 | Q9Y3C6 | PPIL1 | Peptidyl-pı | Color2 [Da | 1 | 4 |

| | | | | | | | | | | |
|---|---|---|---|---|---|---|---|---|---|---|
| | 0.06641 | 0.028888 | Q9Y3C8 | Q9Y3C8 | UFC1 | Ubiquitin-f | Color2 | [Da | 1 | 4 |
| | 0.218814 | 0.102002 | Q9Y3D0 | Q9Y3D0 | CIAO2B | Cytosolic ir | Color2 | [Da | 1 | 4 |
| | 0.033147 | 0.071056 | Q9Y3D3 | Q9Y3D3 | MRPS16 | 28S riboso | Color2 | [Da | 1 | 4 |
| | 0 | NaN | Q9Y3D5 | Q9Y3D5 | MRPS18C | 28S riboso | Color2 | [Da | 1 | 4 |
| + | 3.738884 | 2.316609 | Q9Y3D6 | Q9Y3D6 | FIS1 | Mitochonc | Color2 | [Da | 1 | 4 |
| + | 4.21954 | 2.303985 | Q9Y3D7 | Q9Y3D7 | PAM16 | Mitochonc | Color2 | [Da | 1 | 4 |
| + | 2.148037 | 1.469695 | Q9Y3D9 | Q9Y3D9 | MRPS23 | 28S riboso | Color2 | [Da | 1 | 4 |
| + | 1.254761 | -1.69591 | Q9Y3E0 | Q9Y3E0 | GOLT1B | Vesicle tra | Color2 | [Da | 1 | 4 |
| + | 1.53901 | 1.13096 | Q9Y3E1 | Q9Y3E1 | HDGFL3 | Hepatoma | Color2 | [Da | 1 | 4 |
| | 0.33707 | 0.64613 | Q9Y3E2 | Q9Y3E2 | BOLA1 | BolA-like p | Color2 | [Da | 1 | 4 |
| + | 2.192665 | 2.146591 | Q9Y3E5 | Q9Y3E5 | PTRH2 | Peptidyl-tF | Color2 | [Da | 1 | 4 |
| | 0.534682 | 0.24475 | Q9Y3E7 | Q9Y3E7 | CHMP3 | Charged m | Color2 | [Da | 1 | 4 |
| + | 3.106167 | -1.44125 | Q9Y3F4 | Q9Y3F4 | STRAP | Serine-thr | Color2 | [Da | 1 | 4 |
| + | 1.900427 | -0.98141 | Q9Y3I0 | Q9Y3I0 | RTCB | RNA-splicii | Color2 | [Da | 1 | 4 |
| + | 1.066243 | -0.46397 | Q9Y3I1 | Q9Y3I1 | FBXO7 | F-box only | Color2 | [Da | 1 | 4 |
| | 0.590287 | -0.26766 | Q9Y3L5 | Q9Y3L5 | RAP2C | Ras-relate | Color2 | [Da | 1 | 4 |
| | 0.119428 | -0.39876 | Q9Y3M2 | Q9Y3M2 | CBY1 | Protein ch | Color2 | [Da | 1 | 4 |
| + | 2.343614 | -1.07381 | Q9Y3P9 | Q9Y3P9 | RABGAP1 | Rab GTPas | Color2 | [Da | 1 | 4 |
| | 0 | -1.99556 | Q9Y3Q3 | Q9Y3Q3 | TMED3 | Transmem | Color2 | [Da | 1 | 4 |
| + | 3.272305 | 1.230537 | Q9Y3Q4 | Q9Y3Q4 | HCN4 | Potassium | Color2 | [Da | 1 | 4 |
| + | 1.376741 | 1.112844 | Q9Y3Q8 | Q9Y3Q8 | TSC22D4 | TSC22 don | Color2 | [Da | 1 | 4 |
| + | 1.339438 | 0.420695 | Q9Y3R0 | Q9Y3R0 | GRIP1 | Glutamate | Color2 | [Da | 1 | 4 |
| | 0 | NaN | Q9Y3R5 | Q9Y3R5 | DOP1B | Protein do | Color2 | [Da | 1 | 4 |
| + | 2.372525 | 0.593423 | Q9Y3S1 | Q9Y3S1 | WNK2 | Serine/thr | Color2 | [Da | 1 | 4 |
| + | 1.643905 | -1.74061 | Q9Y3S2 | Q9Y3S2 | ZNF330 | Zinc finger | Color2 | [Da | 1 | 4 |
| | 1.023247 | 0.281143 | Q9Y3T6 | Q9Y3T6 | R3HCC1 | R3H and co | Color2 | [Da | 1 | 4 |
| | 0 | -0.17633 | Q9Y3T9 | Q9Y3T9 | NOC2L | Nucleolar | Color2 | [Da | 1 | 4 |
| + | 1.751588 | 0.465069 | Q9Y3U8 | Q9Y3U8 | RPL36 | 60S riboso | Color2 | [Da | 1 | 4 |
| + | 0.993072 | 1.526415 | Q9Y3X0 | Q9Y3X0 | CCDC9 | Coiled-coil | Color2 | [Da | 1 | 4 |
| + | 3.237572 | 1.567241 | Q9Y3Y2 | Q9Y3Y2 | CHTOP | Chromatin | Color2 | [Da | 1 | 4 |
| + | 1.908823 | -0.88754 | Q9Y3Z3 | Q9Y3Z3 | SAMHD1 | Deoxynucl | Color2 | [Da | 1 | 4 |
| + | 1.409962 | -0.41295 | Q9Y426 | Q9Y426 | C2CD2 | C2 domain | Color2 | [Da | 1 | 4 |
| | 0.558733 | -0.12856 | Q9Y450 | Q9Y450 | HBS1L | HBS1-like | Color2 | [Da | 1 | 4 |
| | 0 | NaN | Q9Y466 | Q9Y466 | NR2E1 | Nuclear re | Color2 | [Da | 1 | 4 |
| | 0.568227 | -0.23867 | Q9Y467 | Q9Y467 | SALL2 | Sal-like prc | Color2 | [Da | 1 | 4 |
| | 0.20427 | 0.117812 | Q9Y478 | Q9Y478 | PRKAB1 | 5'-AMP-ac | Color2 | [Da | 1 | 4 |
| + | 2.235045 | -0.96332 | Q9Y490 | Q9Y490 | TLN1 | Talin-1 | Color2 | [Da | 1 | 4 |
| | 0.619073 | 0.160718 | Q9Y496 | Q9Y496 | KIF3A | Kinesin-lik | Color2 | [Da | 1 | 4 |
| | 0.635322 | -0.28218 | Q9Y4A5 | Q9Y4A5 | TRRAP | Transform | Color2 | [Da | 1 | 4 |
| + | 1.062187 | 0.474475 | Q9Y4B5 | Q9Y4B5 | MTCL1 | Microtubu | Color2 | [Da | 1 | 4 |
| + | 2.573594 | 1.083606 | Q9Y4B6 | Q9Y4B6 | DCAF1 | DDB1- and | Color2 | [Da | 1 | 4 |
| + | 1.935259 | 2.586526 | Q9Y4C0 | Q9Y4C0 | NRXN3 | Neurexin-3 | Color2 | [Da | 1 | 4 |
| | 0 | -0.56373 | Q9Y4C2 | Q9Y4C2 | TCAF1 | TRPM8 cha | Color2 | [Da | 1 | 4 |
| + | 1.247599 | -0.37057 | Q9Y4C8 | Q9Y4C8 | RBM19 | Probable F | Color2 | [Da | 1 | 4 |
| + | 3.180725 | 0.496741 | Q9Y4D1 | Q9Y4D1 | DAAM1 | Disheveled | Color2 | [Da | 1 | 4 |
| + | 0.849494 | -0.49953 | Q9Y4D8 | Q9Y4D8 | HECTD4 | Probable E | Color2 | [Da | 1 | 4 |
| | 0.731177 | 0.332418 | Q9Y4E1 | Q9Y4E1 | WASHC2C | WASH con | Color2 | [Da | 1 | 4 |

| | | | | | | | | | |
|---|---|---|---|---|---|---|---|---|---|
| | 0.088512 | -0.04983 | Q9Y4E6 | Q9Y4E6 | WDR7 | WD repeat | Color2 [Da | 1 | 4 |
| | 0.781267 | -0.25471 | Q9Y4E8 | Q9Y4E8 | USP15 | Ubiquitin c | Color2 [Da | 1 | 4 |
| + | 1.931975 | -0.68458 | Q9Y4F1 | Q9Y4F1 | FARP1 | FERM, ARH | Color2 [Da | 1 | 4 |
| + | 2.033116 | 0.525436 | Q9Y4F5 | Q9Y4F5 | CEP170B | Centrosom | Color2 [Da | 1 | 4 |
| | 0.823344 | 0.087271 | Q9Y4G6 | Q9Y4G6 | TLN2 | Talin-2 | Color2 [Da | 1 | 4 |
| | 0.343283 | 0.087952 | Q9Y4G8 | Q9Y4G8 | RAPGEF2 | Rap guanir | Color2 [Da | 1 | 4 |
| + | 1.92866 | -1.47938 | Q9Y4H2 | Q9Y4H2 | IRS2 | Insulin rec | Color2 [Da | 1 | 4 |
| + | 3.139305 | 0.443875 | Q9Y4I1 | Q9Y4I1 | MYO5A | Unconvent | Color2 [Da | 1 | 4 |
| + | 1.799786 | 0.880065 | Q9Y4J8 | Q9Y4J8 | DTNA | Dystrobrev | Color2 [Da | 1 | 4 |
| | 0 | NaN | Q9Y4K3 | Q9Y4K3 | TRAF6 | TNF recept | Color2 [Da | 1 | 4 |
| | 0 | 0.396181 | Q9Y4K4 | Q9Y4K4 | MAP4K5 | Mitogen-a | Color2 [Da | 1 | 4 |
| + | 1.787311 | -0.20824 | Q9Y4L1 | Q9Y4L1 | HYOU1 | Hypoxia up | Color2 [Da | 1 | 4 |
| | 0.234369 | 0.261803 | Q9Y4L5 | Q9Y4L5 | RNF115 | E3 ubiquiti | Color2 [Da | 1 | 4 |
| + | 2.701096 | -1.70194 | Q9Y4P1 | Q9Y4P1 | ATG4B | Cysteine p | Color2 [Da | 1 | 4 |
| + | 2.697124 | -1.45575 | Q9Y4P3 | Q9Y4P3 | TBL2 | Transducir | Color2 [Da | 1 | 4 |
| | 0.102114 | -0.03274 | Q9Y4P8 | Q9Y4P8 | WIPI2 | WD repeat | Color2 [Da | 1 | 4 |
| | 0 | NaN | Q9Y4R8 | Q9Y4R8 | TELO2 | Telomere l | Color2 [Da | 1 | 4 |
| + | 1.980915 | -0.67543 | Q9Y4W2 | Q9Y4W2 | LAS1L | Ribosomal | Color2 [Da | 1 | 4 |
| | 0.264397 | -0.06008 | Q9Y4W6 | Q9Y4W6 | AFG3L2 | AFG3-like p | Color2 [Da | 1 | 4 |
| + | 2.318073 | -1.7741 | Q9Y4X5 | Q9Y4X5 | ARIH1 | E3 ubiquiti | Color2 [Da | 1 | 4 |
| + | 3.925374 | 1.44478 | Q9Y4Y9 | Q9Y4Y9 | LSM5 | U6 snRNA- | Color2 [Da | 1 | 4 |
| + | 1.586932 | 0.946123 | Q9Y4Z0 | Q9Y4Z0 | LSM4 | U6 snRNA- | Color2 [Da | 1 | 4 |
| | 0.131206 | -0.49904 | Q9Y508 | Q9Y508 | RNF114 | E3 ubiquiti | Color2 [Da | 1 | 4 |
| | 0.71404 | -0.28368 | Q9Y512 | Q9Y512 | SAMM50 | Sorting and | Color2 [Da | 1 | 4 |
| + | 1.396051 | 0.276595 | Q9Y520 | Q9Y520 | PRRC2C | Protein PR | Color2 [Da | 1 | 4 |
| | 0.030617 | -0.02353 | Q9Y530 | Q9Y530 | OARD1 | ADP-ribose | Color2 [Da | 1 | 4 |
| + | 2.418503 | 2.7988 | Q9Y534 | Q9Y534 | CSDC2 | Cold shock | Color2 [Da | 1 | 4 |
| + | 2.508502 | 0.629245 | Q9Y547 | Q9Y547 | HSPB11 | Intraflagell | Color2 [Da | 1 | 4 |
| | 0 | NaN | Q9Y566 | Q9Y566 | SHANK1 | SH3 and m | Color2 [Da | 1 | 4 |
| + | 1.028695 | -0.47482 | Q9Y570 | Q9Y570 | PPME1 | Protein ph | Color2 [Da | 1 | 4 |
| | 0 | NaN | Q9Y575 | Q9Y575 | ASB3 | Ankyrin re | Color2 [Da | 1 | 4 |
| | 0 | NaN | Q9Y576 | Q9Y576 | ASB1 | Ankyrin re | Color2 [Da | 1 | 4 |
| | 0.326266 | 0.120677 | Q9Y580 | Q9Y580 | RBM7 | RNA-bindi | Color2 [Da | 1 | 4 |
| + | 1.146949 | 1.872014 | Q9Y584 | Q9Y584 | TIMM22 | Mitochonc | Color2 [Da | 1 | 4 |
| | 0 | NaN | Q9Y586 | Q9Y586 | MAB21L2 | Protein ma | Color2 [Da | 1 | 4 |
| + | 1.210908 | -0.5588 | Q9Y587 | Q9Y587 | AP4S1 | AP-4 comp | Color2 [Da | 1 | 4 |
| | 0.423487 | -0.63489 | Q9Y5A7 | Q9Y5A7 | NUB1 | NEDD8 ult | Color2 [Da | 1 | 4 |
| + | 1.439858 | -0.4634 | Q9Y5A9 | Q9Y5A9 | YTHDF2 | YTH doma | Color2 [Da | 1 | 4 |
| | 0.360203 | 0.073304 | Q9Y5B0 | Q9Y5B0 | CTDP1 | RNA polym | Color2 [Da | 1 | 4 |
| + | 1.72866 | 1.027526 | Q9Y5B6 | Q9Y5B6 | PAXBP1 | PAX3- and | Color2 [Da | 1 | 4 |
| | 0.448023 | -0.72913 | Q9Y5B8 | Q9Y5B8 | NME7 | Nucleoside | Color2 [Da | 1 | 4 |
| + | 3.267099 | -0.75194 | Q9Y5B9 | Q9Y5B9 | SUPT16H | FACT comp | Color2 [Da | 1 | 4 |
| + | 1.637873 | -0.26898 | Q9Y5E7 | Q9Y5E7 | PCDHB2 | Protocadh | Color2 [Da | 1 | 4 |
| | 0 | 0.135551 | Q9Y5F7 | Q9Y5F7 | PCDHGC4 | Protocadh | Color2 [Da | 1 | 4 |
| + | 1.809556 | 0.744607 | Q9Y5F9 | Q9Y5F9 | PCDHGB6 | Protocadh | Color2 [Da | 1 | 4 |
| | 0 | NaN | Q9Y5H2;Q | Q9Y5H2;Q | PCDHGA1 | Protocadh | Color2 [Da | 1 | 4 |
| | 0 | -1.05548 | Q9Y5H9 | Q9Y5H9 | PCDHA2 | Protocadh | Color2 [Da | 1 | 4 |

| | | | | | | | | | |
|---|---|---|---|---|---|---|---|---|---|
| + | 1.412728 | -1.04358 | Q9Y5I1;Q9 | Q9Y5I1;Q9 | PCDHA11; | Protocadh | Color2 [Da | 1 | 4 |
| | 0.395415 | -0.4323 | Q9Y5J1 | Q9Y5J1 | UTP18 | U3 small n | Color2 [Da | 1 | 4 |
| + | 2.64395 | 2.662644 | Q9Y5J6 | Q9Y5J6 | TIMM10B | Mitochonc | Color2 [Da | 1 | 4 |
| + | 3.890839 | 2.501681 | Q9Y5J7 | Q9Y5J7 | TIMM9 | Mitochonc | Color2 [Da | 1 | 4 |
| + | 2.339117 | 1.43425 | Q9Y5J9 | Q9Y5J9 | TIMM8B | Mitochonc | Color2 [Da | 1 | 4 |
| + | 1.098113 | 0.535453 | Q9Y5K3 | Q9Y5K3 | PCYT1B | Choline-pł | Color2 [Da | 1 | 4 |
| + | 1.241826 | 0.273025 | Q9Y5K5 | Q9Y5K5 | UCHL5 | Ubiquitin c | Color2 [Da | 1 | 4 |
| + | 2.097508 | 1.241335 | Q9Y5K6 | Q9Y5K6 | CD2AP | CD2-assoc | Color2 [Da | 1 | 4 |
| + | 3.164567 | 2.608806 | Q9Y5K8 | Q9Y5K8 | ATP6V1D | V-type pro | Color2 [Da | 1 | 4 |
| + | 2.938446 | -1.58366 | Q9Y5L0 | Q9Y5L0 | TNPO3 | Transporti | Color2 [Da | 1 | 4 |
| + | 1.162457 | 1.367018 | Q9Y5L4 | Q9Y5L4 | TIMM13 | Mitochonc | Color2 [Da | 1 | 4 |
| + | 5.126107 | -1.12693 | Q9Y5M8 | Q9Y5M8 | SRPRB | Signal recc | Color2 [Da | 1 | 4 |
| + | 3.575453 | 1.422587 | Q9Y5N5 | Q9Y5N5 | N6AMT1 | Methyltrar | Color2 [Da | 1 | 4 |
| + | 2.279537 | -0.9364 | Q9Y5P4 | Q9Y5P4 | CERT1 | Ceramide t | Color2 [Da | 1 | 4 |
| + | 2.363294 | -1.30975 | Q9Y5P6 | Q9Y5P6 | GMPPB | Mannose-1 | Color2 [Da | 1 | 4 |
| + | 3.73204 | -1.72702 | Q9Y5Q8 | Q9Y5Q8 | GTF3C5 | General tra | Color2 [Da | 1 | 4 |
| + | 1.907681 | -0.60638 | Q9Y5Q9 | Q9Y5Q9 | GTF3C3 | General tra | Color2 [Da | 1 | 4 |
| | 0 | NaN | Q9Y5R2 | Q9Y5R2 | MMP24 | Matrix me | Color2 [Da | 1 | 4 |
| | 0.789681 | 0.426918 | Q9Y5R8 | Q9Y5R8 | TRAPPC1 | Trafficking | Color2 [Da | 1 | 4 |
| + | 2.026889 | 0.412827 | Q9Y5S2 | Q9Y5S2 | CDC42BPB | Serine/thre | Color2 [Da | 1 | 4 |
| | 0.057008 | -0.02553 | Q9Y5S9 | Q9Y5S9 | RBM8A | RNA-bindi | Color2 [Da | 1 | 4 |
| | 0 | NaN | Q9Y5T5 | Q9Y5T5 | USP16 | Ubiquitin c | Color2 [Da | 1 | 4 |
| | 0.534316 | 1.072377 | Q9Y5U2 | Q9Y5U2 | TSSC4 | Protein TS | Color2 [Da | 1 | 4 |
| + | 2.033453 | 1.154446 | Q9Y5U8 | Q9Y5U8 | MPC1 | Mitochonc | Color2 [Da | 1 | 4 |
| + | 0.943588 | 1.050446 | Q9Y5U9 | Q9Y5U9 | IER3IP1 | Immediate | Color2 [Da | 1 | 4 |
| | 0.127977 | 0.055814 | Q9Y5V3 | Q9Y5V3 | MAGED1 | Melanoma | Color2 [Da | 1 | 4 |
| | 0.969345 | -0.24276 | Q9Y5X1 | Q9Y5X1 | SNX9 | Sorting ne: | Color2 [Da | 1 | 4 |
| + | 1.685222 | -0.6906 | Q9Y5X2 | Q9Y5X2 | SNX8 | Sorting ne: | Color2 [Da | 1 | 4 |
| + | 3.000918 | 0.652246 | Q9Y5X3 | Q9Y5X3 | SNX5 | Sorting ne: | Color2 [Da | 1 | 4 |
| + | 1.344745 | 1.008471 | Q9Y5X9 | Q9Y5X9 | LIPG | Endothelia | Color2 [Da | 1 | 4 |
| + | 2.803352 | -0.96713 | Q9Y5Y2 | Q9Y5Y2 | NUBP2 | Cytosolic F | Color2 [Da | 1 | 4 |
| + | 1.289882 | -1.12337 | Q9Y5Y5 | Q9Y5Y5 | PEX16 | Peroxisom | Color2 [Da | 1 | 4 |
| + | 4.292891 | 2.66144 | Q9Y5Z4 | Q9Y5Z4 | HEBP2 | Heme-binc | Color2 [Da | 1 | 4 |
| | 0 | -0.04778 | Q9Y5Z6 | Q9Y5Z6 | B3GALT1 | Beta-1,3-g | Color2 [Da | 1 | 4 |
| + | 1.239987 | 1.091527 | Q9Y605 | Q9Y605 | MRFAP1 | MORF4 far | Color2 [Da | 1 | 4 |
| + | 1.523395 | -0.87451 | Q9Y606 | Q9Y606 | PUS1 | Pseudouric | Color2 [Da | 1 | 4 |
| + | 3.025334 | 1.831211 | Q9Y608 | Q9Y608 | LRRFIP2 | Leucine-ric | Color2 [Da | 1 | 4 |
| | 0.930867 | -0.31655 | Q9Y617 | Q9Y617 | PSAT1 | Phosphose | Color2 [Da | 1 | 4 |
| | 0 | NaN | Q9Y623 | Q9Y623 | MYH4 | Myosin-4 | Color2 [Da | 1 | 4 |
| + | 1.310668 | -1.19456 | Q9Y624 | Q9Y624 | F11R | Junctional | Color2 [Da | 1 | 4 |
| + | 2.777453 | -1.62215 | Q9Y625 | Q9Y625 | GPC6 | Glypican-6 | Color2 [Da | 1 | 4 |
| | 0.287133 | -0.10194 | Q9Y639 | Q9Y639 | NPTN | Neuroplas | Color2 [Da | 1 | 4 |
| + | 2.020802 | 1.674367 | Q9Y646 | Q9Y646 | CPQ | Carboxype | Color2 [Da | 1 | 4 |
| + | 2.276619 | -0.72969 | Q9Y653 | Q9Y653 | ADGRG1 | Adhesion ( | Color2 [Da | 1 | 4 |
| + | 2.555579 | 1.209493 | Q9Y657 | Q9Y657 | SPIN1 | Spindlin-1 | Color2 [Da | 1 | 4 |
| + | 2.940703 | 2.027451 | Q9Y662 | Q9Y662 | HS3ST3B1 | Heparan su | Color2 [Da | 1 | 4 |
| | 0.256108 | 0.107259 | Q9Y666 | Q9Y666 | SLC12A7 | Solute carr | Color2 [Da | 1 | 4 |

| | | | | | | | | | |
|---|---|---|---|---|---|---|---|---|---|
| | 0 | -1.29526 | Q9Y673 | Q9Y673 | ALG5 | Dolichyl-pl | Color2 [Da | 1 | 4 |
| + | 3.820821 | 0.572235 | Q9Y676 | Q9Y676 | MRPS18B | 28S riboso | Color2 [Da | 1 | 4 |
| + | 4.398063 | -1.92401 | Q9Y678 | Q9Y678 | COPG1 | Coatomer | Color2 [Da | 1 | 4 |
| | 0 | NaN | Q9Y679 | Q9Y679 | AUP1 | Lipid dropl | Color2 [Da | 1 | 4 |
| | 0.448016 | -0.05632 | Q9Y680 | Q9Y680 | FKBP7 | Peptidyl-p | Color2 [Da | 1 | 4 |
| | 0 | NaN | Q9Y689 | Q9Y689 | ARL5A | ADP-ribosy | Color2 [Da | 1 | 4 |
| + | 2.366172 | -0.88524 | Q9Y696 | Q9Y696 | CLIC4 | Chloride ir | Color2 [Da | 1 | 4 |
| | 0.533514 | -0.22468 | Q9Y697 | Q9Y697 | NFS1 | Cysteine d | Color2 [Da | 1 | 4 |
| | 0 | -1.05627 | Q9Y6A1 | Q9Y6A1 | POMT1 | Protein O- | Color2 [Da | 1 | 4 |
| + | 2.442716 | -0.87268 | Q9Y6A4 | Q9Y6A4 | CFAP20 | Cilia- and f | Color2 [Da | 1 | 4 |
| | 0.2923 | -0.37343 | Q9Y6A9 | Q9Y6A9 | SPCS1 | Signal pep | Color2 [Da | 1 | 4 |
| | 0.0781 | 0.03852 | Q9Y6B6 | Q9Y6B6 | SAR1B | GTP-bindir | Color2 [Da | 1 | 4 |
| + | 2.436676 | -0.94969 | Q9Y6C9 | Q9Y6C9 | MTCH2 | Mitochonc | Color2 [Da | 1 | 4 |
| + | 1.172008 | -0.81667 | Q9Y6D5 | Q9Y6D5 | ARFGEF2 | Brefeldin A | Color2 [Da | 1 | 4 |
| | 0.476756 | 0.09788 | Q9Y6D6 | Q9Y6D6 | ARFGEF1 | Brefeldin A | Color2 [Da | 1 | 4 |
| | 0.543665 | 0.086405 | Q9Y6D9 | Q9Y6D9 | MAD1L1 | Mitotic spi | Color2 [Da | 1 | 4 |
| + | 1.853746 | -0.86095 | Q9Y6E0 | Q9Y6E0 | STK24 | Serine/thr | Color2 [Da | 1 | 4 |
| + | 2.106094 | -0.99312 | Q9Y6E2 | Q9Y6E2 | BZW2 | eIF5-mimi | Color2 [Da | 1 | 4 |
| + | 1.734593 | 1.375411 | Q9Y6G5 | Q9Y6G5 | COMMD1( | COMM do | Color2 [Da | 1 | 4 |
| + | 2.304541 | 0.328 | Q9Y6G9 | Q9Y6G9 | DYNC1LI1 | Cytoplasm | Color2 [Da | 1 | 4 |
| + | 2.980052 | 2.397399 | Q9Y6I3 | Q9Y6I3 | EPN1 | Epsin-1 | Color2 [Da | 1 | 4 |
| | 0 | NaN | Q9Y6J9 | Q9Y6J9 | TAF6L | TAF6-like F | Color2 [Da | 1 | 4 |
| + | 1.573994 | -1.2185 | Q9Y6K0 | Q9Y6K0 | CEPT1 | Choline/et | Color2 [Da | 1 | 4 |
| + | 1.574109 | -0.5176 | Q9Y6K1 | Q9Y6K1 | DNMT3A | DNA (cyto: | Color2 [Da | 1 | 4 |
| + | 1.76882 | 2.473777 | Q9Y6K8 | Q9Y6K8 | AK5 | Adenylate | Color2 [Da | 1 | 4 |
| + | 2.837234 | -0.27617 | Q9Y6K9 | Q9Y6K9 | IKBKG | NF-kappa- | Color2 [Da | 1 | 4 |
| + | 3.037982 | 0.413759 | Q9Y6M1 | Q9Y6M1 | IGF2BP2 | Insulin-like | Color2 [Da | 1 | 4 |
| | 0 | 0.842998 | Q9Y6M4 | Q9Y6M4 | CSNK1G3 | Casein kin | Color2 [Da | 1 | 4 |
| + | 1.332601 | -0.38027 | Q9Y6M7 | Q9Y6M7 | SLC4A7 | Sodium bi | Color2 [Da | 1 | 4 |
| + | 0.910578 | 0.93592 | Q9Y6M9 | Q9Y6M9 | NDUFB9 | NADH deh | Color2 [Da | 1 | 4 |
| | 0.348586 | 0.318204 | Q9Y6N5 | Q9Y6N5 | SQOR | Sulfide:qui | Color2 [Da | 1 | 4 |
| + | 1.941641 | 0.755967 | Q9Y6N7 | Q9Y6N7 | ROBO1 | Roundabo | Color2 [Da | 1 | 4 |
| + | 1.832687 | 1.024721 | Q9Y6N8 | Q9Y6N8 | CDH10 | Cadherin-1 | Color2 [Da | 1 | 4 |
| | 0 | -0.86091 | Q9Y6Q2 | Q9Y6Q2 | STON1 | Stonin-1 | Color2 [Da | 1 | 4 |
| + | 1.976736 | 2.010056 | Q9Y6R0 | Q9Y6R0 | NUMBL | Numb-like | Color2 [Da | 1 | 4 |
| + | 3.450893 | 2.645362 | Q9Y6R1 | Q9Y6R1 | SLC4A4 | Electrogen | Color2 [Da | 1 | 4 |
| + | 2.469245 | -0.86501 | Q9Y6R4 | Q9Y6R4 | MAP3K4 | Mitogen-a | Color2 [Da | 1 | 4 |
| + | 4.136158 | 1.444165 | Q9Y6V0 | Q9Y6V0 | PCLO | Protein pic | Color2 [Da | 1 | 4 |
| + | 1.609905 | -0.79924 | Q9Y6V7 | Q9Y6V7 | DDX49 | Probable A | Color2 [Da | 1 | 4 |
| + | 2.167099 | 2.112676 | Q9Y6W5 | Q9Y6W5 | WASF2 | Wiskott-Al | Color2 [Da | 1 | 4 |
| | 0 | NaN | Q9Y6X3 | Q9Y6X3 | MAU2 | MAU2 chr | Color2 [Da | 1 | 4 |
| + | 3.168473 | 2.014215 | Q9Y6X4 | Q9Y6X4 | FAM169A | Soluble lar | Color2 [Da | 1 | 4 |
| + | 2.402114 | 1.115678 | Q9Y6X5 | Q9Y6X5 | ENPP4 | Bis(5'-ader | Color2 [Da | 1 | 4 |
| | 0.526944 | 0.447338 | Q9Y6X9 | Q9Y6X9 | MORC2 | ATPase M( | Color2 [Da | 1 | 4 |
| | 0.796833 | -0.24017 | Q9Y6Y8 | Q9Y6Y8 | SEC23IP | SEC23-inte | Color2 [Da | 1 | 4 |

| Show labe | [1] 1_1_GI | [2] 1_2_GI | [3] 1_3_GI | [5] 2_2_GI | [6] 2_3_GI | [7] 3_1_SP | [8] 3_2_SP | [9] 3_3_SP | [10] 4_1_S |
|---|---|---|---|---|---|---|---|---|---|
| | 9.823007 | 9.842199 | 9.866184 | 9.373689 | 9.447624 | 9.829147 | 9.70195 | 9.849144 | 9.427401 |
| | 7.292034 | 7.515527 | 7.839242 | 9.423004 | 9.349046 | 7.000334 | 7.986549 | 7.551843 | NaN |
| | NaN | NaN | NaN | 7.235868 | 8.51478 | NaN | NaN | 3.873882 | NaN |
| | 5.795576 | 6.154883 | 5.887751 | 4.20122 | 4.882681 | 5.316309 | 4.781431 | 5.260168 | 4.405124 |
| | NaN | 6.919875 | 4.973923 | 7.642953 | 7.619265 | NaN | NaN | 7.14614 | 6.510011 |
| | NaN | 10.20469 | 9.967971 | 7.037113 | 7.930401 | 7.899564 | 8.396096 | 8.60411 | 7.177393 |
| | 7.395829 | 7.147267 | 6.795868 | 6.145801 | 5.562654 | 7.955399 | 5.081549 | 7.200374 | 7.429399 |
| | NaN | 5.240828 | 4.857683 | NaN | NaN | 4.27954 | NaN | NaN | 5.599512 |
| | 7.578902 | 6.675425 | 6.032014 | NaN | 6.365462 | 6.909999 | 6.566643 | 6.715842 | 6.715163 |
| | 7.89498 | 7.756409 | 7.639499 | 6.840949 | 6.96644 | 8.321833 | 8.110454 | 8.314198 | 8.095342 |
| | 6.805365 | 7.018722 | 6.793346 | 6.757805 | 6.442333 | 7.270071 | 7.447229 | 7.18056 | 7.050939 |
| | 4.993548 | 5.060865 | 4.173155 | 4.364157 | 3.946103 | 6.012371 | 5.799213 | 6.31448 | 5.223908 |
| | 10.02197 | 9.861566 | 9.691262 | 10.15128 | 10.1692 | 9.655435 | 9.746737 | 9.713111 | 8.665969 |
| | NaN | 6.758624 | 6.518894 | 4.72125 | 6.020655 | NaN | NaN | NaN | NaN |
| | 7.925488 | 7.950284 | 8.081183 | 7.361043 | 7.579532 | 7.877526 | 7.797285 | 7.696578 | 7.186004 |
| | 7.525835 | 7.514248 | 7.672933 | 6.965853 | 6.541135 | 8.064662 | 7.750246 | 7.45545 | 7.350218 |
| | 4.665512 | 4.859102 | 4.898694 | NaN | NaN | 5.500457 | 4.713401 | 4.57998 | NaN |
| | NaN | NaN | NaN | 7.373893 | 7.038174 | NaN | 6.968749 | 7.136708 | 6.678768 |
| | 6.739266 | 6.662115 | 6.78004 | 6.616837 | 6.660148 | 6.877621 | 6.815917 | 6.692082 | 7.390887 |
| | 6.947547 | 6.876615 | 7.521222 | 5.847274 | NaN | 6.678565 | 6.760338 | 7.298827 | 5.785594 |
| | 6.25023 | 6.69812 | 5.927698 | 7.303876 | 8.240003 | NaN | NaN | NaN | 7.461032 |
| | 7.070277 | 5.551181 | 7.316614 | 6.692136 | 5.946362 | 4.308611 | NaN | NaN | 5.68958 |
| | 7.843351 | 7.494651 | 7.376465 | 6.617204 | 6.722126 | 7.8539 | 7.550991 | 7.720675 | 7.368883 |
| | 6.800285 | 7.262287 | 7.051804 | 6.858072 | 6.697985 | 7.35434 | 7.316253 | 7.225931 | 7.050865 |
| | 6.258414 | 6.093988 | 5.975384 | 5.204422 | 5.255048 | 6.020765 | 5.721019 | 5.89363 | 6.205601 |
| | 6.278679 | 6.618767 | 5.740093 | 6.209013 | 6.599337 | 7.039158 | 7.106237 | 7.0331 | 6.764067 |
| | 6.577173 | 5.923526 | NaN | 7.468723 | 7.232721 | NaN | NaN | NaN | 7.071717 |
| | 5.629169 | 5.012374 | 4.804022 | 5.550603 | 6.485241 | NaN | NaN | NaN | 5.389163 |
| | 6.277208 | 6.722968 | 6.314121 | 6.152754 | 5.774047 | 6.699971 | 6.879263 | 6.789639 | 6.498364 |
| | NaN | NaN | NaN | 4.39252 | NaN | NaN | 5.020737 | NaN | 5.363739 |
| | 9.17468 | 9.055558 | 8.976552 | 9.565311 | 9.021473 | 8.737445 | 8.715682 | 8.638462 | 8.74228 |
| | 6.493459 | 6.826521 | 6.673335 | 6.476251 | 6.147028 | 6.324585 | 6.734576 | 6.788777 | 5.514367 |
| | NaN | 5.614396 | NaN | NaN | NaN | 7.591242 | 7.425884 | 7.543983 | 7.125105 |
| | 8.664354 | 8.592059 | 8.661938 | 7.436653 | 7.605043 | 8.812431 | 8.838002 | 8.918316 | 8.477981 |
| | 9.969637 | 9.825639 | 9.817697 | 11.90447 | 11.96808 | 10.05403 | 9.594458 | 9.480402 | 9.131266 |
| | 13.63303 | 13.62212 | 13.6824 | 13.49738 | 13.61811 | 10.71569 | 10.26586 | 10.75024 | 9.613805 |
| | 7.261961 | 7.490254 | 7.262763 | 7.652247 | 7.739556 | 7.019371 | 6.537949 | 7.203604 | 6.44676 |
| | 6.261034 | 5.090237 | 6.021192 | NaN | 7.612218 | NaN | NaN | NaN | 6.945703 |
| | 7.997441 | 7.033727 | 6.875773 | 6.465006 | 6.142851 | 7.317871 | 7.063457 | 7.375424 | 6.731219 |
| | 8.979691 | 9.092274 | 9.047384 | 8.057817 | 8.291269 | 9.233663 | 9.103438 | 9.261992 | 8.095273 |
| | 6.926777 | 5.159865 | 7.120018 | 7.042208 | 6.007817 | 7.358155 | 7.402829 | 7.23479 | 6.373729 |
| | 5.494322 | 5.510861 | 4.795918 | 7.06566 | 7.592896 | 4.526337 | 5.654936 | 4.945754 | 4.502582 |
| | 5.260765 | 5.016029 | 6.394882 | 6.184634 | 6.750249 | 5.96653 | 5.48804 | 5.595433 | 4.584052 |
| | 9.023524 | 8.948496 | 8.937482 | 7.461709 | 7.542546 | 8.36347 | 8.646164 | 8.983986 | 6.748116 |
| | 11.36747 | 11.53716 | 10.8309 | 11.57511 | 11.65487 | 12.58042 | 12.65526 | 12.55835 | 12.22881 |
| | 8.567659 | 8.644551 | 8.755657 | 9.406602 | 9.524373 | 9.080587 | 9.153568 | 9.086631 | 7.992033 |

-Bruker Confidential-

| | | | | | | | | |
|---|---|---|---|---|---|---|---|---|
| 8.010251 | 7.907418 | 8.001984 | 6.950629 | 7.041677 | 8.063218 | 8.117436 | 8.114638 | 8.025763 |
| 6.932484 | 7.018393 | 7.011103 | 6.883879 | 6.848582 | 7.558143 | 8.201745 | 7.986655 | 6.793255 |
| NaN | NaN | 7.260912 | 8.329029 | 10.97257 | 7.797908 | 7.811027 | 7.832731 | 8.651691 |
| 8.677205 | 7.97545 | 8.179448 | 9.725156 | 9.357608 | NaN | 8.392017 | 8.123038 | 8.191985 |
| 6.21323 | NaN | 6.074907 | 6.683095 | 6.883571 | 5.968605 | 5.860924 | 6.053552 | 6.389235 |
| 6.799144 | 7.754891 | 6.597128 | 7.052069 | 6.699057 | 6.260952 | 6.365346 | 6.177503 | 6.765982 |
| 6.694411 | 5.702507 | 5.873684 | NaN | NaN | 5.781243 | NaN | NaN | 6.181282 |
| NaN | 4.202971 | NaN | 5.414847 | 5.173564 | NaN | NaN | NaN | NaN |
| 8.825735 | 7.118579 | 7.419169 | 8.146933 | 8.184342 | 6.810842 | 7.195264 | 7.219553 | 7.458587 |
| 8.562751 | 8.589179 | 8.827244 | 8.838326 | 9.183201 | 8.661427 | 8.791392 | 8.496695 | 8.458107 |
| 6.85165 | 7.053723 | 6.896684 | 6.366575 | 6.666975 | 6.937371 | 6.02941 | 6.72962 | 6.218894 |
| 6.367876 | 6.36533 | NaN | 5.707093 | 6.08271 | 7.069067 | 6.918996 | 6.635109 | 6.059544 |
| 4.207859 | 5.39341 | NaN | 5.644884 | 5.477682 | 5.234862 | 4.835408 | 5.303421 | 6.012621 |
| 5.873991 | 5.301023 | 6.030182 | NaN | NaN | NaN | NaN | NaN | NaN |
| NaN | NaN | NaN | NaN | NaN | NaN | NaN | NaN | 6.607564 |
| 8.012549 | 7.966332 | 8.205798 | 8.172234 | 7.964019 | 7.961779 | 7.979334 | 7.957519 | 7.649945 |
| 8.689948 | 8.298271 | 8.459286 | 7.957554 | 8.075021 | 8.160925 | 8.247081 | 8.143858 | 7.381877 |
| 10.52317 | 10.2544 | 10.19812 | 10.63426 | 11.25981 | 9.034955 | 8.995854 | 9.207646 | 10.19748 |
| NaN | NaN | 4.667518 | 3.904967 | 3.47721 | 4.89691 | 4.662136 | 4.34275 | NaN |
| NaN | NaN | NaN | 5.724779 | 4.774136 | NaN | NaN | NaN | 6.18855 |
| 7.489555 | 6.423759 | 6.01914 | NaN | NaN | 8.63585 | 8.900074 | 8.776093 | 7.507113 |
| 5.90202 | 6.233912 | 6.14653 | 6.08184 | 6.565244 | 6.162519 | 6.101724 | 6.382645 | 4.731252 |
| 5.956436 | 5.747324 | 6.005474 | NaN | NaN | 6.512954 | 5.954941 | 6.52676 | 6.474083 |
| 6.848135 | 7.108467 | 6.746219 | 7.743521 | 8.466404 | 6.888688 | 6.900675 | 6.976044 | 6.638296 |
| 6.544253 | 6.451301 | 6.147854 | NaN | 5.518218 | 6.998581 | 6.889104 | 7.366612 | 6.41766 |
| NaN | 5.659351 | NaN | 6.544437 | 6.534315 | NaN | NaN | 3.623811 | 4.79085 |
| 8.377732 | 7.766127 | 7.894735 | 9.717791 | 9.782943 | 6.974893 | 6.925814 | 6.953243 | 9.787487 |
| 8.152866 | 8.180993 | 8.111825 | 8.260911 | 6.830616 | 6.620373 | 7.018542 | 6.700588 | 7.262409 |
| 11.37175 | 11.28662 | 11.52354 | 12.80502 | 12.92221 | 10.16265 | 10.03295 | 10.02921 | 10.64422 |
| 5.277977 | 5.61912 | 5.506207 | 7.568818 | 7.87383 | 5.534504 | 5.783248 | 6.030061 | 6.088951 |
| 17.50837 | 15.35707 | 15.7825 | 13.97256 | 11.15256 | 13.16327 | 11.93481 | 11.65496 | 11.82487 |
| 6.938488 | 7.001394 | 7.261048 | NaN | NaN | 7.132599 | 7.184845 | 7.201662 | 6.045266 |
| 7.086949 | 6.29181 | 6.952112 | 8.095405 | 7.481961 | 6.72343 | 6.991404 | 6.885475 | 6.213573 |
| NaN | 7.962151 | 7.833093 | 8.448789 | 8.767513 | 6.73388 | 6.835241 | 6.917683 | 7.13128 |
| 9.653894 | 9.413549 | 9.203018 | 11.40725 | 11.36787 | 8.38732 | 8.068127 | 7.952438 | 9.069731 |
| 8.484215 | 8.644689 | 8.51116 | 7.142232 | 7.086087 | 8.356586 | 8.354299 | 8.505592 | 8.282332 |
| 9.969456 | 10.09313 | 10.06062 | 8.829366 | 9.12415 | 9.792113 | 9.605468 | 9.866359 | 9.388975 |
| 7.939145 | NaN | 6.986869 | NaN | NaN | NaN | NaN | NaN | NaN |
| 7.869971 | 7.629544 | 8.218521 | 6.326271 | 6.249165 | 6.765082 | 6.839258 | 6.922739 | 7.700766 |
| 6.608064 | 5.748431 | 5.489034 | 7.999129 | 8.200044 | NaN | NaN | NaN | 7.840897 |
| 11.47331 | 11.38721 | 11.35708 | 11.27122 | 11.35028 | 11.46503 | 11.31912 | 11.18833 | 10.65733 |
| 6.873991 | 6.688532 | 6.931874 | 6.514575 | 7.061621 | 6.755435 | 6.681977 | 7.30941 | 6.907182 |
| NaN | 4.211459 | NaN | 6.516693 | 6.956218 | 5.700568 | 6.205891 | 5.70779 | 5.313595 |
| 5.949741 | 5.921296 | 5.856667 | 6.607641 | 6.839076 | 5.662014 | 5.271626 | 5.834508 | 5.964806 |
| 8.569898 | 8.287937 | 8.419579 | 8.30296 | 8.218861 | 8.360447 | 8.104087 | 8.545786 | 8.423558 |
| 8.729314 | 8.567287 | 8.661483 | 7.58649 | 7.090237 | 8.399949 | 8.293367 | 8.062801 | 7.774592 |
| 7.483808 | 7.476955 | 7.394673 | 6.955204 | 6.560516 | 7.640745 | 7.415226 | 7.227539 | 7.296144 |

-Bruker Confidential-

| | | | | | | | | |
|---|---|---|---|---|---|---|---|---|
| 7.379285 | 7.410219 | 7.514708 | 7.666395 | 7.789642 | 8.028771 | 7.913021 | 7.983317 | 7.89985 |
| 5.959816 | 6.255335 | 5.961651 | 7.544233 | 7.747955 | 5.328196 | 4.720116 | 5.269217 | 5.324115 |
| 5.227034 | NaN | 5.669678 | 6.791187 | 6.313861 | 6.206841 | 6.06367 | 6.037713 | 5.212415 |
| 6.606956 | 6.990032 | 6.727635 | 6.922576 | 7.01769 | 6.668546 | 6.949239 | 6.906384 | 7.060435 |
| 7.563281 | 7.858353 | 7.527444 | 7.557324 | 7.613606 | 7.413209 | 7.293143 | 7.213837 | 7.4531 |
| 9.486169 | 9.495635 | 9.519166 | 10.56107 | 10.49313 | 9.823492 | 9.417676 | 9.449375 | 9.49806 |
| 10.86975 | 10.91936 | 10.88222 | 10.57912 | 10.71677 | 10.75847 | 10.79314 | 10.97852 | 11.0399 |
| 8.563399 | 8.38489 | 8.501548 | 7.726254 | 7.660928 | 7.968362 | 8.121475 | 7.987371 | 8.412545 |
| 9.246539 | 9.070165 | 9.188532 | 9.891129 | 9.102171 | 8.310419 | 8.271524 | 7.982848 | 8.331422 |
| 8.339551 | 8.165778 | 8.244513 | 10.32586 | 10.64188 | 7.611474 | 7.827715 | 7.76085 | 7.176339 |
| 7.390563 | 6.757224 | 7.368312 | 6.522317 | 6.28981 | 7.15641 | 7.429879 | 7.243919 | 7.469099 |
| 5.526079 | 5.491224 | 6.085062 | 6.024374 | 5.656039 | 5.808649 | 5.710159 | 5.531518 | 5.645311 |
| 5.170199 | NaN | 5.138569 | NaN | NaN | 6.420073 | 6.05654 | 6.440559 | 5.508306 |
| 10.77739 | 10.72055 | 10.79003 | 11.27344 | 11.36539 | 11.44468 | 11.41037 | 11.30045 | 10.34682 |
| 7.559479 | 7.626683 | 7.669301 | 7.707627 | 7.815838 | 7.434497 | 7.550643 | 7.57713 | 7.538108 |
| 7.599605 | 7.373312 | 7.205006 | 7.807015 | 7.821041 | 7.165342 | 7.054753 | 7.296764 | 7.006107 |
| NaN | NaN | NaN | 5.276187 | 5.203531 | NaN | NaN | NaN | 4.57914 |
| 8.376801 | 8.318149 | 8.42284 | 7.909997 | 7.814031 | 8.707686 | 8.747019 | 8.728277 | 8.335732 |
| 9.537169 | 9.635807 | 9.704168 | 9.369826 | 9.422282 | 9.324183 | 9.333862 | 9.593881 | 10.26737 |
| 9.085997 | 8.695965 | 8.817855 | 9.195561 | 8.741191 | 8.631311 | 8.515821 | 8.284735 | 9.115048 |
| 6.889427 | 6.679173 | 6.922283 | 7.862259 | 7.77262 | 7.272919 | 6.802222 | 7.758288 | 7.213449 |
| 6.323798 | 5.728083 | 6.583935 | 6.916208 | 7.400255 | 6.684144 | 6.404913 | 6.443888 | 5.91864 |
| 8.162024 | 8.213924 | 8.388406 | 8.402853 | 8.475236 | 8.155373 | 8.092924 | 8.234401 | 7.735058 |
| 5.159344 | 5.276673 | 4.207485 | NaN | NaN | 6.724457 | 6.556578 | 6.865273 | 6.492203 |
| 8.311706 | 8.125188 | 8.216189 | 7.398906 | 7.795198 | 8.415085 | 8.406365 | 8.252629 | 7.837451 |
| 6.389082 | 6.369711 | 6.321874 | 6.096136 | 6.302009 | 6.371142 | 6.458452 | 6.221274 | 5.853176 |
| 6.570921 | 6.275511 | 4.895428 | 5.121698 | NaN | 6.00334 | 6.423296 | 6.536577 | 5.732849 |
| 9.420125 | 9.265898 | 9.406872 | 7.902775 | 8.132255 | 9.467664 | 9.412749 | 9.355337 | 9.858578 |
| 5.668838 | 5.390361 | 5.410691 | NaN | NaN | 5.557128 | 5.583276 | 5.411784 | 5.382682 |
| 8.434839 | 8.531073 | 8.633678 | 9.359558 | 8.96033 | 7.989891 | 8.277188 | 8.090069 | 7.988385 |
| 5.712508 | 5.662677 | 5.789917 | 5.344211 | 6.162964 | 5.630491 | 6.01373 | 5.873943 | 6.642598 |
| 9.259818 | 9.298967 | 9.218366 | 8.616506 | 8.746992 | 8.559089 | 8.394654 | 8.376698 | 9.756635 |
| 8.516224 | 8.583748 | 8.637874 | 8.692986 | 8.595933 | 8.641953 | 8.614034 | 8.667923 | 7.894152 |
| 6.341339 | 6.201486 | 6.262489 | 5.433274 | 5.838533 | 5.80013 | 6.146691 | 6.216769 | 6.001007 |
| 6.198528 | 6.065926 | 6.489398 | 5.941576 | 5.712231 | 6.053267 | 6.27028 | 5.817582 | 6.193896 |
| 7.044186 | 7.173405 | 6.802535 | 6.601363 | 6.499246 | 7.67241 | 7.02462 | 7.112446 | 4.785614 |
| NaN | 6.625921 | 6.3393 | 4.81162 | 6.240499 | 8.071463 | 7.61551 | 7.45591 | 5.763655 |
| 5.258777 | 5.77128 | 4.949327 | 4.93618 | 5.19681 | 6.565677 | 6.504408 | 6.521452 | 6.623023 |
| 5.915864 | 5.321489 | 5.667032 | 6.221728 | 6.068915 | 5.395461 | 5.552865 | 5.53642 | 5.193673 |
| 8.286669 | 8.352727 | 7.924613 | 7.980972 | 7.918112 | 8.190732 | 8.087553 | 7.962552 | 8.541668 |
| 8.28625 | 8.278906 | 8.232647 | 7.255625 | 7.312968 | 7.973286 | 7.794798 | 7.568376 | 7.13538 |
| 9.146931 | 9.137613 | 9.223418 | 8.275594 | 8.504978 | 8.802519 | 8.956961 | 8.918425 | 8.682159 |
| 6.596883 | NaN | NaN | NaN | NaN | 8.288854 | 8.06223 | 7.60769 | 6.039997 |
| 9.09062 | 9.180496 | 9.288125 | 8.973691 | 9.012356 | 7.835123 | 7.85108 | 7.89185 | 8.444577 |
| 7.180635 | 6.972694 | 6.859951 | 6.341671 | 7.295814 | 7.391293 | 7.374268 | 7.2283 | 7.228503 |
| 8.343713 | 8.139963 | 7.770426 | NaN | 9.418435 | 8.823762 | 8.087709 | 8.822239 | 8.213743 |
| 7.798172 | 6.872883 | 7.896896 | 7.121768 | 7.114532 | 7.058179 | 6.982502 | 7.205976 | 6.788161 |

| | | | | | | | | |
|---|---|---|---|---|---|---|---|---|
| 8.034543 | 8.10463 | 7.824392 | 7.586598 | 7.245071 | 8.168744 | 7.944419 | 7.875497 | 7.262195 |
| 5.992784 | NaN | NaN | 6.542253 | 6.215704 | 5.739361 | NaN | NaN | 6.376911 |
| 7.835305 | 7.970324 | 7.77116 | 9.172365 | 9.119362 | 7.770369 | 7.723691 | 7.587721 | 7.227774 |
| 8.907495 | 8.845728 | 8.491038 | 8.126867 | 8.002232 | 8.323414 | 8.486719 | 8.541751 | 8.832983 |
| 7.238028 | 7.108699 | 7.525344 | 5.869948 | 6.001765 | 7.17946 | 6.631503 | 6.616253 | 7.448358 |
| 6.176208 | 6.122057 | 5.821522 | 6.64477 | 6.106467 | 5.292463 | 5.783107 | 5.374956 | 6.789085 |
| 7.503856 | 6.759978 | 7.406808 | 5.684381 | 5.893588 | 6.491991 | 6.041943 | 6.81175 | 5.555991 |
| 7.667254 | 5.665283 | 5.418045 | NaN | 6.061298 | 6.762649 | 6.636353 | 6.49807 | NaN |
| 7.555285 | 7.35301 | 7.206245 | 7.398965 | 7.614056 | 7.82245 | 7.75526 | 7.917971 | 7.657812 |
| 7.709194 | 7.822443 | 8.115335 | 5.99094 | 7.592911 | 7.673177 | 7.370693 | 7.838927 | 7.804287 |
| 9.953008 | 9.814094 | 9.854719 | 9.540503 | 9.673275 | 8.984642 | 9.367161 | 9.302625 | 8.978493 |
| 6.983563 | 6.814919 | 6.654136 | 6.205885 | 6.580597 | 6.354413 | 6.524303 | 6.485242 | 6.446177 |
| 6.973511 | 7.274733 | 7.114946 | 7.326361 | 7.295586 | 6.938438 | 7.143306 | 6.842102 | 7.934279 |
| 9.374519 | 9.219034 | 9.31956 | 8.503845 | 8.292486 | 9.309174 | 9.523987 | 9.21817 | 8.767144 |
| 9.912495 | 9.908667 | 10.02025 | 8.919049 | 8.779425 | 9.381436 | 9.501868 | 9.641658 | 10.02338 |
| 7.208361 | 6.80904 | 4.733953 | 7.155211 | 7.534097 | NaN | NaN | NaN | 5.934621 |
| 7.348995 | 6.35995 | 6.014647 | 6.635753 | 6.146016 | 6.619591 | 6.885644 | 6.421781 | 7.538693 |
| 6.969501 | 7.145019 | NaN | NaN | 5.981043 | NaN | NaN | NaN | 7.698144 |
| NaN | 8.216259 | 8.168476 | 4.869874 | NaN | 7.794541 | 7.500353 | 8.066977 | 6.048429 |
| 7.411016 | 7.381481 | 7.150813 | 8.293736 | 8.047648 | 6.969497 | 7.057798 | 6.939994 | 7.121846 |
| 7.21757 | 7.234175 | 7.406525 | 5.604872 | 6.51234 | 7.339357 | 7.096579 | 6.93505 | 7.312861 |
| 6.667576 | 6.421307 | 6.58706 | NaN | 4.977629 | 6.442913 | 6.58189 | 6.26373 | NaN |
| 7.506749 | 7.435554 | 7.84091 | 7.490819 | 7.087723 | 7.947069 | 8.021775 | 8.511173 | 7.46365 |
| NaN | 4.407927 | NaN | NaN | NaN | 5.09011 | 5.118367 | 5.4498 | 4.661952 |
| 6.632249 | 6.148746 | 6.70296 | 6.185519 | 5.675939 | 8.061556 | 8.187508 | 8.115132 | 7.116583 |
| 7.548164 | 7.650513 | 7.69943 | 7.934547 | 8.047947 | 7.60174 | 7.536749 | 7.648937 | 7.516267 |
| 6.737699 | 5.957996 | 6.37172 | 6.828081 | 6.337079 | 7.665937 | 6.888969 | 7.034745 | 6.193774 |
| 7.072847 | 6.985689 | 6.7507 | 6.652064 | 7.097423 | 7.128949 | 6.772051 | 7.164947 | 7.264636 |
| 8.295422 | 8.196795 | 8.24552 | 9.060619 | 9.205415 | 8.186969 | 8.072098 | 7.991683 | 8.020252 |
| 7.051587 | 7.113428 | 7.139918 | 5.322139 | 5.707115 | 7.182688 | 7.014194 | 7.193002 | 5.246111 |
| 7.950951 | 7.204171 | 8.247043 | 6.68879 | 7.080602 | 6.001964 | 6.210107 | NaN | 6.600701 |
| 9.341498 | 9.20669 | 9.148725 | 9.199413 | 9.394126 | 9.156308 | 9.293777 | 9.345256 | 8.340283 |
| NaN | NaN | NaN | NaN | NaN | 5.958919 | 6.788281 | 7.000985 | 5.413093 |
| NaN | 7.51602 | 7.719437 | NaN | NaN | NaN | 8.375281 | NaN | 8.05092 |
| 8.22766 | 8.239601 | 8.421567 | 8.279034 | 8.480129 | 7.910938 | 7.873774 | 8.239549 | 8.381933 |
| NaN | NaN | NaN | 3.32584 | NaN | 4.179809 | 3.821977 | 4.725699 | 2.076269 |
| 6.764996 | 6.53428 | 6.901821 | 6.940991 | 7.353802 | 6.692998 | 6.627359 | 6.456675 | 6.259307 |
| NaN | NaN | 6.566195 | NaN | NaN | 8.174465 | 7.842688 | 7.338148 | NaN |
| 12.41012 | 12.36082 | 12.43346 | 6.843334 | 6.989594 | NaN | 6.852626 | NaN | NaN |
| 6.444064 | 6.564739 | 6.059298 | 8.12311 | 8.203469 | 6.075057 | 6.94547 | 6.604513 | 7.017142 |
| 9.342651 | 9.219714 | 9.217192 | 8.804618 | 8.824152 | 10.06897 | 9.902349 | 9.707373 | 9.085108 |
| 8.167505 | 8.337517 | 7.208303 | 8.614847 | 8.171077 | 6.917697 | 7.274129 | 6.01608 | 5.748147 |
| NaN | NaN | NaN | 6.871155 | 7.250287 | NaN | NaN | NaN | NaN |
| 7.613897 | 7.623681 | 7.786988 | 6.509606 | 6.553377 | 7.58035 | 7.528894 | 7.576862 | 6.797738 |
| 8.998981 | 8.729838 | 8.231093 | 9.040054 | 9.083308 | 7.747541 | 7.749509 | 7.853764 | 8.540001 |
| 7.782181 | 7.599136 | 7.95277 | 9.264468 | 9.672517 | 7.678667 | 7.76421 | 8.092756 | 8.507927 |
| 7.029366 | 7.565444 | 6.967235 | 6.993106 | 7.011226 | 7.411093 | 7.534314 | 7.400435 | 5.82903 |

| | | | | | | | | |
|---|---|---|---|---|---|---|---|---|
| 5.399304 | 5.735284 | 5.699439 | NaN | 5.405105 | 6.144016 | 6.276 | 6.447877 | 6.078277 |
| 7.869711 | 7.962312 | 8.064788 | 8.387386 | 8.567158 | 8.156756 | 8.43436 | 8.009337 | 8.245034 |
| 8.514984 | 8.522149 | 8.37557 | 8.222815 | 7.806571 | 8.76421 | 8.98764 | 9.035065 | 7.189672 |
| NaN | NaN | NaN | 6.916317 | 6.781617 | NaN | NaN | 5.920853 | NaN |
| 8.340714 | 8.431155 | 8.246425 | 7.618452 | 7.629399 | 8.50666 | 8.47375 | 8.465836 | 7.938279 |
| 7.311863 | 6.955418 | 7.180457 | 7.955454 | 8.001649 | 8.189291 | 8.206236 | 8.285731 | 7.289122 |
| 7.95437 | 8.108902 | 7.956478 | 9.577043 | 9.49389 | 8.182171 | 7.819777 | 8.101687 | 8.305758 |
| NaN | 5.533405 | 5.245786 | 6.31965 | 5.898788 | NaN | 5.718073 | 5.433774 | 5.080154 |
| 7.883265 | 7.664169 | 7.5976 | 7.97291 | 8.352647 | 6.908696 | 7.107107 | 7.174784 | 7.785364 |
| 7.758922 | 6.768805 | 7.759091 | 5.969576 | 5.971506 | 6.972862 | 7.057825 | 6.792193 | 7.980536 |
| 7.354753 | 7.891869 | 7.44458 | 6.420606 | 7.294372 | 7.720347 | 7.620297 | 7.932727 | 7.152855 |
| 7.302109 | 7.463638 | 7.125067 | 7.542039 | 7.214479 | 7.57743 | 7.532904 | 7.221569 | 5.799779 |
| 7.571421 | 8.260489 | 7.461453 | 6.882813 | 6.255042 | 7.330987 | 7.549513 | 7.744273 | 7.35281 |
| 7.620876 | 7.036418 | 7.515484 | NaN | 7.648119 | 8.146171 | 8.260207 | 8.268196 | 6.482903 |
| 6.921186 | 6.721192 | 6.631495 | 6.751689 | 7.072331 | 6.798546 | 6.882537 | 6.917777 | 6.401855 |
| 10.50379 | 9.960126 | 10.10713 | 9.217563 | 8.802732 | 8.890006 | 9.217067 | 9.24339 | 9.030845 |
| 8.767838 | 8.968955 | 8.442835 | 7.21677 | 7.105854 | 7.445677 | 8.326451 | 7.87473 | 7.241157 |
| 6.184405 | 5.362827 | 6.696926 | 6.837823 | 7.105137 | 6.399747 | 6.599969 | 6.523971 | 6.691295 |
| 6.243296 | 6.483153 | 6.731976 | NaN | NaN | 6.278723 | 6.822395 | 6.546346 | 5.657875 |
| 6.949254 | NaN | NaN | 5.332122 | 5.693157 | NaN | NaN | NaN | 6.377542 |
| 7.674847 | 7.175406 | 7.825961 | 6.602997 | 6.808528 | 8.97004 | 9.14613 | 9.312012 | 8.030364 |
| 8.208404 | 7.934445 | 8.276982 | 6.910472 | 7.014192 | 6.791109 | 6.380894 | 6.505907 | 7.534971 |
| 9.516863 | 9.32674 | 9.317683 | 10.71476 | 10.63176 | 7.881113 | 7.705492 | 7.768087 | 8.44153 |
| 9.359953 | 9.163564 | 9.213384 | 8.066376 | 8.241614 | 9.420274 | 9.738111 | 9.768308 | 9.59068 |
| 7.216872 | 7.197053 | 7.5375 | 8.38054 | 8.449566 | 7.240453 | 6.962917 | 7.443504 | 7.604616 |
| 7.206697 | 6.667007 | 6.728279 | 7.552151 | 7.740388 | 6.514221 | 7.016948 | 7.071102 | 6.820684 |
| 7.856023 | 5.794488 | 8.090809 | 8.099264 | 9.12361 | NaN | 7.763009 | 7.845056 | 7.625751 |
| 8.73712 | 8.722795 | 8.754395 | 9.441209 | 9.547424 | 8.833222 | 8.44137 | 8.716958 | 8.893029 |
| 9.152595 | 8.994143 | 9.142477 | 8.514574 | 8.606727 | 9.60719 | 9.393187 | 9.534097 | 8.266224 |
| 9.057518 | 8.996925 | 8.922934 | 8.796106 | 8.880982 | 8.730208 | 8.944553 | 8.863718 | 9.081288 |
| 7.177551 | 5.440651 | 7.272255 | 6.612699 | 7.082338 | 6.920056 | 6.593498 | 6.7231 | 6.912746 |
| NaN | NaN | NaN | NaN | NaN | NaN | 3.054703 | 4.850719 | NaN |
| 7.891052 | 7.624483 | 7.43632 | NaN | 6.716358 | 8.152417 | 8.248867 | 8.147429 | 7.7759 |
| 8.663876 | 8.798956 | 8.733185 | 7.100139 | 7.538347 | 8.797864 | 8.596981 | 8.414434 | 8.726024 |
| 6.239841 | 6.295227 | 5.597434 | 5.641328 | 5.511617 | 6.190075 | 5.856233 | 5.884204 | 6.555724 |
| 7.328856 | 9.805613 | 7.320525 | NaN | 7.389881 | 8.257794 | 8.091043 | 8.143262 | 7.118439 |
| 8.329807 | 8.096045 | 8.292102 | 8.033621 | 8.162063 | 7.656534 | 8.421244 | 8.078457 | 7.610922 |
| 7.281036 | 7.568193 | 7.804575 | 7.698345 | 7.613007 | 5.836688 | 6.030005 | 6.092426 | NaN |
| 9.200613 | 8.859773 | 9.12254 | 9.813247 | 9.914221 | 8.226091 | 8.340402 | 8.253297 | 7.905347 |
| 8.273193 | 8.036389 | 8.413006 | 7.529669 | 9.133471 | 8.147454 | 8.230987 | 8.574697 | 7.033544 |
| 6.866491 | 7.10913 | 6.961517 | 6.230871 | 6.074605 | 7.097038 | 6.861119 | 6.761009 | 7.025368 |
| 10.3626 | 10.40081 | 10.37125 | 9.959632 | 9.608205 | 9.560676 | 9.753903 | 9.800426 | 10.16689 |
| 8.425083 | 8.365286 | 8.522213 | 8.551837 | 8.772839 | 8.142792 | 7.677967 | 7.658724 | 8.702272 |
| 7.636127 | 7.614698 | 7.841971 | 7.993369 | 8.099585 | 7.347638 | 7.503769 | 7.724859 | 7.052378 |
| 9.058337 | 9.038474 | 9.0527 | 10.0598 | 9.921924 | 8.891442 | 8.653045 | 8.82382 | 8.384601 |
| 5.511541 | 6.326703 | 6.354002 | 5.518052 | 5.984805 | 7.086162 | 6.609993 | 6.932379 | 5.643501 |
| 7.587343 | 7.364794 | 7.751288 | 7.304391 | 7.413916 | 7.982869 | 7.840144 | 8.007194 | 7.26143 |

| | | | | | | | | |
|---|---|---|---|---|---|---|---|---|
| 7.8746 | 6.9234 | 7.441739 | 8.059919 | 7.734406 | 6.146794 | 6.355302 | 6.437747 | 4.888189 |
| 7.309242 | 7.707064 | 7.735958 | 4.572458 | 5.407954 | 7.710183 | 7.755177 | 7.739078 | 7.543272 |
| 7.551953 | 8.303984 | 6.900519 | 7.684911 | 7.903686 | 7.037609 | 7.128925 | NaN | 7.197842 |
| 8.692513 | 8.698567 | 8.675652 | 7.830874 | 7.773728 | 8.798388 | 8.924952 | 8.86001 | 8.137961 |
| 7.879049 | 7.699231 | 7.828703 | 7.845926 | 8.174681 | 8.614327 | 8.437625 | 8.955871 | 6.316021 |
| 10.99616 | 10.89642 | 10.91869 | 11.71497 | 11.67825 | 10.68189 | 10.22766 | 10.38075 | 10.50625 |
| 7.56038 | 7.633914 | 7.663084 | 10.11099 | 9.981444 | 7.149847 | NaN | 7.757823 | 5.732006 |
| 9.096588 | 8.839909 | 9.012525 | 9.697312 | 9.518477 | 8.414375 | 8.280995 | 8.356414 | 8.51554 |
| 7.010105 | 7.1262 | 7.541269 | 6.007202 | 6.944464 | 6.493742 | 6.320232 | 6.75319 | 6.341299 |
| 6.266608 | 6.663927 | 5.717876 | 7.800625 | 7.873045 | 5.984607 | 6.74674 | 6.15455 | 6.409842 |
| 7.381392 | 6.958883 | 7.041334 | 7.910942 | 7.500871 | 6.917896 | 7.027091 | 7.098477 | 7.233587 |
| 7.759214 | 7.562903 | 7.76247 | 7.449679 | 7.66434 | 7.620888 | 7.593283 | 7.071766 | 7.520034 |
| 10.1972 | 10.43574 | 10.35984 | 11.71474 | 11.79594 | 9.901261 | 9.705407 | 9.985128 | 10.12222 |
| 8.577711 | 8.586494 | 8.60079 | 6.868903 | 7.015492 | 8.399853 | 8.288423 | 8.40693 | 8.802032 |
| 7.463128 | 7.384332 | 7.392832 | 7.14541 | 7.044832 | 6.457186 | 6.679842 | 6.46448 | 6.709419 |
| 6.587853 | 7.059821 | 6.864481 | 7.724749 | 7.79096 | 7.626624 | 7.589014 | 7.721943 | 6.300706 |
| NaN | NaN | 5.687088 | 5.87026 | 5.563204 | 6.807918 | 6.225608 | 7.121566 | 5.734285 |
| 6.806295 | 3.040981 | 6.053313 | 7.413181 | 6.91038 | 6.715457 | 6.107307 | 6.559375 | 6.233975 |
| 8.149271 | 8.162625 | 8.30753 | 7.98618 | 7.985726 | 8.3633 | 8.240597 | 8.204237 | 8.107385 |
| 7.85006 | 7.963326 | 7.822341 | 7.472657 | 7.332647 | 7.482358 | 7.562304 | 7.426554 | 7.393335 |
| 7.444127 | 7.58484 | 7.655248 | 6.947155 | 6.883553 | 7.809629 | 7.965741 | 7.945055 | 7.406804 |
| 6.867083 | 6.364667 | 6.778115 | 7.184893 | 7.622827 | 6.993647 | 7.515126 | 7.006295 | 5.581748 |
| 6.726741 | 7.9956 | 6.769557 | 6.243541 | 6.793712 | 8.019714 | 8.503915 | 8.558208 | 6.130367 |
| 6.793203 | 6.078965 | 6.753073 | 6.068711 | 5.854338 | 6.756509 | 6.709062 | 6.770926 | 4.969858 |
| 7.515686 | 7.816873 | 7.616659 | 6.874951 | 7.019485 | 7.793429 | 7.430153 | 7.926682 | 7.482653 |
| 5.232726 | 5.107513 | 5.249248 | 5.379172 | 5.638704 | 4.765871 | 4.579013 | 5.263628 | 4.685042 |
| 8.062004 | 8.05815 | 7.930477 | 8.810324 | 8.687249 | 8.229837 | 8.252615 | 8.279637 | 7.530087 |
| 8.17638 | 8.144386 | 8.056767 | 8.469793 | 8.464402 | 8.05883 | 7.974164 | 8.090744 | 7.710991 |
| 8.354888 | 8.309819 | 8.411681 | 7.152679 | 7.09956 | 8.675161 | 8.499763 | 8.557354 | 8.340899 |
| 5.067922 | 5.690426 | 5.625706 | NaN | NaN | 6.28883 | 6.086582 | 6.062593 | 5.062081 |
| 7.647481 | 7.591373 | 7.218228 | 7.405332 | 7.462756 | 7.161125 | 7.183334 | 7.273662 | 7.804438 |
| 6.749434 | 7.2289 | 7.106638 | NaN | NaN | NaN | 7.647185 | 7.063324 | 6.262783 |
| 11.90696 | 11.97258 | 11.9782 | 11.50659 | 11.53268 | 12.07438 | 12.22508 | 12.18076 | 11.12491 |
| 7.915172 | 6.950599 | 6.131158 | 7.696218 | 7.252148 | NaN | 7.006588 | NaN | 6.947824 |
| 5.733036 | 6.10916 | 5.541825 | 4.985898 | 5.877917 | 5.985583 | 6.001754 | 5.786487 | 5.85545 |
| 6.677094 | 6.614975 | 6.866203 | 6.670603 | 6.876762 | 6.813307 | 6.978724 | 6.946853 | 6.36059 |
| 6.009166 | 5.712768 | 5.239833 | NaN | NaN | 5.886879 | 6.399037 | 6.289354 | 6.319713 |
| 5.136295 | 5.045827 | 4.380802 | NaN | NaN | 7.118073 | NaN | NaN | 5.777594 |
| 8.268814 | 8.124168 | 8.480798 | 10.28936 | 10.55244 | 8.248829 | 8.655166 | 8.151811 | 8.580347 |
| 8.314127 | 8.339147 | 8.305029 | 8.364882 | 8.30124 | 7.130273 | 6.859373 | 6.965114 | 7.688583 |
| 6.415491 | 5.54501 | 5.053798 | NaN | 4.738356 | 6.977938 | 6.730554 | 6.599288 | 6.058088 |
| 7.478646 | 7.624324 | 7.44321 | 6.398242 | 6.920819 | 7.973494 | 7.797855 | 7.923324 | 6.487393 |
| 8.105324 | 8.395239 | 8.140408 | 6.662824 | 6.609107 | 7.572916 | 7.825895 | 7.643514 | 7.776706 |
| 6.864049 | 7.907625 | 5.862619 | NaN | 4.751002 | 6.716812 | 6.398508 | 6.532952 | 7.38977 |
| 9.287488 | 9.281194 | 9.179304 | 9.070057 | 9.133051 | 9.367934 | 9.490442 | 9.542421 | 9.18312 |
| 8.834873 | 9.154729 | 9.019056 | 8.587465 | 8.584723 | 9.968106 | 9.737875 | 9.889013 | 8.428382 |
| 7.306226 | 5.299295 | 5.398194 | 5.98121 | 4.868209 | NaN | NaN | NaN | 5.702127 |

| | | | | | | | | |
|---|---|---|---|---|---|---|---|---|
| 6.754101 | 6.686704 | 6.810063 | 6.450837 | 6.059517 | 6.522213 | 7.08866 | 6.750942 | 6.319194 |
| 7.633909 | 7.487186 | 7.702724 | 6.006371 | 5.383829 | 7.626806 | 7.450127 | 7.643692 | 7.472321 |
| 8.491834 | 8.502086 | 8.43503 | 8.729696 | 8.777929 | 8.695099 | 8.329167 | 8.82714 | 8.305651 |
| 6.924031 | 6.924658 | 7.268201 | 7.655549 | 7.502429 | 7.082839 | 6.95818 | 7.292939 | 8.783337 |
| 7.883381 | 7.666187 | 7.463289 | 7.155528 | 7.294371 | 6.139827 | 6.490342 | 5.762431 | 6.43676 |
| 4.998178 | 6.741344 | 5.640486 | 5.013198 | 5.578168 | 6.296537 | 5.873878 | 6.107636 | 5.997585 |
| 9.675487 | 9.589087 | 9.714145 | 10.56893 | 10.71687 | 8.850444 | 8.79913 | 8.735326 | 9.528079 |
| 7.182911 | 7.608704 | 6.388208 | 4.176239 | 4.338162 | 6.121415 | 6.121663 | 5.81585 | 6.125287 |
| 6.73301 | 6.844256 | 6.620242 | 8.317013 | 8.428428 | 7.787996 | 7.59057 | 7.926908 | 6.958876 |
| 6.14402 | 5.683371 | 5.900989 | 4.753028 | 5.132939 | 5.851266 | 5.762724 | 5.899872 | 5.169418 |
| 7.336924 | 7.358843 | 7.276159 | 5.662517 | 6.063056 | 6.89944 | 7.260592 | 7.052623 | 7.271672 |
| 4.804916 | 4.333924 | 5.08423 | 5.1669 | 3.960471 | 5.348098 | 5.091365 | 5.278053 | 5.506794 |
| NaN | 3.947817 | 4.106888 | 5.541082 | 5.348245 | 4.514533 | 4.247254 | 4.149808 | 5.260738 |
| NaN | NaN | 3.912784 | 6.106631 | 6.488957 | NaN | NaN | NaN | 4.815103 |
| 7.065115 | 5.057763 | 5.642398 | NaN | 5.66696 | 4.883412 | 6.110091 | 6.816062 | NaN |
| 10.36372 | 10.41432 | 10.36829 | 10.44061 | 10.08058 | 8.68122 | 8.91781 | 8.946194 | 9.098273 |
| 7.517497 | 7.23794 | 7.565568 | 5.708789 | 6.086702 | 7.28724 | 7.565698 | 7.578252 | 6.860131 |
| 8.457383 | 8.288395 | 8.510692 | 6.826293 | 6.671231 | 7.983637 | 8.046206 | 8.262499 | 7.147807 |
| 8.682914 | 8.900386 | 8.63895 | 7.495266 | 7.881351 | 8.01665 | 8.253828 | 8.360789 | 8.335262 |
| 7.25123 | 7.143472 | 7.335077 | 5.575379 | 5.667666 | 7.485903 | 7.256307 | 7.344749 | 7.631846 |
| 5.630416 | 5.385266 | 5.884994 | 5.103659 | 3.934745 | 5.749749 | 6.918936 | 6.020375 | 5.847963 |
| 7.909641 | 8.166047 | 7.993107 | 6.122688 | 5.901576 | 7.818202 | 7.907138 | 7.829428 | 7.39936 |
| 3.744766 | NaN | 4.345433 | NaN | NaN | NaN | 5.367372 | NaN | 5.405207 |
| 6.266523 | 6.087596 | 7.027884 | 5.467479 | 6.015438 | 6.347462 | 6.400023 | 6.224602 | 7.006042 |
| 6.979953 | NaN | NaN | NaN | 6.187388 | 6.468923 | 6.960359 | 6.533613 | 7.355744 |
| 10.0969 | 10.14765 | 10.01034 | 10.96599 | 10.62722 | 10.50998 | 10.48879 | 10.24932 | 10.41958 |
| 9.273579 | 9.131461 | 9.235895 | 8.545617 | 8.443033 | 8.996854 | 9.310303 | 9.029794 | 9.228471 |
| 6.482938 | 6.226358 | 6.252097 | 6.442589 | 6.413218 | 6.376857 | 6.026596 | 6.305734 | 6.301971 |
| 8.434933 | 8.333338 | 8.341209 | 7.633507 | 7.690682 | 8.035921 | 8.462455 | 8.417125 | 8.540467 |
| 8.505657 | 8.590566 | 8.5108 | 7.868178 | 7.901032 | 8.404654 | 8.426827 | 8.453901 | 8.524285 |
| 3.900828 | 4.741756 | 5.053645 | NaN | NaN | 5.092357 | 5.159736 | 5.292821 | 5.62329 |
| 6.14232 | 4.876773 | 5.095449 | NaN | NaN | 6.555895 | 5.68157 | 5.822567 | 5.706389 |
| 6.420126 | 5.829686 | 6.395876 | 7.007285 | 6.902905 | 6.341787 | 6.314809 | 6.117857 | 6.555282 |
| 8.137487 | 8.192526 | 7.998841 | 6.980668 | 6.671851 | 8.261286 | 8.170045 | 8.355702 | 7.180384 |
| 6.891134 | 6.931138 | 7.092937 | 5.502181 | 5.555222 | 6.586034 | 6.366868 | 6.540472 | 6.368245 |
| 9.210196 | 9.324474 | 9.197145 | 11.00942 | 10.73938 | 9.287234 | 9.148214 | 9.215425 | 8.895662 |
| NaN | NaN | NaN | 4.98713 | 4.129282 | 4.171205 | NaN | 2.677674 | 4.622656 |
| 8.771079 | NaN | NaN | 6.09878 | NaN | 7.419897 | 5.983987 | 7.46487 | 7.80809 |
| 5.676496 | 6.165517 | 6.13217 | NaN | 4.481138 | 6.385692 | 5.333529 | NaN | 5.755419 |
| NaN | 8.145683 | NaN | 4.121333 | 5.029683 | NaN | NaN | NaN | 4.982378 |
| 5.687449 | NaN | 5.845913 | NaN | 5.672669 | 6.976395 | 6.458189 | 6.218979 | 7.144732 |
| 7.546234 | 7.384456 | 7.379249 | 6.38256 | 6.212825 | 7.285004 | 7.33554 | 7.543633 | 6.350953 |
| 7.759354 | 8.600781 | 8.569739 | 8.336075 | 8.010855 | 8.416004 | 8.238583 | 8.455059 | 7.936669 |
| 5.951721 | 5.813797 | 6.398293 | 6.610946 | 6.287231 | 6.771605 | NaN | 6.895894 | 6.136892 |
| 9.394682 | 8.984107 | 9.24065 | 10.62335 | 10.50073 | 9.449052 | 9.386351 | 9.297221 | 9.475696 |
| 6.598039 | 6.39974 | 6.823456 | 8.57919 | 8.655695 | 7.088582 | 7.261686 | 7.025555 | 6.538752 |
| 7.767582 | 7.549491 | 7.774343 | 6.284849 | 6.971523 | 8.086482 | 7.807707 | 8.241949 | 7.554049 |

-Bruker Confidential-

| | | | | | | | | |
|---|---|---|---|---|---|---|---|---|
| 11.40153 | 11.27279 | 11.24467 | 11.66279 | 11.73094 | 10.66815 | 10.67237 | 10.76226 | 7.551989 |
| NaN | NaN | 4.209021 | NaN | NaN | 4.913926 | 3.830988 | 4.531651 | 3.962771 |
| 9.433752 | 9.337693 | 9.013551 | 7.965845 | 7.964003 | 8.536569 | 8.670602 | 8.777101 | 9.154054 |
| 8.556097 | 8.754274 | 8.444892 | 8.428573 | 7.77952 | 8.191903 | 7.987324 | 8.262869 | 7.784979 |
| 9.289203 | 6.990158 | 7.674094 | 8.563237 | NaN | 9.503668 | 8.610347 | 8.808944 | 8.007777 |
| 6.122929 | 6.169096 | 6.237371 | 5.323781 | 6.649158 | 5.687762 | 5.979904 | 5.856636 | 5.62366 |
| 5.531313 | 4.456081 | 6.491965 | NaN | 4.235787 | 5.907727 | 5.331148 | 5.871301 | 5.166054 |
| 5.749577 | 6.170166 | 6.590921 | 4.843029 | 5.405093 | 6.40181 | 7.024417 | 7.071972 | 5.782494 |
| 6.391703 | 6.927904 | 6.458963 | 5.025421 | 5.776678 | 7.150607 | 6.860177 | 7.088833 | 6.655058 |
| 6.835658 | 6.488848 | 5.995827 | 5.946842 | 6.09994 | 5.358791 | 5.406865 | 5.914016 | 6.740623 |
| 10.95726 | 11.01688 | 10.81717 | 11.57334 | 11.4145 | 10.84394 | 10.81426 | 10.69606 | 10.45644 |
| 7.921824 | 7.953275 | 8.332093 | 6.686696 | 7.206757 | 8.202687 | 8.04655 | 7.798133 | 8.098916 |
| 10.69606 | 10.37332 | 10.493 | 10.0662 | 9.849441 | 10.00086 | 9.984241 | 10.26624 | 10.55431 |
| 7.298418 | 6.577951 | 6.966493 | 8.760706 | 7.49774 | 5.44179 | NaN | NaN | 7.544465 |
| 6.158777 | 5.126084 | 5.605483 | NaN | NaN | 6.946101 | 7.27329 | 6.343646 | 5.695485 |
| NaN | 6.173555 | 6.810409 | 6.31002 | 9.174751 | 6.857473 | 9.466395 | 9.257511 | 7.284709 |
| 9.509658 | 9.4295 | 9.639803 | 9.948151 | 10.13841 | 9.594724 | 9.66795 | 9.637694 | 9.357293 |
| 10.48665 | 10.65536 | 10.31914 | 9.873031 | 9.709082 | 10.92837 | 10.74795 | 10.97851 | 10.24317 |
| 10.297 | 10.21439 | 10.23061 | 10.94363 | 10.70393 | 9.474148 | 9.659509 | 9.599225 | 9.942483 |
| 7.225044 | 7.282954 | 7.297815 | 6.915376 | 6.145205 | 7.034091 | 7.01748 | 7.187707 | 7.415377 |
| 6.525698 | 6.419393 | 6.073955 | 5.912399 | 5.760697 | 6.616 | 6.376879 | 6.459503 | 6.939948 |
| NaN | NaN | NaN | NaN | NaN | 4.774963 | NaN | NaN | 5.632866 |
| NaN | NaN | NaN | 3.450762 | NaN | NaN | NaN | NaN | 4.970853 |
| 7.914549 | 8.030623 | 8.050681 | 8.382189 | 8.310067 | 7.222665 | 7.252794 | 7.239645 | 8.370834 |
| 7.845346 | 6.856061 | 7.465301 | 5.745044 | 5.485224 | 7.242729 | 7.606411 | 7.0464 | 7.089991 |
| 8.091784 | 7.801964 | 8.233346 | 7.429682 | 7.215181 | 8.214147 | 7.990689 | 8.102546 | 7.906564 |
| 2.71303 | 3.856224 | 3.317845 | 4.011152 | NaN | 5.400855 | 5.045926 | 5.628142 | 4.967597 |
| 8.808223 | 8.708249 | 8.710513 | 8.609718 | 8.502264 | 9.300091 | 9.150682 | 9.333994 | 7.712523 |
| 6.903113 | 6.603793 | 6.789867 | 7.673088 | 7.535731 | 6.726916 | 7.223956 | 6.706357 | 6.053329 |
| 4.749553 | NaN | 5.094082 | 6.570125 | NaN | 5.620141 | NaN | NaN | 4.510042 |
| 6.288519 | 7.198339 | 6.604559 | NaN | 5.528439 | 7.506301 | 7.585157 | 6.935241 | 7.341193 |
| 7.042622 | 4.690798 | 6.434315 | 6.95071 | 6.575157 | 6.647406 | 6.16749 | 6.418741 | 6.251076 |
| 5.007405 | NaN | NaN | 5.242936 | 4.80285 | NaN | 4.602535 | 5.173367 | 2.893897 |
| 6.988892 | 7.097169 | 7.400964 | 5.857298 | 6.435082 | 6.772554 | 7.102552 | 6.671052 | 6.630558 |
| 6.083064 | 6.608459 | 5.883616 | 6.507349 | 6.597402 | 6.004895 | 5.668735 | 5.279098 | 5.801939 |
| 9.128237 | 9.090754 | 8.986036 | 10.20934 | 10.41084 | 8.759513 | 8.899421 | 8.708452 | 9.329968 |
| NaN | 5.716414 | 5.243726 | 6.221346 | 6.377322 | 5.393958 | 5.143258 | 5.470213 | 6.022216 |
| 9.425338 | 9.510226 | 9.351748 | 9.819851 | 9.914965 | 9.173654 | 8.880013 | 8.944651 | 9.676712 |
| 12.02174 | 12.04594 | 11.99373 | 12.54154 | 12.56907 | 12.15977 | 12.11346 | 12.07665 | 12.2683 |
| 8.238544 | 8.152128 | 8.38373 | 8.275661 | 7.819563 | 7.235538 | 7.777277 | 7.71082 | 8.054868 |
| 9.556766 | 9.631903 | 9.564809 | 9.05432 | 9.06866 | 9.599582 | 9.261592 | 9.631417 | 9.191335 |
| 7.983164 | 8.425418 | 8.421936 | 10.75588 | 10.77564 | 7.339044 | 7.383666 | 7.483253 | 7.676952 |
| 8.444288 | 8.145287 | 8.351952 | 7.445257 | 7.31278 | 7.10493 | 7.752434 | 7.82386 | 8.05196 |
| 5.638068 | 3.369364 | 4.896829 | 4.111405 | NaN | 5.90735 | 5.394234 | 6.425401 | 3.182775 |
| 6.115916 | 6.203562 | 6.186309 | 5.099611 | 5.321417 | 6.363697 | 6.0674 | 6.308612 | 5.280025 |
| 7.683717 | 7.298878 | 7.386057 | 6.500278 | 6.342867 | 7.727887 | 7.459908 | 7.456897 | 7.443507 |
| 7.223231 | 7.393857 | 7.456737 | 6.564524 | 6.569399 | 7.171733 | 7.228003 | 6.814834 | 7.23719 |

-Bruker Confidential-

| | | | | | | | | |
|---|---|---|---|---|---|---|---|---|
| 8.327209 | 8.376482 | 8.557846 | 8.590517 | 8.26005 | 8.130852 | 7.931607 | 8.136455 | 7.812569 |
| 8.745353 | 8.376461 | 8.580982 | 10.61086 | 10.62827 | 7.600806 | 7.732039 | 7.713029 | 8.228836 |
| 8.945906 | 9.057858 | 9.055696 | 8.570016 | 8.46475 | 8.2964 | 8.841157 | 8.902491 | 8.485428 |
| 6.329117 | 6.299163 | 6.479268 | 7.925183 | 7.896409 | 6.617761 | 6.876317 | 6.67511 | 7.41293 |
| 6.807024 | 6.858832 | 6.744793 | 8.330663 | 7.396392 | 6.514511 | 6.590992 | 6.535243 | 7.504773 |
| 6.383079 | 6.493313 | 6.574203 | 7.0766 | 7.769042 | 4.083666 | 6.29986 | 6.427474 | 6.064881 |
| 7.164281 | 7.241069 | 7.336902 | 5.945979 | 6.455401 | 7.510155 | 7.397497 | 7.342009 | 6.947616 |
| 6.039264 | 6.766023 | 6.78558 | 6.19865 | 5.835607 | 6.595019 | 6.848199 | 6.267053 | 6.350949 |
| NaN | NaN | NaN | 5.787763 | 6.175611 | NaN | NaN | NaN | 4.347664 |
| 3.955901 | NaN | 5.421645 | 5.599182 | 5.952092 | 5.274818 | 5.233327 | 5.811123 | 4.216393 |
| 6.890343 | 6.918823 | 6.779317 | 6.236291 | 6.459975 | 7.057206 | 7.083653 | 6.976742 | 5.847154 |
| 7.331456 | 7.12693 | 7.163374 | 4.788069 | 5.478497 | NaN | NaN | NaN | NaN |
| 6.912162 | 6.256958 | 6.92737 | 8.954927 | 8.957459 | 6.459846 | 7.425598 | 7.447608 | 6.648816 |
| NaN | 7.03758 | NaN | 6.447332 | 6.740285 | NaN | 7.027247 | NaN | NaN |
| 7.116593 | 6.85349 | 7.048919 | 7.524491 | 7.722289 | 6.434413 | 6.708864 | 6.564428 | 5.724984 |
| 7.060168 | 6.806666 | 6.980602 | 7.115249 | 6.919252 | 6.816376 | 6.392999 | 6.478507 | 6.208694 |
| 7.954208 | 7.942652 | 7.748139 | 6.552469 | 6.583654 | 7.755766 | 7.842319 | 7.893598 | 8.124052 |
| 6.804606 | 6.581627 | 6.657103 | 7.972884 | 8.055 | 8.719679 | 8.21773 | 8.075943 | 6.596148 |
| 12.62989 | 12.5146 | 12.53791 | 12.89999 | 12.70485 | 12.78508 | 12.88113 | 12.86203 | 11.76249 |
| 8.923399 | 8.985629 | 8.871795 | 9.109041 | 8.813318 | 8.869073 | 8.870245 | 8.671836 | 8.724444 |
| 8.646857 | 8.28388 | 8.717135 | 8.496811 | 6.996908 | 8.268723 | 7.627686 | 7.659005 | 7.44701 |
| 10.23293 | 9.88084 | 9.885724 | 10.70334 | 10.66841 | 9.408853 | 9.341037 | 9.427994 | 8.909447 |
| 8.797666 | 8.776891 | 8.84349 | 8.829198 | 8.370208 | 8.259285 | 8.359266 | 8.452639 | 8.192959 |
| 9.243052 | 8.739082 | 9.391746 | 6.022716 | 6.727648 | 7.715939 | 7.97264 | 7.871964 | 8.800218 |
| 6.893522 | 6.916866 | 6.844433 | 6.402616 | 6.348567 | 6.682014 | 6.493125 | 6.489203 | 7.364025 |
| 8.660445 | 7.915664 | 8.238943 | 9.507677 | 8.937728 | 7.460453 | 7.724066 | 6.566289 | 7.010298 |
| 7.866107 | 7.149321 | 7.012493 | 7.772949 | 7.670318 | 6.838127 | 6.388413 | 6.331369 | 6.504761 |
| 8.338736 | 9.012614 | 8.704634 | 8.732563 | 8.884066 | 8.091645 | 8.525059 | 8.174829 | 7.145129 |
| NaN | NaN | NaN | 6.542808 | 6.775878 | NaN | NaN | NaN | NaN |
| 4.531714 | 5.696345 | 5.807858 | 8.404628 | 8.211539 | NaN | NaN | NaN | 5.604622 |
| 9.705224 | 9.740446 | 9.79347 | 9.389877 | 9.381661 | 10.30477 | 10.15732 | 10.16314 | 9.418591 |
| 10.0528 | 10.019 | 9.93423 | 9.003847 | 8.845379 | 9.721082 | 9.613837 | 9.629183 | 9.973479 |
| 7.9156 | 8.232317 | 7.522146 | 8.396866 | 8.221815 | 8.157408 | 8.361134 | 7.899392 | 6.288003 |
| 9.809279 | 9.606066 | 9.944448 | 10.26768 | 10.3215 | 10.73387 | 10.47945 | 10.61403 | 9.638012 |
| 9.06722 | 9.056576 | 9.090531 | 8.311074 | 8.42475 | 9.210168 | 9.193308 | 9.336298 | 9.445539 |
| 6.987226 | 6.350468 | 6.923744 | 5.517986 | 5.617033 | 6.901091 | 5.99977 | 5.840549 | 4.978525 |
| 6.032331 | 5.829022 | 5.41404 | NaN | 4.903009 | 6.258507 | 5.730265 | 6.132804 | 5.568284 |
| 6.747035 | 6.919472 | 6.9296 | 7.477603 | 8.056534 | 4.981632 | 5.52831 | 5.111769 | 5.573836 |
| 8.16492 | 8.068776 | 8.156623 | 8.154401 | 8.183376 | 7.962668 | 7.916856 | 7.999603 | 8.581385 |
| 6.972855 | 6.561138 | 6.618655 | 6.536727 | 7.477609 | 6.810971 | 6.719781 | 6.831625 | 6.553001 |
| 8.107794 | 8.020949 | 8.171801 | 7.803316 | 7.624748 | 7.967334 | 7.903749 | 8.076048 | 7.918651 |
| 6.547653 | 7.015666 | 6.957807 | 7.628599 | 7.600634 | 7.025745 | 6.83628 | 6.995234 | 6.773212 |
| 8.97531 | 8.953176 | 8.399653 | 6.75825 | 6.424003 | 7.620946 | 7.488302 | 7.033725 | 6.966072 |
| 10.58661 | 9.701347 | 9.675727 | 6.970778 | 8.08322 | 9.891619 | 8.9775 | 10.04907 | 7.765501 |
| 11.35674 | 11.36654 | 11.41835 | 11.76847 | 11.39505 | 11.51083 | 11.23343 | 11.32433 | 10.96839 |
| 7.068924 | 7.157521 | 7.168476 | 9.633711 | 9.52812 | 5.241933 | 4.747272 | 4.549954 | 5.649028 |
| 6.360975 | 5.595633 | 6.648909 | 5.961759 | 5.747633 | 6.99409 | 6.718504 | 6.861409 | 6.407906 |

| | | | | | | | | |
|---|---|---|---|---|---|---|---|---|
| 9.053199 | 9.0342 | 9.063594 | 7.796086 | 8.138892 | 8.628521 | 8.651753 | 8.502299 | 9.129548 |
| 7.953814 | 5.371021 | 5.485602 | 10.54432 | 7.938224 | NaN | 5.142327 | NaN | NaN |
| 7.35461 | 7.518422 | 7.527831 | 7.445411 | 7.382205 | 7.755981 | 7.646259 | 7.588269 | 8.026258 |
| 5.64467 | 4.506489 | 3.75991 | 7.56626 | 7.093178 | 5.992301 | NaN | NaN | 5.385454 |
| 7.61302 | 7.493532 | 7.436088 | 6.465656 | 7.085638 | 7.330303 | 7.224036 | 7.730725 | 6.997281 |
| 8.843925 | 8.671174 | 8.673825 | 9.506047 | 9.312881 | 8.374058 | 8.844845 | 8.553069 | 8.957364 |
| 9.175163 | 9.878706 | 9.479074 | 7.344291 | 7.237116 | 9.285147 | 9.21771 | 9.129518 | 9.329253 |
| 7.134993 | 7.007732 | 7.046498 | 6.978226 | 7.062334 | 7.171934 | 7.00445 | 7.323673 | 6.902195 |
| 6.77608 | 7.376714 | 7.18747 | 5.817948 | 6.070368 | 6.583083 | 7.018547 | 6.781588 | 6.985266 |
| 4.667953 | 4.041117 | 4.686416 | 4.307792 | 5.082726 | 5.160665 | 4.00757 | 4.89475 | 2.830661 |
| 5.714957 | NaN | 5.279149 | NaN | 4.447763 | 6.973857 | 6.612659 | 7.217371 | 5.718866 |
| 5.739819 | 5.733746 | 5.848461 | 5.698419 | 5.446244 | 5.206904 | 5.149569 | 5.656091 | 6.121842 |
| 7.646804 | 7.745803 | 7.867538 | 7.439399 | 7.371597 | 8.192038 | 7.826705 | 8.037341 | 7.200046 |
| 6.080096 | 6.797345 | 6.607326 | NaN | 6.209418 | 6.398073 | 6.606747 | 6.539027 | 7.517815 |
| 11.48738 | 11.31199 | 11.39156 | 12.46864 | 12.32903 | 10.3279 | 10.37951 | 10.28091 | 11.84072 |
| 7.887866 | 7.623572 | 7.69439 | 8.666486 | 8.524404 | 7.215757 | 7.137026 | 7.491469 | 6.963353 |
| 9.841428 | 9.729749 | 9.805824 | 8.852295 | 8.698013 | 9.906911 | 9.900396 | 9.91951 | 9.574676 |
| 6.853452 | 6.559891 | 7.295943 | 6.643755 | 7.76257 | 7.324293 | 6.702084 | 7.365308 | 7.171019 |
| 6.499439 | 6.847273 | 6.748359 | NaN | NaN | 8.044502 | 7.988375 | 7.921371 | 6.7971 |
| 8.96186 | 8.194577 | 8.904655 | NaN | 9.076842 | 7.719821 | 7.673635 | 7.855939 | 8.11578 |
| 7.694753 | 7.651089 | 8.241504 | 8.531825 | 7.634627 | 6.643984 | 7.073181 | 6.905759 | 7.582032 |
| 8.182106 | 8.201284 | 8.208457 | 9.176513 | 9.427155 | 7.988124 | 8.375802 | 8.395877 | 8.284828 |
| 7.854003 | 7.635805 | 7.791498 | 5.471226 | NaN | 6.464848 | 7.061013 | 7.095148 | 7.387444 |
| 8.13178 | 8.293066 | 8.068801 | 8.392321 | 8.522661 | 8.446366 | 8.244333 | 8.17265 | 8.102343 |
| 8.453683 | 8.423704 | 8.272986 | NaN | NaN | NaN | NaN | NaN | NaN |
| 8.797617 | 8.721552 | 8.848527 | 10.01668 | 10.03494 | 7.58333 | 7.229028 | 7.243553 | 7.775638 |
| 6.987323 | 6.959696 | 6.710174 | 6.247212 | 5.301295 | 7.097443 | 6.229681 | 6.232827 | 6.616487 |
| 6.917962 | 7.087029 | 7.081341 | 6.001181 | 5.920578 | 6.823032 | 6.676503 | 6.781938 | 7.531404 |
| 8.029436 | 9.134979 | 8.581902 | 8.812775 | 8.704011 | 6.292718 | NaN | 6.810103 | 5.900076 |
| 7.906446 | 7.372976 | 7.505725 | 8.984676 | 8.674974 | 7.015473 | 7.284968 | 7.746563 | 5.464595 |
| NaN | 7.58112 | 6.852443 | 7.373597 | 7.620789 | 7.30397 | 7.445464 | 7.783518 | 8.336118 |
| 7.044316 | 7.375214 | 6.626891 | 6.294752 | 8.05769 | 6.969083 | 7.579681 | 7.279078 | 6.048973 |
| 5.885977 | 6.037012 | 6.244205 | 4.119759 | 4.366101 | 6.256935 | 6.333748 | 6.463646 | 6.100688 |
| 7.284467 | 7.344344 | 7.949238 | 6.889873 | 6.893193 | 7.43672 | 6.83991 | 6.976601 | 6.915475 |
| 8.812178 | 8.90036 | 9.025597 | 8.863843 | 8.742605 | 9.487307 | 9.27081 | 8.83587 | 8.68063 |
| 8.584091 | 8.323648 | 8.511489 | 9.247225 | 7.635118 | 8.970426 | 8.413656 | 8.762799 | 7.448545 |
| 10.63794 | 10.65521 | 10.73676 | 9.793865 | 9.767757 | 10.09785 | 10.15863 | 10.13739 | 10.64171 |
| 7.214057 | 7.214133 | 7.106725 | 6.598863 | 6.828887 | 7.193833 | 7.333572 | 7.139675 | 6.326704 |
| 9.094805 | 8.895322 | 8.967258 | 9.372206 | 9.191265 | 9.012483 | 8.885038 | 9.008354 | 9.042742 |
| 11.28178 | 11.19984 | 11.12716 | 10.68464 | 10.61704 | 11.62278 | 11.55642 | 11.69057 | 11.29274 |
| 7.049161 | 5.840647 | 5.45477 | NaN | 8.118307 | 6.725068 | 6.663296 | 6.778458 | 6.235722 |
| 7.437953 | 7.457887 | 7.460277 | 6.546125 | 6.613513 | 7.005731 | 7.527798 | 7.480306 | 7.025766 |
| 7.417719 | 6.151417 | NaN | 6.365708 | 6.192051 | 7.010554 | 6.749068 | 6.292834 | 6.003942 |
| 8.183418 | 7.912619 | 8.181209 | 7.122283 | 7.047916 | 7.617678 | 7.733582 | 7.817288 | 7.230661 |
| 10.33768 | 10.48548 | 10.27132 | 10.06921 | 9.911167 | 10.55684 | 10.56304 | 10.51447 | 9.77925 |
| 6.246641 | 6.381628 | 6.586142 | 6.942235 | 6.995583 | NaN | 6.46632 | 6.363929 | 6.17826 |
| NaN | 6.64579 | 7.253741 | NaN | 6.032582 | 6.982705 | 7.187307 | 6.85203 | 6.582149 |

| | | | | | | | | |
|---|---|---|---|---|---|---|---|---|
| 7.721574 | 7.539265 | 7.711809 | 6.483202 | 7.097618 | 7.616069 | 7.699227 | 7.869175 | 6.376758 |
| 6.462028 | 6.338243 | 6.492158 | 6.24241 | 6.373946 | 6.540451 | 6.210249 | 6.692606 | 6.119391 |
| 8.779244 | 8.622652 | 8.478181 | 7.079556 | 7.263256 | 8.586822 | 8.276045 | 8.53981 | 8.41386 |
| NaN | 5.478757 | 5.862167 | 4.465265 | 5.263462 | 6.499704 | 6.092155 | 6.605575 | 5.903386 |
| 5.093408 | 5.185105 | NaN | 6.811432 | 6.385521 | NaN | 3.976562 | NaN | 4.72785 |
| NaN | 7.145996 | NaN | 9.117604 | 9.36752 | 8.328404 | 8.364289 | 7.974959 | 8.273242 |
| 6.156091 | 5.064627 | 4.814576 | 3.018784 | NaN | 5.203775 | 5.683454 | 5.954019 | 5.692525 |
| NaN | 3.467863 | NaN | 6.576562 | NaN | 5.591128 | 6.393352 | 5.999394 | 7.308574 |
| 8.03062 | 7.937109 | 7.884233 | 7.741634 | 7.522079 | 7.770161 | 7.923128 | 7.642129 | 6.987484 |
| 8.536467 | 8.402998 | 8.482738 | 6.70765 | 6.299246 | 8.229351 | 8.408335 | 8.426633 | 7.021833 |
| 7.178366 | 6.573694 | 6.869193 | 5.489562 | 5.771126 | 7.816572 | 7.853981 | 7.915463 | 7.407099 |
| 6.524117 | 6.500293 | 6.468827 | 5.536278 | 5.123052 | 6.687869 | 6.699271 | 6.781513 | 5.576658 |
| 7.121083 | 7.089827 | 7.446065 | 6.215306 | 6.024692 | 6.940668 | 7.40502 | 7.389927 | 7.082133 |
| 8.337932 | 8.422633 | 8.410751 | 9.081603 | 9.352108 | 8.279699 | 8.29957 | 8.517182 | 8.441726 |
| 9.005013 | 8.92206 | 8.961415 | 8.254941 | 8.381465 | 8.853585 | 8.740531 | 8.993126 | 8.517995 |
| 10.41414 | 10.37596 | 10.53113 | 10.70984 | 10.86776 | 10.34447 | 10.59744 | 10.65633 | 10.47492 |
| 6.46307 | 6.758202 | 6.529594 | 5.205194 | 6.530065 | 7.157814 | 6.790413 | 6.56664 | 5.49517 |
| 7.00693 | 6.579663 | 7.035672 | 6.241889 | 5.704374 | 6.326173 | 6.951307 | 6.283028 | 7.043985 |
| 7.189856 | 7.35511 | 7.659708 | 6.479615 | NaN | 7.205465 | 7.277197 | 7.167336 | 6.765637 |
| 7.026533 | 6.908878 | 6.678438 | 7.534271 | 7.289505 | 7.331537 | 7.638148 | 7.50772 | 7.469597 |
| 8.431994 | 8.08586 | 8.323385 | 6.710851 | 6.781641 | 8.020964 | 8.096433 | 8.11368 | 7.762999 |
| 6.405282 | 6.692583 | 6.520813 | 6.069158 | 5.956633 | 6.821581 | 7.022988 | 6.885028 | 6.850594 |
| 7.250395 | 7.175989 | 7.107902 | 6.233012 | 6.261098 | 7.211333 | 7.256443 | 7.277212 | 7.65202 |
| 7.30862 | 7.13976 | 7.446898 | 7.913162 | 8.161049 | 7.811943 | 7.678917 | 8.095126 | 7.298378 |
| 7.447953 | 7.440771 | 7.478504 | 7.011921 | 6.94301 | 7.483002 | 7.605978 | 7.425743 | 7.430196 |
| 7.992324 | 7.786071 | 8.781702 | 8.708388 | 8.335315 | 7.905043 | 7.894046 | 7.444278 | 7.052619 |
| 9.180005 | 9.269902 | 9.410726 | 8.603238 | 8.498068 | 9.312132 | 8.793073 | 8.849678 | 7.622536 |
| 7.736711 | 7.687484 | 7.557825 | 7.244798 | 7.141737 | 7.923986 | 8.05667 | 7.888798 | 7.323019 |
| 7.632304 | 7.058946 | 7.045043 | 8.322952 | 8.585189 | 7.04743 | 6.844345 | 6.673173 | 6.486936 |
| 6.664597 | 6.10862 | 5.80721 | NaN | NaN | NaN | 7.588623 | 7.362276 | 6.855681 |
| 6.58535 | 5.853316 | 4.781314 | NaN | NaN | 6.480449 | 6.952829 | 6.801009 | 6.612933 |
| 8.233523 | 8.442322 | 8.267628 | 7.640444 | 7.548679 | 8.156378 | 7.550434 | 7.802032 | 7.965741 |
| 9.671724 | 9.520016 | 9.607559 | 7.582224 | 7.242111 | 8.38344 | 8.583334 | 8.621465 | 8.851944 |
| 8.880699 | 8.858121 | 8.864567 | 9.511085 | 9.504992 | 8.981106 | 8.863987 | 9.089049 | 9.020033 |
| 9.583066 | 9.765452 | 9.428219 | 10.31653 | 10.38544 | 7.38175 | 7.545909 | 7.298285 | 8.077799 |
| 9.521483 | 9.505477 | 9.343709 | 9.559505 | 9.581946 | 9.250865 | 9.286429 | 9.215032 | 9.090075 |
| 8.826302 | 8.757358 | 8.727062 | 8.485468 | 8.384899 | 8.544235 | 8.819369 | 8.793467 | 8.296844 |
| 8.646548 | 8.687493 | 8.618155 | 8.140368 | 8.122481 | 8.764479 | 8.649188 | 8.786049 | 9.145491 |
| 7.341922 | 5.636939 | 7.264084 | 7.013847 | 6.531576 | 6.646834 | 7.179694 | 7.225479 | 6.412187 |
| 8.149093 | 7.999395 | 7.986193 | 8.852792 | 8.808935 | 8.490556 | 8.282282 | 8.558956 | 8.342927 |
| 7.027986 | 7.150032 | 6.455399 | 6.700409 | NaN | 6.221711 | 5.965191 | 6.109304 | 6.531263 |
| 7.434104 | 7.135379 | 7.573029 | 7.084108 | 7.228831 | 8.194571 | 8.053574 | 8.073001 | 7.770617 |
| 7.536113 | 7.154355 | 6.438076 | 8.491821 | 8.445803 | 6.346042 | 4.745491 | 5.009408 | 6.31744 |
| 7.639162 | 7.411237 | 7.665571 | 7.17618 | 7.240984 | 6.675206 | 6.651814 | 7.183799 | 5.187608 |
| 8.351067 | 8.258206 | 8.250978 | 7.215025 | 8.605449 | 8.192021 | 7.763707 | 8.096816 | 7.902233 |
| 7.513763 | 7.512974 | 7.516137 | 7.056802 | 6.587007 | 8.462143 | 8.068819 | 8.404162 | 8.462124 |
| 7.827993 | 7.859087 | 7.683303 | 7.295429 | 7.188642 | 7.607297 | 7.788942 | 7.96234 | 8.001726 |

-Bruker Confidential-

| | | | | | | | | |
|---|---|---|---|---|---|---|---|---|
| 5.033031 | 4.880838 | 5.576662 | NaN | 6.017505 | 5.113124 | 5.685267 | 4.382217 | 5.796373 |
| 7.480437 | 7.432797 | 7.346869 | 6.570978 | 6.42086 | 7.467372 | 7.187811 | 7.257922 | 6.064254 |
| 6.174819 | 6.390398 | 6.629261 | 7.666384 | 8.043212 | 5.28554 | 5.685968 | 5.371486 | 6.795439 |
| 7.077581 | 7.092904 | 7.162453 | 7.148128 | 7.117502 | 6.530826 | 6.561225 | 6.646484 | 6.969769 |
| NaN | 6.834745 | 5.216975 | 5.485697 | NaN | 5.602789 | 5.676701 | 6.090895 | 5.345563 |
| 9.143026 | 9.082535 | 8.924426 | 10.03144 | 9.997609 | 9.51965 | 9.387939 | 9.472589 | 9.119154 |
| 7.986669 | 7.693756 | 7.686745 | 8.65944 | 8.328692 | 8.05212 | 7.802813 | 7.861794 | 7.651141 |
| 9.631563 | 9.654193 | 10.15888 | 11.30163 | 11.23955 | 8.013596 | 8.388343 | 8.112448 | 8.577349 |
| 5.007648 | 3.83236 | 3.794743 | NaN | NaN | 5.070803 | 4.877949 | 5.695426 | 4.545797 |
| 7.008015 | 6.875432 | 7.205648 | NaN | 6.179462 | NaN | NaN | NaN | 6.170916 |
| 5.085281 | NaN | 4.195381 | NaN | NaN | 4.738976 | 5.003877 | 5.0091 | 5.639122 |
| 8.480821 | 8.653949 | 8.876898 | 10.4615 | 10.73432 | 8.823431 | 8.546535 | 8.816629 | 8.968564 |
| 6.997999 | 7.058578 | 7.039117 | 7.912282 | 8.014129 | 6.811845 | 6.52633 | 6.897044 | 6.151554 |
| 7.209501 | 6.726901 | 7.128592 | 5.935405 | 6.113423 | 7.380131 | 7.115568 | 6.850208 | 6.555009 |
| 7.074101 | 5.155451 | NaN | 8.304399 | 8.229266 | 6.640274 | 6.161061 | 6.361376 | 5.647154 |
| NaN | 5.849957 | NaN | 7.258328 | 7.284308 | NaN | NaN | NaN | 5.431899 |
| 6.561422 | 6.329119 | 6.19228 | NaN | 5.510451 | 6.124873 | NaN | 6.385184 | 6.033813 |
| 8.107153 | 8.099358 | 7.942381 | 7.46334 | 7.379688 | 7.718866 | 7.447453 | 7.830914 | 7.22265 |
| 8.905705 | 8.848168 | 8.8371 | 8.145719 | 8.354752 | 8.830098 | 9.00549 | 8.869493 | 8.502056 |
| 7.958555 | 8.107133 | 7.922747 | 7.263253 | 7.485842 | 7.980632 | 8.221998 | 8.058715 | 7.255279 |
| 6.483815 | 6.351327 | 6.280257 | 5.454545 | 5.393094 | 6.847888 | 6.908468 | 6.672414 | 5.601603 |
| 8.971436 | 9.1613 | 8.984673 | 6.861028 | 6.960907 | 9.244496 | 9.221533 | 9.356565 | 8.266447 |
| 7.051262 | 6.936887 | 7.34255 | 8.188598 | 7.714013 | 7.08615 | 7.396761 | NaN | 6.950044 |
| 7.917985 | 8.111222 | 8.116658 | 9.2873 | 9.499756 | 8.096617 | 7.871011 | 7.815583 | 7.356127 |
| 4.962144 | 6.904609 | 4.884916 | NaN | 4.655444 | 6.101321 | 6.056644 | 6.171346 | 5.678157 |
| NaN | NaN | NaN | NaN | 5.797153 | 6.733963 | 6.932445 | 6.361929 | 5.516156 |
| 9.957909 | 9.939663 | 9.935087 | 8.762191 | 8.549926 | 10.26431 | 9.884954 | 9.736028 | 9.384914 |
| 8.505274 | 8.490274 | 8.457314 | 7.575571 | 7.644902 | 8.601336 | 8.59139 | 8.622832 | 6.558353 |
| NaN | NaN | NaN | NaN | NaN | 6.09587 | 5.910381 | 6.179464 | 6.494232 |
| 6.464717 | 5.125532 | 6.847508 | NaN | NaN | 6.286182 | 6.481009 | 6.284258 | 6.413257 |
| 9.174786 | 9.037857 | 8.978329 | 7.326335 | 7.572897 | 9.856986 | 9.64938 | 9.879382 | 8.642449 |
| 7.117706 | 7.194334 | 6.872935 | 6.784842 | 7.140402 | 7.014304 | 6.38408 | 6.82847 | 6.781076 |
| 8.095366 | 7.998562 | 8.131785 | 7.444094 | 7.554481 | 8.401398 | 8.092571 | 8.236406 | 7.643351 |
| 8.752453 | 8.908275 | 8.640126 | 8.555939 | 8.324868 | 8.645468 | 8.752177 | 8.833609 | 8.123467 |
| 5.544335 | 5.777452 | 5.438658 | 5.514023 | 5.412024 | 6.09836 | 6.11283 | 6.528753 | 5.823708 |
| 8.651835 | 8.803555 | 8.863684 | 8.161637 | 8.501884 | 9.228629 | 9.396372 | 9.530543 | 8.982347 |
| 6.661448 | NaN | NaN | 8.069651 | 8.42896 | 7.570436 | 7.7051 | 7.472926 | 7.550634 |
| 8.04458 | 7.851374 | 8.045807 | 7.72328 | 7.643739 | 8.223238 | 8.219647 | 8.367183 | 7.185281 |
| 7.673976 | 7.905373 | 7.958628 | 7.892878 | 7.322814 | 8.674901 | 8.011208 | 7.751529 | 7.359812 |
| 7.817189 | 7.793297 | 7.826701 | 7.383623 | 7.415367 | 7.209823 | 7.108469 | 7.266001 | 7.391155 |
| 6.583718 | 6.796301 | 6.655308 | 6.060856 | 6.404328 | 7.151505 | 7.253151 | 6.900242 | 6.556124 |
| 7.249918 | 7.425388 | 7.193986 | 6.923666 | 6.722439 | 6.992167 | 7.049116 | 6.779704 | 7.337523 |
| 7.142664 | 7.253224 | 7.64572 | 7.070441 | 8.179864 | 8.045159 | 7.766528 | 7.683003 | 6.969291 |
| 6.784234 | 5.708861 | 6.983366 | 6.153212 | 6.025203 | 6.700209 | 6.489102 | 6.829573 | 6.291804 |
| 8.937837 | 8.341135 | 8.513935 | 9.000642 | 8.922743 | 7.4972 | 7.55877 | 7.376265 | 8.441347 |
| 6.466468 | 6.564558 | 6.529638 | 5.267584 | 5.558043 | 7.042513 | 7.327364 | 7.268284 | 6.457631 |
| NaN | 8.72771 | 9.719716 | 5.810923 | NaN | NaN | NaN | NaN | 5.588102 |

| | | | | | | | | |
|---|---|---|---|---|---|---|---|---|
| 6.96559 | 6.839092 | 6.660607 | 5.975265 | 6.344914 | 7.258544 | 7.471785 | 7.481165 | 6.641689 |
| 6.361775 | 6.715043 | 6.64287 | 6.712914 | 7.27196 | 5.084198 | 5.437459 | 5.474594 | 6.289077 |
| 7.040101 | 7.253047 | 6.694403 | 7.817574 | 6.772997 | 6.240904 | 6.013473 | 5.963169 | 5.981797 |
| 6.655506 | 6.340469 | 6.555634 | 6.53305 | 6.906677 | 6.066887 | 6.389519 | 6.275198 | 6.636127 |
| 7.169628 | 7.23806 | 7.242839 | 6.286777 | 6.472034 | 7.41227 | 7.319518 | 7.596291 | 7.058177 |
| 9.814919 | 9.85449 | 9.689802 | 10.20173 | 9.906729 | 8.027765 | 8.489568 | 7.951878 | 8.228461 |
| 6.572785 | 6.897827 | 6.850301 | 8.279787 | 8.388013 | 6.955025 | 7.0828 | 7.144608 | 6.609513 |
| 9.690489 | 9.90164 | 9.717282 | 8.441013 | 8.605149 | 7.958538 | 7.504521 | 7.694332 | 6.580384 |
| 7.788645 | 8.189691 | 7.592263 | 6.967818 | 7.038338 | 7.998752 | 7.937883 | 7.679873 | 7.947949 |
| 8.440332 | 8.664026 | 8.518267 | 6.892008 | 7.01613 | 7.8965 | 7.995359 | 7.995832 | 8.387256 |
| 7.399788 | 7.622262 | 7.611118 | 8.078043 | 8.172724 | 7.597694 | 7.534559 | 7.50658 | 7.48618 |
| 9.5384 | 9.529272 | 9.612766 | 9.505277 | 9.376091 | 10.08332 | 10.11373 | 10.28208 | 8.997126 |
| 7.033159 | 5.458789 | 5.695856 | NaN | 7.3688 | 7.761994 | 6.892114 | 7.628561 | 6.98228 |
| 8.953336 | 8.839668 | 8.885653 | 8.629735 | 8.756341 | 8.786132 | 8.817558 | 8.772595 | 7.849566 |
| 8.529212 | 8.648066 | 8.371111 | 8.181287 | 8.188113 | 8.790147 | 8.790338 | 8.808203 | 8.817833 |
| 7.90119 | 7.974972 | 8.2866 | 10.2631 | 9.677254 | 7.361511 | 6.978783 | 7.030766 | 6.68198 |
| 9.565875 | 9.631031 | 9.501457 | 10.93176 | 10.9781 | 9.62707 | 10.05474 | 9.149026 | 9.688612 |
| 7.045501 | 7.084555 | 7.13067 | 6.760379 | 6.999676 | 7.163127 | 7.199707 | 7.004963 | 7.224563 |
| 8.916023 | 9.062733 | 9.016778 | 7.07624 | 7.229276 | 9.647429 | 9.573539 | 9.570935 | 9.260863 |
| 7.102818 | 7.175881 | 7.496246 | NaN | NaN | 6.656674 | 6.299213 | 6.036404 | 5.200251 |
| NaN | NaN | 5.603957 | NaN | NaN | 6.289839 | 6.32622 | 6.16631 | 5.589884 |
| 11.23911 | 11.00829 | 11.0349 | 10.44889 | 10.47378 | 11.52066 | 11.40763 | 11.52392 | 10.95765 |
| 9.667619 | 9.900604 | 9.973951 | 13.03264 | 12.94037 | 8.707248 | 8.528382 | 8.592394 | 8.658162 |
| 9.759236 | 9.735634 | 9.810583 | 8.770944 | 8.775382 | 10.02918 | 10.04322 | 10.04685 | 9.989064 |
| 6.330142 | 6.936231 | 7.151772 | 5.590729 | 6.011692 | 5.934568 | 5.700988 | 5.741992 | 5.916433 |
| NaN | NaN | NaN | 7.097824 | 7.515147 | 5.507892 | NaN | NaN | 6.326703 |
| 8.345388 | 8.208645 | 8.224114 | 10.78789 | 10.61431 | 8.822654 | 8.6828 | 8.69141 | 7.767898 |
| 8.101192 | 7.936766 | 7.935085 | 7.616303 | 7.525819 | 7.729889 | 7.761772 | 7.829328 | 7.356898 |
| 11.53683 | 11.37504 | 11.51538 | 10.66933 | 10.76699 | 11.52244 | 11.4604 | 11.50549 | 10.62344 |
| 6.714268 | 7.161538 | 6.918914 | 7.333867 | 7.167557 | 5.653065 | 5.813193 | 5.992084 | 6.565535 |
| 8.981744 | 8.921231 | 9.129182 | 7.963471 | 7.957987 | 8.903828 | 9.050198 | 9.057711 | 9.246012 |
| 7.736183 | 7.511815 | 7.61049 | 7.361563 | 7.431043 | 7.705958 | 7.685683 | 7.542117 | 7.571411 |
| 5.90276 | 6.173956 | 6.010551 | 6.187074 | 6.310753 | NaN | 5.470721 | 5.616583 | 6.325702 |
| NaN | 6.218953 | 5.836599 | 5.813295 | 6.203972 | 7.618201 | 7.70564 | 7.993083 | 6.731601 |
| 9.523989 | 8.946888 | 9.482101 | 8.553412 | 8.343843 | 8.974742 | 8.819833 | 9.229989 | 8.626385 |
| 8.169032 | 8.182113 | 8.233234 | 7.721807 | 7.883298 | 8.039492 | 8.316866 | 8.155601 | 8.328907 |
| 7.957155 | 7.927624 | 7.458596 | 8.279867 | 8.60681 | 6.465765 | 6.757264 | 6.813559 | 6.633875 |
| 8.096046 | 8.515706 | 8.192993 | 7.776433 | 7.613792 | 8.543633 | 8.301358 | 8.287264 | 8.556085 |
| 7.595455 | 7.477306 | 7.752222 | 7.058669 | NaN | 9.556904 | 7.745541 | 6.963394 | 7.291896 |
| 6.422127 | 5.140477 | 6.042971 | 6.820965 | 6.289578 | 7.537626 | 7.508934 | 7.418012 | 6.288994 |
| 5.923749 | 6.129886 | 5.997437 | 5.816539 | 5.398311 | NaN | NaN | NaN | NaN |
| 9.934825 | 9.840217 | 9.888993 | 10.72451 | 10.60898 | 9.812457 | 9.746725 | 9.850733 | 9.59253 |
| 8.451781 | 8.402786 | 8.660458 | 7.756874 | 8.133931 | 8.60536 | 8.503566 | 8.646169 | 8.621323 |
| 4.752946 | 5.163959 | 4.205299 | NaN | 5.357444 | 5.339591 | 6.180807 | 6.278093 | 5.100986 |
| 7.637764 | 8.425437 | 8.233437 | 6.498969 | 6.52217 | 8.354997 | 8.513894 | 8.627922 | 9.31916 |
| 9.111938 | 9.197682 | 9.284293 | 10.99708 | 10.70103 | 9.203212 | 9.24413 | 9.288875 | 9.240401 |
| 7.689076 | 7.55957 | 7.087152 | 7.489479 | 7.094861 | 7.83358 | 7.235835 | 7.195476 | 7.499334 |

| | | | | | | | | |
|---|---|---|---|---|---|---|---|---|
| 7.885387 | 7.643105 | 7.743306 | 8.586782 | 8.53861 | 8.328773 | 8.178992 | 8.249588 | 7.813547 |
| NaN | NaN | NaN | 6.171881 | 6.095112 | NaN | NaN | NaN | 4.779587 |
| 8.005915 | 7.685327 | 8.145216 | 8.079639 | 8.786613 | 7.394443 | 7.96443 | 8.010885 | 7.599079 |
| 9.469013 | 9.352609 | 9.386758 | 9.403766 | 8.935982 | 7.237477 | 6.880578 | 7.590376 | 8.454909 |
| 7.73133 | 8.008848 | 7.845973 | 7.780398 | 8.067221 | 8.488001 | 8.654763 | 8.539187 | 8.5157 |
| 8.801891 | 9.107587 | 8.964313 | 9.093852 | 9.035455 | 7.773693 | 8.120852 | 8.172653 | 9.164231 |
| 7.04909 | 6.809248 | 7.055212 | 5.935308 | 3.934806 | 6.347685 | 6.675804 | 6.775637 | 6.734423 |
| 11.1487 | 11.24314 | 10.90865 | 11.68302 | 11.91398 | 11.24768 | 10.92755 | 11.23963 | 10.89181 |
| 8.363828 | 8.328023 | 8.494215 | 7.457924 | 7.354074 | 7.867676 | 8.132931 | 7.957617 | 8.46574 |
| 9.186214 | 9.318982 | 9.384314 | 9.079751 | 9.084865 | 9.115183 | 9.131395 | 9.131413 | 9.552377 |
| NaN | 6.139159 | NaN | 5.219964 | 4.548272 | NaN | NaN | NaN | 4.677176 |
| 5.760709 | 4.983251 | 5.742543 | 6.215092 | 6.090704 | 6.263137 | NaN | 6.402728 | 6.565151 |
| 8.404373 | 8.466908 | 8.368128 | 8.141121 | 8.45789 | 8.405719 | 8.44495 | 8.367023 | 8.377577 |
| 7.869282 | 8.279531 | 7.425827 | 5.10067 | 5.763956 | 7.381766 | 7.312707 | 7.448023 | 7.138478 |
| 7.120203 | 6.77752 | 7.018003 | 5.421243 | 4.905312 | 7.563951 | 7.337961 | 7.700243 | 6.469768 |
| 6.757919 | 6.721633 | 6.684928 | 5.92169 | 6.335028 | 6.6137 | 6.79815 | 6.425206 | 6.842911 |
| 8.577786 | 8.113717 | 8.467871 | 7.452488 | 7.340971 | 8.199882 | 8.821757 | 8.037798 | 7.852489 |
| 9.614846 | 9.626742 | 9.477674 | 9.396868 | 10.27186 | 9.629238 | 9.405341 | 9.623073 | 9.304157 |
| 7.587533 | 7.905249 | 6.97861 | 9.890596 | 10.58754 | NaN | 7.287885 | 7.808152 | 6.465384 |
| 6.89123 | 6.75141 | 6.402285 | 6.320305 | NaN | NaN | NaN | 3.237898 | 7.556752 |
| 9.518596 | 9.565999 | 9.464605 | 8.474198 | 8.391388 | 9.215635 | 9.303698 | 9.357531 | 9.475817 |
| 9.137047 | 9.068975 | 9.184179 | 8.095836 | 8.087562 | 9.459036 | 9.44923 | 9.555741 | 8.826823 |
| 6.731978 | 5.569603 | 6.302619 | 7.7886 | 8.149233 | 5.984708 | 6.540563 | NaN | 6.925292 |
| 6.531991 | 7.278485 | 7.187374 | 6.557009 | 6.605173 | NaN | 5.832311 | NaN | 6.260598 |
| 8.138734 | 8.134737 | 8.171171 | 7.16981 | 6.895753 | 8.181321 | 8.002969 | 8.225718 | 7.481361 |
| 7.448571 | 7.078541 | 7.626417 | 7.119923 | 7.065333 | 7.32655 | 7.648238 | 7.533187 | 6.59198 |
| 5.988141 | 5.722827 | 5.882483 | NaN | 7.880445 | 5.305933 | 5.340231 | 4.93303 | 4.801303 |
| 7.649046 | 7.841772 | 7.643836 | 5.761697 | 6.157016 | 7.300499 | 7.46794 | 7.381595 | 7.463414 |
| 8.348691 | 10.20434 | 7.773006 | 8.145741 | 8.06683 | 8.684543 | 8.79406 | 8.688402 | 8.810713 |
| NaN | 5.920092 | 6.276018 | NaN | NaN | 5.230584 | 4.621768 | 4.809232 | NaN |
| 6.474629 | 5.942556 | 6.627988 | 6.547032 | 5.973395 | 6.844074 | 6.243998 | 6.23802 | 6.889278 |
| 8.983873 | 8.782006 | 8.926126 | 7.437358 | 7.355573 | 8.60306 | 8.567458 | 8.59936 | 7.222454 |
| 9.518093 | 9.443913 | 9.528152 | 11.47116 | 11.60111 | 8.855262 | 8.992829 | 9.057389 | 9.294301 |
| 8.87267 | 8.70918 | 8.856345 | 9.563959 | 9.501155 | 8.866831 | 8.76484 | 8.890415 | 8.746507 |
| 6.426319 | 6.134548 | 5.83114 | NaN | NaN | 5.922968 | NaN | 5.940132 | 6.539199 |
| 6.847492 | 6.207763 | 6.634712 | 7.137163 | 8.094851 | 7.653951 | 6.8994 | 7.508199 | 6.57788 |
| 5.31964 | 4.891996 | 5.650725 | NaN | NaN | 5.702997 | 5.794733 | 6.011235 | 6.016285 |
| 6.489702 | 6.540274 | NaN | NaN | 5.558469 | 7.13069 | 7.692019 | 7.756591 | 7.199966 |
| 9.028317 | 9.15612 | 9.373774 | 9.767828 | 9.74541 | 7.186121 | 7.226149 | 7.719298 | 8.909476 |
| 9.340885 | 9.071656 | 9.027957 | 9.423088 | 9.413158 | 8.41457 | 8.184713 | 8.332183 | 7.856459 |
| 7.388334 | 7.251832 | 7.243701 | 8.007322 | 7.992384 | 7.365867 | 7.331687 | 7.470605 | 6.619929 |
| 6.434527 | 7.124813 | 6.489601 | 3.643897 | 7.157703 | 7.386519 | 7.378788 | 6.937594 | 7.199666 |
| NaN | NaN | 6.241271 | NaN | NaN | NaN | NaN | NaN | 6.128427 |
| 11.0906 | 11.18548 | 11.13422 | 10.53757 | 10.68659 | 11.02293 | 10.94881 | 10.97339 | 10.48457 |
| 7.825272 | 8.047378 | 7.920467 | 7.525843 | 8.29543 | 7.508316 | 7.498628 | 7.606689 | 6.785115 |
| 6.938537 | 6.784236 | 6.922056 | 5.672567 | 5.959943 | 6.493063 | 6.656224 | 6.537088 | 6.020536 |
| 8.70975 | 7.72982 | 7.51914 | 5.673417 | 6.229998 | NaN | 6.4966 | 8.812182 | 5.288573 |

| | | | | | | | | |
|---|---|---|---|---|---|---|---|---|
| 8.251547 | 7.926766 | 7.848501 | 8.440225 | 7.935869 | 7.264214 | 6.940971 | 6.866848 | 6.893542 |
| 7.842985 | 6.076629 | 7.085457 | 5.227672 | NaN | 5.718085 | 5.592507 | 6.836247 | 5.585515 |
| 6.515606 | 6.099868 | 6.526189 | 7.508663 | 7.53644 | 7.128857 | 7.38942 | 7.000162 | 6.208445 |
| 5.479607 | 4.749067 | 5.380459 | 4.515535 | 4.321888 | 6.302458 | 6.181732 | 6.395123 | NaN |
| 6.484719 | 6.274652 | 6.259493 | NaN | 4.968169 | 5.834531 | 6.516125 | 5.976762 | 5.674399 |
| 6.138181 | 6.378958 | 6.193266 | 6.077796 | NaN | 6.772752 | 6.59014 | 6.218721 | 6.644175 |
| 6.437893 | 6.700948 | 5.605048 | 5.951168 | 5.773133 | 6.555683 | 6.280791 | 6.375983 | 6.361335 |
| 8.423531 | 8.011238 | 8.151597 | 8.665809 | 8.49406 | 8.848031 | 8.284067 | 8.197327 | 7.035978 |
| 10.55901 | 10.55045 | 10.35947 | 8.915875 | 8.947074 | 10.89459 | 10.71745 | 10.65884 | 9.612057 |
| 6.629612 | 6.584389 | 6.768202 | NaN | NaN | NaN | NaN | NaN | 6.119009 |
| 9.58759 | 9.398049 | 9.194352 | 9.344357 | 9.10255 | 8.368286 | 8.340253 | 8.490849 | 8.511283 |
| 9.399263 | 9.361073 | 8.999413 | 9.337502 | 9.339322 | 8.493873 | 8.743335 | 8.888338 | 8.25736 |
| 8.762651 | 8.615883 | 8.823414 | 9.373322 | 9.51939 | 7.912032 | 8.05302 | 8.054485 | 9.513463 |
| 6.864072 | 6.824146 | 7.077957 | 7.518786 | 8.287062 | 4.592624 | 4.43917 | 4.480793 | 6.149938 |
| 11.81296 | 11.7311 | 11.76392 | 12.50658 | 12.29713 | 11.07121 | 10.95759 | 11.14352 | 10.34988 |
| NaN | 7.834695 | 8.075923 | 7.938154 | 7.575006 | 7.269467 | 7.544411 | 7.463249 | 7.698117 |
| 9.456163 | 9.556833 | 9.498162 | 9.111781 | 9.214633 | 9.918243 | 9.566159 | 9.83404 | 9.499172 |
| 8.095654 | 8.217039 | 8.142341 | 6.601973 | 6.639168 | 8.380934 | 8.357403 | 8.307755 | 7.536997 |
| 9.862342 | 10.44786 | 10.19089 | 11.9912 | 11.69393 | 9.981484 | 9.806746 | 9.874972 | 9.228724 |
| 7.151007 | 7.142437 | 6.922836 | 7.117275 | 7.060923 | 6.963369 | 6.946577 | 6.992126 | 6.953603 |
| 7.718309 | 7.868837 | 7.56079 | 7.131553 | 7.012646 | 6.786336 | 6.671275 | 6.990857 | 7.375966 |
| 9.600988 | 9.594684 | 9.691311 | 9.766825 | 9.827724 | 9.281035 | 9.406884 | 9.167682 | 10.14613 |
| 9.257098 | 9.020888 | 8.877999 | 8.338437 | 7.862947 | 9.449723 | 9.518314 | 9.330806 | 8.989486 |
| 7.075464 | 7.279022 | 7.138753 | 7.286396 | 6.796781 | 7.405339 | 7.493979 | 7.203467 | 6.478902 |
| 7.375793 | 6.867395 | 7.096887 | 6.939528 | 6.519688 | 7.023575 | 6.597004 | 6.27264 | 6.84488 |
| 6.84042 | 7.030519 | 6.70846 | 7.78047 | 7.404854 | 8.51834 | 8.74953 | 8.549397 | 5.76505 |
| 7.187863 | 7.468347 | 7.452284 | 8.647503 | 8.643415 | 7.080666 | 6.66823 | 6.829749 | 6.833576 |
| 6.998752 | 5.345237 | 6.665226 | 5.947182 | 5.685123 | 6.93018 | 6.682683 | 6.789107 | 6.655212 |
| 7.626984 | 6.897254 | 8.09144 | 6.735537 | 6.422295 | 7.593484 | 7.832321 | 7.8014 | 8.049609 |
| 7.230417 | 7.034202 | 6.879858 | 10.22197 | 10.4019 | 6.468227 | 6.145037 | 6.111866 | 7.241802 |
| 5.997058 | 6.607309 | 6.813602 | NaN | 5.182297 | 6.609357 | 6.757397 | 6.217639 | 7.003688 |
| 7.885794 | 7.983588 | 7.918658 | 7.506237 | 7.61615 | 8.202373 | 8.293119 | 8.324375 | 8.30475 |
| 11.88521 | 11.69164 | 11.73956 | 10.62791 | 10.32312 | 8.255197 | 8.716126 | 8.429734 | 9.417737 |
| 10.43236 | 10.3688 | 10.44963 | 11.32239 | 11.4698 | 10.56611 | 10.6359 | 10.82159 | 9.872927 |
| 7.856515 | 8.268974 | 8.152397 | 7.291278 | 7.100889 | 7.475112 | 7.514718 | NaN | 7.203908 |
| 8.299855 | 8.411553 | 8.494667 | 8.26061 | 7.986873 | 8.298551 | 8.541747 | 8.613232 | 8.313477 |
| 8.020131 | 7.728873 | 8.059852 | 6.632086 | 7.791059 | 8.644651 | 9.249961 | 9.256517 | 6.420628 |
| 8.77676 | 8.871757 | 8.603618 | 7.415263 | 7.304022 | 8.489265 | 8.362002 | 8.338221 | 7.990015 |
| NaN | 7.393001 | 6.606795 | NaN | 8.269514 | 8.410044 | 8.636927 | 8.357306 | 7.798303 |
| 8.739041 | 9.157654 | 8.963061 | 7.70986 | 8.511215 | 8.543923 | 9.006052 | 8.740465 | 8.500096 |
| 12.90702 | 12.25094 | 12.57236 | NaN | 9.702751 | NaN | 8.516226 | 8.640821 | 7.944493 |
| 7.730378 | 7.596169 | 7.394596 | 9.540344 | 9.452586 | 6.196369 | 6.614296 | 7.130724 | 6.581611 |
| 7.253133 | 7.163825 | 7.253434 | 6.894371 | 7.13956 | 7.712906 | 7.652491 | 7.597126 | 6.938209 |
| 6.336933 | 6.878471 | 6.337183 | 7.60135 | 7.77291 | 7.135331 | 7.083467 | 6.84266 | 7.679948 |
| 4.928021 | NaN | NaN | 4.641875 | 6.236052 | 5.946422 | 5.723266 | 5.813718 | 5.296773 |
| 9.663579 | 9.342555 | 9.662874 | 8.866194 | 8.759144 | 9.005705 | 9.019664 | 8.781742 | 8.647858 |
| 6.484625 | 6.791245 | 6.56765 | 4.870028 | 4.432813 | 6.177335 | 6.198674 | 6.439869 | 6.928442 |

| | | | | | | | | |
|---|---|---|---|---|---|---|---|---|
| 8.323041 | 8.270594 | 8.104058 | 6.988198 | 7.178995 | 8.924261 | 8.771916 | 8.837714 | 7.51082 |
| 6.647498 | 6.862434 | 6.383145 | 5.125221 | NaN | 6.641445 | 6.363072 | 6.401862 | 6.410799 |
| 6.749126 | 6.409953 | 6.872764 | 7.023287 | 6.695418 | 7.078076 | 7.051923 | 7.210544 | 6.702776 |
| 5.442465 | 5.655332 | 5.685966 | 6.855439 | 6.78208 | 5.694699 | 6.019297 | 5.712903 | 6.771784 |
| NaN | 5.961306 | 5.360054 | 4.296609 | NaN | 6.487731 | 5.834243 | NaN | 7.054831 |
| 8.103488 | 7.913911 | 8.050063 | 6.997558 | 7.020769 | 8.422871 | 8.156834 | 8.44726 | 6.927665 |
| 3.319994 | 4.490288 | NaN | 3.479392 | NaN | 3.579496 | 4.751343 | 4.745749 | 4.509151 |
| 7.909441 | 7.867192 | 7.945315 | 6.680059 | 7.051064 | 7.472061 | 7.662911 | 7.826607 | 7.998264 |
| 8.694349 | 7.640903 | 7.886171 | 8.22905 | 8.569914 | 7.257466 | 7.168008 | 7.28512 | 8.414831 |
| 7.622632 | 7.56234 | 7.670341 | 6.803294 | 6.949062 | 7.367245 | 7.333001 | 7.073342 | 7.080473 |
| 6.973933 | 7.243937 | 6.951605 | 5.557233 | 5.751385 | 7.539994 | 7.687298 | 6.971598 | 6.574959 |
| 7.587522 | 7.506057 | 7.379113 | 7.755624 | 7.403716 | 7.915201 | 7.717136 | 7.512052 | 7.601303 |
| NaN | NaN | NaN | 5.443836 | 6.010247 | NaN | NaN | NaN | NaN |
| 7.747931 | 7.002341 | 7.744673 | 7.418697 | 6.073811 | 8.154116 | 8.146456 | 8.20455 | 7.813798 |
| 8.603443 | 8.598864 | 8.739603 | 8.438224 | 8.421583 | 9.180538 | 9.00454 | 9.258509 | 8.629284 |
| 10.16372 | 10.25957 | 10.28378 | 9.223733 | 9.535547 | 9.844028 | 10.1892 | 9.958251 | 10.0525 |
| 8.535405 | 8.467892 | 8.586277 | 7.272354 | 7.353144 | 8.48125 | 8.517128 | 8.645039 | 8.419282 |
| NaN | NaN | NaN | NaN | NaN | NaN | 4.540272 | NaN | 5.371872 |
| 6.63708 | 6.662541 | 6.703419 | 5.796405 | 5.351586 | 7.005571 | 6.967793 | 7.180417 | 5.730371 |
| 7.615656 | 7.376201 | 7.432539 | 6.978557 | 7.13401 | 7.131159 | 7.019264 | 6.9255 | 7.025722 |
| 5.923471 | 6.051338 | 6.155203 | 5.725385 | 5.529976 | 6.346488 | 6.616669 | 6.579084 | 6.542675 |
| 7.813446 | 7.381269 | 7.738644 | 5.449342 | 5.679656 | 8.409664 | 8.041497 | 7.855278 | 7.175709 |
| 6.354455 | 6.180418 | 6.432836 | 5.728771 | 5.213766 | 6.505724 | 7.383634 | 6.838349 | 6.614707 |
| 6.80535 | 7.37216 | 6.177644 | 6.478577 | NaN | 6.423819 | 5.973268 | 6.317538 | 6.107892 |
| 7.223912 | 7.452817 | 7.42963 | 7.448612 | 7.736023 | 7.319704 | 7.268362 | 7.240339 | 6.973907 |
| 7.153919 | 6.725169 | 6.39185 | NaN | NaN | 6.921116 | 6.668846 | 7.031324 | 6.844548 |
| 6.136525 | 6.843763 | 5.992417 | 6.932662 | 6.437511 | 5.337069 | 6.479054 | 5.855955 | 5.710291 |
| 6.423654 | 6.682601 | 6.226434 | NaN | 5.167742 | 6.58929 | 6.450782 | 6.454795 | 6.117134 |
| 6.036833 | 6.272689 | 6.09577 | 5.509375 | 4.039327 | 5.29744 | 5.00238 | 6.48026 | 5.510886 |
| 11.61393 | 11.40309 | 11.34664 | 11.11477 | 10.58731 | 10.66534 | 11.12859 | 10.64005 | 10.9089 |
| 6.167858 | 5.813742 | 5.933522 | 5.270391 | 5.355699 | 5.985578 | 6.095713 | 5.107249 | 5.869627 |
| 8.761454 | 9.005684 | 8.715923 | 8.004043 | 8.086814 | 8.373644 | 8.714216 | 8.545857 | 8.371735 |
| 6.925093 | 6.76244 | 6.960634 | 5.813243 | 5.731686 | 7.0268 | 6.963927 | 6.984895 | 6.806082 |
| 8.139815 | 8.425377 | 8.431144 | 7.871129 | 8.148698 | 8.498038 | 8.44355 | 8.610846 | 8.863948 |
| 8.371051 | 8.45286 | 7.823117 | 6.796883 | 6.671072 | 7.231368 | 7.246191 | 7.292575 | 6.538936 |
| 5.928936 | 5.626614 | 5.869837 | 5.614484 | 5.979803 | 5.728832 | 5.932272 | 5.153175 | 6.428549 |
| 6.954982 | 7.174757 | 7.456784 | 7.84272 | 8.17228 | 6.311279 | 6.267584 | 7.270149 | 6.61371 |
| 7.531203 | 7.030921 | 8.078958 | NaN | NaN | 7.131202 | 7.466031 | 7.415254 | 6.966319 |
| 7.694299 | 7.760842 | 7.675709 | 6.505524 | 7.392498 | 6.785299 | 6.445619 | 6.89945 | 6.922325 |
| 6.138116 | 6.368375 | 5.647192 | 6.966708 | 7.115454 | 6.755013 | 6.437409 | 6.539029 | 7.034763 |
| 8.318439 | 8.337796 | 8.583097 | 7.292752 | 7.760215 | 7.130479 | 7.212662 | 7.078957 | 7.275303 |
| 7.641152 | 7.881426 | 8.021029 | 8.058933 | 7.706694 | 6.978668 | 6.881713 | 6.910075 | 7.195168 |
| 8.222351 | 8.289742 | 8.049024 | 11.56147 | 11.46953 | 8.944471 | 8.832528 | 8.989104 | 8.416203 |
| 8.050695 | 8.156234 | 8.205467 | 7.238432 | 7.723246 | 8.142344 | 8.019008 | 8.043568 | 7.38177 |
| 5.954317 | 6.878442 | 6.158156 | 5.726565 | 6.400678 | 6.528491 | 6.526012 | 6.705433 | 5.929705 |
| 7.030551 | 6.962203 | 6.977226 | 5.845704 | 6.810376 | 6.893306 | 7.065463 | 7.248917 | 7.276774 |
| 6.54372 | 6.336555 | 6.040146 | 6.331292 | NaN | 6.714138 | 6.518975 | 6.661542 | 6.214509 |

-Bruker Confidential-

| | | | | | | | | |
|---|---|---|---|---|---|---|---|---|
| 7.482808 | 7.862016 | 7.835927 | 7.087742 | 6.799334 | 6.65689 | 7.267755 | 7.007829 | 5.347023 |
| 7.24297 | 6.581957 | 6.812533 | 8.940777 | 9.091478 | 7.539925 | 7.251772 | 7.436368 | 5.903069 |
| 8.029413 | 8.881735 | 8.641641 | 8.557274 | 8.304725 | 7.751038 | 7.321565 | 7.137368 | 6.540678 |
| 7.191884 | 7.291922 | 7.108519 | 7.10654 | 7.651241 | 8.229404 | 8.030886 | 7.836932 | 6.697845 |
| 7.872857 | 7.521945 | 7.409325 | 8.333467 | 7.929975 | 7.183907 | 6.958786 | 6.941753 | NaN |
| 9.909635 | 9.650653 | 9.617116 | 8.265173 | 8.007375 | 9.404613 | 8.899924 | 9.68262 | 8.494655 |
| 9.037601 | 8.92252 | 9.192587 | 8.133261 | 8.380937 | 9.063311 | 9.152033 | 9.335921 | 8.022662 |
| 6.397253 | 6.768154 | 6.734287 | 8.540332 | 8.432036 | 5.305269 | 5.714678 | 6.015827 | 6.564113 |
| 7.086038 | 7.554984 | 7.423389 | 9.363187 | 9.354781 | 9.252664 | 9.143589 | 9.1864 | 8.71081 |
| 9.121256 | 8.699119 | 8.840126 | 8.166361 | 7.979143 | 8.395025 | 8.761229 | 8.608096 | 8.243113 |
| 8.304237 | 8.381017 | 8.562653 | 7.025243 | 7.457109 | 8.186226 | 8.072318 | 8.066146 | 8.199076 |
| 5.534391 | 5.228401 | 5.52986 | NaN | 6.789427 | 5.447736 | 5.340825 | 5.543578 | 5.670128 |
| 7.164126 | 6.464358 | 6.832382 | 6.071739 | 6.2094 | 6.868838 | 6.356512 | 6.546196 | 6.428061 |
| 7.482999 | 7.37215 | 7.465537 | 6.831748 | 6.884519 | 7.765736 | 7.824509 | 7.746293 | 7.09849 |
| 7.786079 | 7.231588 | 7.621177 | 7.696149 | 7.915088 | 7.921434 | 7.839957 | 7.712918 | 7.55324 |
| 7.240767 | 7.515373 | 7.493585 | 6.546213 | 5.846374 | 6.907382 | 6.821906 | 7.082742 | 7.025214 |
| 6.782429 | 6.747812 | 6.793708 | 6.142385 | 6.232562 | 7.07266 | 7.005351 | 7.123659 | 7.057944 |
| 3.898278 | 4.540981 | 4.083236 | 4.705938 | 4.309101 | 4.945996 | 6.016183 | 6.191051 | 6.011338 |
| NaN | 6.445076 | 6.024817 | 5.812269 | 5.3444 | NaN | 7.568301 | 7.403765 | 6.394629 |
| 9.06758 | 8.905844 | 8.939363 | 8.703594 | 8.821225 | 8.684526 | 8.657676 | 8.752147 | 8.129268 |
| 9.449441 | 9.406351 | 9.334116 | 10.23162 | 10.56361 | 9.348845 | 9.513926 | 9.51942 | 8.804069 |
| 9.569328 | 9.401186 | 9.317281 | 7.902033 | 7.605807 | 8.168062 | 8.798902 | 8.977636 | 8.225407 |
| 8.071342 | 8.270151 | 7.970953 | 7.658916 | 8.349841 | 7.920762 | 7.748786 | 7.547546 | 7.425014 |
| 8.082515 | 7.912359 | 7.979739 | 6.907358 | 7.532924 | 8.183022 | 8.251611 | 8.101785 | 8.239178 |
| 6.831719 | 6.551315 | 6.763031 | 7.092192 | 6.644623 | 7.106533 | 6.762878 | 6.644924 | 6.590507 |
| 8.317477 | 8.121527 | 8.377283 | 7.650176 | 7.642199 | 8.095114 | 7.839169 | 7.662802 | 7.242826 |
| 9.354495 | 9.888344 | 9.919528 | 10.66816 | 10.67365 | 10.16307 | 10.19107 | 10.0409 | 9.704311 |
| 7.505133 | 7.247198 | 7.199909 | 6.230968 | 6.274983 | 6.82115 | 6.868756 | 7.125115 | 6.835857 |
| 7.048388 | 6.883373 | 6.877746 | 5.351521 | 5.978329 | 7.045521 | 6.969163 | 7.009453 | 7.035524 |
| 6.25689 | 6.597773 | 5.809865 | 6.910726 | 7.044337 | 6.094241 | 5.814854 | 5.923335 | 6.968177 |
| 6.38187 | 5.775775 | 6.471449 | 5.958138 | 5.654198 | 6.578557 | 6.548979 | NaN | 6.455909 |
| 8.10242 | 7.861389 | 7.939638 | 7.22598 | 7.464701 | 8.387565 | 8.383342 | 8.410555 | 7.658719 |
| 9.156908 | 9.007039 | 8.992434 | 7.772529 | 7.99226 | 9.254595 | 9.423637 | 9.295738 | 9.015132 |
| 7.412501 | 6.849687 | 7.21349 | 5.493289 | 5.600576 | 6.481812 | 6.67256 | 6.628454 | 6.899151 |
| 6.948198 | 7.159338 | 7.311204 | 7.619965 | 7.816145 | 7.074944 | 6.925023 | 7.11123 | 6.305145 |
| 7.168009 | 6.90734 | 6.788399 | 6.420139 | 6.45616 | 7.037671 | 6.168923 | 7.142409 | 6.732666 |
| 8.502142 | 8.371569 | 7.998491 | 8.094258 | 7.864099 | 8.840949 | 8.78078 | 8.768692 | 8.738041 |
| 6.296049 | 6.770923 | 5.699965 | NaN | 6.480217 | 7.081018 | 6.753611 | 6.92801 | 6.545245 |
| 8.042948 | 8.312255 | 8.505883 | 8.881473 | 8.963067 | 7.234666 | 7.892597 | 7.531984 | 7.805099 |
| 6.667632 | 6.334263 | 6.524559 | 6.855423 | 6.754288 | 5.88396 | 6.161503 | 5.909753 | 5.85642 |
| 5.17036 | 3.492441 | 4.345669 | 5.506951 | 4.264624 | NaN | 5.652365 | 4.878773 | 5.811657 |
| 8.548185 | 8.840492 | 8.70864 | 9.19749 | 9.010839 | 8.815025 | 8.864129 | 8.797789 | 9.255781 |
| 6.256321 | 5.840442 | 5.61343 | NaN | 6.209911 | 6.779029 | 6.212786 | 6.644565 | 6.099339 |
| 7.322727 | 7.273821 | 6.811836 | 6.517014 | 6.837529 | 7.382638 | 7.214299 | 6.767131 | 7.610898 |
| 6.173972 | 5.954833 | 6.568829 | 6.534041 | 6.5988 | 6.469884 | 6.619163 | 6.216105 | 6.687605 |
| 8.617328 | 7.834616 | 8.892177 | 7.074861 | 6.592116 | 8.225525 | 8.346987 | 8.073689 | 8.372836 |
| 7.30784 | 6.76981 | 7.511731 | 6.438698 | 6.297559 | 8.288235 | 7.857902 | 8.130555 | 7.143946 |

-Bruker Confidential-

| | | | | | | | | |
|---|---|---|---|---|---|---|---|---|
| 7.164003 | 7.106818 | 7.287106 | 6.671887 | 6.548581 | 7.348206 | 7.370576 | 7.561687 | 7.205197 |
| 6.854709 | 6.888572 | 6.580628 | 6.080715 | 6.000957 | 7.184224 | 6.85038 | 7.231229 | 7.202918 |
| 6.877175 | 6.87171 | 6.780338 | 7.103594 | 7.253286 | 6.270056 | 6.722139 | 6.477464 | 6.517429 |
| 7.997742 | 7.857724 | 7.972776 | 5.750595 | 5.933242 | 8.10726 | 8.013105 | 8.176404 | 7.776831 |
| 6.023992 | 6.056811 | 6.233075 | 6.875085 | 6.936508 | 7.801618 | 7.535585 | 7.468224 | 5.901435 |
| 5.800092 | NaN | 6.080741 | NaN | 5.762508 | 4.758706 | 5.283749 | 5.420496 | 5.510602 |
| 9.089125 | 9.067664 | 8.946628 | 9.921031 | 10.0654 | 9.455502 | 9.32731 | 9.497567 | 7.808418 |
| 7.825333 | 7.332884 | 7.327813 | 7.35851 | 7.216657 | 7.039137 | 6.737213 | 6.806777 | 6.318674 |
| 9.011252 | 8.965169 | 9.206604 | 7.582007 | 7.629437 | 8.942796 | 9.18075 | 8.932539 | 8.379026 |
| 7.578153 | 7.027991 | 4.804576 | NaN | NaN | 7.292562 | NaN | 6.930006 | 6.814714 |
| NaN | 6.451301 | 7.513288 | NaN | 5.516146 | 8.219025 | 7.381346 | 7.58721 | 6.601383 |
| 7.084782 | 6.514373 | 7.547093 | 6.431439 | 6.790946 | 6.398212 | 6.7616 | 6.451024 | 6.000843 |
| 6.890793 | 5.790043 | 6.083769 | 5.342822 | NaN | 6.767907 | 6.722226 | 6.584522 | 5.200947 |
| 7.747019 | 7.654161 | 7.832984 | 7.826988 | 8.12555 | 7.386635 | 7.488778 | 7.495549 | 7.433843 |
| 8.247865 | 8.209155 | 8.209779 | 7.043434 | 6.808526 | 8.881315 | 9.171774 | 9.267734 | 4.314405 |
| 8.032756 | 7.733735 | 7.753773 | 9.247419 | 8.938772 | 8.829523 | 8.591143 | 8.960982 | 7.620784 |
| 8.704877 | 8.693081 | 8.564658 | 7.473866 | 7.762296 | 8.529028 | 8.46676 | 8.511875 | 7.971369 |
| 8.435233 | 7.888252 | 8.24688 | 6.073877 | 6.005199 | 8.605227 | 8.648121 | 8.674277 | 7.757061 |
| 8.696463 | 8.563152 | 8.913583 | 8.030623 | 7.440584 | NaN | NaN | NaN | 7.098239 |
| 7.681646 | 7.62557 | 7.653841 | 8.170733 | 8.3797 | 7.466978 | 7.320497 | 7.232056 | 7.974234 |
| 7.793143 | 7.205634 | 6.988454 | 7.014693 | 6.380538 | 8.925595 | 7.622401 | 8.570745 | 5.752643 |
| 8.760242 | 9.175983 | 8.611345 | 8.102336 | 8.468997 | 9.0963 | 8.99941 | 9.079191 | 8.190404 |
| 6.503865 | 5.354848 | 7.33496 | NaN | 6.736242 | 7.051478 | 6.983876 | 7.084634 | 6.220649 |
| 9.339001 | 9.128247 | 9.13379 | 8.16524 | 8.397304 | 8.874484 | 8.598836 | 8.642883 | 8.289067 |
| 9.526765 | 9.458752 | 9.523896 | 10.27556 | 10.0274 | 8.91567 | 8.920439 | 8.398456 | 9.740659 |
| 5.968729 | 6.248017 | 6.224939 | 1.88726 | NaN | NaN | NaN | NaN | NaN |
| 9.227752 | 9.343875 | 9.170433 | 9.714361 | 9.418303 | 8.106202 | 8.25174 | 7.979481 | 9.774778 |
| 6.358586 | 5.192999 | 5.560606 | 5.971226 | 5.765857 | 6.437112 | 5.797611 | 5.946818 | 5.428474 |
| 9.95803 | 9.849579 | 9.914308 | 9.32043 | 9.140446 | 9.70932 | 9.698751 | 9.632696 | 9.758529 |
| 7.794014 | 7.935533 | 8.03787 | 8.009625 | 7.990998 | 8.322168 | 8.084303 | 8.083621 | 7.86659 |
| 8.525737 | 9.05462 | 9.159976 | 7.374631 | 6.994736 | 8.239912 | 8.581358 | 8.303854 | 9.163574 |
| 10.32523 | 10.19183 | 10.24335 | 10.30545 | 9.579655 | 8.602402 | 8.893718 | 8.92847 | 9.06316 |
| 10.44477 | 10.52947 | 10.39342 | 9.978292 | 9.929877 | 10.35125 | 10.54083 | 10.51804 | 10.30409 |
| 7.282767 | 7.471337 | 7.164167 | 6.934252 | 6.889979 | 6.406262 | 6.427372 | 6.540401 | 5.92516 |
| 7.990705 | 7.949953 | 7.981475 | 9.400295 | 9.556209 | 7.816865 | 7.740805 | 7.665312 | 8.017581 |
| 5.944124 | 5.555583 | 6.265129 | NaN | 4.594882 | 4.683961 | 3.762922 | 4.951827 | 5.965498 |
| 6.002975 | 6.771133 | 7.296597 | 7.631547 | 7.375527 | 7.262975 | 7.142854 | 7.043371 | 7.268233 |
| 6.300341 | 6.633496 | 6.211063 | 9.296559 | 9.231904 | 6.189114 | 4.049034 | 5.03438 | 5.802849 |
| 8.462113 | 8.568884 | 8.633302 | 7.559909 | 7.831846 | 8.424833 | 8.507088 | 8.484633 | 8.314291 |
| 6.44652 | 6.334208 | 7.010931 | 4.968605 | 5.517654 | 6.371998 | 6.144554 | 6.133187 | 6.228493 |
| 7.415513 | 7.664995 | 7.285824 | 8.05334 | 7.939567 | 7.986014 | 8.284009 | 8.159427 | 6.858346 |
| 9.215572 | 9.212272 | 9.142522 | 9.094023 | 9.22541 | 9.117362 | 9.131785 | 9.066319 | 8.252675 |
| NaN | 6.519689 | NaN | NaN | 7.303673 | 6.980152 | 7.020661 | 6.790906 | 6.38606 |
| 7.504856 | 6.64661 | 7.184393 | 6.924862 | 6.902619 | 7.047707 | 7.101527 | 7.128782 | 6.179972 |
| 6.983768 | 7.185389 | 6.934027 | 7.043135 | 7.359726 | 7.444739 | 7.304492 | 7.66528 | 7.006587 |
| 7.572522 | 7.572 | 8.162894 | 6.008441 | 6.657115 | 6.928737 | 6.908883 | 6.889617 | 7.168697 |
| 10.65053 | 10.73768 | 10.55161 | 11.7969 | 11.6126 | 11.32212 | 11.30041 | 11.32091 | 11.32082 |

| | | | | | | | | |
|---|---|---|---|---|---|---|---|---|
| 7.721436 | 7.029377 | 7.175174 | 5.90125 | 5.409814 | 7.343591 | 7.209107 | 7.112849 | 6.805333 |
| 8.554854 | 8.44201 | 8.601832 | 9.994051 | 10.40228 | 7.793093 | 8.041286 | 8.043672 | 8.762807 |
| 7.603444 | 7.746906 | 7.511352 | 6.586698 | 6.968771 | 7.787309 | 7.490666 | 7.232337 | 6.726023 |
| 8.295001 | 7.507944 | 7.588332 | 8.138491 | 7.425224 | NaN | 6.381886 | 6.684712 | 7.955146 |
| NaN | 5.20442 | NaN | 3.92365 | 4.308583 | 4.371575 | 4.282228 | NaN | 5.061143 |
| 7.342213 | 7.24625 | 7.515196 | 7.852305 | 7.756207 | 7.805059 | 7.761562 | 7.910069 | 7.856768 |
| 7.152138 | 6.492972 | 6.609397 | NaN | NaN | 6.778124 | 6.68816 | 6.842824 | 6.478603 |
| 8.874569 | 8.765192 | 8.788474 | 8.504536 | 8.339572 | 8.110686 | 8.578566 | 8.476405 | 7.748675 |
| 3.82142 | 4.666782 | 4.95317 | 5.428327 | 5.813602 | 5.834082 | 5.522004 | 5.947849 | 4.972641 |
| 5.875811 | NaN | 4.957537 | 6.376235 | 6.800641 | NaN | 4.882606 | 5.272635 | 4.604528 |
| 9.158855 | 9.303581 | 9.48767 | 9.789513 | 9.498473 | 8.957424 | 8.957397 | 8.918378 | 8.956714 |
| 6.987901 | 7.714194 | 7.490828 | 6.463212 | 5.935915 | 7.597402 | 7.446976 | 7.578276 | 6.203095 |
| 5.939283 | 6.298744 | 6.046415 | 6.820775 | 6.960352 | 6.715195 | 7.146539 | 6.781416 | 6.159211 |
| 6.832589 | 6.874688 | 6.97539 | 8.596967 | 8.710341 | 8.046778 | 8.117224 | 8.016787 | 7.214127 |
| 9.033116 | 8.886375 | 9.08649 | 8.78534 | 8.393497 | 8.784953 | 8.825014 | 8.668892 | 8.338685 |
| 8.858134 | 8.274702 | 8.344957 | 7.285037 | 7.283603 | 7.862839 | 7.790857 | 7.791801 | 8.000288 |
| 7.091884 | 7.641047 | 7.44871 | NaN | 7.277836 | 7.933953 | 6.830379 | 7.176488 | 6.999411 |
| 8.768248 | 8.922581 | 8.885123 | 8.658621 | 8.592491 | 8.442223 | 8.789923 | 8.788599 | 9.58279 |
| 5.922849 | 5.741964 | 5.879509 | 4.979613 | 4.855567 | 6.018182 | 5.635211 | 5.785394 | 5.672129 |
| 8.664795 | 5.930335 | 8.3501 | 6.81144 | 8.281272 | 7.062199 | 7.253301 | 7.254018 | 6.649912 |
| 9.890727 | 10.13075 | 10.02791 | 11.97616 | 11.8764 | 9.734956 | 9.390227 | 9.777426 | 9.685144 |
| 9.077434 | 9.012635 | 8.925727 | 8.312585 | 8.063573 | 8.540668 | 8.483082 | 8.856465 | 9.501165 |
| 9.369221 | 9.230097 | 9.23391 | 6.142813 | 5.997211 | 7.078492 | 6.760182 | 7.276072 | 6.766481 |
| 10.42223 | 10.48995 | 10.46841 | 9.751446 | 9.504218 | 10.68035 | 10.78401 | 10.82782 | 9.662082 |
| 9.142329 | 9.220921 | 9.372596 | 9.326249 | 9.160483 | 7.916233 | 8.395861 | 7.556941 | 8.324209 |
| 9.795294 | 9.764259 | 9.93378 | 8.400613 | 8.095073 | 10.08116 | 10.09126 | 10.03075 | 9.552328 |
| 9.137506 | 9.068853 | 9.034255 | 11.0752 | 11.06204 | 9.617691 | 9.436299 | 9.601836 | 8.798822 |
| 6.464217 | 6.264707 | 6.565188 | 5.949011 | 5.207077 | 5.799681 | 6.13859 | 5.867593 | 7.17068 |
| 7.281773 | 6.868732 | 7.316101 | 5.846734 | 5.727226 | 7.404193 | 7.248902 | 6.793671 | 6.848069 |
| NaN | 5.596416 | 5.074049 | NaN | NaN | 5.316901 | 6.445271 | 6.233246 | 6.182415 |
| 9.004121 | 8.933826 | 9.153809 | 12.41573 | 12.58946 | 9.380934 | 9.172055 | 9.340733 | 9.354368 |
| 8.647076 | 8.504395 | 8.122654 | 6.824197 | 6.49518 | NaN | NaN | 6.430782 | 6.190522 |
| 6.712778 | 5.564734 | 6.074995 | NaN | NaN | 5.725878 | NaN | 5.21914 | 5.360779 |
| 9.111043 | 8.986567 | 8.348989 | 8.74186 | 8.793726 | 7.915749 | 7.749706 | 7.561323 | 7.356934 |
| 7.806782 | 7.77821 | 7.702177 | 7.861709 | 8.571823 | 7.584453 | 7.309675 | 7.067107 | 6.954982 |
| 9.945389 | 9.868785 | 9.92155 | 10.41531 | 10.25151 | 10.0712 | 10.12245 | 10.12094 | 8.616062 |
| 5.26124 | 5.128574 | 5.56598 | 5.950977 | 6.224567 | 5.885702 | 5.544676 | 5.683122 | 4.76252 |
| 6.03346 | 5.632096 | 6.519619 | 5.815096 | 5.424768 | 8.264249 | 7.762733 | 8.044253 | 7.965206 |
| 11.84581 | 11.82377 | 11.87378 | 11.88031 | 11.91729 | 12.31195 | 12.36446 | 12.49344 | 12.10781 |
| 10.86775 | 10.69045 | 10.60697 | 11.92981 | 11.50966 | 10.78112 | 10.98911 | 10.731 | 10.26851 |
| 10.31359 | 10.43873 | 10.48069 | NaN | 6.604226 | NaN | NaN | NaN | NaN |
| NaN | 6.516035 | 6.963291 | NaN | NaN | NaN | NaN | NaN | 7.849607 |
| NaN | 6.537014 | 6.393386 | NaN | NaN | NaN | NaN | 6.828844 | 7.020154 |
| 6.787905 | 6.108449 | 7.63166 | 8.153566 | 8.20224 | 7.871771 | 7.502769 | NaN | 5.946606 |
| 6.607805 | 10.69785 | 9.42051 | 8.341436 | NaN | 8.033015 | 7.24043 | 7.586029 | 8.271518 |
| NaN | NaN | NaN | NaN | NaN | NaN | NaN | NaN | 6.375514 |
| 7.863199 | 7.595654 | 7.633526 | 11.03734 | 11.14282 | 7.978378 | 7.848037 | 7.711848 | 10.32244 |

| | | | | | | | | |
|---|---|---|---|---|---|---|---|---|
| 7.825033 | 6.668499 | 7.688408 | 6.50541 | 6.698157 | 7.293293 | 7.569884 | 7.280914 | 8.442175 |
| 12.53703 | 12.4586 | 12.51606 | 9.387034 | 9.631047 | 7.285201 | 7.637268 | 7.724103 | 7.235918 |
| NaN | NaN | 2.886288 | 6.365693 | 6.941501 | 7.293132 | 7.680526 | 7.663083 | 3.768558 |
| 7.423914 | 6.147497 | 6.050195 | 6.148581 | 6.300971 | NaN | 6.618183 | 7.014256 | 6.971945 |
| 7.747295 | 6.208861 | 6.150802 | 5.8746 | 6.043086 | 0.661572 | NaN | NaN | 3.226282 |
| 7.096041 | 7.352876 | 6.739733 | 7.479627 | 7.725956 | 7.50888 | 7.818297 | 7.799349 | 7.91275 |
| 10.16342 | 9.860325 | 9.881276 | 8.259261 | 8.478981 | 7.056872 | 7.400212 | 7.406251 | 7.825415 |
| 8.605533 | 8.345137 | 8.579447 | 9.17593 | 8.987415 | 8.321565 | 8.28366 | 8.492193 | 7.960541 |
| 8.090058 | 8.246405 | 8.31396 | 7.465554 | 7.675846 | 8.798159 | 8.287372 | 8.600526 | 7.472056 |
| 9.474364 | 9.22887 | 9.453189 | 7.874199 | 7.321987 | 7.027747 | 6.850202 | 7.294644 | 7.026714 |
| 5.672179 | 5.337652 | 5.34295 | NaN | NaN | 5.755164 | 5.199535 | 5.665158 | 5.432471 |
| 6.280112 | 5.776659 | 6.725995 | 6.368714 | 6.419957 | 6.02664 | 5.92724 | 6.145957 | 4.189035 |
| 6.929572 | 6.639369 | 6.144923 | 7.703908 | 7.345374 | NaN | NaN | NaN | 6.568161 |
| 10.95638 | 9.609069 | 10.33016 | 7.561562 | 7.244236 | NaN | 6.063486 | 6.950103 | 7.705287 |
| 5.25395 | 4.615067 | 6.406353 | 3.919855 | 5.593739 | NaN | NaN | NaN | 5.617336 |
| 8.39395 | 8.316637 | 8.473151 | 7.001904 | 7.012367 | 5.782703 | 5.47906 | 5.631908 | 8.390994 |
| 9.292988 | 8.504006 | 8.950186 | 6.85452 | 7.779214 | 5.431376 | 6.726448 | NaN | 9.949241 |
| 6.571375 | 7.013899 | 4.03041 | 4.589726 | 2.817209 | NaN | NaN | 5.94877 | 8.28738 |
| 3.247257 | 4.438218 | 4.738501 | NaN | NaN | NaN | NaN | NaN | 5.765447 |
| 7.377692 | 6.895674 | 6.507944 | 6.31346 | 6.392523 | NaN | NaN | NaN | 7.557742 |
| 8.746318 | 8.064652 | 7.583323 | 8.01099 | 7.563351 | NaN | NaN | 7.583891 | 9.400825 |
| 7.673221 | 7.701374 | 8.042104 | 6.602595 | 6.21806 | 8.021379 | 7.549493 | 7.912182 | 7.435496 |
| 7.954387 | 8.084354 | 7.801627 | NaN | 7.008232 | NaN | NaN | NaN | 7.070171 |
| 9.586526 | 9.234041 | 9.859199 | 9.282486 | 9.301689 | 7.605572 | 7.823696 | 7.452829 | 7.441695 |
| 8.854155 | 8.274405 | 8.7757 | 7.837751 | 7.028724 | 7.671779 | NaN | 7.683598 | 8.929508 |
| 9.938703 | 9.784872 | 9.982758 | 7.371063 | 7.419881 | 5.862909 | 6.058808 | 6.14466 | 8.566092 |
| NaN | NaN | NaN | NaN | NaN | NaN | NaN | NaN | 6.449278 |
| 8.374104 | 8.42305 | 8.423301 | 11.85896 | 12.18804 | 10.69011 | 10.77539 | 10.54849 | 10.12418 |
| 13.59404 | 13.53448 | 13.58763 | 11.04881 | 10.89167 | 9.422236 | 9.375301 | 9.406851 | 10.00804 |
| 13.42226 | 13.45738 | 13.54633 | 10.95804 | 10.94093 | 9.659657 | 9.533776 | 9.745036 | 10.68415 |
| 9.237228 | 9.311419 | 8.926811 | 10.26822 | 10.17131 | 9.538608 | 9.503643 | 9.64737 | 10.33088 |
| 8.505624 | 9.454956 | 9.290406 | 6.728172 | 7.08092 | 4.925108 | 5.51033 | 5.38295 | 7.414955 |
| 9.552937 | 9.66489 | 9.61114 | 7.886541 | 7.845715 | 9.743085 | 9.525507 | 9.458522 | 9.814863 |
| 3.417414 | 3.327811 | 4.022541 | NaN | NaN | NaN | NaN | NaN | NaN |
| 15.25841 | 15.00311 | 15.20549 | 12.73615 | 12.79379 | 9.763911 | 9.332937 | 9.32644 | 12.05439 |
| 7.009264 | 6.83854 | 6.008736 | NaN | 4.777839 | NaN | NaN | NaN | 7.5841 |
| NaN | NaN | 7.422086 | 6.130929 | 7.54251 | 4.463379 | 5.487813 | 6.705244 | NaN |
| 11.90222 | 9.873096 | 9.676985 | 9.133644 | NaN | NaN | 10.77809 | NaN | NaN |
| 6.542905 | 6.84426 | 6.901082 | 6.286375 | 6.274524 | 5.589641 | 5.743394 | 6.38494 | 7.067062 |
| 8.962149 | 8.938969 | 9.107675 | 7.793433 | 8.186783 | 6.573712 | 6.406642 | 6.323952 | 11.01553 |
| 13.19765 | 13.04258 | 13.09413 | 11.11351 | 10.83932 | 11.35129 | 11.22118 | 11.35855 | 11.9277 |
| 3.688613 | 4.236816 | 3.792241 | NaN | NaN | NaN | NaN | NaN | NaN |
| 7.483099 | 7.560569 | 7.460178 | 6.360898 | 6.390554 | 7.790662 | 7.756283 | 7.599256 | 8.320902 |
| 10.59077 | 10.59279 | 10.61369 | 9.957691 | 9.988574 | 9.068014 | 8.918147 | 9.007467 | 9.409372 |
| 8.428146 | 7.286616 | 7.896154 | 6.871767 | 6.229545 | 7.494505 | 5.949529 | 6.321678 | 7.856176 |
| 7.212695 | 7.380383 | 7.790304 | 5.719742 | NaN | 6.377089 | 6.121323 | 6.606594 | 6.408738 |
| 7.663933 | 8.33804 | 8.245537 | 10.42572 | 10.90504 | 8.788869 | 8.685117 | 8.648737 | 6.715246 |

| | | | | | | | | |
|---|---|---|---|---|---|---|---|---|
| 9.433729 | 9.418659 | 9.431639 | 11.01466 | 11.36769 | 9.927258 | 10.03484 | 9.920612 | 7.982015 |
| 7.192826 | 6.657048 | 6.491138 | NaN | NaN | 7.233698 | 7.407561 | NaN | 7.464083 |
| 9.449706 | 9.381843 | 9.583807 | 9.706978 | 9.297222 | 9.225217 | 9.129652 | 9.221588 | 7.749054 |
| 6.504611 | NaN | 7.019674 | 7.326323 | 6.90188 | NaN | 6.576517 | 6.308645 | 7.474773 |
| 11.66425 | 11.70418 | 11.75346 | 10.29648 | 10.34499 | 10.49346 | 10.45533 | 10.6017 | 10.14078 |
| 6.786324 | 6.874012 | 6.712495 | 5.145577 | 6.173658 | 7.269793 | 6.845228 | 7.006993 | 6.274017 |
| 8.119508 | 7.985754 | 8.175813 | 7.993979 | 7.914675 | 7.867241 | 7.687009 | 7.481677 | 7.8258 |
| 7.739408 | 7.544008 | 7.632391 | 7.063973 | 7.297938 | 7.952886 | 7.777447 | 7.631225 | 6.442103 |
| 11.19645 | 10.96015 | 10.82948 | 10.96918 | 10.89627 | 11.77578 | 11.74522 | 11.72173 | 11.70154 |
| 9.218132 | 8.86246 | 8.913232 | 10.95294 | 10.94433 | 8.435513 | 8.37399 | 8.633439 | 7.482692 |
| 10.23953 | 10.25564 | 10.45316 | 12.00523 | 11.77693 | 10.35664 | 10.40592 | 10.53961 | 10.75969 |
| 8.373516 | 8.207577 | 8.430937 | 8.842237 | 8.200167 | 8.496061 | 8.474661 | 8.367619 | 6.984552 |
| 10.27927 | 10.1555 | 10.16584 | 10.64242 | 11.13458 | 9.590077 | 9.592798 | 9.641191 | 8.84102 |
| 10.0526 | 10.07193 | 9.845178 | 9.374762 | 9.403233 | 10.02819 | 9.933572 | 10.17151 | 9.230068 |
| 5.937368 | 9.359858 | 9.269059 | 8.939441 | 9.371619 | 9.104493 | 8.762055 | 8.864119 | 6.23015 |
| 14.19964 | 13.76786 | 13.8983 | 11.12067 | 11.51678 | 8.930978 | 8.753085 | 8.787359 | 10.20583 |
| 14.21257 | 14.22207 | 14.25673 | 11.19204 | 11.18491 | 9.473204 | 9.718529 | 9.637108 | 9.755473 |
| 7.188443 | 7.297358 | 7.421538 | 7.679305 | 7.739936 | 8.264251 | 8.124627 | 8.279939 | 5.62448 |
| 12.74127 | 12.37497 | 12.62429 | 11.64814 | 11.51863 | 12.83182 | 12.75086 | 12.5979 | 11.33702 |
| 14.23236 | 14.2245 | 14.27347 | 13.87772 | 14.09004 | 14.07384 | 14.16784 | 14.06186 | 13.00324 |
| 8.741119 | 9.059764 | 9.184828 | 7.510731 | 7.497217 | 7.464011 | 7.356374 | 7.489917 | 6.477537 |
| 9.21321 | 9.206243 | 9.001141 | 8.484149 | 8.271453 | 7.734697 | 7.81252 | 7.740123 | 8.403781 |
| NaN | NaN | 7.354987 | NaN | NaN | NaN | 6.925038 | NaN | 6.676351 |
| NaN | NaN | NaN | NaN | NaN | NaN | NaN | NaN | 5.938524 |
| 8.48016 | 8.716407 | 8.598801 | 9.177377 | 9.158256 | 8.317811 | 8.466357 | 8.42404 | 8.390436 |
| 10.84762 | 10.81199 | 11.19341 | 13.09677 | 13.51055 | 10.6598 | 10.75994 | 10.65972 | 11.2178 |
| 6.681781 | 6.626747 | 6.999089 | 5.292709 | 5.462833 | 6.277825 | 6.4492 | 6.613642 | 6.838642 |
| 11.2256 | 10.97387 | 11.12506 | 9.641403 | 9.688254 | 11.06842 | 10.89871 | 10.85186 | 11.16691 |
| 10.29996 | 10.25042 | 10.34541 | 8.930567 | 8.742665 | 9.762091 | 9.946065 | 9.783851 | 10.32797 |
| 10.62197 | 10.50397 | 10.61314 | 9.302011 | 9.151056 | 10.12917 | 10.09256 | 9.863995 | 10.38162 |
| 6.98266 | 4.798768 | 6.616974 | 6.453184 | 5.925564 | 6.406424 | 7.30923 | 8.314337 | 5.103544 |
| 7.037985 | 7.095355 | 7.077164 | 6.390508 | 5.938278 | 6.631173 | 6.654285 | 6.280656 | 7.177561 |
| 8.207457 | 8.258596 | 7.733654 | NaN | NaN | NaN | NaN | NaN | NaN |
| 9.435534 | 9.348768 | 9.388579 | 7.722613 | 7.876286 | 9.306673 | 9.299794 | 9.342947 | 8.829069 |
| 11.95528 | 11.60627 | 11.68558 | 9.892118 | 9.875395 | 11.31005 | 11.41244 | 11.14107 | 10.44892 |
| 9.557684 | 9.52256 | 9.607754 | 9.638077 | 9.470341 | 8.784737 | 8.601769 | 8.819709 | 8.55051 |
| 8.509439 | 7.452472 | 8.75933 | 5.631178 | 5.66866 | 7.42638 | 6.891506 | 5.996461 | 5.862637 |
| 11.08245 | 10.80505 | 10.96704 | 10.21438 | 9.973935 | 9.813638 | 9.817785 | 9.79727 | 9.57758 |
| 10.15132 | 10.04565 | 10.13149 | 7.790874 | 8.111825 | 6.937651 | 7.165573 | 6.997457 | 6.846726 |
| 10.45991 | 10.38583 | 10.33524 | 9.564525 | 9.648263 | 10.07232 | 10.24668 | 10.37619 | 9.756882 |
| 7.94172 | 7.849766 | 8.217208 | 8.02377 | 7.880494 | 6.875376 | 6.604068 | 6.592148 | 7.151059 |
| 6.328438 | 6.345729 | 6.540451 | 5.382275 | 9.307539 | 7.429258 | 6.812796 | 7.486201 | 7.095783 |
| 6.900571 | 6.956609 | 7.049787 | 6.653482 | NaN | 4.39582 | NaN | 8.341633 | 7.281869 |
| 5.494381 | 5.533872 | 5.965938 | NaN | NaN | 7.283882 | 6.537022 | 7.683254 | 5.48059 |
| 11.75835 | 11.80173 | 11.6237 | 10.51702 | 10.54994 | 11.21828 | 11.26033 | 11.17294 | 10.1693 |
| 7.791955 | 7.592906 | 7.767746 | NaN | NaN | NaN | NaN | NaN | NaN |
| 5.121166 | 2.988893 | 5.067191 | 4.85381 | 5.61869 | 5.048268 | 5.411446 | NaN | 5.422427 |

| | | | | | | | | |
|---|---|---|---|---|---|---|---|---|
| 7.87853 | 5.368738 | 6.033207 | NaN | 4.984946 | NaN | NaN | NaN | 8.07519 |
| 8.124817 | 8.162882 | 8.349854 | 6.988564 | 7.155656 | 8.238839 | 8.228406 | 8.21196 | 8.11586 |
| 9.14662 | 9.155705 | 9.157344 | 8.347116 | 8.201295 | 9.51931 | 9.553722 | 9.461625 | 9.004872 |
| 7.011584 | 7.057452 | 6.963712 | NaN | NaN | 6.898537 | 6.781434 | 6.72712 | 7.511855 |
| 10.08072 | 9.998057 | 10.31 | 9.296754 | 9.540899 | 9.599685 | 9.76521 | 9.86496 | 9.799562 |
| 6.720211 | NaN | 7.823624 | 7.874272 | 7.844031 | 6.57613 | 7.199207 | 6.279471 | NaN |
| 5.450912 | NaN | NaN | NaN | NaN | 4.915046 | NaN | NaN | 6.170123 |
| 11.41186 | 11.79862 | 11.8842 | 13.13038 | 13.49849 | 10.27071 | 10.16171 | 10.36228 | 11.3776 |
| 13.30262 | 12.74484 | 12.53764 | 13.3998 | 12.93355 | 9.91793 | 9.446419 | 10.42533 | 10.95704 |
| 9.33438 | 9.4774 | 9.439153 | 9.639633 | 9.482093 | 9.782393 | 9.764584 | 9.712157 | 9.777351 |
| 5.268806 | NaN | 6.303012 | 5.203915 | 5.46659 | 6.204702 | 6.321202 | 6.240884 | 4.919011 |
| 6.645597 | 6.559253 | 6.07081 | 8.413263 | 8.009546 | 6.73366 | 6.503625 | 6.700504 | 3.820778 |
| 7.86455 | 7.655039 | 7.276055 | 6.956022 | 6.321532 | 8.637931 | 7.462544 | 7.844102 | 7.231355 |
| 10.86798 | 10.97481 | 11.02202 | 12.00406 | 11.79607 | 11.16559 | 11.2163 | 11.22887 | 11.27645 |
| 7.871478 | 7.978537 | 8.247629 | 5.102411 | 5.056689 | 7.405401 | 6.552651 | 7.378427 | 6.827541 |
| 9.125572 | 8.301244 | 8.120369 | 7.619946 | 7.212089 | NaN | NaN | NaN | NaN |
| NaN | NaN | NaN | NaN | NaN | NaN | NaN | NaN | 5.727114 |
| 9.052243 | 8.725868 | 8.730934 | 7.472374 | 7.534809 | 8.041661 | 8.515488 | 8.152286 | 8.594938 |
| 7.486147 | 7.498025 | 7.479365 | 6.273277 | 6.098653 | 7.087469 | 7.059549 | 6.89017 | 5.513402 |
| 9.401962 | 9.041784 | 9.425887 | 8.830481 | 8.678533 | 6.729659 | 7.710792 | 7.19102 | 8.190757 |
| 9.771356 | 9.648957 | 9.620587 | 10.018 | 9.848501 | 7.946195 | 7.855338 | 7.403162 | 9.122183 |
| 8.60827 | 8.585161 | 8.512098 | 8.726849 | 8.735984 | 7.781937 | 8.387731 | 8.196833 | 9.982005 |
| 9.428663 | 8.304942 | 9.487289 | 8.420527 | 8.429327 | 7.951876 | 8.184949 | 8.041978 | 8.1675 |
| 7.732965 | 7.943685 | 8.557497 | 8.021312 | 8.177242 | 8.353473 | 8.170105 | 8.344982 | 7.386571 |
| 7.542666 | 7.016901 | 6.904607 | 6.252301 | 6.707662 | 8.068292 | 7.620968 | 8.185139 | 6.969656 |
| 6.018053 | 5.918174 | 6.219325 | NaN | NaN | NaN | 5.839789 | 5.466399 | 5.004451 |
| 10.72052 | 10.76741 | 10.78479 | 10.74151 | 10.8073 | 10.47541 | 10.39291 | 10.27305 | 10.48814 |
| 8.231388 | 8.092816 | 8.622225 | NaN | NaN | 7.859433 | 7.003353 | 7.376272 | 7.168735 |
| 7.942416 | 7.528296 | 7.636597 | 10.24377 | 10.35405 | 7.442613 | 6.861307 | 7.699631 | 8.184045 |
| 9.767603 | 9.583893 | 9.945874 | 8.621246 | 8.443769 | 8.570363 | 8.664139 | 8.848472 | 9.383863 |
| 14.68178 | 14.56796 | 14.70214 | 15.60761 | 15.67298 | 14.27269 | 14.25617 | 14.25692 | 13.43603 |
| 8.42984 | 8.377548 | 8.463968 | 8.238406 | 8.044227 | 6.836905 | 7.0478 | 7.124672 | 8.232661 |
| 9.197252 | 7.635935 | 8.646633 | 8.432999 | 8.559214 | NaN | NaN | 8.055059 | 7.467918 |
| 8.450387 | 8.418156 | 8.289288 | 8.573181 | 8.254217 | 8.589895 | 9.119026 | 8.643509 | 8.34133 |
| 9.532285 | 9.621914 | 9.770803 | 9.490135 | 9.637631 | 9.577662 | 8.639045 | 9.281421 | 9.524483 |
| 10.87212 | 10.76756 | 10.64881 | 11.28622 | 11.36401 | 11.67654 | 11.61461 | 11.54586 | 11.89186 |
| 9.371821 | 9.179488 | 9.01784 | 6.900107 | 7.231984 | 7.813057 | 7.450842 | 8.105028 | 7.930439 |
| 10.36065 | 10.52709 | 10.5421 | 10.27911 | 10.49707 | 11.32557 | 11.35677 | 11.32269 | 11.01436 |
| 6.978848 | 7.239966 | 6.434008 | 7.02647 | 7.164093 | 6.721013 | 6.782219 | 6.771995 | 7.162206 |
| 11.67133 | 11.57125 | 11.66429 | 12.43763 | 12.25454 | 11.50348 | 11.5297 | 11.60989 | 12.46648 |
| 12.0567 | 12.13861 | 12.15454 | 12.40282 | 12.18528 | 10.5106 | 11.08299 | 10.69203 | 11.20257 |
| 8.041222 | 7.973103 | 8.316235 | 6.744363 | 6.295171 | 7.478025 | 7.669876 | 7.442635 | 7.161952 |
| 8.98938 | 9.097075 | 9.000244 | 7.910029 | 7.642231 | 7.119907 | 7.190565 | 7.691742 | 4.751674 |
| 9.035841 | 9.035126 | 8.900608 | 10.21823 | 10.32934 | 9.992454 | 9.861493 | 10.06929 | 7.893709 |
| 8.558174 | 8.505457 | 8.342628 | 8.308569 | 8.177683 | 7.991787 | 8.093403 | 7.897892 | 7.419835 |
| 14.37282 | 14.28718 | 14.32522 | 14.97236 | 14.99844 | 14.93367 | 15.11053 | 15.17197 | 14.17371 |
| 7.798586 | 7.937155 | 8.106138 | 8.777718 | 8.85086 | 7.870719 | 7.905046 | 7.898808 | 8.107086 |

| | | | | | | | | |
|---|---|---|---|---|---|---|---|---|
| 8.769278 | 8.684716 | 9.004362 | 7.273282 | 7.380048 | 10.11439 | 9.99501 | 9.895457 | 8.840858 |
| 9.614711 | 9.633558 | 9.70371 | 12.07128 | 11.54195 | 8.960082 | 9.249595 | 9.248551 | 9.290197 |
| 11.70148 | 11.50152 | 11.67867 | 11.87453 | 11.91082 | 12.21914 | 12.05142 | 12.32105 | 11.41669 |
| 11.63929 | 11.51778 | 11.56433 | 12.97069 | 12.97356 | 11.071 | 10.99381 | 11.10708 | 10.98749 |
| 11.11835 | 11.05017 | 11.1095 | 12.30487 | 12.35249 | 10.47707 | 10.39299 | 10.44538 | 11.04056 |
| 4.999293 | 2.253468 | 5.056076 | 4.616408 | 5.077355 | 5.920387 | 5.529607 | 6.095623 | 4.940968 |
| 9.947584 | 9.929676 | 10.02803 | 10.82641 | 10.85396 | 10.22482 | 10.22368 | 9.961255 | 10.31589 |
| 8.48632 | 8.68328 | 8.933804 | 6.971856 | 7.005723 | 8.603239 | 8.134573 | 8.730568 | 7.372318 |
| 10.93073 | 10.90017 | 10.96318 | 10.9783 | 10.99252 | 11.1913 | 11.15388 | 11.30915 | 11.23569 |
| 9.237689 | 9.195669 | 9.341636 | 9.476658 | 9.505761 | 7.989392 | 8.127304 | 8.051685 | 9.24614 |
| 10.18801 | 10.23878 | 10.34978 | 11.40739 | 11.3778 | 9.14854 | 8.893837 | 9.012763 | 9.606489 |
| 11.20528 | 11.20317 | 11.1636 | 12.0303 | 12.01782 | 10.46416 | 10.17141 | 10.10807 | 10.51896 |
| 9.701483 | 9.738832 | 9.931518 | 11.70965 | 11.50436 | 10.53855 | 10.47217 | 10.53631 | 11.65917 |
| 8.603833 | 8.670635 | 8.610283 | 8.216071 | 8.222681 | 8.927572 | 8.874413 | 8.943212 | 8.356329 |
| 14.45159 | 14.34779 | 14.58004 | 13.26378 | 13.37627 | 14.50605 | 14.66445 | 14.70462 | 13.8131 |
| NaN | 13.05999 | 10.07551 | 11.45442 | 13.89602 | 9.3248 | 12.3159 | 12.2162 | 12.12373 |
| 8.321335 | 8.266204 | 7.678857 | 8.808599 | 8.31115 | 8.267353 | 8.442969 | 8.364784 | NaN |
| 7.348366 | 7.169429 | 7.264316 | 9.187778 | 9.161498 | 7.788477 | 7.67475 | 7.682752 | 7.99916 |
| 9.694945 | 9.70363 | 9.681279 | 10.15912 | 10.08926 | 9.065323 | 9.271149 | 9.142659 | 8.710567 |
| 12.30607 | 12.24874 | 12.37098 | 13.08518 | 13.18859 | 12.28419 | 12.46937 | 12.51122 | 12.95174 |
| 7.619738 | 7.517603 | 7.33682 | 7.704587 | 7.899591 | 8.536233 | 8.530006 | 8.687906 | 7.728814 |
| 8.717157 | 8.894342 | 8.683425 | 10.44694 | 10.20905 | 8.708819 | 8.54964 | 8.299698 | 9.330818 |
| 8.936504 | 8.895981 | 8.905021 | 7.828509 | 8.011278 | 9.198166 | 9.099044 | 9.194437 | 9.370175 |
| 8.833132 | 8.826451 | 8.853454 | 8.479589 | 7.95547 | 8.120228 | 7.963423 | 8.112895 | 7.789957 |
| 13.63656 | 13.65809 | 13.83354 | 12.94118 | 12.78682 | 13.94993 | 14.06966 | 13.98993 | 13.86073 |
| 6.612728 | NaN | 7.878158 | 7.103476 | 7.02933 | 7.729011 | 7.823294 | 7.905204 | 7.676653 |
| 11.61148 | 11.56073 | 11.49624 | 12.9752 | 12.80127 | 11.44241 | 11.38929 | 11.36794 | 11.61907 |
| 6.253478 | 5.985369 | 6.870489 | 7.231871 | 6.924967 | 6.924717 | 7.430085 | 6.479314 | 6.450152 |
| 6.356132 | 4.795838 | 6.19851 | NaN | NaN | NaN | 5.362021 | NaN | 6.615874 |
| 8.975524 | 8.887846 | 8.956261 | 8.204353 | 7.965716 | 8.851859 | 8.927841 | 8.996344 | 9.092792 |
| 11.03872 | 8.677957 | 9.157687 | 10.26889 | 9.75077 | 8.693605 | NaN | NaN | 8.432616 |
| 10.85785 | 10.77366 | 10.63594 | 11.74211 | 11.66394 | 10.91106 | 10.63521 | 10.82525 | 10.1395 |
| 8.219127 | 7.444057 | 7.556575 | 7.178629 | 6.584925 | 7.091025 | 6.462872 | 6.820668 | 8.63307 |
| 6.774121 | 6.915181 | 6.911946 | 5.160997 | 4.996701 | 6.967918 | 6.849991 | 6.775743 | 6.832584 |
| 6.819959 | 6.780657 | 7.071745 | 5.105013 | 5.052759 | 6.397712 | NaN | 4.013206 | 8.616322 |
| 10.36983 | 10.44638 | 10.44545 | 9.8744 | 9.94523 | 10.95592 | 10.84639 | 10.98647 | 10.49589 |
| 6.648189 | 6.162841 | 6.152132 | 5.923765 | 6.34972 | NaN | NaN | 5.537536 | 6.154177 |
| 10.3405 | 10.20538 | 10.19449 | 9.774182 | 9.652075 | 10.33869 | 10.47513 | 10.44849 | 9.948362 |
| 7.405022 | 7.172032 | 7.258933 | 8.636898 | 8.880561 | 7.695458 | 7.472458 | 7.659009 | 6.596254 |
| 9.160933 | 8.864492 | 9.018844 | 8.036946 | 7.79575 | 8.973322 | 9.02633 | 9.06411 | 8.500113 |
| 13.9518 | 13.95703 | 13.98333 | 13.10062 | 12.9325 | 14.09962 | 14.18807 | 14.08153 | 13.95505 |
| 6.376919 | 6.803461 | 6.202612 | 5.551327 | 5.562937 | 6.512648 | 6.437582 | 6.520531 | 6.805845 |
| 9.757447 | 9.666033 | 9.686704 | 8.685734 | 8.645786 | 8.506441 | 8.605887 | 8.719663 | 9.693886 |
| NaN | NaN | NaN | NaN | NaN | NaN | NaN | NaN | 9.772326 |
| 9.480211 | 8.651836 | 8.976606 | 8.487701 | 6.890514 | 8.515461 | 8.679427 | 8.64262 | 7.440876 |
| 7.24254 | 7.469705 | 7.713273 | 9.342571 | 9.011746 | 10.07293 | 9.832889 | 9.753754 | 9.490816 |
| 5.573972 | 6.556661 | 6.875305 | NaN | NaN | NaN | NaN | NaN | 8.078011 |

-Bruker Confidential-

| | | | | | | | | |
|---|---|---|---|---|---|---|---|---|
| NaN | 6.58709 | 7.882813 | 7.242506 | 7.196399 | NaN | NaN | NaN | 6.402322 |
| 9.223863 | 9.269438 | 9.045349 | 8.64446 | 8.515399 | 9.201984 | 9.276961 | 9.184011 | 9.498004 |
| 7.433555 | 6.59614 | 7.194967 | 5.640858 | 5.565943 | 6.348624 | 6.938202 | 6.235064 | 7.616572 |
| 8.336932 | 8.424943 | 8.421665 | 7.091635 | 6.803871 | 8.605173 | 8.61033 | 8.743299 | 8.077184 |
| 8.350078 | 8.23531 | 8.492329 | 9.404692 | 9.64803 | 8.123185 | 8.108225 | 8.107705 | 7.454007 |
| 9.009935 | 8.911993 | 8.901367 | 10.37287 | 10.25212 | 9.728269 | 9.650365 | 9.682547 | 9.843747 |
| 7.46358 | 6.679629 | 6.538706 | NaN | 6.729513 | 6.721379 | 6.714562 | 6.872711 | 8.328889 |
| 13.91764 | 13.96487 | 13.98694 | 15.14286 | 15.1728 | 12.81468 | 12.74305 | 12.75271 | 13.12031 |
| 9.358779 | 9.270592 | 9.026464 | 10.53366 | 10.52892 | 9.115687 | 9.140071 | 9.151854 | 9.027466 |
| 9.276793 | 8.833804 | 9.064951 | 7.432593 | 7.156529 | NaN | 7.2596 | 8.878718 | 7.688324 |
| 9.784272 | 9.722419 | 9.911395 | 7.715749 | 7.502468 | 9.399684 | 9.520807 | 9.719833 | 9.590372 |
| 9.748142 | 9.780537 | 9.803124 | 11.44977 | 11.19581 | 10.45075 | 10.42817 | 10.45423 | 10.65762 |
| 10.40139 | 10.31382 | 10.32084 | 7.428797 | 7.179924 | 7.867992 | 7.286592 | 7.44972 | 7.721027 |
| 6.669077 | 6.347765 | 6.276327 | NaN | NaN | 6.363364 | 6.333338 | 6.838137 | NaN |
| 12.06859 | 12.01037 | 12.03648 | 12.60946 | 12.74176 | 12.12487 | 11.74363 | 12.09377 | 11.63278 |
| NaN | NaN | NaN | NaN | NaN | 6.214464 | 5.216153 | 6.355549 | NaN |
| 5.849123 | 4.267715 | 5.347854 | 5.490253 | 5.263534 | 5.64994 | 5.461841 | 6.112365 | 3.97761 |
| 10.21083 | 9.939079 | 9.856624 | 8.500417 | 8.361389 | 8.599119 | 8.52346 | 8.603302 | 9.830577 |
| 7.055493 | 7.396198 | 7.233951 | 10.01335 | 10.09759 | 8.516955 | 8.515431 | 8.935574 | 7.981818 |
| 7.131747 | 7.178474 | 7.51792 | 6.716789 | 6.643028 | 7.036358 | 6.985813 | 7.118861 | 6.695083 |
| 7.159811 | 7.328166 | 7.157456 | 7.469902 | 8.357579 | 7.052735 | 6.770878 | 6.754132 | 7.784254 |
| 11.95587 | 11.84954 | 11.96526 | 12.5267 | 12.54729 | 11.90375 | 11.86754 | 11.86544 | 11.46834 |
| 9.311448 | 9.26643 | 9.353261 | 8.429296 | 8.383889 | 9.588572 | 9.628745 | 9.520441 | 8.511403 |
| 6.240662 | 6.284632 | 6.059354 | 7.130132 | 5.987586 | 7.596127 | 7.405426 | 7.606823 | 7.406507 |
| 11.73073 | 11.66764 | 11.37855 | 12.41962 | 12.30741 | 11.90702 | 11.51524 | 11.63008 | 11.87482 |
| 6.904303 | 6.463346 | 6.978226 | 8.1832 | 8.5993 | 7.18305 | 6.533627 | NaN | 7.340498 |
| 6.577766 | 6.723639 | 6.789384 | 8.056032 | 8.063201 | 6.839753 | 7.436233 | 6.828489 | 7.440971 |
| 9.316922 | 9.315853 | 9.083969 | 9.64973 | 9.553656 | 9.988725 | 10.06273 | 10.33089 | 8.749521 |
| 11.37221 | 11.52531 | 11.50085 | 11.37525 | 11.45268 | 12.54671 | 12.32502 | 12.42921 | 11.4286 |
| 9.488519 | 9.363276 | 9.640589 | 10.26707 | 10.27526 | 9.679685 | 9.50801 | 9.731878 | 9.889477 |
| 11.37603 | 11.36192 | 11.37721 | 9.508492 | 9.433672 | 11.89423 | 11.8196 | 11.86164 | 9.594867 |
| 10.13668 | 10.35665 | 10.52668 | 8.363629 | 8.538152 | 8.125428 | 8.518108 | 8.62802 | 8.747608 |
| 7.618592 | 7.368447 | 7.419577 | NaN | NaN | NaN | 6.711816 | 6.503433 | 5.802486 |
| 7.042823 | 6.331112 | 6.853842 | 7.834867 | 7.787946 | NaN | 6.087566 | 6.353233 | 6.715918 |
| 9.679623 | 9.891252 | 9.640276 | 11.31391 | 11.47782 | 9.749122 | 9.845024 | 9.611586 | 9.686637 |
| 9.541394 | 9.457102 | 9.410208 | 12.23521 | 11.75871 | 8.706215 | 8.976706 | 8.926655 | 9.089095 |
| 7.763489 | 7.54858 | 7.452125 | 8.870333 | 8.759381 | 8.086515 | 8.104282 | 8.190813 | 8.626332 |
| 10.1135 | 9.891485 | 10.0012 | 8.818477 | 8.891139 | 10.41522 | 10.30118 | 10.40291 | 8.817922 |
| 6.473533 | 7.056833 | 7.816337 | 7.476074 | 7.791608 | 6.631321 | 6.115318 | 6.391934 | 7.195605 |
| NaN | NaN | NaN | NaN | NaN | 7.494621 | 6.284836 | 7.533823 | 5.241471 |
| 9.780922 | 9.818686 | 9.865625 | 10.78443 | 10.72406 | 9.917802 | 9.885674 | 9.987354 | 9.412144 |
| 12.76692 | 12.63614 | 12.6423 | 13.24908 | 12.63536 | 12.4599 | 12.54219 | 12.60028 | 12.19097 |
| 8.666621 | 8.299956 | 8.091071 | 10.47607 | 10.34723 | 7.746251 | 8.193682 | 7.983615 | 7.316534 |
| 6.333239 | 6.268141 | 6.087291 | 7.14698 | 6.969194 | 7.12341 | 6.622846 | 7.066615 | 5.512795 |
| 11.02742 | 10.96008 | 10.96474 | 10.56422 | 10.63426 | 10.90823 | 10.99658 | 11.09247 | 11.13249 |
| 11.55291 | 11.65936 | 11.65353 | 11.63534 | 11.63447 | 12.04595 | 12.07966 | 12.08398 | 10.95995 |
| 7.381861 | 7.420528 | 7.286759 | 6.99657 | 7.24657 | 7.16135 | 7.303314 | 7.514475 | 7.751556 |

| | | | | | | | | |
|---|---|---|---|---|---|---|---|---|
| 8.061635 | 8.364536 | 8.087659 | 9.98312 | 10.22993 | 9.80891 | 9.684754 | 9.792455 | 8.532595 |
| 6.618864 | 6.945487 | 7.582065 | NaN | NaN | 5.456301 | 5.253756 | 6.041639 | 5.124186 |
| 9.661565 | 9.443067 | 9.602426 | 10.56447 | 10.29798 | 9.637806 | 9.380671 | 9.901568 | 8.691314 |
| 15.027 | 15.23061 | 15.33134 | 15.13648 | 15.44127 | 16.14927 | 16.30948 | 16.63535 | 13.92137 |
| NaN | 5.89045 | 5.852071 | 6.595725 | 6.227702 | 5.948091 | 5.894255 | 6.007967 | 5.958244 |
| NaN | 7.145723 | 7.118793 | 6.622675 | 6.752048 | 6.695685 | 6.436254 | 6.253926 | NaN |
| 9.394322 | 9.322479 | 9.346083 | 12.81804 | 12.90512 | 9.833559 | 9.600887 | 9.617906 | 9.398875 |
| 11.76686 | 11.81518 | 11.809 | 10.08367 | 9.77647 | 10.10233 | 9.711698 | 9.438689 | 10.38558 |
| 13.46654 | 13.59991 | 13.85384 | 14.33282 | 14.24029 | 11.82265 | 12.00201 | 11.91217 | 11.54096 |
| 7.430261 | 7.978556 | 5.562081 | NaN | NaN | NaN | NaN | 6.105126 | NaN |
| 6.897503 | 6.935955 | 6.723017 | 7.654842 | 7.093452 | 7.272409 | 7.007976 | 7.216498 | 6.349563 |
| 7.513832 | 8.1469 | 7.650466 | 6.427181 | 6.468383 | 7.78049 | 7.698395 | 7.672295 | 4.719885 |
| 14.51741 | 14.62424 | 14.56837 | 14.823 | 14.83342 | 14.44838 | 14.58116 | 14.47208 | 14.79554 |
| 8.630787 | 8.34499 | 8.327946 | 6.863028 | 7.128253 | NaN | NaN | 7.073822 | 7.884768 |
| 14.27177 | 13.93229 | 13.89353 | 13.1387 | 13.15215 | 12.08202 | 11.91158 | 12.06004 | 11.91459 |
| 6.438681 | 6.368897 | 5.742645 | 4.699752 | 5.192874 | NaN | NaN | NaN | NaN |
| 13.1407 | 12.96761 | 12.56198 | 13.41564 | 13.83547 | 10.82921 | 10.445 | 10.42159 | 11.44066 |
| 5.891912 | 6.041149 | 5.928802 | 6.582386 | 6.612058 | NaN | NaN | 5.738929 | NaN |
| 2.684653 | 4.179532 | 3.977647 | 4.607144 | 5.769207 | 4.018798 | 2.723913 | 2.699637 | 4.772134 |
| 7.656849 | 7.487571 | 7.59381 | 7.86989 | 7.98435 | 7.593644 | 7.352011 | 7.481763 | 7.219064 |
| 8.259246 | 8.301243 | 8.251158 | 8.207697 | 8.213574 | 8.717365 | 8.523788 | 8.694426 | 8.497921 |
| 3.295227 | NaN | 1.817736 | NaN | 1.782199 | NaN | NaN | 2.691604 | 6.798055 |
| 10.71027 | 10.71367 | 10.79466 | 11.18338 | 11.29733 | 11.00738 | 10.91281 | 11.01881 | 10.37487 |
| 6.550923 | 6.530005 | 6.594525 | 6.434114 | 6.177062 | 7.332914 | 7.257145 | 7.236921 | 7.029637 |
| 5.627783 | 5.710393 | 6.638869 | 5.931469 | 4.809877 | 7.904991 | 7.383776 | 7.313622 | 5.068182 |
| 6.580258 | 6.000851 | 6.67991 | NaN | NaN | 6.70356 | 6.377862 | 6.491224 | 6.311632 |
| 13.96497 | 14.0684 | 14.06329 | 13.89196 | 13.90437 | 14.76708 | 14.652 | 14.63334 | 12.97211 |
| 6.869201 | 6.252497 | 5.991102 | NaN | NaN | NaN | 6.98645 | 6.347674 | 7.246132 |
| 8.226626 | 7.54617 | 7.170285 | 5.10631 | 6.220843 | NaN | NaN | 5.985025 | 5.469254 |
| 9.377698 | 9.409115 | 9.393226 | 8.864552 | 9.315217 | 9.182765 | 9.334459 | 9.194401 | 8.955461 |
| 8.112735 | 7.915255 | 7.924497 | 7.227348 | 7.187577 | 7.505953 | 7.274398 | 7.438981 | 7.173775 |
| 8.662724 | 8.633169 | 8.584213 | 6.657202 | 6.38954 | 8.191661 | 8.069345 | 8.307778 | 8.925641 |
| 10.98794 | 11.29603 | 10.85621 | 11.85139 | 11.56921 | 10.43733 | 10.31581 | 10.46463 | 10.08803 |
| 9.916601 | 9.839849 | 9.722329 | 11.40518 | 11.2613 | 9.098612 | 9.247253 | 8.637337 | 9.126904 |
| 7.733299 | 7.62593 | 7.675172 | 7.463362 | 7.523587 | 6.36424 | 6.124064 | 6.019676 | 5.824588 |
| 7.987103 | 7.155855 | 7.099825 | 7.094385 | 7.477755 | 6.252335 | 6.466614 | 5.348041 | 5.91121 |
| 11.16994 | 11.19442 | 11.35538 | 14.01489 | 13.93191 | 11.14864 | 11.0801 | 11.11457 | 10.042 |
| 7.860113 | 7.728148 | 7.789972 | 7.712998 | 7.661645 | 7.924039 | 7.917891 | 7.770919 | 7.473038 |
| 9.587918 | 9.397376 | 9.315423 | 9.003292 | 8.882689 | 8.41888 | 8.464138 | 8.349529 | 8.006677 |
| 9.723816 | 9.611007 | 9.624506 | 8.594387 | 8.626563 | 10.20746 | 10.39046 | 10.41794 | 10.0533 |
| 13.26805 | 13.2024 | 13.20002 | 13.60265 | 13.57098 | 12.78098 | 12.68734 | 12.88051 | 12.47 |
| 13.26313 | 13.32204 | 13.33458 | 13.84013 | 13.83698 | 13.69081 | 13.69271 | 13.83741 | 12.3676 |
| 13.27179 | 13.20312 | 13.30668 | 13.47361 | 13.45382 | 13.06628 | 12.95719 | 12.99245 | 13.34237 |
| 7.214702 | 7.409521 | 7.29494 | 6.167642 | 6.029834 | 6.991658 | 6.89412 | 6.830595 | 8.156387 |
| 7.526981 | 7.643209 | 7.430603 | 6.90969 | 7.535907 | 7.17069 | 7.256631 | 7.224782 | 7.093225 |
| 10.82999 | 10.89463 | 10.85318 | 12.23757 | 12.24734 | 10.03786 | 10.0617 | 10.07995 | 10.78887 |
| 12.97261 | 12.83134 | 12.78595 | 12.67724 | 12.58648 | 12.85572 | 12.73079 | 12.75917 | 12.40677 |

| | | | | | | | | |
|---|---|---|---|---|---|---|---|---|
| 8.10897 | 7.923579 | 8.159035 | 5.746375 | 6.056994 | 7.233099 | 7.487023 | 7.32413 | 8.000066 |
| 8.169735 | 6.987477 | 8.300649 | 5.865123 | 6.254912 | 8.106436 | 8.41301 | 7.924632 | 7.730564 |
| 9.002352 | 9.067573 | 9.159621 | 9.749008 | 9.377604 | 8.280268 | 8.054634 | 8.263137 | 9.962861 |
| 6.503143 | 6.480724 | 6.569949 | 5.551898 | 4.974496 | 7.027982 | 6.816802 | 6.901887 | 6.931508 |
| 9.234848 | 9.236896 | 9.392467 | 9.72278 | 9.473509 | 9.356787 | 9.601043 | 9.415137 | 9.215147 |
| 7.313134 | 7.357732 | 7.44704 | 7.384132 | 7.399369 | 8.044952 | 8.101536 | 8.007115 | 7.530479 |
| 9.376803 | 9.226378 | 9.212923 | 8.460768 | 8.65667 | 9.897491 | 9.919326 | 9.740183 | 9.348338 |
| 6.872625 | 6.709876 | 7.159317 | 6.4233 | 6.725006 | 4.609281 | 6.157363 | 6.219291 | 5.471485 |
| 9.66511 | 9.58285 | 9.55369 | 9.299065 | 9.561373 | 9.097446 | 9.378965 | 9.280619 | 8.533507 |
| 8.380859 | 8.026359 | 8.332997 | 8.388523 | 8.335946 | 8.12264 | 7.89218 | 8.019664 | 7.550145 |
| 9.556315 | 9.619297 | 9.658759 | 9.894016 | 9.80135 | 9.372282 | 9.213869 | 9.332707 | 9.289241 |
| 10.26095 | 10.13111 | 10.10847 | 9.88497 | 9.721786 | 9.13354 | 9.430293 | 9.111945 | 9.138889 |
| 8.542813 | 8.528118 | 8.665009 | 8.152705 | 8.062548 | 8.11028 | 8.225525 | 8.472136 | 8.178798 |
| 6.270817 | 6.040964 | 6.434452 | NaN | NaN | NaN | NaN | NaN | 5.318549 |
| 9.292982 | 9.19306 | 9.113955 | 7.987886 | 7.680093 | 8.997953 | 9.251699 | 9.12868 | 9.091129 |
| 8.737698 | 8.754175 | 8.607648 | 7.121691 | 7.263089 | 8.115518 | 8.114782 | 8.149637 | 8.599437 |
| 8.066858 | 7.912208 | 8.016938 | 7.249937 | 7.335746 | 8.341191 | 8.324783 | 8.47937 | 8.081702 |
| 8.81151 | 8.876163 | 9.180803 | 9.647159 | 9.892446 | 8.205863 | 8.185531 | 7.926382 | 8.368373 |
| 6.50895 | 6.571218 | 6.573616 | 5.843545 | 5.566952 | 6.785148 | 6.804748 | 6.775238 | 5.560623 |
| 6.60533 | 6.515453 | 6.937299 | 7.181895 | 7.201464 | 7.132741 | 6.84934 | 7.019662 | 6.533266 |
| 9.773955 | 9.596865 | 9.665038 | 8.852921 | 8.789153 | 9.798043 | 9.69437 | 9.822628 | 9.670374 |
| 7.393053 | 6.953802 | 7.410252 | 5.363951 | 5.676437 | 7.173093 | 7.201698 | 7.27483 | 7.641714 |
| 7.424607 | 7.506548 | 7.536659 | 6.533472 | 6.68659 | 8.063182 | 8.176359 | 8.489532 | 8.218957 |
| 8.752683 | 8.502726 | 7.490462 | 7.234566 | 6.55547 | 8.918375 | 6.732304 | 7.801142 | 6.504147 |
| 10.16838 | 10.10353 | 10.14731 | 10.09446 | 9.834637 | 9.384204 | 9.6331 | 9.617161 | 9.505131 |
| 9.53991 | 9.508641 | 9.525926 | 8.668654 | 8.314074 | 8.057007 | 8.361127 | 8.301394 | 9.133554 |
| NaN | NaN | NaN | 5.718367 | 6.188428 | 7.626044 | NaN | 6.908659 | 5.70687 |
| 8.891662 | 9.416409 | 8.864284 | 6.922952 | 6.209877 | 7.993377 | 6.627556 | 7.842368 | 5.551111 |
| 7.364506 | 7.23081 | 7.318672 | 6.623256 | 6.901031 | 7.417473 | 6.931172 | 7.590011 | 6.96961 |
| 8.146211 | 9.478015 | 8.026555 | 8.019051 | 7.694525 | 7.096494 | 7.399004 | 7.65275 | 9.039214 |
| 8.16245 | 8.099925 | 8.211903 | 7.201903 | 7.266725 | 7.948409 | 8.160266 | 8.045301 | 8.785229 |
| NaN | 3.563859 | NaN | NaN | NaN | NaN | NaN | NaN | 5.894701 |
| 7.025429 | NaN | NaN | 5.889404 | 5.499453 | 6.329912 | 6.537904 | 5.952425 | 5.918032 |
| 11.53719 | 12.04694 | 11.94954 | 10.92801 | 10.55965 | 10.73088 | 10.8867 | 10.33851 | 10.49549 |
| 9.936561 | 9.885638 | 9.969125 | 6.89539 | 7.726776 | 10.0567 | 9.104209 | 9.907668 | 7.458971 |
| NaN | 7.724918 | 7.867733 | 4.747051 | 4.638512 | 5.519756 | 5.074276 | 5.408651 | 3.058956 |
| 9.836937 | 9.810365 | 9.684159 | 8.65502 | 9.022691 | 9.685079 | 9.90567 | 9.855972 | 9.962825 |
| 10.62314 | 10.49092 | 10.43584 | 10.15082 | 10.06312 | 9.952591 | 10.04074 | 10.03236 | 10.40966 |
| 9.316095 | 9.121323 | 9.374167 | 8.186422 | 8.666836 | 6.671506 | 6.843587 | 6.837726 | 7.494765 |
| 11.85708 | 11.76779 | 11.79094 | 12.1972 | 12.17678 | 12.03335 | 11.9648 | 12.08147 | 11.36439 |
| 6.885015 | 6.105481 | 6.797121 | 7.887794 | 5.25695 | NaN | 6.062361 | 6.445266 | 8.074196 |
| 10.20866 | 10.0677 | 9.965304 | 9.74416 | 10.13231 | 9.86212 | 9.669207 | 10.24183 | 8.135184 |
| 7.041211 | 7.131501 | 5.491594 | 5.114645 | 5.453666 | 8.547434 | 8.343788 | 8.41347 | NaN |
| 7.152548 | 7.08812 | 7.254014 | 7.071949 | 7.186417 | 7.319336 | 7.315542 | 7.125889 | 7.016436 |
| 6.526041 | 7.024408 | 6.82945 | 5.662884 | 5.723794 | 6.509188 | 6.411756 | 6.221115 | 7.237763 |
| NaN | NaN | 4.373765 | 6.319453 | 6.253373 | NaN | 5.978142 | 6.163826 | NaN |
| 8.456806 | 8.771976 | 8.425279 | 5.60092 | 5.885293 | 7.751851 | 6.698675 | 6.378079 | 5.038371 |

| | | | | | | | | |
|---|---|---|---|---|---|---|---|---|
| 9.1115 | 7.778727 | 8.245644 | NaN | NaN | 8.199158 | NaN | NaN | 8.5578 |
| 9.106357 | 9.206295 | 9.137092 | 8.938826 | 9.076745 | 9.155102 | 9.323139 | 9.262366 | 8.574467 |
| 6.719078 | 6.71017 | 6.569735 | 8.044554 | 7.872223 | 7.318413 | 6.960063 | 7.328264 | 6.371523 |
| 11.25603 | 11.22872 | 11.25325 | 10.50348 | 10.35504 | 11.05608 | 11.11889 | 11.10932 | 11.06591 |
| 10.33632 | 10.28879 | 10.27264 | 9.026978 | 9.116584 | 10.58803 | 10.43682 | 10.49704 | 10.53213 |
| 7.051941 | 8.060182 | 8.218242 | 7.320718 | 7.400248 | 7.122339 | 8.037256 | 7.870491 | 6.958068 |
| 9.208676 | 9.298643 | 9.254662 | 9.901582 | 9.657809 | 8.253532 | 8.800518 | 8.62781 | 7.956258 |
| 6.097686 | 6.153722 | 5.165218 | NaN | NaN | 5.704176 | 5.371897 | 5.68473 | 6.29254 |
| 11.78039 | 11.70565 | 11.85568 | 11.6192 | 11.10455 | 9.773263 | 9.927843 | 9.92251 | 10.35805 |
| 9.248819 | 9.040502 | 9.27707 | 8.773557 | 9.094295 | 9.670503 | 9.168225 | 9.250345 | 9.564608 |
| 8.067544 | 7.674882 | 7.435264 | 7.147242 | 5.813542 | 8.335649 | 7.490411 | 7.525533 | 6.537324 |
| 11.73987 | 11.52272 | 11.59706 | 10.90803 | 10.84044 | 10.11286 | 10.13836 | 10.19451 | 11.13126 |
| 11.39461 | 11.30609 | 11.41891 | 9.823231 | 9.730358 | 11.28776 | 11.4257 | 11.4903 | 10.60664 |
| 9.366075 | 9.228442 | 9.244583 | 7.416253 | 7.272972 | 9.029043 | 8.978966 | 9.000398 | 7.836998 |
| 11.89849 | 11.71956 | 11.6967 | 10.06456 | 10.02109 | 11.71827 | 11.70838 | 11.74282 | 10.78437 |
| 11.55308 | 11.41103 | 11.41878 | 10.30207 | 10.17984 | 11.3048 | 11.28312 | 11.34867 | 11.40871 |
| 15.70658 | 15.67106 | 15.70211 | 13.7988 | 13.9003 | 11.52456 | 11.56461 | 11.38339 | 11.99322 |
| 12.10985 | 12.36118 | 11.85886 | 10.34407 | 9.910043 | 7.191711 | 7.855368 | 7.671927 | 8.957165 |
| 12.82951 | 12.66939 | 12.74175 | 10.59587 | 10.31583 | 8.361146 | 8.765418 | 8.435488 | 9.228727 |
| 10.25739 | 10.12407 | 10.22544 | 10.90067 | 10.89405 | 10.77104 | 10.73155 | 10.73634 | 10.89495 |
| 7.987938 | 7.884406 | 7.783446 | 7.438742 | 7.309215 | 8.257767 | 8.282365 | 8.179749 | 9.619398 |
| 8.742498 | 8.736334 | 8.68034 | 8.58713 | 8.397028 | 8.531011 | 8.228572 | 8.493904 | 8.071485 |
| 6.678946 | 6.718072 | 6.568058 | 7.270732 | 7.680806 | 7.205083 | 7.043452 | 7.3353 | 5.854755 |
| 6.879502 | 4.771661 | 8.832179 | NaN | NaN | NaN | NaN | NaN | 6.845362 |
| 6.566209 | 7.517751 | 6.848597 | 6.452915 | 7.363747 | 7.839899 | 7.468843 | 7.476195 | 7.01322 |
| 8.159964 | 8.023396 | 8.121006 | 7.783496 | 7.664474 | 7.696033 | 8.391733 | 8.048568 | 7.486579 |
| 9.720174 | 9.878154 | 9.961319 | 11.25358 | 11.14086 | 10.17507 | 10.28901 | 10.14256 | 10.68035 |
| 7.803458 | 7.68087 | 7.842937 | 6.421209 | 6.70873 | 7.845359 | 7.886896 | 7.809219 | 7.666459 |
| 9.674946 | 9.339136 | 9.407213 | 9.663234 | 9.475653 | 9.403277 | 9.374557 | 9.088794 | 9.349461 |
| 8.900855 | 8.399872 | 7.951973 | 8.174259 | 10.33454 | 7.282943 | 6.418474 | 7.907297 | 7.031438 |
| 7.400361 | 7.410392 | 7.383684 | 7.971926 | 8.082404 | 7.338629 | 7.390361 | 7.337841 | 7.72678 |
| 7.572928 | 7.591704 | 8.127162 | 7.342131 | 7.749926 | 8.030853 | 7.865094 | 8.23244 | 6.897386 |
| 6.592869 | 7.293936 | 6.58378 | 6.789401 | 7.129368 | 6.487052 | 7.167243 | 5.080435 | 7.150918 |
| NaN | 7.295283 | NaN | 7.529899 | 7.099287 | 7.169391 | 7.13115 | 7.127147 | NaN |
| 10.09731 | 9.558269 | 9.624797 | 11.35359 | 11.51017 | 9.934144 | 9.773138 | 9.643469 | 9.477853 |
| 8.260017 | 8.499936 | 8.172267 | 11.03729 | 9.885661 | 7.328058 | 7.841177 | 7.872451 | 6.74123 |
| 11.17224 | 11.03531 | 11.07794 | 12.23596 | 12.29643 | 11.67013 | 11.64812 | 11.61859 | 12.07549 |
| 6.583655 | 6.307812 | 6.524395 | NaN | NaN | NaN | NaN | NaN | NaN |
| 9.344165 | 9.067971 | 9.074533 | 8.705647 | 8.646549 | 8.851058 | 8.756127 | 8.824036 | 8.673738 |
| 10.76192 | 10.73436 | 10.75891 | 11.44591 | 11.41088 | 9.861045 | 10.82677 | 10.1391 | 10.159 |
| 9.330445 | 9.050808 | 9.267556 | 7.713778 | 7.130221 | 9.078738 | 9.420843 | 8.966579 | 8.013712 |
| 8.989676 | 9.117642 | 9.192909 | 8.984038 | 9.23747 | 9.619102 | 9.445437 | 9.408816 | 9.420532 |
| 11.88803 | 11.7289 | 11.79297 | 11.09863 | 11.15073 | 11.95578 | 12.0872 | 12.11641 | 12.34266 |
| 8.033356 | 8.160574 | 7.681645 | 9.377929 | 8.79634 | 7.859894 | 8.336872 | 8.040825 | 6.953749 |
| 11.61658 | 11.44755 | 11.615 | 11.95939 | 11.83154 | 12.69269 | 12.57398 | 12.5076 | 12.58273 |
| 6.251051 | 7.035571 | 7.099338 | NaN | NaN | 5.810111 | 5.354888 | 5.792243 | 6.014262 |
| 11.56399 | 11.81783 | 11.56671 | 11.96548 | 12.01703 | 10.90231 | 11.30482 | 11.10382 | 11.14906 |

| | | | | | | | | |
|---|---|---|---|---|---|---|---|---|
| 8.322838 | 8.229037 | 8.323592 | 8.158837 | 7.84626 | 7.812556 | 7.859315 | 8.124557 | 8.465947 |
| 6.89907 | 7.181992 | 7.20848 | 6.470265 | 7.036023 | 7.329957 | 6.796968 | 7.107975 | 7.460717 |
| 7.434534 | 7.71304 | 7.11559 | 9.407971 | 9.252546 | 7.026939 | 7.04994 | 6.759506 | 8.913278 |
| 6.565691 | 6.32251 | 6.536532 | 6.701663 | 6.739569 | 6.159088 | 6.588043 | 5.914581 | 7.411235 |
| 10.62419 | 10.54143 | 10.63863 | 9.932837 | 9.900889 | 10.98018 | 11.06561 | 11.00797 | 10.76092 |
| 9.657969 | 9.774208 | 9.677321 | 8.876295 | 9.009947 | 9.720336 | 9.606247 | 9.553347 | 9.677217 |
| 9.229681 | 8.784392 | 8.799891 | 6.645411 | 6.480444 | 6.891341 | 6.34563 | 6.811892 | 6.429357 |
| 12.12569 | 12.3934 | 12.47312 | 12.78443 | 12.63217 | 11.07739 | 10.93822 | 10.80213 | 11.2878 |
| 7.655126 | 7.794791 | 7.788975 | 7.102174 | 6.993251 | 7.383521 | 7.43392 | 7.531704 | 7.139308 |
| 8.076435 | 7.705355 | 8.122493 | 6.703075 | 6.426198 | 7.688118 | 7.343234 | 7.724052 | 8.422857 |
| 10.82704 | 10.93207 | 11.04585 | 10.1611 | 10.30544 | 11.14103 | 11.17816 | 11.25456 | 10.60162 |
| 6.326591 | 3.732383 | NaN | NaN | NaN | 5.631229 | 5.870065 | 5.696574 | 4.725525 |
| 10.19561 | 10.35483 | 10.23 | 9.284183 | 9.296835 | 10.09577 | 10.08162 | 10.24965 | 10.02179 |
| 8.969273 | 9.059706 | 8.970543 | 8.899072 | 8.942134 | 9.133302 | 9.169905 | 9.218322 | 8.274481 |
| 8.00192 | 7.859386 | 7.931186 | 8.605096 | 8.424853 | 8.078578 | 8.265728 | 8.32518 | 7.718647 |
| 7.09705 | 7.103167 | NaN | 4.835481 | 8.066635 | NaN | 8.23412 | NaN | 7.549362 |
| NaN | 5.633155 | 4.917936 | 6.734136 | 6.664453 | NaN | NaN | 5.333938 | 5.569243 |
| 8.226286 | 8.423938 | 8.202519 | 8.058559 | 7.819163 | 8.444317 | 8.654187 | 8.557178 | 8.379035 |
| 8.800372 | 8.868818 | 8.717527 | 9.137296 | 8.981192 | 8.561003 | 8.638533 | 8.717668 | 8.632235 |
| 9.239234 | 9.104068 | 8.863804 | 8.183066 | 8.507247 | 9.16983 | 9.041356 | 9.054367 | 7.944939 |
| 10.07628 | 10.04185 | 9.984324 | 10.40012 | 10.36854 | 9.144905 | 9.038098 | 8.986406 | 9.137996 |
| 7.833518 | 7.550439 | 7.366216 | 8.6437 | 8.793243 | 7.980951 | 8.021512 | 7.951275 | 6.882339 |
| 9.684282 | 9.754124 | 10.09629 | 10.31639 | 9.976977 | 9.913136 | 9.931549 | 9.980906 | 9.469658 |
| 7.267735 | 7.175242 | 7.426424 | 6.784141 | 6.98132 | 7.040654 | 7.170794 | 7.246217 | 7.442769 |
| 9.845987 | 9.734128 | 9.841645 | 7.16481 | 7.484153 | 6.276798 | 6.855627 | 6.854705 | 6.875725 |
| 8.111375 | 8.126017 | 8.112856 | 8.325843 | 8.227024 | 7.543244 | 7.35866 | 7.335364 | 7.829862 |
| 8.129996 | 8.320649 | 8.41724 | 7.576489 | 7.559439 | 6.860193 | 7.14255 | 6.995954 | 7.026258 |
| 8.803296 | 8.743788 | 8.554494 | 7.624003 | 7.13043 | 6.818641 | 6.895597 | 7.372306 | 5.936346 |
| 6.765116 | 6.588873 | 6.997822 | 6.765194 | 7.158188 | 5.356208 | 5.728624 | 5.591777 | 5.940796 |
| 9.904599 | 10.02237 | 10.14388 | 11.4831 | 11.21313 | 11.35089 | 11.31538 | 11.47992 | 8.908687 |
| 8.387902 | 7.536646 | 7.744891 | 8.756582 | 8.815275 | 9.173922 | 9.11521 | 9.262733 | 8.522397 |
| 7.157266 | 7.151927 | 7.376782 | 6.669991 | 6.554852 | 7.830584 | 7.431094 | 7.73658 | 4.573721 |
| 7.353338 | 6.62381 | 7.395195 | 8.429634 | 7.897848 | 6.501394 | 6.293228 | 6.689757 | 6.672783 |
| 8.246132 | 8.1054 | 8.15887 | 7.808254 | 7.514072 | 7.48783 | 7.569895 | 7.849988 | 7.746553 |
| 9.653028 | 9.667221 | 9.728235 | 9.093002 | 8.91727 | 9.305098 | 9.337678 | 9.604333 | 8.713942 |
| 8.632909 | 8.432085 | 8.444493 | 8.79209 | 8.685265 | 8.362645 | 8.292439 | 8.339666 | 8.435403 |
| 14.03124 | 13.80225 | 13.9401 | 14.05748 | 13.8739 | 14.50341 | 14.02516 | 14.18469 | 12.51995 |
| 11.74201 | 10.97267 | 10.69727 | 11.23687 | 10.62227 | 10.84409 | 11.17135 | 10.97419 | 9.90384 |
| 11.24333 | 10.92246 | 11.24515 | 9.654089 | 9.606816 | 9.949154 | 10.05574 | 9.853464 | 8.714451 |
| 9.26263 | 9.14153 | 9.155476 | 7.925734 | 7.966738 | 9.411064 | 9.142802 | 9.326238 | 9.202365 |
| 4.574558 | 5.503023 | 5.116467 | 4.958274 | 4.744156 | NaN | 4.223935 | 4.88779 | 5.318361 |
| 6.638605 | 5.68038 | 6.537114 | 6.126888 | 5.102885 | 6.56833 | 6.930057 | 6.9734 | 5.75573 |
| 10.32469 | 10.24029 | 10.31295 | 8.822779 | 8.477592 | 10.06601 | 10.11456 | 10.09291 | 9.913742 |
| 9.726874 | 9.594882 | 9.797358 | 8.651675 | 8.905606 | 8.826857 | 8.773455 | 9.003971 | 8.251743 |
| 7.681113 | 7.991704 | 8.04053 | 7.677453 | 7.291403 | 6.988658 | 7.31111 | 7.175537 | 6.324315 |
| 13.59274 | 13.67914 | 13.76398 | 14.02646 | 13.5555 | 11.1338 | 11.27114 | 11.25953 | 11.14767 |
| 7.793147 | 7.404558 | 7.402805 | 10.82669 | 10.2944 | 7.245944 | 7.111502 | 7.143923 | 7.222212 |

| | | | | | | | | |
|---|---|---|---|---|---|---|---|---|
| 7.998177 | 8.354557 | 7.716697 | 7.170173 | 7.124693 | 6.968692 | 6.915508 | 7.048052 | 7.253637 |
| 6.959353 | 7.081117 | 7.287622 | 6.435245 | 6.153899 | 6.927476 | 6.7542 | 7.09386 | 6.734073 |
| 9.039073 | 9.018817 | 9.088117 | 8.792912 | 8.588286 | 9.324247 | 9.493005 | 9.324985 | 8.823315 |
| NaN | NaN | NaN | NaN | NaN | NaN | NaN | NaN | 5.920344 |
| 9.832285 | 9.734588 | 9.573572 | 12.14596 | 11.58919 | 10.25942 | 10.21352 | 10.51171 | 10.8507 |
| 5.013198 | 4.602492 | 5.102413 | 4.572183 | 5.113165 | 5.028636 | NaN | 4.854714 | 5.865325 |
| 7.613173 | 7.798021 | 7.862883 | 5.946532 | 6.277603 | 7.248257 | 7.083666 | 7.842066 | 6.622652 |
| 8.928535 | 8.914847 | 8.934366 | 10.40196 | 10.07091 | 9.620935 | 9.632021 | 9.567691 | 7.834488 |
| NaN | 5.894162 | NaN | NaN | NaN | NaN | NaN | 4.937473 | 4.511983 |
| 8.699513 | 8.400975 | 8.680225 | 7.155571 | 7.585444 | 9.062972 | 9.08654 | 8.877298 | 7.750062 |
| 6.099165 | 5.245123 | 6.137065 | 3.830898 | NaN | 5.714036 | 6.187854 | 5.598672 | 5.486004 |
| 6.471879 | NaN | 4.738517 | 5.880987 | 5.694024 | 6.327437 | 5.945779 | 6.015992 | 6.437675 |
| 6.886992 | 5.803054 | 7.071757 | 6.734189 | 6.876954 | 7.277742 | 7.076195 | 7.336265 | 6.033982 |
| 9.359675 | 9.260648 | 9.210188 | 9.060298 | 9.125256 | 9.754892 | 9.672283 | 9.859156 | 9.548875 |
| NaN | NaN | NaN | 5.443146 | 4.099159 | 5.116109 | NaN | NaN | 5.328991 |
| NaN | NaN | NaN | NaN | NaN | NaN | 3.826466 | NaN | 5.477343 |
| 5.860771 | 6.003194 | 5.983974 | 6.430066 | 6.258581 | 6.597157 | 6.628671 | 6.673229 | 6.4759 |
| 9.48806 | 9.362264 | 9.269915 | 10.28649 | 10.58257 | 9.668275 | 9.414442 | 9.575864 | 7.922347 |
| 8.571437 | 8.448345 | 8.38514 | 6.944008 | 7.030241 | 8.40901 | 8.411112 | 8.318663 | 7.565012 |
| 8.017145 | 8.088503 | 8.102722 | 7.133576 | 7.262855 | 7.725299 | 7.848263 | 7.717275 | 8.271858 |
| NaN | 0.078165 | 0.462668 | 0.238801 | 2.377481 | NaN | 3.529969 | NaN | NaN |
| 12.02443 | 12.0324 | 12.08818 | 13.84027 | 13.91334 | 11.73626 | 11.8801 | 11.79459 | 11.7373 |
| 9.085741 | 9.133674 | 9.155097 | 7.740713 | 7.756833 | 9.039865 | 9.10533 | 9.151514 | 9.009065 |
| 9.018332 | 8.967162 | 9.04432 | 8.056917 | 7.856574 | 8.93291 | 8.73937 | 9.075761 | 9.130422 |
| 8.452502 | 8.335812 | 8.286325 | 7.305052 | 7.095883 | 8.258938 | 8.490217 | 8.584431 | 8.162484 |
| 8.153014 | 8.227295 | 8.256174 | 8.041529 | 8.209095 | 7.562597 | 7.428949 | 7.434725 | 7.241535 |
| 7.984994 | 8.068014 | 8.030048 | 7.361162 | 7.538634 | 6.933396 | 7.556852 | 7.728898 | 10.10231 |
| 10.79786 | 10.03287 | 9.804283 | 6.686633 | 5.663061 | NaN | 6.997683 | NaN | 6.54001 |
| 11.76231 | 11.7702 | 11.89028 | 12.30295 | 12.23187 | 11.76806 | 11.74987 | 11.84583 | 11.6741 |
| 10.79933 | 10.59635 | 10.74813 | 10.57405 | 10.3875 | 10.717 | 10.68104 | 10.67412 | 11.00089 |
| 8.367208 | 8.274261 | 8.407924 | 6.949194 | 6.994942 | 7.675596 | 7.621668 | 7.631375 | 8.380878 |
| 7.284976 | 6.155207 | 5.981036 | 7.293433 | 8.029027 | NaN | 6.975134 | 6.11488 | 7.363831 |
| 6.100677 | 6.769507 | 6.934933 | NaN | NaN | 6.452317 | 6.059066 | 6.35981 | 6.328976 |
| 6.84651 | 6.907269 | 6.975245 | 5.982996 | 5.710476 | 5.897793 | 7.419956 | 5.967419 | 6.776726 |
| 7.524893 | 7.824258 | 7.709734 | 8.987042 | 8.653517 | 7.214491 | 7.748137 | 7.354732 | 7.611187 |
| 10.68724 | 10.79396 | 10.71613 | 9.969343 | 9.998481 | 9.723284 | 9.832014 | 10.01648 | 10.41874 |
| 10.3596 | 10.25252 | 10.34106 | 9.78516 | 9.710824 | 11.37392 | 10.96297 | 11.13398 | 10.47553 |
| 8.04026 | 7.954984 | 7.981853 | 7.992815 | 8.015059 | 8.409233 | 8.37525 | 8.276461 | 8.226847 |
| 7.332996 | 6.960655 | 7.025629 | 6.352153 | 6.252561 | 6.956439 | 7.203813 | 7.358629 | 7.125807 |
| 6.13687 | 7.862924 | 6.984655 | 6.638125 | 5.26492 | 7.908509 | 8.256136 | 8.501951 | 8.055156 |
| 8.886865 | 7.991389 | 8.109464 | NaN | 6.711117 | 8.663631 | 8.456844 | 8.618109 | NaN |
| 9.631431 | 9.690921 | 9.630361 | 9.726126 | 9.575324 | 9.030466 | 9.13138 | 9.097865 | 9.626916 |
| 6.599155 | 6.302662 | 6.323933 | 7.661884 | 7.734274 | 6.208059 | 6.084088 | 6.086916 | 7.094571 |
| 9.985896 | 9.820652 | 9.869392 | 9.988853 | 10.13865 | 10.31135 | 10.06381 | 10.07556 | 9.970285 |
| 12.6051 | 12.52631 | 12.63313 | 12.53925 | 12.54908 | 12.80031 | 13.10957 | 13.08661 | 12.1254 |
| 9.649309 | 9.757534 | 9.676036 | 8.879632 | 8.816599 | 9.357582 | 9.140511 | 9.31614 | 9.204642 |
| 7.712118 | 8.024212 | 6.226418 | 8.554584 | 8.36747 | 6.394897 | 7.67639 | 7.701566 | 6.350339 |

| | | | | | | | | |
|---|---|---|---|---|---|---|---|---|
| 5.732434 | 6.883354 | 6.258486 | 5.399357 | 6.391175 | 5.271714 | 5.323405 | 5.287353 | 5.33465 |
| 7.749776 | 8.068624 | 8.096659 | 10.56832 | 10.73528 | 7.61161 | 7.305976 | 7.397535 | 5.771855 |
| 7.183541 | 6.975448 | 6.874592 | 7.765233 | 7.349081 | 7.245 | 7.176961 | 7.403742 | 7.241525 |
| 15.68326 | 9.135081 | 9.321843 | 13.38621 | 13.06879 | 10.08024 | 10.25682 | 10.00413 | NaN |
| 7.468043 | 6.880818 | 7.134416 | NaN | 6.611135 | NaN | NaN | NaN | 6.297142 |
| 8.327199 | 7.703953 | 8.47366 | 8.062193 | 8.322195 | 8.366953 | 8.212006 | 8.332646 | 8.144361 |
| 9.585688 | 9.524094 | 8.814649 | 10.00799 | 9.98409 | 9.94532 | 9.725249 | 9.666288 | 9.140342 |
| 8.595761 | 8.618367 | 8.621371 | 7.651043 | 7.605259 | 8.235349 | 8.246037 | 8.362512 | 7.849969 |
| 8.283628 | 8.276642 | 8.260274 | 7.376218 | 6.880629 | 8.636414 | 8.644954 | 8.627428 | 8.375949 |
| 4.260786 | NaN | NaN | NaN | 4.470535 | 6.126239 | 5.685926 | 5.396636 | NaN |
| 11.52664 | 11.32516 | 11.41209 | 12.62796 | 12.60544 | 11.60729 | 11.50767 | 11.61013 | 12.33558 |
| 10.65194 | 10.54135 | 10.61634 | 9.165487 | 9.170956 | 10.494 | 10.5745 | 10.35217 | 9.811351 |
| 9.168051 | 9.153611 | 9.186848 | 9.269327 | 8.736105 | 8.670876 | 8.554615 | 8.609234 | 8.663453 |
| 7.579317 | 7.6337 | 7.334182 | 7.865855 | 7.802447 | 6.821601 | 7.011191 | 7.327827 | 7.144465 |
| 8.313011 | 8.5769 | 8.234003 | 10.49974 | 10.26396 | 7.272772 | 8.016467 | 7.48551 | 8.377232 |
| 5.443847 | 6.1523 | 5.507266 | NaN | 3.678295 | 6.996492 | 7.107607 | 7.196249 | 5.878008 |
| 7.024706 | 6.966321 | 6.938267 | 6.715265 | 6.622254 | 6.671317 | 6.592793 | 6.75017 | 7.040876 |
| 5.954624 | 4.823963 | 7.3073 | 6.457877 | 7.143123 | NaN | 5.83042 | NaN | 7.373093 |
| 7.574578 | 7.149054 | 7.481913 | 6.738657 | 7.143639 | 7.266984 | 7.558333 | 7.475007 | 7.235103 |
| NaN | 4.416899 | 1.530572 | 4.439758 | 4.218054 | 4.78105 | 3.435179 | 3.685253 | 3.135909 |
| 7.549037 | 7.477446 | 7.534373 | 6.396453 | 6.139698 | 7.67304 | 7.580035 | 7.918537 | 7.324393 |
| 8.99802 | 9.086472 | 8.530975 | 7.02859 | 7.511119 | 7.724103 | 7.735264 | 7.540145 | 5.350264 |
| 6.302812 | 5.785465 | 5.86171 | 4.698614 | 5.917227 | 6.244112 | 6.857367 | 6.780693 | 6.62572 |
| 9.285393 | 9.266075 | 9.196276 | 7.81843 | 7.666147 | 9.485461 | 9.337859 | 9.400201 | 8.766713 |
| 8.395265 | 8.119742 | 8.280446 | 8.853518 | 8.800051 | 8.332709 | 8.262867 | 8.259845 | 8.15425 |
| NaN | NaN | NaN | NaN | NaN | NaN | NaN | NaN | 5.817086 |
| 9.822036 | 9.689475 | 9.832463 | 10.54918 | 10.47327 | 10.25095 | 10.32419 | 10.25005 | 9.836205 |
| 6.934825 | 5.418111 | 6.492964 | 5.900121 | 5.565471 | 6.329082 | 6.554728 | 6.288353 | 6.306616 |
| 9.196013 | 9.096793 | 9.258986 | 8.868119 | 8.787467 | 8.509917 | 8.298045 | 8.370851 | 8.068143 |
| 8.009233 | 8.338256 | 8.406107 | 8.31354 | 8.562797 | 8.024074 | 7.410772 | 8.399876 | 8.224888 |
| 6.013706 | 6.047021 | 5.901951 | 7.218055 | 7.062044 | 3.867831 | 5.26983 | 5.025418 | 5.691963 |
| 9.803685 | 9.794864 | 10.1145 | 8.608261 | 8.673183 | 10.12738 | 10.00341 | 10.15082 | 8.757213 |
| 6.813213 | 6.97138 | 7.128293 | 5.81231 | 6.002443 | 6.382909 | 6.381964 | 6.673242 | 7.155548 |
| 7.557782 | 7.876038 | 7.544893 | 7.176064 | 7.428447 | 8.136846 | 7.952528 | 8.136275 | 7.240626 |
| 10.3088 | 10.41376 | 10.34956 | 10.68497 | 10.41694 | 10.23477 | 10.20933 | 10.19733 | 9.666502 |
| 10.06929 | 9.875335 | 9.958365 | 10.03317 | 10.246 | 10.34509 | 10.29893 | 10.37807 | 8.959952 |
| 6.891166 | 7.087448 | 6.674509 | NaN | 6.608525 | 6.853087 | NaN | 7.375383 | 7.532658 |
| 6.0248 | 6.205383 | 6.563956 | 3.880847 | 3.743717 | 7.664132 | 7.148687 | 7.688564 | 7.234488 |
| 8.643124 | 8.572482 | 8.647586 | 8.202897 | 8.207515 | 7.426023 | 7.622193 | 8.07559 | 6.730079 |
| 7.436014 | 7.158326 | 7.03467 | 5.966684 | 5.85885 | 7.74336 | 7.82059 | 7.767107 | 6.515755 |
| 9.051817 | 8.987635 | 9.271813 | 9.378076 | 9.987866 | 9.045058 | 8.737011 | 8.826529 | 7.986322 |
| 12.24138 | 12.14033 | 12.15106 | 11.87209 | 11.7987 | 11.73874 | 11.75506 | 11.79272 | 12.15891 |
| 6.206527 | 6.18007 | 6.129298 | 9.319831 | 9.080693 | NaN | 6.321543 | 6.364957 | 6.201506 |
| 6.880473 | 8.215435 | 7.287325 | 6.198903 | 5.560206 | 6.789154 | 7.253021 | 7.456716 | 6.51658 |
| 9.086562 | 7.868417 | 7.939777 | 7.633998 | 7.390419 | 6.866885 | NaN | NaN | 7.182456 |
| 6.590596 | 6.207777 | 5.292999 | 7.5191 | 8.389639 | NaN | NaN | NaN | NaN |
| 6.90296 | 6.926038 | 7.064148 | 6.179758 | 6.981003 | 6.809568 | 7.074004 | 6.815637 | 6.727241 |

| | | | | | | | | |
|---|---|---|---|---|---|---|---|---|
| 7.766368 | 7.87511 | 8.057897 | 10.33326 | 9.918638 | 8.385345 | 8.155429 | 8.195292 | 7.424352 |
| 7.778348 | 7.489306 | 7.58647 | 7.224892 | 7.160324 | 7.555672 | 7.507188 | 7.501603 | 7.308402 |
| 7.681753 | 7.611659 | 7.240135 | 9.812213 | 9.031077 | 8.073212 | 7.944684 | 7.868331 | 8.734556 |
| 7.980505 | 7.825275 | 7.93081 | 6.610764 | 6.870525 | 8.161625 | 7.891521 | 8.037484 | 8.018105 |
| 10.84546 | 10.7355 | 10.84652 | 10.34494 | 10.31057 | 10.8814 | 10.9715 | 10.88436 | 10.25564 |
| 8.903684 | 8.890587 | 9.009477 | 8.775784 | 8.459266 | 8.831747 | 8.936809 | 8.808264 | 8.453702 |
| NaN | NaN | NaN | 5.300679 | 5.591865 | NaN | NaN | NaN | 5.155582 |
| 10.07315 | 10.18811 | 10.15844 | 10.09074 | 10.17308 | 10.65897 | 10.74611 | 10.73294 | 11.01773 |
| 6.935879 | 6.899897 | 6.993635 | 5.86417 | 6.126933 | 6.971823 | 6.914761 | 7.093368 | 6.56416 |
| NaN | NaN | NaN | 7.599745 | 8.038265 | NaN | 6.148014 | NaN | 7.289237 |
| 7.083527 | 6.795465 | 7.196152 | 7.419466 | 7.298135 | 8.01672 | 7.92191 | 7.846689 | 7.801646 |
| 8.548505 | 8.667731 | 8.708138 | 8.971432 | 9.244098 | 9.106106 | 8.969688 | 8.765357 | 7.680677 |
| NaN | NaN | 6.303004 | 7.383536 | 7.517263 | 7.615112 | 7.467654 | 7.675699 | 5.891148 |
| 6.820945 | 7.201124 | 7.489557 | 8.759001 | 8.698849 | 8.094684 | 8.070244 | 8.235856 | 9.396169 |
| 12.30803 | 12.23441 | 12.25658 | 12.28775 | 12.47849 | 11.99817 | 12.15461 | 12.07227 | 12.1542 |
| NaN | NaN | NaN | NaN | NaN | 4.517459 | 4.880783 | NaN | 5.492059 |
| 6.500955 | 6.315671 | 5.904016 | 6.370093 | 7.035394 | NaN | NaN | NaN | 8.379073 |
| 7.745595 | 7.782429 | 7.80947 | 7.317093 | 7.130141 | 7.998195 | 7.985721 | 7.630149 | 8.455444 |
| 6.797729 | 7.031593 | 6.427613 | NaN | 6.474824 | 7.177585 | 7.887288 | 7.751836 | 6.655958 |
| 5.113671 | 5.278336 | 4.641664 | NaN | 5.11516 | 5.18031 | 5.381671 | 5.841156 | 5.950497 |
| 4.52 | 4.483525 | 4.338075 | 6.265207 | 6.13553 | NaN | 3.886606 | 3.651765 | 6.656079 |
| 8.372383 | 8.291186 | 8.517303 | 6.837304 | 7.180393 | 8.190914 | 8.438046 | 8.294952 | 8.375648 |
| 8.273528 | 8.33129 | 8.327249 | 7.6243 | 7.648731 | 8.256666 | 8.069175 | 8.085979 | 7.769818 |
| 9.867229 | 9.824748 | 9.754043 | 10.79143 | 10.9341 | 10.2768 | 10.48949 | 10.35091 | 8.960543 |
| 9.636055 | 9.585488 | 9.777153 | 9.296903 | 9.101208 | 9.340654 | 9.200109 | 9.35153 | 9.001049 |
| 8.962876 | 8.973371 | 8.936943 | 8.323566 | 8.396385 | 8.922214 | 8.921084 | 9.221759 | 9.056187 |
| 9.103658 | 8.89099 | 8.97715 | 8.086763 | 7.806184 | 8.531828 | 8.450727 | 8.421114 | 9.070611 |
| 7.569401 | 7.380059 | 7.740067 | 7.997231 | 7.951013 | 7.499563 | 7.273959 | 7.544298 | 6.581842 |
| 10.68226 | 10.64131 | 10.85519 | 11.36015 | 11.4347 | 10.65196 | 10.94863 | 10.90111 | 10.44447 |
| 11.27569 | 11.14624 | 11.20407 | 10.5519 | 10.65174 | 11.66045 | 11.54604 | 11.65064 | 11.64892 |
| 11.20237 | 11.20664 | 11.1812 | 12.3014 | 12.19451 | 11.03428 | 11.19817 | 11.30214 | 11.16131 |
| 5.714417 | 4.831929 | 4.991883 | 6.119578 | 5.85786 | NaN | NaN | NaN | 4.296004 |
| 7.209866 | 7.721748 | 7.424675 | 6.406675 | 6.494073 | 8.045622 | 7.625247 | 7.94529 | 4.903236 |
| 8.622343 | 8.61309 | 8.580181 | 8.589205 | 8.532526 | 8.776398 | 8.786926 | 8.976996 | 8.139849 |
| 13.63463 | 13.70229 | 13.83469 | 14.02976 | 14.01282 | 13.23701 | 13.1574 | 13.13303 | 13.57807 |
| 5.564428 | 5.829681 | 4.958656 | 4.96068 | 5.381201 | 5.279215 | 5.314183 | 5.378015 | 5.202266 |
| 7.782174 | 7.837895 | 7.604721 | 9.68407 | 9.444264 | 9.233313 | 8.915707 | 9.508387 | 8.550813 |
| 7.453224 | 7.268293 | 7.27091 | 6.929289 | 7.267084 | 6.779274 | 7.863427 | 7.820782 | 6.994217 |
| 9.831084 | 9.80738 | 9.839248 | 8.321478 | 8.330616 | 9.508661 | 9.556211 | 9.510714 | 9.423194 |
| 11.26996 | 11.44455 | 11.41207 | 10.85351 | 10.73136 | 12.07363 | 11.82573 | 11.91849 | 10.98371 |
| 7.791591 | 6.727849 | 6.890961 | 6.411075 | 6.167453 | NaN | NaN | NaN | 4.930742 |
| 7.213393 | 5.139426 | 8.170609 | 7.850108 | 8.093148 | 9.295461 | 9.057344 | 9.175864 | 7.61163 |
| 6.43645 | 6.11288 | 6.431186 | 6.670533 | 7.102312 | 6.887614 | 5.472873 | 6.702501 | 7.354885 |
| 7.377128 | 7.037982 | 7.268212 | 6.518425 | NaN | 6.973486 | 6.612862 | 6.663241 | 7.563625 |
| 9.047834 | 9.096707 | 9.064238 | 9.418746 | 9.03488 | 7.735177 | 7.613497 | 7.972379 | 8.486725 |
| 6.925523 | 6.794181 | 6.763363 | 5.469762 | 5.420159 | 6.849549 | 6.747738 | 6.8566 | 7.037261 |
| 13.20612 | 13.07631 | 13.13067 | 13.22295 | 13.15798 | 12.77096 | 12.82633 | 12.70513 | 12.82045 |

-Bruker Confidential-

| | | | | | | | | |
|---|---|---|---|---|---|---|---|---|
| 7.574137 | 7.457513 | 7.319613 | 6.252407 | 6.708045 | 7.315612 | 7.016292 | 7.191981 | 7.236119 |
| 10.03538 | 9.970313 | 10.04055 | 11.07787 | 11.25801 | 10.28312 | 10.39559 | 10.24411 | 10.61733 |
| 7.173071 | 7.00099 | 7.154808 | NaN | NaN | NaN | NaN | NaN | NaN |
| 5.804775 | 7.620587 | 6.26945 | 8.900282 | 8.909638 | 7.509001 | 7.275036 | 7.461811 | 8.698854 |
| 7.727721 | 7.283608 | 7.504584 | 6.679945 | 6.747914 | 7.938071 | 7.993811 | 7.697239 | NaN |
| 9.230117 | 9.102469 | 9.061013 | 8.39706 | 8.448149 | 9.139558 | 9.23347 | 9.071448 | 9.531771 |
| 7.628522 | 7.531482 | 7.517107 | 6.441076 | 6.50639 | 8.052821 | 7.786158 | 7.911473 | 7.512722 |
| 7.858136 | 7.977235 | 8.12005 | 7.667856 | 7.854796 | 8.857575 | 8.79771 | 8.77557 | 8.580774 |
| 10.59471 | 10.66073 | 10.66621 | 11.24153 | 11.01389 | 10.94394 | 10.92792 | 11.07207 | 11.08976 |
| 6.676987 | 7.19853 | 6.894282 | 8.293397 | 8.860522 | 4.455816 | 6.093103 | 4.95495 | 3.509572 |
| 7.605167 | 7.670096 | 7.878956 | 7.687383 | 8.114193 | 6.908984 | 7.440852 | 6.380689 | 8.59006 |
| 8.207036 | 7.262936 | 5.734513 | 7.993151 | NaN | 7.539433 | 7.484437 | 8.025554 | 7.590739 |
| 7.334108 | 7.383076 | 7.715802 | 7.093287 | 7.272197 | 7.383751 | 7.378415 | 7.402752 | 6.589969 |
| 6.281618 | 6.492877 | 6.17169 | 4.992777 | NaN | 6.655228 | 6.852521 | 6.515811 | 5.936054 |
| 7.424139 | 7.301143 | 7.320397 | 6.919146 | 6.807479 | 6.987665 | 6.889438 | 6.906963 | 5.750732 |
| NaN | 3.312626 | NaN | 3.938431 | 4.991127 | 6.145896 | 6.004175 | 4.22171 | 3.414774 |
| NaN | NaN | NaN | 7.18142 | 6.645515 | 5.514357 | NaN | 4.869643 | 5.627219 |
| 10.71626 | 10.41066 | 10.36553 | 8.979957 | 8.950042 | 9.931807 | 9.866758 | 10.0159 | 9.843654 |
| 13.77315 | 13.58875 | 13.66464 | 13.71207 | 13.72696 | 13.62254 | 13.38813 | 13.50103 | 12.91151 |
| 9.211473 | 9.112718 | 9.251064 | 10.36335 | 10.36237 | 7.948904 | 7.420437 | 7.125421 | 7.827843 |
| 5.247708 | 5.603597 | 5.414593 | 5.781923 | 5.205943 | 8.361333 | 8.099532 | 6.95831 | 6.631864 |
| 8.473125 | 8.751144 | 8.55193 | 7.848633 | 7.476521 | 8.951349 | 9.181113 | 9.248805 | 8.064353 |
| 5.304189 | 3.190119 | 2.691301 | 4.577829 | NaN | 5.256563 | 5.939264 | 6.055048 | 5.852382 |
| 5.381524 | NaN | 4.772743 | 5.425809 | 6.071996 | 5.05935 | 4.967223 | 5.440946 | 4.344855 |
| 5.058449 | 5.201108 | 5.013534 | 4.896163 | 4.265999 | 5.866543 | 5.805091 | 5.883563 | 6.252392 |
| 8.361039 | 8.202374 | 8.467019 | 7.057319 | 7.113826 | 7.74655 | 7.719127 | 7.739026 | 8.265363 |
| 8.695264 | 8.518876 | 8.464834 | 7.536738 | 7.694643 | 8.45962 | 8.422463 | 8.446404 | 8.393984 |
| 3.018448 | NaN | 3.935461 | 3.32089 | NaN | NaN | NaN | NaN | 6.314183 |
| 5.802229 | 5.390217 | 5.081854 | NaN | NaN | 7.056149 | 5.5436 | 6.806361 | 6.1191 |
| 9.802001 | 9.938269 | 10.04748 | 10.68679 | 10.80357 | 9.997472 | 9.61543 | 9.683804 | 10.50465 |
| 11.37152 | 11.19795 | 11.16393 | 11.252 | 11.4085 | 11.54375 | 11.4602 | 11.36966 | 10.41521 |
| 6.38503 | 5.841452 | 5.861424 | 7.258073 | 7.2158 | 6.546204 | 6.330906 | 6.526371 | 4.97904 |
| 8.529572 | 8.862071 | 8.608465 | 8.600115 | 7.734767 | 7.460795 | 7.598608 | 7.290462 | 6.321541 |
| 10.66267 | 10.61847 | 10.54082 | 10.94445 | 10.97987 | 11.0019 | 10.58667 | 10.66783 | 10.79454 |
| 8.699862 | 8.806748 | 8.54262 | 9.222241 | 9.239882 | 9.401849 | 9.140658 | 9.215237 | 8.985731 |
| 9.126382 | 8.91902 | 9.003227 | 9.566829 | 9.574696 | 9.354201 | 9.572459 | 9.529983 | 7.388687 |
| 8.403827 | 6.283393 | 6.473775 | 5.230327 | 5.389996 | NaN | 6.573123 | 6.027241 | 5.890509 |
| 7.203743 | 7.369714 | 7.435234 | 6.968676 | 6.92785 | 7.191027 | 7.232812 | 7.139198 | 7.255536 |
| 9.146474 | 9.235637 | 9.230569 | 7.76433 | 8.01682 | 8.34282 | 8.795106 | 8.722411 | 9.241702 |
| 6.609344 | 6.502652 | 6.281399 | 6.248514 | 6.506466 | 6.573007 | 6.165071 | 6.241956 | 5.731021 |
| 8.070765 | 8.574563 | 8.396954 | 7.322493 | 7.67254 | 8.244578 | 8.335533 | 7.970201 | 7.757094 |
| 8.152236 | 7.918726 | 8.099592 | 6.841721 | 6.853459 | 7.447411 | 7.177726 | 7.243113 | 7.717445 |
| 6.800385 | 7.179797 | 6.362237 | 6.526731 | 8.14804 | NaN | 7.006971 | 6.337575 | 6.37378 |
| 10.71388 | 9.378644 | 9.460044 | 8.598713 | 8.257257 | NaN | 6.59021 | NaN | 7.452941 |
| 7.448528 | 7.219068 | 7.024738 | 5.999512 | 6.384089 | 7.352589 | 7.342563 | 7.527576 | 7.336707 |
| 5.533493 | 5.505639 | 5.514179 | NaN | 4.395113 | 5.429324 | 5.414845 | 5.85019 | 6.31986 |
| 9.825665 | 9.895912 | 10.18897 | 10.1171 | 10.33604 | 10.10574 | 10.11447 | 9.965621 | 10.75738 |

| | | | | | | | | |
|---|---|---|---|---|---|---|---|---|
| 7.301621 | 7.512959 | 7.103199 | 6.586663 | 6.969887 | 6.620412 | 7.017043 | 6.96438 | 6.599147 |
| NaN | NaN | NaN | 6.153239 | 5.502617 | NaN | NaN | NaN | NaN |
| 8.823556 | 8.621815 | 8.476745 | 8.634764 | 8.88032 | 8.757728 | 8.099731 | 8.136049 | 7.768651 |
| 9.509218 | 9.613873 | 9.655112 | 10.2088 | 10.15962 | 9.827518 | 10.00946 | 10.20587 | 10.00875 |
| 8.641122 | 8.561201 | 8.684923 | 9.416763 | 9.109058 | 8.022276 | 8.079382 | 7.981466 | 7.771954 |
| 13.00106 | 12.91779 | 13.03345 | 13.12861 | 13.08893 | 13.03977 | 12.92297 | 12.8642 | 12.13932 |
| 11.10518 | 11.07028 | 10.76328 | 11.48698 | 11.5979 | 10.37025 | 10.34051 | 10.39383 | 9.764822 |
| 9.276338 | 9.083561 | 9.153318 | 10.04898 | 9.568905 | 8.449274 | 8.609924 | 8.688417 | 7.5879 |
| 9.047379 | 8.90099 | 8.756272 | 9.666642 | 9.066043 | 8.681479 | 8.681143 | 8.349936 | 7.947231 |
| 8.792702 | 8.730681 | 8.71658 | 9.086974 | 8.787214 | 7.653547 | 8.012918 | 8.231711 | 8.016088 |
| 5.128202 | NaN | 4.681652 | NaN | NaN | NaN | NaN | NaN | 6.134105 |
| 5.252676 | 5.488148 | 5.804814 | 3.596315 | NaN | 5.228111 | 5.471417 | 5.575625 | 4.47137 |
| 8.308411 | 8.098261 | 8.300651 | 6.269125 | 6.805502 | 7.814095 | 7.740639 | 7.132237 | 8.143028 |
| 6.799047 | 6.8821 | 6.502905 | NaN | NaN | NaN | 7.158502 | 6.9959 | 7.04513 |
| 8.887632 | 8.888537 | 8.95036 | 9.414276 | 9.399862 | 8.324468 | 8.563355 | 8.485253 | 8.78858 |
| 10.49039 | 10.2316 | 10.3116 | 9.148625 | 9.236553 | 10.21088 | 10.11803 | 10.17927 | 10.03945 |
| 8.348482 | 8.48261 | 8.470781 | 7.90443 | 7.718395 | 8.921005 | 8.921463 | 8.982122 | 8.491871 |
| 7.750566 | 7.845161 | 7.548256 | 6.273134 | 6.672733 | 7.275596 | 6.759556 | 6.535408 | 7.332675 |
| 8.805005 | 8.973941 | 8.758276 | 9.371763 | 9.742228 | 8.279123 | 8.526288 | 8.537663 | 8.888036 |
| 11.05601 | 10.57919 | 10.68283 | 10.81277 | 10.87351 | 11.23552 | 10.84599 | 11.22739 | 10.37644 |
| 7.003365 | 7.078656 | 6.888803 | NaN | 5.402276 | 7.103911 | 7.036879 | 6.558863 | 6.522344 |
| 8.424998 | 8.376952 | 8.330629 | 8.699366 | 8.66308 | 8.303144 | 8.107667 | 8.392709 | 7.507308 |
| 8.635753 | 8.593802 | 8.539519 | 8.666724 | 8.253789 | 8.53756 | 8.620815 | 8.298244 | 8.073738 |
| 9.958976 | 9.89616 | 10.07098 | 10.54865 | 10.21806 | 11.33736 | 11.11973 | 11.02803 | 10.96486 |
| 10.32422 | 10.44875 | 10.52226 | 11.25671 | 11.15639 | 10.27781 | 10.33478 | 10.27908 | 10.98831 |
| 10.31433 | 10.16946 | 10.15403 | 8.970705 | 9.019824 | 9.74236 | 9.703059 | 9.677931 | 10.42271 |
| 8.626338 | 8.605331 | 8.607899 | 7.626534 | 7.64876 | 8.616422 | 8.612612 | 8.724621 | 8.845515 |
| 11.87939 | 11.84185 | 11.69545 | 10.74219 | 10.78135 | 12.0032 | 11.86647 | 11.90059 | 11.91748 |
| 8.431016 | 8.531427 | 8.407397 | 8.742436 | 8.940998 | 7.78172 | 7.722381 | 7.95163 | 7.992394 |
| 8.943319 | 8.824093 | 8.929149 | 7.346856 | 7.310957 | 9.135606 | 9.193526 | 9.104639 | 7.550759 |
| 7.702703 | 7.446846 | 7.848117 | NaN | NaN | 7.060189 | 7.938966 | 7.440946 | 6.667861 |
| NaN | 6.076651 | 5.606363 | 4.278432 | 6.020839 | 6.098402 | 6.154399 | 6.146604 | 5.523427 |
| 7.549883 | 7.477878 | 8.050364 | 8.015779 | 8.061096 | 7.261267 | 7.56399 | 7.348663 | 6.584488 |
| NaN | 6.218513 | NaN | 6.964264 | 6.502588 | NaN | NaN | NaN | 6.351511 |
| 13.21679 | 13.25939 | 13.3362 | 12.75227 | 12.26195 | 12.75447 | 12.8243 | 13.01362 | 12.53038 |
| 8.411859 | 8.641314 | 8.419906 | 7.665842 | 7.888037 | 9.086969 | 8.955071 | 8.830876 | 8.578437 |
| 7.992074 | 7.733219 | 7.955127 | 7.879396 | 7.749011 | 8.021448 | 8.006599 | 8.000937 | 7.581978 |
| 8.587484 | 9.974823 | 8.80528 | 6.08872 | 5.955798 | 9.631103 | 9.719188 | 9.685539 | 7.127824 |
| 11.11588 | 10.94177 | 10.86036 | 10.75726 | 10.65839 | 9.939939 | 9.705806 | 10.00945 | 10.24971 |
| 5.59589 | 6.604825 | 5.721108 | 3.557192 | 3.80813 | 6.263657 | 5.516016 | 5.353402 | 3.876192 |
| 9.467441 | 9.352671 | 9.373403 | 7.359047 | 7.597611 | 9.874487 | 9.943786 | 9.674314 | 9.681039 |
| 6.55635 | 6.147573 | 6.2818 | 6.282856 | 6.177101 | 6.812199 | 6.988729 | 7.119225 | NaN |
| 9.511392 | 9.317496 | 9.522843 | 8.629654 | 8.553423 | 8.730966 | 8.656957 | 8.81093 | 9.148374 |
| 9.915285 | 9.825492 | 9.971853 | 9.814231 | 9.288557 | 9.756737 | 9.570161 | 9.800905 | 9.670952 |
| 6.850542 | 6.18457 | 7.12921 | NaN | NaN | 6.103606 | NaN | 6.953058 | 6.784241 |
| 7.017862 | 7.21685 | 7.488266 | 6.692267 | 5.197597 | 7.539856 | 7.58533 | 7.336241 | 8.062564 |
| 9.520812 | 9.665605 | 9.615014 | 7.866068 | 8.331407 | 9.236981 | 9.230306 | 9.198878 | 9.68099 |

| | | | | | | | | |
|---|---|---|---|---|---|---|---|---|
| 11.1805 | 11.12015 | 11.01845 | 12.7251 | 12.66179 | 12.28355 | 12.15997 | 12.3197 | 12.64906 |
| 9.857374 | 9.783566 | 9.845441 | 12.33763 | 12.24183 | 9.014209 | 9.199762 | 9.336799 | 9.289087 |
| 11.74059 | 11.76851 | 11.82184 | 11.13622 | 11.24195 | 12.71469 | 12.56804 | 12.52601 | 12.44688 |
| 6.555292 | 6.586836 | 6.799186 | 6.379033 | 6.190255 | 6.854748 | 6.342371 | 7.281605 | 5.677273 |
| 8.729901 | 12.0922 | 8.160225 | 7.567834 | 7.715139 | 8.97561 | 8.939973 | 8.922195 | 8.586617 |
| 10.03018 | 10.16939 | 10.14562 | 10.79692 | 11.12136 | 9.506369 | 9.348737 | 9.578285 | 8.96669 |
| 9.726161 | 9.548894 | 9.537271 | 11.07582 | 11.2351 | 8.563803 | 8.89314 | 8.685131 | 8.267001 |
| 11.86099 | 11.57396 | 11.62178 | 12.04309 | 11.47279 | 10.14208 | 10.33036 | 10.2618 | 9.598911 |
| 9.027636 | 9.040016 | 9.01851 | 9.250535 | 8.953306 | 8.326846 | 8.22051 | 7.944281 | 7.301558 |
| 9.278867 | 9.243678 | 9.199586 | 8.318712 | 8.558543 | 8.997056 | 9.132436 | 9.006065 | 8.66531 |
| 8.370101 | 8.785701 | 8.572317 | 7.624275 | 7.671481 | 9.260103 | 9.243602 | 9.050806 | 8.645081 |
| 6.550622 | 6.205035 | 6.467802 | 8.006142 | 7.373019 | 7.119311 | 7.084126 | 6.663556 | NaN |
| NaN | 6.234864 | NaN | 5.154269 | 5.178408 | 6.245002 | 6.432878 | 6.61183 | 6.03888 |
| 7.464993 | 7.43258 | 7.487572 | 7.200035 | 7.076336 | 7.751493 | 7.709785 | 7.744819 | 8.180112 |
| 6.75417 | 7.02377 | 7.217752 | NaN | NaN | 6.556032 | 6.093028 | 6.480595 | 6.597591 |
| 9.024024 | 9.168897 | 9.311261 | 7.677444 | 7.782215 | 8.961468 | 8.955802 | 9.186221 | 9.146359 |
| 8.122232 | 7.314672 | 8.46866 | 7.012402 | 7.86097 | 8.60259 | 8.053628 | 7.873322 | 6.432788 |
| 10.0227 | 9.832301 | 10.00511 | 9.203749 | 9.140855 | 10.47624 | 10.45484 | 10.44296 | 9.754547 |
| 7.859177 | 7.728562 | 7.556423 | 7.929704 | 8.078991 | 7.713943 | 7.596367 | 7.63869 | 7.177685 |
| 6.01076 | 5.250436 | 5.418561 | NaN | 4.434141 | 5.955652 | 5.736641 | 5.455703 | 5.299338 |
| 6.550424 | 6.743983 | 6.011265 | 6.40883 | 6.07628 | 6.167474 | 6.363523 | 5.414287 | 6.522271 |
| 4.800971 | 5.754744 | 6.118952 | 5.149809 | 6.207276 | NaN | NaN | 4.984323 | 5.651143 |
| 8.138761 | 8.027467 | 8.092731 | 7.880294 | 7.791795 | 8.550198 | 8.633645 | 8.512693 | 8.379036 |
| 6.788703 | 6.819311 | 6.008404 | 7.620907 | 7.345943 | 6.257813 | 5.770152 | 6.346245 | 7.539468 |
| 5.967061 | 6.020118 | 6.346684 | 6.090107 | 6.538147 | 6.467479 | 6.618716 | 7.034458 | 6.589826 |
| 6.524439 | 6.939798 | 7.210463 | 6.786237 | 7.098507 | 6.230528 | 6.191326 | 6.553791 | 6.603935 |
| 8.462466 | 8.281376 | 8.273836 | 7.198536 | 7.49141 | 8.461095 | 8.336007 | 8.157 | 8.378668 |
| 8.912797 | 8.869566 | 8.611173 | 9.01828 | 8.830351 | 9.021501 | 9.297582 | 9.132759 | 8.937611 |
| 7.393967 | NaN | 6.381435 | 7.12324 | 7.577342 | NaN | 6.903435 | 7.469188 | 8.074588 |
| 6.765592 | 6.531496 | 7.163705 | 5.028602 | 3.806243 | 6.961789 | 7.166929 | 6.96957 | 6.949439 |
| 7.537148 | 7.414317 | 7.394683 | 6.715025 | 6.094243 | 7.514916 | 7.553456 | 7.737175 | 7.007862 |
| 5.724408 | NaN | 5.317016 | NaN | 6.095453 | NaN | NaN | 5.334403 | 6.443005 |
| 6.482335 | 7.251929 | 6.1597 | 7.023942 | 7.430608 | 6.978656 | 6.806045 | 6.909674 | 6.488145 |
| 8.48158 | 8.290801 | 8.189263 | 8.368124 | 8.342047 | 7.051694 | 6.674844 | 6.921749 | 8.763759 |
| 5.457986 | 6.098874 | 5.424458 | 4.694107 | 7.290636 | 5.747763 | 6.063612 | 6.08262 | 5.775228 |
| 9.841895 | 9.915815 | 9.878894 | 10.5744 | 10.38808 | 11.05133 | 10.8011 | 10.90685 | 10.30382 |
| NaN | NaN | NaN | 5.800405 | 5.318696 | NaN | 6.152072 | NaN | 4.461263 |
| 9.390311 | 8.38565 | 9.331474 | 7.420941 | 7.149611 | 6.956809 | 6.61617 | 6.322457 | 6.953728 |
| NaN | 5.117719 | NaN | 4.90855 | 5.427007 | NaN | 5.063786 | 5.479739 | 5.500005 |
| 5.748517 | 5.812085 | 6.071497 | 4.209266 | 5.460768 | 6.403386 | 6.389333 | 6.565129 | 5.701048 |
| 11.61404 | 11.38398 | 11.59644 | 12.95982 | 12.80473 | 11.43437 | 11.30931 | 11.2624 | 12.12843 |
| 13.11813 | 12.81372 | 12.79732 | 14.56254 | 14.21735 | 12.13644 | 12.07088 | 12.23755 | 13.73696 |
| 12.89575 | 12.67299 | 12.87114 | 14.60454 | 14.52156 | 13.50132 | 13.50107 | 13.47911 | 13.57516 |
| 9.649633 | 9.532586 | 9.793151 | 8.610559 | 8.514495 | 9.187501 | 9.313821 | 9.606281 | 8.613256 |
| 7.764603 | 7.426003 | 7.460635 | 7.721791 | 7.493309 | 8.457503 | 8.267991 | 8.553549 | 6.609188 |
| 7.0561 | 6.521739 | 6.895036 | 6.397709 | 6.266038 | 6.827185 | 6.787578 | 6.761098 | 4.776432 |
| 11.53648 | 11.47364 | 11.56377 | 12.28709 | 12.13901 | 10.8359 | 10.87634 | 11.00803 | 11.13335 |

| | | | | | | | | |
|---|---|---|---|---|---|---|---|---|
| 12.3555 | 12.27341 | 12.37607 | 13.83537 | 13.91075 | 12.8335 | 13.06736 | 12.9462 | 12.51288 |
| 8.971826 | 8.902279 | 8.986009 | 10.35997 | 10.34953 | 9.572278 | 9.555501 | 9.737425 | 9.790783 |
| 10.89827 | 10.98519 | 11.02273 | 11.7969 | 11.92101 | 10.66347 | 10.80492 | 10.84276 | 10.53706 |
| 8.949553 | 9.155818 | 9.128696 | 11.47595 | 11.29699 | 9.883368 | 10.48736 | 10.24507 | 8.25578 |
| 6.172828 | 5.034254 | 5.302358 | 7.427084 | 8.311817 | 5.246413 | 4.665797 | 5.010571 | 4.028903 |
| 11.14645 | 11.08886 | 11.27741 | 12.16183 | 12.32935 | 11.22016 | 10.49577 | 11.05418 | 10.16058 |
| 12.66281 | 12.68052 | 12.7649 | 13.7522 | 13.5589 | 12.12455 | 12.37835 | 12.40665 | 12.47064 |
| 9.290484 | 9.21034 | 9.311044 | 9.766045 | 9.454708 | 9.742896 | 9.475592 | 9.482118 | 9.503386 |
| 9.399884 | 9.275285 | 9.371771 | 9.344476 | 9.472014 | 9.00196 | 9.04759 | 8.831094 | 8.605583 |
| 9.026239 | 9.018152 | 8.958593 | 9.74223 | 9.59342 | 9.424738 | 9.332255 | 9.129786 | 9.013793 |
| 9.094985 | 8.90223 | 8.904131 | 10.69851 | 10.80588 | 9.255968 | 9.096538 | 9.015737 | 8.238735 |
| 12.60559 | 12.5395 | 12.59863 | 12.60252 | 12.63295 | 12.67235 | 12.83881 | 12.81601 | 12.78669 |
| 11.12077 | 10.92992 | 11.02632 | 9.606444 | 9.682465 | 11.21607 | 11.4401 | 11.36028 | 10.92518 |
| 6.209432 | 6.489838 | 4.147551 | NaN | 3.514775 | NaN | 5.893351 | NaN | 6.555716 |
| 4.790873 | 5.38691 | 5.504485 | 4.159655 | 6.819576 | 5.508787 | 4.558349 | 5.44627 | 5.445207 |
| 8.531985 | 8.346344 | 8.050442 | 9.997818 | 10.15687 | 7.836194 | 8.443813 | 8.343316 | 8.030786 |
| 8.701338 | 9.691179 | 9.1483 | 8.659925 | 8.851104 | 9.203519 | 9.229879 | 9.557931 | 8.701743 |
| 7.05623 | 6.830009 | 6.60222 | 5.757839 | 6.312617 | 6.940595 | 6.689868 | 7.1121 | 6.414521 |
| 8.813104 | 8.894532 | 8.728561 | 8.456958 | 8.606069 | 9.019405 | 9.065351 | 8.933599 | 7.92569 |
| 8.457408 | 8.213557 | 8.407242 | 8.543736 | 8.687461 | 9.168813 | 9.010077 | 9.236117 | 8.545745 |
| 5.93483 | 5.822272 | NaN | 5.04802 | 5.567341 | 7.56912 | 6.859103 | 6.963001 | NaN |
| 6.912282 | 6.74565 | 6.750653 | 8.299401 | 8.224424 | 6.990041 | 6.844825 | 7.151222 | 7.645028 |
| 6.939244 | 6.765997 | 6.812057 | 6.522419 | 5.665485 | 7.364107 | 7.58147 | 7.609291 | 7.087844 |
| 6.978428 | 6.924546 | 6.923925 | 7.813272 | 7.810178 | 6.762867 | 6.806433 | 6.825134 | 7.02156 |
| 10.80616 | 10.55668 | 10.72178 | 10.92866 | 10.58104 | 10.20268 | 10.36858 | 10.0028 | 10.50906 |
| 6.034349 | 7.106096 | 7.472013 | NaN | NaN | 7.529397 | 7.230268 | 7.574331 | 7.2189 |
| NaN | NaN | NaN | 6.779819 | 7.14784 | NaN | 7.600166 | 6.382974 | 6.492348 |
| 8.2134 | 8.059332 | 8.22268 | 8.01888 | 8.102799 | 7.947764 | 7.825958 | 7.840463 | 8.061725 |
| 7.534013 | 7.306733 | 7.239532 | 6.358908 | 6.580661 | 7.32338 | 7.190334 | 7.255175 | 7.276597 |
| 9.003323 | 8.120478 | 7.372212 | NaN | 8.206692 | NaN | NaN | NaN | NaN |
| 7.099185 | 7.91652 | 4.741724 | 7.503785 | 7.708793 | NaN | NaN | NaN | 5.934125 |
| 8.913374 | 8.823419 | 8.855826 | 8.023335 | 8.150309 | 8.792746 | 8.499579 | 8.758064 | 8.476922 |
| 10.65255 | 10.87098 | 10.76236 | 10.2689 | 10.314 | 10.89129 | 10.86251 | 10.88771 | 10.25285 |
| 11.51393 | 11.4154 | 11.46093 | 10.87801 | 10.90976 | 12.0943 | 12.22516 | 12.43016 | 11.41346 |
| 8.327341 | 7.957798 | 8.493817 | 8.086893 | 8.095796 | 5.590667 | 6.262971 | 6.171228 | 6.568903 |
| 5.735813 | 5.971343 | 6.485447 | NaN | NaN | NaN | NaN | NaN | 5.826326 |
| 9.181591 | 9.197411 | 9.179115 | 8.643564 | 8.573082 | 9.593811 | 9.825232 | 9.838267 | 9.365918 |
| 10.1525 | 10.07153 | 10.19293 | 9.672909 | 9.929976 | 10.34805 | 10.31399 | 10.34743 | 9.288029 |
| 7.714218 | 7.5895 | 7.928612 | 6.458484 | 7.084189 | 7.328956 | 7.409794 | 7.453957 | 7.546522 |
| 7.281152 | 7.282784 | 6.998553 | 6.742501 | 6.311837 | 6.579405 | 7.290373 | 7.344191 | 7.140399 |
| 8.076989 | 8.133093 | 7.986993 | 7.697926 | 7.898355 | 7.987859 | 7.842294 | 7.957959 | 7.941455 |
| 9.55496 | 9.40871 | 9.454394 | 8.572696 | 8.387535 | 9.492878 | 9.419497 | 9.501282 | 9.415102 |
| 6.307096 | 6.222389 | 6.287225 | 5.596635 | 5.097864 | 6.48822 | 6.366289 | 6.54704 | 6.7522 |
| 6.066444 | 5.235023 | 6.019767 | 5.368943 | NaN | 6.224454 | 6.451477 | 6.746784 | 6.107411 |
| 9.690494 | 9.810318 | 9.786366 | 8.127528 | 8.203662 | 9.930286 | 9.942843 | 9.964025 | 9.536567 |
| 8.766312 | 8.566584 | 8.61598 | 8.027401 | 8.477491 | 9.123663 | 9.193748 | 9.202788 | 8.760612 |
| 8.260394 | 8.299427 | 7.968077 | 8.476736 | 8.341172 | 8.908072 | 8.576591 | 8.662048 | 9.548393 |

-Bruker Confidential-

| | | | | | | | | |
|---|---|---|---|---|---|---|---|---|
| 11.4735 | 11.53933 | 11.72836 | 11.53926 | 11.68602 | 11.19975 | 10.85504 | 10.85337 | 10.42953 |
| 11.96374 | 11.86613 | 12.09463 | 12.30523 | 12.32212 | 11.55801 | 11.65 | 11.36942 | 11.99753 |
| 9.28201 | 9.083993 | 9.081492 | 7.650587 | 7.489956 | 6.173921 | 6.429158 | 6.716278 | 6.590816 |
| 11.92968 | 11.98224 | 11.96858 | 12.5505 | 11.63371 | 11.79131 | 11.71815 | 11.84175 | 11.52326 |
| 10.10171 | 10.22507 | 10.24482 | 9.364056 | 9.723438 | 10.69595 | 10.47174 | 10.46238 | 9.0465 |
| 11.24449 | 11.33211 | 11.3275 | 11.29926 | 11.25708 | 12.0994 | 11.89178 | 11.98132 | 11.28118 |
| 8.970542 | 9.45501 | 9.156622 | 7.45583 | 7.447105 | 5.596166 | 5.684245 | NaN | 6.335428 |
| 9.260943 | 9.238049 | 9.213829 | 8.081988 | 8.268669 | 9.433331 | 9.387345 | 9.345994 | 8.42318 |
| 11.9814 | 11.88486 | 11.91839 | 12.77556 | 12.72579 | 12.53252 | 12.53504 | 12.49138 | 11.73044 |
| 7.858482 | 7.879474 | 7.903476 | 8.141196 | 8.304003 | 8.689581 | 8.398552 | 8.152909 | 8.103697 |
| 8.222144 | 7.816727 | 8.266229 | 6.050209 | 5.996935 | 8.192009 | 7.928882 | 7.795371 | 6.971252 |
| 6.021873 | 5.020945 | 5.274176 | NaN | 6.173061 | NaN | 6.710379 | 6.249989 | 6.80911 |
| 6.147971 | 4.929373 | 5.571548 | NaN | NaN | NaN | 4.70751 | NaN | 6.293097 |
| 5.137966 | 5.047615 | 4.809174 | 4.842023 | 4.814334 | 6.078329 | 6.506747 | 6.362689 | 6.165541 |
| 8.06743 | 8.30667 | 8.141733 | 8.085464 | 7.801426 | 7.952505 | 7.857806 | 7.953634 | 8.336473 |
| 6.385648 | 6.600874 | 5.976497 | 6.436198 | 6.52949 | 6.831471 | 6.810593 | 7.073048 | 4.656206 |
| 6.366343 | 6.664478 | 6.863788 | 6.413363 | 6.854356 | 7.185012 | 6.646457 | 7.021092 | 6.154276 |
| 7.224034 | 6.012356 | 6.96606 | 6.365558 | 6.061806 | 7.154591 | 6.750914 | 6.248557 | 5.803037 |
| 9.417706 | 9.064004 | 8.959518 | 10.02828 | 10.25606 | 7.912322 | 7.972816 | 8.460337 | 8.469372 |
| 6.987524 | 7.323736 | 7.321007 | 6.419244 | 6.478974 | 7.143159 | 7.297293 | 7.298329 | 6.489239 |
| 9.601613 | 9.580014 | 9.577539 | 8.910605 | 8.697652 | 10.19466 | 10.18712 | 10.15247 | 10.01073 |
| 8.335873 | 8.175299 | 8.233214 | 9.162513 | 9.046097 | 8.042988 | 7.479751 | 7.568594 | 7.886615 |
| 8.940561 | 8.725514 | 8.882609 | NaN | NaN | NaN | NaN | NaN | NaN |
| 8.878189 | 8.884604 | 9.004758 | 9.479589 | 9.826816 | 7.79283 | 7.741948 | 7.271579 | 7.586226 |
| NaN | NaN | NaN | NaN | NaN | 8.226201 | 8.049832 | 7.821726 | 6.867601 |
| 5.739236 | 4.838108 | 5.594191 | 4.535863 | 4.959605 | 5.838183 | 5.634204 | 5.699191 | 5.543574 |
| 9.274331 | 9.325281 | 9.478381 | 8.140878 | 8.202496 | 7.407533 | 7.016371 | 7.354877 | 9.129083 |
| 5.177249 | 4.631008 | 5.24579 | NaN | NaN | 5.502617 | 6.014375 | 5.96079 | 5.038448 |
| 5.415353 | NaN | 6.55369 | NaN | NaN | NaN | NaN | NaN | 6.184272 |
| 6.946274 | 7.199645 | 7.282307 | NaN | 6.547307 | NaN | 6.355117 | 6.689999 | 6.198045 |
| 9.196658 | 9.101509 | 9.107252 | 7.705743 | 7.692413 | 8.333182 | 8.50301 | 8.535438 | 8.536994 |
| 9.620857 | 9.509283 | 9.560225 | 7.294208 | 7.340937 | 8.218907 | 8.516703 | 8.421304 | 8.93736 |
| 9.340513 | 9.158998 | 9.238752 | 6.947849 | 6.94158 | 8.223363 | 8.327775 | 8.292996 | 8.646374 |
| 7.882124 | 7.903939 | 8.010447 | 8.532522 | 8.647573 | 7.176827 | 7.343129 | 7.656816 | 6.408834 |
| 7.591593 | 7.712372 | 7.555588 | 6.833933 | 7.136125 | 7.835202 | 7.559673 | 7.601231 | 7.868404 |
| 7.84006 | 7.862326 | 7.958583 | 6.860045 | 7.161446 | 7.827841 | 7.84165 | 7.849465 | 8.464658 |
| 7.436492 | 7.101102 | 7.09909 | 6.314408 | 6.164347 | 7.750268 | 7.738955 | 7.714529 | 5.677634 |
| 7.835603 | 7.28957 | 7.711344 | 7.269261 | 6.49265 | 7.941468 | 8.300256 | 8.284234 | 8.610974 |
| 7.434001 | 7.769056 | NaN | NaN | NaN | 8.938332 | 10.65577 | 10.71237 | 7.139747 |
| 10.81783 | 10.76619 | 10.70037 | 10.25493 | 10.29967 | 10.53026 | 10.61642 | 10.54631 | 10.76783 |
| 7.805731 | 7.668024 | 7.689455 | 6.967512 | 6.350096 | 7.40824 | 7.679013 | 7.968216 | 6.909436 |
| 7.447315 | 7.352228 | 7.541418 | 6.88088 | 6.716271 | 8.493489 | 8.154416 | 8.126476 | 7.479488 |
| 8.115601 | 7.429811 | 7.188366 | 7.156083 | 7.691225 | NaN | 6.838777 | 6.12738 | 6.981789 |
| 8.132538 | 7.797761 | 7.809375 | 6.332343 | 6.039901 | 7.978525 | 7.863505 | 7.747673 | 6.411489 |
| 12.47011 | 12.28318 | 12.26379 | 13.18272 | 13.38204 | 12.93159 | 12.5887 | 13.15383 | 12.23758 |
| 10.00312 | 9.775757 | 9.818234 | 9.399738 | 9.164417 | 10.13938 | 10.41499 | 10.45984 | 9.789471 |
| 8.85432 | 8.939495 | 8.736445 | 8.003839 | 7.807681 | 9.250187 | 9.249722 | 9.089387 | 9.036497 |

-Bruker Confidential-

| | | | | | | | | |
|---|---|---|---|---|---|---|---|---|
| 6.388014 | 6.393508 | 6.087443 | NaN | NaN | 7.568408 | 6.876928 | 7.415824 | 6.418096 |
| 12.16292 | 11.9704 | 12.09807 | 12.82327 | 12.76643 | 12.24502 | 12.29474 | 12.46037 | 11.84871 |
| 6.725586 | 6.708292 | 6.712569 | 8.321671 | 8.576302 | 6.64817 | 6.533943 | 6.908982 | 6.079581 |
| 6.346253 | 6.318636 | 6.195416 | 5.90138 | 6.285447 | 6.235384 | 5.804074 | 6.395202 | 6.87246 |
| 8.738801 | 8.710652 | 8.725603 | 9.713874 | 9.527487 | 9.125968 | 9.017036 | 9.149428 | 8.531137 |
| 9.902394 | 9.910735 | 9.715251 | 10.47845 | 10.55255 | 9.404388 | 9.597452 | 9.164302 | 9.172958 |
| 8.604558 | 8.27809 | 8.646935 | 7.012729 | 7.290944 | 7.809078 | 7.711546 | 7.745979 | 8.054549 |
| 8.249625 | 8.330074 | 7.699657 | 7.108797 | 8.182688 | 8.012171 | 8.032322 | 7.911269 | 8.112162 |
| 7.045454 | 6.832872 | 6.922118 | 6.331585 | 6.482119 | 6.594335 | 6.341602 | 6.59974 | 6.760675 |
| 10.09062 | 9.245479 | 9.540703 | 11.95228 | 11.76411 | 9.649705 | 10.15093 | 10.21595 | 10.39569 |
| 6.855492 | 7.503996 | 6.942785 | 8.207755 | 7.836474 | 6.533369 | 6.641354 | 6.370132 | 7.359774 |
| 6.434049 | 4.269219 | 4.408782 | 5.320046 | 6.432161 | 6.81628 | 6.876439 | 6.966985 | 4.444168 |
| 6.950604 | 7.142832 | 6.806347 | 6.364919 | 5.896759 | 6.791271 | 6.635714 | 6.622116 | 6.802788 |
| 5.586398 | NaN | 6.415399 | NaN | NaN | NaN | NaN | NaN | NaN |
| 6.730053 | 6.854414 | 7.098799 | 7.266387 | 7.290631 | 7.175736 | 7.572654 | 7.721517 | 6.857982 |
| 6.876716 | 6.533824 | 6.620115 | 5.816987 | 5.471264 | 6.615685 | 6.83008 | 6.45911 | 7.159756 |
| 14.06346 | 13.97987 | 14.05094 | 10.43214 | 10.46479 | 8.631973 | 8.698361 | 8.805935 | 8.786265 |
| 8.459543 | 8.681743 | 8.559153 | 5.220727 | 5.703775 | NaN | NaN | 6.068943 | 7.066612 |
| 7.742548 | 7.343988 | 7.22477 | 7.00131 | 6.613924 | 6.577586 | 6.718504 | 6.643832 | 7.6062 |
| 7.677801 | 7.473559 | 7.483505 | 6.615727 | 7.054634 | 7.981861 | 7.728882 | 8.505177 | 7.05376 |
| 8.443622 | 8.456755 | 8.522596 | 8.005511 | 7.915004 | 8.95537 | 8.896587 | 8.905783 | 9.544084 |
| 11.75648 | 11.73833 | 11.77138 | 11.28519 | 11.40699 | 11.91218 | 11.91747 | 11.93239 | 11.76792 |
| 9.722778 | 9.764634 | 9.758973 | 8.504776 | 8.497252 | 9.109008 | 9.131992 | 9.239777 | 9.91666 |
| 6.082542 | 6.190402 | 5.280945 | 4.516205 | NaN | 6.349485 | 6.274929 | 6.14149 | 5.41963 |
| 9.501799 | 9.574586 | 9.589378 | 9.605007 | 9.681368 | 10.11382 | 10.17456 | 10.06942 | 9.584673 |
| 8.677366 | 8.473303 | 8.630481 | 9.142524 | 8.860634 | 8.380274 | 8.451299 | 8.337787 | 7.874583 |
| 9.42137 | 9.351397 | 9.395624 | 8.384588 | 8.489142 | 9.699718 | 9.60105 | 9.664122 | 9.648442 |
| 8.263607 | 8.458578 | 7.824911 | 7.253493 | 7.821693 | 8.906174 | 9.057923 | 9.059042 | 7.462443 |
| 11.57279 | 11.58462 | 11.73307 | 13.01522 | 12.85772 | 10.70295 | 10.62274 | 10.73816 | 12.09671 |
| 8.193674 | 8.325941 | 8.151316 | 8.25491 | 8.444504 | 8.401172 | 8.435985 | 8.464126 | 8.441607 |
| 9.413707 | 9.499408 | 9.475071 | 9.395787 | 9.214166 | 9.989307 | 10.0321 | 10.14269 | 10.08522 |
| 8.340914 | NaN | NaN | 6.031256 | 5.546436 | NaN | NaN | NaN | NaN |
| 7.436695 | NaN | 2.301013 | 6.442282 | 6.840986 | 6.886855 | 6.209314 | 5.578846 | 6.598378 |
| 8.702468 | 6.862772 | 6.923818 | 8.45176 | NaN | NaN | 5.576345 | 5.76999 | 6.061934 |
| 7.792744 | 7.721181 | 7.942219 | 7.438703 | 7.41465 | 7.797982 | 8.098863 | 8.457085 | 7.220077 |
| 14.66121 | 14.48402 | 14.54687 | 12.14602 | 12.16935 | 10.17004 | 10.23122 | 10.25895 | 10.44518 |
| 7.780407 | 7.628069 | 7.504643 | 7.925927 | 7.973966 | 6.612171 | 6.704013 | 6.861052 | 7.644219 |
| 11.23267 | 11.19212 | 11.17673 | 11.39787 | 11.26316 | 10.94198 | 11.06343 | 11.28604 | 11.20726 |
| 6.990506 | 6.855132 | 6.87484 | 4.982976 | NaN | 6.610801 | 6.46474 | 5.650614 | 6.286269 |
| 8.223786 | 8.146653 | 8.107328 | 9.029134 | 8.882904 | 7.326092 | 7.313229 | 7.684504 | 7.434425 |
| 10.51778 | 10.52347 | 10.33591 | 11.54576 | 11.48483 | 9.665419 | 10.31902 | 10.02501 | 10.31438 |
| 8.90729 | 8.732357 | 8.647056 | 8.019016 | 7.754148 | 8.886951 | 8.823781 | 8.832594 | 8.454023 |
| 12.26553 | 12.06251 | 12.166 | 12.8175 | 12.07844 | 11.78215 | 11.64388 | 11.34079 | 10.91176 |
| 10.22104 | 10.3196 | 10.20071 | 11.1341 | 11.15939 | 10.19575 | 10.2866 | 10.20358 | 9.42962 |
| 6.687064 | 6.65561 | 7.160244 | 5.704033 | 6.273822 | 7.254604 | 6.775061 | 6.81758 | 6.624435 |
| 10.85149 | 10.77132 | 10.7249 | 10.3584 | 10.85421 | 10.98625 | 11.4847 | 11.74721 | 11.19514 |
| 8.194949 | 8.212613 | 8.088428 | 7.613111 | 7.9481 | 7.92938 | 7.961396 | 8.023741 | 7.867446 |

-Bruker Confidential-

| | | | | | | | | |
|---|---|---|---|---|---|---|---|---|
| 9.123023 | 9.011561 | 9.028515 | 8.462062 | 8.545296 | 9.171432 | 9.188117 | 9.145459 | 9.036486 |
| 8.800252 | 8.594734 | 8.553804 | 8.778504 | 8.785302 | 8.850465 | 8.741342 | 8.598368 | 7.972967 |
| 8.527761 | 8.274748 | 8.293366 | 7.79347 | 7.343801 | 8.871815 | 8.809994 | 8.514676 | 7.73884 |
| 7.676244 | 7.815279 | 7.242981 | 6.091864 | 6.086857 | 6.778894 | 6.771549 | 6.943286 | 6.139284 |
| 8.507259 | 7.149108 | 6.843573 | 6.409815 | NaN | 7.070133 | 6.151883 | 6.557016 | 8.027512 |
| 8.440949 | 8.432607 | 8.689152 | 7.997149 | 7.994771 | 8.618237 | 8.668345 | 8.68267 | 7.874372 |
| 7.113036 | 7.439318 | 6.611995 | 7.405332 | 8.014474 | 6.50824 | 6.143505 | 6.615039 | 8.166698 |
| 9.992636 | 9.963625 | 9.835539 | 10.21966 | 10.28208 | 10.26466 | 10.08561 | 10.27651 | 8.834519 |
| 8.395744 | 8.433129 | 8.321684 | 7.597737 | 7.551159 | 7.968576 | 8.523338 | 8.707038 | 8.259742 |
| 8.531866 | 8.782419 | 8.698492 | 9.211084 | 8.885875 | 8.309081 | 8.431461 | 8.078265 | 8.804839 |
| 8.992367 | 8.503245 | 8.504878 | 9.492788 | 9.38052 | 8.474585 | 8.523019 | 8.598415 | 9.685744 |
| 8.93075 | 8.815459 | 9.013994 | 10.09385 | 9.834743 | 9.172738 | 8.96182 | 8.9653 | 8.91183 |
| 8.781042 | 8.65862 | 8.69717 | 7.243639 | 7.15162 | 7.940679 | 7.877237 | 7.826933 | 8.656878 |
| 8.884371 | 8.878934 | 8.989635 | 10.81727 | 10.50417 | 8.564229 | 8.701734 | 8.641907 | 8.884301 |
| 9.858088 | 9.615924 | 9.635795 | 10.53401 | 10.20379 | 9.402662 | 9.401542 | 9.32413 | 9.253322 |
| 7.77325 | 8.246233 | 8.308559 | 10.35157 | 10.4869 | 7.569883 | 7.160306 | 6.908285 | 7.170939 |
| 9.747217 | 9.772524 | 9.812965 | 10.05445 | 10.06713 | 10.04481 | 10.0041 | 10.08123 | 10.33109 |
| 11.34652 | 11.46559 | 11.43322 | 12.25533 | 12.61382 | 10.67479 | 10.74828 | 10.76751 | 11.28113 |
| 3.914982 | 3.705657 | 4.225958 | 3.901727 | NaN | 4.468254 | 6.020081 | 4.190462 | 4.837777 |
| 7.102776 | 7.366166 | 7.712851 | 6.859836 | 7.222026 | 6.647662 | 6.563762 | 6.462405 | 6.787767 |
| NaN | NaN | 5.629949 | NaN | NaN | 6.805681 | 6.538973 | 6.620591 | 6.581169 |
| 10.8122 | 11.58703 | 11.22406 | 11.05173 | 11.83073 | 11.66181 | 11.54884 | 11.53741 | 11.77755 |
| 7.713873 | 7.064066 | 7.526094 | 6.721845 | 6.680076 | 6.640746 | 6.386458 | 6.251367 | 7.113245 |
| 9.897362 | 9.802169 | 9.863179 | 9.183475 | 9.170586 | 10.29767 | 10.21644 | 10.29622 | 9.119246 |
| 12.15041 | 12.07831 | 12.07491 | 11.93097 | 11.85605 | 11.94894 | 12.11072 | 12.04078 | 11.90447 |
| 8.96165 | 8.823666 | 8.918451 | 8.105628 | 8.293431 | 8.796687 | 8.715377 | 8.708704 | 9.049879 |
| NaN | 7.429615 | NaN | 7.641491 | 7.414332 | 6.339942 | 6.727247 | NaN | 6.219921 |
| 10.01407 | 9.85517 | 9.858832 | 11.07232 | 11.06321 | 9.852864 | 9.443508 | 10.00517 | 10.16642 |
| 9.700672 | 9.684264 | 9.640671 | 8.67759 | 8.501238 | 9.666238 | 9.745625 | 9.719372 | 9.639351 |
| 7.29398 | 7.553181 | 7.53601 | 6.08687 | 7.28936 | 7.311612 | 7.360404 | 7.280525 | 7.538568 |
| 9.923354 | 9.815403 | 9.859007 | 8.490119 | 8.159449 | 9.384082 | 8.920684 | 9.424193 | 9.499506 |
| 9.940445 | 9.856031 | 10.03517 | 11.88663 | 11.40276 | 10.86944 | 10.47708 | 10.54723 | 10.3665 |
| 9.129839 | 9.142361 | 9.164781 | 7.814268 | 7.734967 | 8.622306 | 8.718837 | 8.781852 | 8.880761 |
| 5.818235 | 5.851586 | 6.636005 | 6.686957 | 6.843801 | 5.996382 | 5.831558 | 5.944163 | 6.756019 |
| NaN | NaN | NaN | 7.484686 | NaN | NaN | NaN | NaN | 6.607074 |
| 9.827087 | 9.983039 | 9.814554 | 9.927218 | 9.846989 | 10.11585 | 10.26665 | 10.20917 | 10.02663 |
| 8.011235 | 9.340754 | 7.916601 | 7.433469 | 7.248067 | 7.524299 | 7.177351 | 7.459633 | 8.206493 |
| 11.57648 | 11.46467 | 11.62815 | 11.15741 | 11.20234 | 11.22414 | 11.27642 | 11.10515 | 11.30699 |
| 7.96636 | 7.85746 | 7.990475 | 7.513216 | 7.424612 | 7.960924 | 7.715271 | 7.809256 | 7.748453 |
| 6.556779 | 6.350352 | 6.301281 | 5.853465 | 5.741655 | 6.090795 | 5.669128 | 5.856013 | 6.323241 |
| 7.298048 | 7.342236 | 7.433425 | 7.385318 | 7.509233 | 7.467802 | 7.36766 | 7.509695 | 6.796617 |
| 8.896149 | 8.706262 | 8.946524 | 8.53542 | 8.280643 | 8.819572 | 9.25061 | 8.832149 | 8.648346 |
| 7.662489 | 7.934564 | 7.410644 | 7.075665 | 7.594264 | 7.152727 | 7.060241 | 6.860423 | 7.846701 |
| 6.847336 | 7.053331 | 6.93977 | 6.329067 | 7.156176 | 6.05444 | 6.383437 | 6.045624 | 6.605435 |
| 7.702168 | 7.952961 | 7.674475 | 6.508575 | 6.891219 | 8.182758 | 7.950453 | 7.980027 | 7.763758 |
| 6.530934 | 6.499466 | 6.464231 | 6.854697 | 6.672616 | 7.359253 | 7.564048 | 7.21067 | 6.739184 |
| 7.11743 | 7.385102 | 7.182016 | 7.903668 | 7.967648 | 6.887409 | 7.067786 | 7.090971 | 7.78547 |

| | | | | | | | | |
|---|---|---|---|---|---|---|---|---|
| 10.62429 | 10.58242 | 10.33836 | 11.3849 | 11.04397 | 10.931 | 11.26416 | 10.9627 | 10.07189 |
| 12.84382 | 12.65468 | 12.70667 | 13.95829 | 13.98623 | 12.88829 | 12.90935 | 12.74518 | 12.30943 |
| 8.076743 | 7.944745 | 8.157938 | 6.77675 | 6.715649 | 7.203566 | 7.127654 | 7.207274 | 7.46006 |
| 7.214746 | 6.858483 | 6.429219 | NaN | 3.960664 | 7.213633 | 6.884104 | 7.092926 | 7.671693 |
| 9.181293 | 9.056126 | 8.951185 | 8.581626 | 8.657208 | 9.871696 | 9.706912 | 9.897639 | 9.849988 |
| 8.94865 | 8.830557 | 8.888927 | 7.491899 | 7.411699 | 8.611217 | 8.506658 | 8.418202 | 8.497005 |
| 8.15686 | 8.272243 | 8.309874 | 9.745747 | 9.663636 | 7.121729 | 7.581758 | 7.519153 | 8.28574 |
| NaN | NaN | 4.274023 | NaN | 5.251169 | NaN | NaN | NaN | 3.610911 |
| 10.05652 | 9.829325 | 9.811955 | 8.392111 | 9.598037 | 8.576556 | 9.089676 | 9.075371 | 8.45007 |
| 9.002357 | 9.152208 | 9.053588 | 9.533798 | 9.484344 | 8.883852 | 9.037805 | 9.227204 | 9.437261 |
| 5.704949 | NaN | NaN | 4.547403 | 5.182287 | 7.304492 | 5.512905 | 6.519777 | NaN |
| 8.462217 | 8.405018 | 8.534639 | 7.620624 | 7.604196 | 8.945255 | 8.820075 | 9.067058 | 9.018382 |
| 5.357419 | 5.478174 | 5.162762 | 6.174963 | 6.71386 | 3.054726 | 2.982652 | 3.306965 | 4.371819 |
| 8.492837 | 8.381847 | 8.599106 | 8.687888 | 8.810698 | 8.373666 | 8.379316 | 8.335203 | 8.167405 |
| 5.269073 | 4.854703 | 5.092545 | NaN | 3.939254 | 4.383676 | 6.011977 | 6.754938 | 6.104645 |
| 8.013364 | 7.950257 | 7.920849 | 8.068575 | 8.430454 | 6.246851 | 6.305538 | 6.579344 | 6.41753 |
| 7.384097 | 7.233519 | 7.012064 | 6.228734 | 5.930812 | 6.951494 | 7.303174 | 7.518033 | 6.533115 |
| 10.58016 | 10.5504 | 10.62801 | 9.721641 | 9.756131 | 10.80707 | 10.94013 | 10.74478 | 10.71045 |
| 8.285198 | 8.218879 | 8.27231 | 7.356953 | 7.499236 | 8.268671 | 8.102322 | 8.145497 | 8.73947 |
| 6.086437 | 6.449896 | 6.666407 | NaN | NaN | NaN | NaN | NaN | NaN |
| 10.15176 | 9.852181 | 9.849202 | 11.15416 | 11.27123 | 9.683178 | 9.500932 | 9.48011 | 9.958484 |
| 7.630214 | 7.392402 | 7.237679 | 5.958252 | 7.393578 | 8.963799 | 8.443454 | 8.283627 | NaN |
| 8.131926 | 8.293524 | 8.40345 | 5.236332 | 5.2802 | 9.569868 | 9.282194 | 9.415743 | 7.17313 |
| 7.814403 | 7.299426 | 8.120266 | 6.376911 | 6.485455 | 7.195391 | 7.52009 | 7.875926 | 7.247538 |
| 8.72261 | 8.54235 | 8.646479 | 8.761502 | 8.687158 | 9.021341 | 9.27444 | 9.208303 | 7.930904 |
| 8.500923 | 8.398033 | 8.421112 | 8.951267 | 8.967606 | 8.055431 | 7.995431 | 7.886399 | 7.709422 |
| 10.38207 | 10.55734 | 10.63754 | 10.67147 | 10.72151 | 9.59433 | 9.588369 | 9.664904 | 10.58757 |
| 9.121675 | 9.325004 | 9.40378 | 8.982782 | 8.723764 | 9.033506 | 8.734787 | 9.449331 | 8.951318 |
| 7.495358 | 7.387 | 7.572024 | 6.285291 | 6.510197 | 7.613714 | 7.653603 | 7.704448 | 7.748302 |
| 4.187817 | 4.351745 | 3.210719 | NaN | NaN | NaN | NaN | NaN | 5.081661 |
| 6.788476 | 5.706159 | 6.932544 | 5.202802 | 5.192972 | 6.476751 | 6.10102 | 6.453103 | 5.637546 |
| 8.170407 | 7.34505 | 8.074575 | 5.770176 | 5.721018 | 7.762581 | 8.009763 | 7.813414 | 6.96877 |
| 7.495194 | 7.300776 | 7.526664 | 6.657343 | 6.565469 | 7.052896 | 7.140038 | 7.151131 | 7.353364 |
| 6.973761 | 7.10806 | 7.101751 | 5.700303 | 5.381553 | 6.119645 | 6.390017 | 6.489028 | 5.970152 |
| 6.093285 | 5.092614 | 6.819347 | NaN | 5.499046 | 7.020663 | 7.150681 | 6.987947 | 6.093589 |
| NaN | NaN | 6.353172 | NaN | NaN | 7.118854 | 6.976255 | 6.926944 | 5.896153 |
| 7.424932 | 7.401546 | 7.508202 | 6.491609 | 6.430696 | 7.305315 | 7.367349 | 7.509435 | 7.188244 |
| 10.3667 | 10.33297 | 10.37682 | 9.602558 | 9.29315 | 9.638269 | 9.930135 | 9.856525 | 9.633129 |
| 8.005573 | 5.504555 | 6.301049 | 4.381802 | 5.58733 | NaN | NaN | NaN | 4.505217 |
| 6.469172 | 5.977427 | 6.092731 | 5.99859 | 6.077124 | 6.212384 | 6.146555 | 6.650382 | 6.641788 |
| 9.529353 | 9.314136 | 9.473979 | 8.056755 | 7.990145 | 8.913647 | 8.792653 | 8.81775 | 9.424358 |
| 7.11329 | 7.23738 | 7.1549 | NaN | NaN | 7.651441 | 7.74702 | 7.638774 | 5.269592 |
| 8.87189 | 9.065735 | 9.282269 | 8.083461 | 8.062846 | 10.23787 | 9.562351 | 10.02722 | 9.351404 |
| 6.407349 | 6.815423 | 6.588799 | 5.353702 | 6.971829 | 7.299071 | 7.206668 | 7.098786 | 6.273931 |
| 6.3093 | 5.51218 | 5.255888 | 6.080529 | 5.678556 | 6.008143 | 5.812459 | 5.776971 | 6.241735 |
| 7.062585 | 7.822727 | 6.489958 | 8.629333 | NaN | 7.137834 | 6.465137 | 7.803249 | 6.595559 |
| 5.552605 | 7.610955 | 5.991838 | 6.828373 | 6.352106 | 5.964553 | 5.983809 | 6.442089 | 4.04831 |

| | | | | | | | | |
|---|---|---|---|---|---|---|---|---|
| 9.116163 | 9.101258 | 9.063102 | 7.920968 | 8.12373 | 8.695911 | 8.743453 | 8.771714 | 9.076659 |
| 7.931271 | 7.85365 | 7.930776 | 7.633658 | 7.823769 | 8.548577 | 7.982198 | 8.350388 | 7.963797 |
| 11.2972 | 10.77982 | 11.02053 | 11.0115 | 10.65833 | 10.82965 | 11.48356 | 11.40341 | 10.9769 |
| 10.07118 | 9.855358 | 9.712833 | 9.167199 | 9.381194 | 10.03075 | 9.908292 | 10.10708 | 9.380075 |
| 6.696989 | 5.873841 | 6.113078 | 6.725045 | 6.145318 | 6.7937 | 6.9507 | 6.664261 | NaN |
| 7.012093 | 6.856545 | 7.082712 | 7.046849 | 7.077202 | 7.805569 | 7.749828 | 8.045794 | 7.539856 |
| 5.641283 | 4.983274 | 5.251565 | 4.19297 | 4.558525 | 5.440809 | 5.252364 | 5.379099 | 5.414326 |
| 6.922736 | 7.425538 | 7.663978 | 3.650435 | 5.972022 | 7.200631 | 7.381346 | 7.347312 | 6.753656 |
| 9.530649 | 9.508216 | 9.742884 | 7.512274 | 7.725276 | 9.851727 | 9.854115 | 9.700232 | 7.560921 |
| 6.431677 | 5.766602 | 5.804928 | 7.302036 | 4.27265 | 5.345829 | 5.967268 | 6.103158 | 4.831467 |
| 7.654803 | 7.478398 | 7.691406 | 6.244605 | 6.422712 | 7.815401 | 7.863055 | 7.946181 | 7.07145 |
| 11.05191 | 10.95684 | 11.05733 | 10.0337 | 9.963421 | 10.88463 | 10.89522 | 11.01113 | 10.40617 |
| 7.798373 | 7.37336 | 7.842077 | 6.049617 | 5.791408 | 7.689404 | 7.396612 | 7.607202 | 6.775989 |
| 7.053609 | 7.097764 | 7.114563 | 6.737921 | 6.962766 | 6.796711 | 7.032906 | 7.003544 | 7.409939 |
| 7.397143 | 7.386735 | 7.207907 | 7.254418 | 7.20456 | 7.578684 | 7.700085 | 7.752107 | 7.544623 |
| 12.64403 | 12.65814 | 12.6264 | 12.69044 | 12.50125 | 12.06383 | 12.04375 | 11.98697 | 12.30045 |
| 8.516791 | 8.504797 | 8.517495 | 7.229035 | 7.382013 | 7.796799 | 7.736947 | 7.976602 | 8.233093 |
| 7.700962 | 7.458508 | 7.568813 | 5.847579 | 5.831256 | 7.450458 | 7.451982 | 7.16746 | 7.33073 |
| 8.968412 | 9.156382 | 9.02995 | 9.713658 | 9.320716 | 7.988757 | 8.367911 | 8.318449 | 9.211239 |
| 7.488949 | 7.180488 | 7.632942 | 6.250485 | 6.532353 | 7.468449 | 7.360397 | 7.397968 | 7.073102 |
| 10.48015 | 10.22033 | 10.31127 | 10.75338 | 11.05934 | 9.753007 | 9.646411 | 9.462559 | 9.861161 |
| 8.472076 | 8.506547 | 8.475551 | 7.807578 | 7.900196 | 8.499578 | 8.698932 | 8.625827 | 9.088858 |
| NaN | 6.063807 | 5.564522 | NaN | NaN | NaN | NaN | NaN | NaN |
| 10.65141 | 10.5349 | 10.56204 | 11.27518 | 11.26775 | 10.22099 | 10.16206 | 10.1572 | 10.36858 |
| 7.895841 | 7.664795 | 7.923265 | 6.112483 | 7.159449 | 7.6768 | 7.750496 | 8.145617 | 7.173718 |
| NaN | 8.088899 | 8.27929 | NaN | NaN | NaN | 6.002654 | NaN | NaN |
| 6.510688 | 1.627734 | 5.652321 | 5.346776 | NaN | 5.721649 | 5.912766 | 5.615754 | 6.202106 |
| 12.42896 | 12.48913 | 12.4712 | 11.50292 | 11.52683 | 12.30936 | 12.39228 | 11.87472 | 12.20185 |
| 7.99905 | 8.167593 | 8.020679 | 7.173143 | 6.915124 | 7.883945 | 7.724714 | 7.66178 | 6.771214 |
| 9.237673 | 9.076348 | 9.037488 | 9.282763 | 9.083447 | 8.512449 | 8.536848 | 8.385908 | 8.309382 |
| 10.50327 | 10.51758 | 10.62551 | 10.52561 | 10.34048 | 10.4362 | 10.35662 | 10.45242 | 10.46891 |
| 7.901343 | 7.774856 | 7.934923 | 6.450924 | 6.774213 | 7.306975 | 7.512956 | 7.263115 | 7.266226 |
| 6.260911 | 6.732151 | 5.455286 | NaN | 5.253317 | 6.400208 | 6.360736 | 6.131609 | 6.079537 |
| 8.327908 | 8.19708 | 8.053634 | 7.775831 | 7.263589 | 9.13178 | 8.913558 | 9.009915 | 8.555144 |
| 8.648319 | 8.576549 | 8.522561 | 8.103271 | 8.164558 | 9.272992 | 8.63349 | 8.946454 | 8.562049 |
| 6.988833 | 7.016204 | 6.677765 | 6.325247 | NaN | 7.841705 | 7.2608 | 7.887528 | 6.860524 |
| 8.421622 | 8.509869 | 8.534503 | 7.999468 | 8.243224 | 8.537643 | 8.642847 | 8.569402 | 8.540308 |
| 7.808362 | 7.440207 | 7.650329 | 6.49227 | 6.495375 | 7.001093 | 7.289777 | 7.36726 | 7.711423 |
| 7.87385 | 7.879186 | 7.924076 | 7.286167 | 7.026888 | 6.686509 | 7.555263 | 7.471232 | 7.272801 |
| 6.175157 | 6.427219 | 6.477184 | 5.783959 | 5.571657 | 5.919399 | 5.810318 | 5.932083 | 5.644299 |
| 8.692039 | 8.616144 | 8.487339 | 9.954041 | 9.923161 | 7.982545 | 8.065563 | 7.938931 | 8.656481 |
| 9.186451 | 9.069181 | 9.115021 | 10.11616 | 9.829925 | 8.636955 | 8.687528 | 8.824931 | 8.711616 |
| 5.398848 | 5.398866 | 5.213655 | 5.915812 | 5.496022 | 5.335538 | 5.578166 | 4.80485 | 4.12791 |
| 7.518783 | 7.647719 | 7.685802 | 8.00508 | 7.906127 | 7.47288 | 7.459302 | 7.670865 | 7.587343 |
| 10.88031 | 10.77936 | 10.79076 | 10.36976 | 10.21408 | 10.883 | 10.85042 | 10.82943 | 9.96498 |
| NaN | 4.645598 | 5.630734 | 7.080312 | 6.711894 | 4.149052 | 5.844638 | 5.666704 | 4.22377 |
| 7.12928 | 7.423033 | 6.772201 | 4.192133 | 4.868091 | 6.049769 | 5.808503 | 5.930198 | 5.088066 |

| | | | | | | | | |
|---|---|---|---|---|---|---|---|---|
| 8.433686 | 8.296615 | 8.575809 | 7.590416 | 7.440536 | 9.034964 | 8.918447 | 8.796112 | 7.996991 |
| 9.607057 | 9.58462 | 9.441623 | 9.470167 | 9.434063 | 9.199934 | 9.379947 | 9.260656 | 9.223362 |
| 7.598933 | 7.697397 | 7.691092 | 7.465356 | 7.608517 | 8.702178 | 8.587306 | 8.632973 | 8.131208 |
| 9.643221 | 9.383278 | 9.208787 | 9.189198 | 9.383956 | 10.16761 | 9.533189 | 9.870426 | 8.981652 |
| 9.454015 | 9.511182 | 9.494947 | 10.28544 | 10.18122 | 10.04976 | 9.884965 | 9.920233 | 9.744849 |
| 11.42858 | 11.15374 | 11.2963 | 11.27485 | 11.15863 | 11.77413 | 11.36672 | 11.63528 | 11.17682 |
| 10.08743 | 10.12195 | 10.09344 | 10.64229 | 10.52561 | 10.54823 | 10.4815 | 10.58482 | 10.12434 |
| 12.16119 | 12.14 | 12.18355 | 13.44746 | 13.18929 | 12.52942 | 12.50121 | 12.42361 | 11.8967 |
| 13.22371 | 13.27503 | 13.28925 | 14.43659 | 14.50611 | 13.22893 | 13.16664 | 13.14551 | 13.01875 |
| 6.999624 | 6.710857 | 7.235344 | 8.598365 | 8.791327 | 7.536008 | 6.775249 | 6.963735 | 6.417323 |
| 6.940613 | 5.930357 | 6.645268 | NaN | 6.320723 | 6.389907 | 6.304916 | 6.166119 | 5.759253 |
| 6.693362 | 7.265542 | 6.737014 | 8.242608 | 8.540056 | 8.261873 | 6.808268 | 7.57559 | 6.296927 |
| 6.501269 | 6.809371 | 6.598134 | 6.56963 | 6.422477 | 6.581713 | 6.834995 | 6.684678 | 6.796041 |
| 6.721705 | 6.853178 | 6.568748 | 6.467591 | 6.390058 | 7.400044 | 7.328227 | 7.243587 | 7.653835 |
| 7.991487 | 8.096266 | 7.943069 | 8.338693 | 8.586143 | 8.721818 | 8.301225 | 8.804023 | 8.551252 |
| 9.336126 | 9.236533 | 9.230066 | 8.425225 | 8.373129 | 8.944743 | 9.337789 | 9.267951 | 9.406896 |
| 7.16645 | 7.33358 | 7.27692 | 6.960789 | 7.594489 | 7.075735 | 7.07077 | 7.035034 | 5.874862 |
| 9.681056 | 9.682384 | 9.383262 | 10.87932 | 11.0693 | 10.28871 | 10.29485 | 10.24493 | 9.484443 |
| 9.814005 | 9.803366 | 9.783777 | 9.250587 | 9.142899 | 10.3035 | 10.27819 | 10.29104 | 10.02012 |
| 10.22907 | 10.366 | 10.31268 | 12.16505 | 11.8918 | 10.18841 | 10.19048 | 9.378945 | 10.42628 |
| 6.442406 | 5.953607 | 6.616317 | NaN | NaN | NaN | NaN | 6.364671 | NaN |
| 7.871892 | 7.903657 | 7.948545 | 7.513945 | 7.483214 | 7.684189 | 7.682492 | 7.67757 | 8.176172 |
| 10.57855 | 11.47185 | 10.45571 | 11.49587 | 11.1808 | 10.64052 | 10.70228 | 10.58245 | 9.865399 |
| 9.276857 | 8.136569 | 8.382627 | 6.35977 | 6.631122 | NaN | NaN | NaN | 6.168818 |
| 4.489739 | 5.479803 | NaN | 5.585773 | 5.794214 | 6.102017 | 5.963939 | 6.184505 | 4.432713 |
| 9.811528 | 9.888681 | 10.03124 | 11.22561 | 11.14567 | 10.08197 | 9.901008 | 9.996522 | 9.35489 |
| 11.28963 | 11.49572 | 11.3568 | 11.81385 | 11.55149 | 11.08141 | 11.22509 | 11.34745 | 10.25022 |
| 7.157818 | 6.876086 | 7.140773 | 6.5192 | 6.197282 | 7.697811 | 6.831687 | 7.257285 | 6.112097 |
| 8.835606 | 8.757599 | 8.769119 | 7.405847 | 7.423601 | 8.403991 | 8.442665 | 8.33838 | 8.334607 |
| 7.496184 | 7.376351 | 6.76249 | 8.824717 | 8.066309 | 7.251035 | 7.405602 | 6.707572 | 7.875834 |
| 7.096636 | 7.246289 | 7.010484 | 6.755866 | 6.82098 | 6.802551 | 7.078481 | 6.815751 | 6.933208 |
| 6.119534 | 6.17988 | 6.682438 | 6.099766 | 5.380682 | 5.442258 | NaN | NaN | 5.484635 |
| 6.64224 | 6.466905 | 6.453318 | 5.501091 | 5.986408 | 7.133088 | 7.459679 | 7.565339 | 6.877399 |
| 8.178883 | 8.809121 | 8.219198 | 10.09553 | 10.09176 | 7.038509 | 7.908757 | 7.828914 | 8.699334 |
| 6.862562 | 6.740119 | 6.560382 | 8.318524 | 8.358129 | 7.464611 | 7.351546 | 7.381589 | 6.60872 |
| 9.803765 | 9.842482 | 9.983654 | 9.799253 | 9.619571 | 9.544018 | 9.403431 | 9.468315 | 10.06291 |
| 9.002597 | 8.98456 | 9.000268 | 7.507763 | 7.767256 | 9.055722 | 8.871806 | 8.819257 | 9.10066 |
| 6.391055 | 5.408016 | 6.558353 | 6.722165 | 6.525246 | 6.949777 | 6.918926 | 6.753519 | 6.349104 |
| 6.121664 | 5.991748 | 6.002971 | 4.812212 | 4.76435 | 6.219907 | 6.769119 | 6.794783 | 6.084323 |
| 7.460106 | 6.666622 | 6.836044 | 6.754452 | 6.49159 | 6.487209 | 6.348417 | 6.461483 | 5.385952 |
| 7.303212 | NaN | 7.189797 | NaN | NaN | NaN | NaN | NaN | 5.601502 |
| 6.80784 | 4.117213 | NaN | 7.680591 | 7.547644 | NaN | 3.912585 | NaN | 4.847213 |
| 5.690269 | 6.672607 | 6.140907 | NaN | NaN | 7.525128 | 7.880105 | 7.858618 | 5.214558 |
| 8.993056 | 8.868895 | 8.952638 | 8.348396 | 8.568508 | 8.358936 | 8.506787 | 8.757289 | 8.694534 |
| 4.461493 | 5.138516 | 4.842663 | 2.530391 | NaN | NaN | NaN | NaN | NaN |
| 6.948579 | 5.856459 | 7.180634 | 8.222154 | 7.591339 | 6.482934 | 6.02498 | 6.337282 | 6.617202 |
| 6.89008 | 6.961387 | 7.065407 | 9.1337 | 8.969663 | 7.181773 | 7.175909 | 7.216949 | 6.517216 |

| | | | | | | | | |
|---|---|---|---|---|---|---|---|---|
| 10.65029 | 10.51834 | 10.44565 | 9.988412 | 10.01534 | 10.61887 | 10.64235 | 10.69323 | 10.67218 |
| 6.647073 | 6.481092 | 6.583062 | NaN | NaN | 7.832387 | NaN | NaN | 6.247428 |
| 7.655766 | 7.369879 | 7.59699 | NaN | 6.383041 | 7.741765 | 6.512494 | 7.753201 | 6.838516 |
| 11.65106 | 11.66209 | 11.59584 | 13.72849 | 13.66957 | 10.46989 | 10.34123 | 10.52287 | 11.35865 |
| 9.257849 | 9.20951 | 9.313816 | 10.07187 | 10.1399 | 9.3489 | 9.227375 | 9.451172 | 9.342944 |
| NaN | NaN | 5.437139 | NaN | NaN | 6.425838 | 6.447839 | 6.173064 | 5.81134 |
| 8.44449 | 8.276275 | 8.139512 | 7.411958 | 7.752889 | 8.021652 | 7.958961 | 8.073692 | 8.068707 |
| 7.993709 | 7.091063 | 8.092086 | 7.142407 | 7.11136 | 7.988986 | 7.967799 | 8.062875 | 8.167929 |
| 10.60319 | 10.59123 | 10.62539 | 9.317361 | 9.539843 | 10.54508 | 10.38323 | 10.52958 | 10.25846 |
| 8.081077 | 7.852571 | 8.129498 | 7.561272 | 7.592876 | 7.41123 | 7.777279 | 7.806664 | 8.028174 |
| NaN | NaN | NaN | 8.041199 | 7.041731 | NaN | NaN | NaN | NaN |
| 13.57228 | 13.28174 | 13.42418 | 14.22561 | 14.3332 | 12.70742 | 13.00776 | 12.5732 | 13.09992 |
| NaN | 5.561 | 5.072167 | 6.397313 | 5.871137 | 5.31015 | 5.567845 | 5.236582 | 5.034755 |
| 7.212337 | 7.640603 | 7.507883 | 7.283086 | 6.886672 | 7.846388 | 7.783883 | 7.847806 | 6.171193 |
| 6.224864 | 6.189837 | 6.155188 | 5.651842 | 5.890481 | 6.968597 | 6.492961 | 6.576717 | 6.514456 |
| 9.836561 | 9.773547 | 9.78296 | 8.851171 | 8.618489 | 10.01471 | 10.18299 | 10.2173 | 9.354868 |
| 8.327243 | 8.349236 | 8.325695 | 7.854275 | 7.951848 | 8.114177 | 8.189343 | 8.3384 | 7.916482 |
| 7.956103 | 8.218534 | 8.345294 | 6.76787 | 7.014213 | 7.17347 | 7.277288 | 6.986107 | 6.696146 |
| 8.853525 | 8.706371 | 8.833175 | 7.247667 | 7.361871 | 8.888508 | 8.833197 | 8.944482 | 9.147203 |
| 8.903933 | 8.872994 | 8.878076 | 9.345093 | 9.028526 | 7.802473 | 7.986659 | 8.026445 | 9.308051 |
| 10.42797 | 10.31596 | 10.4547 | 9.265445 | 9.167744 | 10.46145 | 10.45247 | 10.40822 | 10.45325 |
| 5.689607 | 6.467432 | 6.095111 | 6.055789 | 6.154234 | 6.305333 | 6.421339 | 6.187814 | 6.403854 |
| 7.576771 | 6.594734 | 7.656735 | 6.374187 | 6.747571 | 8.343605 | 7.711862 | 8.21934 | 6.506724 |
| 6.846543 | 6.613233 | 6.701552 | 6.393277 | 6.469585 | 7.121618 | 7.636368 | 7.036717 | 6.528534 |
| 10.55176 | 10.4904 | 10.48153 | 9.972331 | 9.719985 | 10.87244 | 11.02685 | 10.83916 | 10.63868 |
| 8.091324 | 7.316557 | 7.445732 | 7.302301 | 7.476757 | 7.44759 | 7.075755 | 7.359004 | 6.545985 |
| 7.971798 | 8.015559 | 8.144635 | 7.599705 | 7.690555 | 8.290731 | 8.387153 | 8.436824 | 7.249214 |
| 10.10449 | 10.15027 | 10.09479 | 9.838311 | 9.756815 | 10.03842 | 9.886921 | 10.11149 | 9.643561 |
| 9.401853 | 9.157689 | 9.314281 | 10.51297 | 10.53238 | 9.4979 | 9.302835 | 9.442385 | 8.053117 |
| 9.266959 | 9.329775 | 9.341278 | 8.450345 | 8.357958 | 9.303607 | 9.486708 | 9.440941 | 9.142244 |
| 9.548785 | 8.311668 | 7.845916 | 8.28796 | 8.040545 | 6.871966 | 7.366871 | 7.141001 | 8.075685 |
| 6.897896 | 6.736991 | 6.828119 | 6.694964 | 6.682131 | 7.205191 | 7.216891 | 7.057251 | 6.236478 |
| 7.243767 | 7.680279 | 6.98152 | 6.213496 | 5.402979 | 6.889519 | 7.481653 | 6.970172 | 6.923846 |
| 6.874035 | 7.490294 | 6.056433 | NaN | NaN | 6.105172 | 5.705111 | 5.894241 | 6.561997 |
| 8.749373 | 8.794034 | 8.575173 | 9.469389 | 9.033861 | 8.390643 | 8.225685 | 8.52517 | 8.71537 |
| 6.803451 | 6.932336 | 7.070291 | 7.479455 | 7.334006 | 7.026227 | 7.564725 | 7.467205 | 7.070378 |
| 6.747055 | 6.560245 | 6.71197 | 6.675687 | 7.018233 | 7.482959 | 7.633568 | 7.703071 | 6.549844 |
| 9.700769 | 9.581429 | 9.637541 | 9.388599 | 9.401685 | 10.15 | 10.01822 | 10.05251 | 9.721734 |
| 9.04973 | 9.073422 | 9.121595 | 8.241199 | 8.382333 | 9.483912 | 9.5145 | 9.583615 | 9.017101 |
| 9.926782 | 10.25069 | 10.10384 | 9.776844 | 10.03629 | 10.64814 | 10.33212 | 10.65707 | 10.30652 |
| 9.143004 | 8.950243 | 9.011978 | 8.54734 | 8.645676 | 9.622313 | 9.684327 | 9.540068 | 9.644074 |
| 7.274669 | 6.988426 | 6.990726 | 7.429919 | 7.258068 | 7.228477 | 7.342272 | 7.412954 | 7.137722 |
| 6.585002 | 6.657531 | 6.42145 | 5.449054 | 5.990586 | 7.030951 | 6.863294 | 6.900015 | 6.501558 |
| 5.134984 | NaN | 5.275329 | NaN | NaN | 5.444706 | 6.130766 | 6.058368 | 5.474461 |
| 8.686347 | 8.97263 | 8.41726 | 7.866916 | 7.913134 | 7.633376 | 7.41627 | 7.40516 | 8.145391 |
| NaN | NaN | NaN | 5.472186 | 6.347015 | NaN | 4.797615 | 4.824123 | NaN |
| NaN | NaN | 6.708011 | 6.442372 | 5.872307 | 8.919904 | 8.882746 | 8.925273 | 7.328804 |

| | | | | | | | | |
|---|---|---|---|---|---|---|---|---|
| 8.821836 | 9.047072 | 9.160442 | 9.457775 | 9.207427 | 8.340459 | 8.321516 | 8.5505 | 7.792437 |
| 10.37097 | 10.26878 | 10.44985 | 10.54597 | 10.54588 | 9.914985 | 9.91031 | 9.492693 | 8.703918 |
| 8.558281 | 8.508589 | 8.446609 | 6.851401 | 6.893007 | 7.545999 | 7.737548 | 7.703405 | 7.915823 |
| 9.086267 | 7.813852 | 7.851936 | NaN | NaN | NaN | NaN | NaN | NaN |
| 7.336363 | 7.224433 | 7.227043 | 7.611119 | 7.800428 | 7.765961 | 7.795016 | 7.832624 | 8.238635 |
| 7.932735 | 7.84339 | 7.72064 | 8.262234 | 8.18384 | 7.563245 | 7.490066 | 7.582599 | 8.288595 |
| 6.513563 | 6.405511 | 4.405413 | NaN | 2.233528 | 7.078666 | 7.07948 | 6.937575 | 5.957633 |
| 10.45203 | 10.38741 | 10.30072 | 9.36566 | 9.494452 | 10.41235 | 10.60939 | 10.40864 | 10.25628 |
| 7.855299 | 7.761106 | 7.940153 | 8.088057 | 8.074739 | 8.083289 | 8.041423 | 8.034074 | 8.477375 |
| 7.084271 | 7.127705 | 7.293736 | 7.110909 | 7.035143 | 7.103744 | 6.785256 | 7.257809 | 5.472881 |
| 7.595705 | 7.667256 | 7.720149 | 7.016578 | 7.160366 | 7.898002 | 8.062585 | 7.889816 | 8.006157 |
| 9.554512 | 9.233601 | 9.361877 | 9.649505 | 9.550508 | 9.652256 | 9.152792 | 9.518805 | 8.512788 |
| 7.652534 | NaN | 7.17948 | 6.692544 | 6.439591 | 6.011007 | 6.684784 | NaN | 5.833182 |
| 4.963615 | 5.736125 | 6.129605 | 5.791432 | 4.690841 | 5.936259 | 6.204098 | 6.163758 | 6.464582 |
| 7.353106 | 7.416332 | 7.364899 | 6.627776 | 6.65557 | 7.230068 | 7.302804 | 7.240866 | 7.546122 |
| 4.449412 | 4.471984 | 4.164963 | 5.650885 | 5.54611 | 4.668401 | 4.229942 | 4.97838 | 2.461915 |
| 7.339195 | 7.083498 | 7.321918 | 7.97882 | 8.263384 | 6.572963 | 7.378901 | 6.893034 | 5.652601 |
| 6.926138 | 7.072503 | 6.932168 | 5.869797 | 5.717825 | 6.929643 | 6.46869 | 7.061059 | 6.128481 |
| 7.439067 | 7.358647 | 7.49157 | 6.918136 | 6.794931 | 8.240582 | 8.417714 | 8.101725 | 8.650911 |
| 8.434367 | 8.510772 | 8.565324 | 7.507518 | 7.142568 | 8.03659 | 8.135325 | 8.030137 | 7.471977 |
| 8.759007 | 8.70807 | 8.825421 | 7.443663 | 7.138236 | 8.066451 | 8.396367 | 8.017663 | 8.127186 |
| 6.822271 | 6.564001 | 6.933909 | 6.989927 | 7.389613 | 6.851257 | 6.862563 | 7.218102 | 6.938778 |
| 5.65877 | 5.89913 | 5.958512 | 4.385483 | 3.488467 | 6.276268 | 6.926724 | 6.241994 | 5.191512 |
| 7.281414 | 7.038282 | 7.587305 | NaN | NaN | NaN | NaN | NaN | NaN |
| 8.173629 | 8.21806 | 8.07158 | 6.365128 | 6.166464 | 8.290676 | 8.171357 | 8.362124 | 8.152057 |
| 7.115057 | 6.984596 | 7.073412 | 6.744021 | 6.601255 | 7.743034 | 7.446245 | 7.466066 | 7.498981 |
| NaN | NaN | NaN | NaN | NaN | NaN | NaN | NaN | 7.995391 |
| 8.993499 | 8.7449 | 8.830642 | 7.424766 | 7.285443 | 8.929296 | 8.889791 | 8.674834 | 8.87498 |
| 7.618968 | 7.597581 | 7.631405 | 6.942102 | 7.104881 | 7.468072 | 7.171758 | 7.364537 | 7.502114 |
| 8.869395 | 8.912134 | 8.669363 | 8.240036 | 8.722488 | 9.003017 | 8.853406 | 8.949903 | 8.906379 |
| 9.899078 | 9.915536 | 10.0133 | 9.355866 | 9.08881 | 9.133196 | 9.836708 | 9.937985 | 9.383122 |
| 10.26238 | 10.03997 | 10.34951 | 10.43325 | 10.63331 | 9.474419 | 9.233891 | 9.266654 | 8.93414 |
| 6.191508 | 6.025417 | 6.410323 | 8.964034 | 9.15021 | NaN | 6.852479 | NaN | NaN |
| 9.41261 | 9.598434 | 9.262975 | 9.177423 | 9.164618 | 9.691515 | 9.655022 | 9.599982 | 8.544902 |
| 3.506271 | 4.43996 | 3.574134 | 3.925244 | 3.806602 | 7.164377 | 7.05509 | 7.267914 | 6.202028 |
| 8.428126 | 8.157404 | 8.265466 | 6.716607 | NaN | 6.740919 | 7.39532 | NaN | 7.365723 |
| 7.479838 | 7.103159 | 7.379867 | 6.665638 | 5.985934 | 7.497822 | 7.302142 | 7.425736 | 7.561663 |
| 10.56649 | 10.53749 | 10.60132 | 10.15744 | 10.01096 | 11.0567 | 10.88553 | 11.05221 | 10.67981 |
| 9.926373 | 10.08196 | 9.994648 | 10.32249 | 10.26527 | 9.852815 | 9.714314 | 9.949409 | 10.24961 |
| 7.593826 | 7.688499 | 7.657207 | 6.169754 | 6.105723 | 7.978893 | 8.053464 | 8.16795 | 7.775224 |
| 7.060525 | 7.311419 | 7.2684 | 6.973209 | 7.110899 | 7.265269 | 7.477119 | 7.303744 | 6.800429 |
| 7.895821 | 6.988649 | 7.718323 | 7.46153 | 8.247105 | 6.871557 | 7.166366 | 6.96804 | NaN |
| 7.956481 | 8.576187 | 8.664078 | 8.728378 | 8.370965 | 8.370053 | 8.291327 | 8.243337 | 7.960393 |
| 13.02245 | 12.92796 | 12.78317 | 12.65385 | 12.70246 | 12.15736 | 12.02135 | 12.12823 | 12.68312 |
| 11.66855 | 11.81608 | 11.8821 | 12.77014 | 12.81515 | 11.9508 | 11.88404 | 11.97563 | 11.96444 |
| 7.315535 | 7.192223 | 7.216063 | 6.527101 | 6.533098 | 7.216465 | 7.332316 | 7.119067 | 7.260978 |
| 6.58426 | 6.883241 | 6.773996 | 6.50154 | 6.48997 | 6.14987 | 6.178601 | 6.171213 | 6.534538 |

| | | | | | | | | |
|---|---|---|---|---|---|---|---|---|
| 8.814541 | 8.722795 | 8.58042 | 8.647694 | 8.661188 | 8.807523 | 8.848828 | 8.845222 | 8.289386 |
| 9.293285 | 9.237422 | 9.302616 | 8.705414 | 8.853503 | 9.320874 | 9.58871 | 9.505319 | 8.885219 |
| 6.39103 | 6.051375 | 6.26721 | 6.301252 | 6.595757 | 7.298132 | 7.343526 | 6.968284 | 6.165143 |
| 6.116942 | 5.63749 | 5.84886 | 6.663265 | 7.668499 | 7.747427 | 7.057452 | 6.9249 | 5.392116 |
| 5.999835 | 5.090071 | 5.361397 | 5.034726 | 4.17297 | 6.893311 | 6.624207 | NaN | 5.753753 |
| 7.015635 | 6.019628 | 6.36068 | 6.748833 | 6.534556 | 6.474889 | 6.366279 | 6.469911 | 7.173118 |
| 6.701329 | 6.647354 | 6.618883 | 6.648318 | 6.297782 | 7.088122 | 6.936104 | 7.028675 | 6.900177 |
| 7.167526 | 7.293121 | 7.294395 | 7.465878 | 6.695796 | 7.334983 | 7.579938 | 7.843088 | 7.726374 |
| 6.979712 | 6.758341 | 7.204803 | 6.075075 | 6.678956 | 6.646117 | 6.899338 | 7.117369 | 6.479844 |
| 8.987155 | 8.916493 | 8.967495 | 8.803689 | 9.037086 | 8.067389 | 8.265294 | 8.09309 | 8.628242 |
| 9.419638 | 9.501844 | 9.579906 | 11.58786 | 10.82267 | 8.727619 | 8.983612 | 9.00895 | 9.265819 |
| 11.50597 | 11.46966 | 11.40929 | 10.64501 | 10.38822 | 11.54633 | 11.41593 | 11.43091 | 11.47586 |
| 11.26417 | 11.29033 | 11.30707 | 10.60781 | 10.58361 | 10.93671 | 10.91354 | 11.15391 | 10.85978 |
| 8.748079 | 8.589937 | 8.624549 | NaN | NaN | 9.293125 | 9.024321 | 9.04265 | 7.916309 |
| 8.136598 | 7.810096 | 8.112042 | 8.867467 | 8.622773 | 7.226643 | 7.27744 | 7.152338 | 7.027493 |
| 7.018379 | 7.117575 | 7.04789 | 6.191241 | 5.958759 | 6.776959 | 6.848465 | 7.303422 | 7.06226 |
| 7.636157 | 7.643079 | 7.587703 | 7.142514 | 7.365216 | 7.6203 | 7.438919 | 7.406033 | 7.234094 |
| 9.247557 | 9.346951 | 9.356117 | 9.030178 | 9.186642 | 9.433995 | 9.351461 | 9.397816 | 9.340386 |
| 10.23132 | 10.23482 | 10.15761 | 10.59591 | 10.58654 | 10.43452 | 10.40079 | 10.39132 | 9.553868 |
| 11.16966 | 11.15546 | 11.27548 | 11.69607 | 11.45477 | 10.15401 | 10.42953 | 10.30624 | 10.2657 |
| 7.721296 | 7.922819 | 7.410151 | 8.449186 | 8.317583 | 7.455391 | 7.656305 | 7.811916 | 7.716471 |
| 7.19477 | 7.159151 | 7.18649 | 5.381518 | 6.230988 | 6.866531 | 6.508897 | 6.885886 | 7.248657 |
| 5.598782 | 5.986238 | 5.541862 | 5.236664 | 5.35055 | 5.618706 | 6.41267 | 5.816652 | 5.640732 |
| 5.774205 | 5.489047 | NaN | 8.026375 | 8.065719 | NaN | 6.08819 | 6.035244 | 5.963898 |
| 8.758045 | 8.607123 | 8.68231 | 8.411014 | 8.637429 | 9.073681 | 9.056828 | 8.976953 | 9.628182 |
| 7.458718 | 7.633005 | 7.820701 | 6.961671 | 7.380769 | 8.391344 | 8.381617 | 8.257782 | 7.82306 |
| 8.045201 | 7.885948 | 7.648808 | 8.043077 | 8.107409 | 7.748776 | 7.723393 | 7.594201 | 7.430861 |
| 7.781626 | 6.985375 | 7.702363 | 7.390954 | 6.872386 | 7.102695 | 6.976038 | 7.009047 | 6.033577 |
| 10.21106 | 10.06531 | 10.22874 | 11.83805 | 11.67456 | 10.32468 | 10.34627 | 10.38825 | 9.885957 |
| 7.57661 | 6.737145 | 7.485465 | 7.003137 | 7.495492 | 8.054185 | 8.203156 | 8.211729 | 7.208245 |
| 9.120591 | 9.202214 | 9.216174 | 8.567263 | 8.477019 | 8.980394 | 9.048943 | 8.963511 | 9.444611 |
| 7.971532 | 7.873039 | 8.148864 | 7.213479 | 7.580097 | 8.090028 | 8.203115 | 8.143751 | 7.426675 |
| 7.783907 | 6.975399 | 7.126226 | 7.536072 | 7.564007 | 7.769386 | 7.753063 | 7.799155 | 7.789157 |
| 10.25146 | 10.12273 | 10.27031 | 8.683962 | 8.493343 | 9.342068 | 9.165666 | 9.43453 | 9.3782 |
| 9.00138 | 8.800968 | 8.947998 | 8.614633 | 8.30962 | 9.311267 | 9.434266 | 9.33969 | 9.25576 |
| 8.008739 | 7.812607 | 7.795044 | 7.092113 | 7.37163 | 8.341144 | 8.143862 | 8.106252 | 8.266487 |
| 7.630394 | 7.37762 | 7.722868 | 10.82336 | 10.41114 | 7.23524 | 7.033078 | 7.27078 | 8.471133 |
| 8.122148 | 8.249539 | 8.421845 | 8.068033 | 7.983066 | 8.280716 | 8.033111 | 8.171536 | 8.140617 |
| 8.888471 | 8.836501 | 8.738441 | 6.978508 | 7.183448 | 9.245514 | 9.165467 | 9.068451 | 8.737089 |
| 9.005477 | 8.898696 | 9.043083 | 8.515012 | 8.778212 | 9.224835 | 9.167015 | 9.266246 | 7.759389 |
| 10.83335 | 10.80024 | 10.74174 | 10.5748 | 10.50025 | 11.33995 | 11.15157 | 11.02288 | 10.84067 |
| 8.915349 | 8.768681 | 8.737588 | 9.813982 | 9.581675 | 8.891641 | 9.172317 | 8.9737 | 8.453147 |
| 7.514668 | 6.660705 | 6.65468 | 7.932734 | 8.074863 | 6.680908 | 7.16577 | 6.79994 | 8.121843 |
| 9.287418 | 9.21612 | 9.461046 | 7.425446 | 7.370772 | 9.545076 | 9.667914 | 9.38522 | 8.23159 |
| 7.555473 | 7.818694 | 6.524151 | 6.520007 | 6.798969 | 7.010108 | 6.830711 | 7.193895 | 6.635745 |
| 8.272943 | 8.412571 | 8.267137 | 10.93888 | 11.14569 | 8.353084 | 8.603761 | 8.426773 | 7.432568 |
| 7.107422 | 6.892873 | 7.177022 | 6.522653 | 6.26193 | 6.829595 | 6.924705 | 6.70207 | 5.965355 |

-Bruker Confidential-

| | | | | | | | | |
|---|---|---|---|---|---|---|---|---|
| 7.374281 | 7.482096 | 7.576484 | 7.53657 | 6.823567 | 6.923803 | 7.13365 | 7.176901 | 6.966166 |
| 9.32019 | 9.176501 | 9.114865 | 8.713095 | 8.670921 | 8.941895 | 8.87536 | 8.946266 | 9.852232 |
| NaN | NaN | NaN | NaN | NaN | 5.850528 | 6.037766 | 6.605579 | NaN |
| NaN | 5.659517 | NaN | NaN | NaN | NaN | NaN | NaN | 6.395492 |
| 8.19382 | 8.158151 | 8.059333 | 7.315585 | 8.149569 | 8.357394 | 8.056946 | 8.161474 | 7.273279 |
| 6.690179 | 6.069538 | 7.208645 | 5.571251 | 7.245571 | 7.761975 | 7.192748 | 7.373194 | 7.301005 |
| 6.316386 | 6.095072 | 6.119113 | 5.086131 | 5.402854 | 6.319529 | 6.374112 | 6.522504 | 5.69537 |
| 7.434031 | 7.433945 | 7.590581 | 6.754439 | 6.697053 | 7.706926 | 7.524906 | 7.095334 | 7.40802 |
| 7.287862 | 6.963519 | 7.029535 | 6.362118 | 6.207611 | 7.224843 | 7.036456 | 6.986645 | 6.776986 |
| 4.668637 | NaN | 4.462719 | NaN | NaN | 6.164999 | 6.322191 | 5.692572 | 5.812727 |
| NaN | 5.167383 | NaN | 4.642537 | 5.067421 | NaN | NaN | NaN | 5.453586 |
| 8.692436 | 8.226743 | 8.349181 | 9.749914 | 9.363755 | 7.602713 | 7.509576 | 7.965006 | 7.593105 |
| 6.073368 | NaN | 6.50121 | NaN | NaN | 7.349982 | 6.72719 | 6.704741 | 6.96511 |
| 6.185341 | 4.864477 | 6.407405 | 5.569226 | 7.534334 | NaN | NaN | 6.255949 | 6.014844 |
| 6.95812 | 6.784254 | 7.017022 | 7.364015 | 7.422021 | NaN | 5.499984 | NaN | 6.696939 |
| 7.878667 | 7.778395 | 7.623219 | 7.84612 | 7.810964 | 7.15553 | 6.985662 | 7.09257 | 7.311746 |
| 11.94365 | 12.04424 | 12.01259 | 11.7192 | 11.71162 | 12.12849 | 11.94077 | 11.9109 | 11.29766 |
| 10.56564 | 10.57744 | 10.56893 | 9.498631 | 9.261826 | 10.80869 | 10.92405 | 10.81383 | 10.63916 |
| 12.22532 | 12.24828 | 12.28453 | 12.09328 | 11.81583 | 11.66118 | 11.73093 | 11.73681 | 11.83871 |
| 7.05563 | 7.130863 | 7.077911 | 5.800755 | 5.57032 | 6.624334 | 6.569084 | 6.651957 | 7.673219 |
| 8.089934 | 8.159814 | 7.989711 | 7.323775 | 6.585764 | 8.146857 | 7.903948 | 8.152081 | 7.346547 |
| 8.390423 | NaN | NaN | 9.451839 | 8.947431 | NaN | 8.346428 | 8.442735 | 7.576104 |
| 8.751158 | 8.705307 | 8.703167 | 7.547514 | 7.437498 | 8.983144 | 8.88991 | 8.921981 | 8.447411 |
| 7.640661 | 7.688622 | 7.620611 | 5.869443 | 6.421916 | 6.232345 | 6.25188 | 7.606039 | 6.491178 |
| 8.56424 | 8.235356 | 8.640505 | 8.554346 | 8.252658 | 8.048539 | 8.366632 | 8.110687 | 8.553985 |
| NaN | NaN | NaN | 8.857778 | 8.039721 | NaN | 7.424887 | 7.566359 | 7.808374 |
| 8.748797 | 9.038709 | 8.992615 | 7.421118 | 7.257291 | 8.933556 | 9.352472 | 9.381146 | 8.961524 |
| 11.46546 | 11.42082 | 11.50648 | 11.64891 | 11.83829 | 11.49457 | 11.57351 | 11.53632 | 11.26436 |
| NaN | NaN | NaN | NaN | NaN | NaN | NaN | NaN | 3.849805 |
| 8.102461 | 8.409298 | 8.059315 | 9.52038 | 9.587358 | 8.141179 | 8.07413 | 8.177123 | 8.930022 |
| 10.08424 | 10.24722 | 10.05007 | 9.328085 | 9.168303 | 9.344781 | 9.285487 | 9.455632 | 9.46468 |
| 6.165764 | 5.77225 | 5.789457 | 5.137399 | 6.89116 | 6.410813 | 6.133287 | 6.058212 | 6.035325 |
| 5.651511 | 6.272767 | 5.421602 | 8.909967 | 8.934162 | NaN | NaN | NaN | 5.26255 |
| 8.179097 | 7.937539 | 8.072078 | 6.578384 | 6.684573 | 7.712701 | 7.803621 | 7.761687 | 8.220558 |
| 7.748247 | 7.633053 | 7.685867 | 6.200765 | 7.027998 | 7.009272 | 6.820932 | 6.994996 | 7.435141 |
| 7.246117 | 7.474412 | 7.01225 | 5.527919 | 5.491802 | 7.083593 | 7.086424 | 7.030003 | 7.338616 |
| 4.52225 | 5.775383 | 4.780876 | 6.125899 | 4.279774 | 5.544034 | NaN | NaN | 4.019095 |
| NaN | NaN | NaN | 5.52848 | 6.639423 | NaN | NaN | NaN | NaN |
| 7.124008 | 6.910166 | 7.322599 | 7.376774 | 7.15925 | 7.238438 | 7.037262 | 6.960639 | 7.079419 |
| 7.011215 | 7.423765 | 7.316858 | 9.425636 | 9.736297 | 6.874467 | 6.545719 | 7.21039 | 5.959622 |
| 8.33087 | 8.124401 | 7.975962 | 6.937115 | 7.141582 | 7.925035 | 7.723963 | 7.745274 | 7.784715 |
| 5.781527 | 5.192324 | 5.832409 | NaN | NaN | NaN | NaN | 5.576461 | 5.730382 |
| 5.934984 | 5.77204 | NaN | 5.74832 | 4.436961 | NaN | 5.981047 | 5.944672 | 4.931833 |
| 7.650342 | 7.52872 | 7.596634 | 6.631776 | 6.889601 | 7.089625 | 6.880402 | 7.306977 | 7.368802 |
| 9.481706 | 9.296742 | 9.335619 | 11.32438 | 11.25199 | 7.57861 | 7.623416 | 7.499146 | 8.126092 |
| 5.374881 | 4.588112 | 4.936718 | NaN | NaN | 5.701349 | 5.776876 | 5.931583 | 4.629835 |
| 4.944339 | 4.93854 | 3.752262 | 5.98715 | 4.734089 | 5.51741 | 6.346158 | 6.340775 | 5.090571 |

| | | | | | | | | |
|---|---|---|---|---|---|---|---|---|
| 8.216475 | 8.335773 | 8.081209 | 7.056399 | 6.935049 | 8.118734 | 8.192192 | 8.149362 | 8.015648 |
| 8.202619 | 8.711757 | 8.416119 | 6.576927 | 6.903363 | 7.915218 | 7.726724 | 7.810217 | 7.089946 |
| 10.79837 | 10.83176 | 10.78085 | 11.93633 | 11.83136 | 10.89139 | 10.81386 | 10.8402 | 10.68242 |
| 9.156932 | 9.060096 | 9.105645 | 10.7438 | 10.99196 | 8.926965 | 9.143515 | 8.925376 | 10.87128 |
| 7.980309 | 7.949699 | 7.995634 | 8.446467 | 9.135935 | 8.932895 | 8.340018 | 8.740454 | 8.020992 |
| 8.539111 | 8.231107 | 8.267403 | 8.180175 | 8.138814 | 8.008537 | 7.929355 | 8.314906 | 8.012954 |
| 10.43605 | 10.27169 | 10.26655 | 8.942066 | 9.165115 | 10.75842 | 10.97919 | 10.87704 | 9.783358 |
| 9.403594 | 9.276345 | 9.210071 | 8.256425 | 7.726248 | 9.074826 | 9.396654 | 9.290196 | 8.920135 |
| 8.734894 | 8.690863 | 8.642944 | 7.28076 | 7.503176 | 8.363361 | 8.544402 | 8.766739 | 8.14417 |
| 7.320829 | 7.591161 | 6.745415 | 6.381211 | 4.948552 | 7.241409 | 6.914317 | 6.395576 | 6.1298 |
| 8.402936 | 7.958344 | 8.20974 | 8.474153 | 8.497952 | 8.37673 | 8.044795 | 8.188654 | 7.694894 |
| 7.985302 | 7.898076 | 8.087339 | 9.543777 | 9.618347 | 8.247927 | 8.499386 | 8.62593 | 7.779341 |
| 6.027264 | 5.556701 | 6.005141 | 5.929453 | 6.300938 | 6.651953 | 6.847742 | 6.535571 | 7.294823 |
| 9.991638 | 9.904032 | 9.897738 | 8.471263 | 8.424479 | 10.1938 | 10.35288 | 10.33189 | 9.786318 |
| 7.917526 | 8.37582 | 8.125804 | 7.625348 | 7.139025 | 7.913063 | 6.883101 | 7.521336 | 7.566943 |
| 8.763947 | 8.570211 | 8.397079 | 7.69312 | 8.07749 | 8.34997 | 8.259054 | 8.099253 | 7.545501 |
| 7.980709 | 7.634811 | 7.525593 | 7.124627 | 6.599382 | 7.696889 | 7.351292 | 7.669417 | 7.00087 |
| 7.674414 | 7.434977 | 7.785549 | 5.386113 | 5.795925 | 8.607329 | 8.139081 | 8.633792 | 7.493447 |
| 8.249084 | 8.260273 | 8.305659 | 7.203887 | 7.236587 | 8.070748 | 8.11987 | 8.179542 | 8.493914 |
| 9.466482 | 9.584374 | 9.661164 | 7.989858 | 7.916676 | 9.269764 | 9.450786 | 9.515733 | 9.781466 |
| 8.846948 | 8.934097 | 9.025305 | 8.176658 | 8.159346 | 8.143429 | 7.945873 | 8.150537 | 8.328522 |
| NaN | 6.785283 | 7.278738 | NaN | 6.058343 | 7.244403 | NaN | 7.33538 | 6.693145 |
| 6.324503 | 6.442491 | 6.693355 | 6.398486 | 6.595358 | 7.099607 | 7.070801 | 7.233343 | 6.317382 |
| 9.278489 | 9.203399 | 9.222997 | 7.769523 | 7.641675 | 9.438348 | 9.143149 | 8.944439 | 9.297563 |
| 7.343184 | 7.201213 | 7.369826 | 6.34978 | 6.2368 | 6.488692 | 6.51551 | 7.228827 | 7.992515 |
| 8.426636 | 8.139825 | 8.12344 | 8.128368 | 8.601053 | 7.591771 | 7.738117 | 8.006109 | 7.88788 |
| 7.202229 | 7.25878 | 7.164893 | 6.363746 | 6.629984 | 7.490539 | 7.07899 | 7.380245 | 5.691521 |
| 7.659083 | 7.708658 | 7.24573 | 6.463195 | 6.111956 | 8.084575 | 8.346815 | 8.132798 | 7.681515 |
| 7.555376 | 7.663244 | 7.786904 | 8.034797 | 8.10418 | 8.171497 | 7.666838 | 7.876846 | 7.569139 |
| 8.890541 | 8.869312 | 8.686094 | 7.974885 | 8.04044 | 9.458906 | 9.535872 | 9.490565 | 9.976866 |
| 10.37626 | 10.43697 | 10.42837 | 10.92743 | 11.04851 | 9.85775 | 10.03244 | 10.02118 | 10.14626 |
| 7.83804 | 7.531176 | 7.046072 | 7.794071 | 8.123073 | 6.085356 | 6.876965 | 5.746611 | 6.572537 |
| 8.046415 | 8.081684 | 8.089129 | 7.011257 | 6.964699 | 7.911392 | 8.040366 | 8.083162 | 8.23708 |
| 6.464565 | 6.589039 | 6.18739 | NaN | NaN | 5.799292 | 5.623623 | 6.15523 | 6.085756 |
| 7.473015 | 7.662601 | 7.750821 | 6.538351 | 6.499857 | 7.454494 | 7.506067 | 7.424742 | 7.161207 |
| 5.329223 | 3.605392 | 4.510123 | 5.107709 | 4.435755 | 3.716457 | 5.002886 | 3.231672 | 5.418868 |
| 6.451105 | 6.27037 | 6.908349 | 6.689706 | NaN | NaN | 5.448086 | 5.899081 | 6.565066 |
| 6.358335 | 6.148223 | 6.411307 | 7.229818 | 7.360046 | 6.466149 | 6.351365 | 6.084096 | 5.328629 |
| 8.550614 | 8.720422 | 8.759139 | 7.419141 | 7.386545 | 8.2123 | 7.973505 | 8.132254 | 8.793837 |
| 10.55562 | 10.51368 | 10.54746 | 11.07497 | 10.79181 | 10.74434 | 10.76312 | 10.83018 | 10.76916 |
| 9.220257 | 9.13451 | 8.998592 | 8.057177 | 8.423255 | 9.149008 | 9.214563 | 9.113609 | 9.103036 |
| 7.603411 | 7.569155 | 7.462028 | 6.496852 | 6.461591 | 7.206643 | 7.10717 | 7.558955 | 7.610135 |
| 7.063306 | 6.656254 | 8.382671 | 5.866763 | 5.933275 | 7.285637 | 6.665583 | 6.725792 | 6.132122 |
| 7.330412 | 7.110928 | 7.166867 | 7.460234 | 7.437241 | 7.369942 | 7.340568 | 7.084733 | 8.240569 |
| 8.604718 | 8.547499 | 8.610547 | 6.764059 | 6.870291 | 9.072856 | 9.21336 | 9.070383 | 8.473175 |
| 7.206663 | 7.371049 | 7.707146 | 8.996984 | 8.600714 | 6.112983 | 5.93743 | 5.434735 | 5.947072 |
| 8.550039 | 8.324128 | 8.39778 | 10.54584 | 10.53099 | 7.732491 | 7.574529 | 7.7518 | 8.099104 |

| | | | | | | | | |
|---|---|---|---|---|---|---|---|---|
| 6.655612 | 5.930187 | 6.322506 | 6.118721 | 7.321908 | 5.889736 | 5.619822 | 5.706964 | 7.270199 |
| 5.911108 | 6.098909 | 5.367799 | NaN | 5.344909 | 6.13489 | 5.94668 | 6.164267 | 5.066199 |
| 6.076139 | 4.841099 | 5.323499 | 5.27628 | 5.187315 | 6.312492 | 5.764681 | 6.287623 | 4.875324 |
| 7.096172 | 5.541252 | 4.787946 | 5.383608 | NaN | 7.207485 | 6.911599 | 7.197883 | 5.730595 |
| 8.245462 | 8.645123 | 8.382347 | 6.98642 | 6.962069 | 9.053968 | 9.014469 | 9.080615 | 6.783654 |
| 7.485055 | 7.448423 | 7.607353 | 7.40026 | 7.144526 | 8.484301 | 8.259565 | 8.3621 | 7.08636 |
| 9.566866 | 9.464008 | 9.70667 | 8.937377 | 9.151169 | 9.762853 | 9.454175 | 9.830483 | 8.858729 |
| 8.198249 | 8.260136 | 8.282706 | 9.067219 | 8.880383 | 8.083139 | 8.224942 | 8.16707 | 7.291117 |
| 6.329789 | 6.78381 | 6.845731 | NaN | 5.270143 | 6.347812 | 6.391699 | 6.407187 | 5.154539 |
| NaN | NaN | NaN | NaN | NaN | NaN | NaN | NaN | 7.157057 |
| 9.465321 | 9.276544 | 9.36939 | 7.773033 | 7.580996 | 8.653185 | 8.875526 | 8.844148 | 8.938495 |
| 9.70525 | 9.637403 | 9.609382 | 9.113027 | 9.303388 | 9.439087 | 9.295968 | 9.524868 | 9.410204 |
| 9.102939 | 8.357605 | 8.657633 | 8.321306 | 6.735356 | NaN | 6.840858 | NaN | 7.321113 |
| 9.724721 | 9.671046 | 9.79514 | 10.00979 | 9.813494 | 9.924337 | 9.786757 | 9.981164 | 10.5067 |
| 6.951032 | 7.260318 | 7.20505 | 7.921847 | 7.688233 | 7.485256 | 7.425877 | 7.322155 | 7.160052 |
| 10.21585 | 10.29175 | 10.38885 | 10.68763 | 10.81025 | 9.314532 | 9.308498 | 9.291652 | 9.377674 |
| 7.301628 | 7.106393 | 7.259449 | 5.638537 | 6.294813 | 6.776952 | 7.61734 | 7.049995 | 6.906704 |
| 8.916329 | 8.929798 | 8.972331 | 8.039896 | 8.37773 | 9.038095 | 9.09533 | 9.032138 | 8.361934 |
| 9.494058 | 9.30782 | 9.289901 | 8.128611 | 8.165852 | 9.782484 | 9.677627 | 9.620456 | 9.693638 |
| 7.308549 | 7.925939 | 7.887684 | 5.965774 | 6.67644 | 6.846076 | 6.668615 | 6.941232 | 6.539732 |
| 11.31661 | 11.27908 | 11.30618 | 10.35996 | 10.42731 | 11.81969 | 11.81093 | 11.84629 | 11.24002 |
| 7.184951 | 7.217214 | 7.170752 | 8.316751 | 8.331115 | 6.495459 | 6.545162 | 6.462333 | 7.300612 |
| 10.43097 | 10.33838 | 10.32271 | 11.92223 | 12.08954 | 10.91202 | 11.04396 | 11.1692 | 10.64972 |
| NaN | NaN | NaN | NaN | 8.177569 | 6.848843 | 7.582852 | 7.638582 | NaN |
| NaN | 6.380178 | NaN | NaN | NaN | NaN | NaN | NaN | 8.176548 |
| 8.854542 | 8.813272 | 8.754531 | 10.02582 | 10.22402 | 8.211548 | 8.040689 | 8.136394 | 9.428533 |
| 7.069437 | 7.128979 | 7.116853 | 5.546844 | 6.117847 | 6.259754 | 6.319376 | 6.382751 | 5.999957 |
| 8.365979 | 8.444331 | 8.476995 | 9.114167 | 8.918012 | 8.316399 | 8.263663 | 8.267728 | 8.382329 |
| 5.234346 | 4.389311 | 4.406571 | NaN | NaN | 5.859557 | 4.76956 | 5.61874 | 4.986723 |
| 10.3213 | 10.15975 | 10.22272 | 11.47287 | 11.42244 | 10.7663 | 10.9839 | 10.86952 | 11.44281 |
| 5.943357 | 5.732662 | 6.087608 | 6.26706 | 6.302087 | 5.970848 | 6.235599 | 6.495651 | 7.6905 |
| 6.002056 | 6.230307 | 6.045336 | 4.866335 | 4.935285 | 6.159385 | 5.87565 | 5.533345 | 6.566515 |
| 7.653369 | 7.688913 | 7.647941 | 7.356327 | 7.353212 | 7.925623 | 7.953311 | 7.890434 | 7.156748 |
| 9.195063 | 9.192919 | 9.158419 | 7.888817 | 8.132294 | 9.485523 | 9.466677 | 9.537552 | 8.986206 |
| 7.014561 | 6.921938 | 7.024107 | 5.818211 | 5.511919 | 7.120099 | 6.902283 | 6.967278 | 7.166019 |
| 6.979312 | 7.079172 | 6.763807 | 7.778484 | 8.589959 | 6.277946 | 6.466503 | 6.968002 | 6.6601 |
| 8.620429 | 8.755709 | 8.662617 | 8.420098 | 8.195485 | 7.534819 | 8.259346 | 8.151555 | 7.653187 |
| NaN | NaN | NaN | 7.222365 | 6.203668 | NaN | NaN | NaN | NaN |
| 6.403074 | 5.588619 | 6.269251 | 6.141722 | 6.48175 | 7.06626 | 6.581977 | 6.567651 | 4.534227 |
| 8.506771 | 8.492367 | 8.541417 | 9.718593 | 9.821674 | 9.617373 | 9.925606 | 9.865747 | 6.18862 |
| 8.226788 | 6.862122 | 6.792823 | 7.650911 | 7.948462 | 8.507547 | 7.998334 | 8.218554 | 4.122064 |
| 8.876019 | 8.955951 | 8.796997 | 10.57992 | 9.957256 | 7.921225 | 8.123425 | 8.042336 | 8.548778 |
| 6.932596 | 7.417594 | 6.780734 | 5.8344 | 5.871875 | 7.152439 | 7.281516 | 6.970116 | 6.74664 |
| 7.202182 | NaN | NaN | 4.48576 | 6.097539 | NaN | 6.659097 | 6.410334 | 6.835414 |
| 8.267422 | 8.138428 | 8.359715 | 7.826079 | 7.519097 | 7.584435 | 7.635551 | 7.695213 | 8.175663 |
| 9.362539 | 9.686852 | 9.341191 | 8.828542 | 8.949276 | 9.28155 | 9.678648 | 9.716195 | 9.508392 |
| 10.65492 | 10.68342 | 10.64143 | 9.74038 | 9.698114 | 10.78285 | 10.84067 | 10.80748 | 10.12327 |

-Bruker Confidential-

| | | | | | | | | |
|---|---|---|---|---|---|---|---|---|
| 3.800292 | 2.758322 | 1.272862 | NaN | 2.574629 | 4.082312 | 3.769384 | 3.825938 | 2.359097 |
| 8.918477 | 8.542711 | 8.553269 | 8.631076 | 8.830634 | 7.987393 | 7.764078 | 8.042939 | 8.422185 |
| 9.643496 | 9.535185 | 9.625618 | 9.308611 | 9.268009 | 9.786202 | 9.906049 | 9.761078 | 8.568798 |
| 8.451877 | 8.428117 | 8.358526 | 7.7265 | 7.862561 | 8.386167 | 8.410679 | 8.277589 | 8.650723 |
| 6.673341 | 6.347342 | 6.525735 | 6.331038 | 6.818059 | 5.458206 | 5.616297 | 6.24542 | 5.289928 |
| 11.8942 | 11.86814 | 12.01419 | 11.33255 | 11.40149 | 11.5826 | 11.56204 | 11.7247 | 10.78863 |
| 6.869355 | 7.147992 | 6.566303 | 6.294101 | 6.61546 | 6.591854 | 5.784918 | 5.967177 | NaN |
| 5.965965 | NaN | NaN | 9.267914 | 8.679398 | 6.593476 | 7.073155 | 6.903657 | NaN |
| 6.225634 | 5.462172 | 5.952461 | 6.438584 | 6.279004 | 7.904531 | 7.786921 | 7.19501 | NaN |
| 6.962556 | 7.710134 | 7.655036 | 9.263844 | 8.194618 | 6.16238 | 6.847873 | 6.630466 | 8.58731 |
| 6.085851 | 5.336365 | 5.759966 | 5.652651 | 5.749108 | 4.911826 | NaN | 5.242169 | 5.350786 |
| 7.657917 | 7.864612 | 7.740195 | 6.73219 | 6.800482 | 7.934858 | 8.14415 | 7.793791 | 8.359093 |
| 6.536015 | 7.434245 | 7.026542 | 10.15166 | 9.872155 | 4.495377 | 5.141362 | 3.532268 | 5.808772 |
| 7.228284 | 7.150714 | 6.933591 | 6.727512 | 7.0715 | 5.850879 | 5.966955 | 6.3836 | 5.744308 |
| NaN | NaN | NaN | 5.572643 | 5.802143 | NaN | NaN | NaN | 7.279969 |
| 6.17562 | 6.359951 | 6.259926 | 5.301239 | 5.269614 | 6.97529 | 6.804191 | 6.952468 | 6.597638 |
| 7.012625 | 7.245867 | 7.011756 | 8.679899 | 8.672197 | 7.750854 | 7.336154 | 7.417931 | 5.822544 |
| 8.767467 | 7.978915 | 7.975498 | NaN | 8.021097 | 7.95039 | NaN | NaN | 6.373139 |
| 11.61665 | 12.64508 | 12.81643 | 13.96933 | 13.45996 | 10.64297 | 10.96432 | 10.64852 | 9.619794 |
| 4.999213 | 6.251853 | 6.289635 | 5.469274 | NaN | 5.354069 | 5.150906 | 6.150634 | 5.281357 |
| 9.35572 | 9.453106 | 9.299173 | 9.247071 | 9.294858 | 8.969851 | 8.943001 | 9.020459 | 8.96284 |
| 7.822159 | 7.681142 | 7.757843 | 6.367457 | 8.495785 | 7.354499 | 7.517495 | 7.115839 | 7.737142 |
| 7.428648 | 7.497221 | 7.449042 | 8.798589 | 9.123894 | 7.556706 | 7.531689 | 7.828737 | 8.09056 |
| 6.796697 | 5.99539 | 6.028421 | 6.341194 | NaN | 6.228875 | 6.971545 | 6.200086 | 5.384947 |
| 3.740094 | NaN | 3.528109 | NaN | NaN | NaN | NaN | 4.964499 | 5.118956 |
| 6.61797 | 5.230362 | 6.006318 | NaN | 6.134118 | 5.999372 | 6.92978 | NaN | 6.53873 |
| 6.964219 | 7.052579 | 7.286049 | 5.771019 | 6.180113 | NaN | NaN | NaN | NaN |
| 7.314048 | NaN | 7.362583 | NaN | NaN | NaN | 7.25318 | NaN | 6.888069 |
| 6.151029 | 6.000973 | 6.046602 | 6.765814 | 6.765054 | 6.489376 | 6.23322 | 6.051289 | 5.789639 |
| 6.399161 | 6.677968 | 6.483731 | 6.851133 | 6.947263 | 6.09759 | 6.269352 | 6.345468 | 6.077367 |
| NaN | NaN | 5.358295 | 5.234122 | 4.872946 | 3.966851 | NaN | 4.346676 | 5.753104 |
| 6.255507 | 6.100871 | 5.552536 | 6.139178 | 6.205822 | 5.779431 | 5.783524 | 5.968427 | 6.611956 |
| 3.830817 | 7.134478 | 3.944208 | NaN | 4.850237 | NaN | NaN | 3.634054 | 2.940734 |
| 7.794263 | 7.686231 | 7.856486 | 6.56188 | 6.689846 | 7.947323 | 7.690701 | 7.864715 | 7.468241 |
| 7.799608 | 7.319317 | 6.944982 | 9.570002 | 9.695004 | 7.326188 | 7.136114 | 6.774617 | 7.440233 |
| 6.555724 | 6.242646 | 6.749519 | 4.219348 | 5.398657 | 6.074495 | 6.388426 | 6.217088 | 6.231083 |
| NaN | 5.472648 | NaN | 7.989698 | 7.741476 | 6.868673 | 5.453315 | NaN | 5.797564 |
| 5.624151 | 5.225064 | 5.012143 | NaN | 4.087091 | 5.560729 | 5.42982 | 4.839585 | 5.423812 |
| 8.302309 | 8.065414 | 8.128158 | 6.600834 | 6.443395 | 8.643253 | 8.700013 | 8.756533 | 8.440647 |
| 8.673417 | 8.21032 | 8.512942 | 6.710488 | 6.781851 | 8.241586 | 8.349995 | 8.34445 | 8.594801 |
| 8.555825 | 8.296538 | 8.382149 | 7.470117 | 7.33694 | 8.249551 | 8.153765 | 7.971509 | 7.630555 |
| 8.927546 | 8.4291 | 8.632132 | 7.701151 | 7.920553 | 8.367826 | 8.040494 | 8.136912 | 7.762369 |
| 6.720358 | 6.798805 | 6.196718 | 6.294832 | 6.260683 | 6.219714 | 6.255531 | 5.792686 | 6.206108 |
| 3.745111 | NaN | 5.425927 | NaN | NaN | NaN | NaN | NaN | 5.78688 |
| 8.82273 | 8.739115 | 8.681949 | 8.146902 | 7.700091 | 8.421825 | 8.026199 | 8.083005 | 6.815573 |
| NaN | NaN | NaN | 6.897721 | 7.504741 | NaN | NaN | NaN | NaN |
| 5.145394 | 5.587258 | 5.729552 | 6.271316 | 6.369981 | 5.794597 | 6.390769 | 6.32569 | 3.38061 |

| | | | | | | | | |
|---|---|---|---|---|---|---|---|---|
| 6.934427 | 7.036366 | 7.696497 | 8.146821 | 8.365952 | 6.991122 | 7.183819 | 6.724059 | 6.566544 |
| 8.761598 | 8.429155 | 8.325358 | 9.887547 | 10.18395 | 10.44546 | 10.03844 | 10.23477 | 8.715964 |
| 13.28954 | 13.31144 | 13.48023 | 13.17936 | 13.25392 | 13.80293 | 13.99912 | 13.78405 | 12.98718 |
| 6.664352 | 5.412515 | 5.470127 | 6.364372 | 6.119007 | 6.459327 | 6.340119 | 6.288611 | NaN |
| NaN | NaN | NaN | NaN | NaN | NaN | NaN | NaN | 9.007889 |
| 9.760538 | 9.898894 | 9.879771 | 10.36036 | 9.973906 | 8.827782 | 9.167782 | 8.818866 | 8.358577 |
| 6.914934 | 6.977055 | 6.610698 | 8.093001 | NaN | 6.972517 | 7.207752 | 6.975606 | 7.127855 |
| 6.779379 | 6.689481 | 6.270559 | 6.238212 | 5.739116 | 6.828808 | 6.740802 | 6.857914 | 6.36877 |
| 10.1292 | 10.08925 | 10.07841 | 10.35115 | 9.567301 | 9.952641 | 10.24283 | 9.7123 | 9.567039 |
| NaN | 4.158681 | 5.369675 | 7.552421 | 7.55734 | 6.536065 | 6.880013 | 7.016147 | 6.348864 |
| 8.832325 | 8.013347 | 7.640539 | 6.51734 | 6.869467 | 8.100358 | 7.220668 | 7.506642 | 6.641767 |
| 9.293721 | 8.789245 | 9.549762 | 10.89505 | 10.55277 | 6.487732 | 6.321571 | 6.307762 | 7.376961 |
| 11.21394 | 11.06937 | 11.24619 | 13.14794 | 12.94507 | 11.04735 | 11.01906 | 10.99407 | 11.15421 |
| 9.120328 | 9.268172 | 9.108887 | 7.842257 | 7.902744 | 8.333311 | 8.604468 | 8.61285 | 9.069274 |
| 10.06244 | 9.847805 | 9.956277 | 9.570499 | 9.066904 | 8.584102 | 8.702299 | 8.741151 | 8.139088 |
| 2.381263 | NaN | 3.867893 | -0.38444 | 2.51113 | 3.011665 | 2.746246 | 3.125552 | 1.037618 |
| 8.390594 | 8.353677 | 8.298257 | 7.073915 | 6.475914 | 7.165936 | 7.481133 | 7.731146 | 7.116624 |
| 9.096786 | 8.932244 | 9.073806 | 10.68209 | 10.68406 | 9.798271 | 9.189829 | 9.713609 | 8.736825 |
| 4.620237 | 5.44208 | 5.346911 | 6.294693 | 6.505774 | 4.382408 | 4.016535 | 4.795213 | 4.390451 |
| 10.3195 | 10.1296 | 10.2472 | 10.27085 | 10.10561 | 10.4681 | 10.30335 | 10.60409 | 10.22559 |
| 13.99944 | 13.89087 | 14.02636 | 13.19018 | 12.97926 | 14.11008 | 14.24782 | 14.17096 | 13.46308 |
| 9.833385 | 9.356057 | 9.279803 | 7.958677 | 7.523372 | 10.38677 | 10.29049 | 10.44883 | 7.984724 |
| 11.99058 | 11.93599 | 11.94673 | 10.96203 | 10.79256 | 11.9346 | 11.65307 | 11.5573 | 11.79198 |
| 10.74519 | 10.70308 | 10.59416 | 10.61361 | 10.53164 | 10.06886 | 9.238873 | 10.16558 | 10.27815 |
| 9.328933 | 9.255271 | 9.146297 | 9.998838 | 9.962872 | 8.856847 | 8.835161 | 8.872651 | 8.345078 |
| 8.461867 | 8.401919 | 8.330693 | 7.374236 | 7.501772 | 8.606649 | 8.417422 | 8.303808 | 8.271307 |
| 10.20052 | 9.985968 | 10.11508 | 10.24377 | 10.35986 | 9.470224 | 9.491158 | 9.605529 | 8.8104 |
| 7.280286 | 7.270093 | 6.468974 | 7.721723 | 7.407314 | 7.982286 | 7.935256 | 7.901243 | 4.794522 |
| 9.631191 | 9.81472 | 9.610025 | 9.724346 | 8.776458 | 10.30824 | 10.03627 | 10.2469 | 9.643274 |
| 9.67979 | 9.960854 | 10.02482 | 9.84426 | 9.758091 | 9.927335 | 10.01508 | 9.891539 | 9.871695 |
| 8.997566 | 9.034778 | 9.256165 | 9.309587 | 9.385998 | 8.243428 | 8.805661 | 8.730949 | 8.504047 |
| 7.486124 | 7.188423 | 7.216831 | NaN | NaN | NaN | NaN | NaN | NaN |
| 8.092556 | 7.805595 | 7.960431 | 6.967073 | 7.063508 | 7.824462 | 7.996016 | 7.988158 | 7.791006 |
| 9.528498 | 8.472098 | 8.155232 | 9.462687 | 9.208457 | 8.138387 | 8.077449 | 7.696295 | 8.519772 |
| 7.673605 | 7.607081 | 7.672784 | 6.987711 | 7.083155 | 7.628401 | 7.532387 | 7.750375 | 7.766749 |
| 7.881935 | 8.255214 | 8.182785 | 8.077724 | 7.297984 | 8.098942 | 7.813006 | 7.673566 | 7.175464 |
| 11.65524 | 11.43375 | 11.01449 | 11.23978 | 11.44271 | 11.36434 | 11.54934 | 11.51037 | 11.1073 |
| 9.047354 | 8.904428 | 8.98931 | 9.752232 | 9.817003 | 9.141647 | 8.841876 | 9.364959 | 8.59858 |
| 7.525819 | 7.127687 | 7.532054 | 6.616089 | 6.643692 | NaN | 5.775819 | 5.089442 | 8.019139 |
| 10.54882 | 10.89376 | 10.85475 | 9.859873 | 9.701642 | 11.08863 | 11.06852 | 11.13303 | 10.53934 |
| 9.153875 | 8.427123 | 8.985656 | 10.15561 | 9.640656 | 8.326664 | 8.18893 | 8.919807 | 9.13284 |
| 9.318598 | 9.120642 | 9.368031 | 8.965681 | 8.778543 | 8.404829 | 8.788361 | 8.519802 | 8.737234 |
| 9.935499 | 9.53897 | 9.450778 | 10.48343 | 10.43611 | 9.916845 | 9.177561 | 9.661676 | 9.583994 |
| 10.5858 | 10.27245 | 10.55571 | 12.7017 | 12.55474 | 10.85555 | 11.02177 | 10.68896 | 10.69441 |
| 9.470943 | 9.274352 | 9.307491 | 9.68008 | 9.431876 | 9.450215 | 9.766673 | 9.729706 | 8.666588 |
| 5.93189 | 4.688219 | 4.796671 | 5.900031 | 5.684869 | 5.31988 | 5.514503 | 4.776677 | 6.221053 |
| 9.982505 | 10.06922 | 9.928893 | 8.9081 | 8.840127 | 10.32595 | 10.26984 | 10.22496 | 9.840032 |

| | | | | | | | | |
|---|---|---|---|---|---|---|---|---|
| 8.242671 | 8.330606 | 8.2966 | 7.49305 | 7.614763 | 8.339489 | 8.336703 | 8.399774 | 8.297448 |
| 10.69362 | 10.90895 | 10.64593 | 11.11372 | 11.2786 | 11.63773 | 11.60701 | 11.70383 | 10.6139 |
| 9.09503 | 9.227498 | 9.194343 | 8.134557 | 7.95775 | 9.256151 | 9.242717 | 9.31972 | 9.358891 |
| 8.243064 | 8.31316 | 8.328749 | 7.434848 | 7.349011 | 8.171802 | 7.910633 | 8.158934 | 8.464897 |
| 10.52585 | 10.57623 | 10.4089 | 11.51197 | 11.31356 | 9.857861 | 9.902721 | 10.25886 | 9.995937 |
| 7.958894 | 7.910308 | 7.815705 | 7.422697 | 7.809136 | 7.680645 | 7.856305 | 7.831691 | 7.522159 |
| NaN | NaN | NaN | 8.320482 | 6.370976 | NaN | NaN | NaN | NaN |
| 9.444793 | 9.539101 | 9.372359 | 9.267244 | 9.239552 | 9.840258 | 9.744727 | 9.777197 | 9.502361 |
| 9.087898 | 8.945943 | 9.141743 | 10.31955 | 9.804934 | 9.243006 | 9.160837 | 9.065169 | 8.873302 |
| 8.558684 | 8.58853 | 8.609391 | 7.911042 | 8.093769 | 8.643645 | 8.840986 | 8.987839 | 7.407823 |
| 7.473608 | 7.644567 | 7.385912 | 8.369547 | 8.371308 | 7.165299 | 6.939027 | 6.845601 | 7.577285 |
| 10.16085 | 10.15951 | 9.847048 | 8.637893 | 8.212414 | 10.46909 | 10.12304 | 10.40323 | 9.684652 |
| 6.328187 | 6.344213 | NaN | 6.86507 | 6.949155 | NaN | NaN | NaN | NaN |
| 10.22102 | 9.935815 | 9.952943 | 10.574 | 10.66714 | 10.00211 | 10.02476 | 10.19585 | 9.765268 |
| 8.766965 | 8.636312 | 8.680307 | 7.465743 | 7.437536 | 8.523042 | 8.503991 | 8.577755 | 8.194875 |
| 8.315097 | 7.47387 | 7.528195 | 10.10733 | 9.478883 | 7.535271 | 7.543509 | 7.432671 | 7.032745 |
| 9.482677 | 9.663434 | 9.655202 | 9.257998 | 9.134941 | 9.254821 | 8.28518 | 9.343938 | 8.242512 |
| 8.592782 | 8.519984 | 8.470521 | 8.8627 | 7.964973 | 7.828683 | 7.750059 | 8.308302 | 7.813922 |
| 7.375667 | 7.509069 | 7.437008 | 5.809872 | 5.401466 | 6.801138 | 6.985311 | 7.36536 | 6.559985 |
| NaN | NaN | NaN | 6.042093 | 6.317427 | NaN | NaN | NaN | 6.424646 |
| 9.063602 | 8.73037 | 8.759644 | 9.551202 | 9.937566 | 8.351545 | 8.513541 | 8.687707 | 9.229238 |
| 10.77785 | 10.53835 | 10.68682 | 12.02899 | 11.76165 | 9.052501 | 8.980573 | 9.323957 | 9.807662 |
| 8.133488 | 8.323334 | 9.124559 | 7.259078 | NaN | 7.957173 | 7.358739 | 7.452889 | 7.980892 |
| 9.00724 | 8.271995 | 8.343268 | 6.693127 | 6.185194 | NaN | NaN | NaN | 8.262798 |
| 9.493241 | 9.1769 | 9.253205 | 10.0384 | 9.858283 | 9.291761 | 9.413314 | 9.454089 | 9.371214 |
| 11.32062 | 11.21755 | 11.22374 | 10.67294 | 10.7549 | 11.56628 | 11.58685 | 11.44106 | 10.36604 |
| 7.463732 | 7.341216 | 7.283627 | 7.258581 | 7.081502 | 7.239707 | 7.4674 | 6.8055 | 7.36674 |
| 9.594969 | 8.102553 | 9.101321 | 8.088171 | 7.534261 | 7.199172 | 7.271405 | 7.687452 | 8.506067 |
| 7.87535 | 7.344031 | 7.399251 | 9.256948 | 9.23422 | 6.874803 | 6.70527 | 6.818138 | 7.321254 |
| 8.167748 | 7.98722 | 7.743645 | 7.217719 | 7.183081 | 8.078444 | 7.971813 | 8.083497 | 8.619061 |
| 6.31803 | 6.511483 | 7.091515 | 7.82262 | 7.807861 | 7.294766 | 6.545559 | NaN | 6.428004 |
| 10.94206 | 10.83122 | 11.27069 | 11.82867 | 11.84062 | 9.453948 | 9.729576 | 9.157193 | 10.5811 |
| 6.645 | 6.898289 | 7.094299 | 9.796933 | 10.16539 | 6.736342 | 6.438591 | 6.764596 | 7.73563 |
| 9.096614 | 9.352974 | 9.342308 | 7.938568 | 8.157586 | 8.936955 | 8.782017 | 8.824795 | 9.256767 |
| 8.45297 | 8.666734 | 8.599072 | 8.468378 | 8.369417 | 9.094202 | 8.784641 | 8.891017 | 8.652712 |
| 7.30443 | 6.079877 | 7.222488 | 5.527619 | 5.465804 | 6.658062 | 7.687066 | 7.940487 | 6.761614 |
| 9.337908 | 8.941932 | 9.138651 | 8.763902 | 9.059932 | 8.916815 | 8.994051 | 8.865012 | 8.52991 |
| 12.92492 | 12.78886 | 12.86264 | 13.12945 | 13.02026 | 12.16538 | 12.12924 | 12.15591 | 12.48223 |
| 12.78561 | 12.81071 | 12.76204 | 13.04695 | 12.95464 | 12.72738 | 12.57054 | 12.64901 | 12.297 |
| 8.668191 | 7.526249 | 7.120519 | NaN | NaN | 8.563291 | 8.257786 | 8.227564 | 8.354258 |
| 6.984098 | 6.647817 | 7.271425 | 6.176512 | 6.673793 | 6.414593 | 6.517044 | 6.311905 | 5.99288 |
| 8.944717 | 8.379223 | 8.725471 | 9.714837 | 9.866297 | 8.200595 | 7.877793 | 8.49157 | 8.885232 |
| 11.98837 | 11.78706 | 12.09926 | 13.7674 | 13.95949 | 12.74896 | 12.6322 | 12.73724 | 12.18159 |
| 10.74908 | 10.88629 | 10.53499 | 9.779518 | 9.742174 | 9.924332 | 10.1468 | 10.06413 | 10.56043 |
| 9.605113 | 9.53919 | 9.26593 | 9.068238 | 8.446125 | 8.420569 | 8.679445 | 8.417892 | 8.777161 |
| 9.460717 | 9.483201 | 9.510536 | 11.06528 | 10.95494 | 9.550876 | 9.323584 | 9.514502 | 9.40417 |
| 10.29615 | 10.20761 | 10.30393 | 10.73958 | 10.71303 | 10.24673 | 10.13648 | 10.06799 | 10.36248 |

| | | | | | | | | |
|---|---|---|---|---|---|---|---|---|
| 10.82205 | 10.8111 | 10.88839 | 11.20303 | 11.41349 | 11.07782 | 10.95285 | 10.87212 | 10.87521 |
| 10.55926 | 10.46859 | 10.57845 | 10.26135 | 10.6142 | 11.26116 | 11.51401 | 11.35444 | 11.18732 |
| 10.67229 | 10.67346 | 10.79085 | 10.27401 | 10.04568 | 11.22095 | 10.98447 | 10.70725 | 10.53495 |
| 11.13412 | 11.19261 | 11.27165 | 11.21877 | 11.26583 | 11.46319 | 11.54042 | 11.59206 | 11.29983 |
| 7.879318 | 7.632782 | 8.051216 | 7.898244 | 8.423876 | 7.382443 | 6.852108 | 7.381914 | 7.872762 |
| 7.626916 | 7.431822 | 7.833599 | 8.065645 | 8.25434 | 7.455545 | 7.079066 | 7.756106 | 7.414915 |
| 14.12512 | 14.06079 | 14.1576 | 14.57646 | 14.56561 | 13.82399 | 13.84959 | 13.73837 | 13.78915 |
| 11.64506 | 11.82927 | 11.62315 | 11.33637 | 11.56864 | 11.48821 | 11.43659 | 11.35615 | 11.33757 |
| 10.28618 | 10.32943 | 10.16503 | 10.64848 | 10.13867 | 9.931934 | 9.401205 | 9.682133 | 9.478262 |
| 12.60218 | 12.60526 | 12.57459 | 13.7768 | 13.90093 | 13.24569 | 13.26959 | 13.18823 | 12.90237 |
| 7.920402 | 4.522074 | 7.819589 | 7.113699 | 7.589409 | 9.289553 | 8.318266 | 9.315459 | 6.161367 |
| 11.04265 | 10.78976 | 11.08164 | 12.30043 | 12.34073 | 11.6532 | 11.45884 | 11.65356 | 10.71378 |
| 9.0853 | 9.027753 | 9.050668 | 9.228275 | 9.253393 | 9.555655 | 9.15274 | 9.456101 | 9.36216 |
| 10.23559 | 10.43388 | 10.6635 | 10.71034 | 10.96094 | 10.05385 | 9.416591 | 9.960264 | 10.25302 |
| 10.77514 | 10.6722 | 10.8396 | 11.56635 | 11.6482 | 10.78604 | 10.88989 | 10.95613 | 11.07377 |
| 8.548101 | 8.557274 | 8.927534 | 9.449877 | 8.118906 | 7.485642 | 7.107775 | 7.287589 | 7.342279 |
| 8.835035 | 8.379246 | 8.669906 | 8.887617 | 8.121439 | 7.973788 | 8.569983 | 8.715152 | 7.962691 |
| 9.198592 | 8.849599 | 8.948088 | 10.25824 | 10.00962 | 8.708317 | 8.821855 | 9.120021 | 8.748352 |
| 8.773573 | 8.698176 | 8.556685 | 10.26906 | 10.01448 | 9.048608 | 9.399629 | 9.017204 | 10.05053 |
| 11.5657 | 11.55652 | 11.4566 | 12.63235 | 12.47065 | 11.01047 | 10.90084 | 10.69349 | 11.5689 |
| 12.56973 | 11.77713 | 12.55527 | 13.31698 | 12.93707 | 10.31495 | 10.30133 | 10.0613 | 10.52619 |
| 9.381393 | 9.214709 | 9.277887 | 8.811887 | 8.824112 | 8.316705 | 8.770593 | 8.636018 | 8.46583 |
| 10.31723 | 10.12091 | 10.12106 | 9.906005 | 9.930814 | 10.32526 | 10.41169 | 10.26383 | 10.30944 |
| 8.140167 | 8.179415 | 8.054232 | 6.704328 | 5.93701 | 7.654217 | 7.820997 | 8.258664 | 7.297801 |
| 7.585294 | 8.063092 | 7.418582 | 6.40351 | 5.681348 | 7.485864 | 7.098138 | 7.182177 | 6.743402 |
| 10.34927 | 10.40224 | 10.4061 | 10.37827 | 10.42446 | 11.175 | 10.90545 | 11.04941 | 10.01339 |
| 7.556752 | 7.904269 | 7.489606 | 7.445142 | 6.807518 | NaN | 5.467508 | NaN | 6.643482 |
| 8.486576 | 8.51185 | 8.505263 | 8.37756 | 8.458488 | 8.486711 | 8.501034 | 8.443416 | 8.726838 |
| 6.907827 | 6.879129 | 7.27949 | 8.172441 | 8.256471 | NaN | NaN | NaN | 5.88906 |
| 6.710605 | 6.807375 | 6.6954 | 7.166021 | 7.21092 | 6.119719 | 5.709575 | 5.977694 | 6.134282 |
| 9.834201 | 9.674717 | 9.803356 | 8.782131 | 8.553929 | 9.739555 | 9.726198 | 9.793097 | 9.855817 |
| 11.02158 | 10.93343 | 11.16608 | 9.809048 | 9.548546 | 10.18473 | 10.25632 | 10.20455 | 10.27887 |
| 12.56732 | 12.54501 | 12.59506 | 11.77986 | 11.81044 | 12.4858 | 11.82466 | 12.03772 | 12.34376 |
| 8.08028 | 8.117523 | 8.82148 | 6.915103 | 6.90283 | 5.386008 | 5.983379 | 6.239488 | 5.612699 |
| 10.87038 | 10.82121 | 10.83256 | 11.38573 | 11.47469 | 10.968 | 10.90207 | 11.07118 | 11.14472 |
| 8.812205 | 8.842804 | 8.698652 | 8.813375 | 8.837846 | 8.384475 | 8.75026 | 8.968967 | 9.143677 |
| 7.919181 | 7.926045 | 7.878349 | 9.044711 | 8.867585 | 7.846235 | 7.626545 | 7.925264 | 6.168954 |
| 14.29604 | 14.30816 | 14.2781 | 14.0484 | 14.18758 | 14.45113 | 14.48417 | 14.35964 | 13.77209 |
| 8.41117 | 8.849578 | 8.756599 | 7.286202 | 7.525316 | 9.065166 | 8.800529 | 8.858328 | 8.167607 |
| 11.78217 | 11.65875 | 11.86979 | 11.39477 | 11.40003 | 11.50706 | 11.55526 | 11.63291 | 11.98525 |
| 8.422007 | 8.146072 | 8.452345 | 8.901205 | 8.657048 | 8.677159 | 8.659846 | 8.472286 | 8.818392 |
| NaN | NaN | 7.376976 | 10.81853 | 12.22162 | NaN | NaN | NaN | NaN |
| 8.042085 | 8.020763 | 7.9779 | 10.14489 | 10.25224 | 8.987615 | 8.899096 | 8.904849 | 8.743066 |
| 10.05638 | 9.742847 | 9.813603 | 10.1845 | 10.38112 | 10.28011 | 10.39111 | 10.30077 | 10.18352 |
| 9.866013 | 9.963608 | 10.46901 | 10.87535 | 10.84513 | 9.403447 | 10.36909 | 10.21885 | 10.03869 |
| 9.634412 | 9.482777 | 9.666092 | 9.342556 | 9.598863 | 10.15385 | 9.790519 | 10.22923 | 10.09078 |
| 9.606591 | 9.68303 | 9.387402 | 10.9835 | 11.00326 | 8.247966 | 8.607042 | 7.941102 | 9.441978 |

| | | | | | | | | |
|---|---|---|---|---|---|---|---|---|
| 7.894455 | 8.186321 | 8.265742 | 9.131612 | 9.584394 | 8.453707 | 8.548885 | 7.215343 | 6.996302 |
| 9.918045 | 9.926826 | 9.92109 | 8.751291 | 8.843613 | 8.915606 | 9.492723 | 9.614043 | 9.178012 |
| 6.298224 | NaN | 4.21699 | NaN | 3.81643 | NaN | NaN | NaN | NaN |
| 8.342321 | 8.687964 | 8.587981 | 9.847585 | 9.915824 | 9.144473 | 9.331735 | 8.825503 | 9.057706 |
| 10.08364 | 10.1206 | 10.10459 | 9.910604 | 9.37921 | 9.246985 | 9.606718 | 9.482697 | 9.356076 |
| 8.253388 | 8.393557 | 8.410804 | 9.756236 | 9.28269 | 7.749914 | 7.748103 | 8.015863 | 8.580263 |
| NaN | 6.889379 | NaN | 9.913975 | 5.306277 | 6.832375 | 7.279037 | NaN | 6.568634 |
| 11.10791 | 10.84329 | 10.84704 | 12.13303 | 12.13501 | 11.68153 | 11.53198 | 11.73454 | 10.66522 |
| 8.966896 | 9.059547 | 9.102272 | 9.589928 | 9.619602 | 9.735771 | 9.539953 | 9.618431 | 9.324558 |
| 8.71594 | 8.56057 | 8.636831 | 8.699463 | 8.703771 | 8.68191 | 8.6555 | 8.866615 | 8.766832 |
| 8.819572 | 8.850792 | 8.977024 | 7.690137 | 7.547988 | 8.461871 | 8.560193 | 8.552459 | 8.606494 |
| 9.403563 | 9.190038 | 9.35901 | 9.272036 | 9.301008 | 10.42453 | 10.30237 | 10.47173 | 9.322561 |
| 11.77534 | 11.70065 | 11.78675 | 12.48118 | 12.44634 | 11.65549 | 11.87969 | 11.97507 | 11.65621 |
| 8.863621 | 9.124516 | 9.014194 | 10.36826 | 10.20335 | 8.703799 | 8.714977 | 8.541952 | 8.448399 |
| 8.256347 | 8.095031 | 7.820983 | 9.648618 | 9.495685 | 7.582527 | 7.619956 | 7.758246 | 7.281127 |
| 10.69263 | 10.77621 | 10.80222 | 10.48311 | 10.5803 | 10.98292 | 10.47565 | 10.80302 | 10.06132 |
| 8.453384 | 9.367186 | 9.225461 | 8.686083 | 8.029543 | 9.223096 | 8.904639 | 8.821994 | 8.701077 |
| 9.839961 | 9.849618 | 9.733732 | 9.070922 | 9.057147 | 10.12159 | 10.20217 | 10.13228 | 9.842711 |
| 10.83063 | 10.35717 | 10.4112 | 11.3012 | 10.77756 | 11.05999 | 10.33579 | 10.9478 | 10.09807 |
| 11.34691 | 10.63044 | 10.80851 | 9.488471 | 9.301655 | 10.72818 | 10.58224 | 10.61746 | 10.41886 |
| 6.90108 | 6.848622 | 6.787054 | 5.359902 | 4.963484 | 6.447593 | 6.380622 | 6.557772 | 5.645121 |
| 8.611306 | 8.455691 | 8.319678 | 9.382277 | 9.591888 | 7.701528 | 7.459837 | 7.732296 | 7.244146 |
| 13.07374 | 13.05067 | 13.14715 | 12.60854 | 12.40562 | 12.26797 | 12.51587 | 12.27087 | 12.54839 |
| 7.190028 | 6.77879 | 7.494279 | 7.227188 | 7.527692 | 7.821332 | 7.420638 | 7.326848 | 7.356007 |
| 9.701131 | 9.585401 | 9.614471 | 8.951446 | 9.063379 | 9.475467 | 9.349748 | 9.527811 | 9.470263 |
| 7.083802 | 7.582461 | 7.600324 | 7.312704 | 6.921411 | 6.604573 | 6.462147 | 6.219647 | 7.168654 |
| 9.680988 | 9.449081 | 9.494938 | 9.900068 | 9.82513 | 9.421157 | 9.698467 | 9.25704 | 8.234816 |
| 9.621412 | 9.432611 | 9.625923 | 8.942643 | 8.491767 | 7.743361 | 8.868081 | 8.392661 | 7.929863 |
| 11.65561 | 11.61672 | 11.54699 | 11.78344 | 11.36382 | 9.718041 | 9.882026 | 9.118547 | 10.36424 |
| 5.910421 | 7.285674 | 7.259161 | 6.375916 | 5.870693 | 7.196767 | 7.121297 | 6.88681 | 6.858678 |
| 5.201431 | 5.882714 | 4.36398 | 8.040046 | 6.154277 | 6.179007 | 6.856426 | 5.968289 | 6.360028 |
| 8.658337 | 8.83636 | 8.879924 | 9.847001 | 9.725771 | 8.03911 | 8.148082 | 7.890738 | 8.507483 |
| 9.425981 | 9.570837 | 9.480335 | 10.09946 | 9.646426 | 8.937378 | 8.771775 | 8.976569 | 9.881085 |
| 10.19329 | 10.45735 | 10.51578 | 9.766036 | 9.654644 | 9.769746 | 9.838575 | 9.973924 | 10.68917 |
| 10.4215 | 10.82609 | 10.6763 | 12.62991 | 12.64145 | 10.65648 | 10.54114 | 10.60771 | 11.72496 |
| 8.068982 | 7.71701 | 7.74922 | 7.055992 | 7.388195 | 7.961767 | 7.80246 | 6.976243 | 7.777543 |
| NaN | NaN | 7.652203 | 8.034474 | 8.005497 | 7.886444 | 7.897668 | 8.093328 | 7.710357 |
| 9.477574 | 9.987212 | 9.789491 | 11.37855 | 11.51124 | 8.327049 | 7.882003 | 8.012374 | 8.566944 |
| 12.40621 | 11.98074 | 12.04047 | 14.20629 | 14.14631 | 12.13503 | 12.17784 | 12.12712 | 13.06847 |
| 6.870254 | 6.488116 | 7.512774 | 6.919818 | 6.656691 | 6.263913 | 6.122683 | 6.315381 | 6.084464 |
| 8.424352 | 8.45847 | 8.593072 | 8.062073 | 8.272376 | 8.299598 | 8.019832 | 8.485651 | 7.628981 |
| 11.45833 | 11.48555 | 11.41554 | 10.1212 | 10.14667 | 9.059705 | 8.975978 | 9.232648 | 8.74129 |
| 9.654438 | 9.675395 | 9.6374 | 10.56328 | 10.51789 | 10.07548 | 9.960483 | 10.1479 | 9.843244 |
| 13.25933 | 13.18687 | 12.95177 | 12.85933 | 12.77105 | 13.18812 | 13.32805 | 13.2257 | 14.03812 |
| NaN | 7.70058 | NaN | 7.147459 | 7.879846 | NaN | NaN | 7.243231 | NaN |
| 9.35725 | 9.86449 | 10.18231 | 9.70492 | 10.02803 | 12.16085 | 12.22594 | 11.78606 | 9.134487 |
| 14.43363 | 14.28151 | 14.35405 | 13.68118 | 13.55831 | 14.36185 | 14.4244 | 14.5683 | 13.73251 |

-Bruker Confidential-

| | | | | | | | | |
|---|---|---|---|---|---|---|---|---|
| 8.627058 | 8.402281 | 8.381249 | 8.419347 | 8.290907 | 8.198782 | 8.252598 | 8.261614 | 6.465583 |
| 14.56248 | 14.6231 | 14.62072 | 13.9043 | 14.02885 | 15.10667 | 15.14163 | 15.17166 | 14.29346 |
| 10.17983 | 9.996477 | 10.14119 | 9.419724 | 9.017555 | 10.25872 | 10.06407 | 9.881658 | 9.360794 |
| 10.40365 | 10.26406 | 10.48738 | 11.1559 | 11.07962 | 10.66732 | 10.46482 | 10.73278 | 10.16871 |
| 13.92385 | 13.72301 | 13.81834 | 14.37477 | 14.26923 | 14.32988 | 14.27693 | 14.34075 | 13.48486 |
| 5.740627 | 5.115295 | 4.364467 | 6.457603 | 6.187872 | NaN | NaN | 6.134664 | 5.767993 |
| 8.742943 | 8.468273 | 8.733142 | 8.646163 | 9.415976 | 9.443473 | NaN | 8.81038 | 11.57684 |
| 6.740796 | 6.38461 | 6.501556 | 5.737578 | 5.64092 | 6.925911 | 6.341527 | 6.449337 | 7.724118 |
| 10.46201 | 10.40134 | 10.19975 | 9.064482 | 8.673301 | 9.73458 | 9.71813 | 9.883544 | 9.286142 |
| 8.378642 | 8.592101 | 8.459411 | 9.187385 | 9.296379 | 8.153997 | 8.081809 | 8.132334 | 7.857096 |
| 9.408471 | 9.515408 | 9.253469 | 7.109783 | 7.433186 | 9.75573 | 9.717053 | 9.876041 | 6.827965 |
| 8.180157 | 8.156898 | 7.476286 | 9.715269 | 8.891312 | 7.799731 | 7.754028 | 7.9914 | 7.039221 |
| 7.253371 | 7.446076 | 7.076452 | 7.863086 | 7.802971 | 7.009497 | 7.106174 | 6.794773 | 7.244415 |
| 9.51387 | 9.486922 | 9.593418 | 9.045228 | 8.9476 | 9.338708 | 9.508513 | 9.394969 | 9.827777 |
| 9.052023 | 8.89417 | 8.98017 | 8.347307 | 8.305558 | 9.891199 | 9.397196 | 9.716613 | 9.578289 |
| 6.84031 | 7.481676 | 7.261977 | 6.53839 | 6.215466 | 7.328064 | 7.21553 | 7.143746 | 6.782854 |
| 10.23473 | 10.18003 | 10.30731 | 9.789719 | 9.805065 | 9.962226 | 9.902903 | 10.13263 | 9.901386 |
| 6.583485 | 6.330076 | 6.320926 | 6.584877 | 6.801852 | 6.94801 | 7.014705 | 7.108756 | 6.797724 |
| 8.061948 | 7.95962 | 7.938854 | 7.654333 | 7.908671 | 8.049234 | 8.23907 | 7.851372 | 7.808027 |
| 6.655214 | 6.591407 | 6.428132 | 4.950644 | 4.44688 | 5.611873 | 5.455083 | 5.337213 | 5.528274 |
| 9.15083 | 9.263388 | 9.122742 | 8.517937 | 8.57174 | 9.261506 | 9.246915 | 9.493336 | 8.693262 |
| 8.073853 | 7.649466 | 7.736654 | 7.521592 | 7.511218 | 6.004019 | 6.537335 | 6.600595 | 7.317698 |
| 6.305823 | 6.445011 | 5.802612 | NaN | NaN | 6.395508 | 6.318751 | 6.52909 | 5.544273 |
| 11.4248 | 11.19261 | 11.24257 | 11.33884 | 11.19114 | 11.09243 | 11.10486 | 11.20865 | 11.22067 |
| 7.878454 | 7.873292 | 7.999074 | 6.673573 | 6.970287 | 8.087432 | 8.168402 | 8.354386 | 7.912388 |
| 9.624557 | 9.847762 | 9.955743 | 9.966343 | 9.987669 | 10.04557 | 10.29737 | 10.08119 | 9.776082 |
| 11.13743 | 11.09254 | 11.0819 | 9.404821 | 9.455827 | 10.82373 | 10.86382 | 10.86575 | 11.11269 |
| 4.700551 | NaN | NaN | 5.911002 | 7.221374 | NaN | 6.179216 | 6.408741 | 6.448646 |
| 8.158721 | 7.238213 | 7.760085 | 8.428276 | 7.557767 | NaN | 6.901928 | 6.487449 | 7.014434 |
| 5.431206 | 5.569573 | 5.793741 | NaN | NaN | NaN | NaN | NaN | 5.17518 |
| 8.716725 | 8.472593 | 8.497481 | 6.874994 | 7.019599 | 7.420706 | 7.18335 | 7.402619 | 5.868895 |
| 8.264751 | 8.246298 | 8.351986 | 9.396328 | 9.389824 | 8.495605 | 8.192113 | 8.268903 | 7.775263 |
| 6.374166 | 6.897219 | 5.593819 | 7.374265 | 7.277084 | 6.262374 | 6.440969 | 6.481892 | 5.678412 |
| 9.648818 | 9.683808 | 9.626808 | 10.72305 | 10.61082 | 8.951579 | 9.072145 | 9.078749 | 9.689062 |
| 9.219738 | 7.307582 | 6.9653 | NaN | NaN | 6.953935 | 6.715826 | 7.22276 | 7.975869 |
| 9.644019 | 9.469554 | 9.305237 | 9.367366 | 9.442557 | 10.14576 | 10.06538 | 10.00791 | 9.167767 |
| 15.22305 | 15.18842 | 15.11436 | 16.1315 | 16.38925 | 15.66171 | 15.6409 | 15.72994 | 15.28435 |
| 11.61452 | 11.40162 | 11.66098 | 8.801964 | 8.235121 | 7.125096 | 6.963213 | 7.18225 | 7.534951 |
| 5.500679 | 6.06706 | 5.762096 | 6.116842 | 5.958999 | 6.480886 | 6.158446 | 5.939295 | 6.838239 |
| 8.949759 | 9.130937 | 9.099152 | 9.531934 | 9.166544 | 8.759346 | 8.659621 | 8.50726 | 8.520227 |
| 8.125861 | 8.023334 | 7.893483 | 7.293795 | 7.158878 | 6.395524 | 6.828678 | 7.023362 | 6.82517 |
| 7.27896 | 7.099423 | 7.316056 | 8.448483 | 7.993011 | 6.946374 | 6.946432 | 6.915698 | 6.88222 |
| 6.777731 | 6.866675 | 6.810882 | 6.760297 | 6.554737 | 7.345679 | 7.362735 | 7.440442 | 6.921671 |
| 7.264373 | 7.135932 | 7.600519 | 6.256993 | 6.404333 | 7.284475 | 7.181249 | 7.203651 | 6.543945 |
| 7.734603 | 7.693791 | 7.873037 | 7.978776 | 8.138351 | 6.730863 | 7.104355 | 6.898277 | 6.733015 |
| 8.301661 | 6.812143 | 7.942886 | 7.118463 | 7.158552 | 5.821876 | 6.562473 | 6.383157 | 6.149707 |
| 7.911245 | NaN | 7.811677 | 7.329393 | 7.149665 | 6.494715 | 7.146611 | 6.696582 | 6.862314 |

-Bruker Confidential-

| | | | | | | | | |
|---|---|---|---|---|---|---|---|---|
| 7.137598 | 7.517139 | 7.298004 | 6.752929 | 6.770485 | 7.755882 | 7.579451 | 7.473095 | 7.233466 |
| 8.522711 | 8.06128 | 8.495367 | 8.722379 | 8.86888 | 8.037141 | 8.088954 | 8.159637 | 7.501321 |
| 6.947564 | 7.257104 | 7.072375 | 7.469025 | 7.431491 | 7.068064 | 6.499757 | 6.713974 | 7.078829 |
| 7.336082 | 7.89535 | 7.306853 | 6.434398 | 6.995826 | 7.668351 | 7.602971 | 7.822207 | 7.077165 |
| 5.339301 | 5.713143 | 5.701176 | 9.142886 | 9.016946 | 5.542365 | 5.575501 | 6.404505 | 5.208032 |
| 9.503096 | 9.600407 | 9.719695 | 10.58904 | 10.3741 | 8.179997 | 7.877657 | 8.091728 | 8.387593 |
| 8.107157 | 7.7055 | 7.873778 | 8.065825 | 7.861476 | 8.432539 | 7.990992 | 8.347982 | 7.82843 |
| 6.075169 | 5.79451 | 5.871914 | 5.616992 | 5.460714 | 6.391786 | 6.208008 | 5.967569 | 6.166859 |
| 9.460999 | 8.935797 | 8.936091 | 9.152363 | 9.91614 | 8.375475 | 8.905118 | 7.986942 | 8.178461 |
| 7.219926 | NaN | 6.400735 | 7.494485 | 8.118525 | NaN | 6.528676 | NaN | 5.792007 |
| 8.845291 | 9.357591 | 7.928061 | 7.103406 | 7.412683 | 7.567232 | 7.744581 | 7.4745 | 6.995717 |
| 8.158335 | 7.96816 | 7.832414 | 8.385291 | 8.495217 | 7.211183 | 7.088046 | 7.406091 | 7.695061 |
| 6.076366 | 5.427526 | 6.353804 | 5.656365 | 5.424028 | 6.614597 | 6.409057 | 6.543698 | 6.428052 |
| 7.144684 | 7.597261 | 7.522547 | 7.848219 | 7.250908 | 6.663082 | 6.086925 | 6.099189 | 4.137026 |
| 11.63242 | 11.59985 | 11.37461 | 11.3485 | 11.33386 | 11.30577 | 11.17443 | 11.20668 | 11.14749 |
| 9.000943 | 9.059843 | 8.986525 | 8.31414 | 8.297964 | 9.06543 | 9.029978 | 8.863778 | 8.679218 |
| 10.74704 | 10.62621 | 10.66418 | 10.93556 | 10.70262 | 10.94194 | 11.03712 | 11.00194 | 10.23332 |
| 6.785106 | 6.923792 | 6.624758 | 6.621321 | 5.950195 | 6.700812 | 7.007999 | 7.419143 | 6.537887 |
| 8.726217 | 8.504092 | 8.15123 | 8.376337 | 8.291305 | 8.520289 | 8.367338 | 8.594823 | 8.078987 |
| 10.78375 | 10.5566 | 10.93356 | 12.04516 | 11.79865 | 11.27151 | 11.37675 | 11.43009 | 9.421299 |
| 7.336077 | 6.998182 | 7.405884 | 9.131713 | 8.502109 | 7.743101 | 7.806157 | 7.513049 | 7.549482 |
| 7.618816 | 6.311494 | 6.936654 | 8.382383 | 8.473512 | 5.956015 | 6.024347 | 2.692631 | 5.736392 |
| 7.808856 | 7.759754 | 7.592559 | 7.319374 | 7.549697 | 7.570084 | 7.754191 | 7.684614 | 7.237813 |
| 5.606812 | 6.112551 | 5.94323 | 5.815547 | 6.790249 | 4.664703 | 4.09772 | 4.030605 | 3.647404 |
| 8.229897 | 7.644225 | 8.17077 | 6.974643 | 7.139784 | 8.255211 | 8.314636 | 8.135394 | 8.151449 |
| NaN | NaN | NaN | 7.520334 | 7.106496 | 7.433136 | 7.375528 | 7.307075 | 7.301029 |
| 5.634048 | 5.10107 | NaN | NaN | NaN | NaN | 4.968593 | NaN | 5.604771 |
| 5.922745 | 5.545297 | 5.608204 | NaN | NaN | 4.527688 | 4.279935 | 5.143476 | 2.912859 |
| 5.747642 | 5.61975 | 5.278143 | NaN | NaN | 5.624801 | 5.240019 | 5.725771 | 5.725208 |
| 8.650755 | 8.271595 | 8.708219 | 8.29498 | 8.030787 | 7.591279 | 7.69866 | 7.722608 | 8.88474 |
| NaN | 4.470531 | 4.671282 | 4.83414 | 4.965599 | 6.031203 | 5.490892 | 5.806754 | 5.931556 |
| 7.55187 | 7.641619 | 7.756044 | 5.643565 | 5.247809 | 7.113826 | 6.845929 | 6.864951 | 5.406054 |
| 6.346203 | 6.569836 | 6.093608 | 5.152664 | 6.618438 | 6.381574 | 6.223014 | 6.199259 | 5.034384 |
| 8.320849 | 8.401832 | 8.342864 | 9.135064 | 9.010805 | 7.662597 | 7.694386 | 7.757541 | 8.511928 |
| 8.305326 | 8.004899 | 7.938296 | 7.941367 | 8.667864 | 6.762841 | 7.201359 | 6.657748 | 8.034249 |
| 6.241353 | 5.456574 | 5.851014 | 4.760657 | 5.132704 | 6.325992 | 6.344919 | 6.184029 | 6.301336 |
| 8.643183 | 8.543302 | 8.562336 | 9.938519 | 9.886405 | 8.778193 | 8.582682 | 8.429739 | 7.186755 |
| 7.187946 | 6.976171 | 7.207579 | 7.72772 | 7.829677 | 7.750399 | 8.070938 | 7.261756 | 6.600385 |
| 9.400193 | 9.075871 | 9.037774 | 9.793884 | 9.812565 | 9.366353 | 9.761139 | 9.522703 | 8.832312 |
| 7.52052 | 7.541874 | 7.334137 | 7.539411 | 7.544828 | 7.54302 | 7.408768 | 7.715569 | 6.927949 |
| 12.77461 | 12.8488 | 12.85273 | 10.78186 | 10.63469 | 12.20102 | 12.24972 | 12.35344 | 11.78166 |
| 8.159373 | 8.327289 | 8.267802 | 7.801998 | 7.815856 | 8.289883 | 8.233405 | 8.157202 | 8.697234 |
| 7.350973 | 7.401153 | 7.348268 | 7.843168 | 7.899695 | 7.351583 | 7.293975 | 7.248872 | 7.921082 |
| 7.079795 | 6.538213 | 6.821937 | NaN | 6.099388 | 7.153696 | 7.172415 | 7.154367 | 7.943408 |
| 8.70148 | 8.546111 | 8.637098 | 7.275234 | 7.463885 | 9.230187 | 9.078717 | 9.037484 | 8.689588 |
| 6.999763 | 6.492682 | 7.031078 | 6.089866 | 6.268447 | 7.633221 | 7.763194 | 7.608983 | 7.178092 |
| 6.724301 | 5.960783 | 6.232987 | NaN | NaN | 6.123697 | 6.328718 | 6.27141 | 6.009301 |

-Bruker Confidential-

| | | | | | | | | |
|---|---|---|---|---|---|---|---|---|
| 7.585736 | 7.190468 | 7.623408 | 8.965405 | 8.695682 | 7.723718 | 7.551071 | 7.92792 | 6.926587 |
| 5.188012 | 5.374278 | 5.240431 | 7.782425 | 9.286888 | NaN | 6.339651 | 6.725055 | 5.365857 |
| 12.94852 | 12.88316 | 12.93328 | 11.37981 | 11.35719 | 13.00945 | 12.99219 | 13.05798 | 12.65568 |
| 6.21169 | 6.258435 | 6.016841 | 6.444524 | 6.397162 | 6.93648 | 7.042022 | 6.916357 | 6.858408 |
| 5.984529 | 4.850335 | NaN | NaN | NaN | 5.104291 | 4.493288 | 5.003283 | 5.202924 |
| 8.972646 | 8.814933 | 9.004714 | 10.26331 | 9.977296 | 8.796256 | 8.684134 | 8.867769 | 8.679819 |
| 7.49858 | 7.023381 | 7.695992 | 7.436716 | 6.589032 | 7.382645 | 7.391374 | 7.327858 | 6.070585 |
| 7.953334 | 7.872128 | 7.597349 | 7.179274 | 6.95073 | 8.759375 | 8.530826 | 8.152726 | 8.202172 |
| 12.46493 | 12.27946 | 12.35781 | 11.6854 | 11.24023 | 12.23217 | 12.27596 | 12.04906 | 11.76577 |
| 8.591121 | 8.318678 | 8.055736 | 8.16321 | 8.000682 | 7.391837 | 7.530721 | 7.291423 | 8.414534 |
| 9.877615 | 9.943435 | 9.901961 | 9.735066 | 9.750591 | 10.25198 | 10.24315 | 10.34292 | 9.853194 |
| 8.502722 | 8.508797 | 8.583133 | 8.231302 | 7.864284 | 8.502927 | 8.406979 | 8.183976 | 7.843799 |
| 11.19257 | 11.04017 | 11.20138 | 12.84895 | 12.38303 | 11.74305 | 11.78863 | 11.7915 | 12.24318 |
| 9.501494 | 9.444347 | 9.732666 | 11.7993 | 11.56529 | 11.04094 | 10.80489 | 11.17232 | 10.04449 |
| 7.139694 | 7.68803 | 6.465184 | 7.12764 | 6.583651 | 6.687127 | 6.853338 | 6.734438 | 6.090138 |
| 4.592767 | 4.33425 | 4.691015 | NaN | NaN | 4.849035 | NaN | 4.815694 | 4.857476 |
| 8.646684 | 8.191252 | 8.457364 | 7.916072 | 7.577497 | 8.346186 | 8.015433 | 8.065212 | 8.01692 |
| 6.276311 | 6.479621 | 6.610941 | 6.787574 | 6.240563 | 7.842724 | 7.690648 | 7.518083 | 6.478256 |
| 9.222719 | 8.975352 | 9.19333 | 9.141265 | 9.550742 | 8.437323 | 8.368729 | 8.365962 | 7.673949 |
| 9.246602 | 9.235881 | 9.272877 | 9.891853 | 10.00611 | 9.523084 | 9.443223 | 9.64854 | 8.877266 |
| 10.59273 | 10.43819 | 10.42402 | 11.17495 | 11.01571 | 10.91344 | 10.94535 | 10.93528 | 10.74007 |
| 6.096316 | 6.381341 | 5.62473 | 6.343538 | 7.543097 | NaN | 6.582922 | 6.386323 | 6.521346 |
| 6.581533 | 6.699408 | 5.908993 | NaN | NaN | NaN | NaN | NaN | NaN |
| 9.435419 | 9.442533 | 9.196853 | 8.451194 | 8.494236 | 8.878335 | 8.854571 | 8.765724 | 9.032825 |
| 8.644935 | 8.694603 | 8.601242 | 6.962132 | 7.292207 | 8.639513 | 8.462619 | 8.476071 | 7.786874 |
| 6.967884 | 7.52866 | 7.675143 | 9.622082 | 10.34268 | 7.752126 | NaN | NaN | 8.104001 |
| 5.383637 | 3.615898 | NaN | 6.134717 | 5.537103 | 5.460583 | 5.148267 | 5.684623 | 5.312561 |
| 7.196492 | 7.300825 | 7.118931 | 6.567041 | 6.763659 | 6.842828 | 6.893154 | 6.677691 | 6.527275 |
| 8.628777 | 8.557049 | 8.623157 | 7.21698 | 7.250782 | 8.36772 | 8.717603 | 8.582228 | 8.985381 |
| 7.304337 | 7.224027 | 7.616227 | 5.419611 | 6.264916 | 8.328403 | 8.086989 | 8.15724 | 5.685827 |
| 4.732748 | 4.957625 | 4.361303 | 4.588519 | 5.004349 | 5.220762 | 5.423214 | 5.104373 | 5.224123 |
| 9.740657 | 9.799035 | 9.699823 | 11.36335 | 11.41906 | 10.08619 | 9.929473 | 10.03775 | 9.777634 |
| NaN | 5.828226 | 5.008453 | 5.296787 | 5.552292 | 6.153726 | NaN | 5.921436 | 6.117432 |
| 7.446886 | 7.319433 | 7.567612 | 7.417714 | 6.222663 | 7.024823 | 7.25173 | 7.451639 | 7.148746 |
| 7.161871 | 6.833784 | 6.940404 | 7.094782 | 7.683823 | 6.467727 | 6.940815 | 6.793598 | 7.48936 |
| 8.753908 | 6.682739 | 6.910239 | NaN | NaN | NaN | NaN | NaN | 8.462629 |
| 7.491249 | 6.813636 | 6.947986 | 4.696127 | NaN | 7.21572 | 7.257216 | 7.122571 | 7.070007 |
| 6.606582 | 5.756858 | 5.786176 | 5.680657 | 6.10267 | 5.514885 | 5.519377 | 5.069051 | 6.034374 |
| 4.90444 | NaN | 6.627407 | NaN | 5.396388 | 5.150973 | 5.342638 | 5.923714 | 4.2344 |
| 7.851348 | 7.706361 | 7.804322 | 9.129276 | 9.070567 | 7.869412 | 7.551794 | 7.849343 | 7.109045 |
| 6.667765 | 6.631017 | 6.486754 | 5.581763 | 5.497687 | 5.778212 | 5.869806 | 5.803084 | 5.692118 |
| 6.325013 | 6.281007 | 6.248279 | 6.618822 | 6.087708 | 7.803765 | 7.737814 | 7.696671 | 5.104209 |
| 9.904849 | 9.667414 | 9.664648 | 8.169471 | 8.043766 | 9.59293 | 9.755633 | 9.762959 | 9.757509 |
| 6.119036 | 5.922286 | 7.074121 | 4.858104 | 4.909565 | 6.374176 | 6.561102 | 6.716257 | 5.757778 |
| 6.910885 | 6.311872 | 5.976014 | 6.374949 | 6.579475 | 6.603698 | 6.42131 | 6.24264 | 7.005843 |
| 9.024834 | 8.598562 | 8.979023 | 8.954561 | 8.915572 | 8.940954 | 8.811218 | 8.823951 | 8.617991 |
| 6.798577 | 6.159912 | 6.407757 | 5.646751 | 6.264981 | 6.177411 | 6.484936 | 6.73046 | 5.901634 |

| | | | | | | | | |
|---|---|---|---|---|---|---|---|---|
| 9.9192 | 9.835467 | 9.766414 | 7.581425 | 7.532767 | 6.886098 | 6.347438 | 6.373604 | 6.58286 |
| 1.637527 | 5.495268 | 3.85248 | 5.919187 | 6.741642 | 6.474419 | 6.500732 | 7.657572 | 8.133278 |
| 9.62161 | 9.666775 | 9.633431 | 7.737923 | 7.607957 | 9.439272 | 9.411999 | 9.41042 | 9.850049 |
| 6.721063 | 6.945169 | 6.810587 | 6.998351 | 5.379111 | 6.61478 | 6.775047 | 6.650914 | 6.087648 |
| 7.414189 | 7.154798 | 7.342523 | 6.469144 | 6.121248 | 7.17705 | 7.413696 | 7.29974 | 6.825498 |
| 10.20452 | 10.17103 | 10.10849 | 10.31922 | 10.18627 | 10.18102 | 10.18351 | 10.1376 | 10.28492 |
| 7.232014 | 7.193708 | 7.085244 | 5.951026 | 5.98277 | 7.488892 | 7.136795 | 7.247338 | 7.715127 |
| 10.84785 | 10.80419 | 10.81955 | 11.26831 | 11.43482 | 9.593996 | 9.514781 | 9.556787 | 10.24281 |
| 10.89043 | 10.85242 | 10.67609 | 11.23894 | 11.25152 | 10.35083 | 10.60533 | 10.48462 | 10.26019 |
| 8.495665 | 8.532723 | 8.503459 | 7.827596 | 7.979067 | 8.772338 | 8.703003 | 8.713809 | 8.761765 |
| 9.261902 | 9.189015 | 9.205606 | 11.01515 | 10.87873 | 8.831623 | 8.76278 | 8.67908 | 8.593163 |
| 10.39898 | 10.36914 | 10.4781 | 10.46998 | 10.18496 | 10.53191 | 10.54393 | 10.29185 | 9.561963 |
| 6.832182 | 6.860071 | 7.013844 | 6.591006 | 6.810095 | 7.985529 | 7.947584 | 8.017451 | 7.037311 |
| NaN | NaN | NaN | NaN | 6.584622 | NaN | NaN | NaN | 7.185424 |
| 8.011993 | 8.463708 | 8.738203 | 7.430993 | 6.860046 | 7.181784 | 6.998398 | 7.554185 | 6.441787 |
| 7.203469 | 6.781272 | 6.93603 | 5.633134 | 5.942747 | 7.419442 | 7.281584 | 7.350617 | 6.732008 |
| 12.93089 | 13.16496 | 13.03057 | 7.826527 | 8.096782 | 9.215029 | 9.745901 | 7.15143 | 8.847366 |
| 6.738097 | 6.58854 | 6.470227 | NaN | 6.701249 | 6.311474 | 6.244598 | 6.577943 | 5.649264 |
| 12.51685 | 12.53034 | 12.57994 | 12.79416 | 12.59328 | 11.46807 | 11.59497 | 11.45516 | 12.06122 |
| 7.239457 | 7.265848 | 6.840168 | 8.796312 | 9.101361 | 7.815847 | 7.591415 | 7.773627 | 6.807323 |
| 6.948451 | 6.758492 | 6.683402 | 6.265454 | 5.333412 | 7.06319 | 8.495326 | 8.463424 | 6.38375 |
| 7.464527 | 7.781212 | 7.099934 | 6.653621 | 7.106465 | 7.694997 | 7.389752 | 7.808024 | 7.073487 |
| 5.984707 | 5.902799 | 6.25239 | 5.781189 | 5.306777 | 4.984605 | 5.234224 | NaN | 9.102205 |
| 7.879535 | 7.603713 | 7.643151 | 6.952634 | 7.390522 | 7.043437 | 7.183733 | 7.794934 | 6.764996 |
| 9.317618 | 9.525449 | 9.452161 | 10.17628 | 10.10545 | 9.043859 | 9.103675 | 9.011633 | 9.117461 |
| 6.653943 | 6.768877 | 6.949574 | 6.005485 | 6.482269 | 7.147932 | 6.998904 | 7.069883 | 7.480746 |
| 8.705874 | 8.737107 | 8.751184 | 8.107205 | 8.043958 | 8.687165 | 8.552973 | 8.628339 | 8.828793 |
| 8.101098 | 8.06457 | 8.154039 | 5.298214 | 5.82307 | 6.390226 | 6.622238 | 6.51157 | 4.659754 |
| 5.769827 | 4.404615 | 4.878389 | 5.30283 | 5.621898 | 5.385138 | NaN | 5.331593 | 4.155439 |
| 7.828968 | 7.585828 | 7.934464 | 7.028182 | 7.056021 | 8.327288 | 8.2092 | 8.101862 | 7.742442 |
| 6.404088 | 5.881272 | 5.368351 | 7.770102 | 7.617914 | 6.440839 | 6.332047 | 6.442088 | 6.758254 |
| 10.14361 | 10.2641 | 10.26929 | 10.77419 | 10.80289 | 10.4909 | 10.11174 | 10.09478 | 10.17664 |
| 12.17013 | 12.29239 | 12.1134 | 12.57422 | 12.47271 | 11.115 | 10.99404 | 10.96947 | 11.33446 |
| 12.17781 | 12.12028 | 12.15978 | 12.35197 | 12.14463 | 12.1606 | 12.07393 | 12.12773 | 11.64407 |
| 7.461723 | 6.89877 | 6.916137 | 6.581913 | NaN | 7.659695 | 7.496138 | 7.457218 | 7.204241 |
| 8.198224 | 7.958886 | 7.855031 | 6.573094 | 7.061317 | 7.969033 | 7.689469 | 7.829703 | 7.784492 |
| 9.309361 | 9.175666 | 9.32284 | 8.713394 | 8.53186 | 8.997474 | 9.070851 | 9.038192 | 7.622125 |
| 6.306264 | 6.097373 | 6.000859 | 6.684437 | 6.523347 | 5.392073 | 5.481852 | 6.473616 | 5.405751 |
| 5.89403 | 5.81991 | 6.186825 | NaN | 4.69947 | 6.930018 | 7.084051 | 7.388493 | NaN |
| 6.899045 | 7.048867 | 7.124888 | 9.191092 | 9.155749 | 7.007259 | 7.142363 | 7.399825 | 7.622082 |
| 8.866686 | 8.841739 | 8.502042 | NaN | 7.733019 | 10.12919 | 9.840333 | 9.992741 | 8.870039 |
| 7.544472 | 7.483304 | 7.459658 | 8.380557 | 8.74806 | 7.897456 | 7.739661 | 8.307575 | 8.216536 |
| 9.701328 | 9.776253 | 9.643767 | 9.245899 | 9.098036 | 9.65755 | 9.703511 | 9.788905 | 9.624417 |
| 8.94196 | 9.118983 | 8.942796 | 9.454324 | 9.220937 | 6.642109 | 6.74499 | 6.329268 | 7.371642 |
| 7.150795 | 6.152095 | 6.927292 | 7.039276 | 7.941032 | 6.490764 | 6.261065 | 6.544664 | 7.014327 |
| 8.545459 | 8.457221 | 8.770983 | NaN | 6.28055 | 8.156549 | 7.209056 | 7.943454 | 6.577228 |
| 7.955264 | 8.086425 | 8.090537 | 7.233391 | 7.187157 | 8.639071 | 8.684856 | 8.555185 | 7.442152 |

| | | | | | | | | |
|---|---|---|---|---|---|---|---|---|
| 7.674759 | 7.167688 | 6.791944 | 6.991298 | 7.119832 | 6.341628 | 7.376143 | 7.242831 | 7.665414 |
| NaN | NaN | NaN | NaN | NaN | NaN | NaN | NaN | 4.780765 |
| 7.221242 | 7.240286 | 7.174793 | 6.726757 | 6.548097 | 7.649308 | 7.346457 | 7.554599 | 6.755465 |
| 8.653189 | 8.508108 | 8.408088 | 8.117585 | 8.084339 | 8.423735 | 8.417307 | 8.647137 | 8.686555 |
| 7.916839 | 7.355651 | 7.856313 | 6.833671 | 6.862356 | 7.452258 | 7.290097 | 7.090417 | 7.059347 |
| 7.923703 | 7.629256 | 7.763915 | NaN | NaN | NaN | 6.040079 | NaN | NaN |
| 9.388247 | 9.254342 | 9.181928 | 10.70207 | 10.21351 | 9.682911 | 9.744694 | 9.630499 | 9.61947 |
| 8.47295 | 7.906975 | 8.031585 | 7.049991 | 7.083571 | 7.588549 | 7.705688 | 7.936516 | 7.846194 |
| NaN | NaN | 5.838277 | 6.632171 | 7.025122 | NaN | 5.448515 | 5.8366 | 6.821381 |
| 8.776011 | 8.649158 | 8.335519 | 8.402548 | 8.585979 | 8.576278 | 8.43408 | 8.305927 | 7.609107 |
| 6.879173 | 7.04473 | 6.971911 | 5.82262 | 5.755049 | 6.171047 | 6.034266 | 6.096161 | 6.420905 |
| 5.695262 | 5.530671 | 5.581262 | 4.556451 | 6.292694 | 6.435846 | 5.866399 | 6.049978 | 4.680635 |
| 6.818147 | 6.599314 | 6.43387 | 4.730797 | 6.991116 | 6.526741 | 6.424675 | 6.157852 | 6.228604 |
| 7.919102 | 8.128757 | 8.159127 | 7.155735 | 7.305572 | 7.601384 | 7.761859 | 7.600126 | 7.735257 |
| 10.81117 | 10.61446 | 10.78409 | 11.70292 | 11.80076 | 11.14567 | 11.20332 | 11.26705 | 11.18238 |
| 9.152307 | 8.934863 | 9.233947 | 8.745175 | 8.11244 | 8.71113 | 8.540086 | 8.93146 | 8.865098 |
| 11.47225 | 11.78689 | 11.40614 | 11.97402 | 12.18764 | 11.07941 | 10.97263 | 11.13498 | 10.72205 |
| 11.92709 | 11.74325 | 11.74313 | 11.33957 | 11.43377 | 11.78751 | 11.73757 | 11.74835 | 12.01271 |
| 7.98291 | 8.099914 | 8.137681 | 9.106663 | 9.073224 | 8.208271 | 8.043251 | 8.151782 | 8.839848 |
| 6.850867 | 6.425678 | 6.991753 | 4.993194 | 5.788966 | 5.805733 | 5.718378 | 5.871553 | 5.978004 |
| 9.273058 | 9.068094 | 9.537914 | 7.724442 | 8.426031 | 7.589227 | 8.514908 | NaN | 7.815324 |
| 8.91633 | 8.842875 | 8.958002 | 9.087904 | 8.77128 | 8.862718 | 8.9675 | 8.869707 | 8.627224 |
| 8.960847 | 9.093544 | 9.025449 | 7.875355 | 7.487931 | 8.603142 | 8.756249 | 8.809535 | 9.236096 |
| 9.022306 | 8.958536 | 9.042192 | 8.490333 | 8.596575 | 9.072623 | 9.071773 | 9.060636 | 9.217378 |
| 7.58972 | 6.498676 | 7.067621 | NaN | 5.836851 | 8.323911 | 7.902486 | 8.032827 | 6.933185 |
| 8.396637 | 8.319515 | 8.373178 | 6.991144 | 7.000612 | 8.387857 | 8.250858 | 8.406759 | 7.877465 |
| 10.63592 | 10.64124 | 10.5042 | 11.88818 | 11.77841 | 11.59433 | 11.44496 | 11.38896 | 10.91382 |
| 9.065835 | 8.84019 | 9.052271 | 9.696113 | 9.725497 | 9.368319 | 9.314916 | 9.490463 | 8.82709 |
| NaN | 5.21056 | 5.730532 | 7.27965 | 7.980987 | 4.101701 | 6.237173 | 6.016449 | 6.62587 |
| 8.744285 | 8.599344 | 8.741343 | 8.791679 | 8.841635 | 9.166982 | 9.116895 | 9.204403 | 8.725865 |
| 8.05654 | 7.605918 | 8.076555 | 6.974223 | 6.824784 | 8.236567 | 8.169705 | 8.19789 | 7.028667 |
| 8.685694 | 8.703679 | 8.641518 | 7.329865 | 7.133487 | 6.665899 | 6.009933 | 5.913507 | 5.734915 |
| 7.629035 | 7.793318 | 7.894661 | 7.633113 | 7.523752 | 7.985108 | 7.742651 | 7.875073 | 7.143931 |
| 11.43884 | 11.3252 | 11.30524 | 9.867342 | 9.950508 | 11.04801 | 11.16233 | 11.1232 | 11.10963 |
| 9.315942 | 9.489764 | 9.202265 | 10.76898 | 10.55895 | 9.715558 | 9.673233 | 9.734397 | 7.979723 |
| 7.858255 | 7.760087 | 7.701784 | 5.880197 | 5.136943 | 7.157139 | 7.462785 | 7.531639 | 7.336888 |
| 6.191705 | 6.689138 | 6.530018 | 7.256111 | 7.431506 | 6.769373 | 5.68642 | 6.546525 | 6.683703 |
| 7.164835 | 7.15556 | 7.133317 | 7.029812 | 7.134637 | 7.317448 | 7.241453 | 7.164313 | 7.323869 |
| 8.220405 | 8.274697 | 8.070329 | 7.796649 | 7.771305 | 7.9392 | 7.854071 | 8.094624 | 8.350749 |
| 7.977706 | 7.9851 | 8.316283 | 9.051533 | 9.110159 | 8.64578 | 8.840884 | 8.57496 | 8.853482 |
| 8.349173 | 8.282469 | 8.362267 | 7.291094 | 7.155252 | 7.966287 | 8.110459 | 8.240138 | 8.203648 |
| 5.999156 | 5.988456 | NaN | NaN | NaN | 7.16501 | 7.878887 | 7.570543 | 5.56801 |
| 7.615769 | 7.75998 | 7.856676 | 8.785171 | 8.814487 | 7.280655 | 7.20157 | 6.906583 | 7.077707 |
| 10.04575 | 10.07572 | 10.00333 | 7.935352 | 8.114344 | 6.632577 | 6.270789 | 6.345335 | 6.810373 |
| 6.609829 | 6.899218 | 6.98752 | 9.692072 | 7.318653 | 5.314663 | 6.300969 | 5.836312 | 6.224836 |
| 8.915026 | 7.49893 | 7.874179 | 8.311286 | 7.624931 | 6.638155 | 8.164189 | 6.976333 | 6.614883 |
| 8.464137 | 7.762571 | 8.648303 | 7.066527 | 6.20969 | 7.913955 | 8.60218 | 8.651679 | 8.673795 |

| | | | | | | | | |
|---|---|---|---|---|---|---|---|---|
| 6.99284 | 7.0789 | 7.062874 | 7.179471 | 7.10672 | 6.979279 | 6.611382 | 6.864307 | 7.607553 |
| 9.449729 | 9.399446 | 9.291797 | 8.76214 | 8.620549 | 9.437828 | 9.19545 | 9.395013 | 9.435418 |
| 3.508701 | 4.10763 | 3.894721 | 2.879285 | 3.469096 | 4.858301 | 4.596792 | 4.070547 | NaN |
| 7.973495 | 7.962691 | 7.977838 | 8.172719 | 8.180294 | 7.60867 | 7.777718 | 7.720242 | 7.835337 |
| 6.945414 | 7.054774 | 7.1748 | 5.933136 | 6.134996 | 6.911557 | 7.531223 | 6.795435 | 5.562298 |
| 8.287657 | 7.880088 | 8.272865 | 9.096503 | 7.80766 | 7.564643 | 8.387788 | 7.121739 | 7.601381 |
| 8.946559 | 8.846016 | 8.877938 | 7.29665 | 7.34422 | 9.23674 | 9.322382 | 8.976193 | 8.432648 |
| 8.039637 | 8.111011 | 7.971011 | 7.9224 | 7.620745 | 7.824746 | 7.800346 | 8.200756 | 7.749044 |
| NaN | 7.612725 | 7.348401 | NaN | NaN | 8.490417 | 8.53672 | 8.252015 | 7.718151 |
| 9.240295 | 9.160727 | 8.967082 | 8.190894 | 7.79449 | 8.908192 | 8.552057 | 8.80063 | 8.866655 |
| 8.97427 | 8.713223 | 8.865144 | 6.706346 | 6.631342 | 8.87132 | 8.781134 | 8.717263 | 8.787388 |
| 5.914178 | 5.844175 | 5.893301 | 5.818186 | 5.668361 | 7.006281 | 5.866183 | 5.794301 | 5.381827 |
| 6.771688 | 6.733191 | 6.907205 | 7.795311 | 7.856324 | 6.647568 | 5.968289 | 6.600796 | 6.795164 |
| 7.12451 | 7.576888 | 7.382017 | 6.4329 | 6.266714 | 6.683158 | 5.650349 | 6.35463 | 6.745278 |
| 7.409603 | 6.42837 | 7.224175 | 5.930883 | 5.911321 | 7.245297 | 7.435126 | 7.380332 | 6.884077 |
| 5.61783 | 3.689295 | NaN | 5.573874 | 6.083873 | 6.106449 | 6.4807 | 6.470429 | 5.420991 |
| 6.901549 | 6.796126 | 6.70622 | 5.900514 | 6.316946 | 6.002453 | 6.135319 | 6.25417 | 6.676895 |
| 9.033226 | 8.670753 | 9.108986 | 8.988483 | 9.236465 | 9.162891 | 8.993363 | 9.021699 | 9.303987 |
| 8.078074 | 8.394423 | 8.231784 | 6.998991 | 6.97831 | 8.218219 | 8.132335 | 8.422247 | 8.320894 |
| 8.580317 | 8.450995 | 8.569656 | 8.122491 | 8.087191 | 8.144107 | 8.361528 | 8.364119 | 8.586535 |
| 8.335073 | 8.050801 | 8.242794 | 7.797147 | 7.673185 | 8.082425 | 7.857227 | 8.151798 | 8.011601 |
| 10.17958 | 10.19733 | 10.15652 | 9.332802 | 9.172641 | 10.24365 | 10.24999 | 10.19677 | 9.742364 |
| 7.169987 | 7.356731 | 7.38442 | 6.088369 | 5.968038 | 7.615184 | 7.534341 | 7.771972 | 7.360673 |
| 9.139917 | 8.933747 | 9.085564 | 8.062245 | 7.301324 | 8.448306 | 8.449088 | 8.495928 | 8.714086 |
| 6.792699 | 6.71023 | 6.588736 | 6.637679 | 6.282858 | 6.236933 | 5.898192 | 5.744998 | 6.132625 |
| 8.2019 | 7.512051 | 7.822334 | 5.785454 | NaN | 7.516965 | 7.566041 | 7.830717 | 7.740198 |
| 7.719101 | 7.877142 | 7.767433 | 7.639145 | 8.00272 | 8.051782 | 8.116222 | 8.057538 | 7.89037 |
| 7.580935 | 7.699417 | 7.630715 | 7.678363 | 7.676682 | 7.664754 | 7.552095 | 7.794237 | 7.454038 |
| 7.418218 | 7.32706 | 7.384759 | 6.523493 | 6.599478 | 7.35453 | 7.248076 | 7.553791 | 7.063252 |
| 8.74144 | 8.777171 | 8.631558 | 7.917866 | 8.009205 | 8.138263 | 8.363936 | 8.4655 | 8.397499 |
| 5.057189 | 5.248895 | 5.031244 | 4.294071 | 4.471779 | 5.202382 | 4.509028 | 5.515097 | 6.528759 |
| 9.401783 | 9.167747 | 9.299815 | 8.83487 | 8.643268 | 8.175724 | 8.198143 | 8.379275 | 8.269739 |
| 7.769995 | 8.09576 | 7.613538 | 6.772136 | 6.724764 | 6.856888 | 6.860748 | 6.961149 | 7.986218 |
| 7.191282 | 7.288506 | 7.33732 | 6.60632 | 6.614673 | 7.414004 | 7.526966 | 7.519416 | 6.757285 |
| 8.589113 | 8.14344 | 8.621671 | 7.925013 | 7.953825 | 7.027813 | 6.622739 | 7.188876 | 6.906279 |
| 6.530493 | 7.125147 | 7.532852 | 6.589002 | 6.737403 | 7.124935 | 7.149722 | 7.357637 | 7.140056 |
| 7.356467 | 7.12063 | 7.139737 | 7.519748 | 7.478446 | 6.25186 | 6.898745 | 6.183706 | 7.165598 |
| NaN | NaN | 6.226362 | NaN | 6.011245 | 8.283798 | 8.360955 | 8.289076 | 5.814188 |
| 11.1556 | 11.17169 | 11.22384 | 9.818815 | 9.693435 | 11.73853 | 11.6468 | 11.60008 | 9.065163 |
| 6.476606 | NaN | NaN | 4.930627 | 5.705353 | NaN | NaN | 6.453511 | 6.968101 |
| 8.780269 | 8.729886 | 8.73649 | 8.12988 | 7.981988 | 8.488996 | 9.062379 | 8.944154 | 8.371525 |
| 7.032051 | 7.560875 | 7.529479 | 7.230399 | 7.056745 | 8.16429 | 7.955886 | 8.233657 | 7.915482 |
| 10.54277 | 10.58806 | 10.71464 | 10.41496 | 10.28049 | 9.903031 | 10.05101 | 10.14832 | 10.14666 |
| 8.319621 | 8.243553 | 8.17115 | 8.792295 | 8.952827 | 8.281187 | 8.432706 | 8.327996 | 8.637547 |
| 10.80589 | 10.75961 | 10.66049 | 10.20332 | 10.2951 | 10.39392 | 10.48968 | 10.67984 | 10.95565 |
| 10.33665 | 10.22785 | 10.32603 | 10.426 | 10.2939 | 10.06626 | 9.951578 | 9.942797 | 10.64029 |
| 9.808746 | 9.778496 | 9.730783 | 8.365698 | 7.980034 | 9.484541 | 9.454319 | 9.370896 | 9.896531 |

| | | | | | | | | |
|---|---|---|---|---|---|---|---|---|
| 8.059276 | 7.73511 | 7.862606 | 6.504858 | 6.627212 | 8.046185 | 7.888593 | 7.881896 | 7.144295 |
| 8.098166 | 8.468825 | 8.180853 | 9.468936 | 9.631089 | 8.736369 | 8.732761 | 8.67276 | 7.628967 |
| 6.423835 | 6.2554 | 6.380335 | 6.755252 | 6.647095 | 6.223149 | 6.193655 | 6.317369 | 6.511733 |
| 9.279213 | 9.159105 | 9.178991 | 8.424291 | 8.362705 | 8.788306 | 8.664233 | 8.679157 | 8.880049 |
| 6.886289 | 6.85328 | 6.780213 | 6.174905 | 6.307563 | 8.39634 | 8.496097 | 8.566658 | 7.72366 |
| 7.727198 | 7.814083 | 7.766302 | 8.112829 | 7.987908 | 7.826878 | 7.795577 | 7.748685 | 8.250556 |
| 6.952718 | 7.076632 | 7.004291 | 6.831142 | 7.431712 | 6.700981 | 6.988743 | 6.752517 | 6.803385 |
| 4.912647 | 5.082641 | 4.858249 | 6.322968 | 6.77337 | 5.190127 | 4.41539 | 4.914663 | 3.11815 |
| 6.311548 | 5.918313 | 6.091263 | NaN | 6.296695 | 7.473856 | 6.192666 | NaN | 6.728079 |
| 8.035668 | 7.864317 | 8.139562 | 8.344412 | 8.363818 | 7.408721 | 7.550352 | 7.707743 | 7.390448 |
| 6.531441 | 6.195736 | 6.575828 | 6.130984 | 5.819364 | 6.550653 | 6.705414 | 6.538654 | 6.12988 |
| NaN | 1.793521 | 5.542289 | 6.460511 | 6.20326 | NaN | 5.404293 | 5.567911 | 4.60244 |
| 8.404461 | 9.119915 | 8.690998 | 10.60972 | 11.05894 | 9.666368 | 11.3089 | 10.53675 | 10.51204 |
| 7.393408 | 7.410705 | 7.41824 | 7.018679 | 7.226984 | 7.823271 | 7.269239 | 7.758276 | 7.355274 |
| 9.786044 | 9.651977 | 9.492297 | 9.842324 | 9.735625 | 9.66003 | 9.59911 | 9.572865 | 8.409184 |
| 8.769733 | 8.820881 | 8.98285 | 8.435696 | 8.320638 | 6.203181 | 7.533113 | 7.423935 | 9.335952 |
| 6.942147 | 7.113137 | 7.330721 | 6.213857 | 6.351661 | 7.18727 | 7.18028 | 7.373169 | 7.147867 |
| 8.547675 | 8.752971 | 8.258345 | 9.69761 | 9.317565 | 8.72891 | 8.716828 | 8.678062 | 8.800524 |
| 8.036144 | 7.100813 | 6.895689 | NaN | 6.456869 | 7.091325 | 7.154591 | 7.360362 | 7.528897 |
| 7.387068 | 7.525172 | 7.875783 | 7.362939 | 7.631554 | 7.237839 | 7.453933 | 7.388088 | 7.781119 |
| 8.127935 | 8.217476 | 8.257224 | 6.144542 | 6.296231 | 8.373635 | 8.444388 | 8.18714 | 8.038109 |
| 5.812016 | 4.914811 | 6.40009 | 5.266912 | NaN | 6.459801 | 6.173481 | 6.143992 | 6.08174 |
| 8.243712 | 8.483396 | 8.499932 | 8.624923 | 8.204418 | 8.858946 | 8.769314 | 8.726974 | 8.508972 |
| 6.731012 | 7.200288 | 7.206744 | 6.576187 | 6.902177 | 7.258256 | 7.442183 | 7.487069 | 7.432641 |
| 10.42574 | 10.30462 | 10.43227 | 10.00615 | 9.889873 | 10.60312 | 10.50025 | 10.34433 | 10.28558 |
| 8.138916 | 8.259974 | 8.091353 | 7.056047 | 6.758339 | 8.517941 | 8.329085 | 8.427833 | 7.99954 |
| 5.025821 | 4.986115 | 5.540761 | NaN | 4.930234 | 7.272052 | 5.646085 | 6.269465 | 5.729048 |
| 4.336042 | 4.947864 | 4.848709 | NaN | NaN | NaN | NaN | 3.582421 | 4.838743 |
| 5.380882 | 5.288568 | 5.113921 | NaN | 5.17493 | 5.363397 | 5.241001 | 5.591294 | 4.792766 |
| 8.089819 | 8.077796 | 8.148443 | 7.022187 | 6.883767 | 8.043607 | 7.938872 | 7.64435 | 7.476821 |
| 7.700261 | 7.528976 | 7.871208 | 8.20344 | 7.728424 | 7.163257 | 7.369819 | 7.031488 | 7.744381 |
| 7.161113 | 6.700056 | 6.786777 | 6.96672 | 7.166199 | 7.736909 | 7.482274 | 7.470799 | 7.819036 |
| 7.518634 | 7.176635 | 7.706475 | 6.296583 | 6.683435 | 7.660996 | 7.36058 | 7.656999 | 7.297303 |
| 6.405137 | 6.273847 | 5.823396 | 5.513712 | 5.275043 | NaN | 5.913271 | 5.747827 | 5.745504 |
| 6.306804 | 6.552015 | 5.706625 | 5.148911 | 5.904101 | 5.92004 | 5.78228 | 5.650349 | 6.453663 |
| 10.2057 | 10.22548 | 10.38798 | 10.08173 | 10.04681 | 10.19967 | 10.23553 | 10.15128 | 10.37765 |
| 11.86192 | 11.80762 | 12.00676 | 11.70509 | 11.89131 | 11.55255 | 11.09575 | 11.42734 | 11.6022 |
| 8.45285 | 8.494823 | 8.579358 | 8.162449 | 8.408955 | 8.243754 | 8.081691 | 8.184631 | 7.757202 |
| 8.413309 | 8.332572 | 8.368989 | 6.849897 | 7.451293 | 8.961593 | 8.924457 | 8.71711 | 9.157508 |
| NaN | NaN | NaN | 5.386142 | 4.958265 | NaN | NaN | NaN | 5.254172 |
| 9.882541 | 9.715821 | 9.872072 | 9.519547 | 9.461619 | 9.244253 | 9.553337 | 9.270033 | 9.777142 |
| 6.848715 | 7.112394 | 7.002269 | 8.73812 | NaN | 7.243066 | 7.149249 | 7.391418 | 7.259088 |
| 8.547952 | 8.369302 | 8.286185 | 8.608274 | 8.666127 | 8.212453 | 8.148618 | 8.110317 | 8.268437 |
| 9.0889 | 9.186602 | 9.119469 | 9.355056 | 9.66995 | 8.453053 | 8.613761 | 8.695717 | 8.712912 |
| NaN | 5.399983 | 6.084218 | 5.967773 | 7.11332 | 6.705978 | 6.28184 | 6.558677 | 6.267426 |
| 6.919077 | 6.786037 | 6.75389 | 7.100573 | 6.72789 | 8.117919 | 7.647925 | 7.182203 | 6.570709 |
| 5.614042 | NaN | 6.939428 | 6.179939 | 7.092829 | 6.434594 | NaN | 7.344838 | 6.661456 |

| | | | | | | | | |
|---|---|---|---|---|---|---|---|---|
| 9.544442 | 9.439195 | 9.475637 | 8.42569 | 8.303339 | 9.802743 | 9.920905 | 9.761597 | 9.559933 |
| 7.792094 | 6.863258 | 7.525968 | 4.936546 | 5.489496 | 6.734164 | 6.32944 | 6.629435 | 8.395868 |
| 5.589718 | 5.545438 | 5.383889 | 4.984451 | 4.662486 | 5.070072 | 5.225859 | 5.82158 | 5.825321 |
| 7.865222 | 7.676806 | 7.575314 | 7.122034 | 7.117779 | 7.891172 | 7.654984 | 7.806665 | 7.651329 |
| 9.302464 | 9.066719 | 9.217607 | 8.923725 | 8.677782 | 8.18454 | 8.444446 | 8.093743 | 7.518848 |
| 8.201115 | 8.358712 | 8.336907 | 7.760224 | 7.856511 | 8.981059 | 8.63003 | 8.824417 | 8.373635 |
| 10.13021 | 10.0966 | 10.23397 | 10.96651 | 10.74195 | 10.41659 | 10.09611 | 10.2863 | 10.03309 |
| 8.336381 | 8.001251 | 8.028783 | 9.248626 | 8.962234 | 8.577172 | 8.437293 | 8.620353 | 8.101555 |
| 9.866279 | 9.764847 | 9.618331 | 10.63928 | 10.06602 | 9.42719 | 9.511558 | 9.625785 | 9.429471 |
| 7.670528 | 7.260947 | 8.206865 | 7.864388 | 7.373408 | 6.843934 | 6.722079 | 7.05919 | 7.194533 |
| 11.19189 | 11.21623 | 11.16412 | 11.17478 | 11.10475 | 10.60664 | 10.66059 | 10.56628 | 11.02219 |
| 8.223948 | 8.306308 | 8.361962 | 9.124412 | 9.264064 | 7.82544 | 7.524271 | 7.784168 | 8.878016 |
| 5.707037 | 5.31283 | 6.208042 | 4.205602 | NaN | 5.958408 | 6.616218 | 6.523495 | 5.086862 |
| 7.338015 | 7.3995 | 7.853464 | 6.313953 | 6.134502 | 7.446563 | 7.143986 | 7.130458 | 7.281203 |
| 8.333109 | 8.146138 | 8.13063 | 8.14989 | 8.27269 | 8.25525 | 8.059303 | 7.883964 | 7.834361 |
| 6.377883 | 6.539043 | 6.559419 | 6.518073 | NaN | 6.245299 | 6.548658 | 6.683102 | 5.906207 |
| 8.310931 | 6.166109 | 6.719568 | NaN | 5.197547 | 6.255202 | 7.141832 | 6.566942 | 6.788967 |
| 7.138261 | 6.65809 | 6.693083 | 6.049363 | 5.577469 | 7.07582 | 7.559855 | 7.6121 | 6.631679 |
| 8.145356 | 8.323692 | 8.14563 | 6.987679 | 7.158641 | 8.526095 | 8.44593 | 8.244985 | 8.162074 |
| 8.664397 | 8.685223 | 8.538159 | 8.317945 | 8.604689 | 8.301247 | 8.430205 | 8.253773 | 7.412591 |
| 8.631116 | 8.71408 | 8.554996 | 8.548017 | 8.660768 | 8.628873 | 8.761777 | 8.768783 | 9.155816 |
| 5.256544 | 5.332653 | 5.823683 | 4.560971 | 5.312782 | 4.950185 | 5.149579 | 5.362453 | 5.904402 |
| 8.511744 | 8.793639 | 8.585424 | 6.599051 | 6.849751 | 7.788221 | 7.638069 | 7.73836 | 7.824281 |
| 9.048369 | 9.036094 | 9.048371 | 8.079622 | 7.702352 | 9.123118 | 9.164322 | 9.080927 | 8.94337 |
| 8.140386 | 8.217318 | 8.198119 | 7.477688 | 7.490188 | 8.321054 | 8.284982 | 8.294385 | 7.409971 |
| 6.769238 | 5.778647 | 5.959838 | 6.233452 | 5.905159 | 6.278318 | 6.628092 | 6.672789 | 5.265904 |
| 7.214133 | 3.346795 | 6.729297 | 7.600322 | 7.613481 | 6.825468 | 6.873361 | 6.982881 | 8.461973 |
| 9.561256 | 9.244548 | 9.560457 | 10.60471 | 10.23778 | 8.885065 | 8.976013 | 8.792632 | 8.76775 |
| 8.269742 | 8.087606 | 8.003181 | 9.438903 | 9.282478 | 7.9697 | 7.889059 | 7.851827 | 8.476593 |
| 10.22238 | 10.31798 | 10.20886 | 10.4771 | 10.43747 | 10.2737 | 10.2632 | 10.31497 | 10.08468 |
| 5.923004 | 4.051219 | 5.539116 | NaN | 4.908543 | 6.339188 | 6.264737 | 5.804741 | 5.952166 |
| 8.145649 | 8.071699 | 8.174162 | 7.059068 | 7.021812 | 7.62401 | 7.74715 | 7.648698 | 7.913848 |
| 9.761474 | 9.808199 | 9.74438 | 8.816814 | 8.889882 | 10.0489 | 10.2315 | 10.21418 | 9.2962 |
| NaN | NaN | NaN | 5.768259 | 6.078677 | 5.385485 | 5.792326 | 6.214058 | 4.367858 |
| 7.558298 | 8.442883 | 8.799333 | 7.938646 | 7.653107 | 7.386099 | 7.120557 | 6.45243 | 6.369714 |
| 6.520676 | 6.243972 | 6.628136 | 5.279767 | 5.992321 | 5.280772 | 5.185621 | 5.177807 | 6.172387 |
| 6.870425 | 6.900331 | 6.856693 | 6.946429 | 6.755685 | 6.466736 | 6.52575 | 6.902087 | 5.823132 |
| 8.307952 | 7.405645 | 8.191439 | 8.404466 | 7.894611 | 7.267281 | 6.743591 | 6.86548 | 6.836938 |
| 7.461365 | 6.897894 | 7.089577 | 7.466653 | 7.200661 | 6.562265 | 6.665871 | 6.583177 | 6.635469 |
| 8.983957 | 9.090056 | 8.911718 | 9.81926 | 9.777678 | 9.012779 | 8.739629 | 8.907961 | 8.924917 |
| 6.901933 | 6.946417 | 6.968521 | 7.626388 | 7.351794 | 7.385392 | 7.275992 | 7.399396 | 7.935894 |
| 6.729503 | 6.646317 | 6.69649 | 6.641557 | 6.762123 | 6.525238 | 6.640056 | 6.69732 | 6.735363 |
| 8.068398 | 8.079143 | 8.029886 | 9.432138 | 9.319281 | 5.938759 | 6.271764 | 6.146708 | 6.309003 |
| 8.252605 | 8.174892 | 7.763687 | 7.351056 | 6.72998 | 7.404459 | 7.563875 | 7.742826 | 6.981163 |
| 9.398249 | 9.60547 | 9.729747 | 8.667562 | 8.545294 | 10.24699 | 10.07609 | 10.19871 | 7.938992 |
| 6.428426 | 6.401094 | 6.40764 | 4.499352 | 5.137641 | 5.616782 | 4.280452 | 5.715266 | 6.842571 |
| 9.371094 | 8.868351 | 8.796074 | NaN | 8.556641 | 8.227396 | 8.205366 | 8.35443 | 8.518462 |

| | | | | | | | | |
|---|---|---|---|---|---|---|---|---|
| 6.970332 | 7.133583 | 7.264143 | 6.355845 | 5.963602 | 7.251276 | 7.156205 | 7.156048 | 6.649308 |
| 5.305709 | 5.161398 | 6.716499 | 5.988519 | 6.052385 | 6.721593 | 5.833902 | 6.02514 | 6.00723 |
| 8.243505 | 8.780338 | 8.691384 | 6.762676 | 6.65133 | 8.136836 | 8.102129 | 7.652938 | 8.321708 |
| 7.330604 | 7.251338 | 7.271212 | 7.075183 | 6.928928 | 7.416615 | 7.313599 | 7.147506 | 7.238426 |
| 4.884477 | 4.807604 | 5.236135 | 4.503844 | 3.23583 | 5.836834 | 5.572838 | 5.601545 | 6.047224 |
| 8.325498 | 8.580471 | 8.290413 | 8.191758 | 7.837568 | 8.833517 | 8.582582 | 8.571353 | 8.219718 |
| 9.334055 | 9.329141 | 9.386004 | 8.430508 | 8.242558 | 9.279001 | 9.180494 | 9.207313 | 8.57902 |
| 11.93393 | 11.86372 | 11.90455 | 11.43057 | 11.6082 | 12.75636 | 12.90253 | 12.98046 | 11.53576 |
| 11.03927 | 11.00245 | 11.08817 | 13.11593 | 13.08179 | 11.40101 | 11.39212 | 11.50272 | 11.47712 |
| NaN | NaN | 4.907184 | 5.864625 | 6.081717 | NaN | 5.019929 | NaN | NaN |
| 8.843243 | 8.585948 | 8.589293 | 8.239827 | 7.976657 | 8.521454 | 8.437963 | 8.638451 | 8.467669 |
| 8.769345 | 8.680447 | 8.66129 | 9.053709 | 8.842261 | 8.408406 | 8.608231 | 8.973929 | 8.66385 |
| 6.697042 | 8.410864 | 7.637025 | 10.0883 | 11.45417 | 7.840341 | 8.885798 | 8.86259 | 9.687018 |
| NaN | NaN | NaN | 6.924747 | 7.244721 | NaN | NaN | NaN | NaN |
| 5.97403 | 5.141751 | 5.408243 | 5.793133 | 5.930453 | NaN | 6.814646 | 5.633458 | 6.431632 |
| 9.800809 | 9.571269 | 9.476809 | 8.150098 | 7.949884 | 9.428797 | 9.379531 | 9.460587 | 9.577446 |
| 9.757558 | 9.677516 | 9.815247 | 8.630634 | 8.636169 | 9.328636 | 9.530976 | 9.190295 | 8.365529 |
| 8.88184 | 8.80803 | 8.812096 | 8.743208 | 8.096597 | 8.355878 | 8.211605 | 8.410233 | 7.848319 |
| 10.4897 | 10.41498 | 10.38075 | 9.77961 | 9.602344 | 10.10238 | 10.1433 | 10.21619 | 10.08708 |
| 11.8686 | 11.53536 | 11.60686 | 12.56803 | 12.68896 | 10.97688 | 10.90222 | 10.95666 | 10.5109 |
| 8.285423 | 8.35384 | 8.224675 | 7.579841 | 7.638371 | 8.408793 | 8.361233 | 8.235864 | 7.784542 |
| 6.783616 | 6.164868 | 6.182241 | 6.068714 | 6.302527 | 7.996218 | 7.598878 | 7.515718 | 6.542645 |
| 7.566966 | 7.690274 | 7.945728 | 8.085602 | 8.137383 | 7.136245 | 6.857619 | 7.111888 | 8.225946 |
| 6.051216 | 5.638532 | 6.342381 | 5.100435 | 4.951901 | 6.750076 | 6.578298 | 6.445853 | 5.880742 |
| 7.189948 | 7.226488 | 7.194037 | 7.116804 | 7.137644 | 7.969587 | 8.0048 | 7.869359 | 6.640224 |
| 8.253519 | 8.556336 | 8.44679 | 7.029074 | 7.641228 | 7.785144 | 7.776878 | 7.929566 | 7.904239 |
| 10.03439 | 10.03395 | 9.785277 | 10.04309 | 10.2204 | 10.16002 | 9.999822 | 10.44251 | 9.948885 |
| 8.53001 | 8.389492 | 8.452972 | 8.266863 | 8.430955 | 8.263189 | 8.232702 | 8.371943 | 8.49932 |
| 5.371708 | 7.155219 | 7.087101 | 7.025618 | 7.171746 | 6.586636 | 5.881503 | 6.576735 | 5.381481 |
| 6.201878 | 6.194257 | 6.366873 | 5.613372 | 5.858964 | 5.64116 | 6.206075 | 6.058331 | 6.447904 |
| 6.909331 | 6.045579 | 6.917293 | 5.833771 | 5.526456 | 7.340282 | 7.421004 | 7.548655 | 6.979706 |
| 6.711121 | 6.689904 | 6.722925 | 6.251037 | 5.853568 | 7.694105 | 7.757472 | 7.826417 | 6.770026 |
| 5.973005 | 5.638237 | 6.475541 | 5.343942 | 4.942317 | 6.423925 | 6.29513 | 6.136864 | 5.936914 |
| 7.131433 | 7.291625 | 6.98206 | 5.209919 | 5.360953 | 6.938292 | 6.652032 | 6.872717 | 7.014977 |
| 7.70436 | 7.667314 | 7.740929 | 7.112503 | 6.759478 | 7.754623 | 7.61678 | 7.74075 | 7.754126 |
| 9.158378 | 9.217618 | 9.092339 | 7.75769 | 7.841333 | 9.300036 | 9.210257 | 9.280875 | 8.875344 |
| 6.857688 | 7.071337 | 7.075373 | 5.548557 | 5.685703 | 6.84678 | 7.034426 | 6.994865 | 6.933679 |
| 8.496129 | 8.487766 | 8.528212 | 7.325061 | 7.155909 | 8.570104 | 8.584466 | 8.478358 | 8.382313 |
| 6.233551 | 5.046687 | 5.809596 | 5.787396 | 5.70629 | 5.373347 | 5.172915 | 5.423819 | 5.991894 |
| 7.832437 | 7.129303 | 7.710966 | 6.413374 | 6.061527 | 8.036164 | 8.043234 | 7.792922 | 7.358704 |
| 8.782364 | 8.763355 | 8.771297 | 7.930746 | 7.240436 | 8.163532 | 8.211089 | 7.937151 | 8.059732 |
| 6.935922 | 7.069472 | 7.140653 | 7.03281 | 7.090136 | 7.226534 | 7.466649 | 6.597429 | 7.07641 |
| 6.195778 | 5.658996 | 6.074374 | 5.966946 | 6.860905 | 5.988328 | 6.073435 | 6.569625 | 7.183643 |
| 7.44434 | 7.844786 | 7.98252 | NaN | 8.421521 | 8.116783 | 7.842997 | 7.947291 | 8.833179 |
| 10.20996 | 9.548075 | 9.551232 | 10.33727 | 10.35588 | 9.858003 | 9.819561 | 9.834999 | 9.744224 |
| 5.514667 | 5.574759 | 5.268625 | NaN | 3.909785 | 4.663726 | 4.619644 | 4.93117 | 5.775146 |
| 6.416448 | 4.051912 | 6.376166 | 5.589045 | NaN | 7.083884 | 6.938099 | 6.933853 | 6.626226 |

| | | | | | | | | |
|---|---|---|---|---|---|---|---|---|
| 7.41098 | 6.329455 | 7.434875 | 5.614749 | 6.697349 | 6.291444 | 6.346064 | 6.685399 | 6.831097 |
| 9.966516 | 9.995596 | 10.05092 | 9.242863 | 9.118606 | 10.20573 | 10.22822 | 10.2178 | 9.958051 |
| 9.338175 | 9.086876 | 8.90814 | 8.230151 | 8.644524 | 9.471961 | 9.451208 | 9.372811 | 7.925731 |
| 7.301469 | 7.061534 | 7.02296 | 4.798689 | 5.106245 | 7.039347 | 7.122266 | 7.098474 | 6.255591 |
| 6.627687 | 7.064107 | 6.502492 | 7.276903 | 7.139944 | 6.914851 | 6.378901 | 6.775806 | 6.456049 |
| 10.4303 | 10.40388 | 10.39707 | 10.32417 | 10.34012 | 10.905 | 10.84212 | 10.89815 | 9.910078 |
| 7.920077 | 6.810486 | 6.522709 | 5.409932 | 5.388854 | 6.301445 | 6.349488 | 6.524123 | 6.002136 |
| 8.445705 | 8.656449 | 8.352818 | 7.333085 | 7.910839 | 8.008659 | 7.878278 | 8.051216 | 7.469532 |
| 7.915217 | 7.109139 | 7.173274 | 5.912036 | 5.347353 | NaN | NaN | NaN | 5.613295 |
| 11.65746 | 11.61802 | 11.6396 | 10.50219 | 10.41891 | 11.11233 | 11.20092 | 11.26867 | 11.65515 |
| 9.6895 | 9.713481 | 9.605863 | 9.43869 | 9.197319 | 9.266517 | 9.321007 | 9.26917 | 8.855093 |
| 7.698908 | 7.816514 | 7.822617 | 7.714035 | 7.950008 | 7.273337 | 7.549942 | 7.245257 | 7.508269 |
| 5.897139 | NaN | 5.254913 | NaN | 4.185075 | NaN | 6.402737 | 6.680831 | 4.790799 |
| 5.665813 | 5.593626 | 5.919186 | 6.590736 | 7.159481 | 6.649093 | 6.069584 | 6.078773 | 5.770639 |
| 14.27564 | 14.47851 | 14.47132 | 14.22609 | 15.03095 | 14.50757 | 13.77004 | 14.46464 | 13.57684 |
| 6.854156 | 7.119255 | 6.917101 | 6.63745 | 7.083098 | 6.642356 | 6.543449 | 6.507365 | 6.63817 |
| 8.2196 | 8.313264 | 8.247164 | 8.059847 | 8.493166 | 8.182019 | 8.015544 | 8.200955 | 8.432255 |
| NaN | NaN | 7.977502 | 6.337002 | 6.93303 | NaN | NaN | NaN | 8.877585 |
| NaN | NaN | NaN | NaN | NaN | NaN | 7.531227 | 6.480078 | 6.446793 |
| 7.046573 | 7.335849 | 7.417026 | 7.224547 | 6.836176 | 7.593831 | 7.447411 | 7.04569 | 7.692675 |
| 7.306901 | 7.034042 | 7.18777 | 8.220413 | 8.049074 | 7.136526 | 6.805607 | 6.592929 | 6.642749 |
| 6.230502 | 5.911666 | 5.953041 | 5.189236 | 5.447143 | 5.980788 | 6.113663 | 6.551124 | 5.132048 |
| 9.247102 | 9.392385 | 9.424933 | 8.818346 | 8.820286 | 9.616325 | 9.582403 | 9.702737 | 9.595204 |
| 9.655188 | 9.576923 | 10.03029 | 11.84122 | 12.0254 | 9.137312 | 9.854715 | 9.637742 | 9.921787 |
| 7.15419 | 7.797307 | 6.828038 | 7.175052 | 6.750067 | 6.696768 | 6.632023 | 6.934687 | 6.180382 |
| 7.794403 | 7.874883 | 7.935723 | 8.719954 | 8.599856 | 8.144515 | 7.894578 | 8.089886 | 7.878146 |
| 8.409835 | 8.722552 | 8.777207 | 10.655 | 10.68195 | 8.702345 | 8.518575 | 8.362546 | 8.377497 |
| 8.439238 | 7.9767 | 8.082285 | 6.738327 | 8.912248 | 6.1731 | 6.679792 | 6.538233 | 5.781756 |
| 11.23494 | 11.4504 | 11.37952 | 12.35069 | 12.45349 | 11.08117 | 11.33376 | 11.37897 | 10.75985 |
| 10.29287 | 10.12852 | 10.14354 | 8.698843 | 8.506255 | 10.13341 | 10.18193 | 10.05631 | 10.07862 |
| 5.695292 | 3.723859 | 3.089335 | NaN | NaN | 4.251612 | NaN | NaN | 5.951341 |
| 8.91196 | 8.915315 | 9.198141 | NaN | NaN | 7.608557 | 7.28986 | 7.711473 | 7.29427 |
| 8.675008 | 8.476459 | 8.319654 | 9.478199 | 9.420197 | 8.800182 | 8.753771 | 8.54837 | 8.343844 |
| 6.258753 | 6.220282 | 5.741424 | 5.866807 | 5.792581 | 6.361795 | 6.516666 | 6.51184 | 6.650928 |
| 3.953026 | 3.202201 | 4.162058 | NaN | 4.069201 | 4.300227 | 3.885655 | NaN | 5.407162 |
| 7.752523 | 6.553809 | 7.105828 | 6.151399 | 7.371782 | 8.090785 | 7.325185 | 7.830382 | 7.498203 |
| 7.02616 | 6.912126 | 7.262691 | 4.813497 | 5.314966 | 7.795822 | 7.889841 | 7.673084 | 6.676817 |
| 9.187593 | 9.260666 | 9.181715 | 8.169841 | 8.531955 | 9.2033 | 8.779 | 8.690862 | 7.871095 |
| 6.314626 | 6.140435 | 5.969135 | NaN | NaN | 6.494265 | 6.298513 | 6.487059 | 5.503603 |
| 8.143075 | 8.661862 | 8.145117 | 8.123963 | 7.987736 | 6.797441 | 7.306999 | 7.2505 | 7.563136 |
| 9.579044 | 9.396684 | 9.40972 | 8.969895 | 8.895504 | 9.107791 | 9.202719 | 9.157975 | 9.629402 |
| 8.358332 | 8.230384 | 7.996902 | 7.967351 | 8.141059 | 8.331381 | 8.287518 | 8.448302 | 7.54101 |
| 7.525217 | 7.567008 | 7.368513 | 9.076542 | 9.116652 | 6.851036 | 6.865684 | 6.945518 | 8.281392 |
| 8.405249 | 8.279212 | 8.321418 | 8.731189 | 8.564369 | 8.372954 | 8.524158 | 8.440331 | 8.626184 |
| 6.135533 | 6.508821 | 6.447202 | 6.5853 | 6.300291 | 6.47638 | 7.197168 | 6.499502 | 6.269323 |
| 9.336287 | 9.243141 | 9.466103 | 8.530935 | 8.208112 | 9.634252 | 9.69541 | 9.555223 | 9.539359 |
| 8.535282 | 8.444146 | 8.744942 | 7.032599 | 7.069708 | 8.742447 | 8.702681 | 8.751244 | 8.329017 |

-Bruker Confidential-

| | | | | | | | | |
|---|---|---|---|---|---|---|---|---|
| 8.065567 | 7.86617 | 8.010797 | 7.598903 | 7.984074 | 7.07557 | 7.089188 | 6.885893 | 7.787979 |
| 6.792606 | 6.556202 | 5.54879 | NaN | NaN | 6.329662 | 6.520792 | 6.265725 | 5.552279 |
| 6.407265 | 6.840763 | 6.567069 | 5.858926 | 6.052617 | 6.340065 | 6.526559 | 6.631192 | 6.641923 |
| 4.866325 | 4.059835 | 4.569664 | NaN | 3.80848 | 5.665039 | 5.614792 | 5.551813 | 4.660246 |
| 4.730985 | 5.527252 | 4.733968 | NaN | NaN | 5.043219 | 5.398342 | 5.249293 | 4.781895 |
| 4.865094 | 4.594361 | 7.745325 | NaN | NaN | NaN | NaN | NaN | 5.985782 |
| 12.57086 | 12.44624 | 12.45435 | 12.24964 | 12.34839 | 12.47608 | 12.67363 | 12.48553 | 11.76755 |
| 12.65176 | 12.58184 | 12.65487 | 12.31361 | 12.29349 | 12.99248 | 13.11166 | 13.09261 | 12.11563 |
| 8.523973 | 8.159416 | 8.567042 | 8.395133 | 8.510253 | 7.899238 | 8.012677 | 7.843083 | 7.849784 |
| 7.42346 | 6.661838 | 6.901668 | 7.087084 | 5.559609 | 6.48754 | 6.830447 | 7.065917 | 6.059959 |
| 11.10281 | 11.14577 | 11.13272 | 11.05478 | 10.90929 | 11.15762 | 11.12836 | 11.12494 | 10.83356 |
| 12.29256 | 12.22566 | 12.20337 | 10.94731 | 10.82512 | 12.39201 | 12.39598 | 12.35433 | 12.12209 |
| 7.328707 | 7.603395 | 7.091908 | 6.327935 | 6.053315 | 7.835907 | 7.260865 | 7.56988 | 7.655273 |
| 8.602476 | 8.535081 | 8.585442 | 7.758043 | 7.341792 | 7.802041 | 8.174889 | 8.189326 | 8.386311 |
| 6.673124 | 6.602323 | 6.448472 | 5.709543 | 6.017006 | 6.468487 | 6.123331 | 6.233493 | 6.501523 |
| NaN | NaN | 6.769386 | 8.726858 | 8.957401 | 6.483625 | 6.180979 | 6.053056 | 6.004168 |
| 8.925885 | 8.810172 | 8.7364 | 10.81682 | 10.58865 | 8.040377 | 7.864888 | 7.459309 | 9.117682 |
| 7.771401 | 7.739744 | 7.973512 | 7.03309 | 6.80623 | 7.944723 | 7.707037 | 7.714828 | 7.189472 |
| 11.29 | 11.15012 | 11.18328 | 12.55048 | 12.09855 | 10.32323 | 10.44779 | 10.23788 | 11.31998 |
| 7.73647 | 7.188119 | 7.241794 | 6.774949 | 5.784275 | 7.681754 | 7.439658 | 7.25908 | 7.311807 |
| 8.846671 | 8.574459 | 8.771685 | 9.66328 | 9.56112 | 10.29562 | 10.1249 | 10.19849 | 10.14535 |
| 8.009817 | 8.127147 | 7.928958 | 7.052644 | 7.393861 | 7.528386 | 7.566983 | 7.661228 | 8.130428 |
| 8.41062 | 8.369776 | 8.557687 | 8.605985 | 8.762054 | 7.819454 | 7.784357 | 7.694256 | 8.595133 |
| 7.360802 | 7.318337 | 7.058286 | 6.533873 | 7.877119 | 7.833692 | 7.480302 | 7.513786 | 7.677677 |
| 8.747255 | 8.648562 | 8.346128 | 8.092795 | 7.479211 | 8.450989 | 8.902328 | 8.353456 | 8.209114 |
| 8.516961 | 8.359119 | 8.127513 | 7.857109 | 7.679841 | 8.236684 | 8.167486 | 8.024836 | 6.851226 |
| 8.499294 | 8.440935 | 8.602209 | 7.945344 | 7.986282 | 7.897183 | 7.65252 | 7.621771 | 8.168986 |
| 3.626864 | 4.276484 | 2.864062 | 4.003055 | NaN | NaN | 3.182116 | 6.031971 | 4.638799 |
| 10.05114 | 9.872832 | 9.982947 | 10.40183 | 10.31396 | 9.815399 | 9.848402 | 9.854218 | 10.20177 |
| 6.551089 | 6.020763 | 6.283271 | 6.573703 | 6.787372 | 7.050949 | 6.960203 | 6.575102 | 6.551225 |
| 9.482864 | 9.293989 | 9.297097 | 9.102752 | 9.190018 | 9.049574 | 9.093353 | 9.216684 | 9.228189 |
| 7.661114 | 7.703591 | 7.595936 | 6.433602 | 6.203222 | 6.670975 | 6.919435 | 6.462701 | 6.260776 |
| 7.885928 | 7.875881 | 7.960714 | 7.056322 | 7.130084 | 7.389855 | 7.317501 | 7.558943 | 7.827228 |
| 6.707941 | 6.202758 | 6.525203 | 6.112525 | 6.143775 | 6.612441 | 6.775096 | 6.651797 | 6.220873 |
| 8.235756 | 7.913673 | 7.879974 | 7.343791 | 10.47948 | NaN | NaN | 8.146277 | NaN |
| 7.089199 | 6.855748 | 7.090544 | 6.858719 | 7.069061 | 7.294286 | 6.998052 | 7.117694 | 6.576987 |
| 6.457009 | 6.6552 | 7.391979 | 5.809126 | 6.015004 | 6.976125 | 7.331888 | 7.343884 | 7.530715 |
| 7.653544 | 7.844409 | 8.063078 | 7.029993 | 7.213403 | 8.315698 | 8.521885 | 8.462731 | 7.424014 |
| 5.539306 | 5.853702 | 5.722157 | 5.818547 | 5.414023 | 6.281695 | 6.262149 | 6.375987 | 5.830857 |
| 7.406594 | 7.367574 | 7.524477 | 7.194095 | 7.235494 | 6.631264 | 6.715802 | 7.0912 | 7.119502 |
| 8.915626 | 8.838463 | 9.094913 | 7.020216 | 6.736566 | 8.075209 | 8.198378 | 8.210776 | 8.404815 |
| NaN | NaN | NaN | 6.193933 | 5.258227 | 7.113448 | 6.95074 | 7.457201 | 5.928358 |
| 8.63485 | 6.924969 | 7.297522 | 7.424793 | 7.525104 | NaN | NaN | NaN | NaN |
| 8.712423 | 8.5709 | 8.588264 | 9.08954 | 9.098999 | 8.86977 | 8.758491 | 8.990638 | 7.846006 |
| NaN | 6.769301 | NaN | NaN | NaN | NaN | NaN | NaN | 7.089669 |
| 8.306493 | 8.029749 | 7.994504 | 7.243566 | 7.676959 | NaN | NaN | NaN | 6.464102 |
| 6.526402 | 6.794003 | 6.101387 | NaN | 5.324078 | 6.581856 | 6.782909 | 6.503722 | 6.375092 |

-Bruker Confidential-

| | | | | | | | | |
|---|---|---|---|---|---|---|---|---|
| NaN | NaN | NaN | NaN | NaN | 2.604556 | 3.823727 | 4.260885 | NaN |
| 7.057663 | 6.900886 | 7.024037 | NaN | 7.147459 | 6.915135 | 6.567774 | 6.587071 | 7.87377 |
| NaN | 6.688072 | 6.267425 | 6.280615 | 5.338577 | 7.000164 | 6.972935 | 6.964041 | 5.952326 |
| NaN | 5.052961 | 5.765708 | NaN | NaN | NaN | 6.021662 | 6.224783 | 4.263082 |
| 7.987726 | 8.022568 | 8.139156 | 7.309817 | 7.528549 | 7.904192 | 7.932187 | 8.133472 | 8.115366 |
| 9.591773 | 9.432135 | 9.845919 | 8.797859 | 8.738345 | 9.204492 | 9.479041 | 9.30586 | 9.501104 |
| 6.847765 | 6.768709 | 6.9108 | 7.201605 | 6.878613 | 6.700089 | 6.793768 | 6.533813 | 7.084786 |
| 8.545277 | 8.423511 | 8.406354 | 9.87379 | 9.655534 | 8.294413 | 8.115686 | 8.49824 | 9.155144 |
| 2.621869 | NaN | NaN | NaN | NaN | 3.045297 | NaN | 2.620993 | 3.154438 |
| NaN | 6.942338 | 5.853416 | 5.546598 | 5.940045 | 6.770753 | 6.447418 | 6.761811 | 5.523558 |
| 9.047944 | 9.022458 | 8.977314 | 8.500676 | 8.301524 | 9.036414 | 9.077946 | 8.952994 | 8.886924 |
| 6.015646 | 5.808905 | 6.152162 | 5.761517 | 5.457489 | 6.783284 | 6.468967 | 6.615402 | 5.96568 |
| 5.530765 | 6.774759 | 5.683978 | 4.988838 | 5.851691 | 6.285093 | 5.69591 | 6.361135 | 5.894211 |
| 8.016117 | 7.78487 | 7.939453 | 7.024204 | 6.983482 | 7.232447 | 7.253649 | 7.67424 | 7.016623 |
| 7.576811 | 7.643491 | 7.626733 | 6.884919 | 7.389838 | 7.211041 | 7.023694 | 7.240475 | 7.532645 |
| NaN | NaN | 5.594932 | NaN | NaN | NaN | NaN | NaN | 4.873074 |
| 6.059886 | 6.193532 | 5.964681 | 5.135808 | 5.540892 | 6.291742 | 6.526101 | 6.500647 | 6.542071 |
| 6.974524 | 7.110818 | 7.331791 | 6.426936 | 6.46336 | 7.487604 | 7.1916 | 7.46437 | 7.193903 |
| 9.362177 | 9.324854 | 9.383647 | 10.67468 | 10.65784 | 9.499755 | 9.367092 | 9.550658 | 9.579708 |
| 10.75326 | 10.75933 | 10.77331 | 9.724011 | 9.995294 | 11.14837 | 11.15724 | 11.22701 | 11.00599 |
| 8.221758 | 8.432837 | 8.111953 | 6.827117 | 7.001587 | 7.542807 | 7.906684 | 7.878941 | 6.455377 |
| 6.631875 | 6.068289 | 6.343704 | 4.495484 | 4.681113 | 5.957336 | 6.496784 | 5.76801 | 6.771678 |
| 8.791822 | 8.878859 | 9.074526 | 7.956935 | 8.035818 | 8.643995 | 8.142987 | 8.128298 | 8.595087 |
| 7.217957 | 7.58639 | 7.454247 | 7.631167 | 7.107898 | 6.507411 | 6.453067 | 6.759014 | 7.22884 |
| 6.242785 | 6.918496 | 6.241874 | 5.842613 | 6.004006 | 6.747253 | 6.967302 | 6.944546 | 6.926162 |
| 8.037575 | 7.846958 | 8.044985 | 7.038907 | 6.921153 | 8.443316 | 8.355132 | 8.318965 | 7.115351 |
| 7.353379 | 7.438343 | 7.445136 | 7.270648 | 7.31855 | 7.206811 | 7.178985 | 7.452231 | 7.784852 |
| 6.561868 | 6.383756 | 6.779877 | 4.986007 | 4.636082 | 6.901539 | 6.471372 | 6.454133 | 6.491648 |
| 6.013685 | 6.651084 | 5.46298 | NaN | NaN | 6.822601 | 7.727796 | 7.540929 | NaN |
| 9.353728 | 9.301861 | 9.303426 | 9.097914 | 9.147072 | 9.278552 | 9.301015 | 9.412618 | 9.341722 |
| 8.857039 | 8.830775 | 8.696489 | 10.36976 | 10.14152 | 8.209996 | 8.181016 | 8.382854 | 8.320683 |
| 5.185112 | 5.385362 | 5.39199 | 5.31488 | 6.451321 | 5.933427 | 6.302826 | 6.457883 | 4.531548 |
| 6.430902 | 6.855534 | 6.57829 | 6.547709 | 6.557298 | 6.722491 | 6.554338 | 6.346226 | 7.900759 |
| 8.719935 | 8.441133 | 8.11175 | 9.473495 | 9.505347 | 7.796227 | 8.143552 | 7.811043 | 8.332314 |
| 4.453481 | 5.087452 | 4.422074 | 8.958359 | 3.517851 | NaN | 6.86652 | NaN | 6.277618 |
| 8.379494 | 8.180287 | 8.038707 | 7.926157 | 8.058434 | 7.991976 | 8.124638 | 8.061993 | 8.312381 |
| 11.05762 | 11.07599 | 11.01104 | 9.81553 | 9.747982 | 11.20288 | 11.03694 | 11.09655 | 11.34714 |
| 6.147082 | 6.514153 | 6.267839 | 8.099381 | 7.909033 | 5.900928 | 6.198098 | 6.231895 | 5.684324 |
| 8.832085 | 8.741162 | 8.717706 | 8.071106 | 8.019334 | 8.52244 | 8.522977 | 8.626354 | 8.958235 |
| 8.178277 | 8.022653 | 7.749999 | 7.126854 | 7.53775 | 7.949306 | 7.431586 | 8.22797 | 5.289212 |
| 6.089637 | 5.984314 | 6.170013 | 5.751455 | 5.065922 | 6.819174 | 6.49964 | 7.06833 | 5.876773 |
| 6.733817 | 7.31329 | 5.564749 | NaN | 5.719006 | 7.326944 | 7.546519 | 7.350241 | 6.744003 |
| 8.398072 | 8.407922 | 8.352853 | 7.189211 | 6.593389 | 8.837345 | 8.496602 | 8.717229 | 8.10983 |
| 5.256025 | 5.327567 | 5.435465 | 6.43303 | 6.565777 | 5.4803 | 5.232137 | 5.736904 | 5.900514 |
| 6.422143 | 6.167973 | 6.282237 | 5.54528 | 6.053322 | 6.236905 | 6.593718 | 6.57153 | 6.463849 |
| 6.459505 | 6.461375 | 5.765455 | 6.443357 | 6.146138 | 7.658705 | 7.736015 | 7.543778 | NaN |
| 7.574043 | 7.246406 | 7.446486 | 5.293379 | 5.041748 | 7.195836 | 6.517756 | 7.071954 | 6.470828 |

| | | | | | | | | |
|---|---|---|---|---|---|---|---|---|
| 8.166164 | 8.088112 | 8.109317 | 6.357293 | 6.899297 | 8.329552 | 8.821729 | 8.646395 | 8.386104 |
| 8.517787 | 8.576855 | 8.539437 | 7.605688 | 7.548652 | 8.537483 | 8.358398 | 8.382145 | 8.39782 |
| 7.428897 | 7.14021 | 7.234123 | 8.027337 | 7.76915 | 7.158879 | 7.2942 | 7.262108 | 7.53955 |
| 9.607147 | 9.504714 | 9.52829 | 7.235001 | 7.080985 | 9.534286 | 9.587476 | 9.492237 | 9.412845 |
| 4.269859 | 4.664971 | 4.978597 | NaN | 4.950008 | 5.014944 | 4.525646 | 4.721821 | 6.46787 |
| 6.862617 | 6.862694 | 6.980367 | 7.179653 | 6.99352 | 5.881255 | 6.591573 | 6.490218 | 6.227242 |
| 7.572165 | 8.148399 | 8.14628 | 7.544127 | 7.351028 | 8.301152 | 8.359891 | 8.374733 | 6.571934 |
| 10.55984 | 10.3738 | 10.42029 | 11.75437 | 11.72438 | 11.28868 | 11.10179 | 11.10915 | 10.35098 |
| 8.297616 | 8.192116 | 8.161947 | 7.285184 | 7.070166 | 8.145182 | 8.048062 | 8.083016 | 7.933383 |
| 6.681176 | 6.564815 | 6.918231 | 4.981232 | 4.861999 | 5.936813 | 5.66943 | 5.938136 | 6.179051 |
| 7.360325 | 6.6958 | 7.314076 | 6.167531 | 6.613064 | 6.751968 | 6.319375 | 6.846101 | 7.077749 |
| 9.02676 | 9.022379 | 9.023206 | 8.035433 | 7.965607 | 8.764406 | 8.781281 | 8.642414 | 8.822968 |
| 8.828414 | 8.661459 | 8.765672 | 7.499604 | 7.663316 | 8.678824 | 8.734005 | 8.943736 | 8.480372 |
| 7.442499 | 7.253777 | 6.930419 | 7.022265 | 6.773042 | 7.285777 | 7.16623 | 7.307153 | 7.482825 |
| 7.693587 | 8.673275 | 8.225676 | NaN | 5.739074 | 8.466213 | 8.463733 | 8.113437 | 8.463817 |
| 7.492896 | 7.361748 | 7.628835 | 6.561744 | 6.275789 | 7.576608 | 7.501733 | 7.568933 | 7.287917 |
| NaN | 7.40178 | 7.179043 | NaN | NaN | 7.073328 | 7.250679 | 7.211912 | 7.216919 |
| 8.333829 | 8.15009 | 8.029477 | 7.440356 | 7.463735 | 8.198189 | 8.138359 | 8.256322 | 8.141597 |
| 6.465996 | 6.194537 | 5.771904 | NaN | 8.133537 | 7.237466 | 7.049168 | 7.22289 | 7.987374 |
| 7.089206 | 7.066898 | 7.06705 | 5.636833 | 5.248252 | 7.259687 | 6.649615 | 6.771925 | 6.827205 |
| 6.904902 | 6.726943 | 6.903158 | 8.242777 | 8.0028 | 7.009212 | 6.883885 | 6.973 | 7.432967 |
| 5.291379 | 4.61756 | 4.622226 | 6.443498 | NaN | 5.242246 | 4.635665 | 4.588592 | 5.37727 |
| 6.159542 | 5.073047 | 6.116208 | 6.273364 | 6.206275 | 3.646377 | 4.886057 | 4.57694 | 6.025278 |
| 8.489173 | 8.611928 | 8.224406 | 9.789577 | 9.567838 | 8.310074 | 7.905356 | 8.231271 | 8.07619 |
| 7.974033 | 7.696331 | 7.752156 | 7.976328 | 7.671281 | 7.889688 | 7.296114 | 7.681215 | 7.699146 |
| 5.073533 | 6.094372 | 4.955452 | 6.251899 | 5.594521 | 6.26481 | 6.687572 | 6.554924 | 5.538331 |
| 8.087508 | 6.999159 | NaN | NaN | NaN | NaN | NaN | NaN | 6.720945 |
| 8.939969 | 8.822895 | 8.676791 | 7.984467 | 8.108698 | 9.360053 | 9.404034 | 9.388041 | 8.502769 |
| 7.613961 | 7.75517 | 7.619622 | 7.695005 | 7.744661 | 7.722687 | 7.671028 | 7.552496 | 7.422737 |
| 11.09045 | 10.7056 | 10.77814 | 11.78127 | 11.89424 | 10.93951 | 10.81389 | 10.91344 | 10.8372 |
| 9.536783 | 9.350133 | 9.439576 | 9.830111 | 9.928808 | 9.584962 | 9.658827 | 9.652677 | 8.563352 |
| 6.209112 | 6.809748 | 6.194184 | 6.977808 | 6.962698 | 6.931641 | 6.986836 | 7.049981 | 5.997157 |
| 7.190617 | 7.621509 | 7.154227 | 7.363022 | 7.096283 | NaN | 6.381855 | 6.802114 | 7.026964 |
| 7.166709 | 6.764623 | 6.984589 | 6.401593 | 6.922755 | 6.95083 | 6.455228 | 6.919347 | 7.23048 |
| 10.37912 | 10.32332 | 10.01185 | 11.68151 | 11.68603 | 10.33189 | 10.34278 | 10.50536 | 9.925663 |
| 8.542504 | 8.672371 | 9.534321 | 5.937262 | 6.275081 | 9.404492 | 9.401178 | 9.364226 | 7.102808 |
| 8.899489 | 8.949772 | 8.890063 | 9.029808 | 9.058304 | 8.969481 | 8.691629 | 8.329615 | 8.889066 |
| 5.704466 | 5.6595 | 4.818022 | NaN | 5.656968 | 6.560433 | 6.16902 | 6.759425 | 4.780789 |
| 6.861963 | 6.759568 | 6.644977 | 6.699712 | 7.058041 | 7.614324 | 6.790559 | 6.785784 | 6.846634 |
| 9.002287 | 8.961311 | 8.87938 | 9.215367 | 9.126498 | 9.3433 | 9.361564 | 9.412876 | 9.414593 |
| 8.707774 | 8.726038 | 8.476044 | 8.811163 | 8.939112 | 8.17539 | 8.295963 | 8.202006 | 9.020768 |
| 8.520375 | 8.486395 | 8.585634 | 7.026121 | 7.310638 | 8.239455 | 8.221884 | 8.427867 | 8.321998 |
| 9.51145 | 9.607491 | 9.602171 | 8.432979 | 8.682732 | 9.587218 | 9.481873 | 9.708846 | 7.251666 |
| 6.898461 | 7.12025 | 6.980899 | 6.168728 | 5.745099 | NaN | 6.331524 | 6.549726 | 6.640112 |
| 5.061828 | NaN | 4.530747 | NaN | NaN | 6.462777 | 5.318022 | 6.045914 | 4.537937 |
| 7.035387 | 6.617172 | 6.698896 | 6.031011 | 6.437048 | 6.390151 | 6.59444 | 6.303435 | 6.784755 |
| 9.750675 | 9.765825 | 9.697417 | 10.8211 | 10.92776 | 9.933161 | 10.15193 | 10.07962 | 10.13555 |

| | | | | | | | | |
|---|---|---|---|---|---|---|---|---|
| 9.683914 | 9.510146 | 9.46192 | 9.927495 | 10.09041 | 9.33825 | 9.036641 | 9.21431 | 9.942563 |
| 7.563885 | 7.519155 | 7.903629 | 7.690906 | 7.332561 | 7.939374 | 8.157006 | 8.410048 | 7.979652 |
| 7.738562 | 7.977898 | 7.80891 | 7.677302 | 7.873474 | 7.875485 | 7.535596 | 7.584759 | 5.989791 |
| 12.31355 | 12.28523 | 12.274 | 12.53209 | 12.47318 | 11.72139 | 11.59499 | 11.62881 | 11.94933 |
| 7.791938 | 8.016521 | 7.97984 | 8.119115 | 8.194679 | 8.395125 | 8.439031 | 8.618839 | 8.359529 |
| 7.700004 | 7.329043 | 7.740974 | 5.283114 | 5.5368 | 7.302566 | 7.551995 | 7.34979 | 7.412041 |
| 5.889733 | 5.318942 | 5.190518 | 8.683894 | 10.33885 | 6.852982 | NaN | 7.027602 | 7.174716 |
| 8.521127 | 8.453831 | 8.349112 | 8.408602 | 8.356788 | 8.323564 | 8.278949 | 8.367916 | 8.3352 |
| 4.715923 | 4.144297 | 4.980549 | NaN | NaN | 5.918089 | 5.433006 | 5.596345 | 4.510396 |
| 8.708069 | 8.501512 | 9.045885 | 7.174876 | 7.519903 | 8.703754 | 8.364238 | 8.68115 | 7.777597 |
| 9.321051 | 9.281845 | 9.40009 | 10.72358 | 11.15014 | 9.582997 | 9.605536 | 9.636324 | 9.02399 |
| 7.84472 | 6.765115 | 7.685654 | 6.485071 | 6.080853 | 7.923346 | 8.072774 | 7.945091 | 8.14545 |
| 11.77255 | 11.79294 | 11.82386 | 13.31126 | 13.39957 | 11.54423 | 11.46799 | 11.45866 | 13.13285 |
| 6.910448 | 6.717123 | 6.701363 | 6.914811 | 7.252756 | 7.238149 | 7.273204 | 7.524455 | 6.962888 |
| 5.669953 | 5.367667 | 5.389508 | 5.844713 | 5.261934 | 5.450702 | 5.336372 | 5.169487 | 4.979919 |
| 7.893972 | 7.855108 | 7.805208 | 6.300167 | 6.436937 | 7.785982 | 8.013189 | 8.20474 | 7.923423 |
| 5.885451 | 4.662296 | 5.292686 | NaN | NaN | 7.520154 | 7.583478 | 7.446641 | 5.829758 |
| 7.917221 | 7.636477 | 7.782962 | 6.906329 | 6.604487 | 6.836096 | 7.228138 | 7.315146 | 7.349092 |
| 10.35071 | 10.669 | 10.69845 | 10.53369 | 10.65871 | 10.0304 | 10.21542 | 10.11844 | 10.76277 |
| 11.70677 | 11.78024 | 11.84183 | 11.4769 | 11.41332 | 11.44558 | 11.36697 | 11.56271 | 12.07928 |
| 8.078601 | 7.894986 | 7.926327 | 8.695844 | 8.40903 | 7.609972 | 7.456115 | 7.592563 | 7.776613 |
| 10.71196 | 10.91896 | 10.78666 | 11.37797 | 11.44789 | 10.46324 | 10.14621 | 10.45017 | 10.43396 |
| 5.700603 | 5.351522 | 5.737323 | NaN | NaN | 6.820701 | 6.528601 | 6.624842 | 5.178181 |
| 10.45055 | 10.28668 | 10.37257 | 10.43862 | 10.76926 | 9.058 | 9.018641 | 9.219994 | 8.675104 |
| 7.38323 | 6.506427 | 6.705118 | NaN | NaN | 6.368319 | 6.58887 | 6.72878 | 6.507854 |
| 7.42405 | 6.516516 | 7.80285 | 6.805377 | 6.596502 | 6.287478 | 5.792594 | NaN | 5.478942 |
| 8.024229 | 9.140352 | 7.973269 | 6.53267 | 8.930082 | 7.592433 | 7.549623 | 7.850227 | 8.063987 |
| 8.545251 | 8.521694 | 8.552753 | 6.594561 | 6.689301 | 8.629459 | 8.454186 | 8.472996 | 8.779464 |
| 6.52283 | 5.987101 | 6.894314 | 5.260859 | 5.800799 | 6.808934 | 6.674411 | 6.733344 | 6.332968 |
| 9.050677 | 8.775306 | 9.37617 | 11.69886 | 11.32115 | 9.442604 | 9.460054 | 9.223198 | 9.905588 |
| 7.354666 | 7.260387 | 7.203187 | 8.160658 | 8.409714 | 6.833762 | 6.938053 | 6.918429 | 8.238522 |
| 8.546406 | 8.321686 | 8.279526 | 7.045297 | 7.357645 | 8.506214 | 8.604274 | 8.63385 | 8.377139 |
| 7.30162 | 7.4452 | 7.178129 | 7.633745 | 7.179205 | 6.156208 | 6.063359 | 6.247756 | 6.625416 |
| 6.413788 | 6.759114 | 6.54446 | 7.716543 | 7.94162 | 6.506987 | 6.80565 | 6.459836 | 6.822158 |
| 8.706873 | 8.607613 | 8.699952 | 9.427173 | 9.575365 | 8.618638 | 8.791737 | 8.73246 | 8.350934 |
| 10.37915 | 10.1725 | 10.22304 | 11.34945 | 11.47255 | 10.34281 | 10.32662 | 10.36916 | 9.829015 |
| 8.668301 | 8.603538 | 8.61538 | 7.65524 | 7.516684 | 8.76017 | 9.254064 | 9.139512 | 8.985044 |
| 7.939757 | 7.812294 | 7.91888 | 6.99747 | 6.96541 | 8.18309 | 8.181783 | 8.179233 | 8.670298 |
| 8.474628 | 8.446507 | 8.258187 | 9.540951 | 9.533637 | 8.251879 | 8.216203 | 8.328538 | 8.825816 |
| 7.057961 | 5.520416 | 6.064533 | 5.896389 | 5.350383 | 7.687511 | 6.945648 | 7.435249 | NaN |
| 8.182916 | 7.995793 | 7.99047 | 7.310804 | 8.321727 | 7.48844 | 8.099784 | 7.928198 | 7.818111 |
| 6.271903 | 6.574781 | 6.85432 | 7.293414 | 7.362236 | NaN | NaN | NaN | 7.226391 |
| 11.22803 | 11.44902 | 11.32093 | 8.995319 | 8.883409 | 6.728149 | 6.505426 | 6.913212 | 6.287423 |
| 7.276076 | 7.388385 | 6.989902 | 8.050751 | 7.996074 | 7.736382 | 7.024706 | 7.288338 | 5.568301 |
| 7.523591 | 7.44374 | 7.518677 | 7.467189 | 7.298992 | 7.038022 | 6.940624 | 7.314953 | 7.365 |
| 6.632877 | 6.187245 | 6.536636 | 6.418766 | 6.286121 | NaN | NaN | NaN | 5.276224 |
| 5.074813 | 5.359339 | 4.730464 | NaN | NaN | 5.725811 | 6.092951 | 5.980374 | 5.277493 |

| | | | | | | | | |
|---|---|---|---|---|---|---|---|---|
| NaN | 8.010315 | 7.605866 | 7.91015 | 7.636235 | NaN | 6.391696 | NaN | 7.516369 |
| 7.750879 | 7.947484 | 7.843185 | 7.642787 | 7.691816 | 7.448296 | 7.758579 | 7.644731 | 7.610267 |
| 8.105744 | 8.191502 | 8.157725 | 8.457327 | 8.537926 | 8.326514 | 8.461738 | 8.194524 | 7.48793 |
| NaN | NaN | NaN | NaN | NaN | NaN | NaN | NaN | 6.858548 |
| 3.889874 | 4.427786 | NaN | 5.330949 | 6.292664 | NaN | NaN | 4.246678 | 6.94288 |
| 5.945782 | 6.223799 | 5.955901 | 6.739797 | 6.841051 | 5.756687 | 5.609021 | 5.588579 | 5.435047 |
| 5.33789 | NaN | NaN | NaN | NaN | 5.934573 | NaN | NaN | 6.033826 |
| 8.899517 | 8.922499 | 9.007861 | 7.535638 | 7.506926 | 8.545633 | 8.678923 | 8.491476 | 9.172509 |
| 8.020553 | 7.956793 | 7.649647 | 7.154584 | 7.589145 | 6.958959 | 7.004608 | 7.05574 | 7.045805 |
| 8.221428 | 8.164463 | 8.042615 | 9.439852 | 9.569875 | 8.10383 | 7.896715 | 7.851424 | 8.33118 |
| 5.961143 | 5.117431 | 5.800523 | 5.418674 | 5.060117 | 5.746437 | 5.910423 | 5.93079 | 6.179098 |
| 6.816648 | 6.585311 | 6.714983 | 6.30461 | 6.472903 | 6.308857 | 6.131582 | 6.167452 | 6.59612 |
| 7.011193 | 7.075868 | 7.244856 | NaN | NaN | 6.559435 | 6.806355 | 6.705627 | 6.825795 |
| 8.587242 | 8.565717 | 8.524858 | 7.589196 | 7.890844 | 9.366838 | 9.101539 | 9.172141 | 8.894696 |
| 7.460669 | 7.342177 | 7.298857 | 6.855772 | 6.679317 | 6.895345 | 6.896154 | 6.681493 | 6.976453 |
| 6.417384 | 5.492323 | 6.19114 | 7.09407 | 8.349945 | 8.711816 | 8.443898 | 8.584563 | 6.915347 |
| 6.986185 | 7.164352 | 6.733503 | 6.750104 | 7.116067 | 7.041623 | 7.019207 | NaN | 7.120863 |
| NaN | NaN | 5.928589 | 7.024674 | 7.175386 | 6.985858 | 5.784141 | 6.628531 | 6.274208 |
| 9.960094 | 10.04575 | 10.20875 | 10.95619 | 11.19246 | 9.952628 | 9.984913 | 10.12856 | 9.533688 |
| 6.81525 | 6.371805 | 6.838375 | 6.374859 | 6.166151 | 7.398725 | 6.796948 | 6.999429 | 5.361498 |
| 8.518021 | 8.149352 | 8.532938 | 10.68361 | 10.68048 | 7.46521 | 7.739703 | 7.178147 | 7.408294 |
| 10.88181 | 10.57896 | 10.59639 | 10.87638 | 11.06708 | 10.06289 | 10.35518 | 10.42458 | 10.14986 |
| 9.504488 | 9.408663 | 9.577765 | 8.394728 | 9.282508 | 9.066681 | 9.050082 | 9.320792 | 8.678784 |
| 7.717937 | 7.61385 | 7.573539 | 7.290277 | 7.194204 | 6.688477 | 7.074987 | 6.829283 | 7.556489 |
| NaN | NaN | NaN | NaN | NaN | 6.17487 | NaN | NaN | NaN |
| 7.964059 | 7.859296 | 8.044312 | 7.887931 | 7.840885 | 7.942433 | 8.253229 | 7.930623 | 7.944068 |
| 7.50689 | 7.186388 | 7.606398 | 10.35287 | 7.913393 | 7.642084 | 7.833985 | 7.89788 | 7.512628 |
| 6.415559 | 6.731055 | 6.811205 | 5.960648 | 5.240404 | 6.740355 | 6.573247 | 7.091687 | 7.497153 |
| 8.05089 | 7.68287 | 8.025665 | 8.012816 | 7.542705 | 8.621975 | 8.625965 | 8.704285 | 7.157147 |
| 6.781992 | 5.446165 | 7.513046 | 5.859412 | 4.678003 | 6.986117 | 5.693603 | 6.153776 | 5.445134 |
| 9.086674 | 9.055579 | 9.071277 | 8.883705 | 8.703717 | 8.565382 | 8.482224 | 8.729066 | 8.07317 |
| 8.074506 | 7.913083 | 5.507547 | 6.325636 | 5.783348 | 6.552184 | 6.509103 | 6.13999 | 6.662553 |
| 6.876772 | 5.937301 | 5.164234 | 5.912602 | 6.378365 | 6.593823 | 6.597354 | 6.300715 | NaN |
| 7.903211 | 7.686893 | 7.375568 | 8.117021 | 7.941193 | 7.034961 | 7.513206 | 7.306478 | 7.848668 |
| 7.002398 | 6.935569 | 7.120373 | 7.389016 | 6.7547 | 7.405067 | 7.43704 | 7.427651 | 6.836669 |
| 7.71354 | 8.00542 | 8.029485 | 5.849003 | 6.110433 | 7.191383 | 7.737806 | 7.602423 | 7.473092 |
| 8.840291 | 8.921615 | 8.913769 | 9.840837 | 9.913411 | 8.74719 | 8.545955 | 8.706491 | 9.148075 |
| 7.008453 | 7.063673 | 7.007849 | 6.145384 | 6.839388 | 7.17283 | 6.930787 | 6.818485 | 6.889561 |
| 7.403188 | 7.337852 | 7.377232 | 7.495047 | 7.411145 | 7.21383 | 7.31488 | 7.243843 | 6.569229 |
| 9.285865 | 9.17297 | 9.167076 | 10.74078 | 10.70313 | 9.648722 | 9.743775 | 9.737302 | 9.06294 |
| 9.889137 | 8.784271 | 8.625818 | 7.7372 | NaN | NaN | 7.222149 | NaN | NaN |
| NaN | NaN | 5.657884 | NaN | 5.127833 | 5.305334 | NaN | 5.63416 | 6.261804 |
| 5.016626 | 4.768826 | 5.081104 | 6.851878 | 9.652441 | 5.22495 | 4.695293 | 3.228349 | NaN |
| 9.611373 | 9.348516 | 9.648756 | 10.47618 | 10.56561 | 8.443828 | 8.209926 | 8.203164 | 7.625485 |
| 6.975201 | 6.682544 | 6.666308 | 7.837212 | 8.220664 | 6.742377 | 6.594831 | 7.019503 | 7.21711 |
| 8.730603 | 8.552731 | 8.575927 | 9.327504 | 9.454379 | 8.194193 | 8.426651 | 8.270205 | 7.880434 |
| 11.27272 | 11.26376 | 11.32393 | 12.28968 | 12.32279 | 11.13593 | 11.34347 | 11.04186 | 10.63008 |

-Bruker Confidential-

| | | | | | | | | |
|---|---|---|---|---|---|---|---|---|
| 6.254159 | 4.796154 | 6.747095 | NaN | NaN | 6.054593 | 6.182545 | 6.213073 | 5.403514 |
| 5.767644 | 5.652331 | 6.021614 | 5.394217 | 4.967626 | 6.002527 | 5.596334 | 5.626448 | 6.291873 |
| 7.096468 | 6.814172 | 7.061598 | 9.546845 | 9.265941 | 6.599614 | 6.718422 | 6.572 | 6.78817 |
| 7.773187 | 8.14098 | 7.595196 | 7.30279 | 7.610135 | 7.548566 | 7.81264 | 6.371423 | 7.634989 |
| 9.148342 | 8.93738 | 8.785188 | 10.81462 | 10.75127 | 8.054455 | 7.947925 | 7.95504 | 6.245897 |
| 10.93932 | 10.9329 | 10.78918 | 10.51036 | 10.40382 | 11.35359 | 11.35343 | 11.37874 | 11.03983 |
| 8.240605 | 8.511707 | 8.515583 | 9.323668 | 8.899609 | 8.042309 | 8.120753 | 8.175558 | 8.102216 |
| 9.114222 | 8.925926 | 9.194453 | 9.365933 | 9.410533 | 8.642793 | 8.791662 | 8.757372 | 8.814027 |
| 7.338156 | 6.628231 | 7.170677 | 5.657548 | 6.512213 | 7.816274 | 7.587608 | 7.62258 | 7.477573 |
| NaN | NaN | 4.795846 | 5.611776 | 4.991298 | 6.364893 | 6.290572 | 5.859931 | 6.471436 |
| 12.67633 | 12.82354 | 12.83671 | 13.6026 | 13.59491 | 12.36385 | 12.35544 | 12.35146 | 12.31034 |
| 9.699969 | 9.849798 | 9.727933 | 7.790304 | 7.522048 | 7.584265 | 7.55556 | 7.211102 | 9.268262 |
| 6.915055 | 7.099802 | 6.947116 | 6.679681 | 6.708336 | 8.013655 | 8.153018 | 8.20117 | NaN |
| 7.786302 | 7.822265 | 7.618931 | 6.951388 | 6.691758 | 8.08136 | 7.871997 | 8.009248 | 8.043327 |
| 6.304851 | 6.701432 | 6.477494 | 6.087763 | 6.011492 | 5.791245 | 6.040818 | 6.410598 | 6.61245 |
| 8.158281 | 7.952007 | 7.654066 | 8.067451 | 7.779555 | 7.893203 | 8.140519 | 7.801069 | 8.526699 |
| NaN | NaN | NaN | NaN | NaN | NaN | NaN | NaN | 6.38201 |
| 6.599436 | 7.058358 | 7.274776 | 6.536453 | 6.497871 | 7.067109 | 6.800775 | 7.110349 | 7.351651 |
| 8.533023 | 8.297594 | 8.343411 | 7.184134 | 7.06894 | 8.201629 | 8.233615 | 8.025506 | 8.423145 |
| 8.055041 | 7.698054 | 7.787301 | 6.147834 | 6.056482 | 7.815795 | 7.746032 | 7.578749 | 7.486986 |
| 7.135321 | 7.846796 | 7.612564 | NaN | NaN | 6.436749 | 6.495701 | 6.672983 | 6.470144 |
| 7.686284 | 7.719065 | 7.916341 | 9.659708 | 9.75028 | 7.875067 | 7.800803 | 8.147443 | 7.14934 |
| 9.347626 | 8.961793 | 9.056573 | 8.59971 | 8.643171 | 8.588596 | 8.844794 | 8.857021 | 8.762829 |
| 12.96317 | 12.76198 | 12.85998 | 12.5202 | 12.64873 | 12.84007 | 12.92704 | 12.92093 | 12.1809 |
| 6.161742 | 6.062321 | 6.235995 | 6.770052 | 5.953873 | 6.543124 | 6.930164 | 6.548693 | 5.994394 |
| 8.963668 | 8.840205 | 8.99495 | 8.164735 | 7.829984 | 8.792109 | 8.527761 | 8.215561 | 8.068805 |
| 6.300489 | 6.817196 | 6.568319 | 6.592507 | 6.863267 | 6.077224 | 7.433988 | 6.31331 | 6.933658 |
| 8.96485 | 7.870332 | 9.124979 | 9.367718 | 9.128441 | 7.202675 | NaN | NaN | 6.860498 |
| 9.675041 | 7.488325 | 8.036964 | 6.961391 | 6.888161 | NaN | NaN | NaN | NaN |
| 7.895782 | 7.894191 | 8.377927 | 6.091042 | 5.998254 | 7.959591 | 7.865621 | 7.96598 | 6.644009 |
| 8.232836 | 8.135466 | 8.224787 | 7.089042 | 7.464521 | 7.944986 | 7.728051 | 8.095018 | 6.85878 |
| 7.979729 | NaN | NaN | 6.413501 | 6.223954 | 5.833372 | 6.253956 | 6.178187 | 6.232514 |
| 7.531231 | 6.651441 | 7.608069 | 8.524124 | 8.180944 | 5.669314 | 6.782517 | 6.112144 | 5.125866 |
| 10.79336 | 10.42236 | 10.77537 | 10.96807 | 10.69669 | 10.97065 | 11.22813 | 11.01132 | 10.24251 |
| 8.882819 | 8.729402 | 8.981011 | 8.846515 | 8.824828 | 8.375658 | 8.186438 | 8.097704 | 8.131155 |
| 6.669808 | 6.741828 | 7.183655 | 7.156955 | 7.449219 | 7.382382 | 6.796913 | 6.520501 | 6.361722 |
| 12.24489 | 11.99954 | 11.90738 | 12.7467 | 12.82626 | 11.73409 | 11.75183 | 11.87514 | 11.97805 |
| NaN | NaN | NaN | 6.224718 | 8.139369 | 5.470381 | NaN | NaN | 5.694571 |
| 10.97106 | 10.95122 | 10.88591 | 10.26353 | 10.16697 | 10.97742 | 10.90272 | 11.04711 | 10.59964 |
| NaN | NaN | NaN | 6.759921 | 6.838555 | NaN | 5.623843 | NaN | 5.001585 |
| 8.487734 | 7.921597 | 9.204649 | 10.07847 | 9.957656 | 5.92761 | NaN | 7.177157 | 9.780053 |
| 9.624929 | 9.471661 | 9.530243 | 10.91037 | 9.76984 | 9.428705 | 9.372116 | 9.508191 | 9.739614 |
| 7.198897 | 7.178331 | 7.190704 | 5.70152 | 6.212546 | 7.876409 | 7.674426 | 7.864444 | 6.595523 |
| 8.046555 | 8.204603 | 7.989834 | 8.480491 | 8.660511 | 7.855884 | 8.141557 | 8.095862 | 7.599105 |
| 8.111446 | 8.693024 | 8.216483 | 6.487952 | 7.077617 | 7.43264 | 7.387906 | 7.146772 | 7.829127 |
| 7.771063 | 7.884602 | 7.954311 | 7.874528 | 7.783419 | 7.223474 | 6.73146 | 6.605144 | 7.082528 |
| 5.886837 | 5.489247 | 6.25187 | 6.291975 | 6.440017 | 7.046127 | 6.449553 | 6.735723 | 5.792783 |

| | | | | | | | | |
|---|---|---|---|---|---|---|---|---|
| 10.17703 | 10.32798 | 10.53378 | 10.91178 | 10.68613 | 9.462638 | 9.486745 | 9.464616 | 9.487879 |
| 9.559351 | 9.636829 | 9.503166 | 10.19638 | 10.21278 | 9.684118 | 9.529187 | 9.506715 | 8.888214 |
| 6.655506 | 6.480339 | 7.156779 | 6.421257 | 7.89556 | 7.968966 | 9.191274 | 7.583326 | 7.140399 |
| 9.185163 | 8.871155 | 8.843442 | 10.51773 | 10.3167 | 9.419452 | 9.547364 | 9.589564 | 9.62015 |
| 9.762783 | 9.678082 | 9.721373 | 9.754302 | 9.959141 | 9.784697 | 9.613103 | 9.805872 | 8.18704 |
| 7.019414 | 7.2162 | 7.000347 | 5.637679 | 7.153826 | 7.098288 | 7.179653 | 7.043102 | 7.921209 |
| 6.886549 | 5.76085 | 6.801486 | 5.230632 | 4.456174 | 6.382617 | 6.418494 | 6.211174 | 5.706838 |
| 8.169855 | 8.224819 | 8.33465 | 9.363441 | 9.14743 | 8.725353 | 8.529986 | 8.748549 | 7.646699 |
| NaN | 5.260233 | 5.426872 | NaN | 4.165277 | 6.426975 | 6.595708 | 6.507755 | 4.475837 |
| 6.015568 | 6.01832 | 5.906401 | 6.198966 | 6.238969 | 6.094144 | 5.50186 | 5.848197 | 5.655342 |
| 7.514208 | 7.465543 | 7.710519 | 6.177214 | 5.50125 | 7.125955 | 7.072603 | 6.767243 | 7.293772 |
| 6.614269 | 6.618517 | 6.660553 | 7.773729 | 7.970773 | 7.946991 | 7.910833 | 7.751305 | 7.510058 |
| 7.843241 | 6.949909 | 8.087981 | 7.2822 | 7.25165 | 8.52337 | 8.568023 | 8.551835 | 7.616114 |
| 9.349407 | 9.493605 | 9.413939 | 11.01963 | 10.72244 | 8.444614 | 8.443101 | 8.104236 | 8.992851 |
| 7.567587 | 7.45964 | 7.999886 | 8.866612 | 8.958962 | 7.944786 | 7.891069 | 8.03175 | 7.841938 |
| 8.611154 | 8.245404 | 8.558414 | 8.591771 | 8.383707 | 8.507341 | 8.489548 | 8.435043 | 8.557448 |
| 8.02553 | 7.210342 | 7.736683 | 7.781386 | 8.260501 | 7.661591 | 7.417005 | NaN | 8.123896 |
| 11.00122 | 10.97123 | 11.00211 | 11.1077 | 11.12499 | 10.8118 | 10.85002 | 10.84792 | 10.10302 |
| 5.565027 | 3.405315 | 3.91147 | 5.89832 | 6.219203 | 4.777017 | 6.323682 | 6.59105 | 4.977674 |
| 5.399168 | 4.988464 | 4.70818 | NaN | 6.196024 | NaN | NaN | NaN | 4.350793 |
| 5.753278 | 4.852049 | 6.116787 | 5.334733 | 6.082705 | 6.275446 | 6.93349 | 6.947273 | 5.891965 |
| 8.390579 | 8.265518 | 8.332335 | 7.606314 | 7.613757 | 7.674176 | 8.022146 | 8.465425 | 8.042489 |
| 10.83943 | 10.79315 | 10.66971 | 9.798727 | 9.781501 | 10.57719 | 10.38372 | 10.66225 | 10.52436 |
| 7.375177 | 7.187119 | 6.891444 | 5.532533 | 5.877542 | 7.499113 | 7.536758 | 8.677411 | 5.755946 |
| 7.749078 | 7.771422 | 7.580352 | 7.118266 | 6.280132 | 7.737441 | 7.906157 | 7.979956 | 6.421794 |
| 4.189938 | 4.961405 | NaN | 2.62062 | NaN | 5.670348 | 5.487387 | 5.310143 | 5.145255 |
| 7.871153 | 7.392412 | 7.625378 | 5.468848 | 5.514163 | 6.845744 | 7.090545 | 6.843072 | 7.457609 |
| 5.520263 | 4.552715 | 3.180891 | 4.022114 | 4.947891 | 5.413173 | NaN | NaN | 5.782047 |
| 5.152663 | 5.188768 | 4.611227 | 2.374477 | 3.95783 | 5.446226 | 5.058074 | NaN | 5.063959 |
| 8.011584 | 8.08259 | 7.989547 | 7.727976 | 7.885581 | 8.058211 | 7.87536 | 7.985363 | 7.30739 |
| 6.336072 | 5.943715 | 6.108548 | 6.562068 | 6.792561 | 7.062885 | 7.334639 | 7.216574 | 5.183752 |
| 8.206054 | 8.299082 | 8.222768 | 6.87308 | 6.424641 | 8.166435 | 8.650759 | 8.604744 | 8.37956 |
| NaN | 4.951472 | NaN | 6.62964 | 6.386764 | 5.806604 | 5.256441 | 5.652117 | 5.240513 |
| 6.891195 | 5.80144 | 6.41469 | NaN | NaN | 6.177898 | 5.303318 | 5.767692 | 6.281676 |
| 5.726789 | 5.577287 | NaN | NaN | NaN | NaN | NaN | NaN | 5.25702 |
| 7.569895 | 7.404934 | 6.942406 | 6.43617 | 6.89391 | 7.116349 | 6.863871 | 7.360643 | 6.864855 |
| 7.460119 | 7.043589 | 7.296493 | 6.154885 | 6.247916 | 7.232131 | 6.466311 | 6.741272 | 7.157004 |
| 5.824292 | 5.332911 | 6.006354 | NaN | NaN | NaN | 4.805072 | NaN | 6.886142 |
| 3.005017 | 4.474496 | 5.202074 | NaN | NaN | 4.496178 | 4.681405 | 4.396066 | 4.916526 |
| 5.913815 | 6.022559 | 6.228748 | 5.843751 | 6.300172 | 5.778189 | 6.086049 | 6.204544 | 6.68845 |
| 10.00164 | 9.848803 | 9.617806 | 8.496108 | 8.759278 | 8.784563 | 9.330359 | 9.060166 | 9.327548 |
| 6.184507 | 5.96401 | 5.750007 | NaN | NaN | NaN | NaN | NaN | 5.505998 |
| NaN | 3.975677 | 1.442695 | NaN | NaN | NaN | NaN | NaN | 4.406601 |
| 5.235439 | NaN | 4.545426 | NaN | NaN | 5.857521 | 6.42865 | 5.823362 | 5.890602 |
| 6.341373 | 6.0971 | 5.482583 | NaN | NaN | 5.959614 | 5.778124 | 6.07546 | 5.211473 |
| 7.496699 | 7.363284 | 7.216592 | 7.02208 | 7.048218 | 7.657322 | 7.600126 | 7.667903 | 7.691044 |
| 8.101842 | 8.329191 | 8.093085 | 8.728431 | 8.258296 | 6.732553 | 6.485054 | 6.175954 | 7.380119 |

| | | | | | | | | |
|---|---|---|---|---|---|---|---|---|
| 6.636247 | 7.272904 | 6.479974 | 6.440033 | 5.857715 | 7.773159 | 7.236727 | 7.662072 | 5.714756 |
| 7.925876 | 6.63614 | 7.749607 | 6.742942 | 6.866498 | 7.769854 | 7.518067 | 7.338115 | 7.959496 |
| 8.377855 | 8.130763 | 8.474387 | 7.593266 | 7.535683 | 7.349807 | 7.542176 | 7.400152 | 7.243404 |
| 6.084673 | 6.440785 | 6.118601 | 5.510003 | 5.602427 | 5.795243 | 6.048756 | 5.82688 | 6.895573 |
| 6.330353 | 5.023763 | 6.290397 | NaN | NaN | 5.975436 | 6.225557 | 6.391697 | 5.958162 |
| 4.498091 | 5.260731 | 5.343954 | NaN | 3.493991 | 5.120742 | 5.226107 | 5.368791 | 4.673707 |
| 9.124237 | 9.334948 | 9.384996 | 10.09654 | 10.14195 | 9.246138 | 9.353095 | 9.195477 | 8.488545 |
| NaN | NaN | NaN | NaN | NaN | NaN | 6.939291 | 6.768475 | 5.226358 |
| 7.683307 | 8.013614 | 7.917329 | 8.181946 | 8.19386 | 8.681982 | 8.802135 | 8.589318 | 9.055819 |
| 9.582254 | 9.458451 | 8.857183 | 9.291679 | 9.341855 | 8.232347 | 8.321685 | 8.793626 | 8.885942 |
| 10.26843 | 10.25565 | 10.37849 | 8.461188 | 8.029034 | 9.340091 | 9.241902 | 9.459712 | 8.961628 |
| 7.418139 | 7.290839 | 7.480285 | 4.484528 | NaN | NaN | NaN | NaN | NaN |
| 7.233989 | 7.360162 | 7.289401 | 6.850528 | 6.990984 | 7.776134 | 7.666066 | 7.766306 | 6.811805 |
| 7.910968 | 8.028612 | 8.070255 | 8.5707 | 8.548792 | 8.030371 | 7.498871 | 7.780681 | 7.544381 |
| 9.056795 | 6.70557 | 7.564543 | 6.348508 | 7.513523 | NaN | 7.203093 | 7.132317 | 7.132195 |
| 8.186021 | 9.465951 | 9.391358 | 12.89916 | 14.2016 | 8.067565 | 8.19366 | 8.625333 | 9.298017 |
| 7.741832 | 7.987802 | 7.739945 | 7.840239 | 7.411312 | 7.269796 | 7.477007 | 7.542885 | 7.756669 |
| 7.169225 | 7.602631 | 7.459252 | 7.900961 | 8.48816 | 7.153109 | 7.685251 | 7.680955 | 6.976735 |
| 6.164334 | 5.340909 | NaN | 5.794842 | 5.495786 | 6.719064 | 7.108943 | 7.150307 | 6.30227 |
| 4.718732 | 6.161736 | NaN | 8.215213 | 8.035198 | 7.713964 | 8.036551 | 7.665423 | 7.068586 |
| 6.667498 | 6.61407 | 6.823962 | 6.344009 | 6.581825 | 7.449277 | 7.813382 | 7.612097 | 7.181458 |
| 4.584867 | 5.158831 | 5.047176 | NaN | NaN | 4.958782 | 3.830734 | 5.266209 | 4.99979 |
| NaN | 5.88647 | 6.239569 | NaN | 4.156856 | 6.812008 | NaN | 6.457424 | 5.402874 |
| 9.19556 | 9.15698 | 9.225915 | 8.871576 | 8.863069 | 9.019529 | 9.004139 | 8.944692 | 8.235328 |
| 6.089388 | 5.610353 | 4.90432 | 3.981644 | 3.873592 | 6.642393 | 6.333906 | 6.683459 | 4.480708 |
| 7.24421 | 7.194517 | 7.219728 | 6.959095 | 6.59673 | 7.10708 | 7.064551 | 7.050756 | 7.063888 |
| 9.35403 | 9.407455 | 9.461714 | 9.213003 | 8.777205 | 9.134499 | 8.759096 | 8.709775 | 8.195177 |
| 6.758191 | 6.292911 | 6.77137 | 5.845715 | 6.721115 | 7.783075 | 8.468146 | 8.362752 | 6.296342 |
| 5.569475 | 7.414371 | 6.842379 | NaN | 3.417122 | 5.680746 | 5.22346 | 5.715099 | 4.65681 |
| 6.936016 | 6.493737 | 6.854425 | 5.811543 | 5.908246 | 6.375381 | 6.731609 | 6.749969 | 6.148129 |
| 7.934276 | 8.458777 | 8.172084 | 7.172612 | 7.586523 | 7.842122 | 8.367764 | 8.262884 | 8.393254 |
| 7.217555 | 7.152787 | 7.043657 | 8.421675 | 8.708628 | 6.950904 | 6.941306 | 7.362931 | 6.812241 |
| 8.090181 | 6.611703 | 6.197853 | 6.277381 | 7.759047 | 7.737132 | 7.133373 | 6.982821 | 7.916574 |
| 6.504918 | 6.163738 | 6.058339 | 6.299175 | 6.230151 | 6.384886 | 6.312542 | 6.311993 | 4.956751 |
| 7.827175 | 7.809344 | 7.48912 | 8.220323 | 8.167305 | 8.09654 | 8.060273 | 7.952248 | 8.002138 |
| NaN | NaN | 6.115523 | 6.270311 | 6.23608 | 4.823642 | NaN | NaN | 4.437579 |
| 7.649935 | 7.248256 | 7.405467 | 7.067297 | 7.227462 | 6.71478 | 6.204675 | 5.918072 | 5.951104 |
| NaN | NaN | NaN | 5.476131 | 5.499567 | NaN | NaN | 7.241889 | 6.49159 |
| 5.833842 | 5.477524 | 5.826817 | 5.222469 | NaN | NaN | 5.550785 | 5.304758 | 6.092229 |
| 5.218703 | 6.109127 | 5.536355 | NaN | NaN | 6.183305 | 6.474827 | 5.931872 | 6.537408 |
| 9.359118 | 9.239309 | 9.320761 | 8.989052 | 8.795424 | 9.600643 | 9.384753 | 9.373914 | 9.128939 |
| 4.221269 | 3.941414 | NaN | 5.996392 | 5.958568 | NaN | NaN | NaN | NaN |
| 7.492773 | 7.292879 | 7.556627 | 5.944517 | 5.328973 | 7.351127 | 7.418807 | 7.628712 | 7.704194 |
| NaN | NaN | 5.452536 | NaN | NaN | 6.365378 | 6.163737 | 5.732223 | 6.187057 |
| 5.905171 | 5.208385 | 5.390045 | 6.616272 | 6.459347 | NaN | NaN | NaN | 5.106222 |
| 6.343916 | 6.717694 | 6.526079 | 7.468149 | 7.292096 | 6.81823 | NaN | NaN | 6.639734 |
| 9.805818 | 10.36693 | 9.972099 | 8.945617 | 7.90538 | 9.675638 | 9.356779 | 9.783813 | 8.506039 |

| | | | | | | | | |
|---|---|---|---|---|---|---|---|---|
| 9.03395 | 8.565313 | 8.619415 | 9.281188 | 9.048718 | 9.017974 | 9.147865 | 9.0444 | 9.400139 |
| 7.295785 | 6.903483 | 7.346542 | 10.648 | 10.82719 | 7.969676 | 7.913455 | 8.020729 | 6.918668 |
| 5.976412 | 6.51623 | 6.057437 | 6.545648 | 6.545795 | 6.515847 | 6.431764 | 7.072482 | 6.060137 |
| 8.186994 | 8.194802 | 8.181273 | 7.924137 | 7.822731 | 8.957343 | 8.814333 | 8.776649 | 7.835976 |
| 5.516413 | 6.084166 | 5.896486 | 5.188081 | 5.290839 | 5.815647 | 5.702554 | 5.791308 | 5.853075 |
| 7.687551 | 7.362155 | 7.496661 | 6.304768 | 6.52531 | 7.731643 | 7.450185 | 7.447648 | 7.32402 |
| 7.280979 | 7.125959 | 6.099623 | NaN | 4.984866 | 7.372085 | 7.400584 | 7.4757 | 6.764469 |
| 7.355688 | 7.460172 | 7.237255 | 7.081422 | 7.39841 | 7.645118 | 7.739062 | 7.983126 | 6.978034 |
| 6.786064 | 6.680935 | 7.329164 | 6.261219 | 6.141181 | 6.904187 | 6.718187 | 6.780892 | 6.254575 |
| 6.933408 | 7.053634 | 6.839982 | 8.235137 | 8.323133 | 7.21791 | 7.234292 | 7.412934 | 7.486231 |
| 7.348684 | 7.160008 | 7.847855 | NaN | 7.01047 | 7.858911 | 7.708808 | 8.139997 | 7.598078 |
| 7.402716 | 6.544286 | 6.728022 | 7.655396 | 7.248933 | NaN | 6.666717 | 6.32384 | 6.856441 |
| 7.671659 | 7.511859 | 7.333166 | 8.030885 | 8.082272 | 7.628647 | 7.505481 | 7.530249 | 7.323455 |
| 7.081956 | 6.968774 | 7.116277 | 5.433821 | 5.700191 | 8.118831 | 8.096771 | 7.471657 | 5.994463 |
| 7.649116 | 7.153691 | 6.608218 | 7.460333 | 7.274129 | 6.644725 | 6.932917 | 7.120491 | 6.932405 |
| 6.2936 | 6.088915 | 5.313805 | 5.609552 | 6.387565 | NaN | 5.354962 | 5.520382 | NaN |
| 7.318795 | 7.286199 | 5.969603 | NaN | 5.678133 | NaN | NaN | NaN | 6.094963 |
| 10.02493 | 10.05787 | 9.854982 | 9.595328 | 9.382005 | 9.216562 | 9.228725 | 9.250764 | 8.956882 |
| NaN | NaN | NaN | 6.589164 | 5.675848 | NaN | NaN | 5.63717 | 4.969281 |
| NaN | 5.757897 | NaN | 5.958313 | 5.903685 | NaN | NaN | 5.592352 | 6.716398 |
| 7.036045 | 6.759589 | 7.083272 | 7.363765 | 7.419142 | 6.710275 | 6.268445 | 6.385305 | 6.588339 |
| 4.639461 | 4.499581 | NaN | NaN | 3.465979 | 5.224164 | 5.627887 | 5.278682 | 5.068049 |
| 12.84079 | 12.52914 | 12.99373 | 12.21884 | 12.19458 | 11.77412 | 12.51134 | 11.97952 | 12.52389 |
| 8.115371 | 8.087767 | 8.138971 | 7.252551 | 6.954538 | 8.130284 | 7.796432 | 8.32348 | 7.569847 |
| 6.081497 | 6.677479 | 6.590323 | NaN | 5.922777 | 5.446154 | 6.062246 | 5.554735 | 6.537258 |
| 6.867455 | 6.749927 | 6.75841 | NaN | 6.051221 | 7.553826 | 7.10103 | 6.864798 | 5.969734 |
| 6.392254 | 6.515715 | 6.568664 | 6.202823 | 6.580279 | 6.895393 | 7.003144 | 7.11612 | 7.053601 |
| 7.975533 | 7.771929 | 8.233488 | 8.67042 | 8.932878 | 7.973683 | 7.22503 | 6.724707 | 7.734151 |
| 9.098794 | 8.940794 | 9.436926 | 9.171574 | 9.493775 | 10.38642 | 9.580144 | 10.25645 | 9.230665 |
| 8.837808 | 10.34655 | 10.10913 | 12.9817 | 12.27829 | 14.73253 | 14.47108 | 14.57398 | 12.05107 |
| 8.745742 | 7.017488 | 8.351966 | 8.195714 | 7.58871 | 7.80268 | 7.886772 | 7.812802 | 8.886586 |
| 6.93786 | 7.037934 | 7.186755 | 5.967505 | 6.51293 | 7.431406 | 6.810266 | 6.845325 | 6.441996 |
| 6.180301 | 6.276917 | 6.018892 | 4.824741 | NaN | 6.314218 | 6.720809 | 6.617827 | 5.452962 |
| 7.838013 | 7.140669 | 7.291608 | 6.428408 | 6.789461 | 7.176878 | 8.033084 | 7.693024 | 8.31288 |
| 4.936858 | 5.755882 | 4.922315 | NaN | NaN | 6.140144 | 5.716284 | 4.856354 | 5.886367 |
| 6.416073 | 5.980186 | 6.307204 | NaN | 6.418735 | 8.139108 | 8.228937 | 7.730783 | 6.376812 |
| 7.235816 | 7.529423 | 7.407602 | NaN | 6.490402 | 7.015563 | 5.883214 | 6.45062 | 5.963896 |
| 6.770063 | 6.664912 | 5.540205 | 5.591323 | NaN | 6.7564 | NaN | NaN | 6.725796 |
| 7.196337 | 7.41201 | 7.759748 | 7.011667 | 7.173075 | 6.552872 | 7.120191 | 6.791406 | 6.801102 |
| 8.219509 | 8.46025 | 8.189503 | 7.672899 | 7.723039 | 7.720841 | 7.911259 | 7.805302 | 7.992543 |
| 8.066898 | 8.103773 | 8.291766 | 7.138595 | 6.668471 | 5.438206 | 6.079018 | 6.287756 | 6.20651 |
| 8.572002 | 8.642396 | 8.445107 | 7.183373 | 7.613508 | 8.771999 | 8.877387 | 8.862008 | 7.33391 |
| 7.081472 | 7.155635 | 7.260499 | 7.113015 | 7.176518 | 5.994357 | 5.508243 | 6.060472 | 6.851715 |
| 7.302629 | 7.046616 | 7.102204 | 8.552714 | 8.548884 | 6.723857 | 6.597271 | 7.101744 | 7.966957 |
| 6.380224 | 5.213207 | 4.392728 | NaN | NaN | 7.082559 | 6.584766 | 7.185662 | 6.211134 |
| 7.380103 | 7.027838 | 7.194996 | 5.504357 | NaN | 8.075738 | 7.94679 | 7.963413 | 6.991871 |
| 6.907505 | 6.600039 | 7.047 | 5.590281 | 6.83063 | 6.901457 | 7.188379 | 6.93224 | 6.512416 |

| | | | | | | | | |
|---|---|---|---|---|---|---|---|---|
| 0.088654 | 1.277311 | NaN | 6.030233 | 4.925586 | NaN | NaN | NaN | 2.377926 |
| 7.24525 | 7.280365 | 7.144809 | 6.857659 | 6.448625 | 6.981841 | 7.062329 | 7.159372 | 7.420296 |
| 7.161608 | 6.467158 | 7.096064 | 6.894293 | 6.790203 | 7.803648 | 6.469453 | 7.714183 | 6.178213 |
| 6.948837 | 6.531508 | 7.05405 | 7.953202 | 7.309723 | 7.70205 | 7.284056 | 7.595389 | 7.449344 |
| 6.689116 | 6.588668 | 6.876163 | 7.022289 | 7.213254 | 7.438785 | 7.090304 | 7.098152 | 7.479541 |
| 6.290808 | 3.700599 | NaN | 6.891034 | 6.879859 | 6.220289 | 6.351521 | 5.932237 | 5.567924 |
| 7.062923 | 7.077827 | 6.986625 | 5.816265 | 5.592128 | 7.024458 | 6.756519 | 5.726922 | 6.832767 |
| 6.98678 | 6.78489 | 7.258298 | 6.502007 | 5.871459 | 7.273125 | 7.271256 | 7.462656 | 7.398285 |
| 5.948416 | 7.140726 | 5.843193 | NaN | 4.685758 | 6.099159 | 5.61687 | 6.142781 | 5.509744 |
| 5.111178 | 6.379818 | 6.809172 | 3.799935 | 4.448521 | 5.605889 | 5.924397 | 6.114054 | 4.946117 |
| 6.173436 | 5.355904 | NaN | 5.690546 | NaN | 5.807388 | 6.214294 | NaN | 5.875258 |
| 8.534171 | 8.549857 | 8.516878 | 8.266139 | 8.300361 | 8.638716 | 8.56986 | 8.566035 | 8.692401 |
| 7.699827 | 7.692183 | 7.705698 | 7.204515 | 7.788084 | 7.515372 | 7.753213 | 7.785297 | 8.0613 |
| 6.631763 | 6.042375 | 5.974164 | 8.24134 | 8.313445 | 5.795403 | 4.709285 | 6.27132 | 5.862824 |
| 5.230731 | 5.094257 | 5.436059 | 5.883553 | 5.698476 | 4.879136 | 5.125831 | 4.852698 | 5.390004 |
| 8.167674 | 8.377669 | 8.333665 | 8.66642 | 8.776546 | 8.307465 | 7.991117 | 8.389433 | 7.990843 |
| NaN | NaN | 5.776419 | 4.815024 | 6.23707 | NaN | 5.271866 | NaN | 5.89523 |
| 6.776316 | 6.853621 | 6.970353 | 5.095692 | 5.543497 | 6.58967 | 6.507579 | 6.903561 | 6.707134 |
| 7.265508 | 8.128284 | 7.958035 | 9.719534 | 10.03541 | 6.887192 | 6.514642 | 6.640119 | 7.099749 |
| 7.247308 | 7.093112 | 6.804837 | 7.780423 | 7.747241 | 6.818506 | 6.33053 | 6.990196 | 7.505617 |
| 7.243255 | 7.440879 | 7.311926 | 6.248683 | 6.698565 | 7.425473 | 7.274227 | 7.400274 | 6.827966 |
| 7.302315 | 6.817811 | 6.775078 | 6.500364 | 6.056809 | 7.033126 | 6.774449 | 7.009837 | 6.328142 |
| 6.142088 | 7.758721 | NaN | 6.550337 | 5.811692 | 5.605392 | 5.700339 | 5.572587 | NaN |
| 7.10607 | NaN | 6.924625 | 6.140595 | 6.062234 | 6.460468 | NaN | NaN | 5.590983 |
| 6.154282 | 5.850296 | 5.310046 | 4.345719 | NaN | 5.146124 | 5.466098 | 4.877042 | 5.264594 |
| 6.302535 | 6.512533 | 6.936995 | 7.274914 | 8.090716 | 7.411208 | 7.061767 | 6.767686 | 7.283024 |
| 6.383596 | 5.817862 | 6.729466 | 5.373714 | 4.634329 | 7.168195 | 7.317218 | 6.921065 | 6.471417 |
| NaN | NaN | NaN | NaN | NaN | NaN | NaN | NaN | 7.183218 |
| NaN | NaN | NaN | 6.829286 | 6.548245 | NaN | NaN | NaN | 4.662401 |
| 6.988684 | NaN | 7.839481 | 7.154086 | 6.794447 | NaN | 6.31973 | 6.099356 | 7.04173 |
| 6.135827 | 5.790323 | 6.007112 | 5.643338 | 4.67821 | 6.135576 | 6.024891 | 5.983484 | 5.081847 |
| 5.811183 | 6.35556 | 6.170873 | 6.838746 | 7.123838 | 6.844348 | 6.604217 | 6.640553 | 6.687288 |
| 5.453822 | 6.062465 | NaN | 6.843549 | 6.784649 | 7.132155 | 7.216467 | 7.079695 | 6.865046 |
| 7.039766 | 7.181475 | 7.281406 | 6.900517 | 6.445519 | 7.185918 | 7.053961 | 6.989868 | 7.071772 |
| 6.097709 | 6.165782 | 5.835344 | 5.995817 | 5.901193 | 5.833081 | 5.576255 | 5.88328 | 5.767943 |
| 6.626412 | 6.443386 | 6.412724 | 6.069717 | 5.726893 | 7.40746 | 7.074109 | 7.297696 | 6.330941 |
| 5.076396 | 2.91834 | 4.156434 | 4.505942 | 4.410017 | 5.099982 | 5.144948 | 5.292254 | 4.064666 |
| 7.015894 | 6.906033 | 7.103357 | 6.358764 | 6.045901 | 5.387808 | 4.860717 | 5.667704 | 6.911189 |
| 8.337651 | 8.312427 | 8.340016 | 7.513271 | 7.084812 | 7.397881 | 7.949764 | 7.412404 | 6.326518 |
| 8.07894 | 7.840182 | 7.96337 | 6.525425 | 6.799078 | 8.054672 | 7.815796 | 7.667518 | 7.600008 |
| 8.62043 | 8.336658 | 8.098567 | 8.695315 | 8.674474 | 7.621675 | 8.072744 | 7.94751 | 7.673161 |
| 9.417474 | 8.827679 | 8.62242 | 8.238883 | 8.732862 | 8.179665 | 8.136957 | 8.271347 | 7.636083 |
| 4.37153 | 4.458913 | 5.543685 | NaN | NaN | 5.947018 | 4.999689 | 5.965827 | 3.754889 |
| 11.46753 | 11.57553 | 11.62148 | 11.30618 | 11.20335 | 11.43405 | 11.57022 | 11.47664 | 10.4071 |
| 6.837334 | 6.817019 | 6.581233 | 5.326777 | 5.547401 | 6.748919 | 6.660593 | 6.937664 | 6.910807 |
| 10.00856 | 9.810362 | 9.832347 | 9.272231 | 9.232382 | 10.48054 | 10.45851 | 10.42549 | 9.210752 |
| 7.119398 | NaN | 4.954935 | NaN | NaN | 6.747753 | 6.078755 | NaN | 6.185733 |

-Bruker Confidential-

| | | | | | | | | |
|---|---|---|---|---|---|---|---|---|
| 4.181854 | 4.62263 | 5.261935 | NaN | NaN | NaN | NaN | NaN | 4.494491 |
| 9.595963 | 9.71647 | 9.647592 | 10.75917 | 10.84345 | 9.69987 | 9.530119 | 9.768188 | 10.09382 |
| NaN | NaN | NaN | 4.918478 | 6.023066 | NaN | NaN | NaN | 3.83096 |
| 6.489318 | 5.359869 | 6.273293 | 6.277833 | 6.48121 | 6.522496 | 6.263692 | 6.250693 | 6.557043 |
| 7.052989 | 5.690676 | 6.356663 | 9.504244 | 9.05347 | 6.084567 | NaN | NaN | 6.356977 |
| 6.681304 | 5.936639 | 6.74577 | 5.041066 | 6.132343 | 6.42753 | 6.08215 | 5.956086 | 5.689552 |
| 6.789997 | 6.911949 | 6.912057 | 6.875452 | 7.167984 | 8.338486 | 7.933671 | 7.079875 | 6.648012 |
| 5.321872 | 5.043553 | NaN | 5.591288 | 5.109044 | 4.549338 | NaN | 5.19491 | NaN |
| 9.201865 | 8.956306 | 9.173432 | 6.781344 | 6.853063 | 7.435863 | 7.342044 | 7.343266 | 7.643592 |
| 8.064616 | 7.65044 | 7.959099 | 7.293177 | 7.255971 | 7.154571 | 7.178797 | 7.164367 | 7.33207 |
| NaN | NaN | NaN | 4.932482 | 5.299026 | 2.653785 | NaN | 3.969171 | NaN |
| 5.979053 | 5.58241 | 5.81714 | 5.817844 | 6.083395 | 6.312372 | 6.397735 | 5.903842 | 5.194588 |
| 6.845579 | 6.898263 | 6.91475 | 5.947789 | 6.474514 | 7.824443 | 7.297072 | 7.321931 | 6.953697 |
| 9.241881 | 9.229868 | 9.304729 | 9.417951 | 9.399814 | 8.629162 | 8.617702 | 8.456038 | 8.578602 |
| 6.279852 | 5.461161 | 5.425399 | 5.561077 | 5.2767 | NaN | NaN | NaN | 5.371594 |
| NaN | NaN | NaN | 7.821385 | 8.073647 | 5.586773 | 3.828385 | 4.683876 | 4.699606 |
| 7.910181 | 7.958622 | 8.18056 | 7.872048 | 7.937272 | 8.427205 | 8.13072 | 8.221883 | 8.304287 |
| 6.835234 | 6.911925 | 7.146937 | 5.209761 | 5.843365 | 6.048518 | 4.836445 | 5.44987 | 6.495818 |
| 5.675523 | 6.398666 | 5.344088 | NaN | NaN | 7.404279 | 7.303467 | 7.379071 | 6.549073 |
| 6.057064 | 5.459764 | 5.350547 | 6.710409 | 6.105993 | 5.825571 | 6.587274 | 6.547763 | 5.500013 |
| 6.999413 | 6.448101 | 5.413303 | NaN | 6.193462 | 6.588629 | 6.726001 | 6.605844 | 5.978104 |
| 6.224547 | 4.550883 | NaN | NaN | 4.861222 | 5.501698 | 5.6892 | 5.553402 | 5.27348 |
| 8.578605 | 7.339347 | 7.327528 | NaN | 6.187911 | 7.827542 | 7.613362 | 7.393005 | 6.775348 |
| 6.460043 | 6.559294 | 6.603102 | 6.905055 | 7.222327 | 6.085594 | 6.062264 | 5.940935 | 6.864517 |
| 6.787151 | 6.18906 | 6.755272 | 7.782281 | 8.306448 | 6.776263 | 7.187657 | 6.836047 | 6.412379 |
| 6.050396 | 6.123424 | 6.080102 | NaN | 4.527781 | 7.099475 | 6.969548 | 7.007622 | 6.011323 |
| NaN | NaN | NaN | 5.958633 | NaN | 6.671209 | 6.140755 | 6.123714 | 5.938429 |
| 8.744054 | 8.740394 | 8.576383 | 7.490231 | 7.622576 | 9.082349 | 8.735475 | 8.736931 | 8.367869 |
| 8.166585 | 10.75815 | 7.406914 | 6.922509 | 6.754096 | NaN | 9.484613 | 9.179664 | 7.793495 |
| 7.822244 | 7.697762 | 7.669794 | 7.629994 | 7.477245 | 7.551682 | 7.425538 | 7.56073 | 7.743241 |
| NaN | NaN | NaN | NaN | NaN | NaN | NaN | NaN | 6.329587 |
| 6.673567 | 6.856987 | 7.046249 | 7.016698 | 7.25547 | 7.063859 | 7.210169 | 7.178285 | 7.550274 |
| NaN | 5.948067 | 4.828981 | 5.953226 | 6.10644 | 6.765945 | 6.583317 | 6.881721 | 6.175258 |
| 5.807363 | 4.299379 | 6.318615 | 5.779494 | 6.641835 | NaN | NaN | NaN | 6.948541 |
| 7.500847 | 7.972506 | 7.88925 | 7.750675 | 7.594187 | NaN | NaN | NaN | 6.596114 |
| 6.703623 | 6.793941 | 6.260371 | 5.21277 | 5.098404 | NaN | NaN | NaN | 5.949132 |
| NaN | NaN | NaN | 5.55032 | 6.067095 | NaN | NaN | NaN | NaN |
| 8.868287 | NaN | NaN | 6.8606 | 7.012302 | 6.842024 | NaN | 6.654778 | 7.018971 |
| 6.432333 | 6.356863 | 6.230705 | 6.749494 | 5.28141 | 6.610735 | 6.463694 | 6.606102 | 6.720809 |
| 8.116966 | 7.820722 | 7.847795 | 8.387159 | 8.37047 | 8.18404 | 8.250288 | 8.281442 | 7.916574 |
| 9.773784 | 9.637769 | 9.662162 | 10.10905 | 9.402303 | 8.826017 | 9.162099 | 9.124819 | 9.867951 |
| 7.852593 | 7.648602 | 8.004582 | 7.018252 | 6.427237 | 7.032151 | 5.928484 | 6.539617 | 5.274454 |
| 7.15701 | 6.498559 | 7.476512 | 5.694279 | 5.83419 | 5.98947 | 5.791149 | 6.581985 | 7.460307 |
| 7.757166 | 4.315702 | 7.897094 | 6.063025 | 6.087434 | 7.144475 | 7.274864 | 7.340128 | 6.99419 |
| 6.19796 | 6.56984 | 6.557641 | 6.527194 | 6.513899 | 6.563429 | 6.374301 | 6.7964 | 6.967412 |
| 5.305482 | 5.701067 | 5.466054 | 5.054846 | 4.444313 | 6.151274 | 5.985128 | 6.016263 | 6.065823 |
| 6.538354 | 7.739198 | 7.831959 | NaN | 5.27628 | 7.220016 | 7.433731 | 7.183504 | 6.738355 |

| | | | | | | | | |
|---|---|---|---|---|---|---|---|---|
| 5.11659 | 6.4554 | 3.271888 | 5.223247 | 5.955231 | NaN | 5.906535 | 5.217399 | 5.784896 |
| 5.474394 | 3.498932 | 4.630859 | 5.148898 | 5.610422 | 5.082496 | 4.962703 | 4.226254 | 5.441421 |
| 6.708886 | 6.299542 | 6.30363 | 7.502402 | 6.881782 | 5.915672 | 6.020239 | 5.594916 | 6.382611 |
| NaN | 5.683934 | 4.094903 | 6.021106 | 5.259324 | 6.042695 | 5.403235 | 6.146585 | 4.926026 |
| NaN | NaN | NaN | 6.429047 | 5.861051 | NaN | NaN | NaN | 6.3116 |
| 8.332489 | 8.388657 | 8.405658 | 9.221334 | 9.644586 | 9.559402 | 9.086684 | 9.54493 | 8.507025 |
| 9.420681 | 9.311364 | 9.366899 | 9.754593 | 9.194836 | 9.321651 | 9.343432 | 9.306667 | 8.86511 |
| NaN | NaN | 5.189803 | 3.93447 | 5.481811 | 5.93223 | 5.171914 | 5.243709 | 6.045937 |
| 5.956532 | 6.081275 | 6.001544 | 6.528059 | 6.477732 | 5.90015 | 6.009697 | 6.144926 | 6.44638 |
| NaN | NaN | NaN | 5.365499 | 5.390527 | NaN | 2.171228 | NaN | 3.842018 |
| 7.131096 | 5.889836 | 7.015175 | 5.767098 | 6.201265 | 7.174325 | 6.97159 | 6.910307 | 6.582308 |
| 6.759955 | 6.454876 | 6.31277 | 6.295895 | 7.429341 | 6.737658 | 6.16799 | 6.616197 | 5.944729 |
| 6.937788 | 6.893174 | 6.713633 | 6.637136 | 6.090187 | 6.752934 | 6.739532 | NaN | 7.659807 |
| 6.978573 | 7.057459 | 6.953572 | 6.178375 | 6.091308 | 6.336505 | 6.154533 | 6.296467 | 6.330484 |
| 5.746209 | 6.194469 | NaN | 6.603557 | 6.265798 | 6.049152 | 6.458874 | 6.484594 | 5.871902 |
| 8.942293 | 8.933931 | 8.761199 | 9.101184 | 8.992856 | 7.295494 | 7.511756 | 7.44346 | 7.668148 |
| NaN | NaN | NaN | 7.591575 | 8.23645 | NaN | NaN | NaN | 6.266314 |
| 7.510254 | 7.413178 | 7.510678 | 8.05265 | 7.147882 | 6.808095 | 7.181801 | 6.952543 | 7.084085 |
| 5.087576 | 5.42383 | 5.291424 | 4.428113 | 4.393446 | 5.225818 | 5.51173 | 5.338923 | 5.037903 |
| 5.716035 | 5.981993 | 5.76176 | NaN | 4.118893 | 7.609928 | 7.467258 | 7.010775 | 6.227263 |
| NaN | NaN | NaN | 6.366955 | 6.3383 | NaN | NaN | NaN | 6.186131 |
| 6.84165 | 6.833059 | 6.572027 | 6.201878 | 6.756021 | 7.347649 | 7.305758 | 7.360196 | 7.093708 |
| 6.99341 | 6.556847 | 6.755492 | 6.891443 | 7.173381 | 6.799513 | 5.527754 | 5.482873 | 5.897074 |
| 8.432605 | 8.45087 | 8.297471 | 6.160054 | 6.605288 | 7.726839 | 7.993356 | 7.758973 | 7.690892 |
| 9.02946 | 9.097018 | 9.078494 | 8.868246 | 8.890206 | 8.938474 | 8.833321 | 8.815582 | 8.974399 |
| 8.054638 | 7.944342 | 7.823435 | 8.167136 | 8.025215 | 7.709474 | 7.437472 | 7.81536 | 7.877585 |
| 6.172548 | 5.886082 | 5.126562 | 4.897491 | NaN | 6.838155 | 6.538286 | 7.445088 | 6.966557 |
| 5.890688 | 5.890088 | 6.214154 | 6.399564 | 6.896585 | 6.647679 | 6.900569 | 6.951668 | 6.856838 |
| 7.161141 | 6.861833 | 7.330204 | NaN | NaN | NaN | 6.58463 | NaN | 4.727688 |
| 6.422196 | 5.922734 | 6.792694 | 6.778215 | 7.534023 | 7.697041 | 7.824939 | 7.76083 | 7.05414 |
| NaN | NaN | NaN | 8.980496 | 8.793921 | NaN | 8.013983 | NaN | NaN |
| 5.752322 | 5.187307 | 5.465876 | 4.475193 | 4.553571 | 5.360251 | 5.30584 | 5.581769 | 6.038167 |
| 6.325595 | 5.645403 | 6.381189 | NaN | 6.303903 | 5.342147 | 5.913588 | 6.499135 | 5.714641 |
| 8.762002 | 8.714618 | 8.453609 | 6.790774 | 6.192147 | 8.387222 | 7.702969 | 7.74004 | 7.713036 |
| 6.357644 | 6.166274 | 5.715805 | 6.711295 | NaN | 6.548438 | 6.924679 | 6.811675 | 6.099964 |
| 8.633389 | 8.594217 | 8.608968 | 8.680569 | 9.079408 | 7.974706 | 8.43516 | 8.10701 | 8.020309 |
| 7.538526 | 7.214432 | 7.318698 | 7.212148 | 7.447875 | 7.366545 | 7.560717 | 7.276171 | 7.062135 |
| NaN | 2.976364 | NaN | 4.76491 | 3.93329 | NaN | 4.300541 | 4.304449 | 3.325256 |
| 6.418281 | 6.625358 | 6.510444 | 5.593149 | 5.647108 | 6.701748 | 6.339844 | 6.379582 | 6.992277 |
| 6.413439 | 6.538014 | 6.402496 | 6.508919 | 5.992951 | 7.208957 | 6.816499 | 7.419019 | 6.368312 |
| 7.053632 | 6.413744 | 6.791099 | 8.180992 | 8.460153 | NaN | 6.462047 | 6.230399 | 6.533755 |
| 6.836519 | 6.300004 | 6.751283 | 6.205139 | 6.467051 | 7.177458 | 7.075761 | 7.092954 | 6.823726 |
| 7.370433 | 6.94256 | 7.307648 | 6.487469 | 6.552784 | 7.859605 | 7.818238 | 7.680376 | 7.447663 |
| NaN | 6.163933 | 5.473467 | NaN | NaN | NaN | NaN | NaN | 6.056148 |
| NaN | NaN | NaN | NaN | 7.487194 | 7.422029 | 6.491048 | 6.539236 | NaN |
| 7.208203 | 7.921816 | 8.27911 | NaN | NaN | NaN | NaN | NaN | 4.917974 |
| 8.471555 | 8.499703 | 8.156267 | 7.732527 | 7.815822 | 8.470518 | 8.498058 | 8.411226 | 8.438299 |

| | | | | | | | | |
|---|---|---|---|---|---|---|---|---|
| NaN | NaN | NaN | 7.514048 | 6.2262 | 7.235748 | 6.517363 | 6.786566 | 6.624891 |
| 6.344405 | 6.600626 | 6.339965 | 5.273706 | 5.651892 | 5.226236 | 5.842502 | 5.253991 | 6.558675 |
| 7.425415 | 5.962794 | 7.195809 | 6.945051 | 6.354853 | 6.57277 | 6.774693 | 7.069682 | 6.669646 |
| 7.149708 | 7.388141 | 5.571293 | NaN | 5.595669 | 7.408264 | 6.618319 | 6.649276 | 5.402506 |
| 7.076456 | 7.460536 | 7.253293 | 8.265875 | 8.536404 | 5.448527 | 5.995386 | 5.395626 | 5.653547 |
| NaN | 6.787161 | NaN | 9.133121 | 8.988767 | 7.272165 | 6.894353 | 7.155101 | NaN |
| 5.019059 | 5.661033 | 5.18749 | NaN | 6.411788 | 6.003176 | 6.351682 | 6.44006 | 6.102827 |
| 7.107549 | 7.191479 | 7.213025 | 5.737485 | 6.606241 | 7.178977 | 7.015183 | 7.372743 | 6.960581 |
| 9.055511 | 9.177173 | 9.249888 | 7.948994 | 7.043972 | 8.272157 | 8.369361 | 8.131053 | 8.616186 |
| 6.391288 | 5.91433 | 6.666206 | NaN | NaN | 5.587942 | 6.653286 | 6.533409 | 6.491147 |
| 6.280621 | 6.155983 | 6.324671 | 7.118737 | 7.315227 | 6.01877 | 5.982275 | 6.20161 | 6.098955 |
| 7.50852 | 7.09919 | 7.244524 | 6.584989 | 6.330466 | 7.363861 | 7.035496 | 6.552479 | 7.282422 |
| 7.047523 | 6.732409 | 6.385527 | NaN | NaN | 7.069818 | 6.306082 | 6.476827 | 6.439252 |
| 6.443394 | 5.665017 | 6.444485 | 4.374248 | 4.124879 | 6.145041 | 6.339799 | 5.984186 | 5.425961 |
| NaN | NaN | NaN | NaN | 4.464904 | NaN | NaN | NaN | 5.128353 |
| 6.669224 | 6.413686 | 6.552192 | 5.361464 | 5.174789 | 7.209041 | 6.915889 | 6.999467 | 6.678957 |
| 8.272458 | 8.22838 | 8.254262 | 7.553473 | 7.539652 | 8.310376 | 8.291093 | 8.087758 | 8.044081 |
| 7.881914 | 7.630764 | 7.645181 | 6.993446 | 6.910461 | 7.734929 | 7.905046 | 7.822468 | 7.04026 |
| 6.710836 | 6.617495 | 6.827367 | 8.262006 | 8.402807 | 7.662992 | 7.204014 | 7.643063 | 7.852704 |
| 6.196983 | 6.322499 | 6.822408 | 7.983532 | 7.93469 | 6.21014 | 6.184498 | 6.416526 | 6.56438 |
| 8.281601 | 6.208448 | 6.810401 | 6.750731 | 6.757247 | 6.96534 | 6.695975 | 7.190493 | 6.75912 |
| 7.086713 | 7.221481 | 7.19741 | 7.365036 | 6.968124 | 6.874476 | 7.336154 | 7.180222 | 6.366758 |
| NaN | NaN | NaN | 4.875135 | NaN | NaN | 4.963922 | 5.417634 | 6.060624 |
| 7.075292 | 5.77221 | 8.22029 | 8.840159 | 7.920327 | NaN | NaN | NaN | 6.536479 |
| 7.320391 | 7.290424 | 7.373981 | 7.776323 | 7.524883 | 7.145941 | 7.024098 | 7.276018 | 6.989538 |
| 8.449418 | 8.420485 | 8.363741 | 9.317391 | 9.631063 | 8.334744 | 8.192745 | 8.545546 | 7.440077 |
| 4.57504 | NaN | 5.037662 | NaN | NaN | 5.364995 | 5.191138 | 4.873711 | 4.483059 |
| 5.537042 | 5.933379 | 5.513855 | 5.492352 | 5.927521 | 7.323628 | 6.10277 | 6.092472 | 5.114503 |
| 5.120121 | 4.931386 | 5.391855 | 2.527123 | 2.728253 | 6.046587 | 5.847619 | 6.036427 | 5.390512 |
| 7.275054 | 7.43985 | 7.268735 | 6.454756 | 6.741205 | 7.135158 | 7.11542 | 6.952528 | 7.390462 |
| NaN | NaN | NaN | NaN | NaN | NaN | NaN | NaN | 4.050982 |
| NaN | NaN | NaN | 7.057947 | 7.917603 | 7.543498 | NaN | 6.538024 | NaN |
| 6.537924 | 6.816859 | 6.823613 | 5.885083 | 6.48478 | 6.802969 | 6.893328 | 6.88925 | 6.861659 |
| 7.614814 | 7.677759 | 7.819501 | 7.83974 | 7.788129 | 7.235943 | 7.521121 | 7.018155 | 7.364523 |
| 5.890919 | 6.900151 | 6.666005 | NaN | NaN | 6.072199 | 6.172403 | 6.242685 | 6.253839 |
| 10.66604 | 10.53547 | 10.55684 | 11.41197 | 11.39573 | 10.95306 | 10.97865 | 10.93408 | 10.56671 |
| 7.269579 | 6.918378 | 6.678392 | 8.027591 | 7.385422 | 6.880998 | 6.81761 | 7.029942 | 6.643898 |
| 2.755405 | 1.719253 | 4.10787 | NaN | NaN | 5.012808 | 4.339726 | 3.858587 | 3.914057 |
| 7.064433 | 6.97253 | 6.895772 | 6.910306 | 6.596602 | 6.955538 | 6.940455 | 6.874528 | 6.675213 |
| 8.52632 | 8.470453 | 8.599868 | 7.205218 | 7.270348 | 8.193902 | 8.021034 | 8.14384 | 8.707388 |
| 7.183734 | 6.749537 | 6.898657 | 6.777116 | 6.950948 | 6.894815 | 7.188915 | 7.059274 | 6.758655 |
| 7.410673 | 7.606169 | 7.479773 | 7.575794 | 7.69199 | 7.465428 | 7.453756 | 7.437795 | 7.187077 |
| 6.681304 | 6.947299 | 6.487525 | 8.506128 | 8.626617 | 7.00423 | 6.799236 | 7.315031 | 6.407693 |
| NaN | 7.699272 | 7.106786 | NaN | 6.343307 | 5.968312 | 5.897615 | 6.704399 | 8.66732 |
| 8.435675 | 8.348555 | 8.12383 | 9.475581 | 9.187414 | 8.546347 | 8.252935 | 8.481203 | 9.07499 |
| NaN | 3.703005 | 3.240827 | 3.425836 | 4.021293 | NaN | 2.416744 | 2.305487 | NaN |
| 6.864473 | 6.194991 | 6.687964 | 4.846792 | 5.351291 | 5.726444 | 6.295843 | 6.04975 | 6.098512 |

| | | | | | | | | |
|---|---|---|---|---|---|---|---|---|
| 8.061099 | 8.752671 | 8.636642 | 8.359686 | 9.385702 | 7.345087 | 7.693665 | 8.285967 | 8.291806 |
| 5.849021 | 5.776636 | 5.75848 | 5.792578 | 5.5777 | 5.872159 | 5.710715 | 6.687324 | 6.863087 |
| 7.528006 | 7.494622 | 7.626563 | 6.910728 | 6.847027 | 7.845498 | 7.865983 | 7.986586 | 7.273757 |
| 5.742624 | 5.829107 | 5.530133 | 5.557177 | 5.968241 | 5.813247 | 5.646449 | 5.849841 | 3.480925 |
| 6.262254 | 6.440065 | 6.220954 | 6.473812 | 7.042847 | 5.838245 | 5.563584 | 6.421839 | 5.632467 |
| 6.108431 | 6.135059 | 6.185869 | NaN | NaN | 6.47559 | 6.246372 | 6.388158 | 6.36913 |
| 7.489566 | NaN | 6.725027 | NaN | NaN | 7.404222 | 7.293026 | 7.614102 | 7.923392 |
| 8.551642 | 8.602615 | 8.518377 | 9.574011 | 9.789149 | 8.535561 | 8.39485 | 8.02192 | 7.961924 |
| 7.628248 | 7.621661 | 7.751526 | 8.019748 | 8.089089 | 7.557266 | 7.900684 | 7.576816 | 7.43326 |
| 8.838991 | 8.684886 | 8.781847 | NaN | NaN | 7.751389 | NaN | NaN | 7.841726 |
| 7.676363 | 6.543674 | 7.282609 | 6.776742 | 6.438327 | 8.502572 | 8.27374 | 8.348416 | 6.580328 |
| 7.083388 | 6.896163 | 7.12913 | 7.770291 | 7.715103 | 6.742917 | 7.093709 | 7.060296 | 6.575211 |
| NaN | 4.214921 | 2.849908 | NaN | NaN | NaN | 3.847023 | 4.620767 | 3.314873 |
| 4.838246 | 4.351862 | 4.184579 | 4.863863 | 4.403273 | 5.05678 | 5.660466 | 5.172651 | NaN |
| 6.691136 | 6.688598 | 6.713958 | 5.489346 | 6.141649 | 7.048 | 6.860478 | 7.020341 | 6.734642 |
| 6.383674 | NaN | NaN | 7.420055 | 7.66047 | NaN | NaN | NaN | NaN |
| 8.293744 | 8.236188 | 8.136618 | 7.699833 | 7.804307 | 7.173155 | 7.63386 | 7.751069 | 6.958031 |
| 4.362708 | 4.000461 | NaN | 4.431851 | 4.313166 | 5.80748 | NaN | 6.397457 | 5.065691 |
| NaN | 5.899279 | 4.758848 | 6.390269 | 5.9989 | 5.240939 | 5.223568 | 4.480059 | 6.295135 |
| 6.590683 | 6.070331 | 5.730964 | 6.10301 | 5.031744 | 7.132344 | 7.0248 | 7.003803 | 6.601477 |
| NaN | NaN | 4.469408 | NaN | NaN | NaN | NaN | NaN | 5.510912 |
| 8.427709 | 8.38429 | 8.205101 | 6.776334 | 6.581256 | 5.831517 | 6.781762 | 6.646864 | 5.6281 |
| 6.357344 | 6.784625 | 6.279757 | 6.585449 | 6.288394 | 7.137618 | 6.497099 | 6.864498 | 5.940808 |
| 8.033689 | 6.748586 | 6.642406 | 6.796331 | 6.933881 | 7.179677 | 7.4365 | 7.493572 | 6.260665 |
| 7.468719 | 7.305448 | 7.478778 | 7.110382 | 7.479413 | 7.536758 | 7.181946 | 7.199759 | 7.339233 |
| 6.694082 | 5.699204 | 6.167717 | 5.819725 | 5.607184 | 5.143685 | 5.842766 | 3.811126 | 5.473017 |
| 7.496027 | 6.733634 | 6.835095 | 5.359148 | NaN | 7.048528 | 7.080057 | 6.610993 | 5.707786 |
| 5.563493 | 5.543756 | 6.351174 | NaN | NaN | NaN | 5.476995 | NaN | 5.694784 |
| 7.111513 | 7.252859 | 7.241127 | 7.022282 | 7.034743 | 6.999175 | 6.815988 | 6.874654 | 6.715516 |
| 8.433012 | 8.686866 | 8.729141 | 5.956103 | 5.869375 | 7.642272 | 7.851128 | 7.404921 | 8.796533 |
| 7.672996 | 7.606968 | 7.522504 | 7.569879 | 7.57482 | 7.288366 | 7.171041 | 7.149948 | 7.825075 |
| 7.748635 | 7.357634 | 7.374125 | 5.489217 | 5.815041 | 7.588327 | 7.361363 | 7.640666 | 7.545441 |
| 6.052081 | 6.211277 | 6.018705 | NaN | 3.611985 | 6.205427 | 6.255723 | 6.202131 | 6.208852 |
| 4.545962 | 4.205514 | 6.736886 | NaN | NaN | 6.189284 | 5.963667 | 6.396387 | 5.883798 |
| 7.960294 | 7.873153 | 7.837396 | 7.550909 | 7.62853 | 8.14337 | 8.017783 | 8.224471 | 7.692876 |
| 7.900017 | 8.42871 | 8.425059 | 8.88495 | 8.402513 | NaN | 5.646484 | NaN | 6.269858 |
| 6.948977 | 7.229609 | 6.918782 | 6.381432 | 6.705705 | 6.436312 | 6.320199 | 6.537212 | 5.409929 |
| 6.086505 | 5.572263 | 5.874804 | NaN | NaN | 4.116354 | 5.144744 | 5.312663 | 5.269153 |
| 6.926844 | 6.11752 | 6.004889 | NaN | 5.432962 | 7.715515 | 7.735603 | 7.396972 | 6.343956 |
| 7.048604 | 6.982847 | 6.915727 | 6.223331 | 5.995797 | 7.119423 | 7.080509 | 6.970658 | 6.786457 |
| 5.891856 | 5.963439 | 6.175981 | 7.357159 | 7.185274 | 5.513231 | 5.397667 | 5.870899 | 6.776114 |
| 6.349556 | 6.151116 | 6.176969 | 6.831297 | 7.31754 | 6.203193 | 6.184255 | 5.81754 | 6.174981 |
| 0.344843 | 4.343111 | 2.71153 | 5.071844 | 5.813669 | NaN | NaN | 3.070683 | 5.599917 |
| 6.806867 | 5.834806 | 6.831056 | 6.161418 | 5.278976 | 6.941486 | 6.876167 | 6.227769 | 6.591543 |
| 7.130495 | 6.593287 | 6.123628 | NaN | 6.252655 | 6.649475 | 7.098257 | 7.229437 | 7.249907 |
| 8.097816 | 5.330584 | 8.194214 | 8.005437 | 7.20991 | 6.187396 | 6.346384 | NaN | 6.870831 |
| 8.182546 | 7.764335 | 8.170165 | NaN | NaN | 7.25738 | 7.438163 | 6.898561 | 6.858349 |

| | | | | | | | | |
|---|---|---|---|---|---|---|---|---|
| 9.495211 | 9.286453 | 9.522884 | 9.625306 | 9.591826 | 8.473789 | 8.513918 | 8.412854 | 9.0472 |
| 6.80734 | 6.401576 | 6.530521 | 6.893353 | 6.822978 | 6.378028 | NaN | 6.211743 | 6.220062 |
| 5.800387 | 6.812095 | 5.77726 | 7.531438 | 7.437681 | 6.853394 | 6.59597 | 6.64683 | 6.263135 |
| 7.099827 | 7.189187 | 6.91245 | 6.670384 | 6.444823 | 5.966877 | 6.323874 | 6.232725 | 5.09733 |
| 7.103327 | 7.041217 | 6.988233 | 6.312233 | 6.242542 | 6.977572 | 6.99097 | 7.264891 | 6.915714 |
| 6.367442 | 6.126257 | 6.107527 | 5.731495 | 6.10677 | 6.475429 | 6.50191 | 6.303373 | 5.414432 |
| 7.836747 | NaN | 7.591783 | 7.665895 | 7.297902 | 7.588228 | 7.391331 | 7.516719 | 6.894794 |
| 6.60163 | 6.949695 | 6.448847 | NaN | NaN | 6.597883 | 6.774718 | 6.624444 | 7.482929 |
| 6.027337 | 6.744254 | 6.91783 | 5.411361 | 5.73105 | 6.627798 | 6.539217 | 6.450287 | 6.099769 |
| 5.914156 | 6.659182 | 6.451301 | 6.740878 | 6.669822 | 6.654126 | 6.8452 | 6.558363 | 6.547251 |
| 7.623027 | 7.577063 | 7.410211 | 6.725336 | 6.801439 | 7.60326 | 7.589301 | 7.708449 | 7.424728 |
| 5.577263 | 5.21243 | 5.330226 | 6.805781 | 7.042706 | 5.284026 | 5.531714 | 5.844662 | 6.495907 |
| 9.264429 | 9.169739 | 9.116725 | 8.102971 | 7.774592 | 8.938934 | 8.895788 | 8.860302 | 8.947206 |
| 6.766066 | 6.725398 | 6.86922 | 6.79487 | 6.489968 | 6.830107 | 6.946016 | 7.084902 | 6.144338 |
| 7.219927 | 7.237563 | 7.121453 | 6.880976 | 7.045375 | 7.963315 | 8.071733 | 7.915884 | 7.958771 |
| 6.411033 | 6.423728 | 5.645235 | NaN | 5.84012 | 6.323673 | 6.877816 | 6.914104 | 6.235125 |
| 7.201979 | 6.633492 | 7.681156 | 5.683524 | 6.069213 | 7.900136 | 7.582878 | 7.559759 | 7.431887 |
| 7.527446 | 6.986025 | 7.072291 | 6.65466 | 6.895567 | 6.668189 | 6.84124 | 6.933094 | 6.70606 |
| NaN | NaN | NaN | 6.7512 | 5.53591 | NaN | NaN | NaN | NaN |
| 5.995722 | 6.437013 | 6.110516 | 5.943794 | 6.142524 | 6.105779 | 6.665978 | 5.827343 | 5.477026 |
| 10.52626 | 10.19345 | 9.736284 | 9.953825 | 9.976789 | 9.609753 | 9.220715 | 9.7366 | 10.18744 |
| 6.611029 | 6.128387 | 6.200861 | 6.633283 | 7.510063 | 5.722266 | 5.610345 | 6.021784 | 4.969718 |
| 5.750948 | 5.472051 | NaN | NaN | NaN | 7.100863 | 7.29544 | 7.64941 | 5.797232 |
| 7.228945 | 7.460608 | 6.995951 | 7.046613 | 7.463032 | 7.430337 | 7.503601 | 7.32505 | 7.701789 |
| 6.340854 | 6.295099 | 5.91522 | 6.292159 | 6.454989 | 6.463467 | 6.124549 | 6.404919 | 5.714676 |
| 6.033507 | 6.098814 | 6.28532 | 6.156189 | 6.205781 | 5.95781 | 5.716204 | 5.86329 | 6.336285 |
| 7.406342 | 8.597767 | 7.291349 | 6.614222 | 9.521646 | 8.140849 | 7.906062 | 7.861625 | 7.696112 |
| 7.017756 | 7.151888 | 6.769734 | NaN | NaN | 6.641524 | 6.469861 | 6.179255 | 5.623284 |
| 6.260684 | 5.996635 | 6.090378 | 4.984184 | 5.494359 | 5.510122 | 5.723495 | 5.353903 | 5.758945 |
| 6.866926 | 6.731503 | 6.787731 | 6.413881 | 6.594759 | 7.492611 | 7.276647 | 7.266302 | 6.847796 |
| 9.948954 | 9.926701 | 10.02476 | 8.283694 | 8.336703 | 9.794793 | 9.793527 | 9.584149 | 9.072037 |
| 9.395685 | 10.60174 | 9.698073 | 10.21626 | NaN | 9.532046 | 9.627458 | 10.13389 | 9.950228 |
| 7.45395 | 7.669849 | 7.626173 | 5.418683 | 6.540859 | 8.05936 | 7.603826 | 7.693417 | 7.428028 |
| 7.628701 | 7.293755 | 7.443253 | 7.616138 | 7.814646 | 6.684169 | 6.745651 | 6.86647 | 6.722932 |
| 5.792995 | 6.122787 | 6.274016 | 7.902647 | 5.891544 | 6.113156 | 6.014308 | 6.045125 | 5.940978 |
| NaN | 6.169785 | 6.104349 | 6.181396 | 6.342985 | NaN | NaN | NaN | 7.092547 |
| 6.952245 | 6.586143 | 6.680613 | 6.299395 | 6.088262 | 7.509207 | 7.5339 | 7.493708 | 7.164789 |
| 4.65305 | 4.465065 | 4.988044 | NaN | 4.829775 | 5.793859 | 5.901765 | 5.437833 | 4.873821 |
| 6.857213 | 7.376625 | 7.175657 | 6.126371 | 5.941031 | 6.372992 | 6.710951 | 6.496994 | 7.46057 |
| 10.08902 | 9.834795 | 9.761353 | 9.278166 | 9.009364 | 9.263758 | 9.346635 | 9.293295 | 8.721763 |
| 6.121498 | 6.040129 | 6.385324 | 5.683016 | 5.436119 | 5.66519 | 4.846121 | 5.805511 | 5.613276 |
| 6.878653 | 6.872202 | 6.935822 | 6.493064 | 6.087555 | 5.467731 | 5.436322 | 5.872337 | 5.195197 |
| 5.392912 | 6.381408 | 6.538363 | 5.942708 | 5.2459 | 5.249894 | 5.03811 | NaN | 5.399647 |
| 5.305107 | NaN | 4.747622 | 5.420287 | NaN | NaN | NaN | 6.357471 | 5.147504 |
| 7.297331 | 6.778274 | 7.083778 | 6.892086 | 6.161818 | 7.82017 | 7.42705 | 7.188837 | 6.69924 |
| 6.703009 | 7.781398 | 6.861132 | 6.853527 | 7.044997 | 7.842142 | 8.147165 | 8.05551 | 7.27922 |
| 7.742005 | 7.658206 | 7.614197 | 7.791396 | 7.975287 | 7.059529 | 7.063757 | 7.152669 | 8.303101 |

| | | | | | | | | |
|---|---|---|---|---|---|---|---|---|
| 4.714017 | NaN | 4.887828 | NaN | NaN | 5.528526 | 5.813434 | 5.156978 | 5.961434 |
| 8.134043 | 8.166059 | 8.244765 | 8.421649 | 8.319757 | 7.835539 | 7.83011 | 7.88953 | 7.859882 |
| 5.707083 | 4.59548 | 6.055901 | 4.719623 | 5.387482 | 6.092367 | 6.097412 | 6.206713 | 5.677629 |
| 9.030005 | 9.025719 | 9.178022 | 8.545451 | 8.244614 | 7.311008 | 7.317343 | 7.454462 | 7.620679 |
| 7.478174 | 6.873033 | 7.482895 | 6.027809 | 6.105337 | 7.709492 | 7.38336 | 7.629988 | 6.744686 |
| 8.839101 | 8.586291 | 7.773264 | 9.452335 | 9.119193 | NaN | 7.184075 | 7.495061 | NaN |
| 6.343788 | 6.30251 | 5.890471 | 7.257741 | 7.293525 | 6.276533 | 6.063311 | 5.958356 | 6.603302 |
| 8.177445 | 8.671389 | 7.675657 | 8.306799 | 8.681858 | 4.957079 | 6.801825 | 6.179903 | 6.986169 |
| 4.194024 | 3.934715 | 3.442976 | NaN | NaN | 5.310656 | 5.230268 | 5.315263 | 4.504448 |
| 5.634055 | 6.803074 | 5.166731 | 4.794966 | 6.090961 | 6.29372 | 6.578556 | 6.510798 | 5.846838 |
| 5.633055 | 12.35977 | 12.47518 | 5.987195 | 6.382316 | 9.654459 | 9.539564 | 9.604527 | 8.54776 |
| 6.491947 | 6.563791 | 6.543206 | 6.614567 | 6.434467 | 7.002924 | 7.001076 | 7.200484 | 6.005951 |
| 7.625912 | 7.898639 | 8.0947 | 7.421269 | 7.501455 | 8.555341 | 8.542383 | 8.488994 | 7.915722 |
| 9.032146 | 8.970379 | 9.054629 | 7.779661 | 7.866721 | 8.991302 | 8.892342 | 9.022836 | 8.460426 |
| 8.787818 | 8.735641 | 8.882177 | 9.564171 | 9.401113 | 8.179929 | 8.198939 | 8.184325 | 8.26308 |
| 6.779171 | 3.667917 | 6.73848 | 6.740014 | 6.148761 | 6.537843 | 7.4055 | 6.941471 | 6.286263 |
| 7.585416 | 7.749555 | 7.679631 | 7.181371 | 7.373752 | 7.485171 | 7.177087 | 7.332809 | 7.471752 |
| 6.095199 | 6.039675 | 6.356646 | 6.354547 | 6.292964 | 6.414102 | 6.794819 | 6.850255 | 4.786337 |
| 7.118566 | 7.220911 | 6.986331 | 7.895686 | 7.746272 | 7.218433 | 7.156822 | 6.836532 | 8.436321 |
| 7.657282 | 6.98064 | 7.473813 | 6.920435 | 7.022955 | 6.872604 | 6.936734 | 6.681822 | 7.387769 |
| 7.666682 | 6.952913 | 6.92985 | 8.518916 | 8.275218 | 6.603123 | 6.861246 | 6.625578 | 7.469583 |
| 8.863746 | 7.81211 | 8.86017 | 6.289595 | 6.163556 | 7.869822 | 7.199924 | 8.008196 | 4.500089 |
| 6.541868 | 6.190772 | 5.807635 | NaN | 4.854458 | 6.636062 | 6.000118 | 6.259076 | 6.298074 |
| 8.183486 | 8.123561 | 8.028859 | 6.022698 | 6.171079 | 8.375541 | 8.805285 | 8.595963 | 8.432131 |
| 7.634835 | 7.37036 | 7.836568 | 8.758307 | 8.222164 | 7.13293 | 6.719049 | 7.142237 | NaN |
| 7.542013 | 7.812994 | 7.454201 | 7.341383 | 7.507525 | 6.95017 | 7.399666 | 7.330944 | 7.118057 |
| 7.823225 | 7.064926 | 6.885509 | 6.653943 | NaN | 8.805387 | 8.898636 | 8.85213 | 7.64918 |
| 9.724413 | 9.689693 | 9.693089 | 7.17033 | 7.648417 | 7.440258 | 6.866255 | 7.659588 | 7.157186 |
| 5.407314 | 4.507442 | NaN | NaN | NaN | 7.472644 | 7.454027 | 7.69913 | 5.671421 |
| 6.610049 | 6.723627 | 6.745785 | NaN | NaN | NaN | 6.814567 | 5.330564 | 5.729767 |
| 6.903105 | 7.167368 | 7.185521 | 7.077137 | 7.371696 | 6.217263 | 6.449799 | 6.519484 | 6.924445 |
| 5.487497 | 5.603931 | 4.892537 | 5.539696 | 5.351548 | 6.970418 | 6.001144 | 5.76332 | 6.150836 |
| NaN | NaN | NaN | NaN | NaN | 8.332408 | 8.162282 | 8.256172 | NaN |
| 8.277769 | 8.399452 | 8.56825 | 6.615118 | 6.730463 | 7.990417 | 8.157328 | 8.200488 | 7.771843 |
| 9.310491 | 7.788292 | 9.064518 | 9.501593 | 9.048144 | 8.451235 | NaN | 8.427059 | 8.772085 |
| 8.03538 | 8.35041 | 8.041776 | 6.052783 | 6.502083 | 7.866632 | 7.602014 | 7.813448 | 5.430193 |
| 2.16993 | 3.372411 | NaN | 3.565457 | NaN | 2.495549 | NaN | 1.306365 | 1.895383 |
| 8.112016 | 7.991675 | 8.126532 | 7.178005 | 7.210836 | 7.30974 | 7.51686 | 7.631725 | 7.780899 |
| 6.22829 | 6.193531 | 6.670421 | 7.004515 | 7.02989 | 7.026129 | 6.674889 | 6.821312 | 6.25846 |
| 6.271574 | 6.253588 | 6.518057 | 5.669921 | 6.065586 | 5.940229 | 6.043682 | 6.616289 | 4.842372 |
| 9.24167 | 9.123994 | 9.276184 | 9.382015 | 9.543746 | 9.248813 | 9.502468 | 9.312357 | 8.415075 |
| 6.366324 | 6.558241 | 6.622358 | 6.295177 | 6.230133 | 6.612941 | 6.43612 | 6.556633 | 6.823562 |
| 5.126879 | 6.293151 | 5.549414 | 5.72816 | 5.429513 | 5.305724 | 5.270776 | 4.838993 | 6.101233 |
| NaN | NaN | NaN | NaN | NaN | NaN | NaN | NaN | 5.309057 |
| 6.658107 | 6.647653 | 6.798733 | 5.610552 | 6.63623 | 7.464284 | 7.436823 | 7.423414 | 7.397251 |
| 8.238788 | 8.744129 | 8.515471 | 5.429768 | 5.58776 | 10.23464 | 9.787099 | 10.18439 | 7.830766 |
| 7.187984 | 7.1983 | 7.626644 | 6.293659 | 6.840316 | 7.124849 | 6.789285 | 6.939412 | 6.952018 |

| | | | | | | | | |
|---|---|---|---|---|---|---|---|---|
| 9.251578 | 9.319182 | 9.05629 | 9.350476 | 9.672012 | 9.063367 | 9.1446 | 9.13391 | 9.258302 |
| 5.678458 | 3.940595 | 5.604233 | 5.60721 | 5.093113 | NaN | NaN | 6.562229 | NaN |
| 7.235729 | 7.371804 | 7.341032 | 7.529389 | 7.422457 | 7.982613 | 7.92993 | 7.991673 | 7.554399 |
| 8.245349 | 8.182463 | 8.0624 | 6.954686 | 7.569859 | 7.946287 | 7.82894 | 7.900706 | 7.321991 |
| 7.392256 | 8.230956 | 8.0353 | NaN | 6.569841 | 8.331671 | 8.495189 | 8.549098 | 8.14449 |
| 8.180477 | 8.142999 | 8.261909 | 7.715133 | 7.795681 | 8.099901 | 8.558127 | 8.40335 | 8.055678 |
| NaN | 6.852352 | 6.893542 | NaN | NaN | 7.412297 | 7.154336 | 7.517107 | 6.965117 |
| 7.822953 | 6.914905 | 7.063532 | 6.916225 | 7.452709 | 6.616228 | 6.734333 | 7.216864 | 5.207089 |
| 7.808563 | 7.806538 | 8.030293 | 7.902481 | 8.236432 | 7.577559 | 7.595639 | 7.573144 | 8.213967 |
| 9.023691 | 8.914336 | 8.857095 | 11.08173 | 10.34241 | 8.292592 | 9.02849 | 8.751866 | 7.633657 |
| 5.999135 | 4.06322 | NaN | 6.782472 | 6.333811 | 6.495866 | 5.745789 | 6.326607 | 5.158234 |
| 6.998672 | 7.253158 | 6.711295 | 5.740416 | 5.256076 | 6.811253 | 7.028613 | 7.117007 | 7.280495 |
| 7.163345 | 6.954057 | 7.251597 | 5.866593 | NaN | 7.295683 | 7.205102 | 7.020803 | 6.942079 |
| 7.153362 | 5.176583 | 5.803353 | NaN | 6.64316 | 6.699336 | 5.343214 | 5.886652 | 5.438875 |
| 6.660822 | 6.512884 | 6.798244 | 6.554513 | 6.357578 | 6.668234 | 6.999275 | 7.405609 | 6.313201 |
| NaN | 8.020449 | 7.687293 | 8.825171 | 8.566694 | NaN | NaN | NaN | 6.902182 |
| 7.647579 | 7.531307 | 7.629912 | 7.337678 | 7.698803 | 6.721194 | 6.639239 | 6.841529 | 7.912225 |
| NaN | NaN | 6.595269 | 6.664093 | 7.021185 | 6.668274 | 6.384993 | 6.863069 | 6.39344 |
| 6.002906 | 6.140225 | 5.562212 | 5.875126 | 5.723604 | 5.691986 | 4.907921 | 5.094615 | 5.436896 |
| 7.611225 | 6.209869 | 7.402904 | 5.638513 | 5.588534 | 5.85101 | NaN | 6.182265 | 5.348724 |
| 7.363328 | 7.779729 | 7.772448 | 7.354841 | 6.759049 | 7.690202 | 7.539725 | 7.573503 | 6.907144 |
| 5.714335 | 5.146046 | 5.564088 | 5.308463 | 5.274872 | 5.70157 | 5.903724 | 5.44882 | 5.888751 |
| 6.596541 | 5.957232 | 6.155797 | NaN | 6.755975 | 7.016277 | 6.771414 | 6.605368 | 6.504403 |
| 7.172545 | 7.01007 | 6.957747 | 7.551894 | 7.551087 | 6.409833 | 6.500885 | 6.604222 | 6.23907 |
| NaN | 5.902515 | 5.595554 | 6.239141 | 6.102561 | 5.786439 | 5.414472 | 5.619278 | 5.564551 |
| 6.703948 | 6.204933 | 7.619874 | 6.609236 | 7.916125 | 7.247399 | 5.571644 | 6.301879 | 6.579628 |
| 5.650886 | 5.925274 | 5.732183 | 5.740127 | 6.208789 | 6.407724 | 6.688413 | 6.53127 | 5.469761 |
| 5.9935 | 5.584224 | 5.640511 | 7.572289 | 7.248243 | 5.64691 | 5.693695 | 5.748477 | 5.699882 |
| 6.33163 | 7.876735 | NaN | 7.808561 | 7.815332 | 5.38121 | NaN | 5.531635 | 6.067554 |
| 6.475323 | 6.754009 | 6.976293 | 7.901838 | 7.969336 | 7.438231 | 7.652182 | 7.750179 | 6.952738 |
| 9.477635 | 8.726371 | 8.054694 | NaN | 9.036651 | NaN | 7.20736 | NaN | 6.717892 |
| 3.379838 | NaN | NaN | NaN | NaN | NaN | NaN | NaN | 4.451255 |
| 7.24429 | 7.200579 | 7.919372 | 4.627843 | 4.103574 | 6.707376 | 6.240005 | 5.854349 | 6.927376 |
| 8.245835 | 8.296277 | 8.039569 | 7.152321 | 6.947045 | 7.947055 | 8.163429 | 8.149039 | 8.369365 |
| 6.629884 | 6.646395 | 6.721613 | 6.401079 | 6.322355 | 6.938245 | 7.010329 | 7.112765 | 7.018058 |
| 6.338719 | 4.929788 | 6.516572 | NaN | NaN | 6.181146 | 6.115905 | 6.393211 | 6.946893 |
| 7.021287 | 7.29169 | 7.221838 | 7.341962 | 7.101786 | 7.269233 | 7.497858 | 7.205512 | 7.419964 |
| 6.290616 | 5.421774 | 6.001752 | 6.317019 | 6.068474 | 7.475845 | 7.908789 | 7.629969 | 5.454009 |
| 8.130694 | 7.900192 | 8.065924 | 8.204976 | 7.61851 | 7.318884 | 8.622145 | 8.016466 | 7.507913 |
| NaN | NaN | 4.38988 | NaN | NaN | NaN | NaN | NaN | 4.129817 |
| 8.374585 | 8.533593 | 8.368788 | 8.562713 | 8.171014 | 7.890698 | 7.598827 | 7.842831 | 7.951807 |
| 5.725228 | 6.080267 | 5.688644 | 5.753607 | 5.878397 | 4.775308 | 4.999293 | 5.048216 | 4.532738 |
| 5.835204 | 5.904076 | NaN | 5.610019 | 5.87234 | NaN | 6.189747 | 5.924116 | 6.141437 |
| 6.288166 | 5.464986 | 5.717411 | 5.445333 | 5.738817 | NaN | NaN | NaN | 5.269172 |
| 8.187534 | 8.078264 | 8.426517 | NaN | NaN | 7.966118 | 7.825078 | 8.024308 | 6.898039 |
| 9.658163 | 9.454348 | 9.28362 | 7.732383 | 8.240108 | 9.047557 | 9.578492 | 8.96751 | 8.951805 |
| 7.252167 | 7.510568 | 7.093595 | 6.229039 | 6.966166 | 8.363628 | 7.044977 | 7.258653 | 7.185547 |

-Bruker Confidential-

| | | | | | | | | |
|---|---|---|---|---|---|---|---|---|
| 6.012166 | 6.372671 | 6.470981 | 5.670806 | NaN | NaN | NaN | NaN | NaN |
| 8.844655 | 9.102566 | 8.756502 | 6.900339 | 6.926368 | 8.518905 | 8.758013 | 8.50477 | 7.73958 |
| 6.952287 | 7.039485 | 7.12947 | 8.551461 | 8.87328 | 8.272784 | 8.149421 | 8.343425 | 5.987821 |
| 5.040478 | 5.380917 | 5.795974 | 5.912539 | 6.048744 | 4.620849 | 4.78857 | 4.908637 | 4.811853 |
| NaN | NaN | NaN | 7.284241 | 7.134507 | NaN | NaN | NaN | 6.492378 |
| 6.032385 | 7.209985 | 7.262295 | NaN | NaN | NaN | NaN | NaN | 6.359653 |
| NaN | 5.510434 | NaN | 6.276175 | 6.741696 | 5.889256 | 6.268889 | 6.047648 | 7.08088 |
| 6.978914 | 7.36867 | 7.033991 | 5.175184 | 5.567938 | 6.82632 | 7.047779 | 7.053883 | 6.772568 |
| 9.841707 | 9.943076 | 9.927844 | 9.200868 | 9.255474 | 9.696301 | 9.822986 | 9.760861 | 10.18126 |
| 7.618742 | 7.674186 | 7.590867 | 6.640038 | 6.876313 | 8.111304 | 8.04298 | 8.00843 | 6.960908 |
| 7.231771 | 6.920122 | 7.187673 | 5.40752 | 5.552809 | 6.817474 | 6.908741 | 6.515747 | 6.772127 |
| 7.927125 | 7.970716 | 7.966111 | 7.205945 | 7.447727 | 7.697186 | 7.730129 | 7.781799 | 8.023296 |
| 11.20119 | 10.9554 | 10.97314 | 10.99897 | 11.0123 | 10.72737 | 10.87442 | 10.77061 | 10.73709 |
| 9.067029 | 8.682961 | 8.703929 | 6.925382 | 7.179472 | 8.693439 | 8.429745 | 8.319191 | 7.040306 |
| 7.445168 | 7.279373 | 7.628505 | 6.664733 | 6.750554 | 6.570594 | 6.656966 | 7.419728 | 7.077722 |
| 6.533581 | 6.977494 | 7.190142 | 5.102024 | 5.037287 | NaN | NaN | NaN | 6.841816 |
| 7.220132 | 7.340083 | 7.414842 | 5.6719 | 5.970239 | 7.241048 | 7.262095 | 7.354822 | 7.434624 |
| 5.55333 | 4.707376 | 5.393395 | 6.830528 | NaN | 5.821959 | 6.208252 | 6.144565 | 6.138196 |
| 6.992234 | 6.631494 | 7.195934 | 4.388603 | 6.148036 | 8.05092 | 7.148175 | 7.2086 | 6.601319 |
| 7.111522 | 7.517724 | 7.730731 | 9.256495 | 9.146172 | 8.195275 | 8.26469 | 7.867319 | 6.230733 |
| 7.496924 | 7.648229 | 7.631216 | 6.878987 | 6.101979 | 7.607732 | 7.556915 | 7.507049 | 7.519282 |
| 8.150643 | 7.906154 | 7.951863 | 7.145362 | 7.14387 | 8.335794 | 8.126725 | 8.167616 | 8.285007 |
| 7.419428 | 7.381142 | 7.076097 | 5.793944 | 6.726556 | 8.031383 | 7.701919 | 7.865821 | 7.341601 |
| 5.87069 | 4.663542 | 5.924606 | 4.81772 | 5.008126 | 5.278124 | 5.207858 | 5.513693 | 5.00764 |
| 4.542083 | NaN | NaN | NaN | NaN | 6.410527 | 6.532806 | 6.666476 | 5.013534 |
| 6.681102 | 7.515452 | 6.990819 | NaN | 6.38091 | 7.136889 | 6.984306 | 7.324817 | 6.889291 |
| 7.954922 | 8.085697 | 7.805115 | 8.107578 | 7.687623 | 6.483132 | 6.995993 | 6.885292 | 7.631192 |
| 4.648722 | 4.017123 | NaN | 4.535644 | 4.137435 | NaN | NaN | NaN | NaN |
| 9.007571 | 7.548371 | 8.814252 | 6.949032 | 7.316556 | 8.284925 | 8.942725 | 8.390998 | 8.41509 |
| 7.713082 | 7.734033 | 7.669765 | 5.860711 | 5.850155 | 7.804302 | 7.621607 | 7.63284 | 7.4172 |
| 6.976958 | 6.533904 | 6.897648 | 6.066761 | 6.488186 | 6.86067 | 6.949265 | 7.456229 | 6.497692 |
| 7.695284 | 7.774374 | 7.936731 | 6.657163 | 6.75105 | 7.27451 | 7.435588 | 7.442739 | 7.690486 |
| 8.5794 | 8.333388 | 8.456603 | 7.48828 | 7.164668 | 8.760652 | 8.735339 | 8.848861 | 8.282695 |
| NaN | NaN | NaN | NaN | NaN | 4.997363 | NaN | 5.838324 | 5.387892 |
| 6.641345 | 6.545871 | 6.568043 | NaN | NaN | 6.627576 | 6.533689 | 6.64989 | 6.842618 |
| 7.691019 | 7.59915 | 7.473347 | 6.034215 | 6.550455 | 8.092175 | 8.146611 | 8.066126 | 7.209451 |
| 8.177808 | 8.154755 | 8.08707 | 7.568796 | 7.654783 | 8.45441 | 8.14967 | 8.232217 | 7.945206 |
| 5.890002 | 6.462251 | 6.339925 | 8.152329 | 8.361861 | 6.958757 | 7.240417 | 7.213158 | 5.838211 |
| 3.804054 | 4.576678 | NaN | 3.058119 | 2.70561 | 4.829403 | 4.624018 | 5.150307 | NaN |
| 6.941133 | 6.843722 | 6.601659 | 7.263755 | 6.657229 | 6.46563 | 6.70466 | 6.493468 | 6.883658 |
| 4.983041 | NaN | 4.233698 | NaN | NaN | 5.862791 | 5.31044 | 5.237126 | 5.723881 |
| 5.131784 | NaN | NaN | NaN | NaN | 5.604117 | 5.717317 | NaN | NaN |
| 7.464483 | 7.834991 | 7.707999 | 7.365211 | 6.7331 | 7.610409 | 7.413535 | 7.703633 | 6.576235 |
| 7.883266 | 7.799471 | 7.912697 | 7.353207 | 7.362152 | 7.583792 | 7.611285 | 7.803249 | 7.842676 |
| 8.702521 | 8.263773 | 8.360964 | 9.681428 | 9.392439 | 8.435822 | 8.449726 | 8.351675 | 8.798285 |
| NaN | NaN | 5.04924 | 6.99678 | 7.178174 | NaN | NaN | NaN | NaN |
| 6.69316 | 6.280518 | 6.635448 | 5.914215 | 6.391849 | 6.752434 | 6.987012 | 7.078176 | 7.09183 |

| | | | | | | | | |
|---|---|---|---|---|---|---|---|---|
| 7.218479 | 6.915009 | 7.017975 | 6.110011 | 4.886476 | 7.111772 | 7.034091 | 7.086886 | 6.804463 |
| 6.263131 | 5.545936 | 6.197555 | 5.052202 | 6.011036 | 6.514789 | 6.58595 | 6.177349 | 5.187058 |
| 6.422101 | 4.917058 | 5.633053 | NaN | NaN | 6.491321 | 6.44653 | 6.538748 | 6.440305 |
| 8.124344 | 8.209779 | 8.213561 | 8.358684 | 8.077801 | 7.928248 | 8.125758 | 8.020535 | 7.417014 |
| 6.662813 | 6.693974 | 6.331375 | 5.134634 | 5.368402 | 6.457769 | 6.041287 | 6.588693 | 6.2734 |
| 9.500171 | 9.365989 | 9.44666 | 10.43506 | 10.37898 | 8.638553 | 8.402857 | 8.632406 | 8.034611 |
| 7.026794 | 7.140763 | 7.867572 | NaN | 5.852652 | 6.257759 | 7.125154 | 7.131556 | 7.036905 |
| 6.766921 | 6.716154 | 6.530759 | 6.827236 | 6.478117 | 6.170087 | 5.876537 | 6.252966 | 5.791463 |
| 7.831816 | 7.007694 | 7.733082 | 6.627903 | 6.810217 | 7.717528 | 7.614758 | 7.710748 | 7.632405 |
| 6.831337 | 6.348751 | 6.440978 | 6.859332 | 6.55872 | 6.678209 | 6.845441 | 6.769853 | 6.210401 |
| 8.389891 | 8.457437 | 8.699183 | 7.842398 | 7.611231 | 9.888437 | 9.449766 | 9.683057 | 6.621573 |
| 8.480155 | 8.536693 | 8.356769 | 9.363886 | 9.379444 | 8.372716 | 8.140754 | 8.309036 | 8.276729 |
| 6.809917 | 7.522134 | 7.645564 | 8.384739 | 7.97188 | 7.915853 | 7.913038 | 7.909445 | 7.839085 |
| 6.276479 | 6.069819 | 6.797072 | 6.319866 | 6.250727 | 6.822855 | 6.842015 | 6.603559 | 6.099663 |
| 6.177307 | 6.45074 | 5.894886 | NaN | 4.806622 | 6.226043 | 5.950294 | 5.879339 | 6.693743 |
| 5.628989 | 5.884782 | 5.697465 | 5.74075 | 5.880662 | 6.386681 | 6.220323 | 5.878374 | 5.850912 |
| 6.981884 | 6.802255 | 7.237982 | 6.164454 | 6.425928 | 7.1731 | 7.145223 | 7.387265 | 7.658432 |
| 6.522083 | 6.758924 | 6.428733 | 6.270472 | 6.384042 | 6.262995 | 6.839771 | 6.779605 | 6.452067 |
| NaN | 3.64703 | NaN | NaN | 5.794515 | 3.942075 | 4.098743 | 3.559885 | 5.250252 |
| 7.886033 | 7.743769 | 7.782808 | 6.506896 | 7.230396 | 7.664988 | 8.051994 | 8.177287 | 7.526769 |
| 5.407522 | 5.609276 | 4.892166 | 4.30974 | NaN | 5.080073 | 5.379769 | 5.890198 | 5.697672 |
| 12.66791 | 12.24538 | 12.53353 | 11.70939 | 11.12297 | 7.813281 | 7.536272 | 7.508352 | 8.488769 |
| 5.545137 | NaN | 5.194891 | 6.743331 | 6.678312 | NaN | 6.06427 | 6.41345 | 6.274524 |
| 6.602044 | 6.486327 | 6.030123 | 5.289996 | 5.480151 | 7.371918 | 7.284594 | 7.394099 | 6.428086 |
| 7.027028 | 6.496412 | 6.556058 | 7.609185 | 7.602092 | 6.297732 | 6.153591 | 6.071085 | 7.058389 |
| NaN | 6.217552 | 6.476351 | 6.939667 | 7.59522 | NaN | NaN | 8.030145 | 4.291555 |
| 7.640092 | 7.162526 | 7.218855 | 7.615694 | 7.908494 | 5.776058 | 6.044707 | 6.809377 | 6.333979 |
| 5.045914 | 4.297962 | 4.711412 | 2.937132 | NaN | NaN | NaN | NaN | 1.959102 |
| 8.887044 | 8.76181 | 8.73902 | 8.017236 | 7.945379 | 9.114718 | 9.015388 | 8.954487 | 8.833719 |
| 6.773179 | 6.50985 | 6.882055 | 6.341136 | 6.772115 | 7.189202 | 7.454606 | 7.161485 | 6.834328 |
| 8.059608 | 7.916067 | 7.967966 | 8.26667 | 8.874682 | 7.355114 | 7.749463 | 7.896723 | 7.909187 |
| 6.584533 | 5.599628 | 6.713004 | 6.141854 | 5.732244 | 7.34739 | 7.009904 | 7.221004 | 7.077952 |
| 7.445193 | 7.43351 | 7.063295 | 8.193218 | 7.591041 | 6.77494 | 6.876705 | 6.946061 | 6.678721 |
| NaN | 5.220941 | 4.46973 | 5.397202 | 6.013544 | 5.42492 | NaN | 6.309034 | 6.46257 |
| 5.495667 | 5.657979 | 5.168183 | NaN | NaN | 5.835961 | 5.944834 | 5.823983 | 5.668923 |
| 8.252222 | 8.428209 | 8.460141 | 7.028708 | 7.142169 | 8.082172 | 7.640946 | 7.789873 | 6.947369 |
| 5.705616 | 6.194957 | 6.108136 | NaN | 4.800134 | 6.126358 | 6.133902 | 6.050769 | 5.552012 |
| NaN | NaN | NaN | NaN | NaN | NaN | NaN | NaN | 6.43411 |
| 6.037139 | 5.059709 | 6.292085 | 4.640471 | 5.255729 | 6.215981 | 5.534297 | 5.886362 | 6.380613 |
| 6.555533 | 6.446288 | 7.226398 | 4.867859 | 5.235756 | 6.112648 | 6.698244 | 6.446017 | 5.790795 |
| 8.74428 | 8.755316 | 8.828417 | 8.517688 | 8.640649 | 9.205313 | 8.841883 | 9.002991 | 8.368912 |
| 9.217067 | 8.998342 | 9.184861 | 10.12492 | 10.12665 | 9.84635 | 9.715894 | 9.953969 | 10.12019 |
| 9.071748 | 9.077205 | 8.993003 | 9.489612 | 9.198932 | 9.716296 | 9.630089 | 9.872796 | 9.211766 |
| 6.538844 | 6.850479 | 6.722841 | NaN | NaN | 5.350302 | 6.179115 | 6.354765 | 5.913325 |
| 7.723128 | 7.744117 | 7.78688 | 7.758403 | 7.759184 | 7.352172 | 7.431352 | 7.337274 | 7.696514 |
| 8.329168 | 8.129872 | 8.612037 | 7.507033 | 7.539743 | 8.147492 | NaN | 8.245234 | 7.641867 |
| 4.730564 | 5.234148 | 5.586095 | 6.683253 | 5.870229 | 5.251447 | 5.478475 | 4.916245 | 5.385004 |

| | | | | | | | | |
|---|---|---|---|---|---|---|---|---|
| 7.002427 | 6.772554 | 6.880934 | 5.938281 | 5.939262 | 5.760717 | 6.374197 | 6.51262 | 6.065008 |
| NaN | 5.381155 | NaN | 8.390903 | 7.42969 | 6.413397 | 6.443055 | 6.574314 | 6.147989 |
| NaN | 10.42346 | NaN | 7.341883 | 7.553776 | 6.475357 | NaN | NaN | 6.560345 |
| 5.791076 | 5.717938 | 5.585646 | 4.909761 | 5.414542 | 5.738855 | 5.310678 | 5.887665 | 5.748564 |
| 6.565543 | 6.030562 | 6.299248 | 5.806139 | 5.800027 | 6.748867 | 6.756558 | 6.508926 | 5.60792 |
| NaN | 6.63933 | NaN | NaN | NaN | 6.57529 | 6.589921 | 6.362602 | 6.908129 |
| 6.861747 | 6.714853 | 6.530527 | 5.788692 | 5.574125 | 7.232912 | 7.365088 | 7.304244 | 6.677416 |
| 6.414583 | 6.099141 | 6.344695 | 7.03473 | 6.987567 | 6.125074 | 6.44087 | 6.591507 | 6.868728 |
| 9.159424 | 9.65301 | 9.575967 | 9.145411 | 9.400187 | 9.009888 | 9.203341 | 9.167231 | 9.537185 |
| 5.05125 | 5.430121 | 4.555002 | 4.511256 | 5.128222 | NaN | 5.449507 | 5.978221 | 5.972999 |
| 7.210042 | 6.475952 | 7.056911 | 6.438575 | 6.385346 | 6.935239 | 6.901386 | 6.71312 | 6.51554 |
| 8.638928 | 9.363254 | 9.710302 | 10.0522 | 9.363261 | 8.205956 | 8.520907 | 8.579428 | 8.767385 |
| 8.969588 | 8.487183 | 8.591355 | 10.51254 | 10.67924 | 9.129985 | 8.981086 | 8.903465 | 8.486444 |
| 5.795978 | 6.011976 | 6.265579 | 4.750214 | 5.661653 | 6.448395 | 6.43401 | 6.58416 | 5.819895 |
| 7.533739 | NaN | 8.012883 | NaN | 7.643075 | NaN | 8.22035 | 8.300456 | NaN |
| 8.545872 | 8.510505 | 8.383685 | 8.057629 | 7.9804 | 8.607643 | 8.614721 | 8.603466 | 8.502504 |
| 7.905767 | 7.856401 | 8.170639 | 8.425471 | 8.021764 | 7.815408 | 7.866084 | 7.902431 | 8.300529 |
| 13.18107 | 13.01501 | 13.06603 | 10.93733 | 11.10835 | 10.90138 | 9.567257 | 8.484825 | 10.07475 |
| 7.038544 | 6.898413 | 7.158531 | NaN | 6.154478 | 7.378235 | 7.252885 | 6.925727 | 6.420741 |
| 7.369499 | 7.457738 | 7.490365 | 7.759359 | 7.847642 | 7.492596 | 7.306887 | 7.053363 | 7.796853 |
| NaN | 2.862398 | NaN | 6.087264 | 6.146686 | NaN | NaN | NaN | 5.308026 |
| 6.364176 | 6.886443 | 6.481582 | 6.98958 | 6.573679 | 5.685712 | 6.053151 | 6.250118 | 6.548497 |
| 9.470827 | 9.383804 | 9.423555 | 9.673435 | 9.627163 | 9.505464 | 9.546242 | 9.374966 | 9.420514 |
| 6.795495 | 6.719474 | 6.253503 | 6.392229 | NaN | 6.060476 | NaN | 6.300596 | 5.203197 |
| 9.345714 | 9.510701 | 9.355949 | 9.109307 | 8.747747 | 10.33149 | 10.22977 | 10.20086 | 9.701452 |
| 6.322132 | 6.596867 | 6.567141 | NaN | NaN | 6.41826 | NaN | 6.210672 | 6.556465 |
| 5.699755 | 6.49925 | 6.097581 | NaN | 6.027903 | 6.417455 | 6.418006 | 6.300569 | 5.953533 |
| 7.109175 | 7.498148 | 7.369003 | 8.133695 | 8.626346 | 7.860257 | 6.734482 | 7.151232 | 6.411677 |
| 8.298772 | 8.771322 | 8.003428 | 6.486031 | 5.788587 | 7.788199 | 8.031388 | 7.97351 | 8.071281 |
| 4.839314 | 4.522163 | 4.417513 | 3.363334 | 5.875586 | NaN | 4.325111 | 4.853502 | 3.382921 |
| 8.041506 | 8.460328 | 8.196662 | 8.283257 | 8.675882 | 7.327245 | 7.111701 | 7.092441 | 7.403548 |
| 8.237945 | 7.647809 | 7.881481 | 5.798155 | 7.315667 | 7.431833 | 7.390423 | 7.969032 | 7.462744 |
| 6.255054 | 5.936923 | 5.956099 | 5.538936 | 7.060696 | 7.032052 | 7.131579 | 7.008347 | 5.813981 |
| 9.026999 | 8.474293 | 8.847065 | 8.428893 | 8.167617 | 8.845431 | 8.643478 | 8.404248 | 7.933225 |
| 5.257285 | 5.295718 | 5.186552 | 4.753657 | 4.457552 | 5.118763 | 5.408605 | 5.528775 | 4.908607 |
| 6.686131 | 6.464152 | 6.798093 | NaN | NaN | 5.914013 | 6.896143 | 6.41373 | 6.287244 |
| 5.779499 | 5.30032 | 5.52507 | 6.73587 | 6.442896 | NaN | NaN | 6.450701 | 6.328944 |
| 8.157783 | 8.211023 | 8.081556 | 6.726499 | 6.752978 | 8.074189 | 8.201006 | 8.043629 | 7.676178 |
| 5.831744 | 6.364843 | 6.396252 | 6.53286 | 6.51033 | NaN | NaN | NaN | 5.144375 |
| 8.06467 | 6.35305 | 7.740012 | 7.030879 | 6.66354 | 7.160147 | 7.128769 | 6.907137 | 7.472561 |
| 6.744017 | 6.574425 | 6.511609 | 5.383092 | 5.111954 | 7.100568 | 7.200234 | NaN | 6.493453 |
| 6.429692 | 4.116714 | 6.605386 | 4.964064 | 5.227149 | 6.991564 | 6.132502 | 6.377122 | 6.005148 |
| NaN | 5.942873 | 5.772379 | 6.201997 | 5.688323 | 6.91439 | 6.728117 | 7.156318 | 6.567035 |
| 7.037446 | 6.109306 | 6.557903 | 6.883437 | 6.937222 | 6.550798 | 6.407645 | NaN | 6.132301 |
| 7.147457 | 7.255658 | 7.105315 | 6.375193 | 5.532757 | 7.175425 | 7.393712 | 7.466634 | 7.114716 |
| 8.99074 | 8.768312 | 8.984686 | 10.09903 | 10.03404 | 7.426615 | 8.340626 | 7.511832 | 8.763388 |
| 7.28283 | 7.465663 | 7.49922 | 7.191382 | 6.858632 | 7.546485 | 6.845387 | 7.48538 | 7.715463 |

| | | | | | | | | |
|---|---|---|---|---|---|---|---|---|
| 6.385263 | 6.336689 | 7.062432 | 5.417666 | 5.719068 | 6.252666 | 6.69069 | 5.88481 | 5.666118 |
| 8.055843 | 7.933759 | 8.01806 | 7.312107 | 7.396228 | 7.832984 | 8.212246 | 8.055938 | 7.930408 |
| 7.353692 | 7.142174 | 7.341681 | 7.007862 | 6.833111 | 7.132991 | 7.200032 | 7.419372 | 7.255514 |
| 7.144766 | 7.34255 | 7.32024 | 6.421848 | 6.765842 | 7.416998 | 7.233297 | 7.455874 | 6.960513 |
| 8.042203 | 7.892421 | 8.152737 | 7.75528 | 7.895271 | 7.934665 | 8.305194 | 8.529343 | 7.116921 |
| 6.877569 | 5.842143 | 6.339016 | 6.859927 | 4.526627 | 6.704514 | 6.41706 | 6.745718 | 6.010738 |
| 7.498285 | 7.071734 | 6.893366 | NaN | NaN | 7.762209 | 8.02241 | 8.174436 | 7.147293 |
| 7.650649 | 7.322505 | 7.412416 | 7.803935 | 7.428446 | 6.840878 | 6.821185 | 7.057272 | 6.704573 |
| 8.675944 | 8.731441 | 8.628119 | 8.763926 | 8.77275 | 8.658799 | 8.59037 | 8.670211 | 9.377316 |
| 6.246479 | 6.140328 | 6.135396 | 6.806744 | 6.997803 | 6.737176 | 6.789588 | 6.5598 | 5.048505 |
| NaN | 6.402882 | 8.114116 | NaN | 5.439713 | 5.739545 | 6.127848 | 6.094931 | 5.703734 |
| NaN | NaN | 5.848807 | 6.090879 | 6.132684 | NaN | NaN | 4.670902 | 5.859434 |
| 7.334587 | 7.316653 | 7.609358 | 7.837501 | 7.725677 | 7.364186 | 7.515111 | 7.337554 | 7.065735 |
| 7.101511 | 7.300519 | 6.965579 | 5.587023 | 5.205084 | 6.894547 | 6.877569 | 6.810179 | 6.902687 |
| NaN | NaN | NaN | 5.464271 | 6.317319 | 7.560143 | 6.239275 | 7.419347 | NaN |
| 6.294214 | 6.916455 | 6.618628 | 6.586151 | 5.275166 | 7.46722 | 7.41601 | 7.41314 | 6.102114 |
| 7.123794 | 6.446455 | 7.176719 | 5.637102 | 5.730085 | 7.15031 | 6.831966 | 7.224757 | 7.242532 |
| 6.917255 | 6.823484 | 6.87496 | 5.99284 | 6.215088 | 7.371492 | 7.677745 | 7.56173 | 5.560658 |
| NaN | 4.049757 | NaN | NaN | NaN | NaN | 5.810244 | NaN | 6.624759 |
| 8.787066 | 8.569134 | 8.560229 | 8.271239 | 8.317813 | 7.941833 | 7.8601 | 8.487037 | 8.205985 |
| 5.798172 | 5.119312 | 6.188488 | 3.982018 | 3.93174 | 6.330902 | NaN | 6.295142 | 5.523551 |
| 7.896675 | 7.668511 | 7.736679 | 10.67224 | 10.65663 | 8.783411 | 9.131457 | 8.875918 | 9.591593 |
| 6.858626 | 6.185302 | 6.106677 | NaN | NaN | 6.392827 | 6.536328 | 6.37497 | 5.873171 |
| 7.642228 | 7.324463 | 7.557616 | 9.992662 | 9.774106 | 8.126906 | 8.489618 | 7.650503 | 7.2949 |
| 7.027032 | 6.853193 | 7.431043 | 6.247064 | 5.82882 | 7.516376 | 7.533149 | 7.37358 | 6.619514 |
| NaN | NaN | NaN | 6.860152 | 6.786246 | 6.052086 | 6.28987 | 5.996985 | 5.068933 |
| 7.448589 | 7.564228 | 7.671841 | 7.789034 | 7.904162 | 7.361316 | 7.501277 | 7.693783 | 7.960977 |
| 8.853571 | 8.830115 | 8.841619 | 8.267196 | 8.674566 | 8.956869 | 9.04374 | 9.102336 | 8.820312 |
| 8.584586 | 8.250248 | 8.118208 | NaN | 6.592686 | 8.416129 | 8.262224 | 8.411889 | 7.809661 |
| 7.03691 | 6.785413 | 6.38936 | 5.765275 | 6.310692 | 6.849425 | 6.806784 | 7.054924 | 6.295866 |
| 11.21007 | 11.2333 | 11.30985 | 9.72238 | 9.602661 | 11.39311 | 11.42886 | 11.43189 | 10.94618 |
| NaN | NaN | NaN | NaN | 6.314138 | NaN | NaN | NaN | 6.187917 |
| 6.937623 | 6.970297 | 6.890265 | 7.443206 | 7.229299 | 6.001332 | 6.241761 | 5.381725 | 6.295287 |
| 8.740675 | 8.930502 | 8.183968 | 6.341485 | 6.823285 | 7.804577 | 7.607486 | 8.374349 | 7.87385 |
| 6.568963 | 6.748623 | 6.909207 | 5.66023 | 5.502708 | 7.546376 | 7.800069 | 8.021501 | 6.529725 |
| 8.574566 | 8.331947 | 8.523407 | 8.153663 | 8.014439 | 8.167906 | 8.538235 | 8.34399 | 8.285173 |
| 7.765808 | 7.710748 | 7.459655 | 5.57249 | 6.865651 | 8.025726 | 8.085331 | 7.881487 | 7.56765 |
| 6.878891 | 6.034692 | 6.242599 | 6.28174 | NaN | 7.52488 | 6.597435 | 6.409813 | 6.531069 |
| 7.533732 | 7.633346 | 7.743907 | 6.777457 | 6.51724 | 7.169246 | 7.297856 | 7.468455 | 7.274281 |
| 6.422279 | 6.852759 | 7.250743 | 6.589819 | 6.578535 | 7.90581 | 7.289693 | 7.39046 | 7.52184 |
| 6.792974 | 6.250803 | 7.052714 | 5.917376 | 7.0651 | 6.420253 | 6.246522 | 6.65578 | 7.229274 |
| 7.773016 | 7.912402 | 7.960028 | 8.346198 | 8.661188 | 8.423769 | 8.069138 | 8.950892 | 7.640615 |
| 7.43139 | 7.42952 | 7.577982 | 6.920924 | 7.025947 | 7.751433 | 7.753293 | 7.733075 | 6.927955 |
| 6.736415 | 6.821483 | 6.604793 | 5.687888 | 5.922426 | 7.161206 | 6.967135 | 7.309294 | 7.023616 |
| 8.179377 | 6.846281 | 6.888633 | 7.853384 | 7.392296 | 6.460992 | NaN | 6.607092 | 6.795189 |
| 8.567664 | 7.917701 | 8.527744 | 7.522997 | 8.017554 | 8.410312 | 8.513263 | 8.740756 | 8.182792 |
| 7.152339 | 7.290303 | 7.268276 | 6.558468 | 6.569818 | 7.085744 | 7.192727 | 7.159836 | 6.772687 |

| | | | | | | | | |
|---|---|---|---|---|---|---|---|---|
| 5.888011 | 5.372732 | 6.006557 | 4.804509 | 5.200474 | 5.100393 | 3.31596 | 5.632679 | 4.7333 |
| NaN | 4.084517 | 7.090995 | NaN | 3.105755 | 4.57171 | 4.286025 | 4.771079 | 4.611641 |
| NaN | NaN | NaN | 6.280682 | 6.596733 | NaN | NaN | 6.534504 | 7.020124 |
| 7.379828 | 7.261281 | 7.409498 | 7.014393 | 7.161732 | 7.021369 | 7.140635 | 7.035219 | 7.130384 |
| 6.680114 | NaN | NaN | NaN | 5.078989 | NaN | 5.871432 | 5.891088 | 5.82871 |
| 7.25664 | 6.818927 | 7.05933 | 6.620054 | 6.54437 | 7.48528 | 6.317546 | 7.343809 | 6.854004 |
| 7.21114 | 7.22277 | 7.332585 | 6.990823 | 7.073304 | 7.612347 | 7.552453 | 7.708285 | 7.149197 |
| 6.771129 | 6.315021 | 6.743631 | 5.755552 | 6.321565 | 7.053775 | 7.037845 | 7.29024 | 7.197975 |
| 6.529139 | 6.871686 | 7.075567 | 6.146727 | 5.99938 | NaN | 6.070981 | NaN | 6.278534 |
| 6.816062 | 5.099722 | 6.807085 | 6.448339 | 5.965705 | 5.052885 | 5.502609 | 5.52072 | 5.752543 |
| 8.249034 | 7.757643 | 8.251412 | 7.482251 | 7.819302 | 8.233582 | 8.24956 | 8.390719 | 8.521802 |
| 7.128859 | 7.19389 | 7.337987 | 7.316514 | 7.707589 | 7.447194 | 7.18967 | 7.799039 | 7.755035 |
| 7.924599 | 8.329175 | 8.08095 | 6.815439 | 6.626319 | 8.138273 | 8.213293 | 8.13041 | 7.73533 |
| 6.259066 | 5.947429 | 6.379921 | 6.766351 | 6.8268 | 7.040435 | 6.986669 | 7.279492 | 6.404031 |
| 6.061189 | 6.055002 | 5.911504 | NaN | 3.976392 | 6.944008 | 7.064931 | 6.585083 | 5.511749 |
| 6.523438 | 6.888655 | 6.775448 | 6.750249 | 6.312405 | NaN | 6.233933 | NaN | NaN |
| 6.517049 | 6.223715 | 6.828743 | 5.569602 | 5.521073 | 6.942559 | 7.007486 | 7.423545 | 5.206816 |
| 5.533911 | 5.38065 | 5.432193 | NaN | NaN | 5.54586 | 5.921548 | 5.380681 | 5.540698 |
| 6.09674 | 6.345595 | 6.124482 | 5.714387 | 5.370025 | 6.454152 | 6.503881 | 6.404858 | 5.93232 |
| NaN | NaN | NaN | 3.657265 | 3.778264 | NaN | NaN | NaN | NaN |
| 7.800476 | 7.665713 | 7.738282 | 7.689539 | 7.890429 | 7.811161 | 7.571304 | 7.696274 | 8.528754 |
| 7.996156 | 8.005362 | 8.006145 | 7.130142 | 6.857415 | 6.776657 | 7.160644 | 6.190296 | 7.369781 |
| 5.720709 | 5.607788 | 5.887095 | 5.893035 | 6.076838 | 5.904799 | 5.573413 | 6.150329 | 6.718816 |
| 5.872174 | 6.661622 | 6.105655 | 5.91074 | 5.278548 | 6.079096 | 5.614993 | 5.398163 | 6.398368 |
| 4.921386 | 4.497239 | 5.397578 | 6.410055 | 6.583202 | 5.147776 | 5.780834 | 5.604901 | 5.360724 |
| 8.120427 | 7.668198 | 8.158016 | 7.31932 | 6.844317 | 7.271061 | 7.225868 | 7.594587 | 8.416008 |
| 9.051825 | 9.262012 | 9.331893 | 10.96603 | 11.06419 | 9.478503 | 9.671188 | 9.60928 | 8.624839 |
| NaN | NaN | NaN | NaN | 5.770841 | 7.279517 | 7.329202 | NaN | 6.448352 |
| NaN | NaN | NaN | 6.623424 | 6.526823 | NaN | NaN | NaN | 5.505704 |
| 7.952658 | 7.813446 | 7.83389 | 7.770125 | 7.748834 | 7.501064 | 7.315107 | 7.662628 | 6.675534 |
| 6.580364 | 6.844404 | 7.051852 | 5.444601 | NaN | 7.385732 | 7.153425 | 7.427324 | 6.71914 |
| 8.004243 | 7.546901 | 7.931698 | 7.46644 | 7.394457 | 7.151786 | 7.176588 | 7.051469 | 7.852473 |
| 6.294942 | 6.083394 | 6.252335 | 5.693499 | 5.020534 | 5.907769 | NaN | 5.355934 | 5.55366 |
| 8.453631 | 8.468758 | 8.367193 | 8.226204 | 7.927982 | 8.470706 | 8.421038 | 8.358619 | 8.075043 |
| 7.601677 | 7.888752 | 7.465818 | 6.439296 | 6.623177 | 7.489591 | 7.573784 | 7.797012 | 7.466991 |
| 11.05613 | 10.57971 | 10.64588 | 7.647183 | 7.454003 | 6.648427 | 6.768594 | 6.742088 | 6.96369 |
| 8.236405 | 8.020769 | 8.276875 | 7.103409 | 7.33843 | 5.87803 | 6.716868 | 6.833428 | 7.07164 |
| 6.530302 | 7.20175 | 7.194783 | 6.280412 | 5.730949 | 5.446832 | 6.123788 | 5.721798 | 6.465917 |
| 7.001351 | 6.463696 | 6.713881 | 6.832549 | 6.94236 | 6.917721 | 7.331493 | 7.192733 | 6.426181 |
| 6.210968 | 5.464201 | NaN | 6.78824 | 6.890842 | 6.868652 | 6.391771 | 6.920434 | 5.913819 |
| 6.671248 | 6.570154 | NaN | 5.631898 | 6.711994 | 6.859459 | 6.928678 | 6.876613 | 6.627267 |
| 9.018952 | 8.797739 | 8.898582 | 9.354126 | 9.326731 | 8.747237 | 8.603688 | 8.617785 | 8.257617 |
| 11.77235 | 8.366737 | 12.0619 | 13.25174 | 13.34816 | 13.20692 | 13.13093 | 12.7568 | 11.53723 |
| 7.427801 | 6.986415 | 6.998419 | 6.654331 | 6.800282 | 7.404664 | 7.327035 | 7.233201 | 7.199263 |
| 6.854517 | 6.302401 | 7.01324 | 7.317169 | 7.482382 | NaN | NaN | 7.072174 | 7.198578 |
| 5.897463 | 6.208326 | NaN | NaN | NaN | 6.765004 | 7.096242 | 7.544479 | 6.70438 |
| 7.764285 | 7.064565 | 7.35301 | 6.57483 | 7.63202 | 7.926934 | 7.585032 | 7.779272 | 5.530711 |

| | | | | | | | | |
|---|---|---|---|---|---|---|---|---|
| 11.42605 | 10.9849 | 11.20629 | 11.83021 | 11.58256 | 8.471002 | 8.530727 | 7.619278 | 7.14849 |
| 7.300487 | 7.272338 | 7.171462 | 6.606308 | 7.316341 | 7.140019 | 7.01473 | 7.050875 | 7.210424 |
| NaN | NaN | NaN | 5.274176 | 6.072901 | NaN | NaN | NaN | NaN |
| 6.185783 | 5.216405 | 6.861891 | 5.500279 | 5.623812 | 6.17635 | 6.157046 | 5.705309 | 7.98813 |
| 11.28302 | 11.27838 | 11.24747 | 10.89518 | 10.68758 | 10.54792 | 10.4374 | 10.43064 | 9.920565 |
| 7.82517 | 6.016292 | 7.13649 | 6.374732 | 6.113285 | 6.842163 | 6.845794 | 6.950391 | 7.434235 |
| 6.177854 | 5.971461 | 6.182797 | 6.149827 | 6.020456 | 7.10002 | 6.923215 | 6.949196 | 5.985107 |
| 7.817258 | 7.698048 | 7.580051 | 6.203317 | 6.152388 | 7.27032 | 7.823499 | 7.58025 | 7.510235 |
| 9.14526 | 8.750268 | 8.594028 | 9.910274 | 10.27859 | 8.090993 | 8.210989 | 8.302067 | 7.820106 |
| 5.680002 | 6.131721 | 6.152886 | 6.989573 | 6.816312 | 6.092885 | 6.001577 | 5.789292 | 5.875544 |
| NaN | 5.79228 | 5.467397 | NaN | NaN | 7.68011 | 7.924906 | 7.759748 | 5.865043 |
| 6.034926 | 5.493572 | 5.360166 | 8.071556 | 8.247363 | 8.372365 | 8.364733 | 8.60391 | 8.088011 |
| 7.506173 | 6.85928 | 7.663257 | 6.719995 | 6.736181 | 7.694553 | 7.774508 | 7.492047 | 6.552901 |
| 6.196632 | 5.983404 | NaN | NaN | NaN | 6.633801 | 6.526119 | 6.61235 | NaN |
| NaN | 5.988639 | NaN | 6.260492 | 5.893554 | NaN | 6.006024 | 6.547035 | 5.964326 |
| NaN | 5.829876 | 6.240234 | NaN | 6.382738 | NaN | NaN | NaN | 7.540344 |
| 7.476394 | 7.247399 | 7.457532 | 8.710836 | 8.74056 | 6.316246 | 6.998856 | 6.685931 | 7.88631 |
| 6.4927 | 6.489302 | 6.317297 | 5.574563 | 5.923218 | 7.607275 | 7.378634 | 6.940061 | 6.426605 |
| 6.608921 | 6.915936 | 6.626017 | 6.715984 | 6.473833 | 6.17941 | 5.999823 | 6.895961 | 6.211933 |
| 6.69085 | 6.080917 | NaN | NaN | NaN | NaN | NaN | NaN | 7.20453 |
| 7.200331 | 7.085792 | 7.134495 | 6.755403 | 7.530094 | 6.854527 | 7.201824 | 6.878232 | 7.399684 |
| 8.223273 | 7.974263 | 7.970986 | 8.143388 | 8.250946 | 8.883102 | 8.816677 | 8.87917 | 7.350086 |
| NaN | 5.970973 | NaN | 6.215724 | 7.068676 | NaN | NaN | NaN | 5.666322 |
| 8.694203 | 7.508577 | 7.421498 | 6.468502 | 5.692884 | 7.615268 | NaN | 8.353141 | NaN |
| 6.226357 | 5.657159 | 5.626057 | 5.123746 | 5.064677 | 6.173035 | 6.399268 | 6.115307 | 5.793349 |
| 8.316628 | 8.482198 | 8.497965 | 5.64481 | 5.824031 | 6.741867 | 6.594414 | 6.766735 | 7.157136 |
| 8.653187 | 9.445336 | 8.925876 | 8.147108 | 7.806519 | 8.590703 | 8.748325 | 8.469488 | 8.726013 |
| 4.744295 | 4.941562 | 4.87848 | NaN | NaN | 5.454107 | 5.967442 | 6.487878 | 4.980826 |
| 8.084942 | 7.970429 | 7.811477 | 6.823597 | 7.41283 | 7.439662 | 7.334603 | 7.22274 | 7.458807 |
| 7.049651 | 5.893544 | 6.749969 | 6.696314 | 6.545838 | 6.651336 | 5.955984 | 6.312054 | 6.83448 |
| 3.575245 | 3.814864 | 3.609024 | NaN | 5.405639 | 5.662374 | 5.2143 | NaN | 5.137915 |
| 6.710107 | 6.786618 | 6.724505 | 6.253866 | 7.031082 | 6.28154 | 6.138373 | 6.330238 | 6.565403 |
| NaN | NaN | NaN | 7.464993 | 5.258544 | NaN | NaN | NaN | 6.7414 |
| 7.486563 | 7.303182 | 7.31772 | 7.428758 | 7.804045 | 7.202496 | 7.459456 | 7.387632 | 6.919888 |
| 6.929265 | 5.69969 | 7.026789 | 7.049637 | 7.017475 | NaN | 7.120851 | NaN | 6.695906 |
| 7.104117 | 5.597361 | 6.614512 | 5.848293 | 4.499948 | 6.943669 | 6.71894 | 6.852521 | 6.136565 |
| 6.09443 | 6.439843 | NaN | 6.529469 | 6.864608 | 7.066292 | 7.051875 | NaN | 7.447077 |
| 7.407298 | 7.163584 | 7.666808 | 8.120366 | NaN | 7.475485 | 7.439115 | 7.572748 | 6.670472 |
| 7.231027 | 6.875483 | 7.226655 | 6.934847 | 7.017514 | 7.985925 | 7.609397 | 7.937094 | 6.204074 |
| 7.274665 | 7.630701 | 7.331109 | 7.053203 | 7.172783 | 7.388396 | 7.343791 | 7.258027 | 7.154679 |
| 6.598201 | 6.884478 | 6.786783 | 5.153744 | 4.061913 | 6.619039 | 6.673123 | 6.81621 | 6.020029 |
| NaN | NaN | NaN | NaN | NaN | NaN | NaN | NaN | 4.823669 |
| 7.427722 | 6.981053 | 7.114153 | 7.948132 | 7.840115 | 7.12856 | 7.212916 | 7.382766 | 6.978596 |
| NaN | NaN | NaN | NaN | NaN | 5.92291 | 7.098825 | 7.032577 | NaN |
| 7.985621 | 7.978202 | 8.011664 | 8.052626 | 7.878839 | 7.870964 | 8.071024 | 7.960929 | 7.833383 |
| 7.221453 | 6.954414 | 7.059477 | 6.12156 | 6.927538 | 7.667186 | 7.415611 | 7.651435 | 5.808736 |
| 4.95566 | 5.160591 | 5.467359 | 3.762314 | 3.858856 | 4.542842 | 4.576649 | 4.74458 | 4.103963 |

| | | | | | | | | |
|---|---|---|---|---|---|---|---|---|
| 6.789735 | 6.604632 | 6.104127 | 6.622193 | 6.557232 | 6.421966 | 5.629484 | 5.536047 | 6.72609 |
| 7.991876 | 7.953065 | 7.916377 | 7.753753 | 7.615968 | 6.785837 | 6.983738 | 6.971362 | 7.83384 |
| 9.34238 | 9.32693 | 9.36027 | 9.201572 | 9.242969 | 8.875711 | 8.875422 | 8.958794 | 9.457266 |
| 5.725632 | 6.422842 | 6.421712 | 5.693212 | 5.6048 | 6.019072 | 5.534102 | NaN | 5.266988 |
| 8.178782 | 8.245571 | 7.953496 | 8.045324 | 7.70775 | 8.23468 | 7.96153 | 7.93234 | 7.885098 |
| 6.225918 | 6.691531 | 6.39887 | 5.105509 | 5.696793 | 6.153969 | 6.473944 | 6.518323 | 5.271843 |
| 7.596373 | 6.865509 | 7.334416 | 5.859263 | 6.571799 | 7.286244 | 7.528388 | 7.740611 | 6.385226 |
| 6.688386 | 6.964076 | 6.684654 | 5.570867 | 5.597311 | 6.990903 | 6.774021 | 6.875712 | 6.556557 |
| 7.905405 | 6.563138 | 7.402765 | 8.048393 | 7.224047 | 5.435606 | 6.34637 | NaN | 6.109407 |
| 7.785887 | 7.941006 | 8.025688 | 6.716766 | 6.438089 | 7.289102 | 7.199667 | 7.609235 | 7.749162 |
| 6.689902 | 5.776321 | 6.000214 | 5.636254 | 6.161606 | 5.99949 | 5.934213 | 6.31876 | 6.030519 |
| 8.16818 | 7.769238 | 8.183575 | 8.170746 | 6.968751 | 7.528873 | 7.901637 | 7.764138 | 7.596208 |
| 8.162099 | 8.340768 | 8.258777 | 8.160759 | 8.333344 | 8.773308 | 8.591799 | 8.80135 | 8.085158 |
| 7.468739 | 7.306056 | 7.638252 | 7.65857 | 7.478029 | 7.476851 | 7.256181 | 7.426549 | 8.172834 |
| 9.441957 | 9.240818 | 9.221464 | 9.761596 | 9.666849 | 7.953687 | 8.229323 | 8.08174 | 9.015729 |
| 8.844184 | 8.978209 | 9.047276 | 9.868858 | 9.86492 | 8.089454 | 7.738961 | 8.080065 | 8.259622 |
| 7.546033 | 7.530887 | 8.392594 | 5.827534 | NaN | 6.138917 | 6.639804 | 6.636191 | 6.868697 |
| 6.10976 | 6.51244 | 5.837564 | 8.192647 | 6.088737 | 5.822011 | 6.134408 | 5.64423 | 7.558929 |
| 6.994564 | 8.338736 | 7.354229 | 8.768028 | 8.742052 | NaN | 6.836287 | 7.376678 | 6.525319 |
| 7.583174 | 7.75329 | 7.656996 | 8.496095 | 8.729374 | 8.148146 | 7.871559 | 7.497788 | 6.21771 |
| NaN | 6.002028 | NaN | NaN | NaN | 7.232976 | 6.520855 | 6.768693 | NaN |
| 8.117884 | 8.040872 | 8.097745 | 7.426712 | 7.445074 | 8.012265 | 7.937614 | 7.8008 | 7.879857 |
| 5.183225 | NaN | 4.99624 | NaN | NaN | 5.491377 | 5.621264 | 6.009634 | 5.775004 |
| 6.523177 | 7.023539 | 6.645296 | 5.763546 | 6.767362 | 7.425602 | 7.413064 | 7.346277 | 6.963204 |
| 6.912419 | 6.883801 | 6.799256 | 6.109182 | 5.840325 | 6.585856 | 6.292805 | 6.640725 | 6.666062 |
| 7.179577 | 7.560057 | 8.052628 | NaN | 6.340038 | 6.474753 | 6.135178 | 6.161048 | 6.72325 |
| 5.501779 | 4.17743 | 5.471941 | 6.38608 | 6.607528 | 6.019873 | NaN | 5.214307 | 6.105173 |
| 4.543694 | NaN | NaN | 5.133929 | 4.661774 | 5.631825 | NaN | 4.969112 | 5.913317 |
| 6.463221 | 7.050359 | 6.215103 | 6.276929 | 6.498034 | 7.132065 | 6.588474 | 6.690824 | 6.738308 |
| 7.620495 | 7.761789 | 7.757102 | 7.349165 | 6.475915 | 6.774751 | 6.431469 | 6.391637 | 6.880459 |
| 7.929583 | 7.907196 | 8.046435 | 6.528564 | 6.180614 | 8.592591 | 8.530069 | 8.420403 | 7.733305 |
| 6.633957 | 6.786865 | 6.895419 | 7.146908 | 7.153904 | 7.303295 | 6.912858 | 7.334522 | 7.184224 |
| 7.700293 | 6.303821 | 7.182815 | NaN | 6.253159 | 7.860519 | 7.818341 | 8.227431 | 7.135263 |
| 8.463508 | 8.362654 | 7.990802 | 8.950728 | 9.026152 | 9.010387 | 8.955032 | 9.03058 | 8.8049 |
| 8.731382 | 8.385366 | 8.313275 | 7.891351 | 7.203358 | 8.335965 | 8.135063 | 7.697785 | 7.761117 |
| 7.781734 | 7.564497 | 7.642819 | 6.811953 | 6.563015 | 8.074018 | 7.9653 | 7.86724 | 7.307574 |
| 6.723106 | 6.731844 | 7.066051 | 5.725553 | 6.451801 | 7.25024 | 6.81438 | 6.822357 | 7.004347 |
| 6.567113 | 6.729149 | 6.598208 | 6.475691 | 6.06063 | 6.579364 | 6.73752 | 6.997935 | 6.731864 |
| 7.790717 | 7.881099 | 7.973481 | 7.377954 | 7.306803 | 8.281818 | 8.164224 | 8.401364 | 7.585998 |
| 7.452304 | 6.837756 | 7.098362 | NaN | NaN | 7.109073 | 6.803734 | 6.632966 | 7.79342 |
| 6.452173 | 4.589908 | 6.658388 | 6.387301 | 6.555115 | NaN | 6.514555 | 6.542186 | 7.21534 |
| 4.849425 | 5.509066 | 5.36937 | NaN | NaN | 6.033606 | 6.381358 | 6.377954 | 6.759165 |
| 6.229815 | 4.909162 | 5.455922 | NaN | 3.851094 | 5.722819 | 5.38615 | 5.768878 | 5.731895 |
| 7.823124 | 7.17713 | 7.274965 | 6.211771 | NaN | 7.0639 | 7.139771 | 7.407297 | 6.550241 |
| 6.067308 | 5.677827 | 5.851197 | 3.85171 | 3.388412 | 6.636735 | 6.724656 | 6.875652 | 6.525166 |
| NaN | 5.726041 | NaN | NaN | NaN | 6.429767 | 6.516528 | 6.509206 | 6.288421 |
| 7.402761 | 7.250517 | 7.196754 | 6.459334 | 6.460874 | 7.526838 | 6.840173 | 7.197175 | 7.276228 |

| | | | | | | | | |
|---|---|---|---|---|---|---|---|---|
| 5.785537 | NaN | 5.911961 | NaN | NaN | NaN | 5.592951 | 5.125231 | 5.966588 |
| 7.000837 | 6.791044 | 7.123236 | 7.945217 | 7.837306 | 6.652614 | 6.337471 | 7.027336 | 6.00563 |
| 7.580632 | 7.401126 | 7.522797 | 7.644732 | 7.684746 | 7.09479 | 7.422519 | 7.445141 | 7.205711 |
| 6.251722 | 6.473715 | 6.029877 | 5.995344 | 5.978158 | 6.098863 | 6.30907 | 6.265131 | 5.998472 |
| 7.298009 | 6.165345 | 6.318653 | 5.640006 | NaN | 7.230298 | 6.850458 | 7.280291 | 6.302734 |
| 7.505803 | 7.330344 | 7.44534 | 6.21435 | 6.296046 | 7.462893 | 6.580247 | 7.130517 | 7.404701 |
| 7.111837 | 7.370754 | 6.755916 | 6.65473 | 6.312386 | 7.37649 | 7.37102 | 7.178436 | 7.139766 |
| 5.088117 | 3.076089 | 5.957405 | 5.424932 | 5.621666 | 5.796175 | 5.299252 | 4.568489 | 5.8787 |
| 4.294847 | 5.826491 | 3.123214 | 6.221091 | 6.358383 | 4.795449 | 6.2925 | 5.493971 | NaN |
| 6.140022 | 5.958749 | 6.029059 | 4.837259 | 4.973583 | 5.991086 | NaN | 5.759592 | 5.737315 |
| 7.321151 | 7.266003 | 7.337617 | 6.565704 | 5.959065 | 7.368525 | 7.159147 | 7.323286 | 6.515275 |
| 8.886113 | 8.952489 | 9.008862 | 7.80875 | 7.972167 | 7.949534 | 8.174022 | 8.287245 | 8.200804 |
| 4.182761 | NaN | 4.271507 | NaN | NaN | 4.44615 | NaN | NaN | 4.743303 |
| 7.749218 | 7.751812 | 7.666001 | 8.78523 | 8.752632 | 7.280796 | 7.44113 | 7.111521 | 6.804615 |
| NaN | NaN | NaN | 6.021846 | 6.160154 | 5.659173 | 5.594091 | 5.922355 | 5.256019 |
| 5.585435 | 6.207813 | 6.365372 | NaN | 4.231175 | 6.013908 | 5.654151 | 5.702428 | 5.554327 |
| 8.171389 | 8.037748 | 8.27963 | 8.807218 | 9.130639 | 8.265915 | 7.714979 | 8.070951 | 9.021201 |
| 6.665589 | 6.281217 | 6.611488 | 7.351821 | 6.846978 | 7.31067 | 7.283256 | 7.413344 | NaN |
| NaN | 7.613455 | 8.080108 | 9.054731 | NaN | 8.389244 | 8.102554 | 8.043396 | 7.523309 |
| 10.35634 | 6.011851 | 10.01678 | 7.727423 | 7.825656 | 10.42341 | 10.63272 | 10.47423 | 9.981317 |
| 9.070045 | 8.990239 | 9.007213 | 7.702369 | 8.033496 | 7.834513 | 8.545136 | 8.048501 | 7.709194 |
| NaN | 5.408548 | 5.940851 | NaN | NaN | 6.0824 | 6.371914 | 6.558446 | 6.221641 |
| 8.146895 | 7.993311 | 8.235454 | 6.741069 | 6.60533 | 8.334199 | 7.846178 | 8.075455 | 7.044996 |
| 6.034235 | 6.008319 | 6.014875 | 5.298503 | 5.006299 | 5.528896 | 4.886106 | 5.689297 | 5.814682 |
| 7.352495 | 7.267871 | 7.191206 | 7.922011 | 8.339059 | 7.997654 | 7.646036 | 7.967902 | 8.190914 |
| NaN | NaN | NaN | NaN | 6.373628 | NaN | NaN | NaN | 5.844645 |
| 7.52757 | 7.452891 | 7.521385 | 6.726034 | 6.821163 | 7.599097 | 7.666659 | 7.639673 | 7.308862 |
| 10.19898 | 10.4173 | 10.41136 | 11.48801 | 11.34888 | 10.93951 | 10.63611 | 10.59952 | 8.439801 |
| 7.129699 | 7.181112 | 7.261308 | 6.972167 | 7.30577 | 7.480223 | 7.35296 | 7.540204 | 7.145934 |
| 8.29702 | 8.31611 | 8.15967 | 7.429198 | 7.565602 | 8.746162 | 8.88025 | 8.787141 | 7.574789 |
| 7.573179 | 7.376286 | 7.173475 | 7.997808 | 8.068085 | 7.130885 | 6.582258 | 6.448906 | 6.470286 |
| 7.336759 | 7.673329 | 7.343602 | 8.160684 | 8.746255 | 7.526577 | 7.130974 | 7.236406 | 8.558758 |
| 2.330304 | NaN | 1.568245 | 1.812481 | 2.268698 | 5.418114 | 4.863419 | 5.585687 | 3.471311 |
| 5.468336 | 5.278255 | 5.228136 | 6.788777 | 6.468759 | 6.115982 | 5.96033 | 5.957113 | 7.072897 |
| 6.286852 | 7.399021 | 7.295553 | 6.938526 | 7.008309 | 7.703455 | 7.467036 | 7.522357 | NaN |
| 9.892739 | 9.697572 | 9.788108 | 7.179101 | 7.472442 | 9.156806 | 9.218656 | 9.140636 | 8.80691 |
| 9.321353 | 9.323791 | 9.166446 | 9.019378 | 8.90794 | 8.876705 | 8.795082 | 8.745754 | 9.283097 |
| 7.468552 | 7.375791 | 6.903039 | 9.263467 | 9.234322 | 6.74516 | 5.905439 | 6.358006 | 6.373435 |
| 7.869306 | 7.59151 | 7.45279 | 5.992942 | 7.254714 | 8.053426 | 8.248343 | 8.177923 | 7.104808 |
| 10.54923 | 10.52818 | 10.41452 | 10.03317 | 9.567986 | 9.775073 | 9.893639 | 9.857106 | 10.08439 |
| 6.100749 | 6.45331 | 6.540871 | 6.394114 | 5.63493 | 6.965417 | 6.823495 | 7.225194 | 6.733152 |
| 7.929779 | 7.813627 | 7.58015 | 6.472714 | 6.222484 | 7.067583 | 6.771365 | 7.371256 | 6.998011 |
| 6.924998 | 7.097438 | 6.956809 | 4.798211 | 5.65846 | 6.634642 | 6.580655 | 6.629938 | 6.674376 |
| 10.36977 | 10.32984 | 10.22481 | 10.1348 | 9.828806 | 10.14967 | 10.33917 | 10.26793 | 9.92383 |
| 5.821593 | 5.446986 | 5.971274 | 5.836586 | 6.359867 | 5.866137 | 5.786185 | 5.844635 | 5.563239 |
| 12.3416 | 12.46104 | 12.50351 | 10.44086 | 9.545329 | 8.804281 | 9.236267 | 8.899534 | 8.84574 |
| 7.266353 | 7.109464 | 7.258449 | 6.590095 | 6.876358 | 7.468873 | 7.21318 | 6.915984 | 6.899989 |

| | | | | | | | | |
|---|---|---|---|---|---|---|---|---|
| NaN | NaN | NaN | 7.382214 | 7.27588 | 7.75794 | 8.003798 | 7.908545 | 3.8184 |
| 8.143352 | 8.112973 | 8.353242 | 7.657686 | 7.777606 | 8.69759 | 8.683741 | 8.58693 | 7.720104 |
| 10.09855 | 8.538061 | 9.644162 | 9.868598 | 7.173743 | 8.234624 | 8.87893 | 7.942675 | 7.454688 |
| NaN | 3.876305 | 4.771861 | 3.5793 | 5.159221 | 4.391828 | 4.64908 | 4.951346 | 5.450282 |
| 8.320724 | 7.658387 | 7.958576 | 7.004507 | 6.704323 | 8.148086 | 8.186749 | 8.22851 | 7.312993 |
| 6.668333 | 6.579279 | 6.749022 | 7.622274 | 7.510188 | 6.906651 | 5.832989 | 5.934974 | 6.395877 |
| 6.590277 | 7.175753 | 6.853289 | 5.759914 | 6.331838 | 7.117245 | 6.888697 | 7.058079 | 6.389216 |
| 7.67051 | 7.762807 | 7.714602 | NaN | NaN | 7.526553 | 8.864051 | 8.820702 | 8.432267 |
| NaN | NaN | NaN | NaN | NaN | NaN | 3.098089 | NaN | 5.97437 |
| 7.658603 | 6.977469 | 7.370618 | 6.252332 | 7.108579 | 7.137943 | 7.140036 | 7.067587 | 7.539377 |
| 6.574485 | NaN | 6.796851 | NaN | NaN | 7.23764 | 6.982147 | 7.047461 | 6.215449 |
| 7.658511 | 7.947722 | 7.852393 | 9.315474 | 9.40276 | NaN | NaN | NaN | 5.682046 |
| 7.005022 | 6.058255 | 6.864268 | 6.581766 | 7.500414 | 6.605551 | 6.901481 | 6.179515 | 6.405297 |
| 6.763884 | 6.894479 | 7.252607 | 5.525328 | 6.214247 | 6.661855 | 7.259626 | 6.788062 | 6.206362 |
| 7.44648 | 6.605363 | 7.434256 | NaN | 5.597403 | 7.009137 | 7.258348 | 6.810503 | 6.553178 |
| 6.753758 | 6.931757 | 6.955395 | 6.670212 | 6.833741 | 6.67176 | 6.467871 | 6.559638 | 6.451848 |
| 6.97305 | 7.023531 | 6.968019 | 6.610109 | NaN | 6.77036 | 6.885212 | 7.029705 | 7.132861 |
| 6.640437 | 6.806372 | 6.493081 | 6.455543 | 6.629467 | 7.117131 | 7.608304 | 6.549874 | 6.259482 |
| 6.925045 | 6.75504 | 5.760542 | 4.644639 | NaN | 6.640686 | 6.488819 | 6.449229 | 5.427039 |
| 8.576901 | 7.586621 | 8.456231 | 6.95114 | 7.184404 | 7.868417 | 7.972072 | 7.845393 | 7.544046 |
| 6.670457 | 5.978532 | 7.00739 | NaN | NaN | NaN | 6.2907 | 6.141616 | 5.671101 |
| 6.591931 | 5.098299 | 6.312601 | 6.193086 | 6.423557 | 6.58595 | 6.456445 | 6.587917 | 6.31863 |
| 9.975339 | 9.155535 | 10.03906 | 7.809174 | 8.233077 | 10.3844 | 10.28403 | 10.35398 | 8.337087 |
| 6.650379 | 6.437035 | 6.500064 | 7.7778 | 7.465362 | 6.55162 | 6.341832 | 6.189504 | 6.586994 |
| 5.715339 | 6.547655 | 7.055248 | 7.956295 | 7.404685 | 7.604648 | 7.478899 | 7.468045 | 6.727518 |
| 4.847175 | NaN | NaN | 6.155737 | 6.863739 | NaN | 5.219954 | 5.55896 | 4.357805 |
| 7.73495 | 7.917836 | 8.020494 | 8.421584 | 8.533347 | 7.454306 | 7.180179 | 7.481868 | 6.517606 |
| 7.085946 | 6.080473 | 7.326095 | 5.751676 | 5.148039 | 6.905824 | 6.811499 | 6.749762 | 6.993348 |
| 6.84107 | 7.339245 | 6.044379 | 8.568069 | 8.776066 | 6.727069 | 7.063768 | 7.060772 | 6.664445 |
| 7.045004 | 7.905553 | 7.795855 | 7.044307 | 6.570661 | 7.774988 | 7.649073 | 7.538728 | 6.251232 |
| 7.40633 | 6.622265 | 6.406825 | 6.89202 | 7.223276 | 5.778888 | 6.255448 | 6.062713 | 4.186619 |
| 6.306216 | 5.397192 | 5.269525 | 7.175633 | 7.489316 | 7.276136 | 6.698619 | 6.533201 | 5.956456 |
| 7.197287 | 7.47632 | 7.464472 | 6.271656 | 6.588825 | 8.145826 | 7.982342 | 8.04788 | 5.375687 |
| 5.25156 | 6.129636 | 4.834186 | NaN | NaN | 6.578512 | 6.702152 | 6.581939 | NaN |
| 8.165673 | 8.292615 | 8.18643 | 8.127028 | 7.574307 | 7.013635 | 7.566931 | 7.564012 | 7.562381 |
| 8.359769 | 8.245996 | 8.274981 | 7.712439 | 7.881879 | 8.460725 | 8.507675 | 8.637747 | 8.195652 |
| 6.868623 | 7.103763 | 7.130683 | 8.224475 | 8.536954 | 6.762982 | 7.232271 | 7.384126 | 7.438302 |
| NaN | 7.240244 | 7.478895 | NaN | 6.328146 | 6.2282 | 6.681709 | 6.512387 | 6.93186 |
| 6.071692 | 5.636966 | 5.71593 | 5.495022 | 5.626451 | 6.182835 | 5.868332 | 6.201977 | 5.173126 |
| 5.873603 | 5.83989 | 5.641534 | 5.464539 | 5.477993 | 5.827402 | 5.160855 | 5.462514 | 5.332077 |
| 6.81923 | 4.716571 | 6.616514 | 5.520312 | 6.636955 | 6.81305 | 6.716456 | 6.878417 | 6.59457 |
| 6.514519 | 7.551935 | 6.568447 | 7.546892 | 8.584852 | 7.076089 | 7.308853 | 7.048446 | 7.397835 |
| 7.582026 | 7.384797 | 7.059462 | 9.891324 | 9.793231 | 6.749115 | 7.374162 | 7.563313 | 6.2756 |
| NaN | NaN | NaN | NaN | 4.463495 | 5.875149 | 7.11667 | 6.608775 | 5.640567 |
| NaN | NaN | NaN | 6.484083 | 7.00905 | 5.137764 | 5.856183 | NaN | 6.093322 |
| 7.612223 | 7.66434 | 7.443169 | 8.217041 | 8.869411 | 8.349725 | 8.106663 | 8.447831 | 8.000976 |
| 7.668778 | 7.710543 | 7.858746 | 7.309014 | 7.503841 | 6.922634 | 7.047054 | 7.032569 | 7.216359 |

| | | | | | | | | |
|---|---|---|---|---|---|---|---|---|
| 9.277863 | 9.280228 | 9.092542 | 9.123715 | 9.391565 | 8.465838 | 8.576756 | 8.481846 | 8.438643 |
| 8.226573 | 7.916278 | 8.17324 | 6.08642 | 9.310931 | 8.553935 | 8.146463 | 7.87086 | 8.130752 |
| 7.574681 | 7.529308 | 7.427891 | 8.504504 | 8.315825 | 7.430027 | 7.063515 | 7.440829 | 7.105121 |
| 5.761714 | 5.296342 | NaN | 5.588321 | 5.46261 | 6.574561 | 6.644356 | 6.583541 | 5.677243 |
| NaN | NaN | 5.241674 | NaN | NaN | NaN | NaN | NaN | 4.234208 |
| NaN | NaN | NaN | 5.990477 | 6.953598 | 6.170971 | NaN | 6.45738 | 6.205021 |
| 6.220536 | 6.146576 | 6.203437 | 5.620746 | 5.512539 | 7.362751 | 7.636294 | 7.437358 | 5.792339 |
| 7.912886 | 7.693634 | 7.718139 | 8.327014 | 8.59399 | 8.410068 | 8.44077 | 8.452312 | 7.7267 |
| 5.056743 | 5.221714 | 4.863612 | 5.784446 | 5.230035 | 6.264947 | 6.705652 | 6.505369 | 5.614526 |
| 9.967019 | 9.779593 | 9.80437 | 9.762822 | 9.488431 | 8.886996 | 9.087984 | 8.998056 | 8.599738 |
| NaN | 5.208273 | 5.795078 | NaN | NaN | 6.828654 | 6.558433 | 6.6127 | 5.934876 |
| 7.411681 | 7.392035 | 7.519098 | 8.485614 | 8.391414 | 6.96917 | 7.427871 | 7.40716 | 8.624327 |
| 7.651984 | 7.411227 | 7.834138 | 5.267759 | NaN | 6.739099 | 6.632147 | 6.154935 | 6.439736 |
| 6.714292 | 6.417437 | 6.38632 | 6.107041 | 5.904211 | 6.005281 | 6.23686 | 6.431428 | 6.601076 |
| 4.522516 | 4.777757 | 5.03157 | 3.290381 | 3.997286 | 6.921459 | 6.832477 | 6.560248 | 4.51724 |
| 6.565797 | 6.32202 | 6.424166 | 6.525193 | 6.752188 | NaN | 4.516609 | NaN | 4.510841 |
| 7.537096 | 7.297899 | 7.587907 | 8.496553 | 7.998489 | 6.436964 | 6.757383 | 6.601458 | 7.345877 |
| 7.474117 | 7.614307 | 7.618892 | 6.160906 | 6.449394 | 6.703938 | 7.182303 | 7.53202 | 6.97093 |
| 7.79799 | 8.016973 | 7.364484 | 7.693298 | 7.790236 | 7.358344 | 6.619673 | 6.704511 | 6.420468 |
| 8.558845 | 7.653847 | NaN | 6.954739 | NaN | 8.079974 | 8.01318 | 7.937788 | NaN |
| 9.168012 | 9.138918 | 8.989746 | 7.953399 | 7.983032 | 9.174767 | 9.319057 | 9.194911 | 9.246073 |
| 8.20228 | 8.159553 | 8.07199 | 8.940151 | 8.771444 | 8.095499 | 8.077347 | 8.161595 | 8.311062 |
| 7.481701 | 7.27488 | 7.429401 | 8.532326 | 8.035216 | 7.265857 | 7.130045 | 7.385959 | 7.225948 |
| 6.965884 | 6.701608 | 6.870673 | 7.039436 | 5.929694 | NaN | 6.420486 | 6.32812 | 6.817078 |
| 7.046736 | 6.738876 | 6.864418 | NaN | NaN | 6.384308 | 6.744673 | 6.581081 | 5.857707 |
| NaN | NaN | NaN | NaN | 3.684757 | 4.35961 | NaN | 3.907739 | 4.378998 |
| 7.56168 | 7.349695 | 7.749522 | 6.998488 | 7.100292 | 6.653021 | 6.817058 | 6.846917 | 6.073767 |
| 7.549568 | 7.605916 | 7.783447 | 8.167988 | 8.24249 | 8.763529 | 8.890096 | 8.748208 | 8.271688 |
| NaN | 6.306837 | 6.73423 | NaN | 6.674699 | 6.772249 | 7.049229 | 6.825048 | 6.404924 |
| 9.022092 | 9.087175 | 9.156321 | 10.35219 | 10.04974 | 8.729725 | 8.55898 | 8.677094 | 9.082043 |
| 8.980692 | 8.928142 | 9.039574 | NaN | 5.516584 | 8.459785 | 8.140289 | 8.14362 | 7.145393 |
| 5.795373 | 5.393117 | 5.405168 | 6.209573 | 6.767773 | 6.171501 | NaN | NaN | 5.758986 |
| NaN | NaN | 5.709116 | NaN | NaN | NaN | NaN | NaN | 6.095681 |
| 4.481776 | 5.08473 | NaN | NaN | NaN | 5.140878 | 5.562074 | 5.443727 | 5.386478 |
| 10.17015 | 9.447636 | 9.252032 | 8.882876 | 7.180038 | 7.892778 | 7.959193 | 7.641479 | 6.909246 |
| 8.537165 | 8.355712 | 8.345974 | 6.04493 | 6.391035 | 8.279719 | 8.113232 | 8.305674 | 7.559548 |
| 8.177801 | NaN | NaN | 6.735954 | 7.104606 | NaN | NaN | NaN | NaN |
| 7.05654 | 5.959073 | 6.502732 | 6.643842 | 7.438356 | 6.678889 | 6.75878 | 6.638918 | 6.740004 |
| 6.883282 | 7.113101 | 6.737745 | 6.931747 | 7.236611 | 6.644042 | 5.494401 | 6.440763 | 6.843691 |
| 7.769826 | 6.718381 | 7.611479 | 7.2252 | 7.34346 | 7.582335 | 7.378025 | 6.954929 | 6.941496 |
| 7.340503 | 7.70739 | 8.034987 | 8.496435 | 8.145336 | 7.138962 | 7.034006 | 7.900936 | 7.629916 |
| 7.131077 | 6.683854 | 6.651561 | 7.80742 | 7.688216 | 6.023052 | 5.988941 | 5.916208 | 6.659502 |
| NaN | NaN | NaN | NaN | 3.179349 | 3.244325 | 4.510943 | 2.82076 | 3.474175 |
| 7.770077 | 7.368208 | 8.041615 | 6.043372 | 5.788265 | 8.262816 | 8.122687 | 8.239002 | 6.552846 |
| 5.608322 | 5.46485 | 5.121635 | 5.810946 | 5.907778 | 6.334786 | 5.648656 | 6.031464 | 6.98498 |
| 7.309285 | 7.336358 | 7.213567 | 7.151689 | 7.232203 | 7.447686 | 7.395328 | 6.91326 | 7.650956 |
| 7.900023 | 8.017938 | 8.057592 | 7.356707 | 6.584506 | 8.871112 | 8.832986 | 8.563254 | 7.721352 |

| | | | | | | | | |
|---|---|---|---|---|---|---|---|---|
| 6.943433 | 7.18007 | 6.693881 | 6.429537 | 6.6047 | 7.422421 | 7.146515 | 7.531395 | 7.150299 |
| 8.800491 | 8.625453 | 8.419936 | 8.734964 | 8.545206 | 9.030576 | 7.386661 | 8.700657 | 8.539134 |
| 6.348809 | 9.262488 | 3.888438 | NaN | NaN | NaN | 8.21557 | NaN | 6.656594 |
| 7.643632 | 7.622391 | 7.718121 | 5.334953 | 6.491606 | 6.863699 | 6.814195 | 6.713164 | 6.376389 |
| 6.591342 | 6.052213 | 6.19176 | 5.47684 | 5.482818 | 6.073112 | 6.029788 | 5.662073 | 5.588519 |
| 7.66561 | 7.367156 | 7.449917 | 7.258996 | 7.433336 | 7.594332 | 7.47326 | 7.130398 | 7.292054 |
| 8.420006 | 8.333844 | 8.271063 | 8.425021 | 8.253942 | 8.202547 | 8.517304 | 8.562116 | 7.839818 |
| 7.466071 | 6.917931 | 7.599367 | 5.834734 | 6.254374 | 7.807219 | 8.095443 | 8.223458 | 7.328748 |
| 10.15089 | 10.16846 | 10.17083 | 8.890097 | 8.752177 | 8.319077 | 8.239842 | 8.096834 | 8.299631 |
| 8.206415 | 8.227664 | 8.268927 | 7.310713 | 7.623367 | 7.914591 | 8.035188 | 7.808624 | 8.944268 |
| 9.09673 | 9.015377 | 8.966436 | 10.12505 | 10.06508 | 9.131352 | 9.29208 | 9.30181 | 8.730834 |
| NaN | 5.62876 | 5.046375 | 5.121828 | 5.26364 | 4.533453 | NaN | 5.41398 | 5.154649 |
| 7.217435 | 6.810111 | 7.485684 | 6.471053 | 6.288249 | 7.232481 | 7.057798 | 7.112214 | 7.230933 |
| 6.166638 | 6.325805 | 6.159308 | 3.650176 | 3.720347 | 6.444056 | 6.035659 | 5.681495 | 6.0777 |
| 8.452367 | 8.677685 | 8.104449 | 7.964421 | 7.839771 | 8.905591 | 8.790406 | 9.159558 | 8.56928 |
| 6.062966 | 6.084135 | 6.702765 | 4.2556 | 6.883203 | 6.2467 | 6.525331 | 6.477809 | 5.869091 |
| 7.048282 | 6.782879 | 7.468373 | 6.045483 | 6.055604 | 7.450752 | 7.184305 | 7.088002 | 6.480242 |
| 5.97577 | 6.005501 | 5.995156 | 4.83717 | NaN | 6.564168 | 5.736053 | 5.800885 | 6.184595 |
| NaN | NaN | NaN | 7.030911 | 7.182544 | NaN | NaN | NaN | NaN |
| 8.695266 | 8.705096 | 8.746696 | 8.439763 | 8.486982 | 8.589171 | 8.723687 | 8.376461 | 9.347657 |
| 7.456044 | 6.978735 | 7.38689 | 6.868864 | 6.670748 | 6.543554 | 6.65577 | 6.611343 | 6.480213 |
| 8.524126 | 8.462654 | 8.628066 | 7.919805 | 7.881485 | 7.913788 | 7.940563 | 8.286351 | 8.326754 |
| 10.13824 | 9.792466 | 9.845156 | 8.67943 | 8.570752 | 10.09667 | 9.947033 | 9.729773 | 8.664562 |
| 6.22508 | 6.257946 | 6.018559 | 6.850583 | 7.013977 | 5.895354 | 6.2892 | 6.06753 | 6.250766 |
| 8.497885 | 8.6988 | 8.331264 | 6.844003 | 7.232199 | 7.643125 | 7.377068 | 7.878088 | 6.237676 |
| 7.460736 | 7.367914 | 7.170818 | 9.7579 | 9.750101 | 6.248887 | 6.403884 | 6.733588 | 6.804405 |
| NaN | NaN | NaN | NaN | 4.261141 | NaN | 4.612891 | 4.311554 | 5.816642 |
| 6.955955 | 6.975325 | 7.838785 | 5.829895 | 7.094882 | 7.142393 | 7.3616 | 7.047511 | 6.95667 |
| 8.436584 | 8.096742 | 8.149683 | 9.805294 | 10.06401 | 8.182073 | 8.138408 | 7.851737 | 7.600853 |
| 8.104525 | 7.665785 | 8.031389 | 7.661877 | 7.715157 | 8.320088 | 8.367554 | 8.425112 | 8.058948 |
| 8.021825 | 8.169705 | 8.481781 | 9.308613 | 7.704289 | 7.329074 | 7.485267 | 7.501899 | 6.618059 |
| 6.930264 | 6.95128 | 6.818201 | 6.228229 | 6.972302 | 6.893775 | 6.900614 | 6.870258 | 6.542016 |
| 6.893912 | 6.863245 | 6.265345 | 6.945556 | 6.788071 | 6.370648 | 6.1093 | 6.397258 | 6.992398 |
| 5.318381 | 5.736456 | 5.600137 | 6.188803 | 6.547916 | 6.15488 | 5.573395 | 5.834378 | 5.915842 |
| 6.572337 | 6.006866 | 5.936431 | 5.975535 | NaN | 5.826923 | 5.686365 | 5.593238 | 5.234412 |
| 6.497471 | 6.055328 | 6.012896 | NaN | 6.127028 | 4.563381 | 7.114555 | 7.354803 | 6.486015 |
| 6.00237 | 4.093189 | 6.368128 | 7.583164 | 6.458658 | 6.759684 | 6.374118 | 6.382406 | 6.320658 |
| 7.524944 | 7.808878 | 7.669036 | 7.131797 | 6.915366 | 8.175489 | 8.313643 | 7.883766 | 8.068942 |
| 6.530525 | 6.332062 | 6.404424 | 6.883003 | 7.082139 | 7.550431 | 7.548995 | 7.553194 | 6.197933 |
| 7.629582 | 7.367814 | NaN | NaN | NaN | 7.398181 | NaN | 7.472534 | 8.318074 |
| 7.132527 | 7.176076 | 7.078931 | 7.33624 | 7.190472 | 6.729846 | 6.365241 | 6.863536 | 6.700045 |
| 6.900969 | 6.276803 | 6.507428 | 5.536819 | 6.399481 | 6.692427 | 6.831573 | 7.063658 | 5.688866 |
| 5.224945 | 4.658505 | NaN | 4.545796 | 4.388488 | 6.075651 | 6.966112 | 5.235812 | 3.827836 |
| 7.074092 | 7.12983 | 6.729678 | 6.91591 | 7.204233 | 6.953948 | 6.669749 | 6.698892 | 7.157001 |
| 7.264868 | 7.532457 | 7.231202 | 8.392314 | 8.30387 | 7.179456 | 7.137019 | 6.899749 | 8.174593 |
| 6.739003 | 6.627164 | 6.898443 | 5.445684 | 5.619731 | 6.829665 | 6.910344 | 7.235441 | 6.197182 |
| 6.367381 | NaN | NaN | 4.129254 | 4.586444 | NaN | NaN | 5.5895 | 6.803117 |

-Bruker Confidential-

| | | | | | | | | |
|---|---|---|---|---|---|---|---|---|
| NaN | 6.398232 | 5.889162 | 6.294536 | 6.434067 | NaN | 6.671144 | 6.663242 | 6.92644 |
| 6.899317 | 6.632808 | 6.919293 | 6.07456 | 6.521673 | 7.166641 | 6.955945 | 7.02934 | 6.456347 |
| 6.081128 | 6.037705 | 6.133437 | NaN | NaN | 5.510936 | 5.960111 | 5.96097 | 5.603526 |
| 7.888743 | 7.873299 | 8.192179 | 9.890274 | 9.901638 | 4.40169 | 6.083798 | 6.880611 | 6.990736 |
| 7.946339 | 8.143587 | 7.829408 | 7.906753 | 7.13094 | 5.796914 | 6.729667 | 5.95836 | 6.115821 |
| 7.740823 | 7.410398 | 7.610251 | 7.969883 | 7.844206 | 7.29577 | 7.216928 | 7.180063 | 7.070797 |
| 6.766029 | 7.185913 | 7.19303 | 7.047149 | 6.664289 | 7.025516 | 6.837115 | 6.870509 | 6.74852 |
| NaN | NaN | NaN | NaN | 4.639844 | NaN | 3.967237 | NaN | 5.422101 |
| 6.021325 | 6.312638 | 6.585627 | 5.728138 | 5.710906 | 5.448837 | 5.637306 | 5.373812 | 5.739749 |
| 6.672263 | 6.989339 | 6.988631 | 5.311002 | 5.618961 | 6.618535 | 6.82415 | 6.683735 | 6.102081 |
| 6.803658 | 7.431441 | 6.21143 | 7.332749 | 7.711451 | 5.652604 | 5.860936 | 4.654346 | 6.261018 |
| 8.720639 | 8.89128 | 8.959521 | 8.374013 | 8.231651 | 8.956339 | 8.898191 | 8.857863 | 8.984113 |
| 6.992827 | 7.850316 | 6.811232 | 6.709471 | 6.299112 | 7.736898 | 7.641806 | 7.871449 | 6.758219 |
| 6.912049 | 6.379075 | 6.463515 | 5.407036 | 5.086409 | 7.321632 | 7.041604 | 7.261495 | 6.751503 |
| 6.398412 | 6.884958 | 7.003398 | NaN | NaN | 6.236096 | 6.227027 | 6.262733 | 6.586004 |
| 7.401137 | 6.885625 | 7.233511 | 8.124882 | 8.16766 | 6.854708 | 6.416837 | 6.805595 | 7.350996 |
| 7.133962 | 7.0938 | 7.033448 | 7.206623 | 7.322976 | 6.94567 | 6.928621 | 7.027908 | 6.96559 |
| 7.250818 | 6.82167 | 7.010891 | 6.588698 | 6.755466 | 7.040732 | NaN | NaN | 6.207781 |
| 7.250038 | 6.982971 | 7.266368 | 7.216694 | 7.418718 | 7.340871 | 7.31419 | 7.236753 | 7.214305 |
| 8.206006 | 8.468125 | 8.065349 | 7.938731 | 8.010199 | 8.295838 | 8.39888 | 8.017261 | 8.137595 |
| 7.582697 | 7.754472 | 7.771266 | 7.160862 | 7.465816 | 7.923813 | 7.720985 | 7.992234 | 7.147718 |
| 8.053124 | 8.212214 | 8.173706 | 7.900524 | 7.78745 | 7.646573 | 7.580189 | 7.607371 | 7.656067 |
| 6.039494 | 5.861739 | 6.073195 | 7.846048 | 7.988106 | 6.819036 | 6.772526 | 7.08822 | 7.381555 |
| NaN | 5.161301 | 5.919263 | NaN | NaN | NaN | NaN | 2.86846 | 5.679234 |
| 6.86739 | 6.295517 | 7.317667 | 6.208582 | 5.392285 | 6.313289 | 6.212606 | NaN | 6.496274 |
| 7.954407 | 8.041076 | 7.82597 | 6.270845 | 6.214134 | 7.818444 | 7.778488 | 7.722633 | 7.841738 |
| 5.871012 | 5.661198 | 6.171896 | 5.775029 | 5.218185 | 6.455175 | 6.401046 | 6.616223 | 5.717286 |
| 7.41858 | 5.4203 | 7.540949 | 7.521455 | 6.335945 | 7.065775 | 7.346443 | 7.184776 | 6.788274 |
| 8.008303 | 8.006394 | 8.026121 | 7.954978 | 7.817131 | 7.963863 | 7.72635 | 7.507251 | 7.970532 |
| 6.818779 | 6.471675 | 6.897943 | 5.942962 | 6.794373 | 7.577306 | 7.62566 | 7.679998 | 7.07162 |
| 5.85028 | 5.934152 | 5.795582 | NaN | 5.663406 | NaN | NaN | NaN | 6.353786 |
| 8.758493 | 8.779496 | 8.830681 | 8.154391 | 8.229198 | 8.40445 | 8.237618 | 8.332221 | 8.728179 |
| 7.109648 | 6.594361 | 6.890938 | 6.27379 | 5.937605 | 6.428249 | 6.378778 | 6.325293 | 6.747865 |
| 8.097947 | 8.094131 | 8.248716 | 9.09156 | 8.767331 | 8.037442 | 7.862144 | 7.927058 | 7.134986 |
| 7.454418 | 7.255595 | 7.282588 | 8.039407 | 7.744184 | 8.692264 | 6.918143 | 8.3528 | 6.692044 |
| 4.259051 | 5.607514 | 5.518209 | 4.769264 | 5.797992 | 6.267563 | 5.906365 | 5.738128 | 5.869533 |
| 5.988405 | 6.715874 | 7.349464 | NaN | NaN | 6.477567 | 6.768637 | 6.540648 | 6.784611 |
| 7.855455 | 7.720058 | 7.436787 | 6.982949 | 6.832722 | 7.233103 | 7.010071 | 6.662347 | 6.898514 |
| 8.606246 | 8.602344 | 8.616589 | 7.823222 | 7.620248 | 8.741052 | 8.63449 | 8.716147 | 8.349265 |
| 6.716815 | 6.792833 | 6.591323 | 6.543702 | 6.509621 | 7.342221 | 7.373702 | 7.309682 | 6.629208 |
| 7.080588 | 7.065833 | 7.302155 | 7.308106 | 7.111238 | 6.278822 | 5.976777 | 6.327992 | 6.450532 |
| 8.376225 | 8.482497 | 8.376569 | 7.132812 | 7.400621 | 8.258532 | 8.040718 | 8.332887 | 7.860212 |
| 8.67519 | 8.62111 | 8.42456 | 7.51094 | 7.464692 | 8.244419 | 8.261933 | 8.025711 | 7.237319 |
| 8.169965 | 8.09168 | 8.163888 | 6.954732 | 6.829775 | 7.525155 | 7.657749 | 7.460326 | 6.492402 |
| NaN | NaN | NaN | NaN | NaN | NaN | NaN | NaN | 6.258217 |
| NaN | NaN | NaN | 5.787257 | 5.811643 | 4.750723 | 4.815306 | NaN | 6.02846 |
| 7.251131 | 7.366296 | 7.562038 | NaN | NaN | 6.30315 | 7.069174 | 6.450131 | 6.278861 |

| | | | | | | | | |
|---|---|---|---|---|---|---|---|---|
| 8.374065 | NaN | NaN | 7.68399 | 7.537651 | NaN | 6.764393 | 6.669928 | 6.647042 |
| NaN | 4.402917 | 4.505626 | 4.885231 | 6.939759 | 5.294322 | 5.347862 | 6.005432 | 5.24214 |
| 6.184349 | 5.007593 | 6.054522 | 5.093858 | 5.096296 | 6.216791 | 5.909436 | 5.924354 | 5.862665 |
| 6.824753 | 6.434139 | 7.008233 | 5.75667 | 5.072411 | 7.01827 | 7.296299 | 7.305165 | 6.83487 |
| 8.499681 | 8.441826 | 8.555159 | 9.883471 | 9.852819 | 8.434798 | 8.351609 | 8.384577 | 7.819513 |
| NaN | 6.369023 | 7.362325 | 5.881771 | 6.212581 | NaN | 5.92227 | 5.999652 | 5.99585 |
| 7.1979 | 6.716387 | 4.84085 | 6.850537 | 7.341964 | NaN | 6.013458 | NaN | 6.477212 |
| 6.217295 | 5.319767 | 6.150851 | 6.455639 | 6.910349 | 6.600267 | 6.263458 | 6.297152 | 4.746951 |
| 7.006287 | 7.283812 | 7.242988 | 7.095859 | 7.474741 | 6.07339 | 6.309861 | 6.304324 | 8.793518 |
| 8.014692 | 7.772917 | 7.825078 | 7.042487 | 6.56777 | 7.858777 | 7.940151 | 8.114189 | 7.723744 |
| 8.048634 | 8.29617 | 8.062643 | 6.100993 | 6.689602 | 7.32748 | 7.30078 | 7.689937 | 7.678327 |
| 7.319944 | 7.715805 | 7.894558 | 7.107126 | 6.78801 | 8.665507 | 9.36659 | 9.114315 | 7.478578 |
| 2.917068 | NaN | NaN | 4.811254 | 5.222169 | NaN | NaN | NaN | NaN |
| 5.476329 | 5.79602 | 5.560624 | 5.847637 | 6.011264 | 5.621219 | 5.923226 | 5.74945 | 5.64152 |
| 6.676011 | 7.02663 | 7.00793 | 8.132048 | 8.029178 | 3.958029 | 4.35782 | 5.164201 | 5.563845 |
| 6.1679 | 6.34826 | 6.351897 | 7.516515 | 8.057327 | 6.808564 | 6.407609 | 6.282649 | 5.829875 |
| 7.8992 | 7.726645 | 8.299382 | 6.375657 | 6.374148 | 7.152003 | 7.369287 | 7.595019 | 7.314836 |
| 5.924013 | 5.037332 | 6.147323 | 6.012304 | 5.456181 | 7.056988 | 6.56779 | 7.166388 | 6.02267 |
| 9.107984 | 9.061505 | 8.954212 | 9.286825 | 9.252986 | 8.7575 | 8.636672 | 8.732216 | 8.840618 |
| 9.073034 | 9.131623 | 8.966287 | 10.87343 | 11.88449 | 10.29577 | 9.994232 | 10.1073 | 11.5727 |
| 7.219573 | 7.19762 | 7.24335 | 6.391207 | 6.241214 | 7.566004 | 7.579255 | 7.556835 | 7.269777 |
| NaN | NaN | 8.203911 | NaN | 6.413939 | NaN | 5.64973 | NaN | 6.649191 |
| NaN | 5.775692 | NaN | NaN | NaN | 6.573615 | NaN | 6.439065 | 6.944701 |
| 5.254606 | 5.38868 | 5.952322 | NaN | 4.333023 | 7.105938 | 6.820628 | 6.642375 | 5.707705 |
| 9.880515 | 9.637115 | 9.880583 | 8.251521 | 8.392677 | 9.690218 | 9.945914 | 10.04279 | 9.186986 |
| 7.86484 | 7.951878 | 8.13231 | 7.399301 | 7.623553 | 7.977023 | 7.722617 | 7.658583 | 6.327658 |
| 6.270519 | 6.267901 | 6.520961 | 6.290075 | 4.377454 | 5.916001 | 6.218688 | 6.633181 | 5.306863 |
| 7.455509 | 7.677483 | 7.591019 | 6.38653 | 6.051391 | 6.546805 | 6.106699 | 6.354173 | 6.851943 |
| 8.04406 | 7.808539 | 7.858266 | 7.466596 | 7.466375 | 8.181555 | 8.216846 | 8.378886 | 7.394781 |
| 6.776869 | 6.391998 | 6.032965 | 6.711195 | 7.145277 | 5.950774 | 6.286694 | 6.235144 | 7.529359 |
| 6.150626 | 6.542574 | 6.794388 | 4.938599 | 4.611393 | 6.445823 | 6.146399 | 6.609136 | 4.774888 |
| 6.943406 | 7.323821 | 7.111147 | 6.311918 | 6.38573 | 7.627679 | 7.627727 | 7.606093 | 7.477039 |
| 8.798113 | 8.746824 | 8.713437 | 9.337617 | 9.276034 | 8.858016 | 8.909472 | 8.807091 | 7.684504 |
| 7.158845 | 7.412389 | 7.144984 | 7.830304 | 7.491055 | 7.131127 | 7.349328 | 7.462376 | 7.465014 |
| NaN | NaN | 5.087861 | 5.272185 | 6.71064 | NaN | 4.856078 | NaN | 4.999683 |
| NaN | 7.541904 | 7.675326 | 6.515498 | 6.339036 | 6.657764 | 6.528037 | 6.888604 | 4.899563 |
| 6.176507 | 5.650732 | 5.622088 | NaN | NaN | 5.652022 | NaN | NaN | 4.457078 |
| 8.385558 | 7.103667 | 8.219255 | 6.564866 | 6.639034 | 8.643249 | 8.315738 | 8.200084 | 7.966912 |
| 6.784819 | 6.739166 | 7.092645 | 6.211018 | 6.184888 | 7.175047 | 7.299173 | 7.414554 | 6.690175 |
| 7.915757 | 7.988388 | 7.986772 | 6.914008 | 7.32656 | 7.988812 | 7.779561 | 8.038261 | 8.198091 |
| 9.456138 | 9.449118 | 9.411283 | 7.628082 | 7.645448 | 8.988869 | 8.974277 | 9.320734 | 8.33721 |
| 7.399039 | 7.349642 | 7.326809 | 7.756423 | 8.04767 | 8.159142 | 7.868445 | 8.126556 | 6.665784 |
| 8.273865 | 7.679822 | 7.981575 | 8.958181 | 8.727409 | 7.268541 | 7.150271 | 7.030574 | 6.268793 |
| 6.894533 | 6.554385 | 6.527013 | 4.905598 | 5.785534 | 6.305408 | 6.093234 | 5.892466 | 4.820772 |
| 7.580704 | 7.858829 | 8.004891 | 8.74582 | 8.870396 | 8.796986 | 8.579636 | 8.872414 | 8.108624 |
| 6.91231 | 6.94499 | 6.445609 | 6.832122 | 6.964223 | 7.065081 | 7.102876 | 7.172107 | 7.314279 |
| 6.343506 | 5.600091 | 5.622847 | 4.395525 | NaN | 6.080902 | 6.076033 | 6.882873 | 6.244009 |

| | | | | | | | | |
|---|---|---|---|---|---|---|---|---|
| NaN | NaN | NaN | NaN | NaN | 6.442269 | 6.771908 | 6.805805 | 3.056408 |
| 9.052544 | 9.209818 | 9.17575 | 7.086605 | 7.454713 | 8.960332 | 9.232139 | 8.950642 | 8.728843 |
| 7.487154 | 7.251704 | 7.547587 | 6.903075 | 6.867818 | 7.32488 | 7.402554 | 7.529981 | 7.640703 |
| 6.727803 | 5.635566 | 6.37387 | 5.534554 | 5.801516 | 6.146258 | 6.336746 | 6.31659 | 3.1967 |
| 7.751549 | 7.867091 | 7.727373 | 6.097098 | 6.081393 | 7.645168 | 7.67891 | 7.804651 | 7.903935 |
| 5.292052 | 3.746866 | NaN | 5.050067 | 4.605268 | 6.192741 | 4.609897 | 4.608979 | 6.128564 |
| 6.256779 | 6.172372 | 6.011997 | 4.876095 | 5.061398 | 5.484381 | 5.943372 | 5.777411 | 6.42641 |
| 4.318627 | 3.735403 | 4.907818 | NaN | 4.105975 | 4.596831 | 4.413865 | 4.344284 | 4.912442 |
| 7.22687 | 7.271231 | 7.388093 | 8.355567 | 8.375065 | 7.143228 | 7.17381 | 7.073171 | 6.447345 |
| 7.735392 | 7.740427 | 7.719904 | 7.981466 | 7.623072 | 6.697344 | 6.799299 | 6.991878 | 7.27508 |
| 7.055107 | 7.08026 | 6.91802 | 6.591755 | 6.347561 | 6.347973 | 6.904586 | 7.170805 | 8.145893 |
| 7.121938 | 5.3056 | 6.666919 | 5.777417 | 6.938046 | 8.028474 | 7.639603 | 8.411893 | 8.13794 |
| 6.9287 | 6.334351 | 5.71729 | 5.717694 | 6.057448 | 6.985188 | 6.607907 | 6.525165 | 5.906605 |
| NaN | NaN | 4.645154 | NaN | NaN | 6.299006 | NaN | 5.830209 | 6.293462 |
| NaN | NaN | NaN | 5.460195 | 6.099533 | NaN | NaN | 5.47102 | 4.436403 |
| 8.173656 | 8.238383 | 8.265429 | 9.153624 | 9.270537 | 8.791687 | 8.530316 | 8.625549 | 7.79483 |
| 6.879228 | 6.853542 | 7.062921 | 6.875357 | 6.672016 | 7.413562 | 7.039015 | 7.264496 | 7.017172 |
| NaN | 6.904667 | 6.065996 | 6.42637 | 6.665692 | 6.9463 | 6.755358 | 7.071058 | 6.896105 |
| 10.74426 | 12.02469 | 12.14554 | 8.60579 | 10.46934 | 9.308147 | 8.668959 | 8.907355 | 9.757154 |
| 6.446518 | 6.421366 | 6.328684 | 6.25415 | 6.332353 | 6.277459 | 6.316741 | 6.327168 | 5.655538 |
| NaN | NaN | NaN | 5.16907 | NaN | NaN | NaN | NaN | 7.266178 |
| 5.861992 | 5.814128 | 6.3656 | 6.563836 | 6.523811 | 5.747799 | 5.834669 | 6.25132 | 6.265341 |
| 7.337371 | 7.240579 | 7.574582 | 6.312889 | 7.006944 | 7.473667 | 7.25484 | 7.315765 | 7.286846 |
| 5.242322 | 5.959231 | 5.276764 | 7.493228 | 7.425848 | 6.637888 | 6.421939 | 6.548372 | 5.366089 |
| 4.947726 | NaN | 5.264771 | 6.491772 | 6.432537 | 6.880001 | NaN | NaN | 5.396398 |
| 7.220119 | 7.430605 | 7.41224 | 6.737265 | 6.559709 | 6.756559 | 6.985592 | 6.856273 | 6.414921 |
| 7.003088 | 7.87359 | 7.288407 | 5.464981 | 5.269679 | 6.301628 | 6.628693 | 6.780853 | 6.535213 |
| 6.442315 | 6.031611 | 5.665102 | 5.315928 | 5.663922 | 6.873612 | 6.633842 | 6.624136 | 5.80868 |
| 6.885989 | 6.51701 | 6.224309 | NaN | 5.033173 | 6.340562 | 6.551909 | 6.639575 | 6.844414 |
| 6.102198 | 6.527387 | 6.850093 | 6.71123 | 6.840973 | 7.245236 | 7.287836 | 7.170499 | 7.371981 |
| 4.981429 | NaN | NaN | NaN | 3.588807 | 6.131252 | 5.904564 | 4.920358 | 4.795929 |
| NaN | NaN | NaN | NaN | NaN | NaN | NaN | 4.9818 | 6.012503 |
| 7.200316 | 7.52986 | 6.994882 | NaN | NaN | 7.992036 | 8.107413 | 7.518111 | 7.007871 |
| 5.571944 | 4.848826 | 5.541991 | 5.766599 | NaN | 6.545268 | 6.208653 | 5.91494 | 5.949175 |
| 8.593768 | 8.753277 | 8.263505 | 9.911804 | 9.822894 | 8.844534 | 8.714947 | 8.780079 | 8.094966 |
| 8.004618 | 7.76599 | 7.837489 | 8.330175 | 7.427569 | 8.36734 | 8.020298 | 7.830054 | 7.848598 |
| 8.617022 | 7.707169 | 9.820539 | 8.483239 | 8.783948 | 7.710833 | 7.756067 | 7.78735 | 7.899859 |
| 6.712752 | 5.523434 | 7.0624 | 6.629064 | 6.622329 | 7.279185 | 7.252234 | 6.978141 | 7.51265 |
| 8.568235 | 8.31178 | 8.480529 | 6.804236 | 6.58917 | 7.77697 | 7.81285 | 7.807773 | 7.864777 |
| NaN | NaN | NaN | 6.98933 | 7.718092 | NaN | NaN | NaN | NaN |
| 6.565199 | 6.434692 | 6.558175 | 5.487232 | 4.747698 | 6.630857 | 6.381745 | 6.377058 | 6.123982 |
| 6.105264 | 6.127004 | 6.390545 | 5.874949 | 6.19542 | 6.423921 | 7.347055 | 6.498675 | 6.353297 |
| 6.6191 | 6.939523 | 6.654676 | 6.843131 | 6.824162 | 6.915878 | 6.294704 | 6.677935 | 6.519181 |
| NaN | NaN | NaN | 7.717831 | 5.513405 | NaN | 6.908004 | NaN | 6.532662 |
| 6.811372 | 6.779117 | 6.66239 | 5.188054 | 5.994413 | 6.764979 | 6.593733 | 6.832374 | 7.530128 |
| 7.622164 | 7.628879 | 7.064128 | 7.573275 | 7.689386 | 6.284636 | 5.428789 | 6.486632 | 6.787049 |
| 7.416034 | 7.005468 | 7.687567 | 7.610865 | 7.554443 | 6.750096 | 7.599061 | 7.295657 | 6.998212 |

| | | | | | | | | |
|---|---|---|---|---|---|---|---|---|
| 6.769173 | 6.298791 | 6.968986 | 5.374821 | 5.168759 | 7.155671 | 7.230033 | 6.770076 | 5.723896 |
| 8.265147 | 8.320081 | 8.058504 | 8.714661 | 8.86977 | 7.904693 | 7.758248 | 7.955337 | 7.974423 |
| 7.575656 | 7.26916 | 7.812862 | 7.277914 | 7.081201 | 7.664277 | 7.02677 | 7.397854 | 5.927427 |
| 6.958574 | 6.745613 | 5.795833 | NaN | 5.931582 | 6.693166 | 6.65331 | 7.100758 | 5.99885 |
| NaN | 5.997741 | 5.416777 | 9.322489 | 3.112258 | 5.33092 | 5.905104 | NaN | 5.17537 |
| 7.341471 | 4.924492 | 7.959022 | 8.174148 | 8.013155 | NaN | NaN | NaN | 6.130099 |
| 11.64715 | 9.317064 | 11.6027 | 11.22859 | 11.12755 | 12.59432 | 12.66081 | NaN | 11.82719 |
| 7.550023 | 6.480312 | 6.797947 | 6.732369 | 6.591572 | 7.125979 | 6.751593 | 7.176615 | 6.750817 |
| 7.244057 | 6.157457 | 7.325199 | 6.273764 | 5.771712 | 6.679031 | 6.783227 | 6.863764 | 7.10852 |
| 7.177282 | 6.748109 | 6.617688 | 6.022344 | 6.071029 | 7.297706 | 7.366447 | 7.888542 | 6.966283 |
| 6.013823 | 5.963344 | 6.713805 | 6.949932 | 6.881679 | 6.340682 | 6.090061 | 6.169656 | 6.344053 |
| 7.158046 | 7.794947 | 7.378172 | 6.714972 | 6.97762 | 7.785339 | 7.992106 | 8.004403 | 7.503617 |
| 8.698704 | 8.465694 | 8.349219 | 8.388658 | 8.66145 | 7.666997 | 7.982486 | 8.530116 | 8.359822 |
| 6.711497 | 6.619351 | 6.815038 | 7.41845 | 7.557104 | 6.995633 | 6.788671 | 7.064869 | 7.406929 |
| 8.244286 | 7.945889 | 8.085878 | 7.720072 | 7.411988 | 7.683774 | 7.271596 | 7.596231 | 7.979823 |
| 6.498403 | 6.618197 | 6.832447 | 6.99997 | 7.344615 | 6.990565 | 6.772279 | 7.13914 | 6.774412 |
| 6.171669 | 6.925673 | 5.985311 | 5.829127 | 5.492208 | 6.502646 | 6.011756 | 6.396438 | 7.032807 |
| 6.498571 | 7.499073 | 7.191558 | 7.330163 | 7.307218 | 5.948769 | 6.126553 | 6.574091 | 6.095046 |
| 7.442337 | 7.681228 | 7.414239 | 6.802797 | 6.548292 | 8.231645 | 7.651554 | 7.856302 | 7.459984 |
| NaN | NaN | NaN | 7.169208 | 7.4086 | NaN | NaN | NaN | NaN |
| 7.811246 | 6.762875 | 7.196825 | 7.899427 | 7.891687 | 7.401342 | 7.589214 | 7.9189 | 7.238996 |
| 7.422057 | 5.798375 | 6.860775 | NaN | NaN | 7.647635 | 7.839829 | 7.63314 | 6.311449 |
| 7.213833 | 6.915142 | 6.968206 | NaN | 5.851772 | 6.632322 | 6.232173 | 6.003448 | 5.41349 |
| 8.322102 | 8.427793 | 8.443205 | 9.58705 | 9.859275 | 8.161967 | 7.225811 | 7.43381 | 8.44203 |
| 5.935275 | 6.297503 | 5.612204 | 7.58012 | 7.823733 | 6.920604 | 6.908211 | 7.049631 | 6.451785 |
| 7.79011 | 7.567516 | 7.644864 | 6.809618 | 6.621744 | 7.46518 | 7.641037 | 7.811723 | 7.887159 |
| 8.517712 | 8.411043 | 8.423718 | 7.591481 | 7.368941 | 8.486166 | 8.371511 | 8.506694 | 8.464516 |
| 6.630368 | 6.21999 | 6.877971 | 6.80626 | 6.533639 | 6.379594 | 6.611932 | 6.339501 | 6.173829 |
| 6.925748 | 5.632865 | 5.909055 | 6.779706 | 6.671084 | 7.340187 | 6.883595 | 7.399285 | 6.662354 |
| 7.182357 | 7.009264 | 6.806402 | 6.775122 | 6.52794 | NaN | 7.083472 | 7.03078 | 7.501573 |
| 8.534862 | 8.460983 | 8.59321 | 9.250829 | 9.480932 | 8.594114 | 8.469892 | 8.690848 | 7.951676 |
| 6.763703 | 6.815315 | 6.671203 | 6.92276 | 7.093941 | 5.943652 | 6.280634 | 6.65556 | 6.131162 |
| 5.612003 | 5.666821 | NaN | NaN | NaN | 6.012716 | 6.381574 | 6.174499 | 5.464808 |
| 7.519074 | 8.478902 | 6.211589 | 6.478564 | 9.266212 | 7.158378 | NaN | 5.986973 | 7.320276 |
| 7.529892 | 6.789815 | 7.56814 | 9.936467 | 9.453248 | 7.528743 | 8.163333 | 6.735031 | 7.628517 |
| 7.282662 | NaN | NaN | NaN | NaN | NaN | NaN | 5.853626 | 7.522024 |
| NaN | NaN | NaN | NaN | NaN | NaN | NaN | NaN | 6.183743 |
| 8.094398 | 8.332767 | 8.315066 | 9.267936 | 9.048081 | 8.19578 | 8.239029 | 8.148414 | 8.344909 |
| 8.399165 | 8.306049 | 8.204793 | 7.788637 | 7.546177 | 8.303636 | 8.227287 | 8.225307 | 8.451421 |
| 5.579601 | 5.247368 | 5.496916 | 6.132496 | 6.179729 | 5.966144 | 5.330184 | 5.693519 | 6.082212 |
| 6.011696 | 6.513454 | 6.461443 | 6.638845 | 7.041389 | 5.966159 | 6.853892 | 6.808718 | 6.430686 |
| 9.119407 | 8.544323 | 8.758752 | 9.57248 | 9.498335 | 8.248523 | 8.576878 | 8.387431 | 8.759154 |
| 7.393046 | 8.136703 | 8.360921 | 8.535504 | 8.396008 | 7.661282 | 7.867094 | 8.314851 | 8.15405 |
| 7.696975 | 7.192621 | 7.49855 | 6.378201 | 5.940273 | NaN | NaN | NaN | 6.089926 |
| 6.200585 | 6.70193 | 6.532131 | 3.823201 | 4.828794 | 5.930417 | 5.53968 | 5.59742 | 5.675376 |
| 7.179303 | 7.115269 | 7.203451 | 6.618458 | 6.645382 | 6.590838 | 6.572048 | 6.787021 | 6.712283 |
| 7.27176 | 7.422195 | 7.414851 | 6.991373 | 7.106775 | 7.930357 | 7.788991 | 7.878591 | 7.764242 |

| | | | | | | | | |
|---|---|---|---|---|---|---|---|---|
| 6.464637 | 6.543644 | 6.682431 | 6.493392 | 5.957097 | 6.169599 | 6.362041 | 6.906825 | 6.540096 |
| 6.493038 | 5.432762 | 5.671972 | NaN | 5.805769 | 6.305526 | 6.440024 | 6.547037 | 6.530373 |
| 8.021391 | 7.176232 | 7.315272 | NaN | 4.905181 | NaN | NaN | 7.446306 | NaN |
| 7.816819 | 7.721707 | 7.531135 | 6.280583 | 6.459093 | 7.684573 | 7.665131 | 7.667905 | 7.492813 |
| 8.257676 | 8.088586 | 8.304602 | 8.468678 | 8.143266 | 7.506695 | 8.125813 | 7.811816 | 8.082114 |
| 9.126354 | 8.97089 | 9.117283 | 10.08744 | 9.98728 | 7.050772 | 7.398278 | 7.479767 | 8.372598 |
| 7.64707 | 7.701371 | 7.8627 | 6.333443 | 6.624906 | 8.735095 | 8.429317 | 8.554755 | 7.666571 |
| 3.632611 | 4.081007 | 2.852068 | NaN | 2.894564 | 3.11813 | 3.105453 | NaN | 4.297109 |
| 9.119563 | 9.177841 | 8.922452 | 9.012957 | 9.094241 | 8.280343 | 8.413852 | 8.218099 | 7.891405 |
| 7.66727 | 7.165859 | 7.735226 | 7.12464 | 6.919676 | 6.859735 | 7.040049 | 6.851416 | 7.081797 |
| 7.253222 | 6.638364 | 7.244392 | 7.690612 | 7.640163 | NaN | 7.336983 | 6.865901 | 7.149057 |
| 6.413575 | 4.70023 | 6.951139 | NaN | NaN | 7.434478 | 7.700027 | 7.305944 | 6.042021 |
| 6.847784 | 6.541894 | 6.766502 | 5.575311 | 5.387906 | 6.046978 | 6.375375 | 6.587268 | 7.184155 |
| 9.212538 | 9.246977 | 9.241443 | 9.474963 | 9.230936 | 9.186555 | 9.003276 | 9.184508 | 8.800254 |
| 6.215647 | 7.69537 | 7.671041 | 7.109056 | 6.227259 | 6.693258 | 6.694906 | 7.233191 | NaN |
| 7.125874 | 7.336951 | 7.16033 | 6.028825 | 5.987214 | 6.990937 | 7.047758 | 7.017359 | 6.359044 |
| NaN | NaN | NaN | NaN | NaN | NaN | 7.580163 | 8.232064 | 5.987357 |
| 7.107507 | 6.336915 | 6.227369 | 7.525092 | 7.72025 | 6.673312 | 7.066047 | NaN | 7.564744 |
| 8.579027 | 8.661336 | 8.732544 | 7.785956 | 7.897643 | 8.718182 | 8.547142 | 8.455832 | 8.412106 |
| 4.259589 | 3.786087 | NaN | NaN | 6.257954 | 6.118486 | 6.076756 | 4.945275 | 7.724486 |
| 8.394927 | 8.45343 | 8.473795 | 6.424447 | 7.041839 | 8.083318 | 8.014201 | 7.854175 | 7.740795 |
| 7.281446 | 7.508852 | 7.406144 | 7.094226 | 6.641428 | 7.528156 | 6.775456 | 7.325459 | 7.447217 |
| 10.48489 | 10.42899 | 10.59431 | 10.24609 | 10.25244 | 8.431766 | 7.868071 | 7.993774 | 10.45515 |
| 6.795803 | 6.702577 | 7.229032 | 6.243757 | 6.099593 | 6.857703 | 6.628375 | 6.577477 | 6.263224 |
| 6.779655 | 6.878767 | 6.182898 | 8.172657 | 7.73044 | 6.793808 | 7.137968 | 6.844831 | 7.06592 |
| 5.829342 | NaN | NaN | NaN | NaN | 6.609077 | 6.535317 | 6.466723 | 6.65556 |
| 5.099132 | 4.687902 | 4.987008 | 5.277102 | 4.17582 | 0.65026 | 3.848341 | NaN | 6.218405 |
| 9.369856 | 9.230883 | 9.19815 | 9.393471 | 9.300361 | 8.413608 | 8.326019 | 8.411917 | 8.47154 |
| NaN | NaN | NaN | 7.805559 | 7.84312 | 7.374062 | NaN | NaN | NaN |
| 6.442592 | 7.006682 | 6.407565 | 6.910599 | 7.087979 | 6.713156 | 6.817306 | 6.750894 | 5.978866 |
| 4.81662 | 4.382312 | NaN | 4.238999 | 4.178766 | 5.604369 | 6.216353 | 6.422089 | 4.307186 |
| 6.981081 | 7.300341 | 6.926551 | 6.695813 | 6.192308 | 6.826663 | 6.969296 | 7.055287 | 6.20179 |
| 10.53479 | 10.41449 | 10.45786 | 11.01218 | 10.8154 | 10.05542 | 10.22769 | 10.15035 | 10.37561 |
| 8.269978 | 8.192445 | 8.328711 | 7.161821 | 7.009542 | 8.173175 | 8.207973 | 8.567725 | 7.35982 |
| 5.782322 | 5.958752 | 5.816668 | 5.608604 | 5.799818 | 5.558027 | 5.851816 | 6.030088 | 5.476694 |
| 8.122191 | 8.070618 | 8.084966 | 7.233085 | 7.286665 | 7.739573 | 7.747856 | 7.728538 | 7.753787 |
| NaN | 4.94255 | 4.572445 | 3.779741 | 3.364339 | 5.973975 | 5.11215 | 5.829811 | NaN |
| 7.428708 | 7.666924 | 7.580708 | 7.599722 | 7.317519 | 7.611581 | 7.280401 | 7.405736 | 7.461522 |
| 6.657753 | 6.865696 | 6.527322 | 6.638346 | 6.407807 | 6.079111 | 6.429144 | 6.059681 | 6.835637 |
| 6.880171 | 7.093524 | 6.674184 | NaN | NaN | NaN | NaN | NaN | NaN |
| 7.911571 | 7.867338 | 7.999303 | 4.40546 | 4.159062 | 7.420061 | 8.10975 | 8.06191 | 6.816963 |
| 9.245752 | 9.094295 | 9.136138 | 7.585634 | 7.622764 | 9.122027 | 9.04174 | 9.148641 | 8.103984 |
| 6.644065 | 6.623124 | 6.35678 | 6.847024 | 7.121162 | 6.710229 | 6.742311 | 6.381506 | 6.446879 |
| 8.331953 | 7.596934 | 7.957947 | 7.328194 | 7.570179 | 8.115453 | 8.247133 | 7.949733 | 7.778432 |
| NaN | 6.195042 | 5.901056 | 6.21022 | 7.206956 | NaN | 6.247412 | 6.428244 | 6.355973 |
| 5.412815 | 5.72682 | 5.730388 | 5.851082 | 5.67667 | 5.714494 | 5.426416 | 5.32795 | 5.817233 |
| 7.242513 | 7.391925 | 7.126798 | 7.090876 | 7.197108 | 7.495108 | 7.3729 | 7.756259 | 7.070495 |

-Bruker Confidential-

| | | | | | | | | |
|---|---|---|---|---|---|---|---|---|
| 7.419662 | 6.355433 | 5.611053 | NaN | 6.351446 | 6.369453 | NaN | 6.980933 | 6.453114 |
| NaN | NaN | NaN | 5.65473 | 6.496053 | NaN | NaN | 4.28753 | 5.197005 |
| 7.590194 | 7.839662 | 7.451285 | 6.790833 | 6.849598 | 7.037077 | 7.28256 | 7.117968 | 6.97322 |
| 7.447687 | 7.466581 | 7.146842 | 7.001031 | 7.398099 | 7.182617 | 7.264028 | 7.167437 | 6.218992 |
| 5.759887 | 5.679267 | 5.241201 | NaN | 5.237997 | 6.692072 | 6.730883 | 6.892579 | 6.735164 |
| 9.088971 | 9.293094 | 8.99859 | 9.701657 | 9.517985 | 7.491567 | 7.765795 | 7.448871 | 7.840324 |
| 9.336608 | 9.404967 | 9.308411 | 9.551105 | 9.273125 | 9.686352 | 9.624969 | 9.515097 | 8.926402 |
| 7.771995 | 7.530846 | 7.942205 | 7.14795 | 7.234285 | 7.448202 | 7.159313 | 7.464706 | 5.792895 |
| 6.477805 | 6.821601 | 6.626708 | 6.308863 | 5.89482 | 6.840044 | 6.716884 | 7.051301 | 5.806506 |
| 9.386999 | 9.235937 | 9.265786 | 8.696503 | 8.72837 | 9.211713 | 9.179447 | 9.324026 | 9.38018 |
| 5.808327 | 5.862581 | 5.328927 | NaN | 4.158156 | 6.416932 | 5.935298 | 6.004613 | 6.897875 |
| 8.558523 | 8.394326 | 7.943464 | 8.209561 | 8.348237 | 7.925041 | 7.88092 | 8.221311 | 7.705743 |
| 7.713296 | 7.761145 | 7.866867 | 6.804339 | 6.797108 | 7.476586 | 7.390198 | 7.377287 | 7.141588 |
| 11.77848 | 11.59737 | 11.56395 | 12.55517 | 12.2498 | 11.72602 | 11.58983 | 11.66318 | 10.61614 |
| 7.932063 | 7.26624 | 7.600454 | 8.938112 | 8.677981 | 7.16158 | 7.472075 | 7.323831 | 7.476895 |
| 5.241479 | 5.632153 | 4.771832 | NaN | NaN | 6.151657 | 6.460822 | 6.011003 | 4.529691 |
| 7.804014 | 7.472549 | 7.538272 | 7.171172 | 6.914088 | 7.793626 | 7.666604 | 7.934407 | 7.870984 |
| 7.298794 | 7.223073 | 7.183641 | 7.736166 | 7.094118 | 7.00828 | 7.22151 | 7.189502 | 6.636755 |
| 6.077127 | 6.385721 | 6.030881 | 6.067329 | 6.25267 | 6.354307 | 6.159395 | 5.927922 | 6.298578 |
| 6.850371 | 7.077003 | 6.470874 | 7.281883 | 7.309703 | 6.603607 | 6.54219 | 6.554915 | 6.550932 |
| 10.13699 | 10.17809 | 10.06829 | 10.50287 | 10.44284 | 9.554096 | 9.665913 | 9.632656 | 9.675931 |
| 7.74341 | 8.476163 | 7.306541 | 6.665552 | 5.795703 | 7.256711 | 7.012258 | 7.16451 | 7.290065 |
| NaN | 4.181206 | NaN | NaN | NaN | 5.979574 | 4.542221 | 5.387891 | 4.420041 |
| NaN | NaN | 6.308593 | NaN | NaN | NaN | 5.906263 | 5.706098 | 6.014121 |
| 4.89809 | 5.23296 | 5.838129 | 5.652384 | NaN | 6.885254 | 6.553618 | 6.190966 | 6.30083 |
| 8.48223 | 8.481287 | 8.509519 | 7.64311 | 7.688536 | 8.118039 | 8.32274 | 8.144221 | 8.162992 |
| 7.478355 | 7.412621 | 7.557213 | 7.395906 | 7.024387 | 7.74493 | 7.603631 | 7.572463 | 7.252573 |
| 6.950833 | 6.836506 | 6.761065 | 6.984888 | 6.835829 | 7.234729 | 7.150625 | 7.254851 | 6.361061 |
| 4.096479 | 4.385114 | 4.183676 | 3.786537 | 5.245917 | 5.617135 | 5.32303 | 5.220136 | 4.388182 |
| 7.495203 | 6.981826 | 7.188493 | 6.347324 | 6.165935 | 7.318295 | 7.196313 | 7.322916 | 6.769775 |
| 6.772319 | 6.941644 | 7.145237 | 5.811132 | 5.956066 | 5.298051 | 5.292808 | 5.210987 | 5.069796 |
| 9.466957 | 9.419475 | 9.776843 | 9.579283 | 9.168206 | 7.056962 | 7.153231 | 7.85348 | 7.472212 |
| 8.304889 | 8.638206 | 8.685778 | 6.774099 | 7.72633 | 8.918967 | 8.972416 | 8.815168 | 8.639304 |
| 6.851414 | 6.843256 | 6.636744 | 6.384999 | 6.39182 | 7.159017 | 7.131706 | 7.012514 | 6.292778 |
| 7.09138 | 7.265245 | 7.124886 | 8.696002 | 8.614805 | 7.303667 | 7.023168 | 7.404905 | 6.928129 |
| 6.045832 | 4.98241 | 5.654409 | 4.778371 | 4.725622 | 6.100873 | 5.753913 | 5.810651 | 6.048879 |
| 8.258474 | 7.389157 | 8.359255 | 6.80559 | 6.542098 | 8.733943 | 7.790181 | 8.25995 | 6.963834 |
| 7.458684 | 7.39566 | 7.487457 | 6.221009 | 6.347549 | 7.230338 | 7.229037 | 7.337705 | 6.585641 |
| 9.985163 | 9.97482 | 9.983161 | 9.574165 | 9.615506 | 9.602347 | 9.740299 | 9.724165 | 10.17753 |
| 7.190646 | 7.70604 | 7.885624 | 6.944431 | 6.528095 | 7.287492 | 7.541871 | 7.635897 | 6.944957 |
| 7.824029 | 7.898973 | 7.699013 | 7.131813 | 7.726556 | 7.821856 | 7.649731 | 8.013831 | 7.683699 |
| 7.153366 | 6.692659 | 6.881722 | NaN | NaN | 7.460652 | 7.680656 | 7.585299 | 6.410896 |
| 5.995883 | 6.120356 | 6.423413 | 6.816725 | 7.065363 | 6.234698 | 6.230862 | 5.869612 | 6.464522 |
| 8.564857 | 8.497314 | 8.474558 | 8.051813 | 8.037931 | 9.116341 | 9.037939 | 9.003202 | 8.739458 |
| 7.219926 | 7.914029 | 7.602569 | 7.352103 | 7.172431 | NaN | 6.380963 | 6.25735 | 6.553183 |
| 7.755642 | 7.696952 | 7.864668 | 6.678953 | 6.651247 | 7.420969 | 7.458687 | 7.739239 | 7.771993 |
| 7.660731 | 8.022228 | 7.461966 | 6.616958 | 6.57424 | 7.533732 | 7.345473 | 7.551916 | 7.747277 |

| | | | | | | | | |
|---|---|---|---|---|---|---|---|---|
| 11.57384 | 11.51149 | 11.47393 | 10.54313 | 10.42664 | 11.16775 | 11.17528 | 11.14144 | 11.33268 |
| 5.783301 | 5.743536 | 5.283802 | 7.247888 | 6.070914 | 5.833353 | 7.193984 | 7.088565 | 5.639062 |
| 7.642043 | 7.687032 | 7.882694 | 7.369374 | 7.51316 | 7.636951 | 7.567621 | 7.547581 | 7.615267 |
| 8.474477 | 7.858461 | 8.345247 | 6.151407 | 6.445488 | 7.603382 | 7.84961 | 7.776834 | 8.447877 |
| 8.249097 | 8.355021 | 8.226597 | 7.334063 | 6.860188 | 8.126296 | 8.756496 | 8.722836 | 8.252081 |
| 8.042223 | 8.081113 | 7.51081 | 9.05379 | 8.977404 | 7.786263 | 7.860825 | 7.971944 | 7.591419 |
| 6.796919 | 6.708421 | 6.460455 | 7.16769 | 7.050169 | 6.320185 | 6.119844 | 6.224731 | 5.964102 |
| 6.608281 | 6.915498 | 6.786767 | NaN | NaN | 7.301367 | 7.250728 | 6.863185 | 7.008612 |
| 8.23906 | 8.677464 | 8.543086 | 9.321206 | 9.112794 | 8.795222 | 9.185422 | 8.745981 | 9.838756 |
| 7.295747 | 7.566984 | 7.30628 | 6.31821 | 6.323948 | 6.474702 | 6.236555 | 6.325122 | 6.511035 |
| 7.013228 | 6.873894 | 6.872996 | 6.498812 | 6.358173 | 7.205803 | 7.280465 | 7.397858 | 7.040549 |
| 7.286924 | 6.945794 | 7.184587 | 7.276525 | 7.83157 | 6.001014 | 6.231482 | 6.137882 | 7.523256 |
| 10.40095 | 10.00555 | 9.953775 | 11.22166 | 11.01335 | 9.369587 | 9.370659 | 9.217962 | 9.35465 |
| NaN | NaN | NaN | NaN | 3.700114 | NaN | NaN | NaN | 3.973207 |
| 8.811351 | 8.683259 | 8.481363 | 9.380814 | 9.484421 | 9.199196 | 9.300968 | 9.292775 | 8.570232 |
| 8.488989 | 8.213007 | 7.916638 | 7.217621 | 7.288169 | 8.775846 | 8.642174 | 8.686945 | 8.245193 |
| 6.799135 | 6.552418 | 6.851224 | 5.621875 | 6.440906 | 6.9679 | 6.914145 | 7.029384 | 6.296726 |
| 7.903288 | 7.876905 | 7.811577 | 6.082028 | 6.104991 | 7.977423 | 7.805666 | 7.791305 | 7.497424 |
| 6.608072 | 6.574612 | 6.699953 | 5.668248 | 6.095869 | 6.80633 | 6.918807 | 6.890911 | 4.603518 |
| 7.986553 | 5.643474 | 8.101934 | 6.492315 | 7.256556 | 7.878154 | 7.698794 | 7.839211 | 5.993822 |
| 7.534475 | 7.569647 | 7.577325 | 8.62684 | 8.62509 | 7.450864 | 7.093764 | 6.984246 | 6.960472 |
| NaN | NaN | 5.820762 | 6.057617 | 6.466048 | NaN | NaN | 5.964716 | 6.024452 |
| 6.863412 | 6.767477 | 6.399697 | 6.987976 | 7.157753 | 7.314645 | 7.432009 | 7.264126 | 7.381363 |
| 6.864417 | 6.482967 | 6.501481 | 6.362316 | 6.541182 | 6.74999 | 6.688971 | 6.759223 | 6.099338 |
| 7.463548 | 7.488335 | 7.423404 | 6.817893 | 6.620136 | 7.443124 | 7.571198 | 7.642365 | 7.70598 |
| 6.507484 | 6.951869 | 6.608078 | 8.569434 | 8.260982 | 6.279216 | 6.59053 | 6.969953 | 6.826633 |
| 5.777322 | 6.662767 | 5.454877 | 5.512607 | 5.880036 | NaN | NaN | 6.174276 | 5.969275 |
| 10.67548 | 10.35321 | 10.40709 | 11.20357 | 11.0462 | 9.372237 | 9.514429 | 9.326715 | 8.969648 |
| 8.444077 | 8.51814 | 8.444849 | 7.593628 | 7.875316 | 8.534078 | 8.583029 | 8.460743 | 7.597423 |
| 6.592463 | 7.570014 | 7.09615 | 4.421806 | 4.286489 | 6.96672 | 6.804776 | 6.983178 | 5.109887 |
| NaN | 5.823722 | NaN | 6.378145 | 6.195904 | NaN | NaN | NaN | NaN |
| 11.07807 | 10.83232 | 10.89257 | 9.806242 | 9.709047 | 10.70568 | 10.96968 | 10.88571 | 10.70502 |
| 8.598473 | 7.232478 | 8.609491 | 7.26975 | 7.537733 | 8.828837 | 8.867963 | 8.84613 | 8.093307 |
| 7.544821 | 7.656066 | 7.683799 | 7.333994 | 7.220499 | 7.546921 | 7.614221 | 7.776499 | 7.366977 |
| 7.402638 | 7.400232 | 7.298351 | 6.085217 | 5.713391 | 7.461241 | 7.223371 | 7.283145 | 6.98234 |
| 8.952027 | 8.818429 | 8.984646 | 8.105757 | 7.896335 | 7.931552 | 8.138249 | 8.007895 | 8.292897 |
| 6.766747 | 6.588012 | 6.635906 | 8.266161 | 8.497564 | 6.76113 | 6.741921 | 6.517124 | 6.735106 |
| 6.183619 | 6.176308 | 5.891504 | 6.486365 | 6.182685 | 6.391691 | 6.172185 | 6.883616 | 6.537845 |
| 6.927521 | 6.359556 | 6.846582 | NaN | 5.272699 | 6.777893 | 7.294669 | 7.240047 | 7.11139 |
| 7.975989 | 7.924215 | 8.07628 | 8.316598 | 8.516095 | 8.410538 | 8.30605 | 8.413173 | 7.640011 |
| 8.09643 | 8.059245 | 7.993152 | 6.522835 | 6.396195 | 7.620085 | 7.536523 | 7.555737 | 8.090238 |
| 8.78799 | 8.730848 | 8.595552 | 7.608737 | 7.573047 | 8.66219 | 8.541027 | 8.607544 | 8.73409 |
| 9.459816 | 9.353676 | 8.825742 | 8.434528 | 8.45694 | 9.257246 | 9.315171 | 9.396271 | 8.531033 |
| 7.18322 | 7.046623 | NaN | 8.787174 | 8.031796 | 6.530712 | 5.700794 | 5.925675 | 7.215053 |
| 8.561465 | 7.815132 | 8.011531 | 8.520806 | 8.434762 | 6.804066 | 7.690283 | 7.321596 | 7.722859 |
| 9.472231 | 9.345131 | 9.315116 | 9.580328 | 9.512021 | 9.160307 | 9.162152 | 9.082697 | 9.742472 |
| 5.356861 | 5.810072 | 5.661568 | 5.223002 | 4.935242 | NaN | NaN | 4.510582 | 5.720565 |

-Bruker Confidential-

| | | | | | | | | |
|---|---|---|---|---|---|---|---|---|
| 8.772293 | 8.224524 | 8.177869 | 7.613841 | 7.130044 | 8.258931 | 8.315891 | 8.089163 | 7.261011 |
| 6.487551 | 6.622925 | 6.480136 | 6.847878 | 6.940557 | 6.554651 | 6.27143 | 6.292685 | 6.07744 |
| 7.991891 | 7.774031 | 7.92506 | 8.539682 | 8.629632 | 6.192707 | 5.581004 | 6.331736 | 6.214559 |
| 10.90928 | 10.80086 | 11.00065 | 10.74044 | 10.78447 | 10.28452 | 10.29857 | 10.42731 | 9.801337 |
| 6.479992 | 6.168631 | 6.299977 | 5.674561 | 5.835053 | 5.78403 | 6.557948 | 6.286001 | 6.203093 |
| 6.08278 | 5.363673 | 5.261416 | 5.130056 | 4.728499 | 6.620485 | 5.933232 | 6.19436 | 5.447244 |
| 9.111534 | 9.077943 | 8.963476 | 7.395088 | 7.507361 | 8.871884 | 8.553227 | 8.750817 | 9.112113 |
| 7.40344 | 7.749107 | 7.31876 | 6.675585 | 6.684416 | 7.354118 | 7.245202 | 7.398508 | 7.323308 |
| 7.903199 | 7.433113 | 7.797274 | 6.65767 | NaN | 7.273969 | NaN | NaN | 6.346361 |
| 6.672459 | 7.046532 | 7.064709 | 5.930195 | 6.224183 | 6.954892 | 7.271431 | 7.087581 | 6.620752 |
| NaN | NaN | NaN | NaN | NaN | NaN | 6.652178 | NaN | 5.473892 |
| 7.063302 | 4.84936 | 6.544868 | 7.431067 | 6.634199 | 6.398116 | 6.035153 | 6.27056 | 6.536101 |
| 8.265236 | 8.375413 | 8.195682 | 6.810931 | 6.871614 | 8.345012 | 8.216779 | 8.282602 | 8.298062 |
| 8.408373 | 8.392977 | 8.301156 | 7.030907 | 7.304133 | 8.728179 | 8.686007 | 8.696657 | 8.425884 |
| 4.551917 | 3.982239 | 4.668603 | 4.685544 | 5.289595 | 5.189011 | 3.80608 | 4.853496 | 4.237681 |
| 8.103043 | 8.166883 | 8.388005 | 7.718725 | 7.647885 | 8.57849 | 8.855345 | 8.588966 | 8.582186 |
| 7.673622 | 7.71905 | 7.570443 | 7.895888 | 6.958933 | 7.866418 | 8.298317 | 8.117235 | 6.387779 |
| 8.495376 | 8.391618 | 8.395397 | 7.144706 | 6.952441 | 8.351908 | 8.226621 | 8.337711 | 7.426651 |
| 6.212055 | 6.856434 | 6.119705 | 6.239128 | 6.321774 | 6.936533 | 6.608772 | 6.664749 | 6.090837 |
| 6.085387 | 6.089107 | 6.322734 | 6.147099 | 6.605213 | 5.025043 | NaN | 5.420041 | 7.006689 |
| 5.557254 | 5.532339 | 5.467826 | NaN | 5.03229 | 5.168576 | 5.440581 | 5.759627 | 5.715837 |
| 10.5358 | 10.38235 | 10.65993 | 8.776548 | 8.907571 | 8.607774 | 8.434459 | 8.022973 | 8.453134 |
| 7.133545 | 6.876029 | 6.5754 | 6.812893 | 6.742456 | 6.30796 | 6.364342 | 6.243504 | 6.97658 |
| 1.627125 | NaN | NaN | NaN | NaN | NaN | NaN | NaN | 6.063656 |
| 6.106533 | 6.331595 | 6.744246 | 5.208769 | 7.110582 | 7.318825 | 7.14673 | 6.616679 | 6.611722 |
| 7.828401 | 7.789438 | 7.858956 | NaN | NaN | 8.735959 | 8.803015 | 8.578655 | 8.030918 |
| 6.502971 | 6.8685 | 6.765264 | 9.24138 | 9.145692 | 5.808049 | 5.793668 | 6.001104 | 5.506316 |
| 6.088885 | 6.340288 | 6.977015 | 7.396329 | 7.44747 | 6.963996 | 6.885576 | 7.429394 | 7.672408 |
| 9.758292 | 9.730783 | 9.717181 | 10.51555 | 10.33415 | 9.582654 | 9.437498 | 9.396707 | 9.718738 |
| 8.244535 | 8.074595 | 8.258795 | 8.770917 | 8.311954 | 6.908254 | 7.046803 | 7.022723 | 6.86294 |
| 7.223703 | 7.958135 | 7.395606 | 9.055766 | 8.943561 | 7.688935 | 7.238955 | 7.123645 | 7.365583 |
| 7.874071 | 7.924144 | 7.978048 | 8.862408 | 8.903329 | 7.246284 | 6.969995 | 7.416933 | 6.753469 |
| 6.799503 | 5.773914 | 3.535014 | NaN | NaN | 5.202834 | NaN | 6.555285 | NaN |
| 6.003717 | 5.305877 | 5.246166 | 6.170665 | 6.40647 | 6.529445 | 6.851331 | 6.791237 | 4.788149 |
| 8.220447 | 6.902169 | 8.168987 | 6.38087 | 6.488206 | 7.910154 | 7.852957 | 7.851301 | 7.268959 |
| 7.349621 | 7.705033 | 7.529384 | 5.576939 | 5.246522 | 6.456036 | 5.912088 | 7.0467 | 7.179552 |
| 8.026829 | 7.345988 | 7.934003 | 6.106589 | 6.029526 | 7.45386 | 7.525125 | 7.736925 | 7.425484 |
| 7.96771 | 7.174665 | 7.442917 | 6.878772 | 7.329715 | 8.119937 | 8.388573 | 8.41228 | 7.111988 |
| 7.649118 | 7.984497 | 7.712902 | 6.736371 | 7.026983 | 7.874937 | 7.683507 | 7.667037 | 7.384863 |
| NaN | NaN | NaN | 8.559211 | 6.516436 | NaN | NaN | NaN | NaN |
| 6.655777 | 6.793691 | 6.825354 | 7.580662 | 7.75267 | 6.874881 | 6.968204 | 6.915919 | 6.478452 |
| 5.800298 | 6.407728 | 6.186024 | 5.10665 | 5.468471 | 6.797303 | 6.03537 | 4.714633 | 5.400165 |
| 5.678207 | 5.882069 | 5.67222 | 6.54746 | 6.350615 | 6.545385 | 6.769214 | 6.345537 | 5.729771 |
| 5.647403 | 6.177098 | 6.37533 | 7.425225 | 7.605602 | 6.89773 | 7.20975 | 7.30408 | 7.788178 |
| 8.860443 | 8.697826 | 8.897968 | 9.228846 | 9.254093 | 8.32063 | 8.255656 | 8.326317 | 8.05957 |
| 7.124082 | 7.061991 | 6.816925 | 6.631896 | 6.941638 | 7.625791 | 7.722915 | 7.717994 | 7.080422 |
| 6.30813 | 6.226835 | NaN | NaN | NaN | 7.945848 | NaN | 8.004796 | NaN |

| | | | | | | | | |
|---|---|---|---|---|---|---|---|---|
| 6.683156 | 6.976231 | 6.713241 | 7.046871 | 6.894537 | 6.544991 | 6.845749 | 6.619827 | 7.038573 |
| 8.983415 | 9.056489 | 8.979746 | 7.807903 | 7.066236 | 8.619316 | 8.572227 | 8.800841 | 8.259459 |
| 6.830086 | 6.577901 | 6.644705 | 5.563548 | 4.825917 | 7.201355 | 7.29337 | 7.155108 | 6.371218 |
| 5.553818 | NaN | 6.252646 | 5.485717 | 5.226393 | 6.456882 | 6.69101 | 6.776785 | 5.905687 |
| 6.975226 | 6.497652 | 6.633092 | 5.693941 | 6.318443 | 7.025678 | 6.779172 | 6.664762 | 6.405501 |
| 6.543365 | 6.656876 | 6.784291 | 4.842463 | 5.346402 | 6.825387 | 7.249043 | 6.747449 | 5.96896 |
| 8.340583 | 8.013419 | 8.422793 | 9.047209 | 9.317054 | 7.691227 | 8.236423 | 8.060588 | 7.660347 |
| 8.790794 | 8.692637 | 8.631742 | 7.789177 | 7.464824 | 8.273406 | 8.273306 | 8.032099 | 8.150398 |
| 8.144968 | 7.433883 | 7.525828 | 7.849984 | 7.432694 | 7.276222 | 7.748074 | 7.59221 | 7.191147 |
| 6.173164 | 6.158687 | 6.275767 | 7.77373 | 7.883229 | 5.300245 | 5.05792 | 5.289447 | 6.64087 |
| 8.152063 | 8.131479 | 8.196133 | 6.568397 | 6.791397 | 8.198363 | 8.363773 | 8.488994 | 8.141037 |
| 9.292665 | 9.48818 | 9.313312 | 9.313144 | 9.353436 | 9.717527 | 9.604787 | 9.651267 | 9.611548 |
| 6.601106 | 6.698381 | 6.466372 | 7.964201 | 7.911087 | 4.851877 | 5.028022 | 5.250427 | 5.416806 |
| 9.334301 | 9.228074 | 9.131622 | 8.845916 | 8.815772 | 7.997043 | 8.168434 | 7.933311 | 7.730216 |
| 8.541961 | 8.25663 | 8.389391 | 8.418274 | 8.510809 | 7.552421 | 7.349575 | 7.42739 | 6.923328 |
| 7.547514 | 7.335413 | 7.597648 | 8.162388 | 8.075973 | 7.299172 | 7.532859 | 7.847754 | 6.841297 |
| 7.30953 | 7.832379 | 7.423291 | 6.593727 | 6.558476 | 7.80452 | 7.618759 | 7.602438 | 7.772135 |
| 10.09355 | 10.01813 | 9.914302 | 11.72557 | 11.85935 | 9.889525 | 9.758632 | 9.805027 | 10.26977 |
| 9.512593 | 9.34842 | 9.270794 | 9.697673 | 9.840327 | 9.306084 | 9.402987 | 9.335606 | 9.144177 |
| NaN | 7.387503 | 6.631224 | 7.036006 | 7.040527 | 6.098235 | 6.194854 | 6.195779 | 6.345207 |
| 8.480441 | 8.847416 | 8.062209 | 8.485395 | 7.843074 | 7.382525 | 7.127837 | 7.238129 | 7.269496 |
| 7.504847 | 7.246835 | 7.284576 | 7.931052 | 7.848703 | 8.552147 | 8.619023 | 8.034548 | 8.358669 |
| 7.13368 | 6.58564 | 6.507017 | 7.707669 | 7.527892 | 7.216055 | 6.732833 | 6.322133 | 5.781094 |
| 8.602873 | 8.625648 | 8.565509 | 7.349874 | 7.215415 | 8.980591 | 8.842819 | 9.490203 | 8.631644 |
| 9.236897 | 9.416946 | 9.376196 | 7.788658 | 7.83122 | 8.687035 | 8.864787 | 8.911172 | 9.760345 |
| 6.473241 | 6.259129 | 6.41748 | 4.986372 | 5.3738 | 5.980782 | 6.016876 | 5.864066 | 6.456239 |
| 6.012169 | 6.342509 | 6.057973 | 6.640937 | 6.7349 | 6.248486 | 5.625433 | 5.89963 | 5.691693 |
| 8.586061 | 6.703439 | 8.721372 | 7.479004 | 6.767714 | 7.036914 | 7.311358 | 7.304709 | 8.408202 |
| 6.428538 | 6.270099 | 6.716803 | 8.465238 | 8.019253 | 6.469615 | 7.098111 | 7.073184 | 6.659831 |
| 7.532154 | 7.547887 | 7.473377 | 8.275884 | 8.393474 | 7.936488 | 7.79215 | 7.613239 | 7.711774 |
| 12.05967 | 10.07573 | 12.16974 | 12.01216 | 12.18362 | 6.729383 | NaN | 5.949505 | 9.366066 |
| 8.300576 | 8.945066 | 8.390612 | 10.09608 | 10.0786 | 7.15867 | 7.053755 | 7.25429 | 6.902472 |
| 7.541918 | 7.418668 | 7.568229 | 6.580098 | 6.644732 | 7.781495 | 7.219088 | 7.178395 | NaN |
| 7.911542 | 6.815796 | 7.469643 | 7.692924 | 7.707538 | 6.605677 | 5.934127 | 5.562922 | 6.599111 |
| 7.53774 | 7.521321 | 7.163716 | 7.415465 | 7.638531 | 7.447848 | 6.740533 | 7.04241 | 7.346245 |
| 5.435722 | 5.56795 | 5.834538 | NaN | NaN | NaN | NaN | NaN | 5.570674 |
| 7.124786 | 3.35026 | 5.807194 | 4.801962 | 4.321618 | 5.708889 | 6.371906 | 5.958948 | 4.838059 |
| 7.753795 | 7.571361 | 7.56371 | 7.865196 | 7.681598 | 7.598821 | 7.637926 | 7.946454 | 7.954871 |
| NaN | 6.731345 | NaN | 6.553393 | 7.36411 | 6.439064 | 7.154594 | 6.635284 | 7.598025 |
| 10.13794 | 10.28487 | 10.23471 | 8.990534 | 9.079361 | 10.4477 | 10.25966 | 10.40175 | 9.635097 |
| NaN | NaN | NaN | 5.417527 | 5.973516 | NaN | NaN | NaN | NaN |
| 6.541167 | 5.932416 | 6.486383 | 6.09035 | 5.936598 | 6.55051 | 6.385365 | 6.509797 | 6.79581 |
| 7.382892 | 7.368442 | 7.127791 | 6.799284 | 6.932341 | 7.409488 | 7.353361 | 7.02618 | 6.960277 |
| 7.209479 | 6.881012 | 7.183726 | 6.258673 | 6.109388 | 7.424452 | 7.623796 | 6.866159 | 6.831357 |
| 6.83052 | 6.414517 | 6.400982 | 6.840953 | 6.659494 | 6.70875 | 6.608179 | 6.383788 | 6.058108 |
| 6.637947 | 6.959124 | 6.758343 | 5.747878 | 6.167726 | 7.498161 | 7.0813 | 7.253436 | 7.745362 |
| 9.649881 | 9.302667 | 9.009988 | 8.83138 | 7.663444 | 7.751466 | 7.841815 | 7.692517 | 6.988401 |

-Bruker Confidential-

| | | | | | | | | |
|---|---|---|---|---|---|---|---|---|
| 7.292324 | 7.436907 | 7.583353 | 10.49259 | 10.59626 | 7.301652 | 7.119524 | 7.239615 | 7.44095 |
| 8.043464 | 7.461882 | 7.793715 | 7.381243 | 7.325161 | 7.36247 | 7.452403 | 6.848843 | 5.279993 |
| 4.422813 | 4.644666 | 4.830048 | 5.646324 | 5.609868 | 3.023222 | 3.690366 | 3.520455 | 4.674815 |
| 8.371067 | 8.61002 | 8.160188 | 8.176992 | 7.896796 | 8.472744 | 8.580282 | 8.445672 | 8.352291 |
| 5.140012 | 5.100746 | NaN | 5.819688 | 6.040939 | 4.11249 | NaN | 4.557213 | 3.800502 |
| 7.398826 | 7.319691 | 7.198491 | 7.204029 | 7.182241 | 7.664195 | 7.385078 | 8.035663 | 7.687458 |
| 8.763157 | 8.865159 | 8.896797 | 8.797152 | 8.800438 | 6.882533 | 7.036442 | 6.787489 | 6.750712 |
| 6.243017 | 6.738909 | 6.364769 | 8.240256 | 8.63986 | 8.60116 | 8.316524 | 8.527109 | 8.791554 |
| 6.731761 | 6.295604 | 6.510854 | 4.69549 | 5.864145 | 6.647775 | 6.277884 | 5.941949 | 6.362186 |
| 7.365336 | 7.256102 | 7.145111 | 7.674016 | 7.875926 | 7.456262 | 7.553715 | 7.511402 | 7.451185 |
| 8.300079 | 7.93264 | 8.028793 | 8.161539 | 8.214436 | 8.030483 | 8.265224 | 8.389333 | 7.608034 |
| 6.011624 | 7.007079 | 5.605639 | 7.428792 | 7.362602 | 7.166318 | 7.62471 | 7.436631 | 6.708573 |
| 8.247993 | 8.462523 | 8.247738 | 9.122352 | 9.102134 | 7.388551 | 7.692717 | 7.979393 | 7.830735 |
| 7.509657 | 7.106358 | 7.973984 | 6.368385 | NaN | 8.258423 | 8.139485 | 8.028512 | 7.085411 |
| 8.672411 | 8.742405 | 8.66243 | 6.976311 | 7.061078 | NaN | 8.45954 | 8.702223 | 7.27068 |
| 6.746926 | 5.298076 | 6.808893 | 6.821481 | 6.62367 | 6.624723 | 6.409204 | 6.409312 | 6.714937 |
| 8.305492 | 8.392053 | 8.394744 | 7.207623 | 7.257304 | 8.674783 | 8.559517 | 8.751158 | 8.354813 |
| 6.954535 | 6.779049 | 6.60125 | 8.054837 | 7.807914 | 5.114846 | 7.388757 | 6.210125 | 6.18054 |
| 6.37894 | 5.988776 | 5.501434 | 5.34535 | 5.278291 | 6.065517 | 6.714145 | NaN | 6.087749 |
| 7.007289 | 7.412494 | 7.120047 | 6.665907 | 7.191465 | 7.22242 | 7.573814 | 7.406611 | 7.556653 |
| 9.016212 | 9.001338 | 8.842495 | 7.735595 | 7.445896 | 9.381643 | 9.154455 | 9.077319 | 8.834743 |
| 7.05628 | 6.778259 | 6.936573 | 6.3563 | 6.696452 | 6.943584 | 6.744777 | 6.740928 | 6.814859 |
| NaN | NaN | NaN | 8.008446 | 6.502556 | NaN | NaN | 7.20914 | 7.328303 |
| 8.740874 | 8.889348 | 8.658288 | 9.144703 | 9.054173 | 9.842437 | 9.674642 | 9.590044 | 9.424927 |
| 8.022815 | 8.265784 | 8.201548 | 7.642596 | 8.303965 | 8.099933 | 8.207724 | 8.220315 | 7.232394 |
| 4.457788 | 5.301529 | 5.385023 | 8.310657 | 8.25577 | 5.724152 | 4.692743 | 5.871898 | 5.028925 |
| 10.65976 | 10.01667 | 10.25822 | 7.225883 | 7.629887 | 7.677197 | NaN | NaN | 8.022365 |
| 7.264266 | 5.688577 | 6.649382 | NaN | 5.562083 | 7.077625 | 6.646235 | 7.007455 | 7.448834 |
| 6.076669 | 6.116041 | 6.402514 | 6.032931 | 6.097802 | 5.619006 | NaN | NaN | 6.599702 |
| 7.816357 | 7.794418 | 7.214819 | 7.100727 | 6.911468 | 6.906641 | 6.898093 | 7.253107 | 6.471315 |
| 8.001227 | 6.056332 | 8.108904 | 5.996866 | 6.22086 | 6.556798 | 6.320355 | 6.504737 | 8.723377 |
| 7.764606 | 7.734911 | 7.952159 | 5.681252 | 5.848107 | 7.760916 | 7.676173 | 7.894042 | 7.075454 |
| 8.364487 | 7.74102 | 7.552664 | 9.443577 | 9.531334 | 8.253325 | 7.721984 | 8.205353 | 7.112491 |
| 7.377455 | 7.212885 | 7.253911 | 8.297905 | 7.988616 | 6.816753 | 6.469603 | 6.670537 | 5.968053 |
| NaN | NaN | NaN | 5.355663 | NaN | NaN | NaN | NaN | 7.336456 |
| 7.677495 | 7.669698 | 7.863877 | 8.640143 | 9.03264 | 8.24829 | 7.809437 | 7.960328 | 8.014274 |
| 5.615473 | 5.912022 | 5.833502 | 5.930986 | 6.103071 | 5.323064 | 5.508804 | 5.167348 | 5.604568 |
| 7.750537 | 7.504598 | 7.291316 | 7.121514 | 6.844646 | 7.961808 | 8.252346 | 8.027301 | 7.399272 |
| 9.178494 | 9.114043 | 8.988286 | 9.788526 | 9.73837 | 8.738155 | 8.705276 | 8.770454 | 8.779945 |
| 8.286439 | 8.45081 | 8.399978 | 7.410418 | 7.539537 | 8.117902 | 8.154791 | 8.051128 | 8.514774 |
| 6.741059 | 7.054073 | 7.028377 | 6.081143 | 6.033358 | 7.405049 | 6.940473 | 6.992751 | 6.674845 |
| NaN | 7.994 | 6.841587 | 7.246098 | 6.27458 | 5.854198 | 5.260608 | 6.139331 | 5.513753 |
| 9.027932 | 8.657013 | 8.711348 | 6.929205 | 6.968474 | 8.770467 | 8.460365 | 8.671585 | 7.931228 |
| 6.475073 | 6.982696 | 6.662108 | 7.039989 | 7.432878 | NaN | 6.28252 | 6.564552 | 7.137001 |
| 7.784807 | 7.168563 | 7.505158 | 7.585867 | 7.784552 | 7.268422 | 7.214124 | 7.442208 | 7.012852 |
| 9.150211 | 9.274717 | 8.901371 | 8.242475 | 8.310627 | 8.883512 | 8.868405 | 8.781501 | 8.293211 |
| 8.752084 | 8.537158 | 8.290374 | 9.083735 | 8.722471 | 8.306562 | 8.176054 | 8.204925 | 7.307492 |

| | | | | | | | | |
|---|---|---|---|---|---|---|---|---|
| 7.171708 | 7.47822 | 7.26415 | 7.151389 | 7.021856 | 5.503731 | NaN | NaN | 5.957355 |
| NaN | NaN | NaN | 5.958801 | 6.281158 | NaN | 5.97485 | NaN | 6.085253 |
| 6.965704 | 6.121342 | 5.260611 | NaN | 6.071486 | 5.389025 | 4.580024 | 4.20084 | 6.405766 |
| 7.771296 | 7.79144 | 7.853201 | 6.28501 | 6.431053 | 7.811946 | 7.305199 | 7.540973 | 7.727732 |
| 5.62323 | 5.356309 | 5.6959 | 6.567653 | 6.830032 | 5.002493 | 5.543689 | 5.675308 | 5.971542 |
| NaN | 8.276626 | NaN | 7.87867 | 8.499743 | NaN | 7.052184 | 6.408309 | 6.056922 |
| 6.174725 | 6.264435 | 6.2951 | 7.205801 | 7.117762 | 5.284467 | 4.521273 | 5.27891 | 5.163037 |
| 6.858294 | 6.934485 | 6.584259 | 6.616793 | 7.822001 | 7.379076 | 6.195633 | 5.883007 | 5.684644 |
| 7.672304 | 7.042966 | 7.480573 | 6.620676 | 7.39437 | 7.437393 | 7.652495 | 7.461283 | 7.678826 |
| 7.038651 | 6.90731 | 7.250026 | 6.161924 | 6.047164 | 6.36167 | 6.398451 | 6.293586 | 6.332887 |
| 7.389807 | 7.198089 | 7.417428 | 6.375088 | 6.168183 | 7.644222 | 7.433438 | 7.515897 | 6.951184 |
| 5.61549 | 3.846415 | 5.055052 | 6.519199 | 5.242296 | 5.198349 | 5.090854 | 5.153536 | 5.660256 |
| 7.222557 | 7.015739 | 7.465246 | 5.974502 | 6.206313 | 8.214043 | 7.377264 | 7.763391 | 6.097974 |
| 6.728633 | 5.812941 | 6.18048 | 6.685324 | 6.716926 | 7.630777 | 8.052046 | 7.929514 | 5.772049 |
| 7.478166 | 7.653884 | 6.134033 | 7.553904 | 7.575029 | NaN | NaN | 6.518536 | 5.800875 |
| 3.921441 | 3.072388 | 4.608654 | 5.19165 | 4.72861 | 6.178049 | 5.774249 | 5.771523 | 4.63953 |
| 9.231976 | 9.401299 | 9.506179 | 9.35704 | 8.880644 | 9.074052 | 9.445084 | 9.479271 | 8.648025 |
| 8.350581 | 7.867894 | 8.126195 | 8.977098 | 8.790514 | 8.187347 | 8.214572 | 8.132346 | 8.034452 |
| 6.446429 | 6.822262 | 6.329876 | 7.254638 | 6.698359 | 6.436686 | 6.404301 | 6.422703 | 6.895573 |
| 8.414173 | 8.381259 | 8.350076 | 8.43335 | 8.418366 | 8.551869 | 8.468906 | 8.456518 | 7.820578 |
| 5.752854 | 4.801948 | 6.282122 | 5.544077 | 5.664653 | 6.314429 | 5.933857 | 6.104732 | 5.420947 |
| 6.642496 | 6.363252 | 5.935994 | 6.993426 | 7.149678 | 6.23892 | 6.271199 | 6.32065 | 6.878029 |
| 6.432203 | 6.518364 | 5.777415 | 6.567392 | 6.510635 | 6.366306 | 6.561846 | 6.430477 | 6.118385 |
| 4.236766 | 4.385086 | 4.584218 | NaN | NaN | 5.354591 | 4.556253 | 5.130477 | 5.5879 |
| 8.162529 | 8.231422 | 7.811713 | 9.755493 | 9.651349 | 8.586791 | 8.935183 | 8.915143 | 8.297072 |
| 6.361573 | 3.825868 | 6.317027 | 5.590467 | 5.805499 | 6.438646 | 6.470436 | 6.399532 | 6.585057 |
| 7.172604 | 7.167411 | 7.221313 | 6.217621 | 6.185989 | 7.120767 | 7.307594 | 7.411126 | 7.15714 |
| 7.759669 | 7.798191 | 7.353501 | 7.971754 | 7.896188 | 7.892547 | 7.860276 | 7.81918 | 7.081841 |
| 6.602914 | 5.59802 | 6.483734 | 4.553753 | NaN | 6.629436 | 6.877179 | 7.503324 | 6.432277 |
| 10.13795 | 9.900004 | 9.865692 | 8.147105 | 8.272987 | 10.2272 | 10.13049 | 10.12769 | 9.425292 |
| 7.725033 | 7.150874 | 7.603373 | 6.180306 | 6.809798 | 7.556541 | 6.557174 | 7.556909 | 6.438392 |
| 6.86978 | 5.856584 | 5.822961 | 6.263095 | 6.032841 | 7.476701 | 7.241042 | 6.916232 | 5.628382 |
| 9.948691 | 10.05143 | 10.01545 | 11.70079 | 11.51539 | 11.48491 | 11.24954 | 11.28071 | 9.848709 |
| 8.283858 | 8.2373 | 8.255013 | 7.119977 | 7.225512 | 8.211249 | 7.816458 | 7.716781 | 7.831071 |
| 9.09656 | 8.999781 | 9.148026 | 8.506026 | 8.497079 | 9.481058 | 9.554709 | 9.375994 | 8.568428 |
| NaN | 6.924962 | 6.627485 | 6.392144 | 4.506187 | 5.463446 | 5.202266 | 5.95107 | 7.295476 |
| NaN | 7.185586 | 7.194588 | 8.386289 | 8.14657 | 7.300694 | 7.305785 | 6.758235 | 7.164554 |
| 10.43925 | 10.08782 | 9.911091 | 9.784997 | 9.005886 | 9.724202 | 8.670617 | 9.498127 | 8.803323 |
| 8.313819 | 8.263683 | 8.493599 | 7.840916 | 7.997923 | 8.814807 | 8.728937 | 8.64217 | 7.9497 |
| NaN | NaN | NaN | 7.505213 | 6.291791 | 6.112957 | NaN | NaN | 4.722457 |
| 7.206508 | 8.035945 | 7.878752 | 7.169422 | 6.493967 | NaN | NaN | NaN | 5.127851 |
| NaN | NaN | NaN | 7.251809 | 6.855596 | NaN | NaN | NaN | 6.142096 |
| NaN | NaN | NaN | 5.417839 | 5.469078 | NaN | 7.317246 | NaN | 6.767935 |
| 7.241552 | 7.115285 | 7.469064 | 5.586293 | 6.231085 | 6.682504 | 6.797545 | 6.165355 | 6.635371 |
| 9.6235 | 9.730102 | 9.678073 | 8.7472 | 8.880282 | 9.436863 | 9.623939 | 9.578162 | 9.769946 |
| 7.249654 | 7.714176 | 7.626951 | 7.607921 | 6.612436 | NaN | 6.466403 | 6.397758 | 7.478754 |
| 6.46442 | 6.224333 | 6.06131 | NaN | NaN | 7.625706 | 7.468683 | 7.941916 | 6.945129 |

| | | | | | | | | |
|---|---|---|---|---|---|---|---|---|
| 9.528014 | 9.578711 | 9.582783 | 9.084868 | 8.789177 | 8.056116 | 8.958112 | 8.391369 | 8.603924 |
| 8.691769 | 8.350771 | 8.06904 | 8.804757 | 8.786206 | 7.887885 | 7.818302 | 7.733782 | 7.816332 |
| 7.560301 | 7.541746 | 7.508359 | 6.986226 | 7.001883 | 7.804017 | 7.596264 | 7.974703 | 7.744548 |
| 9.075216 | 9.118923 | 8.627686 | 8.355927 | 8.468555 | 8.839668 | 8.655993 | 8.53618 | 7.910476 |
| 7.102239 | 7.120304 | 7.270319 | NaN | 4.70728 | 8.141196 | 8.060732 | 7.746459 | 7.50668 |
| 6.747599 | 6.551546 | 6.350928 | 6.215294 | 6.307841 | 6.804425 | 6.688902 | 6.742645 | 6.294557 |
| 5.22249 | 6.004426 | 6.334784 | 6.962976 | 7.073105 | 5.470169 | 5.330047 | 5.207233 | 6.082164 |
| 4.728653 | 4.598331 | 3.611894 | NaN | 5.003593 | 5.156079 | 4.7241 | 4.858585 | 5.163268 |
| 9.085849 | 8.581905 | 8.457427 | 8.625386 | 8.371939 | 7.73886 | 8.178247 | 8.128255 | 8.985554 |
| 7.246596 | 7.302503 | 7.086721 | 6.515924 | 5.737431 | 6.489128 | 6.392593 | 6.353248 | 6.647727 |
| 10.03475 | 8.238935 | 9.734189 | 7.191635 | 8.026525 | 8.661372 | 8.90526 | 8.601525 | 9.622383 |
| 6.233681 | 7.488289 | 7.766424 | 4.858158 | 5.269335 | NaN | 6.471849 | NaN | 6.758967 |
| 6.831334 | 7.039804 | 6.692026 | 6.748004 | 6.903908 | 7.749581 | 7.72212 | 7.478605 | 7.367441 |
| 6.83812 | 6.926901 | 7.144237 | 7.457335 | 7.336286 | 6.874111 | 7.236479 | 6.603849 | 6.492249 |
| NaN | NaN | 6.566898 | NaN | NaN | 7.809452 | 7.710906 | 7.785582 | 8.13534 |
| 8.031164 | 7.969226 | 7.690773 | 7.916958 | 8.019315 | 8.418687 | 8.276089 | 8.323765 | 7.568549 |
| 7.146134 | 6.29676 | 6.592445 | NaN | 5.854001 | 6.365875 | 6.544072 | 6.228693 | 5.822473 |
| 6.737079 | 6.120355 | 5.708217 | 7.532532 | 7.344125 | 7.084282 | 6.828533 | 7.085555 | 6.699986 |
| NaN | 6.72581 | NaN | NaN | NaN | 7.597115 | 7.664596 | 7.570733 | NaN |
| 6.658143 | 6.786818 | 6.641305 | 6.829741 | 7.218752 | 6.738132 | 6.898717 | 6.473213 | 6.970897 |
| 7.862887 | 7.792754 | 7.711949 | 8.263566 | 7.37784 | 7.456324 | 7.389962 | 7.522852 | 7.174977 |
| 6.812092 | 6.959426 | 6.798874 | 6.530961 | 6.485126 | 6.769905 | 6.546637 | 6.752631 | 7.18553 |
| NaN | NaN | 8.828882 | 7.99694 | 7.404055 | NaN | NaN | NaN | NaN |
| 8.19278 | 6.34613 | 7.852015 | 7.277589 | 6.34101 | 8.9702 | 8.825038 | 8.852847 | 7.697319 |
| 6.761147 | 5.860787 | 6.522661 | 5.786434 | 6.178168 | 5.983372 | 5.602755 | 6.078671 | 6.869332 |
| 4.875684 | 5.001343 | 4.847297 | 5.488894 | 6.321421 | 5.157258 | 4.48843 | 5.243511 | 4.369285 |
| 6.265903 | 5.601717 | NaN | 5.89819 | 6.107493 | NaN | NaN | NaN | 6.360773 |
| 8.26065 | 7.34545 | 8.031006 | 6.597858 | 6.267507 | 8.178209 | 8.095084 | 7.966788 | 5.9532 |
| 6.720485 | 6.046979 | 6.49122 | 5.313546 | 6.397415 | 6.891182 | 7.021368 | 6.938931 | 6.337228 |
| 8.141777 | 7.534689 | 7.950277 | 8.36753 | 7.804811 | NaN | 6.762747 | NaN | 8.086232 |
| 6.328054 | 6.664334 | 6.358099 | 7.854967 | 7.142273 | 5.751189 | 6.048412 | 6.525205 | 7.131223 |
| 8.620538 | 8.382361 | 8.346477 | 8.264226 | 8.289609 | 8.09203 | 8.263701 | 8.30767 | 7.134138 |
| 6.324341 | 6.368408 | 6.54504 | 5.896726 | 5.204243 | 6.686305 | 5.750256 | 6.544671 | 6.303106 |
| 8.165414 | 8.122938 | 8.087905 | 8.465152 | 8.336779 | 7.771833 | 7.87818 | 7.866052 | 8.002796 |
| 6.844699 | 6.703184 | 6.679655 | 8.877249 | 8.496563 | 7.266129 | 7.398123 | 7.595307 | 8.225129 |
| 6.079681 | 6.31713 | 5.943609 | 5.621156 | 5.647349 | 5.710291 | 6.133259 | 5.599766 | 6.321911 |
| 8.799304 | 8.696867 | 8.913481 | 7.491179 | 7.842561 | 8.95885 | 9.155513 | 9.162527 | 8.855342 |
| 8.72576 | 8.273942 | 8.314897 | 7.598099 | 7.336069 | 6.500089 | 6.582828 | 6.323325 | 8.604261 |
| NaN | 6.843595 | NaN | NaN | NaN | 5.878271 | 5.737463 | 5.747973 | 6.440159 |
| 8.031837 | 8.251253 | 8.38667 | 8.047452 | 8.493677 | 7.459891 | 7.146096 | 7.105721 | 7.268139 |
| 9.847142 | 9.256545 | 9.443827 | 10.0956 | 9.606158 | 10.00736 | 9.355165 | 9.760957 | 9.288915 |
| 7.013245 | 7.918175 | 7.350935 | 10.26408 | 10.6296 | 7.253386 | 6.607311 | 6.815117 | 6.280204 |
| 5.459222 | 5.219528 | 5.568198 | 5.674134 | 5.735433 | 5.771496 | 5.330518 | 5.267331 | 6.093396 |
| NaN | 2.310091 | NaN | 2.534388 | NaN | NaN | NaN | NaN | NaN |
| 7.851298 | 7.364424 | 7.4895 | 7.657875 | 7.731946 | 7.407182 | 6.882599 | 7.495498 | 7.591052 |
| NaN | NaN | NaN | NaN | 3.703658 | NaN | 4.781652 | NaN | 4.834141 |
| 9.882808 | 10.02967 | 10.07312 | 9.31754 | 9.089792 | 9.534798 | 9.759884 | 9.131989 | 9.369222 |

| | | | | | | | | |
|---|---|---|---|---|---|---|---|---|
| 8.26034 | 8.169586 | 7.770089 | 7.848487 | 7.677772 | 7.799998 | 7.667439 | 7.69714 | 6.713681 |
| 5.921076 | 5.767033 | 6.30582 | 4.568655 | 5.186808 | 5.429508 | 5.775579 | 5.795118 | 5.441604 |
| 8.591364 | 8.403512 | 8.323634 | 9.835236 | 9.388168 | 8.144422 | 7.981739 | 7.775334 | 8.373884 |
| 6.349007 | 6.700844 | 6.291493 | 5.111688 | 5.695461 | 7.014271 | 6.909135 | 6.678753 | 6.024148 |
| 6.5432 | 6.372379 | 6.889253 | 6.82666 | 6.916864 | 7.023525 | 7.251459 | 7.32274 | 7.834077 |
| 8.532394 | 8.33461 | 8.531823 | 8.289816 | 8.27622 | 8.08447 | 8.037561 | 8.192198 | 8.172259 |
| 7.719782 | 7.261213 | 7.196864 | 7.93527 | 7.967847 | 7.497293 | 6.314473 | 6.633462 | 7.204611 |
| 6.315533 | 7.016008 | NaN | NaN | NaN | 7.260674 | 7.620672 | 7.664934 | 7.430644 |
| 8.746495 | 8.324831 | 8.569847 | 6.904984 | 6.875522 | 8.698638 | 8.382484 | 8.730618 | 8.332506 |
| 6.642409 | 6.367966 | 6.512268 | 7.583098 | 7.733181 | 6.821066 | 6.933868 | 7.073498 | 7.664143 |
| 6.380795 | 5.897454 | 6.261004 | NaN | NaN | 6.695273 | 5.617309 | 5.766764 | 4.047668 |
| 7.541253 | 7.695767 | 7.319639 | NaN | NaN | 8.32585 | 8.003039 | 7.695569 | 7.297017 |
| 6.763002 | 6.594908 | 7.124494 | 6.656849 | 6.20183 | 7.298985 | 7.654421 | 7.419655 | 7.058728 |
| 6.329494 | 7.532417 | 6.654282 | 8.159894 | 8.469048 | NaN | 4.589873 | 5.095174 | 6.806427 |
| 6.475994 | 6.506868 | 6.338014 | 7.763535 | 7.841379 | 6.765617 | 6.963581 | 6.879208 | 6.76559 |
| 6.521739 | 6.359567 | 6.004142 | 6.850653 | 6.628055 | 6.924531 | 5.9209 | 6.278276 | 6.158752 |
| 4.586986 | 4.473129 | 4.272762 | 5.576654 | 5.788867 | 4.079793 | 4.256393 | 4.169585 | 4.902282 |
| 7.421463 | 7.31779 | 7.239944 | 7.063273 | 6.58532 | 5.844158 | 5.632554 | 6.118421 | 6.541494 |
| 10.26647 | 10.34385 | 10.12502 | 10.36849 | 10.55814 | 10.26878 | 9.980654 | 9.96298 | 8.632063 |
| 6.622566 | 6.508742 | 6.473197 | 8.058254 | 8.099011 | 5.413853 | 4.663738 | 5.00743 | 6.020299 |
| 5.719368 | 5.16535 | 5.700729 | 6.685696 | 6.335214 | 5.065708 | 4.894818 | 5.021465 | 7.105376 |
| 7.757847 | 7.762693 | 7.950179 | 7.049191 | 7.317971 | 8.616695 | 8.613452 | 7.99313 | 8.268926 |
| 6.620489 | 6.471007 | 6.492587 | 5.623127 | 6.091253 | 6.530928 | 6.413927 | 6.682672 | 6.595509 |
| 5.756674 | 5.922074 | 6.333102 | 4.618529 | 4.997647 | 5.63085 | 5.732203 | 5.696587 | 5.427526 |
| 6.522877 | 6.896357 | 6.672927 | 6.39197 | 6.42172 | 6.966522 | 7.098346 | 7.138231 | 6.620209 |
| NaN | 5.803429 | 7.45686 | 6.257227 | 7.173282 | 7.538317 | 7.27113 | 7.383661 | 6.183461 |
| 12.84853 | 12.82798 | 10.89258 | 8.864916 | 8.795918 | 8.752808 | 8.45142 | 8.537547 | 8.712787 |
| 5.543561 | 5.315278 | 5.945268 | 5.353552 | 4.642844 | 6.121374 | 6.011116 | 6.017074 | 6.164847 |
| 7.615084 | 7.819525 | 7.474304 | 7.966069 | 7.810209 | 7.664774 | 7.469608 | 7.588469 | 7.558065 |
| 6.859053 | 6.369979 | 7.259486 | NaN | NaN | 7.370255 | 7.437719 | 7.360682 | 6.805071 |
| NaN | 3.922667 | 1.617877 | 4.304874 | 4.543629 | 4.245454 | NaN | NaN | 5.868765 |
| 8.961337 | 9.262482 | 9.318513 | 11.48083 | 11.22038 | 9.030088 | 9.097887 | 9.02213 | 8.632639 |
| 7.330643 | 7.85172 | 7.79124 | 7.100352 | 6.460481 | 7.968142 | 7.936827 | 7.897115 | 7.116782 |
| 8.433391 | 8.637752 | 8.452589 | 8.325924 | 8.383012 | 8.189168 | 7.808807 | 7.981065 | 8.246235 |
| 7.608478 | 7.56498 | 7.720516 | 7.511951 | 7.292413 | 7.731136 | 7.699906 | 8.294001 | 7.829521 |
| 5.086687 | 4.856226 | 5.237046 | 5.176948 | 5.367795 | 5.322051 | 5.984084 | 5.818381 | 4.714798 |
| 4.414861 | 5.718002 | NaN | 5.854771 | NaN | NaN | NaN | NaN | 5.623194 |
| 7.327359 | 7.434834 | 7.229209 | 5.48024 | 5.120434 | 7.496476 | 7.35898 | 7.55663 | 7.327268 |
| 7.536445 | 7.335151 | 7.794096 | 6.342109 | 6.781118 | 8.372493 | 8.296182 | 8.410776 | 7.891755 |
| 6.753453 | 6.638699 | 6.935007 | 6.591967 | 6.879669 | 6.819303 | 6.903344 | 6.890934 | 6.074027 |
| 7.800912 | 7.697516 | 7.774395 | 6.46321 | 6.802587 | 7.222093 | 7.42493 | 7.31697 | 7.190678 |
| 8.368102 | 8.253708 | 8.40138 | 7.826278 | 8.04043 | 7.793062 | 7.550652 | 7.754794 | 7.832409 |
| NaN | 5.607807 | 2.616121 | 5.274934 | 5.052736 | 5.781742 | NaN | NaN | 4.943383 |
| 6.163084 | 6.440982 | 6.696564 | 5.01926 | 5.985421 | 5.932611 | NaN | NaN | NaN |
| NaN | NaN | NaN | 7.285027 | 7.210513 | 6.225797 | 6.173468 | 4.862687 | 5.523718 |
| 7.705486 | 7.835837 | 7.764801 | 7.123367 | 7.111971 | 7.989739 | 7.771397 | 7.897938 | 7.770042 |
| NaN | NaN | NaN | NaN | NaN | NaN | NaN | 4.131757 | 6.109561 |

| | | | | | | | | |
|---|---|---|---|---|---|---|---|---|
| 6.456257 | 6.363577 | 6.453055 | NaN | NaN | 6.436727 | 5.969208 | 6.50206 | 6.391277 |
| NaN | NaN | NaN | 6.432374 | 6.371624 | NaN | NaN | NaN | NaN |
| 8.614256 | 8.487126 | 8.733696 | 9.536295 | 9.385695 | 8.024493 | 7.889876 | 7.97926 | 8.676799 |
| 7.668602 | 7.32056 | 8.088511 | 6.797974 | 7.343684 | 8.052513 | 6.987426 | 7.320622 | 6.939279 |
| 6.607546 | 6.570557 | 7.28883 | 7.541628 | 7.350161 | 6.229281 | 6.061367 | 6.103151 | 7.121062 |
| 7.905808 | 7.949172 | 7.93781 | 7.187778 | 7.32919 | 7.93721 | 7.888112 | 7.899932 | 7.728152 |
| 6.805269 | 6.347579 | 6.848688 | 4.55672 | 5.229708 | 6.098334 | 5.71051 | 6.15151 | 6.543559 |
| 6.813508 | 6.633337 | 6.716233 | NaN | 5.616447 | 7.443762 | 7.513778 | 7.301398 | 6.269618 |
| 7.753125 | 8.422601 | 7.937056 | 6.766296 | 6.684786 | 6.49217 | 7.196289 | 7.153909 | 6.79565 |
| NaN | NaN | NaN | 6.650856 | 6.265548 | NaN | NaN | NaN | NaN |
| 6.628165 | 6.851181 | 6.553395 | 6.165132 | 6.305261 | 7.338788 | 7.094313 | 7.198169 | 7.21357 |
| 6.869663 | 7.165615 | 6.859262 | 6.989488 | 6.960324 | 7.740119 | 7.823443 | 7.691838 | 6.703389 |
| 7.423813 | 7.435647 | 6.210637 | 6.425267 | 6.30775 | 7.606853 | 7.396566 | 8.002435 | 7.293344 |
| 6.074405 | NaN | NaN | 7.167313 | 6.73165 | NaN | NaN | NaN | 5.952195 |
| 8.476632 | 8.233902 | 8.450488 | 7.500829 | 7.43199 | 8.504778 | 8.605817 | 8.530412 | 8.216977 |
| 10.12788 | 9.994584 | 10.14205 | 9.637248 | 9.518515 | 10.23518 | 10.15457 | 10.12477 | 9.745735 |
| 6.895629 | 6.775616 | 6.385925 | NaN | NaN | 6.260201 | 6.348777 | 6.070112 | 6.252817 |
| 7.529616 | 7.536954 | 7.423818 | 6.838515 | 6.778362 | 8.13211 | 7.857922 | 7.888259 | 7.850764 |
| 6.991786 | 6.955349 | 7.031363 | 8.661619 | 8.827331 | 4.191074 | NaN | 4.222901 | 6.3446 |
| 5.10618 | NaN | NaN | 7.07672 | 7.682827 | NaN | NaN | NaN | NaN |
| 6.829908 | 5.630888 | 5.96825 | 5.861361 | NaN | 7.398196 | 7.472563 | 7.372619 | 7.187643 |
| 6.866289 | 6.905396 | 6.771219 | 6.712675 | 6.208761 | 6.14112 | 6.331761 | 6.542465 | 6.743441 |
| 8.255821 | 8.141007 | 8.187099 | 7.441679 | 7.761474 | 8.021378 | 8.052016 | 8.042438 | 8.121077 |
| 7.357564 | 7.42918 | 7.352425 | 7.279015 | 7.446912 | 7.533562 | 7.135086 | 7.301959 | 7.560432 |
| 6.155285 | NaN | 4.902265 | 6.229697 | 6.57657 | 6.70276 | 6.368627 | 6.245797 | 6.233028 |
| 6.48534 | 7.002992 | 7.166851 | 5.191974 | 6.383427 | 7.463242 | 7.302829 | 7.583874 | 7.158425 |
| 6.448882 | 6.255341 | 6.435046 | 6.175095 | NaN | NaN | 5.970162 | NaN | 5.840478 |
| 5.490174 | 5.827329 | 6.308295 | 4.246732 | 3.59223 | 7.118313 | 7.010716 | 6.971842 | 5.765962 |
| 5.745702 | 5.919742 | NaN | 5.912291 | 4.941358 | 6.074256 | 6.063006 | NaN | 6.148262 |
| 8.056087 | 8.943488 | 8.439021 | 8.812686 | 8.706364 | 7.873114 | 7.651657 | 7.379797 | 8.665103 |
| 7.403776 | 7.522217 | 7.528281 | 6.514595 | 6.382308 | 7.123266 | 7.677146 | 7.709423 | 6.802391 |
| 7.92502 | 7.967451 | 7.884069 | 8.239992 | 8.173073 | 6.767035 | 7.294716 | 7.513028 | 6.486289 |
| 7.821366 | 8.053539 | 6.603983 | NaN | 6.590392 | 7.074543 | 6.39273 | 6.805919 | 6.91824 |
| 8.445347 | 8.344338 | 8.349215 | 7.766198 | 7.536035 | 8.919609 | 8.732389 | 8.884821 | 7.359393 |
| 6.364364 | 5.93627 | 6.351015 | 6.77478 | 7.096387 | 5.476021 | 5.612753 | 5.785687 | 5.618494 |
| 7.657454 | 7.186458 | 7.327063 | 5.506481 | 4.914731 | 7.920412 | 7.612329 | 7.596948 | 7.044043 |
| NaN | NaN | NaN | NaN | NaN | 6.912118 | 6.562853 | 6.349001 | NaN |
| 6.987564 | 7.449477 | 7.602012 | 5.866658 | 5.599116 | 6.713032 | 7.1922 | 7.156832 | NaN |
| 7.244321 | 7.558992 | 7.439811 | 6.853985 | 6.242548 | 7.366413 | 7.25227 | 6.858035 | 6.314251 |
| NaN | NaN | NaN | NaN | NaN | 5.445092 | 4.983848 | NaN | 4.430809 |
| 7.432547 | 7.025995 | 7.418339 | 6.881056 | 5.88319 | 8.003059 | 7.263202 | 6.970586 | 7.065291 |
| 6.938657 | 6.976383 | 6.993925 | 8.277903 | 8.018084 | 7.12009 | 7.029629 | 6.918991 | 8.21402 |
| 6.536377 | 6.385777 | 6.546288 | NaN | 4.043455 | 6.50109 | 6.549949 | 7.082676 | 6.54975 |
| 7.603775 | 7.536539 | 7.70161 | NaN | NaN | NaN | 6.116786 | NaN | 6.162434 |
| 5.492237 | 5.615317 | 5.921898 | 4.746612 | 5.236002 | 6.202416 | 6.359153 | 6.409718 | 5.566251 |
| 6.907559 | 6.971045 | 7.133494 | 7.579735 | 7.671417 | 6.603348 | 6.515859 | 6.610975 | 7.071975 |
| 9.056238 | 8.865893 | 8.984218 | 6.987916 | 7.615753 | 9.128122 | 9.03809 | 9.125233 | 8.235843 |

| | | | | | | | | |
|---|---|---|---|---|---|---|---|---|
| NaN | NaN | NaN | 6.226394 | 6.099905 | 4.370688 | 4.342773 | NaN | NaN |
| 9.327581 | 6.767306 | 9.629202 | 9.604418 | 8.453507 | NaN | 8.035728 | 7.708346 | 7.568454 |
| 4.796625 | 5.073855 | NaN | NaN | NaN | 4.633391 | 4.455971 | 5.468952 | 5.272205 |
| 6.657999 | 6.422006 | 6.511877 | 6.394628 | 6.585305 | 6.498518 | 6.15413 | 6.467058 | 5.344427 |
| 6.482678 | 5.112596 | 6.452317 | 6.30591 | 6.311506 | 5.887114 | 6.866845 | 7.079995 | 5.802433 |
| 7.957982 | 7.803045 | 7.683607 | 6.077592 | 6.508062 | 7.87902 | 7.833888 | 7.902431 | 8.039838 |
| 6.552905 | 6.696811 | 6.520548 | 5.812336 | 6.110947 | 6.296419 | 6.629727 | 6.627102 | 6.813293 |
| 6.243426 | NaN | NaN | 4.445856 | 6.393283 | NaN | NaN | NaN | 6.269248 |
| 5.75443 | 5.433791 | 6.12861 | 7.321903 | 7.672494 | 6.011805 | NaN | 6.443605 | 5.99314 |
| 7.027889 | 6.453866 | 6.672057 | NaN | NaN | 7.273948 | 7.058571 | 7.382885 | 6.296085 |
| NaN | 6.628504 | 6.117857 | 6.628425 | 6.72151 | 6.271488 | 6.13835 | 6.348912 | NaN |
| NaN | NaN | NaN | 7.189831 | 7.462715 | NaN | 6.985594 | NaN | NaN |
| 7.390635 | 7.107615 | 7.184951 | 8.072751 | 8.12327 | 7.689183 | 7.475858 | 7.915573 | 7.167822 |
| 7.665913 | 9.970496 | 9.311302 | NaN | 8.961214 | NaN | NaN | NaN | 8.0068 |
| 6.20314 | 5.033954 | 6.133331 | 5.384153 | 5.433226 | 5.863599 | 6.009528 | 5.980858 | 5.860754 |
| 8.083054 | 8.545171 | 8.267536 | 8.18419 | 8.151784 | 8.447752 | 7.843743 | 8.289767 | 8.085932 |
| 8.11814 | 7.870948 | 7.919641 | 7.373961 | 7.220185 | 8.41327 | 8.30145 | 8.367948 | 7.484726 |
| 6.416241 | 7.290417 | 6.638022 | 5.969779 | 6.307876 | NaN | 6.636662 | 5.985202 | 6.649141 |
| 9.047811 | 9.046408 | 8.853264 | 6.448855 | 6.334527 | 5.77297 | 6.409472 | 5.034692 | 5.170884 |
| 10.26491 | 10.2044 | 10.11731 | 8.812667 | 8.824855 | 9.735439 | 10.14091 | 9.837658 | 10.26478 |
| 7.11272 | 7.261403 | 7.222782 | 7.020588 | 6.969306 | 6.758688 | 6.640707 | 6.479535 | 7.224408 |
| 7.869838 | 8.010065 | 8.130271 | 7.00644 | 6.668811 | 8.299661 | 8.486341 | 8.267738 | 7.605447 |
| NaN | NaN | NaN | 6.000589 | 5.121367 | NaN | 6.196476 | NaN | 6.410694 |
| 10.01241 | 9.872162 | 10.0352 | 10.08897 | 10.00965 | 9.584266 | 9.233389 | 9.501875 | 9.811483 |
| 5.290908 | 6.304472 | 5.765548 | 6.53881 | 6.445189 | 5.58931 | NaN | NaN | 6.676493 |
| 7.062411 | 7.057484 | 6.438849 | 6.642862 | 6.435334 | 6.567249 | 6.838231 | 5.904179 | 6.046717 |
| NaN | NaN | NaN | NaN | NaN | NaN | 5.737159 | 5.806423 | 6.665975 |
| 10.21659 | 10.59356 | 10.37452 | 10.66819 | 10.93677 | 10.41436 | 10.32131 | 10.36896 | 10.03608 |
| 7.729294 | 7.778883 | 7.780589 | 8.857217 | 9.254334 | 8.078117 | 8.33568 | 8.0807 | 8.215403 |
| 6.367424 | 6.459744 | 6.497892 | 6.674636 | 6.49293 | 5.973033 | 6.009861 | 5.11473 | 5.187418 |
| 7.280528 | 7.542728 | 7.532309 | 7.655244 | 7.152025 | 7.146876 | 7.460135 | 7.606257 | 7.491228 |
| 8.922548 | 8.689124 | 8.731655 | 7.536096 | 7.678318 | 7.165313 | 8.021102 | 8.322683 | 8.705871 |
| 6.693899 | 6.887366 | 6.917617 | 7.145956 | 7.438909 | 6.870078 | 6.712818 | 6.931189 | 6.821278 |
| 6.784785 | 6.662612 | 6.950462 | 5.952064 | 6.982864 | 7.205538 | 7.422073 | 7.388434 | 6.788405 |
| 8.150346 | 8.162703 | 8.075646 | 9.567847 | 9.834355 | 8.412177 | 8.830493 | 8.982691 | 8.159888 |
| 7.282304 | 6.798878 | 7.327486 | 5.282267 | 5.499303 | 7.628541 | 7.721267 | 7.647676 | 6.946268 |
| 5.465611 | 4.875331 | 5.518582 | NaN | 4.402723 | 5.522242 | 5.853587 | 5.774103 | 5.225475 |
| 8.239177 | 7.812519 | 7.61224 | 7.517728 | 6.444836 | NaN | 6.355343 | NaN | 5.570577 |
| 5.791817 | 6.175912 | 6.183379 | 5.763138 | 5.557525 | 4.978307 | 5.140125 | 5.930849 | 5.481606 |
| 7.094202 | 7.438054 | 7.640049 | 8.588322 | 8.101587 | 7.013599 | 7.27012 | 7.021393 | 7.69454 |
| 6.093945 | 5.696715 | NaN | NaN | NaN | 5.909991 | 5.392445 | 5.679275 | NaN |
| 7.386172 | 7.060537 | 7.322926 | 5.268911 | 5.684793 | 7.520384 | 7.332142 | 7.300148 | 7.318941 |
| 10.49124 | 10.30183 | 10.29725 | 8.984628 | 8.759387 | 10.22763 | 10.07369 | 10.10369 | 10.00936 |
| 7.944115 | 8.189328 | 7.631381 | 8.71995 | 8.529861 | 8.112617 | 8.336027 | 8.449853 | 8.223156 |
| 7.095634 | 6.303933 | 6.42609 | NaN | 4.794505 | 6.685813 | 6.950218 | 6.805644 | 6.781444 |
| 7.019391 | 6.841309 | 7.138954 | 6.74215 | 6.881813 | 6.590522 | 6.71819 | 6.762938 | 6.162006 |
| 7.745895 | 7.541325 | 7.506547 | NaN | 6.50991 | 7.860023 | 7.668903 | 8.10276 | 7.424876 |

| | | | | | | | | |
|---|---|---|---|---|---|---|---|---|
| 7.126241 | 4.646808 | 7.653128 | 6.78055 | 5.944442 | 7.273946 | 7.01767 | 7.064143 | 7.239896 |
| 7.00783 | 6.735641 | 6.820241 | 7.061659 | 8.04251 | 6.83002 | 7.004016 | 6.403762 | 6.757367 |
| NaN | NaN | NaN | 3.879052 | 4.393241 | 4.788871 | 5.204414 | 4.78933 | 5.117771 |
| 8.096591 | 8.576545 | 9.106045 | 8.588047 | 9.218473 | 7.571253 | 8.178469 | 8.411099 | 7.777728 |
| 6.303764 | 6.223624 | 5.71022 | 6.913964 | 6.748833 | 5.952276 | 6.207737 | 7.103088 | 6.83428 |
| 7.215811 | 7.173572 | 7.378111 | 6.572564 | 6.757668 | 7.680614 | 7.752151 | 7.377725 | 7.670554 |
| 7.431322 | 8.127077 | 8.488894 | 5.853272 | 6.268968 | 7.410417 | 7.274577 | 7.411666 | 7.083293 |
| 7.006011 | 7.334427 | 6.954603 | 6.627081 | NaN | 7.596605 | 7.520608 | 7.664904 | 6.850643 |
| 7.06694 | 7.339119 | 7.508222 | 6.291928 | 6.220848 | 7.518165 | 7.886938 | 7.780688 | 7.245264 |
| 7.2893 | 6.959235 | 7.091204 | 5.512151 | 5.466999 | 6.551443 | 6.735442 | 6.688738 | 6.949121 |
| 6.937632 | 7.273119 | 6.974768 | 7.263018 | 7.273504 | 6.87196 | 6.853435 | 6.80799 | 6.96003 |
| 4.473589 | 4.139261 | 4.480412 | NaN | 4.761071 | 4.901446 | NaN | 4.735614 | 4.274197 |
| 6.44382 | NaN | 5.556109 | NaN | NaN | NaN | NaN | NaN | 7.174419 |
| 7.925994 | 7.245358 | 7.741795 | NaN | NaN | NaN | NaN | NaN | 5.662162 |
| 6.285643 | 5.923371 | 6.333539 | 4.76133 | 4.587054 | 8.120319 | 7.801251 | 8.045056 | 6.227983 |
| 7.280619 | 7.380133 | 7.360134 | 6.263554 | 6.760049 | 7.66119 | 7.9902 | 7.52643 | 6.968266 |
| 7.233607 | 7.297011 | 6.933653 | 7.441239 | 7.483531 | 6.929147 | 7.232105 | 6.560918 | 7.285326 |
| 7.361484 | 7.518269 | 7.44363 | 6.550931 | 6.784419 | 6.679872 | 7.322813 | 7.585548 | 6.802302 |
| 8.118804 | 7.752932 | 7.830163 | 6.537537 | 6.816747 | 7.573853 | 7.875075 | 7.424309 | 7.289282 |
| 8.160196 | 8.270631 | 8.252517 | 6.759348 | 7.147707 | 7.584354 | 8.670318 | 8.673427 | 7.399784 |
| 8.221276 | 8.583851 | 8.520101 | 7.691363 | 8.203576 | 8.65259 | 8.81574 | 8.627399 | 8.404128 |
| 6.434605 | 6.149063 | 6.385975 | 7.222197 | 7.517301 | 6.728393 | 6.656151 | 6.568421 | 6.551785 |
| 6.628761 | 5.260104 | NaN | 7.027408 | 6.974419 | 7.028755 | NaN | 6.666107 | 5.187887 |
| 5.205547 | 4.862132 | 4.314728 | NaN | NaN | 6.805979 | 6.840145 | 6.853666 | 6.196101 |
| 8.779604 | 8.622819 | 8.669234 | 9.727048 | 9.704003 | 8.968633 | 8.815435 | 8.791059 | 8.901804 |
| 5.686556 | 6.193337 | 6.060355 | 5.261854 | 5.515008 | 6.153476 | 6.446241 | 6.685084 | 6.159481 |
| 6.264323 | 6.863379 | 6.373379 | 4.472504 | 4.945009 | 6.870116 | 6.643362 | 6.441765 | 7.170719 |
| 5.262445 | 5.510193 | 4.897174 | 3.797792 | 4.618093 | 6.591024 | 5.620403 | 6.274969 | 5.790624 |
| 7.676023 | 7.641839 | 7.972422 | 7.356802 | 6.794246 | 7.451311 | 7.457427 | 7.491868 | 6.91166 |
| 4.945691 | 3.464323 | 6.132517 | 4.506027 | 4.68309 | 5.141346 | 4.966235 | 5.252949 | 4.905598 |
| 7.505175 | 7.053535 | 7.436865 | 6.45057 | 6.151288 | 6.322029 | 6.5294 | 6.450877 | 7.140814 |
| 7.359332 | 6.960371 | 7.051609 | 8.278025 | 8.40495 | 6.972548 | 6.891386 | 6.936527 | 7.501454 |
| 7.213185 | 7.276058 | 7.245529 | 6.763602 | 6.9461 | 7.066542 | 7.305433 | 7.336831 | 7.447992 |
| 7.644786 | 7.688336 | 7.984803 | 7.478762 | 7.311562 | 7.649789 | 7.68766 | 7.263864 | 8.133265 |
| 6.655543 | 6.229007 | 6.352738 | 5.144176 | 5.520668 | 6.341819 | 6.233616 | 6.161751 | 6.181056 |
| 5.94458 | 5.441423 | 5.579374 | 4.369054 | 4.472399 | 6.399127 | 6.272897 | 6.337085 | 6.01629 |
| 8.272936 | 7.647913 | 7.417001 | 7.966723 | 8.099007 | 8.759792 | 8.891479 | 8.968209 | 7.783669 |
| 6.942083 | 6.758039 | 6.747151 | 6.214817 | 5.823963 | 6.394069 | 6.539269 | 6.527058 | 6.827178 |
| 2.76511 | NaN | 2.469456 | NaN | NaN | NaN | NaN | NaN | 3.797875 |
| 9.367762 | 9.343321 | 9.229921 | 9.859894 | 9.757855 | 8.907401 | 8.919066 | 8.994831 | 8.956923 |
| 7.622874 | 6.8763 | 7.592055 | 7.3719 | 6.982704 | 7.636752 | 7.51296 | 7.332338 | 8.108472 |
| 6.562945 | 7.003216 | 6.499128 | 7.384638 | 6.731204 | 6.1351 | 6.34649 | 5.967573 | 6.840448 |
| 6.546329 | 4.563904 | 6.409817 | NaN | NaN | 6.259691 | 6.53082 | 6.550078 | 6.73851 |
| 6.687182 | 7.266062 | 7.131455 | 4.919866 | 6.725851 | 7.073598 | 7.080636 | 7.200437 | 7.19045 |
| 7.372951 | 7.371403 | 7.441639 | 6.773707 | 6.685628 | 6.620071 | 6.381769 | 6.524357 | 6.914111 |
| 6.581833 | 6.400188 | 6.579895 | 6.309703 | 5.896467 | 7.393594 | 7.247534 | 7.483936 | 7.138697 |
| 7.45649 | 7.683698 | 7.329043 | 7.743542 | 7.49794 | 6.738454 | 6.800996 | 6.746582 | 6.915131 |

| | | | | | | | | |
|---|---|---|---|---|---|---|---|---|
| 7.22178 | 7.240575 | 7.41123 | 7.496018 | 7.853352 | 7.108151 | 7.243497 | 7.089666 | 7.045051 |
| 5.375581 | 4.944524 | 5.458901 | 5.405443 | 5.358231 | 5.001931 | 4.688604 | 4.987479 | 5.563193 |
| 5.649337 | 5.495025 | 5.767578 | 5.353652 | 4.134727 | 6.335656 | 6.397414 | 6.192215 | 4.728377 |
| 7.809163 | 7.609218 | 7.280363 | 9.293236 | 9.139167 | 6.955643 | 7.258437 | 7.647626 | 7.946546 |
| 7.411281 | 7.183999 | 7.157381 | 6.220724 | 7.193745 | 7.973934 | 7.507749 | 7.52707 | 6.861099 |
| 10.395 | 10.2766 | 10.19657 | 10.82413 | 10.22812 | 10.54435 | 10.36933 | 10.7348 | 10.23461 |
| 7.068839 | 7.371494 | 7.054986 | 7.838254 | 7.536915 | 6.929635 | 6.618066 | 6.552357 | 5.74912 |
| 9.624922 | 9.497206 | 9.305517 | 9.451826 | 9.310343 | 9.439669 | 9.304728 | 8.89795 | 10.09733 |
| 7.973228 | 7.160859 | 7.532128 | 7.526939 | 6.202413 | 8.234057 | 7.99505 | 8.161044 | 7.679259 |
| 8.474755 | 8.821645 | 8.53792 | 8.270597 | 8.456203 | 8.75304 | 8.812728 | 8.808006 | 8.297672 |
| 7.4658 | 7.384551 | 7.700578 | 7.73603 | 7.845326 | 7.285401 | 7.641468 | 7.65869 | 8.025685 |
| 8.973053 | 9.070725 | 9.012496 | 7.99786 | 8.017298 | 9.347868 | 9.455616 | 9.353077 | 9.046701 |
| 9.478516 | 9.484536 | 9.664372 | 5.628627 | 5.713459 | NaN | NaN | NaN | 5.185289 |
| 8.24019 | 8.102621 | 8.312989 | 7.520144 | 7.271318 | 7.737793 | 7.650969 | 7.759347 | 7.422523 |
| 9.570541 | 9.383204 | 9.463469 | 10.9828 | 10.74068 | 9.581537 | 9.445712 | 9.378222 | 9.603045 |
| 5.880357 | 5.930345 | NaN | NaN | 4.7098 | 6.024123 | NaN | 5.982022 | 6.337793 |
| 6.19033 | 3.889984 | 6.377036 | 5.761014 | 5.930328 | 6.287943 | 6.5393 | 6.074378 | 5.080777 |
| 6.188494 | 5.776725 | 6.036616 | 6.44242 | 6.219772 | 5.332553 | 5.030972 | 5.325914 | 5.336953 |
| 9.79297 | 10.04614 | 10.03798 | 12.20533 | 12.61223 | 10.25029 | 10.06561 | 10.13749 | 9.823905 |
| 4.316873 | 2.443722 | 2.324616 | NaN | NaN | 2.770473 | 3.82385 | 3.270558 | 3.83906 |
| 8.110666 | 8.238484 | 7.670963 | 7.995087 | 7.972064 | 7.538298 | 7.335694 | 7.294669 | 7.217314 |
| 7.142921 | 7.296647 | 7.432654 | 6.355155 | 6.684258 | 7.437589 | 7.215716 | 7.447356 | 7.376964 |
| 12.25666 | 12.30462 | 12.34583 | 11.38993 | 11.1562 | 12.59113 | 12.71836 | 12.51752 | 11.88785 |
| 4.729568 | 4.727687 | NaN | 5.881817 | 6.089269 | 5.341167 | 5.27338 | 4.194334 | 4.849009 |
| 7.827179 | 8.201186 | 7.947986 | 7.193434 | 7.03416 | 7.513109 | 7.061386 | 7.212927 | 7.288559 |
| 7.057692 | 6.57781 | 6.751545 | 5.561878 | 6.056271 | 7.250306 | 7.220109 | 7.123492 | 6.875084 |
| 8.527542 | 7.190086 | 7.315995 | 6.589871 | 6.86402 | 6.765287 | NaN | 7.133637 | 7.292984 |
| 7.833703 | 7.855863 | 7.839875 | 6.287258 | 6.296786 | 7.771487 | 7.761602 | 7.610806 | 7.523674 |
| NaN | NaN | NaN | 4.08015 | 5.105092 | 6.310142 | 5.079754 | 5.897665 | 6.607944 |
| 6.031742 | 6.22378 | 6.110029 | 5.348314 | 5.775162 | 6.138623 | 6.522979 | 6.663917 | 6.022034 |
| 7.460008 | 7.380632 | 7.119219 | 9.363652 | 9.697609 | 7.247689 | 7.609282 | 7.618483 | 6.245914 |
| 6.766943 | 6.746655 | 6.346167 | 4.902295 | 5.41944 | 6.805541 | 7.135419 | 6.518075 | 6.961343 |
| 8.857719 | 8.975471 | 8.650003 | 10.87989 | 10.64879 | 9.398258 | 9.08122 | 9.156521 | 8.7968 |
| 7.043388 | 8.342548 | 7.104758 | 11.41704 | 11.34683 | 6.526773 | 7.107339 | 6.54835 | 8.282222 |
| NaN | NaN | NaN | 6.495584 | NaN | NaN | NaN | NaN | 7.267962 |
| 8.019031 | 7.98074 | 7.923866 | 9.170935 | 8.704288 | 7.881469 | 7.950517 | 7.817369 | 7.617971 |
| 6.469145 | 6.306511 | 6.619261 | 6.197563 | 6.192357 | 5.9485 | 5.829728 | 6.069612 | 6.322436 |
| 5.279315 | 4.310064 | 5.116871 | 5.1798 | 3.622619 | 6.075246 | 5.067385 | 6.013531 | 4.643767 |
| 9.595695 | 9.546134 | 9.536821 | 8.456219 | 8.41207 | 9.450153 | 9.33943 | 9.261259 | 9.881357 |
| 7.504897 | 7.440539 | 7.792302 | 7.978485 | 7.86811 | 7.143105 | 7.133787 | 7.242154 | 6.244765 |
| 7.956506 | 8.169852 | 7.639079 | 7.338362 | 6.139825 | 7.32653 | 7.365417 | 7.45373 | 7.738158 |
| 8.584953 | 8.387087 | 8.526338 | 8.235143 | 8.346774 | 7.793594 | 7.579771 | 7.580742 | 7.200241 |
| 12.07527 | 12.09387 | 12.18257 | 13.99747 | 13.96427 | 12.68209 | 12.52036 | 12.69779 | 11.92343 |
| 9.859281 | 9.763711 | 9.54213 | 10.62376 | 10.74227 | 10.09174 | 10.22467 | 10.07843 | 9.891731 |
| 6.551992 | 6.746356 | 6.45327 | 6.509831 | 6.619351 | 6.45338 | 6.411743 | 5.911622 | 6.269483 |
| NaN | 4.762433 | 5.85441 | 6.822197 | 5.861322 | 5.115644 | NaN | 6.659464 | 6.257456 |
| 6.940914 | 5.946439 | 6.845614 | 5.909752 | 6.305257 | 6.716657 | 7.183662 | 6.326465 | 6.572266 |

| | | | | | | | | |
|---|---|---|---|---|---|---|---|---|
| 11.3859 | 11.36808 | 11.21909 | 11.5684 | 11.05494 | 9.967479 | 9.738269 | 9.819816 | 9.489094 |
| 7.337316 | 7.037418 | 7.639109 | 6.049067 | 6.902371 | 6.39784 | 6.12489 | 6.613735 | 7.173209 |
| 5.387619 | 5.643739 | 6.202176 | 6.78999 | 6.494643 | 5.366146 | 5.383301 | 5.08125 | 5.344843 |
| 7.316786 | 7.043797 | 6.999593 | 7.817024 | 7.84846 | 6.842555 | 7.176952 | 6.627011 | 6.835822 |
| 7.694 | 8.057194 | 7.745393 | 7.950785 | 8.417661 | 6.767846 | 7.262592 | 7.73507 | 8.217041 |
| 7.553264 | 5.492826 | 7.922336 | 6.671006 | NaN | 6.904363 | 7.276258 | 6.981592 | 6.828098 |
| 9.614416 | 9.565593 | 9.643501 | 10.10664 | 9.753758 | 9.275973 | 9.317358 | 9.251049 | 9.165079 |
| NaN | 6.572427 | 5.63472 | NaN | 5.080504 | 6.3748 | NaN | NaN | 6.966848 |
| 6.124358 | 6.261662 | 6.353709 | 6.612789 | 7.152447 | 7.385332 | 6.196085 | 6.272316 | 5.264088 |
| 9.055942 | 8.891229 | 8.955972 | 7.976545 | 8.148895 | 9.32819 | 9.290942 | 9.485308 | 8.171461 |
| 7.721854 | 7.709406 | 7.725768 | 6.067685 | 6.239394 | 8.040177 | 7.815094 | 7.705606 | 7.673795 |
| 8.742079 | 8.457698 | 8.544034 | 9.871936 | 9.789448 | 8.42726 | 8.435719 | 8.439915 | 7.820271 |
| 9.759814 | 9.455854 | 9.483321 | 11.20741 | 10.86575 | 9.731755 | 9.904095 | 9.499206 | 9.835851 |
| 9.845547 | 10.01159 | 10.11558 | NaN | 8.003129 | 10.14666 | 10.03622 | 9.785208 | 9.373528 |
| 9.660817 | 9.480804 | 9.369362 | 9.069879 | 9.081041 | 9.846613 | 9.624858 | 9.66938 | 8.986724 |
| 7.871768 | 8.057954 | 7.6339 | 7.159269 | 6.897346 | 7.364798 | 7.593626 | 7.937425 | 6.94596 |
| 8.1218 | 8.385691 | 7.395193 | 9.795668 | 10.06263 | 7.044636 | 7.232079 | 6.260707 | 7.313358 |
| 6.355684 | 6.132005 | 6.14968 | NaN | 5.609015 | 6.87477 | 7.068286 | 6.892026 | 6.176881 |
| 7.767619 | 7.420088 | 7.763402 | 7.927361 | 8.172937 | 7.524527 | NaN | 8.416907 | 7.425216 |
| 9.622498 | 7.617306 | 7.598804 | 6.252118 | 5.657464 | 6.959129 | 6.412161 | 6.566077 | 6.220998 |
| 6.372051 | 7.374134 | 6.953881 | 6.148793 | 6.249291 | 6.713222 | 6.548492 | 7.209318 | 6.205528 |
| 10.83904 | 10.87548 | 10.95085 | 9.927869 | 9.822163 | 11.10393 | 11.14759 | 11.17826 | 10.62345 |
| 8.419209 | 7.048211 | 8.292718 | 6.090168 | 6.500819 | 7.200231 | 6.841962 | 7.923233 | 7.437859 |
| 7.576248 | 6.977337 | 7.482448 | 8.396133 | 8.009341 | 7.509431 | 7.683359 | 7.420281 | 7.151226 |
| 8.654407 | 8.911963 | 8.854318 | 8.734734 | 8.939849 | 8.310789 | 8.159584 | 8.390904 | 7.893427 |
| 10.18563 | 9.969995 | 10.12385 | 8.505459 | 8.582996 | 10.20428 | 10.59533 | 10.52048 | 9.77065 |
| 11.0232 | 10.82934 | 10.95023 | 9.834772 | 9.760284 | 10.89765 | 11.06338 | 10.96042 | 11.001 |
| 8.168203 | 8.000274 | 7.753388 | 8.923321 | 8.668172 | 7.261965 | 7.379222 | 6.988162 | 7.847943 |
| 6.823446 | NaN | NaN | NaN | 4.936706 | NaN | NaN | NaN | 6.424543 |
| 8.024701 | NaN | 6.99935 | 8.417872 | 8.231705 | 7.815447 | 7.191983 | 7.725049 | 8.268689 |
| 8.125444 | 7.882482 | 7.900514 | 5.775426 | 6.211231 | 7.353076 | 7.178373 | 6.971403 | 7.945647 |
| 7.520602 | 7.090402 | 7.57362 | 7.841215 | 7.396048 | 7.317973 | 7.189753 | 7.19383 | 6.773754 |
| 6.71707 | 5.944147 | 6.492902 | 5.035227 | 5.595392 | 6.010381 | 6.191957 | 5.998926 | 6.660109 |
| 9.816796 | 9.983708 | 9.745419 | 8.585613 | 8.165862 | 8.647485 | 8.357306 | 9.358571 | 8.353973 |
| 7.275447 | 7.393526 | 7.184433 | 7.693795 | 7.986181 | 7.782581 | 7.617774 | 7.876223 | NaN |
| NaN | 2.930158 | 2.966115 | 5.424934 | 5.361801 | 5.110112 | 5.092985 | 5.169481 | 6.79019 |
| 8.866379 | 8.954067 | 9.012054 | 9.433083 | 9.312257 | 8.058664 | 8.019604 | 8.233356 | 8.176272 |
| 9.187157 | 9.29041 | 8.955884 | 10.30989 | 10.27674 | 9.382502 | 9.384708 | 9.43594 | 8.43018 |
| 8.543487 | 8.606186 | 8.517952 | 9.371446 | 9.562592 | 7.663872 | 7.538431 | 7.578328 | 7.12898 |
| NaN | NaN | NaN | NaN | NaN | NaN | NaN | NaN | 6.171664 |
| 8.066042 | 7.866748 | 7.892343 | 6.21553 | 5.638246 | 6.905344 | 6.610893 | 6.763984 | 7.000474 |
| 7.178943 | 7.056027 | 7.238264 | 5.92691 | 5.892951 | 6.57482 | 6.815198 | 6.812055 | 6.592819 |
| 7.869535 | 10.32411 | NaN | 10.32469 | 10.3934 | 7.559308 | 7.470325 | 7.658743 | 7.915985 |
| 7.764639 | 7.7458 | 7.743382 | 7.529753 | 7.711462 | 7.62805 | 7.597388 | 7.634048 | 7.782566 |
| 10.75244 | 10.60235 | 10.591 | 9.920608 | 9.868102 | 10.82128 | 10.90372 | 10.93815 | 9.821207 |
| 10.61754 | 10.61362 | 10.75851 | 11.10025 | 11.14954 | 10.76195 | 10.7704 | 10.63224 | 9.89575 |
| 6.921693 | 7.123072 | 6.911376 | NaN | 4.115482 | 5.950651 | 6.842584 | 6.453929 | 6.860384 |

| | | | | | | | | |
|---|---|---|---|---|---|---|---|---|
| 9.64613 | 9.695928 | 9.904197 | 10.81129 | 11.10852 | 9.472267 | 9.339202 | 9.518606 | 9.276485 |
| 6.667994 | 7.060504 | 7.050887 | 6.295162 | 7.230523 | NaN | 6.722087 | 6.360618 | 6.493353 |
| 6.743072 | 6.201998 | 6.752452 | 6.468046 | 6.597612 | NaN | NaN | NaN | 6.26094 |
| 5.549724 | 5.107793 | 5.079957 | 4.952763 | 4.987079 | 5.538442 | NaN | NaN | 5.33234 |
| 9.284111 | 9.216095 | 9.066592 | 9.117254 | 8.943501 | 8.651587 | 8.572935 | 8.705499 | 8.954499 |
| 8.847728 | 8.931738 | 8.903248 | 7.287328 | 6.75912 | 6.84205 | 6.500406 | 6.950519 | 6.04231 |
| 7.646456 | 7.642767 | 7.739265 | 6.374319 | 6.52657 | 7.578012 | 7.401086 | 7.421234 | 6.520154 |
| NaN | NaN | NaN | 6.335181 | 6.300868 | NaN | NaN | NaN | NaN |
| 5.758222 | 5.88801 | 5.873816 | NaN | NaN | 5.532965 | 5.312452 | 5.596475 | 5.302252 |
| 6.663249 | 6.509719 | 6.321555 | 8.664951 | 8.53585 | 6.587352 | 6.190979 | 6.484821 | 5.871325 |
| 6.712418 | 6.520628 | 6.483245 | 5.963099 | 6.14644 | 6.772175 | 6.864858 | 6.785981 | 6.743766 |
| 7.999938 | 7.995089 | 7.515989 | 6.545626 | 6.711726 | 8.427119 | 7.809707 | 7.728071 | 7.21388 |
| 7.173321 | 6.83676 | 7.094166 | 5.304376 | 6.251488 | 6.591273 | 6.894069 | 6.732935 | 6.872514 |
| 6.82411 | 6.250167 | 7.185716 | 6.900557 | 6.945801 | 7.213179 | 7.022864 | 7.070043 | 6.584467 |
| 7.854254 | 7.676301 | 7.38909 | 7.42658 | 7.468246 | 7.70637 | 7.650462 | 7.500647 | 7.947811 |
| 8.592566 | 8.359282 | 8.122717 | NaN | 6.77934 | 8.49706 | 8.055281 | 8.083857 | 7.36587 |
| 6.492232 | 6.238382 | 6.479625 | 7.192801 | 7.171588 | 6.807637 | 6.036077 | 6.341812 | 5.771362 |
| 5.123194 | 4.969295 | 4.743694 | 5.245334 | 6.070253 | 4.581431 | 5.297105 | 4.597143 | 6.784423 |
| 6.850002 | 7.494086 | 7.78197 | 7.207299 | 6.421788 | 6.29117 | 6.304539 | 6.031946 | 6.719014 |
| 7.891827 | 7.877176 | 7.966298 | 6.996135 | 6.804855 | 7.485135 | 7.485559 | 7.550207 | 7.61094 |
| 6.732488 | 5.157699 | 5.969752 | 5.355443 | 5.807283 | 6.173833 | 7.081269 | 6.714082 | 6.119668 |
| 6.960229 | 6.417451 | 6.562907 | 7.012609 | 7.099326 | 5.223205 | 6.035465 | 7.102961 | 6.949462 |
| 7.443081 | 7.29091 | 7.39096 | 7.05807 | 7.004208 | 7.622812 | 7.089213 | 7.135256 | 6.063877 |
| 6.440384 | 6.574773 | 6.560743 | 6.62256 | 7.115439 | 4.94984 | NaN | NaN | 6.132328 |
| 6.204927 | 6.622015 | 6.211474 | NaN | 4.6567 | 5.805999 | 5.957833 | 5.844079 | 5.554696 |
| NaN | NaN | NaN | NaN | NaN | 5.363146 | 5.217828 | 4.695353 | 5.14217 |
| NaN | 6.227998 | 5.876609 | NaN | NaN | 6.445518 | 6.60595 | 6.986221 | 5.719357 |
| 6.670894 | 6.598053 | 7.011279 | 5.308013 | 6.629247 | 6.826472 | 6.841451 | 6.803591 | 5.743244 |
| 7.765552 | 6.749531 | 7.875528 | 6.590807 | 6.873394 | 8.004582 | 8.088624 | 8.205812 | 7.539639 |
| 6.789235 | 7.084799 | 7.183112 | 6.556734 | 7.438731 | 7.562642 | 7.560647 | 6.817424 | 6.429218 |
| 8.165425 | 8.032392 | 8.016816 | 8.206952 | 8.027512 | 7.693125 | 7.747185 | 7.457285 | 7.896618 |
| 6.571111 | 7.149946 | 7.689565 | 7.882795 | 7.746988 | 8.639958 | 8.541225 | 8.584161 | 7.141399 |
| 8.50088 | 8.621243 | 8.697385 | 8.449202 | 8.046884 | 8.537163 | 8.292972 | 8.106581 | 8.799933 |
| 7.276311 | 7.088338 | 7.767781 | 7.497365 | 6.888875 | 6.1194 | 6.534778 | 6.344306 | 6.907571 |
| 7.504305 | 7.445939 | 7.287465 | 8.498082 | 8.39987 | 7.632194 | 7.539388 | 7.567376 | 7.25924 |
| 7.566865 | 7.228248 | 7.886333 | 7.786381 | 7.595106 | 7.838935 | 7.664816 | 7.644531 | 7.106136 |
| 6.806023 | 7.174358 | 6.717102 | 6.066711 | 5.978347 | 6.271319 | 6.642436 | 6.527267 | 5.20716 |
| 8.319934 | 8.450933 | 8.448048 | 8.612396 | 8.77489 | 9.397038 | 9.266154 | 9.175097 | 8.427009 |
| 8.652836 | 8.487998 | 8.626379 | 8.30352 | 8.28595 | 8.658025 | 8.585899 | 8.661554 | 8.674759 |
| 7.811182 | 6.381104 | 7.615645 | 7.034664 | 7.715725 | 7.081817 | 7.381917 | 6.489532 | 5.886301 |
| 8.71399 | 8.819279 | 8.552139 | 9.129983 | 9.196198 | 7.43402 | 7.362945 | 7.4826 | 8.641357 |
| 6.706343 | 6.726646 | 6.25615 | 6.185054 | 6.373372 | 6.122295 | 5.675797 | 5.756586 | 5.629733 |
| 7.62367 | 6.824474 | 7.490808 | 6.42911 | 6.540721 | 7.409169 | 7.762567 | 6.979987 | 7.5179 |
| 5.816313 | 6.1699 | 5.537817 | 10.41066 | 10.20381 | 5.166131 | 5.000573 | 5.469999 | 5.707263 |
| 9.604811 | 9.594464 | 9.614864 | 9.30211 | 9.302094 | 8.351014 | 8.19011 | 8.218218 | 8.10534 |
| 7.462694 | 7.394034 | 7.405908 | 5.594207 | 5.731167 | 7.61741 | 7.443229 | 7.368266 | 6.565 |
| 7.176688 | 7.121633 | 7.119004 | 6.6276 | 7.513937 | 7.454391 | 7.475834 | 7.311359 | 6.549027 |

| | | | | | | | | |
|---|---|---|---|---|---|---|---|---|
| 8.557202 | 8.634908 | 8.560905 | 8.226563 | 9.461289 | 7.394756 | 7.153689 | 7.324836 | 6.827021 |
| 8.317032 | 8.35998 | 8.227832 | 10.10044 | 9.724086 | 7.330971 | 6.743254 | 7.505326 | 8.332591 |
| NaN | NaN | NaN | 5.796654 | 8.201333 | NaN | 6.146156 | 6.202458 | 6.313375 |
| 7.4942 | 7.489433 | 7.528871 | 7.441891 | 7.312276 | 5.360073 | 6.015699 | 6.39587 | 6.19373 |
| 6.35041 | 6.689784 | NaN | 6.540007 | 6.680581 | 6.250747 | 6.186359 | 6.12366 | 6.369561 |
| 7.019729 | 6.752487 | 7.108884 | 5.64797 | 6.54242 | 7.222136 | 7.363702 | 7.411835 | 6.732439 |
| 6.753457 | 6.721824 | 7.002701 | 6.996281 | 6.805305 | 6.353314 | 6.321805 | 6.252982 | 7.022882 |
| 10.71255 | 10.67174 | 10.67822 | 11.45755 | 11.37914 | 10.7995 | 10.87966 | 10.6083 | 10.10712 |
| 6.574965 | 6.865471 | 6.304772 | NaN | 5.900781 | 7.011986 | 6.766069 | 6.814357 | 6.072762 |
| 6.518163 | 5.42652 | 6.472356 | 7.229056 | 6.862889 | 6.182254 | 6.035695 | NaN | 7.708436 |
| 9.52747 | 9.849268 | 9.677664 | 8.793061 | 8.731663 | 9.43198 | 9.442805 | 9.332829 | 9.429859 |
| 6.581764 | 6.328859 | 5.434524 | 6.41355 | 6.306422 | 6.59205 | 6.505231 | NaN | 6.187402 |
| 5.733834 | 4.101233 | 5.785284 | 5.103985 | 6.755885 | 5.567787 | 6.070945 | 5.905949 | 5.429673 |
| 8.656731 | 8.57691 | 8.718617 | 8.26912 | 8.342713 | 7.893284 | 8.028064 | 7.780711 | 8.744202 |
| 7.372771 | 7.656219 | 6.568291 | 7.782018 | 7.603641 | 6.524976 | 6.29164 | 6.487159 | 7.105056 |
| 6.444713 | 4.902028 | 7.039733 | 7.067435 | 5.751083 | 8.353728 | 8.510564 | 8.590431 | 6.361838 |
| 7.063933 | 6.242565 | 6.043501 | 6.196043 | 6.350526 | 4.219452 | 6.602565 | 6.010959 | 5.810883 |
| 7.748382 | 6.034231 | 5.061367 | 7.632547 | 7.789005 | NaN | NaN | 7.667852 | NaN |
| 6.737383 | 6.57966 | 6.92426 | 6.292828 | 6.356123 | 6.765064 | 6.761848 | 6.981803 | 7.188607 |
| 6.302865 | 4.912558 | 3.229011 | NaN | 4.771016 | 5.624215 | 5.566786 | 5.474138 | 5.644044 |
| 8.605785 | 8.774927 | 8.772804 | 7.868131 | 7.620046 | 8.060247 | 8.129208 | 8.17318 | 9.091423 |
| 7.151248 | 7.327891 | 7.032789 | 7.422231 | 7.5385 | 7.277889 | 7.286304 | 7.364319 | 6.464147 |
| 5.660787 | 5.70436 | 6.060345 | NaN | 5.724179 | 5.801968 | 6.181892 | 6.192485 | 6.200434 |
| 4.318851 | 5.015916 | NaN | NaN | NaN | 6.121322 | NaN | 5.953032 | 5.400963 |
| 8.123446 | 7.755394 | 7.903448 | 8.31272 | 8.109657 | 6.977488 | 6.446183 | 6.86518 | 7.277703 |
| NaN | 5.726737 | 5.32866 | 6.835055 | 6.881488 | NaN | NaN | NaN | NaN |
| 4.79539 | 4.795695 | 4.923279 | 5.512507 | 5.921309 | 4.07214 | 4.913967 | 4.127879 | 4.751704 |
| 8.993167 | 8.84778 | 8.83166 | 9.987773 | 9.957448 | 8.353352 | 8.403619 | 8.453226 | 9.408934 |
| 5.491369 | 4.876964 | 4.701865 | 2.995394 | NaN | 5.776712 | 5.638612 | 5.459976 | 5.590323 |
| 5.691088 | 5.400638 | 5.411798 | 6.709006 | 7.435662 | 6.1034 | 5.966465 | 6.248382 | 7.249604 |
| 6.342708 | 5.856654 | 6.341274 | NaN | 4.964994 | NaN | 5.712631 | 5.67074 | 4.676095 |
| 3.381777 | 5.816006 | NaN | 8.412867 | 8.759042 | 6.845005 | 7.195626 | 7.044854 | 8.165582 |
| 5.471806 | 5.923602 | 5.671196 | 6.21094 | 5.009609 | 5.781556 | 5.637023 | 5.840939 | 5.141884 |
| 9.249162 | 9.190819 | 9.261488 | 8.567998 | 8.608385 | 9.412454 | 9.404583 | 9.34253 | 8.830415 |
| 8.124813 | 8.207788 | 8.072238 | 7.551616 | 7.468283 | 7.229073 | 7.231587 | 7.013238 | 6.628431 |
| 6.810479 | 6.614306 | 7.100437 | 7.095768 | 7.064631 | 7.029918 | 6.296095 | 6.258364 | 7.253672 |
| 6.691319 | 6.87658 | 6.97808 | 6.964483 | 6.89876 | 7.228895 | 6.669925 | 6.916776 | 6.541466 |
| 7.571806 | 8.324052 | 6.737321 | NaN | 6.682994 | NaN | 7.34576 | 7.070556 | 6.772704 |
| 7.196841 | 7.112473 | 7.716002 | 8.9629 | 8.791535 | 5.329337 | NaN | 5.672739 | 7.132769 |
| 8.909645 | 9.077965 | 9.034077 | 9.733506 | 8.999025 | 10.08224 | 9.806382 | 9.939483 | 9.115336 |
| 6.318346 | 6.401485 | 6.581239 | NaN | 5.671931 | 6.822233 | 6.420156 | 6.527903 | 6.580583 |
| 7.033741 | 7.111267 | 7.770983 | 7.613689 | 7.704836 | 7.685834 | 7.656044 | 7.727334 | 7.476612 |
| 7.941638 | 6.492486 | 8.22294 | 7.331287 | 7.233196 | 6.286341 | 6.486957 | 6.373845 | 7.074115 |
| 5.044031 | 5.440236 | 5.350855 | 8.56573 | 8.749279 | 3.776391 | 2.234841 | 4.128645 | 5.34056 |
| 7.56071 | 6.006783 | 7.894506 | 8.308238 | 8.846981 | 8.40132 | 8.331868 | 8.168152 | 8.177444 |
| 8.484668 | 8.461724 | 8.469406 | 7.892378 | 7.965614 | 8.127887 | 7.943954 | 7.912787 | 8.483327 |
| 5.737093 | 3.933276 | 6.50477 | 5.994126 | 6.402909 | 5.48169 | 5.192875 | 5.764648 | 5.843918 |

| | | | | | | | | |
|---|---|---|---|---|---|---|---|---|
| 7.217775 | 7.362846 | 7.095468 | 6.730021 | 6.516344 | 7.237083 | 7.34919 | 7.214496 | 7.263698 |
| 8.925545 | 9.228531 | 9.068765 | 8.433884 | 8.228012 | 8.702396 | 8.965267 | 8.980527 | 9.088112 |
| 7.644323 | 7.681842 | 7.728374 | 6.6971 | 6.542095 | 7.43525 | 7.308216 | 7.508791 | 7.175008 |
| 6.581498 | 7.213686 | 6.684269 | 7.065571 | 6.803414 | 7.155021 | 6.607212 | 6.416699 | 6.151803 |
| 7.804011 | 7.859561 | 7.982307 | 6.35558 | 6.153335 | 7.765338 | 7.761425 | 7.571353 | 7.741567 |
| 7.407031 | 7.382381 | 7.507876 | 7.432927 | 7.495612 | 6.850838 | 6.859103 | 6.933283 | 6.707862 |
| 5.795 | 6.003242 | 5.704061 | 5.451228 | 5.225555 | NaN | NaN | 5.857943 | 6.195139 |
| 7.146205 | 7.045923 | 6.997559 | 5.405281 | 6.283689 | 7.273146 | 7.051538 | 7.104404 | 6.30223 |
| 7.411276 | 6.905057 | 6.840312 | 7.369601 | 6.207345 | 6.582568 | 6.498114 | 6.344181 | 6.222951 |
| 7.658521 | 7.116287 | 6.852715 | 8.368403 | NaN | 6.84772 | 5.892156 | 7.06781 | 5.767012 |
| 7.891614 | 7.306244 | 7.00846 | 6.674242 | 6.537732 | 8.845621 | 8.923137 | 9.163576 | 5.571934 |
| 9.010248 | 9.060567 | 8.985304 | 9.469312 | 9.33743 | 8.687648 | 8.922853 | 8.686888 | 8.371119 |
| NaN | 6.627504 | NaN | 6.8427 | 5.213673 | 6.225403 | 6.25789 | 6.034504 | 6.461853 |
| 7.344228 | 7.035355 | 7.165047 | 8.314169 | 8.19324 | 8.185119 | 8.373213 | 8.469323 | 7.010113 |
| 5.890098 | 7.138443 | NaN | 6.537667 | 6.867166 | 5.358572 | 5.238505 | 5.102107 | 6.618458 |
| 7.914332 | 8.117815 | 7.541738 | 8.688166 | 8.400992 | 7.375415 | 7.90322 | 7.885409 | 7.097872 |
| 6.624822 | 6.280367 | NaN | 7.016057 | 7.453239 | 8.026233 | 8.200403 | 8.459226 | 8.072987 |
| 6.937782 | 7.242106 | 7.176046 | 6.567451 | 5.744771 | 6.667049 | 6.281816 | 6.794783 | 6.445353 |
| 13.15847 | 13.10021 | 13.17659 | 12.33301 | 12.48254 | 13.43295 | 13.54659 | 13.57382 | 12.86889 |
| 7.031538 | 7.123889 | 7.35151 | 6.71904 | 7.386189 | 7.736525 | 7.518119 | 7.589046 | 7.050294 |
| 6.446332 | 7.030288 | 6.443236 | 5.376499 | 5.726672 | 7.802691 | 6.861406 | 7.147357 | 6.583735 |
| 9.250277 | 9.403429 | 9.35498 | 9.32281 | 9.396605 | 9.280628 | 8.934132 | 8.98043 | 8.793122 |
| 7.440959 | 7.943413 | 6.320902 | NaN | NaN | 7.019269 | 6.843574 | 7.176712 | 6.839984 |
| 8.755278 | 8.619755 | 9.066028 | 10.73309 | 10.85843 | 8.605766 | 9.096788 | 8.671293 | 8.735609 |
| 4.809312 | 3.243329 | 3.54163 | 3.936581 | 4.292845 | 5.202045 | 5.955466 | 5.434449 | 2.527138 |
| 7.529972 | 7.294544 | 7.494652 | 8.00857 | 7.997769 | 7.758599 | 7.592305 | 7.632122 | 3.960587 |
| 9.341091 | 7.843943 | 9.135978 | 7.698862 | 7.501582 | 8.674736 | 9.479712 | 8.828808 | 7.949063 |
| NaN | NaN | NaN | 5.527459 | 7.651962 | 5.876882 | NaN | 7.211861 | 6.160551 |
| 8.601545 | 8.494201 | 8.376942 | 7.72128 | 7.809917 | 8.2409 | 8.194169 | 8.246026 | 7.970539 |
| 6.9934 | 6.190353 | 6.95505 | 6.435282 | 6.859632 | 7.722137 | 7.578403 | 7.789371 | 6.668273 |
| 9.115857 | 9.008098 | 8.778989 | 10.05884 | 9.868762 | 8.344332 | 7.902753 | 7.576313 | 7.370114 |
| 8.135446 | 8.046926 | 8.334035 | 7.378179 | 7.37426 | 9.054916 | 9.186047 | 9.013621 | 7.755458 |
| 6.747964 | 6.551761 | 6.344049 | 7.51326 | 7.543142 | 6.897726 | 6.764173 | 6.681095 | 6.999739 |
| 7.295279 | 7.734894 | 8.418672 | 7.159868 | 6.948859 | 7.404966 | NaN | 7.044701 | 8.341551 |
| 7.806376 | 7.933298 | 7.409764 | 5.818875 | 5.555665 | 7.354781 | 7.473941 | 6.853539 | 7.204222 |
| NaN | NaN | NaN | 6.415189 | 6.222174 | NaN | NaN | NaN | 6.054075 |
| 5.841271 | 5.61622 | 6.149319 | 6.076499 | 5.970992 | 6.98715 | 6.246714 | 5.764686 | 5.427573 |
| 8.119661 | 8.158707 | 7.956239 | 7.551629 | 7.185161 | 7.32193 | 7.700886 | 7.591802 | 7.825503 |
| 5.771541 | 6.223571 | 6.284676 | 5.281607 | NaN | 5.375714 | 5.095391 | NaN | 5.238873 |
| 7.41434 | 7.353827 | 7.367518 | 7.675557 | 7.623584 | 7.394614 | 7.305016 | 7.548714 | 7.348276 |
| 5.981091 | 6.043301 | 5.654204 | 6.20235 | 5.436492 | 4.987164 | 5.084661 | 4.944901 | 5.368821 |
| 8.087713 | 7.619623 | 7.999362 | 7.008036 | 7.167398 | 8.042086 | 7.72252 | 8.116487 | 7.421682 |
| 6.852465 | 4.831707 | 6.024381 | 6.752028 | 6.300496 | 5.708496 | 5.574036 | 5.733622 | 6.278334 |
| 7.225796 | 6.85541 | 7.149659 | 5.358213 | 5.65799 | 5.72529 | 5.774272 | 5.787035 | 4.691329 |
| 12.25215 | 13.7214 | 13.84711 | 13.98104 | 11.48397 | 15.59693 | 15.75904 | 15.5744 | 11.7641 |
| 8.029129 | 6.021828 | 7.484536 | 7.566588 | 7.672117 | 6.977962 | 6.737062 | 6.572978 | 7.298403 |
| 6.252944 | 6.116039 | 5.642026 | 5.783815 | 5.642242 | 5.616662 | 5.201582 | 5.830636 | 5.105712 |

| | | | | | | | | |
|---|---|---|---|---|---|---|---|---|
| 9.019097 | 9.378322 | 9.712938 | 7.397818 | 8.686138 | 7.375333 | NaN | 7.913282 | 7.796409 |
| 9.254711 | 9.389995 | 9.154743 | 10.13243 | 10.45642 | 8.18117 | 8.633315 | 8.616564 | 8.365103 |
| 6.574244 | 7.072793 | 6.40449 | 4.522232 | 4.221583 | 6.914822 | 7.986068 | 7.980506 | 6.469 |
| 7.663917 | 6.922635 | 7.353741 | 5.354425 | 5.787912 | 8.125081 | 8.119773 | 8.349669 | 6.181109 |
| 6.148458 | 5.993607 | 6.485956 | 6.094507 | NaN | 8.020904 | 7.968503 | 8.117865 | 7.250111 |
| 6.612835 | 6.082438 | 5.990611 | 6.039153 | 6.304857 | 5.933463 | 6.537538 | 6.022107 | 6.234849 |
| 7.351494 | 6.991882 | 7.18407 | 7.456168 | 7.553054 | 6.540705 | 6.283484 | 6.635954 | 7.861763 |
| 10.31835 | 10.02611 | 10.1334 | 8.99961 | 8.763366 | 9.900219 | 10.21544 | 9.994328 | 9.435252 |
| 7.757382 | 7.79171 | 7.845911 | 8.421949 | 8.599656 | 7.936479 | 8.287365 | 8.124942 | 7.977406 |
| NaN | 4.809345 | NaN | NaN | NaN | NaN | NaN | NaN | 4.132322 |
| 8.515004 | 8.036201 | 7.410308 | 9.140633 | 8.650161 | 8.352585 | 8.236951 | 8.432894 | 8.379707 |
| NaN | 5.724042 | 3.699273 | 7.066909 | 6.627924 | 6.835797 | 6.057176 | 6.582499 | 6.099251 |
| 7.315621 | 7.610104 | 7.53029 | 6.471132 | 6.619021 | 7.25948 | 7.44947 | 7.085597 | 7.646252 |
| NaN | NaN | NaN | 5.80104 | NaN | 7.467377 | 7.720654 | NaN | 7.069263 |
| 7.864054 | 6.223099 | 7.976716 | 6.831176 | 7.114223 | 8.729306 | 7.57427 | 7.192074 | 6.573459 |
| NaN | NaN | 6.573869 | NaN | NaN | 6.073264 | 6.055565 | 6.300565 | NaN |
| 6.579749 | 6.494764 | 6.935632 | 4.927917 | NaN | 6.252074 | 6.487231 | 6.781868 | 6.269686 |
| 5.641107 | 5.647793 | 5.973261 | 4.065383 | 4.250076 | 5.569735 | 5.140308 | 5.319355 | 5.809225 |
| 6.854612 | 6.993598 | 6.868128 | 7.997044 | 7.990275 | 6.827952 | 6.822902 | 7.427027 | 7.051304 |
| 7.296457 | 8.51285 | 8.442112 | 8.637423 | 8.768383 | 6.933002 | 7.101539 | 6.713202 | 8.375841 |
| 7.198112 | 6.247509 | 6.906828 | 6.415691 | 6.072963 | 6.081816 | 6.445389 | 6.956371 | 6.682901 |
| NaN | 4.936498 | NaN | 4.270061 | 5.294796 | 5.929801 | 5.350705 | 5.879928 | 5.536708 |
| 4.451338 | NaN | NaN | 9.267281 | 9.25099 | 5.366624 | 5.402839 | 5.312988 | 3.818049 |
| 7.960598 | 8.372885 | 8.078817 | 8.738579 | 9.373283 | 8.399956 | 8.35838 | 8.243609 | 8.795453 |
| 7.012111 | 7.008033 | 7.271392 | 8.057307 | 8.149722 | 6.744539 | 6.230374 | 6.466004 | 6.947723 |
| 7.770221 | 7.728037 | 7.684487 | 8.136039 | 8.070592 | 7.375381 | 7.355335 | 7.713949 | 7.355245 |
| 6.741104 | 6.533413 | 6.630994 | 5.734624 | NaN | 6.692467 | 6.466308 | 6.685469 | 6.535652 |
| 7.797833 | 7.617991 | 7.799695 | 8.087255 | 8.194313 | 7.806464 | 7.822292 | 7.938401 | 7.858935 |
| 8.636538 | 8.950888 | 8.813333 | 8.43198 | 8.407999 | 9.631555 | 9.561303 | 9.48 | 9.605574 |
| NaN | NaN | NaN | 5.644512 | NaN | NaN | 5.107075 | 5.087336 | 5.149641 |
| 7.407712 | 7.373046 | 7.450969 | 8.308912 | 8.170437 | 6.961505 | 7.049982 | 7.246689 | 6.759192 |
| 6.626702 | 6.161942 | 6.276595 | 6.634356 | 6.623141 | 6.719977 | 6.641325 | 6.074786 | 7.046313 |
| 7.943314 | 8.050117 | 8.18614 | 8.665922 | 8.46769 | 7.328319 | 7.451249 | 7.603805 | 8.555754 |
| 9.702213 | 9.578109 | 9.62128 | 8.410656 | 8.579911 | 9.557288 | 9.490742 | 9.856396 | 8.331288 |
| 4.094977 | NaN | 5.998581 | 6.408286 | 6.268848 | 6.224267 | 6.232337 | 6.440814 | 6.542887 |
| 8.208356 | 8.142317 | 8.137144 | 8.124034 | 7.961861 | 8.31357 | 8.346701 | 8.465956 | 8.408916 |
| 5.859849 | 6.730628 | 6.430745 | NaN | 6.64272 | NaN | 5.616001 | 5.602486 | 7.730269 |
| 7.319484 | 6.937338 | 6.908844 | 6.696758 | 6.948155 | 6.366781 | 6.778201 | 7.077492 | 6.864681 |
| NaN | NaN | 3.032979 | NaN | NaN | 1.481253 | NaN | NaN | 4.373459 |
| 7.649767 | 7.813482 | 7.509492 | 7.652927 | 6.969345 | 7.49719 | 7.642961 | 7.543243 | 7.404371 |
| 6.5884 | 6.842772 | 6.817239 | 8.193089 | 8.268716 | 6.671457 | 6.506356 | 6.459045 | 6.989009 |
| 6.986627 | 7.225069 | 7.409109 | 8.6003 | 8.84094 | 6.870242 | 6.474007 | 7.128965 | 7.642137 |
| 7.949868 | 7.666655 | 7.499233 | 7.62349 | 7.231628 | 6.976202 | 7.015716 | 7.456299 | 6.902646 |
| 9.354893 | 9.358022 | 9.393603 | 10.33501 | 10.39602 | 9.377678 | 9.399095 | 9.356542 | 8.647989 |
| 6.663984 | NaN | 6.445639 | NaN | NaN | 6.386867 | 6.925336 | 6.572263 | 6.447044 |
| 5.072957 | 5.529538 | 5.763631 | NaN | 5.3836 | NaN | NaN | NaN | 5.67074 |
| 6.472658 | 6.116244 | 6.386535 | 5.178178 | 4.712603 | 5.890079 | 5.796463 | 5.716316 | 6.220058 |

| | | | | | | | | |
|---|---|---|---|---|---|---|---|---|
| 6.895937 | 7.025892 | 7.302411 | 6.130405 | 5.851466 | 7.048441 | 7.157988 | 7.134784 | 7.380178 |
| 5.387445 | 6.332476 | 5.669031 | 5.954587 | 6.793326 | 7.94087 | 7.628696 | 7.999535 | 7.221603 |
| 5.572357 | 4.644738 | 5.097703 | NaN | 3.470199 | 5.566752 | 5.600741 | 5.411835 | 6.725356 |
| 7.139395 | 6.511518 | 8.62848 | 7.623239 | 7.070384 | NaN | 7.629757 | 7.267832 | 6.908473 |
| 8.093895 | NaN | 7.895519 | 9.427647 | 8.174637 | NaN | NaN | NaN | 7.066561 |
| NaN | NaN | 7.756276 | 8.422926 | 8.514252 | 7.902663 | 7.918386 | 7.896757 | 7.394567 |
| 8.606326 | 8.613845 | 8.687399 | 7.871412 | 7.823031 | 7.461005 | 7.644233 | 7.342957 | 5.712472 |
| 7.294457 | 7.230894 | 7.318866 | NaN | NaN | 7.639063 | 7.606896 | 7.762649 | 7.864691 |
| 7.233042 | 7.400832 | 7.166125 | 5.472502 | 6.148954 | 7.3399 | 7.182236 | 7.373434 | 7.293717 |
| 8.219151 | 8.049659 | 8.118287 | 9.582647 | 9.489777 | 8.11421 | 8.020153 | 7.970319 | 7.957055 |
| 7.921668 | 7.834895 | 7.667679 | 6.811288 | 7.198574 | 8.456 | 8.235879 | 8.198781 | 7.847419 |
| 7.218674 | 6.290939 | 7.267184 | 5.931034 | 6.763452 | 7.022114 | 7.405448 | 7.34314 | 6.540144 |
| 9.273987 | 9.178796 | 9.263579 | 9.33179 | 9.328295 | 9.025911 | 8.889786 | 9.12706 | 9.083064 |
| 7.440496 | 6.305509 | 7.444995 | 5.665604 | 6.381788 | 7.799887 | 7.574265 | 7.532776 | 7.156753 |
| 8.771027 | 8.764263 | 8.766278 | 8.204765 | 8.160741 | 9.097701 | 9.105212 | 9.075047 | 9.35753 |
| 6.315413 | 6.541014 | 6.368528 | 8.067738 | 8.262935 | 6.261436 | 5.898396 | 5.58784 | 5.826664 |
| 6.866076 | 6.268585 | 6.792931 | 5.699745 | 5.440014 | 7.0404 | 6.854828 | 6.77563 | 7.07901 |
| 6.43215 | 6.232363 | 6.370957 | 5.523261 | 5.661422 | 6.278938 | 6.540014 | 6.30602 | 5.686962 |
| 6.535184 | 6.445108 | 6.49807 | 5.714661 | 5.324066 | 6.607856 | 6.202596 | 6.477774 | 6.126869 |
| 7.783949 | 7.977303 | 7.301258 | 6.939271 | 6.862846 | 6.983019 | 6.55758 | 6.887285 | 5.130144 |
| 6.080122 | 5.094263 | 5.636462 | 6.028573 | 5.783556 | 5.499294 | 6.071543 | 6.819311 | 6.128383 |
| 8.277988 | 9.678736 | 8.767366 | 8.339986 | 7.975291 | 7.918762 | 8.04879 | 8.488797 | 8.471837 |
| 6.57984 | 6.529117 | 6.020399 | 6.371507 | 5.879188 | 5.805107 | 6.401731 | 6.192022 | 6.515494 |
| 8.742269 | 8.919641 | 9.038403 | 10.07933 | 9.874925 | 7.895724 | 8.208146 | 8.12305 | 8.51195 |
| 7.495962 | 7.714941 | 7.157654 | 7.360832 | 6.876531 | 7.447569 | 7.709416 | 7.645855 | 7.688451 |
| 4.424028 | 4.251112 | 3.980769 | NaN | 4.832244 | 4.13028 | 4.490283 | 4.988585 | 4.840745 |
| 6.622578 | 6.618416 | 7.185943 | 10.00755 | 10.10767 | 6.800446 | 6.663881 | 7.107488 | 6.610296 |
| 7.778926 | 7.890229 | 7.664556 | 7.645683 | 7.698894 | 7.996776 | 7.989364 | 8.107165 | 7.514003 |
| 6.132456 | 6.002091 | 6.009548 | 5.752326 | 6.129676 | 6.657037 | 6.353506 | 6.660172 | 5.533973 |
| 7.945229 | 8.307903 | 7.866569 | 8.20927 | 8.403832 | 7.637366 | 7.756439 | 7.61798 | 7.920878 |
| 6.895478 | 7.006373 | 7.443676 | 6.05213 | 4.280285 | 6.828579 | 6.907795 | 7.336706 | 6.139178 |
| 7.417861 | 7.663504 | 7.546664 | 6.648228 | NaN | 7.785769 | 7.809024 | 7.745059 | 8.125375 |
| 7.311014 | 7.107716 | 7.092961 | 7.915522 | 7.768689 | 6.948359 | 6.945853 | 7.236989 | 7.060227 |
| 7.52626 | 7.809575 | 7.802966 | 8.065146 | 8.082789 | 8.072495 | 7.919218 | 7.503308 | 7.464634 |
| 7.260202 | 7.067901 | 7.254207 | 7.744987 | 7.791759 | 7.476777 | 7.279411 | 7.306429 | 6.457817 |
| 8.036667 | 7.928246 | 8.037216 | 9.593374 | 9.244762 | 7.297018 | 7.667484 | 7.453754 | 7.411985 |
| 6.909383 | 6.420747 | 6.785654 | 6.069334 | 6.463076 | 6.488029 | 6.379823 | 6.712798 | 6.895713 |
| 8.327552 | 8.267202 | 8.877991 | 9.55403 | 8.734081 | 7.344694 | 5.573606 | 5.607887 | 6.942491 |
| 6.668638 | 6.284059 | 6.204404 | 6.245936 | 6.442481 | 6.196095 | 5.930509 | 5.720678 | 6.128012 |
| 8.141456 | 8.086083 | 8.203959 | 7.937193 | 7.946379 | 8.812545 | 8.701406 | 8.753032 | 8.381106 |
| NaN | NaN | 4.896379 | NaN | NaN | NaN | NaN | NaN | 4.442592 |
| 6.429055 | 6.69367 | 6.424672 | 5.3144 | 6.238189 | 6.609302 | 6.680613 | 6.451295 | 6.297724 |
| 8.361732 | 7.952505 | 8.763293 | 9.641395 | 9.140072 | 6.698661 | 7.429893 | 6.389283 | 7.515408 |
| 8.286963 | 8.333834 | 8.337123 | 8.579968 | 8.40652 | 7.605539 | 7.175603 | 7.141363 | 7.800099 |
| 6.627028 | 6.837685 | 6.701926 | 6.757151 | 6.509148 | 6.580588 | 7.049764 | 7.108641 | 6.791997 |
| 7.270146 | 7.143987 | 7.258137 | 6.759991 | 6.396603 | 7.465141 | 7.334551 | 7.375242 | 7.559695 |
| 7.378293 | 7.259295 | 7.188174 | 7.70115 | 7.675596 | 7.445455 | 7.133311 | 7.436482 | 6.891793 |

-Bruker Confidential-

| | | | | | | | | |
|---|---|---|---|---|---|---|---|---|
| NaN | NaN | 6.079903 | 4.239334 | 4.808419 | NaN | 5.274495 | NaN | 5.828344 |
| 7.78252 | 7.675832 | 7.674295 | NaN | 5.636369 | 6.86145 | 6.976716 | 6.868196 | 6.807275 |
| 4.108963 | 3.948961 | NaN | 2.966742 | 2.070233 | 4.881693 | 5.0902 | 5.214683 | NaN |
| 7.414844 | 7.261236 | 7.700379 | 7.230555 | 7.892624 | 6.78564 | 7.462016 | 7.351295 | 7.406478 |
| 5.661376 | 5.710307 | 5.71732 | NaN | 4.280111 | 5.107802 | 5.63098 | 5.472973 | 5.538897 |
| 7.826478 | 7.670263 | 7.721594 | 8.506274 | 8.46775 | 8.215755 | 8.104066 | 8.001629 | 7.673354 |
| 7.891179 | 6.805178 | 5.633606 | 6.842107 | NaN | 7.442852 | 7.38173 | 6.883642 | 7.710098 |
| 7.827792 | 7.909374 | 8.106669 | 6.491058 | 7.185306 | 7.288584 | 7.367115 | 7.157067 | 6.627975 |
| 8.248025 | 8.450768 | 8.196736 | 6.668732 | 6.584848 | 7.846945 | 7.886442 | 7.742688 | 7.979259 |
| 8.746209 | 8.074218 | 8.966652 | 7.188052 | 7.025988 | 6.733493 | 6.749642 | 6.313568 | 6.810017 |
| 7.996661 | 8.083938 | 7.859079 | 6.452329 | 6.254977 | 8.065999 | 7.712113 | 7.909182 | 7.493632 |
| 6.676013 | 6.682698 | 6.704002 | 6.84352 | 6.361807 | 6.703464 | 6.705998 | 6.877823 | 6.22503 |
| 5.610851 | 4.410204 | 4.462442 | NaN | 5.321128 | 6.375813 | 6.329376 | 6.220973 | 5.776727 |
| 5.279811 | NaN | 4.748665 | NaN | NaN | 4.764148 | NaN | NaN | 5.53325 |
| 7.772587 | 7.672709 | 7.613671 | 7.852152 | 7.798344 | 7.443421 | 7.174752 | 7.580129 | 6.893786 |
| 6.871297 | 7.009462 | 6.786659 | 7.546865 | 7.630589 | 6.368725 | 6.122078 | 6.199363 | 7.228904 |
| 7.972849 | 8.145653 | 7.709599 | 6.415453 | 6.408678 | 7.334947 | 7.726781 | 7.487866 | 7.271624 |
| 6.861759 | 6.386416 | 6.310514 | 7.027425 | 7.792656 | 7.350457 | 7.018341 | 7.455457 | 6.792994 |
| 7.456159 | 7.308342 | 7.249966 | 8.014114 | 8.04525 | 7.478126 | 7.661794 | 7.555565 | 7.544776 |
| 5.328371 | 5.022673 | 5.290989 | 2.322189 | NaN | 6.348014 | 4.871385 | 4.592623 | 4.008402 |
| 6.327768 | 5.996915 | 7.462036 | NaN | 5.264275 | 6.774731 | 6.806684 | 6.655631 | 6.023723 |
| NaN | NaN | NaN | NaN | 4.490261 | 5.813063 | 5.549078 | 6.450472 | 6.297229 |
| 8.013563 | 7.754929 | 7.626385 | 8.10572 | 8.115582 | 8.374064 | 8.307709 | 8.399119 | 7.944225 |
| 6.303747 | 6.83303 | 6.068552 | 5.088502 | 6.936724 | 7.935205 | 6.986846 | 8.023113 | 5.964474 |
| 9.032192 | 8.976515 | 9.087212 | 8.044347 | 8.000765 | 9.394802 | 9.126222 | 9.341082 | 9.593418 |
| 9.02672 | 8.005147 | 8.637543 | 9.116858 | 8.512017 | 6.94631 | 6.091958 | 6.388844 | 5.819851 |
| NaN | 8.836185 | NaN | 7.431653 | 7.861747 | NaN | NaN | NaN | NaN |
| 6.661605 | 6.721029 | 6.926477 | 6.621926 | 6.199163 | 6.501276 | 6.687924 | 5.951012 | 5.836486 |
| 4.687953 | 4.879129 | 4.668386 | NaN | 4.526114 | NaN | NaN | NaN | NaN |
| 8.241658 | 8.091498 | 8.200129 | 7.630399 | 7.504258 | 8.269412 | 8.09229 | 8.326854 | 7.567704 |
| 8.343622 | 7.429872 | 7.338446 | 6.374601 | 6.5664 | NaN | NaN | NaN | NaN |
| 6.819245 | NaN | 6.475493 | 6.870292 | 6.638002 | 6.596662 | 6.945159 | 6.481411 | 6.535961 |
| 8.552606 | 8.458436 | 8.445987 | 8.064106 | 7.317671 | 8.045416 | 8.097176 | 8.171682 | 7.948092 |
| 5.041601 | 6.353325 | 5.87277 | 6.722188 | 6.715376 | 5.484503 | 5.570167 | 5.446596 | 5.432156 |
| 6.469332 | 6.986586 | 6.897031 | 7.89526 | 7.956141 | 6.510417 | 6.656841 | 6.661738 | 7.24237 |
| 3.903691 | 4.549348 | 4.57607 | 6.457422 | NaN | 6.288648 | 7.479082 | 7.446485 | 6.939338 |
| 6.896792 | 7.605888 | 6.921256 | 6.137667 | 6.059124 | 5.380164 | 5.488842 | 5.54123 | 5.896543 |
| 7.69133 | 7.6932 | 7.514716 | 7.464009 | 7.694109 | 7.578779 | 7.678822 | 7.778696 | 7.242801 |
| 6.844078 | 6.794755 | 6.476323 | 6.795746 | 6.563615 | 6.173904 | 5.918277 | 5.813962 | 6.13486 |
| 5.844509 | 6.010448 | 5.593446 | NaN | 5.408162 | 6.1376 | 5.599863 | 5.837401 | 5.321423 |
| 9.272308 | 7.989517 | 9.683773 | 6.729702 | 6.984319 | 8.52668 | 9.189543 | 7.750577 | 8.196378 |
| 6.456487 | 6.306098 | 6.493885 | 6.360417 | 5.130981 | 6.671852 | 6.558817 | 6.593563 | 6.750328 |
| 8.561463 | 8.768341 | 8.66577 | 9.141184 | 8.626132 | 9.047626 | 8.971758 | 9.08893 | 8.918777 |
| 11.78977 | 11.98214 | 11.94212 | 11.80462 | 12.15381 | 8.645364 | 8.188228 | 8.688814 | 8.729103 |
| 8.795488 | 8.456993 | 8.539002 | 7.615964 | 6.783508 | 7.6409 | 7.775952 | 7.699748 | 7.326545 |
| 6.109517 | 6.557422 | 6.482922 | 5.927239 | 6.242733 | 6.808987 | 6.818007 | 7.170762 | 6.284197 |
| 10.36202 | 10.37028 | 10.41516 | 10.67039 | 10.73461 | 9.77753 | 9.620951 | 9.878779 | 9.213697 |

| | | | | | | | | |
|---|---|---|---|---|---|---|---|---|
| 4.280352 | 4.906059 | 5.22846 | NaN | 4.631083 | 4.573277 | 5.218417 | 5.882529 | 5.981393 |
| NaN | 3.916552 | NaN | 5.943908 | 7.169195 | 6.207278 | 6.087205 | 6.123625 | 5.940528 |
| 8.463404 | 8.415497 | 8.092628 | 8.016973 | 7.362831 | 8.966548 | 8.572658 | 9.045495 | NaN |
| 6.079249 | 6.113335 | 6.422338 | 5.200597 | 5.871786 | 6.891041 | 6.23812 | 6.644102 | 7.105218 |
| 7.765724 | 7.59729 | 7.547141 | 8.484881 | 8.302554 | 7.496962 | 7.231188 | 7.374239 | 6.996364 |
| 6.901742 | 7.383088 | 7.209312 | 6.19988 | 6.226518 | 7.891028 | 7.887395 | 7.693372 | 6.931339 |
| 8.112753 | 8.025298 | 7.964054 | 7.610251 | 8.167415 | 7.175312 | 7.437613 | 7.376075 | 7.607548 |
| 7.810433 | 7.792839 | 7.810073 | 8.516163 | 8.695317 | 8.076305 | 7.92607 | 7.828692 | 6.468521 |
| 7.584166 | 3.647349 | 4.816274 | 6.185432 | NaN | 7.177444 | 7.215405 | 7.362428 | 7.07421 |
| 5.643383 | NaN | 5.088802 | 8.195555 | 7.670187 | 6.59655 | NaN | NaN | 6.167621 |
| 6.989365 | 6.640319 | 7.018969 | 6.888316 | 6.475459 | 7.651731 | 7.669861 | 7.732077 | 7.217647 |
| 7.959184 | 7.899079 | 7.888667 | 8.213485 | 8.547399 | 7.501335 | 7.588036 | 7.688961 | 7.386975 |
| 8.544254 | 8.432771 | 8.72212 | 8.881166 | 8.802104 | 8.360523 | 8.271514 | 8.132383 | 8.138817 |
| 8.053087 | 7.954576 | 8.055811 | 8.069414 | 7.826969 | 7.75342 | 8.060707 | 7.600161 | 7.957205 |
| 7.580541 | 7.542604 | 7.49932 | 8.238217 | 8.328353 | 8.196595 | 8.045665 | 8.081296 | 7.252851 |
| 6.973398 | 6.492005 | 6.317253 | 7.008713 | 7.554877 | 7.689359 | 7.673476 | 7.498678 | 6.667922 |
| 6.750686 | 6.651075 | 6.555092 | 6.385802 | 6.049156 | 7.01507 | 6.460805 | 6.961818 | 6.120532 |
| 8.042986 | NaN | NaN | 7.799338 | 8.825438 | NaN | NaN | NaN | NaN |
| 6.344862 | 6.373652 | 6.014922 | 7.282499 | 7.044849 | 6.302488 | 6.502632 | NaN | 4.951173 |
| 6.279642 | 6.871264 | 6.815125 | 6.174087 | NaN | 6.144378 | 5.556194 | 5.601315 | 6.445325 |
| 5.92375 | 5.234397 | 3.285149 | 6.192953 | 6.439844 | NaN | NaN | NaN | NaN |
| 7.351757 | 7.187396 | 6.927054 | 7.659692 | 7.666831 | 7.564381 | 7.193913 | 7.662095 | 7.857874 |
| 8.46172 | 8.371876 | 8.177869 | 7.352969 | 7.74864 | 7.691539 | 7.700538 | 7.762462 | 7.6514 |
| 6.091719 | 6.531183 | 6.658651 | NaN | 7.383967 | NaN | NaN | NaN | NaN |
| 7.945466 | 8.046851 | 8.036006 | 7.175656 | 7.084426 | 7.428529 | 7.420097 | 7.494629 | 7.211367 |
| 8.399414 | 7.372416 | 7.571306 | 7.549511 | 8.813355 | 7.550835 | 7.419257 | NaN | 7.923006 |
| NaN | 5.882944 | NaN | 6.614756 | 7.449471 | NaN | NaN | 6.761094 | 5.322048 |
| 7.96097 | 7.968144 | 8.098042 | 8.325809 | 8.33383 | 8.088414 | 7.839214 | 7.758866 | 7.58074 |
| 7.068925 | 6.902493 | 7.046622 | 7.141932 | 6.911739 | 6.699931 | 6.765827 | 6.951131 | 5.99318 |
| 7.750319 | 7.274348 | 7.361236 | 7.398709 | 7.618701 | 6.590174 | 7.216966 | 6.943161 | 7.142882 |
| 8.691772 | 8.784774 | 8.950497 | 8.294374 | 8.209718 | 8.286217 | 8.36875 | 8.327575 | 8.064034 |
| 6.834586 | 7.00737 | 7.210598 | 5.92945 | 6.6796 | 6.760724 | 6.590548 | 6.612662 | 4.970097 |
| 4.98846 | 6.341734 | 6.024469 | 9.029337 | 9.143465 | NaN | 3.849697 | 4.225706 | 5.56909 |
| 7.490217 | 7.503815 | 7.439683 | 5.755559 | 6.487207 | 7.331307 | 7.603376 | 7.407618 | 6.749183 |
| 7.701977 | 7.341561 | 7.218248 | 7.807478 | 7.677276 | 6.821088 | 6.832078 | 6.689038 | 6.989946 |
| 8.708538 | 8.663531 | 8.818295 | 8.366297 | 8.030426 | 8.632338 | 8.602438 | 8.623693 | 8.533393 |
| 7.804907 | 8.028234 | 7.629924 | 6.839054 | 7.017416 | 7.170135 | 7.833437 | 7.589432 | 7.935474 |
| 7.148115 | 7.480571 | 7.50872 | 6.840578 | 6.802114 | 7.322366 | 7.540959 | 7.43204 | 7.436237 |
| 8.482764 | 8.794479 | 8.806143 | 9.182524 | 9.211387 | 8.176422 | 7.813845 | 7.974651 | 7.659053 |
| 6.803083 | 7.831165 | 6.859212 | 7.322587 | 7.330203 | 7.026554 | 7.045259 | 6.952969 | 6.879122 |
| 6.289859 | 6.739948 | 6.984445 | 6.155396 | 6.763632 | 8.768861 | 8.279087 | 7.864996 | 6.872319 |
| NaN | NaN | NaN | NaN | NaN | NaN | NaN | NaN | 4.761607 |
| 5.825736 | 6.012727 | 5.709074 | 4.773379 | 4.897357 | 5.490753 | 5.155322 | 5.422174 | 4.470634 |
| 7.45077 | 6.866258 | 7.372716 | 7.293798 | 7.472795 | 6.303029 | 6.253444 | 7.104743 | 6.984323 |
| 8.148374 | 8.19028 | 8.166508 | 8.156639 | 8.428638 | 7.404172 | 7.206515 | 7.455768 | 7.117129 |
| 7.407047 | 7.278461 | 7.220737 | 6.763375 | 6.943992 | 6.85675 | 7.014486 | 7.274951 | 6.107074 |
| 8.151488 | 8.38754 | 8.139112 | 6.028075 | 6.029142 | 8.160604 | 7.881213 | 8.030178 | 7.712319 |

| | | | | | | | | |
|---|---|---|---|---|---|---|---|---|
| 7.971349 | 7.698644 | 7.782453 | 6.99009 | 6.819013 | 7.064875 | 6.963776 | 7.459037 | 6.38385 |
| 7.075179 | 6.266726 | 7.152066 | 6.324781 | NaN | 6.670472 | 6.992824 | 6.230406 | 5.841114 |
| 7.753839 | 7.746362 | 7.732811 | 8.056644 | 8.404468 | 7.891623 | 8.21637 | 8.075053 | 7.114344 |
| 7.98543 | 6.929521 | 7.29724 | 7.283695 | 7.236698 | 6.927567 | 6.645735 | 7.219497 | 7.032278 |
| 8.130594 | 8.012136 | 7.755132 | 6.542145 | 6.482987 | NaN | 7.11191 | 7.370646 | 7.443417 |
| 11.42272 | 11.47779 | 11.52349 | 10.51629 | 10.39739 | 11.33609 | 11.27611 | 11.05117 | 10.45326 |
| 8.643394 | 8.943936 | 8.704903 | 9.020325 | 8.662082 | 6.790806 | 7.5021 | 7.100242 | 6.432099 |
| 8.134429 | 8.343035 | 8.411534 | 7.188698 | 6.534504 | 7.442273 | 7.35628 | 7.638461 | 7.673811 |
| 6.321239 | 6.584731 | 6.655534 | 5.626976 | 6.109724 | 5.599372 | 5.443758 | 5.650549 | 5.998591 |
| 7.420276 | 8.093405 | 7.488953 | 7.173276 | 7.289299 | 8.152642 | 7.77795 | 7.859817 | 6.58205 |
| 6.638499 | 6.551346 | NaN | NaN | 6.999491 | 8.084912 | 7.522692 | 7.879968 | 6.815736 |
| 6.271683 | 5.800899 | 6.504279 | 5.195275 | 5.262344 | 5.871397 | 5.809237 | 6.124471 | 6.020558 |
| 9.804826 | 9.708097 | 9.71056 | 9.286391 | 9.266364 | 9.377136 | 9.497868 | 9.430635 | 9.058 |
| 6.374124 | 5.723482 | 5.929024 | 7.09819 | 6.746048 | NaN | NaN | 6.195576 | 6.199943 |
| 8.108705 | 8.194195 | 7.747377 | 7.912953 | 7.977881 | 7.67186 | 7.529845 | 7.626391 | 6.60503 |
| NaN | 5.017603 | NaN | 6.013951 | 5.396055 | 5.75637 | 6.295995 | 5.850582 | 6.694726 |
| 6.201003 | 5.712034 | 7.273746 | 5.628868 | 6.147128 | 6.370039 | 6.301582 | 6.675557 | 5.74452 |
| 5.11491 | 5.561418 | 4.403627 | NaN | NaN | 5.481602 | 5.42869 | 5.31561 | 5.625859 |
| 7.814085 | 7.909331 | 7.930318 | 7.859385 | 7.683341 | 7.581586 | 7.675723 | 7.706658 | 7.349809 |
| 7.502385 | 7.70611 | 7.437832 | NaN | 5.369804 | NaN | NaN | 5.163508 | 5.187762 |
| 8.697346 | 8.199396 | 8.120176 | 9.733999 | 9.226782 | 8.011846 | 7.720488 | 7.84271 | 7.912668 |
| 8.437184 | 8.684826 | 8.247663 | 7.494282 | 7.404924 | 9.248304 | 9.239159 | 9.246414 | 8.840827 |
| 5.035906 | NaN | 4.285796 | 3.4558 | NaN | 5.537217 | 5.924115 | 6.31772 | 4.930237 |
| 7.355306 | 5.829492 | 6.570621 | 6.597987 | NaN | 6.639905 | 6.572709 | 7.147724 | 6.640281 |
| 6.813126 | 7.003001 | 6.975809 | 7.158863 | 6.956219 | 6.946064 | 7.080399 | 6.877882 | 6.779938 |
| 8.127185 | 7.964507 | 8.12071 | 7.676427 | 8.546763 | 7.777812 | 5.918071 | 7.71047 | 7.704336 |
| 6.737545 | 6.68575 | 7.172746 | 5.957617 | 6.278191 | 6.752857 | 7.011674 | 7.561931 | 6.451561 |
| 6.584532 | 5.934236 | 6.97826 | 5.513129 | NaN | 7.022141 | 7.490702 | 6.887764 | 6.035783 |
| 7.571174 | 7.247232 | 7.391157 | 7.669097 | 6.832363 | 8.099065 | 8.286882 | 8.218884 | 7.554797 |
| 7.806743 | 7.723065 | 7.577147 | 8.861805 | 8.897374 | 7.815341 | 8.000562 | 7.308731 | 7.728421 |
| 6.836217 | 6.591274 | 6.816602 | 6.775534 | 6.95295 | 6.645447 | 6.638396 | 6.655429 | 6.90356 |
| 7.284067 | 6.219237 | 6.893102 | NaN | NaN | 4.826894 | 5.558368 | 4.863731 | NaN |
| NaN | 6.403638 | NaN | 5.397091 | 5.54457 | 6.375473 | 6.612208 | 6.924123 | 4.672005 |
| 7.865855 | 7.936632 | 7.503823 | 7.125391 | 6.554441 | 8.457072 | 8.548073 | 7.908751 | 7.202081 |
| 7.899019 | 7.609879 | 7.724592 | 6.949404 | 6.713137 | 7.701298 | 7.600041 | 7.740397 | 7.509431 |
| 8.147655 | 8.353025 | 8.167286 | 6.713558 | 6.560639 | 8.220978 | 8.234241 | 8.283871 | 7.45706 |
| 9.714445 | 9.485492 | 9.606586 | 10.14652 | 10.63607 | 7.945461 | 7.866596 | 7.909886 | 7.996583 |
| 7.658689 | 7.107575 | 7.644238 | 6.103964 | 6.153467 | 7.734403 | 7.2604 | 7.4867 | 6.693007 |
| 7.561234 | 6.954192 | 7.124089 | 5.697318 | 6.275448 | 6.253704 | 6.069057 | 6.627056 | 6.922544 |
| 5.96438 | 5.303566 | 5.737251 | 5.678398 | 6.425131 | 5.069555 | 4.913784 | 5.474786 | 5.529119 |
| 6.68186 | 6.6904 | 6.617616 | 6.899216 | 6.785452 | 5.75714 | 6.555161 | 6.225811 | 6.628967 |
| NaN | 4.83884 | NaN | NaN | 4.633636 | 6.158373 | 5.801786 | 5.677903 | 4.418061 |
| 7.528466 | 7.210578 | 7.400771 | 8.07945 | 8.132748 | 7.39769 | 7.508329 | 7.399757 | 7.507264 |
| 7.829144 | 7.628979 | 8.116176 | 8.465984 | 8.651683 | 7.872333 | 8.045436 | 8.164426 | 8.123418 |
| 6.950171 | 6.602058 | 6.334202 | 6.269918 | 6.382283 | 6.759125 | 7.235569 | 6.613596 | 5.903281 |
| 5.402045 | NaN | 5.409268 | NaN | NaN | 4.675005 | 3.562222 | 3.606294 | NaN |
| 8.650039 | 7.569591 | 7.989383 | 6.263438 | NaN | 7.769283 | 6.698834 | 6.875607 | NaN |

| | | | | | | | | |
|---|---|---|---|---|---|---|---|---|
| 6.774843 | 6.970588 | 7.060817 | 5.770226 | 5.69501 | 6.590209 | 6.975737 | 6.730533 | 6.793024 |
| 7.354671 | 7.195632 | 7.165475 | 7.037269 | 6.885414 | 6.844929 | 6.89798 | 6.887689 | 7.154509 |
| 7.07453 | 6.7901 | 7.275368 | 6.775434 | 6.618235 | 8.176331 | 8.009597 | 8.01456 | 6.525967 |
| 7.293698 | 6.915647 | 7.132864 | 5.425971 | 4.691467 | 6.98317 | 7.164139 | 7.092687 | 7.485453 |
| 8.462125 | 8.520701 | 8.68962 | 8.680522 | 8.527509 | 9.091187 | 9.076977 | 9.264038 | 8.998521 |
| 8.972831 | 7.89119 | 9.148021 | 7.388833 | 5.380134 | 9.32148 | 9.423744 | 9.623365 | 8.954082 |
| 6.93985 | 7.026608 | 6.694709 | 7.28576 | 7.742561 | 6.864172 | 6.877869 | 6.563615 | 6.795676 |
| NaN | NaN | NaN | 7.972721 | 7.434896 | NaN | NaN | NaN | NaN |
| 8.44614 | 8.429187 | 8.826431 | 9.577724 | 9.714655 | 7.96158 | 7.880564 | 7.813911 | 7.54086 |
| 8.071095 | 8.011041 | 7.995597 | 7.379506 | 7.908716 | 7.233205 | 6.323541 | 5.922192 | 3.750363 |
| 9.321629 | 9.392348 | 9.296497 | 10.02676 | 9.826087 | 8.857298 | 8.818302 | 8.764465 | 9.290531 |
| 6.787022 | 6.122092 | 6.093487 | NaN | NaN | 8.247704 | 8.63617 | 8.402933 | 6.150461 |
| 7.821424 | 7.872546 | 7.836379 | 6.309318 | 6.509202 | 7.280022 | 7.047363 | 7.487576 | 7.406466 |
| 7.7037 | 7.711728 | 7.398424 | 6.423616 | 6.982343 | 7.598996 | 7.810672 | 7.958456 | 6.720862 |
| 5.581109 | 5.594238 | 4.716806 | NaN | NaN | 5.86475 | 6.057968 | 6.031616 | 6.031696 |
| 7.976118 | 8.217412 | 8.046917 | 8.139522 | 8.115951 | 8.629382 | 8.920111 | 8.566223 | 7.653033 |
| 8.612976 | 8.558831 | 8.606768 | 9.51946 | 9.458075 | 8.301248 | 8.218179 | 8.898939 | 8.140261 |
| 6.521397 | 7.024274 | 6.270741 | NaN | NaN | 5.811985 | 5.449074 | 6.013771 | 5.588845 |
| 7.090299 | 7.127372 | 6.85328 | 7.698419 | 7.444771 | 6.487292 | 6.734608 | 6.785037 | 6.114369 |
| 8.172056 | 7.180389 | 7.061447 | NaN | 6.980889 | 7.058888 | 7.316295 | 7.4141 | 7.086602 |
| 8.181835 | 8.454549 | 8.506676 | 8.69775 | 8.888084 | 7.687033 | 8.049451 | 8.117195 | 8.636023 |
| 6.780906 | 6.335886 | 6.901846 | 5.092879 | 5.125034 | 5.140127 | 5.467046 | 6.045787 | 7.199082 |
| 9.545152 | 9.591208 | 9.554493 | 7.231876 | 7.190583 | 9.269883 | 9.044854 | 9.116103 | 9.196652 |
| 5.777378 | 5.775443 | 6.017108 | 6.95199 | 7.256049 | 4.940202 | 4.922032 | 5.617478 | 6.68231 |
| 7.512035 | 7.617111 | 7.568846 | 7.808703 | 7.556009 | 7.382055 | 7.116685 | 7.478662 | 8.123595 |
| 8.028316 | 8.00173 | 7.471232 | 10.18138 | 10.08868 | 8.034605 | 7.954185 | 8.095817 | 8.595384 |
| NaN | 5.466796 | NaN | 4.834964 | 6.802866 | 6.318619 | 6.492721 | NaN | 6.583727 |
| 6.033199 | 6.014042 | 6.049538 | NaN | NaN | 6.056186 | 5.682199 | 5.788854 | 5.146716 |
| 7.310211 | 7.045164 | 7.162954 | 6.742315 | 7.05361 | 7.224261 | 7.579659 | 7.459728 | 7.419372 |
| 7.538892 | 6.954249 | 7.642663 | 6.784375 | 7.008557 | 7.817777 | 7.764473 | 7.671311 | 7.320667 |
| 5.372624 | 5.856131 | 5.763849 | 5.232529 | 6.132406 | 6.235797 | 5.40133 | 4.398072 | 4.714742 |
| 6.973174 | 6.997198 | 6.996766 | 6.432098 | 5.110271 | 7.903088 | 7.896134 | 8.116262 | 6.196043 |
| 7.070601 | 7.093356 | 8.028688 | NaN | NaN | 6.979675 | 7.143359 | 7.622302 | 6.37236 |
| 8.394072 | 8.343826 | 8.296532 | 10.07074 | 9.77636 | 8.074671 | 8.037575 | 7.966282 | 8.37972 |
| 11.2471 | 11.07753 | 11.1134 | 13.00087 | 12.94064 | 11.9348 | 11.76356 | 12.03511 | 11.72036 |
| 7.036288 | 6.987045 | 6.898049 | 8.633915 | 7.767614 | 7.366745 | 7.245636 | 6.900595 | 7.305144 |
| 7.333672 | 7.449094 | 7.547617 | 7.875716 | 7.622586 | 7.276035 | 6.942684 | 7.867223 | 7.863812 |
| NaN | NaN | NaN | 5.447523 | 5.173438 | NaN | 5.344439 | NaN | 5.667869 |
| 9.038895 | 9.035501 | 8.899229 | 8.170125 | 8.184548 | 9.491757 | 9.306891 | 9.51446 | 8.650886 |
| 8.044547 | 8.160263 | 7.829082 | 6.858222 | 7.306107 | 7.934939 | 8.15933 | 7.936258 | 8.025948 |
| 6.427595 | 7.066465 | 6.547205 | 4.658493 | NaN | 6.603354 | 6.695485 | 7.176626 | 7.378562 |
| 8.990341 | 9.042319 | 8.913239 | 10.95842 | 10.86501 | 9.592729 | 9.801308 | 9.660889 | 9.443185 |
| 5.644072 | 5.202379 | 5.482875 | 6.334372 | 5.845558 | 4.880681 | 3.860769 | 4.353397 | 5.45432 |
| NaN | 6.162773 | NaN | NaN | NaN | 7.831558 | 8.699659 | 7.924343 | 6.921446 |
| 6.762435 | 6.897528 | 6.927607 | 6.676846 | 6.118797 | 6.890744 | 6.819117 | 6.716936 | 6.758825 |
| 8.215713 | 7.998241 | 8.133548 | 7.378253 | 7.100711 | 8.427481 | 8.468723 | 8.114722 | 8.134564 |
| 6.947566 | 7.438322 | 6.861944 | 7.240546 | 8.09802 | 7.205041 | 6.439659 | 7.106045 | 6.894578 |

-Bruker Confidential-

| | | | | | | | | |
|---|---|---|---|---|---|---|---|---|
| 7.976098 | 7.019879 | 7.399606 | NaN | 6.215607 | 8.6547 | 9.081031 | 8.907918 | 7.151312 |
| 9.982536 | 9.993583 | 10.01447 | 10.49893 | 10.21835 | 9.876391 | 9.911224 | 9.964177 | 9.633745 |
| 8.177997 | 8.143683 | 7.873685 | 9.959719 | 9.921307 | 9.140295 | 9.152103 | 9.140589 | 7.981507 |
| 7.732034 | 8.638978 | NaN | 6.218545 | 6.464587 | NaN | 6.673499 | 7.055856 | NaN |
| 7.076233 | 6.893413 | 7.270712 | 6.262997 | 5.865443 | 7.391514 | 7.686957 | 7.578969 | 6.91105 |
| 6.008576 | 5.235089 | 6.217977 | NaN | 4.697963 | 5.657527 | 4.596909 | 4.748673 | 5.882087 |
| 6.765466 | 7.077572 | 7.002784 | 6.986842 | 7.269351 | 6.824265 | 6.658478 | 7.123911 | 6.901713 |
| 3.848651 | NaN | 3.884442 | NaN | NaN | NaN | NaN | NaN | 4.141037 |
| 7.190147 | 7.292279 | 7.161674 | 7.292374 | 7.282189 | 7.654684 | 7.654881 | 7.609698 | 7.074492 |
| 6.100319 | 7.351349 | NaN | 6.181436 | 6.293233 | NaN | 5.449092 | NaN | 6.991137 |
| 7.092169 | 5.813373 | 6.878151 | 5.925971 | 6.145969 | 7.318369 | 6.957345 | 7.244082 | 6.99396 |
| 6.356821 | 4.276027 | NaN | 7.556213 | 7.637452 | 6.242937 | 7.013694 | 6.990842 | 7.113564 |
| 7.391971 | 7.069141 | 7.314791 | 5.478773 | 6.693201 | 6.889426 | 7.508095 | 7.362102 | 7.519047 |
| 8.954193 | 8.555805 | 8.74447 | 9.797683 | 10.4691 | 9.454885 | 9.273767 | 9.822314 | 10.10054 |
| 4.895869 | 5.163475 | 5.600204 | 4.91615 | 4.419348 | 5.763807 | 5.29645 | 4.994182 | 4.679718 |
| 7.718888 | 7.360485 | 7.726764 | 6.773898 | 6.662788 | 6.946269 | 6.868401 | 6.979067 | 7.235349 |
| 9.094627 | 7.938039 | 8.366739 | 8.560928 | 6.654923 | 7.02042 | 8.660541 | 8.088838 | NaN |
| 7.621027 | 7.41551 | 7.573111 | 5.82794 | 5.543368 | 7.907391 | 7.937558 | 7.960066 | 7.146581 |
| 6.745235 | 6.916293 | 7.165287 | 6.461855 | 6.073116 | 6.73702 | 6.842071 | 6.91287 | 6.600798 |
| 5.807296 | 6.012255 | NaN | NaN | 5.121613 | NaN | NaN | NaN | 4.523593 |
| 7.47189 | 7.634207 | 7.88557 | 6.869712 | 6.435888 | NaN | 7.115231 | 7.450546 | 6.726906 |
| 4.576935 | 5.253766 | 4.629347 | NaN | NaN | 4.873043 | NaN | 3.665085 | 4.982606 |
| 7.464802 | 7.629336 | 7.559581 | 8.930882 | 8.720834 | 7.010191 | 6.891124 | 6.869296 | 6.184687 |
| 5.837834 | 5.58926 | 5.774893 | 6.717961 | 6.766905 | 6.07191 | 5.490758 | 6.200406 | 5.708239 |
| 4.964911 | 4.774652 | 4.92275 | 2.713645 | 4.121468 | 3.0285 | NaN | 3.42186 | 4.586311 |
| 8.674202 | 8.87069 | 9.118677 | 10.23003 | 10.27138 | 8.619318 | 8.736121 | 8.791024 | 8.737443 |
| 7.931915 | 7.343358 | 7.99271 | 6.314281 | 6.100877 | 7.40482 | 7.753901 | 7.513084 | 7.578232 |
| 8.105117 | 7.312194 | 8.007736 | 6.167304 | 6.370408 | 6.633642 | 6.658293 | 6.626757 | 6.34155 |
| 6.976892 | 7.005946 | 6.815436 | 6.749182 | 6.608144 | 6.837304 | 6.650211 | 7.326684 | 6.734251 |
| NaN | 6.220049 | NaN | 5.593084 | 6.801593 | NaN | 6.994869 | 6.185783 | NaN |
| 5.936471 | 5.99019 | 5.703831 | 6.303441 | 6.772707 | 5.737878 | 5.542112 | 5.59134 | 6.40324 |
| 6.733875 | 7.272083 | 8.10802 | 6.860713 | 6.888167 | 7.529428 | 7.219226 | 7.1384 | 7.290958 |
| 7.228508 | 7.773973 | 8.044978 | 5.53314 | NaN | 7.552794 | 7.683258 | 7.714131 | 7.297419 |
| 8.777983 | 9.04323 | 9.029114 | 7.982822 | 7.963669 | 8.677272 | 8.736221 | 8.783408 | 8.566101 |
| 5.990113 | 4.862793 | 6.046419 | 5.099652 | 5.61847 | 6.402779 | 5.82336 | 6.484714 | 5.570052 |
| 6.56679 | 6.57483 | 6.199177 | 6.516433 | 6.840493 | 6.510999 | 6.1387 | 6.864217 | 6.031381 |
| 7.126396 | 7.100567 | 6.769902 | 11.26306 | 10.56073 | 9.177935 | 9.164544 | 8.956595 | 10.72732 |
| 7.634866 | 7.538098 | 7.503155 | 7.77049 | 7.697191 | 6.868014 | 6.964141 | 6.651842 | 6.536502 |
| 5.683336 | 6.27382 | 5.46085 | 6.147701 | 6.126848 | 6.363106 | 5.957646 | 6.032062 | 5.425161 |
| 7.766693 | 7.390661 | 7.896913 | 8.523224 | 8.645713 | 7.713329 | 7.696919 | 7.669057 | 8.420326 |
| 8.30423 | 7.804828 | 8.345434 | 6.775211 | 7.036894 | 8.168119 | 8.429435 | 8.372665 | 8.318797 |
| 7.230218 | 7.088903 | 6.025112 | 5.393661 | 5.950356 | 7.195148 | 6.949172 | 6.386498 | 7.232917 |
| 7.360587 | 7.37959 | 7.428916 | 8.705152 | 8.171363 | 6.868142 | 7.161595 | 6.705936 | 6.908192 |
| 8.335417 | 8.024229 | 8.255535 | 7.015166 | 9.259901 | 8.031757 | 7.428499 | 7.870318 | 6.737582 |
| 8.264357 | 7.623434 | 8.478661 | 7.364806 | 6.654725 | 7.92337 | 8.210952 | 8.447334 | 7.759494 |
| 6.085649 | 6.539517 | 6.410545 | 6.079313 | 6.073201 | 6.869084 | 7.239414 | 7.283996 | 6.61389 |
| 5.592226 | 5.847636 | 6.463957 | 6.158053 | 6.285472 | 6.352292 | 6.282313 | 6.46707 | 6.621622 |

| | | | | | | | | |
|---|---|---|---|---|---|---|---|---|
| 5.077322 | 5.5363 | 4.973469 | 5.326068 | 5.643296 | 5.840507 | 5.533307 | 5.75888 | 5.551597 |
| 6.768662 | 6.876813 | 6.274771 | 7.419542 | 7.345922 | 7.580616 | 7.663806 | 7.583044 | 6.722087 |
| 7.116005 | 7.020088 | 6.939148 | 7.361984 | 7.138658 | 6.821174 | 6.86691 | 7.13895 | 5.966489 |
| 6.243013 | 6.685874 | NaN | 7.141872 | 6.703915 | 5.697899 | NaN | 5.575074 | 6.415968 |
| 7.097141 | 7.288767 | 7.673068 | 6.516478 | 5.901555 | 7.142413 | 7.050311 | 7.283225 | 7.099775 |
| 5.391988 | 5.376423 | 5.423782 | 7.221493 | 7.336496 | 5.483395 | 5.672688 | 5.553459 | 4.803362 |
| 7.851992 | 7.767904 | 7.707017 | 8.374641 | 8.272799 | 7.545713 | 7.289256 | 7.154788 | 7.576183 |
| 7.651293 | 7.365984 | 7.653728 | 6.650048 | 6.625959 | 7.433844 | 7.227216 | 7.577854 | 7.081594 |
| 5.335804 | 5.849893 | 5.952224 | 8.263947 | 7.955752 | NaN | 4.594865 | NaN | 6.751525 |
| 8.95666 | 9.120758 | 8.889705 | 7.444179 | 7.316861 | 8.215091 | 8.429368 | 8.208502 | 8.023635 |
| 10.65821 | 10.57602 | 10.67483 | 10.52176 | 9.128006 | 10.15608 | 10.51248 | NaN | 9.675982 |
| 7.4028 | 7.466731 | 7.527546 | 6.645871 | 6.14805 | 7.173685 | 7.277448 | 7.292602 | 6.875175 |
| 5.835822 | 5.123238 | 5.807063 | NaN | 5.18252 | 6.08393 | 5.890249 | NaN | 5.908283 |
| 5.933494 | 6.813176 | 6.80475 | 6.198373 | 6.419736 | 5.987198 | 6.423286 | 6.158559 | 6.973622 |
| 3.375391 | 5.119003 | 6.075853 | 5.166837 | 5.609407 | 5.505635 | 5.400998 | 6.05501 | 6.153962 |
| 5.378136 | 5.080839 | 6.233142 | NaN | NaN | 6.068164 | 5.992319 | 6.204646 | 5.956208 |
| 7.146397 | 6.185633 | 7.30379 | 5.377923 | 6.512202 | 7.985401 | 7.63604 | 7.492472 | 6.566111 |
| 6.693907 | 7.395402 | 6.822034 | 6.743856 | 7.787746 | 7.833542 | 7.561505 | 6.449358 | 6.756428 |
| 6.192289 | 6.321367 | 6.32472 | 6.219928 | 5.973066 | 6.179099 | 5.789751 | 6.392334 | 5.587367 |
| 7.397794 | 7.086334 | 7.191844 | 5.70003 | 5.634707 | 6.795015 | 6.203023 | 7.582964 | 6.957234 |
| 6.416768 | 6.887856 | 6.845875 | 5.791115 | 5.948383 | 7.001128 | 6.828771 | 6.950592 | 7.015717 |
| 5.903441 | 6.155727 | 5.242915 | 6.381386 | 6.07072 | NaN | 7.43126 | NaN | 6.197836 |
| 8.633808 | 8.779061 | 8.44867 | 9.737496 | 9.638983 | 7.899452 | 7.870204 | 7.781182 | 7.89977 |
| NaN | 4.618742 | 5.157275 | NaN | NaN | NaN | NaN | NaN | 4.658787 |
| 7.360661 | 7.705976 | 7.700291 | 6.667039 | 6.650363 | 7.584711 | 7.097871 | 7.568368 | 5.528141 |
| 5.973472 | 5.835781 | 4.654134 | 6.903002 | 6.890968 | 6.86258 | 6.460294 | 6.79135 | 5.964046 |
| 7.802642 | 8.172939 | 8.02397 | 10.36727 | 10.44183 | 6.979865 | 6.223943 | 6.503068 | 6.891718 |
| 7.524318 | 6.506495 | 7.340444 | 6.83694 | NaN | 5.862851 | 6.540198 | 6.867757 | 6.358851 |
| 9.23106 | 9.522082 | 9.316147 | 8.387124 | 7.790846 | 8.458425 | 8.212674 | 8.510136 | 7.987344 |
| 6.264767 | 6.420393 | 6.234685 | NaN | 4.618609 | 6.004909 | 6.219452 | 6.129085 | 5.995598 |
| 6.20758 | 5.003287 | 6.463418 | NaN | 6.926867 | 7.279104 | 6.980105 | 7.635105 | 6.451148 |
| 7.684235 | 7.354462 | 7.242767 | 6.579466 | 6.417939 | 7.567871 | 7.768119 | 7.955469 | 8.012437 |
| 7.950459 | 7.458333 | 7.834119 | 6.32583 | 6.2834 | 7.550616 | 7.153982 | 7.575027 | 7.438803 |
| 8.243586 | 8.351316 | 8.189857 | 7.534312 | 7.422018 | 7.830151 | 7.617968 | 7.719891 | 7.887854 |
| 6.26219 | 6.386872 | 6.380714 | 5.089419 | 5.653317 | 6.963854 | 6.912745 | 6.586625 | 6.697322 |
| 11.34424 | 11.2996 | 11.34305 | 10.82184 | 10.56867 | 11.7816 | 11.69235 | 11.7077 | 10.76928 |
| 7.871202 | 7.167209 | 7.280265 | 7.996235 | 6.777667 | 6.85788 | 6.576173 | 6.578687 | 7.038073 |
| 7.212519 | 7.268116 | 7.179053 | 5.46882 | 5.839448 | 7.308535 | 6.787867 | 7.005976 | 7.070457 |
| 6.272572 | 5.040366 | 6.271708 | NaN | NaN | 7.054478 | 7.040573 | 7.081564 | 5.796672 |
| 9.289412 | 9.382323 | 9.268167 | 10.23813 | 10.21013 | 7.56908 | 7.604885 | 7.775648 | 7.884468 |
| 7.03919 | 6.545264 | 6.817133 | 5.39533 | 5.536907 | 6.823399 | 7.522966 | 6.336683 | 5.773347 |
| 8.236113 | 7.988062 | 8.045462 | 7.097082 | 6.928405 | 7.475332 | 7.466481 | 7.264396 | 7.849332 |
| 9.403776 | 9.304336 | 9.183263 | 8.492712 | 8.621905 | 9.905046 | 9.750315 | 9.969675 | 9.597898 |
| 6.151125 | 5.107518 | 5.434101 | 4.676791 | 5.369779 | 5.99311 | 5.610858 | 5.311468 | 6.294989 |
| NaN | 7.729089 | 7.518174 | 6.207577 | 6.723943 | NaN | NaN | NaN | NaN |
| 6.809108 | 6.963376 | 7.004355 | 6.179746 | 6.418252 | 7.285996 | 7.268615 | 7.181224 | 7.249326 |
| NaN | NaN | NaN | 6.257221 | 6.540956 | 6.416083 | 6.827374 | 6.908676 | 6.196862 |

| | | | | | | | | |
|---|---|---|---|---|---|---|---|---|
| 6.428796 | 6.570139 | 8.562767 | 4.919704 | 5.087211 | 8.409531 | 7.941662 | 7.556043 | 7.796472 |
| 4.356259 | 3.461825 | 2.857209 | NaN | NaN | NaN | NaN | NaN | 4.13132 |
| 8.762797 | 8.621979 | 8.345854 | 7.468437 | 7.945131 | 7.904603 | 7.81149 | 7.399981 | 9.034014 |
| 7.030382 | 5.890083 | 7.003759 | 5.885864 | 6.027167 | 6.643775 | 6.476704 | 7.049596 | 5.719502 |
| 6.628384 | 6.53364 | 6.288199 | 7.437036 | 6.96959 | 7.205862 | 7.438816 | 7.113891 | 6.711061 |
| 8.297771 | 7.99941 | 8.214238 | 8.096336 | 7.660908 | 8.532759 | 8.526489 | 8.617309 | 8.000204 |
| 7.665713 | 8.097612 | 8.075448 | 6.898989 | 6.54811 | 8.095276 | 8.161294 | 7.954221 | 7.759606 |
| 8.535267 | 8.48819 | 8.568696 | 10.49318 | 10.22258 | 9.247265 | 9.412439 | 8.875099 | 8.39948 |
| 2.694195 | 3.820707 | 2.688269 | 4.289082 | 4.067604 | 4.611741 | 3.266102 | 4.841311 | 4.495232 |
| 7.353595 | 7.244336 | 7.520293 | 6.914006 | 5.704373 | NaN | 6.634327 | 6.1692 | 6.682674 |
| 7.303517 | 6.926259 | 7.193961 | 7.733657 | 6.423972 | 6.110744 | 7.310449 | 6.576795 | 7.66447 |
| 7.462515 | 6.740394 | 6.572937 | 6.705964 | 6.743906 | 6.394392 | 6.224512 | 5.697795 | 7.432139 |
| 7.039331 | 7.055006 | 7.008364 | 7.871364 | 7.662897 | 7.363442 | 6.880392 | 7.136189 | 4.34548 |
| 7.884695 | 8.063761 | 8.251524 | 6.418351 | 6.284791 | 7.98844 | 7.905412 | 7.83131 | 7.326061 |
| 6.611746 | 7.521438 | 6.369957 | 5.486598 | 5.924873 | 6.961334 | 6.907313 | 6.733155 | 5.943306 |
| 8.35959 | 8.381738 | 8.369508 | 11.1783 | 11.14766 | 8.053199 | 8.315768 | 8.440712 | 8.412817 |
| 6.800248 | 7.069117 | 7.097468 | 7.006288 | 6.838363 | 7.328237 | 7.312645 | 7.205451 | 7.083848 |
| 7.526515 | 7.658565 | 7.551919 | 7.17634 | 6.784399 | 8.052597 | 8.133355 | 8.089972 | 7.984005 |
| 8.371427 | 8.48213 | 8.164852 | 6.944113 | 7.87905 | 7.576993 | 7.525591 | 6.966738 | 7.754232 |
| 7.958753 | 7.758989 | 7.934961 | 6.53581 | 6.453965 | 7.491794 | 7.528053 | 7.561949 | 7.966186 |
| 8.953843 | 9.050952 | 8.907582 | 9.590885 | 9.728981 | 8.502089 | 8.314099 | 8.471186 | 8.364844 |
| 3.734637 | 4.457256 | NaN | NaN | NaN | 3.872638 | 4.465425 | 4.618535 | 3.92325 |
| 5.094048 | NaN | 5.143133 | NaN | NaN | 6.493437 | 6.647375 | 6.403648 | 4.781708 |
| 8.895841 | 9.054868 | 9.172034 | 8.826036 | 9.02313 | 9.192003 | 9.150877 | 8.977489 | 9.152528 |
| 6.133656 | 6.342484 | 4.864346 | 5.580317 | 5.854996 | 5.958383 | 6.082729 | 5.554516 | 5.115153 |
| 7.886675 | 6.70811 | 7.990743 | 6.690734 | 7.463983 | NaN | NaN | NaN | 6.939304 |
| 10.10828 | 9.940089 | 10.02684 | 10.913 | 10.63596 | 10.28312 | 10.16937 | 10.08638 | 10.16616 |
| 7.593441 | 7.616253 | 7.77286 | 6.192687 | 6.156938 | 7.403491 | 7.384839 | 7.376304 | 7.044052 |
| 6.574452 | 6.859218 | 7.024464 | 6.822115 | 6.66712 | 6.238342 | 6.419474 | 6.224994 | 6.568088 |
| 6.828352 | 5.524611 | 6.318451 | 5.83685 | NaN | 5.787725 | 5.895977 | 5.740213 | 5.946648 |
| 6.360518 | 6.461825 | 6.974237 | 5.53651 | 4.396852 | 7.159697 | 7.420655 | 7.309568 | 6.355779 |
| 4.885982 | 5.508755 | 5.807523 | NaN | NaN | NaN | NaN | NaN | 5.390497 |
| 8.803517 | 8.530835 | 8.577173 | 9.023866 | 8.579771 | 8.492284 | 8.443542 | 8.164966 | 8.089962 |
| 7.164918 | 6.635873 | 7.1661 | 5.448543 | 5.400133 | 7.267805 | 6.937638 | 6.83602 | 6.809644 |
| 6.378429 | 6.244317 | 6.385914 | 6.362119 | 6.639872 | 7.530063 | NaN | 7.987195 | 6.044714 |
| 6.941969 | 7.047867 | 5.997744 | 8.151383 | 8.100204 | 7.306871 | 7.3632 | 7.095121 | 8.422232 |
| NaN | NaN | 2.908278 | NaN | NaN | 4.604884 | 4.749776 | 5.17072 | 4.743849 |
| 6.256362 | 5.282074 | 5.837126 | NaN | 3.933006 | 6.839444 | 6.986518 | 6.94302 | 6.261019 |
| 8.423493 | 8.441964 | 8.412681 | 7.68678 | 7.758903 | 8.398922 | 8.137563 | 7.727993 | 7.983677 |
| 7.897022 | 7.761431 | 7.870721 | 8.680702 | 9.074718 | 7.930118 | 7.904663 | 7.871584 | 7.439135 |
| 7.616325 | 7.557857 | 7.568431 | 7.444783 | 7.435999 | 6.541871 | 6.745161 | 6.294325 | 6.563953 |
| 7.053063 | 7.785278 | 7.116647 | 6.996934 | 6.933146 | 7.204627 | 7.310954 | 6.900155 | 6.51165 |
| 5.385929 | 4.612771 | 6.138617 | NaN | 4.085349 | 6.104314 | 6.421402 | 6.358355 | 5.206635 |
| 7.157093 | 5.320906 | 5.802468 | NaN | NaN | NaN | NaN | NaN | NaN |
| 9.022234 | 8.959646 | 9.118216 | 7.768399 | 8.120111 | 8.465732 | 8.1683 | 8.146439 | 5.413084 |
| 8.465952 | 8.024839 | 8.416146 | 6.79203 | 6.682789 | 7.985862 | 8.086643 | 8.119125 | 7.829892 |
| 9.033843 | 8.812726 | 8.969311 | 8.435394 | 8.209105 | 8.832619 | 8.846426 | 9.05889 | 8.418704 |

-Bruker Confidential-

| | | | | | | | | |
|---|---|---|---|---|---|---|---|---|
| 5.605998 | 5.805176 | 5.80432 | 5.518823 | 5.436182 | 5.982017 | 6.000812 | 5.450098 | 5.20437 |
| NaN | 4.283236 | NaN | NaN | NaN | NaN | NaN | NaN | 2.467758 |
| 7.21405 | 7.286229 | 7.185702 | 6.944628 | 6.930399 | 6.918229 | 6.720305 | 7.233748 | 7.162352 |
| NaN | 7.573677 | 7.832407 | 5.974688 | 6.597804 | 7.240657 | NaN | 7.647315 | 7.878973 |
| 8.801561 | 8.912137 | 9.391876 | 8.118785 | 8.561733 | 8.178708 | NaN | 8.618805 | 7.688996 |
| 7.563024 | 7.5229 | 7.364422 | 4.889967 | 5.48349 | 7.171249 | 7.215976 | 7.296165 | 6.79383 |
| 9.456359 | 9.484134 | 9.320058 | 9.615214 | 9.634805 | 8.603149 | 7.595725 | 8.77831 | 8.310875 |
| NaN | NaN | NaN | NaN | NaN | 5.120546 | 4.81381 | 5.474246 | 4.935067 |
| 5.859922 | 5.469061 | 6.271111 | NaN | NaN | 6.191749 | 6.253213 | 6.282871 | 5.065704 |
| 9.724074 | 9.656829 | 9.640461 | 8.966526 | 8.881673 | 9.733622 | 9.531693 | 9.730022 | 8.609822 |
| 6.8996 | 6.829236 | 6.894013 | 5.699255 | 5.957378 | 6.877918 | 6.791208 | 7.040691 | 6.930998 |
| 6.577655 | 6.282217 | 6.534938 | 7.015334 | 6.377747 | 6.480822 | 7.010247 | 7.233311 | 6.042698 |
| 5.905746 | 6.220333 | 6.442774 | 5.564912 | 5.774815 | 5.449739 | 5.241747 | 5.395666 | 6.036714 |
| 7.341393 | 6.686265 | 7.473231 | 6.4959 | 5.456469 | 7.513651 | 7.268835 | 7.258566 | 4.870979 |
| 7.500726 | 6.678504 | 7.511484 | 6.794878 | 6.733307 | 7.451166 | 7.284276 | 7.420266 | 6.01241 |
| 7.467095 | 7.486157 | 7.689058 | 6.277956 | 6.612794 | 7.76834 | 7.91653 | 7.967795 | 7.712269 |
| 6.793238 | 6.429196 | 5.992674 | 5.119725 | NaN | 6.230685 | 6.170928 | 6.62487 | 6.226432 |
| 7.790168 | 7.550603 | 7.658555 | 7.005969 | 7.048785 | 7.762571 | 7.758499 | 7.494416 | 7.69059 |
| 4.960464 | 5.992882 | 5.340426 | 3.636886 | 4.308581 | NaN | NaN | NaN | 5.67295 |
| 7.178264 | 5.026928 | 4.478167 | NaN | NaN | NaN | 6.429365 | 6.463853 | 6.835013 |
| 7.595208 | 7.361761 | 7.252587 | 5.383193 | 6.295838 | 7.195868 | 7.075487 | 6.828057 | 6.7068 |
| 8.616828 | 8.505135 | 8.46785 | 7.843932 | 7.88678 | 7.941216 | 7.481441 | 7.804925 | 7.567203 |
| 8.287718 | 8.292449 | 8.241853 | 7.534057 | 7.450097 | 8.498762 | 8.559061 | 8.547995 | 8.588211 |
| 6.423431 | 6.23405 | 6.417897 | 7.847791 | 7.745981 | 5.709312 | 5.686282 | 5.99398 | 6.141656 |
| 8.068293 | 7.504746 | 7.871329 | 6.715688 | 6.513676 | 7.743558 | 7.446566 | 7.49825 | 7.698097 |
| 7.922884 | 6.978337 | 7.072521 | 7.483623 | 7.196708 | 7.132022 | 7.145402 | 6.709091 | NaN |
| 8.33849 | 7.803786 | 7.308347 | 8.225598 | 7.977635 | 8.194221 | 8.78972 | 8.524161 | 7.130924 |
| NaN | 7.17722 | NaN | 6.46156 | 6.172678 | NaN | NaN | 5.888076 | 4.868567 |
| NaN | 6.689836 | 5.805071 | NaN | NaN | NaN | 6.323182 | NaN | 6.02953 |
| 8.117023 | 8.380305 | 8.216556 | 9.482248 | 9.301332 | 8.597463 | 8.544066 | 8.626069 | 7.98937 |
| 7.816329 | 8.253497 | 8.031258 | 6.485978 | 6.261057 | 7.754612 | 7.467463 | 7.664753 | 7.92439 |
| 10.44473 | 10.52264 | 10.36837 | 10.4781 | 10.55173 | 10.92449 | 10.77155 | 10.94503 | 10.52632 |
| 7.615957 | 6.445475 | 7.32526 | 4.30477 | 3.65607 | 6.350037 | 6.15701 | 6.055247 | 6.849702 |
| NaN | NaN | 7.533031 | 8.328372 | 7.741827 | 8.745168 | 9.012642 | 6.974996 | 7.038561 |
| 6.572249 | 6.728816 | NaN | 5.518745 | 5.890491 | NaN | NaN | NaN | 8.491305 |
| 6.861042 | 6.970451 | 6.878158 | 6.52348 | 6.483042 | 7.422507 | 7.597854 | 7.649924 | 7.791008 |
| 5.82684 | 6.035934 | 6.289421 | 6.40842 | 6.618764 | 5.435026 | 5.864187 | 5.892602 | 7.297997 |
| 5.849557 | 6.708253 | 6.569664 | 6.955644 | 7.50648 | 7.517474 | 7.506084 | 7.348061 | 7.385845 |
| 8.741347 | 8.485086 | 8.633017 | 9.66216 | 9.733248 | 7.873102 | 8.162296 | 8.378999 | 8.406532 |
| 9.576859 | 9.272655 | 9.453069 | 9.350714 | 9.159056 | 8.942506 | 8.462001 | 8.847578 | 8.966793 |
| 6.827353 | 6.250885 | 6.730833 | 5.441037 | 5.376781 | 6.405293 | 6.497481 | 6.671586 | 6.284834 |
| 7.628308 | 7.664473 | 7.789876 | 9.699059 | 9.02127 | 7.06216 | 7.649068 | 7.73177 | 7.189287 |
| 7.748159 | 6.955581 | 7.451473 | 6.501167 | 6.974944 | 7.988039 | 7.380584 | 7.80848 | 7.834944 |
| 9.740148 | 9.701785 | 9.864477 | 9.748445 | 9.416569 | 9.46871 | 9.667862 | 9.699525 | 10.04546 |
| 7.354808 | 7.528786 | 7.247764 | 5.957268 | 6.582862 | 7.18256 | 6.928588 | 7.27788 | 6.441544 |
| 6.936397 | 7.282316 | 7.134537 | 6.920776 | 7.211742 | 7.63882 | 7.590249 | 7.428956 | 7.119141 |
| 7.341198 | 7.437823 | 7.451888 | 6.911545 | 6.896184 | 7.092293 | 7.259787 | 7.066511 | 6.790988 |

| | | | | | | | | |
|---|---|---|---|---|---|---|---|---|
| 6.757217 | 5.312118 | 8.004475 | NaN | 5.144182 | 6.701746 | 7.189835 | 7.270102 | 6.597697 |
| 10.63727 | 10.52662 | 10.59109 | 11.4729 | 11.75902 | 9.797536 | 9.907318 | 9.910213 | 10.21732 |
| 8.0326 | 7.612325 | 7.767796 | 9.948636 | 9.192855 | 7.841787 | 7.975024 | 7.78929 | 8.237509 |
| 7.868146 | 7.880038 | 7.05871 | 8.75201 | 9.028454 | 7.28613 | 7.11237 | 7.617429 | 7.369563 |
| 6.378221 | 6.334137 | 5.380569 | 6.95696 | 7.055942 | NaN | NaN | NaN | NaN |
| 7.436752 | 7.002616 | 7.459536 | 6.655615 | 6.117263 | 6.836675 | 6.865031 | 6.256326 | 7.054118 |
| 7.840597 | 7.738331 | 8.378037 | 7.394831 | 7.981303 | 8.445594 | 8.431683 | 8.329234 | 8.349162 |
| NaN | NaN | NaN | 4.260115 | NaN | NaN | NaN | 4.382783 | 5.898607 |
| 7.562041 | 7.711525 | NaN | NaN | 6.176578 | NaN | 6.634444 | 6.776201 | 7.470429 |
| NaN | NaN | NaN | 5.643792 | 5.886563 | NaN | NaN | NaN | 5.761369 |
| NaN | NaN | NaN | 4.254199 | 5.155385 | NaN | NaN | NaN | NaN |
| 6.735399 | 6.648516 | 6.245807 | 5.439665 | 6.330331 | 6.336789 | 6.627011 | 6.450363 | 6.419914 |
| 8.465431 | 8.427027 | 8.427465 | 8.882197 | 8.771737 | 8.06316 | 7.75952 | 8.161135 | 7.568692 |
| 6.333971 | 6.689105 | 6.163749 | NaN | NaN | 6.26248 | NaN | NaN | 6.272317 |
| 4.66384 | 3.54097 | 4.411657 | 3.097139 | 1.940101 | 3.804383 | 4.611424 | 3.571421 | 5.644205 |
| 8.782668 | 8.540263 | 8.298609 | 9.652752 | 9.636708 | 8.893904 | 9.143174 | 9.087435 | 9.755911 |
| 6.347799 | 6.319704 | 6.283778 | 5.580546 | 6.075385 | 5.889077 | 5.990465 | 6.244692 | 5.751423 |
| 6.74797 | 7.122026 | 6.787121 | 7.320411 | 7.597412 | 6.454476 | 6.205903 | 6.403928 | 7.383896 |
| 8.199472 | 8.189647 | 8.298525 | 8.549915 | 8.609768 | 7.304236 | 6.406087 | 7.146402 | 6.938169 |
| 8.046167 | 8.169731 | 8.238115 | 8.789808 | 8.747133 | 7.474841 | 7.833388 | 7.960244 | 7.813052 |
| 6.269523 | 5.916277 | 5.950364 | 7.017898 | 6.801787 | 6.053263 | 5.71094 | 6.196903 | 5.472751 |
| 7.425493 | 7.423261 | 7.26147 | 6.219619 | 6.364535 | 7.697194 | 7.627437 | 7.622076 | 7.439374 |
| NaN | 6.255847 | NaN | NaN | NaN | 6.297556 | 6.939645 | 6.686953 | 6.164439 |
| 9.151571 | 9.123416 | 9.104695 | 8.791257 | 8.732486 | 9.543889 | 9.513822 | 9.538835 | 9.575492 |
| 8.853082 | 8.690654 | 8.580334 | 9.884202 | 10.09443 | 8.604353 | 8.972617 | 8.873251 | 8.92889 |
| 8.011787 | 7.944417 | 8.099751 | 9.586767 | 9.58263 | 7.630861 | 8.133756 | 7.864422 | 8.388321 |
| NaN | 7.314317 | 7.340767 | 7.71155 | 6.991207 | 7.668797 | 7.532285 | 7.314758 | 7.493667 |
| 8.190891 | 8.097507 | 7.984645 | 7.518 | 7.643801 | 7.945323 | 7.854894 | 8.100391 | 6.606096 |
| NaN | NaN | NaN | NaN | NaN | NaN | NaN | NaN | 8.034503 |
| 5.73645 | 4.994446 | 6.031622 | 4.567464 | 4.272105 | 6.192644 | 6.212627 | 6.268626 | 6.10496 |
| 6.807921 | 6.984398 | 6.770977 | 6.77197 | 6.910909 | 7.163896 | 6.849272 | 7.548739 | 7.386786 |
| 5.504615 | 4.995958 | 5.663131 | 5.840929 | 6.33512 | 6.39774 | 6.043863 | 6.011958 | 5.819096 |
| 5.259746 | 5.59236 | 4.203185 | 6.287217 | 6.838759 | NaN | NaN | NaN | NaN |
| NaN | 3.408918 | NaN | 4.819881 | 5.035775 | 3.390732 | NaN | NaN | 2.692785 |
| 10.82526 | 10.69105 | 10.64077 | 11.80185 | 11.58113 | 10.77917 | 10.37383 | 10.50073 | 10.33551 |
| 5.471533 | 5.503832 | 6.13724 | 6.254406 | 5.889879 | 6.818999 | 5.917208 | 6.602532 | 6.39056 |
| 7.262213 | 6.801691 | 6.80479 | 6.793363 | 6.373227 | 6.920891 | 6.582414 | 7.090011 | 6.338435 |
| 9.49641 | 9.395555 | 9.519807 | 9.527984 | 9.521806 | 9.403932 | 9.268133 | 9.191009 | 9.260097 |
| 4.701708 | 5.653743 | 5.448062 | 6.288929 | 5.788974 | 5.349901 | 5.224581 | 4.932638 | 4.622718 |
| 6.565284 | 6.196812 | 6.822842 | 5.963412 | 5.865293 | 6.767728 | 6.860754 | 7.296649 | 6.373168 |
| 9.421352 | 9.682195 | 9.874674 | 10.13917 | 9.56262 | 8.8136 | 8.593035 | 8.56511 | 9.348782 |
| 7.842319 | 7.774991 | 7.918348 | 7.169095 | 7.174447 | 8.575891 | 8.065356 | 7.848305 | 7.085831 |
| 6.172037 | 4.561479 | 5.675476 | 6.254664 | 5.994522 | NaN | 4.351637 | NaN | 5.308803 |
| 6.092336 | 4.846853 | 4.547807 | 5.233692 | 5.000192 | NaN | NaN | NaN | 6.236115 |
| 7.743526 | 7.704593 | 8.50589 | 5.185093 | 5.540138 | 7.73977 | 7.676348 | 7.363686 | 7.384603 |
| 8.465786 | 6.497573 | 8.33885 | 5.261746 | 5.881005 | 6.881001 | 6.458937 | 6.985136 | 6.05431 |
| 7.182326 | 7.22016 | 5.981999 | NaN | 5.566905 | 7.451457 | 7.404445 | 7.498972 | 6.177387 |

| | | | | | | | | |
|---|---|---|---|---|---|---|---|---|
| NaN | 2.195314 | NaN | 2.73784 | 3.290652 | NaN | NaN | 4.511761 | 3.898905 |
| 9.386959 | 9.408719 | 9.310081 | 8.768364 | 8.649751 | 9.367196 | 9.135224 | 9.14837 | 9.397537 |
| 8.738802 | 8.711071 | 8.801082 | 9.154299 | 9.350683 | 8.585838 | 8.715011 | 8.435096 | 7.598024 |
| 7.396461 | 8.005087 | 7.71259 | 7.505063 | 7.220106 | 6.84547 | 7.145803 | 6.881362 | 7.030716 |
| 8.665151 | 8.501823 | 8.601703 | 7.539944 | 8.011489 | 8.545443 | 8.808745 | 8.628331 | 7.832326 |
| 7.6141 | 8.047598 | 7.532702 | 8.296585 | 8.344169 | 6.91866 | 8.058391 | NaN | 7.282488 |
| 7.049691 | 5.835157 | 7.902875 | 7.23515 | 7.314215 | 6.902763 | 6.885335 | 6.814471 | 7.362738 |
| 6.004415 | 5.911477 | 5.819764 | 6.273983 | 4.123651 | 6.458239 | 6.440425 | 6.587773 | 6.561889 |
| NaN | NaN | NaN | 4.482296 | 4.799973 | NaN | 5.741529 | NaN | 6.631196 |
| 6.610704 | 6.119585 | 6.456547 | 5.004934 | 5.904369 | 6.847221 | 6.423079 | 6.925354 | 5.689987 |
| 9.256572 | 9.330904 | 9.375931 | 9.844059 | 9.474165 | 8.166143 | 8.014157 | 8.307393 | 7.831547 |
| 8.643672 | 8.524646 | 8.620795 | 7.699737 | 7.589659 | 8.40938 | 8.514524 | 8.579721 | 8.77837 |
| 8.973698 | 9.027061 | 8.993077 | 9.529019 | 9.415372 | 8.62992 | 8.152086 | 8.644432 | 9.050584 |
| 4.310941 | 4.204282 | 3.165065 | 4.723883 | 4.863244 | 3.53907 | 3.756686 | 3.577355 | 4.560482 |
| 8.46374 | 8.470302 | 8.510877 | 8.424241 | 8.460495 | 8.283399 | 8.187326 | 8.586317 | 8.586344 |
| 9.505937 | 9.526266 | 9.454851 | 9.824109 | 9.901654 | 9.475533 | 9.518355 | 9.622898 | 9.329403 |
| 10.85372 | 10.67531 | 10.7738 | 11.58908 | 11.6449 | 9.853637 | 10.21719 | 9.938783 | 10.97655 |
| 6.532233 | 6.517378 | 6.520664 | 5.509505 | 5.994682 | 6.437312 | 7.181015 | 6.6714 | 6.606797 |
| 6.248971 | 6.423895 | 6.88231 | NaN | NaN | 6.091216 | 6.002129 | 6.269706 | 5.09415 |
| NaN | 4.664056 | 4.699566 | 4.527776 | NaN | 5.809978 | 7.354464 | 7.842631 | 6.738946 |
| 5.736415 | 5.515319 | 6.006563 | 5.554682 | 5.587381 | 6.154009 | 6.080501 | 6.195326 | 5.382401 |
| 7.305723 | 7.157609 | 7.125878 | 7.815844 | 8.045473 | 6.859458 | 6.195447 | 6.369098 | 7.270453 |
| 6.523823 | 6.82527 | 4.631928 | NaN | NaN | 7.158947 | 6.510455 | 6.910424 | 5.845774 |
| 5.976485 | 5.946784 | 6.132007 | NaN | NaN | NaN | 6.941332 | 6.521112 | 6.281739 |
| 7.254137 | 7.55791 | 7.089367 | 5.392759 | 5.755887 | 7.694261 | 7.729486 | 7.579937 | 6.686732 |
| 6.81356 | 6.724884 | 6.680354 | 6.707014 | 6.910061 | 7.444004 | 7.206288 | 7.273739 | 6.434041 |
| 8.472144 | 8.25849 | 8.396335 | 7.894099 | 7.414285 | 7.855688 | 7.819393 | 8.108733 | 8.220202 |
| 4.819246 | 4.508165 | 5.149328 | 5.794686 | 5.671607 | 5.376109 | 5.125586 | 5.195133 | 4.511343 |
| NaN | 4.120176 | 4.254275 | 5.147861 | 6.873952 | 4.345877 | 2.883682 | 4.538593 | 5.771054 |
| 7.655262 | 7.68247 | 8.043935 | 8.68158 | 8.317896 | 7.792639 | 7.737694 | 8.11189 | 7.882858 |
| 7.625013 | 7.46382 | 7.170348 | 6.211814 | 6.097349 | 6.83952 | 6.947841 | 7.281547 | 6.930041 |
| 7.220673 | 8.05059 | 7.203758 | 6.30398 | 6.326104 | 7.604203 | 7.384462 | 7.367741 | 6.580068 |
| 8.435238 | 8.218032 | 8.464584 | 8.419954 | 8.566684 | 8.182664 | 8.133968 | 8.315179 | 7.740145 |
| 8.87698 | 8.681387 | 8.490347 | 8.546023 | 8.507345 | 8.472941 | 8.516782 | 8.47494 | 8.904744 |
| NaN | 7.532185 | 8.934793 | 8.387961 | 8.021662 | NaN | NaN | 6.473332 | 8.426888 |
| 8.297139 | 8.191025 | 8.223824 | 9.4284 | 9.270028 | 7.61554 | 7.597762 | 7.577083 | 8.07326 |
| 7.3702 | 7.295193 | 7.587167 | 8.503249 | 8.336889 | 7.510429 | 6.982004 | 7.75594 | 7.452903 |
| 7.394051 | 7.733402 | 7.828891 | 8.574344 | 8.133596 | 7.993227 | 7.704809 | 7.722964 | 7.587543 |
| 11.06295 | 11.2254 | 11.20416 | 11.51999 | 11.75952 | 10.58218 | 10.68228 | 10.29369 | 10.38684 |
| 8.687078 | 8.540858 | 8.720341 | 7.643063 | 7.398037 | 8.430616 | 8.48616 | 8.332706 | 8.416218 |
| 5.656922 | 6.617378 | 5.242991 | 5.224423 | 4.684646 | 5.087705 | 5.417328 | 5.77821 | 5.823195 |
| 5.997061 | 5.677521 | 5.133338 | 6.793742 | 8.142256 | 6.942327 | 6.81536 | 6.797607 | 5.781727 |
| 10.24365 | 10.51186 | 10.46176 | 10.2915 | 10.37078 | 10.34288 | 10.66195 | 10.416 | 9.39938 |
| 7.678063 | 7.011122 | 7.411791 | 5.797656 | 5.58119 | 6.195499 | 6.732378 | 6.7364 | 5.65242 |
| NaN | NaN | NaN | 5.732668 | 6.296734 | NaN | NaN | NaN | 5.907067 |
| 7.934341 | 7.916853 | 7.854001 | 6.93154 | 7.25837 | 7.941559 | 8.144982 | 8.165483 | 7.592004 |
| 6.758849 | 6.871295 | 6.835014 | 7.625411 | 7.463708 | 7.404397 | 7.153425 | 6.802585 | 7.631964 |

| | | | | | | | | |
|---|---|---|---|---|---|---|---|---|
| NaN | NaN | NaN | NaN | NaN | 4.785145 | NaN | NaN | 5.451326 |
| 9.422807 | 9.17471 | 9.225109 | 6.440485 | 6.814731 | 8.478468 | 8.694691 | 8.40484 | 9.072566 |
| 7.88214 | 8.076015 | 7.699234 | 5.763618 | 6.692485 | 8.397565 | 8.604693 | 8.844189 | 7.756262 |
| 2.39152 | 1.326909 | 2.861032 | 2.373728 | 0.032145 | 1.050211 | NaN | 2.79109 | 1.57969 |
| 11.30537 | 11.2517 | 11.23209 | 12.47458 | 12.42166 | 11.35779 | 11.34829 | 11.60264 | 10.53277 |
| 8.844961 | 8.815339 | 8.880334 | 7.537038 | 7.629886 | 8.853183 | 8.993807 | 9.149631 | 8.387983 |
| 5.942411 | 4.696033 | 4.44193 | NaN | 5.521958 | 5.356496 | 5.303566 | 5.301631 | 5.600848 |
| 7.172992 | 7.380412 | 7.551544 | 7.930596 | 7.643684 | 6.350205 | 6.243104 | 7.137839 | 7.498549 |
| 6.79341 | 6.503071 | 6.278091 | 7.78303 | 8.182487 | NaN | NaN | NaN | NaN |
| 6.728035 | 6.416278 | 6.856307 | 7.644472 | 7.681107 | 6.60765 | 6.921022 | 7.171532 | NaN |
| 8.257977 | 8.058879 | 8.339336 | 8.788947 | 8.886706 | 7.286642 | 7.450037 | 7.67892 | 8.158583 |
| 9.807675 | 9.691236 | 9.635839 | 8.922454 | 8.752279 | 9.907331 | 9.906382 | 9.927756 | 9.547295 |
| 8.551729 | 8.384637 | 8.652215 | 7.402171 | 7.485939 | 8.462774 | 8.552317 | 8.718266 | 8.535047 |
| 5.324293 | 5.814279 | 6.049809 | 6.585154 | 6.314917 | NaN | 4.480149 | NaN | 6.470423 |
| 7.119639 | 6.952857 | 6.390166 | 5.96262 | 6.318852 | 6.539343 | 6.271487 | 6.830119 | 6.027617 |
| 8.123086 | 8.097515 | 8.099105 | 8.023834 | 7.836839 | 8.185382 | 8.24141 | 8.312261 | 7.959682 |
| 6.553297 | 6.212754 | 7.403477 | 7.383269 | 6.923599 | 6.339121 | 6.446106 | 5.581281 | 5.550147 |
| 6.717006 | 7.028439 | 7.000767 | 6.642338 | 6.437214 | 7.19391 | 7.436166 | 7.37031 | 6.893265 |
| 7.552884 | 6.707481 | 7.367582 | 6.255025 | 6.315288 | 7.478764 | 6.720418 | 7.757325 | 7.005231 |
| 8.915772 | 9.111474 | 9.28343 | 9.107665 | 8.604926 | 8.061715 | 8.60763 | 7.866529 | 8.307488 |
| 7.049107 | 7.052258 | 6.764694 | 6.922041 | 7.059066 | 7.480914 | 7.352599 | 7.509756 | 6.188713 |
| 7.493015 | 7.365817 | 7.270939 | 5.007051 | 6.69 | 7.707991 | 7.451775 | 7.617069 | 6.533834 |
| 10.23367 | 10.24981 | 8.332989 | 9.464545 | 9.761715 | 10.08053 | 9.962124 | 9.936178 | 9.397403 |
| 6.435327 | 6.494454 | 7.096363 | NaN | NaN | 7.882337 | 8.034895 | 6.810693 | 6.08804 |
| 8.358963 | 8.451529 | 8.432759 | 7.580182 | 7.929712 | 8.511321 | 8.441022 | 8.393449 | 8.529735 |
| 8.004923 | 7.87773 | 7.994653 | 7.711618 | 6.041901 | 7.567496 | 7.626636 | 7.597683 | 7.198078 |
| 5.976312 | 6.191738 | 6.661273 | 5.171881 | 5.228389 | 6.618147 | 6.635299 | 6.335759 | 5.699772 |
| 9.045075 | 9.216307 | 8.940609 | 7.510228 | 7.8912 | 8.942962 | 9.027292 | 9.087088 | 8.673775 |
| 9.466565 | 9.440434 | 9.24211 | 9.603117 | 9.360574 | 9.541862 | 9.433293 | 9.607162 | 9.438044 |
| 7.237868 | 7.418275 | 7.128742 | 7.89321 | 7.903884 | 7.733991 | 7.307528 | 7.689458 | 7.140646 |
| NaN | NaN | NaN | 6.595144 | 6.539222 | 6.697683 | 6.131216 | NaN | 6.767667 |
| 8.753651 | 8.629924 | 8.619216 | 8.838626 | 8.951989 | 8.156542 | 8.072968 | 8.286922 | 8.76497 |
| 8.382576 | 8.418003 | 8.712039 | 7.085026 | 7.061176 | 7.409327 | 7.59633 | 7.51168 | 8.335534 |
| 7.443281 | 7.510206 | 7.178241 | 8.629629 | 8.772997 | 6.585987 | 5.759879 | 6.91841 | 6.178392 |
| 6.067148 | 5.700581 | 5.371943 | NaN | 4.363165 | 6.095047 | 5.987034 | 5.781409 | 5.983488 |
| 8.492353 | 7.932238 | 8.442889 | 7.550486 | 7.93172 | 7.711354 | 7.492727 | 7.58698 | 8.169648 |
| NaN | 5.415315 | NaN | NaN | NaN | 6.430301 | 6.388145 | 6.412377 | 5.192641 |
| 7.185206 | 7.167454 | 7.233685 | 8.116181 | 7.186357 | 7.339926 | 7.286684 | 7.148344 | 6.524711 |
| 7.183943 | 6.921786 | 5.781512 | 4.953532 | 5.597227 | 7.798842 | 7.955056 | 7.813607 | 6.859773 |
| 9.123996 | 9.072569 | 9.166791 | 10.46306 | 10.50331 | 9.376869 | 9.158274 | 9.427097 | 8.506448 |
| 6.176234 | 6.285991 | 5.988346 | 5.939225 | 5.833567 | 6.263206 | 6.066822 | 6.370121 | 6.679987 |
| 8.251315 | 8.287193 | 8.246804 | 9.047631 | 8.819002 | 8.134623 | 8.012861 | 8.207378 | 8.12883 |
| NaN | 7.032743 | NaN | 6.861921 | 8.42811 | 7.369678 | NaN | 7.692534 | 5.469441 |
| 5.539071 | 5.659179 | 6.139473 | NaN | NaN | NaN | NaN | NaN | 5.45816 |
| 9.537025 | 9.646404 | 9.798304 | 10.02502 | 8.364263 | 6.8494 | 6.940279 | 6.796196 | 7.389648 |
| NaN | 5.482635 | 3.669186 | NaN | NaN | 6.982503 | 4.805366 | 6.731273 | 4.676131 |
| 7.197827 | 6.803681 | 7.09581 | 8.249987 | 8.130581 | 7.313165 | 7.353616 | 7.141018 | 7.400232 |

| | | | | | | | | |
|---|---|---|---|---|---|---|---|---|
| 8.205159 | 8.247195 | 8.23957 | 8.990396 | 9.276486 | 7.643844 | 7.506695 | 7.715429 | 7.353327 |
| 6.454897 | 6.378323 | 6.60109 | 5.178106 | 4.611161 | 5.709207 | 5.718535 | 6.04884 | 5.764482 |
| 7.679643 | 6.92415 | 7.469021 | 7.603921 | 7.47113 | 7.605277 | 7.653752 | 7.716297 | 7.750295 |
| 6.38302 | 6.635377 | 6.707395 | 5.447724 | 5.794101 | 6.669102 | 6.59461 | 6.551737 | 6.445181 |
| 7.227612 | 7.331191 | 7.44361 | 6.767027 | 6.645959 | 6.86494 | 6.924971 | 6.695416 | 6.618851 |
| 8.637671 | 8.407809 | 8.559925 | 8.220769 | 8.208963 | 8.415732 | 8.468491 | 8.224319 | 8.332158 |
| 6.697414 | 5.599523 | 6.739683 | 5.8777 | 5.516567 | 6.622204 | 7.018867 | 6.813473 | 6.386563 |
| 7.765275 | 7.610559 | 7.398394 | 7.410301 | 7.354633 | 7.656499 | 7.532047 | 7.56468 | 8.008208 |
| 5.997265 | 6.932548 | 6.080164 | 6.679564 | 6.724492 | NaN | NaN | 7.036602 | 5.836773 |
| 4.619264 | 4.130609 | 5.324252 | 3.305425 | NaN | 3.921439 | NaN | NaN | 4.261807 |
| 7.113317 | 6.842648 | 6.861607 | 7.337267 | 7.484557 | 7.417088 | 7.580688 | 7.042241 | 6.948187 |
| 6.429836 | 6.729151 | 6.438002 | 7.252396 | 5.82563 | 6.101737 | 5.966508 | 6.446237 | 6.528436 |
| 5.410837 | 5.670669 | 5.609838 | 6.228129 | 6.694494 | 5.577307 | 4.594994 | 5.63073 | 4.753754 |
| 8.742103 | 8.715183 | 8.622089 | 7.17572 | 7.295732 | 8.816554 | 8.678425 | 8.805256 | 8.483656 |
| 6.634684 | 6.888541 | 6.541055 | 6.176606 | 6.794527 | 6.699792 | 6.630544 | 6.805237 | 6.104515 |
| 8.754431 | 9.084302 | 8.954823 | 8.341265 | 7.984893 | 9.099224 | 8.994193 | 8.537578 | 8.635917 |
| 7.53723 | 7.401899 | 7.887726 | 6.306505 | 7.505013 | 7.679003 | 8.15461 | 8.043208 | 7.985023 |
| 10.7169 | 10.55925 | 10.68514 | 10.83597 | 11.08693 | 11.42009 | 11.23253 | 11.34589 | 10.67874 |
| 7.12639 | 7.443686 | 7.27278 | 7.629236 | 7.544492 | NaN | NaN | NaN | NaN |
| 6.548218 | 4.948853 | 5.331906 | NaN | NaN | 6.497159 | 6.152434 | 6.033267 | 5.905494 |
| 6.691222 | 6.570521 | 6.2363 | 6.273726 | 6.835235 | 7.389432 | 7.478192 | 7.449807 | 7.938272 |
| 8.019312 | 8.197478 | 8.177069 | 7.172764 | 7.064603 | 8.199724 | 8.505831 | 8.708045 | 7.746943 |
| 7.18591 | 8.322918 | 8.043597 | 7.547698 | 7.49046 | 8.56622 | 7.660784 | 7.657573 | 7.695261 |
| 8.231138 | 8.213724 | 8.325605 | 6.704059 | 7.01693 | 8.088658 | 7.643256 | 8.103987 | 7.404932 |
| 9.587568 | 9.554222 | 9.572714 | 8.902103 | 9.150336 | 9.549216 | 9.591632 | 9.238407 | 9.293792 |
| 5.598617 | 4.304006 | 5.207055 | 5.179996 | 5.386416 | 6.170048 | 5.447533 | 5.190269 | 5.573733 |
| 7.00618 | 6.221697 | 7.231995 | 4.722456 | 5.982636 | 7.33408 | 6.973711 | 7.153504 | 7.143771 |
| 9.081105 | 9.032276 | 9.170974 | 9.41256 | 9.456537 | 9.343254 | 9.278543 | 9.382987 | 8.270591 |
| 8.088104 | 7.815948 | 7.585655 | 7.806833 | 7.646914 | 8.486792 | 8.29362 | 7.913819 | 8.128706 |
| 7.142978 | 6.818281 | 5.662713 | 5.704393 | 5.750761 | 5.781191 | 4.895024 | 4.762929 | 4.999914 |
| 6.651031 | 5.809674 | 6.632763 | 7.093951 | 5.576602 | 6.811851 | 6.283142 | NaN | 5.883135 |
| 6.306518 | 6.123231 | 6.111361 | 6.544581 | 5.787986 | 6.693622 | NaN | 5.506515 | 5.56791 |
| 6.26029 | 5.699107 | 6.666611 | 5.072056 | 4.334211 | 5.933065 | 5.259498 | 5.500578 | 6.747985 |
| 8.70947 | 8.646365 | 8.516913 | 8.093904 | 7.877616 | 8.251431 | 8.311674 | 8.184804 | 8.195354 |
| 5.559681 | 6.153228 | 6.169713 | 4.841826 | 5.224402 | 5.1672 | 5.164101 | 5.468546 | 5.236797 |
| 8.014035 | 7.928171 | 7.568127 | 7.074944 | 6.36569 | 5.796096 | 7.195229 | 5.988933 | 6.648315 |
| 7.188021 | 7.203692 | 7.367324 | 8.815011 | 8.790736 | 7.637464 | 7.291226 | 7.363242 | 7.332624 |
| 8.228689 | 7.624083 | 7.821945 | 8.256139 | 8.000589 | 7.744635 | 7.917739 | 7.797106 | 7.342724 |
| 5.850828 | 5.993545 | 5.431119 | NaN | NaN | 6.244076 | 6.289923 | 6.827925 | 6.560865 |
| 8.709611 | 8.263686 | 5.461165 | NaN | 6.896313 | 6.388474 | 8.529554 | NaN | 7.039366 |
| 5.449548 | 3.65722 | 3.70796 | NaN | 4.197574 | 6.194331 | 6.905872 | 6.679093 | 6.607091 |
| 5.070373 | 4.773041 | 5.183639 | 5.269247 | 6.228055 | NaN | NaN | NaN | 4.991307 |
| 7.203342 | 7.171515 | 8.205911 | NaN | NaN | 6.927741 | 7.302483 | 6.886779 | 7.229167 |
| 6.583723 | 6.052467 | 6.424942 | 6.089106 | 6.692341 | 7.722946 | 7.601292 | 7.902937 | 6.710839 |
| 7.230306 | 7.155331 | 6.961599 | 6.864803 | 6.776496 | 7.092011 | 7.158165 | 6.904837 | 6.984439 |
| 5.453369 | 5.703866 | 5.723377 | 4.862153 | 3.511385 | 5.748631 | 5.42003 | 5.02962 | 5.546167 |
| 8.411365 | 8.369328 | 8.072901 | 7.792684 | 7.846593 | 7.61217 | 7.371981 | 7.335232 | 7.770506 |

| | | | | | | | | |
|---|---|---|---|---|---|---|---|---|
| 6.489897 | 4.651027 | 7.692613 | 6.186573 | 6.26382 | NaN | NaN | NaN | 6.78377 |
| 6.586067 | 6.20201 | 6.254676 | 5.35009 | 4.529174 | 5.421242 | 5.626451 | 6.407442 | 6.036833 |
| 5.063956 | 5.660626 | 5.51392 | 5.259637 | 5.730155 | 6.776422 | 6.701365 | 6.716527 | 5.381441 |
| 3.921639 | 5.461791 | 5.758897 | 5.460742 | 5.218469 | 4.632385 | 4.033499 | 4.633759 | 4.619016 |
| 8.948386 | 8.951504 | 9.331373 | 10.64994 | 10.56169 | 9.266067 | 9.491505 | 9.656761 | 10.00338 |
| NaN | NaN | 4.378197 | 6.139047 | 5.949489 | 5.038865 | 5.550359 | 5.273945 | 5.287971 |
| 8.639403 | 8.68892 | 8.515518 | 8.765049 | 8.12156 | 7.735894 | 6.973458 | 7.118269 | 6.691292 |
| NaN | 5.77869 | 5.756012 | 6.856666 | 6.390887 | 5.423507 | 5.441134 | 4.965983 | 6.4135 |
| 7.102034 | 6.807477 | 6.909419 | 8.26242 | 8.313696 | 6.233663 | 6.439173 | 6.431635 | 5.966279 |
| 6.844463 | 6.699409 | 6.628467 | 7.342972 | 7.217261 | 7.815809 | 6.090221 | 7.426293 | 6.952178 |
| 6.379229 | 6.036798 | 6.161664 | NaN | NaN | 6.553428 | 6.345479 | 6.513399 | 5.3969 |
| 6.286379 | 6.863451 | 7.036711 | 6.041021 | 5.811902 | 6.463742 | 6.400688 | 6.547707 | 6.039685 |
| 7.275101 | 6.488125 | 6.911231 | 6.248059 | 5.521935 | 6.252252 | 5.712314 | 5.903421 | 6.321594 |
| 6.287884 | 7.009119 | 6.769058 | 5.741952 | 6.23257 | 6.621417 | 6.405894 | 6.605528 | 6.722542 |
| 5.347232 | 5.220203 | 5.279457 | 5.419694 | 5.188898 | 6.98217 | 5.94293 | 6.743391 | 5.616951 |
| 7.774915 | 7.696 | 7.837109 | 8.071994 | 8.030235 | 7.828452 | 7.57684 | 7.898277 | 7.527845 |
| 8.588617 | 8.543449 | 8.403456 | 7.419515 | 7.990223 | 7.521747 | 7.566317 | 7.319454 | 7.593102 |
| 7.703263 | 7.251566 | 7.214568 | 7.193536 | 8.139695 | 8.088669 | 7.950391 | 7.895617 | 8.005816 |
| 6.666035 | 7.418848 | 6.733724 | 5.908734 | 6.539752 | 6.53286 | 6.474329 | 6.232249 | 5.580136 |
| 7.613953 | 7.207932 | 7.104762 | 5.373013 | 6.896114 | 5.509821 | 5.886537 | 5.908215 | 6.943331 |
| 7.269724 | 7.344942 | 7.466517 | 7.072429 | 6.391441 | 7.896224 | 7.996136 | 7.675464 | 7.578578 |
| 7.718317 | 7.282647 | 7.589791 | 6.70994 | 6.910463 | 7.504912 | 7.16938 | 7.149168 | 7.138671 |
| 8.701604 | 9.33595 | 8.786947 | NaN | 7.210491 | 8.725044 | 8.432011 | 8.588806 | 7.532869 |
| 6.404433 | 5.685007 | 6.12944 | 6.749433 | 6.895845 | 7.210977 | 7.272954 | 7.362106 | 4.973573 |
| NaN | NaN | 8.289247 | NaN | NaN | NaN | NaN | NaN | 7.088871 |
| 7.840807 | 6.185409 | 6.411108 | NaN | 6.248054 | 8.053442 | 7.53006 | 8.17776 | 6.269863 |
| 7.093514 | 7.240597 | 7.21135 | 7.654534 | 7.869141 | 6.811961 | 7.308536 | 6.850423 | 7.234428 |
| 8.248914 | 8.408486 | 8.674745 | 7.361264 | 7.565646 | 8.8686 | 9.035744 | 9.174295 | 9.473751 |
| 6.098819 | 6.60369 | 5.899714 | 7.384986 | 8.648068 | 7.124619 | 7.101974 | 7.33545 | 5.847167 |
| 6.788918 | 6.977854 | 6.791359 | 7.586951 | 7.774169 | 7.288759 | 6.972858 | 7.055309 | 6.373138 |
| 7.765688 | 7.692451 | 7.644688 | 7.078286 | 4.481276 | 6.21422 | 6.281921 | 6.214611 | 5.876744 |
| 9.221519 | 9.255761 | 9.112044 | 10.02106 | 9.907979 | 8.946604 | 8.832279 | 8.885462 | 8.458918 |
| 6.802849 | 6.696364 | 6.719853 | 6.745392 | 6.913575 | 7.56642 | 7.510761 | 7.277776 | 6.848183 |
| 8.688869 | 8.791065 | 8.836813 | 8.931103 | 8.489313 | 7.72786 | 7.715079 | 7.541111 | 6.957086 |
| 6.993032 | 7.153885 | 7.154147 | 9.324942 | 9.398802 | 5.158764 | 5.755967 | 5.533894 | 6.536114 |
| 8.036898 | 7.976951 | 8.019043 | 8.320726 | 8.10147 | 8.202292 | 7.874551 | 8.311899 | 8.476157 |
| 8.717783 | 9.173044 | 9.099163 | 9.177586 | 7.697989 | 7.664229 | 8.063956 | 7.410095 | 7.820731 |
| 10.89286 | 11.0136 | 11.34641 | 11.25302 | 11.0308 | 10.61501 | 10.44561 | 10.48008 | 10.45632 |
| NaN | NaN | NaN | 4.697326 | 5.771424 | 5.241318 | 3.895359 | 5.014823 | NaN |
| 7.165992 | 7.234839 | 7.419553 | 6.414904 | 6.386447 | 8.16231 | 7.653358 | 8.314658 | 7.080842 |
| 6.266838 | 5.954482 | 6.041138 | NaN | NaN | 7.723343 | 8.518817 | 8.097676 | NaN |
| 5.59306 | NaN | 6.072343 | 5.955154 | 5.792324 | 6.72605 | 5.444389 | 6.339413 | 5.546719 |
| 5.401883 | 4.955518 | 5.52237 | 5.302735 | 5.151558 | 6.022159 | NaN | 5.506158 | 5.880439 |
| 7.400221 | 7.278081 | 7.454782 | 6.984163 | 7.269322 | 8.195323 | 7.949169 | 8.014283 | 7.903584 |
| NaN | 4.446288 | NaN | 5.01749 | 4.967525 | 6.253531 | 6.435111 | 6.547798 | 6.022892 |
| 7.459122 | 7.041849 | 7.485416 | 6.280523 | 5.814976 | 7.844107 | 7.477332 | 7.809622 | 7.26643 |
| 8.983833 | 8.782069 | 8.978193 | 8.026743 | 7.837949 | 8.518042 | 8.380476 | 8.575942 | 8.333606 |

| | | | | | | | | |
|---|---|---|---|---|---|---|---|---|
| 6.700904 | 6.572495 | 6.532535 | 6.697253 | 6.516735 | 7.480223 | 7.065318 | 7.162092 | 7.564179 |
| 7.57478 | 7.445478 | 7.361324 | 6.917768 | 7.381908 | 7.266888 | 6.814729 | 7.051943 | 6.845795 |
| 6.089861 | 5.48409 | 5.944258 | 4.733036 | 4.884652 | 5.667434 | 5.994071 | 6.14549 | 6.274963 |
| 6.11969 | 6.582174 | 7.104447 | 5.653276 | 5.549017 | 6.109128 | 6.780001 | 6.088195 | 6.259367 |
| 5.285215 | NaN | 5.059692 | 5.838142 | 5.725457 | NaN | 4.938312 | NaN | 5.842306 |
| 8.011063 | 7.489174 | 7.397808 | 6.138095 | 6.224842 | 7.992187 | 8.080731 | 7.892362 | 7.782408 |
| 4.990993 | 4.98701 | 5.071868 | 4.690804 | 3.995604 | 4.141393 | 4.059262 | 4.444065 | 4.224083 |
| NaN | 7.568242 | 7.759587 | 7.309073 | 8.3257 | 8.188682 | 7.609707 | 7.946332 | 7.5693 |
| NaN | 2.611915 | NaN | 5.642998 | 6.396101 | NaN | NaN | NaN | 5.022173 |
| 7.417837 | 6.740961 | 7.062459 | 6.070124 | 6.388876 | 6.521877 | 6.526939 | 6.288368 | 6.460495 |
| 5.754188 | 5.966122 | 6.0796 | 5.664196 | 5.886884 | 5.596388 | 5.712847 | 5.778628 | 5.460222 |
| 7.079719 | 7.091035 | 7.144782 | 7.289862 | 7.232868 | 6.722683 | 6.725715 | 6.565645 | 6.742795 |
| 6.836014 | 7.652878 | 6.986526 | 7.831676 | 8.138847 | 7.857257 | 7.402834 | 7.808645 | 7.787193 |
| 6.70481 | 7.364959 | 4.943339 | 5.932265 | 6.359899 | 6.923338 | 6.449166 | 7.19266 | 6.233982 |
| 7.048868 | 6.846012 | 6.829787 | 6.60272 | 6.777149 | 6.704827 | 6.915412 | 6.693647 | 6.424299 |
| 7.490058 | 7.657593 | 7.968831 | NaN | NaN | 7.005647 | 5.993692 | 7.605762 | 6.444666 |
| 7.098923 | 7.31163 | 7.635068 | 6.427627 | 6.454113 | 7.247147 | 7.230569 | 7.235362 | 6.233979 |
| 7.799125 | 7.874807 | 7.775487 | 7.333026 | 7.535824 | 8.615475 | 8.743893 | 8.713787 | 7.337275 |
| 5.255203 | 5.313186 | 6.09009 | 5.99394 | 5.825761 | 6.779023 | 6.429586 | 6.551551 | 4.626949 |
| 6.484516 | 6.675866 | 6.364752 | 7.495579 | 7.368526 | 6.875045 | 6.787326 | 6.69482 | 6.842624 |
| 8.303369 | 6.88838 | 8.738323 | 7.127201 | 6.33193 | 7.429137 | 7.068642 | 8.433125 | 6.188777 |
| 7.180193 | 5.369198 | 6.426596 | 6.333186 | NaN | 7.246074 | 7.641135 | 7.721318 | 6.719334 |
| 8.824286 | 8.812525 | 8.931161 | 10.02869 | 9.775362 | 8.984813 | 9.099576 | 9.089237 | 9.301791 |
| 8.89516 | 8.745344 | 8.628334 | 10.06665 | 9.86242 | 8.580162 | 8.577902 | 8.854901 | 7.952107 |
| 9.363423 | 9.510613 | 9.254155 | 9.899195 | 9.952303 | 8.988679 | 8.8215 | 8.94667 | 9.117315 |
| 8.645292 | 8.104263 | 8.795436 | NaN | NaN | NaN | NaN | 5.703952 | 4.659719 |
| 7.236424 | 6.368476 | 6.302783 | 7.154024 | NaN | 7.363892 | 6.897191 | 7.11137 | 7.09224 |
| 7.360327 | 7.312924 | 7.301004 | 7.674201 | 7.938505 | 7.123358 | 6.977912 | 7.229928 | 6.735282 |
| 5.488814 | 3.858018 | 3.445704 | 7.352682 | 7.909987 | NaN | NaN | 4.573818 | 4.418405 |
| NaN | 4.938909 | 5.418871 | 3.765323 | NaN | 5.950982 | 6.710915 | 6.633749 | 6.440228 |
| 7.656375 | 7.561069 | 7.702095 | 6.697231 | 6.508853 | 7.702445 | 7.55236 | 7.669157 | 7.363657 |
| 7.120418 | 7.317024 | 7.110974 | 8.364834 | 8.440181 | 6.586561 | 6.760815 | 6.624966 | 7.945659 |
| 7.05472 | 6.879037 | 6.534424 | 5.550783 | 6.207207 | 7.381445 | 6.874122 | 7.25348 | 6.779901 |
| 7.651451 | 7.688308 | 7.660101 | 7.249979 | 7.342617 | 7.520092 | 7.548578 | 7.527676 | 7.562263 |
| 8.346873 | 8.439271 | 8.705789 | 7.517943 | 7.568116 | 8.016168 | 7.601805 | 7.952105 | 7.52314 |
| NaN | NaN | 7.018034 | 7.660163 | 7.495555 | 6.632677 | 6.600566 | 6.759088 | 7.543976 |
| 6.335959 | 6.585191 | 6.425198 | NaN | NaN | 6.751741 | 6.191804 | 6.404086 | 5.337597 |
| 9.405829 | 9.34907 | 9.377119 | 8.566113 | 8.515985 | 9.603557 | 9.684739 | 9.79636 | 9.702593 |
| 5.79616 | 5.699559 | 5.695994 | 5.162644 | 5.674798 | 5.291872 | 7.185416 | 6.412172 | 6.89976 |
| 6.450377 | 6.501844 | 6.889763 | NaN | NaN | 6.60032 | 6.629562 | 7.059628 | 6.449625 |
| 8.524965 | 8.210149 | 8.341882 | 6.28078 | 6.691279 | 7.79852 | 7.865447 | 7.655199 | 8.251898 |
| 7.522998 | 7.195003 | 7.561547 | 5.66268 | 5.866082 | 7.019892 | 7.081642 | 7.134971 | 7.771882 |
| 9.356036 | 9.269146 | 9.393577 | 8.020098 | 7.77274 | 8.899612 | 9.093542 | 8.709438 | 8.48335 |
| NaN | 5.007554 | 5.24944 | NaN | NaN | NaN | 5.330535 | 5.998378 | 5.509896 |
| 6.596504 | 6.776294 | 6.394199 | 6.641443 | 6.506554 | 6.973875 | 6.286685 | 6.401711 | 6.719058 |
| 8.375729 | 8.442882 | 8.350568 | 8.150987 | 8.131185 | 8.231824 | 8.05387 | 8.089741 | 6.807172 |
| 6.338637 | 5.428034 | 6.177386 | 6.154355 | 6.894639 | 6.590765 | 6.802002 | 6.753242 | 5.59449 |

| | | | | | | | | |
|---|---|---|---|---|---|---|---|---|
| 8.799809 | 8.805316 | 7.543301 | 7.443982 | 8.008102 | 7.78178 | 8.056918 | 8.171707 | 6.865128 |
| 6.205276 | 6.923146 | 6.452582 | 6.399919 | 6.320834 | 5.693861 | 5.813699 | 5.526196 | 5.37289 |
| 6.981373 | 6.664922 | 7.168216 | 7.340743 | 5.354792 | 5.682025 | 6.915764 | NaN | 8.864503 |
| 7.480735 | 7.155656 | 7.156794 | 8.409027 | 8.252924 | 7.994715 | 7.423517 | 7.806177 | 5.160912 |
| 7.18445 | 7.08938 | 7.334859 | 5.561043 | 5.659029 | 6.904939 | 7.171154 | 6.801095 | 5.763335 |
| 6.957913 | 6.894857 | 6.648565 | 6.606817 | 6.580999 | 7.598367 | 7.373224 | 7.46569 | 6.598135 |
| 5.444037 | 6.124704 | 6.242885 | NaN | NaN | NaN | NaN | NaN | 5.64099 |
| 7.387034 | 5.778053 | 6.954346 | 6.840333 | 6.622658 | 8.304199 | 8.253838 | 8.594524 | 7.099125 |
| 6.982489 | 6.661899 | 6.656998 | 7.463724 | 5.385679 | 7.386652 | 7.106928 | 7.076712 | 7.728605 |
| 7.265471 | 7.282154 | 7.232868 | NaN | 6.170601 | NaN | NaN | NaN | NaN |
| 7.684159 | 7.03809 | 7.246437 | NaN | 7.232107 | 6.805945 | 6.84267 | 6.679917 | 7.421685 |
| 11.73472 | 11.60884 | 11.57338 | 11.52796 | 11.49022 | 10.57457 | 10.36127 | 10.52043 | 11.95758 |
| 6.464487 | 6.006933 | 6.333865 | 5.822669 | 5.883709 | 7.395654 | 6.589969 | 7.225991 | 6.551819 |
| 5.386985 | 5.140336 | 5.353453 | 4.480648 | NaN | 5.289855 | 5.04408 | 5.233983 | 5.159104 |
| 8.475184 | 8.45118 | 8.315866 | 7.435619 | 7.550986 | 8.17677 | 7.478996 | 8.027604 | 7.688065 |
| 8.367532 | 7.07202 | 7.93552 | 7.548607 | 7.851292 | NaN | 6.920801 | 7.507823 | 7.742176 |
| 7.596361 | 7.663149 | 7.593993 | 6.584709 | 6.727671 | 8.580948 | 8.127306 | 8.061431 | 7.528237 |
| 8.936451 | 8.544589 | 8.238775 | 8.643602 | 8.250636 | 8.010428 | 8.023802 | 8.481778 | 8.621185 |
| 7.56192 | 6.203196 | 7.085672 | 6.134247 | 5.551435 | 7.088499 | 7.054064 | 7.257799 | 7.462055 |
| 6.627239 | 6.820917 | 7.146192 | 6.774498 | 7.23555 | 6.923989 | 6.321617 | 6.77445 | 6.338125 |
| 5.879531 | 6.260235 | 4.700445 | 5.297626 | 6.226901 | NaN | 4.719426 | NaN | 5.268816 |
| NaN | NaN | NaN | 4.576356 | 4.635514 | NaN | 5.695 | 5.754111 | 5.919463 |
| 7.533358 | 7.448714 | 7.35657 | 7.762398 | 7.66844 | 7.594405 | 7.649314 | 7.620558 | 7.125963 |
| 8.025817 | 8.136532 | 7.987264 | 7.512166 | 7.825389 | 7.798325 | 7.735689 | 8.134166 | 7.614828 |
| 7.828944 | 7.434728 | 7.56375 | 7.389818 | 7.425045 | 7.91446 | 7.93016 | 7.84234 | 7.504005 |
| 7.25916 | 6.78186 | 6.970882 | 7.462422 | 7.160744 | 7.887725 | 7.990126 | 7.997931 | 7.472823 |
| 8.991856 | 8.931661 | 9.034783 | 10.26344 | 10.31081 | 8.658318 | 8.471853 | 8.634258 | 7.584944 |
| 9.608823 | 9.725121 | 9.771558 | 7.246083 | 7.487678 | 5.679264 | 5.377537 | 5.553666 | 6.156836 |
| 6.368981 | 6.217513 | 6.299695 | 7.483689 | 7.33417 | 6.00605 | 6.137227 | 6.027953 | 5.885811 |
| 5.641799 | 5.057243 | NaN | 4.036554 | 4.661142 | 6.057857 | 6.765168 | 6.539111 | 6.730518 |
| 6.622172 | 6.924206 | 6.788366 | 7.317959 | 7.098471 | 7.434475 | 7.241214 | 7.263746 | 6.675051 |
| 5.092248 | 4.433436 | 3.919679 | NaN | NaN | 6.053428 | 6.240372 | 6.133643 | 3.440015 |
| 8.272342 | 8.035376 | 8.028063 | 7.838459 | 7.682248 | 8.145477 | 8.24789 | 8.140826 | 7.430709 |
| 7.250336 | 7.106361 | 7.027175 | 8.265874 | 8.47664 | 6.680255 | 6.346756 | 7.008051 | 6.065504 |
| 6.290526 | 6.030099 | 6.354453 | 6.787321 | 7.102395 | 6.388323 | 6.539928 | 6.74091 | 6.017086 |
| 7.195297 | 6.773232 | 7.13476 | 5.544386 | NaN | 7.504816 | 7.533319 | 7.572676 | 8.046011 |
| 6.848146 | NaN | NaN | 7.065418 | 6.162408 | NaN | NaN | NaN | 6.479519 |
| 7.887175 | 8.14256 | 7.938389 | 7.672791 | 7.815732 | 7.947731 | 7.994425 | 7.719803 | 7.419321 |
| 8.268409 | 8.289006 | 8.063727 | 8.346053 | 8.508898 | 8.132589 | 8.394295 | 8.39068 | 8.236869 |
| 7.218247 | 7.183104 | 7.433495 | 7.9547 | 7.827748 | 6.9162 | 6.995845 | 6.976809 | 7.202159 |
| 10.19753 | 10.38613 | 10.45472 | 8.718787 | 8.820072 | 10.18738 | 10.23985 | 10.33698 | 10.30736 |
| 7.509009 | 7.432731 | 7.234601 | 6.794635 | 6.80067 | 6.940917 | 6.495864 | 7.072977 | 7.180485 |
| 8.263897 | 8.127462 | 7.973976 | 7.894944 | 8.257822 | 8.48029 | 7.66057 | 7.689573 | 7.73827 |
| 8.913985 | 8.986854 | 9.007827 | 10.52858 | 10.44197 | 9.287437 | 9.417288 | 9.474385 | 8.448181 |
| 7.160069 | 7.196876 | 7.292448 | 7.790053 | 7.76557 | 7.843084 | 7.868858 | 7.662373 | 7.309057 |
| 4.79614 | NaN | 3.841801 | 4.837451 | 5.292697 | 4.905236 | 3.872812 | 4.763061 | 4.627732 |
| 6.690984 | 6.621626 | 6.259957 | 6.024779 | 6.384569 | 6.890666 | 7.2955 | 6.761202 | 5.161227 |

| | | | | | | | | |
|---|---|---|---|---|---|---|---|---|
| 6.529374 | 6.917215 | 6.888949 | 6.142529 | 6.360274 | 6.336099 | 6.506278 | 6.753634 | 7.145093 |
| NaN | 7.198139 | 7.423978 | NaN | NaN | 7.104275 | 7.364011 | 7.221928 | NaN |
| 10.24609 | 10.11493 | 10.19898 | 10.89486 | 10.84023 | 10.59018 | 10.69646 | 10.5069 | 9.940027 |
| 6.605002 | 6.180534 | 6.251155 | 6.5795 | 6.333577 | 6.800064 | 6.928648 | 7.033089 | 6.435689 |
| 11.48554 | 11.4949 | 11.35247 | 10.94309 | 11.04691 | 12.03285 | 11.5622 | 11.69308 | 10.58279 |
| 7.483345 | 7.829488 | 7.791604 | 7.127791 | 7.699586 | 7.378008 | 6.914473 | 7.024523 | 6.638145 |
| 7.895678 | 7.02492 | 6.913105 | NaN | 5.982277 | 8.074984 | 7.620367 | 8.276224 | 5.505398 |
| NaN | 6.283722 | NaN | 6.720587 | 7.112402 | 5.893733 | 6.051447 | 6.537484 | 5.893865 |
| 6.068035 | 6.458532 | 5.990345 | 6.376424 | 6.605675 | 6.052347 | 5.414477 | 5.752335 | 4.329412 |
| 7.567822 | 7.028409 | 6.796008 | 7.909967 | 7.553354 | NaN | NaN | 6.361919 | 5.434106 |
| 6.973187 | 6.995764 | 6.705438 | 7.822531 | 7.959189 | 7.562688 | 7.161084 | 7.079575 | 6.216955 |
| 6.945486 | 6.751454 | 6.749619 | 8.691083 | 8.943167 | 6.894616 | 7.201025 | 7.249252 | 6.099085 |
| 5.41402 | 3.942653 | 4.960225 | 5.841095 | 7.224947 | 4.224044 | 4.717172 | NaN | 4.779031 |
| 6.777624 | 6.341049 | 6.424285 | 5.191233 | 5.855026 | 6.446891 | 6.758828 | 6.024819 | 6.474819 |
| 6.878739 | 6.377717 | 7.270149 | 6.462769 | 6.196469 | 6.020222 | 6.312323 | 6.216722 | 7.402103 |
| 9.337171 | 9.360088 | 9.600985 | 9.120771 | 8.897531 | 9.531156 | 9.253135 | 9.459534 | 9.547991 |
| 4.542645 | 3.580232 | 4.330213 | 3.231896 | 4.981347 | NaN | NaN | NaN | 5.02998 |
| 5.564428 | 4.341612 | 5.441749 | 4.390432 | 3.807068 | 6.212061 | 6.198012 | 6.351687 | 5.232512 |
| 9.247676 | 9.191704 | 8.929373 | 9.187016 | 8.228931 | 9.496525 | 9.60537 | 9.257157 | 6.611646 |
| 8.882192 | 8.809422 | 8.896464 | 9.582942 | 9.313101 | 8.957207 | 9.089173 | 9.357769 | 8.776596 |
| 6.74841 | 5.371608 | 6.184345 | 5.672691 | 4.370713 | 6.5949 | 6.09 | 6.060242 | 6.007472 |
| 6.875633 | 6.496066 | 6.433776 | 6.244984 | 6.450116 | 6.511389 | 6.490539 | 6.544592 | 5.652532 |
| 6.246241 | 6.551715 | 6.201345 | 6.443061 | 6.278835 | 6.252898 | 5.841097 | 6.171254 | 6.253112 |
| NaN | 5.911836 | NaN | 4.231662 | 5.335422 | 6.113523 | 4.98487 | 6.046474 | 4.437559 |
| 6.522024 | 6.572802 | 6.57639 | 7.201962 | 6.681799 | 6.850286 | 6.384059 | 6.591755 | 5.41824 |
| NaN | NaN | NaN | 5.926493 | 6.615639 | 5.221601 | 5.739622 | 5.545608 | 4.970648 |
| 6.361323 | 6.56152 | 6.898661 | 6.134859 | 6.263444 | 7.374373 | 6.937548 | 7.35395 | 5.579603 |
| 6.621948 | NaN | 4.458408 | 5.025202 | 5.182046 | 6.242435 | 6.373521 | 6.031078 | 5.928653 |
| 7.402261 | 7.334334 | 7.25081 | 5.841086 | 6.384958 | 7.156095 | 7.450636 | 7.462257 | 7.046765 |
| 8.494948 | 8.46666 | 8.342444 | 7.149268 | 7.137601 | 8.31891 | 8.144146 | 8.178278 | 8.417068 |
| 8.214231 | 8.29633 | 8.169917 | 8.470323 | 8.57426 | 7.676581 | 8.027347 | 7.991457 | 7.940105 |
| 7.882623 | 7.964071 | 8.281411 | 7.291451 | 7.019928 | 7.929726 | 7.799917 | 7.891842 | 7.34563 |
| 6.5966 | 6.613064 | 6.283365 | 7.340029 | 7.783574 | 5.996443 | 5.827322 | 5.65109 | 6.357809 |
| 7.009108 | 6.844018 | 6.978873 | 6.413525 | 6.523493 | 7.282654 | 7.104871 | 7.249238 | 6.217649 |
| 6.79468 | 6.815767 | 6.966444 | 7.207072 | 7.104593 | 7.350682 | 7.527235 | 7.422582 | 7.649291 |
| 7.095278 | 7.063488 | 7.747791 | NaN | NaN | 7.48477 | 7.14682 | 7.122841 | 7.084101 |
| 7.877758 | 7.83905 | 7.80245 | 6.005741 | 6.214072 | 8.116916 | 8.083161 | 8.193044 | 6.880643 |
| 8.382919 | 8.421001 | 8.453608 | 7.953997 | 7.801956 | 8.158287 | 8.108987 | 8.210876 | 8.08053 |
| 7.652714 | 7.747948 | 7.728169 | 7.659384 | 7.832273 | 7.474969 | 8.065379 | 7.376873 | 8.269309 |
| 5.63846 | 5.145042 | 5.410369 | 5.402604 | 4.554523 | 5.762709 | 5.795905 | 6.269382 | 5.153367 |
| 6.028824 | 5.9409 | 6.846054 | 5.769369 | 5.889097 | 5.872659 | 6.206569 | 6.444011 | 4.933664 |
| 5.431153 | 5.668056 | 5.758855 | 5.805432 | 5.602846 | 5.776802 | 5.621668 | 5.664733 | 6.204252 |
| 6.337252 | 6.152214 | 6.351692 | 6.874466 | 6.590741 | 7.221469 | 6.95059 | 7.018209 | 6.591474 |
| 6.281501 | 6.315357 | 6.343934 | 5.411527 | 4.910107 | 6.293712 | 6.602992 | 6.486646 | 6.165592 |
| 7.467816 | 7.222987 | 7.261981 | 6.885562 | 6.625955 | 6.751018 | 6.787122 | 6.511188 | 7.407224 |
| 9.075533 | 8.854547 | 8.941617 | 7.901957 | 7.558785 | 8.521381 | 8.532997 | 8.609431 | 9.143682 |
| 7.995264 | 8.040664 | 8.056315 | 8.71335 | 8.740992 | 8.287934 | 7.898016 | 8.292928 | 7.727848 |

-Bruker Confidential-

| | | | | | | | | |
|---|---|---|---|---|---|---|---|---|
| 10.46923 | 10.40495 | 10.31206 | 10.76336 | 10.50835 | 9.702288 | 9.820129 | 9.818599 | 9.81547 |
| 6.975543 | 6.857252 | 7.3429 | 4.934544 | 5.132915 | 7.252383 | 7.413132 | 7.252581 | 6.605176 |
| 6.213087 | 6.540642 | 6.639647 | 5.981503 | 7.564003 | 6.83203 | 6.905044 | 6.852578 | 7.14627 |
| 6.171856 | 4.726749 | 6.507987 | 6.321999 | NaN | NaN | NaN | NaN | 5.429859 |
| 6.577556 | 5.615449 | 6.221235 | 6.84135 | 6.910547 | 6.184326 | 6.120311 | 5.984968 | 7.046216 |
| 7.120876 | 6.855683 | 6.890919 | NaN | 4.700297 | 2.726367 | NaN | NaN | 5.483181 |
| NaN | 5.052023 | 4.423767 | 6.255473 | 7.085284 | 5.901486 | 4.559976 | 5.129728 | 4.355901 |
| 4.754934 | 4.914713 | 3.973515 | 4.784855 | 3.372519 | 5.914999 | 6.048521 | 5.884184 | 4.372216 |
| 7.842717 | 7.867137 | 7.83593 | 8.045656 | 8.292442 | 7.520808 | 7.478735 | 7.571928 | 7.525442 |
| 7.896445 | 7.847608 | 7.957904 | 6.658251 | 6.672203 | 7.788388 | 7.75742 | 7.894114 | 7.859197 |
| 9.509549 | 9.364586 | 9.343636 | 9.187386 | 9.34457 | 9.698272 | 9.527218 | 9.767511 | 9.655211 |
| 7.409697 | 7.334005 | 7.440978 | 6.331815 | 7.281463 | 7.783892 | 7.622993 | 7.640217 | 6.49545 |
| 7.97567 | 7.555581 | 7.382412 | 7.735822 | 7.412047 | 7.478802 | 7.885459 | 7.303925 | 7.027153 |
| 7.072679 | NaN | 6.314968 | NaN | 5.734787 | NaN | NaN | NaN | 5.876591 |
| 4.83972 | 2.616642 | 3.841081 | 4.142567 | 3.881984 | 4.592951 | 4.648068 | 4.85064 | 5.686795 |
| 7.481815 | 7.455202 | 7.250337 | 7.340567 | 7.640316 | 7.226633 | 7.19453 | 7.174789 | 7.554627 |
| 8.069237 | 8.440495 | 8.264672 | 6.739682 | 5.659143 | 8.092413 | 8.35062 | 7.505972 | 8.069696 |
| 7.68432 | 7.916671 | 6.519532 | 6.592271 | 6.818584 | 7.385044 | 7.287846 | 7.533995 | 6.825909 |
| 7.560476 | 7.229184 | 7.789148 | 7.165995 | 7.16863 | 7.271236 | 7.212003 | 7.087957 | 7.284362 |
| 9.397148 | 9.55015 | 9.442067 | 8.338625 | 8.484322 | 9.612305 | 9.46975 | 9.699578 | 9.148878 |
| 9.968374 | 9.894622 | 9.952834 | 8.078257 | 8.107 | 9.538807 | 9.870611 | 9.728751 | 8.706147 |
| 6.872989 | 6.769795 | 6.505448 | 6.675575 | 6.452468 | 6.709966 | 6.635526 | 7.000332 | 6.837237 |
| 7.891053 | 7.980406 | 7.849116 | 9.489838 | 9.48651 | 7.907896 | 7.965498 | 8.025617 | 8.043428 |
| NaN | 4.35372 | 5.534956 | NaN | NaN | NaN | NaN | NaN | 5.245762 |
| 5.07608 | 5.81271 | 5.551438 | 4.473227 | 4.793407 | 6.153461 | 6.126114 | 6.176783 | 5.553137 |
| 9.268587 | 9.360769 | 9.516973 | 9.132189 | 9.140107 | 9.334616 | 9.184443 | 9.084869 | 9.496197 |
| 9.213088 | 9.3063 | 9.15295 | 8.122539 | 7.593964 | 9.064082 | 9.101127 | 8.841949 | 9.13354 |
| 6.428174 | 7.127783 | 6.848165 | NaN | NaN | 6.273984 | 6.286811 | 6.417688 | 5.695239 |
| 7.649918 | 7.633054 | 7.500273 | 5.928997 | 6.718009 | 8.198624 | 8.316262 | 8.441676 | 7.57407 |
| 8.007407 | 7.491214 | 7.719569 | 6.608082 | 6.720631 | 6.784292 | 6.584266 | 6.806631 | 6.25504 |
| 7.485643 | 7.266231 | 6.892723 | 7.489578 | 7.515088 | 6.985967 | 7.048751 | 6.846738 | 7.519746 |
| 9.636448 | 9.153657 | 8.9315 | 10.81891 | 10.28887 | 9.24974 | 9.172798 | 9.167451 | 8.485646 |
| 6.328781 | 7.630032 | 6.987802 | 6.600744 | 6.646032 | 7.574592 | 7.132806 | 7.447945 | 6.281004 |
| 7.069548 | 6.616681 | 6.830746 | 6.269541 | 6.342746 | 6.340683 | 6.694697 | 6.795057 | 6.213778 |
| 6.794552 | 6.862801 | 6.994996 | 7.698372 | 8.117268 | 6.649929 | 6.708066 | 6.310741 | 6.785015 |
| 8.127012 | 8.345372 | 7.82216 | 7.650855 | 8.233938 | 7.915011 | 7.874895 | 7.874387 | 7.840463 |
| 8.416423 | 8.561748 | 8.53375 | 6.478702 | 5.384996 | 6.930843 | 7.032108 | 6.606473 | 7.257546 |
| 5.170637 | 6.01543 | NaN | 4.591645 | NaN | 6.10437 | 6.693915 | 6.963091 | 5.908178 |
| 7.627149 | 7.056783 | 7.230529 | 5.415711 | 5.801537 | 7.589117 | 7.488247 | 7.148987 | 7.064783 |
| 7.421336 | 7.394468 | 7.4103 | 7.541337 | 7.682178 | 7.593766 | 7.714153 | 7.739169 | 6.987423 |
| 10.46255 | 10.58035 | 10.6636 | 9.926188 | 9.544228 | 7.935825 | 7.963609 | 8.022491 | 7.676764 |
| 7.066664 | 5.517597 | 6.479884 | 6.087389 | 5.048823 | 6.409643 | 6.161386 | 6.592412 | 5.804289 |
| 5.33056 | 6.22534 | 5.237739 | NaN | 5.723418 | 6.540462 | 5.950145 | 6.055048 | 6.777937 |
| 10.24465 | 10.25474 | 10.33495 | 10.02577 | 10.08566 | 10.55895 | 10.50956 | 10.7681 | 9.682343 |
| 8.036098 | 8.091091 | 8.166212 | 8.264706 | 7.546597 | 7.843463 | 7.919452 | 8.252424 | 8.475647 |
| 7.363492 | 7.386467 | 7.283985 | 7.409867 | 7.491811 | 7.341896 | 6.776622 | 6.998697 | 6.885113 |
| 8.850021 | 8.679941 | 8.788951 | 7.901557 | 7.882693 | 8.981207 | 9.133772 | 8.940605 | 8.858142 |

| | | | | | | | | |
|---|---|---|---|---|---|---|---|---|
| 6.85213 | 6.804707 | 7.214031 | 6.823295 | 6.587934 | 6.397245 | 6.491531 | 6.445862 | 6.28442 |
| 9.405194 | 9.505668 | 9.627165 | 11.01486 | 11.21364 | 9.901414 | 9.870914 | 10.07827 | 9.892842 |
| 9.360027 | 9.551929 | 9.60866 | 8.751573 | 9.184566 | 9.710804 | 9.452458 | 9.417885 | 9.786115 |
| 6.457931 | 6.408149 | 6.699329 | 4.876449 | 5.020129 | 6.705582 | 6.632644 | 6.545179 | 5.46405 |
| 11.72635 | 11.76606 | 11.50435 | 12.02168 | 12.03137 | 12.19096 | 11.95765 | 12.0601 | 11.99634 |
| 8.137474 | 8.193472 | 8.112143 | 7.227038 | 7.204567 | 7.586894 | 7.599901 | 7.937288 | 8.035583 |
| 8.735343 | 8.840852 | 8.691067 | 7.741847 | 7.305662 | 9.181001 | 9.45882 | 9.226099 | 8.4115 |
| 8.252418 | 7.908038 | 8.407355 | 7.949246 | 8.124062 | 7.741061 | 7.895184 | 7.895407 | 8.435206 |
| 5.672631 | 5.923201 | 6.309289 | 5.493642 | 4.867975 | 5.355933 | 5.423305 | 5.636579 | 5.650254 |
| 8.087014 | 8.546103 | 8.138073 | 8.516406 | 8.246159 | 8.485771 | 8.686109 | 8.736191 | 7.804515 |
| 7.598084 | 7.186525 | 7.391906 | 7.530314 | 6.890698 | 7.526706 | 7.404608 | 7.232386 | 7.134112 |
| 5.316887 | 5.635612 | 5.817486 | NaN | NaN | NaN | 5.181863 | 5.217122 | 6.325845 |
| 6.500184 | 5.88765 | 6.563838 | 5.120001 | 5.596141 | 6.942402 | 7.028211 | 6.96532 | 5.92539 |
| 7.665643 | 7.691214 | 7.688427 | 8.312902 | 8.088193 | 7.587101 | 7.675285 | 7.58253 | 7.772751 |
| 8.78011 | 8.793835 | 8.686193 | 9.525841 | 9.368875 | 8.355675 | 8.170315 | 7.992783 | 7.41358 |
| NaN | NaN | NaN | NaN | NaN | NaN | NaN | 4.67146 | 5.302348 |
| 7.178057 | 7.393891 | 7.724509 | 6.456195 | 7.131114 | NaN | NaN | NaN | NaN |
| 5.076207 | 5.598623 | 5.041938 | 5.456886 | 6.233544 | NaN | NaN | NaN | 5.689066 |
| 7.809038 | 7.910609 | 7.814631 | 7.632971 | 7.419645 | 7.734136 | 7.678591 | 7.813234 | 7.691924 |
| 5.555192 | 4.526671 | 4.674287 | NaN | NaN | 5.487926 | 5.414827 | 6.233464 | 6.921317 |
| 6.387801 | 6.589125 | 5.802926 | 8.011099 | 8.791103 | 7.227273 | 6.381351 | 5.922474 | 7.769612 |
| 7.869393 | 8.259924 | 8.18155 | 7.708547 | 7.083572 | 6.686627 | 7.538972 | 7.311812 | 7.41283 |
| 6.662625 | 4.335502 | NaN | 3.929758 | 6.127649 | 5.75736 | NaN | 4.983982 | 5.089622 |
| 7.078432 | 7.496366 | 7.833165 | 7.3124 | 7.626858 | 6.883693 | 6.625788 | 6.554487 | 7.074247 |
| 8.142253 | 8.356308 | 8.040159 | 7.758928 | 8.09573 | 8.093194 | 8.098051 | 8.105096 | 7.956843 |
| 8.928867 | 8.958952 | 8.82938 | 9.611806 | 9.626501 | 9.545552 | 9.364939 | 9.257948 | 9.013561 |
| 6.696786 | 7.087903 | 6.824608 | 6.211454 | 6.395992 | 6.668093 | 6.566322 | 6.463809 | 6.496537 |
| 6.646364 | 6.934741 | 6.557278 | 8.446971 | 8.650817 | 6.677639 | 6.771285 | 6.769342 | 6.756659 |
| 8.266273 | 8.338426 | 8.044197 | 9.864369 | 9.883749 | 8.423912 | 8.632839 | 8.388211 | 8.072548 |
| 3.578858 | 4.268049 | NaN | NaN | NaN | NaN | NaN | 3.266816 | 4.460884 |
| NaN | NaN | 3.996957 | NaN | 3.311228 | 3.842661 | 4.436452 | 4.347609 | 5.429162 |
| 7.137928 | 7.059008 | 7.419912 | 6.085485 | NaN | 7.897147 | 7.480364 | 7.480544 | 7.307065 |
| 5.024537 | 4.473189 | 4.967944 | NaN | 3.787552 | NaN | 4.772126 | 5.138988 | 5.330783 |
| NaN | NaN | 5.578165 | 5.289334 | 5.865036 | 5.574706 | NaN | 5.476941 | 4.259435 |
| 5.806304 | 5.316174 | 5.937186 | 5.094989 | 3.545483 | 6.457061 | 6.701159 | 6.525463 | 5.997652 |
| 6.335145 | 6.569967 | 6.134729 | NaN | 5.77401 | 6.8261 | 6.738856 | 5.799084 | 6.155032 |
| 6.883586 | 7.022734 | 7.192057 | 6.000084 | 6.376747 | NaN | NaN | NaN | 6.996561 |
| 7.654824 | 8.184723 | 8.848795 | 7.760998 | 7.98299 | 8.347522 | 8.035126 | 8.387191 | 7.852153 |
| 6.837132 | 6.752693 | 6.758973 | 5.461603 | 5.131029 | 6.924503 | 6.349401 | 7.143931 | 6.261444 |
| 6.231441 | 6.383119 | 6.46678 | 5.115968 | 5.347806 | 6.208385 | 6.947701 | 6.334624 | 7.970937 |
| 6.67173 | 6.680196 | 6.508303 | 6.016601 | 5.850935 | 7.082701 | 6.989043 | 7.119005 | 6.83938 |
| NaN | 5.73002 | 5.693568 | NaN | 4.917109 | NaN | 6.920353 | 7.130955 | NaN |
| 7.187644 | 7.32048 | 6.860125 | 7.297768 | 6.913117 | 7.226186 | 6.987095 | 7.106204 | 6.768745 |
| 5.95272 | 6.189456 | 6.098656 | 6.189112 | 5.986338 | 5.649222 | 5.615253 | 5.546433 | 6.245019 |
| NaN | 5.786762 | 6.191944 | 3.608964 | 5.114431 | 5.822781 | NaN | NaN | 4.919956 |
| 8.719163 | 8.40475 | 8.365925 | 7.069918 | 6.801548 | 8.30372 | 8.466463 | 8.430901 | 7.474603 |
| 9.370785 | 9.555907 | 9.417358 | 9.256477 | 9.064141 | 9.757621 | 9.614033 | 9.646473 | 8.727787 |

| | | | | | | | | |
|---|---|---|---|---|---|---|---|---|
| 8.154227 | 7.753217 | 7.893616 | 6.889788 | 7.379206 | 8.094054 | 8.211551 | 8.156006 | 7.903056 |
| 6.853906 | 6.686059 | 6.832922 | 5.881976 | 6.332049 | 6.48805 | 6.569518 | 6.790777 | 7.214707 |
| 8.30572 | 7.906058 | 8.240808 | 6.865404 | 7.020976 | 8.534919 | 8.397632 | 8.695741 | 8.500179 |
| 6.187869 | 6.401564 | 5.96982 | 6.195928 | 6.214791 | 5.83395 | 6.115196 | NaN | 5.705142 |
| 9.112291 | 8.847953 | 8.961061 | 9.980932 | 10.60084 | 8.551497 | 9.212807 | 8.318741 | 8.348047 |
| 6.630616 | 6.781241 | 6.715293 | NaN | NaN | NaN | 7.168458 | NaN | 6.181951 |
| 8.415174 | 8.487218 | 8.777359 | 8.583081 | 8.16455 | 8.199565 | 8.164004 | 8.467734 | 8.149762 |
| 7.381582 | 7.341504 | 7.234099 | 6.404757 | 6.285484 | 7.204259 | 6.060049 | 6.561533 | 6.837413 |
| 9.109221 | 8.994909 | 8.554449 | 8.029413 | 7.949205 | 8.364929 | 8.316642 | 8.929281 | 9.046501 |
| 7.898502 | 6.570889 | 7.104086 | 6.38096 | 5.964274 | 6.813867 | 7.339684 | 7.072714 | 6.199064 |
| 7.893866 | 7.845662 | 7.947477 | 7.584208 | 7.280493 | 8.206164 | 8.069104 | 8.138797 | 7.693732 |
| 7.327209 | 6.507949 | 7.126484 | 4.819786 | 5.546358 | 7.249129 | 7.297101 | 7.026162 | 5.777318 |
| 8.863314 | 8.941561 | 8.804059 | 8.800113 | 8.456557 | 8.999318 | 9.099004 | 8.986381 | 8.743781 |
| 8.670137 | 8.750931 | 8.757218 | 10.42942 | 10.71488 | 8.126604 | 8.384963 | 8.49044 | 8.636142 |
| 7.797915 | 7.819121 | 7.842263 | NaN | NaN | NaN | 9.034524 | 9.125892 | 7.712047 |
| NaN | NaN | NaN | 6.873234 | 6.430806 | NaN | NaN | NaN | NaN |
| 13.06034 | 13.08482 | 13.06031 | 12.65287 | 12.21598 | 11.35317 | 11.64645 | 11.43557 | 11.222 |
| 8.884933 | 8.597635 | 8.577539 | 8.808705 | 8.986862 | 8.618229 | 8.49229 | 8.55042 | 7.789759 |
| 5.617676 | 5.180361 | 5.454853 | 5.626549 | 4.194353 | 6.369512 | 6.939132 | 6.693573 | 6.555085 |
| 8.48024 | 7.98704 | 8.248404 | 7.213688 | 7.751056 | 8.600577 | 8.792169 | 8.815611 | 8.139888 |
| 7.085837 | 7.308129 | 6.560693 | 8.048916 | 8.150937 | 7.681634 | 7.831474 | 8.045984 | 6.672989 |
| 5.306002 | 4.106225 | NaN | 5.050349 | 5.629875 | 5.86111 | 6.121366 | 6.322318 | 6.323781 |
| 8.632819 | 8.615877 | 8.484963 | 8.566576 | 8.424212 | 9.132513 | 9.047758 | 8.809171 | 8.299755 |
| 5.432477 | 4.358258 | 3.632854 | NaN | NaN | NaN | NaN | NaN | 5.583119 |
| 6.575953 | 6.423632 | 6.609944 | 7.596099 | 7.68205 | 5.963558 | 6.705606 | 6.669893 | 7.161231 |
| 5.083478 | 4.715652 | 4.194073 | NaN | 5.512024 | 5.745708 | 5.849515 | 5.822961 | 5.15488 |
| 7.655551 | 7.396791 | 7.316277 | 8.983789 | 9.041434 | 7.235291 | 7.329985 | 7.628288 | 8.098546 |
| 6.637445 | 6.891531 | 6.436496 | 6.666819 | 6.411653 | 6.09103 | 6.337601 | 6.551301 | 4.801569 |
| 9.679135 | 9.552704 | 9.583689 | 9.989801 | 10.06934 | 9.324032 | 9.264591 | 9.279337 | 10.23529 |
| 8.53969 | 8.3982 | 8.443127 | 7.804257 | 8.351155 | 8.853225 | 9.212212 | 9.342973 | 8.647175 |
| 9.092926 | 9.022871 | 9.140492 | 9.230866 | 9.402659 | 7.886725 | 9.046906 | 8.958183 | 7.504824 |
| 7.772253 | 8.357909 | 8.115244 | 7.745797 | 7.732196 | 7.316329 | 7.459455 | 7.28447 | 7.447618 |
| 7.886133 | 7.833891 | 7.793957 | 8.761604 | 8.91642 | 7.747043 | 7.678735 | 7.725958 | 7.558615 |
| 6.634038 | 6.70162 | 6.866521 | 8.70118 | 8.790735 | 7.378626 | 7.386956 | 7.428714 | 7.930723 |
| 7.719876 | 7.380539 | 7.515828 | NaN | 5.959559 | 7.324414 | 7.39818 | 7.537505 | 7.032948 |
| 9.005029 | 8.594688 | 8.605061 | 7.844972 | 7.896692 | 8.466952 | 8.38826 | 8.403625 | 8.229967 |
| 9.958568 | 9.770797 | 9.806117 | 10.8099 | 10.53009 | 9.577745 | 9.604653 | 9.448745 | 10.28358 |
| 7.31314 | 6.770267 | 7.034411 | 5.036329 | 5.351215 | 7.126864 | 6.742792 | 7.0067 | 7.467161 |
| 8.58147 | 8.497385 | 8.327485 | 8.837311 | 8.676445 | 8.177405 | 8.244948 | 7.868597 | 8.498208 |
| 7.620638 | 7.19452 | 7.433029 | 7.707571 | 7.704347 | 6.364146 | 5.61814 | 6.02198 | 7.416249 |
| 9.115609 | 7.911787 | 7.515755 | 4.853849 | 7.000508 | NaN | NaN | 7.452276 | NaN |
| NaN | 6.830859 | 7.23498 | 6.71981 | 6.483598 | 8.435835 | 7.64445 | 6.973297 | 7.215121 |
| 9.110568 | 8.806442 | 8.925536 | 6.661967 | 6.828055 | 8.788203 | 8.778285 | 8.767015 | 8.716125 |
| 8.154124 | 7.969641 | 8.154966 | 7.586002 | 7.578535 | 7.205863 | 7.759461 | 7.654053 | 7.14346 |
| 7.803677 | 8.068647 | 7.745225 | 6.634084 | 6.735007 | 8.318724 | 8.357791 | 8.328572 | 7.412447 |
| NaN | NaN | NaN | NaN | 3.6502 | NaN | NaN | NaN | 4.73872 |
| 5.451569 | 5.875239 | 5.938119 | 5.864741 | 6.457911 | 6.819645 | 6.291249 | 6.580637 | 6.280903 |

| | | | | | | | | |
|---|---|---|---|---|---|---|---|---|
| 7.612268 | 7.444676 | 7.642065 | 6.2373 | 6.574856 | 7.610646 | 7.711225 | 7.861902 | 7.71835 |
| 6.932796 | 7.75527 | 7.717863 | 7.26309 | 5.924934 | 4.660596 | NaN | 4.492994 | 3.662987 |
| 9.97441 | 10.01784 | 9.981589 | 11.55774 | 11.47072 | 10.27444 | 10.24082 | 10.34451 | 10.49288 |
| 7.287004 | 6.991101 | 7.185708 | 9.065588 | 9.149752 | 6.97309 | 6.243979 | 7.168757 | 7.842391 |
| 6.717802 | 6.439077 | 6.284114 | 7.040888 | 6.77679 | 6.12401 | 5.991412 | 5.864088 | 5.805248 |
| 7.209946 | 6.456632 | 7.239985 | NaN | 6.480142 | 7.752286 | 7.550472 | 7.830557 | 7.333485 |
| 7.129898 | 7.252695 | 7.064441 | 6.340919 | 5.577223 | 6.541762 | 6.358765 | 6.840117 | 7.467971 |
| 7.293838 | 7.211423 | 7.168673 | 6.285424 | 6.468958 | 7.357065 | 7.352433 | 7.37312 | 6.735912 |
| 7.345379 | 7.351781 | 7.676887 | 6.882987 | 6.606241 | 8.149974 | 8.13968 | 8.381572 | 7.198395 |
| 7.732519 | 7.990693 | 7.741591 | 7.098838 | 7.469905 | 7.609925 | 7.961365 | 7.724985 | 7.591389 |
| 7.275387 | 6.733524 | 7.232587 | 6.790006 | 6.654824 | 7.484442 | 6.420552 | 6.497571 | 6.508448 |
| 8.394114 | 8.173412 | 8.336029 | 7.810132 | 7.920429 | 8.175908 | 8.485799 | 8.40531 | 8.139795 |
| 7.236046 | 7.397316 | 6.860198 | NaN | NaN | 6.465283 | 6.476535 | 5.693334 | 6.131608 |
| 7.741611 | 7.672409 | 7.624502 | 8.081841 | 8.004217 | 7.866131 | 7.522108 | 7.864816 | 7.684417 |
| NaN | NaN | 6.503577 | 6.993712 | 7.817353 | NaN | NaN | NaN | NaN |
| 7.095636 | 7.074786 | 6.649058 | 6.857224 | 6.925255 | 6.54078 | 6.337241 | 5.980366 | 6.214602 |
| 6.393499 | 6.47968 | 6.022208 | 7.46816 | 7.293109 | 6.304736 | 4.894503 | 5.849911 | 5.395639 |
| 7.815054 | 7.467178 | 8.15312 | 8.185358 | 8.077592 | 7.454 | 7.974369 | 7.23352 | 6.748277 |
| 8.225126 | 8.302715 | 8.220224 | 7.286281 | 7.647025 | 7.540076 | 7.130424 | 7.451855 | 7.520243 |
| 7.862791 | 7.78878 | 7.861699 | 6.836238 | 6.463415 | 7.414397 | 7.751555 | 7.71778 | 7.277025 |
| 7.786064 | NaN | NaN | 6.241495 | 6.718128 | NaN | NaN | NaN | NaN |
| 8.974602 | 8.946788 | 9.198081 | 7.526814 | 7.610684 | 8.832918 | 8.888651 | 8.753391 | 8.628204 |
| 8.118175 | 8.386225 | 8.266013 | 8.871585 | 8.91786 | 7.319129 | 7.562276 | 7.385035 | 8.092088 |
| 7.099621 | 6.957235 | 6.496718 | 7.37856 | 7.129673 | 7.122873 | 7.202929 | 7.1926 | 6.475968 |
| 7.05022 | 6.806691 | 7.324567 | 6.172508 | 5.682029 | 7.014519 | 7.131735 | 7.085896 | 7.127807 |
| 8.310019 | 8.359353 | 8.435373 | 8.102773 | 7.986536 | 8.937831 | 8.596143 | 8.681214 | 8.103638 |
| 6.890624 | 7.415709 | 7.118949 | 5.810305 | 6.13987 | 6.646121 | 6.648872 | 6.989878 | 6.795273 |
| 6.712817 | 4.810119 | 8.101758 | NaN | NaN | 7.723436 | 6.841612 | 7.436779 | 7.447577 |
| 6.823622 | 5.687575 | 5.620669 | 5.826976 | 6.073416 | 6.273493 | 5.944716 | 6.167857 | 6.389848 |
| 5.572677 | 5.424789 | 5.505098 | NaN | NaN | 6.893333 | 6.997617 | 6.9379 | 6.904008 |
| 5.208475 | 4.362513 | 4.093428 | NaN | NaN | NaN | 3.204424 | 4.432724 | 4.877447 |
| NaN | NaN | NaN | 7.03411 | 6.919285 | NaN | 6.973695 | 7.059702 | 6.246778 |
| 5.950825 | 6.022085 | 6.117377 | 5.512978 | 6.122263 | 5.346762 | 5.883641 | 5.717844 | 6.127857 |
| 6.752225 | 6.466947 | 6.853806 | 7.036281 | 7.051877 | 6.900084 | 6.955615 | 7.002071 | 6.425318 |
| 11.64683 | 11.59455 | 11.6195 | 12.47599 | 12.3162 | 11.11896 | 11.19476 | 11.15049 | 11.54727 |
| 7.484178 | 7.674514 | 7.652653 | 7.74965 | 7.414349 | 7.379431 | 7.30329 | 7.405241 | 7.438714 |
| 8.487685 | 8.231718 | 8.457392 | 7.963691 | 7.616497 | 7.639481 | 7.652849 | 7.689987 | 7.391601 |
| 7.140491 | 7.558397 | 7.669219 | 8.191389 | 8.075559 | 7.442395 | 7.159959 | 5.702978 | 5.708552 |
| 6.679125 | 6.461596 | 6.845486 | 7.378311 | 7.45679 | 6.387393 | 6.076343 | 6.658707 | 6.794103 |
| 7.410944 | 7.44918 | 7.730762 | NaN | 5.354609 | NaN | NaN | NaN | NaN |
| 7.986475 | 6.479075 | 7.847167 | 6.338159 | NaN | 9.345614 | 9.063192 | 9.018589 | 6.893085 |
| 8.010118 | 7.660651 | 7.211339 | 7.56879 | 8.176836 | 6.940941 | 7.511131 | 6.854496 | 6.943887 |
| 9.281297 | 9.165812 | 9.390716 | 12.33484 | 12.36543 | 8.565502 | 7.966918 | 8.372273 | 8.188704 |
| 6.529122 | 5.889645 | 6.177138 | NaN | NaN | 5.006922 | 6.658074 | NaN | 6.665833 |
| 9.720139 | 9.824953 | 9.772892 | 10.9979 | 11.01902 | 9.750504 | 9.795966 | 9.945455 | 10.28372 |
| 5.954046 | 5.931925 | 5.39889 | 7.682065 | 8.324968 | NaN | NaN | NaN | 5.626304 |
| 12.84506 | 12.76196 | 12.96092 | 12.2186 | 11.84973 | 12.93456 | 12.79594 | 13.31842 | 12.46007 |

| | | | | | | | | |
|---|---|---|---|---|---|---|---|---|
| 7.044826 | 6.99318 | 6.958053 | 6.944936 | 7.114283 | 7.063519 | 7.014299 | 7.141691 | 6.507971 |
| 7.125583 | 7.395592 | 7.092053 | 5.362678 | 6.045767 | 7.193913 | 6.941738 | 7.097941 | 7.131107 |
| 8.876204 | 8.787233 | 8.857498 | 7.36964 | 7.839457 | NaN | 8.547106 | 8.846017 | 8.678739 |
| 7.885259 | 7.972549 | 8.346169 | 7.884798 | 8.102585 | 8.467791 | 8.475205 | 8.481503 | 8.065272 |
| 6.76203 | 6.787455 | 6.76895 | 5.981495 | 5.752615 | 6.77252 | 7.063844 | 6.820744 | 5.929479 |
| NaN | NaN | NaN | 6.443577 | 4.727393 | NaN | NaN | NaN | 5.489074 |
| 7.413712 | 6.540301 | 7.375677 | 7.265635 | 7.153512 | 5.584336 | 6.285271 | 6.474253 | 7.274345 |
| 8.287571 | 8.422551 | 8.235061 | 9.235883 | 9.002406 | 8.060033 | 8.380838 | 8.66557 | 7.932307 |
| 6.593091 | 6.290792 | 6.367029 | 6.964828 | 6.572673 | 6.414912 | 6.367352 | 6.441159 | 5.87818 |
| 6.25934 | 6.100449 | 5.262111 | NaN | 5.155717 | 6.549646 | 6.52755 | 6.976726 | 6.271631 |
| 9.295993 | 9.288962 | 9.319927 | 9.66681 | 9.840738 | 8.923071 | 8.724909 | 9.103472 | 9.173692 |
| 8.268596 | 8.668056 | 8.011215 | 7.358253 | 7.401391 | 7.626251 | 7.785198 | 7.836837 | 8.390919 |
| 8.825048 | 8.544937 | 7.523166 | NaN | 7.310066 | 8.150507 | 8.466797 | 8.399762 | 7.811476 |
| 6.218227 | 6.28969 | 5.980381 | 5.495275 | 6.094473 | 6.62101 | 6.406589 | 6.538228 | 5.849851 |
| 5.957406 | 6.234081 | 5.463321 | 5.019895 | 4.727774 | 7.318963 | 6.371939 | 6.76309 | 4.664607 |
| 7.92417 | 7.471117 | 7.795553 | 7.102995 | 7.614002 | NaN | NaN | NaN | 6.424438 |
| 8.860955 | 8.793095 | 8.644607 | 9.036774 | 8.950292 | 9.041859 | 8.75631 | 9.103744 | 8.075863 |
| 7.273243 | 6.89305 | 7.375308 | 4.545134 | 4.400612 | 7.443496 | 7.367786 | 7.226117 | 7.060338 |
| 9.195616 | 8.694932 | 8.880773 | NaN | NaN | NaN | NaN | NaN | 6.429548 |
| 7.025988 | 7.26469 | 7.416755 | 7.715885 | 7.731076 | 7.091625 | 7.264411 | 7.321015 | 7.062225 |
| 7.717594 | 8.415963 | 7.93964 | 7.341175 | 7.033522 | 7.417397 | 7.963803 | 8.07138 | 8.249202 |
| 8.033426 | 8.133725 | 7.835973 | 10.16696 | 10.16545 | 5.576305 | 5.432667 | 5.9115 | 7.225053 |
| 8.431499 | 8.271131 | 8.244291 | 8.759418 | 9.102568 | 8.501466 | 8.153245 | 8.364603 | 8.018233 |
| 6.182416 | 5.748917 | 5.296484 | NaN | NaN | 7.137034 | 7.397911 | 6.909519 | NaN |
| 6.986038 | 7.083114 | 7.160782 | 5.959832 | 6.443625 | 7.211504 | 7.270748 | 7.228481 | 7.267943 |
| 7.131717 | 7.164703 | 7.334298 | 7.208655 | 6.261056 | 7.603566 | 7.393715 | 7.691772 | 7.576067 |
| 9.167246 | 9.044209 | 9.245833 | 8.331652 | 8.90355 | 8.55453 | 8.82318 | 8.689434 | 8.531875 |
| 7.840966 | 7.862779 | 8.104549 | 6.672915 | 6.200981 | 7.626175 | 7.788908 | 7.621238 | 7.194513 |
| 8.255204 | 8.372484 | 8.025228 | 6.70939 | 6.889081 | 7.718854 | 7.681817 | 7.676474 | 7.170369 |
| 6.831412 | 7.273606 | 7.078247 | 6.301761 | 6.422576 | 6.755341 | 6.898615 | 7.079128 | 6.855586 |
| 6.972442 | 7.052735 | 7.103256 | 6.342283 | 5.773169 | 6.708933 | 6.146245 | 6.272559 | 6.667168 |
| 5.475687 | 6.211382 | 4.889376 | 6.705959 | 6.731381 | 6.964061 | 6.820518 | 6.978902 | 6.604417 |
| 7.58877 | 7.333709 | 7.417742 | 7.437902 | 7.21773 | 7.742141 | 7.976328 | 7.885003 | 7.563921 |
| 7.077709 | 7.117384 | 6.884011 | 7.008183 | 7.083017 | 7.358852 | 7.288168 | 7.305202 | 6.523858 |
| 6.15238 | 6.844348 | 6.205493 | NaN | 4.693215 | 6.625066 | 6.488659 | 6.336019 | 6.057801 |
| 7.650298 | 7.842271 | 7.6957 | 6.853483 | 6.598446 | 7.510338 | 7.235201 | 7.646147 | 7.670621 |
| 7.339334 | 7.127828 | 7.146242 | 6.195578 | 6.517583 | 6.862739 | 7.238926 | 6.818124 | 5.831635 |
| 6.799793 | 6.723536 | 6.13279 | 7.017083 | 6.121059 | 6.855622 | 6.86484 | 6.746214 | 5.772539 |
| 8.761635 | 8.665788 | 8.712916 | 8.473404 | 8.362714 | 9.193596 | 8.917499 | 8.811653 | 9.080608 |
| 7.226083 | 6.480336 | 7.293653 | NaN | 5.542119 | 7.088106 | 7.213707 | 7.690411 | 7.088926 |
| NaN | NaN | NaN | NaN | NaN | NaN | NaN | NaN | 6.524654 |
| NaN | NaN | NaN | 5.453105 | 5.116107 | 7.383428 | 7.119165 | 7.303813 | 5.648335 |
| NaN | NaN | NaN | 5.885447 | 6.899022 | NaN | NaN | NaN | 5.526023 |
| 7.825127 | 7.976987 | 8.925341 | 10.75201 | 8.90591 | 7.316657 | 7.633759 | 7.607572 | 8.750791 |
| 6.238196 | 7.756169 | NaN | 6.960351 | NaN | 6.899395 | 6.765878 | 6.861859 | 6.083121 |
| 6.460689 | 6.074721 | 6.426722 | 7.158467 | 6.92612 | 6.307182 | 6.755029 | 6.804286 | 5.893322 |
| 8.796664 | 8.500532 | 8.646943 | 7.708798 | 7.93352 | 8.321903 | 8.225245 | 8.237906 | 7.78222 |

-Bruker Confidential-

| | | | | | | | | |
|---|---|---|---|---|---|---|---|---|
| 7.150506 | 6.861873 | 7.341075 | 6.778683 | 6.516598 | 7.345295 | 7.194105 | 7.677582 | 6.979238 |
| 7.030716 | 6.979926 | 6.919205 | 7.126917 | 7.105778 | 6.542328 | 6.447504 | 6.434999 | 7.63786 |
| 5.703422 | 5.359771 | 6.332797 | 6.927215 | NaN | 5.919871 | 6.185442 | 6.565524 | 5.98186 |
| 5.086807 | 5.001543 | 5.390411 | 4.503826 | NaN | 6.326965 | 7.235849 | 6.774672 | 6.471407 |
| 6.196529 | 6.114396 | 6.187687 | 5.15865 | 4.960499 | 6.571417 | 5.763083 | 5.126323 | 6.150417 |
| 8.806403 | 8.40493 | 8.507924 | 9.414783 | 9.357082 | 8.298126 | 8.599108 | 8.531269 | 8.546844 |
| 6.79548 | 6.609534 | 6.866025 | 4.724358 | 5.028586 | 6.166189 | 5.972837 | 6.482829 | 6.617176 |
| 7.072659 | 7.154268 | 7.183444 | 6.759966 | 6.509833 | 7.340136 | 6.744328 | 7.340362 | 7.163025 |
| NaN | NaN | NaN | NaN | NaN | NaN | NaN | 5.28351 | 5.418315 |
| 6.685047 | 5.917027 | 7.026978 | 5.609987 | 6.220499 | 5.767015 | 5.938402 | 4.770784 | 6.871072 |
| 6.780376 | 6.909205 | 6.777531 | 6.647475 | 6.362044 | 6.957141 | 6.620014 | 7.165092 | 6.919793 |
| 5.171011 | 5.228636 | 5.373869 | 5.383297 | 6.384944 | 5.328993 | 5.370907 | 4.381482 | 5.219998 |
| 8.713015 | 8.645613 | 8.43259 | 8.208693 | 8.345343 | 9.028443 | 8.958169 | 8.996713 | 7.498781 |
| 8.437144 | 8.229332 | 8.207089 | 6.481875 | 6.592643 | 7.888342 | 8.223373 | 8.074852 | 6.687805 |
| 8.134039 | 8.28917 | 7.893931 | 9.016026 | 8.700685 | 7.530131 | 7.865985 | 7.879002 | 8.392876 |
| 7.654716 | 7.457128 | 8.146484 | 6.754008 | 6.39026 | 8.932397 | 8.884015 | 8.3114 | 6.810463 |
| 5.613692 | 4.615827 | 5.360656 | NaN | NaN | 5.637629 | 4.876605 | 5.697732 | 4.37332 |
| 5.987136 | 6.467832 | 6.201188 | 5.170571 | 5.045579 | 5.960812 | 5.742571 | 5.818408 | 6.111255 |
| 8.48901 | 8.422027 | 8.768143 | 8.608849 | 8.430026 | 8.651471 | 8.554671 | 8.855615 | 8.11711 |
| 8.407195 | 8.291783 | 8.417258 | 7.64636 | 7.673821 | 8.459046 | 8.607646 | 8.411348 | 7.886007 |
| NaN | NaN | NaN | NaN | NaN | NaN | NaN | NaN | 5.596083 |
| 4.684797 | 4.051658 | 4.271195 | 3.819615 | 3.990466 | 4.684875 | 4.667997 | 4.315376 | 5.410405 |
| 8.098498 | 8.09124 | 8.117953 | 6.583547 | 6.281357 | 8.183873 | 8.102623 | 8.068855 | 7.828723 |
| 7.280437 | 6.951967 | 6.786653 | 7.233695 | NaN | NaN | NaN | NaN | NaN |
| 6.607003 | 6.874333 | 6.870095 | 7.575016 | 7.842147 | 7.144022 | 7.02276 | 7.052619 | 7.592415 |
| 7.157584 | 7.47017 | 7.283518 | 6.227522 | 5.323821 | 7.693092 | 7.673792 | 7.858335 | 8.252903 |
| 8.369395 | 8.656517 | 8.445848 | 5.960833 | 6.199261 | 9.098954 | 9.154998 | 9.149734 | 7.814254 |
| 7.523696 | 7.358832 | 7.629791 | 7.664109 | 7.344502 | 7.284503 | 7.432142 | 7.384626 | 6.270669 |
| 8.816916 | 8.689718 | 8.508449 | 9.46756 | 9.573554 | 7.436066 | 7.086686 | 7.312882 | 7.234159 |
| 7.572979 | 7.227574 | 7.499915 | 7.041607 | 6.873917 | 7.563852 | 7.184579 | 7.506804 | 7.209327 |
| 7.607895 | 7.607763 | 7.614775 | 6.615819 | 6.504901 | 7.289724 | 7.291788 | 7.368124 | 8.180366 |
| 6.429697 | 7.566223 | 6.377578 | 7.194737 | 7.327077 | 6.833557 | NaN | 6.280498 | 5.80453 |
| 7.663557 | 7.773975 | 7.886643 | 8.199623 | 9.546838 | 7.084905 | 7.075053 | 6.406972 | 7.002085 |
| 7.899552 | 7.775612 | 7.862951 | 9.560767 | 9.555501 | 6.31553 | 5.888262 | 6.741746 | 7.356244 |
| 8.872422 | 8.745328 | 8.934844 | 7.740387 | 8.159965 | 8.098611 | 8.095905 | 8.410132 | 8.170168 |
| 9.402158 | 9.26056 | 9.047215 | 11.17815 | 11.53155 | 9.035929 | 8.840013 | 9.051558 | 9.442155 |
| 6.959631 | 7.18874 | 7.172582 | 7.654627 | 7.749903 | 6.041161 | 6.011056 | 5.991604 | 6.770885 |
| 6.640762 | 6.724299 | 6.39616 | 7.631988 | 8.361766 | 6.72559 | 6.822347 | 6.770411 | 6.713821 |
| NaN | NaN | NaN | 7.010938 | 6.884072 | NaN | 6.305216 | NaN | 6.875303 |
| 8.738999 | 8.220081 | 8.267006 | 8.58287 | 8.50429 | 7.714299 | 8.188056 | 8.157922 | 8.886155 |
| 6.065648 | 6.278441 | 5.718648 | 7.269884 | 7.283102 | 6.070902 | 5.958505 | 5.619564 | 6.382764 |
| 7.420857 | 7.376076 | 7.33907 | 6.781002 | 6.821522 | 9.212422 | 8.561071 | 8.941237 | 6.215229 |
| 8.86949 | 8.551058 | 8.743633 | 9.278905 | 8.831567 | 8.927461 | 8.928153 | 8.913342 | 8.569649 |
| NaN | NaN | NaN | NaN | NaN | NaN | NaN | NaN | 5.984957 |
| 8.137675 | 7.982927 | 8.252326 | 7.053022 | 7.075483 | 7.647187 | 8.053318 | 8.104796 | 5.836778 |
| 7.052189 | 7.341968 | 7.522724 | 7.53238 | 6.408059 | 6.516016 | 5.690639 | 6.229952 | 6.978572 |
| NaN | NaN | 6.346336 | 3.655307 | NaN | 6.262918 | 5.855042 | 5.851367 | 7.358181 |

| | | | | | | | | |
|---|---|---|---|---|---|---|---|---|
| 8.781004 | 8.765238 | 8.75169 | 10.05182 | 9.951842 | 8.670497 | 8.578739 | 8.55699 | 9.062077 |
| NaN | NaN | NaN | NaN | 6.126054 | NaN | NaN | NaN | 5.722207 |
| 8.510681 | 8.431995 | 8.651978 | 7.968503 | 7.994585 | 8.240815 | 8.465993 | 8.25529 | 7.963598 |
| 8.515469 | 8.436669 | 8.260726 | 8.456166 | 8.314624 | 8.284518 | 8.150196 | 8.220971 | 7.879632 |
| 8.134648 | 8.145148 | 8.204173 | 8.374118 | 8.042774 | 8.355083 | 8.276842 | 8.338505 | 8.189167 |
| 4.058966 | 4.229143 | 6.651802 | 5.081199 | 4.35657 | 6.94108 | 6.718397 | 6.820433 | 5.233392 |
| 8.528992 | 8.749385 | 8.564843 | 8.276447 | 8.273231 | 8.036815 | 8.373899 | 8.441081 | 8.214997 |
| 7.237744 | 7.211119 | 7.326962 | 7.076788 | 7.068695 | 7.377334 | 6.951051 | 7.481006 | 7.148871 |
| 6.982082 | 5.734952 | 6.837475 | 5.996436 | 5.646759 | 6.728053 | 7.110395 | 7.278032 | 6.401521 |
| 8.700074 | 8.425079 | 8.637048 | NaN | 6.819992 | 7.848502 | 8.458643 | 8.285894 | 8.178894 |
| 8.306091 | 8.270168 | 8.21077 | 7.097201 | 7.305501 | 8.381441 | 8.289193 | 8.258142 | 8.130309 |
| 8.309997 | 8.5658 | 8.588019 | 8.581468 | 8.399001 | 8.021602 | 7.978893 | 8.006409 | 7.947351 |
| 7.156032 | 7.161791 | 7.167915 | 7.610179 | 7.528699 | 6.786364 | 6.917834 | 6.762993 | 7.709613 |
| 6.715706 | 6.560489 | 6.74539 | 6.431578 | 6.452012 | 6.701885 | 6.870274 | 6.696034 | 6.468507 |
| 9.733933 | 9.774187 | 9.671485 | 8.413663 | 8.244115 | 9.881154 | 9.940192 | 9.992302 | 9.513307 |
| 7.139132 | 7.5306 | 6.985242 | 6.433791 | 6.480005 | 7.288476 | 7.387889 | 7.292836 | 7.078739 |
| 8.237064 | 8.195158 | 8.245677 | 8.521712 | 8.559578 | 7.890568 | 7.693166 | 7.826159 | 8.208769 |
| 7.903767 | 5.192658 | 7.875352 | 7.916044 | 8.221266 | 8.006786 | 7.783067 | 7.858115 | 6.769256 |
| 7.349687 | 6.582131 | 7.083137 | 6.882498 | NaN | 6.586329 | 6.964897 | 6.766867 | 7.215685 |
| 10.14757 | 10.17199 | 10.09548 | 11.52103 | 11.33388 | 8.478914 | 8.355422 | 8.279366 | 9.31087 |
| NaN | 3.4508 | NaN | 4.889856 | NaN | 3.285768 | 4.159629 | NaN | 5.238021 |
| 6.472126 | 6.363506 | 6.52332 | 5.330267 | 6.052607 | 6.418986 | 6.497621 | 6.382188 | 6.602282 |
| 7.447819 | 7.561169 | 7.446079 | 8.04114 | 8.333022 | 7.225623 | 7.314576 | 7.384069 | 6.73841 |
| NaN | NaN | 4.625323 | NaN | 4.971943 | NaN | NaN | 5.722989 | 4.851266 |
| 9.380359 | 9.363405 | 9.393168 | 8.603399 | 8.6803 | 9.633099 | 9.509379 | 9.610349 | 9.027946 |
| NaN | 4.914565 | 6.165941 | 4.481863 | 4.694413 | 3.507857 | 4.137379 | 3.97569 | 4.588253 |
| 8.238433 | 7.860187 | 8.312027 | 7.970147 | 8.15554 | 7.489202 | 7.294811 | 7.632079 | 8.192181 |
| 7.168447 | 6.869378 | 6.685529 | 6.748391 | 7.957314 | 6.524597 | 6.442797 | 6.337977 | 6.490992 |
| 7.627571 | 7.384054 | 7.393237 | 7.189498 | 7.238034 | 7.699512 | 7.259586 | 7.201465 | 7.503604 |
| 6.245183 | 6.269433 | 6.317038 | 7.292354 | 7.362 | 6.961861 | 6.831135 | 6.779408 | 7.447099 |
| 8.056843 | 8.046698 | 7.939971 | 7.948886 | 8.0661 | 8.318252 | 7.982965 | 8.098875 | 7.839212 |
| NaN | 6.940653 | NaN | NaN | 5.236442 | 6.875446 | 6.8297 | 6.954453 | 5.449039 |
| 5.240477 | 7.064968 | 5.795971 | NaN | NaN | 6.698185 | 7.063496 | 6.978133 | 6.611548 |
| 5.105012 | 4.382121 | 4.627548 | 4.228425 | 3.604776 | 4.138797 | 4.643935 | 4.948054 | 4.92924 |
| 7.361473 | 7.278842 | 6.869554 | 7.277106 | 7.463154 | 7.31076 | 6.856495 | 6.989972 | 6.554848 |
| NaN | 6.138628 | 7.577181 | 4.921592 | NaN | 6.162624 | 5.834343 | 6.231177 | 6.214554 |
| 6.943137 | 6.969765 | 7.058942 | 6.250057 | 6.230743 | 7.100167 | 6.946632 | 7.324936 | 5.989665 |
| 3.952573 | 3.996047 | 4.692873 | 5.686862 | 4.703301 | 5.418977 | 4.786484 | 5.426743 | 4.090368 |
| 7.653337 | 7.896923 | 7.701949 | 7.535632 | 7.384184 | 8.216991 | 8.20945 | 8.322281 | 7.6699 |
| 8.97462 | 8.988556 | 9.054384 | 8.564761 | 8.406168 | 8.977938 | 8.802319 | 8.815031 | 8.461884 |
| 6.417401 | 6.717546 | 6.891224 | 6.44421 | 6.275476 | 5.943094 | 6.059664 | 6.444379 | 6.119883 |
| 6.604375 | 6.540952 | 6.84033 | 5.021155 | 6.23549 | 6.586665 | 6.622094 | 6.878595 | 6.654725 |
| 8.174468 | 7.876082 | 8.12384 | 6.372896 | 7.118051 | 8.200665 | 8.029947 | 8.146932 | 8.317 |
| 5.739803 | 5.945353 | 5.881569 | 7.066572 | 7.274781 | 7.056756 | 7.090713 | 7.053384 | 4.108031 |
| NaN | NaN | 5.714652 | 5.912225 | 8.04063 | 6.687428 | 6.954383 | 7.421146 | 5.441659 |
| 6.322053 | 6.078043 | 6.651548 | 6.333565 | 6.098639 | 6.106489 | 6.31117 | 6.012001 | 5.709867 |
| 7.232603 | 7.108625 | 7.090022 | 7.724009 | 6.614043 | 7.150819 | 7.350664 | 5.509879 | 6.072147 |

-Bruker Confidential-

| | | | | | | | | |
|---|---|---|---|---|---|---|---|---|
| 7.369229 | 7.46829 | 7.572666 | 7.518256 | 7.857205 | 7.699262 | 7.762975 | 7.58309 | 6.710101 |
| 7.266568 | 7.303559 | 7.354448 | 7.616185 | 7.322198 | 7.358886 | 7.115804 | 6.845631 | 7.308944 |
| 6.347622 | 6.615462 | 6.461388 | 8.114688 | 8.239799 | 6.360318 | 6.60435 | 6.456249 | 5.626766 |
| 9.323603 | 9.330518 | 9.401145 | 9.682142 | 9.897413 | 9.509414 | 9.358393 | 9.649969 | 8.60166 |
| 7.50518 | 3.792229 | 6.498023 | NaN | NaN | 4.284264 | NaN | NaN | 5.896484 |
| 7.072989 | 6.382339 | 6.737967 | 6.063756 | 6.619636 | 6.732971 | 7.421268 | 7.739988 | 5.755118 |
| 7.702895 | 8.004461 | 7.057455 | 7.101027 | 7.304657 | 6.908137 | 7.161759 | 7.203249 | 6.864341 |
| 7.870286 | 8.151711 | 7.835637 | 7.654501 | 7.82712 | 7.946912 | 8.161057 | 8.051675 | 7.269456 |
| 8.14217 | 8.291526 | 8.199457 | 8.557832 | 8.593287 | 8.030439 | 7.938011 | 8.057067 | 8.152607 |
| 6.685726 | 6.59606 | 6.65274 | NaN | 5.220381 | 7.546602 | 7.65152 | 7.64327 | 5.942309 |
| 10.84378 | 10.65015 | 10.53208 | 10.30542 | 10.26113 | 10.42142 | 10.42549 | 10.34894 | 10.56663 |
| 7.266613 | 6.411826 | 7.036868 | 6.751951 | 6.762355 | 6.923698 | 7.420152 | 7.159242 | 6.378621 |
| 9.436451 | 9.537305 | 9.324294 | 9.622527 | 9.761395 | 9.210464 | 9.113338 | 9.16394 | 9.784084 |
| 5.917215 | 6.598894 | 6.369635 | 7.173498 | 7.091212 | 6.79375 | 6.995052 | 6.928605 | 7.48823 |
| 5.040441 | 3.596139 | 4.859001 | 3.950979 | 4.657988 | 5.000099 | 4.794384 | 4.811745 | 4.26703 |
| 8.917987 | 8.966246 | 8.973696 | 8.344339 | 8.352126 | 9.187578 | 8.989327 | 9.117976 | 8.829255 |
| 5.021265 | 6.031586 | 5.499124 | 5.647155 | 6.012198 | 5.069402 | 5.469715 | 5.01613 | 5.352498 |
| 6.192174 | 6.447289 | 5.931552 | 7.198785 | 7.562217 | 6.031839 | 5.691594 | 5.359279 | 6.873239 |
| 8.784703 | 8.784789 | 8.765285 | 7.419109 | 7.501498 | 8.166127 | 8.49146 | 8.001306 | 6.525764 |
| 9.6345 | 9.555532 | 9.576097 | 10.13307 | 9.390777 | 8.787625 | 8.64048 | 8.838555 | 8.726933 |
| 7.010466 | 7.451411 | 7.55099 | 6.00697 | 7.451432 | 6.639341 | 6.732897 | 6.591215 | 7.391852 |
| 6.290748 | 5.115763 | 6.58694 | 6.559209 | 6.587139 | 6.947757 | 6.899196 | 7.085347 | 7.150006 |
| 6.842991 | 6.771452 | 7.20932 | 7.213126 | NaN | 7.130192 | 7.225626 | 7.057664 | 6.520709 |
| 6.884078 | 6.505214 | 6.911771 | NaN | 6.520121 | 7.315449 | 7.143898 | 7.442731 | 6.521273 |
| 5.936914 | 6.2666 | 6.160775 | 5.317667 | 4.633492 | 6.368512 | 6.352438 | 6.246012 | 7.041875 |
| 10.08584 | 9.942274 | 9.911053 | 11.05581 | 10.9817 | 9.748878 | 9.570898 | 9.852937 | 10.1121 |
| 10.45833 | 10.34627 | 10.37632 | 10.04855 | 9.982198 | 10.00215 | 10.05249 | 10.19757 | 10.1484 |
| 6.217857 | 6.424773 | 5.815472 | NaN | 5.049892 | 6.605152 | 7.059554 | 6.962027 | 3.983195 |
| 8.297052 | 8.099099 | 8.73556 | 5.216078 | NaN | NaN | NaN | NaN | 6.597503 |
| 6.678452 | 7.261068 | 6.893986 | 5.899694 | 6.187116 | 7.79319 | 7.76299 | 7.532342 | 6.449714 |
| 8.397241 | 7.398588 | 7.148623 | 7.921082 | 7.851732 | 7.706879 | 7.632562 | 7.497838 | 7.566761 |
| 7.715071 | 6.686637 | 6.465468 | 9.425188 | 8.872516 | 5.103295 | 5.269071 | 8.288129 | 1.270609 |
| 7.138072 | 7.212281 | 7.297487 | 6.903267 | 6.910778 | 8.2561 | 8.053455 | 8.210659 | 7.652419 |
| 5.061831 | 5.333788 | 5.371208 | 6.899414 | 8.205608 | NaN | NaN | NaN | 6.711661 |
| 8.072687 | 7.074032 | 6.361694 | 8.142118 | 8.235936 | NaN | NaN | NaN | 7.089828 |
| 5.854377 | 5.196348 | 5.922641 | NaN | 5.991919 | 5.759145 | 5.852956 | 5.708913 | 5.538024 |
| 5.198086 | NaN | 5.033279 | NaN | NaN | 6.55163 | 6.251995 | 6.32929 | 5.244216 |
| 8.285721 | 7.937517 | 8.093098 | 7.539872 | 7.416124 | 7.968068 | 7.887943 | 8.022941 | 8.203162 |
| 7.938798 | 7.692395 | 7.651908 | 7.077286 | 7.08279 | 7.968177 | 7.825856 | 7.824717 | 7.883901 |
| 10.32478 | 10.33504 | 10.30692 | 9.967829 | 10.02832 | 10.24988 | 10.36446 | 10.3102 | 10.47848 |
| 9.845488 | 9.716509 | 9.891188 | 10.9074 | 10.81511 | 9.294931 | 9.147662 | 9.267923 | 9.383645 |
| 8.166718 | 8.157921 | 8.163113 | 7.919114 | 7.877697 | 8.126649 | 7.953128 | 8.170319 | 7.887288 |
| 11.93938 | 11.97318 | 11.79595 | 11.17992 | 11.38476 | 11.77905 | 11.5251 | 11.73608 | 11.25075 |
| 8.999969 | 8.860025 | 8.991784 | 9.533532 | 9.595958 | 8.767467 | 8.227888 | 8.449777 | 8.146869 |
| 7.47354 | 7.676826 | 7.474886 | 8.211632 | 8.107806 | 6.422197 | 6.540152 | 6.420619 | 8.006097 |
| 10.30122 | 10.3725 | 10.27467 | 11.0984 | 11.09886 | 10.49919 | 10.58504 | 10.64544 | 9.894555 |
| 9.35252 | 9.780832 | 9.868456 | 8.046611 | 8.604991 | 8.536437 | 8.603567 | 8.722231 | 8.738256 |

| | | | | | | | | |
|---|---|---|---|---|---|---|---|---|
| 5.772328 | NaN | NaN | 5.937673 | 6.233319 | 4.346444 | NaN | 5.173608 | NaN |
| 7.016745 | 6.118337 | 7.649663 | 7.014437 | 6.434052 | 6.234736 | 6.709697 | 6.287772 | 8.044856 |
| 7.972768 | 8.034257 | 7.898776 | 7.261147 | 7.342482 | 7.898376 | 7.936369 | 8.090751 | 8.070784 |
| 8.392387 | 8.144098 | 8.263044 | 9.208943 | 9.135753 | 8.215392 | 8.253778 | 8.468269 | 8.20703 |
| 9.674524 | 9.687176 | 9.703453 | 7.893481 | 8.021254 | 9.647462 | 9.667936 | 9.580241 | 9.019498 |
| 11.02404 | 11.00587 | 11.10367 | 10.03046 | 10.21489 | 10.04424 | 9.888749 | 9.86241 | 10.5988 |
| 5.796689 | 5.902022 | 5.809823 | 5.895141 | 6.333243 | 6.168873 | 6.295513 | 6.776456 | 5.194605 |
| 6.66851 | 6.829976 | 6.966216 | 7.330954 | 7.771396 | 6.944674 | 6.613603 | 7.079301 | 5.995608 |
| 6.778052 | 6.696892 | 6.778909 | 5.987985 | 5.791033 | 6.817811 | 7.200571 | 7.06874 | 6.168287 |
| 5.813022 | 5.127601 | 5.943456 | 5.699446 | 5.787057 | 4.253747 | 5.386096 | 5.350021 | 7.177548 |
| 7.780084 | 8.007403 | 7.907412 | 6.801936 | 7.567209 | 7.833117 | 8.051728 | 8.01509 | 7.130877 |
| 6.443959 | 6.728945 | 6.745103 | 6.359961 | 6.646673 | 6.705368 | 6.613882 | 6.628565 | 5.907716 |
| 6.970369 | 7.222807 | 7.364788 | 7.424795 | 6.678797 | 6.719397 | 7.188869 | 7.298177 | 6.826038 |
| NaN | NaN | NaN | 5.866538 | 4.564339 | NaN | NaN | NaN | 5.549456 |
| 8.335474 | 8.003715 | 8.180552 | 7.593867 | 7.475281 | 7.969353 | 7.866539 | 8.006081 | 8.281135 |
| 6.129863 | 5.456217 | 6.169805 | NaN | NaN | 5.947423 | 5.849588 | 6.033595 | 5.781428 |
| 7.306473 | 7.301655 | 7.749531 | 8.503017 | 8.482561 | 7.819918 | 7.635707 | 7.67708 | 7.483462 |
| 7.021808 | 7.428411 | 7.461582 | 6.686568 | 7.175673 | 7.907824 | 7.996175 | 7.883833 | 7.358364 |
| 6.058178 | 5.967171 | 5.78045 | 6.507887 | 6.803176 | 6.798538 | 6.998851 | 6.857179 | 6.382988 |
| 7.970612 | 7.59268 | 7.85924 | 8.236159 | 8.344625 | 7.030309 | 7.587208 | 7.457068 | 7.97686 |
| 6.608972 | 6.769603 | 7.049415 | 6.136742 | 6.588953 | 6.56562 | 6.231713 | 6.181643 | 6.509004 |
| 6.947464 | 6.956203 | 6.869259 | 7.189234 | 6.922966 | 6.786119 | 6.527395 | 6.814947 | 6.026962 |
| 6.593659 | 6.525875 | 6.628907 | 5.652779 | 6.198142 | 6.936143 | 7.000143 | 7.276571 | 6.280213 |
| 4.676453 | 5.428363 | 4.747614 | 5.500025 | 5.510155 | 5.986385 | 6.184026 | 6.312416 | 5.651169 |
| 10.32195 | 10.19041 | 10.20909 | 9.480507 | 9.702938 | 9.80761 | 9.683942 | 9.677612 | 9.216814 |
| 7.328309 | NaN | 6.773015 | 7.402375 | 7.493385 | 7.291711 | 7.04553 | 7.534829 | NaN |
| 9.870787 | 9.908536 | 9.88474 | 10.54354 | 10.44092 | 9.30023 | 9.341738 | 9.306801 | 10.21256 |
| 5.49354 | 5.153959 | 5.617219 | 5.54019 | 5.665266 | 4.958422 | 5.213978 | 5.266828 | 5.540426 |
| 6.624337 | 7.05338 | 6.891619 | 3.601997 | 4.794933 | 7.750958 | 7.674161 | 7.174093 | 6.099686 |
| 7.610592 | 6.084337 | 7.508747 | 7.359164 | 7.68852 | 7.863629 | 7.420875 | 7.489612 | 7.178045 |
| 7.67158 | 7.770494 | 7.74771 | 6.387787 | 6.243058 | 7.382003 | 7.308946 | 7.339354 | 7.206604 |
| 6.474456 | 5.729457 | 6.539651 | 6.266964 | 5.68426 | 5.536129 | 5.786174 | 5.577677 | 5.776538 |
| NaN | NaN | NaN | 6.109318 | NaN | 6.766614 | 7.190268 | 7.03414 | 6.609336 |
| 6.452203 | 5.737407 | 5.930926 | 4.014062 | NaN | 6.323231 | 6.268371 | 6.07571 | 6.259811 |
| 8.345497 | 8.285478 | 8.451826 | 7.075688 | 7.274701 | 8.154621 | 8.066484 | 7.961342 | 7.618824 |
| 6.715327 | 7.0092 | 7.107677 | 6.796273 | 7.240664 | 7.521137 | 7.220594 | 7.363385 | 6.866066 |
| 8.306864 | 8.341701 | 8.28973 | 10.2859 | 10.06934 | 8.425543 | 8.688781 | 8.42223 | 8.019938 |
| 8.384059 | 8.072878 | 8.398042 | 8.130566 | 7.884811 | 6.92093 | 7.190388 | 7.268597 | 7.339551 |
| 8.848066 | 8.622653 | 8.81433 | 9.997091 | 9.61139 | 8.783348 | 8.512824 | 7.896993 | 6.65649 |
| 7.178804 | 8.178465 | 6.845196 | 6.581494 | 6.883482 | 7.451129 | 7.625075 | 7.416672 | 7.174426 |
| 7.953626 | 7.905484 | 8.253334 | 7.52125 | 7.971217 | 7.758196 | 7.59028 | 7.197804 | 7.438162 |
| 7.490771 | 7.499698 | 7.246294 | 6.861157 | 6.878327 | 7.678395 | 7.526346 | 7.77213 | 7.543361 |
| 8.22411 | 8.374377 | 8.493466 | 8.43504 | 8.45546 | 7.931284 | 7.659435 | 8.137743 | 7.406441 |
| 7.761159 | 7.920676 | 7.788967 | 7.199014 | 7.613444 | 7.370892 | 8.00389 | 7.999883 | 7.87788 |
| 7.184006 | 7.093656 | 7.13556 | 7.148906 | 7.424883 | 6.430524 | 6.520482 | 6.591174 | 5.836278 |
| 7.42703 | 7.36976 | 7.690381 | NaN | NaN | 7.939311 | 8.457876 | 8.633213 | 7.740742 |
| 10.72634 | 10.34836 | 10.46877 | 10.49682 | 10.65223 | 9.677453 | 9.685963 | 9.492918 | 9.862796 |

| | | | | | | | | |
|---|---|---|---|---|---|---|---|---|
| 5.984319 | 6.232224 | 6.135486 | 4.246036 | 5.057621 | 6.95501 | 6.04645 | 6.515022 | 5.740941 |
| 9.752624 | 9.872489 | 9.937159 | 10.21141 | 10.25849 | 9.458021 | 9.47385 | 9.455905 | 9.672981 |
| 7.313323 | 7.362508 | 7.309764 | 8.941708 | 9.473426 | 7.308476 | 7.297423 | 7.476515 | 7.601698 |
| 5.784491 | 5.816131 | 5.919024 | 5.55725 | 6.340203 | 6.487142 | 6.289687 | 5.614086 | 5.79045 |
| 6.208869 | 6.310086 | 6.10498 | NaN | NaN | 5.63138 | 5.682105 | NaN | 5.822076 |
| 8.584873 | 8.619332 | 8.687863 | 7.596307 | 7.715855 | 9.129552 | 9.001727 | 9.092941 | 8.708431 |
| 9.074619 | 9.144239 | 8.873091 | 9.376846 | 9.731747 | 9.215075 | 9.329761 | 8.935954 | 8.787468 |
| 6.443616 | 5.58182 | 6.522568 | 5.303751 | 5.374255 | 5.087571 | NaN | 5.167525 | 5.559119 |
| 8.356045 | 8.183021 | 8.199915 | 8.777515 | 8.451349 | 8.400501 | 8.582705 | 8.153035 | 9.101969 |
| NaN | NaN | NaN | NaN | 5.601366 | NaN | 5.31696 | 5.686382 | 6.690443 |
| 8.857325 | 8.984485 | 8.861411 | 9.658 | 9.537794 | 8.564661 | 8.676164 | 8.660136 | 9.069599 |
| 7.660813 | 7.671357 | 7.798321 | 7.290605 | 6.694444 | 8.018003 | 8.304378 | 8.173502 | 7.591685 |
| 7.246226 | 7.197957 | 7.188115 | NaN | 8.42431 | 8.422204 | 9.271074 | 9.559137 | 6.718493 |
| 7.428164 | 8.331604 | 8.102811 | 6.712471 | NaN | 7.685688 | 7.512915 | 7.436376 | 7.216907 |
| 9.343023 | 8.343366 | 8.982384 | 8.281911 | 7.524334 | 8.39111 | 8.429019 | 8.582323 | 9.118895 |
| 5.92747 | 5.976824 | 6.075934 | 5.653227 | 5.270292 | 6.312943 | 5.990131 | 6.23545 | 5.884051 |
| 6.973778 | 7.049852 | 7.100802 | 6.383584 | 6.913051 | 6.986925 | 7.054845 | 6.915701 | 7.448087 |
| 6.388434 | 6.653572 | 6.944833 | 6.124511 | 6.176366 | 7.144027 | 7.048644 | 6.756392 | 7.032483 |
| 6.674441 | 5.898703 | 6.941541 | 6.236471 | 5.299727 | 6.733621 | 6.57112 | 6.904345 | 6.052348 |
| 7.602973 | 7.368569 | 7.094074 | 5.977312 | 6.506237 | 7.453288 | 7.545958 | 7.454571 | 6.739285 |
| NaN | 6.640375 | 4.028424 | 7.671994 | NaN | 8.959891 | 8.744308 | 8.318581 | 6.20797 |
| 9.62705 | 9.621016 | 9.651674 | 10.86096 | 10.4702 | 9.118999 | 9.09203 | 9.44584 | 9.561651 |
| 7.260895 | 7.496969 | 6.858083 | 8.714808 | 8.656984 | 7.149494 | 8.28127 | 8.090506 | 7.496168 |
| 5.376243 | 5.21867 | NaN | 6.583951 | 7.033738 | NaN | NaN | NaN | 6.243793 |
| 8.12564 | 7.969292 | 7.932676 | 7.898776 | 8.085744 | 7.783585 | 7.598477 | 7.770685 | 7.296738 |
| 8.374654 | 8.170252 | 7.570682 | 7.489324 | 7.731807 | 7.321014 | 7.072654 | 7.558248 | 6.925899 |
| 9.633925 | 9.406816 | 9.538342 | 10.32642 | 10.31259 | 9.739767 | 9.702088 | 10.00269 | 9.345179 |
| 10.33786 | 9.834492 | 9.183553 | 9.184714 | 9.118141 | 9.739293 | 10.15024 | 9.989785 | 10.16576 |
| NaN | NaN | NaN | 6.807352 | 7.257136 | NaN | NaN | 9.150025 | 8.891344 |
| NaN | NaN | NaN | 6.164789 | 6.288718 | 6.716616 | 6.573461 | 6.308103 | 6.344303 |
| 8.565103 | 8.671133 | 8.490238 | 7.897425 | 7.703949 | 8.199192 | 8.145775 | 7.731445 | 8.272685 |
| 7.313706 | 7.272074 | 7.399652 | 5.338767 | 5.909274 | 7.677251 | 7.368132 | 7.459565 | 6.204004 |
| 6.947719 | 6.895818 | 6.735081 | 6.22539 | 6.54931 | 7.05853 | 7.14731 | 6.97049 | 7.443501 |
| 9.084805 | 8.568354 | 8.59043 | 9.268023 | 9.097887 | 8.231856 | 8.038279 | 7.996444 | 8.341359 |
| 8.068893 | 7.941375 | 7.759376 | 7.786653 | 7.641249 | 7.63547 | 8.076213 | 7.604104 | 7.59634 |
| 9.056392 | 9.322588 | 9.326043 | 8.000999 | 8.145365 | 7.94111 | 7.958335 | 7.431257 | 9.240615 |
| 8.049983 | 8.177575 | 8.077027 | 6.777614 | 6.757675 | 8.143897 | 7.382785 | 7.634832 | 6.405655 |
| 8.632737 | 8.807758 | 8.759632 | 7.744556 | 7.863459 | 8.333492 | 8.348258 | 8.021361 | 8.813103 |
| 8.389836 | 8.662393 | 8.236784 | 8.347485 | 8.708673 | 7.391695 | 7.298057 | 7.64379 | 7.600493 |
| 7.956691 | 7.69755 | 7.980415 | 8.091392 | 7.860587 | 7.469573 | 7.175285 | 7.711183 | 7.037027 |
| 7.092216 | 7.225354 | 6.464916 | 7.986536 | 8.082168 | 6.594434 | 6.444906 | 6.872997 | 5.86287 |
| 5.930718 | NaN | NaN | 6.456306 | 6.519846 | 5.713247 | NaN | 6.435112 | 5.920797 |
| 6.168973 | 6.233164 | 6.025442 | 6.939141 | 7.428272 | 6.845541 | 6.581193 | 6.710838 | 5.513439 |
| 7.206728 | 10.1673 | NaN | 7.701659 | 7.867252 | 7.312047 | NaN | 7.338293 | 7.635066 |
| 6.122153 | 6.507505 | 6.736374 | 6.245765 | 6.88337 | 7.195316 | 6.968381 | 6.854289 | 6.875275 |
| 6.988966 | 6.717498 | 7.089166 | 6.49745 | 6.251325 | 7.148968 | 6.860592 | 7.211178 | 6.585054 |
| 9.235182 | 9.322498 | 9.317937 | 9.245206 | 9.157088 | 9.459855 | 9.265931 | 9.473469 | 8.988126 |

| | | | | | | | | |
|---|---|---|---|---|---|---|---|---|
| 7.007957 | 6.855846 | 6.99303 | 6.430055 | 6.487397 | 7.36905 | 7.236202 | 7.161605 | 6.605858 |
| 7.968972 | 8.348206 | 7.998721 | 8.206699 | 8.514841 | 8.266707 | 8.116753 | 8.19919 | 8.161523 |
| 6.881856 | 6.988194 | 7.075789 | 6.753479 | 7.050732 | 8.255233 | 7.923184 | 5.182947 | 7.34489 |
| 6.177909 | 6.061088 | NaN | 6.672843 | 6.894949 | NaN | NaN | NaN | NaN |
| 9.221896 | 9.0665 | 9.067474 | 11.64647 | 11.65201 | 9.637152 | 9.478684 | 9.425066 | 9.373381 |
| 8.09992 | 7.792972 | 8.139912 | 9.238042 | 9.311117 | 7.913035 | 7.955317 | 8.135482 | 7.04504 |
| 7.481298 | 7.134933 | 7.330157 | 8.356947 | 8.144556 | 7.320091 | 7.330736 | 7.183993 | 6.988828 |
| 7.47909 | 6.422138 | 7.541335 | 4.022473 | 5.339437 | 7.465567 | 6.81421 | 7.188606 | 6.046151 |
| 9.947828 | 9.814966 | 10.03307 | 11.55033 | 10.89665 | 10.18686 | 10.19295 | 10.10889 | 10.09884 |
| 8.150446 | 9.200187 | 7.709475 | 7.550274 | 7.994649 | NaN | 7.257498 | 7.02222 | 7.750822 |
| 8.359921 | 6.208159 | 8.590943 | 8.179708 | 7.650517 | 7.067238 | 7.229716 | 6.308218 | 6.043502 |
| 7.852243 | 7.514302 | 7.761217 | 8.04378 | 8.139495 | 7.75807 | 8.0767 | 7.965358 | 7.55766 |
| 8.410944 | 8.652486 | 8.75591 | 7.690174 | 7.832955 | 9.053001 | 8.777304 | 9.060667 | 9.145385 |
| 9.496456 | 9.470689 | 9.527919 | 8.808649 | 8.319704 | 9.191071 | 9.446911 | 9.348721 | 9.545011 |
| 7.756574 | 7.560667 | 7.895195 | 6.151425 | 5.872052 | 6.724526 | 6.639353 | 7.107221 | 6.661818 |
| 6.4478 | 3.153282 | 5.878297 | 5.921988 | 5.448062 | 6.150026 | 6.117063 | 6.15731 | 6.067432 |
| NaN | 5.804892 | 6.067721 | 4.205192 | 7.380124 | NaN | NaN | NaN | 6.204195 |
| 7.61761 | 7.704697 | 7.832922 | 6.091307 | 6.291902 | 7.745572 | 7.76699 | 7.787619 | 7.282806 |
| 7.023706 | 7.302578 | 7.199465 | 5.569571 | NaN | 7.628012 | 8.067593 | 7.718489 | 7.808192 |
| 6.716329 | 6.729456 | 6.487194 | 6.952176 | 7.023188 | 6.347701 | 6.384217 | 6.621367 | 5.868608 |
| 7.171103 | 6.91314 | 6.558144 | 8.288093 | 7.567609 | 6.339561 | 6.211802 | 6.495505 | 6.77331 |
| NaN | NaN | 6.426505 | 7.942461 | 7.927287 | NaN | 6.842264 | 6.76371 | 7.421255 |
| 5.478212 | 4.857495 | 5.232945 | NaN | NaN | 6.198011 | 6.764024 | 5.979522 | 5.810851 |
| 6.551358 | 6.783643 | 7.03018 | 7.401256 | 7.518664 | 6.953655 | 6.925817 | 6.809455 | 6.831119 |
| NaN | NaN | NaN | 2.579122 | 1.885696 | NaN | NaN | 3.240679 | 3.576621 |
| 7.043246 | 7.089005 | 7.30879 | 7.848001 | 7.749197 | 7.142212 | 7.244543 | 7.227916 | 7.434091 |
| 8.171729 | 6.924812 | 7.430124 | 6.110093 | NaN | 6.501575 | 6.679967 | 7.051985 | 6.350264 |
| 9.262898 | 9.231951 | 8.896087 | 9.390706 | 9.523604 | 9.36659 | 9.184199 | 9.142057 | 8.943168 |
| 3.907875 | 3.128854 | 3.881478 | 4.94305 | 5.060889 | NaN | 3.14926 | 2.86665 | 2.465475 |
| 7.0632 | 7.114162 | 6.834269 | 8.546481 | 8.516739 | 6.789483 | 6.357976 | 6.515337 | 6.918614 |
| 7.505984 | 7.748798 | 7.622293 | 6.513507 | 6.125444 | 7.670975 | 7.806536 | 7.822466 | 7.08845 |
| 4.824663 | NaN | NaN | 4.887939 | 4.618854 | NaN | NaN | NaN | 5.206123 |
| 7.763144 | 7.868235 | 7.608793 | 7.309644 | 7.528463 | 7.442538 | 7.616045 | 7.636202 | 7.467232 |
| 6.868918 | 6.663104 | 7.04133 | 5.342432 | NaN | 7.250144 | 6.517909 | 7.262376 | NaN |
| 7.645929 | 7.136021 | 7.422816 | 6.911419 | 7.089156 | 7.219842 | 7.301499 | 6.752205 | 7.259505 |
| 8.133797 | 8.282051 | 8.240029 | 8.196805 | 8.669637 | 8.005154 | 8.287372 | 8.232869 | 8.370362 |
| 6.786748 | 6.851348 | 6.822383 | 6.52919 | 6.306397 | 7.189823 | 7.15694 | 7.146984 | 7.488527 |
| 8.655595 | 8.616864 | 8.514299 | 8.205276 | 7.983463 | 8.344347 | 8.286714 | 8.134891 | 8.04465 |
| 7.060638 | 6.938148 | 6.973269 | 6.326985 | 6.816604 | 7.41331 | 7.232023 | 7.05513 | 6.814518 |
| 5.111234 | 5.939301 | 6.256579 | 6.315956 | 6.172361 | 5.312745 | NaN | NaN | 5.883584 |
| 7.454186 | 7.443627 | 7.581075 | 8.177951 | 7.916092 | 7.06991 | 7.068189 | 7.234942 | 7.061256 |
| 8.144524 | 6.844086 | 8.035459 | 7.257698 | 7.340137 | 8.029858 | 7.98374 | 8.250136 | 4.990474 |
| 7.415925 | 7.513569 | 7.46587 | 6.327353 | NaN | 7.030079 | 7.357546 | 7.145344 | 6.839179 |
| 6.534364 | 6.918066 | 6.604984 | 6.700922 | 6.41634 | 7.157356 | 7.363959 | 7.28283 | 6.98378 |
| 8.426193 | 8.44149 | 8.425688 | 8.322269 | 8.280164 | 8.845407 | 8.541967 | 8.803917 | 7.778488 |
| 7.21064 | 7.047462 | 7.152754 | 6.178874 | 6.85223 | 7.110012 | 7.341174 | 7.354167 | 6.989388 |
| 4.585774 | 5.206593 | NaN | 6.26355 | 6.432376 | 5.771982 | NaN | 6.169862 | 6.300303 |

| | | | | | | | | |
|---|---|---|---|---|---|---|---|---|
| 7.908147 | 7.885512 | 7.89923 | 6.746493 | 7.235422 | 7.899439 | 7.578696 | 7.854403 | 7.05163 |
| 6.060809 | 6.517732 | 6.262829 | 6.135986 | 6.147889 | 6.755298 | 6.985785 | 6.849245 | 6.383427 |
| 8.134804 | 8.111427 | 8.174108 | 6.783681 | 7.011004 | 7.963513 | 8.127712 | 8.140723 | 7.632877 |
| 6.549881 | 6.589238 | 6.352872 | 7.734406 | 7.634395 | 6.944005 | 6.69954 | 6.961601 | 7.087538 |
| 6.856578 | 6.937455 | 6.803376 | 6.579512 | 6.615024 | 7.11025 | 7.029302 | 7.045333 | 6.574571 |
| 6.783644 | 6.880244 | 6.710207 | 6.645762 | 6.445227 | 6.815445 | 7.208526 | 7.233289 | 6.401512 |
| NaN | 5.375034 | 4.615663 | 4.890085 | 5.573779 | 6.013428 | 6.489403 | 6.495313 | 6.825954 |
| 10.966 | 10.96966 | 10.97741 | 10.64964 | 10.66789 | 10.97637 | 10.93363 | 10.92525 | 10.26098 |
| 5.587359 | 6.050858 | 5.845822 | 7.028526 | 6.751152 | 5.761885 | 6.760028 | 6.042989 | 5.928701 |
| 7.255044 | 6.60636 | 7.744253 | NaN | NaN | 6.780956 | 6.544495 | 6.340931 | 7.420905 |
| 6.186873 | 5.213061 | 6.709161 | NaN | 6.551788 | 6.174716 | 6.027413 | 6.058227 | 6.25156 |
| 11.91904 | 11.83323 | 11.92425 | 11.95194 | 11.92174 | 11.96278 | 11.87668 | 11.83626 | 12.20164 |
| 4.908093 | 4.490501 | 3.269113 | 5.056173 | 5.215505 | 5.088993 | 4.400827 | 4.471684 | 4.367044 |
| 7.523338 | 6.882586 | 7.671978 | 5.730042 | 6.171781 | 7.33572 | 7.363135 | 7.476639 | 7.675842 |
| 7.12719 | 7.193861 | 6.582881 | 5.937106 | 5.953923 | 7.426271 | 7.080065 | 7.009407 | 7.20611 |
| 7.39554 | 7.23428 | 7.356796 | 6.379326 | 6.376381 | 6.602726 | 6.786624 | 6.957153 | 6.308723 |
| 6.210885 | 5.845004 | 6.384447 | NaN | NaN | 6.393291 | 6.221338 | 6.177688 | 5.732326 |
| 8.265376 | 8.119152 | 8.316887 | 7.339561 | 7.078764 | 8.000069 | 8.05911 | 7.904553 | 7.90779 |
| 9.257185 | 9.217871 | 9.21458 | 8.400252 | 8.633375 | 8.944515 | 8.953013 | 8.973579 | 8.565545 |
| 7.212064 | 7.274681 | 7.224236 | 6.707923 | 6.145968 | 7.294015 | 7.446439 | 7.400303 | 8.089454 |
| 9.615989 | 9.749523 | 9.613759 | 11.91626 | 11.82212 | 9.918144 | 10.06284 | 9.879529 | 10.40445 |
| 9.849975 | 9.949552 | 9.866711 | 10.37143 | 10.33372 | 10.32223 | 10.08448 | 10.29094 | 9.15272 |
| 6.883837 | 6.641301 | NaN | 5.788264 | 6.527256 | 7.386911 | 7.260983 | 7.44781 | 7.84047 |
| 9.1711 | 8.923118 | 8.89311 | 8.103101 | 8.365305 | 8.480648 | 8.361771 | 8.447383 | 8.456351 |
| 7.390483 | 7.191706 | 7.540236 | 8.293603 | 8.151896 | 6.538997 | 6.240067 | 6.117771 | 7.881253 |
| 7.399088 | 8.362194 | 7.040502 | 6.411407 | 6.440783 | 6.686348 | 6.767255 | 6.578998 | 6.345298 |
| 6.099125 | 6.14037 | 5.931674 | 7.281135 | 7.669667 | 6.732355 | 6.260144 | 6.507977 | 4.273022 |
| 7.072905 | NaN | 7.620138 | 8.58129 | 8.765995 | 8.105049 | 8.004036 | 7.812086 | 8.054611 |
| 6.116219 | 5.128457 | 4.248912 | 5.890397 | 6.825139 | 5.435963 | 6.166694 | NaN | NaN |
| 8.57856 | 8.53247 | 8.52641 | 8.016152 | 7.524822 | 9.389617 | 9.242909 | 9.389287 | 8.332059 |
| 7.095816 | 7.254698 | 6.937043 | NaN | NaN | 6.228091 | 6.552889 | 6.57496 | 7.276947 |
| NaN | 6.985528 | NaN | NaN | NaN | 6.450649 | 6.684763 | 6.763888 | 5.99123 |
| 7.497918 | 7.374563 | 7.22013 | 7.21984 | 6.967363 | 6.706464 | 7.043731 | 6.908533 | 6.991484 |
| NaN | 5.828641 | 5.28501 | 7.087458 | 8.77384 | NaN | 6.808951 | 6.658071 | 6.278298 |
| NaN | NaN | NaN | NaN | NaN | 6.189551 | 6.713199 | 6.21174 | 7.791558 |
| 7.550614 | 7.471171 | 7.743229 | 7.465054 | 7.063291 | 8.23886 | 8.393955 | 8.521585 | 7.667436 |
| 6.665168 | 5.342032 | 6.089252 | 4.187071 | 5.250662 | 5.366844 | 5.299502 | 5.909013 | 5.699471 |
| 7.68576 | 7.795238 | 7.830383 | 7.50441 | 7.299721 | 6.820968 | 7.081385 | 6.811353 | 7.80626 |
| 6.580308 | 6.87564 | 6.896157 | 6.332254 | 6.321877 | 6.802781 | 6.23915 | 6.571654 | 6.287644 |
| 6.297355 | 5.774805 | NaN | 6.764212 | 6.706686 | 6.350208 | 5.985703 | 5.967443 | 5.964921 |
| 6.570079 | 6.405041 | 6.574355 | 5.740156 | 4.913343 | 6.576728 | 6.286972 | 5.961229 | 6.503115 |
| 9.545918 | 9.457634 | 9.434933 | 8.400456 | 8.512416 | 9.088627 | 9.180988 | 8.974202 | 9.242783 |
| 7.981844 | 8.034895 | 8.278346 | 7.713854 | 7.733457 | 7.633244 | 7.267179 | 7.074212 | 7.954093 |
| 7.267764 | 6.712394 | 5.527887 | 6.164214 | 6.156772 | 7.468824 | 7.33259 | 7.256865 | NaN |
| 8.008142 | 7.905885 | 7.877191 | 7.508487 | 7.544026 | 7.676504 | 7.945155 | 7.807804 | 6.5704 |
| 7.226217 | 6.035737 | 5.285513 | NaN | NaN | 6.951633 | 6.743262 | 6.921484 | 5.640597 |
| NaN | NaN | NaN | NaN | 5.060313 | 5.998945 | 5.810126 | 6.081221 | 5.894952 |

| | | | | | | | | |
|---|---|---|---|---|---|---|---|---|
| NaN | 7.758718 | NaN | 5.190697 | 4.402524 | NaN | NaN | NaN | 5.788397 |
| 6.75039 | 7.182578 | 6.383653 | 6.200703 | 5.45606 | 6.652306 | 6.165624 | 6.424261 | 6.421273 |
| 8.090146 | 8.161983 | 8.19706 | 8.257722 | 8.443749 | 6.766003 | 6.933524 | 6.763585 | 6.033337 |
| 9.207072 | 9.163007 | 9.165427 | 9.562245 | 9.821138 | 8.693589 | 8.306373 | 8.733009 | 7.207781 |
| 7.557628 | 7.15878 | 6.104173 | 7.526463 | 7.595793 | 6.957213 | 5.545706 | 6.435648 | 6.268715 |
| 8.176812 | 8.209373 | 7.803008 | 8.145084 | 8.603641 | 8.783237 | 8.749184 | 8.519732 | 7.700359 |
| 8.824825 | 8.887275 | 8.888486 | 8.575354 | 8.645689 | 8.236058 | 8.756537 | 8.372569 | 8.303338 |
| 5.916786 | 6.325814 | 4.828454 | 6.960356 | 7.173878 | NaN | NaN | NaN | 5.710779 |
| 7.865516 | 8.129158 | 7.941896 | 10.28747 | 9.860679 | 8.995225 | 8.896534 | 8.719453 | 7.562648 |
| 7.18351 | 7.087448 | 6.960197 | 6.10895 | 6.403236 | 7.643502 | 7.637906 | 7.416222 | 7.825403 |
| 8.525113 | 9.337558 | 9.460462 | 10.23709 | 8.941913 | 7.245587 | 7.521646 | 7.309235 | 8.360501 |
| 7.968635 | 8.290747 | 8.353552 | 7.253275 | 7.29842 | 8.072842 | 8.574363 | 8.215641 | 8.401171 |
| 8.489451 | 8.556748 | 8.728856 | 9.733248 | 9.746956 | 8.485654 | 8.034654 | 8.154469 | 8.32701 |
| 6.797055 | 6.889866 | 6.740835 | 5.976233 | 5.956114 | 6.351274 | 6.427651 | 6.47308 | 6.733219 |
| 8.378022 | 7.58615 | 8.35588 | 7.069791 | 6.639105 | 7.580961 | 7.752447 | 7.498083 | 8.236734 |
| 8.167863 | 8.282694 | 8.242658 | 6.239767 | 6.439471 | 8.34274 | 8.056525 | 8.357927 | 8.049909 |
| 6.430046 | 6.599064 | 6.671646 | 6.334801 | 6.064794 | 6.380475 | 6.178405 | 6.242457 | 6.848411 |
| 6.514627 | 6.711792 | 6.681786 | 6.385827 | NaN | 7.237152 | 7.529751 | 7.313776 | NaN |
| 8.095663 | 7.840315 | 7.500708 | 7.231449 | 7.114023 | 7.267392 | 7.278189 | 7.195038 | 6.953704 |
| 6.50293 | 6.564173 | 6.67751 | 6.637697 | 6.589307 | 6.632357 | 6.648167 | 6.754932 | 6.241449 |
| 8.194559 | 8.387679 | 8.444715 | 8.899655 | 8.556324 | 8.396826 | 8.166753 | 8.664434 | 8.722614 |
| 5.639676 | NaN | NaN | NaN | NaN | 6.52362 | 6.167286 | 6.353446 | 4.165726 |
| 7.428325 | 7.223769 | 7.342437 | 7.886812 | 8.309362 | 6.532545 | 6.360359 | NaN | 7.446617 |
| 7.415361 | 6.783687 | 7.256697 | 6.531674 | 6.590866 | 7.265059 | 8.00006 | 8.391822 | 5.479043 |
| 9.359036 | 9.023009 | 8.979441 | 7.693737 | 7.998497 | 7.278198 | 7.877642 | 7.375295 | 7.501388 |
| 7.500791 | 7.33708 | 7.138275 | 6.958753 | 6.937055 | 6.578504 | 6.941654 | 7.139328 | 6.769038 |
| 7.514187 | 7.552838 | 7.64086 | 7.053433 | 6.870297 | 6.702448 | 6.909463 | 7.172855 | 7.279982 |
| 6.429671 | 5.121919 | 6.093571 | 4.965466 | 4.766242 | 6.624025 | 6.371253 | 6.52784 | 5.462323 |
| 8.369362 | 8.288739 | 8.477489 | 8.61317 | 8.518935 | 8.132631 | 8.06788 | 8.204615 | 7.91868 |
| 7.585456 | 7.46024 | 7.893193 | 7.114724 | 7.805424 | 6.669869 | 6.694436 | 7.525625 | 6.228355 |
| 6.690759 | 6.917511 | 6.97803 | 6.134334 | 6.05752 | 7.54091 | 7.538677 | 7.228131 | 7.162896 |
| 5.539073 | 5.689446 | 5.765358 | 4.862431 | 5.497005 | 6.816666 | 6.169095 | 6.475756 | 6.249851 |
| 7.661684 | 7.952388 | 8.018842 | 10.69525 | 10.69306 | 6.765557 | 7.24676 | 7.237405 | 8.041709 |
| 7.975045 | 6.095416 | 7.040643 | NaN | 6.270005 | 6.956898 | 7.863486 | 7.113913 | NaN |
| 7.340123 | 7.044676 | 6.867821 | 7.767804 | 6.892796 | 5.833083 | 6.437169 | 6.356521 | 6.355591 |
| 6.218563 | 6.358737 | 6.432339 | 5.61112 | 5.456632 | 7.130414 | 5.954203 | 6.213668 | 6.239296 |
| 9.062737 | 9.026901 | 9.139279 | 9.72223 | 10.05645 | 7.82741 | 7.606703 | 8.210819 | 8.114659 |
| 10.1881 | 9.946153 | 10.04255 | 10.01809 | 9.820345 | 9.762173 | 10.05644 | 9.880522 | 10.30668 |
| 7.53323 | 7.663529 | 7.360119 | NaN | NaN | NaN | NaN | NaN | NaN |
| 7.228888 | 7.348541 | 7.376297 | 6.364396 | 5.455699 | 8.050762 | 7.32024 | 7.052515 | 7.330057 |
| 7.825189 | 7.696938 | 7.795517 | 5.98124 | 6.017375 | 8.245936 | 8.212808 | 8.49928 | 7.441144 |
| 8.39229 | 8.586405 | 8.651914 | 7.69084 | 7.334939 | 7.986415 | 7.506666 | 7.962103 | 7.616716 |
| 7.836583 | 8.556907 | 8.775076 | 5.56987 | 5.712608 | 3.947989 | NaN | 3.593343 | 4.336798 |
| 9.59841 | 9.501745 | 9.561292 | 8.135606 | 8.338958 | 9.875135 | 9.708203 | 9.711733 | 9.067604 |
| 8.911791 | 8.834883 | 9.005738 | 9.435464 | 9.198181 | 8.472838 | 8.282396 | 8.473831 | 8.042809 |
| 8.931221 | 9.03826 | 9.054053 | 9.309264 | 9.574142 | 7.965457 | 8.114707 | 8.110853 | 7.539464 |
| 5.652279 | 5.826267 | 6.043016 | 5.841462 | 6.134587 | 5.520547 | 5.352395 | 5.143752 | 6.002007 |

| | | | | | | | | |
|---|---|---|---|---|---|---|---|---|
| 5.106048 | 4.819643 | 6.208659 | NaN | 5.16186 | 6.640856 | 7.29256 | 6.760241 | 6.35827 |
| 7.582524 | 7.581288 | 7.508216 | 7.57159 | 7.621829 | 7.56666 | 7.842429 | 8.273261 | 7.004013 |
| 8.734683 | 8.805557 | 8.757368 | 7.295915 | 7.320919 | 8.89254 | 9.02566 | 9.015346 | 9.261448 |
| NaN | NaN | NaN | NaN | NaN | NaN | 7.008439 | 7.304812 | 6.60036 |
| 6.553343 | 6.76917 | 6.199063 | 7.149427 | 7.258586 | 5.633266 | 5.770525 | 5.256144 | 7.272769 |
| 6.597427 | 6.764936 | 6.418293 | NaN | NaN | NaN | NaN | NaN | 6.366599 |
| 6.779721 | 6.813066 | 7.053753 | 6.687738 | 6.583916 | 6.829583 | 6.93194 | 7.45515 | 7.357013 |
| 8.532042 | 7.700059 | 8.172344 | 7.045293 | 7.260579 | 7.370189 | 7.197398 | 7.296128 | 7.162022 |
| 7.98676 | 7.161676 | 7.506713 | 5.431144 | NaN | 7.619987 | 7.451826 | 7.510324 | 6.569297 |
| 9.679156 | 9.876413 | 9.97376 | 8.239191 | 8.149304 | 9.607686 | 9.328903 | 8.927814 | 8.947676 |
| 9.036378 | 8.641499 | 9.031207 | 7.990043 | 6.812214 | 8.801903 | 8.995269 | 8.772382 | 7.996077 |
| 8.59893 | 8.661963 | 8.793171 | 8.626347 | 8.337926 | 8.477876 | 8.499693 | 8.429056 | 8.523457 |
| 9.095221 | 8.668773 | 8.584677 | 6.942812 | 7.072974 | 8.066079 | 8.210711 | 7.941473 | 8.030484 |
| 6.062376 | 5.831945 | 6.193624 | 4.955009 | 5.712721 | 6.130037 | 6.1482 | 6.322724 | 6.072799 |
| 6.647225 | 6.684908 | 6.631089 | 6.793744 | 6.68539 | 6.631457 | 6.669525 | 6.628242 | 6.750649 |
| 8.598424 | 8.501751 | 8.601349 | 8.174446 | 8.309829 | 7.8708 | 7.809523 | 7.890868 | 8.216141 |
| 8.422406 | 8.372561 | 8.264777 | 6.975449 | 7.09452 | 8.199907 | 8.128705 | 8.144791 | 7.818721 |
| 7.260943 | 7.058184 | 7.240205 | 6.732569 | 6.37398 | 7.587099 | 6.745149 | 7.030713 | 7.622249 |
| 7.80868 | 6.930255 | 7.279165 | 8.169501 | 8.334957 | 7.854576 | 8.032781 | 7.969303 | 7.231635 |
| 9.776084 | 9.808612 | 9.846424 | 9.407214 | 9.400808 | 9.674318 | 9.764452 | 9.632539 | 9.13178 |
| 7.309938 | 7.870129 | 7.189095 | 8.767824 | 8.476961 | 6.369961 | 6.587028 | 6.454163 | 6.44549 |
| 5.976927 | 6.098179 | 5.424905 | NaN | NaN | 5.083231 | 5.457636 | 5.259856 | 5.649006 |
| 6.894952 | 4.520825 | 7.21863 | 5.394062 | 4.804086 | 6.833268 | 6.67693 | 7.069217 | 6.617308 |
| 7.907804 | 8.063369 | 7.828586 | 6.809664 | 6.982132 | 7.226851 | 6.871379 | 7.37539 | 7.579067 |
| 5.48496 | 5.406255 | 5.913206 | 5.66344 | 5.676726 | 5.235361 | 5.147195 | 5.079545 | 3.56299 |
| 7.745078 | 6.720582 | 7.451867 | 6.781357 | 6.78374 | 6.58907 | 7.147434 | 6.65316 | 7.040228 |
| 12.03851 | 11.92452 | 11.94762 | 12.22815 | 12.19733 | 10.88636 | 10.89766 | 10.76261 | 11.81025 |
| 7.711454 | 8.050455 | 7.469072 | NaN | 7.539546 | 6.974887 | 7.077656 | 7.399643 | 6.652448 |
| 7.408595 | 6.907144 | 7.280024 | 7.562571 | 7.828336 | 7.343068 | 7.507145 | 7.140632 | 8.008748 |
| 8.323823 | 8.750467 | 8.216631 | 7.921502 | 8.822615 | 8.224349 | 10.11677 | 8.827883 | 7.696321 |
| NaN | 7.277194 | NaN | 6.841931 | 7.079156 | 7.540625 | 7.861784 | 7.506328 | 6.093865 |
| 5.732544 | 6.247048 | 6.275294 | 5.998177 | 5.97295 | 6.009809 | 6.577054 | 5.594101 | 5.094586 |
| 7.719026 | 6.908876 | 6.098347 | 6.797194 | 6.640862 | 6.137615 | 6.221562 | 7.005533 | 5.595956 |
| 6.885612 | 6.846436 | 6.131056 | NaN | 5.524167 | NaN | NaN | 6.98932 | 6.613715 |
| 3.322988 | 4.783251 | 4.502917 | 5.822133 | 5.360314 | 5.05279 | 4.347134 | 4.601964 | 3.547069 |
| 6.076218 | 6.536596 | 6.283587 | 8.157245 | 8.360994 | 7.663406 | 7.851826 | 8.487227 | 5.587368 |
| 7.878464 | NaN | 4.525149 | 6.770345 | 7.000314 | 7.785934 | 7.578513 | 7.54278 | 7.815385 |
| 9.802144 | 9.938199 | 10.00396 | 10.5511 | 10.57815 | 9.862387 | 9.727294 | 9.915155 | 9.178252 |
| 6.433617 | 5.82997 | 6.262991 | 5.282831 | 5.473564 | 6.112974 | 6.088646 | 6.156488 | 6.190372 |
| 5.743479 | 6.096555 | 6.059891 | 7.686623 | 7.730706 | 7.11324 | 6.674223 | 7.069755 | 5.865093 |
| 5.768903 | 5.943076 | 5.657786 | NaN | NaN | 5.889207 | NaN | 5.740045 | 5.442902 |
| 8.72892 | 8.791136 | 8.675568 | 8.459164 | 8.257154 | 7.298214 | 7.082258 | 7.110787 | 6.486414 |
| 7.633344 | 7.78534 | 7.512387 | 8.347136 | 8.631321 | 7.477465 | 7.547877 | 7.749053 | 7.319229 |
| 6.170533 | 6.447608 | 6.549006 | 6.026574 | 6.228098 | 5.853748 | 6.065035 | 6.357527 | 6.127104 |
| 8.447464 | 8.055701 | 8.486412 | 7.869198 | 7.615133 | 7.927351 | 7.780552 | 7.890035 | 8.087003 |

-Bruker Confidential-

| [11] 4_2_S | [12] 4_3_SP_CDKL5_S6-C12_1_12206.d.PG.Quantity |
|---|---|
| 9.38518 | 9.247501 |
| NaN | NaN |
| NaN | NaN |
| 4.320309 | 4.091077 |
| 6.876291 | 6.512304 |
| 7.357651 | NaN |
| 7.445229 | 7.552409 |
| 5.834313 | 5.254733 |
| 6.594919 | 6.907578 |
| 8.042384 | 8.123834 |
| 7.043981 | 7.04834 |
| 4.417755 | 4.980915 |
| 8.622231 | 8.478333 |
| NaN | 3.783441 |
| 6.928082 | 7.614986 |
| 7.157159 | 7.111979 |
| 5.020511 | 5.189949 |
| 6.469814 | NaN |
| 7.156692 | 7.079915 |
| 5.511058 | 5.633136 |
| 6.983566 | 6.005487 |
| 4.768034 | 5.440777 |
| 7.222181 | 7.356377 |
| 6.891258 | 6.835426 |
| 5.98448 | 5.873837 |
| 6.775175 | 6.732234 |
| 7.336741 | 7.134357 |
| 5.355481 | 5.1982 |
| 6.430319 | 6.207212 |
| 5.120088 | NaN |
| 8.817739 | 8.756616 |
| 5.546694 | 5.423263 |
| NaN | 7.113843 |
| 8.323703 | 8.371634 |
| 8.986219 | 8.589777 |
| 11.02198 | 10.84904 |
| 7.149379 | 6.771018 |
| NaN | 7.199502 |
| 7.150961 | 7.293262 |
| 8.098818 | 7.977431 |
| 6.15032 | 6.531079 |
| NaN | 4.591942 |
| NaN | 4.361043 |
| 6.88635 | 7.240417 |
| 12.21738 | 12.07696 |
| 7.857573 | 8.102899 |

| | |
|---|---|
| 7.933952 | 7.735065 |
| 6.770821 | 6.573628 |
| 6.935966 | 8.29602 |
| 8.24079 | 8.383064 |
| 5.907951 | 5.397764 |
| 6.561642 | 6.778534 |
| 6.281372 | 6.000458 |
| NaN | NaN |
| 7.098437 | 7.315251 |
| 8.524393 | 8.348822 |
| 6.102235 | 6.820247 |
| 5.586654 | 5.908372 |
| 6.567816 | 6.083786 |
| NaN | NaN |
| 6.192465 | 6.561138 |
| 7.620372 | 7.717474 |
| 7.669851 | 7.165139 |
| 10.00574 | 9.63203 |
| NaN | NaN |
| 5.856081 | 6.084018 |
| 7.309594 | 6.352077 |
| NaN | NaN |
| 6.488903 | 6.361782 |
| 6.150304 | 6.567137 |
| 6.31482 | 6.533065 |
| 5.147794 | 4.528179 |
| 10.3247 | 9.476647 |
| 6.503942 | 4.667392 |
| 10.86195 | 10.8795 |
| 6.4183 | 6.146116 |
| 12.15229 | 12.06324 |
| 5.603945 | 5.783256 |
| 6.416131 | 5.461838 |
| 7.2898 | 7.268856 |
| 9.079413 | 8.848425 |
| 8.328904 | 8.228125 |
| 9.365475 | 9.478935 |
| NaN | NaN |
| 7.512682 | 7.705879 |
| 7.521587 | 7.332856 |
| 10.71491 | 10.68252 |
| 6.806306 | 7.031466 |
| 5.380089 | 5.438745 |
| 5.784119 | 5.990499 |
| 8.672527 | 8.367212 |
| 7.70613 | 8.060017 |
| 6.947604 | 6.83255 |

-Bruker Confidential-

| | |
|---|---|
| 7.948541 | 7.8837 |
| 4.86002 | 5.825249 |
| 5.046706 | 5.415348 |
| 7.086155 | 6.952029 |
| 7.448139 | 7.368306 |
| 9.500043 | 9.599315 |
| 11.0391 | 11.02376 |
| 8.443486 | 8.3336 |
| 8.243559 | 8.201912 |
| 7.657189 | 7.505503 |
| 7.268703 | 7.338744 |
| 5.559541 | 5.504433 |
| 5.107023 | 4.820444 |
| 10.58232 | 10.41644 |
| 7.601902 | 7.46532 |
| 7.244396 | 7.30863 |
| 4.723431 | 4.334153 |
| 8.304503 | 8.196424 |
| 10.355 | 10.27518 |
| 9.027295 | 8.988505 |
| 6.979337 | 6.637704 |
| 5.580063 | 6.148098 |
| 7.367903 | 7.563131 |
| 6.666461 | 6.640913 |
| 7.58817 | 7.451556 |
| 5.823086 | 5.849075 |
| 5.539568 | 5.612463 |
| 9.8903 | 9.829286 |
| 4.70517 | 4.770216 |
| 7.932884 | 7.758839 |
| 6.536226 | 6.821852 |
| 9.582896 | 9.624914 |
| 7.948182 | 7.956258 |
| 5.824331 | 6.064712 |
| 6.226013 | 6.395427 |
| 6.897698 | 6.731112 |
| NaN | NaN |
| 6.430965 | 6.775787 |
| 5.260202 | 4.983314 |
| 8.401829 | 8.675783 |
| 7.45645 | 7.264992 |
| 8.765762 | 8.727546 |
| 7.418937 | 5.812089 |
| 8.399858 | 8.26022 |
| 7.223708 | 7.025391 |
| 8.222942 | 9.273795 |
| 6.333371 | 6.811185 |

| | |
|---|---|
| 7.107363 | 7.242222 |
| NaN | 6.171438 |
| 7.122729 | 7.107951 |
| 8.832813 | 8.448828 |
| 7.620697 | 7.229604 |
| 6.529238 | 6.216477 |
| 5.5685 | 4.817676 |
| NaN | NaN |
| 7.436616 | 7.700951 |
| 7.882872 | 7.870208 |
| 8.717931 | 9.183562 |
| 6.345781 | 6.365911 |
| 7.571165 | 7.841413 |
| 9.032187 | 8.891129 |
| 10.08763 | 10.08507 |
| 5.61923 | 5.740213 |
| 7.614637 | 7.604403 |
| 7.002687 | 7.079141 |
| 7.140276 | 7.572656 |
| 7.304477 | 7.348605 |
| 7.521112 | 7.336794 |
| NaN | NaN |
| 7.424237 | NaN |
| 4.892071 | 4.631238 |
| 6.638177 | 5.685942 |
| 7.568162 | 7.623082 |
| 3.943981 | 6.216049 |
| 7.223176 | 6.54836 |
| 7.989067 | 8.120744 |
| 5.00358 | 5.157576 |
| 6.348558 | 6.968267 |
| 8.259564 | 8.112448 |
| 5.65266 | 4.943898 |
| 7.68187 | 8.136767 |
| 8.382147 | 8.253709 |
| NaN | NaN |
| 6.570199 | 6.496218 |
| NaN | NaN |
| 6.222774 | NaN |
| 6.529131 | 7.049978 |
| 9.13573 | 8.82219 |
| 5.864231 | 4.493363 |
| NaN | 6.005323 |
| 6.399909 | 6.680296 |
| 8.157688 | 8.462872 |
| 8.311428 | 8.331351 |
| 5.789374 | 5.495103 |

| | |
|---|---|
| 5.913787 | 5.628582 |
| 8.193995 | 8.321589 |
| 6.739549 | 7.351501 |
| NaN | NaN |
| 8.096067 | 8.022276 |
| 7.405931 | 7.387836 |
| 8.386818 | 8.196653 |
| NaN | 4.809462 |
| 7.82278 | 7.714081 |
| 7.80355 | 7.929779 |
| 7.354932 | 7.311633 |
| 7.600442 | 7.428854 |
| 7.243206 | 7.021834 |
| 7.368376 | 6.312493 |
| 6.493003 | 6.175649 |
| 9.305141 | 8.698071 |
| 7.224523 | 7.223922 |
| 6.588488 | 6.499506 |
| 5.741704 | 5.31275 |
| 6.433903 | 6.522733 |
| 8.145601 | 7.969267 |
| 7.834415 | 6.948462 |
| 8.554125 | 8.475075 |
| 9.589342 | 9.542622 |
| 7.437761 | 7.336707 |
| 6.700843 | 7.167346 |
| 8.110187 | 6.979932 |
| 8.792308 | 8.75092 |
| 8.183947 | 8.295326 |
| 9.03738 | 9.034787 |
| 6.713657 | 6.909781 |
| NaN | NaN |
| 7.950365 | 8.042557 |
| 8.740159 | 8.427383 |
| 6.094162 | 6.460223 |
| 7.149693 | 6.789066 |
| 7.575643 | 7.263484 |
| NaN | NaN |
| 7.622109 | 7.959678 |
| 7.084633 | 7.277837 |
| 6.958158 | 6.927972 |
| 10.39931 | 9.838049 |
| 8.319353 | 8.524714 |
| 6.937295 | 6.695933 |
| 8.737451 | 8.526803 |
| 5.226925 | 4.580132 |
| 7.262465 | 7.049449 |

| | |
|---|---|
| 4.578594 | 6.020633 |
| 7.676395 | 7.54811 |
| 7.119731 | 6.993942 |
| 8.227688 | 8.371677 |
| 6.471404 | 6.302292 |
| 10.31709 | 10.55472 |
| 5.800292 | 6.032331 |
| 8.611217 | 8.545356 |
| 5.975413 | 5.805569 |
| 6.406573 | 6.386708 |
| 7.115026 | 7.207713 |
| 7.283288 | 7.393271 |
| 10.0508 | 9.971888 |
| 8.593548 | 8.753243 |
| 6.679771 | 6.854128 |
| 6.18929 | 5.998685 |
| 5.776182 | 6.30333 |
| 7.577778 | 6.678012 |
| 8.171816 | 8.179612 |
| 7.288341 | 7.460435 |
| 7.421274 | 7.276642 |
| 4.128454 | 4.927441 |
| 6.712322 | 6.732569 |
| 5.000749 | 4.659385 |
| 7.81423 | 7.719672 |
| 4.915974 | 5.049493 |
| 7.561756 | 7.296536 |
| 7.500192 | 7.688087 |
| 8.264898 | 8.207064 |
| 5.146117 | 4.80264 |
| 7.578492 | 7.541945 |
| 5.845924 | 5.467435 |
| 11.22693 | 11.13917 |
| 6.12371 | 6.632934 |
| 5.632665 | 5.571232 |
| 6.549798 | 6.49767 |
| 6.574591 | 6.24311 |
| 6.169165 | 6.028431 |
| 8.454686 | 8.553776 |
| 7.910736 | 7.483885 |
| 6.253386 | 5.843053 |
| 6.67468 | 6.508453 |
| 7.698798 | 7.496328 |
| 7.323386 | 7.190338 |
| 9.39163 | 9.298049 |
| 8.731162 | 8.850863 |
| 6.706462 | 5.688247 |

| | |
|---|---|
| 6.827852 | 5.605412 |
| 7.251456 | 7.36578 |
| 8.081346 | 7.983073 |
| 8.587547 | 8.63242 |
| 7.057438 | 6.929429 |
| 5.640059 | 5.890423 |
| 10.0025 | 9.345568 |
| 6.731166 | 6.030968 |
| 7.100901 | 6.832089 |
| 5.118243 | 4.449746 |
| 7.088731 | 7.085372 |
| 5.655062 | 5.836008 |
| 4.84114 | 5.158588 |
| 4.435575 | 4.818969 |
| 5.102549 | 5.329669 |
| 9.151896 | 9.159744 |
| 6.883983 | 6.923563 |
| 7.022346 | 7.458082 |
| 8.399916 | 8.270082 |
| 7.433156 | 7.730548 |
| 5.90145 | 5.764396 |
| 7.335626 | 7.427026 |
| 5.373006 | 5.299304 |
| 7.121555 | 6.773172 |
| 7.201067 | 6.968249 |
| 11.10681 | 10.86516 |
| 9.249427 | 9.067065 |
| 6.079853 | 6.114302 |
| 8.230433 | 7.976317 |
| 8.546799 | 8.469445 |
| 5.31848 | 5.044277 |
| 6.102301 | 5.72167 |
| 6.528424 | 6.371176 |
| 7.01687 | 7.199036 |
| 6.717172 | 6.27604 |
| 8.942937 | 9.051292 |
| 5.082326 | 4.442795 |
| 7.691347 | 8.003112 |
| 5.665853 | 5.498391 |
| 4.295584 | 5.532571 |
| 6.797601 | 6.746752 |
| 6.426635 | 6.377268 |
| 7.394928 | 7.809769 |
| 6.174276 | 5.423916 |
| 9.537389 | 9.487094 |
| 6.242783 | 6.257626 |
| 7.742341 | 7.587743 |

| | |
|---|---|
| 7.60471 | 7.392665 |
| 3.972236 | 3.542738 |
| 9.186895 | 9.077273 |
| 8.267741 | 8.050837 |
| 7.864115 | 7.649921 |
| 4.872647 | 4.760282 |
| 4.790081 | 5.876214 |
| 5.968459 | 5.494039 |
| 6.192571 | 6.144478 |
| 6.74849 | 6.431949 |
| 9.483736 | 10.81527 |
| 7.92663 | 7.841766 |
| 10.63956 | 10.40643 |
| 7.76823 | 7.533809 |
| 5.395214 | 5.708293 |
| 4.569874 | 5.040829 |
| 9.561602 | 9.252643 |
| 10.46961 | 10.23645 |
| 9.539937 | 9.776804 |
| 7.421938 | 7.001907 |
| 6.793159 | 6.941801 |
| 5.682694 | 5.61386 |
| 5.487749 | 4.472487 |
| 8.208103 | 8.367922 |
| 7.1292 | 6.87755 |
| 7.735901 | 7.796999 |
| 5.188208 | 4.814993 |
| 7.782825 | 7.671189 |
| 5.568765 | 5.844284 |
| 4.637094 | 4.462768 |
| 6.808944 | 7.284808 |
| 6.50039 | 6.774871 |
| NaN | NaN |
| 6.616 | 6.779206 |
| 5.82135 | 5.369555 |
| 9.493529 | 9.583631 |
| 6.209152 | 6.052061 |
| 9.890102 | 9.950242 |
| 12.29671 | 12.3525 |
| 7.988137 | 7.687146 |
| 8.937231 | 8.955085 |
| 8.015061 | 8.130093 |
| 7.910412 | 7.701437 |
| 5.066256 | 3.941172 |
| 5.456698 | 5.231412 |
| 7.448805 | 7.349486 |
| 7.31147 | 7.127209 |

| | |
|---|---|
| 8.052182 | 8.078618 |
| 8.489072 | 8.438607 |
| 8.801711 | 9.218807 |
| 7.201588 | 7.389506 |
| 7.363784 | 7.26166 |
| 6.377499 | 5.375983 |
| 7.434536 | 7.085604 |
| NaN | 6.882474 |
| NaN | NaN |
| 4.016515 | 4.152131 |
| 6.039153 | 6.058626 |
| NaN | NaN |
| 6.685357 | 6.713475 |
| NaN | 5.927777 |
| 5.520944 | 5.44322 |
| 6.804268 | 5.954969 |
| 7.944067 | 7.717925 |
| 6.124963 | 6.09446 |
| 11.86218 | 11.90348 |
| 8.827947 | 8.992318 |
| 7.090673 | 7.590843 |
| 8.846488 | 8.936507 |
| 8.061683 | 7.656717 |
| 8.796115 | 8.570011 |
| 7.402486 | 7.550579 |
| 7.528093 | 7.371244 |
| 6.040816 | 6.397614 |
| 7.181248 | 5.760672 |
| NaN | NaN |
| 6.225246 | 6.019908 |
| 9.341483 | 9.000188 |
| 9.652303 | 9.791506 |
| 6.306719 | 5.814145 |
| 9.583509 | 9.557487 |
| 9.537281 | 9.443093 |
| 5.132823 | 5.265293 |
| 6.15956 | 5.676507 |
| 5.635872 | 5.118637 |
| 8.533736 | 8.552009 |
| 6.231351 | 6.648852 |
| 7.836629 | 7.81216 |
| 6.856653 | 6.448092 |
| 7.361374 | 6.998028 |
| 7.961367 | 6.757428 |
| 10.99156 | 11.03515 |
| 4.85444 | 5.067616 |
| 6.514314 | 6.623971 |

| | |
|---|---|
| 9.097231 | 9.06172 |
| NaN | 5.64676 |
| 8.140615 | 8.161865 |
| 6.134378 | 6.009096 |
| 7.147157 | 6.467658 |
| 8.446926 | 8.582782 |
| 9.057189 | 9.296693 |
| 6.880701 | 6.740522 |
| 6.358409 | 6.949932 |
| 3.611692 | 3.017457 |
| 5.864273 | 5.71769 |
| 5.649708 | 6.090759 |
| 7.349919 | 7.12158 |
| 7.791162 | 7.410834 |
| 11.97748 | 11.8596 |
| 7.1188 | 7.238933 |
| 9.314607 | 9.54789 |
| 7.545139 | 7.385411 |
| 7.205224 | 7.124402 |
| 8.103994 | 8.09729 |
| 7.338275 | 7.236901 |
| 8.130849 | 7.909321 |
| 7.65473 | 7.715033 |
| 7.922798 | 8.034606 |
| NaN | NaN |
| 7.750291 | 7.813025 |
| 6.865291 | 6.754187 |
| 7.613461 | 7.225703 |
| 5.924313 | 6.410819 |
| 6.685243 | 6.439596 |
| 8.387073 | 8.22897 |
| 6.177787 | 6.066124 |
| 5.746984 | 6.056539 |
| 6.947921 | 6.747821 |
| 8.639773 | 8.709252 |
| 7.600806 | 7.242951 |
| 10.46737 | 10.39282 |
| 6.45515 | 6.190022 |
| 9.028112 | 9.070726 |
| 11.33463 | 11.34719 |
| 6.345885 | 6.057223 |
| 6.930267 | 6.642542 |
| 5.970307 | 5.929935 |
| 7.324198 | 7.470313 |
| 9.70789 | 9.581655 |
| 6.158284 | 6.102057 |
| 6.168425 | 6.387668 |

| | |
|---|---|
| 6.499397 | 6.303561 |
| 5.911288 | 6.096141 |
| 8.422529 | 8.229666 |
| 5.320131 | 6.260677 |
| 4.221935 | 5.476183 |
| 8.812812 | 8.559362 |
| 5.513339 | 5.435503 |
| 7.297013 | 7.138455 |
| 6.988099 | 7.189429 |
| 6.836736 | 6.951843 |
| 7.311182 | 7.311382 |
| 5.386833 | 5.54951 |
| 7.099212 | 6.868193 |
| 8.413465 | 8.396643 |
| 8.497949 | 8.233106 |
| 10.17608 | 10.32951 |
| 5.796373 | 4.721133 |
| 7.40565 | 6.964435 |
| 6.740107 | 6.751672 |
| 7.393558 | 7.389877 |
| 7.993464 | 7.976978 |
| 6.789976 | 6.76741 |
| 7.514654 | 7.797311 |
| 7.337644 | 6.975384 |
| 7.303617 | 7.268916 |
| 7.348527 | 6.951427 |
| 7.63292 | 7.694055 |
| 7.465318 | 7.220895 |
| 6.734545 | 6.605266 |
| 6.286968 | 6.662685 |
| 7.05817 | 7.126008 |
| 7.97866 | 7.873852 |
| 8.887156 | 8.860777 |
| 9.220793 | 9.113602 |
| 8.314839 | 8.243579 |
| 8.970774 | 9.05619 |
| 8.327477 | 8.369352 |
| 9.350462 | 9.173918 |
| 6.383178 | 5.950861 |
| 8.418022 | 8.297496 |
| 6.348482 | 6.52546 |
| 7.497328 | 7.652062 |
| 5.659416 | 5.80087 |
| 5.798894 | 5.086541 |
| 7.120223 | 7.954668 |
| 8.389548 | 8.417279 |
| 8.145848 | 7.953725 |

| | |
|---|---|
| 5.937497 | 5.52705 |
| 5.998391 | 6.486146 |
| 6.69074 | 6.682552 |
| 6.998379 | 6.658195 |
| 5.683188 | 5.526629 |
| 9.205207 | 9.16018 |
| 7.748497 | 7.766815 |
| 8.854295 | 8.385523 |
| 3.747434 | 2.75323 |
| NaN | 6.126257 |
| 5.445092 | 5.837249 |
| 8.90569 | 8.927878 |
| 5.693202 | 5.595363 |
| 6.896613 | 6.922808 |
| 5.836189 | 5.204818 |
| NaN | NaN |
| 5.997565 | 6.089414 |
| 7.179611 | 7.149058 |
| 8.357199 | 8.30987 |
| 7.109802 | 7.241833 |
| 5.475242 | 5.411667 |
| 8.066662 | 8.209093 |
| 6.600929 | 6.47139 |
| 7.584211 | 7.445628 |
| 5.938634 | 5.797936 |
| 5.933461 | 5.193519 |
| 9.53734 | 9.408158 |
| 6.48294 | 6.346629 |
| 6.374561 | 6.400014 |
| 6.66265 | 6.579941 |
| 8.767649 | 8.553461 |
| 6.741808 | 6.960392 |
| 7.876231 | 7.866307 |
| 8.24447 | 8.09429 |
| 5.688 | 5.995638 |
| 8.812683 | 8.753745 |
| 7.608118 | 7.511806 |
| 7.153725 | 7.165009 |
| 7.407259 | 7.300983 |
| 7.362009 | 7.240214 |
| 6.521548 | 6.329869 |
| 7.322367 | 7.06071 |
| 7.22125 | 7.135335 |
| 6.429471 | 6.353797 |
| 8.44944 | 8.293044 |
| 6.4723 | 6.43317 |
| 6.153102 | 5.665141 |

| | |
|---|---|
| 6.782422 | 6.532884 |
| 6.298275 | 5.920045 |
| 5.969273 | 6.204616 |
| 6.707219 | 6.843534 |
| 6.872686 | 7.216284 |
| 8.405239 | 8.221684 |
| 6.629992 | 6.711183 |
| 6.74443 | 6.952263 |
| 7.756888 | 7.991518 |
| 8.357555 | 8.337412 |
| 7.279189 | 7.801747 |
| 8.964911 | 8.726101 |
| 7.766508 | 7.56989 |
| 7.900432 | 7.785749 |
| 9.036728 | 8.559111 |
| 6.94441 | 6.82205 |
| 9.791279 | 9.794065 |
| 6.665361 | 7.146718 |
| 9.383302 | 9.327774 |
| 5.45676 | 5.445148 |
| 5.359433 | 5.700763 |
| 10.89393 | 10.57163 |
| 8.938223 | 8.71788 |
| 9.917631 | 10.04797 |
| 6.071575 | 5.789289 |
| 6.289269 | 6.14984 |
| 7.740033 | 7.858065 |
| 7.391394 | 7.088229 |
| 10.7159 | 10.73667 |
| 6.414405 | 6.316311 |
| 9.44447 | 9.360504 |
| 7.640718 | 7.65913 |
| 5.90057 | 5.508348 |
| 6.846827 | 6.747439 |
| 8.494394 | 8.772124 |
| 8.19975 | 8.068701 |
| 6.167525 | 6.520789 |
| 8.826985 | 8.802257 |
| 8.092236 | NaN |
| 5.952818 | 6.137488 |
| NaN | NaN |
| 9.688419 | 9.715627 |
| 8.638284 | 8.56146 |
| 4.90835 | 5.045894 |
| 9.359551 | 9.422759 |
| 9.288601 | 9.145761 |
| 7.551601 | 7.372104 |

-Bruker Confidential-

| | |
|---|---|
| 7.704134 | 8.180031 |
| NaN | 5.224196 |
| 7.525133 | 7.889226 |
| 8.509535 | 8.379174 |
| 8.678349 | 8.546003 |
| 9.274495 | 8.830705 |
| 6.643095 | 6.808583 |
| 10.87552 | 10.91312 |
| 8.513618 | 8.427642 |
| 9.460401 | 9.400872 |
| 4.552001 | 4.789321 |
| 6.816922 | 6.831251 |
| 8.191692 | 8.298067 |
| 6.728736 | 7.158445 |
| 6.386599 | 5.927968 |
| 6.770522 | 6.637128 |
| 7.877984 | 7.792299 |
| 9.310709 | 9.356577 |
| 6.628364 | 6.726867 |
| 8.013411 | 7.250299 |
| 9.454203 | 9.482116 |
| 8.858425 | 8.893297 |
| 7.245895 | 7.070443 |
| 5.90908 | 5.727839 |
| 7.159878 | 7.283266 |
| 6.730966 | 6.340841 |
| 4.519139 | 4.30966 |
| 7.514536 | 7.38838 |
| 8.894237 | 8.389283 |
| 5.56157 | 5.436286 |
| 6.930949 | 6.990602 |
| 7.098428 | 7.17915 |
| 9.390429 | 9.337375 |
| 8.810229 | 8.756495 |
| 6.423579 | 6.52893 |
| 6.548514 | 6.581023 |
| 5.919156 | 5.934809 |
| 6.886134 | 7.142652 |
| 8.757191 | 8.68884 |
| 7.885118 | 7.930942 |
| 7.202373 | 7.03795 |
| 7.206068 | 7.189074 |
| 6.277878 | 6.591505 |
| 10.53606 | 10.18617 |
| 6.557557 | 6.931297 |
| 6.336707 | 6.019686 |
| 5.460809 | NaN |

| | |
|---|---|
| 6.799004 | 6.496832 |
| 5.654759 | NaN |
| 5.964857 | 5.916836 |
| 4.546875 | NaN |
| 6.301732 | 5.689355 |
| 6.594809 | 6.547541 |
| 6.340003 | 6.35795 |
| 6.824141 | 6.4182 |
| 9.50182 | 9.595513 |
| 5.768126 | 5.793387 |
| 8.51578 | 8.458234 |
| 8.48615 | 8.408729 |
| 9.582356 | 9.543762 |
| 6.391813 | 6.7885 |
| 10.2968 | 10.42472 |
| 7.363434 | 7.293324 |
| 9.340189 | 9.34157 |
| 7.493444 | 7.370075 |
| 9.039252 | 9.276693 |
| 6.799445 | 6.92957 |
| 7.308605 | 7.351254 |
| 9.988724 | 10.12361 |
| 9.032189 | 9.213135 |
| 6.776032 | 6.341957 |
| 6.851844 | 6.777748 |
| 5.697319 | 5.43047 |
| 7.317241 | 7.515595 |
| 6.31998 | 6.15485 |
| 7.819239 | 8.028115 |
| 6.859538 | 6.666672 |
| 7.131328 | 7.034136 |
| 8.650177 | 8.522428 |
| 9.127706 | 9.279693 |
| 9.723592 | 9.762968 |
| 7.155265 | 7.245562 |
| 8.403751 | 8.408531 |
| 6.936873 | 6.633736 |
| 8.084275 | 8.07051 |
| 7.721384 | 7.517667 |
| 7.734942 | 8.684491 |
| 7.634983 | 8.072174 |
| 6.164917 | 6.528148 |
| 6.803692 | 6.928689 |
| 7.612786 | 7.31303 |
| 5.809874 | 5.666286 |
| 8.339611 | 8.009429 |
| 6.635703 | 6.687775 |

| | |
|---|---|
| 7.727373 | 7.582654 |
| 6.616786 | 6.150192 |
| 6.951088 | 6.801636 |
| 6.921991 | 6.96044 |
| 6.54631 | 6.462276 |
| 7.106787 | 7.714726 |
| 3.754238 | 4.77384 |
| 8.006266 | 8.082482 |
| 8.22798 | 8.37109 |
| 7.00491 | 7.01008 |
| 7.029861 | 6.451348 |
| 7.504404 | 7.557151 |
| NaN | NaN |
| 7.903445 | 8.23997 |
| 8.301291 | 8.594205 |
| 10.11218 | 9.999446 |
| 8.326811 | 8.188503 |
| 3.638029 | 5.458731 |
| 5.69725 | 5.903154 |
| 7.235416 | 7.153409 |
| 6.322847 | 6.467144 |
| 6.524549 | 6.835793 |
| 6.103035 | 3.690614 |
| 5.627633 | 5.98079 |
| 6.8998 | 6.721714 |
| 6.775886 | 6.815997 |
| 5.711267 | 5.739119 |
| 5.890077 | 5.81084 |
| 5.397594 | 6.545955 |
| 10.43523 | 10.63825 |
| 5.969305 | 6.112892 |
| 8.256117 | 8.378959 |
| 6.752838 | 6.825253 |
| 8.99768 | 8.940736 |
| 6.215156 | 5.91636 |
| 5.884035 | 6.376254 |
| 6.327262 | 6.580327 |
| 6.725879 | 6.958996 |
| 6.677491 | 6.631085 |
| 7.106747 | 6.890068 |
| 7.362735 | 7.064765 |
| 6.585149 | 6.779636 |
| 8.356236 | 8.272858 |
| 7.113431 | 7.302274 |
| 5.935039 | 5.629209 |
| 7.32335 | 7.168486 |
| NaN | NaN |

| | |
|---|---|
| 4.929454 | NaN |
| 6.822346 | 6.599224 |
| 6.882585 | 7.309824 |
| 6.990246 | 7.631017 |
| NaN | NaN |
| 8.688195 | 8.452401 |
| 7.887979 | 7.85036 |
| 6.67204 | 6.470432 |
| 8.795597 | 8.792167 |
| 8.23783 | 8.414284 |
| 8.315331 | 8.062398 |
| 5.411752 | 5.338846 |
| 6.521574 | 6.199742 |
| 7.041224 | 7.176188 |
| 7.669429 | 6.061267 |
| 6.995656 | 6.992572 |
| 6.934429 | 6.962486 |
| 6.195865 | 5.921921 |
| 6.379037 | 6.680181 |
| 8.290681 | 8.204752 |
| 7.962294 | 8.614189 |
| 6.831416 | 8.578664 |
| 7.433483 | 7.382102 |
| 8.228812 | 8.518492 |
| 6.887 | 6.710315 |
| 7.028494 | 7.397712 |
| 9.788075 | 9.888596 |
| 6.798879 | 6.892689 |
| 6.964747 | 6.993555 |
| 7.094996 | 7.053753 |
| 6.7219 | 6.809795 |
| 7.665372 | 7.452882 |
| 8.873345 | 8.894092 |
| 6.930473 | 6.774473 |
| 6.706134 | 5.759816 |
| 6.833137 | 7.004204 |
| 8.638485 | 8.66892 |
| 6.341753 | 6.240775 |
| 7.582218 | 7.524884 |
| 5.497564 | 5.952297 |
| 5.105697 | 4.96374 |
| 8.714179 | 8.814608 |
| 6.367013 | 5.932364 |
| 7.522538 | 7.857237 |
| 6.251462 | 6.266264 |
| 8.299787 | 8.384962 |
| 7.543941 | 7.27355 |

| | |
|---|---|
| 7.27585 | 7.130079 |
| 7.290704 | 7.06068 |
| 6.195299 | 6.703608 |
| 7.799723 | 7.869989 |
| 5.772548 | 6.174831 |
| 5.418557 | 5.118981 |
| 7.800412 | 7.510824 |
| 6.222334 | 5.540477 |
| 8.488852 | 8.334871 |
| 6.978787 | 6.866067 |
| 5.970632 | 5.70619 |
| 5.686852 | 6.181803 |
| 5.11132 | 5.16715 |
| 7.517841 | 7.23631 |
| 6.994083 | 4.842399 |
| 7.333305 | 7.686012 |
| 8.202687 | 8.204173 |
| 7.886882 | 7.744873 |
| 7.486927 | 7.187671 |
| 7.959826 | 8.053487 |
| 7.05441 | 6.30679 |
| 9.406213 | 8.948121 |
| 6.458344 | 6.015063 |
| 8.391458 | 8.610252 |
| 9.709921 | 9.570194 |
| NaN | 4.170716 |
| 9.914778 | 9.769728 |
| 5.665573 | 5.645511 |
| 9.583337 | 9.630317 |
| 7.974149 | 7.831535 |
| 8.693137 | 9.589183 |
| 8.74754 | 8.702827 |
| 10.1999 | 10.12805 |
| 7.06642 | 6.461018 |
| 8.113986 | 7.816591 |
| 5.67994 | 5.48615 |
| 6.972267 | 7.14353 |
| 5.757652 | 5.480879 |
| 8.260937 | 8.357186 |
| 6.064177 | 6.102596 |
| 6.993723 | 7.250233 |
| 8.163485 | 8.057794 |
| 4.046101 | 6.122299 |
| 6.311679 | NaN |
| 6.790833 | 7.121676 |
| 7.371607 | 6.912666 |
| 10.91793 | 11.10028 |

| | |
|---|---|
| 6.929323 | 6.432055 |
| 8.607968 | 8.44803 |
| 6.722089 | 6.696209 |
| 6.559899 | 8.347682 |
| 4.843205 | 5.121391 |
| 7.851207 | 7.83451 |
| 5.899799 | 6.247778 |
| 7.622458 | 7.461215 |
| 2.710994 | 4.616781 |
| 6.280241 | 5.873399 |
| 8.955468 | 9.136781 |
| 6.700747 | 6.375619 |
| 6.079725 | 5.869171 |
| 6.998241 | 7.192998 |
| 8.487797 | 8.607038 |
| 7.831033 | 8.180356 |
| 7.205008 | 7.272717 |
| 9.464697 | 9.505989 |
| 5.505076 | 5.46014 |
| 7.064028 | 6.99052 |
| 9.925925 | 9.705508 |
| 9.268754 | 9.552027 |
| 6.954603 | 6.07751 |
| 9.69367 | 9.819272 |
| 8.550249 | 8.487767 |
| 9.644463 | 9.249975 |
| 8.746454 | 8.652992 |
| 6.995061 | 7.044552 |
| 6.976699 | 7.051695 |
| 6.560513 | 5.536652 |
| 8.855034 | 9.348585 |
| 5.582885 | 5.980363 |
| NaN | 5.837261 |
| 6.833809 | 7.420287 |
| 6.238319 | 7.369255 |
| 8.82018 | 8.55418 |
| 5.685437 | 4.624165 |
| 7.987999 | 7.813525 |
| 12.12696 | 11.94572 |
| 10.44952 | 10.65237 |
| NaN | 6.84011 |
| 7.935436 | 7.783095 |
| 6.34523 | 6.984593 |
| 6.713258 | 6.196256 |
| NaN | 8.479801 |
| 6.507545 | 6.399885 |
| 10.3734 | 10.21191 |

-Bruker Confidential-

| | |
|---|---|
| 8.424223 | 8.35365 |
| 7.244438 | 7.138149 |
| 4.248877 | 4.173021 |
| 6.73565 | 6.95965 |
| 5.640681 | 5.387447 |
| 8.09882 | 7.341078 |
| 7.442333 | 7.655141 |
| 8.072787 | 8.39095 |
| 7.679567 | 7.761017 |
| 6.940827 | 6.737036 |
| 5.763416 | 5.607903 |
| NaN | NaN |
| 6.500677 | 6.901496 |
| 7.631503 | 7.78403 |
| 5.766219 | 5.678589 |
| 8.517179 | 8.057075 |
| 9.979448 | 9.687653 |
| 8.547525 | 8.378642 |
| 5.981192 | 5.76055 |
| 7.444067 | 7.61286 |
| 9.421744 | 9.472208 |
| 7.548372 | 7.631663 |
| 7.284582 | 6.464906 |
| 7.257538 | 7.109959 |
| 9.472433 | 8.579818 |
| 8.195295 | 8.154418 |
| 5.978478 | 6.288521 |
| 9.954515 | 10.01479 |
| 9.95708 | 9.9996 |
| 10.71553 | 10.67372 |
| 10.36153 | 10.19973 |
| 7.6191 | 7.558516 |
| 9.710027 | 9.774258 |
| NaN | NaN |
| 8.676187 | 12.1706 |
| 7.638551 | 7.520145 |
| NaN | NaN |
| NaN | NaN |
| 6.920003 | 7.200233 |
| 11.07279 | 11.13741 |
| 11.91157 | 11.85206 |
| NaN | NaN |
| 8.05859 | 8.102674 |
| 9.325442 | 9.230663 |
| 7.574907 | 7.770768 |
| 6.455072 | 6.377506 |
| 7.446304 | 7.069301 |

| | |
|---|---|
| 8.129261 | 7.797599 |
| 7.50519 | 7.727632 |
| 7.481641 | 7.70444 |
| 7.273172 | 7.266438 |
| 10.0018 | 9.942118 |
| 6.35275 | 6.281971 |
| 7.83865 | 7.916072 |
| 5.973756 | 6.701194 |
| 11.61626 | 11.67517 |
| 8.881987 | 8.873271 |
| 10.75458 | 10.51837 |
| 7.453783 | 7.036009 |
| 8.90946 | 8.40554 |
| 9.046874 | 9.041616 |
| 8.67913 | 7.905352 |
| 10.42542 | 10.30461 |
| 9.960628 | 9.826797 |
| 5.498505 | 5.036892 |
| 11.23712 | 11.36859 |
| 12.94307 | 13.0123 |
| 6.113894 | 6.168336 |
| 8.459499 | 8.575459 |
| 7.09368 | 6.974256 |
| 5.88729 | 5.805066 |
| 8.38159 | 8.399261 |
| 11.24967 | 11.22791 |
| 5.938021 | 6.793812 |
| 11.26004 | 11.05816 |
| 10.34564 | 10.4136 |
| 10.38511 | 10.63336 |
| 5.151042 | 5.44545 |
| 7.084109 | 7.178562 |
| NaN | NaN |
| 8.973099 | 8.811905 |
| 10.07183 | 9.862547 |
| 8.367313 | 8.347192 |
| 6.830348 | 5.702048 |
| 9.538201 | 9.403312 |
| 6.73276 | 6.832897 |
| 9.70795 | 9.894133 |
| 8.2129 | 8.22888 |
| 7.527859 | 7.471503 |
| NaN | NaN |
| 5.395646 | 4.606554 |
| 10.52634 | 10.44549 |
| NaN | NaN |
| 5.022748 | 5.245427 |

| | |
|---|---|
| 8.201993 | 8.268439 |
| 8.257946 | 8.100493 |
| 9.086359 | 9.044501 |
| 7.673392 | 7.381167 |
| 9.764893 | 9.807001 |
| NaN | NaN |
| 6.12044 | 5.929131 |
| 11.1405 | 10.97679 |
| 11.16318 | 11.21659 |
| 9.746604 | 10.04486 |
| 5.831125 | 5.863732 |
| 3.919109 | 2.842744 |
| 7.138127 | 7.089784 |
| 11.24665 | 11.11504 |
| 7.591074 | 7.811001 |
| NaN | NaN |
| 5.51802 | 5.555073 |
| 8.213444 | 8.348938 |
| 6.414219 | 5.855575 |
| 8.21864 | 8.543855 |
| 9.212543 | 8.999807 |
| 10.34913 | 10.35091 |
| 8.257621 | 6.505441 |
| 7.39864 | 7.092541 |
| 7.089423 | 6.759502 |
| 5.673801 | NaN |
| 10.46888 | 10.64795 |
| 6.909842 | 7.451453 |
| 8.211854 | 8.310421 |
| 9.469077 | 9.25598 |
| 13.45782 | 13.49931 |
| 8.379594 | 7.936384 |
| NaN | 7.627495 |
| 8.054028 | 8.441401 |
| 9.67631 | 9.598454 |
| 11.73609 | 11.61177 |
| 7.780476 | 7.852764 |
| 10.9455 | 11.07533 |
| 7.018873 | 7.241424 |
| 12.55193 | 12.51027 |
| 11.0989 | 10.97642 |
| 6.927793 | 6.926944 |
| 5.194043 | 4.809894 |
| 7.969293 | 7.921004 |
| 7.538375 | 7.550048 |
| 14.04556 | 14.1553 |
| 8.190045 | 8.106809 |

| | |
|---|---|
| 9.09653 | 8.829761 |
| 9.244449 | 8.937595 |
| 11.48355 | 11.53744 |
| 10.98296 | 11.0183 |
| 11.11818 | 10.99025 |
| 5.374084 | 4.518112 |
| 9.918515 | 10.18882 |
| 7.698836 | 7.698203 |
| 11.15724 | 11.17074 |
| 9.331043 | 9.72544 |
| 9.26131 | 9.094321 |
| 10.64036 | 10.49862 |
| 11.65452 | 11.6664 |
| 8.387444 | 8.448227 |
| 13.52568 | 13.72871 |
| 8.635303 | 12.03501 |
| 6.170256 | NaN |
| 7.929813 | 7.835938 |
| 8.530713 | 8.82117 |
| 12.90963 | 13.04304 |
| 7.147897 | 6.751051 |
| 9.336219 | 9.405508 |
| 9.331228 | 9.374484 |
| 7.649169 | 7.790286 |
| 13.91418 | 13.94475 |
| 7.874772 | 7.48482 |
| 11.50518 | 11.75798 |
| 6.376743 | 6.815147 |
| 6.562168 | 6.299043 |
| 8.96326 | 8.875777 |
| 8.826735 | 7.687612 |
| 10.28913 | 10.1899 |
| 8.478575 | 8.7595 |
| 6.625652 | 7.095265 |
| 8.614981 | 8.625388 |
| 10.47834 | 10.51013 |
| 6.566108 | 6.53094 |
| 9.892816 | 9.822858 |
| 6.483765 | 6.261736 |
| 8.525116 | 8.645903 |
| 13.95359 | 13.86857 |
| 6.650786 | 6.772583 |
| 9.219805 | 9.364003 |
| 9.78183 | 9.769766 |
| 6.570707 | 7.195588 |
| 9.19664 | 7.670311 |
| 7.673244 | 7.830382 |

-Bruker Confidential-

| | |
|---|---|
| 6.09645 | 5.68945 |
| 9.697955 | 9.411341 |
| 7.163781 | 7.827547 |
| 7.880752 | 7.562028 |
| 7.702875 | 7.382673 |
| 9.721114 | 9.687088 |
| 8.087577 | 8.290346 |
| 13.15056 | 13.19328 |
| 9.042681 | 8.681129 |
| 7.87533 | 7.87927 |
| 9.631875 | 9.6344 |
| 10.81798 | 10.61947 |
| 7.711927 | 7.711125 |
| NaN | NaN |
| 11.77384 | 11.70906 |
| NaN | NaN |
| 4.061577 | 3.291184 |
| 9.93934 | 9.82678 |
| 7.694183 | 7.725554 |
| 7.331668 | 7.27204 |
| 7.343981 | 7.854294 |
| 11.38156 | 11.35458 |
| 8.242064 | 8.509056 |
| 6.234582 | 6.128514 |
| 12.06567 | 11.52063 |
| 7.092023 | 6.211502 |
| 7.679212 | 7.778025 |
| 8.779222 | 8.785021 |
| 11.73167 | 11.72706 |
| 9.818939 | 9.639643 |
| 9.390553 | 9.468719 |
| 7.852156 | 8.736265 |
| NaN | 5.814901 |
| 7.193947 | 6.179839 |
| 10.02525 | 10.08951 |
| 8.92462 | 8.876433 |
| 8.609628 | 8.591398 |
| 9.022464 | 9.000034 |
| 7.460626 | 6.889341 |
| NaN | 5.132748 |
| 9.559103 | 9.436857 |
| 12.02582 | 12.19455 |
| 7.705903 | 7.586084 |
| NaN | 5.502206 |
| 11.17377 | 11.02982 |
| 10.89434 | 11.02378 |
| 7.922605 | 7.810223 |

| | |
|---|---|
| 8.557148 | 8.545517 |
| 4.665009 | 5.17366 |
| 9.030725 | 8.794757 |
| 14.39774 | 14.35101 |
| 5.9524 | 5.848815 |
| 7.160492 | 6.928364 |
| 9.777445 | 9.611087 |
| 9.263284 | 9.294246 |
| 11.72251 | 11.50831 |
| NaN | NaN |
| 6.354725 | 6.152117 |
| 5.021112 | 4.712466 |
| 14.93175 | 14.8279 |
| 7.573788 | 7.741977 |
| 12.01737 | 12.23597 |
| NaN | NaN |
| 11.70756 | 11.79575 |
| NaN | 4.663036 |
| 4.875282 | 4.156342 |
| 7.231745 | 7.19496 |
| 8.626281 | 8.557484 |
| 6.788654 | 6.704655 |
| 10.22744 | 10.51484 |
| 7.212673 | 6.787395 |
| 5.254691 | 5.225793 |
| 6.262777 | 6.521169 |
| 13.03111 | 13.34376 |
| 7.599236 | 7.125149 |
| 3.877829 | NaN |
| 8.867165 | 8.673828 |
| 7.071874 | 7.100883 |
| 8.951463 | 8.93756 |
| 10.29828 | 10.27822 |
| 8.372919 | 9.130747 |
| 7.353838 | 6.999253 |
| 6.648616 | 6.302129 |
| 10.14808 | 10.12954 |
| 7.726931 | 7.71075 |
| 8.026443 | 7.962489 |
| 10.1762 | 9.90319 |
| 12.37362 | 12.59624 |
| 12.4062 | 12.1577 |
| 13.28516 | 13.22774 |
| 8.186966 | 7.846375 |
| 6.89719 | 7.125481 |
| 10.69654 | 10.58767 |
| 12.36228 | 12.49751 |

| | |
|---|---|
| 7.944826 | 7.933413 |
| 7.76641 | 7.733648 |
| 9.735763 | 10.01287 |
| 6.979187 | 6.734258 |
| 9.193167 | 9.162155 |
| 7.640451 | 7.571959 |
| 9.471396 | 9.215355 |
| 5.800223 | 5.640118 |
| 8.474348 | 8.27564 |
| 7.421904 | 7.636437 |
| 9.184968 | 9.082456 |
| 8.844418 | 8.932594 |
| 8.00668 | 8.067872 |
| 5.235213 | 5.094013 |
| 9.110018 | 9.140723 |
| 8.487537 | 8.615268 |
| 8.132616 | 8.054222 |
| 8.559856 | 8.293264 |
| 5.562245 | 5.70688 |
| 6.490289 | 6.361255 |
| 9.569959 | 9.568144 |
| 7.674468 | 7.519514 |
| 7.985551 | 8.225052 |
| 6.859041 | 6.422249 |
| 9.405501 | 9.479959 |
| 9.097815 | 9.101161 |
| NaN | 5.65309 |
| 5.979966 | 5.401826 |
| 7.34321 | 7.129915 |
| 9.148526 | 9.567676 |
| 8.899576 | 8.976261 |
| 6.048934 | 5.239832 |
| 6.207873 | 6.25457 |
| 9.973387 | 10.06539 |
| 7.40188 | 7.406973 |
| 3.14622 | NaN |
| 10.03117 | 10.02518 |
| 10.45112 | 10.49345 |
| 7.655242 | 7.295428 |
| 11.28773 | 11.34372 |
| 8.2826 | 8.145436 |
| 8.292111 | 8.226646 |
| NaN | NaN |
| 7.261363 | 7.162049 |
| 7.300012 | 7.407064 |
| NaN | 5.441844 |
| 5.310404 | 3.343308 |

| | |
|---|---|
| NaN | NaN |
| 8.584425 | 8.624854 |
| 6.204025 | 5.929607 |
| 11.10796 | 11.19271 |
| 10.52992 | 10.356 |
| 8.523283 | NaN |
| 7.7655 | 7.790311 |
| 5.912584 | 5.792576 |
| 10.33114 | 10.45229 |
| 9.459901 | 9.386352 |
| 6.300488 | 6.516634 |
| 11.00569 | 11.12495 |
| 10.46808 | 10.60302 |
| 7.580697 | 7.514589 |
| 10.66812 | 10.67467 |
| 11.35429 | 11.39769 |
| 11.89448 | 11.896 |
| 9.170033 | 8.919417 |
| 9.31194 | 9.249399 |
| 10.88829 | 10.87958 |
| 9.564349 | 9.764112 |
| 8.33008 | 8.390941 |
| 5.951174 | 5.825271 |
| 6.842887 | 7.474006 |
| 6.995376 | 7.253211 |
| 7.615183 | 7.468791 |
| 10.73988 | 10.87832 |
| 7.762901 | 7.810591 |
| 9.418253 | 9.332627 |
| NaN | 6.883866 |
| 7.602587 | 7.907644 |
| 7.048566 | 6.879105 |
| 6.870738 | 6.903422 |
| NaN | NaN |
| 9.278556 | 9.443942 |
| 6.809337 | 6.277578 |
| 11.86875 | 11.91355 |
| NaN | NaN |
| 9.47664 | 8.607356 |
| 9.577498 | 9.788615 |
| 7.828681 | 8.157169 |
| 9.569118 | 9.513314 |
| 12.22782 | 12.38089 |
| 6.870696 | 6.691176 |
| 12.4676 | 12.51397 |
| 6.00685 | 6.067404 |
| 10.60891 | 10.99636 |

| | |
|---|---|
| 8.377611 | 8.463821 |
| 7.306086 | 7.138326 |
| 8.327874 | 8.557264 |
| 7.096804 | 7.164768 |
| 10.82445 | 10.86219 |
| 9.442656 | 9.684799 |
| 6.332657 | 6.6079 |
| 10.96474 | 11.13916 |
| 7.463428 | 7.042339 |
| 8.139923 | 6.504704 |
| 10.31523 | 10.36293 |
| 6.032604 | NaN |
| 10.23783 | 10.24989 |
| 8.216243 | 8.221184 |
| 7.732422 | 7.746065 |
| 7.256661 | NaN |
| 5.971157 | 6.086983 |
| 8.524117 | 8.509911 |
| 8.483253 | 8.477112 |
| 7.429254 | 8.025791 |
| 9.248654 | 9.008477 |
| 6.79249 | 6.683113 |
| 9.679715 | 9.785326 |
| 7.40183 | 7.345428 |
| 6.82774 | 6.636439 |
| 7.7796 | 7.770898 |
| 6.969694 | 7.139338 |
| 5.902487 | 6.050029 |
| 6.178496 | 6.112406 |
| 8.910511 | 8.512489 |
| 8.202243 | 8.369688 |
| 5.45455 | 4.594168 |
| 7.215566 | 7.196523 |
| 7.699823 | 7.72421 |
| 8.799515 | 8.788704 |
| 8.474336 | 8.616497 |
| 12.66644 | 12.32341 |
| 9.632298 | 9.727969 |
| 8.520299 | 8.946199 |
| 9.114778 | 8.973247 |
| 5.159718 | 4.55904 |
| 5.468882 | 5.820335 |
| 9.901193 | 9.766701 |
| 8.313479 | 8.400851 |
| 5.997899 | 6.173557 |
| 11.26603 | 11.16387 |
| 7.392702 | 7.352738 |

| | |
|---|---|
| 7.515224 | 7.275251 |
| 6.860455 | 6.998796 |
| 9.082566 | 8.91573 |
| 6.266595 | 6.324141 |
| 11.05913 | 10.92614 |
| 5.177308 | 6.415385 |
| 6.743123 | 6.574232 |
| 7.816782 | 7.755127 |
| 4.214769 | 5.07662 |
| 7.79459 | 7.86877 |
| 5.547165 | 5.559869 |
| 6.427526 | 5.969391 |
| 6.180473 | 5.412421 |
| 9.645485 | 9.549603 |
| 4.774423 | 4.836354 |
| 4.980393 | 4.990158 |
| 6.295879 | 6.029899 |
| 7.733598 | 7.958355 |
| 7.672776 | 7.445058 |
| 8.166248 | 8.196886 |
| NaN | 1.678528 |
| 11.54462 | 11.58709 |
| 9.006082 | 9.039661 |
| 9.237652 | 9.2462 |
| 8.204547 | 8.09933 |
| 7.409799 | 7.283916 |
| 9.028812 | 9.133085 |
| 6.392794 | 5.979079 |
| 11.70796 | 11.65385 |
| 11.05447 | 10.79434 |
| 8.522259 | 8.06984 |
| 7.255676 | 7.160651 |
| 6.321064 | 6.34545 |
| 6.731778 | 7.227983 |
| 7.355332 | 6.9876 |
| 10.45823 | 10.53743 |
| 10.15346 | 10.47203 |
| 8.327923 | 8.341064 |
| 7.059466 | 7.04405 |
| 8.050591 | 8.225362 |
| 6.159375 | 6.574839 |
| 9.515779 | 9.671073 |
| 7.268229 | 7.339402 |
| 10.22725 | 10.24492 |
| 11.81073 | 12.17368 |
| 9.002871 | 8.997123 |
| 6.327456 | 7.359547 |

| | |
|---|---|
| 5.59952 | 5.546491 |
| 6.097324 | 5.494147 |
| 7.124033 | 6.957366 |
| 11.2693 | 10.75285 |
| 6.254562 | 5.779192 |
| 8.014702 | 8.111654 |
| 9.250291 | 9.121474 |
| 7.692048 | 7.793173 |
| 8.214611 | 8.160101 |
| NaN | NaN |
| 12.22017 | 11.87159 |
| 9.794072 | 9.807471 |
| 8.611231 | 8.581936 |
| 6.662313 | 6.661146 |
| 8.619688 | 8.294932 |
| 6.200868 | 6.017913 |
| 7.096393 | 7.127154 |
| NaN | 7.135379 |
| 7.501487 | 7.246062 |
| 4.059308 | 3.367378 |
| 7.262087 | 7.111756 |
| NaN | 6.048082 |
| 6.707639 | 6.035077 |
| 8.768795 | 8.800043 |
| 8.125362 | 8.255005 |
| NaN | 6.288297 |
| 9.899367 | 9.953798 |
| 6.189387 | 5.962455 |
| 7.94037 | 8.332704 |
| 8.349249 | 8.153147 |
| 6.086224 | 6.23706 |
| 9.09691 | 8.829144 |
| 6.851361 | 6.748838 |
| 6.970676 | 7.295511 |
| 9.736115 | 9.702876 |
| 9.366148 | 9.349871 |
| 7.166187 | 7.270617 |
| 7.063368 | 7.225603 |
| 7.070132 | 7.212118 |
| 6.392797 | 6.597606 |
| 8.588222 | 7.613708 |
| 12.08243 | 12.04782 |
| 5.615295 | 5.70566 |
| 6.631259 | 6.415937 |
| 7.399197 | 7.298892 |
| 4.098828 | 4.586542 |
| 6.762276 | 6.794042 |

| | |
|---|---|
| 7.783003 | 7.562992 |
| 7.539892 | 7.468109 |
| 8.595185 | 8.85626 |
| 7.919841 | 7.989882 |
| 10.23626 | 10.27423 |
| 8.336688 | 8.298479 |
| 5.312746 | 4.881866 |
| 11.01475 | 11.00165 |
| 6.798953 | 6.685333 |
| 7.703397 | 7.310583 |
| 7.875129 | 7.959232 |
| 7.451805 | 7.54397 |
| 5.744609 | NaN |
| 9.301917 | 9.131305 |
| 11.61582 | 12.03529 |
| 5.132156 | 5.22515 |
| 8.395146 | 8.647688 |
| 8.132563 | 8.249735 |
| 6.980669 | NaN |
| 5.978477 | 6.060015 |
| 6.596714 | 6.547227 |
| 8.358038 | 8.07119 |
| 7.968182 | 7.851613 |
| 9.110211 | 8.792247 |
| 8.703113 | 8.620979 |
| 9.198961 | 9.090921 |
| 9.191852 | 9.143759 |
| 6.602167 | 6.455047 |
| 10.44548 | 10.52191 |
| 11.70371 | 11.68328 |
| 11.13055 | 11.20838 |
| 4.865606 | 4.264632 |
| 5.510949 | 5.532226 |
| 8.198482 | 7.963697 |
| 13.49153 | 13.52465 |
| 4.896097 | 4.898532 |
| 8.702826 | 8.81783 |
| 7.039253 | 7.153674 |
| 9.338626 | 9.273809 |
| 11.11795 | 11.00691 |
| 4.949989 | 5.387957 |
| 8.046689 | 7.348294 |
| 6.82739 | 6.382295 |
| 7.828963 | 7.741015 |
| 8.432636 | 8.320809 |
| 6.669974 | 6.651206 |
| 12.83749 | 12.87654 |

| | |
|---|---|
| 6.734614 | 7.131448 |
| 10.22148 | 10.76042 |
| NaN | NaN |
| 8.265043 | 8.72219 |
| NaN | NaN |
| 9.644246 | 9.741091 |
| 7.590128 | 7.481658 |
| 8.55673 | 8.568334 |
| 11.1198 | 10.9387 |
| 4.680738 | 4.04115 |
| 8.299839 | 8.515388 |
| 6.600414 | 7.186182 |
| 6.655167 | 6.419692 |
| 6.008618 | 6.413755 |
| 5.85987 | 5.91674 |
| 4.473084 | 5.080359 |
| 5.384741 | 5.838359 |
| 9.809318 | 9.99941 |
| 13.1739 | 13.14314 |
| 7.750648 | 7.748373 |
| 6.419232 | 6.689162 |
| 8.220232 | 8.115391 |
| 5.773098 | 5.866459 |
| 4.969037 | NaN |
| 6.300293 | 5.967609 |
| 8.499192 | 8.102253 |
| 8.509865 | 8.406624 |
| 5.930161 | 5.813316 |
| 6.580596 | 6.207895 |
| 10.67356 | 10.49392 |
| 10.50992 | 10.39241 |
| 4.183889 | 4.569696 |
| 6.334717 | 6.22753 |
| 11.0858 | 11.04824 |
| 9.047421 | 8.930168 |
| 7.359261 | 7.369993 |
| NaN | 4.3766 |
| 7.150576 | 7.144509 |
| 9.274387 | 9.039761 |
| 6.150587 | 6.061609 |
| 7.85121 | 7.592891 |
| 7.523024 | 7.580219 |
| 5.848605 | 7.04198 |
| 7.353186 | 7.388959 |
| 6.646832 | 7.084608 |
| 5.721123 | 5.524148 |
| 9.508256 | 10.80681 |

| | |
|---|---|
| 6.46547 | 6.262934 |
| NaN | NaN |
| 7.831286 | 8.00057 |
| 10.04569 | 10.05027 |
| 7.528985 | 7.705945 |
| 12.17265 | 11.8674 |
| 9.918988 | 9.848777 |
| 7.757773 | 7.777673 |
| 7.699252 | 7.924678 |
| 7.875851 | 7.865437 |
| 5.668081 | 6.161543 |
| 4.548124 | 4.437676 |
| 7.958304 | 7.642848 |
| 7.376604 | 6.866277 |
| 8.868807 | 8.764579 |
| 10.03333 | 10.08513 |
| 8.244725 | 8.333843 |
| 7.307327 | 7.34049 |
| 8.772223 | 8.900787 |
| 11.01136 | 10.81861 |
| 7.227768 | 4.115969 |
| 7.641038 | 7.605477 |
| 8.124898 | 8.181443 |
| 10.98152 | 11.05166 |
| 11.14186 | 10.8262 |
| 10.59631 | 10.36086 |
| 8.812377 | 8.888021 |
| 11.67081 | 11.88017 |
| 7.967977 | 7.738309 |
| 7.591643 | 7.70949 |
| 6.511246 | 5.999116 |
| 6.711538 | 6.681313 |
| 7.366373 | 6.968421 |
| 6.755905 | 6.879932 |
| 12.7207 | 12.68791 |
| 8.530784 | 8.308475 |
| 7.493531 | 7.436502 |
| 8.333173 | 7.894854 |
| 10.14238 | 10.28358 |
| 3.875058 | 4.508385 |
| 9.727436 | 9.601536 |
| NaN | NaN |
| 9.18561 | 9.071349 |
| 9.478252 | 9.200856 |
| 6.77073 | 6.435713 |
| 7.890378 | 7.698282 |
| 9.68662 | 9.682262 |

| | |
|---|---|
| 12.61625 | 12.42889 |
| 9.340855 | 9.201992 |
| 12.43023 | 12.42196 |
| 5.592976 | 5.748643 |
| 8.384313 | 8.636838 |
| 8.946952 | 8.898686 |
| 7.932195 | 8.159939 |
| 9.496748 | 9.59976 |
| 7.590448 | 7.681068 |
| 8.740415 | 8.637799 |
| 8.738312 | 8.627579 |
| 6.588572 | 6.634776 |
| NaN | NaN |
| 7.879587 | 7.992968 |
| 6.079292 | 6.001286 |
| 8.909703 | 8.837118 |
| 6.279447 | 6.010164 |
| 9.594714 | 9.455762 |
| 7.646948 | 7.502464 |
| 4.952342 | 5.183649 |
| 5.898087 | 6.368471 |
| 5.623824 | 4.122371 |
| 8.352272 | 8.30284 |
| 7.798625 | 7.726487 |
| 6.377622 | 6.530478 |
| 6.515849 | 6.469731 |
| 8.444844 | 8.145031 |
| 9.117525 | 8.989706 |
| 7.549585 | 8.003676 |
| 7.031939 | 6.664464 |
| 6.999937 | 6.867502 |
| 6.561589 | 6.319633 |
| 6.9931 | 6.081842 |
| 8.805226 | 9.040578 |
| 4.87516 | 5.286002 |
| 10.43906 | 10.55745 |
| 5.061734 | 3.892331 |
| 6.996102 | 6.810997 |
| 6.198979 | NaN |
| 5.865775 | 5.517385 |
| 12.26575 | 12.29289 |
| 13.75474 | 13.84573 |
| 13.46357 | 13.41588 |
| 8.691139 | 8.559953 |
| 6.900095 | 6.596937 |
| 5.180441 | 5.505942 |
| 10.94151 | 11.14875 |

-Bruker Confidential-

| | |
|---|---|
| 12.61626 | 12.55825 |
| 9.130954 | 9.820662 |
| 10.09984 | 10.50672 |
| 8.396235 | 8.621209 |
| NaN | 3.456684 |
| 10.61139 | 10.27111 |
| 12.38419 | 12.42284 |
| 9.536314 | 8.987157 |
| 8.909849 | 8.861228 |
| 9.164141 | 9.085797 |
| 8.329708 | 7.782324 |
| 12.59748 | 12.66037 |
| 10.99723 | 10.77275 |
| 6.321539 | 6.418789 |
| 4.194036 | 5.574025 |
| 7.947319 | 7.859076 |
| 7.863188 | 8.67872 |
| 6.497691 | 6.580849 |
| 8.056425 | 7.757663 |
| 8.481727 | 8.657575 |
| NaN | NaN |
| 7.470689 | 7.68497 |
| 6.973968 | 7.163862 |
| 6.904816 | 7.137043 |
| 10.57827 | 10.1391 |
| 7.356397 | 7.036166 |
| 6.295973 | 5.866071 |
| 8.334977 | 8.083806 |
| 7.390725 | 7.448966 |
| NaN | NaN |
| NaN | NaN |
| 8.269072 | 8.428763 |
| 10.28658 | 10.11247 |
| 11.5293 | 11.51359 |
| 7.152729 | 6.891251 |
| 6.318191 | 6.217613 |
| 9.122641 | 8.589378 |
| 9.28903 | 9.3217 |
| 7.469757 | 7.263083 |
| 7.057089 | 7.079247 |
| 7.46471 | 7.796347 |
| 9.250488 | 9.556772 |
| 6.780393 | 6.627509 |
| 6.361241 | 6.09621 |
| 9.340279 | 9.309716 |
| 8.435348 | 8.422684 |
| 9.549533 | 9.483102 |

| | |
|---|---|
| 10.52658 | 10.5261 |
| 11.82883 | 11.87334 |
| 6.279529 | 6.43091 |
| 11.56283 | 11.46748 |
| 9.430648 | 8.983874 |
| 11.34423 | 11.31511 |
| 6.48913 | 6.176536 |
| 8.590426 | 8.760943 |
| 11.66902 | 11.76964 |
| 8.212318 | 8.194068 |
| 7.261133 | 7.001846 |
| 6.881348 | 6.871532 |
| 6.400178 | 6.537698 |
| 6.401622 | 6.070937 |
| 8.693667 | 8.452809 |
| 4.725683 | 4.91889 |
| 6.316798 | 6.218558 |
| 5.782413 | 6.616657 |
| 8.768473 | 8.751944 |
| 6.848101 | 6.554925 |
| 10.03586 | 10.02096 |
| 7.558132 | 7.766626 |
| NaN | NaN |
| 7.723363 | 7.239949 |
| 6.104142 | 6.494662 |
| 5.575864 | 5.560401 |
| 9.101765 | 9.251536 |
| 5.924621 | 5.258173 |
| 6.318159 | 6.136189 |
| 6.360572 | 6.517812 |
| 8.685302 | 8.513579 |
| 8.982346 | 8.976882 |
| 8.626451 | 8.594633 |
| 6.498058 | 6.39727 |
| 7.853053 | 7.993486 |
| 8.265444 | 8.15607 |
| 5.771767 | 5.512933 |
| 8.796717 | 8.478759 |
| 7.010086 | NaN |
| 10.78463 | 10.73551 |
| 6.891839 | 6.581735 |
| 7.627754 | 7.314724 |
| NaN | NaN |
| 6.18041 | 5.974484 |
| 12.31021 | 12.28823 |
| 9.884541 | 9.779353 |
| 9.154377 | 8.919346 |

| | |
|---|---|
| 6.552577 | 6.3783 |
| 11.81793 | 11.80379 |
| 5.268695 | 4.745783 |
| 6.730291 | 6.730666 |
| 8.505963 | 8.525732 |
| 9.297053 | 9.400514 |
| 8.02878 | 8.228336 |
| 8.006074 | 8.169526 |
| 6.501417 | 6.792722 |
| 10.57967 | 10.7298 |
| 7.554462 | 7.464229 |
| 4.593655 | NaN |
| 6.724641 | 6.754087 |
| NaN | NaN |
| 7.083073 | 7.018753 |
| 7.22699 | 7.383249 |
| 8.810193 | 8.701151 |
| 6.165427 | 7.261553 |
| 7.621347 | 7.27548 |
| 7.383914 | 7.334305 |
| 9.577892 | 9.65565 |
| 11.7917 | 11.73447 |
| 9.962122 | 9.848283 |
| 5.26538 | 5.003771 |
| 9.721387 | 9.64655 |
| 8.005943 | 7.769861 |
| 9.403732 | 9.241933 |
| 7.604702 | 7.860735 |
| 12.08139 | 11.64527 |
| 8.353709 | 8.270989 |
| 9.982746 | 9.792423 |
| 6.117147 | NaN |
| 5.606676 | 6.314738 |
| 6.136316 | 5.978727 |
| 7.211058 | 7.421515 |
| 10.4468 | 10.2838 |
| 7.766222 | 7.011879 |
| 11.33962 | 11.44164 |
| 5.854966 | 5.932774 |
| 7.591031 | 7.23326 |
| 10.45795 | 10.36849 |
| 8.257263 | 8.370386 |
| 10.62905 | 11.01633 |
| 9.482032 | 9.426147 |
| 6.635457 | 6.381795 |
| 11.1432 | 11.08881 |
| 7.367211 | 7.805383 |

-Bruker Confidential-

| | |
|---|---|
| 8.793779 | 8.963272 |
| 8.156986 | 8.147304 |
| 7.407807 | 7.798464 |
| 5.702633 | 5.392679 |
| 7.504955 | 7.57363 |
| 8.247585 | 8.116318 |
| 8.209605 | 8.088985 |
| 9.003347 | 8.882656 |
| 8.542481 | 8.379829 |
| 8.765019 | 8.665478 |
| 9.534928 | 9.557735 |
| 9.293633 | 9.20099 |
| 8.263291 | 8.3746 |
| 8.950156 | 8.83976 |
| 9.474328 | 9.229807 |
| 7.881257 | 7.271145 |
| 10.03232 | 9.877626 |
| 11.12575 | 11.16326 |
| 4.500088 | 3.590865 |
| 7.079984 | 7.157521 |
| 6.203285 | 6.30292 |
| 11.56521 | 10.55205 |
| 7.144724 | 6.76123 |
| 9.490644 | 9.319635 |
| 11.86143 | 11.87286 |
| 9.035725 | 8.84949 |
| NaN | NaN |
| 10.20978 | 9.6886 |
| 9.687417 | 9.74339 |
| 6.754776 | 7.581573 |
| 9.560562 | 9.380467 |
| 10.52443 | 10.62785 |
| 9.188849 | 9.107556 |
| 6.849833 | 6.696639 |
| NaN | 6.490231 |
| 9.811477 | 9.836699 |
| 7.97717 | 7.664261 |
| 11.07505 | 10.98938 |
| 7.715497 | 7.741874 |
| 6.421599 | 6.750459 |
| 6.586868 | 6.482458 |
| 8.577985 | 8.384787 |
| 6.937457 | 7.758053 |
| 6.650889 | 6.554552 |
| 7.793602 | 7.813806 |
| 6.466577 | 6.663652 |
| 8.011285 | 8.090959 |

| | |
|---|---|
| 9.731387 | 10.00225 |
| 12.48224 | 12.27668 |
| 7.202143 | 7.462275 |
| 7.894745 | 7.554303 |
| 9.944094 | 9.944415 |
| 8.488126 | 8.422096 |
| 8.295372 | 8.262566 |
| 3.524609 | 3.199456 |
| 8.551757 | 8.325104 |
| 9.354827 | 9.457799 |
| NaN | NaN |
| 9.015464 | 8.795506 |
| 4.652775 | 3.957836 |
| 8.465982 | 8.246701 |
| 5.640711 | 6.164237 |
| 6.275347 | 6.025195 |
| 6.344676 | 6.232224 |
| 10.66102 | 10.5361 |
| 8.957364 | 8.69844 |
| NaN | NaN |
| 10.4331 | 10.05133 |
| 6.844396 | NaN |
| 7.131469 | 7.083313 |
| 6.968086 | 7.403511 |
| 8.002956 | 8.118775 |
| 7.681026 | 7.848175 |
| 10.49045 | 10.48645 |
| 9.089218 | 9.059546 |
| 7.569336 | 7.522897 |
| 5.220715 | 4.504512 |
| 6.114853 | 5.81698 |
| 7.024999 | 6.85071 |
| 7.071959 | 7.014701 |
| 6.267879 | 5.927629 |
| 5.671903 | 6.020499 |
| 5.705282 | 5.785186 |
| 6.861249 | 7.020664 |
| 9.472992 | 9.583248 |
| 3.764155 | 4.593487 |
| 6.647248 | 6.369139 |
| 8.66199 | 8.989699 |
| 5.247059 | 4.759489 |
| 9.533887 | 9.377575 |
| NaN | 6.541424 |
| 6.0917 | 6.237102 |
| 6.70861 | 6.655267 |
| NaN | 6.030434 |

| | |
|---|---|
| 9.005355 | 9.087468 |
| 8.095078 | 8.2057 |
| 10.90173 | 10.92917 |
| 9.300434 | 9.503401 |
| 6.135187 | 6.275857 |
| 7.715421 | 7.532552 |
| 5.100461 | 4.748684 |
| 6.693047 | 6.501935 |
| 7.70841 | 7.596872 |
| 4.769945 | 4.887667 |
| 6.966211 | 6.745399 |
| 10.52971 | 10.32674 |
| 6.596056 | 6.391375 |
| 7.437148 | 7.531521 |
| 7.368938 | 7.628185 |
| 12.33689 | 12.20814 |
| 8.279977 | 8.130293 |
| 7.353552 | 7.307937 |
| 9.084138 | 9.058789 |
| 6.894566 | 7.064074 |
| 10.24493 | 10.24218 |
| 9.06477 | 8.800368 |
| NaN | 5.049274 |
| 10.49517 | 10.45522 |
| 6.879988 | 6.921789 |
| 5.846582 | NaN |
| 6.07038 | 4.942626 |
| 12.21343 | 12.01811 |
| 6.678527 | 6.493727 |
| 8.232894 | 8.197822 |
| 10.49609 | 10.54938 |
| 7.041521 | 7.051665 |
| 5.759913 | 5.914873 |
| 8.309966 | 8.533869 |
| 8.946848 | 8.712276 |
| 6.790322 | 6.719711 |
| 8.51144 | 8.186555 |
| 7.402664 | 7.492289 |
| 6.757793 | 6.401604 |
| 5.839548 | 5.864383 |
| 8.378772 | 8.608916 |
| 8.954661 | 8.861021 |
| 4.800246 | 4.550384 |
| 7.604989 | 7.568455 |
| 9.832397 | 10.01712 |
| 4.2829 | 4.719099 |
| 5.83525 | 5.90167 |

| | |
|---|---|
| 8.087763 | 7.992132 |
| 9.776043 | 9.746934 |
| 8.370511 | 8.433886 |
| 8.633919 | 8.844728 |
| 9.402 | 9.611485 |
| 11.64327 | 11.49254 |
| 10.27143 | 10.34323 |
| 11.99296 | 11.97968 |
| 13.04766 | 13.06397 |
| 6.124036 | 6.291753 |
| 5.996554 | 5.797957 |
| 6.324995 | 6.40343 |
| 6.726479 | 6.7672 |
| 7.847373 | 7.874426 |
| 8.460624 | 8.589262 |
| 9.536449 | 9.697912 |
| 6.204562 | 5.795941 |
| 9.314012 | 8.962819 |
| 10.04627 | 10.08933 |
| 10.14089 | 9.96883 |
| NaN | NaN |
| 8.325535 | 8.389009 |
| 9.972783 | 9.681541 |
| 5.903396 | 6.588286 |
| 5.363428 | 4.448906 |
| 9.399772 | 9.408216 |
| 10.12948 | 10.09617 |
| 6.455554 | 5.402832 |
| 8.127232 | 8.481508 |
| 7.942315 | 7.944951 |
| 6.672055 | 6.747879 |
| 5.147195 | 5.791221 |
| 6.52136 | 6.611047 |
| 7.871479 | 7.817367 |
| 6.471624 | 6.029209 |
| 10.10727 | 10.15094 |
| 9.181678 | 9.027811 |
| 7.50249 | 6.300865 |
| 6.089696 | 6.13271 |
| 6.297331 | 5.572334 |
| NaN | NaN |
| 5.37143 | 4.592088 |
| 5.38363 | 5.115808 |
| 8.708743 | 8.381203 |
| NaN | NaN |
| 6.463612 | 7.135452 |
| 6.552923 | 6.476171 |

| | |
|---|---|
| 10.60604 | 10.68731 |
| 6.05162 | 5.995941 |
| 6.574338 | 6.800808 |
| 11.39782 | 11.3518 |
| 8.963784 | 9.519988 |
| 6.310899 | 6.388824 |
| 7.942727 | 7.804452 |
| 8.28096 | 8.250069 |
| 10.45524 | 10.20084 |
| 8.026497 | 8.091609 |
| NaN | NaN |
| 13.11103 | 12.8709 |
| 5.642698 | 5.658248 |
| 6.446078 | 6.525411 |
| 6.483028 | 6.551522 |
| 9.177768 | 9.249029 |
| 7.558645 | 7.95794 |
| 6.541343 | 6.539646 |
| 9.034299 | 9.186461 |
| 9.200773 | 9.219512 |
| 10.34355 | 10.30159 |
| 6.584996 | 6.476041 |
| 7.105352 | 6.42313 |
| 6.487863 | 6.550378 |
| 10.59189 | 10.57401 |
| 6.706841 | 6.514716 |
| 7.363653 | 7.400515 |
| 9.624229 | 9.829064 |
| 8.327173 | 8.119511 |
| 9.019763 | 9.060383 |
| 7.904933 | 7.713487 |
| 6.239885 | 6.227157 |
| 6.349031 | 6.03044 |
| 6.524661 | 6.210298 |
| 8.323326 | 8.30934 |
| 8.62526 | 8.060585 |
| 7.138173 | 6.9991 |
| 9.611143 | 9.572289 |
| 9.190475 | 8.994675 |
| 10.48591 | 10.12147 |
| 9.442602 | 9.669645 |
| 6.841674 | 6.870542 |
| 6.105098 | 6.11799 |
| 5.542791 | 5.021218 |
| 7.45045 | 8.008138 |
| NaN | NaN |
| 7.225813 | 7.375475 |

| | |
|---|---|
| 7.421015 | 7.443157 |
| 8.887257 | 8.79827 |
| 7.982749 | 7.983039 |
| NaN | NaN |
| 8.191783 | 7.98879 |
| 8.267484 | 8.102051 |
| 6.372704 | 6.317263 |
| 9.926311 | 10.02445 |
| 8.411053 | 8.529558 |
| 5.497965 | 5.236999 |
| 8.03178 | 7.921753 |
| 8.383357 | 8.451608 |
| 6.538175 | 5.735357 |
| 6.365082 | 6.181343 |
| 7.412927 | 7.531874 |
| 3.716309 | 3.404253 |
| 5.666666 | 5.314214 |
| 6.010274 | 6.675471 |
| 8.481842 | 8.609149 |
| 7.413508 | 7.228917 |
| 7.944117 | 8.092622 |
| 6.898883 | 7.072316 |
| 5.406953 | 4.937415 |
| NaN | NaN |
| 8.328853 | 8.284307 |
| 7.483782 | 7.688755 |
| 8.107246 | 8.289382 |
| 9.131377 | 8.744994 |
| 7.512537 | 7.40308 |
| 8.545925 | 8.61997 |
| 8.996168 | 9.483981 |
| 8.830955 | 8.885833 |
| NaN | NaN |
| 8.528784 | 8.895123 |
| 6.027352 | 6.042665 |
| 7.415329 | 7.23828 |
| 7.414783 | 7.334394 |
| 10.64902 | 10.87166 |
| 10.36479 | 10.33504 |
| 7.943635 | 7.910037 |
| 6.886799 | 7.044527 |
| 6.608501 | 5.279941 |
| 8.151358 | 8.050921 |
| 12.78272 | 12.50776 |
| 11.93205 | 11.81998 |
| 7.125063 | 7.203689 |
| 6.208017 | 6.627439 |

-Bruker Confidential-

| | |
|---|---|
| 8.456724 | 8.200301 |
| 9.096699 | 8.615825 |
| 6.284704 | 6.143366 |
| NaN | 5.358257 |
| 5.937773 | 5.945366 |
| 6.847324 | 7.401625 |
| 6.677787 | 6.422767 |
| 7.840039 | 7.786228 |
| 7.137125 | 5.792835 |
| 7.846103 | 8.786448 |
| 9.209279 | 9.033753 |
| 11.56585 | 11.56263 |
| 10.72295 | 10.93996 |
| 8.144165 | 7.696115 |
| 6.937185 | 7.075714 |
| 7.050689 | 6.904862 |
| 7.169258 | 7.25201 |
| 9.408444 | 9.126759 |
| 9.493159 | 9.287944 |
| 10.61022 | 10.69085 |
| 7.586191 | 7.662618 |
| 7.035733 | 7.189324 |
| 3.77939 | 4.85399 |
| 4.882008 | 4.964973 |
| 9.637006 | 9.7258 |
| 7.909854 | 7.639817 |
| 7.193543 | 7.642662 |
| 6.066436 | 5.948058 |
| 9.755315 | 9.94442 |
| 7.363411 | 7.091092 |
| 9.356495 | 9.313123 |
| 7.351456 | 7.264388 |
| 7.585768 | 7.649272 |
| 9.421033 | 9.49333 |
| 9.261439 | 9.281892 |
| 8.266744 | 8.491637 |
| 8.452733 | 8.6591 |
| 8.106269 | 8.094415 |
| 8.556992 | 8.577289 |
| 7.68861 | 7.830149 |
| 10.82614 | 10.72974 |
| 8.419613 | 8.810461 |
| 8.177866 | 8.107347 |
| 8.642083 | 8.705088 |
| 6.680264 | 7.048588 |
| 7.391743 | 7.427401 |
| 6.270998 | 6.386981 |

| | |
|---|---|
| 6.781464 | 6.802037 |
| 9.729039 | 9.472096 |
| NaN | 4.358809 |
| 6.369821 | 6.485991 |
| 7.296066 | 7.289426 |
| 6.422322 | 6.378097 |
| 5.817092 | 5.983466 |
| 7.564624 | 7.230474 |
| 6.620274 | 6.758795 |
| 4.522161 | 4.882442 |
| 5.601668 | 5.78654 |
| 7.445613 | 7.278581 |
| 7.074074 | 7.033892 |
| 6.158816 | 6.301486 |
| 6.258095 | 6.326948 |
| 7.343132 | 7.064144 |
| 11.40631 | 11.47529 |
| 10.4547 | 10.49629 |
| 11.95835 | 11.903 |
| 7.385882 | 7.653165 |
| 7.835737 | 8.042087 |
| 10.0086 | 8.239073 |
| 8.364315 | 8.558366 |
| 6.344484 | 6.383927 |
| 8.619727 | 8.195469 |
| NaN | 8.603206 |
| 9.079857 | 8.756729 |
| 11.20641 | 11.07544 |
| 4.83904 | 4.269297 |
| 8.819141 | 8.716447 |
| 9.525165 | 9.186376 |
| 5.857091 | 5.397093 |
| 6.058517 | 5.777412 |
| 8.309201 | 8.172821 |
| 7.264276 | 7.328826 |
| 7.44859 | 7.304309 |
| 4.633753 | 5.663605 |
| 5.639206 | 5.788417 |
| 7.29719 | 7.038937 |
| 6.11394 | 6.209542 |
| 7.812219 | 7.583459 |
| 5.686351 | 5.877597 |
| NaN | NaN |
| 7.334593 | 7.175529 |
| 7.899822 | 7.720757 |
| 4.664155 | 4.751771 |
| 4.802571 | 5.365622 |

| | |
|---|---|
| 8.058356 | 8.292026 |
| 7.408434 | 7.296745 |
| 10.27569 | 10.80536 |
| 10.88474 | 10.89443 |
| 7.918747 | 8.150448 |
| 8.074399 | 8.017674 |
| 9.612267 | 9.35234 |
| 8.9855 | 8.744396 |
| 8.219936 | 8.185623 |
| 6.109072 | 5.721673 |
| 7.665765 | 7.701104 |
| 8.05656 | 7.81969 |
| 6.447975 | 7.282839 |
| 9.775375 | 9.779222 |
| 7.696234 | 7.634394 |
| 7.57138 | 7.532197 |
| 7.377144 | 7.241414 |
| 7.212157 | 6.89881 |
| 8.491444 | 8.473452 |
| 9.640285 | 9.756543 |
| 8.34816 | 8.230543 |
| 6.857282 | 6.51615 |
| 6.029906 | 5.566553 |
| 9.214767 | 9.312497 |
| 8.06213 | 7.225503 |
| 8.058747 | 8.167713 |
| 6.240216 | 5.968689 |
| 7.855267 | 7.729441 |
| 7.382337 | 7.50174 |
| 9.826777 | 10.00208 |
| 10.12635 | 10.23817 |
| 6.367587 | 7.282497 |
| 8.329638 | 8.346408 |
| 5.484063 | 5.572649 |
| 7.26054 | 7.197174 |
| 5.053684 | 4.802134 |
| 6.214752 | 6.708071 |
| 5.561637 | 5.689726 |
| 9.045635 | 8.70593 |
| 10.73628 | 10.93423 |
| 8.972141 | 9.064163 |
| 7.745837 | 7.673753 |
| 6.090594 | 7.732672 |
| 7.988815 | 8.147749 |
| 8.664831 | 8.50974 |
| 6.038334 | 5.661882 |
| 8.042408 | 8.161959 |

| | |
|---|---|
| 7.076622 | 7.103291 |
| 5.372746 | 5.300664 |
| 5.07394 | 4.073936 |
| 6.483525 | 5.896961 |
| 6.780499 | 7.153112 |
| 7.110108 | 7.224011 |
| 9.058671 | 8.805533 |
| 7.620393 | 7.569044 |
| 4.51595 | 5.400466 |
| 7.004723 | 6.881684 |
| 8.970397 | 8.744364 |
| 9.263453 | 9.258543 |
| 6.709604 | 7.202678 |
| 10.4602 | 10.50216 |
| 7.217519 | 6.989566 |
| 9.196136 | 9.660724 |
| 6.844813 | 6.943142 |
| 8.43372 | 8.394014 |
| 9.760406 | 9.664348 |
| 6.772243 | 5.93828 |
| 11.33175 | 11.37432 |
| 7.106192 | 7.251448 |
| 10.79957 | 10.72114 |
| 6.255798 | NaN |
| 8.317461 | 7.992015 |
| 9.4617 | 9.421503 |
| 6.05742 | 5.999085 |
| 8.359113 | 8.345793 |
| 5.331812 | 4.685422 |
| 11.12939 | 11.35083 |
| 7.604404 | 7.639973 |
| 6.672075 | 6.540885 |
| 7.53471 | 7.509453 |
| 8.557783 | 8.858017 |
| 7.093409 | 7.331016 |
| 6.384725 | 6.633608 |
| 7.959258 | 7.597526 |
| NaN | NaN |
| NaN | 4.674553 |
| 6.32502 | 6.156653 |
| NaN | NaN |
| 8.452759 | 8.250336 |
| 6.903253 | 6.97095 |
| 6.63462 | NaN |
| 8.007733 | 7.962075 |
| 9.549577 | 9.469182 |
| 10.21048 | 10.15698 |

| | |
|---|---|
| 1.182722 | 2.174435 |
| 8.083158 | 8.070654 |
| 8.558723 | 8.515434 |
| 8.822174 | 8.674102 |
| 4.858928 | 4.757223 |
| 10.66689 | 10.79045 |
| NaN | NaN |
| NaN | NaN |
| NaN | NaN |
| 7.920652 | 8.696764 |
| 5.152272 | 4.85374 |
| 8.354807 | 8.321726 |
| 6.224079 | 5.791492 |
| NaN | 5.423368 |
| 7.377199 | 6.901447 |
| 7.039053 | 6.203381 |
| 6.215191 | 6.406212 |
| NaN | 6.786578 |
| 9.68249 | 9.32095 |
| 5.604913 | 5.83202 |
| 9.05147 | 9.071725 |
| 7.786497 | 7.761566 |
| 8.070497 | 7.679991 |
| 6.125216 | 5.706841 |
| 5.002419 | 5.189455 |
| 6.414619 | 6.738944 |
| NaN | 5.546845 |
| 6.796154 | 6.380158 |
| 5.776655 | 5.553501 |
| 5.680024 | 5.633065 |
| 5.450898 | 5.916536 |
| 6.716396 | 6.382359 |
| 3.94423 | 2.332828 |
| 7.683631 | 7.38469 |
| 7.916595 | 7.836124 |
| 6.325646 | 6.276068 |
| 6.170948 | 5.264921 |
| 5.571434 | 6.172263 |
| 8.348637 | 8.386442 |
| 8.590245 | 8.603947 |
| 7.301977 | 7.440098 |
| 7.766486 | 7.924921 |
| 6.330608 | 6.318592 |
| 6.327347 | 6.016624 |
| 6.996008 | 6.761917 |
| NaN | NaN |
| 3.970918 | 4.010364 |

| | |
|---|---|
| 6.490841 | 6.469596 |
| NaN | 8.982397 |
| 13.09021 | 12.99703 |
| 6.088079 | 4.733116 |
| 8.447577 | 8.633774 |
| 8.387815 | 8.407692 |
| 6.174645 | 7.031609 |
| 6.142006 | 6.009391 |
| 9.470164 | 9.532291 |
| 6.004302 | 6.103758 |
| 6.592418 | 5.622936 |
| 7.255663 | 7.350835 |
| 11.19765 | 11.16254 |
| 8.751412 | 8.928872 |
| 7.930427 | 8.26093 |
| 1.065923 | 1.577438 |
| 7.474888 | 7.473225 |
| 9.187985 | 9.039347 |
| 3.724574 | 3.605955 |
| 10.23351 | 10.31668 |
| 13.46524 | 13.66368 |
| 8.032045 | 7.450053 |
| 11.86576 | 11.83548 |
| 10.2352 | 10.3631 |
| 8.835923 | 8.542103 |
| 8.091307 | 7.652264 |
| 8.80167 | 8.852266 |
| 4.854194 | 5.113446 |
| 9.490986 | 9.591739 |
| 9.805637 | 9.427661 |
| 8.729795 | 8.191859 |
| NaN | NaN |
| 7.179394 | 7.322422 |
| 7.985261 | 8.613503 |
| 7.750309 | 7.825239 |
| 7.283448 | 7.336295 |
| 11.13559 | 11.05292 |
| 8.436003 | 8.577852 |
| 7.913452 | 8.088836 |
| 10.6125 | 10.50492 |
| 9.506919 | 9.670719 |
| 8.735222 | 8.696736 |
| 9.667194 | 9.576238 |
| 10.53664 | 10.56131 |
| 8.607065 | 8.739145 |
| 6.483733 | 6.258186 |
| 9.99422 | 10.09496 |

-Bruker Confidential-

| | |
|---|---|
| 8.070211 | 8.332002 |
| 10.61701 | 10.58428 |
| 9.47949 | 9.210235 |
| 8.170165 | 8.641246 |
| 10.31085 | 10.25439 |
| 7.435074 | 7.242579 |
| NaN | NaN |
| 9.615209 | 9.500218 |
| 9.031216 | 9.239136 |
| 7.50533 | 7.311083 |
| 7.699071 | 7.530646 |
| 9.842451 | 9.887936 |
| NaN | NaN |
| 9.672662 | 9.93441 |
| 8.276532 | 8.278516 |
| 6.729875 | 7.244512 |
| 8.130733 | 8.970425 |
| 7.683802 | 7.812035 |
| 6.791846 | 6.797361 |
| 6.188424 | 5.457489 |
| 9.00438 | 9.179851 |
| 9.81553 | 9.891859 |
| 7.891182 | 7.608991 |
| 8.328628 | 8.306223 |
| 9.47172 | 9.392141 |
| 10.37979 | 10.36946 |
| 7.461827 | 7.624991 |
| 7.60027 | 8.594551 |
| 7.442606 | 7.2674 |
| 8.682789 | 8.706058 |
| 6.303905 | 6.44492 |
| 10.63748 | 10.31585 |
| 7.907753 | 7.842036 |
| 8.266356 | 9.114274 |
| 8.974456 | 8.832952 |
| 6.522438 | 6.467448 |
| 8.629536 | 8.633984 |
| 12.56694 | 12.43849 |
| 11.96097 | 12.36056 |
| 8.239806 | 8.008251 |
| 6.606977 | 6.367291 |
| 8.980105 | 8.61032 |
| 12.11916 | 12.35128 |
| 10.63973 | 10.16035 |
| 8.895354 | 8.601234 |
| 9.520131 | 9.666802 |
| 10.46413 | 10.37931 |

| | |
|---|---|
| 10.99361 | 10.88424 |
| 10.22388 | 11.42537 |
| 10.08503 | 10.59839 |
| 11.15711 | 10.93688 |
| 7.862821 | 7.814751 |
| 7.434842 | 7.540452 |
| 13.8367 | 13.7408 |
| 11.93603 | 11.50642 |
| 9.380364 | 9.141038 |
| 12.86709 | 12.77596 |
| 6.183342 | 6.453388 |
| 10.77337 | 10.45544 |
| 9.176866 | 9.15685 |
| 10.34651 | 10.20999 |
| 11.16187 | 11.29617 |
| 7.66467 | 7.456515 |
| 7.659003 | 7.860858 |
| 8.769222 | 8.308392 |
| 10.10567 | 9.944222 |
| 11.61328 | 11.46243 |
| 10.73776 | 10.62352 |
| 8.239121 | 8.629463 |
| 10.40112 | 10.25384 |
| 7.373358 | 7.474133 |
| 6.782114 | 6.697228 |
| 10.20914 | 9.963293 |
| 5.675475 | NaN |
| 8.742729 | 8.816645 |
| 5.485803 | 5.710618 |
| 6.136004 | 6.33101 |
| 10.05566 | 10.1656 |
| 10.41751 | 10.38201 |
| 12.41316 | 12.31639 |
| 5.554185 | 5.494689 |
| 11.48597 | 11.19121 |
| 9.050958 | 9.049366 |
| 6.196657 | 6.699941 |
| 13.69973 | 13.50752 |
| 7.80588 | 8.096726 |
| 11.95541 | 11.92262 |
| 9.064285 | 9.166332 |
| NaN | 8.487865 |
| 8.788186 | 8.832252 |
| 10.35358 | 10.4829 |
| 9.454079 | 10.10958 |
| 10.25333 | 10.04062 |
| 9.390773 | 9.351542 |

| | |
|---|---|
| 8.69773 | 7.685387 |
| 9.102052 | 8.800259 |
| NaN | NaN |
| 8.618087 | 8.441482 |
| 9.504218 | 9.461367 |
| 8.499674 | 8.505051 |
| NaN | 6.425818 |
| 10.80164 | 10.76511 |
| 9.461094 | 9.246617 |
| 8.638093 | 8.763792 |
| 8.900475 | 8.9209 |
| 9.347105 | 9.342192 |
| 11.64329 | 11.51005 |
| 8.360871 | 8.545464 |
| 7.547194 | 7.217528 |
| 10.01794 | 9.545854 |
| 8.415398 | 8.766043 |
| 9.921706 | 9.824808 |
| 9.318369 | 9.839745 |
| 10.65172 | 10.42217 |
| 5.693908 | 5.911165 |
| 7.21218 | 6.894657 |
| 12.42903 | 12.52588 |
| NaN | 5.95885 |
| 9.518165 | 9.438346 |
| 7.14944 | 7.047695 |
| 8.292823 | 8.296593 |
| 7.549998 | 6.854678 |
| 9.846227 | 10.17593 |
| 6.985167 | 6.905652 |
| 6.136788 | 5.702189 |
| 8.482273 | 8.398604 |
| 9.685336 | 9.82136 |
| 10.63394 | 10.40046 |
| 11.7244 | 11.67924 |
| 8.235344 | 7.860815 |
| 7.286257 | 7.769122 |
| 7.252074 | 8.664731 |
| 13.20147 | 13.19636 |
| 6.007622 | 6.19399 |
| 7.848826 | 7.234335 |
| 8.655664 | 8.746298 |
| 9.610049 | 9.657105 |
| 14.05841 | 14.04652 |
| NaN | NaN |
| 9.092162 | 8.485207 |
| 13.70518 | 13.72338 |

| | |
|---|---|
| 5.726278 | 5.909338 |
| 14.19505 | 14.26341 |
| 9.565535 | 9.223727 |
| 10.15188 | 10.16335 |
| 13.53509 | 13.4569 |
| 5.538397 | 5.273751 |
| 11.58387 | 11.58717 |
| 6.881331 | 7.580355 |
| 9.123219 | 8.992602 |
| 8.027157 | 7.783661 |
| 7.287335 | 6.807962 |
| 7.167688 | 7.233984 |
| 7.212354 | 7.178932 |
| 9.803239 | 9.593775 |
| 8.886985 | 7.3803 |
| 6.992231 | 6.786422 |
| 9.766233 | 9.832356 |
| 6.798292 | 6.730597 |
| 7.93813 | 7.913131 |
| 5.101628 | 5.298974 |
| 8.941697 | 8.701317 |
| 7.406335 | 7.184839 |
| 5.145306 | 4.968419 |
| 11.31955 | 11.26819 |
| 7.698399 | 7.713106 |
| 9.772768 | 9.630919 |
| 11.11206 | 11.09699 |
| 5.355877 | 6.227098 |
| 7.149905 | 6.00359 |
| 5.230803 | 5.144866 |
| 4.998099 | 5.730677 |
| 7.902767 | 7.885508 |
| 6.048947 | 6.100268 |
| 9.720549 | 9.729489 |
| 7.715719 | 6.629436 |
| 9.266655 | 9.076259 |
| 15.2266 | 15.20428 |
| 7.0967 | 7.078353 |
| 6.821065 | 6.602693 |
| 8.599692 | 7.972013 |
| 7.441973 | 6.560713 |
| 6.188004 | 6.602759 |
| 6.875007 | 6.685251 |
| 6.733961 | 6.388168 |
| 7.073845 | 7.320309 |
| 6.871675 | 6.532583 |
| 6.904299 | 6.731263 |

| | |
|---|---|
| 7.253521 | 7.287222 |
| 7.446049 | 7.190016 |
| 6.633269 | 6.406267 |
| 7.03515 | 7.008961 |
| NaN | 6.140103 |
| 8.526754 | 8.201467 |
| 7.022811 | 6.966367 |
| 6.441365 | 5.969846 |
| 9.059064 | 7.831834 |
| 6.131562 | 5.903124 |
| 7.208955 | 6.333287 |
| 7.448768 | 7.668368 |
| 6.508753 | 6.508872 |
| 6.381763 | 4.643274 |
| 11.20031 | 11.14069 |
| 8.302204 | 8.932673 |
| 10.69625 | 10.14823 |
| 7.790305 | 6.512603 |
| 8.011042 | 8.274139 |
| 10.34024 | 10.25675 |
| 7.377891 | 7.545527 |
| 5.123557 | 4.722783 |
| 7.146152 | 7.088076 |
| 4.248574 | 3.610132 |
| 8.284101 | 7.859782 |
| 7.570405 | 7.391088 |
| 5.416349 | 5.337474 |
| 3.538758 | 3.627103 |
| 5.43926 | 5.531927 |
| 8.734118 | 8.756534 |
| 6.01927 | 5.705221 |
| 5.694709 | 5.448845 |
| 5.102119 | 5.732574 |
| 8.566351 | 8.442646 |
| 7.990737 | 7.831102 |
| 6.200966 | 6.498002 |
| 6.956865 | 7.226038 |
| 7.03379 | 6.850291 |
| 8.868687 | 8.745537 |
| 6.813255 | 7.038688 |
| 11.89646 | 11.88035 |
| 8.515376 | 8.475396 |
| 7.873 | 7.886919 |
| 8.074876 | 8.062099 |
| 8.656609 | 8.768116 |
| 7.329259 | 7.256256 |
| 5.72578 | 5.97495 |

| | |
|---|---|
| 7.112839 | 7.496672 |
| 5.034712 | 5.344238 |
| 12.70518 | 12.66763 |
| 6.715802 | 6.52584 |
| 5.115763 | 5.500279 |
| 8.700561 | 8.786748 |
| 5.536052 | 5.972554 |
| 7.562616 | 8.112053 |
| 11.75861 | 11.61073 |
| 8.179175 | 8.389376 |
| 9.810423 | 9.814809 |
| 8.161188 | 7.79128 |
| 12.36112 | 12.22618 |
| 9.986532 | 10.01847 |
| 6.762396 | 6.254728 |
| 5.607956 | 5.074339 |
| 8.270254 | 8.097843 |
| 6.430888 | 6.348271 |
| 7.671581 | 7.436541 |
| 9.026648 | 8.888525 |
| 10.70006 | 10.82736 |
| 6.633551 | 6.551041 |
| NaN | NaN |
| 9.268489 | 9.095057 |
| 7.580926 | 7.530759 |
| 8.74108 | 7.339295 |
| 4.823418 | 4.682526 |
| 6.642274 | 6.1556 |
| 8.946171 | 8.857201 |
| 5.472162 | 5.608646 |
| 5.174255 | 5.353584 |
| 9.864822 | 9.696388 |
| 5.495131 | 6.21844 |
| 7.057958 | 6.989893 |
| 7.660798 | 7.552413 |
| 8.102842 | 8.007275 |
| 5.814537 | 5.617124 |
| 6.198739 | 6.714108 |
| 4.547308 | 4.38506 |
| 6.936982 | 7.040715 |
| 5.751166 | 5.916356 |
| 5.100454 | 5.235137 |
| 9.89176 | 9.760272 |
| 6.084247 | 5.835423 |
| 7.211064 | 6.8726 |
| 8.660808 | 8.575946 |
| 5.921969 | 5.900741 |

| | |
|---|---|
| 6.186566 | 6.43006 |
| 6.990177 | 5.427081 |
| 9.693765 | 9.155663 |
| 6.291663 | 5.853873 |
| 6.848858 | 6.563653 |
| 10.3887 | 10.42906 |
| 7.599703 | 7.671454 |
| 10.20602 | 10.2739 |
| 10.2754 | 10.40868 |
| 8.820328 | 8.843405 |
| 8.653064 | 8.587023 |
| 9.291555 | 9.664266 |
| 7.120578 | 7.079768 |
| 7.072018 | 7.174959 |
| 6.411571 | 6.737824 |
| 6.885591 | 6.53902 |
| 8.046418 | 8.977841 |
| 5.984777 | 5.626024 |
| 12.169 | 12.10933 |
| 5.136058 | 6.815952 |
| 6.381604 | 6.503621 |
| 7.038735 | 7.104527 |
| 9.08903 | 9.338491 |
| 6.514959 | 6.928183 |
| 9.576575 | 9.194748 |
| 7.509803 | 7.23919 |
| 8.725498 | 8.882459 |
| 4.817062 | 4.561302 |
| NaN | 4.891428 |
| 7.998977 | 8.047027 |
| 6.604572 | 6.704066 |
| 10.25462 | 10.1775 |
| 11.34757 | 11.40059 |
| 11.63657 | 11.51711 |
| 7.172812 | 7.147042 |
| 7.93031 | 7.696804 |
| 7.747997 | 7.637672 |
| 4.825475 | 5.336629 |
| NaN | NaN |
| 7.493804 | 7.469193 |
| 9.093715 | 8.933838 |
| 8.192905 | 8.463679 |
| 9.650161 | 9.622738 |
| 7.546468 | 7.577964 |
| 6.314707 | 6.924214 |
| 5.544241 | 7.23398 |
| 7.825426 | 7.754864 |

| | |
|---|---|
| 8.32305 | 8.13042 |
| 7.359038 | 5.819135 |
| 7.134598 | 7.246679 |
| 8.554567 | 8.674785 |
| 7.903813 | 7.72727 |
| NaN | NaN |
| 9.69771 | 9.6419 |
| 8.121922 | 7.894193 |
| 6.806462 | 6.739464 |
| 7.602423 | 7.666363 |
| 6.429102 | 6.34607 |
| 4.658326 | 4.622032 |
| 6.083816 | 6.058203 |
| 7.789395 | 7.999666 |
| 11.0903 | 10.94087 |
| 8.683692 | 8.55807 |
| 10.76272 | 10.66217 |
| 12.05917 | 11.9712 |
| 9.000453 | 9.006733 |
| 5.628369 | 6.241183 |
| 7.804629 | 7.476762 |
| 8.415809 | 8.259564 |
| 9.177687 | 9.396815 |
| 9.157486 | 9.271143 |
| 6.946383 | 6.877132 |
| 7.694272 | 7.633224 |
| 10.99359 | 10.88346 |
| 8.809831 | 8.784893 |
| NaN | 6.059203 |
| 8.648476 | 8.453038 |
| 6.539417 | 6.478489 |
| 5.823516 | 6.117155 |
| 7.119946 | 6.969899 |
| 11.02656 | 11.1108 |
| NaN | 5.651186 |
| 7.741347 | 7.367256 |
| 6.146103 | 6.480963 |
| 7.318213 | 7.352252 |
| 8.36777 | 8.347384 |
| 8.962467 | 8.737426 |
| 8.216527 | 8.188996 |
| 5.73762 | 5.29861 |
| 7.00817 | 7.43304 |
| 6.798729 | 6.925367 |
| 6.835925 | 6.611584 |
| NaN | NaN |
| 8.789841 | 8.8655 |

| | |
|---|---|
| 7.406284 | 7.574318 |
| 9.458625 | 9.435268 |
| 4.027427 | 3.499433 |
| 7.809828 | 7.65137 |
| 5.842098 | 5.509689 |
| 7.333551 | 7.215927 |
| 8.49502 | 8.372541 |
| 7.838665 | 7.928873 |
| 7.431684 | 7.185388 |
| 8.50386 | 8.731702 |
| 8.837766 | 8.995029 |
| 5.462718 | 5.92215 |
| 6.683199 | 6.621964 |
| 6.623441 | 6.793127 |
| 6.740179 | 7.017563 |
| 5.235032 | 4.261833 |
| 7.027437 | 6.701938 |
| 9.146809 | 9.19406 |
| 8.474193 | 8.431715 |
| 8.793642 | 8.726327 |
| 8.030588 | 7.873064 |
| 9.756927 | 9.708256 |
| 7.435087 | 7.389071 |
| 8.866851 | 8.767609 |
| 6.069001 | 6.059785 |
| 7.673336 | 7.413398 |
| 7.751565 | 7.892208 |
| 7.49754 | 7.53315 |
| 7.219635 | 7.229069 |
| 8.642055 | 8.099469 |
| 6.529441 | 6.328014 |
| 8.249872 | 8.268172 |
| 8.054835 | 7.98017 |
| 6.89833 | 6.505223 |
| 7.005138 | 7.06137 |
| 7.780437 | 7.643709 |
| 7.175867 | 6.998284 |
| 6.297917 | 6.193053 |
| 9.117151 | 9.059125 |
| 6.271243 | 6.150582 |
| 8.368862 | 8.183742 |
| 7.014113 | 7.897682 |
| 10.1031 | 10.0434 |
| 9.149155 | 8.412248 |
| 10.89988 | 10.70171 |
| 10.54496 | 10.71479 |
| 9.808739 | 9.841589 |

| | |
|---|---|
| 6.660841 | 6.909348 |
| 8.042769 | 7.506692 |
| 6.673112 | 6.682998 |
| 8.786471 | 8.774279 |
| 7.650835 | 7.788719 |
| 8.261454 | 8.177992 |
| 7.032892 | 6.939454 |
| 4.242167 | 4.138932 |
| 6.740649 | 6.793388 |
| 7.657957 | 7.712089 |
| 5.973252 | 6.003209 |
| 4.394746 | 4.832286 |
| 8.613717 | 8.604277 |
| 7.33634 | 7.509986 |
| 8.472824 | 8.542248 |
| 9.649723 | 9.439567 |
| 6.972417 | 6.934727 |
| 8.645829 | 8.529091 |
| 8.125945 | 7.891002 |
| 7.848104 | 7.707036 |
| 8.09103 | 8.168152 |
| 5.477155 | 5.941202 |
| 8.69879 | 8.512317 |
| 7.45104 | 7.297074 |
| 10.18459 | 10.08018 |
| 7.978465 | 7.833573 |
| 5.657122 | 5.431724 |
| 5.128731 | 4.00202 |
| 5.141944 | 5.600181 |
| 7.575149 | 7.304204 |
| 7.844699 | 7.711977 |
| 7.572236 | 7.619113 |
| 7.311876 | 7.33297 |
| 5.662724 | 5.554878 |
| 5.693082 | 6.409051 |
| 10.56586 | 10.52985 |
| 11.38856 | 11.34958 |
| 7.687086 | 7.700407 |
| 9.216154 | 9.409284 |
| 4.237729 | 4.172489 |
| 9.422387 | 9.450101 |
| 7.265201 | 7.534206 |
| 8.042045 | 8.187015 |
| 8.517542 | 8.634899 |
| 5.349144 | 5.940043 |
| 6.465853 | 6.498224 |
| 6.635683 | 6.727387 |

-Bruker Confidential-

| | |
|---|---|
| 9.462334 | 9.33569 |
| 8.583565 | 8.430818 |
| 5.701611 | 5.860609 |
| 7.805223 | 7.465197 |
| 7.560059 | 7.349347 |
| 8.484694 | 7.981595 |
| 10.13826 | 9.776721 |
| 7.718256 | 7.993779 |
| 9.20067 | 9.720806 |
| 7.231884 | 7.466616 |
| 11.13972 | 11.02267 |
| 8.975923 | 8.70904 |
| 5.612428 | 4.842062 |
| 7.321668 | 7.412259 |
| 8.130637 | 8.113792 |
| 6.194033 | 5.816624 |
| 6.625895 | 6.963821 |
| 5.940221 | 6.208093 |
| 7.899678 | 8.106853 |
| 7.482409 | 7.239061 |
| 9.184312 | 9.042456 |
| 5.792699 | 5.941317 |
| 7.706209 | 7.569444 |
| 8.860728 | 8.935879 |
| 7.389238 | 7.314012 |
| 5.314274 | 5.180632 |
| 8.492482 | 8.729047 |
| 8.560752 | 8.388312 |
| 8.694155 | 8.472078 |
| 10.00502 | 10.0164 |
| 6.077139 | 5.949213 |
| 8.141187 | 7.885395 |
| 9.29691 | 9.221148 |
| 5.232831 | 3.961398 |
| 6.136702 | 6.566451 |
| 5.838199 | 5.216138 |
| 6.048446 | 5.80446 |
| 6.673015 | 6.444696 |
| 6.757074 | 6.620656 |
| 9.059738 | 8.812068 |
| 7.820655 | 7.808749 |
| 6.845411 | 6.790069 |
| 6.569858 | 6.282303 |
| 7.490704 | 7.089503 |
| 7.814032 | 8.062955 |
| 6.700127 | 6.73921 |
| 7.36543 | 7.07629 |

| | |
|---|---|
| 6.535818 | 6.6899 |
| 6.167968 | 5.62377 |
| 8.246747 | 7.989206 |
| 7.136419 | 7.303965 |
| 5.846653 | 5.925146 |
| 8.05665 | 7.80233 |
| 8.270069 | 8.615498 |
| 11.54015 | 11.44107 |
| 11.42286 | 11.47626 |
| 3.599949 | 3.378194 |
| 8.374213 | 8.199109 |
| 8.64056 | 8.732345 |
| 8.232636 | 8.37842 |
| NaN | NaN |
| 6.471847 | 6.24162 |
| 9.67768 | 9.786168 |
| 8.188002 | 8.441054 |
| 7.940236 | 7.174905 |
| 9.85879 | 10.07731 |
| 10.52162 | 10.48303 |
| 7.917132 | 7.82787 |
| 6.224453 | 6.123066 |
| 8.34886 | 8.210673 |
| 6.242109 | 6.23442 |
| 6.63673 | 6.519479 |
| 7.920239 | 7.435288 |
| 9.808976 | 9.691523 |
| 8.398744 | 8.46221 |
| 5.526001 | 5.187067 |
| 6.279521 | 6.032856 |
| 6.852777 | 6.665785 |
| 6.733881 | 6.876009 |
| 6.189927 | 5.704279 |
| 6.899999 | 6.597444 |
| 7.516945 | 7.641151 |
| 8.915622 | 9.024058 |
| 7.010076 | 6.710296 |
| 8.251231 | 8.09655 |
| 5.925788 | 5.977626 |
| 7.501566 | 7.75645 |
| 8.178276 | 7.337658 |
| 6.897554 | 6.670915 |
| 7.277478 | 7.30475 |
| 8.772817 | 8.514475 |
| 9.863589 | 9.874114 |
| 6.003337 | 5.999904 |
| 6.362379 | 6.167426 |

| | |
|---|---|
| 7.061718 | 6.96447 |
| 10.03557 | 9.791047 |
| 8.015211 | 7.851484 |
| 6.698339 | 6.252414 |
| 6.504779 | 6.68457 |
| 9.954792 | 9.939895 |
| 5.806247 | 6.031704 |
| 7.311601 | 7.349068 |
| 5.536824 | 5.892132 |
| 11.76523 | 11.68002 |
| 8.953808 | 8.811341 |
| 7.260866 | 7.386229 |
| 5.03951 | 3.727554 |
| 6.06237 | 5.576435 |
| 13.12179 | 13.71649 |
| 6.879096 | 6.522131 |
| 8.322051 | 8.383044 |
| 9.156739 | 8.921937 |
| 6.785732 | 6.092366 |
| 7.536968 | 7.747766 |
| 6.659267 | 6.279484 |
| 6.801786 | 6.145733 |
| 9.61739 | 9.452127 |
| 10.15101 | 10.31896 |
| 6.427581 | 6.448789 |
| 8.118054 | 7.980527 |
| 8.116501 | 7.973352 |
| 6.606943 | 6.784517 |
| 10.81435 | 10.55629 |
| 10.1137 | 9.976544 |
| 6.023019 | 6.096268 |
| 6.570413 | 6.579947 |
| 8.428355 | 8.279182 |
| 6.66744 | 6.361204 |
| 4.883216 | 5.156608 |
| 7.595056 | 7.405926 |
| 7.109154 | 6.90925 |
| 7.800834 | 7.947021 |
| 5.958361 | 5.9038 |
| 7.762187 | 7.755566 |
| 9.720009 | 9.632096 |
| 7.644472 | 7.830303 |
| 8.271144 | 8.157188 |
| 8.593986 | 8.751346 |
| 6.380165 | 6.229289 |
| 9.677917 | 9.667474 |
| 8.467259 | 8.004545 |

-Bruker Confidential-

| | |
|---|---|
| 7.553846 | 7.502538 |
| 5.638042 | 5.71586 |
| 6.529776 | 6.347426 |
| 4.72037 | 4.292453 |
| 4.808815 | 4.816535 |
| 5.027362 | 5.259808 |
| 11.90635 | 11.91915 |
| 12.10246 | 11.77515 |
| 7.516112 | 7.85089 |
| 6.237798 | 5.537542 |
| 10.74878 | 10.73877 |
| 12.16107 | 12.15628 |
| 7.785154 | 7.544384 |
| 8.082604 | 8.191402 |
| 6.61649 | 6.62187 |
| 6.954074 | 5.967414 |
| 9.405399 | 9.405147 |
| 7.109684 | 7.380167 |
| 11.34748 | 11.37946 |
| 7.195897 | 7.05817 |
| 10.31462 | 10.27962 |
| 8.148142 | 7.776839 |
| 8.737365 | 8.358987 |
| 7.785366 | 7.326781 |
| 8.297527 | 8.426 |
| 7.116516 | 6.870712 |
| 8.18462 | 8.174474 |
| 4.75807 | 4.525223 |
| 10.68486 | 10.45187 |
| 6.403507 | 6.457043 |
| 9.19878 | 9.356989 |
| 6.205004 | 6.632649 |
| 7.584625 | 7.703066 |
| 6.431894 | 6.462612 |
| 10.22832 | NaN |
| 6.349498 | 7.216938 |
| 7.155811 | 7.469886 |
| 7.601688 | 7.355675 |
| 6.001239 | 5.893189 |
| 6.878396 | 7.215428 |
| 8.206879 | 8.54628 |
| 6.916528 | 6.871707 |
| 5.825789 | NaN |
| 8.00052 | 7.643445 |
| 7.31034 | 7.310571 |
| 6.64275 | 6.188383 |
| 6.159015 | 6.431397 |

| | |
|---|---|
| 4.239614 | NaN |
| 7.809809 | 7.537487 |
| 5.909239 | 5.74547 |
| 4.415916 | 4.554732 |
| 8.024523 | 8.199142 |
| 9.269415 | 9.248005 |
| 7.207701 | 7.160853 |
| 8.882142 | 8.723414 |
| 2.30382 | 3.949584 |
| 5.43257 | 5.269942 |
| 8.898132 | 8.896596 |
| 6.192674 | 6.319093 |
| 5.960724 | 6.088074 |
| 7.115904 | 7.045329 |
| 7.452487 | 7.424259 |
| 6.211846 | 5.962009 |
| 6.336301 | 6.440083 |
| 7.126261 | 6.973467 |
| 9.466157 | 9.835485 |
| 10.93257 | 11.04802 |
| 6.158341 | 6.349809 |
| 6.689117 | 6.77 |
| 8.675937 | 8.437134 |
| 6.987929 | 7.000067 |
| 6.886203 | 6.932154 |
| 7.840803 | 7.494277 |
| 7.88379 | 7.710744 |
| 6.332056 | 6.356087 |
| NaN | NaN |
| 9.638742 | 9.418822 |
| 8.178021 | 8.265763 |
| 4.614457 | 5.322854 |
| 7.635402 | 7.920783 |
| 7.91839 | 8.124133 |
| 5.737324 | 5.810691 |
| 8.274395 | 8.445409 |
| 11.11224 | 11.38764 |
| 5.746563 | 6.121192 |
| 8.752241 | 8.975594 |
| 4.540555 | 5.639308 |
| 5.861958 | 5.648489 |
| 6.308887 | 6.69116 |
| 8.036701 | 7.98461 |
| 5.909704 | 5.792829 |
| 6.437795 | 6.543532 |
| NaN | NaN |
| 6.34882 | 6.840118 |

| | |
|---|---|
| 8.179296 | 8.313803 |
| 8.407809 | 8.315607 |
| 7.769419 | 7.802732 |
| 9.15502 | 9.28695 |
| 6.215287 | 6.155415 |
| 6.203681 | 6.30234 |
| 7.200586 | 6.756505 |
| 10.32234 | 10.24911 |
| 7.90466 | 8.156822 |
| 6.064433 | 6.112725 |
| 8.046494 | 6.82167 |
| 8.946386 | 8.906497 |
| 8.61311 | 8.293052 |
| 7.511776 | 7.435287 |
| 8.293863 | 8.484786 |
| 7.296953 | 7.167897 |
| 6.691502 | NaN |
| 8.359197 | 8.355592 |
| 7.84217 | 7.754127 |
| 6.820607 | 7.125012 |
| 7.398732 | 7.396364 |
| 5.637858 | 5.342986 |
| 6.557654 | 5.992806 |
| 7.872351 | 7.972066 |
| 7.73006 | 7.726015 |
| 5.37262 | 5.710121 |
| 6.518297 | 6.318418 |
| 8.658505 | 8.325243 |
| 7.448364 | 7.335986 |
| 11.04834 | 10.89454 |
| 8.519083 | 8.184047 |
| 5.85073 | 5.976735 |
| 7.09138 | 7.133513 |
| 7.004425 | 7.360952 |
| 10.09564 | 10.02273 |
| 8.566839 | 8.313807 |
| 8.796063 | 8.560339 |
| 5.052958 | 4.554949 |
| 6.813927 | 6.827106 |
| 9.417132 | 9.461185 |
| 8.976636 | 8.90113 |
| 8.159894 | 8.155784 |
| 7.534492 | 7.943205 |
| 6.352714 | 5.832864 |
| 4.291883 | 4.176061 |
| 6.195548 | 6.432583 |
| 10.04798 | 9.984289 |

| | |
|---|---|
| 9.893227 | 10.05029 |
| 8.036777 | 8.027692 |
| 6.332509 | 6.024338 |
| 12.03366 | 12.07491 |
| 8.28611 | 8.505067 |
| 7.255929 | 7.380473 |
| 6.995118 | 6.970006 |
| 8.249533 | 8.007685 |
| 3.993423 | 4.863096 |
| 7.951 | 7.778347 |
| 9.158131 | 8.819562 |
| 7.987365 | 8.167887 |
| 13.14911 | 13.03191 |
| 6.817221 | 7.109955 |
| 4.938717 | 5.337061 |
| 8.091022 | 7.721577 |
| NaN | 5.116583 |
| 7.151474 | 7.298374 |
| 10.70209 | 10.50632 |
| 11.79509 | 11.98842 |
| 7.992822 | 7.783079 |
| 10.69107 | 10.50768 |
| 6.408769 | 5.399173 |
| 8.458107 | 8.745611 |
| 6.287181 | 6.082915 |
| 7.764522 | 8.072521 |
| 7.677054 | 7.601453 |
| 8.643083 | 8.849771 |
| 6.435375 | 6.783275 |
| 10.17615 | 10.04522 |
| 8.118745 | 8.33555 |
| 8.432234 | 8.272976 |
| 6.372518 | 6.719738 |
| 6.580471 | 6.696253 |
| 8.478007 | 8.3077 |
| 9.733934 | 9.631877 |
| 9.084554 | 9.009369 |
| 8.672264 | 8.544823 |
| 8.868682 | 8.693087 |
| NaN | NaN |
| 7.528296 | 7.686225 |
| 7.448102 | 7.392646 |
| 6.865108 | 7.035679 |
| 5.856509 | 5.668206 |
| 7.114866 | 7.162882 |
| 5.762394 | 5.61679 |
| 5.152774 | 5.083558 |

| | |
|---|---|
| 7.290092 | 7.407948 |
| 7.588603 | 7.448978 |
| 7.509214 | 7.475586 |
| 6.984905 | 7.204848 |
| 6.316663 | 6.556313 |
| 5.104305 | 5.61878 |
| 6.054007 | 5.822396 |
| 9.619275 | 9.451638 |
| 7.384719 | 7.067026 |
| 8.249801 | 8.334259 |
| 5.957513 | 5.859505 |
| 6.66225 | 6.307056 |
| 6.902824 | 6.703383 |
| 8.779147 | 8.676094 |
| 7.071423 | 7.027296 |
| 6.282183 | 7.713964 |
| 7.35088 | 7.00194 |
| 6.083204 | 5.436079 |
| 9.838867 | 9.386079 |
| 5.667808 | 5.611822 |
| 7.758779 | 7.147999 |
| 10.27649 | 10.01607 |
| 8.335047 | 8.862742 |
| 7.84324 | 6.945151 |
| 8.794284 | 7.647871 |
| 8.072979 | 7.88164 |
| 7.588236 | 7.186474 |
| 7.426506 | 7.441875 |
| 7.019367 | 7.110865 |
| 4.962512 | 5.229792 |
| 8.200568 | 7.978117 |
| 6.187405 | 6.37998 |
| NaN | NaN |
| 7.769068 | 7.338928 |
| 6.896715 | 6.649796 |
| 7.850996 | 7.589756 |
| 9.287263 | 9.044367 |
| 7.073012 | 6.990481 |
| 6.521976 | 6.420807 |
| 9.290942 | 9.005097 |
| NaN | NaN |
| 6.261652 | 6.300488 |
| NaN | 3.607911 |
| 8.077643 | 8.257688 |
| 7.33451 | 7.19826 |
| 8.141173 | 8.060536 |
| 10.69253 | 10.54213 |

| | |
|---|---|
| 4.247257 | 5.350936 |
| 5.922265 | 6.228763 |
| 6.552978 | 6.864775 |
| 7.159697 | 7.621497 |
| 7.246907 | 6.743492 |
| 10.92611 | 11.11566 |
| 8.215005 | 8.384243 |
| 8.918932 | 8.811011 |
| 7.453997 | 7.313087 |
| 6.472314 | 6.583296 |
| 12.15886 | 12.20673 |
| 9.234444 | 9.199981 |
| 6.120548 | NaN |
| 8.18222 | 8.068723 |
| 6.667187 | 6.66557 |
| 8.353401 | 8.185153 |
| 5.908617 | 5.832006 |
| 7.232192 | 7.220115 |
| 8.379318 | 8.327027 |
| 7.46873 | 7.350722 |
| 6.744834 | 6.527316 |
| 7.468043 | 7.605718 |
| 8.754498 | 8.843533 |
| 12.12176 | 12.07878 |
| 4.813056 | 5.276006 |
| 7.670076 | 7.791745 |
| 6.400496 | 6.731606 |
| 7.111437 | 7.021663 |
| NaN | 5.816326 |
| 6.595672 | 6.404345 |
| 6.928913 | 6.807111 |
| 5.808256 | 5.996268 |
| 5.405179 | 5.463985 |
| 10.05743 | 9.984776 |
| 8.183523 | 8.022715 |
| 6.404957 | 6.670263 |
| 11.96358 | 12.00741 |
| NaN | 5.602201 |
| 10.88779 | 10.79771 |
| 4.967244 | 5.696203 |
| 9.693513 | 9.833918 |
| 9.86549 | 10.08213 |
| 6.842457 | 6.68375 |
| 7.613267 | 7.63184 |
| 7.565362 | 7.45892 |
| 7.181606 | 7.352912 |
| 5.716324 | 6.00947 |

| | |
|---|---|
| 9.520075 | 9.43103 |
| 8.824995 | 8.756233 |
| 7.016492 | 5.737006 |
| 9.711406 | 9.757027 |
| 8.020804 | 8.351057 |
| 7.779575 | 7.957623 |
| 5.558247 | 5.331229 |
| 7.651896 | 7.447755 |
| NaN | 4.717362 |
| 4.931585 | 4.933637 |
| 6.822451 | 7.101132 |
| 7.449317 | 7.462209 |
| 7.513126 | 7.13956 |
| 9.187108 | 9.239601 |
| 7.67157 | 8.800092 |
| 8.484818 | 8.417235 |
| 7.939223 | 7.466882 |
| 10.05071 | 10.06124 |
| 4.833587 | 5.067165 |
| 4.641047 | NaN |
| 5.688925 | 5.9319 |
| 7.987964 | 8.415386 |
| 10.42779 | 10.47795 |
| 5.131574 | 5.890277 |
| 5.968133 | 6.226415 |
| 5.528099 | 4.766774 |
| 7.496181 | 7.434705 |
| 6.245008 | 5.786033 |
| 5.725889 | 5.233623 |
| 7.614077 | 7.508398 |
| 5.011645 | 4.669056 |
| 8.151978 | 8.216666 |
| 5.146119 | 4.822931 |
| 6.026145 | 5.732251 |
| 4.405497 | 4.249859 |
| 6.831789 | 6.977225 |
| 7.421864 | 7.535708 |
| 6.640147 | 6.557822 |
| 3.610171 | 4.070096 |
| 6.665189 | 6.31235 |
| 9.096378 | 9.167293 |
| 5.585521 | 5.280396 |
| 3.3331 | NaN |
| 5.786605 | 5.967876 |
| 4.846809 | 4.775407 |
| 7.577094 | 7.782332 |
| 7.456819 | 7.307188 |

| | |
|---|---|
| 5.456654 | 5.614111 |
| 7.724842 | 8.056644 |
| 6.287953 | 6.900331 |
| 6.634192 | 6.236813 |
| 5.759598 | 6.415553 |
| 4.364402 | 4.382767 |
| 8.435053 | 8.679795 |
| NaN | 6.223394 |
| 8.769534 | 8.822525 |
| 8.692815 | 7.544984 |
| 9.129141 | 9.105137 |
| NaN | NaN |
| 7.033152 | 6.913037 |
| 7.500902 | 7.690795 |
| 7.190256 | 7.197764 |
| 8.728792 | 9.156126 |
| 7.699079 | 7.522131 |
| 7.067797 | 7.131921 |
| 6.203032 | 5.886312 |
| 7.613614 | 6.736882 |
| 7.166436 | 7.229222 |
| 5.186814 | 4.267104 |
| 6.03371 | 5.805454 |
| 8.057769 | 8.03666 |
| 4.314185 | 5.041854 |
| 6.66491 | 6.765812 |
| 7.944435 | 8.116097 |
| 6.853518 | 6.835635 |
| 5.046258 | 5.003305 |
| 5.350509 | 6.116921 |
| 8.126377 | 7.802473 |
| 6.911938 | 6.965712 |
| 6.243906 | 6.241005 |
| 5.683213 | 5.796166 |
| 8.136636 | 8.056342 |
| 5.158571 | 4.187118 |
| 6.193588 | 6.565718 |
| 5.640639 | 6.505667 |
| 6.185922 | 6.154295 |
| 6.706805 | 5.885631 |
| 8.988196 | 9.206254 |
| NaN | NaN |
| 7.892718 | 7.535522 |
| NaN | NaN |
| 5.357929 | 4.899031 |
| 7.257479 | 6.622571 |
| 8.467325 | 8.694754 |

| | |
|---|---|
| 9.307073 | 8.968085 |
| 7.137906 | 6.917649 |
| 6.206565 | 6.272374 |
| 7.945592 | 7.9367 |
| 5.673446 | 5.957808 |
| 7.581492 | 7.895589 |
| 6.883086 | 6.86446 |
| 6.983148 | 6.622889 |
| 6.20365 | 6.106811 |
| 7.70672 | 7.683913 |
| 7.799918 | 7.330013 |
| 6.985079 | 6.862052 |
| 7.093651 | 7.26505 |
| 5.620848 | 6.198682 |
| 7.413732 | 7.118859 |
| 5.424092 | NaN |
| 6.24374 | 6.070217 |
| 8.848613 | 8.694661 |
| 4.921624 | 5.589626 |
| 7.147329 | 6.349349 |
| 6.599814 | 6.139731 |
| 5.308125 | 3.592477 |
| 12.47792 | 12.46833 |
| 7.279933 | 7.43735 |
| 6.466151 | 6.072899 |
| 6.129268 | NaN |
| 7.017841 | 6.87492 |
| 9.514106 | 9.087956 |
| 9.558614 | 9.521671 |
| 12.00273 | 14.12934 |
| 9.063275 | 9.06365 |
| 5.936263 | 5.83523 |
| 5.356218 | 4.478555 |
| 8.21146 | 8.359071 |
| 5.746078 | 5.593329 |
| 6.433665 | 5.28541 |
| 6.587822 | 5.414167 |
| 6.687182 | 5.894139 |
| 6.413311 | 6.700358 |
| 8.254995 | 8.224638 |
| 5.973024 | 5.869371 |
| 7.392229 | 7.238844 |
| 6.821819 | 6.91724 |
| 8.304049 | 8.11992 |
| 5.684137 | 5.553578 |
| 5.112526 | 5.550364 |
| 6.502758 | 6.594444 |

| | |
|---|---|
| NaN | NaN |
| 7.433579 | 7.193042 |
| 6.322564 | 6.130501 |
| 7.577549 | 7.48785 |
| 7.216766 | 7.252909 |
| 5.829181 | 5.774159 |
| 6.892295 | 6.889698 |
| 7.338509 | 7.435303 |
| 5.634884 | 5.526731 |
| 4.24725 | 4.755632 |
| 5.990183 | 5.781936 |
| 8.721129 | 8.771776 |
| 8.2546 | 7.549949 |
| 4.982257 | 5.434972 |
| 5.566879 | 5.112409 |
| 7.897251 | 7.894902 |
| 5.687963 | 5.735125 |
| 6.104577 | 6.587373 |
| 7.262602 | 7.010241 |
| 7.624234 | 7.494097 |
| 6.898185 | 6.651558 |
| 6.42396 | 6.146928 |
| NaN | NaN |
| 5.25199 | 5.381396 |
| 5.159357 | 5.057104 |
| 6.999279 | 5.05078 |
| 6.333619 | 6.663435 |
| 7.970002 | 7.68207 |
| 4.418861 | 4.778005 |
| 4.444661 | 6.879628 |
| 5.119741 | 5.289792 |
| 6.669879 | 6.701404 |
| 7.226591 | 7.020355 |
| 6.513233 | 6.540258 |
| 5.368586 | 5.650143 |
| 6.011873 | 5.904863 |
| NaN | NaN |
| 7.063785 | 7.131546 |
| 6.686706 | 6.34949 |
| 7.403116 | 7.612373 |
| 7.791231 | 7.488214 |
| 7.493366 | 7.871731 |
| 2.939897 | 3.23215 |
| 10.54482 | 10.56541 |
| 6.762126 | 6.732597 |
| 9.220808 | 9.212487 |
| 6.086297 | 6.124758 |

| | |
|---|---|
| 3.761279 | 4.308908 |
| 10.15816 | 10.16683 |
| 3.960939 | 3.081682 |
| 6.621999 | 6.363571 |
| 5.961359 | 6.230451 |
| 6.012853 | 5.827957 |
| 6.632856 | 5.440145 |
| NaN | NaN |
| 7.452192 | 7.708661 |
| 7.436143 | 7.431742 |
| 3.523734 | NaN |
| 5.454786 | 5.408901 |
| 7.007185 | 6.996815 |
| 8.35612 | 8.454367 |
| 5.536754 | 5.801069 |
| 5.78561 | 5.784482 |
| 8.137847 | 8.264561 |
| 6.928113 | 6.503201 |
| 6.431543 | 6.528789 |
| 6.004487 | 5.709202 |
| 6.091264 | 6.177123 |
| 6.514021 | 5.667622 |
| 6.656525 | 6.671005 |
| 6.503502 | 6.817325 |
| 5.995621 | 6.06229 |
| 5.59847 | 6.093886 |
| 5.119513 | 5.117565 |
| 8.398191 | 8.48086 |
| 7.663807 | 7.482959 |
| 7.677049 | 7.686523 |
| 6.568057 | 6.040763 |
| 7.221318 | 7.411331 |
| 6.265285 | 5.761698 |
| 7.287324 | 7.184265 |
| 7.058963 | 6.428818 |
| 5.646486 | 5.912624 |
| NaN | 5.121361 |
| NaN | 6.89841 |
| 6.674013 | 6.772871 |
| 7.686481 | 7.665646 |
| 9.74045 | 9.840678 |
| 5.277039 | 5.0108 |
| 6.942511 | 7.554813 |
| 7.587035 | 7.598391 |
| 7.112392 | 6.726184 |
| 6.125696 | 6.111669 |
| 6.613205 | 6.751786 |

| | |
|---|---|
| 5.63562 | 5.612156 |
| 5.537553 | 5.723955 |
| 6.307536 | 5.82604 |
| 4.900743 | 4.560457 |
| 6.359304 | 6.105208 |
| 8.94168 | 8.613518 |
| 8.797867 | 8.884459 |
| 4.966394 | 5.372516 |
| 6.471249 | 6.067075 |
| 3.696172 | NaN |
| 6.504183 | 6.329219 |
| 6.090979 | 6.203967 |
| 7.76425 | 7.891578 |
| 6.309422 | 6.44699 |
| 5.446559 | 6.365042 |
| 7.887253 | 7.902324 |
| 6.276671 | 7.343487 |
| 6.989939 | 6.976074 |
| 5.220852 | 4.959152 |
| 5.910637 | 5.347569 |
| 6.403805 | 6.039698 |
| 7.025676 | 6.90391 |
| 5.499073 | 6.112429 |
| 8.079977 | 8.045898 |
| 8.878165 | 9.029273 |
| 7.656957 | 7.402547 |
| 6.915697 | 7.066541 |
| 7.124233 | 6.781278 |
| 6.367176 | 5.976636 |
| 7.850411 | 7.321494 |
| NaN | NaN |
| 6.145974 | 6.118192 |
| 5.846886 | 5.557801 |
| 7.40584 | 7.569026 |
| 5.938445 | 5.739813 |
| 8.227828 | 7.453645 |
| 6.783141 | 6.685394 |
| 2.698122 | 2.404503 |
| 6.637299 | 6.474783 |
| 5.983897 | 6.137813 |
| 6.306742 | 6.229292 |
| 6.708311 | 7.00468 |
| 7.53837 | 7.426682 |
| 5.96045 | 6.05578 |
| NaN | NaN |
| NaN | 5.161388 |
| 8.320219 | 7.783965 |

| | |
|---|---|
| NaN | 6.560322 |
| 6.615589 | 6.61126 |
| 6.596861 | 6.657196 |
| 6.277933 | 6.782285 |
| 5.37468 | 5.698688 |
| NaN | NaN |
| 6.382719 | 6.120939 |
| 7.0145 | 7.053957 |
| 8.201869 | 8.493305 |
| 6.496369 | 6.171865 |
| 6.254337 | 6.159002 |
| 5.924618 | 7.285206 |
| 6.279288 | 6.457458 |
| 5.812881 | 5.397345 |
| 4.373107 | 4.117781 |
| 6.463345 | 6.832726 |
| 7.94148 | 8.016673 |
| 7.339547 | 7.035983 |
| 7.152043 | 7.652027 |
| 6.647808 | 6.291378 |
| 7.159012 | 7.130498 |
| 6.06431 | 6.495986 |
| 5.608775 | 5.513674 |
| 5.299257 | 6.313078 |
| 6.874015 | 7.32429 |
| 7.206499 | 7.458601 |
| 3.589795 | 5.537156 |
| 4.652636 | NaN |
| 5.258114 | 4.854521 |
| 7.309746 | 7.482348 |
| 4.198716 | 3.864619 |
| NaN | 7.044333 |
| 6.858411 | 6.697404 |
| 7.204895 | 7.460801 |
| 6.532584 | 6.257168 |
| 10.42647 | 10.40019 |
| 6.83059 | 6.780172 |
| 3.813237 | 4.770268 |
| 6.8759 | 6.646169 |
| 8.5958 | 8.369678 |
| 6.499808 | 6.850748 |
| 7.138058 | 6.839997 |
| 7.103361 | 6.187551 |
| 9.06234 | 8.709049 |
| 9.199706 | 9.006611 |
| NaN | 2.960948 |
| 6.241862 | 5.707709 |

-Bruker Confidential-

| | |
|---|---|
| 8.580062 | 8.537396 |
| 6.759501 | 6.494076 |
| 7.285219 | 7.453134 |
| 4.595183 | 4.07021 |
| 4.994834 | 5.438296 |
| 6.326781 | 6.203722 |
| 7.678368 | 7.833947 |
| 8.200591 | 7.981593 |
| 7.649951 | 7.903337 |
| 7.831752 | 7.860736 |
| 6.662062 | 6.633461 |
| 6.553215 | 6.318303 |
| NaN | 4.527673 |
| 3.414148 | 3.708135 |
| 6.693673 | 6.939457 |
| NaN | NaN |
| 6.80646 | 6.648609 |
| 6.031404 | 5.963537 |
| 6.389232 | 6.304101 |
| 5.906303 | 6.155859 |
| 5.278291 | 5.435455 |
| 5.569979 | 5.775019 |
| 6.143404 | 6.108189 |
| 6.660585 | 6.346882 |
| 7.65336 | 8.047505 |
| 5.931184 | 5.986415 |
| 5.772123 | 4.790101 |
| 6.200921 | 5.709528 |
| 6.590008 | 6.021782 |
| 8.664313 | 8.293252 |
| 7.743903 | 7.782921 |
| 7.833569 | 7.177816 |
| 6.12693 | 6.184117 |
| 5.766397 | 5.964249 |
| 7.778569 | 7.724096 |
| 6.084183 | 6.504436 |
| 6.782541 | 6.192225 |
| 5.112658 | 5.295272 |
| 6.74817 | 6.304716 |
| 6.69125 | 6.740024 |
| 6.566467 | 6.514243 |
| 6.462815 | 5.686155 |
| 5.775172 | 5.745992 |
| 6.220468 | 6.365716 |
| 7.194511 | 7.421298 |
| 6.991918 | 6.786678 |
| 7.591098 | 7.102025 |

-Bruker Confidential-

| | |
|---|---|
| 8.999715 | 9.134881 |
| 6.149753 | 6.097366 |
| 6.271939 | 5.115393 |
| 5.626623 | 5.797748 |
| 6.908688 | 6.666796 |
| 5.345625 | 5.519477 |
| 6.82023 | 6.749861 |
| 7.172743 | 7.37224 |
| 6.420231 | NaN |
| 6.189665 | 5.812313 |
| 7.444715 | 7.337909 |
| 6.157515 | 6.758607 |
| 9.137488 | 9.071341 |
| 6.431739 | 6.397323 |
| 8.20505 | 8.12107 |
| 5.942947 | 6.004547 |
| 7.044569 | 6.233534 |
| 7.159311 | 6.570074 |
| NaN | NaN |
| 5.512278 | 5.088368 |
| 10.3023 | 9.969224 |
| 4.808122 | 4.524657 |
| 5.405176 | 6.200926 |
| 7.178596 | 7.407022 |
| 5.829685 | 5.895357 |
| 6.700165 | 6.371886 |
| 6.787487 | 7.733921 |
| 7.434875 | 7.152057 |
| 6.05083 | 5.670686 |
| 6.814795 | 6.858526 |
| 8.76201 | 8.900935 |
| 7.662444 | 7.212576 |
| 7.213246 | 7.342165 |
| 7.151676 | 7.455016 |
| 5.930217 | 5.952038 |
| 7.029729 | 6.957922 |
| 7.253342 | 7.08889 |
| 5.137496 | 4.550539 |
| 7.673151 | 7.329656 |
| 9.576012 | 9.225227 |
| 4.99343 | 4.917284 |
| 4.899259 | 4.800211 |
| 5.583432 | 5.881673 |
| 5.139696 | 5.338288 |
| 6.377406 | 6.729119 |
| 6.972077 | 7.166149 |
| 8.490778 | 8.178238 |

| | |
|---|---|
| 5.787915 | 6.007355 |
| 7.781902 | 7.687986 |
| 5.325733 | 5.401413 |
| 8.063787 | 7.483679 |
| 6.294383 | 6.449996 |
| 5.900585 | 5.867487 |
| 6.432428 | 5.713939 |
| 6.739839 | 7.303642 |
| 4.283746 | 3.84588 |
| 6.889882 | 6.503749 |
| 4.874596 | 8.429603 |
| 5.980981 | 6.357116 |
| 7.755074 | 8.02462 |
| 8.593214 | 8.268573 |
| 8.291199 | 8.14178 |
| 6.027297 | 5.927982 |
| 7.472943 | 7.477347 |
| 5.327785 | 5.339122 |
| 8.373496 | 8.461299 |
| 7.543877 | 7.206412 |
| 7.180304 | 7.545513 |
| 5.549283 | 4.941452 |
| 6.074734 | 5.928408 |
| 8.284425 | 8.335454 |
| 5.7569 | 5.712171 |
| 6.925998 | 7.009527 |
| 7.342826 | 8.138863 |
| 7.238302 | 7.325949 |
| 5.911993 | 5.772544 |
| 6.445241 | 5.839272 |
| 6.767427 | 6.802171 |
| 6.128114 | 5.970802 |
| NaN | NaN |
| 7.794697 | 7.799296 |
| 8.55234 | 8.337279 |
| 5.379041 | 5.428775 |
| 3.092341 | 1.226928 |
| 7.707749 | 7.711493 |
| 5.988799 | 6.017147 |
| 4.964583 | 5.103199 |
| 8.309926 | 8.355771 |
| 6.853041 | 6.641044 |
| 6.033869 | 5.648485 |
| 5.461836 | 5.910196 |
| 6.909687 | 7.321257 |
| 7.921699 | 7.84423 |
| 6.881422 | 6.749104 |

-Bruker Confidential-

| | |
|---|---|
| 9.129055 | 9.069334 |
| NaN | NaN |
| 7.505484 | 7.242215 |
| 7.531742 | 7.357279 |
| 7.825623 | 8.353773 |
| 8.163773 | 7.975934 |
| 7.372097 | 7.166761 |
| 5.266255 | 4.58789 |
| 7.594631 | 7.794577 |
| 7.259071 | 7.242992 |
| 6.026271 | 5.700415 |
| 7.536364 | 7.196757 |
| 7.048627 | 6.80955 |
| 5.299738 | 5.712685 |
| 6.651265 | 6.310886 |
| 7.821022 | 7.65392 |
| 7.750744 | 7.891008 |
| 5.514112 | 6.466121 |
| 5.286152 | 5.712438 |
| 5.885029 | 6.09546 |
| 7.154572 | 7.057134 |
| 6.081843 | 5.556802 |
| 6.64979 | 6.483017 |
| 6.250308 | 6.155702 |
| 5.148389 | 5.335958 |
| 6.394342 | 6.352781 |
| 5.570092 | 5.566388 |
| 6.024003 | 5.613901 |
| 6.453782 | NaN |
| 7.083187 | 6.871094 |
| 6.850089 | NaN |
| 5.283593 | 4.743154 |
| 6.814461 | 7.407148 |
| 8.315888 | 8.235754 |
| 7.057327 | 6.93912 |
| 6.817894 | 6.49315 |
| 7.38799 | 7.474712 |
| 6.242471 | 6.010817 |
| 8.038502 | 8.125043 |
| 4.771985 | 4.823925 |
| 7.458086 | 7.897542 |
| 3.852726 | 3.349187 |
| 6.14464 | 5.293958 |
| 5.133746 | 5.636472 |
| 7.012906 | 6.974374 |
| 8.360435 | 8.612062 |
| 7.157963 | 5.29135 |

| | |
|---|---|
| NaN | NaN |
| 7.731244 | 7.793621 |
| 6.260776 | 5.95121 |
| 4.883201 | 4.804338 |
| 6.384478 | 6.731915 |
| 6.453363 | 6.390628 |
| 6.631777 | 6.847726 |
| 6.795607 | 6.599637 |
| 9.990086 | 10.10992 |
| 7.250222 | 6.975102 |
| 6.481382 | 7.050148 |
| 7.832428 | 7.957753 |
| 10.73816 | 10.7169 |
| 6.576302 | 6.663049 |
| 7.036915 | 6.965672 |
| 6.944231 | 7.225711 |
| 7.342371 | 7.383616 |
| 4.754746 | 4.517827 |
| 6.273279 | 6.733625 |
| 6.195963 | 5.914057 |
| 7.23498 | 7.356315 |
| 8.153084 | 8.120472 |
| 6.829299 | 7.52556 |
| 5.32723 | 5.555964 |
| NaN | NaN |
| 6.973449 | 6.636635 |
| 7.738605 | 7.662056 |
| 4.447623 | 5.063258 |
| 7.919788 | 8.070695 |
| 7.334948 | 7.440504 |
| 6.685238 | 5.891664 |
| 7.661904 | 7.742551 |
| 8.579318 | 8.664562 |
| 5.314951 | 5.168561 |
| 6.873111 | 6.447042 |
| 7.101334 | 7.179459 |
| 7.97032 | 7.737346 |
| 5.819483 | 5.215297 |
| 3.363037 | 3.518502 |
| 6.952969 | 6.454287 |
| 5.610803 | 5.316342 |
| NaN | NaN |
| 6.837996 | 6.872168 |
| 7.774082 | 7.907807 |
| 8.693893 | 8.922447 |
| NaN | NaN |
| 7.207958 | 7.101982 |

| | |
|---|---|
| 6.739401 | 6.67036 |
| 5.61526 | 5.290661 |
| 6.494565 | 6.595633 |
| 7.427149 | 7.455995 |
| 6.189775 | 6.152273 |
| 8.045903 | 8.042728 |
| 7.614645 | 7.165346 |
| 5.35031 | 5.683104 |
| 7.460203 | 7.51979 |
| 6.463015 | 6.128872 |
| 6.632703 | 7.034457 |
| 8.44863 | 8.501741 |
| 7.584809 | 7.601579 |
| 5.834434 | 5.695135 |
| 6.773407 | 6.590022 |
| 5.164691 | 5.498333 |
| 7.511968 | 7.284488 |
| 6.479967 | 6.65887 |
| 5.073771 | 4.939279 |
| 7.495296 | 7.358626 |
| 5.839631 | 5.733361 |
| 7.264004 | 8.455306 |
| 6.40627 | 6.111994 |
| 6.475903 | 6.44084 |
| 7.351357 | 6.614861 |
| NaN | 4.667679 |
| 6.582793 | 6.434953 |
| 2.470325 | NaN |
| 8.733356 | 8.76938 |
| 6.710653 | 6.182198 |
| 8.039415 | 8.158755 |
| 6.889743 | 6.758608 |
| 6.848509 | 6.520817 |
| 7.223978 | 6.774472 |
| 5.805926 | 5.719918 |
| 7.35284 | 7.110036 |
| 5.78732 | 6.173711 |
| 6.578742 | 6.231276 |
| 6.169684 | 6.632048 |
| 6.3087 | 6.691435 |
| 8.40936 | 8.374505 |
| 10.1992 | 10.35042 |
| 9.478848 | 9.629395 |
| 6.050565 | 5.935019 |
| 7.588115 | 7.664639 |
| 7.851715 | 7.660141 |
| 5.142186 | 5.107601 |

| | |
|---|---|
| 5.795626 | 6.082592 |
| NaN | 6.013919 |
| 6.350593 | 6.652568 |
| 5.561622 | 6.004954 |
| 5.618414 | 5.055364 |
| 6.720991 | 6.450533 |
| 6.807489 | 6.577875 |
| 6.871683 | 6.985585 |
| 9.437253 | 9.079443 |
| NaN | 5.434062 |
| 6.290189 | 6.773588 |
| 8.565668 | 8.684049 |
| 7.877154 | 8.297166 |
| 6.057195 | 5.884492 |
| 7.396633 | 7.245525 |
| 8.428204 | 8.361885 |
| 8.516343 | 8.229639 |
| 9.93047 | 9.286915 |
| 6.536927 | 6.760612 |
| 7.803661 | 7.769977 |
| 5.422104 | 5.01875 |
| 6.713056 | 6.181744 |
| 9.293025 | 9.40042 |
| NaN | NaN |
| 9.681509 | 9.760113 |
| 6.441876 | 6.490961 |
| 5.947039 | 6.160233 |
| 6.707116 | 7.138397 |
| 7.869188 | 8.049127 |
| 3.905521 | 3.284681 |
| 7.195862 | 4.03542 |
| 7.529202 | 7.002342 |
| 6.256571 | 6.167484 |
| 7.803084 | 8.029336 |
| 4.645062 | 4.715052 |
| 5.876262 | 5.893455 |
| 6.174539 | 6.061566 |
| 7.660906 | 7.61177 |
| 4.745271 | 5.162999 |
| 7.506358 | 7.535073 |
| 6.53966 | 6.404585 |
| 5.526844 | 4.888576 |
| 6.407835 | 6.40413 |
| 6.217651 | 6.402683 |
| 7.13945 | 6.987467 |
| 8.66441 | 8.673943 |
| 7.549575 | 7.908936 |

| | |
|---|---|
| 6.445394 | 6.244316 |
| 7.690409 | 7.668515 |
| 7.270088 | 7.301393 |
| 7.008451 | 6.985062 |
| 7.447329 | 7.709474 |
| 5.989075 | 6.549629 |
| 6.958647 | 6.76891 |
| 6.462749 | 6.068054 |
| 9.392873 | 9.325385 |
| 5.209797 | 5.140958 |
| 5.882956 | 5.834384 |
| 5.395944 | 5.065941 |
| 7.057535 | 7.195583 |
| 6.742645 | 6.862872 |
| 4.808622 | 4.47152 |
| 6.12593 | 5.95301 |
| 7.152642 | 7.105924 |
| 5.616055 | 5.381701 |
| 6.445833 | 6.437431 |
| 7.857424 | 8.334246 |
| 5.519855 | 5.598298 |
| 9.504935 | 9.673745 |
| 6.133498 | 5.500412 |
| 7.541213 | 5.261625 |
| 6.648387 | 6.740971 |
| 4.324245 | 4.746857 |
| 7.901675 | 7.741745 |
| 8.513307 | 8.686426 |
| 7.726341 | 8.102934 |
| 6.358296 | 5.801748 |
| 11.20586 | 10.97923 |
| 6.129782 | 6.449977 |
| 6.499745 | 6.46877 |
| 7.745255 | 7.743086 |
| 5.963382 | 6.201488 |
| 8.414371 | 8.290542 |
| 6.99503 | 7.335755 |
| 6.465114 | 6.183324 |
| 7.370693 | 7.310256 |
| 6.998778 | 7.597795 |
| 7.037928 | 7.281578 |
| 7.287024 | 7.083368 |
| 6.772344 | 6.767686 |
| 6.650395 | 6.988602 |
| 7.24803 | 7.948565 |
| 8.218728 | 7.437807 |
| 6.529883 | 6.779103 |

| | |
|---|---|
| 4.456779 | 4.747265 |
| 4.438077 | 4.157229 |
| 6.413303 | 6.009198 |
| 7.091985 | 7.300853 |
| 6.446788 | 6.01438 |
| 6.538821 | 6.58673 |
| 7.202981 | 7.111096 |
| 7.106967 | 7.160696 |
| 6.862613 | 6.530475 |
| 5.081884 | 5.475317 |
| 8.61678 | 8.544503 |
| 8.380834 | 8.216605 |
| 8.067445 | 8.047439 |
| 6.742178 | 6.909125 |
| 5.55596 | 5.546896 |
| NaN | 5.82669 |
| 5.383272 | 5.571893 |
| 5.750396 | 4.902552 |
| 6.268757 | 6.579157 |
| NaN | NaN |
| 8.381287 | 8.451509 |
| 7.320358 | 6.985408 |
| 6.50735 | 6.570035 |
| 7.154696 | 6.643347 |
| 5.27886 | 4.69989 |
| 8.304876 | 8.376371 |
| 8.563507 | 8.255465 |
| 6.451844 | 6.453184 |
| 5.719983 | 5.500975 |
| 6.667784 | 6.522198 |
| 6.824142 | 6.428547 |
| 7.780951 | 7.794271 |
| 5.846355 | 5.118637 |
| 7.822098 | 7.995547 |
| 7.38062 | 7.32759 |
| 7.020281 | 7.077381 |
| 6.931531 | 6.738693 |
| 7.04092 | 6.959273 |
| 6.633433 | 6.764041 |
| 5.742203 | 5.868778 |
| 6.768903 | 6.715355 |
| 8.216579 | 8.288466 |
| 11.14175 | 10.65118 |
| 6.935311 | 6.898861 |
| 7.254831 | 7.487595 |
| 6.53102 | 6.792859 |
| 6.518507 | 5.528194 |

| | |
|---|---|
| 9.38258 | 9.465888 |
| 7.18749 | 6.948826 |
| NaN | NaN |
| 7.763066 | 7.7238 |
| 9.948044 | 10.08835 |
| 7.536431 | 7.175725 |
| 6.011883 | 5.864432 |
| 6.72671 | 6.754227 |
| 7.33681 | 7.951365 |
| 5.801342 | 6.123931 |
| 5.931375 | 6.071176 |
| NaN | 8.229469 |
| 6.945224 | 6.889365 |
| NaN | 5.188064 |
| 6.001305 | 6.263299 |
| 7.27364 | 7.495277 |
| 7.979557 | 7.77317 |
| 6.740104 | 6.483663 |
| 5.922297 | 5.842798 |
| 8.253387 | 7.769019 |
| 7.326677 | 7.260793 |
| 7.378073 | 7.505222 |
| 5.536161 | NaN |
| 5.023761 | 5.19071 |
| 5.65889 | 5.464599 |
| 6.23503 | 6.911863 |
| 8.976876 | 8.737702 |
| 5.175784 | 5.037295 |
| 7.800986 | 7.583264 |
| 6.745314 | 6.74151 |
| 5.219667 | NaN |
| 6.732716 | 5.945999 |
| 6.909323 | 6.95503 |
| 6.744882 | 7.108894 |
| 6.863353 | 6.512153 |
| 6.15135 | 6.196469 |
| 7.920911 | 7.373169 |
| 6.850471 | 6.72583 |
| 5.831972 | 6.432352 |
| 6.594398 | 6.580844 |
| 6.509134 | 6.265804 |
| NaN | 4.451607 |
| 6.981141 | 7.185574 |
| NaN | 4.858928 |
| 7.862318 | 7.837869 |
| 5.440806 | 5.906142 |
| 4.216321 | 4.224449 |

-Bruker Confidential-

| | |
|---|---|
| 6.568245 | 5.361795 |
| 7.86523 | 7.625255 |
| 9.320038 | 9.210321 |
| 5.336397 | 4.942718 |
| 7.792112 | 7.662056 |
| 5.784627 | 5.914267 |
| 6.160342 | 6.078055 |
| 6.325815 | 6.278866 |
| 6.464873 | 5.778578 |
| 7.700389 | 7.707731 |
| 6.608299 | 6.038883 |
| 7.459556 | 7.665264 |
| 7.717502 | 7.756579 |
| 7.933802 | 8.338067 |
| 9.355192 | 8.256561 |
| 8.269381 | 8.326426 |
| 6.917717 | 6.033079 |
| 7.868823 | 8.342326 |
| 7.070735 | 6.956462 |
| 6.268055 | 6.557396 |
| 5.873271 | 5.892248 |
| 7.692222 | 8.045884 |
| 4.83839 | 5.971537 |
| 7.012585 | 7.169829 |
| 6.647914 | 6.574549 |
| 6.650012 | 6.382745 |
| 6.838304 | 7.071247 |
| NaN | 3.636889 |
| 6.500897 | 6.380719 |
| 6.515801 | 6.735433 |
| 7.823985 | 7.506081 |
| 7.159541 | 7.330167 |
| 7.370181 | 7.154482 |
| 8.765886 | 8.62495 |
| 7.390516 | 7.738139 |
| 7.424521 | 7.059953 |
| 6.943254 | 7.135225 |
| 6.628767 | 6.65282 |
| 7.235682 | 7.34784 |
| 8.190421 | 7.649338 |
| 7.094481 | 6.820374 |
| 6.76901 | 6.742746 |
| 5.762008 | 5.767745 |
| 6.461183 | 6.297246 |
| 6.229434 | 6.244898 |
| 6.118822 | 6.246504 |
| 7.292334 | 7.244163 |

| | |
|---|---|
| 5.844616 | 5.933392 |
| 6.363184 | 6.380374 |
| 7.264768 | 6.765407 |
| 5.809085 | 5.721052 |
| 6.494533 | 6.347262 |
| 7.207073 | 6.767206 |
| 7.008317 | 7.275932 |
| 5.822205 | 5.979953 |
| NaN | NaN |
| 5.615246 | 5.66355 |
| 6.435987 | 6.557306 |
| 8.176772 | 8.103846 |
| 4.435595 | 4.18855 |
| 6.867204 | 6.86213 |
| NaN | 5.861823 |
| 5.619242 | 5.11916 |
| 8.804443 | 8.838509 |
| NaN | NaN |
| NaN | 7.193829 |
| 10.01882 | 10.05182 |
| 7.771806 | 7.350558 |
| 4.938013 | 6.120915 |
| 6.87833 | 7.30222 |
| 5.823945 | 5.752608 |
| 8.052312 | 8.226971 |
| 5.861065 | 5.595329 |
| 7.67431 | 7.563226 |
| 8.797979 | 8.645019 |
| 6.927862 | 7.004894 |
| 7.492021 | 7.475821 |
| 6.610176 | 6.495371 |
| 8.402253 | 8.206534 |
| 3.845571 | 3.172482 |
| 6.646128 | 6.916878 |
| NaN | NaN |
| 8.807286 | 8.665213 |
| 8.882304 | 9.021752 |
| 6.670587 | 6.476991 |
| 7.222456 | 6.998444 |
| 9.88229 | 10.04601 |
| 6.959543 | 6.689568 |
| 6.949666 | 6.909411 |
| 6.604197 | 6.553062 |
| 9.833945 | 10.01119 |
| 5.224934 | 5.740898 |
| 7.708202 | 8.924145 |
| 6.617902 | 6.605992 |

| | |
|---|---|
| 7.009465 | 6.146553 |
| 7.808096 | 7.691302 |
| 7.552125 | 7.776547 |
| 4.67089 | 5.50075 |
| 7.46185 | 7.251932 |
| 6.85314 | 6.808124 |
| 6.028053 | 6.16583 |
| 8.868789 | 8.379827 |
| 5.976007 | 5.785535 |
| 7.148049 | 7.293181 |
| 6.251833 | 5.914443 |
| 5.444947 | 5.209108 |
| 6.344778 | 6.490456 |
| 6.317421 | 5.816023 |
| 6.503971 | 6.326931 |
| 6.481542 | 6.533252 |
| 7.036625 | 7.155051 |
| 6.242927 | 5.908072 |
| 5.146039 | 5.86376 |
| 7.623861 | 7.505334 |
| 5.694681 | 4.89003 |
| 6.300796 | 6.579041 |
| 7.441387 | 7.927955 |
| 6.335437 | 6.407058 |
| 6.511216 | 6.045573 |
| 4.559516 | 5.044906 |
| 6.466967 | 6.351501 |
| 7.269874 | 6.854946 |
| 6.518733 | 5.94624 |
| 6.150787 | 5.775057 |
| 3.923739 | 4.612341 |
| 6.340707 | 6.46443 |
| 5.488209 | 6.017982 |
| NaN | NaN |
| 7.434437 | 7.889254 |
| 8.388526 | 8.12388 |
| 7.097503 | 7.238456 |
| 7.298685 | 7.003531 |
| 6.090897 | 5.791924 |
| 4.858407 | 4.646581 |
| 6.656351 | 5.944859 |
| 6.816606 | 7.365471 |
| 5.945489 | NaN |
| 4.81988 | 5.289265 |
| 6.284321 | 6.218853 |
| 8.069087 | 7.068225 |
| 7.242775 | 7.287972 |

| | |
|---|---|
| 8.285172 | 8.619046 |
| 8.20529 | 8.201069 |
| 7.022284 | 7.152189 |
| 4.637252 | 5.877048 |
| 4.155659 | 4.216908 |
| 6.005477 | 5.568598 |
| 4.650404 | 5.490929 |
| 7.85491 | 7.795943 |
| 5.491754 | 5.130254 |
| 8.369619 | 8.723819 |
| 3.755569 | 4.271863 |
| 8.642014 | 8.605712 |
| 7.627588 | 6.568648 |
| 6.218209 | 6.442339 |
| NaN | 4.210584 |
| 4.80114 | 4.386558 |
| 7.650641 | 7.429945 |
| 6.721046 | 6.570174 |
| 7.167653 | 6.75978 |
| NaN | NaN |
| 9.170643 | 9.07445 |
| 8.592728 | 8.515899 |
| 6.889265 | 7.186856 |
| 6.702084 | 6.311592 |
| 5.964761 | 6.279497 |
| 4.563226 | 5.085855 |
| 6.31293 | 6.20076 |
| 8.45437 | 7.909012 |
| 5.843583 | 5.514948 |
| 9.147463 | 8.962983 |
| 7.121828 | 7.10236 |
| 5.845842 | 5.894597 |
| 6.418543 | 6.400975 |
| 5.506217 | 5.239606 |
| 5.957022 | 6.46467 |
| 7.344959 | 7.589794 |
| NaN | NaN |
| 6.759379 | 6.711746 |
| 7.123577 | 7.478783 |
| 6.568733 | 7.002394 |
| 7.688018 | 7.30476 |
| 6.880075 | 6.949957 |
| 2.825589 | 3.660488 |
| 6.481274 | 6.838992 |
| 6.911411 | 6.586616 |
| 7.522071 | 7.675465 |
| 7.626862 | 7.750529 |

-Bruker Confidential-

| | |
|---|---|
| 6.69318 | 6.586298 |
| 8.64294 | 8.381299 |
| 6.523382 | 6.274484 |
| 6.443587 | 6.722078 |
| 6.095313 | 6.044402 |
| 7.070107 | 7.034198 |
| 7.736737 | 7.917289 |
| 7.405627 | 7.380849 |
| 8.34578 | 8.437372 |
| 8.913513 | 8.82627 |
| 9.043741 | 8.850434 |
| NaN | NaN |
| 7.124407 | 7.244356 |
| 5.85795 | 6.106342 |
| 8.614433 | 8.096685 |
| 5.740499 | 5.73172 |
| 6.286978 | 6.364865 |
| 5.545614 | 4.831251 |
| NaN | NaN |
| 9.426708 | 9.444874 |
| 6.571285 | 6.271938 |
| 8.167399 | 7.722398 |
| 8.829727 | 8.948539 |
| 5.664787 | 6.34747 |
| 5.137279 | 6.619535 |
| 7.023837 | 7.015581 |
| 5.767664 | 5.634821 |
| 7.282011 | 6.864575 |
| 7.636996 | 7.297817 |
| 8.509459 | 8.393567 |
| 6.580627 | 6.393876 |
| 6.707193 | 6.360443 |
| 6.98407 | 7.072006 |
| 6.17709 | 5.879171 |
| 5.494499 | 5.285867 |
| NaN | 6.389975 |
| 5.919321 | 6.298095 |
| 8.096437 | 8.226493 |
| 6.286362 | 6.276553 |
| 8.273172 | 8.525042 |
| 7.300325 | 7.001852 |
| 6.088246 | 6.1791 |
| NaN | 4.406034 |
| 7.09442 | 6.842955 |
| 7.94748 | 7.827175 |
| 6.251444 | 6.450854 |
| 7.096643 | 7.126076 |

| | |
|---|---|
| NaN | 6.592875 |
| 6.471397 | 6.555298 |
| 5.498863 | 5.660327 |
| 7.001675 | 7.124387 |
| 6.409808 | 6.55529 |
| 6.920099 | 7.002034 |
| 6.720366 | 6.744456 |
| 6.344845 | 5.231277 |
| 5.591938 | 4.973897 |
| 6.393242 | 6.69027 |
| 6.648603 | 6.203413 |
| 9.064525 | 8.877238 |
| 6.847383 | 6.564268 |
| 6.881097 | 6.994748 |
| 6.26275 | 6.083971 |
| 6.982074 | 6.741652 |
| 6.535411 | 6.625983 |
| 6.425344 | 6.399712 |
| 6.316987 | 7.021961 |
| 8.018326 | 8.368667 |
| 6.871126 | 6.916609 |
| 7.695177 | 7.673445 |
| 7.378262 | 7.239769 |
| 5.455743 | 5.144376 |
| 6.54834 | 6.647679 |
| 7.772206 | 7.854768 |
| 5.967772 | 5.968069 |
| 6.508677 | 5.490027 |
| 8.03599 | 7.71541 |
| 6.923449 | 6.984092 |
| 6.636467 | 6.619279 |
| 8.631228 | 8.203592 |
| 6.848744 | 6.733689 |
| 7.223406 | 7.256644 |
| 6.677123 | 6.610712 |
| 6.076746 | 5.817302 |
| 5.404326 | 6.159943 |
| 6.80079 | 7.04872 |
| 8.206859 | 8.465271 |
| 6.673625 | 6.491631 |
| 6.359888 | 6.012888 |
| 8.007488 | 8.054875 |
| 7.237954 | 7.139417 |
| 6.446489 | 5.348985 |
| 6.526027 | 6.429434 |
| 5.70646 | 6.055333 |
| 6.034565 | 5.932947 |

| | |
|---|---|
| 6.713877 | 6.518283 |
| 5.569977 | 4.964902 |
| 5.956054 | 5.751464 |
| 6.907504 | 5.777486 |
| 7.70593 | 7.512897 |
| 6.143199 | 5.271506 |
| 6.348651 | 6.518441 |
| 4.308239 | 5.264734 |
| 8.600809 | 8.807367 |
| 7.806231 | 7.785417 |
| 7.779509 | 7.622742 |
| 9.532567 | 8.595077 |
| NaN | NaN |
| 5.398091 | 5.486944 |
| 5.928867 | 5.685871 |
| 5.314076 | 5.758979 |
| 7.462009 | 7.361361 |
| 5.830859 | 6.340854 |
| 8.820306 | 8.831222 |
| 11.64763 | 11.55563 |
| 7.275491 | 7.107246 |
| 6.907861 | 6.418607 |
| 6.73385 | 6.494716 |
| 5.88288 | 5.82602 |
| 8.966221 | 8.877681 |
| 6.481217 | 6.413966 |
| 5.448864 | 5.644004 |
| 6.349496 | 6.491199 |
| 7.304242 | 7.220142 |
| 7.412083 | 7.275337 |
| 5.207895 | 4.724714 |
| 7.427328 | 7.469542 |
| 7.881385 | 7.728615 |
| 7.484529 | 7.521797 |
| 5.268803 | NaN |
| 6.055967 | 5.16937 |
| NaN | 4.355556 |
| 8.105228 | 8.066455 |
| 6.591465 | 6.791504 |
| 8.182364 | 8.37964 |
| 8.229194 | 8.316563 |
| 7.101658 | 6.818181 |
| 5.922485 | 6.305472 |
| 5.65833 | 4.788473 |
| 8.254307 | 7.869219 |
| 6.892876 | 7.28169 |
| 6.28657 | 6.448692 |

-Bruker Confidential-

| | |
|---|---|
| 3.212608 | 3.452625 |
| 8.786658 | 8.592629 |
| 7.283105 | 7.428582 |
| 5.503649 | 5.682318 |
| 7.472936 | 8.147141 |
| 6.053433 | 6.183894 |
| 6.473928 | 6.540726 |
| 4.981018 | 4.995911 |
| 6.537313 | 5.961544 |
| 7.679585 | 7.02343 |
| 8.010528 | 7.726125 |
| 7.342207 | 7.585329 |
| 5.662656 | 6.158807 |
| 6.654664 | 6.790942 |
| 4.49783 | 4.287847 |
| 7.74318 | 7.95483 |
| 6.781286 | 6.692605 |
| 6.403986 | 6.888274 |
| NaN | 9.983408 |
| 5.505763 | 5.552113 |
| 7.091407 | 7.226686 |
| 6.42226 | 6.573586 |
| 7.434498 | 7.404205 |
| 5.026519 | 5.312137 |
| 5.593088 | 5.229778 |
| 6.567488 | 6.326672 |
| 6.680219 | 6.765779 |
| 5.771868 | 5.095485 |
| 7.041337 | 7.157345 |
| 7.027485 | 7.109488 |
| 5.351671 | 5.30184 |
| 6.136357 | 5.844565 |
| 7.185058 | 7.140441 |
| 5.758238 | 5.941064 |
| 8.189522 | 8.233827 |
| 7.298393 | 7.749208 |
| 8.03347 | 8.088465 |
| 7.235241 | 7.402225 |
| 7.781304 | 7.867118 |
| NaN | NaN |
| 6.306222 | 6.279595 |
| 6.448247 | 6.754518 |
| 6.652827 | 6.165572 |
| 6.668859 | 6.660627 |
| 7.498914 | 7.691381 |
| 7.01545 | 6.752897 |
| 7.401083 | 6.551535 |

| | |
|---|---|
| 5.370693 | 4.386859 |
| 7.840678 | 7.756472 |
| 5.933828 | 5.981444 |
| NaN | 4.31702 |
| 5.006474 | 4.055412 |
| 8.170548 | 6.813867 |
| 12.03562 | 8.358283 |
| 6.370828 | 6.728849 |
| 7.270842 | 7.291155 |
| 7.257936 | 7.171263 |
| 6.222509 | 6.335 |
| 7.639266 | 7.245392 |
| 8.291925 | 8.2256 |
| 7.569658 | 7.431956 |
| 8.349366 | 7.914356 |
| 6.678489 | 6.457443 |
| NaN | 6.814478 |
| 6.324974 | 5.93197 |
| 7.396124 | 7.839111 |
| 6.012741 | NaN |
| 7.290395 | 7.281609 |
| 6.651537 | 6.327699 |
| 5.343229 | 5.246556 |
| 8.543844 | 8.450098 |
| 6.559334 | 6.494219 |
| 7.545874 | 7.743586 |
| 8.497647 | 8.510658 |
| 6.302226 | 6.163126 |
| 6.558228 | 6.516224 |
| 7.183315 | 7.380657 |
| 7.979583 | 7.982712 |
| 6.270826 | 6.266634 |
| 5.653844 | 5.026533 |
| 7.768997 | 7.566401 |
| 7.262259 | 7.494151 |
| 6.558402 | 6.915362 |
| 4.993752 | 5.78493 |
| 8.38951 | 8.253887 |
| 8.454705 | 8.313895 |
| 6.240359 | 6.403413 |
| NaN | 6.777017 |
| 8.620157 | 8.860332 |
| 7.230234 | 8.100591 |
| 6.318521 | NaN |
| 5.755914 | 6.175945 |
| 6.612151 | 6.652351 |
| 7.748477 | 7.735693 |

-Bruker Confidential-

| | |
|---|---|
| 6.919282 | 6.784543 |
| 6.682245 | 6.019837 |
| 5.130725 | NaN |
| 7.676847 | 7.660771 |
| 8.159361 | 8.18169 |
| 8.598462 | 8.590402 |
| 7.578824 | 7.403035 |
| 3.957387 | 4.533074 |
| 8.123158 | 7.999239 |
| 7.031005 | 7.189291 |
| 7.012169 | 7.052349 |
| 5.925807 | 6.043331 |
| 7.040549 | 7.010626 |
| 8.901501 | 8.813354 |
| NaN | 3.975645 |
| 6.402519 | 6.352121 |
| 7.539961 | 7.586416 |
| 7.486597 | 7.541104 |
| 8.186618 | 8.493572 |
| 7.646376 | 6.477397 |
| 8.069955 | 7.726751 |
| 7.06345 | 7.163891 |
| 10.46502 | 10.52652 |
| 5.886381 | 6.235294 |
| 7.556115 | 7.064544 |
| 6.522928 | 6.397365 |
| 5.980232 | 5.852027 |
| 8.461575 | 8.400743 |
| NaN | 7.506748 |
| 5.836911 | 5.634018 |
| 5.384689 | 4.839334 |
| 5.975382 | 6.198019 |
| 10.23868 | 10.10524 |
| 7.650778 | 7.667117 |
| 5.430513 | 5.199747 |
| 7.667563 | 7.68042 |
| 3.969442 | 3.744806 |
| 7.59547 | 7.196157 |
| 6.844723 | 6.581454 |
| NaN | NaN |
| 7.027296 | 6.987946 |
| 8.087946 | 7.664614 |
| 6.425623 | 6.679183 |
| 8.201158 | 7.775993 |
| 6.394884 | 6.525057 |
| 6.11265 | 6.183073 |
| 6.98292 | 6.901855 |

| | |
|---|---|
| 6.723592 | 6.364368 |
| 4.971548 | 4.976947 |
| 6.880455 | 6.958526 |
| 6.151656 | 6.39051 |
| 6.99578 | 6.67634 |
| 7.622622 | 7.429789 |
| 8.932305 | 9.071239 |
| 6.193612 | 6.017487 |
| 6.137998 | 5.93399 |
| 9.412873 | 9.451924 |
| 6.376392 | 6.542442 |
| 7.34583 | 7.587463 |
| 7.172904 | 7.265747 |
| 10.39256 | 10.43583 |
| 7.370213 | 7.460378 |
| NaN | 4.860532 |
| 8.010264 | 7.715295 |
| 6.682324 | 6.463285 |
| 6.220359 | 6.247099 |
| 6.789833 | 6.765227 |
| 9.6677 | 9.711693 |
| 7.572421 | 6.971566 |
| 3.586518 | NaN |
| 5.658071 | 5.206773 |
| 4.863568 | 5.382043 |
| 8.268731 | 8.267749 |
| 7.294672 | 7.224434 |
| 6.238739 | 6.076943 |
| 4.599379 | 4.601511 |
| 6.874182 | 6.810122 |
| 5.342381 | 4.968824 |
| 6.308594 | 6.897336 |
| 8.454618 | 7.544166 |
| 6.280941 | 6.262345 |
| 6.647849 | 6.674365 |
| 6.172095 | 5.908044 |
| 6.972575 | 6.823611 |
| 6.844223 | 6.475539 |
| 10.10069 | 10.15439 |
| 6.937173 | 6.961597 |
| 8.14736 | 7.803742 |
| 6.240562 | 6.408848 |
| 6.872006 | 6.493537 |
| 8.608185 | 8.557502 |
| 6.590023 | 6.500342 |
| 7.950716 | 7.784583 |
| 7.470459 | 7.478386 |

| | |
|---|---|
| 11.35319 | 11.24226 |
| 6.053336 | 5.541263 |
| 7.627398 | 7.603888 |
| 8.339836 | 8.551032 |
| 8.130908 | 8.236428 |
| 7.672743 | 7.817295 |
| 5.959806 | 6.091124 |
| 7.113262 | 7.199858 |
| 9.504837 | 9.956346 |
| 6.700487 | 6.525891 |
| 6.610917 | 7.144409 |
| 8.021125 | 7.869872 |
| 9.360036 | 9.007272 |
| 3.394052 | 3.792749 |
| 8.558042 | 8.517864 |
| 7.878508 | 8.425688 |
| 6.058185 | 5.668062 |
| 7.456592 | 7.644809 |
| 4.561762 | 3.997375 |
| 7.315691 | 6.375815 |
| 6.807926 | 6.970365 |
| 5.41416 | 5.811239 |
| 7.525405 | 7.405917 |
| 6.282841 | 6.118041 |
| 7.541506 | 7.884867 |
| 6.77025 | 6.574888 |
| 5.796258 | 5.800723 |
| 8.831861 | 8.877961 |
| 7.276742 | 7.255548 |
| 5.382135 | 5.525954 |
| NaN | NaN |
| 10.50123 | 10.64151 |
| 7.905017 | 8.063641 |
| 7.278833 | 7.274744 |
| 7.057988 | 7.010839 |
| 8.230603 | 8.055669 |
| 6.790246 | 6.678046 |
| 6.786424 | 6.499186 |
| 7.128935 | 7.13067 |
| 7.218346 | 7.2288 |
| 7.57239 | 7.410638 |
| 8.792236 | 8.84334 |
| 8.812012 | 8.681099 |
| 6.82181 | 6.892237 |
| 6.849119 | 7.43369 |
| 9.269031 | 9.796834 |
| 5.89698 | 6.326756 |

| | |
|---|---|
| 7.444236 | 7.270742 |
| 6.155741 | 5.899471 |
| 6.246237 | 6.071316 |
| 10.03005 | 9.611247 |
| 6.163805 | 6.023579 |
| 5.609132 | 5.567533 |
| 9.257702 | 9.300838 |
| 7.411698 | 7.204643 |
| 6.467286 | 5.831028 |
| 6.810114 | 6.981355 |
| 5.666122 | 5.800919 |
| 6.487742 | 6.320045 |
| 8.287682 | 8.345266 |
| 8.582163 | 8.418479 |
| 4.777108 | 4.31157 |
| 8.604591 | 8.535234 |
| 5.73328 | 5.999212 |
| 7.519207 | 7.482179 |
| 7.446443 | 6.669372 |
| 6.154126 | 6.627397 |
| 5.255284 | 4.716885 |
| 8.387709 | NaN |
| 7.393348 | 7.010108 |
| 5.680059 | 6.116189 |
| 6.570283 | 6.453367 |
| 8.326735 | 8.022385 |
| 5.758183 | 5.673999 |
| 7.818841 | 7.733007 |
| 9.362905 | 9.724698 |
| 6.747149 | 6.895498 |
| 7.441278 | 7.441731 |
| 7.148292 | 6.926676 |
| NaN | 3.117246 |
| 4.549438 | 4.938808 |
| 6.620242 | 6.773689 |
| 5.886337 | 7.064297 |
| 7.641155 | 7.212985 |
| 7.451427 | 7.537837 |
| 7.063384 | 7.267508 |
| NaN | NaN |
| 6.532276 | 5.429548 |
| 5.418681 | NaN |
| 5.98207 | 6.173494 |
| 7.748378 | 7.525469 |
| 8.226999 | 8.225656 |
| 7.094228 | 7.064292 |
| 5.76169 | NaN |

| | |
|---|---|
| 7.297466 | 6.993463 |
| 8.403426 | 8.705013 |
| 6.390201 | 6.210249 |
| 5.802858 | 5.681559 |
| 6.906139 | 6.718428 |
| 6.073114 | 6.140278 |
| 7.655021 | 7.822561 |
| 7.864067 | 8.198308 |
| 7.033806 | 6.889754 |
| 7.209707 | 7.010767 |
| 8.180074 | 8.007157 |
| 9.555491 | 9.420792 |
| 5.543496 | 5.27887 |
| 7.274433 | 7.703036 |
| 6.920879 | 7.097221 |
| 6.782598 | 6.565577 |
| 7.788104 | 7.719545 |
| 10.25555 | 10.23145 |
| 9.311298 | 9.058637 |
| 5.689903 | 6.284834 |
| 7.193909 | 7.278728 |
| 8.28589 | 8.338179 |
| 6.620591 | 6.435727 |
| 8.545873 | 8.663872 |
| 9.556416 | 9.760322 |
| 6.526194 | 6.327129 |
| 5.527775 | 5.555209 |
| 8.162236 | 8.307783 |
| 6.537803 | 6.433853 |
| 6.891004 | 7.475367 |
| 9.367216 | 9.320215 |
| 6.609756 | 6.790719 |
| 4.820167 | 5.318738 |
| 6.170603 | 6.458949 |
| 7.586839 | 7.385784 |
| 5.613136 | 5.105675 |
| 5.514436 | 4.864655 |
| 7.658919 | 7.683002 |
| 8.491838 | NaN |
| 9.695676 | 9.785467 |
| NaN | NaN |
| 7.027795 | 6.780965 |
| 6.771087 | 6.828854 |
| 6.984914 | 7.030026 |
| 6.070731 | 6.130501 |
| 7.848649 | NaN |
| 7.017001 | 7.061031 |

| | |
|---|---|
| 7.67172 | 7.532277 |
| 5.951576 | 5.838173 |
| 5.06734 | 5.010769 |
| 8.252691 | 7.067596 |
| 4.371864 | 4.55433 |
| 7.736824 | 7.555961 |
| 7.012367 | 7.265088 |
| 8.993179 | 8.813937 |
| 6.223117 | 6.424026 |
| 7.303405 | 7.229894 |
| 7.717748 | 7.906236 |
| 6.663128 | 6.499034 |
| 7.799965 | 7.686163 |
| 6.991676 | 6.759303 |
| 7.408528 | 6.24466 |
| 6.78334 | 6.545548 |
| 8.33361 | 8.389677 |
| 6.970617 | 6.670473 |
| 6.2954 | 6.539566 |
| 7.450615 | 7.432924 |
| 8.967311 | 9.012909 |
| 6.99368 | 6.697488 |
| 7.085742 | 7.103381 |
| 9.52405 | 9.560295 |
| 7.370514 | 7.452682 |
| 4.961044 | 5.206663 |
| 7.798948 | 7.811605 |
| 7.579599 | 7.602595 |
| 6.7015 | 6.861849 |
| 6.09254 | 6.24966 |
| 8.878648 | 8.812023 |
| 7.349489 | 7.120815 |
| 7.174194 | 7.171818 |
| 6.014352 | 5.788102 |
| 6.93961 | 7.253985 |
| 7.853789 | 7.987547 |
| 5.644297 | 5.540392 |
| 7.666404 | 7.789306 |
| 8.89315 | 8.880715 |
| 8.205341 | 8.376604 |
| 6.619106 | 6.702434 |
| 6.085293 | 5.277855 |
| 7.431707 | 7.91917 |
| 7.200692 | NaN |
| 6.072414 | 6.855717 |
| 8.272242 | 8.167916 |
| 8.380104 | 7.826954 |

-Bruker Confidential-

| | |
|---|---|
| 6.885969 | 5.821693 |
| 5.833862 | 6.051993 |
| 6.379452 | 5.918306 |
| 7.533444 | 7.496099 |
| 5.430151 | 5.741415 |
| 6.116652 | NaN |
| 4.966092 | 5.448362 |
| 6.037969 | 5.365477 |
| 7.520264 | 7.354269 |
| 6.867526 | 6.621397 |
| 7.332577 | 6.773586 |
| 5.562607 | 5.183563 |
| 5.857344 | 6.558595 |
| 4.090394 | 5.438821 |
| NaN | 5.611778 |
| 4.521929 | 4.452627 |
| 8.665258 | 8.797826 |
| 7.570227 | 7.903724 |
| 7.754714 | 7.421991 |
| 7.871293 | 7.948522 |
| 5.151475 | 4.649752 |
| 6.836621 | 6.639055 |
| 6.237562 | 6.643963 |
| 6.981225 | 5.96901 |
| 8.177534 | 8.202432 |
| 7.040491 | 6.657594 |
| 6.538894 | 7.152027 |
| 7.168539 | 7.092435 |
| 6.493392 | 5.954221 |
| 9.505227 | 9.417656 |
| 7.337811 | 6.491524 |
| 5.99474 | 6.240388 |
| 9.804172 | 9.647048 |
| 8.071624 | 7.843696 |
| 8.667551 | 8.411674 |
| 7.246799 | 7.098546 |
| 6.746633 | 6.503993 |
| 8.599042 | 9.19805 |
| 7.294481 | 7.328654 |
| NaN | NaN |
| 4.550296 | 5.394998 |
| NaN | 5.953129 |
| 6.631829 | 6.46001 |
| 5.838393 | 6.410103 |
| 9.785181 | 9.661605 |
| 7.591248 | 7.662354 |
| 6.719107 | 6.844043 |

| | |
|---|---|
| 7.549693 | 7.964277 |
| 7.544514 | 7.492556 |
| 7.768566 | 7.314857 |
| 8.010371 | 6.712994 |
| 7.216622 | 7.395832 |
| 5.948027 | 5.805749 |
| 6.161295 | 6.302958 |
| 5.196069 | 5.090183 |
| 6.640537 | 9.023877 |
| NaN | NaN |
| 9.367413 | 9.742082 |
| 6.678212 | 6.427278 |
| 7.376847 | 7.380101 |
| 6.115123 | 5.948716 |
| 8.019606 | 8.526345 |
| 7.278248 | 7.479248 |
| 6.040148 | 6.053724 |
| 6.552755 | 6.691107 |
| NaN | NaN |
| 6.980069 | 6.614235 |
| 7.279983 | 7.251282 |
| 7.060293 | 7.166462 |
| NaN | NaN |
| 7.760991 | 7.607747 |
| 7.146433 | 5.728101 |
| 4.047445 | 4.211754 |
| 7.291106 | 6.778988 |
| NaN | 5.48326 |
| 6.305931 | 6.217003 |
| 7.833132 | 8.206973 |
| 6.762946 | 7.089946 |
| 7.39658 | 7.011435 |
| 6.457549 | 6.452377 |
| 7.920606 | 7.781742 |
| 8.137129 | 8.174582 |
| 4.797269 | 5.580298 |
| 8.891055 | 8.834578 |
| 8.917261 | 8.490017 |
| 5.988022 | 6.109022 |
| 7.273377 | 7.377417 |
| 9.489193 | 8.824479 |
| 6.635002 | 6.850749 |
| 6.04604 | 5.605974 |
| 6.139766 | 5.835548 |
| 7.411602 | 7.468755 |
| 4.212458 | 3.515382 |
| 9.445242 | 9.388383 |

| | |
|---|---|
| 6.687462 | 6.981392 |
| 5.871678 | 5.942721 |
| 8.302798 | 8.306981 |
| 5.887239 | 5.761803 |
| 7.758393 | 7.776087 |
| 8.429612 | 8.505197 |
| 7.255976 | 7.193976 |
| 7.385617 | 6.592184 |
| 8.302152 | 8.447304 |
| 6.891815 | 7.684689 |
| NaN | NaN |
| 7.322546 | 7.414917 |
| 7.014331 | 7.035406 |
| 6.885139 | 6.521087 |
| 6.801846 | 6.798324 |
| 6.380776 | 6.259261 |
| 4.942787 | 4.757693 |
| 6.342496 | 6.215242 |
| 8.652507 | 7.65159 |
| 5.792063 | 5.666314 |
| 7.351702 | 7.032657 |
| 8.268601 | 7.852954 |
| 6.447736 | 6.365407 |
| 5.928957 | 5.901298 |
| 6.673725 | 6.612069 |
| 6.393946 | 5.82452 |
| 8.789309 | 8.891421 |
| 6.171015 | 5.961758 |
| 7.588037 | 7.671098 |
| 6.861846 | 6.957078 |
| 4.712071 | 5.91943 |
| 8.726942 | 8.506008 |
| 7.234833 | 6.93072 |
| 8.091683 | 8.18611 |
| 7.671942 | 7.884284 |
| 4.699501 | 4.037041 |
| 5.413533 | 5.384474 |
| 7.440863 | 7.355193 |
| 7.99436 | 8.041629 |
| 5.770008 | 5.660686 |
| 7.254438 | 7.173694 |
| 7.320498 | 7.859919 |
| 5.643862 | 5.374198 |
| NaN | NaN |
| 6.062781 | 5.978572 |
| 7.844077 | 7.867557 |
| 5.037478 | 5.984305 |

| | |
|---|---|
| 5.995023 | 6.353468 |
| NaN | NaN |
| 8.284608 | 8.455664 |
| 6.949405 | 7.12794 |
| 7.20837 | 7.057229 |
| 7.96459 | 7.863065 |
| 6.5848 | 6.669576 |
| 6.131067 | 6.267268 |
| 7.28905 | 6.643536 |
| NaN | NaN |
| 7.338994 | 7.218906 |
| 6.476777 | 6.631108 |
| 7.424232 | 7.555625 |
| 6.25183 | 6.698959 |
| 8.111819 | 8.253983 |
| 9.76951 | 9.746978 |
| 6.194303 | 5.979619 |
| 7.979526 | 7.811662 |
| 6.542408 | 6.491812 |
| NaN | NaN |
| 7.392774 | 6.883407 |
| 6.645585 | 6.673796 |
| 8.037266 | 8.095314 |
| 7.393301 | 7.188016 |
| 3.591276 | 4.250189 |
| 7.220463 | 7.374171 |
| 6.163019 | 5.859858 |
| 5.44332 | 5.585717 |
| 6.411998 | 5.587078 |
| 8.30258 | 8.382636 |
| 6.884801 | 6.664148 |
| 6.882387 | 7.256624 |
| 7.227906 | 6.848708 |
| 6.866901 | 6.881736 |
| 5.71819 | 5.93573 |
| 6.874915 | 6.596807 |
| NaN | NaN |
| 4.213119 | NaN |
| 6.599547 | 5.765995 |
| 4.787114 | 4.859403 |
| 6.764371 | 6.985178 |
| 8.440872 | 7.674513 |
| 6.702606 | 6.680351 |
| 5.59867 | 5.474986 |
| 5.816004 | 5.691806 |
| 6.731227 | 7.374886 |
| 7.837135 | 7.915096 |

| | |
|---|---|
| NaN | 4.943445 |
| 7.797554 | 7.898182 |
| 5.05958 | 4.727298 |
| 5.110576 | 5.578383 |
| 5.896128 | 5.941679 |
| 7.866851 | 7.653314 |
| 6.792165 | 6.693974 |
| 5.362407 | 5.55983 |
| 5.819158 | NaN |
| 6.056115 | 5.881019 |
| NaN | NaN |
| NaN | NaN |
| 7.201549 | 7.394024 |
| 7.727304 | 7.728404 |
| 6.051877 | 6.002327 |
| 7.868796 | 8.220388 |
| 7.298683 | 7.640335 |
| 6.846153 | 6.522221 |
| 4.838046 | 5.82532 |
| 10.21636 | 10.21117 |
| 7.40528 | 7.361155 |
| 7.5042 | 7.5913 |
| 6.101794 | 7.477148 |
| 9.548349 | 9.644724 |
| 6.826774 | 6.418954 |
| 4.765257 | 5.224082 |
| 6.328035 | 6.66584 |
| 10.1709 | 10.19421 |
| 8.203983 | 8.125754 |
| 5.834989 | 5.770748 |
| 7.455808 | 7.391436 |
| 8.426152 | 7.892992 |
| 6.806704 | 6.823347 |
| 6.533973 | 6.501538 |
| 8.017147 | 8.202005 |
| 6.953463 | 6.986317 |
| 4.785075 | 4.902034 |
| 6.137671 | 6.550297 |
| 5.462508 | 5.330468 |
| 7.823556 | 7.725163 |
| NaN | NaN |
| 7.305469 | 7.46313 |
| 9.96452 | 10.1228 |
| 8.241789 | 8.368502 |
| 6.540448 | 6.505084 |
| 5.883416 | 6.210322 |
| 7.401069 | 7.508559 |

| | |
|---|---|
| 7.123004 | 7.263804 |
| 6.469913 | 6.14083 |
| 4.965008 | 4.496112 |
| 8.141501 | 7.816257 |
| 6.923285 | 6.74036 |
| 7.704813 | 7.56955 |
| 6.902718 | 6.991305 |
| 6.109596 | 6.285511 |
| 7.192019 | 7.128998 |
| 7.187767 | 6.976985 |
| 6.774121 | 7.305862 |
| 4.243091 | 4.822423 |
| 6.899122 | 6.697136 |
| NaN | NaN |
| 5.972785 | 6.251451 |
| 6.944139 | 7.079457 |
| 6.896433 | 7.448276 |
| 6.995877 | 6.743697 |
| 7.220325 | 7.09806 |
| 7.169587 | 7.693262 |
| 8.13525 | 8.157804 |
| 6.58944 | 6.11483 |
| 6.026753 | 5.415252 |
| 5.72713 | 6.170656 |
| 8.772436 | 8.929471 |
| 5.880711 | 5.969799 |
| 7.131364 | 6.855572 |
| 5.305216 | 5.193494 |
| 7.137002 | 6.797893 |
| 4.894794 | 4.443082 |
| 6.950394 | 6.792037 |
| 7.509033 | 7.375899 |
| 7.313771 | 7.347081 |
| 8.161857 | 7.947171 |
| 6.139044 | 6.25884 |
| 6.194945 | 5.770501 |
| 7.17443 | 7.711069 |
| 7.02324 | 6.934212 |
| 3.661572 | 3.457388 |
| 8.987097 | 8.941008 |
| 7.832406 | 7.688253 |
| 6.928639 | 6.61386 |
| 6.746546 | 6.672134 |
| 7.216105 | 7.164001 |
| 6.993899 | 6.520057 |
| 6.923306 | 6.829937 |
| 7.171625 | 6.931828 |

-Bruker Confidential-

| | |
|---|---|
| 7.222092 | 7.14193 |
| 5.661501 | 5.124325 |
| 4.657634 | 4.522587 |
| 7.766848 | 7.663883 |
| 6.940756 | 6.786688 |
| 10.39133 | 10.54649 |
| 5.903317 | 5.866377 |
| 10.10626 | 9.580999 |
| 7.703122 | 7.236291 |
| 8.267881 | 8.278993 |
| 8.28496 | 8.180283 |
| 8.99894 | 8.986163 |
| NaN | 5.016724 |
| 7.383201 | 7.369722 |
| 9.628962 | 9.605409 |
| 6.372744 | 5.399411 |
| 4.845447 | 5.064549 |
| 5.861951 | 5.588792 |
| 9.787287 | 9.677075 |
| 4.122949 | 3.865296 |
| 7.990727 | 7.563636 |
| 7.161704 | 7.369688 |
| 11.98724 | 11.76214 |
| 4.892118 | 4.923065 |
| 7.069554 | 7.260211 |
| 7.009001 | 6.798999 |
| 7.213268 | 7.527838 |
| 7.664542 | 7.597005 |
| 5.774995 | 5.860131 |
| 6.055144 | 6.001902 |
| 6.527031 | 6.480687 |
| 6.879929 | 6.979058 |
| 9.096626 | 8.954318 |
| 7.710581 | 8.167898 |
| 6.847833 | 7.477503 |
| 7.467324 | 7.645508 |
| 6.208666 | 6.123967 |
| 5.084774 | 4.755436 |
| 9.98826 | 9.873981 |
| 6.848127 | 6.771282 |
| 7.728212 | 7.776777 |
| 7.326872 | 7.318804 |
| 11.94236 | 11.86875 |
| 9.954111 | 9.664096 |
| 6.323262 | 6.297996 |
| 6.267678 | 6.35322 |
| 6.152224 | 6.815515 |

| | |
|---|---|
| 9.730234 | 9.980143 |
| 7.307884 | 6.848688 |
| 4.976532 | 5.334355 |
| 6.680478 | 6.75821 |
| 7.332929 | 7.383255 |
| 6.971444 | 7.159584 |
| 9.220319 | 9.309425 |
| 7.204268 | 6.84186 |
| 5.778617 | 5.679304 |
| 8.083405 | 7.659073 |
| 7.647654 | 7.283017 |
| 7.897461 | 7.944777 |
| 9.840904 | 9.668791 |
| 9.711513 | 9.600277 |
| 9.023635 | 9.26339 |
| 8.096501 | 7.238614 |
| 7.157941 | 6.962552 |
| 6.23478 | 6.113651 |
| 7.417567 | 7.372228 |
| 6.308496 | 6.577393 |
| 5.739297 | 5.949407 |
| 10.50007 | 10.52571 |
| 7.819961 | 7.584014 |
| 6.917503 | 6.942207 |
| 7.909312 | 7.771604 |
| 9.596849 | 9.460695 |
| 10.74418 | 10.88317 |
| 7.760905 | 7.977082 |
| 5.450675 | 6.214919 |
| 8.123365 | 8.240947 |
| 8.179113 | 8.189132 |
| 6.753421 | 6.914225 |
| 6.360888 | 6.566585 |
| 7.936174 | 8.206621 |
| NaN | NaN |
| 6.70497 | 6.692122 |
| 7.917347 | 8.180089 |
| 8.306659 | 8.524387 |
| 6.999214 | 6.919908 |
| 6.402654 | 5.245015 |
| 7.159797 | 6.68198 |
| 6.720929 | 6.149939 |
| 8.146408 | 7.8332 |
| 7.802677 | 7.740824 |
| 9.770455 | 9.626392 |
| 9.828274 | 9.728316 |
| 6.883821 | 6.753128 |

| | |
|---|---|
| 9.570086 | 9.37834 |
| 6.57579 | 6.722552 |
| 6.315046 | 6.157726 |
| 5.434808 | 5.263365 |
| 9.139845 | 9.002136 |
| 5.937947 | 5.738677 |
| 6.63529 | 6.565947 |
| NaN | NaN |
| 5.47422 | 5.404056 |
| 5.520705 | 6.075129 |
| 6.809102 | 6.734651 |
| 6.333082 | 5.781424 |
| 6.248431 | 6.81859 |
| 6.68912 | 6.132079 |
| 7.915752 | 8.161937 |
| 7.633144 | 6.806807 |
| 5.756688 | 5.80556 |
| 6.859312 | 6.832533 |
| 6.518939 | 6.023624 |
| 7.764534 | 7.865554 |
| 5.760068 | 6.027506 |
| 6.74019 | 6.720123 |
| 6.362923 | 6.085666 |
| 6.011297 | 5.085562 |
| 6.070666 | 6.07825 |
| 4.982119 | 5.161797 |
| 5.895622 | 6.018423 |
| 5.730679 | 5.485329 |
| 7.191991 | 7.447958 |
| 7.703291 | 7.300267 |
| 8.068513 | 7.910234 |
| 7.015796 | 7.053236 |
| 8.566726 | 7.961017 |
| 6.871644 | 5.872108 |
| 7.245883 | 7.130912 |
| 6.999554 | 7.001056 |
| 6.898159 | 6.810833 |
| 8.510698 | 8.36977 |
| 8.677306 | 8.404452 |
| 7.000759 | 6.088817 |
| 8.657186 | 8.62196 |
| 5.604752 | 5.67684 |
| 7.34501 | 7.276596 |
| 6.106165 | 5.946122 |
| 7.90184 | 8.005481 |
| 6.151056 | 6.803412 |
| 6.404426 | 6.248045 |

-Bruker Confidential-

| | |
|---|---|
| 7.077913 | 6.616915 |
| 8.552331 | 8.350528 |
| 7.148923 | 5.549872 |
| 6.408242 | 6.248893 |
| 6.805068 | 6.329594 |
| 6.713928 | 6.513432 |
| 6.945721 | 7.192152 |
| 10.23881 | 10.04624 |
| 6.265522 | 6.475388 |
| 7.905009 | 7.287864 |
| 9.425113 | 9.179213 |
| 6.199814 | 6.187497 |
| 5.182675 | 5.147024 |
| 8.632742 | 8.77374 |
| 7.040214 | 7.049604 |
| 6.218667 | 6.436024 |
| 5.357162 | 5.455003 |
| NaN | NaN |
| 7.032473 | 6.811994 |
| 5.537213 | 5.970146 |
| 9.095411 | 9.258452 |
| 6.903511 | 6.710833 |
| 6.20153 | 6.12142 |
| 5.521393 | 4.866283 |
| 7.27834 | 7.436643 |
| 5.518394 | 5.041312 |
| 4.787501 | 5.062253 |
| 9.544412 | 9.600576 |
| 5.451725 | 5.406662 |
| 2.511209 | 7.584826 |
| 4.232718 | 4.876792 |
| 7.991703 | 8.05845 |
| 6.244002 | 5.171305 |
| 8.896868 | 8.749461 |
| 6.020958 | 6.50116 |
| 7.035232 | 7.365073 |
| 6.976541 | 6.720942 |
| 7.069912 | 6.470781 |
| 7.063274 | 7.521753 |
| 9.048702 | 8.956758 |
| 6.468784 | 6.768665 |
| 7.292634 | 7.815815 |
| 6.513978 | 6.901016 |
| 5.327577 | 4.826456 |
| 7.117373 | 7.990097 |
| 8.566125 | 8.634583 |
| 5.686789 | 5.446939 |

| | |
|---|---|
| 7.048758 | 7.008742 |
| 9.149605 | 9.072105 |
| 7.349232 | 7.266783 |
| 5.826573 | 6.397713 |
| 7.708351 | 7.871945 |
| 6.539925 | 6.374927 |
| 6.28183 | 6.292411 |
| 6.299316 | 6.426826 |
| 5.519551 | 5.786027 |
| 6.165603 | 6.139226 |
| NaN | 5.73878 |
| 8.104447 | 8.61729 |
| 6.561629 | 5.984842 |
| 6.913286 | 7.331559 |
| 6.570531 | 6.387018 |
| 7.210522 | 7.460533 |
| NaN | 8.082266 |
| 6.618307 | 6.282078 |
| 12.65842 | 12.82291 |
| 7.164171 | 7.072506 |
| 6.44294 | 6.500659 |
| 9.031272 | 9.179306 |
| 7.052125 | 6.491841 |
| 8.849547 | 8.629311 |
| NaN | 5.070646 |
| 6.747693 | 6.601084 |
| 7.787972 | 8.152141 |
| 6.067334 | 5.748483 |
| 7.952586 | 8.181655 |
| 6.8141 | 6.540096 |
| 7.428174 | 7.616143 |
| 7.85764 | 7.930749 |
| 6.845731 | 7.163961 |
| 8.284588 | 8.333417 |
| 6.437692 | 7.125098 |
| 5.495836 | 5.955716 |
| 5.087307 | 5.337073 |
| 7.736987 | 7.652815 |
| 5.513938 | 5.560636 |
| 7.44272 | 7.359666 |
| 5.165296 | 5.017798 |
| 7.018901 | 7.24297 |
| 6.206253 | 5.403019 |
| 5.200526 | 4.536696 |
| 13.59113 | 13.03926 |
| 7.466227 | 7.323127 |
| 6.151484 | 6.028953 |

-Bruker Confidential-

| | |
|---|---|
| 8.016479 | 8.381635 |
| 8.411293 | 8.405107 |
| 7.006945 | 5.483078 |
| 6.291937 | 5.912899 |
| 7.571199 | 6.89389 |
| 6.220131 | 5.795132 |
| 8.028507 | 7.946493 |
| 9.049879 | 9.233125 |
| 7.474142 | 7.852983 |
| 4.465665 | 4.822462 |
| 8.537504 | 8.145309 |
| 4.989913 | 5.126499 |
| 7.795569 | 7.643212 |
| 6.975563 | 6.814523 |
| 6.860983 | 6.766269 |
| 5.10684 | 5.745952 |
| 6.224438 | 6.406744 |
| 5.751304 | 5.828363 |
| 7.192713 | 7.028427 |
| 8.37248 | 8.180723 |
| 6.546397 | 6.534238 |
| NaN | 4.636497 |
| 4.156533 | 4.283221 |
| 8.951362 | 7.455787 |
| 6.72984 | 6.949077 |
| 7.604838 | 7.466922 |
| 6.558466 | 6.433939 |
| 7.85066 | 7.246993 |
| 9.56925 | 9.946306 |
| 4.203317 | 4.913862 |
| 6.707129 | 6.747768 |
| 7.047285 | 7.092493 |
| 8.646367 | 8.7591 |
| 8.282263 | 8.368983 |
| 6.461316 | 6.46595 |
| 8.502573 | 8.376819 |
| NaN | 7.674277 |
| 6.892784 | 6.947134 |
| 3.828864 | 4.241707 |
| 7.765342 | 7.692155 |
| 7.210911 | 6.958004 |
| 7.344147 | 7.533352 |
| 7.020653 | 7.065537 |
| 8.869333 | 8.381235 |
| 6.049045 | 6.331167 |
| 5.391437 | 5.857385 |
| 5.751289 | 5.822057 |

-Bruker Confidential-

| | |
|---|---|
| 7.582602 | 7.384633 |
| 7.410905 | 7.419078 |
| 6.711096 | 6.822441 |
| 6.480992 | 5.887627 |
| NaN | NaN |
| 7.723892 | 7.574383 |
| 6.525593 | 5.929821 |
| 7.977419 | 7.95013 |
| 7.45803 | 7.436713 |
| 8.029172 | 7.940397 |
| 8.16884 | 7.574398 |
| 6.783542 | 6.777515 |
| 9.182213 | 9.208692 |
| 7.223594 | 7.179049 |
| 9.592328 | 9.297247 |
| 5.839179 | 5.163607 |
| 6.905503 | 6.950794 |
| 5.713503 | 5.588334 |
| 6.460406 | 5.870389 |
| 6.218549 | NaN |
| 6.142793 | 6.353381 |
| 8.632892 | 8.689799 |
| 6.469008 | 6.522561 |
| 8.608565 | 8.820108 |
| 7.683349 | 7.587061 |
| 4.980579 | 4.888181 |
| 6.509585 | 6.41963 |
| 7.552174 | 7.049374 |
| 6.115112 | 4.927457 |
| 7.913281 | 7.915839 |
| 6.162771 | 6.049365 |
| 8.029987 | 7.771126 |
| 6.912373 | 6.872802 |
| 7.20014 | 7.382952 |
| 6.676282 | 6.679698 |
| 7.222245 | 7.321237 |
| 7.060602 | 6.707884 |
| 6.268692 | 5.991018 |
| 5.884079 | 5.666797 |
| 8.054428 | 8.554102 |
| 4.90307 | 4.653329 |
| 5.694898 | 5.880452 |
| 7.888771 | 7.524526 |
| 7.846497 | 7.530067 |
| 6.776337 | 6.756716 |
| 7.506518 | 7.398324 |
| 6.826959 | 7.02304 |

| | |
|---|---|
| 6.143519 | 4.773085 |
| 6.907044 | NaN |
| NaN | NaN |
| 7.191847 | 7.189169 |
| 5.411469 | 5.668936 |
| 7.407154 | 7.51536 |
| 7.310287 | 7.744833 |
| 7.420276 | 7.35409 |
| 7.896356 | 7.847221 |
| 7.29537 | 6.168494 |
| 7.6392 | 7.542863 |
| 6.345861 | 6.241218 |
| 5.784654 | 5.721028 |
| 5.064213 | 5.365528 |
| 7.042393 | 7.036546 |
| 7.36889 | 7.260313 |
| 7.285068 | 7.402786 |
| 7.092431 | 7.148825 |
| 7.308193 | 7.058562 |
| 4.982777 | 4.791912 |
| 7.095567 | 6.922241 |
| 6.254507 | 5.926904 |
| 7.928574 | 7.976641 |
| 5.479451 | 5.436214 |
| 9.686644 | 9.769382 |
| 6.659656 | 6.427706 |
| NaN | NaN |
| 5.495006 | 6.345843 |
| 4.124767 | 4.76001 |
| 7.737689 | 7.616737 |
| NaN | 4.906597 |
| 6.24757 | 6.328346 |
| 7.687354 | 7.712811 |
| 5.056356 | 5.421404 |
| 6.840132 | 6.903368 |
| 5.351373 | 4.925255 |
| 6.115783 | 5.69547 |
| 7.281172 | 7.388308 |
| 5.958273 | 5.928225 |
| 5.127399 | 5.138945 |
| 7.601503 | 8.068789 |
| 6.803514 | 6.711248 |
| 8.446705 | 8.446528 |
| 7.554321 | 8.873736 |
| 7.385756 | 7.498784 |
| 6.26743 | 6.179454 |
| 9.327621 | 9.283865 |

| | |
|---|---|
| 5.43054 | 5.534583 |
| 4.467131 | 5.522517 |
| NaN | NaN |
| 7.06905 | 6.994009 |
| 6.994901 | 7.025106 |
| 6.520933 | 6.371223 |
| 6.718906 | 6.978548 |
| 6.329914 | 6.396626 |
| 7.177472 | 7.286614 |
| 6.832226 | 6.271416 |
| 7.142086 | 7.193307 |
| 7.730364 | 7.832206 |
| 7.921992 | 7.778462 |
| 8.064525 | 8.173767 |
| 7.405183 | 7.135547 |
| 6.099963 | 6.65349 |
| 6.137213 | 6.273894 |
| NaN | NaN |
| 5.591072 | NaN |
| 6.413074 | 6.835352 |
| 6.468505 | 5.890476 |
| 7.960584 | 7.909935 |
| 7.787773 | 7.737947 |
| NaN | NaN |
| 7.227401 | 7.366035 |
| 7.47527 | 7.190539 |
| 4.685453 | NaN |
| 7.387372 | 7.378154 |
| 6.177282 | 6.134428 |
| 7.137097 | 6.664011 |
| 8.041903 | 8.115786 |
| NaN | 5.473375 |
| 5.801228 | 5.413222 |
| 7.040502 | 7.16066 |
| 6.953797 | 6.502877 |
| 8.652992 | 8.512677 |
| 8.024449 | 7.886491 |
| 7.300014 | 7.2805 |
| 6.986547 | 7.147452 |
| 5.431733 | 6.523059 |
| 6.671012 | 6.887061 |
| 4.211147 | 4.402918 |
| 4.874718 | 4.847429 |
| 7.450775 | 6.832777 |
| 6.864447 | 6.769296 |
| 5.991297 | 6.490269 |
| 7.949419 | 7.972353 |

| | |
|---|---|
| 5.930849 | 5.828269 |
| 5.816634 | 5.593251 |
| 7.196224 | 6.942428 |
| 7.322196 | 7.096214 |
| 6.831442 | 7.356323 |
| 10.18761 | 10.40623 |
| 6.291058 | NaN |
| 7.74097 | 7.614322 |
| 5.846239 | 5.534462 |
| 5.838428 | 6.362814 |
| 6.560151 | 6.389534 |
| 6.71033 | 5.983091 |
| 9.151638 | 9.013515 |
| 4.910889 | 5.428491 |
| 6.510606 | 6.66278 |
| 6.272533 | 6.387376 |
| 5.894338 | 5.798367 |
| 5.430161 | 5.507734 |
| 7.368664 | 7.455548 |
| 5.510197 | 4.265159 |
| 8.383415 | 8.265712 |
| 8.539151 | 8.708355 |
| 4.803366 | 4.940049 |
| 6.968836 | 6.92309 |
| 6.892416 | 6.896949 |
| 7.701272 | 7.274101 |
| 6.621089 | 6.586946 |
| 6.163815 | 6.083109 |
| 7.569976 | 7.194511 |
| 7.347189 | 7.768083 |
| 7.105425 | 7.102729 |
| 4.258957 | 3.401211 |
| 5.167672 | NaN |
| 7.564228 | 7.668998 |
| 7.638237 | 7.639008 |
| 7.690901 | 7.678329 |
| 7.448117 | 7.959733 |
| 6.808454 | 6.837569 |
| 6.991323 | 6.620274 |
| 5.93167 | 5.964266 |
| 6.397512 | 6.407061 |
| 4.297777 | 4.531177 |
| 7.237822 | 7.501035 |
| 8.096107 | 8.021778 |
| 5.251818 | 5.326021 |
| 4.725871 | 4.59704 |
| NaN | 9.902918 |

| | |
|---|---|
| 7.09183 | 6.973921 |
| 6.886175 | 7.011382 |
| 6.424205 | 6.474297 |
| 7.333501 | 7.294267 |
| 8.823071 | 8.739116 |
| 9.39291 | 7.73751 |
| 6.876915 | 6.69367 |
| NaN | NaN |
| 7.613385 | 6.912451 |
| NaN | 4.769974 |
| 9.295133 | 9.343038 |
| NaN | 5.703663 |
| 7.210423 | 6.505028 |
| 6.980116 | 6.69779 |
| 6.519515 | 6.48248 |
| 7.160807 | 7.446397 |
| 8.21503 | 7.763779 |
| 5.619478 | 5.671056 |
| 5.888874 | 5.912102 |
| 6.979277 | 6.952004 |
| 8.978702 | 8.471354 |
| 7.195793 | 7.304727 |
| 9.121274 | 9.23724 |
| 6.658874 | 6.533441 |
| 7.745433 | 8.005247 |
| 8.688054 | 8.569239 |
| 6.365431 | 6.012178 |
| 5.531981 | 5.548524 |
| 7.235936 | 7.170821 |
| 7.606473 | 7.74926 |
| NaN | NaN |
| 5.770355 | 6.167259 |
| 6.618589 | 6.674394 |
| 8.317307 | 8.339252 |
| 11.70028 | 11.67099 |
| 7.268432 | 6.920002 |
| 7.651055 | 7.883512 |
| 5.498516 | NaN |
| 8.537875 | 8.557538 |
| 8.131664 | 7.883477 |
| 7.271467 | 7.155259 |
| 9.702829 | 9.710038 |
| 5.391376 | 5.281052 |
| 7.10847 | 6.229805 |
| 6.580179 | 6.580888 |
| 8.080789 | 8.185862 |
| 7.090868 | 7.117312 |

| | |
|---|---|
| 6.644793 | 7.023122 |
| 9.660524 | 9.505667 |
| 7.891747 | 7.462587 |
| 2.958729 | NaN |
| 6.952219 | 6.76783 |
| 5.465408 | 5.258262 |
| 7.056415 | 7.070999 |
| 4.045569 | 4.454224 |
| 7.144771 | 7.242792 |
| 6.124275 | 6.619442 |
| 6.694028 | 6.548532 |
| 7.108311 | 6.931555 |
| 7.575693 | 7.331923 |
| 10.11468 | 9.936021 |
| 4.618527 | 4.819277 |
| 7.406854 | 7.167352 |
| NaN | 8.15103 |
| 7.032794 | 6.819247 |
| 6.934966 | 6.840929 |
| 4.081984 | NaN |
| 6.941325 | 7.056787 |
| 4.600027 | 4.840777 |
| 6.453636 | 6.746178 |
| 5.977252 | 5.85202 |
| 4.494631 | 4.819949 |
| 8.422322 | 8.942908 |
| 7.664104 | 7.964779 |
| 6.372985 | 6.57417 |
| 6.642523 | 6.55037 |
| 7.288785 | 7.004509 |
| 6.374145 | 5.757604 |
| 7.246472 | 7.04092 |
| 7.929024 | 8.140953 |
| 8.509176 | 8.582844 |
| 5.463834 | 5.597801 |
| 5.889799 | 5.658905 |
| 10.5677 | 10.74568 |
| 6.600785 | 6.766062 |
| 6.023377 | 5.466588 |
| 8.392187 | 8.284248 |
| 8.372603 | 8.338784 |
| 7.209002 | 7.058388 |
| 6.528692 | 6.392297 |
| 7.684534 | 6.927967 |
| 7.665284 | 7.77387 |
| 6.359569 | 6.261661 |
| 6.576862 | 6.579995 |

| | |
|---|---|
| 5.552709 | 5.353802 |
| 6.520121 | 6.679564 |
| 5.864162 | 5.931046 |
| 6.59569 | 6.331948 |
| 6.915472 | 6.896584 |
| 4.715227 | 4.916054 |
| 7.751496 | 7.542651 |
| 7.088102 | 6.910178 |
| 6.404246 | 6.449693 |
| 8.129041 | 8.13024 |
| 9.709367 | 10.25406 |
| 6.832889 | 6.943378 |
| 5.920234 | 5.333961 |
| 6.851485 | 5.752464 |
| 5.842415 | 5.577326 |
| 5.649505 | 5.830477 |
| 6.490319 | 6.775858 |
| 6.644237 | 6.795669 |
| 5.222707 | 5.782597 |
| 6.751272 | 7.223598 |
| 6.890409 | 6.832535 |
| 6.389575 | 5.752205 |
| 7.579252 | 7.924319 |
| 4.620549 | 4.969007 |
| 5.295169 | 5.078891 |
| 5.460333 | 5.617079 |
| 6.434773 | 6.951262 |
| 6.386671 | 7.181581 |
| 7.790332 | 7.51204 |
| 6.188822 | 6.201369 |
| 6.613802 | 6.965194 |
| 8.180074 | 7.958597 |
| 6.999375 | 7.32875 |
| 7.978239 | 7.945046 |
| 6.338673 | 6.422495 |
| 10.74775 | 10.66456 |
| 7.096151 | 7.123058 |
| 7.38423 | 7.172153 |
| 6.26028 | 6.06645 |
| 7.994364 | 7.971148 |
| 5.881758 | 5.629953 |
| 7.881394 | 7.439832 |
| 9.543141 | 8.862421 |
| 6.178451 | 6.168884 |
| NaN | 4.589136 |
| 7.61815 | 7.434911 |
| 5.904803 | 6.511287 |

| | |
|---|---|
| 7.419208 | 7.091938 |
| 4.330572 | 4.645206 |
| 8.894161 | 9.236048 |
| 5.795111 | 5.447324 |
| 6.863552 | 6.514962 |
| 7.868954 | 7.817519 |
| 7.827565 | 7.767933 |
| 8.517773 | 8.461217 |
| 3.842346 | 4.42309 |
| 7.044577 | 6.823396 |
| 6.784087 | 6.82091 |
| 7.22616 | 5.583273 |
| 3.979713 | 4.930707 |
| 7.408186 | 7.187463 |
| 6.11599 | 5.822162 |
| 8.359779 | 8.3637 |
| 6.975668 | 7.239559 |
| 8.363605 | 8.148665 |
| 7.082055 | 7.065141 |
| 7.909269 | 7.871845 |
| 8.382708 | 8.390748 |
| 4.505733 | 3.991625 |
| 4.487164 | 4.512729 |
| 9.289854 | 9.279796 |
| 5.218808 | 4.822043 |
| 7.22975 | 6.354967 |
| 10.33333 | 10.28694 |
| 6.939877 | 6.978018 |
| 7.12582 | 6.917828 |
| 5.2666 | 5.781176 |
| 5.155077 | 5.514833 |
| 5.5446 | 4.305999 |
| 7.814587 | 8.05638 |
| 6.797744 | 6.842636 |
| 6.419349 | 6.133346 |
| 8.222188 | 8.443683 |
| 4.767033 | 3.914995 |
| 4.121493 | 6.483944 |
| 7.875269 | 7.368822 |
| 7.376994 | 7.359118 |
| 6.096571 | 6.405159 |
| 6.711788 | 6.425911 |
| 4.464992 | 4.637337 |
| 5.553787 | NaN |
| 5.251952 | 4.799104 |
| 7.818416 | 7.739866 |
| 8.479447 | 8.361047 |

| | |
|---|---|
| 5.451758 | 5.321654 |
| 2.96988 | 3.046763 |
| 7.373428 | 7.23839 |
| 8.066894 | 7.885504 |
| 8.500237 | 8.035892 |
| 6.521558 | 6.529078 |
| 8.90748 | 7.724314 |
| 5.333229 | 5.186247 |
| 5.135112 | 5.073416 |
| 8.630969 | 8.58986 |
| 6.957873 | 7.111587 |
| 6.327156 | 5.425732 |
| 6.336794 | 6.337669 |
| 5.024498 | 5.123451 |
| 5.863726 | 6.147186 |
| 7.499997 | 7.465766 |
| 6.321651 | 6.451775 |
| 7.857761 | 7.833488 |
| NaN | 5.95021 |
| 6.753496 | 6.838875 |
| 6.717899 | 6.899086 |
| 7.324487 | 7.439266 |
| 8.698642 | 8.577888 |
| 6.28043 | 6.207769 |
| 7.717168 | 7.617777 |
| NaN | NaN |
| 7.008751 | 6.873722 |
| 5.697473 | 4.905626 |
| 6.547102 | 5.959908 |
| 8.001779 | 7.878051 |
| 7.844584 | 7.588858 |
| 10.46342 | 10.45483 |
| 6.164463 | 6.964073 |
| NaN | 5.756667 |
| 8.299932 | 8.471168 |
| 7.625443 | 7.599796 |
| 7.02415 | 7.260669 |
| 7.125739 | 7.460348 |
| 8.441937 | 8.592607 |
| 8.90002 | 9.088081 |
| 5.798298 | 5.891322 |
| 7.137463 | 7.223428 |
| 7.959787 | 7.613651 |
| 9.720508 | 9.768926 |
| 6.626882 | 6.211599 |
| 7.180254 | 6.984542 |
| 6.933433 | 6.746128 |

| | |
|---|---|
| 6.74873 | 6.626696 |
| 10.29513 | 10.00159 |
| 8.058289 | 7.686718 |
| 7.467581 | 7.087391 |
| NaN | NaN |
| 7.169161 | 6.995677 |
| 7.68547 | 7.622393 |
| 5.403012 | 6.024287 |
| 6.307904 | 7.706385 |
| 5.730783 | 5.735199 |
| NaN | NaN |
| 7.118915 | 6.4931 |
| 7.898208 | 7.005495 |
| 5.313359 | 6.121008 |
| 4.786613 | 5.946822 |
| 9.516908 | 9.582957 |
| 5.673942 | 5.7858 |
| 7.169739 | 7.311294 |
| 7.186893 | 7.153802 |
| 7.876036 | 8.014164 |
| 5.473886 | 5.254704 |
| 7.452794 | 7.560774 |
| 5.685142 | 5.665943 |
| 9.528608 | 9.517298 |
| 8.627573 | 9.233335 |
| 8.219063 | 8.160849 |
| 6.918759 | 7.504086 |
| 6.53909 | 6.611009 |
| 7.626818 | 8.057057 |
| 6.1374 | 6.19417 |
| 7.679135 | 7.396255 |
| 5.541958 | 5.786543 |
| NaN | NaN |
| 3.287279 | 3.452362 |
| 10.36942 | 10.2166 |
| 6.059743 | 5.842861 |
| 6.805044 | 6.558405 |
| 9.362378 | 9.651572 |
| 5.046482 | 4.90146 |
| 6.013624 | 5.720645 |
| 9.089496 | 9.447226 |
| 7.437157 | 6.630066 |
| 5.156128 | 5.106193 |
| 5.823193 | 6.332008 |
| 7.142518 | 6.121795 |
| 5.670375 | 5.785046 |
| 6.294613 | 6.134523 |

| | |
|---|---|
| 4.219407 | 4.032806 |
| 9.389829 | 9.48 |
| 7.596225 | 7.373501 |
| 6.096507 | 5.835359 |
| 6.991134 | 7.808675 |
| 6.936816 | 7.182522 |
| 7.208126 | 7.404205 |
| 6.382859 | 6.725895 |
| 6.519505 | 6.558554 |
| 5.767037 | 5.969121 |
| 7.768197 | 7.739775 |
| 8.817578 | 8.638099 |
| 8.772769 | 8.784448 |
| 4.896222 | 4.558256 |
| 8.619676 | 8.62314 |
| 8.955067 | 9.130929 |
| 10.92474 | 10.69352 |
| 6.455211 | 6.152322 |
| 4.594037 | 4.724895 |
| 6.070467 | 5.952752 |
| 5.879124 | 5.618967 |
| 7.404809 | 7.231409 |
| 5.913641 | 5.690533 |
| 6.489504 | 6.595531 |
| 6.707043 | 6.696196 |
| 6.48704 | 6.158089 |
| 8.333388 | 8.191899 |
| 4.836926 | 4.49823 |
| NaN | 6.015285 |
| 7.516833 | 7.06939 |
| 7.061668 | 6.948653 |
| 7.044295 | 6.94984 |
| 7.951658 | 7.708751 |
| 8.744945 | 8.762995 |
| 8.325307 | 8.051188 |
| 8.091254 | 8.202037 |
| 7.369378 | 7.343685 |
| 7.651017 | 7.782715 |
| 10.26474 | 10.20691 |
| 8.448331 | 8.377722 |
| 5.742463 | 6.179089 |
| 6.133048 | 5.270611 |
| 9.74579 | 9.03478 |
| 6.099312 | 6.302064 |
| 5.514692 | NaN |
| 7.577457 | 7.278196 |
| 7.459271 | 7.59916 |

| | |
|---|---|
| 5.990642 | 5.715692 |
| 9.305617 | 9.196764 |
| 8.155099 | 7.794427 |
| 1.406861 | 1.499534 |
| 10.36351 | 10.31689 |
| 8.456052 | 8.289068 |
| 6.045886 | 5.327307 |
| 7.822322 | 7.802752 |
| NaN | 5.920996 |
| 4.126256 | 5.659808 |
| 8.065779 | 8.359674 |
| 9.421566 | 9.461916 |
| 8.390841 | 8.488138 |
| 6.621195 | 6.090952 |
| 6.082301 | 6.161778 |
| 8.272534 | 8.030888 |
| 5.254646 | 5.010841 |
| 7.077155 | 6.720448 |
| 6.657348 | 6.947581 |
| 8.098401 | 8.322682 |
| 6.440314 | 6.177706 |
| 6.770602 | 6.830389 |
| 9.217363 | 9.571532 |
| 6.002856 | 5.69514 |
| 8.481236 | 8.514463 |
| 7.396324 | 7.156813 |
| 5.623521 | 6.131688 |
| 8.989976 | 8.619376 |
| 9.511732 | 9.397927 |
| 7.065533 | 7.029984 |
| 7.050305 | 6.939935 |
| 9.025918 | 8.876896 |
| 7.817083 | 7.742504 |
| 6.513445 | 5.720789 |
| 5.824658 | 5.927879 |
| 8.099771 | 8.08793 |
| 6.396869 | 5.541103 |
| 6.469155 | 6.306623 |
| NaN | 6.62723 |
| 8.977494 | 8.96177 |
| 6.482137 | 6.376086 |
| 8.040778 | 8.165831 |
| 6.116727 | 5.981636 |
| 5.381298 | 5.380773 |
| 7.015953 | 6.975595 |
| NaN | 5.145811 |
| 7.189894 | 7.426023 |

| | |
|---|---|
| 6.91503 | 7.385488 |
| 5.951411 | 5.977415 |
| 8.065586 | 7.901967 |
| 6.331329 | 6.638568 |
| 6.910352 | 6.780808 |
| 8.248734 | 8.358205 |
| 6.322935 | 5.882075 |
| 7.915518 | 8.258492 |
| 6.034949 | NaN |
| 5.290687 | NaN |
| 7.015644 | 6.841295 |
| 6.706399 | NaN |
| 4.704886 | 4.925496 |
| 8.402626 | 7.996659 |
| 6.32685 | 6.364542 |
| 8.336663 | 8.679413 |
| 7.957531 | 7.996283 |
| 10.76951 | 10.66059 |
| NaN | NaN |
| 5.829926 | 5.4386 |
| 8.001814 | 7.67588 |
| 7.794755 | 7.686991 |
| 7.890687 | 7.797357 |
| 7.541635 | 7.469321 |
| 9.293879 | 9.280872 |
| 5.329338 | 5.327064 |
| 7.293055 | 7.390058 |
| 8.882051 | 8.542258 |
| 8.192662 | 8.375425 |
| 5.335385 | 5.999294 |
| 6.237177 | 6.253706 |
| 5.984695 | 5.947407 |
| 7.025239 | 6.720117 |
| 8.051014 | 8.250865 |
| 5.407478 | 5.300761 |
| 7.041136 | 7.297705 |
| 7.140821 | 6.520535 |
| 7.285148 | 7.37254 |
| 6.567235 | 6.593032 |
| 6.733437 | 6.008534 |
| 6.318114 | 5.580281 |
| 5.171528 | 5.631292 |
| 7.020699 | 7.231188 |
| 6.741516 | 7.037679 |
| 5.834583 | 5.932139 |
| 5.585365 | 5.85061 |
| 8.407714 | 8.196986 |

-Bruker Confidential-

| | |
|---|---|
| 6.711206 | 6.692947 |
| 6.104462 | 6.042114 |
| 5.541309 | 3.884752 |
| 2.285815 | 4.978139 |
| 9.547606 | 10.32667 |
| 5.531077 | 5.625125 |
| 6.695146 | 7.548399 |
| 6.604122 | 4.362216 |
| 6.467522 | 5.290643 |
| 6.886749 | 7.288883 |
| 5.065891 | 4.965143 |
| 6.04772 | 6.223641 |
| 6.266355 | 6.112566 |
| 6.750573 | 6.582579 |
| 5.773724 | 5.414109 |
| 7.241498 | 7.597834 |
| 7.546976 | 7.72445 |
| 7.253485 | 6.800507 |
| 5.285858 | 5.652539 |
| 6.156205 | 6.665586 |
| 7.55772 | 7.438744 |
| 7.144059 | 7.498054 |
| 7.52789 | 7.466675 |
| 5.280118 | 5.614266 |
| 7.512581 | 7.315311 |
| 6.507336 | 6.520962 |
| 7.225549 | 6.754886 |
| 8.84372 | 6.268384 |
| 5.199522 | 6.04685 |
| 5.819854 | 6.607133 |
| 5.581544 | 5.902886 |
| 8.647955 | 8.344236 |
| 6.938805 | 6.921412 |
| 7.539352 | 7.798162 |
| 6.225614 | 6.383149 |
| 8.634914 | 8.620368 |
| 7.547915 | 7.844211 |
| 10.45047 | 10.54628 |
| 4.763774 | 4.931976 |
| 7.116116 | 7.047174 |
| 6.581523 | 6.454975 |
| NaN | 4.551258 |
| 5.89928 | 5.893528 |
| 8.109157 | 7.976569 |
| 6.033872 | 5.848149 |
| 7.445116 | 7.463545 |
| 8.309524 | 8.102601 |

| | |
|---|---|
| 7.375358 | 7.433828 |
| 6.814336 | 6.860808 |
| 6.203729 | 6.186367 |
| 5.526055 | 5.546292 |
| 6.229943 | 4.721025 |
| 7.605818 | 6.443855 |
| NaN | 4.808068 |
| 7.689033 | 7.59924 |
| 4.759912 | 4.855608 |
| 6.390255 | 6.468093 |
| 5.315505 | 5.610829 |
| 6.561625 | 6.831884 |
| 7.891162 | 7.122298 |
| 6.039789 | 6.399968 |
| 6.498542 | 6.314178 |
| 7.103626 | 7.11714 |
| 6.672629 | 6.56599 |
| 7.154666 | 7.144172 |
| 4.465966 | 4.073269 |
| 6.974161 | 6.797754 |
| 6.50912 | 7.191952 |
| 6.346607 | 6.772157 |
| 9.391706 | 9.273866 |
| 8.059553 | 8.114012 |
| 9.310065 | 9.172772 |
| 3.995669 | 5.45855 |
| 7.652867 | 7.233732 |
| 6.855449 | 6.54525 |
| 4.758685 | NaN |
| 5.662805 | 5.659489 |
| 7.547544 | 7.671988 |
| 7.730962 | 8.031483 |
| 6.870799 | 6.644026 |
| 7.355084 | 7.385444 |
| 7.344707 | 7.392903 |
| NaN | 6.803312 |
| NaN | 5.880653 |
| 9.610625 | 9.611542 |
| 6.567761 | 6.94231 |
| 6.615903 | 6.375499 |
| 8.338572 | 8.266307 |
| 7.966918 | 7.904538 |
| 8.64881 | 8.73572 |
| NaN | 5.68888 |
| 6.645641 | 6.278361 |
| 7.061615 | 6.926694 |
| 5.998098 | 6.430412 |

| | |
|---|---|
| 7.62963 | 7.018906 |
| 5.124957 | 4.767426 |
| 8.982631 | 8.873796 |
| 5.351094 | 5.51457 |
| 5.798379 | 5.734372 |
| 6.382544 | 6.665677 |
| 5.654212 | 6.070195 |
| 7.414716 | 7.101606 |
| 7.404937 | 7.311894 |
| NaN | NaN |
| 7.135185 | 6.9795 |
| 12.25547 | 12.07079 |
| 6.776646 | 6.641534 |
| 6.278585 | 5.23361 |
| 8.279435 | 8.174006 |
| 7.590309 | 5.501954 |
| 7.588288 | 7.36893 |
| 8.729032 | 8.657484 |
| 7.549313 | 7.239527 |
| 6.243719 | 6.018272 |
| 4.779562 | 4.766927 |
| 5.579936 | 4.223572 |
| 7.24385 | 6.954178 |
| 7.619526 | 7.485044 |
| 7.423922 | 7.381451 |
| 7.293615 | 7.383529 |
| 7.652263 | 7.757871 |
| 6.451752 | 6.537137 |
| 5.637733 | 6.085897 |
| 6.650795 | 6.204634 |
| 6.588791 | 6.218959 |
| 3.028173 | 3.360691 |
| 7.51001 | 7.681108 |
| 6.181205 | 5.931341 |
| 5.944982 | 5.894333 |
| 7.828317 | 7.962823 |
| 6.059789 | 5.668931 |
| 7.096087 | 7.277726 |
| 8.091605 | 8.009827 |
| 7.281016 | 7.136588 |
| 10.16153 | 10.27073 |
| 7.233698 | 6.535545 |
| 7.308692 | 7.413947 |
| 8.735214 | 8.93983 |
| 7.383448 | 7.421251 |
| 4.869198 | 4.687 |
| 6.009212 | 6.188932 |

| | |
|---|---|
| 6.972326 | 7.005584 |
| 5.923574 | 5.958678 |
| 10.20809 | 10.07367 |
| 6.348446 | 6.312141 |
| 10.91618 | 10.68951 |
| 6.647644 | 6.439599 |
| 6.570013 | 6.122933 |
| 6.622526 | 6.682039 |
| 4.822295 | 4.676614 |
| 6.019573 | 5.672382 |
| 6.185463 | 5.78027 |
| 6.328824 | 6.190849 |
| 6.496036 | 7.105551 |
| 6.211658 | 6.699643 |
| 7.627755 | 7.550676 |
| 9.732459 | 8.956166 |
| 5.195231 | 5.07102 |
| 6.339119 | 5.607221 |
| 6.685872 | 6.237213 |
| 8.864828 | 8.948197 |
| 6.041326 | 5.176742 |
| 5.736982 | 5.564952 |
| 6.62256 | 6.384429 |
| 4.777335 | 4.208365 |
| 5.46576 | 5.405725 |
| NaN | 5.013877 |
| 5.641691 | 5.406949 |
| 5.736706 | 5.382749 |
| 6.912213 | 6.951689 |
| 8.420427 | 8.383948 |
| 7.885808 | 7.822923 |
| 7.032848 | 7.073548 |
| 5.940912 | 5.927151 |
| 6.311024 | 6.116707 |
| 7.530253 | 7.388751 |
| 6.792663 | 6.784977 |
| 6.671346 | 6.811424 |
| 8.086138 | 8.185148 |
| 8.346391 | 8.059768 |
| 5.324557 | 5.083708 |
| 4.704244 | 5.240506 |
| 6.504176 | 6.495369 |
| 6.650976 | 6.606308 |
| 6.096958 | 6.330879 |
| 6.85076 | 7.240698 |
| 9.143033 | 9.243722 |
| 7.41898 | 7.833615 |

| | |
|---|---|
| 9.776822 | 9.62107 |
| 6.787912 | 6.56527 |
| 6.776066 | 7.105202 |
| 4.94519 | 5.739498 |
| 6.984944 | 6.89203 |
| 5.005636 | 5.202466 |
| 7.422313 | 7.206577 |
| 4.857677 | 4.528522 |
| 7.849347 | 7.520705 |
| 7.903011 | 7.784017 |
| 9.72071 | 9.700043 |
| 6.257765 | 6.207993 |
| 6.926158 | 7.298798 |
| 5.975597 | 5.922592 |
| 4.379673 | 4.819079 |
| 7.4334 | 7.390267 |
| 7.792589 | 8.339755 |
| 6.882088 | 7.119581 |
| 7.733712 | 7.437467 |
| 9.148525 | 8.977169 |
| 8.870428 | 8.863733 |
| 6.943645 | 6.799047 |
| 7.855969 | 8.130642 |
| 5.615491 | 5.683351 |
| 5.583634 | 5.653158 |
| 9.43034 | 9.348971 |
| 9.050297 | 9.10053 |
| 5.471833 | 5.63003 |
| 7.52796 | 7.584345 |
| 6.537171 | 6.757838 |
| 7.615133 | 7.696406 |
| 8.901141 | 8.880738 |
| 6.121112 | 6.185416 |
| 6.239409 | 6.378856 |
| 6.583697 | 6.930366 |
| 7.878231 | 7.591915 |
| 7.398668 | 7.623438 |
| 4.908929 | 5.47434 |
| 7.526114 | 7.556088 |
| 6.714441 | 6.766834 |
| 7.895291 | 7.753764 |
| 5.755295 | 5.568635 |
| 6.266861 | 5.575427 |
| 9.642914 | 9.841514 |
| 8.510945 | 8.181368 |
| 6.940353 | 6.915696 |
| 8.834397 | 9.067599 |

| | |
|---|---|
| 6.630391 | NaN |
| 9.669827 | 9.746785 |
| 9.967715 | 9.72609 |
| 4.952112 | 5.658771 |
| 12.09409 | 12.18177 |
| 7.941532 | 8.083987 |
| 8.66399 | 8.42456 |
| 8.191335 | 8.358862 |
| 5.709957 | 5.793869 |
| 7.822104 | 7.781654 |
| 7.3981 | 7.107178 |
| 6.616867 | 6.006804 |
| 5.305394 | 5.756227 |
| 7.751299 | 7.744277 |
| 8.357589 | 8.339779 |
| 5.087442 | 4.422306 |
| 5.742866 | 6.023761 |
| 5.288785 | 5.799736 |
| 7.735505 | 7.729348 |
| 6.609245 | 6.653689 |
| 7.803327 | 7.384904 |
| 8.038731 | 7.81981 |
| 2.835316 | 5.072994 |
| 7.138076 | 6.869102 |
| 8.078353 | 7.852524 |
| 9.168976 | 8.959936 |
| 6.667648 | 6.667867 |
| 6.580192 | 6.773502 |
| 8.017492 | 7.870003 |
| 3.419762 | 3.438759 |
| 5.080525 | 5.38877 |
| 7.061284 | 7.020338 |
| 5.076316 | 5.102958 |
| 4.157791 | NaN |
| 5.810981 | 5.803916 |
| 6.310484 | 6.187716 |
| 7.114824 | 7.134457 |
| 7.50435 | 7.652515 |
| 6.287609 | 6.35304 |
| 7.986254 | 7.841354 |
| 6.957635 | 6.685796 |
| NaN | 4.280349 |
| 6.981072 | 6.922042 |
| 5.9722 | 6.432695 |
| 4.932019 | 5.399824 |
| 7.410589 | 8.156757 |
| 8.964966 | 8.806187 |

| | |
|---|---|
| 7.733431 | 8.079251 |
| 6.99612 | 7.293046 |
| 8.653594 | 8.484066 |
| 5.244712 | 5.432308 |
| 8.290851 | 7.965205 |
| 5.699894 | 6.086274 |
| 7.414555 | 7.573335 |
| 7.467197 | NaN |
| 9.053437 | 8.992814 |
| 5.78914 | 5.830718 |
| 7.817642 | 7.732139 |
| 5.62668 | 5.291456 |
| 8.675493 | 8.685023 |
| 8.550482 | 8.293406 |
| 7.415375 | 7.954328 |
| NaN | NaN |
| 11.36664 | 11.23536 |
| 7.867436 | 7.985213 |
| 6.383499 | 5.955009 |
| 8.228207 | 8.073737 |
| 6.595095 | 6.648208 |
| 6.696972 | 6.622733 |
| 8.714917 | 8.690318 |
| 5.634049 | 5.58115 |
| 7.112258 | 6.923205 |
| 5.294301 | 5.267315 |
| 7.784185 | 8.005181 |
| 5.241905 | 5.794046 |
| 10.12541 | 10.21659 |
| 8.677866 | 8.639475 |
| 7.383206 | 7.383546 |
| 7.14828 | 7.08354 |
| 7.425263 | 7.356944 |
| 7.928449 | 7.918418 |
| 7.146081 | 7.102686 |
| 7.814683 | 8.171961 |
| 10.43865 | 10.12775 |
| 7.577832 | 7.502646 |
| 8.740112 | 8.688825 |
| 7.497614 | 7.425455 |
| 6.078656 | 6.338546 |
| 7.701857 | 6.765615 |
| 8.775917 | 8.717504 |
| 7.367188 | 7.020542 |
| 7.499022 | 7.658012 |
| 5.2263 | 5.228287 |
| 6.467987 | 6.340564 |

| | |
|---|---|
| 7.737531 | 7.676693 |
| NaN | NaN |
| 10.45275 | 10.6216 |
| 7.657755 | 7.972283 |
| 6.336825 | 5.759929 |
| 7.134265 | 7.100906 |
| 7.369842 | 7.458497 |
| 6.595604 | 6.503428 |
| 7.428464 | 7.207114 |
| 7.568084 | 7.679093 |
| 6.43332 | 6.283502 |
| 8.268139 | 8.092244 |
| 5.887501 | 5.847874 |
| 7.465971 | 7.197692 |
| NaN | NaN |
| 6.36438 | 5.442759 |
| 4.973229 | 5.586326 |
| 7.145872 | 7.628762 |
| 7.47443 | 7.32022 |
| 7.358506 | 7.944342 |
| NaN | NaN |
| 8.794217 | 8.662704 |
| 7.870715 | 7.841992 |
| 6.309222 | 6.307352 |
| 7.068076 | 6.930746 |
| 7.912679 | 7.930643 |
| 7.082147 | 6.82657 |
| 7.40612 | 6.764457 |
| 6.29004 | 6.382924 |
| 6.800688 | 6.993613 |
| 5.117629 | 5.502927 |
| NaN | 6.413799 |
| 6.665599 | 6.576231 |
| 6.370099 | 6.537858 |
| 11.50272 | 11.45608 |
| 7.424496 | 7.351022 |
| 7.672779 | 7.436211 |
| 6.77633 | 6.971748 |
| 6.543839 | 6.686414 |
| NaN | 3.312588 |
| 7.148457 | 6.737241 |
| 6.882501 | 6.87896 |
| 7.958858 | 8.342501 |
| 6.377394 | 7.007135 |
| 10.04287 | 10.24512 |
| NaN | NaN |
| 12.24461 | 12.44179 |

| | |
|---|---|
| 6.585167 | 6.567801 |
| 7.018274 | 6.947571 |
| 8.947522 | NaN |
| 8.174315 | 7.695471 |
| 5.809058 | 5.922506 |
| 6.468625 | 5.197634 |
| 7.376268 | 7.180862 |
| 7.813801 | 7.877322 |
| 5.928869 | 5.78289 |
| 6.667046 | 6.64844 |
| 9.112578 | 9.087376 |
| 8.468101 | 8.386544 |
| 8.08192 | 7.970445 |
| 5.858281 | 5.936123 |
| 5.076557 | 4.577603 |
| 6.359707 | 5.760802 |
| 8.372738 | 8.180811 |
| 6.917751 | 7.12583 |
| NaN | NaN |
| 6.663975 | 6.144377 |
| 7.910121 | 7.696796 |
| 7.03941 | 7.123863 |
| 7.914164 | 7.442614 |
| 4.329592 | 4.167359 |
| 7.310033 | 7.477185 |
| 7.542067 | 7.594727 |
| 8.740145 | 8.510499 |
| 7.312678 | 7.458564 |
| 7.40408 | 7.308606 |
| 6.855412 | 6.731771 |
| 6.994995 | 6.597781 |
| 6.40882 | 6.503697 |
| 7.514423 | 7.639951 |
| 6.556468 | 6.444543 |
| 5.906131 | 6.060486 |
| 7.468645 | 7.455629 |
| 6.012547 | 5.475272 |
| 5.86296 | 6.173941 |
| 8.849547 | 8.823092 |
| 7.095829 | 6.102335 |
| 6.54353 | 6.341121 |
| 6.124674 | NaN |
| 5.979786 | 4.691166 |
| 9.340409 | NaN |
| 6.36881 | 5.804501 |
| 5.941482 | 5.348438 |
| 7.82583 | 7.822986 |

-Bruker Confidential-

| | |
|---|---|
| 6.808414 | 6.887738 |
| 7.448539 | 7.502434 |
| 6.480025 | 6.192055 |
| 5.865472 | 6.458317 |
| 6.09204 | 6.599709 |
| 8.478377 | 8.522882 |
| 6.449186 | 6.800575 |
| 7.292954 | 7.157141 |
| 5.687432 | 5.152981 |
| 6.707342 | 6.551763 |
| 6.751788 | 6.642698 |
| 5.129739 | 5.804204 |
| 7.841985 | 7.74899 |
| 6.459962 | 6.889346 |
| 8.225986 | 8.226018 |
| 7.09093 | 6.98027 |
| 5.195304 | 4.776068 |
| 6.265336 | 6.203505 |
| 8.144825 | 7.936056 |
| 8.064611 | 8.04168 |
| 6.812303 | 5.872785 |
| 5.459931 | 5.254614 |
| 7.771041 | 7.74169 |
| 5.694474 | 5.750381 |
| 7.519089 | 7.321997 |
| 8.184506 | 7.994958 |
| 8.064555 | 8.005175 |
| 6.209473 | 6.0559 |
| 7.192281 | 6.8733 |
| 7.02326 | 7.127646 |
| 8.180692 | 8.135291 |
| 5.624843 | 5.798021 |
| 6.7758 | 7.458344 |
| 7.213518 | 7.524411 |
| 8.219358 | 7.902651 |
| 9.559844 | 9.522449 |
| 6.499374 | 6.547648 |
| 6.966561 | 6.737384 |
| 6.646921 | 6.549977 |
| 8.578024 | 8.374625 |
| 6.299812 | 6.505961 |
| 6.224321 | 6.227657 |
| 8.523446 | 8.665115 |
| 5.456644 | 5.393332 |
| 5.506892 | 5.915672 |
| 6.693471 | NaN |
| 7.182587 | 6.861898 |

| | |
|---|---|
| 8.935278 | 8.929463 |
| 5.350836 | 5.851872 |
| 7.887708 | 8.224268 |
| 8.025495 | 7.95143 |
| 7.960381 | 8.224638 |
| 5.14085 | 5.640432 |
| 8.200584 | 8.292294 |
| 7.444002 | 7.387077 |
| 6.403401 | 6.508011 |
| 8.108564 | 7.945208 |
| 8.035682 | 8.06985 |
| 8.055614 | 8.085451 |
| 7.381666 | 7.536999 |
| 6.308867 | 6.181393 |
| 9.668417 | 9.650826 |
| 7.330678 | 7.263057 |
| 8.098662 | 8.147563 |
| 6.606771 | 7.008783 |
| 7.125552 | 7.092729 |
| 9.378695 | 9.403355 |
| 4.633095 | 4.908366 |
| 6.568976 | 6.57177 |
| 6.908065 | 7.004829 |
| 4.585553 | 4.24114 |
| 9.024404 | 9.011057 |
| 4.914236 | 5.056587 |
| 8.226922 | 8.386967 |
| 6.321322 | 6.213317 |
| 7.517183 | 6.868584 |
| 7.196694 | 7.609871 |
| 7.867299 | 7.600112 |
| 5.182504 | 5.357592 |
| 6.743374 | 6.787714 |
| 5.268297 | 4.68739 |
| 6.661468 | 6.416354 |
| 5.442914 | 5.435328 |
| 5.703406 | 6.457598 |
| 4.230247 | 3.915621 |
| 7.910411 | 7.609006 |
| 8.59504 | 8.600455 |
| 6.184855 | 6.479696 |
| 6.665808 | 6.415624 |
| 8.233357 | 8.256942 |
| 4.069772 | 4.141672 |
| 5.865421 | 5.136655 |
| 6.011993 | 6.196937 |
| 5.78373 | 6.059669 |

| | |
|---|---|
| 7.410381 | 7.109684 |
| 7.350234 | 7.121623 |
| 5.718194 | 5.295324 |
| 8.592152 | 8.460394 |
| 5.709944 | 5.693367 |
| 6.196106 | 6.483384 |
| 6.978503 | 7.070702 |
| 7.469907 | 7.040278 |
| 8.224288 | 8.216516 |
| 6.08592 | 6.285502 |
| 10.63139 | 10.58765 |
| 6.593479 | 6.511512 |
| 9.860097 | 9.732638 |
| 7.05178 | 7.281684 |
| 4.416704 | 4.179071 |
| 8.813207 | 8.765973 |
| 4.95838 | 5.387625 |
| 6.12244 | 6.742734 |
| 6.52589 | 6.535697 |
| 8.715754 | 8.578222 |
| 7.361747 | 7.329527 |
| 6.898151 | 7.431561 |
| 6.631579 | 6.645642 |
| NaN | NaN |
| 6.874116 | 6.855383 |
| 10.07997 | 10.0437 |
| 10.12821 | 10.12172 |
| NaN | 4.875388 |
| 6.610152 | NaN |
| 6.079014 | 6.298828 |
| 7.224669 | 7.498971 |
| 0.576284 | 1.865809 |
| 7.480268 | 7.373801 |
| 6.64003 | 6.751606 |
| 7.936218 | NaN |
| 5.186346 | 5.939532 |
| 6.18556 | 5.570271 |
| 8.095433 | 8.094813 |
| 7.941965 | 7.777799 |
| 10.48954 | 10.45256 |
| 9.47863 | 9.380876 |
| 7.885962 | 7.900742 |
| 11.35285 | 11.176 |
| 8.447594 | 7.676207 |
| 8.032488 | 7.884144 |
| 9.668111 | 9.846719 |
| 8.749676 | 8.453601 |

-Bruker Confidential-

| | |
|---|---|
| NaN | NaN |
| 8.196479 | 8.131092 |
| 8.004504 | 8.087568 |
| 8.342582 | 7.975067 |
| 9.050927 | 9.10241 |
| 10.65859 | 10.15597 |
| 5.198248 | 5.059157 |
| 5.908699 | 6.071096 |
| 5.928789 | 6.127599 |
| 7.093405 | 6.700218 |
| 7.087429 | 7.131691 |
| 6.177977 | 5.788679 |
| 6.833748 | 6.886189 |
| 5.290744 | 5.563194 |
| 8.543411 | 8.314974 |
| 5.744181 | 5.698009 |
| 7.717707 | 7.457263 |
| 7.181436 | 7.019526 |
| 6.361429 | 6.482678 |
| 8.179744 | 8.135061 |
| 6.438836 | 6.709172 |
| 6.38814 | 6.10801 |
| 6.54589 | 6.382844 |
| 5.249675 | 5.754454 |
| 9.328953 | 9.435261 |
| NaN | NaN |
| 10.07134 | 10.13062 |
| 5.680311 | 5.443571 |
| 6.198932 | 6.179699 |
| 7.264086 | 7.414574 |
| 7.346694 | 7.226213 |
| 5.781516 | 5.605396 |
| 7.173807 | 6.68022 |
| 5.394011 | 4.860883 |
| 7.458786 | 7.502687 |
| 6.572592 | 6.922123 |
| 8.084199 | 7.781939 |
| 7.394268 | 7.15684 |
| 6.743578 | 6.520681 |
| 7.48356 | 7.039194 |
| 7.388811 | 7.579901 |
| 7.441993 | 7.328214 |
| 7.652568 | 7.66539 |
| 7.882131 | 7.561734 |
| 5.735531 | 5.669079 |
| 7.480566 | 7.654291 |
| 10.06935 | 9.798266 |

-Bruker Confidential-

| | |
|---|---|
| 5.272202 | 5.886258 |
| 9.781861 | 9.562523 |
| 7.768859 | 7.701666 |
| 5.472211 | 5.479811 |
| 5.802433 | 6.203049 |
| 8.676043 | 8.588746 |
| 8.816975 | 8.574502 |
| 5.744634 | 5.643903 |
| 8.733791 | 8.649051 |
| 6.774197 | 6.994324 |
| 9.134836 | 9.060927 |
| 7.6803 | 7.302905 |
| 6.912745 | 4.428109 |
| 6.667293 | 7.287708 |
| 9.036672 | 9.053275 |
| 5.716004 | 5.526435 |
| 7.32372 | 7.138678 |
| 7.19699 | 7.182472 |
| 6.512169 | 6.665787 |
| 7.077355 | 6.677404 |
| NaN | 6.058208 |
| 9.284788 | 9.348035 |
| 7.466351 | 7.205082 |
| 5.71593 | 5.361022 |
| 7.235745 | 7.288646 |
| 6.711595 | 6.886814 |
| 9.353389 | 9.493488 |
| 9.898638 | 9.755779 |
| 8.92517 | 8.885934 |
| 5.996494 | 5.604717 |
| 8.535906 | 8.547724 |
| 6.057487 | 6.059098 |
| 7.527092 | 7.001831 |
| 8.099602 | 7.387219 |
| 7.778501 | 7.498517 |
| 9.406684 | 9.321758 |
| 6.83986 | 6.079529 |
| 8.461221 | 8.667964 |
| 7.495904 | 7.474082 |
| 7.371871 | 7.232414 |
| 6.15433 | 6.197917 |
| 6.165005 | 5.58694 |
| 5.167427 | 4.999466 |
| 7.503981 | 7.075081 |
| 6.299065 | 6.031029 |
| 6.898256 | 6.397933 |
| 9.30617 | 9.061818 |

| | |
|---|---|
| 6.466905 | 6.216751 |
| 8.466644 | 8.148138 |
| 7.401957 | 5.7463 |
| NaN | NaN |
| 9.443299 | 9.18122 |
| 6.989953 | 6.876791 |
| 6.462981 | 6.89136 |
| 6.336065 | 6.748375 |
| 9.887982 | 10.29076 |
| 7.65708 | 5.971092 |
| 5.856822 | 5.405239 |
| 7.956115 | 8.026889 |
| 9.338338 | 9.124727 |
| 9.55727 | 9.534485 |
| 6.258876 | 6.506444 |
| 5.903481 | 5.887134 |
| 6.018047 | 6.352005 |
| 7.099821 | 7.413614 |
| 7.856375 | 7.030826 |
| 5.690828 | 5.711997 |
| 7.080971 | 6.59074 |
| NaN | 7.607102 |
| 5.72585 | 5.734274 |
| 6.808141 | 6.960351 |
| 3.941666 | 4.400778 |
| 7.429568 | 7.688708 |
| 6.338222 | 6.170768 |
| 9.121684 | 8.911407 |
| 3.983044 | 3.978143 |
| 7.115736 | 6.858757 |
| 7.255479 | 7.27712 |
| 5.220482 | 5.07243 |
| 7.580209 | 7.595413 |
| NaN | NaN |
| 7.009213 | 7.448163 |
| 8.450809 | 8.125055 |
| 7.438593 | 7.21621 |
| 7.89458 | 7.861723 |
| 6.819392 | 6.928009 |
| 5.490812 | 5.934656 |
| 6.933311 | 6.895679 |
| NaN | 4.434308 |
| 6.921058 | 6.913005 |
| 6.969277 | 6.83456 |
| 7.849999 | 7.78494 |
| 7.041517 | 7.01433 |
| 5.866576 | 5.879757 |

| | |
|---|---|
| 7.131309 | 6.939425 |
| 6.216465 | 6.590051 |
| 7.443341 | 7.669559 |
| 7.108247 | 7.281106 |
| 6.461442 | 6.493979 |
| 6.566028 | 6.405089 |
| 6.59776 | 6.71023 |
| 10.19765 | 10.18605 |
| 5.869134 | 6.231487 |
| 7.749434 | 7.384803 |
| 6.098359 | 6.116902 |
| 12.09167 | 12.14195 |
| 4.768784 | 5.48628 |
| 7.666357 | 7.616347 |
| 7.588463 | 7.40923 |
| 6.584424 | 6.338633 |
| 5.26741 | 5.867274 |
| 7.786802 | 7.959191 |
| 8.566038 | 8.599098 |
| 8.152186 | 8.361493 |
| 10.48789 | 10.38089 |
| 9.74826 | 9.318372 |
| 7.539895 | 4.590022 |
| 8.72699 | 8.370301 |
| 8.035337 | 7.921871 |
| 6.174089 | 6.829482 |
| 5.10322 | 4.653559 |
| 8.064094 | 8.014487 |
| NaN | NaN |
| 8.255944 | 8.14792 |
| 7.276011 | 7.221139 |
| 5.732448 | 5.179842 |
| 6.813018 | 7.114271 |
| 5.644508 | 6.253099 |
| 7.218031 | 7.757096 |
| 7.814404 | 7.987084 |
| 5.734136 | 4.627662 |
| 7.766849 | 8.023277 |
| 6.34247 | 6.131171 |
| 5.386398 | 5.772449 |
| 6.531145 | 5.133385 |
| 9.189589 | 9.192746 |
| 7.935704 | 8.088097 |
| 6.024941 | NaN |
| 6.96693 | 6.807619 |
| NaN | NaN |
| 6.007434 | 6.444984 |

-Bruker Confidential-

| | |
|---|---|
| 5.857745 | 5.87442 |
| 5.736113 | 6.62466 |
| 5.703379 | 5.327558 |
| 7.183601 | 7.178649 |
| 6.271425 | 5.840494 |
| 8.002379 | 7.813989 |
| 8.241804 | 8.467347 |
| 6.102396 | 5.66417 |
| 7.352249 | 7.480915 |
| 7.936046 | 7.757806 |
| 8.130979 | 8.175972 |
| 8.405791 | 8.401384 |
| 8.218821 | 8.406713 |
| 7.07554 | 6.898959 |
| 8.129359 | 8.126502 |
| 8.087956 | 8.062047 |
| 6.840104 | 6.730029 |
| NaN | NaN |
| 6.88797 | 6.395782 |
| 6.262579 | 6.097998 |
| 8.663152 | 8.874804 |
| 4.758706 | 4.362919 |
| 5.768093 | 7.862421 |
| 5.619266 | 5.122163 |
| 6.347318 | 6.538308 |
| 7.134002 | 6.773228 |
| 7.078453 | 7.255454 |
| 5.764959 | 5.442092 |
| 7.862994 | 7.959745 |
| 6.463701 | 6.662755 |
| 7.082048 | 6.944218 |
| 5.974385 | 6.685043 |
| 7.926722 | 8.129701 |
| NaN | 6.317787 |
| 5.990059 | 6.370667 |
| 6.74432 | 6.241545 |
| 8.096328 | 7.963404 |
| 10.04418 | 10.35645 |
| NaN | NaN |
| 7.058948 | 6.924813 |
| 7.587404 | 7.835823 |
| 7.669383 | 7.558403 |
| 3.969598 | 3.594221 |
| 8.919409 | 8.913905 |
| 8.019459 | 8.259721 |
| 7.490523 | 7.212769 |
| 5.699541 | 5.940748 |

| | |
|---|---|
| 6.982558 | 6.030528 |
| 7.041472 | 7.027937 |
| 9.279711 | 9.15612 |
| 6.765028 | 7.074007 |
| 7.212462 | 7.295753 |
| 6.558544 | 6.798609 |
| 7.604374 | 7.601823 |
| 7.387262 | 7.58356 |
| 6.465508 | 6.427434 |
| 9.21023 | 9.042866 |
| 7.402613 | 7.924979 |
| 8.56652 | 8.240871 |
| 7.797365 | 8.044911 |
| 6.126307 | 6.252495 |
| 6.552464 | 6.621948 |
| 8.151881 | 8.099174 |
| 7.730081 | 8.139011 |
| 7.423044 | 7.593888 |
| 6.466693 | 6.932126 |
| 9.014607 | 9.081647 |
| 6.193982 | 6.035508 |
| 5.705569 | 5.9409 |
| 6.248393 | 6.087022 |
| 7.362864 | 7.298571 |
| 3.624141 | 2.401788 |
| 7.039644 | 7.096274 |
| 11.75584 | 11.83084 |
| 6.747727 | 6.689469 |
| 8.05263 | 8.16578 |
| 7.000528 | 7.611566 |
| 6.94652 | 6.886634 |
| 5.436256 | 5.157948 |
| NaN | 5.792657 |
| 6.052908 | 6.488605 |
| 4.007226 | 3.189207 |
| 5.463932 | 5.789971 |
| 7.671434 | 7.764204 |
| 9.056993 | 9.126123 |
| 5.934748 | 6.407203 |
| 5.814409 | 5.108465 |
| 5.593405 | 4.107187 |
| 6.36339 | 6.182029 |
| 7.50989 | 7.291532 |
| 5.715874 | 5.197017 |
| 7.855035 | 8.004958 |

-Bruker Confidential-
