## Supplementary material for "Dormant viral pathways underlie space-induced neural senescence": Table_S4

| Name | Descriptio | Gene ID | Alias | Fold Chang | Log Fold C | p-Value | p-Adj | Average Lo | Cluster |
|---|---|---|---|---|---|---|---|---|---|
| ABL1 | ABL proto- | 25 | ABL, JTK7, | 22.4414 | 4.48809 | 0.048039 | 0.998331 | 5.1375 | |
| ACTG1 | actin gamr | 71 | ACT, ACTG | 44.5724 | 5.47808 | 0.012814 | 0.998331 | 9.3456 | |
| ADAR | adenosine | 103 | ADAR1, AG | 22.0108 | 4.46014 | 0.049538 | 0.998331 | 5.0534 | |
| AHNAK | AHNAK nu | 79026 | AHNAKRS | 26.5537 | 4.73084 | 0.036414 | 0.998331 | 5.826 | |
| ALDOA | aldolase A, | 226 | ALDA, GSD | 22.9314 | 4.51925 | 0.047395 | 0.998331 | 5.7999 | |
| ARF1 | ADP-ribosy | 375 | - | 26.3752 | 4.72111 | 0.036714 | 0.998331 | 5.722 | |
| ATP5F1 | ATP syntha | 515 | PIG47, ATF | 23.7501 | 4.56986 | 0.044389 | 0.998331 | 5.6864 | |
| B2M | beta-2-mic | 567 | -, B2-M | 25.7959 | 4.68907 | 0.039848 | 0.998331 | 7.3979 | |
| BRD2 | bromodon | 6046 | D6S113E, I | 23.7586 | 4.57038 | 0.043743 | 0.998331 | 5.3332 | |
| BSG | basigin (Ok | 682 | 5F7, CD14 | 23.0702 | 4.52796 | 0.045977 | 0.998331 | 5.2588 | |
| CBX5 | chromobo | 23468 | HEL25, HP | 27.1432 | 4.76252 | 0.035052 | 0.998331 | 5.8955 | |
| CCNI | cyclin I | 10983 | CCNI1, CYC | 25.6016 | 4.67816 | 0.039123 | 0.998331 | 5.936 | |
| CFL1 | cofilin 1 (n | 1072 | CFL, HEL-S | 23.4061 | 4.54881 | 0.04506 | 0.998331 | 5.3926 | |
| CLTC | clathrin, he | 1213 | CHC, CHC1 | 23.5973 | 4.56055 | 0.044216 | 0.998331 | 5.2982 | |
| CNN2 | calponin 2 | 1265 | - | 23.0955 | 4.52954 | 0.045807 | 0.998331 | 5.2222 | |
| COTL1 | coactosin- | 23406 | CLP | 22.9301 | 4.51917 | 0.046415 | 0.998331 | 5.2297 | |
| PTGS1 | prostaglan | 5742 | COX1, COX | 1.29961 | 0.378084 | 0.872494 | 0.998331 | | |
| PTGS1 | prostaglan | 5742 | COX1, COX | 1.29961 | 0.378084 | 0.872494 | 0.998331 | | |
| CREB1 | cAMP resp | 1385 | CREB | 22.5494 | 4.49502 | 0.048869 | 0.998331 | 5.8614 | |
| CSDE1 | cold shock | 7812 | D1S155E, U | 25.3294 | 4.66274 | 0.039544 | 0.998331 | 5.7024 | |
| CYFIP2 | cytoplasm | 26999 | PIR121 | 22.198 | 4.47236 | 0.048916 | 0.998331 | 5.1054 | |
| DDX5 | DEAD (Asp | 1655 | G17P1, HL | 22.9482 | 4.52031 | 0.04644 | 0.998331 | 5.2701 | |
| DPYSL2 | dihydropyr | 1808 | CRMP-2, C | 27.997 | 4.8072 | 0.033502 | 0.998331 | 6.2298 | |

| | | | | | | | | |
|---|---|---|---|---|---|---|---|---|
| EEF1A1 | eukaryotic | 1915 | CCS-3, CCS | 47.6099 | 5.57319 | 0.012492 | 0.998331 | 9.9008 |
| EEF1G | eukaryotic | 1937 | EF1G, GIG3 | 37.7674 | 5.23907 | 0.019543 | 0.998331 | 7.7384 |
| EEF2 | eukaryotic | 1938 | EEF-2, EF-2 | 37.6255 | 5.23364 | 0.013252 | 0.998331 | 9.0997 |
| EIF3F | eukaryotic | 8665 | EIF3S5, EIF | 25.219 | 4.65644 | 0.039648 | 0.998331 | 5.5759 |
| EIF4B | eukaryotic | 1975 | EIF-4B, PRO | 30.5352 | 4.9324 | 0.028487 | 0.998331 | 6.3307 |
| ENSA | endosulfin | 2029 | ARPP-19E | 23.64 | 4.56316 | 0.044456 | 0.998331 | 5.5044 |
| FLNA | filamin A, a | 2316 | ABP-280, A | 28.2003 | 4.81764 | 0.026912 | 0.998331 | 7.2053 |
| GAPDH | glyceraldeh | 2597 | G3PD, GAP | 21.6234 | 4.43452 | 0.038698 | 0.998331 | 8.2695 |
| GLTSCR2 | glioma tum | 29997 | PICT-1, PIC | 24.3082 | 4.60337 | 0.042143 | 0.998331 | 5.43 |
| GNAS | GNAS com | 2778 | AHO, C20O | 25.2745 | 4.65961 | 0.040777 | 0.998331 | 6.6077 |
| GNB2 | guanine nu | 2783 | - | 22.891 | 4.51671 | 0.046478 | 0.998331 | 5.1902 |
| GNB2L1 | guanine nu | 10399 | GNB2-RS1, | 35.1821 | 5.13677 | 0.02227 | 0.998331 | 7.2846 |
| GPX1 | glutathione | 2876 | GPXD, GSH | 22.2905 | 4.47836 | 0.048753 | 0.998331 | 5.2014 |
| H3F3B | H3 histone | 3021 | H3.3B | 30.0493 | 4.90926 | 0.030033 | 0.998331 | 7.0375 |
| HIST1H3B | histone clu | 8358 | H3/L, H3FL | 23.8096 | 4.57347 | 0.043962 | 0.998331 | 5.5461 |
| HLA-A | major histo | 3105 | HLAA, HLA | 22.8637 | 4.51499 | 0.047711 | 0.998331 | 5.8584 |
| HLA-B | major histo | 3106 | AS, HLAB, S | 24.998 | 4.64374 | 0.041371 | 0.998331 | 6.3879 |
| HLA-C | major histo | 3107 | D6S204, H | 22.4303 | 4.48738 | 0.049255 | 0.998331 | 5.8153 |
| HMGN2 | high mobil | 3151 | HMG17 | 22.4386 | 4.48791 | 0.048023 | 0.998331 | 5.123 |
| HNRNPDL | heterogen | 9987 | HNRNP, HN | 22.7654 | 4.50877 | 0.046912 | 0.998331 | 5.1789 |
| HNRNPH1 | heterogen | 3187 | HNRPH, HN | 26.9927 | 4.7545 | 0.03535 | 0.998331 | 5.8514 |
| HSP90AA1 | heat shock | 3320 | EL52, HSP8 | 29.595 | 4.88728 | 0.030571 | 0.998331 | 6.6278 |
| HSP90AB1 | heat shock | 3326 | D6S182, H | 25.9573 | 4.69807 | 0.03911 | 0.998331 | 6.8862 |

| | | | | | | | | |
|---|---|---|---|---|---|---|---|---|
| HSPA1A | heat shock | 3303 | HEL-S-103, | 24.6151 | 4.62147 | 0.041813 | 0.998331 | 5.8021 |
| HSPA1B | heat shock | 3304 | HSP70-1B, | 22.5014 | 4.49194 | 0.049915 | 0.998331 | 6.7533 |
| HSPA8 | heat shock | 3312 | HEL-33, HE | 33.3152 | 5.05811 | 0.024354 | 0.998331 | 6.7072 |
| KLF13 | Kruppel-lik | 51621 | BTEB3, FKL | 24.292 | 4.60241 | 0.042458 | 0.998331 | 5.5797 |
| LDOC1L | leucine zip | 84247 | MAR6, MA | 23.4976 | 4.55444 | 0.045075 | 0.998331 | 5.584 |
| LENG8 | leukocyte | 114823 | PP13842 | 27.768 | 4.79535 | 0.033626 | 0.998331 | 5.9322 |
| LINC01305 | long interg | 285084 | - | -22.5529 | -4.49524 | 0.049827 | 0.998331 | 5.9118 |
| MCL1 | myeloid ce | 4170 | BCL2L3, EA | 27.2186 | 4.76652 | 0.034989 | 0.998331 | 5.9822 |
| MID1IP1 | MID1 inter | 58526 | G12-LIKE, I | 23.4322 | 4.55042 | 0.045897 | 0.998331 | 6.0012 |
| MSN | moesin | 4478 | HEL70 | 25.309 | 4.66158 | 0.039445 | 0.998331 | 5.6117 |
| MYH9 | myosin, he | 4627 | BDPLT6, D | 26.32 | 4.71809 | 0.036873 | 0.998331 | 5.733 |
| MYL6 | myosin, lig | 4637 | ESMLC, LC | 24.7696 | 4.6305 | 0.040858 | 0.998331 | 5.5076 |
| NACA | nascent po | 4666 | HSD48, NA | 23.033 | 4.52563 | 0.046505 | 0.998331 | 5.4649 |
| NAP1L1 | nucleosom | 4673 | NAP1, NAF | 23.2164 | 4.53707 | 0.045751 | 0.998331 | 5.4077 |
| NCOA4 | nuclear rec | 8031 | ARA70, ELI | 22.0244 | 4.46103 | 0.049613 | 0.998331 | 5.114 |
| NPM1 | nucleopho | 4869 | B23, NPM | 27.394 | 4.77579 | 0.0344 | 0.998331 | 5.8641 |
| PABPC1 | poly(A) bir | 26986 | PAB1, PAB | 27.5408 | 4.7835 | 0.034335 | 0.998331 | 6.0606 |
| PDIA4 | protein dis | 9601 | ERP70, ERI | 23.7249 | 4.56833 | 0.04427 | 0.998331 | 5.5611 |
| PFN1 | profilin 1 | 5216 | ALS18 | 23.9591 | 4.5825 | 0.043329 | 0.998331 | 5.4662 |
| PKM | pyruvate k | 5315 | CTHBP, HE | 27.7343 | 4.7936 | 0.033872 | 0.998331 | 6.0505 |
| POLR2A | polymeras | 5430 | POLR2, PO | 22.4431 | 4.4882 | 0.048459 | 0.998331 | 5.3426 |
| PPIA | peptidylpr | 5478 | CYPA, CYPI | 29.1742 | 4.86662 | 0.03087 | 0.998331 | 6.1401 |
| PPIB | peptidylpr | 5479 | CYP-S1, CY | 22.9882 | 4.52282 | 0.046243 | 0.998331 | 5.2484 |
| PRKAR1A | protein kir | 5573 | ACRDYS1, . | 25.921 | 4.69605 | 0.038094 | 0.998331 | 5.8346 |

| | | | | | | | | |
|---|---|---|---|---|---|---|---|---|
| PRRC2B | proline-ric | 84726 | BAT2L, BA | 22.9584 | 4.52095 | 0.047191 | 0.998331 | 5.7252 |
| PSAP | prosaposir | 5660 | GLBA, SAP | 25.564 | 4.67604 | 0.038695 | 0.998331 | 5.5957 |
| PTGS1 | prostaglan | 5742 | COX1, CO | 1.29961 | 0.378084 | 0.872494 | 0.998331 | |
| PTMA | prothymos | 5757 | TMSA | 25.0184 | 4.64492 | 0.040125 | 0.998331 | 5.515 |
| RALY | RALY heter | 22913 | HNRPCL2, | 23.5623 | 4.55841 | 0.044466 | 0.998331 | 5.367 |
| RBBP4 | retinoblast | 5928 | NURF55, R | 23.0257 | 4.52517 | 0.046111 | 0.998331 | 5.2494 |
| RMRP | RNA comp | 6023 | CHH, NME | 48.8325 | 5.60977 | 0.011765 | 0.998331 | 9.4063 |
| RPL10 | ribosomal | 6134 | AUTSX5, D | 20.7581 | 4.3756 | 0.046416 | 0.998331 | 6.5347 |
| RPL10A | ribosomal | 4736 | CSA19, CS | 32.1302 | 5.00586 | 0.025998 | 0.998331 | 6.5398 |
| RPL11 | ribosomal | 6135 | DBA7, GIG | 27.1577 | 4.76329 | 0.035244 | 0.998331 | 6.0577 |
| RPL12 | ribosomal | 6136 | L12 | 30.0945 | 4.91143 | 0.029504 | 0.998331 | 6.5163 |
| RPL13 | ribosomal | 6137 | BBC1, D16 | 34.0759 | 5.09068 | 0.023682 | 0.998331 | 7.2178 |
| RPL15 | ribosomal | 6138 | DBA12, EC | 28.4359 | 4.82964 | 0.032589 | 0.998331 | 6.2762 |
| RPL18 | ribosomal | 6141 | L18 | 31.9741 | 4.99883 | 0.026442 | 0.998331 | 6.7399 |
| RPL18A | ribosomal | 6142 | L18A | 31.6306 | 4.98325 | 0.026865 | 0.998331 | 6.5913 |
| RPL19 | ribosomal | 6143 | L19 | 32.2977 | 5.01336 | 0.025869 | 0.998331 | 6.6777 |
| RPL24 | ribosomal | 6152 | HEL-S-310, | 22.6909 | 4.50404 | 0.047245 | 0.998331 | 5.2025 |
| RPL26 | ribosomal | 6154 | DBA11, L2 | 23.7039 | 4.56705 | 0.043981 | 0.998331 | 5.3598 |
| RPL27 | ribosomal | 6155 | L27 | 25.4478 | 4.66947 | 0.039007 | 0.998331 | 5.5867 |
| RPL27A | ribosomal | 6157 | L27A | 31.0724 | 4.95756 | 0.027647 | 0.998331 | 6.4316 |
| RPL29 | ribosomal | 6159 | HIP, HUM | 26.8684 | 4.74784 | 0.036319 | 0.998331 | 6.37 |
| RPL3 | ribosomal | 6122 | ASC-1, L3, | 36.0733 | 5.17286 | 0.021462 | 0.998331 | 7.808 |
| RPL31 | ribosomal | 6160 | L31 | 24.8672 | 4.63617 | 0.040626 | 0.998331 | 5.542 |

| | | | | | | | | |
|---|---|---|---|---|---|---|---|---|
| RPL32 | ribosomal | 6161 | L32, PP993 | 29.8794 | 4.90108 | 0.029644 | 0.998331 | 6.2735 |
| RPL34 | ribosomal | 6164 | L34 | 23.1511 | 4.53301 | 0.045628 | 0.998331 | 5.2311 |
| RPL37 | ribosomal | 6167 | L37 | 30.1603 | 4.91458 | 0.029105 | 0.998331 | 6.2681 |
| RPL37A | ribosomal | 6168 | L37A | 27.5773 | 4.78541 | 0.034356 | 0.998331 | 6.1429 |
| RPL38 | ribosomal | 6169 | L38 | 32.0388 | 5.00175 | 0.026388 | 0.998331 | 6.7994 |
| RPL4 | ribosomal | 6124 | L4 | 38.6536 | 5.27253 | 0.018646 | 0.998331 | 7.749 |
| RPL41 | ribosomal | 6171 | L41 | 23.9257 | 4.58049 | 0.043239 | 0.998331 | 5.3585 |
| RPL6 | ribosomal | 6128 | L6, SHUJUI | 31.0248 | 4.95535 | 0.027944 | 0.998331 | 6.6419 |
| RPL7 | ribosomal | 6129 | L7, HUML7 | 30.1831 | 4.91567 | 0.029096 | 0.998331 | 6.296 |
| RPL7A | ribosomal | 6130 | L7A, SURF3 | 33.0725 | 5.04756 | 0.025063 | 0.998331 | 7.145 |
| RPL8 | ribosomal | 6132 | L8 | 32.1367 | 5.00615 | 0.026267 | 0.998331 | 6.8386 |
| RPL9 | ribosomal | 6133 | L9, NPC-A- | 28.404 | 4.82802 | 0.032317 | 0.998331 | 6.0104 |
| RPLP0 | ribosomal | 6175 | L10E, LP0, | 36.0049 | 5.17012 | 0.021424 | 0.998331 | 7.5631 |
| RPLP2 | ribosomal | 6181 | D11S2243I | 25.8124 | 4.68999 | 0.038258 | 0.998331 | 5.7454 |
| RPS11 | ribosomal | 6205 | S11 | 34.5281 | 5.1097 | 0.023024 | 0.998331 | 7.1461 |
| RPS12 | ribosomal | 6206 | S12 | 28.4158 | 4.82862 | 0.032309 | 0.998331 | 6.023 |
| RPS13 | ribosomal | 6207 | S13 | 22.1061 | 4.46637 | 0.049186 | 0.998331 | 5.0664 |
| RPS14 | ribosomal | 6208 | EMTB, S14 | 30.4472 | 4.92824 | 0.028673 | 0.998331 | 6.3519 |
| RPS15 | ribosomal | 6209 | RIG, S15 | 23.1053 | 4.53015 | 0.046836 | 0.998331 | 5.8445 |
| RPS15A | ribosomal | 6210 | S15A | 25.1481 | 4.65238 | 0.03984 | 0.998331 | 5.5678 |
| RPS18 | ribosomal | 6222 | D6S218E, I | 35.3285 | 5.14276 | 0.021917 | 0.998331 | 7.0417 |
| RPS19 | ribosomal | 6223 | DBA, DBA1 | 30.481 | 4.92984 | 0.029146 | 0.998331 | 6.9166 |
| RPS2 | ribosomal | 6187 | LLREP3, S2 | 34.4514 | 5.10649 | 0.023456 | 0.998331 | 7.7204 |
| RPS20 | ribosomal | 6224 | S20 | 23.4868 | 4.55378 | 0.045126 | 0.998331 | 5.5866 |

| | | | | | | | | |
|---|---|---|---|---|---|---|---|---|
| RPS23 | ribosomal | 6228 | S23 | 26.7152 | 4.73959 | 0.036256 | 0.998331 | 5.9972 |
| RPS25 | ribosomal | 6230 | S25 | 24.1602 | 4.59456 | 0.042528 | 0.998331 | 5.3857 |
| RPS27 | ribosomal | 6232 | MPS-1, MF | 31.7649 | 4.98936 | 0.026537 | 0.998331 | 6.4882 |
| RPS3 | ribosomal | 6188 | S3 | 30.01 | 4.90737 | 0.029558 | 0.998331 | 6.4154 |
| RPS3A | ribosomal | 6189 | FTE1, MFT | 28.9935 | 4.85766 | 0.031205 | 0.998331 | 6.1118 |
| RPS4X | ribosomal | 6191 | CCG2, DXS | 32.1291 | 5.00581 | 0.026066 | 0.998331 | 6.6054 |
| RPS4Y1 | ribosomal | 6192 | RPS4Y, S4 | 26.1446 | 4.70844 | 0.037347 | 0.998331 | 5.7401 |
| RPS5 | ribosomal | 6193 | S5 | 19.7032 | 4.30036 | 0.049356 | 0.998331 | 6.2992 |
| RPS6 | ribosomal | 6194 | S6 | 33.4698 | 5.06479 | 0.024195 | 0.998331 | 6.7777 |
| RPS7 | ribosomal | 6201 | DBA8, S7 | 24.4064 | 4.60919 | 0.042035 | 0.998331 | 5.5393 |
| RPS8 | ribosomal | 6202 | S8 | 33.3857 | 5.06116 | 0.024276 | 0.998331 | 6.7327 |
| RPS9 | ribosomal | 6203 | S9 | 29.7626 | 4.89543 | 0.029864 | 0.998331 | 6.2691 |
| RPSA | ribosomal | 3921 | 37LRP, 67L | 32.7888 | 5.03513 | 0.025096 | 0.998331 | 6.6631 |
| SCARNA10 | small Cajal | 692148 | U85 | 26.9494 | 4.75218 | 0.035632 | 0.998331 | 5.9722 |
| SET | SET nuclea | 6418 | 2PP2A, I2F | 22.6502 | 4.50145 | 0.04737 | 0.998331 | 5.1898 |
| SETD1B | SET domai | 23067 | KMT2G, SE | 22.2033 | 4.4727 | 0.048935 | 0.998331 | 5.1267 |
| SHISA9 | shisa famil | 729993 | CKAMP44 | -20.1672 | -4.33394 | 0.046536 | 0.998331 | 6.3255 |
| SMARCA4 | SWI/SNF r | 6597 | BAF190, B/ | 22.7944 | 4.51061 | 0.047842 | 0.998331 | 5.765 |
| SNORD17 | small nucle | 692086 | HBI-43 | 22.0627 | 4.46354 | 0.049637 | 0.998331 | 5.1975 |
| SPARC | secreted p | 6678 | BM-40, ON | 24.7475 | 4.62921 | 0.040925 | 0.998331 | 5.5078 |
| SPTAN1 | spectrin, a | 6709 | EIEE5, NEA | 25.0075 | 4.64429 | 0.040365 | 0.998331 | 5.6321 |
| TMSB10 | thymosin b | 9168 | MIG12, TB | 27.7749 | 4.79571 | 0.033617 | 0.998331 | 5.935 |
| TMSB4X | thymosin b | 7114 | FX, PTMB4 | 27.6796 | 4.79075 | 0.033854 | 0.998331 | 5.9485 |

| | | | | | | | | |
|---|---|---|---|---|---|---|---|---|
| TRA2B | transforme | 6434 | HTRA2-BET | 23.5623 | 4.55841 | 0.044589 | 0.998331 | 5.4319 |
| TUBA1A | tubulin, alp | 7846 | B-ALPHA-1 | 25.0155 | 4.64475 | 0.041973 | 0.998331 | 7.2779 |
| TUBA1B | tubulin, alp | 10376 | K-ALPHA-1 | 29.3683 | 4.87619 | 0.030966 | 0.998331 | 6.5732 |
| TUBB | tubulin, be | 203068 | CDCBM6, I | 33.8844 | 5.08255 | 0.024207 | 0.998331 | 7.6792 |
| TXNIP | thioredoxi | 10628 | EST01027, | 29.0736 | 4.86164 | 0.031272 | 0.998331 | 6.3013 |
| UBA52 | ubiquitin A | 7311 | CEP52, HU | 30.5886 | 4.93492 | 0.028379 | 0.998331 | 6.3214 |
| UBB | ubiquitin B | 7314 | - | 25.7821 | 4.6883 | 0.038274 | 0.998331 | 5.7038 |
| UBC | ubiquitin C | 7316 | HMG20 | 26.0474 | 4.70307 | 0.038371 | 0.998331 | 6.3174 |
| UBE2D3 | ubiquitin-c | 7323 | E2(17)KB3, | 23.9358 | 4.5811 | 0.043372 | 0.998331 | 5.448 |
| VIM | vimentin | 7431 | CTRCT30, I | 32.645 | 5.02879 | 0.022531 | 0.998331 | 8.0999 |
| YBX1 | Y box bind | 4904 | BP-8, CSDA | 27.3697 | 4.77451 | 0.034851 | 0.998331 | 6.1479 |
| YWHAZ | tyrosine 3- | 7534 | 14-3-3-ZET | 26.1555 | 4.70904 | 0.03727 | 0.998331 | 5.7083 |
| ZEB2 | zinc finger | 9839 | HSPC082, S | 23.579 | 4.55943 | 0.044987 | 0.998331 | 5.7024 |

| Ground 1 | Ground 2 | Space 1 | Space 2 |
|---|---|---|---|
| 0 | 0 | 6.50621 | 5.69778 |
| 1.90133 | 0 | 6.4078 | 11.2973 |
| 0 | 0 | 5.8564 | 6.26477 |
| 0 | 0 | 7.39928 | 5.90912 |
| 0 | 0 | 3.08605 | 7.75723 |
| 0 | 0 | 6.50621 | 6.9334 |
| 0 | 0 | 7.5796 | 4.06194 |
| 0 | 0 | 9.39286 | 2.12986 |
| 0 | 0 | 6.03184 | 6.61218 |
| 0 | 0 | 5.56828 | 6.75094 |
| 0 | 0 | 6.09645 | 7.4237 |
| 0 | 0 | 4.64231 | 7.79392 |
| 0 | 0 | 5.0859 | 7.0882 |
| 0 | 0 | 6.24634 | 6.3833 |
| 0 | 0 | 6.09645 | 6.37306 |
| 0 | 0 | 6.68491 | 5.62067 |
| 0 | 2.69402 | 0 | 0 |
| 0 | 2.69402 | 0 | 0 |
| 0 | 0 | 7.8376 | 2.539 |
| 0 | 0 | 5.37347 | 7.39938 |
| 0 | 0 | 5.56828 | 6.52758 |
| 0 | 0 | 5.32036 | 6.86295 |
| 0 | 0 | 5.37347 | 8.02656 |

| | | | |
|---|---|---|---|
| 0 | 0 | 9.6048 | 11.5732 |
| 0 | 0 | 7.20786 | 9.4683 |
| 1.90133 | 3.03031 | 8.01521 | 10.9128 |
| 0 | 0 | 6.03184 | 6.99293 |
| 0 | 0 | 7.00403 | 7.61165 |
| 0 | 0 | 7.29302 | 4.74134 |
| 1.90133 | 0 | 7.61311 | 8.62132 |
| 1.90133 | 3.40051 | 5.14819 | 10.215 |
| 0 | 0 | 5.9642 | 6.80749 |
| 0 | 0 | 2.80598 | 8.5891 |
| 0 | 0 | 6.06451 | 6.34186 |
| 0 | 0 | 6.78631 | 9.00921 |
| 0 | 0 | 6.88105 | 4.96838 |
| 0 | 0 | 4.72637 | 8.9687 |
| 0 | 0 | 7.34713 | 4.70445 |
| 0 | 0 | 7.82812 | 2.7708 |
| 0 | 0 | 8.36253 | 2.97047 |
| 0 | 0 | 7.78956 | 2.58031 |
| 0 | 0 | 5.89324 | 6.3568 |
| 0 | 0 | 6.45784 | 5.88253 |
| 0 | 0 | 7.30674 | 6.22135 |
| 0 | 0 | 5.26522 | 8.48825 |
| 0 | 0 | 2.80598 | 8.87089 |

| | | | |
|---|---|---|---|
| 0 | 0 | 4.35602 | 7.67738 |
| 0 | 0 | 1.32096 | 8.75164 |
| 0 | 0 | 7.62411 | 7.79885 |
| 0 | 0 | 7.33379 | 4.99917 |
| 0 | 0 | 7.44961 | 4.2529 |
| 0 | 0 | 7.20786 | 6.62086 |
| 7.91725 | 0 | 0 | 0.116922 |
| 0 | 0 | 7.65661 | 5.73027 |
| 0 | 0 | 7.97281 | 2.80606 |
| 0 | 0 | 7.11735 | 5.87633 |
| 0 | 0 | 7.10169 | 6.27424 |
| 0 | 0 | 5.9642 | 6.9254 |
| 0 | 0 | 4.35602 | 7.30549 |
| 0 | 0 | 4.80581 | 7.16828 |
| 0 | 0 | 6.68491 | 5.2341 |
| 0 | 0 | 6.8439 | 6.90827 |
| 0 | 0 | 5.65664 | 7.7718 |
| 0 | 0 | 7.38642 | 4.55703 |
| 0 | 0 | 7.13283 | 5.2661 |
| 0 | 0 | 5.8564 | 7.70857 |
| 0 | 0 | 4.35602 | 7.16828 |
| 0 | 0 | 7.44961 | 6.77227 |
| 0 | 0 | 6.7466 | 5.54444 |
| 0 | 0 | 7.5796 | 5.28497 |

| | | | |
|---|---|---|---|
| 0 | 0 | 3.32048 | 7.66962 |
| 0 | 0 | 6.553 | 6.66594 |
| 0 | 2.69402 | 0 | 0 |
| 0 | 0 | 6.43304 | 6.6221 |
| 0 | 0 | 6.9526 | 5.43327 |
| 0 | 0 | 6.72633 | 5.59319 |
| 0 | 0 | 10.0503 | 10.6934 |
| 1.90133 | 0 | 5.32036 | 8.36697 |
| 0 | 0 | 7.46193 | 7.62837 |
| 0 | 0 | 5.37347 | 7.82647 |
| 0 | 0 | 5.8564 | 8.27717 |
| 0 | 0 | 6.38212 | 9.00566 |
| 0 | 0 | 5.52199 | 8.05568 |
| 0 | 0 | 6.35597 | 8.44138 |
| 0 | 0 | 6.62046 | 8.17637 |
| 0 | 0 | 6.76659 | 8.2414 |
| 0 | 0 | 5.47416 | 6.71197 |
| 0 | 0 | 5.65664 | 6.85561 |
| 0 | 0 | 6.82496 | 6.33735 |
| 0 | 0 | 6.89927 | 7.83236 |
| 0 | 0 | 4.24641 | 8.29395 |
| 0 | 0 | 6.45784 | 9.66265 |
| 0 | 0 | 5.8186 | 7.04366 |

| | | | |
|---|---|---|---|
| 0 | 0 | 6.64227 | 7.72478 |
| 0 | 0 | 6.09645 | 6.38913 |
| 0 | 0 | 7.05369 | 7.47114 |
| 0 | 0 | 5.37347 | 7.92598 |
| 0 | 0 | 8.54143 | 6.22787 |
| 0 | 0 | 7.56826 | 9.39386 |
| 0 | 0 | 6.06451 | 6.63196 |
| 0 | 0 | 6.06451 | 8.38602 |
| 0 | 0 | 6.82496 | 7.66483 |
| 0 | 0 | 6.03184 | 8.97355 |
| 0 | 0 | 6.18824 | 8.59573 |
| 0 | 0 | 6.9526 | 7.08731 |
| 0 | 0 | 6.70577 | 9.35328 |
| 0 | 0 | 5.61314 | 7.38924 |
| 0 | 0 | 6.7466 | 8.84914 |
| 0 | 0 | 6.7466 | 7.27357 |
| 0 | 0 | 6.21758 | 5.94515 |
| 0 | 0 | 6.7466 | 7.79007 |
| 0 | 0 | 3.08605 | 7.80322 |
| 0 | 0 | 5.99842 | 6.99771 |
| 0 | 0 | 7.53368 | 8.42517 |
| 0 | 0 | 5.14819 | 8.81304 |
| 0 | 0 | 5.89324 | 9.61871 |
| 0 | 0 | 4.24641 | 7.45319 |

| | | | |
|---|---|---|---|
| 0 | 0 | 5.26522 | 7.77514 |
| 0 | 0 | 6.50621 | 6.29144 |
| 0 | 0 | 7.41203 | 7.57571 |
| 0 | 0 | 6.1277 | 8.09541 |
| 0 | 0 | 6.76659 | 7.40729 |
| 0 | 0 | 6.93504 | 8.07207 |
| 0 | 0 | 7.25107 | 5.988 |
| 1.90133 | 0 | 6.18824 | 7.91492 |
| 0 | 0 | 7.23681 | 8.18015 |
| 0 | 0 | 5.32036 | 7.21001 |
| 0 | 0 | 7.43719 | 7.98918 |
| 0 | 0 | 6.553 | 7.75892 |
| 0 | 0 | 7.23681 | 8.00301 |
| 0 | 0 | 7.6884 | 5.54965 |
| 0 | 0 | 5.52199 | 6.67312 |
| 0 | 0 | 6.64227 | 5.38745 |
| 7.70669 | 6.7936 | 0 | 1.98355 |
| 0 | 0 | 3.08605 | 7.72133 |
| 0 | 0 | 6.93504 | 4.72301 |
| 0 | 0 | 5.92916 | 6.94333 |
| 0 | 0 | 5.4247 | 7.29854 |
| 0 | 0 | 6.59832 | 7.22718 |
| 0 | 0 | 7.38642 | 6.35234 |

| | | | |
|---|---|---|---|
| 0 | 0 | 7.14815 | 5.03676 |
| 0 | 0 | 1.9988 | 9.27327 |
| 0 | 0 | 5.26522 | 8.428 |
| 0 | 0 | 5.69886 | 9.5885 |
| 0 | 0 | 8.01521 | 5.87633 |
| 0 | 0 | 7.38642 | 7.272 |
| 0 | 0 | 5.8186 | 7.26726 |
| 0 | 0 | 3.85649 | 8.25979 |
| 0 | 0 | 7.08587 | 5.34912 |
| 1.90133 | 0 | 5.32036 | 10.0454 |
| 0 | 0 | 5.20789 | 7.95817 |
| 0 | 0 | 6.24634 | 7.07922 |
| 0 | 0 | 7.61311 | 3.85867 |
