## Supplementary material for "Dormant viral pathways underlie space-induced neural senescence": Table_S5

| Bank | Pathway | Description | Log | Count |
|---|---|---|---|---|
| Reactome Gene Sets | R-HSA-2262752 | Cellular Responses | -17.871 | 46 |
| Reactome Gene Sets | R-HSA-9824446 | Viral Infection Pathv | -9.975 | 34 |
| Reactome Gene Sets | R-HSA-422475 | Axon Guidance | -9.577 | 28 |
| CORUM | CORUM:3055 | Nop56p-Associatec | -9.34 | 13 |
| Reactome Gene Sets | R-HSA-9675108 | Nervous System De | -9.145 | 28 |
| WikiPathways | WP477 | Cytoplasmic Ribosc | -9.113 | 12 |
| Reactome Gene Sets | R-HSA-2408557 | Selenocysteine Syn | -8.774 | 12 |
| CORUM | CORUM:306 | Ribosome, Cytoplas | -8.442 | 11 |
| GO Biological Processes | GO:0002181 | Cytoplasmic Transl | -8.428 | 13 |
| Reactome Gene Sets | R-HSA-9711097 | Cellular Response t | -8.064 | 14 |
| Reactome Gene Sets | R-HSA-156902 | Peptide Chain Elon | -7.89 | 11 |
| Reactome Gene Sets | R-HSA-192823 | Viral mRNA Translat | -7.89 | 11 |
| Reactome Gene Sets | R-HSA-1799339 | SRP-Dependent Co | -7.849 | 12 |
| GO Biological Processes | GO:0043604 | Amide Biosynthetic | -7.754 | 25 |
| Reactome Gene Sets | R-HSA-156842 | Eukaryotic Translati | -7.689 | 11 |

Viral Infection Pathways
CALR
CANX
CCNH
EGFR
ERCC3
FYN
GTF2E2
HLAC
RPL10A
PIK3C3
PTPN11
RPL5
RPL7
RPL7A
RPL8
RPL31
RPS7
RPS8
RPS15
RPS18
RPS27
H2AC14
H2AC17
H2BC11

SEC24C
AKT3
TUBB3
TUBB4B
MAN1B1
H2AC12
CHMP4B
VPS37A
H2AC21
TUBB2B

| Total from pathway | Ratio |
|---|---|
| 788 | 0.058376 |
| 767 | 0.044329 |
| 553 | 0.050633 |
| 104 | 0.125 |
| 578 | 0.048443 |
| 88 | 0.136364 |
| 94 | 0.12766 |
| 80 | 0.1375 |
| 123 | 0.105691 |
| 157 | 0.089172 |
| 90 | 0.122222 |
| 90 | 0.122222 |
| 113 | 0.106195 |
| 545 | 0.045872 |
| 94 | 0.117021 |
