## Supplementary material for "Dormant viral pathways underlie space-induced neural senescence": Table_S6

| p_val | avg_log2FC | pct.1 | pct.2 | p_val_adj | cluster | gene |
|---|---|---|---|---|---|---|
| 2.7775E-69 | -0.4193406 | 0.569 | 0.65 | 1.3904E-65 | KO | Grm3 |
| 3.9934E-66 | -0.2121697 | 0.975 | 0.982 | 1.9991E-62 | KO | Gja1 |
| 9.8383E-60 | -0.3757787 | 0.512 | 0.611 | 4.925E-56 | KO | Tubb2b |
| 1.4651E-44 | -0.1372959 | 0.989 | 0.993 | 7.3343E-41 | KO | Gnao1 |
| 4.6884E-38 | -0.2267369 | 0.793 | 0.836 | 2.347E-34 | KO | Msmo1 |
| 6.403E-32 | -0.3947879 | 0.292 | 0.365 | 3.2053E-28 | KO | Idi1 |
| 6.2739E-31 | -0.3333891 | 0.389 | 0.459 | 3.1407E-27 | KO | Hapln1 |
| 2.5883E-29 | -0.5820813 | 0.311 | 0.374 | 1.2957E-25 | KO | Ccn1 |
| 7.191E-29 | -0.9347425 | 0.051 | 0.092 | 3.5998E-25 | KO | Eif2a |
| 3.4714E-28 | -0.116879 | 0.993 | 0.995 | 1.7378E-24 | KO | Atp1b2 |
| 2.5718E-27 | -0.1879486 | 0.724 | 0.771 | 1.2875E-23 | KO | Hmgcr |
| 8.661E-27 | -0.5164137 | 0.229 | 0.293 | 4.3357E-23 | KO | Gfap |
| 2.7898E-26 | -1.35424 | 0.04 | 0.074 | 1.3966E-22 | KO | Gpr88 |
| 4.4877E-25 | -1.5846723 | 0.018 | 0.044 | 2.2465E-21 | KO | Tac1 |
| 1.0294E-24 | -0.4209862 | 0.195 | 0.253 | 5.1531E-21 | KO | Prelp |
| 2.2884E-24 | -0.1460549 | 0.853 | 0.88 | 1.1456E-20 | KO | Gabbr1 |
| 1.3478E-22 | -0.5412104 | 0.114 | 0.16 | 6.747E-19 | KO | Clk1 |
| 3.1213E-21 | -0.4489066 | 0.174 | 0.225 | 1.5625E-17 | KO | Fam214a |
| 3.6816E-21 | -1.1217016 | 0.045 | 0.077 | 1.843E-17 | KO | Pde10a |
| 6.9173E-21 | -0.2777427 | 0.469 | 0.519 | 3.4628E-17 | KO | Syne1 |
| 4.3222E-20 | -0.1731818 | 0.772 | 0.807 | 2.1637E-16 | KO | Nsmf |
| 1.2443E-19 | -0.1890969 | 0.644 | 0.685 | 6.2289E-16 | KO | Rsrp1 |
| 6.2268E-19 | -1.3621002 | 0.014 | 0.034 | 3.1172E-15 | KO | Six3 |
| 3.5001E-18 | -0.2556704 | 0.453 | 0.501 | 1.7521E-14 | KO | Ivd |
| 5.6672E-18 | -0.2564968 | 0.411 | 0.462 | 2.837E-14 | KO | Gm2a |
| 6.6402E-18 | -3.1021418 | 0.002 | 0.013 | 3.3241E-14 | KO | Pmch |
| 7.0739E-18 | -0.2947373 | 0.35 | 0.402 | 3.5412E-14 | KO | Acss2 |
| 9.8521E-17 | -1.0943364 | 0.015 | 0.033 | 4.9319E-13 | KO | Rpe65 |
| 2.0397E-16 | -0.1907409 | 0.567 | 0.608 | 1.0211E-12 | KO | Dst |
| 3.0981E-16 | -0.1401092 | 0.79 | 0.817 | 1.5509E-12 | KO | Ncam1 |
| 5.4707E-16 | -0.2839928 | 0.328 | 0.374 | 2.7387E-12 | KO | Sorbs1 |
| 5.9083E-16 | -0.3064672 | 0.226 | 0.273 | 2.9577E-12 | KO | Apln |
| 1.2427E-15 | -0.2499252 | 0.412 | 0.457 | 6.221E-12 | KO | Gabbr2 |
| 1.8454E-15 | -0.3048788 | 0.359 | 0.404 | 9.2382E-12 | KO | Ndrg4 |
| 2.5257E-15 | -0.49966 | 0.092 | 0.126 | 1.2643E-11 | KO | Slc32a1 |
| 4.8252E-15 | -0.2374246 | 0.431 | 0.478 | 2.4155E-11 | KO | Gria2 |
| 5.8383E-15 | -1.1012182 | 0.017 | 0.034 | 2.9227E-11 | KO | Otof |
| 6.4064E-15 | -0.2816896 | 0.323 | 0.37 | 3.2071E-11 | KO | Pou3f3 |
| 8.513E-15 | -0.4272933 | 0.128 | 0.165 | 4.2616E-11 | KO | Cd38 |
| 9.4765E-15 | -0.3975539 | 0.135 | 0.174 | 4.744E-11 | KO | Noct |
| 2.5694E-14 | -0.2871821 | 0.351 | 0.392 | 1.2862E-10 | KO | Ppp3r1 |
| 2.8274E-14 | -0.4189521 | 0.115 | 0.151 | 1.4154E-10 | KO | Igfbp2 |
| 4.6264E-14 | -0.4122416 | 0.138 | 0.175 | 2.316E-10 | KO | Hsd17b7 |
| 5.256E-14 | -0.3490868 | 0.161 | 0.201 | 2.6311E-10 | KO | Jam2 |
| 7.245E-14 | -0.1837853 | 0.467 | 0.514 | 3.6269E-10 | KO | Atp6v0a1 |
| 7.9328E-14 | -0.6926775 | 0.044 | 0.069 | 3.9712E-10 | KO | Gad2 |
| 8.4008E-14 | -0.22606 | 0.38 | 0.426 | 4.2054E-10 | KO | Ogt |
| 8.7416E-14 | -0.4718241 | 0.1 | 0.133 | 4.376E-10 | KO | Mecp2 |
| 1.0944E-13 | -1.1508103 | 0.105 | 0.137 | 5.4786E-10 | KO | Crym |

| | | | | | | |
|---|---|---|---|---|---|---|
| 1.7092E-13 | -1.0385592 | 0.017 | 0.034 | 8.5562E-10 | KO | A2m |
| 2.493E-13 | -0.2051228 | 0.45 | 0.492 | 1.248E-09 | KO | Hnrnpa1 |
| 2.8438E-13 | -0.3384334 | 0.165 | 0.203 | 1.4236E-09 | KO | Cbx5 |
| 5.168E-13 | -1.422355 | 0.009 | 0.021 | 2.5871E-09 | KO | Pdyn |
| 6.3124E-13 | -0.1228452 | 0.835 | 0.855 | 3.16E-09 | KO | Cd81 |
| 8.26E-13 | -0.5597961 | 0.058 | 0.084 | 4.1349E-09 | KO | Adamts20 |
| 1.2342E-12 | -0.4021211 | 0.134 | 0.168 | 6.1785E-09 | KO | Arhgef1 |
| 5.9193E-12 | -0.3837147 | 0.163 | 0.198 | 2.9632E-08 | KO | Ptpn5 |
| 9.3383E-12 | -0.1995198 | 0.621 | 0.655 | 4.6748E-08 | KO | Hpcal4 |
| 9.9389E-12 | -0.2105179 | 0.418 | 0.457 | 4.9754E-08 | KO | Washc2 |
| 2.2849E-11 | -0.5894671 | 0.057 | 0.08 | 1.1438E-07 | KO | Serpina3n |
| 2.305E-11 | -0.4462824 | 0.08 | 0.107 | 1.1539E-07 | KO | Efemp1 |
| 2.4133E-11 | -0.3062584 | 0.213 | 0.25 | 1.2081E-07 | KO | Fstl1 |
| 2.6577E-11 | -0.8058089 | 0.055 | 0.078 | 1.3305E-07 | KO | Thbs4 |
| 4.5443E-11 | -0.4746028 | 0.049 | 0.072 | 2.2749E-07 | KO | Gdpd2 |
| 1.0555E-10 | -0.2711518 | 0.178 | 0.213 | 5.2837E-07 | KO | Cyp7b1 |
| 1.1545E-10 | -0.3673014 | 0.161 | 0.193 | 5.7796E-07 | KO | Rbp1 |
| 1.1668E-10 | -0.1779322 | 0.385 | 0.424 | 5.8408E-07 | KO | Zeb1 |
| 1.4075E-10 | -0.3224212 | 0.178 | 0.211 | 7.0459E-07 | KO | Gmfb |
| 2.0928E-10 | -0.2346301 | 0.335 | 0.372 | 1.0476E-06 | KO | Tardbp |
| 2.3656E-10 | -0.2185797 | 0.391 | 0.424 | 1.1842E-06 | KO | Vcam1 |
| 2.7712E-10 | -0.2704693 | 0.226 | 0.261 | 1.3873E-06 | KO | Tceal5 |
| 2.7718E-10 | -0.1507661 | 0.612 | 0.639 | 1.3876E-06 | KO | Dio2 |
| 3.0187E-10 | -0.2556491 | 0.274 | 0.311 | 1.5112E-06 | KO | Ldlr |
| 3.1727E-10 | -1.0196191 | 0.015 | 0.027 | 1.5883E-06 | KO | Rxrg |
| 3.7361E-10 | -0.1031878 | 0.714 | 0.739 | 1.8703E-06 | KO | Mertk |
| 3.8616E-10 | -1.0321106 | 0.016 | 0.029 | 1.9331E-06 | KO | Rgs9 |
| 3.8947E-10 | -0.3217112 | 0.125 | 0.155 | 1.9497E-06 | KO | Cideb |
| 5.5159E-10 | -0.2331422 | 0.327 | 0.363 | 2.7612E-06 | KO | Zbtb16 |
| 6.2053E-10 | -0.2660222 | 0.218 | 0.253 | 3.1064E-06 | KO | Prkcz |
| 6.2568E-10 | -0.2564858 | 0.213 | 0.247 | 3.1322E-06 | KO | Hnrnph1 |
| 6.6751E-10 | -0.3447652 | 0.088 | 0.114 | 3.3415E-06 | KO | Cers4 |
| 7.2477E-10 | -0.1198483 | 0.75 | 0.772 | 3.6282E-06 | KO | Rtn3 |
| 1.353E-09 | -0.414104 | 0.023 | 0.038 | 6.7731E-06 | KO | Calb2 |
| 1.65E-09 | -0.2677406 | 0.173 | 0.205 | 8.2597E-06 | KO | Slc14a1 |
| 2.0645E-09 | -0.1098709 | 0.761 | 0.781 | 1.0335E-05 | KO | Gpm6a |
| 2.3417E-09 | -0.1304551 | 0.582 | 0.617 | 1.1723E-05 | KO | Acly |
| 2.6663E-09 | -0.6893729 | 0.029 | 0.044 | 1.3348E-05 | KO | Ints11 |
| 2.8594E-09 | -0.2077746 | 0.251 | 0.287 | 1.4314E-05 | KO | Cdh20 |
| 4.1402E-09 | -0.3858541 | 0.052 | 0.073 | 2.0726E-05 | KO | Ednra |
| 7.0629E-09 | -0.2585622 | 0.303 | 0.335 | 3.5357E-05 | KO | Gap43 |
| 7.5523E-09 | -0.2922454 | 0.126 | 0.154 | 3.7807E-05 | KO | Cd151 |
| 8.0746E-09 | -0.1700241 | 0.366 | 0.403 | 4.0421E-05 | KO | Rab5a |
| 9.4555E-09 | -0.156674 | 0.391 | 0.428 | 4.7334E-05 | KO | Fam20a |
| 1.0032E-08 | -0.1683706 | 0.414 | 0.449 | 5.022E-05 | KO | Hist1h2bc |
| 1.2063E-08 | -0.527995 | 0.047 | 0.065 | 6.0385E-05 | KO | Fgf2 |
| 1.2218E-08 | -0.1869692 | 0.386 | 0.42 | 6.1166E-05 | KO | Epb41l1 |
| 1.3594E-08 | -0.8214606 | 0.015 | 0.026 | 6.8051E-05 | KO | Npas4 |
| 1.3754E-08 | -0.1596024 | 0.408 | 0.444 | 6.8853E-05 | KO | Gpt2 |
| 1.4751E-08 | -0.1204617 | 0.898 | 0.912 | 7.3845E-05 | KO | Map2 |

| | | | | | | |
|---|---|---|---|---|---|---|
| 1.542E-08 | -0.2160423 | 0.217 | 0.25 | 7.7191E-05 | KO | Exoc4 |
| 1.6102E-08 | -0.3046738 | 0.159 | 0.188 | 8.0607E-05 | KO | C4b |
| 1.6132E-08 | -1.0737691 | 0.008 | 0.017 | 8.0757E-05 | KO | Lhcgr |
| 1.7288E-08 | -0.2328581 | 0.186 | 0.217 | 8.6543E-05 | KO | Lgr4 |
| 2.7063E-08 | -0.4818297 | 0.054 | 0.073 | 0.00013548 | KO | Ifi27 |
| 2.711E-08 | -0.2129328 | 0.261 | 0.293 | 0.00013571 | KO | Tiparp |
| 3.2921E-08 | -0.1409839 | 0.595 | 0.628 | 0.0001648 | KO | Cdk5r2 |
| 3.3795E-08 | -0.1940116 | 0.346 | 0.377 | 0.00016918 | KO | Smarca2 |
| 3.4544E-08 | -0.3894784 | 0.126 | 0.151 | 0.00017293 | KO | Sez6 |
| 3.9321E-08 | -1.0623821 | 0.01 | 0.019 | 0.00019684 | KO | Tctn1 |
| 4.0277E-08 | -0.3291125 | 0.081 | 0.103 | 0.00020162 | KO | Map3k7 |
| 4.882E-08 | -0.2532547 | 0.183 | 0.211 | 0.00024439 | KO | Apc2 |
| 5.5042E-08 | -0.1470196 | 0.241 | 0.277 | 0.00027554 | KO | Fabp7 |
| 6.3906E-08 | -0.9918275 | 0.01 | 0.02 | 0.00031991 | KO | Bbox1 |
| 7.4333E-08 | -0.308821 | 0.102 | 0.126 | 0.00037211 | KO | Col1a2 |
| 8.7688E-08 | -0.4794256 | 0.039 | 0.055 | 0.00043897 | KO | Erg28 |
| 9.2472E-08 | -0.1634429 | 0.464 | 0.496 | 0.00046291 | KO | App |
| 9.4717E-08 | -0.177067 | 0.265 | 0.298 | 0.00047416 | KO | Eif4g1 |
| 1.0575E-07 | -0.4377773 | 0.054 | 0.072 | 0.0005294 | KO | Scn3b |
| 1.1668E-07 | -0.3556721 | 0.065 | 0.084 | 0.0005841 | KO | Tet1 |
| 1.2948E-07 | -0.1186343 | 0.57 | 0.599 | 0.00064818 | KO | Rnaset2a |
| 1.9813E-07 | -0.6588294 | 0.02 | 0.032 | 0.00099185 | KO | Dach1 |
| 2.2135E-07 | -0.3536768 | 0.071 | 0.091 | 0.00110808 | KO | Elk1 |
| 2.222E-07 | -0.1999327 | 0.209 | 0.239 | 0.00111236 | KO | Nde1 |
| 2.2375E-07 | -0.1911248 | 0.287 | 0.318 | 0.00112009 | KO | Bex3 |
| 2.3309E-07 | -0.5026747 | 0.062 | 0.08 | 0.00116685 | KO | Spock3 |
| 2.3937E-07 | -0.2866448 | 0.122 | 0.146 | 0.00119828 | KO | Paxbp1 |
| 2.5626E-07 | -0.3117853 | 0.122 | 0.146 | 0.00128283 | KO | Ilk |
| 2.7414E-07 | -0.1434548 | 0.582 | 0.604 | 0.00137235 | KO | Mapk8ip1 |
| 3.9882E-07 | -0.2094573 | 0.238 | 0.266 | 0.00199652 | KO | Itgb8 |
| 4.2514E-07 | -0.2174295 | 0.347 | 0.373 | 0.00212825 | KO | Itsn1 |
| 4.3183E-07 | -0.2731312 | 0.142 | 0.166 | 0.00216176 | KO | Galnt18 |
| 4.3564E-07 | -0.2022529 | 0.256 | 0.286 | 0.00218084 | KO | Ctsl |
| 4.3582E-07 | -0.2816596 | 0.138 | 0.163 | 0.0021817 | KO | Emx2 |
| 4.5004E-07 | -0.1141001 | 0.498 | 0.531 | 0.00225292 | KO | Vegfa |
| 4.5755E-07 | -0.2010215 | 0.257 | 0.286 | 0.00229049 | KO | Cit |
| 4.8324E-07 | -0.3083768 | 0.097 | 0.119 | 0.0024191 | KO | Arnt |
| 5.2849E-07 | -0.179596 | 0.34 | 0.371 | 0.00264562 | KO | Rcn2 |
| 5.4086E-07 | -0.1255788 | 0.442 | 0.476 | 0.00270755 | KO | Adcyap1r1 |
| 5.5016E-07 | -0.2009689 | 0.267 | 0.295 | 0.00275412 | KO | Camkv |
| 5.8362E-07 | -0.8255468 | 0.014 | 0.023 | 0.00292159 | KO | Kcna4 |
| 6.1304E-07 | -0.4248702 | 0.046 | 0.062 | 0.00306888 | KO | Efnb3 |
| 7.3889E-07 | -0.5120052 | 0.031 | 0.045 | 0.00369889 | KO | Kcng1 |
| 7.6338E-07 | -0.2209226 | 0.226 | 0.253 | 0.00382146 | KO | Cic |
| 7.98E-07 | -0.2648187 | 0.146 | 0.17 | 0.0039948 | KO | Cyld |
| 8.8437E-07 | -0.107084 | 0.699 | 0.717 | 0.00442717 | KO | Dpysl2 |
| 9.0005E-07 | -0.1959401 | 0.254 | 0.282 | 0.00450565 | KO | Rab5c |
| 9.7836E-07 | -0.3170045 | 0.064 | 0.081 | 0.00489766 | KO | Cdk5rap2 |
| 9.9824E-07 | -0.2440647 | 0.145 | 0.17 | 0.00499718 | KO | Atp2b4 |
| 1.4577E-06 | -0.4802296 | 0.033 | 0.046 | 0.00729735 | KO | Cd2ap |

| | | | | | | |
|---|---|---|---|---|---|---|
| 1.6952E-06 | -0.517457 | 0.039 | 0.053 | 0.00848633 | KO | Ugt8a |
| 1.7819E-06 | -0.6605177 | 0.021 | 0.031 | 0.00892023 | KO | Actn2 |
| 1.8401E-06 | -0.2221816 | 0.183 | 0.209 | 0.00921151 | KO | Rgcc |
| 1.8648E-06 | -0.3263017 | 0.077 | 0.095 | 0.00933504 | KO | Pfas |
| 1.9194E-06 | -0.4531219 | 0.033 | 0.046 | 0.00960872 | KO | Tgfbr3 |
| 2.0408E-06 | -0.3284832 | 0.068 | 0.086 | 0.01021646 | KO | Pcdh8 |
| 2.1541E-06 | -0.1800087 | 0.318 | 0.344 | 0.01078332 | KO | St6galnac5 |
| 3.211E-06 | -0.2655373 | 0.083 | 0.102 | 0.01607438 | KO | Stxbp4 |
| 3.4676E-06 | -0.4316517 | 0.043 | 0.057 | 0.01735889 | KO | Senp3 |
| 3.541E-06 | -0.1489225 | 0.324 | 0.352 | 0.01772614 | KO | Bin1 |
| 3.9135E-06 | -0.2485636 | 0.108 | 0.128 | 0.01959123 | KO | Nectin3 |
| 4.031E-06 | -0.2450314 | 0.134 | 0.156 | 0.02017928 | KO | Cpt1c |
| 5.1451E-06 | -0.6273595 | 0.028 | 0.039 | 0.0257566 | KO | Syn3 |
| 5.5518E-06 | -0.1412451 | 0.387 | 0.414 | 0.02779226 | KO | Mprip |
| 5.6914E-06 | -0.1740539 | 0.271 | 0.297 | 0.02849115 | KO | Rnd2 |
| 6.1916E-06 | -0.9641517 | 0.009 | 0.016 | 0.03099526 | KO | Dlx6 |
| 6.2957E-06 | -0.1845544 | 0.184 | 0.209 | 0.03151603 | KO | Ank2 |
| 8.2611E-06 | -0.2613803 | 0.091 | 0.11 | 0.04135498 | KO | Gtf2a2 |
| 8.5653E-06 | -0.2291981 | 0.106 | 0.126 | 0.04287814 | KO | C1qtnf5 |
| 8.7095E-06 | -0.2375369 | 0.118 | 0.138 | 0.04359957 | KO | Epb41l2 |
| 9.5959E-06 | -0.1801163 | 0.28 | 0.306 | 0.04803726 | KO | Bcr |
| 9.6678E-06 | -0.8424035 | 0.007 | 0.014 | 0.04839678 | KO | Abca8a |
| 9.9222E-06 | -0.2661525 | 0.12 | 0.14 | 0.04967032 | KO | Gopc |
| 9.9242E-06 | -0.2449201 | 0.143 | 0.164 | 0.04968053 | KO | Rragd |
